# *Plasmodium falciparum* SET10 is a histone H3 lysine K18 methyltransferase that participates in a chromatin modulation network crucial for intraerythrocytic development

**DOI:** 10.1101/2024.07.05.602231

**Authors:** Jean-Pierre Musabyimana, Sherihan Musa, Janice Manti, Ute Distler, Stefan Tenzer, Che Julius Ngwa, Gabriele Pradel

## Abstract

Lifecycle progression of the malaria parasite *Plasmodium falciparum* requires precise tuning of gene expression including histone methylation. The histone methyltransferase *Pf*SET10 was previously described as a H3K4 methyltransferase involved in *var* gene regulation, making it a prominent antimalarial target. In this study, we investigate the role of *Pf*SET10 in the blood stages of *P. falciparum* in more detail, using tagged *Pf*SET10-knockout (KO) and -knockdown (KD) lines. We demonstrate a nuclear localization of *Pf*SET10 with peak protein levels in schizonts. *Pf*SET10 deficiency results in reduced intraerythrocytic growth, but has no effect on gametocyte formation. When the *Pf*SET10-KO line is screened for histone methylation variations, lack of *Pf*SET10 renders the parasites unable to mark H3K18me1, while no significant changes in the H3K4 methylation status are observed. Comparative transcriptomic profiling of *Pf*SET10-KO schizonts demonstrates the upregulation of transcripts particularly encoding proteins linked to erythrocyte invasion and multigene family proteins, suggesting a repressive function of the histone methylation mark. TurboID coupled with mass spectrometry further reveals an extensive nuclear *Pf*SET10 interaction network with roles in transcriptional regulation, DNA replication and repair, chromatin remodeling and mRNA processing. Main interactors of *Pf*SET10 include ApiAP2 transcription factors, chromatin modulators like *Pf*MORC and *Pf*ISWI, mediators of RNA polymerase II, and DNA replication licensing factors. The combined data pinpoint *Pf*SET10 as a histone H3 lysine K18 methyltransferase of the *P. falciparum* blood stages that regulates nucleic acid metabolic processes as part of a comprehensive chromatin modulation network.

**Importance:** The fine-tuned regulation of DNA replication and transcription is particularly crucial for the rapidly multiplying blood stages of malaria parasites and proteins involved in these processes represent important drug targets. This study demonstrates that contrary to previous reports the histone methyltransferase *Pf*SET10 of the malaria parasite *Plasmodium falciparum* methylates histone 3 at lysine K18, a histone mark to date not well understood. Deficiency of *Pf*SET10 due to genetic knockout affects genes involved in intraerythrocytic development. Furthermore, in the nuclei of blood stage parasites, *Pf*SET10 interacts with various protein complexes crucial for DNA replication, remodeling and repair, as well as for transcriptional regulation and mRNA processing. In summary, this study highlights *Pf*SET10 as a H3K18 methyltransferase with critical functions in chromatin maintenance during the development of *P. falciparum* in red blood cells.

## Introduction

Understanding the epigenetic mechanisms of gene regulation is critical for identifying drivers of disease as a first step towards therapies targeting transcriptional mechanisms of disease-onset. The use of anti-epigenetic agents is among others currently being explored for cancer and pain therapy with the first epigenome-targeting drugs having received approval (reviewed in, e.g., Weinhold, 2006; Lee et al., 2014; Ghasemi, 2020; Zohourian and Brown, 2024). The importance of epigenetics has also been acknowledged as a main driver of lifecycle progression of malaria parasites, which are responsible for 249 million infections and 608,000 deaths per year (WHO World Malaria Report 2023). When the 23-Mb genome of *Plasmodium falciparum*, the causative agent of malaria tropica, was sequenced in 2002, roughly 5,300 genes coding for core proteins including a varying number of subtelomeric multigene families like *var*, *rif*, *stevor* and *pfmc-2tm* were identified (Gardner et al., 2002). Subsequent proteomic and transcriptomic analyses demonstrated lifecycle stage specificity for the majority of the gene products (e.g., Florens et al., 2002; Lasonder et al., 2002, 2016; Le Roch et al., 2003; López-Barragán et al., 2011) and permitted a first glimpse on the tightly regulated transcriptional program that is based on well-coordinated sequences of gene activation and silencing needed by the parasite to progress from one lifecycle stage to another.

A significant part of epigenetic control in *Plasmodium* involves the posttranslational modification (PTM) of its histones, e.g. via histone acetylation and methylation (reviewed in, e.g., Cui and Miao, 2010; Jabeena and Rajavelu, 2019). Histone PTMs in malaria parasites have particularly been studied during expression of the 60 *var* genes encoding the *P. falciparum* erythrocyte membrane protein *Pf*EMP1 (Lopez-Rubio et al., 2007, 2009; Petter et al., 2011; e.g. reviewed in Llinás et al., 2008; Cui and Miao, 2010; Duffy et al., 2014; Voss et al., 2014; Duraisingh and Horn, 2016). The switch of *var* gene expression and thus *Pf*EMP1 structure alters the antigenic type of the infected red blood cells (RBCs) and in consequence pathogenesis of malaria. Only the active *var* gene copy exhibits a euchromatic state characterized by the two histone PTMs H3K9ac and H3K4me3, while *var* gene silencing is linked to H3K9me3 and H3K36me3 and the binding of heterochromatin protein HP1 (e.g., Duraisingh et al., 2005; Freitas-Junior et al., 2005; Lopez-Rubio et al., 2007, 2009; Pérez-Toledo et al., 2009; Salcedo-Amaya et al., 2009; Tonkin et al., 2009; Jiang et al., 2013; Brancucci et al., 2014; Coleman et al., 2014). Overarching regulatory structures like reader complexes or nucleosome composition modifications support *var*-regulating PTMs (e.g., Herrera-Solorio et al., 2019; Hoeijmakers et al., 2019). Additional factors regulating the monoallelic expression of *var* genes include long noncoding RNAs (lncRNA) (Amit-Avraham et al., 2015; Bacons-Simon et al., 2020).

The genome of *P. falciparum* encodes ten SET (Su(var)3-9-’Enhancer of zeste-Trithorax)-domain-containing lysine-specific histone methyltransferases (HMTs), termed *Pf*SET1 to *Pf*SET10, which are abundantly transcribed in the asexual and sexual blood stages (Cui et al., 2008; Volz et al., 2010; Ngwa et al., 2019). Treatment of blood stage parasites with the commercially available inhibitor BIX-01294, known to target G9a lysine-specific HMTs, resulted in impaired intraerythrocytic replication, gametocyte development and gametogenesis (Ngwa et al., 2019). Comparative transcriptomics between BIX-01294-treated and untreated immature, mature and activated gametocytes further demonstrated a deregulation of various genes, particularly affecting antigenic variation, translation and host cell remodeling.

To date, the *Pf*SET proteins have not been investigated in detail. Previous studies indicated that *Pf*SET2 and *Pf*SET10 are important for immune evasion of *P. falciparum*. While *Pf*SET2 (also termed *Pf*SETvs) is responsible for H3K36me3 marks needed to repress *var* gene expression (Jiang et al., 2013), *Pf*SET10 was described as a H3K4 methyltransferase with suggested essential functions in maintaining the active *var* gene in a poised state during cellular division (Volz et al., 2012). However, a subsequent report demonstrated that *Pf*SET10 deficiency had no significant effect either on parasite viability or on *var* gene expression (Ngwa et al., 2021), leaving its precise function unclear.

Here, we investigated the role of *Pf*SET10 for the *P. falciparum* blood stages in further detail. We demonstrate that *Pf*SET10 is a H3K18 methyltransferase with no activity in marking H3K4. Lack of *Pf*SET10 deregulates the expression of genes encoding for RBC invasion and exported proteins as well as multigene family proteins. Further, *Pf*SET10 forms extensive networks with proteins important for the DNA replication, RNA synthesis and chromatin modulation.

## Results

### *Pf*SET10 localizes to the nuclei of the *P. falciparum* blood stages

*Pf*SET10 is a 271-kDa protein (PF3D7_1221000; 2329 aa) comprising a central SET and PHD zinc finger domain (Fig. 1A). For functional characterization, we generated a conditional *Pf*SET10-HA-KD line (using the pSLI-HA-*glmS* vector) by fusing the 3′-region of *pfset10* with the sequences coding for a hemagglutinin A (HA)-tag and for the *glmS* element (Fig. S1A; Prommana et al., 2013; Musabyimana et al., 2022). Vector integration was confirmed by diagnostic PCR (Fig. S1A, B). To prove the conditional down-regulation of *Pf*SET10-HA synthesis, asexual blood stage parasites of line *Pf*SET10-HA-KD were treated with 2.5 and 5 mM glucosamine hydrochloride (GlcN) for 72 h and lysates were subjected to Western blotting. In untreated cultures, a prominent *Pf*SET10-HA band was detected at ~275 kDa. Upon addition of GlcN, *Pf*SET10-HA levels were significantly reduced to 68.6 ± 19.91% (2.5 mM) and 15.8 ± 7.51% (5 mM), respectively, compared to the untreated *Pf*SET10-HA-KD control, as shown by quantitative Western blotting (Fig. S2A, B).

**Figure 1.**
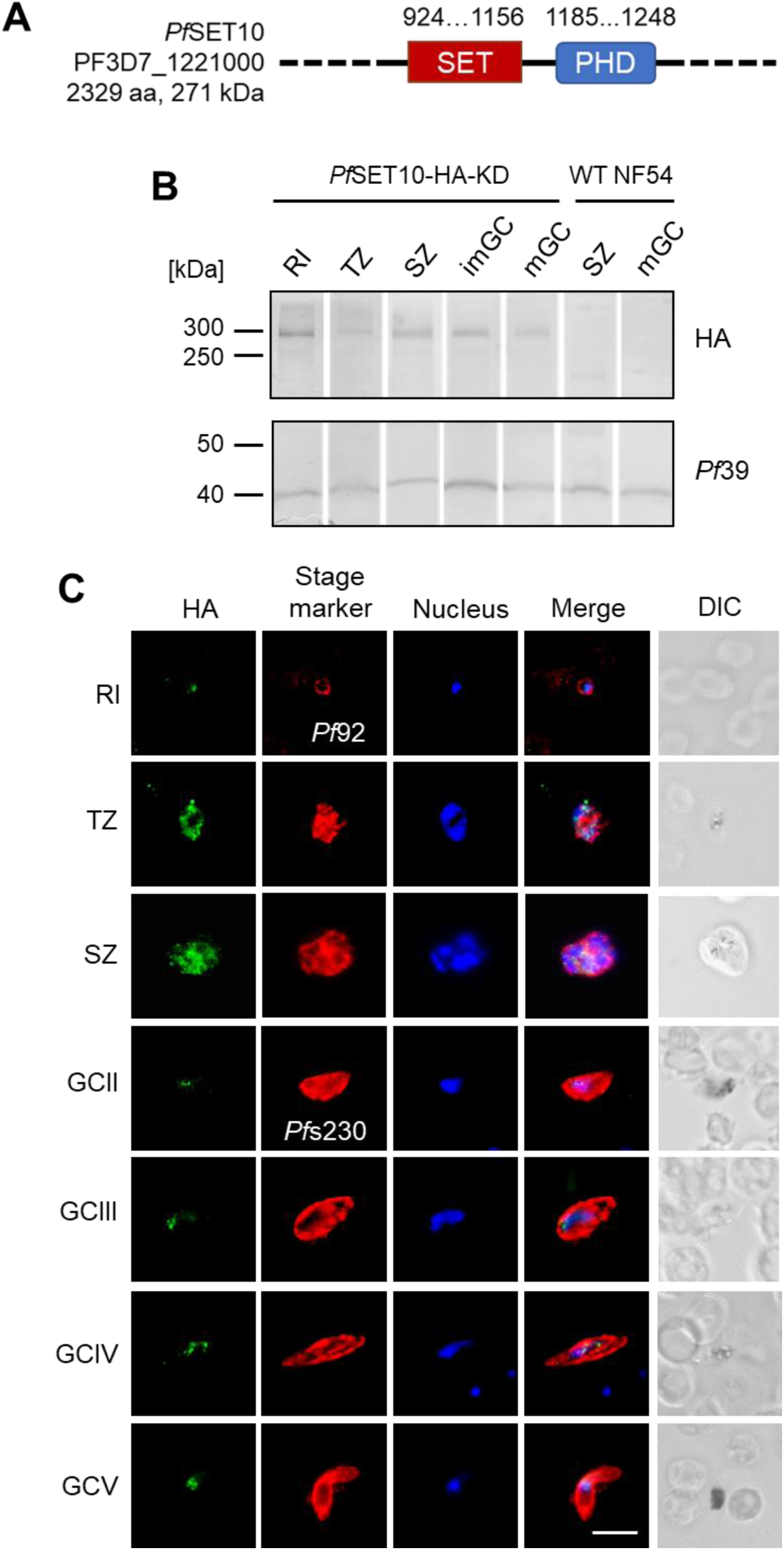
*Pf*SET10 is expressed in the *P. falciparum* blood stages. **(A)** Schematic depicting *Pf*SET10. The positions of the SET and PHD domains are indicated. **(B)** Protein expression of *Pf*SET10-HA in blood stage parasites. Western blot analysis of lysates from rings (RI), trophozoites (TZ), schizonts (SZ), immature and mature gametocytes (imGC, mGC) of line *Pf*SET10-HA-KD was employed, using rat anti-HA antibody to detected *Pf*SET10-HA (~275 kDa). As negative control, WT NF54 lysate was used. Immunoblotting with rabbit antisera against *Pf*39 (~39 kDa) served as loading control. (**C**) Localization of *Pf*SET10-HA in the blood stages. Methanol-fixed RI, TZ, SZ, and GC stages II-V of line *Pf*SET10-HA-KD were immunolabeled with rat anti-HA antibody (green). Asexual blood stages and gametocytes were highlighted with rabbit antisera directed against *Pf*92 and *Pf*s230, respectively (red); nuclei were highlighted with Hoechst 33342 nuclear stain (blue). Bar, 5 μm. Results (B, C) are representative of three independent experiments.

The untreated *Pf*SET10-HA-KD line was initially used for in-depth expression analysis. Western blotting using anti-HA antibody demonstrated the presence of *Pf*SET10-HA in asexual blood stages and immature and mature gametocytes, whereas no protein band was detected in the WT NF54 (Fig. 1B). Immunofluorescence assays further assigned *Pf*SET10 to the parasites’ nuclei with particular intense signals detected in schizonts (Fig. 1C). No labeling was observed in the WT NF54 blood stages (Fig. S3A).

### *Pf*SET10 deficiency affects intraerythrocytic growth, but not gametocyte formation

We tested the effect of *Pf*SET10 deficiency on intraerythrocytic growth and gametocyte development, using an established *Pf*SET10-KO line, which was previously generated via selection-linked integration-mediated targeted gene disruption (using vector pSLI-TGD) (Ngwa et al., 2021). The transgenic parasites express a truncated N-terminal fragment of *Pf*SET10 fused with green fluorescent protein (GFP), which lacks the SET and PHD domains. We previously showed that the asexual blood stage parasites of *Pf*SET10-KO displayed normal morphologies (Ngwa et al., 2021); in addition, no morphological differences were observed in *Pf*SET10-KO gametocytes (Fig. S3B). In conformation with our previous reports, *Pf*SET10-KO parasites exhibited reduced parasitemia compared to WT NF54 with normal stage progression through the replication cycles (Fig. S4A, B). No significant differences in gametocyte development were observed between *Pf*SET10-KO and WT NF54 (Fig. S4C).

Intraerythrocytic development was also investigated in the *Pf*SET10-HA-KD line. Here, growth assays revealed no differences in parasitemia, when synchronized asexual blood stages of line *Pf*SET10-HA-KD were treated with 2.5 mM GlcN over a period of 72 h; compared to untreated cultures; GlcN-treated and untreated WT NF54 parasites served as controls (Fig. S5A, B). Furthermore, no differences in gametocyte numbers were observed between line *Pf*SET10-HA-KD and WT NF54 (Fig. S5C). To investigate both gametocyte commitment and gametocyte maturation, we treated gametocytes of line *Pf*SET10-HA-KD with GlcN either prior to or after induction and treatment was continued for 7 days, while untreated transgenic parasites and treated and untreated WT NF54 gametocytes served as controls.

The combined results assign *Pf*SET10 to the blood stage nuclei with peak expression in schizonts and indicate functions of *Pf*SET10 during intraerythrocytic replication of the parasite.

### *Pf*SET10 deficiency abolishes H3K18 mono-methylation, but not H3K4 methylation

To determine the methylation activity of *Pf*SET10, we investigated the effect of *Pf*SET10 deficiency on seven histone 3 methylation marks, for which commercial antibodies were available, i.e. H3K4me1, H3K4me2, H3K4me3, H3K36me1, H3K36me2, H3K36me3, and H3K18me1. Lysates generated from asexual blood stages of line *Pf*SET10-KO and WT NF54 were immunoblotted with rabbit antibodies directed against the above H3 lysine methylation marks. No differences in band intensities were identified between *Pf*SET10-KO and WT NF54 for any of the methylation marks concerning lysines H3K4 and H3K36 (Fig. 2A). In contrast, no signal was detected for the methylation mark H3K18me1 in the *Pf*SET10-KO line (Fig. 2B). Immunoblotting with antibodies directed against histone H3 and *Pf*39 served as loading controls. For the methylation mark H3K4me3, band signals from three independent Western blots were quantified and demonstrated similar signal intensities for line *Pf*SET10-KO and WT NF54 (Fig. 2C).

**Figure 2.**
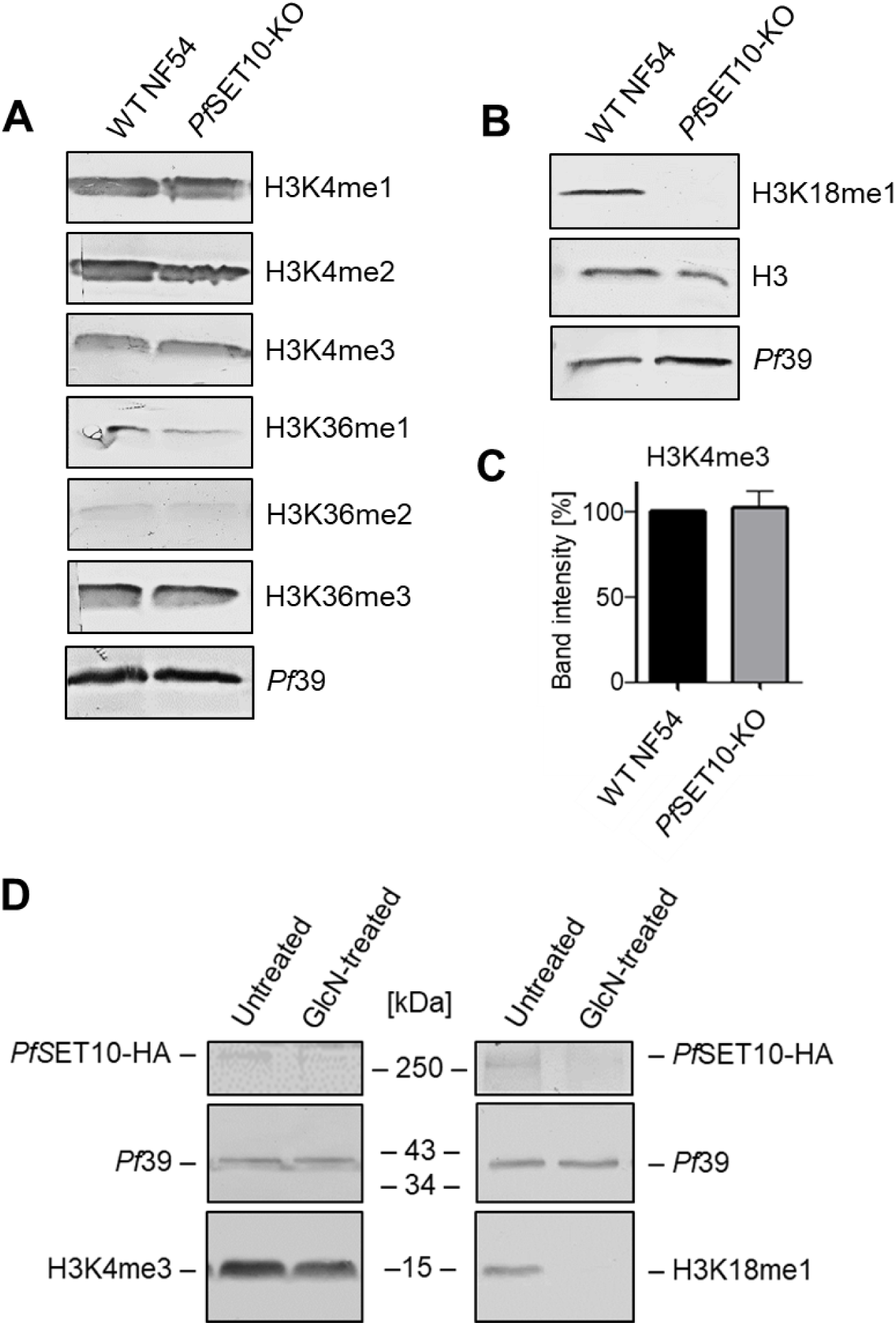
*Pf*SET10 deficiency affects H3K18me1 methylation. (**A**, **B**) Lysates of asexual blood stage parasites of line *Pf*SET10-KO and WT NF54 were immunoblotted with rabbit antibodies directed against selected histone 3 (H3) methylation marks as indicated (~15 kDa). Rabbit antisera directed against H3 and *Pf*39 (~39 kDa) were used as loading controls. (**C**) Quantification of the H3K4me3 signal. Western blots were performed, using rabbit anti-H3K4me3 antibody as described in (A). H3K4me3 signals were evaluated by measuring the band intensities of three independent immunoblots using Image J; the values were normalized to the respective *Pf*39 protein band. Results are shown as mean ± SD (WT NF54 set to 100%). (**D**) Asexual blood stage parasites of line *Pf*SET10-HA-KD were treated or not with 2.5 μM GlcN and the lysates were immunoblotted with H3K4me3 and H3K18me1 antibodies to detect the respective methylation mark (~15 kDa). *Pf*SET10-HA (~250 kDa) was detected with anti-HA antibodies; blotting with anti-*Pf*39 antisera was used as loading control. The results (A, B, D) are representative of two to three independent experiments.

The methylation effects of *Pf*SET10 on H3K4me3 and H3K18me1 were confirmed using line *Pf*SET10-HA-KD. Asexual blood stages of line *Pf*SET10-HA-KD were treated with 2.5 mM GlcN for 72 h. Lysates were immunoblotted with anti-HA antibodies to highlight *Pf*SET10-HA in the samples, while the H3K4me3 and H3K18me1 methylation marks were detected with the respective antibodies. Treatment of line *Pf*SET10-HA-KD with GlcN reduced the *Pf*SET10-HA levels, as expected, and further abolished the H3K18me1 mark, while the H3K4me3 signal did not alter (Fig. 2D). The levels of *Pf*39, used for loading and viability control, where not affected by the GlcN treatment.

We then investigated the H3K18me1 mark in the asexual and sexual blood stages during stage development. Western blotting demonstrated particularly prominent signals in the asexual blood stages, while the methylation mark vanished in the maturing gametocytes (Fig. 3A). Immunofluorescence assays further assigned the H3K18me1 mark to the blood stage nuclei and confirmed strong signals in schizonts (Fig. 3B). Furthermore, the H3K18me1 methylation mark overlapped with the *Pf*SET10-HA signal, when the *Pf*SET10-HA-KD line was employed in immunofluorescence assays (Fig. 3C).

**Figure 3.**
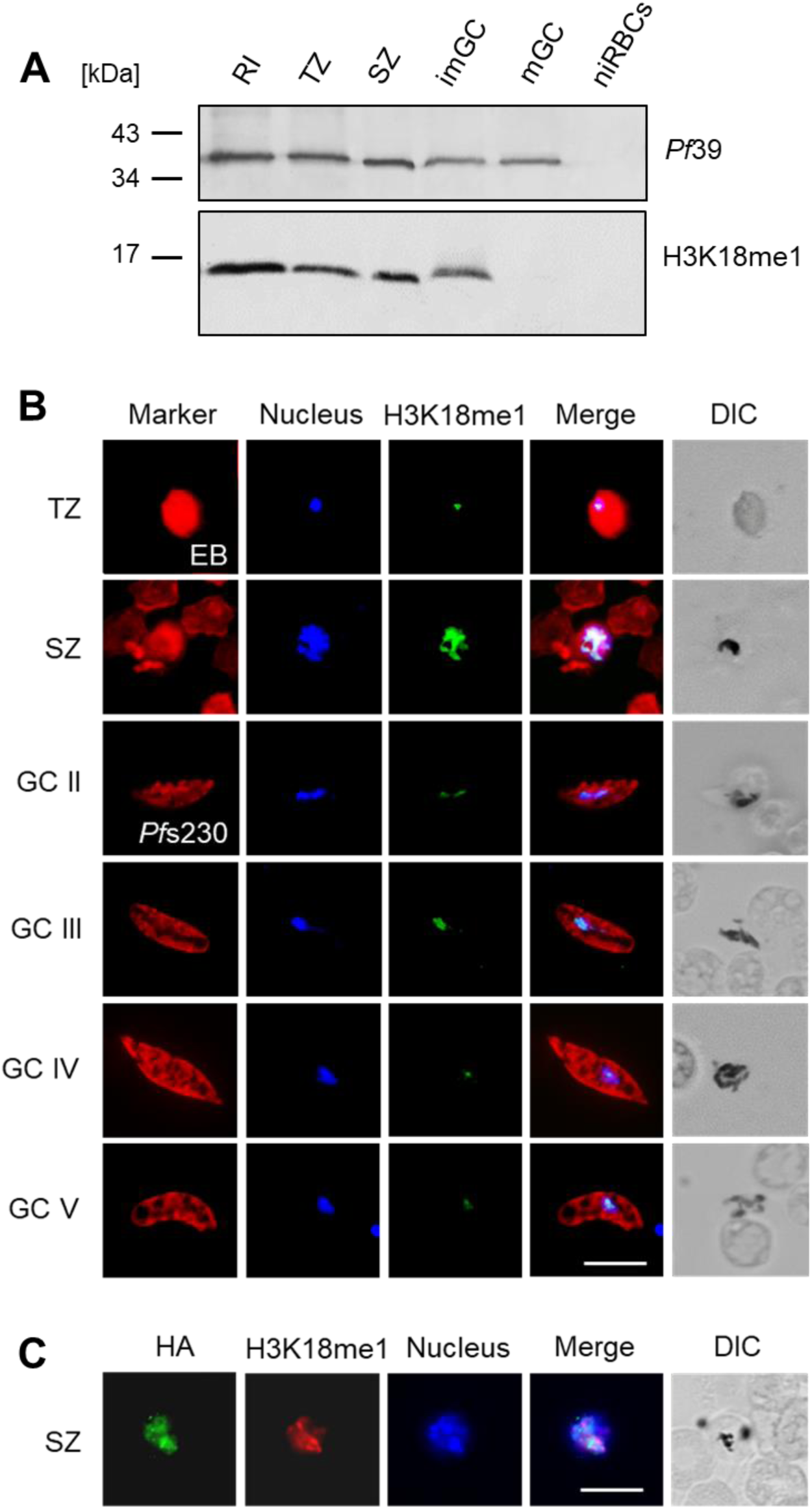
The H3K18me1 methylation mark associates with blood stage nuclei. (**A**) H3K18me1 levels in the parasite blood stages. Lysates of WT NF54 rings (RI), trophozoites (TZ), schizonts (SZ), and immature and mature gametocytes (imGC, mGC) were immunoblotted with rabbit anti-H3K18me1 antibodies to detect the methylation mark (~15 kDa). Immunoblotting with rabbit antisera directed against *Pf*39 (~39 kDa) was used as loading control. (**B**) Localization of H3K18me1 in the blood stage nuclei. Methanol-fixed rings RI, TZ, SZ, and GC stages II-V of WT NF54 were immunolabeled with rabbit anti-H3K18me1 antibody (green). Asexual blood stages and gametocytes were highlighted with Evans blue and mouse anti-*Pf*s230 antisera, respectively (red); nuclei were highlighted with Hoechst 33342 nuclear stain (blue). (**C**) The H3K18me1 mark colocalizes with *Pf*SET10. Schizonts of line *Pf*SETS10-HA-KD were immunolabelled as described in (B), using anti-H3K18me1 antibody (red), while rat anti-HA antibody was used to detect *Pf*SET10-HA (green). Bar, 5 μm. The results (A-C) are representative of two to three independent experiments.

The combined data pinpoint *Pf*SET10 as a H3K18 methyltransferase with particular methylation activity in the asexual blood stages, but no detectable effect towards H3K4me3 methylation.

### *Pf*SET10 deficiency results in the upregulation of genes linked to antigenic variation and invasion

We carried out a comparative transcriptomics analysis using RNA isolated from schizonts of line *Pf*SET10-KO and WT NF54. A total of 139 deregulated genes with a log-fold change greater than two were identified by RNA-seq, of which the majority (121 genes) were transcriptionally upregulated in the *Pf*SET10-KO, while 18 genes were transcriptionally downregulated (Fig. 4A; Table S1). Genes that were transcriptionally upregulated had peak expression assigned to the ring and ookinete stages (PlasmoDB database), whereas downregulated genes exhibited peak expression in rings and trophozoites (Fig. 4B).

**Figure 4.**
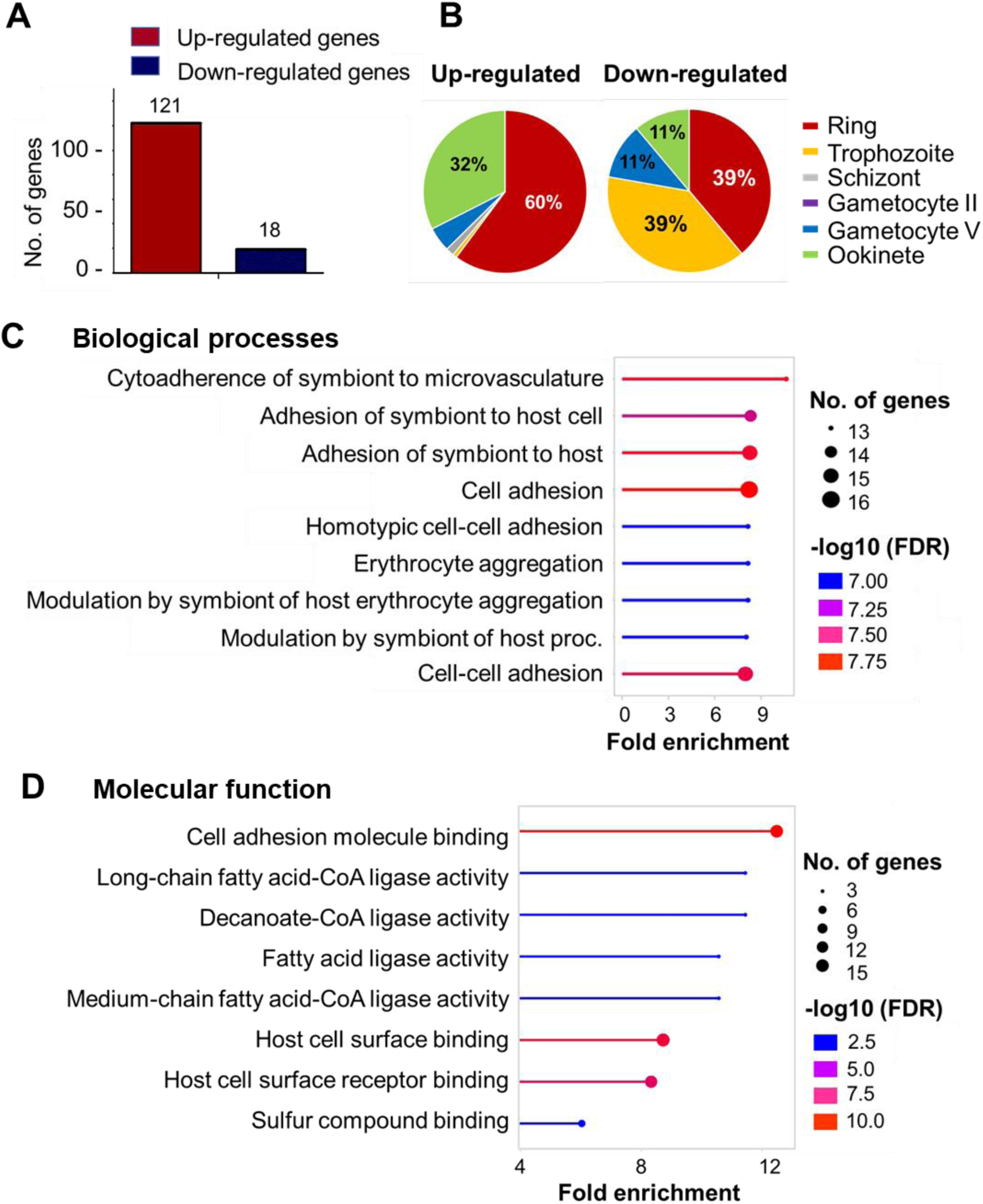
*Pf*SET10 deficiency deregulates genes involved in invasion and cytoadherence. (**A**) Numbers of transcriptionally deregulated genes in line *Pf*SET10-KO. RNA-seq-based comparative transcriptomics analyses were performed on transcripts of schizonts of line *Pf*SET10-KO and WT NF54, resulting in the deregulation of 139 genes. (**B**) Pie chart depicting the deregulated genes in line *Pf*SET10-KO (percentage of total numbers) grouped by peak expression (PlasmoDB). (**C**, **D**) Functional prediction analysis of the transcriptionally up-regulated genes in the *Pf*SET10-KO. A GO enrichment analysis was performed (ShinyGO 0.77 program; p < 0.05) and the enriched GO terms were based on biological process **(C)** and molecular function **(D)**.

Gene ontology (GO) enrichment analysis assigned the transcriptionally upregulated genes to biological processes such as cytoadherence, cell adhesion and erythrocyte aggregation with molecular functions in cell adhesion molecule and host cell surface binding. Further molecular functions included coenzyme A ligase and fatty acid ligase activities (Fig. 4C, D). Upregulated RNAs included those coding for members of the *var*, *rif*, and *stevor* multigene families, RBC invasion-related proteins such as EBA175, GAMA, and MSP8, proteins of the infected RBC membrane and the Maureŕs clefts like CLAG3.2, SBP1, HRP, RESA, RESA3, REX1, MAHRP2, MaTrA as well as lysophospholipase LPL120 and three acyl-CoA synthetases; further three non-coding RNAs were identified. Down-regulated genes included mainly those coding for chaperones and ribosomes units (Table S1).

We subsequently investigated the impact of *Pf*SET10 deficiency on protein synthesis of four STEVOR proteins (PF3D7_0300400, PF_0324600, PF3D7_0631900, PF3D7_1254100), which were transcriptionally upregulated in the comparative RNA-seq analysis. Lysates of *Pf*SET10-KO and WT NF54 schizonts were subjected to Western blotting, using rabbit antibodies directed against the four STEVOR proteins (Wichers et al., 2019), while rabbit antibodies directed against *Pf*92 and *Pf*39 served as stage and loading control, respectively (Fig. 5). Immunoblotting detected increased band intensities for all of the STEVOR proteins in the absence of *Pf*SET10. Immunoblotting with anti-H3K18me1 antibodies confirmed the absence of this methylation mark in the *Pf*SET10-KO line.

**Figure 5.**
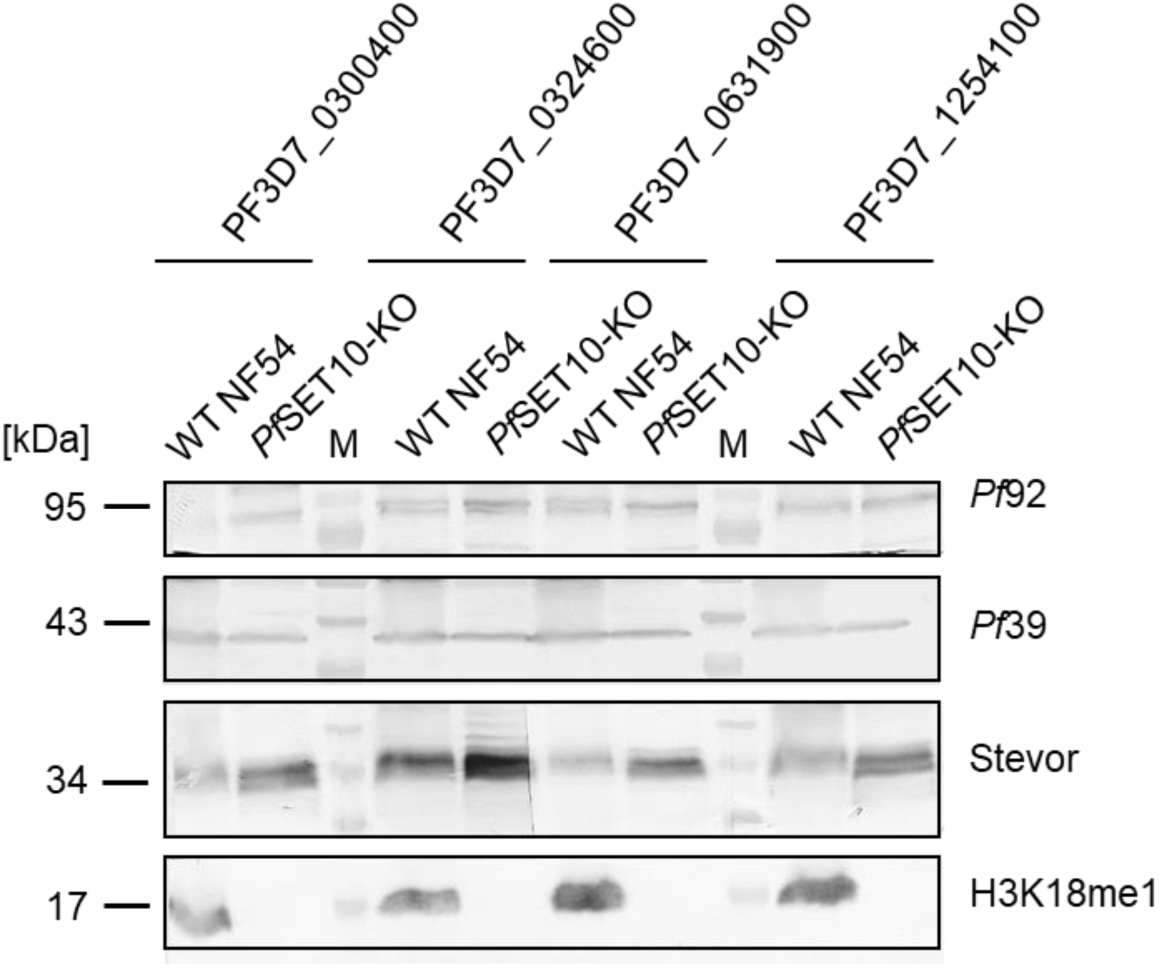
*Pf*SET10 deficiency affects STEVOR expression. Lysates of asexual blood stage parasites of line *Pf*SET10-KO and WT NF54 were immunoblotted with rabbit antibodies directed against selected STEVOR proteins (~35 kDa; indicated by geneID). Rabbit antisera directed against *Pf*92 (~92 kDa) and *Pf*39 (~39 kDa) were used as loading and stage controls; rabbit anti-H3K18me1 antibodies highlight the methylation mark (~15 kDa). M, molecular weight marker. The results are representative of two independent experiments.

The transcriptomics analyses demonstrate that lack of *Pf*SET10 leads to an upregulation of genes assigned to RBC invasion, antigenic variation and host cell remodeling, suggesting that *Pf*SET10 acts as a repressive HMT during regulation of intraerythrocytic replication.

### The *Pf*SET10 interactome comprises proteins of DNA and RNA metabolic processes

To determine the *Pf*SET10 interaction network, we generated a transgenic line that endogenously expresses *Pf*SET10 fused with the enhanced TurboID version of BirA in addition to GFP, using vector pSLI-TurboID-GFP (Fig. S6A; Branon et al. 2018; Sassmannshausen et al., 2024). Successfull vector integration was demonstrated by diagnostic PCR (Fig. S6B).

Western blot analyses of asexual blood stage lysates prepared from the transgenic line, using mouse anti-GFP antibody, demonstrated the expression of *Pf*SET10-TurboID-GFP with a molecular weight of ~300 kDa (Fig. S6C). No protein band was detected in lysates of non-infected RBCs; immunoblotting with anti-*Pf*39 served as loading control. Immunofluorescence analyses, using anti-GFP antibodies, confirmed the expression of *Pf*SET10-TurboID-GFP in the transgenic blood stage parasites and demonstrated the nuclear localization of the fusion protein (Fig. S6D).

Subsequent Western blotting was employed to confirm protein biotinylation in the transgenic line. Asexual blood stages of line *Pf*SET10-TurboID-GFP were treated with 50 µM biotin for 10 min. Immunoblotting of the respective lysates using streptavidin conjugated to alkaline phosphatase detected multiple protein bands indicative of biotinylated proteins, including a protein band of ~300 kDa, likely representing biotinylated *Pf*SET10-TurboID-GFP (Fig. S7A). In lysates of transgenic parasites that were not treated with biotin, only weak protein bands were detected, indicative of endogenous biotin, and no biotin-positive protein bands were detected in the biotin-treated WT NF54. Immunofluorescence assays, using fluorophore-conjugated streptavidin, confirmed the presence of biotinylated proteins in the nuclei of schizonts and the signal overlapped with the *Pf*SET10-TurboID-GFP signal, while no biotinylated proteins were detected in biotin-treated WT NF54 parasites (Fig. S7B).

The biotinylated proteins were purified from schizonts of line *Pf*SET10-TurboID-GFP via streptavidin-coated beads; WT NF54 lysates served as control. The biotinylated proteins were then prepared for mass spectrometry to identify potential *Pf*SET10 interactors. A total of 1,203 proteins were identified (Tables S2, S3). Among the top hits sorted by peptide count (≥ 50) were the asparagine and aspartate rich protein 1 (AARP1), three ApiAP2 transcription factors (PF3D7_0516800, PF3D7_0613800, PF3D7_1239200), and the transcriptional regulator *Pf*MORC, as well as the epigenetic regulators *Pf*PHD1 and *Pf*PHD2, *Pf*SET1 and *Pf*HDA1 in addition to *Pf*SET10 (Table S4).

The hits were subjected to two curation steps (Fig. S7C). Initially, we excluded proteins with predicted signal peptides, leading to a total of 1,139 proteins. Subsequently, we focused on proteins with predicted nuclear localization (PlasmoDB database), resulting in 801 hits (Figs. 6A, S7C). The nuclear proteins exhibited peak expression profiles assigned to the ring and ookinete stages as well as to trophozoites and mature gametocytes (Fig. 6B). Further, GO term analysis highlighted the involvement of *Pf*SET10 interactors in metabolic processes of nucleic acids and cellular macromolecules as well as of nitrogen compounds (Fig. 6C). Molecular functions that involve the potential *Pf*SET10 interactors comprise binding of DNA and RNA, ATP and proteins (Fig.6D).

**Figure 6.**
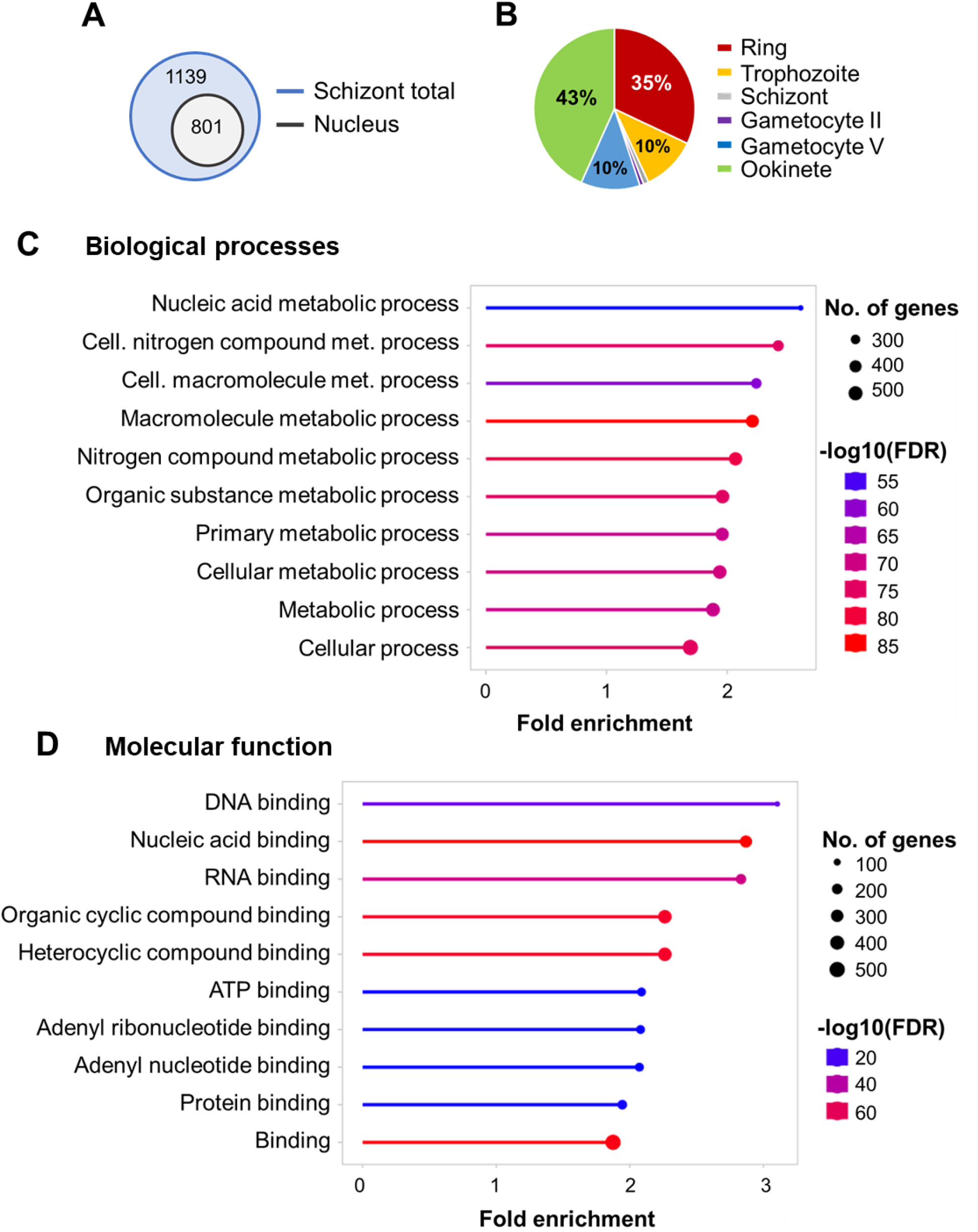
*Pf*SET10 interacts with nuclear proteins involved in nucleic acid metabolic processes. (**A**) Numbers of *Pf*SET10 interactors in schizonts. TurboID analysis was performed using schizonts of line *Pf*SET10-TurboID-GFP, resulting in the identification of 1,139 interactors (w/o signal peptide), of which 801 were assigned to the nucleus. (**B**) Pie chart depicting the nuclear *Pf*SET10 interactors (percentage of total numbers) grouped by peak expression (PlasmoDB). (**C**, **D**) Functional prediction analysis of the nuclear *Pf*SET10 interactors. A GO enrichment analysis was performed (ShinyGO 0.77 program; p < 0.05) and the enriched GO terms were based on biological process **(C)** and molecular function **(D)**.

A STRING-based protein-protein interaction analysis was conducted, using the K-means algorithm with a confidence level of 0.9 and a maximal cluster number of 43. Based on the physical interaction among the query proteins, 15 main clusters were identified (Fig. S8; Table S5). The most extensive clusters could be assigned to ribosome and translation (cluster red), the proteasome (cluster brown) and the spliceosome (cluster salmon). Further important clusters included transcription (cluster fire brick), DNA replication (cluster sienna) and chromatin remodeling (cluster dark golden rod). Other clusters included proteins involved in mRNA processing, the replication factor C complex, DNA repair proteins, the prefoldin & T-complex, chromatin condensation regulators, the ribonucleotide reductase (RNR) complex, the CCR-NOT complex, a chaperone complex and proteins involved in nuclear export and import (Fig. S8; Table S5).

A second STRING analysis was performed after excluding proteins from the two most prominent protein complexes of the nucleus, i.e. ribosomal and proteasomal proteins (Fig. S7C). Following this curation step, various sub-clusters were found (K means; confidence level of 0.9; cluster number of 29). Several of these were assigned to three megaclusters (Fig. S9; Table S6). The most extensive megacluster (cluster red) comprised subclusters like the spliceosome complex, the ribosomal subunit biogenesis complex, the exosome complex, the CCR-NOT complex and proteins of chromatin remodeling. A second megacluster (cluster salmon) involved proteins of transcription and mRNA processing as well as DNA replication and repair, including the replication factor C complex. The third megacluster (cluster fire brick) comprised proteins of translation and nuclear export-import. Further smaller clusters included a V-type proton ATPase cluster, a prefoldin and a T-complex cluster, chromatin condensation proteins, a ribonuclease P and a RNR complex, as well as a ubiquitin fusion degradation complex (Fig. S9; Table S6).

The curated nuclear proteins were further investigated for proteins that are assigned to the two main functions of *Pf*SET10, i.e. transcription and DNA replication (Tables 1, S4). A total of 15 ApiAP2 transcription factors and eight other transcription factors were identified. In addition, 22 epigenetic regulators (including *Pf*SET10) were found. Furthermore, 17 RNA polymerases, 10 mediators of RNA polymerase II (MED proteins), 21 RNA helicases and further proteins are related to transcription. In addition, various proteins could be assigned to DNA replication, including three DNA polymerase subunits, 12 DNA helicases, six replication factor A and C subunits, eight DNA replication licensing factors (MCM proteins), further proteins have assigned functions in, e.g., DNA repair and condensation as well as in the regulation of replication (Tables 1, S4). Noteworthy, we further identified chromatin remodeling proteins, which in addition to *Pf*MORC, include *Pf*ISWI, *Pf*SWIP, *Pf*SNF2L, *Pf*CHD1, and *Pf*CAF1A and *Pf*CAF1B.

**Table 1.**
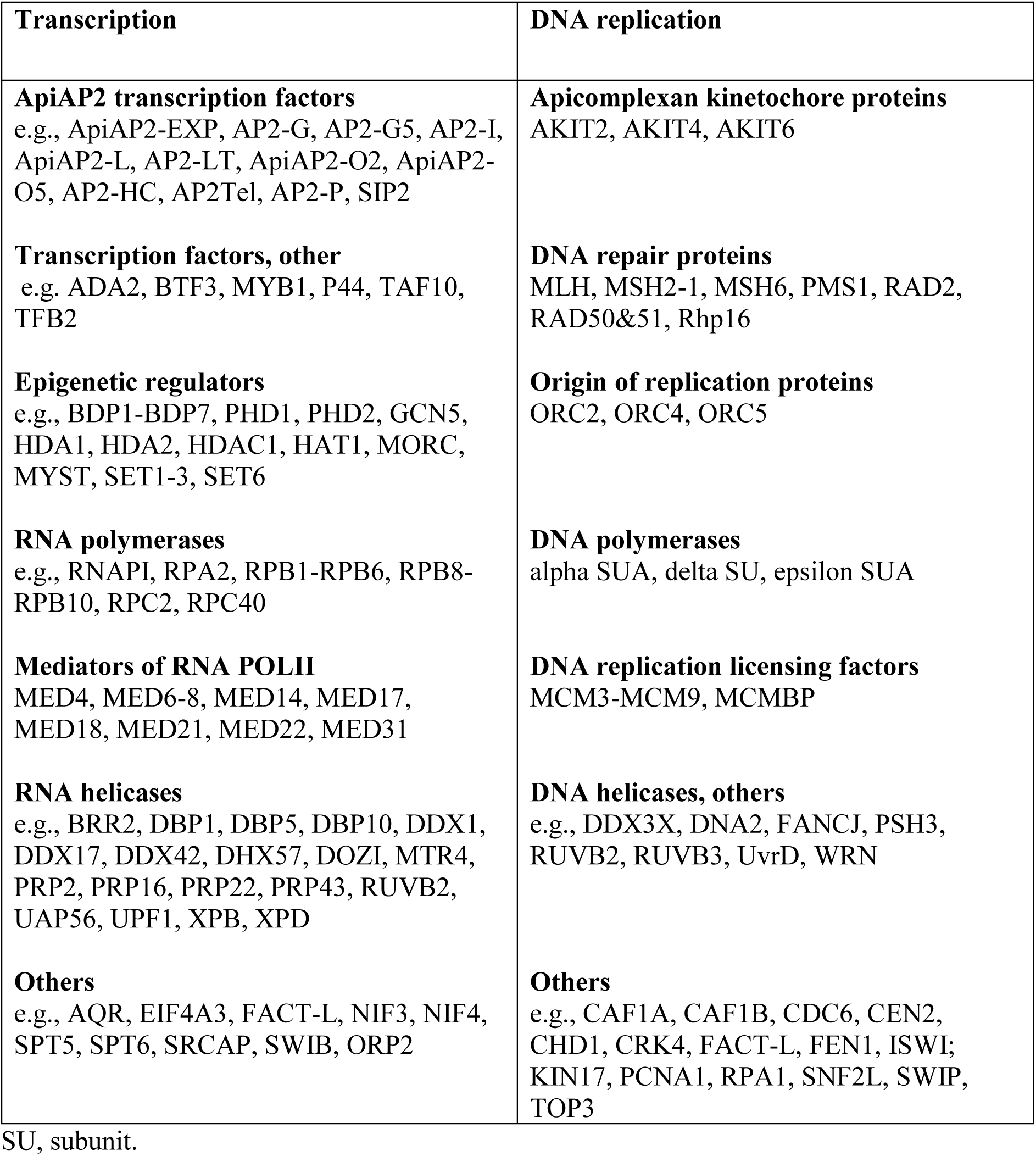
Selected interactors of *Pf*SET10 with involvement in transcription and DNA replication.

In summary, the protein-protein interaction analyses revealed main functions of *Pf*SET10 in nuclear metabolic processes, in particular of DNA replication and RNA synthesis and processing.

### *Pf*SET10 interacts with *Pf*AP2-P, *Pf*ISWI, and *Pf*MORC as part of a chromatin modulation network

Among the top hits of nuclear interactors of *Pf*SET10 are three proteins recently identified as chromatin modifiers, i.e. the chromatin-associated microrchidia protein *Pf*MORC (hit no. 2), ApiAP2 transcription factor *Pf*AP2-P (hit no. 40), and the *var* gene promotor-associated chromatin remodeling protein *Pf*ISWI (hit no. 43, see Table S6), for which protein interactomics data are available (Bryant et al., 2020; Chahine et al., 2023; Subudhi et al., 2023; Singh et al., 2024). We thus compared the interactors of *Pf*AP2-P, *Pf*ISWI, *Pf*MORC, and *Pf*SET10, and identified a total of 328 proteins, with which at least two of the bait proteins interacted (Fig. 7A; Table S7; bait proteins included). The majority of interactions were found between *Pf*SET10 and *Pf*AP2-P as well as between *Pf*SET10 and PfMORC. Two interactors were shared by all four bait proteins, i.e. HDAC1 and MCM5 (bait proteins excluded).

**Figure 7.**
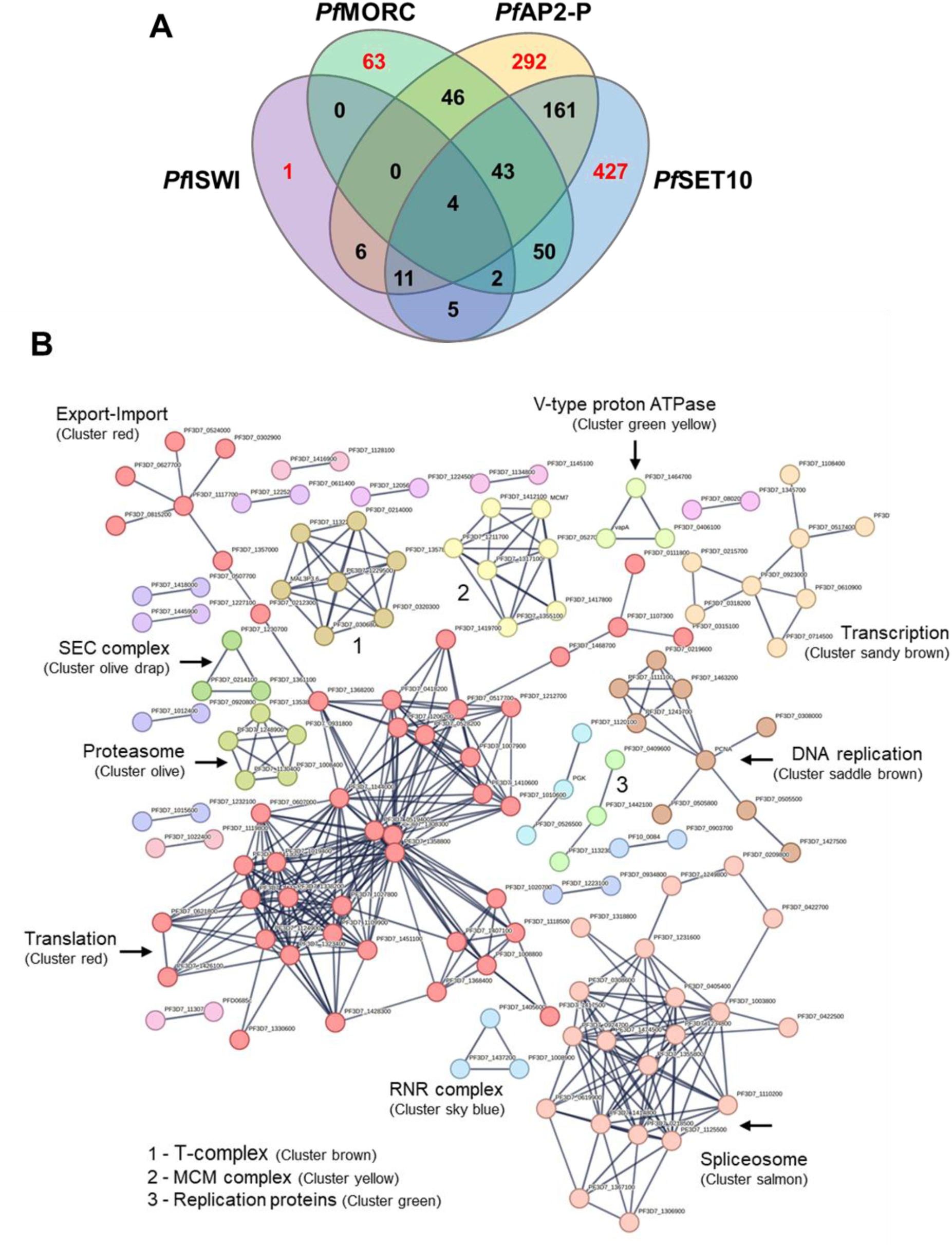
*Pf*SET10 forms a comprehensive interaction network with *Pf*AP2-P, *Pf*ISWI, and *Pf*MORC. (**A**) Venn diagram depicting interactors shared by *Pf*AP2-P, *Pf*ISWI, *Pf*MORC and *Pf*SET10. The list of interactors was extracted from (Bryant et al., 2020; Chahine et al., 2023; Subudhi et al., 2023; Singh et al., 2024). (**B**) Network analysis of the shared interactors. A protein-protein network analysis of the 328 interactors was performed (STRING program, highest interaction confidence of 0.9). Clustering of the interactors was employed by the K-means algorithm with a maximal cluster number of 25; disconnected nodes were excluded. Clusters ≥ 3 proteins were considered.

The 328 interactors were subjected to STRING analysis (Fig. 6B; K means; confidence level of 0.9; cluster number of 25). One megacluster was identified that was assigned to translation and which exhibited a nuclear export-import branch (cluster red). A second extensive cluster belonged to the spliceosome (cluster salmon). Three further clusters could be associated with DNA replication, replication factors and DNA replication licensing factors (MCM proteins). Addition clusters included a transcription cluster, a proteasome cluster, a V-type proton ATPase cluster, a SEC protein transport cluster, a RNR cluster and a T-complex cluster (Fig. 7B; Table S7).

The combined interactor analyses pinpoints *Pf*SET10 as a component of an overarching protein-protein network involved in chromatin modulation during replication and transcription.

## Discussion

*Pf*SET10, one of ten SET proteins encoded in the *P. falciparum* genome, has originally been designated a H3K4 methyltransferase crucial for maintaining the active *var* gene in a poised state during cellular division (Volz et al., 2012). In a later study, the ability to knockout the *pfset10* gene without altering *var* gene expression questioned the essentiality of *Pf*SET10 in *var* gene regulation (Ngwa et al., 2021). Here, we show that *Pf*SET10 is a H3K18me1 methyltransferase without detectable H3K4 methylation activity. In accord with previous reports (Saraf et al., 2016; Coetzee et al., 2017), this methylation mark is particularly present in the nuclei of schizonts, but vanishes in gametocytes upon maturation. We further demonstrate that the lack of *Pf*SET10 completely abolishes the H3K18me1 mark in blood stage parasites. *Pf*SET10 deficiency and hence the absence of H3K18me1 methylation in schizonts leads to the transcriptional upregulation of *rif* and *var* genes as well as of genes encoding exported proteins and RBC invasion-related proteins, suggesting that *Pf*SET10 is involved in gene repression during schizont-to-merozoite transition. While the H3K18me1 methylation mark to date is not well understood, it was found on the gene bodies of repressed genes of *Theileria* macroschizonts (Cheeseman et al., 2021). In accord with our findings, peak H3K18me1 levels were observed in schizonts, whereas the methylation mark decreased during the differentiation of *T. annulata* from schizonts to merozoites, and chemical inhibition of the H3K18me1 methylation activity affected genes associated to macroschizont-to-merozoite transition.

A previous quantitative chromatin proteomics analysis revealed a highly dynamic histone PTM landscape in the blood stages of *P. falciparum* with euchromatic histone PTMs being abundant during schizogony and late gametocytes and heterochromatic PTMs marking early gametocytes (Coetzee et al., 2017). The histone prevalence increased significantly from early asexual development to schizonts in accord with higher histone gene expression, transcriptional activity and DNA synthesis during schizogony (e.g., Miao et al., 2006; Otto et al., 2010). The same study discovered the H3K18me1 methylation mark and assigned it to the late stages of intraerythrocytic development (Coetzee et al., 2017). Later investigations found H3K18me1 in the centromeric regions of chromosomes, in particular associated with internal *var* gene clusters (Reers et al., 2023). Because H3K18me1 was enriched in coding sequences and did not correlate with known activating marks, a function of this mark in chromosome organization during schizogony rather than a role in transcriptional regulation was suggested (Reers et al., 2023).

We applied loss-of-function studies, using a *Pf*SET10-KO and a *glmS* ribozyme-based *Pf*SET10-HA-KD line. The complete loss of *Pf*SET10 resulted in reduced intraerythrocytic growth (Ngwa et al., 2021 and this study), while the partial reduction of *Pf*SET10 levels had no detectable effect on parasitemia. Similar results were obtained during loss-of-function approaches of the related chromatin-modifying protein *Pf*MORC (see detailed discussion of *Pf*MORC below). While partial *Pf*MORC knockdown using the *glmS*-ribozyme system had no significant effect on parasite survival (Singh et al., 2021), transposon mutagenesis identified *Pf*MORC as likely essential (Zhang *et al*. 2018) and a TetR-DOZI knockdown system resulted in reduced intraerythrocytic growth and death of the parasite (Chahine et al., 2023). We thus conclude that *Pf*SET10 has a crucial function for blood stage survival in a dose-dependent manner.

To unveil the *Pf*SET10 interaction network, we performed TurboID analyses and evaluated the hits via STRING analyses according to known protein-protein interaction patterns. We show multitude interactions of *Pf*SET10 with proteins involved in gene regulation, transcriptional control and DNA replication and repair, confirming the proposed function of *Pf*SET10 in chromatin modulation. Prominent interactors include, in addition to polymerases and helicases, epigenetic regulators like SET and bromodomain proteins and histone acyltransferases and deacetylases, as well as MED and MCM proteins and chromatin remodeling proteins like *Pf*ISWI, *Pf*SWIP and *Pf*SNF2L. In addition, *Pf*SET10 binds to 15 out of the 28 known ApiAP2 transcription factors (Painter et al., 2011).

Large-scale protein interaction networks in *Plasmodium* schizonts were previously reported, with more than 20,000 putative protein interactions, organized into 600 protein clusters (Hillier et al., 2019). Among others, chromatin regulator networks were identified that included the interaction of *Pf*MORC and an EELM2 domain-containing protein (PF3D7_1141800) with *Pf*AP2-P and the histone deacetylase HDAC1. The authors hypothesized that *Pf*MORC and EELM2 proteins form a scaffold connecting transcription factor complexes with epigenetic regulators to effect nucleosome reorganization and regulate gene expression (Hillier et al., 2019). Noteworthy, *Pf*AP2-P has been shown to bind to promoters of genes controlling trophozoite development, host cell remodeling, and antigenic variation and to also target a genetic element in the *var* gene introns that is important for tethering members of this multigene family to the nuclear periphery as part of their epigenetic silencing (Zhang et al., 2011; Subudhi et al., 2023). *P. falciparum* blood stages lacking PfAP2-P exhibited de-repression of most *var* genes (Subudhi et al., 2023), and similar effects were observed by us for *Pf*SET10-KO parasites, indicating common functions of the two interactors in repressing genes assigned to antigenic variation and pathogenicity. Worth mentioning in this context, while many malarial ApiAP2 proteins act as transcription factors, some of them bind heterochromatin independent of DNA motif recognition, where they act as activators or repressors of heterochromatic gene expression (Shang et al., 2022).

One of our top hits of *Pf*SET10 interactors is *Pf*MORC. MORC proteins belong to a highly conserved nuclear protein superfamily that is involved in signaling-dependent chromatin remodeling and epigenetic regulation (reviewed in, e.g., Lorković 2012; Li et al., 2013). In plants, MORC proteins function in gene repression and heterochromatin compaction (Koch *et al*. 2017, Zhong *et al*. 2023). They can also form protein complexes with immune-responsive proteins, SWI chromatin remodeling proteins, and histone deacetylases (Iyer et al., 2008; Kang et al., 2012; Moissiard et al., 2012; Bordiya et al., 2016; Kim et al., 2019). The *Toxoplasma gondii* orthologue *Tg*MORC was identified in a complex with HDAC1 and ApiAP2 proteins to support the epigenetic rewiring of sexual gene transcription (Farhat et al., 2020, Srivastava et al., 2023, Antunes et al., 2024). In *P. falciparum*, *Pf*MORC has been shown to associate with several ApiAP2 proteins, with which they share DNA binding sites (Hillier et al., 2019; Bryant et al., 2020; Singh et al., 2021, 2024). In addition, *Pf*MORC locates together with *Pf*ISWI at *var* gene promoter regions, suggesting a role of the two proteins in *var* gene silencing (Bryant et al., 2020). Recently, two independent interactomics analyses on *Pf*MORC have been published (Chahine et al., 2023; Singh et al., 2024). Among the reported interaction partners of *Pf*MORC were ApiAP2 transcription factors, EELM2 proteins, *Pf*CHD1, the RNA helicase *Pf*DBP5, the epigenetic regulators *Pf*BDP3, *Pf*HDAC1, *Pf*SET3 and *Pf*PHD2, as well as *Pf*ISWI (Chahine et al., 2023; Singh et al., 2024), all of which were also identified by us as *Pf*SET10 interactors.

Chromatin immunoprecipitation followed by deep sequencing has revealed that *Pf*MORC localizes to subtelomeric regions on all chromosomes across the *P. falciparum* genome, with additional occupancy at internal heterochromatic islands (Chahine et al., 2023; Singh et al., 2024). Within the subtelomeric regions of chromosomes, *Pf*MORC was bound upstream and within the gene bodies of multigene families, including *var, rif,* and *stevor* genes as well as genes encoding exported proteins. The majority of reads mapped to antigenic genes in the trophozoite and schizont stages. In addition, *Pf*MORC colocalized with the depletion of H3K36me2, which is demarcated by the SET2 methyltransferase (Jiang et al., 2013); a potential colocalization with H3K18me1 has not been investigated, though (Singh et al., 2024). Phenotypical analyses by two independent loss-of-function approaches unveiled a significant upregulation of genes upon *Pf*MORC deficiency that are related to RBC invasion and host cell remodeling as well as to *var* genes (Chahine et al., 2023; Singh et al., 2024), These observations are comparable to our reports on the transcriptional changes in the *Pf*SET10-KO line, suggesting roles of both proteins in repressing genes linked to schizont-to-merozoite transition. Importantly, parasites lacking *Pf*MORC failed to maintain their overall chromatin structure with a significant weakening of the tightly controlled heterochromatin cluster (Chahine et al., 2023).

We also performed a meta-analysis on the interactors of *Pf*SET10 and of three of its own interactors, i.e. *Pf*MORC, *Pf*AP2-P, and *Pf*ISWI. We demonstrated an extensive network with 328 shared binding partners, forming clusters particularly assigned to DNA replication and transcription including RNA processing via the spliceosome. Further, proteins involved in nuclear export-import processes as well as in proteostasis, like proteasomal proteins and T-complex chaperonin subunits were found. Noteworthy, a weakness of the STRING analysis is the missing information on the interactions between the chromatin proteins identified here, like *Pf*MORC, *Pf*AP2-P, EELM2, *Pf*CHD1, *Pf*BDP3, *Pf*HDAC1, *Pf*SET3, *Pf*SET10, *Pf*PHD2, and *Pf*ISWI, which were therefore not correctly mapped.

Our data provide a first glimpse on the impact of *Pf*SET10 on regulating the nucleic acid metabolism during intraerythrocytic replication and unveiled an extensive chromatin modulation network in the blood stage nuclei. The combined data on *Pf*SET10 and its top interacting proteins led us propose two related critical functions of this network, i.e. heterochromatin organization and gene repression during schizont-to-merozoite transition. Noteworthy, SET proteins present promising drug targets and HMT inhibitors are currently in clinical trials to test their use in cancer therapy. The most advanced clinical trials include tazemetostat, an Enhancer of zeste homolog 2 (EZH2) inhibitor approved for follicular lymphoma (Marsh and Jimeno, 2020; Marzochi et al., 2023). In *Plasmodium*, HMT inhibitors exhibit rapid antimalarial activities *in vitro* and *in vivo* (e.g., Malmquist et al., 2012, 2015; Sundriyal et al., 2014; Ngwa et al., 2019; Coetzee et al, 2020) and even a reversion of epigenetically acquired drug resistance by the HMT inhibitor chaetocin has been reported (Chan et al., 2020). Hence, information on proteins crucial during histone methylation in the *Plasmodium* blood and sexual stages may advance the current search for antiepigenetic targets in malaria therapy.

## Materials and Methods

### Gene identifiers

The following PlasmoDB gene identifiers (gene IDs) are assigned to the genes and proteins investigated in this study: *Pf*SET10 [PF3D7_1221000]; *Pf*39 [PF3D7_1108600]; *Pf*92 [PF3D7_1364100]; *Pf*s230 [PF3D7_0209000]; STEVOR proteins [PF3D7_0300400; PF3D7_0324600; PF3D7_0631900; PF3D7_1254100]; histone H3 [PF3D7_0610400].

### Antibodies

The following antibodies were used: mouse and rabbit polyclonal anti-*Pf*39 antisera (Simon et al., 2009); rabbit polyclonal anti-*Pf*92 antisera (Musabyimana et al., 2022); mouse and rabbit polyclonal anti-*Pf*s230 antisera (Williamson et al., 1995); rabbit polyclonal anti-STEVOR antisera (Wichers et al., 2019); mouse monoclonal anti-GFP antibody (Roche; Basel, CH); rat monoclonal anti-HA antibody (Roche);); rabbit monoclonal anti-H3K18me1 antibody (Abclonal, Dusseldorf, DE); rabbit polyclonal anti-H3K4me1, anti-H3K4me2, anti-H3K4me3, anti-H3K36me1, anti-H3K36me2, anti-H3K36me3 antibody, and anti-H3 antibody (Abclonal, Dusseldorf, DE).

The following dilutions were used for (a) immunolabeling: mouse or rabbit anti-*Pf*s230 (1:200), rabbit anti-*Pf*92 (1:200), rabbit anti-*Pf*39 (1:200) mouse anti-GFP (1:200), rat anti-HA (1:50), rabbit anti-H3K18me1 (1:500); (b) immunoblotting: rabbit anti-*Pf*39 (1:10,000), mouse anti-GFP (1:1,000), rat anti-HA (1:500), rabbit anti-H3K4me3 (1:4,000), rabbit anti-H3K18me1 (1:4,000), rabbit anti-H3K4me1 (1:4,000), rabbit anti-H3K4me2 (1:4,000), rabbit anti-H3K4me3 (1:4,000), rabbit anti-H3K36me1 (1:4,000), rabbit anti-H3K36me2 (1:4,000), rabbit anti-H3K36me3 (1:4,000), rabbit anti-H3 (1:4,000), rabbit anti-STEVOR (1:2,000).

### Bioinformatics

Predictions of gene expression and protein properties and function were made using the database PlasmoDB (http://plasmoDB.org; Aurrecoechea et al. 2009); the peak transcript expression of candidate genes was analyzed using table “Transcriptomes of 7 sexual and asexual life stages” (López-Barragán et al. 2011) and sex specificity was predicted using table “Gametocyte transcriptomes” (Lasonder et al. 2016) of the PlasmoDB database. The gene ontology enrichment (GO) analysis was performed using the ShinyGO 0.77 (Ge et al., 2020) with a p-value cut-off of 0.05. The network analyses were conducted using the STRING database (version 11.0) (Szklarczyk et al. 2019) with parameters as indicated (see Tables S5-S7). For the meta-analysis of shared interactors, hits as provided by (Bryant et al., 2020, Table EV11), (Chahine et al. 2023, Table S3a), (Singh et al., 2024; Table S2), (Subudhi et al., 2023, Table S6) were compared to the *Pf*SET10 curated nuclear interactors (Table S7).

### Parasite culture

The gametocyte-producing strain *P. falciparum* NF54 (WT NF54) was cultured *in vitro* in RPMI1640/HEPES medium (Gibco; Thermo Fisher Scientific; Waltham, US) supplemented with 10% (v/v) heat inactivated human A+ serum, 50 μg/ml hypoxanthine (Sigma-Aldrich; Taufkirchen, DE) and 10 μg/ml gentamicin (Gibco). For cultivation of the transgenic lines, the selection drug WR99210 (Jacobus Pharmaceutical Company; Princeton, US) was added at a final concentration of 4.0 nM. All cultures were kept at 37°C in an atmosphere of 5% O_2_ and 5% CO_2_ in N_2_. Human serum and erythrocyte concentrate were obtained from the Department of Transfusion Medicine, University Hospital Aachen, Germany. Donor sera and blood samples were pooled and anonymized. The work with human blood was approved by the Ethics Committee of the RWTH University Hospital (EK 007/13). To synchronize the asexual parasite blood stages, parasite cultures with 4% ring stages were centrifuged, the pellet was resuspended in 5× pellet’s volume of 5% (w/v) sorbitol (AppliChem, Darmstadt, DE)/ddH2O and incubated for 10 min at room temperature (RT). Cells were washed once with RPMI and cultivated as described. Schizonts and gametocytes were enriched via Percoll gradient centrifugation (GE Healthcare Life Sciences, Chicago, IL, US) as previously described (Karuiki et al., 1998; Farrukh et al., 2024). Gametocytogenesis was then induced by the addition of lysed RBCs (0.5 ml of 50% haematocrit lysed RBC in 15 ml of culture medium) followed by washing with RPMI the next day. To kill the asexual blood stages in gametocyte cultures, these were maintained in cell culture medium supplemented with 50 mM N-acetylglucosamine (GlcNAc; Carl Roth, Karlsruhe, DE) for 5 consecutive days (Fivelman et al., 2007).

### Generation of the *Pf*SET10-HA-KD parasite line

The *Pf*SET10-HA-KD line was generated by single crossover homologous recombination, using the pSLI-HA-*glmS* vector (Musabyimana et al., 2022). A 906-bp gene fragment homologous to the 3′-coding region of the *pfset10* gene was amplified using forward primer *Pf*SET10 pSLI-HA-*glmS* SacII and reverse primer *Pf*SET10 pSLI-HA-*glmS* XhoI. The stop codon was excluded from the homologous gene fragment. The insert was ligated to the vector backbone via SacII and XhoI restriction sites. A WT NF54 culture with at least 5% ring stages was transfected with 100 μg plasmid DNA in transfection buffer by electroporation (310 V, 950 μF, 12 ms; Bio-Rad gene-pulser Xcell) as described (e.g., Wirth et al., 2014; Ngwa et al., 2017). At 6 hours post-transfection, WR99210 was introduced at a final concentration of 4 nM. After 3-5 weeks, WR99210-resistant parasites emerged in the cultures. WT NF54 parasites were eliminated from the culture by addition of neomycin (G418; Sigma-Aldrich) at a final concentration of 550 μg/ml for a maximum of 7 days. Successful integration into the gene locus was confirmed by diagnostic PCR using primers 5’ Int *Pf*SET10 pSLI-HA-*glmS* (1), 3’ Int *Pf*SET10 pSLI-HA-*glmS* (2), pSLI-HA-*glmS* FP (3) and pSLI-HA-*glmS* RP (4) (Fig. S1A, B; for primer sequences, see Table S8).

### Generation of the *Pf*SET10-TurboID-GFP line

The *Pf*SET10-TurboID-GFP line was generated by single crossover homologous recombination, using the pSLI-TurboID-GFP vector (Sassmannshausen et al., 2024). A 1,144-bp gene fragment homologous to the 3′-coding region of the *pfset10* gene was amplified using forward primer *Pf*SET10 pSLI-TurboID-GFP NotI FP and reverse primer *Pf*SET10 pSLI-TurboID-GFP SpeI RP. The stop codon was excluded from the full-length gene sequence. The fragment was ligated to the vector via NotI and SpeI restriction sites. Transfection and selection were carried out as described above. Successful integration into the gene locus was confirmed by diagnostic PCR using primers 5’ Int *Pf*SET10 pSLI-TurboID-GFP (1), 3’ Int *Pf*SET10 pSLI-TurboID-GFP (2), pSLI-HA-*glmS* FP (3) and pSLI-TurboID-GFP (4) (Fig. S6A, B; for primer sequences, see Table S8).

### Indirect Immunofluorescence Assay

Parasite cultures containing mixed asexual blood stages and gametocytes of WT NF54 as well as of lines *Pf*SET10-HA-*glmS* and *Pf*SET10-TurboID-GFP were coated on glass slides as cell monolayers and then air-dried. After fixation in methanol at −80 °C for 10 min, the cells were serially incubated in 0.01% (w/v) saponin/0.5% (w/v) BSA/PBS and 1% (v/v) neutral goat serum (Sigma-Aldrich)/PBS for 30 min at RT to facilitate membrane permeabilization and blocking of non-specific binding. The primary antibodies were diluted in 3% (w/v) BSA/PBS and were added to the slide for 2 h incubation at 37°C. The slides were washed 3x with PBS and incubated with the secondary antibody for 1 h at 37°C. Following 2x washing with PBS, the incubation with the second primary antibody and the corresponding visualization with the second secondary antibody were carried out as described above. The nuclei were stained with Hoechst 33342 staining solution for 2 min at RT (1:5,000 in 1x PBS). The cells were mounted with anti-fading solution (Citifluor Limited; London, UK), covered with a coverslip and sealed airtight with nail polish. The parasites were visualized by conventional fluorescence microscopy using a Leica DM5500 B (Leica; Wetzlar, DE) microscope. The following secondary antibodies were used: Anti-mouse Alexa Fluor 488, anti-rabbit Alexa Fluor 488, anti-rat Alexa Fluor 488, anti-mouse Alexa Fluor 594, anti-rabbit Alexa Fluor 594, (1:1,000; all fluorophores from Invitrogen Molecular Probes; Eugene, US, or Sigma-Aldrich); further Alexa Fluor 594 streptavidin (1:500; Invitrogen Molecular Probes) was used. Alternatively, the asexual blood stages were stained with 0.01% (w/v) Evans Blue (Sigma-Aldrich)/PBS for 3 min at RT followed by 5 min washing with PBS. Images were processed using the Adobe Photoshop CS software.

## Western Blotting

Asexual blood stage parasites of WT NF54 as well as lines *Pf*SET10-HA-*glmS* and *Pf*SET10-TurboID-GFP were harvested from mixed or synchronized cultures, while gametocytes were enriched by Percoll purification. The infected RBCs were lysed with 0.05% (w/v) saponin/PBS for 10 min at 4°C to release the parasites, then washed with PBS, and resuspended in lysis buffer (0.5% (v/v) Triton X-100, 4% (w/v) SDS in PBS) supplemented with protease inhibitor cocktail. After adding 5x SDS-PAGE loading buffer containing 25 mM DTT, the lysates were heat-denatured for 10 min at 95°C. Lysates were separated via SDS-PAGE and transferred to Hybond ECL nitrocellulose membranes (Amersham Biosciences; Buckinhamshire, UK) following the manufacturer’s protocol. Non-specific binding sites were blocked by incubation with 5% (w/v) skim milk and 1% (w/v) BSA in Tris buffer (pH 7.5) for 1 h at 4°C. For immunodetection, membranes were incubated overnight at 4°C with the primary antibody in 3% (w/v) skim milk/TBS. The membranes were washed 3x each with 3% (w/v) skim milk/TBS and 3% (w/v) skim milk/0.1 % (v/v) Tween/TBS and then incubated for 1 h at RT with goat anti-mouse, anti-rabbit or anti-rat alkaline phosphatase-conjugated secondary antibodies (dilution 1:10,000, Sigma-Aldrich) in 3% (w/v) skim milk/TBS. The membranes were developed in a NBT/BCIP solution (nitroblue tetrazolium chloride/5-bromo-4-chloro-3-indoxyl phosphate; Roche) for up to 20 min at RT. For the detection of biotinylated proteins, the blocking step was performed overnight at 4°C in 5% (w/v) skim milk/TBS and the membrane was washed 5x with 1x TBS before incubation with streptavidin-conjugated alkaline phosphatase (dilution 1:1,000, Sigma-Aldrich) in 5% (w/v) BSA/TBS for 1 h at RT. Blots were scanned and processed using the Adobe Photoshop CS software. Band intensities were measured using the ImageJ program version 1.51f.

### Asexual blood stage replication assay

#### *Pf*SET10-KO

The asexual blood stage replication assay was performed as described (Ngwa et al., 2021). In short, tightly synchronized asexual blood stage cultures of line *Pf*SET10-KO and WT NF54 were set to an initial parasitemia of 0.25% at the ring stage and cultivated as described. Giemsa-stained thin blood smears were prepared every 12 hours over an 84-hour period. The parasitemia at each time point was determined microscopically at 1,000x magnification by counting the percentage of parasites in 1,000 RBCs. Two experiments were performed, each in triplicate.

#### *Pf*SET10-HA-KD

Asexual blood stage cultures of line *Pf*SET10-HA-*glmS* and WT NF54 were tightly synchronized and set to an initial parasitemia of 0.25% ring stages. The cultures were then continuously treated with 2.5 mM GlcN for transcript knockdown. Untreated cultures were used for control. The parasitemia was assessed every 12 hours during a period of 96 hours employing the LUNA-FX7™ automated cell counter. Two experiments were performed, each in triplicate. Data analysis was performed using MS Excel 2016 and GraphPad Prism 5.

### Gametocyte commitment and development assays

#### *Pf*SET10-KO

The gametocyte development assay was performed as described (Hanhsen et al., 2022). In short, tightly synchronized cultures of line *Pf*SET10-KO and WT NF54 were set to a parasitemia of 5% ring stages. Gametocytogenesis was then induced by the addition of lysed RBCs followed by washing with RPMI the next day. The cultures were then maintained in cell culture medium supplemented with 50 mM GlcNAc to kill the asexual blood stages for 5 days (Fivelman et al., 2007) and then maintained with normal cell culture medium until day 10 post-induction. Giemsa-stained blood smears were prepared every 48 hours between day 5 and day 13 post-induction. The gametocytemia at each time point was determined microscopically by counting the percentage of gametocytes in 1,000 RBCs. Two experiments were performed, each in triplicate. Data were analyzed using MS Excel 2016 and GraphPad Prism 5.

#### *Pf*SET10-HA-KD

The gametocyte commitment and development assay was performed as described (Flammersfeld et al., 2020). In short, tightly synchronized cultures of line *Pf*SET10-HA-KD and WT NF54 were each divided into three groups (untreated, pre-induction and post-induction group) and set to an initial parasitemia of 0.5% schizonts. The pre-induction group was cultured in medium supplemented with 2.5 mM GlcN (Sigma-Aldrich) for 72 hours. Then, all cultures were adjusted to a parasitemia of 5.3% and gametocytogenesis was induced by the addition of lysed RBC for 24 hours. Afterwards, cells were washed and the parasites were cultivated in medium for 4 days. Heparin was added to the medium at a final concentration of 20 U/ml to kill the asexual blood stages. The post-induction group was treated with 2.5 mM GlcN for 7 consecutive days after addition of lysed RBC for 24 hours. Giemsa-stained thin blood smears were prepared on days 3 and 7 post-induction and gametocytemia was determined in 1,000 RBCs. Three experiments were performed, each in triplicate. Data were analyzed using MS Excel 2016 and GraphPad Prism 5.

### RNA isolation and sequencing

*Pf*SET10-KO and WT NF54 schizonts were enriched by Percoll gradient centrifugation (GE Healthcare Life Sciences, Chicago, IL, USA) as described above. Purified samples were stored at −80°C in TRIzol LS reagent (Life Technologies, Thermo Fisher Scientific, Waltham, USA) and RNA was isolated according to the manufacturer’s protocol. RNA preparations were treated with RNAse-free DNAse I (Qiagen, Hilden, DE) to remove gDNA contamination, followed by phenol/chloroform extraction and ethanol precipitation. The quality of the RNA was assessed by ND-1000 (NanoDrop Technologies) and agarose gel electrophoresis. RNA purity was confirmed by a A260/A280 ratio above 2.1. Purified RNA underwent mRNA sequencing on the Illumina NextSeq sequencer (Genomic Facility of the Interdisciplinary Center for Clinical Research, IZKF) using standard paired-end sequencing protocols provided by Illumina. The data underwent analysis using the NextGen pipeline, an in-house adapted pipeline integrated into the workflow management system of the QuickNGS-Environment, as described (Hanhsen et al., 2022). Significantly deregulated genes were defined as ≥ 2-fold deregulated in their transcript levels log2-fold change ≥ 1).

### Preparation of samples for TurboID analysis

Highly synchronized schizont cultures of line *Pf*SET10-TurboID-GFP and WT NF54 as control were treated with biotin at a final concentration of 50 µM for 24 h to induce the biotinylation of proximal proteins by the BirA ligase. After treatment, the RBCs were lysed with 0.05% (w/v) saponin and the parasites were resuspended in 100 µl binding buffer (Tris buffer containing 1% (v/v) Triton X-100 and protease inhibitor). The sample was sonicated on ice (2 x 60 pulses at 30% duty cycle) and another 100 µl of ice-cold binding buffer was added. After a second session of sonification, cell debris was pelleted by centrifugation (5 min, 16,000x *g*, 4°C). The supernatant was mixed with pre-equilibrated Cytiva Streptavidin Mag Sepharose Magnet-Beads (Cytiva; Washington DC, US) in a low-binding reaction tube. Incubation was performed with slow end-over-end mixing over night at 4°C. The beads were washed 6x with 500 μl washing buffer (3x: RIPA buffer containing 0.03% (w/v) SDS, followed by 3x 25 mM Tris buffer (pH 7.5)) and were resuspended in 45 μl elution buffer (1% (w/v) SDS/5 mM biotin in Tris buffer (pH 7.5)), followed by an incubation for 5 min at 95°C. The supernatant was transferred into a new reaction tube and stored at −20°C. For each culture, three independent samples were collected.

### Proteolytic digestion

Samples were processed by single-pot solid-phase-enhanced sample preparation (SP3) as described before (Hughes et al. 2014; Sielaff et al. 2017). In brief, proteins bound to the streptavidin beads were released by incubating the samples for 5 min at 95° in an SDS-containing buffer (1% (w/v) SDS, 5 mM biotin in water/Tris buffer, pH 8.0). After elution, proteins were reduced and alkylated, using DTT and iodoacetamide (IAA), respectively. Afterwards, 2 µl of carboxylate-modified paramagnetic beads (Sera-Mag Speed Beads, GE Healthcare; Chicago, US; 0.5 μg solids/μl in water as described by Hughes et al. 2014) were added to the samples. After adding acetonitrile to a final concentration of 70% (v/v), samples were allowed to settle at RT for 20 min. Subsequently, beads were washed twice with 70% (v/v) ethanol in water and once with acetonitrile. The beads were resuspended in 50 mM NH_4_HCO_3_ supplemented with trypsin (Mass Spectrometry Grade, Promega; Madison, US) at an enzyme-to-protein ratio of 1:25 (w/w) and incubated overnight at 37°C. After overnight digestion, acetonitrile was added to the samples to reach a final concentration of 95% (v/v) followed by incubation at RT for 20 min. To increase the yield, supernatants derived from this initial peptide-binding step were additionally subjected to the SP3 peptide purification procedure (Sielaff et al. 2017). Each sample was washed with acetonitrile. To recover bound peptides, paramagnetic beads from the original sample and corresponding supernatants were pooled in 2% (v/v) dimethyl sulfoxide (DMSO) in water and sonicated for 1 min. After 2 min of centrifugation at 14,000xg and 4°C, supernatants containing tryptic peptides were transferred into a glass vial for MS analysis and acidified with 0.1% (v/v) formic acid.

### Liquid chromatography-mass spectrometry (LC–MS) analysis

Tryptic peptides were separated using an Ultimate 3000 RSLCnano LC system (Thermo Fisher Scientific;) equipped with a PEPMAP100 C18 5 µm 0.3 x 5 mm trap (Thermo Fisher Scientific) and an HSS-T3 C18 1.8 μm, 75 μm x 250 mm analytical reversed-phase column (Waters Corporation; Milford, US). Mobile phase A was water containing 0.1% (v/v) formic acid and 3% (v/v) DMSO. Peptides were separated by running a gradient of 2–35% mobile phase B (0.1% (v/v) formic acid, 3% (v/v) DMSO in ACN) over 40 min at a flow rate of 300 nl/min. Total analysis time was 60 min including wash and column re-equilibration steps. Column temperature was set to 55°C. Mass spectrometric analysis of eluting peptides was conducted on an Orbitrap Exploris 480 (Thermo Fisher Scientific) instrument platform. Spray voltage was set to 1.8 kV, the funnel RF level to 40, and heated capillary temperature was at 275°C. Data were acquired in data-dependent acquisition (DDA) mode targeting the 10 most abundant peptides for fragmentation (Top10). Full MS resolution was set to 120,000 at *m/z* 200 and full MS automated gain control (AGC) target to 300% with a maximum injection time of 50 ms. Mass range was set to *m/z* 350–1,500. For MS2 scans, the collection of isolated peptide precursors was limited by an ion target of 1 × 10^5^ (AGC tar-get value of 100%) and maximum injection times of 25 ms. Fragment ion spectra were acquired at a resolution of 15,000 at *m/z* 200. Intensity threshold was kept at 1E4. Isolation window width of the quadrupole was set to 1.6 *m/z* and normalized collision energy was fixed at 30%. All data were acquired in profile mode using positive polarity. Each sample was analyzed in three technical replicates.

### Data analysis and label-free quantification

DDA raw data acquired with the Exploris 480 were processed with MaxQuant (version 2.0.1; Cox & Mann 2008; Cox et al. 2014), using the standard settings and label-free quantification (LFQ) enabled for each parameter group, i.e. control and affinity-purified samples (LFQ min ratio count 2, stabilize large LFQ ratios disabled, match-between-runs). Data were searched against the forward and reverse sequences of the *P. falciparum* proteome (UniProtKB/TrEMBL, 5,445 entries, UP000001450, released April 2020) and a list of common contaminants. For peptide identification, trypsin was set as protease allowing two missed cleavages. Carbamidomethylation was set as fixed and oxidation of methionine as well as acetylation of protein N-termini as variable modifications. Only peptides with a minimum length of 7 amino acids were considered. Peptide and protein false discovery rates (FDR) were set to 1%. In addition, proteins had to be identified by at least two peptides. Statistical analysis of the data was conducted using Student’s t-test, which was corrected by the Benjamini-Hochberg (BH) method for multiple hypothesis testing (FDR of 0.01). In addition, proteins in the affinity-enriched samples had to be identified in all three biological replicates and show at least a two-fold enrichment compared to the controls. The datasets of protein hits were further edited by verification of the gene IDs and gene names via the PlasmoDB database (www.plasmodb.org; Aurrecoechea et al. 2009). PlasmoDB gene IDs were extracted from the fasta headers provided by mass spectrometry and verified manually. Following the generation of an initial list of significantly enriched proteins, those with a putative signal peptide were excluded. For STRING-based analyses, only proteins with predicted nuclear localization (according to PlasmoDB) were considered. In a further curation step, known ribosomal and proteasomal proteins were excluded.

## Data availability

The mass spectrometry proteomics data have been deposited to the ProteomeXchange Consortium (http://proteomecentral.proteomexchange.org) via the jPOST partner repository (Vizcaino et al., 2013) with the dataset identifiers PXD052931 (ProteomeXchange) and JPST003166 (jPOST).

## Supplemental Figure Legends

**Figure S1.**
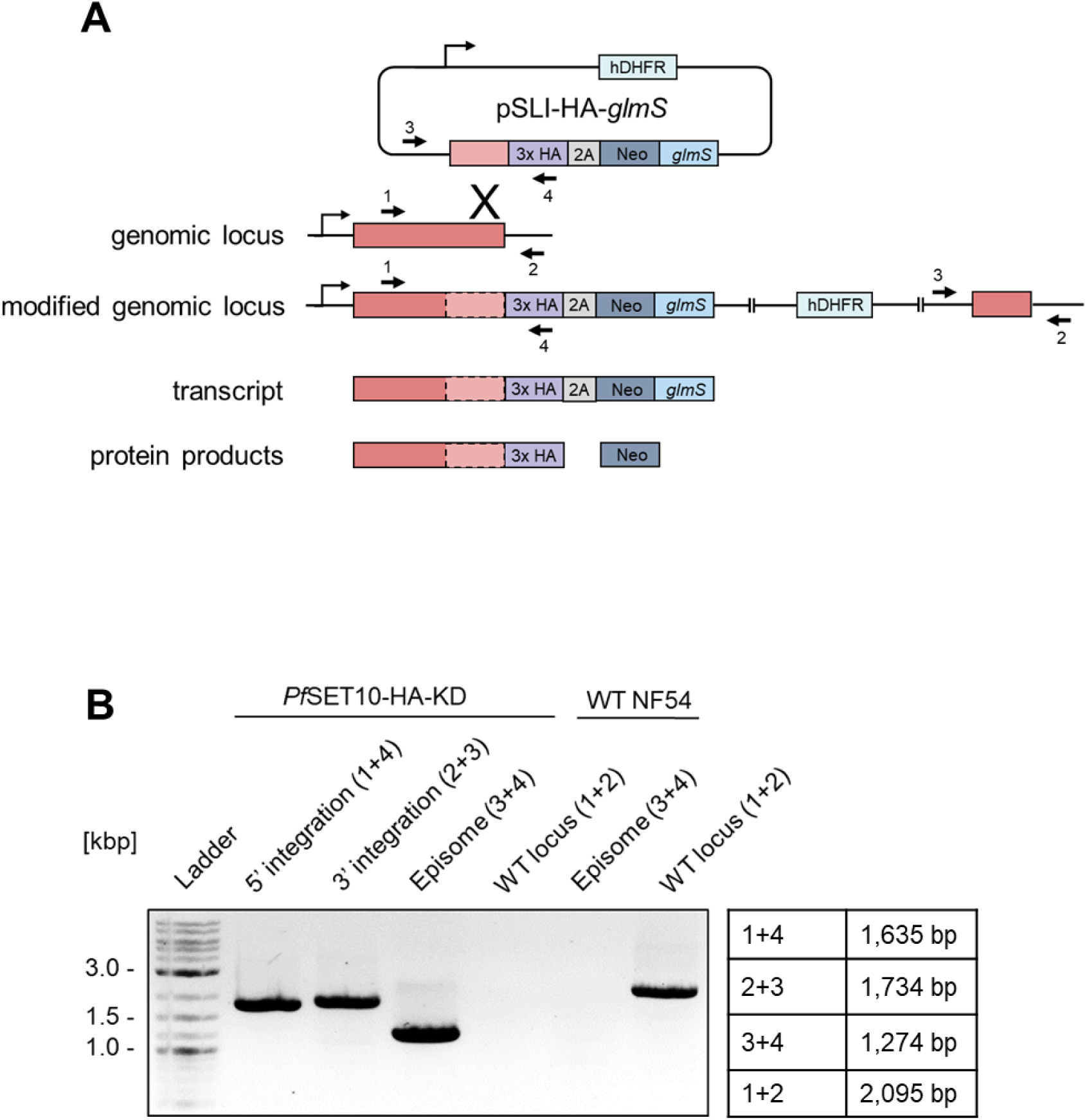
Generation of the *Pf*SET10-HA-KD line. **(A)** Schematic depicting the single-crossover homologous recombination strategy for the generation of the pSLI-HA-*glmS*-based *Pf*SET10-HA-KD line. The coding region of the gene of interest was fused at the 3’-end with a HA-encoding sequence followed by the 2A-skip peptide sequence and the Neo and *glmS*-ribozyme sequences. The numbered arrows indicate the positions of primers used to confirm vector integration. *glmS*, glucosamine-6-phosphate-activated ribozyme; HA, hemagglutinin; hDHFR, human dihydrofolate reductase-encoding gene conferring resistance to WR99210; Neo, gene conferring resistance to neomycin. **(B)** Confirmation of vector integration into the *pfset10* gene locus. Diagnostic PCR demonstrates successful 5′ (primers 1 and 4) and 3′ (primers 3 and 2) integration in line *Pf*SET10-HA-KD. WT NF54 gDNA served as control, demonstrating the original gene locus (primers 1 and 2). Episomal DNA was further detected (primers 3 and 4). Band sizes are indicated.

**Figure S2.**
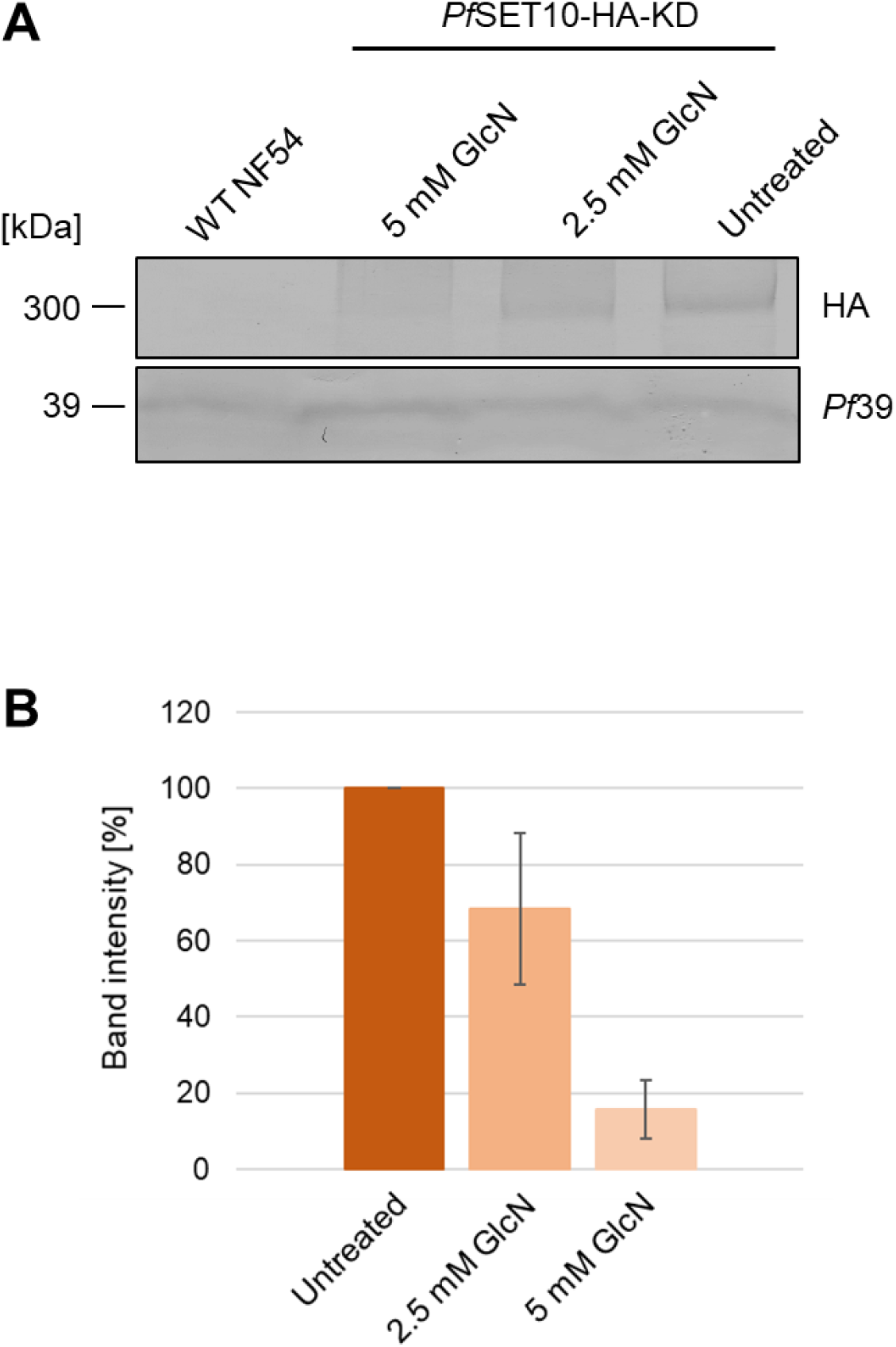
Verification of *Pf*SET10-HA knockdown. **(A)** Confirmation of *Pf*SET10-HA knockdown. Lysates of mixed asexual blood stages of line *Pf*SET10-HA-KD treated with 2.5 and 5 mM GlcN for 72 h were subjected to Western blotting using rat anti-HA antibody to detect *Pf*SET10-HA (~275 kDa). Untreated *Pf*SET10-HA-KD parasites and WT NF54 served as controls. Equal loading was confirmed by immunoblotting with rabbit antisera against *Pf*39 (~39 kDa). **(B)** Quantification of *Pf*SET10-HA levels following knockdown. *Pf*SET10-HA band intensities of three independent Western blots as described in (A) were quantified using Image J and normalized to the respective *Pf*39 protein band intensities (untreated set to 100%). Results are shown as mean ± SD.

**Figure S3.**
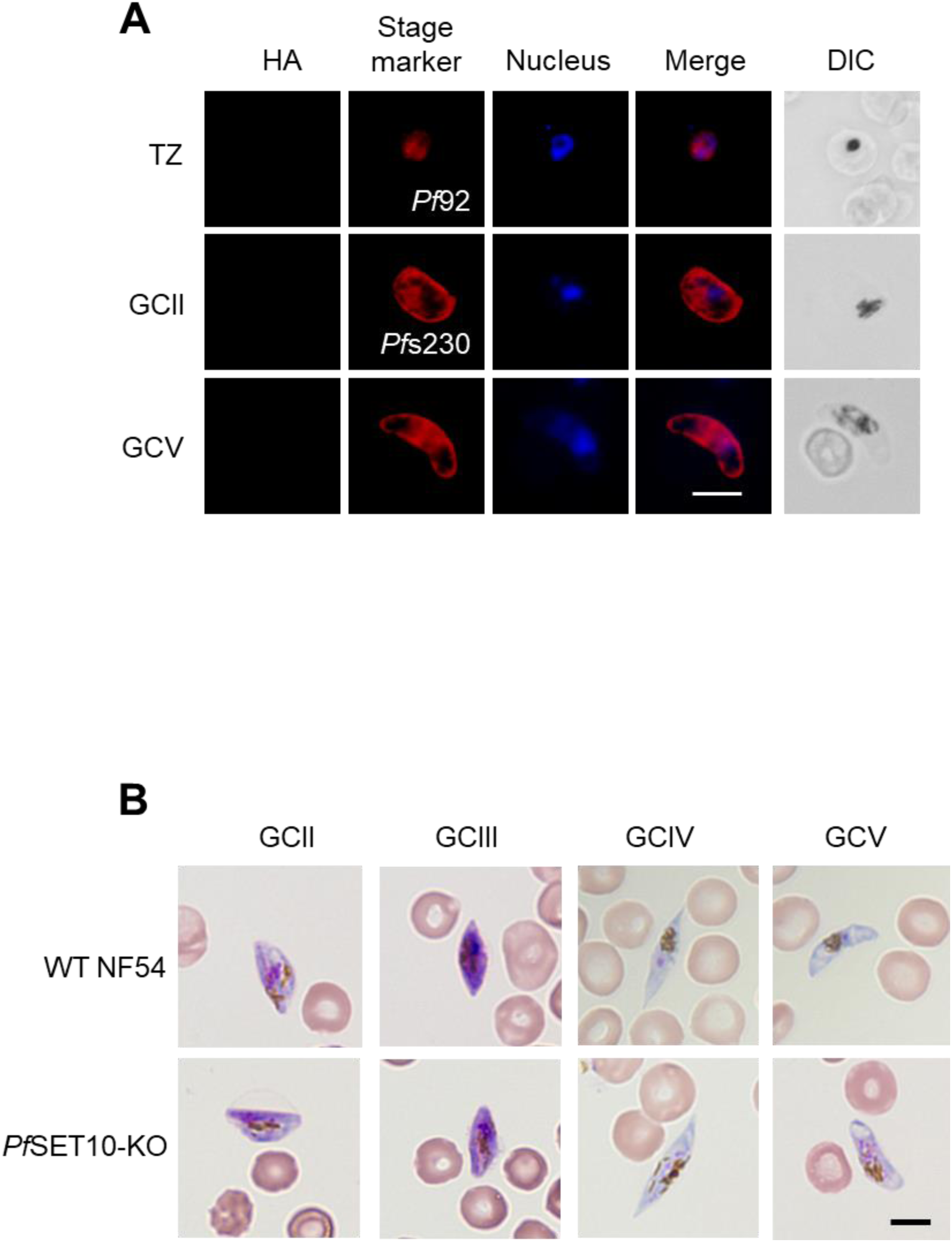
*Pf*SET10-HA immunolabelling control and *Pf*SET10-KO gametocyte morphology. (**A**) Immunofluorescence assay control for line *Pf*SET10-HA-KD. Methanol-fixed WT NF54 trophozoites (TZ) and gametocyte (GC) stages II and V were immunolabeled with rat anti-HA antibody (green). Asexual blood stages and gametocytes were highlighted by rabbit antisera directed against *Pf*92 and *Pf*s230, respectively (red); nuclei were highlighted by Hoechst 33342 nuclear stain (blue). Bar, 5 μm. **(B)** Morphology of *Pf*SET10-KO gametocytes. Gametocyte (GC) stages II-V of line *Pf*SET10-KO and WT NF54 were Giemsa-stained to compare their morphologies. Bar, 5 μm. Results (A, B) are representative of two independent experiments.

**Figure S4.**
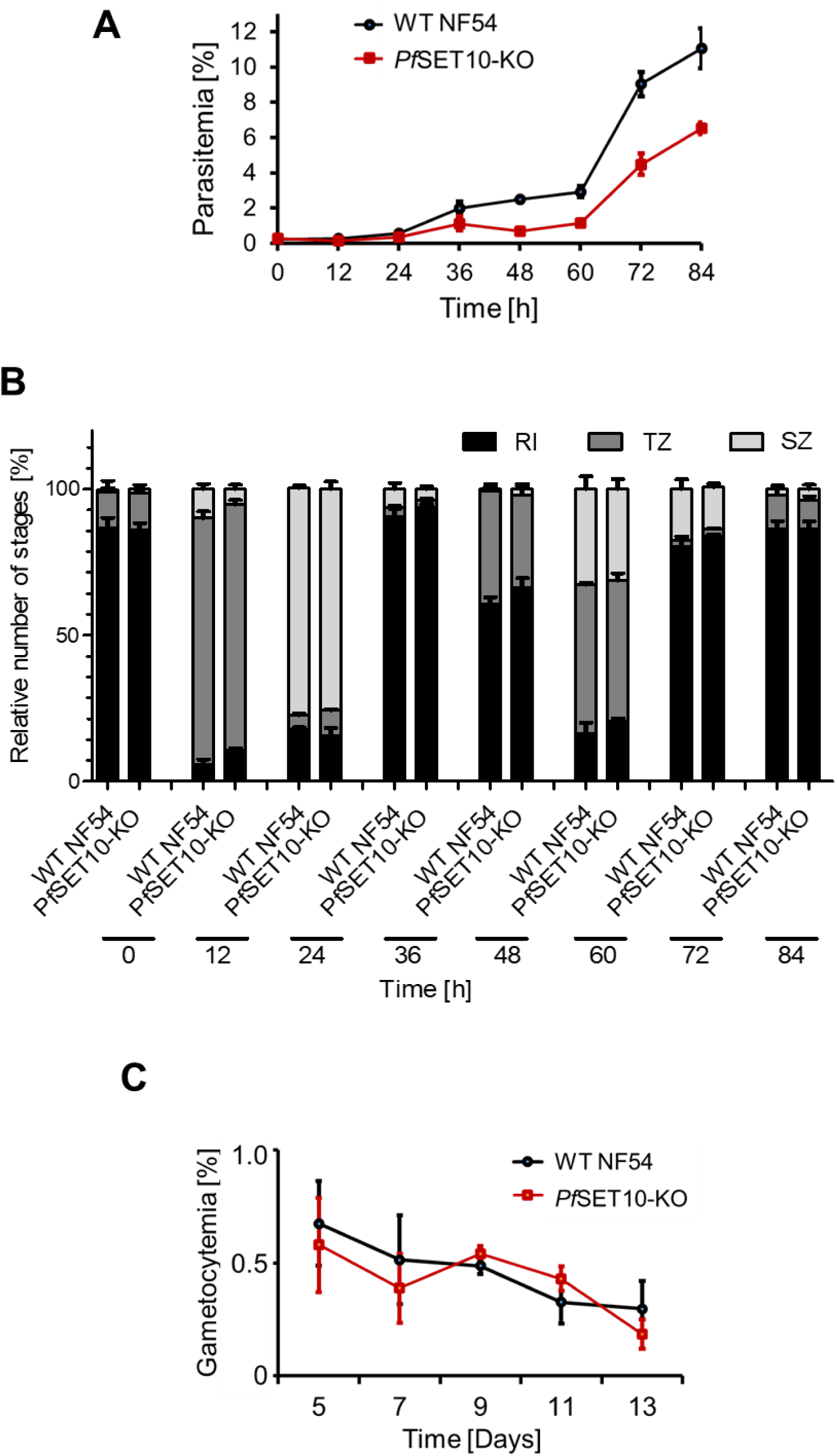
Intraerythrocytic growth and gametocyte development of the *Pf*SET10-KO line. (**A**) Intraerythrocytic growth of line *Pf*SET10-KO. Synchronized ring stage cultures of line *Pf*SET10-KO and WT NF54 were set up with an initial parasitemia of 0.25%. Parasitemia was followed via Giemsa smears over a time-period of 0 - 84 h. The experiment was performed in triplicate (mean ± SD). (**B**) Stage development of the *Pf*SET10-KO blood stages. Parasites were set up as described in (A) and the numbers of rings (RI), trophozoites (TZ), and schizonts (SZ) were determined in a total number of 50 infected RBCs every 24 h via Giemsa-stained blood smears. The experiment was performed in triplicate. (**C**) Gametocyte development of line *Pf*SET10-KO. Gametocyte production was induced in synchronized ring stage cultures of line *Pf*SET10-KO and WT NF54 at a parasitemia of 5.3% by addition of lysed RBCs and the gametocytemia was followed via Giemsa smears every 48 h between day 5 - 13 post-induction. The experiment was performed in triplicate (mean ± SD).

**Figure S5.**
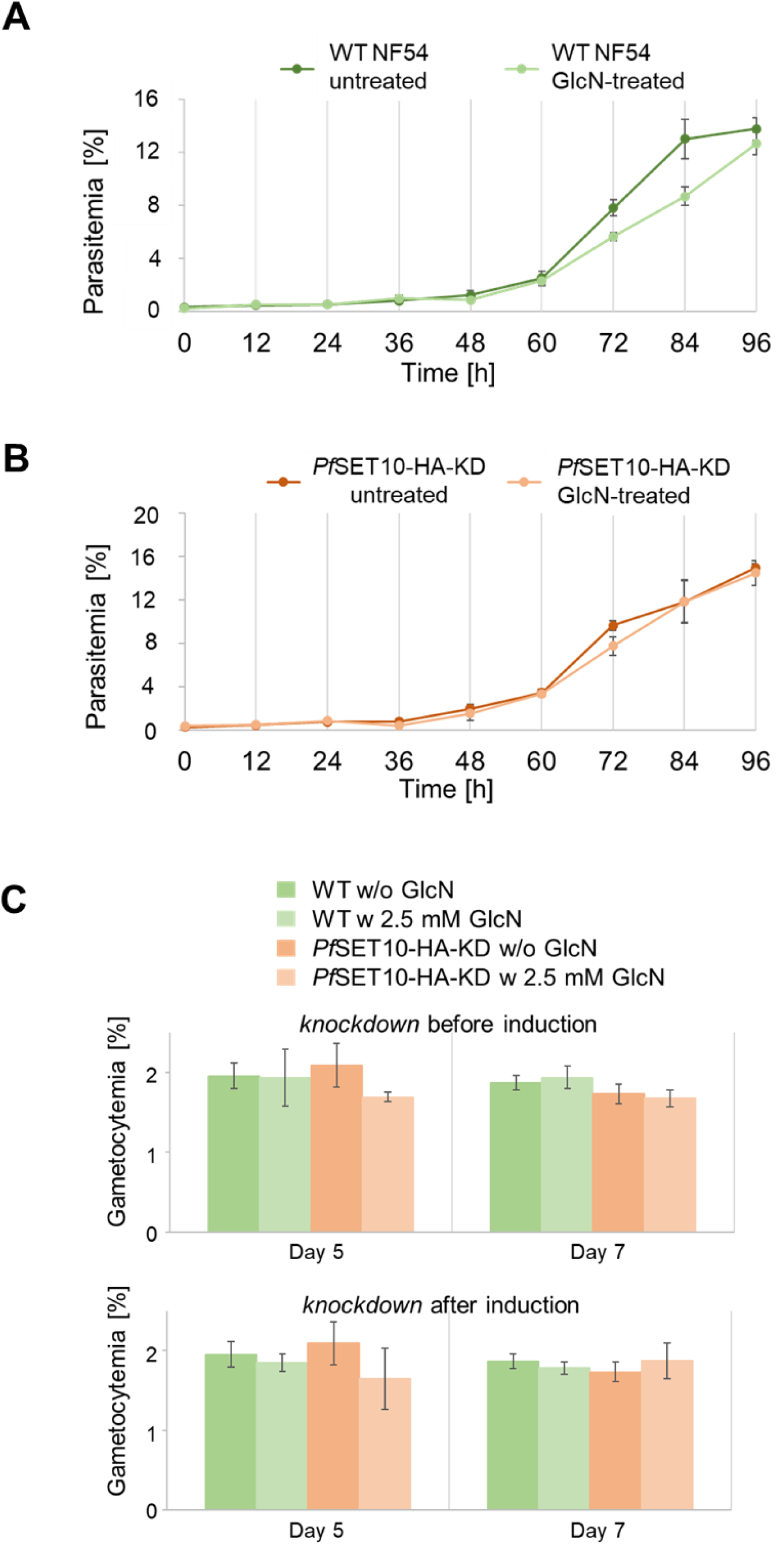
Intraerythrocytic growth and gametocyte formation following *Pf*SET10-HA knockdown. (**A**, **B**) Intraerythrocytic growth following *Pf*SET10-HA knockdown. Synchronized ring stage cultures of line *Pf*SET10-HA-KD (**A**) and WT NF54 (**B**) with a starting parasitemia of 0.25% were maintained in cell culture medium supplemented or not with 2.5 mM GlcN for transcript knockdown. The parasitemia was followed via Giemsa smears over a time-period of 0 - 96 h. The experiment was performed in triplicate (mean ± SD). (**C**) Gametocyte formation in dependence of *Pf*SET10-HA knockdown. Synchronized parasites of line *Pf*SET10-HA-KD and WT NF54 were cultivated in the presence (knockdown before induction) or absence (knockdown after induction) of 2.5 mM GlcN. Gametocyte production was then induced by addition of lysed RBCs. The cultures were maintained in cell culture medium supplemented or not with 2.5 mM GlcN and the gametocytemia was determined via Giemsa smears on days 5 and 7 post-induction. The experiment was performed in triplicate (mean ± SD).

**Figure S6.**
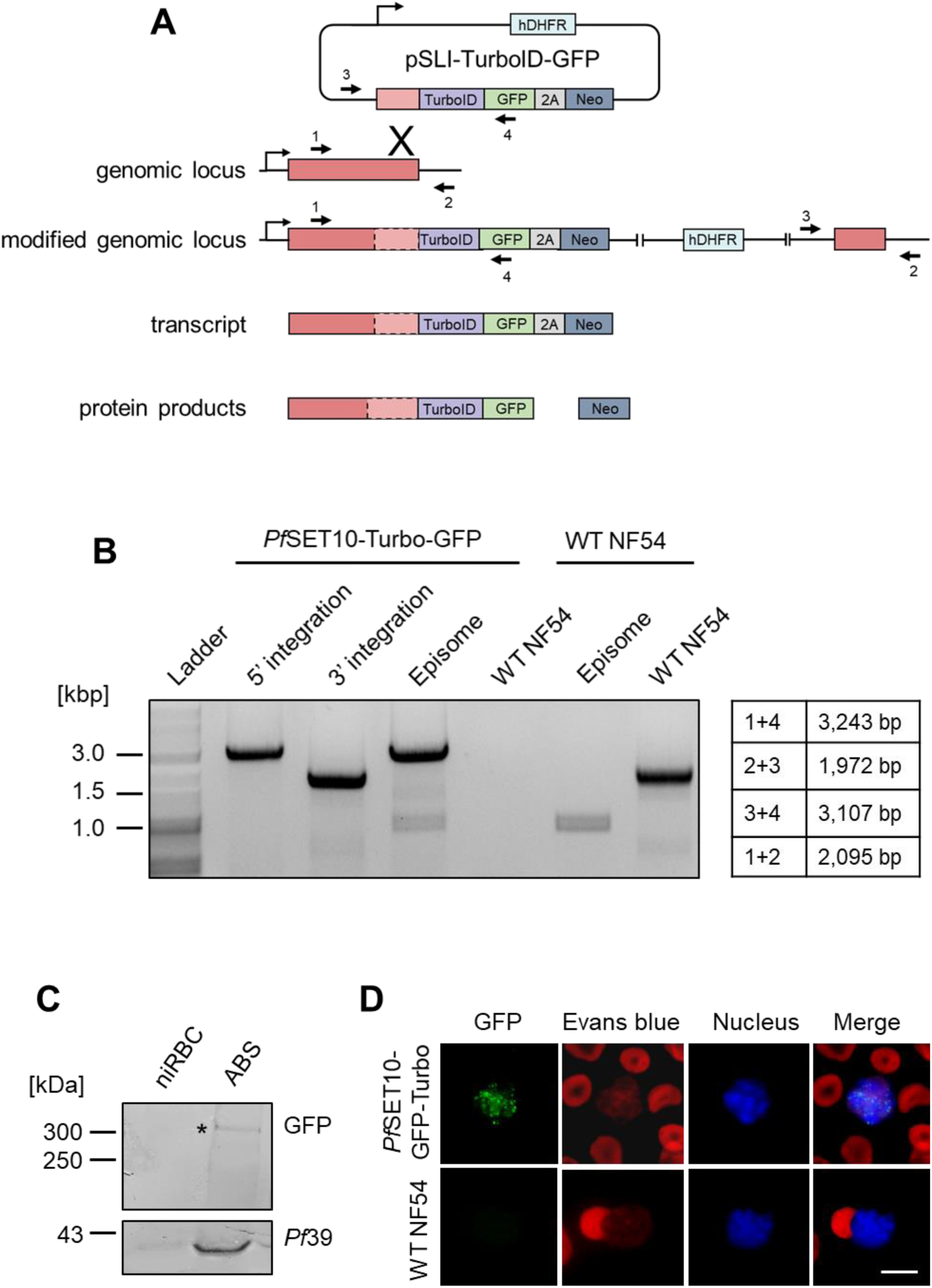
Generation of the *Pf*SET10-TurboID-GFP line. (**A**) Schematic depicting the single-crossover homologous recombination strategy for the generation of the pSLI-TurboID-GFP-based line. The coding region of the gene of interest was fused at the 3’-end with a sequence coding for an advanced *E. coli* biotin ligase and a GFP-encoding sequence followed by the 2A-skip peptide sequence and the Neo sequence. The numbered arrows indicate the positions of primers used to confirm vector integration. HA, hemagglutinin; hDHFR, human dihydrofolate reductase-encoding gene conferring resistance to WR99210; Neo, gene conferring resistance to neomycin. **(B)** Confirmation of vector integration into the *pfset10* gene locus. Diagnostic PCR demonstrates successful 5′ (primers 1 and 4) and 3′ (primers 3 and 2) integration. As a control, WT NF54 gDNA was used, demonstrating the original gene locus (primers 1 and 2). Episomal DNA was further detected (primers 3 and 4). Band sizes are indicated. (**C**) Expression of *Pf*SET10-TurboID-GFP. Lysates of asexual blood stages (ABS) were immunoblotted with mouse anti-GFP antibody to detect *Pf*SET10-TurboID-GFP (~300 kDa). Non-infected RBCs (niRBC) served as negative control; equal loading was confirmed by immunoblotting with rabbit antisera against *Pf*39 (~39 kDa). Asterisk (*) highlights the *Pf*SET10-TurboID-GFP protein. (**D**) Localization of *Pf*SET10-TurboID-GFP in transgenic schizonts. Methanol-fixed schizonts of line *Pf*SET10-TurboID-GFP and WT NF54 were immunolabeled with anti-GFP antibody (green); schizonts were highlighted by Evans Blue staining (red) and nuclei were highlighted by Hoechst 33342 nuclear stain (blue). Bar, 5 μm. Results (B-D) are representative of two independent experiments.

**Figure S7.**
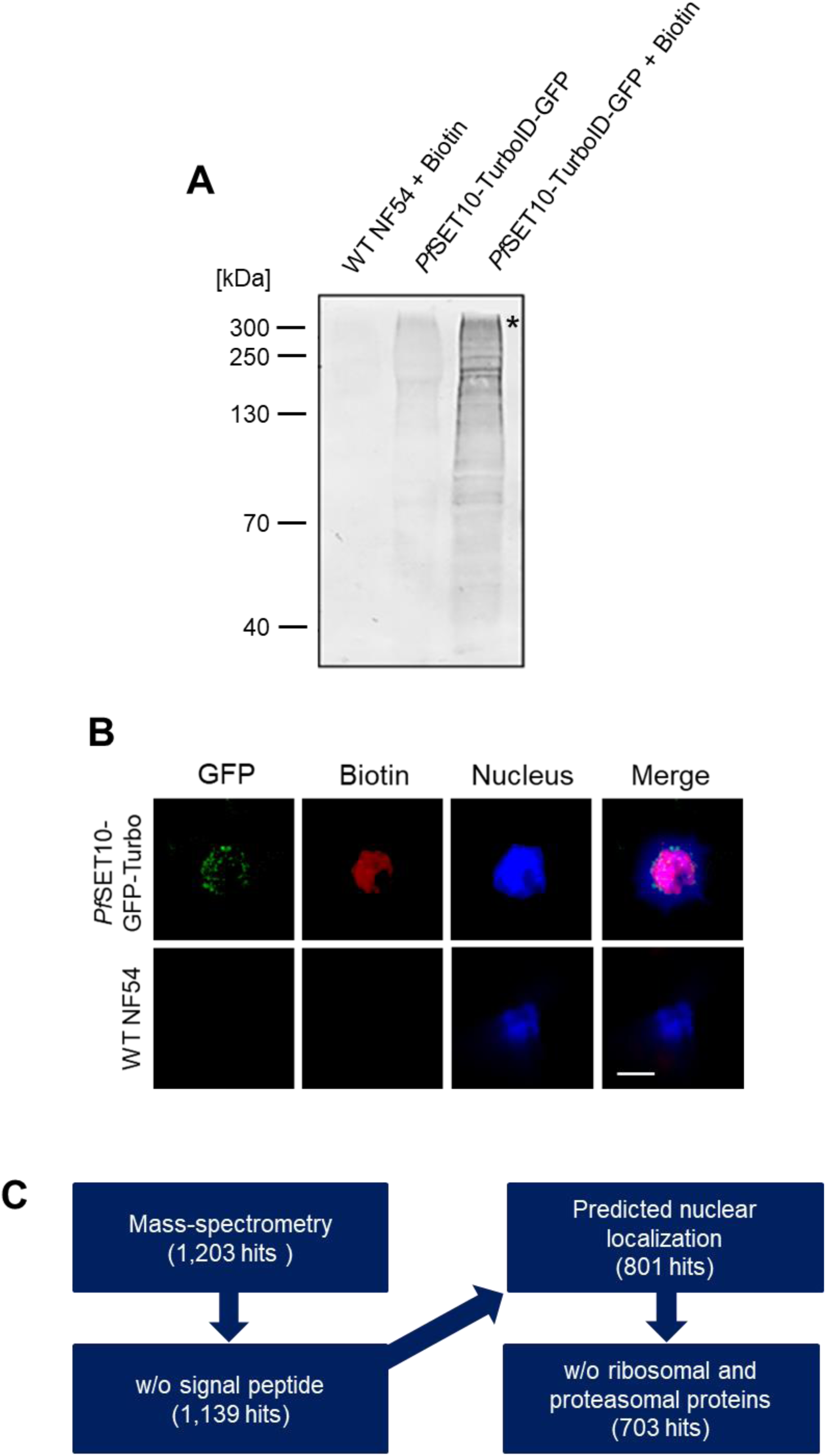
Verification of biotinylated proteins in the *Pf*SET10-TurboID-GFP line. (**A**) Detection of biotinylated proteins. Asexual blood stages of line *Pf*SET10-TurboID-GFP were treated or not with 50 μM biotin for 10 min. Lysates were prepared and immunoblotted with streptavidin coupled to alkaline phosphatase to detect biotinylated proteins. Asterisks (*) indicate biotinylated *Pf*SET10-TurboID-GFP (~300 kDa). Biotin-treated WT NF54 served as negative control. (**B**) Localization of biotinylated proteins in schizonts. Methanol-fixed schizonts of line *Pf*SET10-TurboID-GFP and WT NF54 were immunolabeled with mouse anti-GFP antibody (green). Biotinylated proteins were highlighted by fluorophore-conjugated streptavidin (red) and nuclei were highlighted by Hoechst 33342 nuclear stain (blue). Bar; 5 µm. (**C**) Schematic depicting the curation steps during analysis of the *Pf*SET10 interactors. Results (A, B) are representative of two independent experiments.

**Figure S8.**
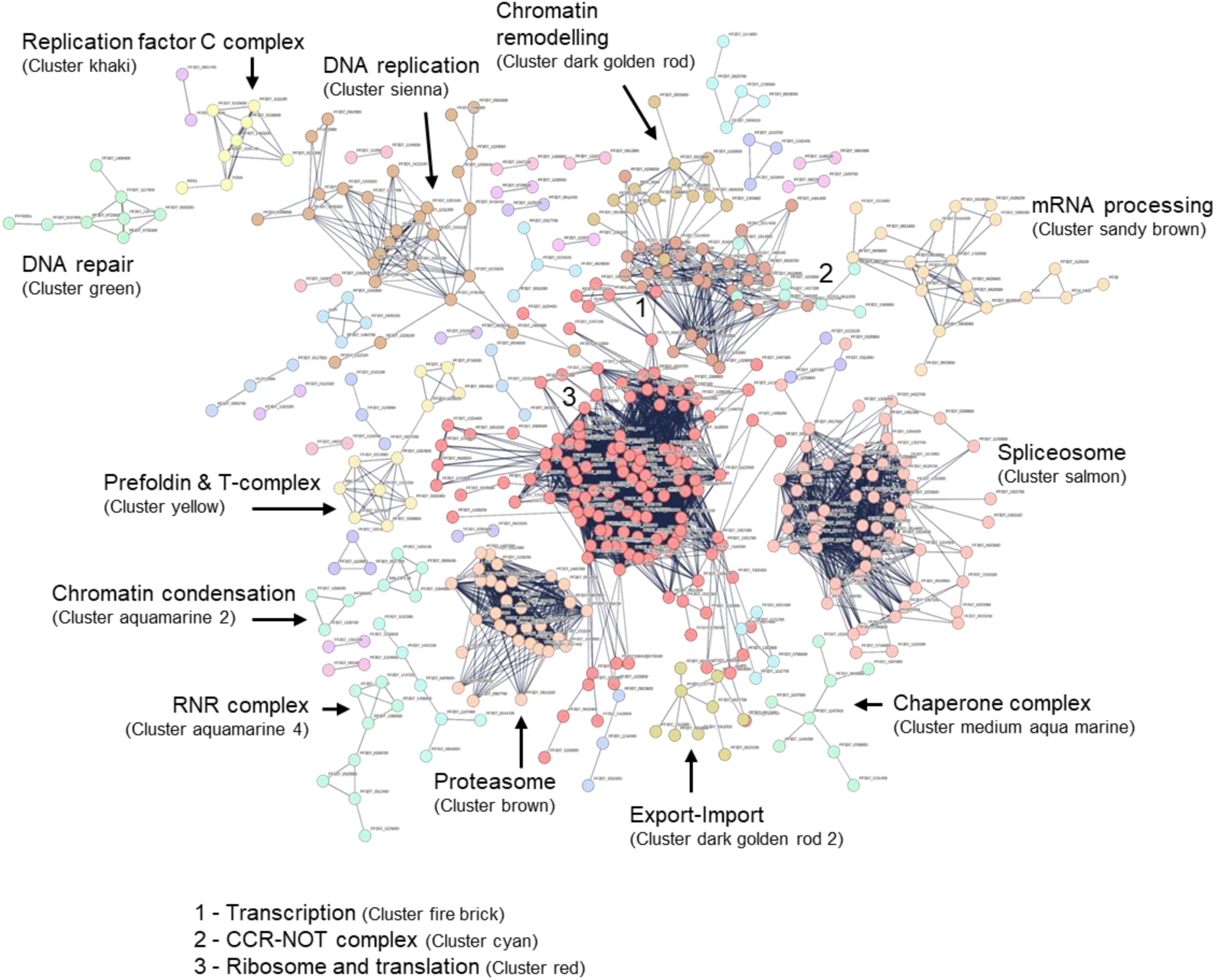
Network analysis of the *Pf*SET10 nuclear interactors. A protein-protein network analysis of the 801 nuclear *Pf*SET10 interactors in schizonts was performed (STRING program, highest interaction confidence of 0.9). Clustering of the interactors was employed by the K-means algorithm with a maximal cluster number of 43; disconnected nodes were excluded. Clusters ≥ 7 proteins were considered.

**Figure S9.**
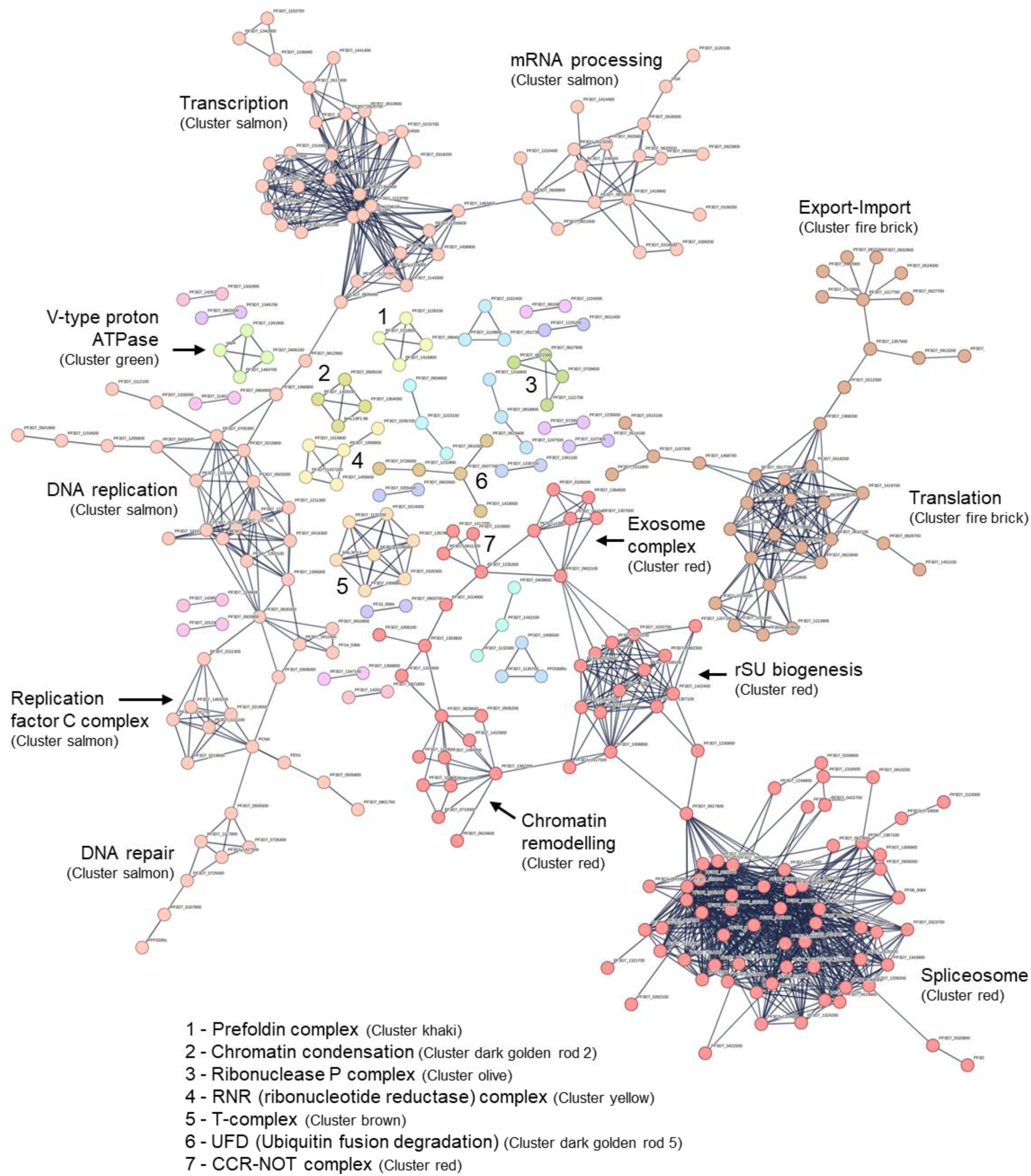
Network analysis of the *Pf*SET10 curated nuclear interactors. A protein-protein network analysis of the 703 nuclear *Pf*SET10 interactors in schizonts was performed (STRING program, highest interaction confidence of 0.9). Clustering of the interactors was employed by the K-means algorithm with a maximal cluster number of 29; disconnected nodes were excluded. Clusters ≥ 4 proteins were considered.

## References

Amit-Avraham I, Pozner G, Eshar S, Fastman Y, Kolevzon N, Yavin E, Dzikowski R. 2015. Antisense long noncoding RNAs regulate var gene activation in the malaria parasite Plasmodium falciparum. Proc Natl Acad Sci U S A 112:E982–91.

Antunes AV, Shahinas M, Swale C, Farhat DC, Ramakrishnan C, Bruley C, Cannella D, Robert MG, Corrao C, Couté Y, Hehl AB, Bougdour A, Coppens I, Hakimi M-A. 2024. In vitro production of cat-restricted Toxoplasma pre-sexual stages. Nature 625:366–376.

Aurrecoechea C, Brestelli J, Brunk BP, Dommer J, Fischer S, Gajria B, Gao X, Gingle A, Grant G, Harb OS, Heiges M, Innamorato F, Iodice J, Kissinger JC, Kraemer E, Li W, Miller JA, Nayak V, Pennington C, Pinney DF, Roos DS, Ross C, Stoeckert CJ, Treatman C, Wang H. 2009. PlasmoDB: a functional genomic database for malaria parasites. Nucleic Acids Res 37:D539–43.

Barcons-Simon A, Cordon-Obras C, Guizetti J, Bryant JM, Scherf A. 2020. CRISPR Interference of a Clonally Variant GC-Rich Noncoding RNA Family Leads to General Repression of var Genes in Plasmodium falciparum. mBio 11.

Bordiya Y, Zheng Y, Nam J-C, Bonnard AC, Choi HW, Lee B-K, Kim J, Klessig DF, Fei Z, Kang H-G. 2016. Pathogen Infection and MORC Proteins Affect Chromatin Accessibility of Transposable Elements and Expression of Their Proximal Genes in Arabidopsis. MPMI 29:674–687.

Brancucci NMB, Bertschi NL, Zhu L, Niederwieser I, Chin WH, Wampfler R, Freymond C, Rottmann M, Felger I, Bozdech Z, Voss TS. 2014. Heterochromatin protein 1 secures survival and transmission of malaria parasites. Cell Host Microbe 16:165–176.

Branon TC, Bosch JA, Sanchez AD, Udeshi ND, Svinkina T, Carr SA, Feldman JL, Perrimon N, Ting AY. 2018. Efficient proximity labeling in living cells and organisms with TurboID. Nat Biotechnol 36:880–887.

Bryant JM, Baumgarten S, Dingli F, Loew D, Sinha A, Claës A, Preiser PR, Dedon PC, Scherf A. 2020. Exploring the virulence gene interactome with CRISPR/dCas9 in the human malaria parasite. Mol Syst Biol 16:e9569.

Chahine Z, Gupta M, Lenz T, Hollin T, Abel S, Banks C, Saraf A, Prudhomme J, Florens L, Le Roch KG. 2023. PfMORC protein regulates chromatin accessibility and transcriptional repression in the human malaria parasite, P. falciparum. bioRxiv.

Chan A, Dziedziech A, Kirkman LA, Deitsch KW, Ankarklev J. 2020. A Histone Methyltransferase Inhibitor Can Reverse Epigenetically Acquired Drug Resistance in the Malaria Parasite Plasmodium falciparum. Antimicrob Agents Chemother 64.

Cheeseman K, Jannot G, Lourenço N, Villares M, Berthelet J, Calegari-Silva T, Hamroune J, Letourneur F, Rodrigues-Lima F, Weitzman JB. 2021. Dynamic methylation of histone H3K18 in differentiating Theileria parasites. Nat Commun 12:3221.

Coetzee N, Grüning H von, Opperman D, van der Watt M, Reader J, Birkholtz L-M. 2020. Epigenetic inhibitors target multiple stages of Plasmodium falciparum parasites. Sci Rep 10.

Coetzee N, Sidoli S, van Biljon R, Painter H, Llinás M, Garcia BA, Birkholtz L-M. 2017. Quantitative chromatin proteomics reveals a dynamic histone post-translational modification landscape that defines asexual and sexual Plasmodium falciparum parasites. Sci Rep 7:607.

Coleman BI, Skillman KM, Jiang RHY, Childs LM, Altenhofen LM, Ganter M, Leung Y, Goldowitz I, Kafsack BFC, Marti M, Llinás M, Buckee CO, Duraisingh MT. 2014. A Plasmodium falciparum histone deacetylase regulates antigenic variation and gametocyte conversion. Cell Host Microbe 16:177–186.

Cox J, Hein MY, Luber CA, Paron I, Nagaraj N, Mann M. 2014. Accurate proteome-wide label-free quantification by delayed normalization and maximal peptide ratio extraction, termed MaxLFQ. Mol Cell Proteomics 13:2513–2526.

Cox J, Mann M. 2008. MaxQuant enables high peptide identification rates, individualized p.p.b.-range mass accuracies and proteome-wide protein quantification. Nat Biotechnol 26:1367–1372.

Cui L, Fan Q, Cui L, Miao J. 2008. Histone lysine methyltransferases and demethylases in Plasmodium falciparum. Int J Parasitol 38:1083–1097.

Cui L, Miao J. 2010. Chromatin-mediated epigenetic regulation in the malaria parasite Plasmodium falciparum. Eukaryot Cell 9:1138–1149.

Duffy MF, Selvarajah SA, Josling GA, Petter M. 2014. Epigenetic regulation of the Plasmodium falciparum genome. Brief Funct Genomics 13:203–216.

Duraisingh MT, Horn D. 2016. Epigenetic Regulation of Virulence Gene Expression in Parasitic Protozoa. Cell Host Microbe 19:629–640.

Duraisingh MT, Voss TS, Marty AJ, Duffy MF, Good RT, Thompson JK, Freitas-Junior LH, Scherf A, Crabb BS, Cowman AF. 2005. Heterochromatin silencing and locus repositioning linked to regulation of virulence genes in Plasmodium falciparum. Cell 121:13–24.

Farhat DC, Swale C, Dard C, Cannella D, Ortet P, Barakat M, Sindikubwabo F, Belmudes L, Bock P-J de, Couté Y, Bougdour A, Hakimi M-A. 2020. A MORC-driven transcriptional switch controls Toxoplasma developmental trajectories and sexual commitment. Nat Microbiol 5:570–583.

Farrukh A, Musabyimana JP, Distler U, Mahlich VJ, Mueller J, Bick F, Tenzer S, Pradel G, Ngwa CJ. 2024. The Plasmodium falciparum CCCH zinc finger protein MD3 regulates male gametocytogenesis through its interaction with RNA-binding proteins. Mol Microbiol 121:543–564.

Fivelman QL, McRobert L, Sharp S, Taylor CJ, Saeed M, Swales CA, Sutherland CJ, Baker DA. 2007. Improved synchronous production of Plasmodium falciparum gametocytes in vitro. Mol Biochem Parasitol 154:119–123.

Flammersfeld A, Panyot A, Yamaryo-Botté Y, Aurass P, Przyborski JM, Flieger A, Botté C, Pradel G. 2020. A patatin-like phospholipase functions during gametocyte induction in the malaria parasite Plasmodium falciparum. Cell Microbiol 22:e13146.

Florens L, Washburn MP, Raine JD, Anthony RM, Grainger M, Haynes JD, Moch JK, Muster N, Sacci JB, Tabb DL, Witney AA, Wolters D, Wu Y, Gardner MJ, Holder AA, Sinden RE, Yates JR, Carucci DJ. 2002. A proteomic view of the Plasmodium falciparum life cycle. Nature 419:520–526.

Freitas-Junior LH, Hernandez-Rivas R, Ralph SA, Montiel-Condado D, Ruvalcaba-Salazar OK, Rojas-Meza AP, Mâncio-Silva L, Leal-Silvestre RJ, Gontijo AM, Shorte S, Scherf A. 2005. Telomeric heterochromatin propagation and histone acetylation control mutually exclusive expression of antigenic variation genes in malaria parasites. Cell 121:25–36.

Gardner MJ, Hall N, Fung E, White O, Berriman M, Hyman RW, Carlton JM, Pain A, Nelson KE, Bowman S, Paulsen IT, James K, Eisen JA, Rutherford K, Salzberg SL, Craig A, Kyes S, Chan M-S, Nene V, Shallom SJ, Suh B, Peterson J, Angiuoli S, Pertea M, Allen J, Selengut J, Haft D, Mather MW, Vaidya AB, Martin DMA, Fairlamb AH, Fraunholz MJ, Roos DS, Ralph SA, McFadden GI, Cummings LM, Subramanian GM, Mungall C, Venter JC, Carucci DJ, Hoffman SL, Newbold C, Davis RW, Fraser CM, Barrell B. 2002. Genome sequence of the human malaria parasite Plasmodium falciparum. Nature 419:498–511.

Ge SX, Jung D, Yao R. 2020. ShinyGO: a graphical gene-set enrichment tool for animals and plants. Bioinformatics 36:2628–2629.

Ghasemi S. 2020. Cancer’s epigenetic drugs: where are they in the cancer medicines? Pharmacogenomics J 20:367–379.

Hanhsen B, Farrukh A, Pradel G, Ngwa CJ. 2022. The Plasmodium falciparum CCCH Zinc Finger Protein ZNF4 Plays an Important Role in Gametocyte Exflagellation through the Regulation of Male Enriched Transcripts. Cells 11.

Herrera-Solorio AM, Vembar SS, MacPherson CR, Lozano-Amado D, Meza GR, Xoconostle-Cazares B, Martins RM, Chen P, Vargas M, Scherf A, Hernández-Rivas R. 2019. Clipped histone H3 is integrated into nucleosomes of DNA replication genes in the human malaria parasite Plasmodium falciparum. EMBO Rep 20.

Hillier C, Pardo M, Yu L, Bushell E, Sanderson T, Metcalf T, Herd C, Anar B, Rayner JC, Billker O, Choudhary JS. 2019. Landscape of the Plasmodium Interactome Reveals Both Conserved and Species-Specific Functionality. Cell Rep 28:1635–1647.e5.

Hoeijmakers WAM, Miao J, Schmidt S, Toenhake CG, Shrestha S, Venhuizen J, Henderson R, Birnbaum J, Ghidelli-Disse S, Drewes G, Cui L, Stunnenberg HG, Spielmann T, Bártfai R. 2019. Epigenetic reader complexes of the human malaria parasite, Plasmodium falciparum. Nucleic Acids Res 47:11574–11588.

Hughes CS, Foehr S, Garfield DA, Furlong EE, Steinmetz LM, Krijgsveld J. 2014. Ultrasensitive proteome analysis using paramagnetic bead technology. Mol Syst Biol 10:757.

Iyer LM, Anantharaman V, Wolf MY, Aravind L. 2008. Comparative genomics of transcription factors and chromatin proteins in parasitic protists and other eukaryotes. Int J Parasitol 38:1–31.

Jabeena CA, Rajavelu A. 2019. Epigenetic Players of Chromatin Structure Regulation in Plasmodium falciparum. Chembiochem 20:1225–1230.

Jiang L, Mu J, Zhang Q, Ni T, Srinivasan P, Rayavara K, Yang W, Turner L, Lavstsen T, Theander TG, Peng W, Wei G, Jing Q, Wakabayashi Y, Bansal A, Luo Y, Ribeiro JMC, Scherf A, Aravind L, Zhu J, Zhao K, Miller LH. 2013. PfSETvs methylation of histone H3K36 represses virulence genes in Plasmodium falciparum. Nature 499:223–227.

Kang H-G, Woo Choi H, Einem S von, Manosalva P, Ehlers K, Liu P-P, Buxa SV, Moreau M, Mang H-G, Kachroo P, Kogel K-H, Klessig DF. 2012. CRT1 is a nuclear-translocated MORC endonuclease that participates in multiple levels of plant immunity. Nat Commun 3.

Kariuki MM, Kiaira JK, Mulaa FK, Mwangi JK, Wasunna MK, Martin SK. 1998. Plasmodium falciparum: purification of the various gametocyte developmental stages from in vitro-cultivated parasites. Am J Trop Med Hyg 59:505–508.

Kim H, Yen L, Wongpalee SP, Kirshner JA, Mehta N, Xue Y, Johnston JB, Burlingame AL, Kim JK, Loparo JJ, Jacobsen SE. 2019. The Gene-Silencing Protein MORC-1 Topologically Entraps DNA and Forms Multimeric Assemblies to Cause DNA Compaction. Molecular Cell 75:700–710.e6.

Koch A, Kang H-G, Steinbrenner J, Dempsey DA, Klessig DF, Kogel K-H. 2017. MORC Proteins: Novel Players in Plant and Animal Health. Front Plant Sci 8:1720.

Lasonder E, Ishihama Y, Andersen JS, Vermunt AMW, Pain A, Sauerwein RW, Eling WMC, Hall N, Waters AP, Stunnenberg HG, Mann M. 2002. Analysis of the Plasmodium falciparum proteome by high-accuracy mass spectrometry. Nature 419:537–542.

Lasonder E, Rijpma SR, van Schaijk BCL, Hoeijmakers WAM, Kensche PR, Gresnigt MS, Italiaander A, Vos MW, Woestenenk R, Bousema T, Mair GR, Khan SM, Janse CJ, Bártfai R, Sauerwein RW. 2016. Integrated transcriptomic and proteomic analyses of P. falciparum gametocytes: molecular insight into sex-specific processes and translational repression. Nucleic Acids Res 44:6087–6101.

Le Roch KG, Zhou Y, Blair PL, Grainger M, Moch JK, Haynes JD, La Vega P de, Holder AA, Batalov S, Carucci DJ, Winzeler EA. 2003. Discovery of gene function by expression profiling of the malaria parasite life cycle. Science 301:1503–1508.

Lee HJ, Hore TA, Reik W. 2014. Reprogramming the methylome: erasing memory and creating diversity. Cell Stem Cell 14:710–719.

Li D-Q, Nair SS, Kumar R. 2013. The MORC family: new epigenetic regulators of transcription and DNA damage response. Epigenetics 8:685–693.

Llinás M, Deitsch KW, Voss TS. 2008. Plasmodium gene regulation: far more to factor in. Trends Parasitol 24:551–556.

López-Barragán MJ, Lemieux J, Quiñones M, Williamson KC, Molina-Cruz A, Cui K, Barillas-Mury C, Zhao K, Su X. 2011. Directional gene expression and antisense transcripts in sexual and asexual stages of Plasmodium falciparum. BMC Genomics 12:587.

Lopez-Rubio JJ, Gontijo AM, Nunes MC, Issar N, Hernandez Rivas R, Scherf A. 2007. 5’ flanking region of var genes nucleate histone modification patterns linked to phenotypic inheritance of virulence traits in malaria parasites. Mol Microbiol 66:1296–1305.

Lopez-Rubio J-J, Mancio-Silva L, Scherf A. 2009. Genome-wide analysis of heterochromatin associates clonally variant gene regulation with perinuclear repressive centers in malaria parasites. Cell Host Microbe 5:179–190.

Lorković ZJ. 2012. MORC proteins and epigenetic regulation. Plant Signal Behav 7:1561–1565.

Malmquist NA, Moss TA, Mecheri S, Scherf A, Fuchter MJ. 2012. Small-molecule histone methyltransferase inhibitors display rapid antimalarial activity against all blood stage forms in Plasmodium falciparum. Proc. Natl. Acad. Sci. U.S.A. 109:16708–16713.

Malmquist NA, Sundriyal S, Caron J, Chen P, Witkowski B, Menard D, Suwanarusk R, Renia L, Nosten F, Jiménez-Díaz MB, Angulo-Barturen I, Martínez MS, Ferrer S, Sanz LM, Gamo F-J, Wittlin S, Duffy S, Avery VM, Ruecker A, Delves MJ, Sinden RE, Fuchter MJ, Scherf A. 2015. Histone Methyltransferase Inhibitors Are Orally Bioavailable, Fast-Acting Molecules with Activity against Different Species Causing Malaria in Humans. Antimicrob Agents Chemother 59:950–959.

Marsh S, Jimeno A. 2020. Tazemetostat for the treatment of multiple types of hematological malignancies and solid tumors. Drugs of Today 56:377.

Marzochi LL, Cuzziol CI, Nascimento Filho CHVD, dos Santos JA, Castanhole-Nunes MMU, Pavarino ÉC, Guerra ENS, Goloni-Bertollo EM. 2023. Use of histone methyltransferase inhibitors in cancer treatment: A systematic review. European Journal of Pharmacology 944:175590.

Miao J, Fan Q, Cui L, Li J, Li J, Cui L. 2006. The malaria parasite Plasmodium falciparum histones: organization, expression, and acetylation. Gene 369:53–65.

Moissiard G, Cokus SJ, Cary J, Feng S, Billi AC, Stroud H, Husmann D, Zhan Y, Lajoie BR, McCord RP, Hale CJ, Feng W, Michaels SD, Frand AR, Pellegrini M, Dekker J, Kim JK, Jacobsen SE. 2012. MORC Family ATPases Required for Heterochromatin Condensation and Gene Silencing. Science 336:1448–1451.

Musabyimana JP, Distler U, Sassmannshausen J, Berks C, Manti J, Bennink S, Blaschke L, Burda P-C, Flammersfeld A, Tenzer S, Ngwa CJ, Pradel G. 2022. Plasmodium falciparum S-Adenosylmethionine Synthetase Is Essential for Parasite Survival through a Complex Interaction Network with Cytoplasmic and Nuclear Proteins. Microorganisms 10.

Ngwa CJ, Gross MR, Musabyimana J-P, Pradel G, Deitsch KW. 2021. The Role of the Histone Methyltransferase PfSET10 in Antigenic Variation by Malaria Parasites: a Cautionary Tale. mSphere 6.

Ngwa CJ, Kiesow MJ, Papst O, Orchard LM, Filarsky M, Rosinski AN, Voss TS, Llinás M, Pradel G. 2017. Transcriptional Profiling Defines Histone Acetylation as a Regulator of Gene Expression during Human-to-Mosquito Transmission of the Malaria Parasite Plasmodium falciparum. Front Cell Infect Microbiol 7:320.

Ngwa CJ, Kiesow MJ, Orchard LM, Farrukh A, Llinás M, Pradel G. 2019. The G9a Histone Methyltransferase Inhibitor BIX-01294 Modulates Gene Expression during Plasmodium falciparum Gametocyte Development and Transmission. Int J Mol Sci 20.

Painter HJ, Campbell TL, Llinás M. 2011. The Apicomplexan AP2 family: integral factors regulating Plasmodium development. Mol Biochem Parasitol 176:1–7.

Otto TD, Wilinski D, Assefa S, Keane TM, Sarry LR, Böhme U, Lemieux J, Barrell B, Pain A, Berriman M, Newbold C, Llinás M. 2010. New insights into the blood-stage transcriptome of Plasmodium falciparum using RNA-Seq. Mol Microbiol 76:12–24.

Pérez-Toledo K, Rojas-Meza AP, Mancio-Silva L, Hernández-Cuevas NA, Delgadillo DM, Vargas M, Martínez-Calvillo S, Scherf A, Hernandez-Rivas R. 2009. Plasmodium falciparum heterochromatin protein 1 binds to tri-methylated histone 3 lysine 9 and is linked to mutually exclusive expression of var genes. Nucleic Acids Res 37:2596–2606.

Petter M, Lee CC, Byrne TJ, Boysen KE, Volz J, Ralph SA, Cowman AF, Brown GV, Duffy MF. 2011. Expression of P. falciparum var genes involves exchange of the histone variant H2A.Z at the promoter. PLoS Pathog 7:e1001292.

Prommana P, Uthaipibull C, Wongsombat C, Kamchonwongpaisan S, Yuthavong Y, Knuepfer E, Holder AA, Shaw PJ. 2013. Inducible knockdown of Plasmodium gene expression using the glmS ribozyme. PLoS One 8:e73783.

Reers AB, Bautista R, McLellan J, Morales B, Garza R, Bol S, Hanson KK, Bunnik EM. 2023. Histone modification analysis reveals common regulators of gene expression in liver and blood stage merozoites of Plasmodium parasites. Epigenetics Chromatin 16:25.

Salcedo-Amaya AM, van Driel MA, Alako BT, Trelle MB, van den Elzen AMG, Cohen AM, Janssen-Megens EM, van de Vegte-Bolmer M, Selzer RR, Iniguez AL, Green RD, Sauerwein RW, Jensen ON, Stunnenberg HG. 2009. Dynamic histone H3 epigenome marking during the intraerythrocytic cycle of Plasmodium falciparum. Proc Natl Acad Sci U S A 106:9655–9660.

Sassmannshausen J, Bennink S, Distler U, Küchenhoff J, Minns AM, Lindner SE, Burda P-C, Tenzer S, Gilberger TW, Pradel G. 2024. Comparative proteomics of vesicles essential for the egress of Plasmodium falciparum gametocytes from red blood cells. Mol Microbiol 121:431–452.

Saraf A, Cervantes S, Bunnik EM, Ponts N, Sardiu ME, Chung D-WD, Prudhomme J, Varberg JM, Wen Z, Washburn MP, Florens L, Le Roch KG. 2016. Dynamic and Combinatorial Landscape of Histone Modifications during the Intraerythrocytic Developmental Cycle of the Malaria Parasite. J Proteome Res 15:2787–2801.

Shang X, Wang C, Fan Y, Guo G, Wang F, Zhao Y, Sheng F, Tang J, He X, Yu X, Zhang M, Zhu G, Yin S, Mu J, Culleton R, Cao J, Jiang M, Zhang Q. 2022. Genome-wide landscape of ApiAP2 transcription factors reveals a heterochromatin-associated regulatory network during Plasmodium falciparum blood-stage development. Nucleic Acids Res 50:3413–3431.

Shrestha S, Lucky AB, Brashear AM, Li X, Cui L, Miao J. 2022. Distinct Histone Post-translational Modifications during Plasmodium falciparum Gametocyte Development. J Proteome Res 21:1857–1867.

Sielaff M, Kuharev J, Bohn T, Hahlbrock J, Bopp T, Tenzer S, Distler U. 2017. Evaluation of FASP, SP3, and iST Protocols for Proteomic Sample Preparation in the Low Microgram Range. J Proteome Res 16:4060–4072.

Simon V, Czobor P, Bálint S, Mészáros A, Bitter I. 2009. Prevalence and correlates of adult attention-deficit hyperactivity disorder: meta-analysis. Br J Psychiatry 194:204–211.

Singh MK, Bonnell VA, Da Silva IT, Santiago VF, Moraes MS, Adderley J, Doerig C, Palmisano G, Llinás M, Garcia CRS. 2024. A Plasmodium falciparum MORC protein complex modulates epigenetic control of gene expression through interaction with heterochromatin.

Singh S, Santos JM, Orchard LM, Yamada N, van Biljon R, Painter HJ, Mahony S, Llinás M. 2021. The PfAP2-G2 transcription factor is a critical regulator of gametocyte maturation. Mol Microbiol 115:1005–1024.

Srivastava S, Holmes MJ, White MW, Sullivan WJ. 2023. Toxoplasma gondii AP2XII-2 Contributes to Transcriptional Repression for Sexual Commitment. mSphere 8.

Subudhi AK, Green JL, Satyam R, Salunke RP, Lenz T, Shuaib M, Isaioglou I, Abel S, Gupta M, Esau L, Mourier T, Nugmanova R, Mfarrej S, Shivapurkar R, Stead Z, Rached FB, Ostwal Y, Sougrat R, Dada A, Kadamany AF, Fischle W, Merzaban J, Knuepfer E, Ferguson DJP, Gupta I, Le Roch KG, Holder AA, Pain A. 2023. DNA-binding protein PfAP2-P regulates parasite pathogenesis during malaria parasite blood stages. Nat Microbiol 8:2154–2169.

Sundriyal S, Malmquist NA, Caron J, Blundell S, Liu F, Chen X, Srimongkolpithak N, Jin J, Charman SA, Scherf A, Fuchter MJ. 2014. Development of Diaminoquinazoline Histone Lysine Methyltransferase Inhibitors as Potent Blood-Stage Antimalarial Compounds. ChemMedChem 9:2360–2373.

Szklarczyk D, Gable AL, Lyon D, Junge A, Wyder S, Huerta-Cepas J, Simonovic M, Doncheva NT, Morris JH, Bork P, Jensen LJ, Mering C von. 2019. STRING v11: protein-protein association networks with increased coverage, supporting functional discovery in genome-wide experimental datasets. Nucleic Acids Res 47:D607–D613.

Tonkin CJ, Carret CK, Duraisingh MT, Voss TS, Ralph SA, Hommel M, Duffy MF, Da Silva LM, Scherf A, Ivens A, Speed TP, Beeson JG, Cowman AF. 2009. Sir2 paralogues cooperate to regulate virulence genes and antigenic variation in Plasmodium falciparum. PLoS Biol 7:e84.

Vizcaíno JA, Côté RG, Csordas A, Dianes JA, Fabregat A, Foster JM, Griss J, Alpi E, Birim M, Contell J, O’Kelly G, Schoenegger A, Ovelleiro D, Pérez-Riverol Y, Reisinger F, Ríos D, Wang R, Hermjakob H. 2013. The PRoteomics IDEntifications (PRIDE) database and associated tools: status in 2013. Nucleic Acids Res 41:D1063–9.

Volz J, Carvalho TG, Ralph SA, Gilson P, Thompson J, Tonkin CJ, Langer C, Crabb BS, Cowman AF. 2010. Potential epigenetic regulatory proteins localise to distinct nuclear sub-compartments in Plasmodium falciparum. Int J Parasitol 40:109–121.

Volz JC, Bártfai R, Petter M, Langer C, Josling GA, Tsuboi T, Schwach F, Baum J, Rayner JC, Stunnenberg HG, Duffy MF, Cowman AF. 2012. PfSET10, a Plasmodium falciparum methyltransferase, maintains the active var gene in a poised state during parasite division. Cell Host Microbe 11:7–18.

Voss TS, Bozdech Z, Bártfai R. 2014. Epigenetic memory takes center stage in the survival strategy of malaria parasites. Curr Opin Microbiol 20:88–95.

Weinhold B. 2006. Epigenetics: the science of change. Environ Health Perspect 114:A160–7.

Wichers JS, Scholz JAM, Strauss J, Witt S, Lill A, Ehnold L-I, Neupert N, Liffner B, Lühken R, Petter M, Lorenzen S, Wilson DW, Löw C, Lavazec C, Bruchhaus I, Tannich E, Gilberger TW, Bachmann A. 2019. Dissecting the Gene Expression, Localization, Membrane Topology, and Function of the Plasmodium falciparum STEVOR Protein Family. mBio 10.

Williamson KC, Keister DB, Muratova O, Kaslow DC. 1995. Recombinant Pfs230, a Plasmodium falciparum gametocyte protein, induces antisera that reduce the infectivity of Plasmodium falciparum to mosquitoes. Mol Biochem Parasitol 75:33–42.

Wirth CC, Glushakova S, Scheuermayer M, Repnik U, Garg S, Schaack D, Kachman MM, Weißbach T, Zimmerberg J, Dandekar T, Griffiths G, Chitnis CE, Singh S, Fischer R, Pradel G. 2014. Perforin-like protein PPLP2 permeabilizes the red blood cell membrane during egress of Plasmodium falciparum gametocytes. Cell Microbiol 16:709–733.

World Health Organization. 2023. WHO word malaria report 2023. Geneva, Switzerland: WHO.

Zhang M, Wang C, Otto TD, Oberstaller J, Liao X, Adapa SR, Udenze K, Bronner IF, Casandra D, Mayho M, Brown J, Li S, Swanson J, Rayner JC, Jiang RHY, Adams JH. 2018. Uncovering the essential genes of the human malaria parasite Plasmodium falciparum by saturation mutagenesis. Science 360.

Zhang Q, Huang Y, Zhang Y, Fang X, Claes A, Duchateau M, Namane A, Lopez-Rubio J-J, Pan W, Scherf A. 2011. A critical role of perinuclear filamentous actin in spatial repositioning and mutually exclusive expression of virulence genes in malaria parasites. Cell Host Microbe 10:451–463.

Zhong Z, Xue Y, Harris CJ, Wang M, Li Z, Ke Y, Liu M, Zhou J, Jami-Alahmadi Y, Feng S, Wohlschlegel JA, Jacobsen SE. 2023. MORC proteins regulate transcription factor binding by mediating chromatin compaction in active chromatin regions. Genome Biol 24:96.

Zohourian N, Brown JA. 2024. Current trends in clinical trials and the development of small molecule epigenetic inhibitors as cancer therapeutics. Epigenomics.

