## Supplementary material for "*Plasmodium falciparum* SET10 is a histone H3 lysine K18 methyltransferase that participates in a chromatin modulation network crucial for intraerythrocytic development": Table S1

Table S1. List of transcriptionally deregulated genes in *Pf* SET10-KO schizonts

### All genes

| gene_id | gene_name | baseMean | log2FoldChange | lfcSE | stat | pvalue | padj | 01_JP_S1 | 02_JP_S2 | 03_JP_S3 | 04_JP_S4 | sig |
|---|---|---|---|---|---|---|---|---|---|---|---|---|
| PF3D7_0222200 | EPF3 | 9.38287743 | 6.775961438 | 2.26972854 | 2.98536205 | 0.00283243 | 0.08850955 | 0 | 37.5315097 | 0 | 0 | Non-sig. |
| PF3D7_0324000 | EPF1 | 6.18678037 | 6.174912419 | 2.43564212 | 2.53522977 | 0.01123736 | 0.21636979 | 0 | 0 | 0 | 24.7471215 | Non-sig. |
| PF3D7_0631300 | EPF1 | 5.65487159 | 6.045042317 | 2.15379083 | 2.80669888 | 0.0050052 | 0.12701704 | 0 | 4.27746625 | 0 | 18.3420201 | Non-sig. |
| PF3D7_0701500 | EPF3 | 5.62972646 | 6.039015436 | 2.48154863 | 2.43356723 | 0.01495086 | 0.26055806 | 0 | 22.5189058 | 0 | 0 | Non-sig. |
| PF3D7_1255100 | RIF | 147.464972 | 5.921610151 | 0.72648659 | 8.15102472 | 3.6085E-16 | 3.1863E-13 | 8.3817014 | 339.507833 | 1.64123839 | 240.329117 | Sig. genes |
| PF3D7_0101600 | RIF | 37.1331716 | 5.661533899 | 1.1104928 | 5.09821755 | 3.4287E-07 | 6.2379E-05 | 3.14328293 | 88.0567073 | 0 | 57.3326961 | Sig. genes |
| PF3D7_1101600 | EPF3 | 4.03485676 | 5.558218907 | 2.67654688 | 2.07663798 | 0.03783498 | 0.43374033 | 0 | 0 | 0 | 16.1394271 | Non-sig. |
| PF3D7_0222000 | EPF1 | 3.48506876 | 5.347150627 | 2.77869214 | 1.92434079 | 0.05431188 | 0.50202873 | 0 | 13.940275 | 0 | 0 | Non-sig. |
| PF3D7_0901300 | RIF | 2.6853769 | 4.970961195 | 2.58036206 | 1.9264588 | 0.05404711 | 0.50202873 | 0 | 5.35140499 | 0 | 5.39010261 | Non-sig. |
| PF3D7_API06700 | N/A | 2.68082212 | 4.968643171 | 2.99184738 | 1.66072748 | 0.09676819 | 0.64895933 | 0 | 10.7232885 | 0 | 0 | Non-sig. |
| PF3D7_0901600 | N/A | 2.41571404 | 4.818342671 | 2.67404797 | 1.80189089 | 0.07156257 | 0.58133781 | 0 | 5.34814807 | 0 | 4.31470807 | Non-sig. |
| PF3D7_0412400 | VAR | 324.642616 | 4.676104275 | 0.6733819 | 6.94420842 | 3.8059E-12 | 1.8331E-09 | 42.1632811 | 549.15281 | 7.39343837 | 699.860934 | Sig. genes |
| PF3D7_1400500 | RIF | 2.1462242 | 4.647754629 | 2.79075725 | 1.66540985 | 0.09583101 | 0.64772846 | 0 | 5.34996435 | 0 | 3.23493244 | Non-sig. |
| PF3D7_0631900 | N/A | 163.528712 | 4.120673442 | 0.6666414 | 6.18124445 | 6.3598E-10 | 1.982E-07 | 6.27671638 | 362.226339 | 28.7163033 | 256.895489 | Sig. genes |
| PF3D7_0413100 | VAR | 71.2906139 | 4.064505801 | 0.90283001 | 4.50196132 | 6.7329E-06 | 0.00075896 | 0 | 162.986639 | 15.6035626 | 106.572254 | Sig. genes |
| PF3D7_0425200 | N/A | 143.660193 | 4.02169277 | 0.58422322 | 6.88382907 | 5.8265E-12 | 2.3745E-09 | 23.0781103 | 285.6895 | 10.6703764 | 255.202784 | Sig. genes |
| PF3D7_0201300 | N/A | 4.03292777 | 3.983187904 | 2.18870588 | 1.81988267 | 0.06877687 | 0.57367529 | 1.04571193 | 9.66907302 | 0 | 5.41692615 | Non-sig. |
| PF3D7_0712900 | VAR | 197.609312 | 3.965676312 | 0.51356184 | 7.72190605 | 1.146E-14 | 8.6738E-12 | 26.3480798 | 347.525513 | 21.3582618 | 395.205392 | Sig. genes |
| PF3D7_0425100 | N/A | 64.4411399 | 3.856353739 | 0.78292742 | 4.92555711 | 8.412E-07 | 0.00012045 | 14.631373 | 138.6929 | 2.461023 | 101.979264 | Sig. genes |
| PF3D7_1479000 | ACS1a | 484.348242 | 3.851159185 | 0.45233968 | 8.51386556 | 1.6823E-17 | 4.2116E-14 | 67.3819947 | 973.137637 | 58.3140639 | 838.559272 | Sig. genes |
| PF3D7_0632400 | RIF | 3.48929017 | 3.750592208 | 2.29942208 | 1.63110211 | 0.10286878 | 0.66002937 | 1.04794096 | 9.66441366 | 0 | 3.24480604 | Non-sig. |
| PF3D7_1148200 | N/A | 747.804978 | 3.718372609 | 0.45093968 | 8.24583148 | 1.6402E-16 | 1.7379E-13 | 117.201143 | 1188.36933 | 94.2194419 | 1591.43 | Sig. genes |
| PF3D7_1370500 | N/A | 959.806226 | 3.636647033 | 0.5012759 | 7.25478128 | 4.0231E-13 | 2.6643E-10 | 194.490947 | 1633.14664 | 91.8068248 | 1919.78049 | Sig. genes |
| PF3D7_0935600 | GIG | 3296.12516 | 3.499472447 | 0.51301391 | 6.82139878 | 9.0158E-12 | 3.4118E-09 | 758.026194 | 6160.75015 | 313.744555 | 5951.97973 | Sig. genes |
| PF3D7_1372700 | RIF | 2.95494456 | 3.486081916 | 2.37889561 | 1.4654203 | 0.14280628 | 0.74065302 | 1.04724102 | 6.44384163 | 0 | 4.32869558 | Non-sig. |
| PF3D7_1479200 | N/A | 373.606948 | 3.452431013 | 0.68866087 | 5.0132528 | 5.3518E-07 | 8.4674E-05 | 107.672908 | 742.870353 | 18.0481843 | 625.836345 | Sig. genes |
| PF3D7_0202200 | PTP1 | 368.896774 | 3.445386779 | 0.48718199 | 7.07207333 | 1.5264E-12 | 8.9852E-10 | 76.8446618 | 610.519869 | 47.6354415 | 740.587123 | Sig. genes |
| PF3D7_0631800 | RIF | 2.6268725 | 3.417478368 | 2.45869665 | 1.38995527 | 0.16454246 | 0.77284241 | 0 | 4.28773692 | 0.81866927 | 5.40108382 | Non-sig. |
| PF3D7_0102200 | RESA | 27786.8823 | 3.382108993 | 0.75800095 | 4.46187962 | 8.1244E-06 | 0.00086224 | 8832.8789 | 49243.0973 | 895.384546 | 52176.1686 | Sig. genes |
| PF3D7_0712600 | VAR | 154.696898 | 3.350530969 | 0.53838204 | 6.22333352 | 4.867E-10 | 1.6116E-07 | 31.6180094 | 232.755635 | 23.8227178 | 330.59123 | Sig. genes |
| PF3D7_1000400 | RIF | 2.68794358 | 3.331486091 | 2.44344741 | 1.36343679 | 0.17274489 | 0.78315911 | 1.04780684 | 4.29540985 | 0 | 5.40855764 | Non-sig. |
| PF3D7_1040000 | RIF | 2.68420852 | 3.329119037 | 2.45681426 | 1.35505524 | 0.17539996 | 0.78744529 | 1.04752247 | 6.44348011 | 0 | 3.24583148 | Non-sig. |
| PF3D7_0200700 | RIF | 2.68275992 | 3.32787714 | 2.48657841 | 1.33833589 | 0.18078697 | 0.79699299 | 1.04655646 | 7.51908567 | 0 | 2.16539756 | Non-sig. |
| PF3D7_0702000 | N/A | 827.790246 | 3.325777239 | 0.72998409 | 4.5559585 | 5.2147E-06 | 0.00061395 | 267.246392 | 1683.72172 | 33.6704458 | 1326.52243 | Sig. genes |
| PF3D7_0829800 | N/A | 2864.00317 | 3.228717914 | 0.46900598 | 6.88417216 | 5.8125E-12 | 2.3745E-09 | 388.354932 | 4811.47247 | 715.32748 | 5540.85779 | Sig. genes |
| PF3D7_0712800 | VAR | 67.6344179 | 3.193437918 | 0.62550554 | 5.10537112 | 3.3015E-07 | 6.2379E-05 | 12.6473318 | 130.857634 | 13.9650661 | 113.06764 | Sig. genes |
| PF3D7_0831800 | HRP2 | 966.038595 | 3.192153975 | 0.4595747 | 6.94588704 | 3.7609E-12 | 1.8331E-09 | 143.219698 | 1782.00322 | 237.372417 | 1701.55904 | Sig. genes |
| PF3D7_0702300 | STARP | 312.402092 | 3.18043644 | 0.66581758 | 4.77673848 | 1.7816E-06 | 0.00024203 | 104.251081 | 538.576957 | 20.5337937 | 586.246537 | Sig. genes |
| PF3D7_0711700 | VAR | 120.406062 | 3.110292221 | 0.65708317 | 4.73348334 | 2.207E-06 | 0.00028519 | 38.9955449 | 173.762271 | 11.5006224 | 257.365811 | Sig. genes |
| PF3D7_0412900 | VAR | 131.211613 | 3.09499195 | 0.55789041 | 5.54767013 | 2.895E-08 | 6.9717E-06 | 34.7811876 | 263.858284 | 20.5369672 | 205.670012 | Sig. genes |
| PF3D7_0900600 | RIF | 8.5408864 | 2.974073642 | 1.38337812 | 2.14986315 | 0.03156604 | 0.39723726 | 3.14433033 | 13.9656101 | 0.82067433 | 16.2329308 | Non-sig. |
| PF3D7_0401200 | RIF | 2.14714106 | 2.959439338 | 2.62180444 | 1.12877959 | 0.25899083 | 0.85030738 | 1.04771452 | 4.29548893 | 0 | 3.24536079 | Non-sig. |
| PF3D7_0221700 | N/A | 269.10043 | 2.92073452 | 0.83567451 | 3.49506237 | 0.00047395 | 0.02262155 | 117.876903 | 582.652903 | 8.21274226 | 367.659174 | Sig. genes |
| PF3D7_1200700 | ACS7 | 2200.68667 | 2.909515877 | 0.59711539 | 4.8726191 | 1.1013E-06 | 0.00015354 | 818.615753 | 3753.4258 | 216.034047 | 4014.67107 | Sig. genes |
| PF3D7_0412700 | VAR | 128.012664 | 2.902326347 | 0.57917125 | 5.0111713 | 5.41E-07 | 8.4674E-05 | 41.1111922 | 254.194216 | 19.716002 | 197.029247 | Sig. genes |
| PF3D7_0935900 | REX1 | 1819.53833 | 2.862709247 | 0.7125015 | 4.01782909 | 5.8737E-05 | 0.00426284 | 774.333682 | 3031.52527 | 105.962691 | 3366.33168 | Sig. genes |
| PF3D7_0207300 | SERA8 | 85.6024445 | 2.830283474 | 0.57284061 | 4.94078706 | 7.7808E-07 | 0.00011778 | 19.9790859 | 148.114803 | 22.1709303 | 152.144958 | Sig. genes |
| PF3D7_1352900 | N/A | 225.334622 | 2.82814138 | 0.55534556 | 5.09257942 | 3.5322E-07 | 6.2379E-05 | 78.9441105 | 372.328361 | 32.8516007 | 417.214414 | Sig. genes |
| PF3D7_0315200 | CTRP | 472.749512 | 2.773315653 | 0.58559553 | 4.73588934 | 2.181E-06 | 0.00028519 | 184.43005 | 724.018682 | 57.5026593 | 925.046657 | Sig. genes |
| PF3D7_1401100 | N/A | 1616.07888 | 2.756782246 | 0.41446593 | 6.65140864 | 2.903E-11 | 1.0253E-08 | 446.471004 | 2873.15914 | 386.853879 | 2757.83149 | Sig. genes |
| PF3D7_0301100 | N/A | 1.8809471 | 2.739832745 | 2.73635968 | 1.00126923 | 0.31669666 | 0.88585961 | 1.0433895 | 3.22454678 | 0 | 3.2558521 | Non-sig. |
| PF3D7_1479700 | RIF | 1.88302102 | 2.733661556 | 2.80184783 | 0.97566382 | 0.32923109 | 0.88909228 | 1.04724102 | 1.0739736 | 0 | 5.41086947 | Non-sig. |

| | | | | | | | | | | | | |
|---|---|---|---|---|---|---|---|---|---|---|---|---|
| PF3D7_0424800 | N/A | 67.3481893 | 2.729785986 | 0.72733117 | 3.75315415 | 0.00017462 | 0.01051312 | 28.3311069 | 138.537731 | 7.38755065 | 95.1363691 | Sig. genes |
| PF3D7_1407700 | N/A | 44.5078722 | 2.705923137 | 0.82046792 | 3.2980243 | 0.00097368 | 0.03968109 | 19.9286162 | 94.5171167 | 4.10390078 | 59.4818553 | Sig. genes |
| PF3D7_1221400 | IMC1h | 230.614839 | 2.69702061 | 0.47172626 | 5.71734254 | 1.082E-08 | 2.8663E-06 | 59.9672014 | 418.52654 | 63.2337406 | 380.731874 | Sig. genes |
| PF3D7_0408800 | N/A | 8.74701147 | 2.681771571 | 1.31110796 | 2.04542391 | 0.04081308 | 0.45141486 | 3.13959087 | 16.1209674 | 1.64104451 | 14.0864431 | Non-sig. |
| PF3D7_0801300 | WARP | 148.670449 | 2.678988781 | 0.68466481 | 3.9128472 | 9.1214E-05 | 0.00623873 | 66.9194756 | 269.823378 | 13.9468891 | 243.992053 | Sig. genes |
| PF3D7_0632000 | RIF | 3.42930241 | 2.67393564 | 1.98926222 | 1.3441846 | 0.1788887 | 0.79309819 | 1.04817573 | 7.51981197 | 0.82068343 | 4.32853853 | Non-sig. |
| PF3D7_1201300 | LSAP1 | 8.54706051 | 2.666542147 | 1.32589995 | 2.01111867 | 0.04431292 | 0.47237398 | 3.07993183 | 17.3548912 | 1.62951529 | 12.1239038 | Non-sig. |
| PF3D7_0701900 | N/A | 2660.76804 | 2.633547752 | 0.83888172 | 3.13935529 | 0.0016932 | 0.06061199 | 1394.64025 | 4462.73518 | 82.9608195 | 4702.73591 | Non-sig. |
| PF3D7_1115800 | N/A | 3664.66186 | 2.63131269 | 0.60042556 | 4.38241282 | 1.1737E-05 | 0.00119584 | 1620.52272 | 6523.12582 | 417.241846 | 6097.75704 | Sig. genes |
| PF3D7_0902500 | FIKK9.6 | 666.287426 | 2.614373756 | 0.45092069 | 5.79785718 | 6.7168E-09 | 1.8729E-06 | 224.226334 | 1166.30606 | 150.299009 | 1124.31831 | Sig. genes |
| PF3D7_1243300 | N/A | 27.024299 | 2.606037482 | 0.87742958 | 2.97008164 | 0.00297721 | 0.09065079 | 11.4155798 | 57.7728012 | 4.05844586 | 34.8503691 | Non-sig. |
| PF3D7_0301800 | N/A | 1186.03502 | 2.594848708 | 0.5034771 | 5.15385649 | 2.5518E-07 | 5.4078E-05 | 460.836355 | 2241.75094 | 213.519426 | 1828.03335 | Sig. genes |
| PF3D7_1401400 | ETRAMP14 | 3110.5344 | 2.568849662 | 0.50235174 | 5.11364737 | 3.16E-07 | 6.2006E-05 | 1247.43619 | 5330.6744 | 547.728765 | 5316.29822 | Sig. genes |
| PF3D7_1479800 | RIF | 3.16553146 | 2.539351401 | 2.03533323 | 1.24763423 | 0.21216504 | 0.83017014 | 1.04756107 | 4.29774174 | 0.82059814 | 6.49622489 | Non-sig. |
| PF3D7_0700500 | RIF | 3.16306877 | 2.537897948 | 2.03034519 | 1.24998348 | 0.21130558 | 0.82768725 | 1.04788182 | 5.37171641 | 0.82064267 | 5.41203417 | Non-sig. |
| PF3D7_0500400 | RIF | 6.31347084 | 2.533318111 | 1.55593039 | 1.62816931 | 0.10348899 | 0.6615241 | 2.09652255 | 17.1877485 | 1.64139056 | 4.32822178 | Non-sig. |
| PF3D7_1002100 | PTP5 | 2209.43167 | 2.514083058 | 0.42930503 | 5.85616956 | 4.7366E-09 | 1.3942E-06 | 771.666522 | 3710.24614 | 545.351318 | 3810.46268 | Sig. genes |
| PF3D7_1148900 | N/A | 120.348396 | 2.483204816 | 0.58070709 | 4.27617442 | 1.9013E-05 | 0.00170065 | 50.4690058 | 195.344681 | 22.9917186 | 212.588178 | Sig. genes |
| PF3D7_1149600 | N/A | 2033.23576 | 2.481025169 | 0.54753283 | 4.53128112 | 5.8627E-06 | 0.00067523 | 918.254836 | 3414.9182 | 317.8633 | 3481.90669 | Sig. genes |
| PF3D7_0501300 | SBP1 | 1674.19272 | 2.476218915 | 0.53938723 | 4.59080007 | 4.4155E-06 | 0.00053167 | 747.329038 | 2989.36642 | 273.48903 | 2686.5864 | Sig. genes |
| PF3D7_1149200 | RESA3 | 21673.4427 | 2.452373106 | 0.73178906 | 3.35120221 | 0.00080462 | 0.03523018 | 11953.5 | 33672.3987 | 1439.92714 | 39627.9451 | Sig. genes |
| PF3D7_1479100 | N/A | 89.392952 | 2.447873942 | 0.74105882 | 3.3032114 | 0.00095584 | 0.03925625 | 46.8294351 | 179.757863 | 9.01759753 | 121.966912 | Sig. genes |
| PF3D7_0420700 | VAR | 34.1347111 | 2.442372336 | 1.1643372 | 2.09765035 | 0.03593604 | 0.42592653 | 14.7546625 | 3.21784493 | 6.57175254 | 111.994584 | Non-sig. |
| PF3D7_1404800 | N/A | 2671.08521 | 2.442105647 | 0.46027026 | 5.30580807 | 1.1217E-07 | 2.5839E-05 | 1036.71186 | 3920.75927 | 624.281871 | 5102.58783 | Sig. genes |
| PF3D7_0108500 | N/A | 3058.68094 | 2.440882895 | 0.66413853 | 3.67526171 | 0.00023761 | 0.01258835 | 1607.79036 | 5764.69544 | 295.680094 | 4566.55788 | Sig. genes |
| PF3D7_0424600 | N/A | 5168.90345 | 2.436398106 | 0.43699565 | 5.57533714 | 2.4705E-08 | 6.2327E-06 | 1954.68867 | 8994.56266 | 1269.83748 | 8456.52499 | Sig. genes |
| PF3D7_0223300 | N/A | 3.21745597 | 2.424881418 | 2.14847472 | 1.12865253 | 0.25904444 | 0.85030738 | 2.09790613 | 5.36803448 | 0 | 5.40388327 | Non-sig. |
| PF3D7_1312450 | ARA1 | 2.90270214 | 2.400530073 | 2.08903606 | 1.14910897 | 0.25051105 | 0.8494288 | 1.05090745 | 4.38340776 | 0.80885441 | 5.36763895 | Non-sig. |
| PF3D7_1300200 | RIF | 29.8242955 | 2.389674601 | 0.78366345 | 3.04936336 | 0.00229327 | 0.0749984 | 8.38130271 | 56.9435283 | 10.6679596 | 43.3043915 | Non-sig. |
| PF3D7_0900900 | N/A | 2.88969368 | 2.385169469 | 2.13491408 | 1.11722036 | 0.26390012 | 0.85144857 | 1.04600195 | 7.52421516 | 0.82045421 | 2.16810339 | Non-sig. |
| PF3D7_0112100 | PIMMS1 | 24.8944049 | 2.370670003 | 0.84751962 | 2.79718598 | 0.00515498 | 0.12795001 | 6.20600582 | 49.601315 | 9.82609714 | 33.9442017 | Non-sig. |
| PF3D7_0605900 | N/A | 942.656415 | 2.360429394 | 0.47914723 | 4.92631337 | 8.3796E-07 | 0.00012045 | 223.170009 | 1343.35036 | 390.940576 | 1813.16471 | Sig. genes |
| PF3D7_0425400 | N/A | 7.46541391 | 2.357681757 | 1.42268529 | 1.65720541 | 0.09747795 | 0.65042594 | 4.16959672 | 12.9109618 | 0.81999271 | 11.9611045 | Non-sig. |
| PF3D7_0324300 | EPF4 | 188.982343 | 2.352752651 | 0.60154697 | 3.91117033 | 9.185E-05 | 0.00623873 | 88.9380615 | 404.121613 | 35.2813086 | 227.588388 | Sig. genes |
| PF3D7_1029700 | N/A | 41.4947026 | 2.345737688 | 0.69582316 | 3.37116934 | 0.0007485 | 0.0333239 | 11.5718603 | 66.5362131 | 15.602952 | 72.2677848 | Sig. genes |
| PF3D7_0506400 | N/A | 2.96271424 | 2.314968845 | 2.2134181 | 1.0458796 | 0.29561662 | 0.87191643 | 2.04777857 | 5.40196621 | 0 | 4.40111219 | Non-sig. |
| PF3D7_0901500 | RIF | 36.9911835 | 2.29435731 | 0.72512009 | 3.16410667 | 0.0015556 | 0.05683832 | 13.6248534 | 70.9031713 | 11.4893543 | 51.9473549 | Non-sig. |
| PF3D7_1401000 | GBPH | 6.48087951 | 2.287652045 | 1.47548773 | 1.55043786 | 0.12103646 | 0.70312626 | 1.0462602 | 9.67330954 | 3.28194355 | 11.9220047 | Non-sig. |
| PF3D7_0805200 | GAMER | 2003.94472 | 2.266209458 | 0.66792068 | 3.39293202 | 0.00069149 | 0.03113077 | 1170.8524 | 3524.60945 | 209.275543 | 3111.0415 | Sig. genes |
| PF3D7_1150400 | VAR | 57.629348 | 2.257157066 | 0.77489354 | 2.91286087 | 0.00358134 | 0.10092524 | 33.7350403 | 91.1645852 | 6.57209487 | 99.0456714 | Non-sig. |
| PF3D7_0601000 | EPF4 | 66.2130177 | 2.255006856 | 2.80041045 | 0.80524155 | 0.4206803 | 0.90969969 | 0 | 45.0186307 | 45.8229467 | 174.010493 | Non-sig. |
| PF3D7_0201400 | N/A | 4.29729504 | 2.239243047 | 1.90974597 | 1.17253451 | 0.24098253 | 0.84185407 | 3.09523802 | 8.61941172 | 0 | 5.47453042 | Non-sig. |
| PF3D7_1040900 | RIF | 2.62518869 | 2.217482432 | 2.15747041 | 1.02781592 | 0.30403643 | 0.87568674 | 1.04639931 | 4.29908625 | 0.82050423 | 4.33476495 | Non-sig. |
| PF3D7_1040400 | RIF | 2.62408476 | 2.215526141 | 2.1577437 | 1.0267791 | 0.30452449 | 0.87587989 | 1.04766284 | 4.29762478 | 0.82061227 | 4.33043915 | Non-sig. |
| PF3D7_0222600 | RIF | 2.62364876 | 2.214756715 | 2.15785121 | 1.02637138 | 0.30471657 | 0.87595679 | 1.04813212 | 4.29708551 | 0.82067738 | 4.32870002 | Non-sig. |
| PF3D7_0631500 | EPF3 | 11.1619908 | 2.209615644 | 1.17757206 | 1.87641649 | 0.06059811 | 0.53330365 | 2.08464877 | 13.9871558 | 5.73988405 | 22.8362746 | Non-sig. |
| PF3D7_0730800 | N/A | 3903.99137 | 2.202953809 | 0.58825352 | 3.74490544 | 0.00018046 | 0.01054778 | 2185.11733 | 6690.15551 | 602.002904 | 6138.68975 | Sig. genes |
| PF3D7_0702600 | N/A | 12.6508076 | 2.194220863 | 1.33706511 | 1.64107256 | 0.10078235 | 0.65652696 | 9.26083061 | 18.4304726 | 0 | 22.911927 | Non-sig. |
| PF3D7_1102700 | ETRAMP11.1 | 5302.52957 | 2.194062438 | 0.57199968 | 3.83577565 | 0.00012517 | 0.00780169 | 2921.70795 | 9348.03272 | 882.816541 | 8057.56108 | Sig. genes |
| PF3D7_0402100 | N/A | 4354.06502 | 2.182185039 | 0.48957644 | 4.4572918 | 8.3002E-06 | 0.00086224 | 2138.36585 | 6896.56955 | 1006.86097 | 7374.4637 | Sig. genes |
| PF3D7_1100500 | RIF | 20.0614476 | 2.175790986 | 0.90501003 | 2.4041623 | 0.01620957 | 0.27349786 | 6.28141616 | 39.7611489 | 8.20570219 | 25.9975231 | Non-sig. |
| PF3D7_0731500 | EBA175 | 60363.9877 | 2.17062345 | 0.42282718 | 5.13359485 | 2.8426E-07 | 5.7923E-05 | 24181.2409 | 85027.8241 | 19702.7014 | 112544.185 | Sig. genes |
| PF3D7_0202500 | ETRAMP2 | 10880.5003 | 2.15942263 | 0.5254957 | 4.10930598 | 3.9685E-05 | 0.00300359 | 5702.91389 | 19971.3541 | 2258.03638 | 15589.6969 | Sig. genes |
| PF3D7_1201200 | N/A | 2396.51096 | 2.151157889 | 0.41065202 | 5.23839604 | 1.6198E-07 | 3.5757E-05 | 838.231737 | 4145.52536 | 923.201462 | 3679.08527 | Sig. genes |

| | | | | | | | | | | | |
|---|---|---|---|---|---|---|---|---|---|---|---|
| PF3D7_0402900 | N/A | 89.8922552 | 2.14527275 | 0.57247622 | 3.74735698 | 0.00017871 | 0.01054778 | 37.7243107 | 171.894204 | 28.7225697 | 121.227937 Sig. genes |
| PF3D7_0302200 | CLAG3.2 | 2772.12371 | 2.141431293 | 0.51797564 | 4.13423165 | 3.5614E-05 | 0.00279136 | 617.377873 | 3783.47561 | 1430.91429 | 5256.72706 Sig. genes |
| PF3D7_0219700 | GEXP20 | 194.73734 | 2.136944463 | 0.49910188 | 4.28157969 | 1.8557E-05 | 0.00170065 | 87.1400005 | 332.868571 | 57.464318 | 301.476471 Sig. genes |
| PF3D7_0100400 | RIF | 41.5906177 | 2.136083065 | 0.74115049 | 2.88211787 | 0.00395012 | 0.10787491 | 8.38575019 | 71.9745147 | 22.1586136 | 63.8435923 Non-sig. |
| PF3D7_0814500 | N/A | 525.611109 | 2.135293946 | 0.45910328 | 4.65100996 | 3.3031E-06 | 0.00040698 | 231.651564 | 769.258746 | 158.518796 | 943.01533 Sig. genes |
| PF3D7_0102700 | MaTrA | 340.975532 | 2.130169047 | 0.50427782 | 4.22419735 | 2.3979E-05 | 0.00202067 | 167.065942 | 591.512663 | 87.0297809 | 518.293743 Sig. genes |
| PF3D7_1478800 | N/A | 328.849101 | 2.111019499 | 0.4468021 | 4.72473046 | 2.3042E-06 | 0.00029066 | 120.783731 | 551.856515 | 126.430269 | 516.325888 Sig. genes |
| PF3D7_1353200 | MAHRP2 | 155.248386 | 2.110064882 | 0.52099204 | 4.05009044 | 5.1198E-05 | 0.00376731 | 44.1922939 | 252.179544 | 72.2688306 | 252.352874 Sig. genes |
| PF3D7_0925100 | N/A | 51.2422923 | 2.104568574 | 0.65653987 | 3.20554571 | 0.00134807 | 0.050985 | 24.0950831 | 81.6559741 | 14.7708946 | 84.4472175 Non-sig. |
| PF3D7_1249700 | N/A | 90.710989 | 2.094626486 | 0.64259961 | 3.25961371 | 0.00111564 | 0.04444108 | 49.5120349 | 113.711072 | 19.7137684 | 179.907081 Sig. genes |
| PF3D7_1101100 | RIF | 14.0391058 | 2.090273434 | 1.07877896 | 1.93762903 | 0.0526685 | 0.50193661 | 8.38150274 | 20.4135812 | 2.46184724 | 24.899492 Non-sig. |
| PF3D7_1218500 | DRPC | 2143.9741 | 2.076481931 | 0.41444108 | 5.01031879 | 5.434E-07 | 8.4674E-05 | 801.606784 | 3134.63724 | 841.92996 | 3797.72242 Sig. genes |
| PF3D7_0712300 | VAR | 121.741369 | 2.071024695 | 0.51628393 | 4.0114065 | 6.0358E-05 | 0.0043191 | 48.4819979 | 207.011813 | 45.1811868 | 186.29048 Sig. genes |
| PF3D7_0401900 | ACS6 | 139.190289 | 2.060818381 | 0.77652398 | 2.6539018 | 0.0079567 | 0.16929552 | 95.8136312 | 178.102032 | 12.3199817 | 270.525509 Non-sig. |
| PF3D7_0900200 | RIF | 19.0238496 | 2.059365292 | 0.94118268 | 2.18806118 | 0.02866515 | 0.37778095 | 7.33812264 | 41.8946669 | 7.38630985 | 19.476299 Non-sig. |
| PF3D7_0215300 | ACS8 | 354.165023 | 2.049524335 | 0.47898478 | 4.27889243 | 1.8783E-05 | 0.00170065 | 171.717492 | 556.750097 | 104.319278 | 583.873227 Sig. genes |
| PF3D7_0201700 | N/A | 392.7173 | 2.035311804 | 0.49247013 | 4.13286347 | 3.5827E-05 | 0.00279136 | 200.1126 | 605.064838 | 108.421717 | 657.270043 Sig. genes |
| PF3D7_0324400 | RIF | 4.70682973 | 2.02123362 | 1.64094425 | 1.23175033 | 0.21804236 | 0.83527883 | 2.09537569 | 8.59519209 | 1.64123149 | 6.49551964 Non-sig. |
| PF3D7_0702200 | LPL20 | 2826.06953 | 2.01253598 | 0.5664898 | 3.55264295 | 0.00038138 | 0.01870889 | 1713.56397 | 4469.34981 | 532.250343 | 4589.11398 Sig. genes |
| PF3D7_1255200 | VAR | 334.087504 | 2.006021682 | 0.46264515 | 4.33598339 | 1.4511E-05 | 0.00145055 | 123.312958 | 613.527169 | 142.936856 | 456.573033 Sig. genes |
| PF3D7_1350600 | N/A | 99.8833485 | 2.004972262 | 0.5359206 | 3.74117407 | 0.00018316 | 0.01054778 | 38.6373644 | 153.77935 | 41.0121085 | 166.104571 Sig. genes |
| PF3D7_1002000 | N/A | 47.5695731 | 1.993594823 | 0.84127692 | 2.36972484 | 0.01780133 | 0.29198587 | 33.6527952 | 81.5678765 | 4.92697114 | 70.1306495 Non-sig. |
| PF3D7_0814600 | PIMMS22 | 125.132769 | 1.991141988 | 0.51053193 | 3.90013215 | 9.614E-05 | 0.00644748 | 51.3833399 | 199.787365 | 49.2448597 | 200.115509 Sig. genes |
| PF3D7_0201200 | RIF | 10.4829309 | 1.983423423 | 1.16222637 | 1.70657238 | 0.08790156 | 0.62914105 | 5.23929074 | 19.3383015 | 3.28255754 | 14.071574 Non-sig. |
| PF3D7_0623800 | TKL4 | 5944.58975 | 1.968293526 | 0.53793843 | 3.65895691 | 0.00025324 | 0.01328402 | 3478.32349 | 7608.21066 | 1362.09742 | 11329.7274 Sig. genes |
| PF3D7_1312500 | N/A | 1202.12762 | 1.966880499 | 0.55479006 | 3.54526992 | 0.00039221 | 0.01906362 | 727.141349 | 2131.12533 | 252.945343 | 1697.29846 Sig. genes |
| PF3D7_0401400 | RIF | 4.760065 | 1.960843126 | 1.69639297 | 1.15588968 | 0.24772629 | 0.84730586 | 3.14088513 | 10.7460959 | 0.82057775 | 4.33270125 Non-sig. |
| PF3D7_0830500 | TryThrA | 4094.15717 | 1.952388383 | 0.76691203 | 2.54577881 | 0.01090343 | 0.21237628 | 3066.68432 | 6708.46844 | 296.511832 | 6304.96409 Non-sig. |
| PF3D7_0221800 | N/A | 26.138257 | 1.944521889 | 0.80842427 | 2.40532348 | 0.01615816 | 0.27349786 | 12.6165738 | 35.4201745 | 9.0323758 | 47.483904 Non-sig. |
| PF3D7_0900100 | VAR | 524.613941 | 1.944129271 | 0.45403043 | 4.28193607 | 1.8527E-05 | 0.00170065 | 246.626116 | 944.960337 | 186.47512 | 720.394193 Sig. genes |
| PF3D7_1016700 | N/A | 755.814635 | 1.942750759 | 0.48467501 | 4.00835764 | 6.1142E-05 | 0.0043191 | 407.715219 | 1093.10472 | 216.854328 | 1305.58427 Sig. genes |
| PF3D7_0501400 | FIRA | 2513.79821 | 1.941565798 | 0.66327725 | 2.92723111 | 0.00341995 | 0.09793987 | 1753.92943 | 3600.63409 | 323.666277 | 4376.96303 Non-sig. |
| PF3D7_1416500 | GDH1 | 909.765071 | 1.93621152 | 0.43417922 | 4.45947528 | 8.2161E-06 | 0.00086224 | 433.138229 | 1370.68082 | 321.055898 | 1514.18534 Sig. genes |
| PF3D7_0712400 | VAR | 31.1722341 | 1.917343622 | 0.83971926 | 2.28331506 | 0.02241182 | 0.32982723 | 5.26930319 | 48.2682393 | 20.536578 | 50.6148157 Non-sig. |
| PF3D7_1416600 | N/A | 801.369583 | 1.90410352 | 0.57523842 | 3.31011189 | 0.00093259 | 0.03921306 | 511.329475 | 1076.79519 | 165.125724 | 1452.22795 Sig. genes |
| PF3D7_0707500 | N/A | 1532.83349 | 1.90240123 | 0.45093726 | 4.21877139 | 2.4564E-05 | 0.00203342 | 766.126145 | 2036.94257 | 528.19712 | 2800.06814 Sig. genes |
| PF3D7_0413300 | RIF | 6.52731034 | 1.877808648 | 1.41058175 | 1.33122993 | 0.18311337 | 0.80052251 | 3.14283692 | 8.5953657 | 2.46181577 | 11.909223 Non-sig. |
| PF3D7_0828800 | GAMA | 4593.01103 | 1.874758937 | 0.41975212 | 4.46634777 | 7.9566E-06 | 0.00086224 | 2225.58299 | 6722.12521 | 1710.94368 | 7713.39223 Sig. genes |
| PF3D7_0700100 | VAR | 44.2343015 | 1.870673604 | 0.73842138 | 2.53334162 | 0.01129808 | 0.21636979 | 28.4570429 | 72.9366473 | 9.85772663 | 65.6857893 Non-sig. |
| PF3D7_0219900 | N/A | 601.524935 | 1.863898364 | 0.45644133 | 4.08354427 | 4.4354E-05 | 0.00330968 | 315.322037 | 915.633156 | 203.62856 | 971.515988 Sig. genes |
| PF3D7_1355200 | N/A | 65.5037825 | 1.86243243 | 0.60899152 | 3.05822394 | 0.00222653 | 0.07372601 | 33.6993034 | 113.721537 | 22.9980852 | 91.5962047 Non-sig. |
| PF3D7_0320400 | Cap380 | 323.460567 | 1.860492725 | 0.45173038 | 4.11859107 | 3.812E-05 | 0.00292692 | 133.884967 | 462.259021 | 145.407415 | 552.290863 Sig. genes |
| PF3D7_0407600 | N/A | 83.7814706 | 1.855959114 | 0.57002975 | 3.25589868 | 0.00113034 | 0.04469064 | 43.176699 | 125.523265 | 29.5689029 | 136.857016 Sig. genes |
| PF3D7_0223400 | RIF | 23.412137 | 1.853274999 | 0.82943911 | 2.23437138 | 0.02545865 | 0.35967977 | 10.4690269 | 40.8357745 | 9.84684263 | 32.4969039 Non-sig. |
| PF3D7_0701100 | RIF | 5.89046905 | 1.851734515 | 1.69264534 | 1.09398849 | 0.27396003 | 0.85618956 | 5.23262747 | 9.66745667 | 0 | 8.66179208 Non-sig. |
| PF3D7_1147000 | SLARP | 764.523505 | 1.846348325 | 0.43713616 | 4.22373739 | 2.4028E-05 | 0.00202067 | 373.178863 | 1076.82684 | 292.456174 | 1315.63214 Sig. genes |
| PF3D7_1213400 | N/A | 85.785717 | 1.844676875 | 0.60029407 | 3.07295536 | 0.0021195 | 0.07164324 | 50.4412066 | 133.112587 | 24.6316024 | 134.957472 Non-sig. |
| PF3D7_0632800 | VAR | 11.6942768 | 1.844030031 | 1.16020521 | 1.58939989 | 0.11197014 | 0.68406239 | 5.26971939 | 8.58082944 | 4.92884649 | 27.9977118 Non-sig. |
| PF3D7_0731200 | N/A | 619.239074 | 1.840748832 | 0.42649621 | 4.31597937 | 1.589E-05 | 0.00155895 | 259.68524 | 923.077115 | 280.823655 | 1013.37029 Sig. genes |
| PF3D7_1125800 | N/A | 11658.8946 | 1.838727504 | 0.49985991 | 3.67848564 | 0.00023462 | 0.01255588 | 7069.63304 | 19009.0685 | 3120.19673 | 17436.68 Sig. genes |
| PF3D7_0101000 | RIF | 3.097605 | 1.838330917 | 1.96228644 | 0.93683108 | 0.34884546 | 0.8930363 | 1.04790534 | 5.37168263 | 1.64129187 | 4.32954018 Non-sig. |
| PF3D7_0200500 | RIF | 3.09734384 | 1.837932462 | 1.96233941 | 0.93660274 | 0.34896294 | 0.8930363 | 1.04817573 | 5.37129426 | 1.64136685 | 4.32853853 Non-sig. |
| PF3D7_1240100 | ETRAMP12 | 1207.14812 | 1.836063179 | 0.6246444 | 2.93937349 | 0.00328877 | 0.09573559 | 858.451185 | 1966.2972 | 198.648073 | 1805.196 Non-sig. |
| PF3D7_0632100 | RIF | 3.48110397 | 1.829284476 | 2.05379311 | 0.89068586 | 0.37309773 | 0.89654416 | 3.13900829 | 5.37101047 | 0 | 5.41439712 Non-sig. |

| | | | | | | | | | | | | |
|---|---|---|---|---|---|---|---|---|---|---|---|---|
| PF3D7_0731000 | N/A | 576.18588 | 1.824124902 | 0.58307826 | 3.12843921 | 0.00175737 | 0.06146994 | 389.899778 | 914.591673 | 118.233358 | 882.018712 | Non-sig. |
| PF3D7_1240400 | VAR | 814.502693 | 1.820426623 | 0.4377688 | 4.15842024 | 3.2046E-05 | 0.00257239 | 403.701937 | 1409.36049 | 315.45181 | 1129.49653 | Sig. genes |
| PF3D7_0732500 | RIF | 8.33400783 | 1.796894403 | 1.26413368 | 1.42144334 | 0.15518791 | 0.75842353 | 4.1928751 | 15.0394508 | 3.28275757 | 10.8209478 | Non-sig. |
| PF3D7_0408600 | SIAP1 | 207.439476 | 1.793879089 | 0.48367736 | 3.70883414 | 0.00020822 | 0.01158266 | 77.8955982 | 311.163213 | 107.589976 | 333.109115 | Sig. genes |
| PF3D7_1479400 | RIF | 2.08275344 | 1.792222338 | 2.64316766 | 0.67805852 | 0.49773459 | 0.93170321 | 1.05000286 | 6.4587374 | 0.82227348 | 0 | Non-sig. |
| PF3D7_0937600 | VAR | 2.08518109 | 1.791634149 | 2.64251076 | 0.67800449 | 0.49776885 | 0.93170321 | 1.05339504 | 0 | 0.8215004 | 6.46582893 | Non-sig. |
| PF3D7_0502400 | MSP8 | 709.076547 | 1.784147081 | 0.53953899 | 3.30679916 | 0.00094369 | 0.03925625 | 463.035181 | 1112.76314 | 175.74919 | 1084.75868 | Sig. genes |
| PF3D7_1102800 | ETRAMP11.2 | 19973.0398 | 1.781952219 | 0.67812435 | 2.62776617 | 0.00859476 | 0.17927175 | 15449.2648 | 32199.7643 | 2549.47861 | 29693.6516 | Non-sig. |
| PF3D7_0324100 | MC-2TM | 70.7482225 | 1.773160127 | 0.59482669 | 2.98096935 | 0.00287338 | 0.08850955 | 38.8150181 | 113.844684 | 25.444956 | 104.888232 | Non-sig. |
| PF3D7_0718300 | CRMP2 | 89.4061867 | 1.768704994 | 0.57639148 | 3.06858284 | 0.00215077 | 0.07211875 | 48.4868357 | 113.692412 | 32.8596155 | 162.585884 | Non-sig. |
| PF3D7_0625400 | N/A | 1119.99268 | 1.764454151 | 0.4556105 | 3.87272492 | 0.00010763 | 0.00695364 | 613.434524 | 1486.24734 | 405.691328 | 1974.59755 | Sig. genes |
| PF3D7_0424500 | FIKK4.1 | 1109.00757 | 1.758921684 | 0.54297593 | 3.23941003 | 0.00119777 | 0.0463197 | 740.468011 | 1582.36383 | 271.878961 | 1841.3195 | Sig. genes |
| PF3D7_0927200 | N/A | 1086.50218 | 1.754079356 | 0.45474147 | 3.85731115 | 0.00011464 | 0.00729089 | 612.242759 | 1548.90999 | 381.974889 | 1802.88108 | Sig. genes |
| PF3D7_1105600 | PTEX88 | 3623.11959 | 1.751719293 | 0.45900712 | 3.81632274 | 0.00013546 | 0.00834468 | 1230.5072 | 4971.01327 | 2087.21902 | 6203.73888 | Sig. genes |
| PF3D7_1039800 | EPF3 | 28.1498954 | 1.750303701 | 1.19590961 | 1.46357525 | 0.14331004 | 0.74073814 | 26.0409627 | 35.4824931 | 0 | 51.0761259 | Non-sig. |
| PF3D7_0609500 | N/A | 64.4565299 | 1.749795697 | 0.62538445 | 2.79795202 | 0.00514278 | 0.12795001 | 21.0565872 | 90.1255503 | 37.7808532 | 108.863129 | Non-sig. |
| PF3D7_0501100 | HSP40 | 2784.02104 | 1.741910913 | 0.40935306 | 4.25527757 | 2.0879E-05 | 0.00181339 | 1194.89426 | 4436.15333 | 1368.05572 | 4136.98086 | Sig. genes |
| PF3D7_0800400 | RIF | 15.0595488 | 1.740635544 | 0.9940301 | 1.75108938 | 0.07993052 | 0.60514927 | 7.32893736 | 29.0142309 | 6.56465282 | 17.3303741 | Non-sig. |
| PF3D7_0315900 | N/A | 4.92036463 | 1.728714137 | 1.58438341 | 1.09109584 | 0.27523072 | 0.85802262 | 2.09010208 | 6.45092753 | 2.4610005 | 8.67942842 | Non-sig. |
| PF3D7_1026500 | N/A | 315.578242 | 1.727110229 | 0.48082514 | 3.59197158 | 0.00032819 | 0.01655931 | 181.742062 | 479.261264 | 111.45126 | 489.858383 | Sig. genes |
| PF3D7_1219700 | RKIP | 37.4179271 | 1.724874878 | 0.72887622 | 2.36648533 | 0.01795789 | 0.29274117 | 21.8082125 | 68.9280024 | 13.1118158 | 45.8236776 | Non-sig. |
| PF3D7_1216600 | CelTOS | 91.5889157 | 1.723679441 | 0.79376673 | 2.17151887 | 0.02989197 | 0.38815605 | 75.7315756 | 149.747406 | 9.83206324 | 131.044618 | Non-sig. |
| PF3D7_1114200 | N/A | 830.16875 | 1.721901106 | 0.436721 | 3.94279441 | 8.0538E-05 | 0.00561432 | 410.424379 | 1084.8585 | 362.175788 | 1463.21633 | Sig. genes |
| PF3D7_0411700 | N/A | 2.14631627 | 1.713898408 | 2.49012211 | 0.68827886 | 0.49127719 | 0.93170321 | 2.05540482 | 3.23547614 | 0 | 3.29438413 | Non-sig. |
| PF3D7_0202100 | LSAP2 | 7033.35463 | 1.70502735 | 0.46021201 | 3.70487364 | 0.0002115 | 0.01158266 | 4229.55899 | 11437.1001 | 2374.47816 | 10092.2812 | Sig. genes |
| PF3D7_1455300 | N/A | 4332.106 | 1.681283943 | 0.43609909 | 3.85527967 | 0.0001156 | 0.00729089 | 2462.47775 | 6084.9508 | 1656.72134 | 7124.27411 | Sig. genes |
| PF3D7_1000500 | RIF | 25.7544285 | 1.675353696 | 0.85558515 | 1.95813788 | 0.05021384 | 0.4954291 | 6.28892398 | 44.0448689 | 18.0549691 | 34.6289521 | Non-sig. |
| PF3D7_0600500 | RIF | 2.13860887 | 1.673469834 | 2.50760438 | 0.667358 | 0.50454349 | 0.93170321 | 2.0956591 | 4.2953904 | 0 | 2.16338596 | Non-sig. |
| PF3D7_1226100 | HAD3 | 404.71533 | 1.669024682 | 0.44436071 | 3.75601316 | 0.00017264 | 0.01051312 | 189.341092 | 682.549615 | 197.908086 | 549.062526 | Sig. genes |
| PF3D7_0911100 | N/A | 20608.1451 | 1.65337583 | 0.46856778 | 3.52857345 | 0.00041781 | 0.02012306 | 13053.5237 | 31258.935 | 6830.9401 | 31289.1815 | Sig. genes |
| PF3D7_1345100 | TRX2 | 396.804341 | 1.652792553 | 0.460919 | 3.58586333 | 0.00033596 | 0.01679191 | 224.913083 | 550.327846 | 158.349595 | 653.626842 | Sig. genes |
| PF3D7_1427100 | N/A | 1175.33757 | 1.642611139 | 0.54101263 | 3.03617893 | 0.00239597 | 0.07610251 | 835.476573 | 1757.10301 | 305.569317 | 1803.2014 | Non-sig. |
| PF3D7_0424900 | N/A | 357.040227 | 1.640805647 | 0.44967178 | 3.64889621 | 0.00026337 | 0.01367972 | 177.671127 | 602.154671 | 169.149713 | 479.185396 | Sig. genes |
| PF3D7_1422900 | N/A | 206.327101 | 1.637897333 | 0.66703616 | 2.45548506 | 0.01406946 | 0.25182432 | 165.088914 | 328.429276 | 36.1303502 | 295.659865 | Non-sig. |
| PF3D7_0700200 | RIF | 13.4885723 | 1.63654737 | 1.08013932 | 1.51512619 | 0.12974045 | 0.71812182 | 3.14034497 | 24.717169 | 9.84666708 | 16.2501082 | Non-sig. |
| PF3D7_0931600 | N/A | 11.8152965 | 1.634248997 | 1.12020242 | 1.45888723 | 0.14459615 | 0.74131203 | 8.34959082 | 21.5108204 | 3.28067679 | 14.1200981 | Non-sig. |
| PF3D7_0917200 | N/A | 12.5070976 | 1.633549975 | 1.0585983 | 1.54312544 | 0.12280034 | 0.7079393 | 7.32768641 | 17.1944112 | 4.92335769 | 20.5829349 | Non-sig. |
| PF3D7_0818600 | PBLP | 888.088709 | 1.632913715 | 0.4214782 | 3.87425428 | 0.00010695 | 0.00695364 | 440.081382 | 1452.64168 | 426.087873 | 1233.54391 | Sig. genes |
| PF3D7_0323400 | RIPR | 15444.2683 | 1.62889597 | 0.41887884 | 3.88870439 | 0.00010078 | 0.00667421 | 6405.2536 | 23923.035 | 8688.66346 | 22760.1209 | Sig. genes |
| PF3D7_0600600 | VAR | 5.71071773 | 1.625160103 | 1.47662005 | 1.10059463 | 0.27107312 | 0.8556331 | 3.14823938 | 9.66513139 | 2.46256361 | 7.56693655 | Non-sig. |
| PF3D7_0304600 | CSP | 254.827785 | 1.624596799 | 0.51217986 | 3.17192635 | 0.00151431 | 0.05571414 | 162.985587 | 417.513237 | 87.0412718 | 351.771043 | Non-sig. |
| PF3D7_0424700 | FIKK4.2 | 688.753346 | 1.612673504 | 0.4751541 | 3.394001 | 0.00068879 | 0.03113077 | 427.921594 | 909.555719 | 251.373255 | 1166.16281 | Sig. genes |
| PF3D7_0532900 | RIF | 3.95431667 | 1.611641039 | 1.78949251 | 0.90061346 | 0.36779388 | 0.89615989 | 3.13987604 | 7.52291789 | 0.82053659 | 4.33393616 | Non-sig. |
| PF3D7_1353100 | N/A | 200.010387 | 1.600179296 | 0.51682991 | 3.09614301 | 0.00196056 | 0.06744828 | 79.9272985 | 236.115951 | 118.247802 | 365.750496 | Non-sig. |
| PF3D7_0400900 | RIF | 3.57226343 | 1.598985064 | 1.83804441 | 0.86993821 | 0.38433417 | 0.90292925 | 1.04696171 | 6.44765752 | 2.46173326 | 4.33270125 | Non-sig. |
| PF3D7_0827700 | N/A | 11.8620403 | 1.597339932 | 1.15075984 | 1.3880741 | 0.16511448 | 0.77284241 | 9.47284529 | 13.9490867 | 2.46384969 | 21.5623794 | Non-sig. |
| PF3D7_0605700 | N/A | 242.281475 | 1.583590358 | 0.53719957 | 2.94786231 | 0.0031998 | 0.09470679 | 164.122782 | 417.125466 | 78.7557133 | 309.121938 | Non-sig. |
| PF3D7_0926600 | N/A | 231.46143 | 1.579646572 | 0.47579143 | 3.32004 | 0.00090005 | 0.03845517 | 133.771439 | 329.344377 | 98.5667352 | 364.163168 | Sig. genes |
| PF3D7_0613600 | N/A | 3709.1296 | 1.578712473 | 0.47685094 | 3.31070436 | 0.00093061 | 0.03921306 | 1255.65592 | 5826.15329 | 2465.02412 | 5289.68508 | Sig. genes |
| PF3D7_0814700 | N/A | 74.9867568 | 1.571626788 | 0.72733766 | 2.16079393 | 0.03071126 | 0.39012163 | 61.9685001 | 107.364499 | 13.9566355 | 116.657392 | Non-sig. |
| PF3D7_1373400 | RIF | 9.70778989 | 1.568458116 | 1.18930335 | 1.31880408 | 0.18723462 | 0.80191512 | 3.14497205 | 12.8905951 | 6.56563179 | 16.2299606 | Non-sig. |
| PF3D7_1370900 | N/A | 2603.90198 | 1.567429876 | 0.76623352 | 2.04562947 | 0.04079284 | 0.45141486 | 2393.77633 | 4074.39093 | 234.412771 | 3713.02788 | Non-sig. |
| PF3D7_1400300 | RIF | 32.4221409 | 1.56070276 | 0.76400372 | 2.04279472 | 0.04107276 | 0.45239813 | 10.475091 | 50.4991672 | 22.1559548 | 46.5583507 | Non-sig. |
| PF3D7_0600700 | RIF | 6.8903155 | 1.556903828 | 1.50043794 | 1.03763294 | 0.29944098 | 0.87358937 | 6.28612706 | 15.0415861 | 0.82061575 | 5.41293304 | Non-sig. |

| | | | | | | | | | | | | |
|---|---|---|---|---|---|---|---|---|---|---|---|---|
| PF3D7_0424200 | RH4 | 6237.08386 | 1.556729456 | 0.74849228 | 2.07982034 | 0.03754202 | 0.43374033 | 5710.2286 | 8938.48838 | 619.371063 | 9680.24739 | Non-sig. |
| PF3D7_0936300 | REX3 | 223.827904 | 1.556698468 | 0.5566881 | 2.79635663 | 0.00516823 | 0.12795001 | 161.081733 | 358.35708 | 66.5270709 | 309.345731 | Non-sig. |
| PF3D7_0502500 | N/A | 12.6140479 | 1.534792712 | 1.13560996 | 1.35151396 | 0.17653085 | 0.78924933 | 2.10198931 | 18.2486419 | 10.6739373 | 19.431623 | Non-sig. |
| PF3D7_0936000 | REX2 | 252.462043 | 1.529430331 | 0.48708738 | 3.13995066 | 0.00168976 | 0.06061199 | 159.966244 | 380.930488 | 100.19441 | 368.757029 | Non-sig. |
| PF3D7_0909200 | POFUT2 | 48.5809164 | 1.527286632 | 0.64269341 | 2.37638444 | 0.01748324 | 0.2885551 | 22.0458387 | 68.7226807 | 27.9108764 | 75.6442698 | Non-sig. |
| PF3D7_1351700 | IMC1f | 23806.516 | 1.522817589 | 0.40804107 | 3.73202039 | 0.00018995 | 0.01070591 | 11083.2052 | 34607.3263 | 13500.4704 | 36035.0622 | Sig. genes |
| PF3D7_1205500 | N/A | 7989.33905 | 1.517681285 | 0.4433238 | 3.42341487 | 0.0006184 | 0.02848924 | 3345.28081 | 10126.3977 | 4926.3982 | 13559.2795 | Sig. genes |
| PF3D7_0600200 | VAR | 10.1884087 | 1.51371727 | 1.16560878 | 1.29864951 | 0.19406424 | 0.81267745 | 7.37913322 | 15.0155767 | 3.28601592 | 15.0729088 | Non-sig. |
| PF3D7_1418500 | N/A | 14.2314243 | 1.513532012 | 4.58216574 | 0.33030931 | 0.74116626 | 0.98954916 | 0 | 42.174961 | 14.750736 | 0 | Non-sig. |
| PF3D7_1254600 | N/A | 4.38121678 | 1.506019118 | 1.66722606 | 0.90330829 | 0.3663623 | 0.89567376 | 2.09215994 | 4.29946857 | 2.46138564 | 8.67185297 | Non-sig. |
| PF3D7_1404300 | SOAP | 32.6168806 | 1.501753679 | 0.72369886 | 2.07510853 | 0.03797648 | 0.43374033 | 17.6774797 | 48.4480876 | 16.393547 | 47.9484079 | Non-sig. |
| PF3D7_0720300 | N/A | 37.792003 | 1.499321093 | 0.89329044 | 1.6784251 | 0.09326414 | 0.64025761 | 34.9161433 | 66.190894 | 4.88685746 | 45.1741172 | Non-sig. |
| PF3D7_0307300 | EB1 | 96.3255731 | 1.498736149 | 0.55360415 | 2.7072343 | 0.00678464 | 0.15295744 | 59.8651636 | 128.839036 | 41.0486012 | 155.549491 | Non-sig. |
| PF3D7_0805100 | N/A | 513.896912 | 1.48799776 | 0.43146884 | 3.44867958 | 0.00056333 | 0.02641192 | 279.959394 | 804.767787 | 260.344214 | 710.516253 | Sig. genes |
| PF3D7_1102200 | ECIJP | 106.426702 | 1.476239816 | 0.52643466 | 2.8042223 | 0.00504381 | 0.12724816 | 52.5389315 | 167.467797 | 59.9360992 | 145.763979 | Non-sig. |
| PF3D7_0723200 | N/A | 23.8611489 | 1.474162906 | 0.81550904 | 1.80765979 | 0.07065947 | 0.58041649 | 14.6706748 | 33.3044323 | 10.6683971 | 36.8010916 | Non-sig. |
| PF3D7_1040700 | RIF | 3.68676302 | 1.471776056 | 1.82444191 | 0.80669933 | 0.41983973 | 0.90957507 | 3.14465632 | 5.37123244 | 0.82068939 | 5.41047392 | Non-sig. |
| PF3D7_0808200 | PMX | 10218.1918 | 1.468265335 | 0.42716535 | 3.43722949 | 0.0005877 | 0.02731246 | 6380.27537 | 15866.634 | 4470.60515 | 14155.2529 | Sig. genes |
| PF3D7_1430500 | N/A | 94.55358 | 1.467567971 | 0.54739241 | 2.68101632 | 0.00733989 | 0.16135584 | 41.977673 | 141.754173 | 58.2809532 | 136.201521 | Non-sig. |
| PF3D7_1408300 | N/A | 23.7190607 | 1.458867861 | 0.81320216 | 1.79397932 | 0.07281647 | 0.58363337 | 11.35006 | 36.6919729 | 13.9116677 | 32.9225422 | Non-sig. |
| PF3D7_0410000 | EVP1 | 4045.54152 | 1.457792137 | 0.40729594 | 3.57919637 | 0.00034465 | 0.01706513 | 2261.37734 | 5593.55824 | 2057.56516 | 6269.66535 | Sig. genes |
| PF3D7_0932900 | N/A | 24.6362274 | 1.455546483 | 0.81639092 | 1.78290381 | 0.07460196 | 0.59167844 | 15.7540932 | 41.8726102 | 10.6726485 | 30.2455578 | Non-sig. |
| PF3D7_0423900 | N/A | 334.379542 | 1.44906183 | 0.54361187 | 2.66561845 | 0.00768469 | 0.16526913 | 254.862467 | 521.524754 | 104.174606 | 456.956341 | Non-sig. |
| PF3D7_0414900 | ARO | 6107.41942 | 1.44844614 | 0.43536338 | 3.32698201 | 0.00087792 | 0.03781482 | 3949.14932 | 9367.89703 | 2602.25494 | 8510.37637 | Sig. genes |
| PF3D7_0200400 | N/A | 3.29982687 | 1.44790592 | 1.93467034 | 0.7483993 | 0.45421935 | 0.92203286 | 1.04615601 | 7.52391166 | 2.46140993 | 2.16782989 | Non-sig. |
| PF3D7_0800300 | VAR | 251.495093 | 1.447049248 | 0.45861869 | 3.1552339 | 0.00160369 | 0.05819435 | 140.217825 | 390.393782 | 129.79993 | 345.568835 | Non-sig. |
| PF3D7_1371000 | N/A | 41.8277333 | 1.446119559 | 0.72801405 | 1.98638963 | 0.04699008 | 0.48052729 | 12.6206504 | 62.2479873 | 32.0256913 | 60.4166041 | Non-sig. |
| PF3D7_0505300 | N/A | 548.238968 | 1.442038456 | 0.43180903 | 3.33952824 | 0.00083921 | 0.03644365 | 319.933391 | 773.663842 | 270.196533 | 829.162105 | Sig. genes |
| PF3D7_1244500 | PIMMS57 | 89.1733904 | 1.440016379 | 0.5738515 | 2.50938854 | 0.01209404 | 0.22637855 | 61.0229239 | 127.702094 | 35.3127199 | 132.655823 | Non-sig. |
| PF3D7_1417000 | IMC1l | 46.9858559 | 1.434277243 | 0.68718111 | 2.08718957 | 0.036871 | 0.43217382 | 34.5980579 | 65.5254049 | 16.4143619 | 71.4055989 | Non-sig. |
| PF3D7_0831500 | N/A | 1835.70501 | 1.434076144 | 0.56069042 | 2.55769689 | 0.01053679 | 0.20675521 | 1494.45691 | 2889.11446 | 489.536582 | 2469.71208 | Non-sig. |
| PF3D7_1000100 | VAR | 1096.99964 | 1.414599286 | 0.43211943 | 3.2736304 | 0.00106175 | 0.04292299 | 694.549885 | 1626.06661 | 502.742451 | 1564.63961 | Sig. genes |
| PF3D7_0300500 | RIF | 1.87016816 | 1.407740487 | 2.63035264 | 0.53519078 | 0.59251794 | 0.95322751 | 2.09504494 | 3.22174006 | 0 | 2.16388766 | Non-sig. |
| PF3D7_1310400 | N/A | 1918.81756 | 1.406976445 | 0.56981681 | 2.46917328 | 0.01354256 | 0.24576103 | 1603.72616 | 2580.85503 | 498.614593 | 2992.07448 | Non-sig. |
| PF3D7_1342600 | MyoA | 31953.5556 | 1.406548463 | 0.43234987 | 3.25326445 | 0.00114087 | 0.04477292 | 21025.3396 | 44217.2146 | 13982.8513 | 48588.8168 | Sig. genes |
| PF3D7_1100400 | RIF | 1.86809 | 1.406302692 | 2.67910482 | 0.52491514 | 0.59964218 | 0.95655566 | 2.09267219 | 4.29680838 | 0 | 1.08287942 | Non-sig. |
| PF3D7_0617700 | RIF | 6.36401806 | 1.40079984 | 1.50659583 | 0.92977812 | 0.35248598 | 0.8930363 | 6.28212684 | 7.52214707 | 0.82058517 | 10.8312132 | Non-sig. |
| PF3D7_1133400 | AMA1 | 21789.1908 | 1.400775761 | 0.43515011 | 3.21906331 | 0.0012861 | 0.04901987 | 9482.44733 | 33414.5172 | 14458.4847 | 29801.3138 | Sig. genes |
| PF3D7_1230100 | TLAP1 | 307.439943 | 1.400251797 | 0.46095477 | 3.03772059 | 0.00238375 | 0.07610251 | 182.845722 | 388.605055 | 155.178551 | 503.130446 | Non-sig. |
| PF3D7_0400400 | VAR | 45.230974 | 1.400127318 | 0.6563306 | 2.13326532 | 0.03290297 | 0.40615121 | 28.4658231 | 63.2777682 | 21.359554 | 67.8207508 | Non-sig. |
| PF3D7_1230700 | SEC13 | 4196.99985 | 1.394452286 | 0.57141408 | 2.4403534 | 0.0146729 | 0.25826254 | 1069.96488 | 6178.49305 | 3555.6752 | 5983.86626 | Non-sig. |
| PF3D7_0112900 | N/A | 324.848533 | 1.387031191 | 0.50849103 | 2.72773975 | 0.00637699 | 0.14818111 | 239.101637 | 489.189988 | 120.744106 | 450.358399 | Non-sig. |
| PF3D7_0209500 | N/A | 905.023324 | 1.379123206 | 0.78227905 | 1.76295557 | 0.07790798 | 0.60344518 | 920.356623 | 1469.69353 | 85.3863703 | 1144.65677 | Non-sig. |
| PF3D7_1370300 | MAHRP1 | 121.839482 | 1.3769056 | 0.5362812 | 2.56750677 | 0.01024328 | 0.20249586 | 63.1350761 | 137.355393 | 72.2694769 | 214.597981 | Non-sig. |
| PF3D7_0500500 | RIF | 4.85465848 | 1.375301034 | 1.58242666 | 0.86910886 | 0.38478759 | 0.90292925 | 2.09266027 | 5.37395762 | 3.28198219 | 8.67003383 | Non-sig. |
| PF3D7_1017500 | ELC | 4567.98501 | 1.36879103 | 0.43883518 | 3.11914608 | 0.00181376 | 0.06280589 | 3069.04704 | 6018.6672 | 2031.59303 | 7152.63275 | Non-sig. |
| PF3D7_1131700 | N/A | 12.9054932 | 1.363404552 | 1.0944396 | 1.24575587 | 0.21285403 | 0.83163767 | 11.3189641 | 20.5281913 | 3.2698281 | 16.5049892 | Non-sig. |
| PF3D7_1125700 | N/A | 9724.77413 | 1.359969937 | 0.61677528 | 2.2049683 | 0.02745632 | 0.3691969 | 8817.86137 | 15320.5822 | 2088.59776 | 12672.0552 | Non-sig. |
| PF3D7_0421700 | N/A | 132.817638 | 1.35609184 | 0.79870983 | 1.69785295 | 0.08953552 | 0.62981332 | 134.885062 | 180.20485 | 14.7861625 | 201.394477 | Non-sig. |
| PF3D7_0507400 | SUB1-ProM | 722.433532 | 1.351233473 | 0.43144166 | 3.13190308 | 0.00173677 | 0.06146994 | 357.039892 | 1075.84872 | 456.449071 | 1000.39644 | Non-sig. |
| PF3D7_0611800 | N/A | 7992.45088 | 1.345976948 | 0.40724756 | 3.30505838 | 0.00094957 | 0.03925625 | 4366.93535 | 10543.7465 | 4658.85981 | 12400.2619 | Sig. genes |
| PF3D7_1436300 | PTEX150 | 8885.0481 | 1.336792422 | 0.64291702 | 2.07926122 | 0.03759335 | 0.43374033 | 8280.44833 | 9968.55302 | 1799.87561 | 15491.3155 | Non-sig. |
| PF3D7_1016000 | N/A | 2460.07255 | 1.335453829 | 0.41685197 | 3.20366442 | 0.00135691 | 0.050985 | 1563.44035 | 3453.74683 | 1229.51883 | 3593.5842 | Non-sig. |
| PF3D7_1340900 | PiT | 3012.71061 | 1.335282559 | 0.47200013 | 2.82898772 | 0.00466955 | 0.12247164 | 1276.50255 | 3493.88408 | 2143.48275 | 5136.97305 | Non-sig. |

| | | | | | | | | | | | |
|---|---|---|---|---|---|---|---|---|---|---|---|
| PF3D7_0318600 | N/A | 1576.6099 | 1.334162927 | 0.41974166 | 3.17853347 | 0.00148022 | 0.05484065 | 1001.80808 | 2220.59697 | 789.36932 | 2294.66526 | Non-sig. |
| PF3D7_0620000 | PIMMS43 | 89.7298024 | 1.326333912 | 0.64395681 | 2.05966284 | 0.03943078 | 0.44209722 | 75.6154076 | 144.945932 | 27.0918406 | 111.266029 | Non-sig. |
| PF3D7_0801900 | LSD2 | 3003.65224 | 1.326062495 | 0.70093453 | 1.89184928 | 0.05851106 | 0.52454723 | 2980.1426 | 3707.39587 | 446.104801 | 4880.96568 | Non-sig. |
| PF3D7_1146100 | N/A | 153.215508 | 1.322375001 | 0.71035815 | 1.86156097 | 0.062665 | 0.54515462 | 146.788914 | 263.171145 | 28.7254544 | 174.176518 | Non-sig. |
| PF3D7_1224300 | PABP1 | 24772.3412 | 1.317519026 | 0.40892152 | 3.22193612 | 0.00127328 | 0.0488827 | 12950.2511 | 33237.8334 | 15422.6709 | 37478.6096 | Sig. genes |
| PF3D7_1318900 | N/A | 133.196186 | 1.314580072 | 0.57459778 | 2.28782658 | 0.02214762 | 0.32776007 | 106.359098 | 179.167114 | 46.8171982 | 200.441332 | Non-sig. |
| PF3D7_0524200 | N/A | 192.908653 | 1.312970143 | 0.50441958 | 2.60293257 | 0.00924301 | 0.18834417 | 129.772538 | 322.416836 | 91.8987522 | 227.546485 | Non-sig. |
| PF3D7_1328600 | N/A | 99.1055554 | 1.31292675 | 0.6153722 | 2.13354903 | 0.03287971 | 0.40615121 | 79.6495656 | 168.665636 | 34.4678006 | 113.639219 | Non-sig. |
| PF3D7_0628300 | CEPT | 15569.0733 | 1.311974026 | 0.47894666 | 2.73929047 | 0.00615719 | 0.1443399 | 5907.06065 | 22316.0715 | 11973.4886 | 22079.6726 | Non-sig. |
| PF3D7_0924900 | ATAT | 3.79007418 | 1.306938675 | 1.78159457 | 0.73357805 | 0.46320594 | 0.92509266 | 1.0376955 | 5.38620709 | 3.27749333 | 5.45890081 | Non-sig. |
| PF3D7_0314100 | N/A | 1175.29207 | 1.305087119 | 0.61423829 | 2.12472447 | 0.03360961 | 0.41314087 | 1084.8638 | 1832.76423 | 270.097123 | 1513.44311 | Non-sig. |
| PF3D7_0800800 | N/A | 107.184599 | 1.303660612 | 0.55530917 | 2.34763027 | 0.01889327 | 0.30332278 | 78.7474924 | 149.250499 | 45.1511948 | 155.589209 | Non-sig. |
| PF3D7_1477800 | ACBP | 1065.39528 | 1.303226467 | 0.74567137 | 1.74772228 | 0.08051213 | 0.6065657 | 1102.16594 | 1650.91187 | 127.216821 | 1381.2865 | Non-sig. |
| PF3D7_0931300 | N/A | 247.092819 | 1.302595362 | 0.56665283 | 2.29875383 | 0.02151892 | 0.3235449 | 201.009134 | 426.624112 | 84.5190013 | 276.219029 | Non-sig. |
| PF3D7_0812300 | SSP3 | 133.970385 | 1.300735293 | 0.52305522 | 2.48680301 | 0.01288967 | 0.2379425 | 60.8826127 | 180.401938 | 93.5819302 | 201.015058 | Non-sig. |
| PF3D7_1411000 | N/A | 7852.84716 | 1.297578783 | 0.60656247 | 2.13923355 | 0.03241676 | 0.40505657 | 7283.31528 | 11049.0763 | 1800.56151 | 11278.4356 | Non-sig. |
| PF3D7_0808700 | VAR | 747.06066 | 1.287475817 | 0.43232277 | 2.97804304 | 0.00290095 | 0.08883957 | 398.389617 | 1173.4294 | 469.883125 | 946.540502 | Non-sig. |
| PF3D7_0918700 | N/A | 2608.31854 | 1.286253902 | 0.73180636 | 1.75764242 | 0.07880837 | 0.60514927 | 2705.85756 | 3740.58625 | 328.508654 | 3658.32169 | Non-sig. |
| PF3D7_1001500 | ETRAMP10 | 593.354623 | 1.281907921 | 0.43671364 | 2.93535121 | 0.0033317 | 0.0959571 | 391.310268 | 867.135174 | 300.558227 | 814.414825 | Non-sig. |
| PF3D7_1008600 | N/A | 84.1560128 | 1.281716678 | 0.5689838 | 2.25264179 | 0.02428174 | 0.34675116 | 59.7484701 | 114.944374 | 38.5724014 | 123.358805 | Non-sig. |
| PF3D7_0909500 | SPM1 | 452.94114 | 1.279995379 | 0.43127923 | 2.96790405 | 0.00299838 | 0.09077379 | 259.887836 | 654.597208 | 268.543511 | 628.736003 | Non-sig. |
| PF3D7_1030200 | CLAMP | 7181.22523 | 1.278733065 | 0.42036017 | 3.04199389 | 0.00235017 | 0.07610251 | 3648.67829 | 11049.3282 | 4734.82942 | 9292.06501 | Non-sig. |
| PF3D7_1131900 | N/A | 51.4213352 | 1.276767067 | 0.70169952 | 1.81953532 | 0.0688298 | 0.57367529 | 43.1314795 | 60.0972247 | 17.245368 | 85.2112686 | Non-sig. |
| PF3D7_1370800 | N/A | 631.130681 | 1.271691101 | 0.80276966 | 1.5841295 | 0.11316425 | 0.68498781 | 684.006293 | 940.150672 | 55.7999344 | 844.565824 | Non-sig. |
| PF3D7_0811300 | CAF1 | 17879.657 | 1.26467638 | 0.44538159 | 2.83953446 | 0.00451794 | 0.12028168 | 9011.78004 | 20092.7331 | 12005.9784 | 30408.1363 | Non-sig. |
| PF3D7_1114100 | ROM1 | 2660.80103 | 1.262637975 | 0.42356767 | 2.9809593 | 0.00287347 | 0.08850955 | 1698.41952 | 4245.69918 | 1432.67799 | 3266.40743 | Non-sig. |
| PF3D7_1209400 | CIA1 | 5703.2633 | 1.25689874 | 0.42554494 | 2.95362163 | 0.00314069 | 0.09347958 | 3949.09029 | 7885.43476 | 2781.11984 | 8197.40833 | Non-sig. |
| PF3D7_0301000 | ACS2 | 6.13816019 | 1.254150762 | 1.72651049 | 0.72640784 | 0.46758878 | 0.92538923 | 7.36394959 | 12.8754557 | 0 | 4.31323545 | Non-sig. |
| PF3D7_0421300 | VAR | 194.088493 | 1.252677724 | 0.47776968 | 2.62192805 | 0.00874339 | 0.18165676 | 101.176671 | 275.660636 | 128.149581 | 271.367084 | Non-sig. |
| PF3D7_0308400 | N/A | 10.142741 | 1.251194748 | 1.14317385 | 1.0944921 | 0.27373921 | 0.85618956 | 7.13509625 | 16.6063154 | 4.90459741 | 11.9249551 | Non-sig. |
| PF3D7_0201000 | RIF | 3.46755368 | 1.247298688 | 2.10224748 | 0.59331677 | 0.55296918 | 0.94170064 | 4.18387046 | 7.51998749 | 0 | 2.16635678 | Non-sig. |
| PF3D7_1413100 | N/A | 17.6570396 | 1.246413547 | 1.04837375 | 1.1889019 | 0.23447827 | 0.83828834 | 17.8641924 | 32.2048615 | 3.28420068 | 17.2749039 | Non-sig. |
| PF3D7_1041300 | VAR | 2014.20796 | 1.246284992 | 0.42088273 | 2.96112172 | 0.00306521 | 0.09174842 | 1125.59095 | 3180.28671 | 1263.42352 | 2487.53064 | Non-sig. |
| PF3D7_1252200 | CHT1 | 294.935743 | 1.24154816 | 0.44954719 | 2.76177491 | 0.00574881 | 0.13844179 | 186.605351 | 421.090448 | 164.141367 | 407.905806 | Non-sig. |
| PF3D7_0829900 | N/A | 7898.20185 | 1.231722597 | 0.58598217 | 2.10197966 | 0.03555506 | 0.42313505 | 7389.64864 | 11358.0347 | 2045.90756 | 10799.2165 | Non-sig. |
| PF3D7_1033800 | N/A | 57.268528 | 1.229782845 | 0.62058907 | 1.98163793 | 0.04751978 | 0.48229845 | 39.8830401 | 90.2045029 | 28.7305874 | 70.2559816 | Non-sig. |
| PF3D7_1420500 | N/A | 48.3214957 | 1.228616154 | 0.64020107 | 1.91910982 | 0.05497044 | 0.50446754 | 27.3788262 | 60.0804497 | 30.389967 | 75.4367397 | Non-sig. |
| PF3D7_1310300 | N/A | 323.765645 | 1.227188674 | 0.4742664 | 2.58755136 | 0.00966608 | 0.19600817 | 156.607845 | 402.536608 | 230.723753 | 505.194372 | Non-sig. |
| PF3D7_0113700 | HSP40 | 439.683221 | 1.224874738 | 0.47431473 | 2.58240923 | 0.00981132 | 0.19689529 | 329.428827 | 651.32377 | 197.927612 | 580.052676 | Non-sig. |
| PF3D7_0109000 | PHIL1 | 4435.21375 | 1.223094634 | 0.46317248 | 2.64068933 | 0.00827376 | 0.17325832 | 1954.05809 | 5414.53225 | 3365.97042 | 7006.29426 | Non-sig. |
| PF3D7_0421900 | N/A | 8.20187942 | 1.220844126 | 1.32868725 | 0.91883483 | 0.35818197 | 0.89552149 | 8.35416398 | 12.0126016 | 1.62087828 | 10.8198738 | Non-sig. |
| PF3D7_1478900 | N/A | 26028.2755 | 1.219721732 | 0.41772533 | 2.91991326 | 0.00350129 | 0.09973024 | 17747.3704 | 34018.5123 | 13527.3241 | 38819.8952 | Non-sig. |
| PF3D7_0420900 | VAR | 193.371203 | 1.217075662 | 0.5133204 | 2.37098636 | 0.01774069 | 0.29189488 | 147.549823 | 291.749493 | 85.4331053 | 248.75239 | Non-sig. |
| PF3D7_0701800 | RIF | 3.09364913 | 1.215154908 | 1.91616263 | 0.63416063 | 0.52597599 | 0.93247207 | 2.09404228 | 3.22377732 | 1.64117035 | 5.41560658 | Non-sig. |
| PF3D7_0621350 | SYS1 | 14.002175 | 1.212942545 | 1.00581701 | 1.20592766 | 0.22784539 | 0.83828834 | 6.17624288 | 21.604921 | 10.6289124 | 17.5986237 | Non-sig. |
| PF3D7_0304000 | IMC1a | 178.071747 | 1.210371176 | 0.55897947 | 2.16532314 | 0.03036295 | 0.39012163 | 150.478396 | 257.538251 | 64.878768 | 239.391571 | Non-sig. |
| PF3D7_1001900 | PfJ23 | 371.632512 | 1.200481591 | 0.45981104 | 2.61081504 | 0.00903267 | 0.18548489 | 199.358415 | 444.614863 | 251.174803 | 591.381966 | Non-sig. |
| PF3D7_1253500 | N/A | 98.6467128 | 1.199615608 | 0.67222905 | 1.78453401 | 0.07433693 | 0.59046038 | 90.5369078 | 176.52868 | 29.5121374 | 98.0091257 | Non-sig. |
| PF3D7_1036900 | N/A | 11475.0638 | 1.198401079 | 0.43775812 | 2.73758732 | 0.00618917 | 0.14445029 | 8429.71172 | 14808.7845 | 5501.56437 | 17160.1945 | Non-sig. |
| PF3D7_0309800 | YTH2 | 369.361086 | 1.189980224 | 0.4460901 | 2.66757819 | 0.00764001 | 0.16526913 | 250.08402 | 524.947894 | 200.334177 | 502.078251 | Non-sig. |
| PF3D7_1217600 | APC10 | 80.0319389 | 1.189972056 | 0.56312054 | 2.11317465 | 0.03458582 | 0.41644474 | 53.3422578 | 98.8917308 | 44.3039667 | 123.589801 | Non-sig. |
| PF3D7_1372200 | HRPIII | 209.758246 | 1.189021664 | 0.4643388 | 2.56067698 | 0.01044684 | 0.20575234 | 129.361269 | 292.99509 | 126.460354 | 290.216271 | Non-sig. |
| PF3D7_1332000 | SYN5 | 1033.47856 | 1.187088601 | 0.41323797 | 2.87265133 | 0.00407043 | 0.11059047 | 630.887558 | 1429.64955 | 630.634374 | 1442.74278 | Non-sig. |
| PF3D7_1337800 | CDPK5 | 2975.42191 | 1.186959345 | 0.535502 | 2.21653577 | 0.02665483 | 0.36318813 | 2652.00721 | 3890.01374 | 980.722984 | 4378.94371 | Non-sig. |

| | | | | | | | | | | | | |
|---|---|---|---|---|---|---|---|---|---|---|---|---|
| PF3D7_0209700 | N/A | 2166.15035 | 1.184361704 | 0.41432433 | 2.8585377 | 0.00425599 | 0.11445792 | 1387.05334 | 3281.38231 | 1260.63922 | 2735.52652 | Non-sig. |
| PF3D7_0401800 | PfD80 | 1776.98545 | 1.180670405 | 0.41937483 | 2.81531058 | 0.00487301 | 0.12532628 | 1009.43203 | 2712.52477 | 1166.23183 | 2219.75318 | Non-sig. |
| PF3D7_1328500 | N/A | 3629.15393 | 1.17779321 | 0.42896627 | 2.74565458 | 0.00603903 | 0.14347438 | 1903.05556 | 4465.49167 | 2546.48517 | 5601.58332 | Non-sig. |
| PF3D7_1433500 | TOP2 | 11494.4787 | 1.172161307 | 0.42747823 | 2.74203742 | 0.00610594 | 0.1443399 | 8044.97729 | 13916.9238 | 6087.15487 | 17928.859 | Non-sig. |
| PF3D7_0900700 | RIF | 2.350332 | 1.171059672 | 2.24445193 | 0.52175752 | 0.60183917 | 0.95693578 | 2.09681171 | 1.07419275 | 0.82071531 | 5.40960824 | Non-sig. |
| PF3D7_1400800 | RIF | 2.3449054 | 1.170571717 | 2.1916775 | 0.53409852 | 0.59327337 | 0.95391876 | 2.09312509 | 4.2988801 | 0.82052626 | 2.16709016 | Non-sig. |
| PF3D7_0919300 | TrxL1 | 1478.87915 | 1.170448582 | 0.58529746 | 1.99974997 | 0.04552727 | 0.47763064 | 1410.66545 | 2250.18348 | 409.565666 | 1845.102 | Non-sig. |
| PF3D7_0102500 | EBA181 | 19844.6394 | 1.16962419 | 0.55325125 | 2.11409226 | 0.03450739 | 0.41644474 | 18163.5056 | 23256.184 | 6264.74388 | 31694.1241 | Non-sig. |
| PF3D7_1475700 | N/A | 55.9171389 | 1.167463952 | 0.66568908 | 1.75376762 | 0.07947033 | 0.60514927 | 21.0071367 | 81.5960755 | 47.6171016 | 73.4482418 | Non-sig. |
| PF3D7_0503600 | MyoB | 7385.83778 | 1.165683705 | 0.40140115 | 2.9040368 | 0.00368385 | 0.1021834 | 4575.85977 | 10032.7425 | 4532.92662 | 10401.8222 | Non-sig. |
| PF3D7_1200100 | VAR | 8960.81935 | 1.162542659 | 0.41286527 | 2.81579184 | 0.00486572 | 0.12532628 | 5941.98246 | 13593.2051 | 5125.97735 | 11182.1125 | Non-sig. |
| PF3D7_1018300 | N/A | 1757.62881 | 1.160097219 | 0.44563311 | 2.60325634 | 0.00923429 | 0.18834417 | 1229.93785 | 1973.00053 | 943.731241 | 2883.84563 | Non-sig. |
| PF3D7_0604500 | N/A | 19407.0994 | 1.1592334 | 0.53592086 | 2.16306827 | 0.03053593 | 0.39012163 | 17620.8318 | 25540.8548 | 6388.11395 | 28078.5969 | Non-sig. |
| PF3D7_1461800 | N/A | 767.316496 | 1.156978682 | 0.51732349 | 2.23647042 | 0.02532097 | 0.35869122 | 648.398315 | 868.724861 | 302.320662 | 1249.82214 | Non-sig. |
| PF3D7_0408700 | PLP1 | 35.9095975 | 1.156825877 | 0.75814604 | 1.52586154 | 0.12704435 | 0.71411411 | 31.5669959 | 56.8739923 | 13.1398654 | 42.0575362 | Non-sig. |
| PF3D7_1121400 | N/A | 68.7041932 | 1.154642943 | 0.59937083 | 1.92642499 | 0.05405133 | 0.50202873 | 52.5452454 | 101.980151 | 32.8423914 | 87.4489844 | Non-sig. |
| PF3D7_0808600 | VAR | 242.345051 | 1.154265323 | 0.50152559 | 2.30150834 | 0.02136291 | 0.3235449 | 190.788593 | 311.0437 | 110.080057 | 357.467854 | Non-sig. |
| PF3D7_1103800 | NOT1-G | 6017.8649 | 1.153433368 | 0.54064345 | 2.13344556 | 0.03288819 | 0.40615121 | 5456.40751 | 7171.66825 | 2009.47045 | 9433.91339 | Non-sig. |
| PF3D7_0415800 | N/A | 1446.6311 | 1.150703036 | 0.41080909 | 2.80106515 | 0.00509342 | 0.12789079 | 895.082687 | 1906.47801 | 901.843804 | 2083.11988 | Non-sig. |
| PF3D7_0413200 | RIF | 3.14569294 | 1.145682834 | 1.94326903 | 0.58956471 | 0.55548252 | 0.94243456 | 3.1449097 | 5.37111114 | 0.82070109 | 3.24604983 | Non-sig. |
| PF3D7_0110800 | N/A | 1816.75678 | 1.143150624 | 0.4094929 | 2.79162499 | 0.00524441 | 0.12863372 | 1084.66857 | 2584.13399 | 1180.08916 | 2418.13541 | Non-sig. |
| PF3D7_0410800 | N/A | 12406.0524 | 1.140857182 | 0.42813277 | 2.66472751 | 0.00770507 | 0.16526913 | 9050.4958 | 15600.2741 | 6432.60945 | 18540.8302 | Non-sig. |
| PF3D7_1321900 | N/A | 5281.60756 | 1.140353714 | 0.41277187 | 2.76267307 | 0.00573302 | 0.13844179 | 3011.27118 | 6707.39043 | 3581.60763 | 7826.16101 | Non-sig. |
| PF3D7_1027000 | N/A | 337.243066 | 1.135455251 | 0.58840974 | 1.92970166 | 0.05364381 | 0.50202873 | 321.424021 | 449.432307 | 101.039774 | 477.076162 | Non-sig. |
| PF3D7_1240600 | VAR | 149.142313 | 1.133937654 | 0.52817202 | 2.14690975 | 0.03180047 | 0.39923904 | 118.047366 | 188.775702 | 69.0044258 | 220.741758 | Non-sig. |
| PF3D7_0711200 | N/A | 203.799466 | 1.13259967 | 0.49602696 | 2.28334298 | 0.02241017 | 0.32982723 | 98.9909719 | 289.668668 | 156.058234 | 270.47999 | Non-sig. |
| PF3D7_1133700 | N/A | 1309.84423 | 1.128672435 | 0.57646682 | 1.95791397 | 0.05024011 | 0.4954291 | 1260.29785 | 1957.59752 | 384.438394 | 1637.04317 | Non-sig. |
| PF3D7_1351800 | N/A | 288.08211 | 1.126062925 | 0.48142296 | 2.33903036 | 0.01933386 | 0.30809506 | 214.429935 | 338.586835 | 147.891718 | 451.419954 | Non-sig. |
| PF3D7_1251200 | N/A | 38825.6249 | 1.123518425 | 0.43699564 | 2.57100602 | 0.01014036 | 0.20121201 | 29549.4454 | 57227.1221 | 19307.0016 | 49218.9306 | Non-sig. |
| PF3D7_0411000 | N/A | 724.760604 | 1.122626799 | 0.42108891 | 2.66600894 | 0.00767577 | 0.16526913 | 468.863814 | 949.334762 | 443.572922 | 1037.27092 | Non-sig. |
| PF3D7_1332100 | N/A | 28.2258105 | 1.122420124 | 0.78988563 | 1.4209907 | 0.15531946 | 0.75842353 | 11.5542495 | 37.5768439 | 23.8086683 | 39.9634804 | Non-sig. |
| PF3D7_1116000 | RON4 | 14672.4043 | 1.121921569 | 0.41734117 | 2.68825996 | 0.00718255 | 0.15922925 | 9597.53478 | 17266.6011 | 8879.51105 | 22945.9701 | Non-sig. |
| PF3D7_0106600 | N/A | 69.8691606 | 1.12070197 | 0.90740423 | 1.23506364 | 0.21680682 | 0.83527883 | 82.6312195 | 91.3663791 | 5.74314531 | 99.7358983 | Non-sig. |
| PF3D7_0906500 | N/A | 1184.98497 | 1.120103246 | 0.41666841 | 2.68823657 | 0.00718305 | 0.15922925 | 686.098716 | 1605.32083 | 807.289264 | 1641.23109 | Non-sig. |
| PF3D7_1240300 | VAR | 1539.71203 | 1.116928575 | 0.41120812 | 2.71621237 | 0.00660335 | 0.15144837 | 948.685629 | 2216.98219 | 994.830361 | 1998.34995 | Non-sig. |
| PF3D7_1471500 | TMEM43 | 49.9403434 | 1.114961454 | 0.64192318 | 1.73690791 | 0.08240343 | 0.60888892 | 25.1869455 | 70.8768165 | 37.7595812 | 65.9380303 | Non-sig. |
| PF3D7_0624800 | N/A | 75.5488328 | 1.109010675 | 0.5764013 | 1.92402528 | 0.05435141 | 0.50202873 | 38.8910046 | 107.341693 | 56.6550494 | 99.3075839 | Non-sig. |
| PF3D7_1103900 | N/A | 29.4097153 | 1.107804773 | 0.76675802 | 1.4447906 | 0.14851671 | 0.74754886 | 13.4153628 | 42.0858198 | 23.732859 | 38.4048196 | Non-sig. |
| PF3D7_1211300 | MCM8 | 1951.02807 | 1.107372307 | 0.53430906 | 2.07253139 | 0.03821591 | 0.43374033 | 1801.71421 | 2594.84315 | 672.753099 | 2734.80181 | Non-sig. |
| PF3D7_1224200 | N/A | 1513.49069 | 1.103968194 | 0.41514939 | 2.65920708 | 0.00783248 | 0.16732452 | 1041.12603 | 2136.26068 | 881.2614 | 1995.31466 | Non-sig. |
| PF3D7_0416000 | N/A | 3323.15069 | 1.103827109 | 0.40713021 | 2.71123852 | 0.00670324 | 0.15241957 | 1984.1829 | 4403.94221 | 2236.59062 | 4667.88703 | Non-sig. |
| PF3D7_1118600 | MYST | 1835.62029 | 1.101352669 | 0.41563641 | 2.64979835 | 0.00805398 | 0.17068 | 1168.06465 | 2763.16046 | 1166.17804 | 2245.07802 | Non-sig. |
| PF3D7_0420300 | ApiAP2 | 34077.4234 | 1.097674863 | 0.44213163 | 2.48268793 | 0.01303953 | 0.23935 | 23433.0829 | 35981.4252 | 19976.4897 | 56918.6958 | Non-sig. |
| PF3D7_0724800 | N/A | 1921.11647 | 1.094606813 | 0.46795361 | 2.33913531 | 0.01932843 | 0.30809506 | 1556.03085 | 2952.2006 | 895.130982 | 2281.10344 | Non-sig. |
| PF3D7_1246400 | MTIP | 10239.8815 | 1.089010045 | 0.40853079 | 2.66567432 | 0.00768341 | 0.16526913 | 6125.38376 | 14969.8072 | 6971.98851 | 12892.3464 | Non-sig. |
| PF3D7_1400200 | RIF | 247.114335 | 1.088858231 | 0.47355337 | 2.29933585 | 0.02148588 | 0.3235449 | 147.774091 | 388.895486 | 168.236336 | 283.551426 | Non-sig. |
| PF3D7_0207900 | SERA2 | 878.389318 | 1.088225552 | 0.43187107 | 2.51979265 | 0.0117424 | 0.22060719 | 485.496936 | 1263.79906 | 638.200614 | 1126.06066 | Non-sig. |
| PF3D7_1252600 | N/A | 929.762084 | 1.087705205 | 0.42162206 | 2.57981092 | 0.00988544 | 0.19763424 | 538.392365 | 1269.00494 | 651.392774 | 1260.25826 | Non-sig. |
| PF3D7_0222800 | N/A | 2.76240461 | 1.087576429 | 2.04321541 | 0.53228672 | 0.59452743 | 0.9542761 | 1.04615601 | 5.37422261 | 2.46140993 | 2.16782989 | Non-sig. |
| PF3D7_1447500 | IMC20 | 97.688731 | 1.084778685 | 0.74433932 | 1.4573712 | 0.14501394 | 0.74131203 | 105.117418 | 164.110642 | 20.4508911 | 101.075973 | Non-sig. |
| PF3D7_1040500 | RIF | 7.56866641 | 1.083079144 | 1.32624544 | 0.81665061 | 0.41412812 | 0.90793155 | 7.33398872 | 11.8183106 | 2.46185758 | 8.66050871 | Non-sig. |
| PF3D7_0104300 | UBP1 | 19219.8172 | 1.079788419 | 0.66331239 | 1.62787314 | 0.10355179 | 0.6615241 | 20815.8009 | 21943.9176 | 3875.18116 | 30244.3692 | Non-sig. |
| PF3D7_1247600 | N/A | 82.2629235 | 1.077302366 | 0.60845329 | 1.77055888 | 0.07663409 | 0.59794902 | 72.456576 | 112.739302 | 33.65889 | 110.196926 | Non-sig. |
| PF3D7_0900300 | RIF | 2.40052712 | 1.077299343 | 2.38319195 | 0.45204053 | 0.65123979 | 0.97317009 | 3.14256742 | 4.29565341 | 0 | 2.16388766 | Non-sig. |

| | | | | | | | | | | | | |
|---|---|---|---|---|---|---|---|---|---|---|---|---|
| PF3D7_0517600 | CPbeta | 92.1720316 | 1.076758765 | 0.60083644 | 1.79209964 | 0.07311701 | 0.58427436 | 81.9636696 | 123.452582 | 36.947143 | 126.324732 | Non-sig. |
| PF3D7_0114200 | EPF3 | 4.0062908 | 1.076320282 | 1.957838 | 0.54974941 | 0.58249126 | 0.95129458 | 5.23607181 | 4.29591724 | 0 | 6.49317414 | Non-sig. |
| PF3D7_0625500 | N/A | 304.860766 | 1.07461085 | 0.44655227 | 2.40646149 | 0.0161079 | 0.27349786 | 186.355116 | 418.437325 | 206.152881 | 408.497739 | Non-sig. |
| PF3D7_1254000 | RIF | 2.39842637 | 1.074593571 | 2.43947708 | 0.44050161 | 0.65957385 | 0.97707593 | 3.14156256 | 5.36992528 | 0 | 1.08221764 | Non-sig. |
| PF3D7_0518900 | N/A | 161.475131 | 1.072935291 | 0.69322844 | 1.54773697 | 0.12168564 | 0.70612322 | 172.467488 | 253.064869 | 36.0606121 | 184.307556 | Non-sig. |
| PF3D7_0301300 | EH1 | 601.027922 | 1.071677299 | 0.47216766 | 2.26969651 | 0.023226 | 0.3380532 | 286.072447 | 753.412057 | 488.598393 | 876.02879 | Non-sig. |
| PF3D7_1321400 | DHHC8 | 198.49818 | 1.070854773 | 0.48010931 | 2.23043952 | 0.02571828 | 0.36142026 | 143.779352 | 248.066984 | 112.458417 | 289.687967 | Non-sig. |
| PF3D7_0935700 | N/A | 678.861739 | 1.07041282 | 0.44978585 | 2.37982769 | 0.01732074 | 0.28676642 | 525.645939 | 959.587629 | 350.615184 | 879.598203 | Non-sig. |
| PF3D7_0402300 | RH1 | 14733.603 | 1.069365668 | 0.80169555 | 1.33388001 | 0.18224318 | 0.79993735 | 17612.6228 | 16060.8433 | 1408.08277 | 23852.8632 | Non-sig. |
| PF3D7_0909700 | N/A | 2335.51522 | 1.066325377 | 0.43891444 | 2.42946068 | 0.01512131 | 0.26180613 | 1562.0636 | 2502.33079 | 1457.3149 | 3820.35159 | Non-sig. |
| PF3D7_1101700 | MC-2TM | 51.4886954 | 1.065984587 | 0.63031134 | 1.69120325 | 0.09079799 | 0.63212583 | 31.2719754 | 79.6175975 | 35.2596866 | 59.8055223 | Non-sig. |
| PF3D7_0712000 | VAR | 314.406201 | 1.064749729 | 0.45832608 | 2.32312708 | 0.02017233 | 0.3154509 | 193.92862 | 484.814788 | 212.762436 | 366.118958 | Non-sig. |
| PF3D7_0632500 | VAR | 43.255343 | 1.064230722 | 0.66219116 | 1.60713519 | 0.10802471 | 0.67668385 | 26.3591602 | 66.4939195 | 29.575141 | 50.5931511 | Non-sig. |
| PF3D7_0316900 | N/A | 3332.76405 | 1.058560074 | 0.40505395 | 2.61338042 | 0.00896514 | 0.18481454 | 2255.54342 | 4527.04128 | 2068.80504 | 4479.66648 | Non-sig. |
| PF3D7_1140000 | CA | 146.3176 | 1.058363594 | 0.64701114 | 1.63577337 | 0.10188703 | 0.65895255 | 149.247401 | 213.613978 | 41.0539024 | 181.355118 | Non-sig. |
| PF3D7_1254200 | RIF | 6.38736665 | 1.057330834 | 1.37499822 | 0.76896888 | 0.44191179 | 0.91813673 | 4.19094934 | 8.59507833 | 4.10311308 | 8.66032584 | Non-sig. |
| PF3D7_1473700 | NUP116 | 473.62106 | 1.056579561 | 0.66615574 | 1.58608489 | 0.11272005 | 0.68406239 | 515.36986 | 646.783093 | 100.219584 | 632.111703 | Non-sig. |
| PF3D7_0622100 | N/A | 336.57395 | 1.055161695 | 0.50905473 | 2.07278635 | 0.03819217 | 0.43374033 | 290.775879 | 419.43378 | 147.034002 | 489.052138 | Non-sig. |
| PF3D7_0629500 | AAT1 | 1602.09149 | 1.052327895 | 0.40857512 | 2.57560445 | 0.0100065 | 0.19930249 | 1045.76524 | 2115.63051 | 1039.02648 | 2207.94374 | Non-sig. |
| PF3D7_1243700 | N/A | 12906.6172 | 1.051932569 | 0.48922401 | 2.15020633 | 0.0315389 | 0.39723726 | 11447.4232 | 17980.4681 | 5351.43482 | 16847.1425 | Non-sig. |
| PF3D7_0929600 | N/A | 569.637566 | 1.051509406 | 0.58590658 | 1.79467076 | 0.07270617 | 0.58363337 | 564.724707 | 886.982372 | 177.312356 | 649.530829 | Non-sig. |
| PF3D7_0212600 | SPATR | 5299.66368 | 1.049655726 | 0.45781944 | 2.29272863 | 0.02186363 | 0.32547388 | 4401.21109 | 7513.24324 | 2504.23448 | 6779.96593 | Non-sig. |
| PF3D7_1218000 | TRAMP | 1191.19736 | 1.04923925 | 0.52663665 | 1.99233998 | 0.04633377 | 0.47851132 | 1111.85449 | 1553.98028 | 440.982024 | 1657.97263 | Non-sig. |
| PF3D7_0404100 | ApiAP2 | 342.994625 | 1.044001505 | 0.56801646 | 1.83797757 | 0.06606571 | 0.56240843 | 327.780996 | 393.644045 | 120.757024 | 529.796434 | Non-sig. |
| PF3D7_0300200 | RIF | 13.6237152 | 1.043474639 | 1.02308254 | 1.01993202 | 0.3077607 | 0.87709318 | 10.4687204 | 23.641939 | 7.38510116 | 12.9991003 | Non-sig. |
| PF3D7_1107900 | MSCS | 1049.53913 | 1.040062751 | 0.50830366 | 2.04614451 | 0.04074215 | 0.45141486 | 949.393414 | 1305.44 | 424.685916 | 1518.63721 | Non-sig. |
| PF3D7_1252500 | N/A | 52.3731915 | 1.039432989 | 0.62066905 | 1.67469765 | 0.09399357 | 0.64421466 | 31.5478996 | 66.5431613 | 36.9519559 | 74.4497491 | Non-sig. |
| PF3D7_1300100 | VAR | 1573.07533 | 1.039257725 | 0.52378533 | 1.98412911 | 0.04724145 | 0.48052729 | 1455.44863 | 2402.6545 | 604.601708 | 1829.59648 | Non-sig. |
| PF3D7_1373000 | RIF | 3.17758825 | 1.037897603 | 2.10539981 | 0.49296936 | 0.62203424 | 0.96429926 | 0 | 3.21577647 | 4.09302618 | 5.40155034 | Non-sig. |
| PF3D7_1008800 | NOP5 | 3084.79459 | 1.036622083 | 0.41077132 | 2.52359898 | 0.01161604 | 0.21979199 | 1899.18674 | 4436.61469 | 2144.45092 | 3858.92601 | Non-sig. |
| PF3D7_0829500 | N/A | 206.402814 | 1.035199674 | 0.81963744 | 1.26299705 | 0.20659024 | 0.82391338 | 249.79461 | 298.180491 | 21.3584823 | 256.277672 | Non-sig. |
| PF3D7_1016500 | N/A | 189.27168 | 1.034660796 | 0.52144877 | 1.98420413 | 0.04723309 | 0.48052729 | 155.826947 | 296.122422 | 92.8103401 | 212.327012 | Non-sig. |
| PF3D7_1327100 | N/A | 8.48344284 | 1.029194464 | 1.39409703 | 0.73825167 | 0.46036153 | 0.92336312 | 10.4756107 | 12.8932252 | 0.82059814 | 9.74433734 | Non-sig. |
| PF3D7_0614000 | N/A | 4114.94936 | 1.029010572 | 0.42108971 | 2.44368489 | 0.01453812 | 0.25674317 | 2294.79649 | 5593.05202 | 3118.22481 | 5453.72414 | Non-sig. |
| PF3D7_1019300 | N/A | 2984.75537 | 1.028947887 | 0.49835328 | 2.06469575 | 0.0389518 | 0.43907792 | 1221.74553 | 3827.92431 | 2704.38028 | 4184.97135 | Non-sig. |
| PF3D7_0829400 | N/A | 1892.6532 | 1.028505453 | 0.41005841 | 2.50819253 | 0.01213505 | 0.22637855 | 1177.87732 | 2625.59283 | 1312.42752 | 2454.71511 | Non-sig. |
| PF3D7_1313100 | N/A | 111.509754 | 1.025350528 | 0.56260292 | 1.82251193 | 0.06837735 | 0.57292315 | 93.8213078 | 126.559519 | 53.3978478 | 172.260341 | Non-sig. |
| PF3D7_0531100 | N/A | 3006.17484 | 1.023726598 | 0.5232052 | 1.95664456 | 0.05038927 | 0.4954291 | 2826.30242 | 3599.41437 | 1138.61113 | 4460.37146 | Non-sig. |
| PF3D7_1116800 | HSP101 | 7092.22911 | 1.022497303 | 0.59889828 | 1.7072971 | 0.08776685 | 0.62914105 | 7345.46694 | 7739.61569 | 2013.30828 | 11270.5255 | Non-sig. |
| PF3D7_0732900 | RIF | 10.8394577 | 1.019068668 | 1.12283388 | 0.90758632 | 0.36409683 | 0.89567376 | 5.23714877 | 18.2660438 | 9.02637882 | 10.8282594 | Non-sig. |
| PF3D7_0727000 | VPS53 | 1090.35413 | 1.018599484 | 0.50844952 | 2.00334437 | 0.04514034 | 0.47730745 | 1000.73078 | 1510.44518 | 441.092156 | 1409.1484 | Non-sig. |
| PF3D7_0625000 | SMS1 | 1242.39442 | 1.016745801 | 0.41207843 | 2.46735994 | 0.01361135 | 0.24576103 | 846.05514 | 1709.97259 | 797.723897 | 1615.82606 | Non-sig. |
| PF3D7_1222600 | AP2-G | 5106.95327 | 1.015379143 | 0.48290982 | 2.10262684 | 0.0354984 | 0.42313505 | 4499.01828 | 6107.10486 | 2262.40931 | 7559.28064 | Non-sig. |
| PF3D7_1369300 | N/A | 118.317042 | 1.014487474 | 0.69890091 | 1.45154694 | 0.14662762 | 0.7426703 | 128.373969 | 181.352315 | 28.7435151 | 134.79837 | Non-sig. |
| PF3D7_1475800 | N/A | 355.547792 | 1.013857051 | 0.4782498 | 2.119932 | 0.03401178 | 0.41400122 | 259.489874 | 582.537006 | 211.687851 | 368.476438 | Non-sig. |
| PF3D7_1137800 | SPELD | 111.850711 | 1.011773763 | 0.63287913 | 1.59868403 | 0.10989083 | 0.67975706 | 109.39398 | 179.129742 | 39.2828405 | 119.596283 | Non-sig. |
| PF3D7_0729900 | N/A | 699.524784 | 1.009974331 | 0.52501434 | 1.92370809 | 0.05439118 | 0.50202873 | 292.092476 | 745.351673 | 635.878012 | 1124.77697 | Non-sig. |
| PF3D7_0910000 | SET4 | 321.7026 | 1.002976273 | 0.50020766 | 2.00511978 | 0.04495024 | 0.47724727 | 280.077191 | 437.741861 | 148.661722 | 420.329624 | Non-sig. |
| PF3D7_1232600 | N/A | 1568.69317 | 1.002889936 | 0.42603285 | 2.35402023 | 0.0185716 | 0.29997665 | 910.304861 | 2261.65797 | 1178.27106 | 1924.5388 | Non-sig. |
| PF3D7_1469500 | N/A | 61.2764137 | 1.000600371 | 0.59596142 | 1.67896838 | 0.09315821 | 0.64025761 | 38.9348492 | 90.1384662 | 42.7049887 | 73.3273507 | Non-sig. |
| PF3D7_0315600 | N/A | 168.423756 | 0.997384178 | 0.61195202 | 1.62984047 | 0.10313522 | 0.66071388 | 169.421124 | 252.172288 | 55.8450957 | 196.256514 | Non-sig. |
| PF3D7_0628400 | N/A | 74.9101459 | 0.997046332 | 0.64564439 | 1.54426546 | 0.12252404 | 0.7079393 | 72.4383211 | 97.7143067 | 27.9110117 | 101.576944 | Non-sig. |
| PF3D7_1321600 | PDEgamma | 231.499757 | 0.996362344 | 0.51287195 | 1.94271171 | 0.052051 | 0.50193661 | 201.97604 | 317.659201 | 107.577733 | 298.786054 | Non-sig. |
| PF3D7_1039900 | EPF4 | 19.9255819 | 0.990377309 | 0.88324417 | 1.12129504 | 0.2621623 | 0.85030738 | 17.7870143 | 25.795252 | 9.02539506 | 27.0946663 | Non-sig. |

| | | | | | | | | | | | | |
|---|---|---|---|---|---|---|---|---|---|---|---|---|
| PF3D7_1200500 | RIF | 728.960116 | 0.986517264 | 0.44496024 | 2.21709082 | 0.02661689 | 0.36318813 | 469.723175 | 1148.64006 | 508.230056 | 789.247172 | Non-sig. |
| PF3D7_1102900 | N/A | 34.9791766 | 0.983400498 | 0.71318547 | 1.37888465 | 0.16793032 | 0.77528056 | 28.24589 | 49.4430062 | 18.8707733 | 43.3570369 | Non-sig. |
| PF3D7_1302000 | PTP6 | 480.778607 | 0.983164306 | 0.44215098 | 2.22359409 | 0.02617577 | 0.36318813 | 338.947955 | 556.876316 | 307.164775 | 720.125382 | Non-sig. |
| PF3D7_0113300 | N/A | 159.529773 | 0.981006069 | 0.4923567 | 1.99247024 | 0.04631949 | 0.47851132 | 93.6840797 | 221.033954 | 120.730474 | 202.670584 | Non-sig. |
| PF3D7_0919200 | N/A | 3030.63555 | 0.980464189 | 0.44165134 | 2.21999598 | 0.02641904 | 0.36318813 | 2432.88991 | 4494.59916 | 1644.8374 | 3550.21571 | Non-sig. |
| PF3D7_0100300 | VAR | 7.27429663 | 0.979605314 | 1.3394638 | 0.73134139 | 0.46457066 | 0.92538923 | 3.15976401 | 6.43680758 | 6.57104632 | 12.9295686 | Non-sig. |
| PF3D7_0608800 | OAT | 1857.7428 | 0.977006606 | 0.4516051 | 2.16340917 | 0.03050972 | 0.39012163 | 1025.94337 | 2036.23749 | 1477.15232 | 2891.63801 | Non-sig. |
| PF3D7_1149300 | FIKK11 | 172.120118 | 0.976382647 | 0.50332119 | 1.93987989 | 0.05239429 | 0.50193661 | 110.412444 | 181.386182 | 121.529873 | 275.151974 | Non-sig. |
| PF3D7_0732400 | RIF | 3.23505655 | 0.97099456 | 1.92915252 | 0.503327 | 0.61473438 | 0.96217474 | 1.048426 | 6.4451218 | 3.28287239 | 2.16380603 | Non-sig. |
| PF3D7_1230300 | SPM2 | 229.431647 | 0.970952565 | 0.46650411 | 2.08133763 | 0.03740301 | 0.43374033 | 161.52935 | 335.104587 | 148.561642 | 272.531007 | Non-sig. |
| PF3D7_1020300 | N/A | 577.518998 | 0.970847252 | 0.47779329 | 2.03193989 | 0.04215974 | 0.46054085 | 499.946755 | 807.9781 | 280.877984 | 721.273153 | Non-sig. |
| PF3D7_0316000 | MA | 1262.61502 | 0.963825122 | 0.60772149 | 1.58596519 | 0.11274721 | 0.68406239 | 1363.24648 | 1548.61792 | 349.007615 | 1789.58808 | Non-sig. |
| PF3D7_1141000 | N/A | 61.5495552 | 0.962714924 | 0.66790211 | 1.44140122 | 0.14947137 | 0.74848704 | 59.9735057 | 87.9952864 | 23.815743 | 74.4136858 | Non-sig. |
| PF3D7_1111600 | N/A | 129.479084 | 0.959144515 | 0.50556599 | 1.89716976 | 0.05780554 | 0.52083968 | 78.8888262 | 177.078631 | 96.9003458 | 165.048533 | Non-sig. |
| PF3D7_1233200 | N/A | 2000.96437 | 0.956597041 | 0.57532492 | 1.66270746 | 0.09637101 | 0.64875933 | 2087.35843 | 2377.89527 | 634.845248 | 2903.75853 | Non-sig. |
| PF3D7_1449000 | GEST | 113.458138 | 0.955261697 | 0.56341459 | 1.69548625 | 0.08998322 | 0.62981332 | 98.0909894 | 176.396255 | 56.5901909 | 122.755116 | Non-sig. |
| PF3D7_0115000 | SURF1.3 | 56.782885 | 0.954118181 | 0.80436718 | 1.18617243 | 0.23555421 | 0.83828834 | 65.3345451 | 55.7777896 | 12.3217887 | 93.6974165 | Non-sig. |
| PF3D7_0111000 | N/A | 2846.13876 | 0.95340709 | 0.44141072 | 2.15990921 | 0.0307797 | 0.39012163 | 1534.60978 | 3444.08775 | 2342.04149 | 4063.81603 | Non-sig. |
| PF3D7_0206200 | PAT | 3247.5675 | 0.951001902 | 0.42810514 | 2.22142139 | 0.02632243 | 0.36318813 | 1945.80586 | 3720.29566 | 2482.66391 | 4841.50457 | Non-sig. |
| PF3D7_1203000 | ORC1 | 3140.49862 | 0.95074227 | 0.47718572 | 1.99239462 | 0.04632778 | 0.47851132 | 2838.80253 | 4307.17696 | 1444.83377 | 3971.18123 | Non-sig. |
| PF3D7_1306000 | N/A | 1289.43021 | 0.949124894 | 0.53442287 | 1.77598104 | 0.07573604 | 0.59646408 | 1275.76106 | 1616.6153 | 484.632528 | 1780.71196 | Non-sig. |
| PF3D7_0900400 | RIF | 5.3136905 | 0.946120561 | 1.49839542 | 0.63142249 | 0.52776432 | 0.93270419 | 3.14260552 | 8.59554299 | 4.10297271 | 5.4136408 | Non-sig. |
| PF3D7_1245700 | N/A | 17.6892787 | 0.944496769 | 0.94950747 | 0.99472284 | 0.31987108 | 0.88680115 | 17.7848545 | 23.6465041 | 6.56377238 | 22.761984 | Non-sig. |
| PF3D7_1243900 | DOC2 | 3444.49891 | 0.944376425 | 0.6209533 | 1.52084935 | 0.12829765 | 0.71411411 | 3829.73019 | 4439.60519 | 882.251416 | 4626.40886 | Non-sig. |
| PF3D7_1014200 | HAP2 | 14.2762609 | 0.940737831 | 1.0030521 | 0.93787535 | 0.34830847 | 0.8930363 | 6.3143067 | 18.2418638 | 13.1401957 | 19.4086775 | Non-sig. |
| PF3D7_0424400 | SURF4.2 | 661.858951 | 0.940493681 | 0.67051947 | 1.40263441 | 0.16072589 | 0.7667698 | 765.27638 | 810.854617 | 142.11873 | 929.186079 | Non-sig. |
| PF3D7_1217900 | N/A | 1497.17401 | 0.939583681 | 0.41054931 | 2.28860127 | 0.02210253 | 0.32776007 | 1067.08225 | 1918.09129 | 985.338812 | 2018.18367 | Non-sig. |
| PF3D7_0722900 | N/A | 59.7617998 | 0.939494752 | 0.61745053 | 1.5215709 | 0.12811663 | 0.71411411 | 32.5170672 | 88.0712816 | 49.2473328 | 69.2115175 | Non-sig. |
| PF3D7_0214100 | SEC31 | 11113.3639 | 0.938194457 | 0.45192794 | 2.07598244 | 0.03789558 | 0.43374033 | 5602.37872 | 14324.5869 | 9641.21347 | 14885.2766 | Non-sig. |
| PF3D7_0802600 | ACbeta | 10322.2745 | 0.933332251 | 0.53526703 | 1.74367596 | 0.08121561 | 0.60860013 | 10374.9552 | 12486.9179 | 3815.81175 | 14611.4131 | Non-sig. |
| PF3D7_1204900 | N/A | 1369.86554 | 0.931730052 | 0.47295234 | 1.97002946 | 0.04883499 | 0.48633045 | 1203.54524 | 2031.3904 | 681.400839 | 1563.12567 | Non-sig. |
| PF3D7_1138700 | KIC5 | 2721.39036 | 0.929408069 | 0.56604543 | 1.64193194 | 0.10060411 | 0.65652696 | 2830.76021 | 3066.61451 | 917.579414 | 4070.60728 | Non-sig. |
| PF3D7_0104200 | N/A | 3423.36558 | 0.928552581 | 0.46056194 | 2.01612966 | 0.04378642 | 0.47123681 | 1958.38351 | 3455.2752 | 2757.77298 | 5522.03064 | Non-sig. |
| PF3D7_0600300 | RIF | 5.73394266 | 0.927306997 | 1.4723618 | 0.62980919 | 0.52881943 | 0.93270419 | 2.09639464 | 7.51976855 | 5.74480493 | 7.57480251 | Non-sig. |
| PF3D7_0317100 | B9 | 135.383419 | 0.925977076 | 0.55341331 | 1.67321072 | 0.09428582 | 0.64455004 | 67.3667811 | 143.780296 | 119.087405 | 211.299196 | Non-sig. |
| PF3D7_1305800 | N/A | 6.60288671 | 0.923769416 | 1.37921608 | 0.6697786 | 0.50299894 | 0.93170321 | 4.18059612 | 5.37562987 | 4.92206621 | 11.9332547 | Non-sig. |
| PF3D7_1211100 | N/A | 103.751741 | 0.923037543 | 0.64412275 | 1.43301497 | 0.15185356 | 0.75391062 | 108.409515 | 149.551974 | 35.2596866 | 121.785791 | Non-sig. |
| PF3D7_1132500 | N/A | 48.4734269 | 0.922183405 | 0.63213298 | 1.45884399 | 0.14460805 | 0.74131203 | 35.8152082 | 61.1474853 | 31.2131615 | 65.7178526 | Non-sig. |
| PF3D7_0100900 | RIF | 4.51033677 | 0.921971988 | 1.64707805 | 0.55976217 | 0.57564167 | 0.94896603 | 2.09542535 | 8.59513501 | 4.10309597 | 3.24769077 | Non-sig. |
| PF3D7_1426500 | ABCG2 | 190.569374 | 0.921507286 | 0.56712951 | 1.62486217 | 0.10419193 | 0.6615241 | 75.7443477 | 231.802495 | 187.23597 | 267.494683 | Non-sig. |
| PF3D7_1341800 | IMC1k | 994.084619 | 0.921140167 | 0.44069345 | 2.09020615 | 0.03659928 | 0.42994016 | 817.673996 | 1341.32677 | 556.812375 | 1260.52534 | Non-sig. |
| PF3D7_1202300 | N/A | 477.20991 | 0.921056083 | 0.52558158 | 1.75245122 | 0.07969625 | 0.60514927 | 455.5657 | 538.359806 | 204.56759 | 710.346544 | Non-sig. |
| PF3D7_0525600 | N/A | 2159.16479 | 0.920720489 | 0.45007615 | 2.04569936 | 0.04078596 | 0.45141486 | 1786.38805 | 3253.7713 | 1199.22393 | 2397.27587 | Non-sig. |
| PF3D7_0803500 | N/A | 3660.63428 | 0.917477594 | 0.43572817 | 2.1056192 | 0.03523744 | 0.42141755 | 2450.74859 | 3806.98751 | 2617.90414 | 5766.8969 | Non-sig. |
| PF3D7_1026400 | CDC20 | 263.505314 | 0.916232468 | 0.51017706 | 1.79591076 | 0.07250871 | 0.58363337 | 239.803956 | 358.457673 | 125.640315 | 330.119312 | Non-sig. |
| PF3D7_1327600 | NMNAT | 678.580938 | 0.913859005 | 0.43675553 | 2.09238107 | 0.03640444 | 0.42955618 | 539.066013 | 922.138428 | 402.312976 | 850.806334 | Non-sig. |
| PF3D7_1414100 | N/A | 955.475707 | 0.913146125 | 0.43813581 | 2.08416228 | 0.03714541 | 0.43374033 | 579.526529 | 1086.62282 | 745.866813 | 1409.88667 | Non-sig. |
| PF3D7_0806500 | N/A | 4247.70701 | 0.911345945 | 0.42938108 | 2.12246414 | 0.03379878 | 0.41377036 | 3496.69657 | 5447.12146 | 2401.57816 | 5645.43184 | Non-sig. |
| PF3D7_0307700 | N/A | 11498.1192 | 0.910007048 | 0.40145944 | 2.26674718 | 0.02340567 | 0.33880673 | 8267.04846 | 15137.7008 | 7707.88026 | 14879.8472 | Non-sig. |
| PF3D7_0808800 | RIF | 25.6519576 | 0.908661768 | 0.84501398 | 1.07532159 | 0.28223078 | 0.86390383 | 25.1650626 | 41.8919668 | 10.6695481 | 24.8812528 | Non-sig. |
| PF3D7_1016800 | N/A | 2818.77645 | 0.907852935 | 0.71114416 | 1.27660885 | 0.20174037 | 0.81901954 | 3440.86138 | 4047.71944 | 479.683424 | 3306.84154 | Non-sig. |
| PF3D7_0725300 | N/A | 1164.0165 | 0.906352495 | 0.51844533 | 1.74821229 | 0.08042727 | 0.6065657 | 1143.20025 | 1599.79641 | 477.176031 | 1435.89332 | Non-sig. |
| PF3D7_1213500 | N/A | 15.6756529 | 0.905237755 | 0.96102545 | 0.94194982 | 0.34621833 | 0.8930363 | 9.46468053 | 15.0257365 | 12.3173895 | 25.8948049 | Non-sig. |
| PF3D7_1413700 | N/A | 4765.95372 | 0.904868336 | 0.67096794 | 1.34860145 | 0.17746502 | 0.79075665 | 5658.32987 | 6318.39101 | 979.125098 | 6107.9689 | Non-sig. |

| | | | | | | | | | | | | |
|---|---|---|---|---|---|---|---|---|---|---|---|---|
| PF3D7_1003600 | IMC1c | 24875.085 | 0.903701272 | 0.40725292 | 2.21901732 | 0.02648555 | 0.36318813 | 18960.9329 | 31866.7514 | 15697.9882 | 32974.6676 | Non-sig. |
| PF3D7_1219300 | VAR | 26.4221511 | 0.903373523 | 0.78326713 | 1.15334027 | 0.24877075 | 0.8482655 | 22.1329865 | 37.54105 | 14.7865587 | 31.2280092 | Non-sig. |
| PF3D7_0916700 | HoMu | 2929.18546 | 0.902074607 | 0.4949804 | 1.8224451 | 0.06838748 | 0.57292315 | 1345.86187 | 3219.97273 | 2737.96556 | 4412.94169 | Non-sig. |
| PF3D7_0417100 | PUF2 | 36.4524612 | 0.900181793 | 0.71701586 | 1.25545591 | 0.2093133 | 0.82566351 | 30.4588981 | 37.5776424 | 20.5244409 | 57.2488634 | Non-sig. |
| PF3D7_0607600 | SAS6 | 5906.46454 | 0.897664372 | 0.41483547 | 2.16390456 | 0.03047168 | 0.39012163 | 3598.98771 | 7759.35507 | 4652.77918 | 7614.73622 | Non-sig. |
| PF3D7_0530900 | FRM1 | 3520.4984 | 0.895474015 | 0.50294071 | 1.78047629 | 0.07499805 | 0.59393071 | 3281.62032 | 3575.84792 | 1642.18771 | 5582.33765 | Non-sig. |
| PF3D7_0828400 | OXA1 | 404.30404 | 0.895297265 | 0.46165337 | 1.93932791 | 0.05246142 | 0.50193661 | 225.173243 | 533.33092 | 339.992786 | 518.71921 | Non-sig. |
| PF3D7_1238400 | N/A | 73.4337348 | 0.895287064 | 0.60387711 | 1.482565 | 0.13819002 | 0.72945925 | 66.0130589 | 106.361563 | 36.9285736 | 84.4317437 | Non-sig. |
| PF3D7_0107500 | NCR1 | 10022.4171 | 0.89508257 | 0.42958552 | 2.08359577 | 0.03719695 | 0.43374033 | 7490.44995 | 10670.8826 | 6528.42778 | 15399.9082 | Non-sig. |
| PF3D7_1116900 | N/A | 136.113937 | 0.894507217 | 0.57268531 | 1.56195243 | 0.1182992 | 0.697844 | 130.105087 | 203.428001 | 60.6656527 | 150.25701 | Non-sig. |
| PF3D7_0400300 | RIF | 12.8729836 | 0.89346886 | 1.04401759 | 0.85579866 | 0.39210916 | 0.90601068 | 11.529933 | 20.4109182 | 6.56546741 | 12.9856156 | Non-sig. |
| PF3D7_1004100 | N/A | 49.6997069 | 0.890187789 | 0.64365 | 1.38303082 | 0.16665542 | 0.77315271 | 36.8569231 | 51.4972095 | 32.8541446 | 77.5905504 | Non-sig. |
| PF3D7_1211000 | N/A | 616.060154 | 0.888255163 | 0.5905569 | 1.5040975 | 0.13255626 | 0.72037933 | 670.173651 | 794.844299 | 194.682626 | 804.54004 | Non-sig. |
| PF3D7_0504200 | DDX27 | 926.225243 | 0.887283837 | 0.43433723 | 2.04284544 | 0.04106774 | 0.45239813 | 750.991506 | 1262.10193 | 549.37183 | 1142.43571 | Non-sig. |
| PF3D7_1036400 | LSA1 | 271.503367 | 0.884399295 | 1.97201219 | 0.44847557 | 0.65381001 | 0.97317009 | 375.923863 | 344.391908 | 5.74943058 | 359.948268 | Non-sig. |
| PF3D7_1335000 | MSRP1 | 1117.11986 | 0.880431489 | 0.4169974 | 2.11135967 | 0.03474141 | 0.41736958 | 772.342109 | 1335.4387 | 800.520002 | 1560.17862 | Non-sig. |
| PF3D7_1036300 | DBLMSP2 | 1630.92068 | 0.879710024 | 0.5059666 | 1.73867212 | 0.08209245 | 0.60888892 | 1597.88046 | 2129.19188 | 699.662451 | 2096.9479 | Non-sig. |
| PF3D7_1028700 | MTRAP | 18462.5559 | 0.879540259 | 0.4782575 | 1.83905168 | 0.06590758 | 0.56240843 | 17389.7701 | 24369.9577 | 8616.23974 | 23474.2559 | Non-sig. |
| PF3D7_1101300 | RIF | 8.69631957 | 0.879083358 | 1.27971138 | 0.68693877 | 0.49212132 | 0.93170321 | 3.14097315 | 16.1190172 | 9.02639763 | 6.49889028 | Non-sig. |
| PF3D7_1113400 | DSK2 | 3258.48122 | 0.87466768 | 0.44157593 | 1.9807866 | 0.04761521 | 0.48234299 | 1780.44982 | 4224.30667 | 2818.99458 | 4210.1738 | Non-sig. |
| PF3D7_1232900 | N/A | 289.372452 | 0.87414405 | 0.46687004 | 1.87234984 | 0.06115822 | 0.53711809 | 238.831099 | 393.837551 | 169.993791 | 354.827365 | Non-sig. |
| PF3D7_0818100 | N/A | 7471.78991 | 0.872675368 | 0.40110157 | 2.1756967 | 0.02957795 | 0.38502203 | 5312.03829 | 9627.61472 | 5244.87512 | 9702.63153 | Non-sig. |
| PF3D7_0115600 | RIF | 12.0491622 | 0.872673 | 1.07116331 | 0.8146965 | 0.41524606 | 0.90793155 | 6.29019115 | 19.3356798 | 10.6692258 | 11.901552 | Non-sig. |
| PF3D7_1342500 | SPECT1 | 480.32047 | 0.872445228 | 0.70770914 | 1.23277371 | 0.2176602 | 0.83527883 | 587.214139 | 706.428385 | 91.9520461 | 535.687309 | Non-sig. |
| PF3D7_1147200 | N/A | 997.618037 | 0.870519094 | 0.50921625 | 1.70952732 | 0.08735331 | 0.62914105 | 907.656501 | 936.321742 | 503.584171 | 1642.90973 | Non-sig. |
| PF3D7_1251000 | N/A | 9.71290833 | 0.867188606 | 1.24335287 | 0.69745977 | 0.48551511 | 0.93170321 | 11.4177343 | 9.69437659 | 2.45824393 | 15.2812785 | Non-sig. |
| PF3D7_0730900 | PTP4 | 3376.22978 | 0.86656769 | 0.43031024 | 2.01382077 | 0.04402835 | 0.47123681 | 2656.63873 | 3700.20795 | 2126.8948 | 5021.17766 | Non-sig. |
| PF3D7_1030900 | P28 | 35.6719749 | 0.865093184 | 0.74072585 | 1.16789928 | 0.2428474 | 0.8422984 | 34.3654478 | 51.6568776 | 16.3973878 | 40.2681863 | Non-sig. |
| PF3D7_1037400 | N/A | 991.434159 | 0.86506111 | 0.43502652 | 1.98852502 | 0.04675366 | 0.48052729 | 584.711206 | 1310.86538 | 820.597305 | 1249.56274 | Non-sig. |
| PF3D7_1329500 | N/A | 3173.45786 | 0.859372931 | 0.53075393 | 1.61915508 | 0.1054139 | 0.66724354 | 1355.57208 | 3131.87241 | 3154.54285 | 5051.8441 | Non-sig. |
| PF3D7_0524800 | UFD1 | 953.518095 | 0.858391994 | 0.50918441 | 1.68581751 | 0.09183095 | 0.6368068 | 415.826926 | 1185.61278 | 939.577342 | 1273.05533 | Non-sig. |
| PF3D7_0210500 | N/A | 371.692793 | 0.857350408 | 0.4393113 | 1.95157832 | 0.05098829 | 0.49786962 | 279.160259 | 464.500568 | 249.707212 | 493.403135 | Non-sig. |
| PF3D7_1011500 | N/A | 385.783708 | 0.857104707 | 0.45440517 | 1.8862125 | 0.05926633 | 0.52683388 | 229.21838 | 481.918825 | 319.421878 | 512.575751 | Non-sig. |
| PF3D7_1332500 | N/A | 379.105218 | 0.85641794 | 0.45425268 | 1.88533383 | 0.05938478 | 0.52700264 | 309.159367 | 525.337565 | 230.605113 | 451.318828 | Non-sig. |
| PF3D7_1466100 | PPKL | 415.510994 | 0.856400969 | 0.54308723 | 1.57691235 | 0.11481569 | 0.68949152 | 169.557392 | 481.701049 | 421.361146 | 589.424388 | Non-sig. |
| PF3D7_1474000 | N/A | 661.802344 | 0.855560539 | 0.55033264 | 1.55462438 | 0.12003555 | 0.70192976 | 690.521062 | 784.013208 | 252.204582 | 920.470522 | Non-sig. |
| PF3D7_1301600 | EBA140 | 26570.4346 | 0.851314511 | 0.48270054 | 1.76364938 | 0.07779103 | 0.60344518 | 24199.5686 | 26618.3039 | 13702.4916 | 41761.3742 | Non-sig. |
| PF3D7_0404500 | P52 | 51.4828095 | 0.850361336 | 0.64164086 | 1.32529174 | 0.18507441 | 0.80052251 | 31.4973458 | 53.6880199 | 41.867085 | 78.8787873 | Non-sig. |
| PF3D7_0724600 | N/A | 2740.40806 | 0.850121125 | 0.40599465 | 2.093922 | 0.03626693 | 0.42888881 | 1947.81142 | 3370.06076 | 1963.34545 | 3680.41463 | Non-sig. |
| PF3D7_0822700 | TRP1 | 163.852687 | 0.8496257 | 0.48064193 | 1.76768953 | 0.07711282 | 0.59991737 | 114.783994 | 202.769763 | 119.096456 | 218.760537 | Non-sig. |
| PF3D7_1207700 | N/A | 86.8992459 | 0.849431865 | 0.70803143 | 1.19970927 | 0.23025227 | 0.83828834 | 100.635103 | 112.793965 | 23.8002477 | 110.367668 | Non-sig. |
| PF3D7_0408000 | N/A | 2582.49768 | 0.846569165 | 0.56981543 | 1.48569015 | 0.1373611 | 0.7277391 | 2827.11779 | 3360.67044 | 865.000107 | 3277.20239 | Non-sig. |
| PF3D7_1114800 | G3PDH | 389.88032 | 0.846123099 | 0.46826327 | 1.80693888 | 0.07077181 | 0.58041649 | 308.829648 | 599.073499 | 248.761811 | 402.856322 | Non-sig. |
| PF3D7_0611500 | N/A | 133.742251 | 0.845793432 | 0.53879259 | 1.56979412 | 0.11646302 | 0.69328209 | 102.699287 | 218.087599 | 88.6304584 | 125.551661 | Non-sig. |
| PF3D7_1301200 | GBPH2 | 478.757945 | 0.845736966 | 0.43929649 | 1.92520765 | 0.05420339 | 0.50202873 | 361.973664 | 681.411931 | 322.748793 | 548.89739 | Non-sig. |
| PF3D7_1138800 | N/A | 2964.82763 | 0.84532698 | 0.43876434 | 1.92660821 | 0.05402847 | 0.50202873 | 2451.5754 | 3239.22517 | 1789.18422 | 4379.32571 | Non-sig. |
| PF3D7_0515000 | CWC2 | 3157.32254 | 0.844799065 | 0.52848201 | 1.59853894 | 0.10992309 | 0.67975706 | 3271.35871 | 4058.42475 | 1246.03132 | 4053.47541 | Non-sig. |
| PF3D7_1444800 | FBPA | 41231.0417 | 0.844648826 | 0.416162 | 2.02961546 | 0.04239564 | 0.46216484 | 26012.4637 | 57351.4885 | 32976.7487 | 48583.4659 | Non-sig. |
| PF3D7_1139700 | N/A | 859.377948 | 0.843946752 | 0.43697202 | 1.93135192 | 0.05343955 | 0.50202873 | 516.774926 | 1167.44486 | 712.868682 | 1040.42332 | Non-sig. |
| PF3D7_0205300 | N/A | 24.3704749 | 0.842566392 | 0.8560008 | 0.98430561 | 0.32496527 | 0.88694295 | 25.2275794 | 36.4957766 | 9.85311937 | 25.9054244 | Non-sig. |
| PF3D7_0902300 | FIKK9.4 | 118.729758 | 0.840171969 | 0.55769837 | 1.50649888 | 0.13193915 | 0.72037933 | 110.540325 | 135.190651 | 59.9559312 | 169.232125 | Non-sig. |
| PF3D7_1304200 | MIT2 | 424.840401 | 0.83782521 | 0.5534532 | 1.51381402 | 0.13007301 | 0.71812182 | 176.550904 | 434.392022 | 432.970745 | 655.447934 | Non-sig. |
| PF3D7_1231300 | N/A | 732.335787 | 0.834011686 | 0.51037067 | 1.63412935 | 0.10223169 | 0.65895255 | 725.458339 | 999.941588 | 327.712482 | 876.230738 | Non-sig. |
| PF3D7_0808900 | RIF | 181.020521 | 0.833852599 | 0.50843097 | 1.64005075 | 0.10099462 | 0.65652696 | 157.085673 | 264.333965 | 103.389704 | 199.27274 | Non-sig. |

| | | | | | | | | | | | | |
|---|---|---|---|---|---|---|---|---|---|---|---|---|
| PF3D7_0422900 | N/A | 226.913839 | 0.832938959 | 0.52367 | 1.59057988 | 0.11170416 | 0.68406239 | 215.906118 | 319.062368 | 110.79062 | 261.896249 | Non-sig. |
| PF3D7_0300800 | RIF | 10.0913136 | 0.832642054 | 1.13271939 | 0.73508237 | 0.46228933 | 0.92422976 | 6.28031348 | 15.0454928 | 8.20547972 | 10.8339686 | Non-sig. |
| PF3D7_0800600 | N/A | 95.3906616 | 0.829834827 | 0.54945641 | 1.51028327 | 0.13097116 | 0.72037933 | 82.6068948 | 123.622932 | 54.9675043 | 120.365315 | Non-sig. |
| PF3D7_0903800 | CCp4 | 37.8395775 | 0.8294758 | 0.68811194 | 1.2054373 | 0.22803453 | 0.83828834 | 31.6042618 | 47.2004712 | 22.9995024 | 49.5540746 | Non-sig. |
| PF3D7_1124200 | N/A | 1636.8469 | 0.828976335 | 0.43858391 | 1.89012026 | 0.05874188 | 0.5248136 | 1014.42214 | 2404.75786 | 1343.5673 | 1784.64028 | Non-sig. |
| PF3D7_0307800 | N/A | 1457.65397 | 0.827364192 | 0.41007132 | 2.01761049 | 0.04363184 | 0.47123681 | 1077.3522 | 1814.16728 | 1024.24073 | 1914.85568 | Non-sig. |
| PF3D7_0618400 | N/A | 19.3115994 | 0.827307021 | 0.88686999 | 0.93283912 | 0.35090304 | 0.8930363 | 10.5102368 | 28.9829122 | 17.2425977 | 20.5106508 | Non-sig. |
| PF3D7_0506500 | N/A | 4441.00625 | 0.826478595 | 0.63986618 | 1.29164287 | 0.19648085 | 0.81437577 | 5301.60875 | 5264.74576 | 1104.14781 | 6093.52269 | Non-sig. |
| PF3D7_1335900 | TRAP | 46.4148779 | 0.825891755 | 0.71038391 | 1.16259919 | 0.24499216 | 0.84331907 | 48.3387665 | 56.8951351 | 18.8841789 | 61.5414309 | Non-sig. |
| PF3D7_1457100 | N/A | 1138.7073 | 0.825429324 | 0.42860426 | 1.92585422 | 0.05412258 | 0.50202873 | 699.153229 | 1454.84024 | 943.717573 | 1457.11818 | Non-sig. |
| PF3D7_1131500 | N/A | 29.6492587 | 0.823569706 | 0.75608184 | 1.08926 | 0.27603925 | 0.85889199 | 24.0116041 | 30.1128828 | 18.8647001 | 45.6078479 | Non-sig. |
| PF3D7_1035300 | GLURP | 11705.376 | 0.822863759 | 0.48512094 | 1.69620333 | 0.08984738 | 0.62981332 | 11214.37 | 12787.7313 | 5695.76675 | 17123.636 | Non-sig. |
| PF3D7_0812900 | N/A | 151.533886 | 0.821117913 | 0.50249018 | 1.63409744 | 0.10223839 | 0.65895255 | 110.653854 | 160.894751 | 108.432874 | 226.154064 | Non-sig. |
| PF3D7_1236500 | N/A | 32.1910872 | 0.820165291 | 0.72551658 | 1.130457 | 0.25828371 | 0.85030738 | 26.1167554 | 36.5589873 | 20.5072654 | 45.5813407 | Non-sig. |
| PF3D7_0629600 | N/A | 285.34943 | 0.819239345 | 0.45484343 | 1.80114581 | 0.0716799 | 0.58133781 | 228.266179 | 372.395071 | 184.769665 | 355.966806 | Non-sig. |
| PF3D7_1236600 | N/A | 35.3638889 | 0.819138289 | 0.69442904 | 1.17958531 | 0.23816519 | 0.84066259 | 25.8054718 | 46.3975895 | 25.3715449 | 43.8809492 | Non-sig. |
| PF3D7_1131800 | N/A | 5637.65283 | 0.818851717 | 0.44310326 | 1.84799301 | 0.06460335 | 0.55653426 | 3144.88934 | 6782.12117 | 5013.43638 | 7610.16443 | Non-sig. |
| PF3D7_0210700 | SYN17 | 1236.49072 | 0.817301411 | 0.41159617 | 1.98568759 | 0.04706802 | 0.48052729 | 897.672734 | 1525.24063 | 892.980006 | 1630.0695 | Non-sig. |
| PF3D7_0216800 | N/A | 956.101633 | 0.816562817 | 0.43696369 | 1.86872006 | 0.06166177 | 0.53908262 | 570.796572 | 1218.74278 | 813.971582 | 1220.8956 | Non-sig. |
| PF3D7_0321300 | N/A | 412.452414 | 0.81549369 | 0.5994065 | 1.36050191 | 0.17367115 | 0.78387252 | 464.371708 | 540.729063 | 133.879355 | 510.829531 | Non-sig. |
| PF3D7_0200200 | RIF | 2.13403647 | 0.815394682 | 2.48902676 | 0.32759579 | 0.74321731 | 0.98983542 | 3.14055051 | 2.14811451 | 0 | 3.24748085 | Non-sig. |
| PF3D7_0409900 | ALP5b | 358.674869 | 0.814905149 | 0.45095867 | 1.80705064 | 0.07075438 | 0.58041649 | 256.399172 | 519.574345 | 263.556287 | 395.169674 | Non-sig. |
| PF3D7_0719200 | NEK4 | 45.2277737 | 0.813124684 | 0.65642855 | 1.23871011 | 0.21545287 | 0.83503497 | 37.8108029 | 64.4189318 | 27.913208 | 50.7681521 | Non-sig. |
| PF3D7_1476600 | N/A | 356.829231 | 0.813097725 | 0.46175988 | 1.76086696 | 0.07826092 | 0.60381404 | 307.690363 | 449.446136 | 210.290162 | 459.890262 | Non-sig. |
| PF3D7_1425400 | N/A | 2260.38259 | 0.812939193 | 0.44954205 | 1.80837188 | 0.07054864 | 0.58041649 | 1303.98665 | 2481.35859 | 1975.43596 | 3280.74916 | Non-sig. |
| PF3D7_0813000 | KIC7 | 2444.83448 | 0.812588686 | 0.45039337 | 1.80417549 | 0.0712038 | 0.58066864 | 2188.43022 | 2832.35651 | 1359.83825 | 3398.71293 | Non-sig. |
| PF3D7_0717500 | CDPK4 | 12987.4623 | 0.811839331 | 0.4072214 | 1.99360672 | 0.04619505 | 0.47851132 | 10244.7666 | 17059.0475 | 8608.9057 | 16037.1293 | Non-sig. |
| PF3D7_0815100 | N/A | 237.712364 | 0.810234337 | 0.45606301 | 1.77658419 | 0.07563668 | 0.59646408 | 173.682466 | 301.508214 | 171.65055 | 304.008224 | Non-sig. |
| PF3D7_1150200 | RIF | 2.12985663 | 0.809957506 | 2.53823769 | 0.31910231 | 0.74964893 | 0.99028237 | 3.14089755 | 4.29612999 | 0 | 1.08239896 | Non-sig. |
| PF3D7_1146700 | N/A | 62.1703358 | 0.809833009 | 0.6435479 | 1.25838809 | 0.20825144 | 0.82398515 | 61.0211846 | 89.0702419 | 29.5639902 | 69.0259267 | Non-sig. |
| PF3D7_1300300 | VAR | 19.4816659 | 0.809122537 | 0.93563441 | 0.86478493 | 0.38715684 | 0.90421052 | 21.0867708 | 19.3048301 | 7.39375127 | 30.1413113 | Non-sig. |
| PF3D7_0911900 | ICP | 4905.58927 | 0.808831711 | 0.42757093 | 1.89169012 | 0.05853228 | 0.52454723 | 2986.24548 | 6754.45344 | 4144.24227 | 5737.4159 | Non-sig. |
| PF3D7_0822900 | PIC2 | 30207.507 | 0.806658689 | 0.56583166 | 1.42561603 | 0.15397921 | 0.75535355 | 33556.1272 | 35776.1268 | 10396.1042 | 41101.6696 | Non-sig. |
| PF3D7_1334400 | MSRP4 | 54.672516 | 0.806527188 | 0.61149834 | 1.31893601 | 0.18719051 | 0.80191512 | 37.6678849 | 75.2354067 | 41.8452729 | 63.9414995 | Non-sig. |
| PF3D7_1331000 | N/A | 1272.27892 | 0.805793038 | 0.41580965 | 1.93788921 | 0.05263674 | 0.50193661 | 993.017169 | 1683.82351 | 858.974657 | 1553.30034 | Non-sig. |
| PF3D7_1335400 | RH2a | 28149.4637 | 0.802631684 | 0.68725797 | 1.1678754 | 0.24285703 | 0.8422984 | 35326.7006 | 30195.1451 | 5703.83968 | 41372.1694 | Non-sig. |
| PF3D7_0303400 | DHHC1 | 1580.79946 | 0.800383466 | 0.51958631 | 1.54042447 | 0.12345692 | 0.70887094 | 666.540136 | 2013.79257 | 1639.40308 | 2003.46206 | Non-sig. |
| PF3D7_0404600 | N/A | 3108.77551 | 0.79980925 | 0.45551573 | 1.75583234 | 0.07911704 | 0.60514927 | 1841.1366 | 3183.02458 | 2695.49966 | 4715.44122 | Non-sig. |
| PF3D7_1129600 | N/A | 431.513362 | 0.794673043 | 0.55521573 | 1.43128697 | 0.15234799 | 0.75391062 | 462.424103 | 546.040783 | 169.207334 | 548.381227 | Non-sig. |
| PF3D7_1007700 | AP2-I | 11572.0985 | 0.793004042 | 0.59391681 | 1.33521064 | 0.18180741 | 0.79970932 | 3664.31042 | 12381.3045 | 13274.0459 | 16968.7334 | Non-sig. |
| PF3D7_1364200 | NUP205 | 2863.08113 | 0.791189683 | 0.52738228 | 1.50022045 | 0.13355731 | 0.72178914 | 1203.5912 | 3109.48732 | 2990.16137 | 4149.08463 | Non-sig. |
| PF3D7_1221900 | N/A | 926.932306 | 0.787961833 | 0.45294537 | 1.73963991 | 0.08192227 | 0.60888892 | 759.541258 | 937.578035 | 600.449297 | 1410.16063 | Non-sig. |
| PF3D7_1117400 | N/A | 2820.05143 | 0.787141885 | 0.41285801 | 1.90656805 | 0.05657655 | 0.51325777 | 1941.74156 | 3867.81281 | 2196.68644 | 3273.9649 | Non-sig. |
| PF3D7_0925900 | LCN | 5177.39781 | 0.786883127 | 0.40327146 | 1.95124921 | 0.05102741 | 0.49786962 | 3658.52702 | 6663.95993 | 3940.3006 | 6446.8037 | Non-sig. |
| PF3D7_0830600 | N/A | 420.148956 | 0.786524851 | 0.56358136 | 1.39558352 | 0.16283995 | 0.76900226 | 457.048145 | 533.203368 | 160.165318 | 530.178993 | Non-sig. |
| PF3D7_1466400 | ApiAP2 | 2210.14749 | 0.785112055 | 0.44732913 | 1.75511052 | 0.0792404 | 0.60514927 | 1265.52599 | 2534.70948 | 1980.50381 | 3059.85068 | Non-sig. |
| PF3D7_0412300 | N/A | 197.566889 | 0.784242044 | 0.61585535 | 1.27341922 | 0.20286932 | 0.82045929 | 67.3304286 | 227.507842 | 222.549244 | 272.880042 | Non-sig. |
| PF3D7_0412200 | N/A | 54.841243 | 0.783927412 | 0.69225869 | 1.13241976 | 0.257458 | 0.85030738 | 22.0365541 | 61.2133729 | 58.2801489 | 77.834896 | Non-sig. |
| PF3D7_0102600 | FIKK1 | 125.866643 | 0.783909623 | 0.51462948 | 1.52325053 | 0.12769604 | 0.71411411 | 102.137105 | 140.54804 | 82.9554074 | 177.82602 | Non-sig. |
| PF3D7_1105900 | N/A | 4.32594368 | 0.783448165 | 1.64999565 | 0.47481832 | 0.63491647 | 0.9668494 | 3.08474878 | 3.242355 | 3.26876594 | 7.70790499 | Non-sig. |
| PF3D7_1440400 | N/A | 2883.57046 | 0.783379815 | 0.40875709 | 1.91649229 | 0.05530246 | 0.5051594 | 2278.95025 | 3662.19942 | 1959.92858 | 3633.20361 | Non-sig. |
| PF3D7_1344100 | N/A | 1242.28098 | 0.782796516 | 0.44532198 | 1.75782142 | 0.0787779 | 0.60514927 | 732.151885 | 1426.74893 | 1094.11567 | 1716.10742 | Non-sig. |
| PF3D7_1434900 | N/A | 112.649176 | 0.782184116 | 0.52539993 | 1.48874043 | 0.13655573 | 0.72522545 | 95.2681451 | 151.522216 | 70.5690389 | 133.237306 | Non-sig. |
| PF3D7_0621400 | ALV7 | 771.798235 | 0.782095375 | 0.49180277 | 1.59026224 | 0.11177571 | 0.68406239 | 758.62762 | 940.041846 | 376.947063 | 1011.57641 | Non-sig. |

| | | | | | | | | | | | | |
|---|---|---|---|---|---|---|---|---|---|---|---|---|
| PF3D7_0903500 | NUP138 | 1059.9282 | 0.779804194 | 0.45646219 | 1.70836538 | 0.08756857 | 0.62914105 | 969.413083 | 1260.38814 | 591.438711 | 1418.47287 | Non-sig. |
| PF3D7_0930300 | MSP1 | 235467.363 | 0.776110566 | 0.46244815 | 1.67826503 | 0.09329537 | 0.64025761 | 130841.596 | 245918.305 | 216390.286 | 348719.266 | Non-sig. |
| PF3D7_1316300 | N/A | 11.3126052 | 0.775976693 | 1.19272549 | 0.65059119 | 0.51531042 | 0.93247207 | 14.3772355 | 14.1062561 | 2.44342873 | 14.3235006 | Non-sig. |
| PF3D7_0937000 | N/A | 16.610336 | 0.772700468 | 0.91989347 | 0.83998908 | 0.40091451 | 0.90667178 | 10.5213291 | 20.3893363 | 13.9608742 | 21.5698046 | Non-sig. |
| PF3D7_1359400 | CELF1 | 3177.40607 | 0.771103128 | 0.41084561 | 1.87686838 | 0.06053614 | 0.53330365 | 2539.9341 | 3819.39094 | 2156.04814 | 4194.25109 | Non-sig. |
| PF3D7_1001400 | XL1 | 240.792879 | 0.771075057 | 0.48046578 | 1.60484907 | 0.10852702 | 0.6779986 | 145.353716 | 278.925859 | 210.274636 | 328.617306 | Non-sig. |
| PF3D7_0410700 | RbgA | 7884.88329 | 0.771043715 | 0.52490368 | 1.46892419 | 0.14185335 | 0.73825055 | 8417.37136 | 10038.6475 | 3236.36545 | 9847.14888 | Non-sig. |
| PF3D7_1029800 | HPR2 | 1458.85715 | 0.770482826 | 0.41902099 | 1.83876906 | 0.06594916 | 0.56240843 | 1045.49494 | 2039.11192 | 1111.06085 | 1639.7609 | Non-sig. |
| PF3D7_1442200 | YihA2 | 1233.3484 | 0.768206535 | 0.5742431 | 1.33777234 | 0.18097066 | 0.79699299 | 1394.32566 | 1685.43322 | 431.215014 | 1422.41971 | Non-sig. |
| PF3D7_1460000 | HCS2 | 144.112397 | 0.765965043 | 0.49397858 | 1.55060377 | 0.12099667 | 0.70312626 | 101.773237 | 195.461121 | 111.632379 | 167.582851 | Non-sig. |
| PF3D7_0803600 | N/A | 5250.54629 | 0.765610295 | 0.46549349 | 1.64472826 | 0.10002586 | 0.65652696 | 5057.2684 | 6543.10408 | 2721.48775 | 6680.32492 | Non-sig. |
| PF3D7_1366600 | SRPR-alpha | 4745.52655 | 0.76536291 | 0.56780142 | 1.34794116 | 0.17767731 | 0.79103729 | 5373.93808 | 6319.07099 | 1657.50229 | 5631.59484 | Non-sig. |
| PF3D7_1208200 | CRMP3 | 149.947685 | 0.76185088 | 0.49967885 | 1.52468107 | 0.12733867 | 0.71411411 | 114.920111 | 160.874685 | 107.619703 | 216.376243 | Non-sig. |
| PF3D7_1344400 | N/A | 450.683284 | 0.761530776 | 0.50019512 | 1.52246741 | 0.127892 | 0.71411411 | 228.292235 | 504.380594 | 440.171198 | 629.889109 | Non-sig. |
| PF3D7_0301200 | FIKK3 | 182.462304 | 0.761313265 | 0.49901449 | 1.52563357 | 0.12710115 | 0.71411411 | 106.300087 | 226.408174 | 164.254948 | 232.886005 | Non-sig. |
| PF3D7_0919800 | N/A | 4858.45766 | 0.760860626 | 0.47113761 | 1.6149435 | 0.10632293 | 0.67059391 | 4738.81777 | 6276.87629 | 2474.01943 | 5944.11714 | Non-sig. |
| PF3D7_0805600 | PAP2 | 2790.76259 | 0.759240531 | 0.42087941 | 1.80393842 | 0.07124096 | 0.58066864 | 2276.87073 | 3133.49771 | 1869.1226 | 3883.5593 | Non-sig. |
| PF3D7_0406700 | N/A | 1104.40757 | 0.756222395 | 0.4496862 | 1.68166691 | 0.09263345 | 0.64025761 | 1004.18286 | 1462.48299 | 638.965166 | 1311.99927 | Non-sig. |
| PF3D7_1117900 | N/A | 2594.679 | 0.753289536 | 0.57175321 | 1.31750819 | 0.18766834 | 0.80247527 | 2966.36552 | 3186.85272 | 898.66735 | 3326.83039 | Non-sig. |
| PF3D7_1231400 | ICM1 | 4684.5463 | 0.75027148 | 0.57416522 | 1.30671705 | 0.19130883 | 0.80801527 | 5368.56657 | 5307.30525 | 1618.29287 | 6444.0205 | Non-sig. |
| PF3D7_0632200 | RIF | 3.46729652 | 0.749901472 | 2.07370988 | 0.36162314 | 0.71763367 | 0.98797939 | 5.2397048 | 3.22147122 | 0 | 5.40801007 | Non-sig. |
| PF3D7_1000200 | RIF | 13.3342839 | 0.749308788 | 1.05702775 | 0.7088828 | 0.47839721 | 0.92880667 | 12.5783678 | 23.6335583 | 7.38617777 | 9.7390318 | Non-sig. |
| PF3D7_1413800 | DPH1 | 350.997962 | 0.748555138 | 0.48113682 | 1.55580513 | 0.11975444 | 0.70181652 | 230.500374 | 340.145409 | 293.196884 | 540.149179 | Non-sig. |
| PF3D7_0934600 | N/A | 987.823982 | 0.748142526 | 0.47507996 | 1.57477179 | 0.11530913 | 0.69029127 | 938.942601 | 1400.48819 | 535.989595 | 1075.87555 | Non-sig. |
| PF3D7_1456800 | VP1 | 1964.63236 | 0.74619418 | 0.60573998 | 1.23187209 | 0.21799687 | 0.83527883 | 601.253664 | 2203.69341 | 2333.43804 | 2720.14433 | Non-sig. |
| PF3D7_0821400 | N/A | 2081.10783 | 0.74545592 | 0.44058961 | 1.69195074 | 0.09065537 | 0.63196333 | 1584.39269 | 2048.63517 | 1525.82129 | 3165.58217 | Non-sig. |
| PF3D7_1420000 | SF3B4 | 823.392242 | 0.744586323 | 0.46897771 | 1.58767955 | 0.11235882 | 0.68406239 | 444.799843 | 1043.24188 | 785.837021 | 1019.69022 | Non-sig. |
| PF3D7_0220000 | LSA3 | 24076.045 | 0.74311886 | 0.50426553 | 1.4736658 | 0.1405716 | 0.73498918 | 24787.0887 | 25679.0773 | 11231.2326 | 34606.7815 | Non-sig. |
| PF3D7_1322400 | N/A | 70.6166961 | 0.74141738 | 0.85370104 | 0.86847426 | 0.38513476 | 0.90292925 | 94.5491785 | 115.946441 | 11.4941479 | 60.4770172 | Non-sig. |
| PF3D7_1028100 | PPO | 55.4998435 | 0.740852813 | 0.61885744 | 1.19713001 | 0.23125588 | 0.83828834 | 46.2484584 | 79.4329996 | 36.9489027 | 59.3690132 | Non-sig. |
| PF3D7_0403100 | N/A | 873.835564 | 0.740698342 | 0.42244663 | 1.75335364 | 0.07954132 | 0.60514927 | 653.306574 | 1197.64026 | 655.327928 | 989.067497 | Non-sig. |
| PF3D7_1409100 | N/A | 2915.65387 | 0.739964212 | 0.62560376 | 1.18280014 | 0.23688837 | 0.83828834 | 3556.72878 | 3990.52342 | 811.561659 | 3303.80164 | Non-sig. |
| PF3D7_1230900 | RIO1 | 1292.52547 | 0.738078366 | 0.54125716 | 1.36363715 | 0.17268179 | 0.78315911 | 1420.86312 | 1624.21385 | 517.471332 | 1607.55356 | Non-sig. |
| PF3D7_0722300 | N/A | 1226.42654 | 0.736822104 | 0.46435672 | 1.58675879 | 0.11256729 | 0.68406239 | 1174.78397 | 1552.19274 | 665.358433 | 1513.37103 | Non-sig. |
| PF3D7_1473400 | N/A | 3627.71269 | 0.736609461 | 0.41484994 | 1.77560459 | 0.07579811 | 0.59646408 | 3037.60762 | 4384.53017 | 2405.00588 | 4683.7071 | Non-sig. |
| PF3D7_0206500 | N/A | 1993.00321 | 0.73611453 | 0.49348192 | 1.49167477 | 0.13578443 | 0.72505481 | 1137.96671 | 1789.23251 | 1852.34186 | 3192.47178 | Non-sig. |
| PF3D7_1136200 | N/A | 5484.76997 | 0.735341543 | 0.41072469 | 1.79035143 | 0.07339743 | 0.58563192 | 3716.71881 | 7128.44004 | 4516.05663 | 6577.86441 | Non-sig. |
| PF3D7_0200100 | VAR | 334.646461 | 0.734362096 | 0.44742283 | 1.64131565 | 0.10073191 | 0.65652696 | 229.693058 | 440.876164 | 272.715715 | 395.300906 | Non-sig. |
| PF3D7_1248200 | RBM22 | 960.901204 | 0.733852674 | 0.49810334 | 1.47329403 | 0.14067177 | 0.73498918 | 461.004711 | 1185.36395 | 981.859275 | 1215.37688 | Non-sig. |
| PF3D7_1222400 | ApiAP2 | 3033.99544 | 0.73344596 | 0.42979787 | 1.70649046 | 0.0879168 | 0.62914105 | 2375.23323 | 3112.4383 | 2182.76908 | 4465.54116 | Non-sig. |
| PF3D7_1366700 | N/A | 359.993557 | 0.733194516 | 0.43963376 | 1.66773933 | 0.09536748 | 0.64772846 | 278.931523 | 469.972723 | 261.990804 | 429.079179 | Non-sig. |
| PF3D7_0407200 | PTH2 | 229.888093 | 0.733147195 | 0.52478771 | 1.39703574 | 0.16240282 | 0.76890988 | 229.233723 | 315.954468 | 116.518006 | 257.846176 | Non-sig. |
| PF3D7_0100200 | RIF | 14.6364625 | 0.733003612 | 1.00577933 | 0.72879169 | 0.4661291 | 0.92538923 | 6.28212684 | 19.3426639 | 15.5911183 | 17.3299411 | Non-sig. |
| PF3D7_0623700 | SUV3 | 602.899455 | 0.732258472 | 0.4232997 | 1.72988188 | 0.08365138 | 0.61468103 | 460.219213 | 730.601377 | 446.000796 | 774.776435 | Non-sig. |
| PF3D7_0730300 | ApiAP2 | 16701.1925 | 0.729909277 | 0.47681082 | 1.53081524 | 0.12581506 | 0.71290716 | 16595.0601 | 18793.8053 | 8533.79536 | 22882.1091 | Non-sig. |
| PF3D7_0723000 | N/A | 221.2092 | 0.729256783 | 0.50038667 | 1.4573865 | 0.14500972 | 0.74131203 | 124.057055 | 260.824784 | 208.564023 | 291.390939 | Non-sig. |
| PF3D7_1404900 | N/A | 4555.86892 | 0.726231631 | 0.43718864 | 1.6611402 | 0.09668529 | 0.64895933 | 4149.76195 | 5384.79659 | 2716.17898 | 5972.73816 | Non-sig. |
| PF3D7_1346800 | P47 | 48.0009808 | 0.725270703 | 0.66021215 | 1.09854189 | 0.27196794 | 0.8556331 | 46.2752187 | 57.9551984 | 26.2774406 | 61.4960653 | Non-sig. |
| PF3D7_1464600 | UIS2 | 16443.3455 | 0.724524844 | 0.42808308 | 1.69248651 | 0.09055325 | 0.63196333 | 14572.816 | 18962.9709 | 10225.5767 | 22012.0184 | Non-sig. |
| PF3D7_1428400 | WDTC1 | 292.281238 | 0.723811037 | 0.59952831 | 1.20730086 | 0.2273163 | 0.83828834 | 336.211247 | 316.417251 | 105.148897 | 411.347556 | Non-sig. |
| PF3D7_1136900 | SUB2 | 8087.61453 | 0.723324186 | 0.47488294 | 1.52316313 | 0.1277179 | 0.71411411 | 8046.29373 | 9282.8391 | 4157.33613 | 10863.9892 | Non-sig. |
| PF3D7_0622900 | AP2Tel | 4878.34521 | 0.720876698 | 0.5458649 | 1.32061376 | 0.18663018 | 0.80156944 | 5438.53517 | 5356.23908 | 1930.54139 | 6788.06521 | Non-sig. |
| PF3D7_0631600 | EPF4 | 93.9219164 | 0.720633391 | 3.18933567 | 0.22595094 | 0.82123958 | 0.99096399 | 104.885 | 233.791446 | 37.0112193 | 0 | Non-sig. |
| PF3D7_0508100 | SET9 | 10687.0679 | 0.720499586 | 0.43317076 | 1.66331538 | 0.09624932 | 0.64875933 | 9560.68746 | 12040.5953 | 6584.72963 | 14562.2591 | Non-sig. |

| | | | | | | | | | | | | |
|---|---|---|---|---|---|---|---|---|---|---|---|---|
| PF3D7_1141300 | APR1 | 3872.19641 | 0.717300823 | 0.52414763 | 1.36850915 | 0.17115276 | 0.78237042 | 4214.71655 | 4994.74761 | 1643.6268 | 4635.69467 | Non-sig. |
| PF3D7_1106800 | pTKL | 741.079026 | 0.716575423 | 0.59014061 | 1.21424523 | 0.22465409 | 0.83828834 | 871.393822 | 908.559251 | 250.538795 | 933.824235 | Non-sig. |
| PF3D7_0703500 | N/A | 9374.31773 | 0.715982583 | 0.51305632 | 1.39552433 | 0.16285778 | 0.76900226 | 4252.8435 | 10089.1581 | 9936.23468 | 13219.0347 | Non-sig. |
| PF3D7_1303700 | N/A | 1046.92733 | 0.715445448 | 0.55212573 | 1.29580168 | 0.19504382 | 0.81344649 | 1177.32271 | 1282.06371 | 408.213187 | 1320.10971 | Non-sig. |
| PF3D7_1431200 | N/A | 1053.96447 | 0.715229751 | 0.54612655 | 1.30964107 | 0.19031729 | 0.805996 | 1174.93388 | 1270.02776 | 421.40304 | 1349.4932 | Non-sig. |
| PF3D7_0109100 | CCp5 | 1104.79879 | 0.714306111 | 0.44734687 | 1.59676117 | 0.11031895 | 0.68040724 | 653.015471 | 1339.93018 | 1020.15428 | 1406.09525 | Non-sig. |
| PF3D7_1319600 | N/A | 584.50642 | 0.714210007 | 0.60789841 | 1.17488381 | 0.2400412 | 0.84181962 | 697.766823 | 657.491199 | 188.120268 | 794.647389 | Non-sig. |
| PF3D7_1035500 | MSP6 | 24730.2182 | 0.712720609 | 0.46297637 | 1.53943195 | 0.12369888 | 0.70887094 | 24195.572 | 32829.5044 | 13290.6195 | 28605.1768 | Non-sig. |
| PF3D7_1005300 | N/A | 18.1891581 | 0.711378337 | 0.91288841 | 0.779261 | 0.43582598 | 0.91522571 | 9.42781656 | 25.786629 | 18.0530799 | 19.4891069 | Non-sig. |
| PF3D7_0523000 | MDR1 | 5725.79947 | 0.710893147 | 0.61165052 | 1.16225381 | 0.24513238 | 0.84331907 | 1734.88779 | 6077.27757 | 6950.56208 | 8140.47043 | Non-sig. |
| PF3D7_1132000 | N/A | 2804.83292 | 0.710775001 | 0.42661995 | 1.66606133 | 0.0957012 | 0.64772846 | 2481.33249 | 3361.53946 | 1774.03419 | 3602.42554 | Non-sig. |
| PF3D7_0506200 | TBP | 132.29817 | 0.710005762 | 0.50458433 | 1.4071102 | 0.15939471 | 0.7667698 | 88.1704213 | 166.444594 | 112.461507 | 162.116158 | Non-sig. |
| PF3D7_0414800 | N/A | 2996.99817 | 0.709193727 | 0.52142665 | 1.36010258 | 0.17379746 | 0.78387252 | 3219.21154 | 4216.50424 | 1330.95895 | 3221.31793 | Non-sig. |
| PF3D7_0215500 | N/A | 1349.5884 | 0.708996953 | 0.58956085 | 1.20258486 | 0.22913701 | 0.83828834 | 1598.53078 | 1787.3844 | 450.922808 | 1561.51561 | Non-sig. |
| PF3D7_0103800 | ARP1 | 250.840946 | 0.708085101 | 0.45493878 | 1.5564404 | 0.1196034 | 0.70181652 | 199.605544 | 318.85194 | 181.443611 | 303.462689 | Non-sig. |
| PF3D7_1417600 | N/A | 9049.9822 | 0.707470344 | 0.49446965 | 1.43076597 | 0.1524973 | 0.75391062 | 8868.39972 | 8416.52813 | 4880.83251 | 14034.1684 | Non-sig. |
| PF3D7_0526600 | N/A | 3386.4765 | 0.707295465 | 0.4904565 | 1.4421166 | 0.14926948 | 0.74818327 | 1893.95032 | 3125.07088 | 3250.89288 | 5275.99192 | Non-sig. |
| PF3D7_1423300 | PP7 | 9641.30323 | 0.70674532 | 0.40363672 | 1.75094406 | 0.07995555 | 0.60514927 | 7181.74476 | 11273.0295 | 7470.0036 | 12640.4351 | Non-sig. |
| PF3D7_0207700 | SERA4 | 9456.04982 | 0.705890758 | 0.44908995 | 1.57182489 | 0.11599117 | 0.6917189 | 5341.33041 | 11703.2448 | 9033.82712 | 11745.797 | Non-sig. |
| PF3D7_1036800 | ACT | 998.732022 | 0.704892125 | 0.48441852 | 1.45513041 | 0.14563316 | 0.7422476 | 512.512212 | 1191.08576 | 1006.04657 | 1285.28354 | Non-sig. |
| PF3D7_1022500 | CS | 2571.27135 | 0.704741329 | 0.40497119 | 1.74022583 | 0.08181937 | 0.60888892 | 1990.71132 | 3185.59413 | 1920.22109 | 3188.55887 | Non-sig. |
| PF3D7_1240000 | N/A | 217.394089 | 0.703880387 | 0.53459272 | 1.31666662 | 0.1879504 | 0.80303323 | 106.227676 | 254.35555 | 224.181269 | 284.81186 | Non-sig. |
| PF3D7_0729700 | N/A | 416.467806 | 0.703628789 | 0.45034046 | 1.56243743 | 0.11818497 | 0.697844 | 293.638203 | 589.091949 | 340.007209 | 443.133864 | Non-sig. |
| PF3D7_1229900 | N/A | 325.881731 | 0.70313917 | 0.45683246 | 1.53916203 | 0.12376475 | 0.70887094 | 270.974584 | 451.858821 | 225.169552 | 355.523968 | Non-sig. |
| PF3D7_1033100 | AdoMetDC/OI | 31673.9029 | 0.702368784 | 0.42822104 | 1.64020148 | 0.10096328 | 0.65652696 | 28542.2891 | 37025.6521 | 19683.0851 | 41444.5854 | Non-sig. |
| PF3D7_1401500 | N/A | 134.922557 | 0.701533311 | 0.54733332 | 1.28172958 | 0.19993753 | 0.81595849 | 70.5070719 | 174.909157 | 134.686109 | 159.587892 | Non-sig. |
| PF3D7_0214700 | N/A | 1688.68492 | 0.699948728 | 0.5219666 | 1.34098374 | 0.17992573 | 0.79619661 | 1812.45942 | 2384.02036 | 761.762124 | 1796.49776 | Non-sig. |
| PF3D7_1371500 | N/A | 112.592567 | 0.699496402 | 0.52138386 | 1.34161498 | 0.17972087 | 0.79612136 | 90.408758 | 156.71232 | 81.2898018 | 121.959388 | Non-sig. |
| PF3D7_0909100 | N/A | 566.907733 | 0.699323324 | 0.43012433 | 1.6258632 | 0.10397876 | 0.6615241 | 468.528782 | 727.433327 | 395.880545 | 675.78828 | Non-sig. |
| PF3D7_1134300 | EMMA1 | 6401.95953 | 0.696155722 | 0.53405507 | 1.30352798 | 0.19239457 | 0.81090408 | 7077.67286 | 6870.97893 | 2696.10818 | 8963.07815 | Non-sig. |
| PF3D7_0502000 | VPS11 | 979.109367 | 0.695250053 | 0.44035684 | 1.57883333 | 0.11437429 | 0.68780363 | 890.210671 | 1212.18473 | 605.3826 | 1208.65947 | Non-sig. |
| PF3D7_0208000 | SERA1 | 5512.45357 | 0.695051696 | 0.40860678 | 1.70102829 | 0.08893767 | 0.62981332 | 4593.62952 | 6874.79159 | 3825.89476 | 6755.49839 | Non-sig. |
| PF3D7_1148700 | GEXP12 | 228.001155 | 0.693921629 | 0.61705943 | 1.12456207 | 0.26077465 | 0.85030738 | 267.505764 | 329.357239 | 81.3155889 | 233.826028 | Non-sig. |
| PF3D7_1367300 | N/A | 7.85066208 | 0.689664734 | 1.31307466 | 0.52522888 | 0.59942409 | 0.95655566 | 6.28392764 | 15.0430607 | 5.74395369 | 4.33170632 | Non-sig. |
| PF3D7_0103500 | N/A | 354.906188 | 0.688163274 | 0.47130669 | 1.46011778 | 0.14425771 | 0.74129712 | 225.525731 | 373.274592 | 317.906118 | 502.918312 | Non-sig. |
| PF3D7_1247700 | N/A | 954.46913 | 0.687130895 | 0.49839608 | 1.37868438 | 0.16799209 | 0.77528056 | 996.01062 | 1181.97868 | 467.162648 | 1172.72457 | Non-sig. |
| PF3D7_0602800 | JmjC2 | 50.5915214 | 0.686452267 | 0.63751659 | 1.07675985 | 0.28158757 | 0.86390383 | 43.1443773 | 72.9690655 | 34.4925406 | 51.760102 | Non-sig. |
| PF3D7_0403800 | N/A | 3798.61693 | 0.686125242 | 0.50124927 | 1.36883041 | 0.17105229 | 0.78237042 | 3788.7261 | 3479.57863 | 2035.59469 | 5890.56828 | Non-sig. |
| PF3D7_1407200 | N/A | 1013.54056 | 0.685586554 | 0.4789715 | 1.43137234 | 0.15232353 | 0.75391062 | 1017.68444 | 1188.20309 | 537.010937 | 1311.26377 | Non-sig. |
| PF3D7_1002700 | N/A | 2001.44098 | 0.684463515 | 0.4519461 | 1.51448041 | 0.12990404 | 0.71812182 | 1920.33451 | 2422.29603 | 1150.78074 | 2512.35264 | Non-sig. |
| PF3D7_1148000 | N/A | 2125.05015 | 0.684052698 | 0.52345993 | 1.30679095 | 0.19128372 | 0.80801527 | 2338.14594 | 2652.72637 | 923.299048 | 2586.02924 | Non-sig. |
| PF3D7_0906600 | N/A | 1756.21287 | 0.683374259 | 0.49810675 | 1.37194338 | 0.17008106 | 0.77948916 | 863.653735 | 1931.05177 | 1831.67358 | 2398.4724 | Non-sig. |
| PF3D7_1428100 | WBP11 | 1280.4099 | 0.683199771 | 0.51425534 | 1.32852247 | 0.18400556 | 0.80052251 | 1379.63028 | 1656.8059 | 586.377836 | 1498.8256 | Non-sig. |
| PF3D7_0423800 | CyRPA | 3263.34945 | 0.683123063 | 0.43140358 | 1.58348954 | 0.11330993 | 0.68498781 | 2974.55771 | 4164.88862 | 2035.4669 | 3878.48459 | Non-sig. |
| PF3D7_1431900 | ISC3 | 306.536079 | 0.682620574 | 0.56603806 | 1.20596232 | 0.22783202 | 0.83828834 | 125.291496 | 395.904397 | 344.957925 | 359.990497 | Non-sig. |
| PF3D7_1204000 | N/A | 5.73228888 | 0.681389635 | 1.59387596 | 0.42750481 | 0.66901168 | 0.97883902 | 7.24034012 | 11.861184 | 1.63768286 | 2.1899485 | Non-sig. |
| PF3D7_1235500 | MT-A70.2 | 514.202362 | 0.681360511 | 0.55657238 | 1.22420827 | 0.22087367 | 0.83527883 | 581.829499 | 598.802612 | 208.591885 | 667.585454 | Non-sig. |
| PF3D7_1237900 | N/A | 10980.6837 | 0.680368372 | 0.4536518 | 1.49975899 | 0.13367684 | 0.72178914 | 10692.1535 | 12917.4077 | 6185.05439 | 14128.119 | Non-sig. |
| PF3D7_1423500 | N/A | 1113.60691 | 0.679828617 | 0.71441922 | 0.95158221 | 0.34130891 | 0.8930363 | 1503.75117 | 1324.7218 | 208.649024 | 1417.30565 | Non-sig. |
| PF3D7_0530400 | N/A | 158.848412 | 0.677993494 | 0.57174376 | 1.18583454 | 0.23568765 | 0.83828834 | 74.8200151 | 164.11454 | 169.219792 | 227.239302 | Non-sig. |
| PF3D7_1448500 | N/A | 10648.9131 | 0.677940169 | 0.56380628 | 1.20243458 | 0.2291952 | 0.83828834 | 12394.1718 | 11385.4505 | 3990.18474 | 14825.8452 | Non-sig. |
| PF3D7_1445600 | N/A | 3529.57011 | 0.677141969 | 0.41016294 | 1.65090969 | 0.09875702 | 0.65401834 | 2928.01522 | 4126.00666 | 2504.38514 | 4559.87342 | Non-sig. |
| PF3D7_0308700 | N/A | 2513.74729 | 0.676980548 | 0.41240364 | 1.64154842 | 0.10068362 | 0.65652696 | 1750.27266 | 3037.17044 | 2118.667 | 3148.87907 | Non-sig. |
| PF3D7_0220100 | N/A | 771.513345 | 0.676622433 | 0.44958947 | 1.5049784 | 0.13232963 | 0.72037933 | 672.032998 | 790.635812 | 515.834702 | 1107.54987 | Non-sig. |

| | | | | | | | | | | | | |
|---|---|---|---|---|---|---|---|---|---|---|---|---|
| PF3D7_0202900 | DHHC12 | 204.030668 | 0.676453922 | 0.47132483 | 1.43521808 | 0.15122496 | 0.75299798 | 141.615677 | 259.913003 | 172.36762 | 242.226372 | Non-sig. |
| PF3D7_1245800 | EpsL | 871.522977 | 0.67576322 | 0.48821359 | 1.38415488 | 0.16631103 | 0.77284241 | 891.888128 | 1013.80094 | 450.639066 | 1129.76378 | Non-sig. |
| PF3D7_0937100 | N/A | 6.5986399 | 0.675569597 | 1.57034419 | 0.43020479 | 0.66704668 | 0.97883902 | 9.44216963 | 12.8878175 | 0.82081545 | 3.24375699 | Non-sig. |
| PF3D7_1014100 | MSA180 | 8215.80254 | 0.675286246 | 0.48466311 | 1.39331061 | 0.16352588 | 0.77052723 | 4294.336 | 8573.59408 | 8359.98065 | 11635.2994 | Non-sig. |
| PF3D7_0517500 | N/A | 86.708954 | 0.675211397 | 0.60107052 | 1.12334805 | 0.2612897 | 0.85030738 | 89.4832095 | 92.2735107 | 44.3508056 | 120.72829 | Non-sig. |
| PF3D7_0721200 | N/A | 380.043706 | 0.675101677 | 0.44868054 | 1.50463773 | 0.13241724 | 0.72037933 | 316.961385 | 416.272123 | 268.581426 | 518.359891 | Non-sig. |
| PF3D7_0525300 | N/A | 517.154409 | 0.674553447 | 0.4391715 | 1.53596819 | 0.12454618 | 0.7110406 | 445.581515 | 595.779593 | 351.413288 | 675.843238 | Non-sig. |
| PF3D7_1217400 | N/A | 4421.11658 | 0.671090545 | 0.51138417 | 1.31230216 | 0.1894182 | 0.80547152 | 4800.24668 | 5822.25909 | 2022.20364 | 5039.75692 | Non-sig. |
| PF3D7_1304700 | CYC4 | 140.971863 | 0.669661329 | 0.49150361 | 1.36247489 | 0.17304806 | 0.78315911 | 106.854721 | 167.610393 | 110.781391 | 178.640946 | Non-sig. |
| PF3D7_1201100 | N/A | 193.563805 | 0.667611363 | 0.46998538 | 1.42049391 | 0.15546394 | 0.75842353 | 142.174724 | 245.678618 | 156.88091 | 229.520968 | Non-sig. |
| PF3D7_0919500 | MFS3 | 60.8681925 | 0.667589876 | 0.66448055 | 1.00467933 | 0.31505128 | 0.88486757 | 27.2970767 | 81.6063131 | 66.4945148 | 68.0748653 | Non-sig. |
| PF3D7_1023800 | N/A | 508.95248 | 0.666778091 | 0.73726572 | 0.90439318 | 0.36578695 | 0.89567376 | 697.683277 | 679.254203 | 89.5165152 | 569.355925 | Non-sig. |
| PF3D7_0615400 | N/A | 13775.0671 | 0.666042884 | 0.66799454 | 0.99707833 | 0.31872649 | 0.88651753 | 18129.015 | 17366.9304 | 3172.73059 | 16431.5925 | Non-sig. |
| PF3D7_0314800 | N/A | 750.623231 | 0.665209141 | 0.52723357 | 1.26169723 | 0.20705775 | 0.82391338 | 825.657611 | 885.089419 | 335.938068 | 955.807824 | Non-sig. |
| PF3D7_0315800 | N/A | 356.936379 | 0.664870314 | 0.45485416 | 1.46172195 | 0.14381742 | 0.7412338 | 318.439447 | 419.723183 | 234.004008 | 455.57888 | Non-sig. |
| PF3D7_0914500 | N/A | 4145.07812 | 0.664580367 | 0.50682519 | 1.31126152 | 0.18976942 | 0.80574162 | 4476.43674 | 4702.5066 | 1937.80412 | 5463.56503 | Non-sig. |
| PF3D7_1122800 | CDPK6 | 1927.19462 | 0.664516584 | 0.49392252 | 1.34538629 | 0.17850052 | 0.79309819 | 1988.52047 | 1997.26488 | 993.960405 | 2729.0327 | Non-sig. |
| PF3D7_1133500 | N/A | 112.50383 | 0.663904215 | 0.52419789 | 1.26651447 | 0.20532895 | 0.82391338 | 75.5006945 | 126.747461 | 98.4894068 | 149.277756 | Non-sig. |
| PF3D7_0500100 | VAR | 1892.06876 | 0.662851 | 0.41349065 | 1.60306163 | 0.10892105 | 0.6779986 | 1596.704 | 2366.17247 | 1333.24993 | 2272.14862 | Non-sig. |
| PF3D7_0813400 | N/A | 3780.56717 | 0.661875983 | 0.40788734 | 1.62269313 | 0.10465502 | 0.66323241 | 3110.66372 | 4416.56184 | 2745.93797 | 4849.10514 | Non-sig. |
| PF3D7_0113400 | N/A | 31.2693839 | 0.661132706 | 0.783495 | 0.84382505 | 0.39876718 | 0.90645858 | 34.6968696 | 36.4931314 | 13.9592174 | 39.9283173 | Non-sig. |
| PF3D7_0214300 | N/A | 705.949669 | 0.660396723 | 0.65815671 | 1.00340346 | 0.31566623 | 0.88486757 | 914.002887 | 836.570647 | 180.732407 | 892.492734 | Non-sig. |
| PF3D7_1302200 | UIS3 | 40.6412402 | 0.659788626 | 0.84779098 | 0.77824445 | 0.43642492 | 0.91522571 | 53.4403613 | 40.8253897 | 9.84765942 | 58.4515502 | Non-sig. |
| PF3D7_1009400 | ZNF2 | 250.065811 | 0.659715575 | 0.46540987 | 1.41749374 | 0.15633862 | 0.75894163 | 169.166504 | 290.925899 | 218.395285 | 321.775558 | Non-sig. |
| PF3D7_0819700 | N/A | 254.237499 | 0.659509049 | 0.54009708 | 1.22109353 | 0.22205061 | 0.83671702 | 122.032871 | 279.021322 | 271.820762 | 344.075039 | Non-sig. |
| PF3D7_1341500 | ISC1 | 1585.57522 | 0.659231187 | 0.43392113 | 1.51924196 | 0.1287016 | 0.71411411 | 1015.05476 | 2061.61396 | 1443.65553 | 1821.97663 | Non-sig. |
| PF3D7_1013000 | N/A | 705.510194 | 0.657227692 | 0.43689468 | 1.50431609 | 0.1325 | 0.72037933 | 631.304349 | 839.172957 | 463.996924 | 887.566544 | Non-sig. |
| PF3D7_1307500 | N/A | 2457.30472 | 0.65671309 | 0.52388813 | 1.25353687 | 0.21001039 | 0.82566351 | 2697.23964 | 3462.4927 | 1118.17531 | 2551.31123 | Non-sig. |
| PF3D7_1222500 | DPH6 | 863.204357 | 0.656178135 | 0.45161728 | 1.45295178 | 0.14623714 | 0.7426703 | 759.124279 | 1248.86674 | 581.501199 | 863.325209 | Non-sig. |
| PF3D7_1121300 | TKL2 | 4220.60687 | 0.655923614 | 0.45324229 | 1.44718098 | 0.14784624 | 0.74652201 | 2663.14297 | 4184.3036 | 3891.28024 | 6143.70065 | Non-sig. |
| PF3D7_0823500 | IMC1i | 121.076644 | 0.655422955 | 0.5390563 | 1.21587107 | 0.22403404 | 0.83828834 | 117.713304 | 156.722919 | 70.6124007 | 139.257952 | Non-sig. |
| PF3D7_0937500 | RIF | 2.66249775 | 0.655223477 | 2.29558215 | 0.28542802 | 0.77531624 | 0.99028237 | 4.1910439 | 4.29544914 | 0 | 2.16349797 | Non-sig. |
| PF3D7_0709900 | N/A | 996.871112 | 0.653838265 | 0.52793283 | 1.23848761 | 0.21553531 | 0.83503497 | 1108.85007 | 1174.47086 | 441.138762 | 1263.02475 | Non-sig. |
| PF3D7_0507800 | N/A | 11505.9683 | 0.652955063 | 0.56739767 | 1.15078911 | 0.249819 | 0.84896798 | 13704.1553 | 14394.4233 | 4187.86183 | 13737.4327 | Non-sig. |
| PF3D7_0508800 | SSB | 4143.90868 | 0.652542271 | 0.41718515 | 1.56415509 | 0.11778115 | 0.69721177 | 3574.58933 | 5467.96924 | 2870.40861 | 4662.66755 | Non-sig. |
| PF3D7_1142100 | N/A | 6205.74113 | 0.65211045 | 0.47786788 | 1.36462499 | 0.17237095 | 0.78315911 | 6310.03651 | 6545.4624 | 3343.60208 | 8623.86352 | Non-sig. |
| PF3D7_0926800 | N/A | 1806.96794 | 0.652027157 | 0.55537413 | 1.17403228 | 0.24038209 | 0.84181962 | 2107.45975 | 2266.86021 | 703.917345 | 2149.63445 | Non-sig. |
| PF3D7_1358600 | N/A | 1062.62865 | 0.651566263 | 0.60313975 | 1.08029069 | 0.28001275 | 0.86300615 | 1306.39251 | 1182.40435 | 347.411748 | 1414.30597 | Non-sig. |
| PF3D7_1111900 | N/A | 298.195579 | 0.651285885 | 0.47308465 | 1.37667938 | 0.1686114 | 0.77543678 | 183.795954 | 371.486229 | 279.951299 | 357.548834 | Non-sig. |
| PF3D7_0107300 | N/A | 92.1617439 | 0.651212792 | 0.55261759 | 1.17841487 | 0.23863125 | 0.84066259 | 86.1348534 | 114.876591 | 57.469572 | 110.16596 | Non-sig. |
| PF3D7_0527200 | USP14 | 6786.83822 | 0.650433553 | 0.54758184 | 1.18782893 | 0.23490081 | 0.83828834 | 7866.5378 | 8781.11648 | 2698.60241 | 7801.09618 | Non-sig. |
| PF3D7_0830300 | SIAP2 | 45.0412836 | 0.650183375 | 0.64876569 | 1.0021852 | 0.31625415 | 0.88585961 | 39.8446825 | 54.7818027 | 30.3671755 | 55.1714737 | Non-sig. |
| PF3D7_0529700 | N/A | 273.859502 | 0.646756246 | 0.46428718 | 1.39300906 | 0.16361705 | 0.77052723 | 181.165573 | 343.293327 | 245.59422 | 325.38489 | Non-sig. |
| PF3D7_1372600 | RIF | 15.9223121 | 0.645468123 | 0.94119208 | 0.6857985 | 0.49284018 | 0.93170321 | 12.537765 | 23.6549211 | 12.3046329 | 15.1919293 | Non-sig. |
| PF3D7_1249600 | LRR12 | 678.570177 | 0.644732618 | 0.42595603 | 1.51361309 | 0.130124 | 0.71812182 | 520.032845 | 755.364452 | 538.776083 | 900.107329 | Non-sig. |
| PF3D7_0311400 | PKRP | 844.295582 | 0.64467664 | 0.63256988 | 1.01913901 | 0.30813698 | 0.87724478 | 1073.12402 | 942.766871 | 244.807679 | 1116.48376 | Non-sig. |
| PF3D7_1359100 | N/A | 668.802825 | 0.644411464 | 0.42318185 | 1.52277671 | 0.12781458 | 0.71411411 | 523.060689 | 867.013112 | 520.684715 | 764.452784 | Non-sig. |
| PF3D7_0705100 | N/A | 1825.7303 | 0.644218814 | 0.46516166 | 1.38493531 | 0.16607225 | 0.77284241 | 1779.6028 | 1893.61329 | 1070.21746 | 2559.48766 | Non-sig. |
| PF3D7_1148500 | N/A | 10.3427606 | 0.644158912 | 1.35221435 | 0.47637337 | 0.6338084 | 0.9668494 | 15.4658951 | 15.2224937 | 0.81342545 | 9.86922802 | Non-sig. |
| PF3D7_1136000 | N/A | 3368.69814 | 0.643591667 | 0.49085366 | 1.31116812 | 0.18980097 | 0.80574162 | 3445.49759 | 3299.00275 | 1813.92625 | 4916.36598 | Non-sig. |
| PF3D7_0708200 | N/A | 61.8635276 | 0.642848878 | 0.63463685 | 1.01293972 | 0.31108895 | 0.8804206 | 63.1540464 | 64.3798998 | 33.672775 | 86.2473891 | Non-sig. |
| PF3D7_1459600 | N/A | 2045.92619 | 0.642744197 | 0.40843471 | 1.57367673 | 0.11556221 | 0.69102547 | 1518.64659 | 2545.82659 | 1676.40996 | 2442.82162 | Non-sig. |
| PF3D7_1211800 | PfpUB | 23940.3722 | 0.6411729 | 0.44779001 | 1.43186065 | 0.15218371 | 0.75391062 | 23148.637 | 31986.4726 | 14264.4594 | 26361.9197 | Non-sig. |
| PF3D7_1326500 | N/A | 830.724873 | 0.640797039 | 0.45179363 | 1.41834015 | 0.15609148 | 0.75894163 | 763.066243 | 869.878093 | 535.602647 | 1154.35251 | Non-sig. |

| | | | | | | | | | | | | |
|---|---|---|---|---|---|---|---|---|---|---|---|---|
| PF3D7_1146200 | N/A | 176.803113 | 0.640567217 | 0.65056931 | 0.98462563 | 0.324808 | 0.88694295 | 218.551022 | 251.871527 | 58.2050795 | 178.584822 | Non-sig. |
| PF3D7_1203400 | MFS4 | 2405.57319 | 0.639129512 | 0.46069442 | 1.38731768 | 0.16534491 | 0.77284241 | 2401.8581 | 2800.94047 | 1361.0712 | 3058.423 | Non-sig. |
| PF3D7_1337500 | N/A | 5048.01093 | 0.63883924 | 0.49329076 | 1.29505616 | 0.19530086 | 0.81344649 | 5291.03377 | 5181.10175 | 2605.91764 | 7113.99057 | Non-sig. |
| PF3D7_0612700 | P12 | 5956.91461 | 0.638574 | 0.41360277 | 1.54393067 | 0.12260513 | 0.7079393 | 4180.35496 | 7731.75175 | 5138.84098 | 6776.71076 | Non-sig. |
| PF3D7_0806300 | FLP | 4152.72546 | 0.63845473 | 0.42678191 | 1.49597421 | 0.13466038 | 0.72209582 | 2739.59039 | 4643.6181 | 3757.26549 | 5470.42784 | Non-sig. |
| PF3D7_0718600 | N/A | 1238.65206 | 0.638375777 | 0.61157113 | 1.04382916 | 0.29656444 | 0.87299304 | 1545.0184 | 1685.40339 | 393.436681 | 1330.74978 | Non-sig. |
| PF3D7_1328300 | N/A | 8617.04861 | 0.638273896 | 0.40240233 | 1.58615856 | 0.11270335 | 0.68406239 | 6446.76886 | 10558.6036 | 7035.90921 | 10426.9127 | Non-sig. |
| PF3D7_1335300 | RH2b | 14370.121 | 0.637450761 | 0.5615018 | 1.1352604 | 0.25626623 | 0.85030738 | 16140.5552 | 11925.0878 | 6352.03242 | 23062.8088 | Non-sig. |
| PF3D7_1361500 | N/A | 4812.04886 | 0.637232604 | 0.42934204 | 1.48420734 | 0.13775392 | 0.72905594 | 3231.41655 | 6584.02215 | 4300.80528 | 5131.95147 | Non-sig. |
| PF3D7_1339700 | N/A | 8686.65236 | 0.636257556 | 0.60024025 | 1.06000483 | 0.2891424 | 0.86724628 | 10832.0822 | 10397.9155 | 2771.63895 | 10744.9728 | Non-sig. |
| PF3D7_1218300 | AP2-MU | 1586.16745 | 0.635137114 | 0.47237771 | 1.34455354 | 0.17876946 | 0.79309819 | 888.820869 | 1705.82211 | 1595.90339 | 2154.12344 | Non-sig. |
| PF3D7_0516500 | MFS1 | 66.897144 | 0.63489178 | 0.57721987 | 1.09991325 | 0.27136992 | 0.8556331 | 55.6204168 | 77.3204578 | 49.2515098 | 85.3961918 | Non-sig. |
| PF3D7_0816500 | HSP20 | 12.9743593 | 0.634817986 | 1.07932676 | 0.58816107 | 0.55642418 | 0.94243456 | 15.5385218 | 18.3225576 | 4.91453999 | 13.1218179 | Non-sig. |
| PF3D7_1319100 | N/A | 504.694946 | 0.634785143 | 0.48096787 | 1.31980779 | 0.1868992 | 0.80177487 | 422.150796 | 443.127265 | 368.768827 | 784.732895 | Non-sig. |
| PF3D7_1119700 | N/A | 662.40168 | 0.633745464 | 0.51645199 | 1.227114 | 0.21977975 | 0.83527883 | 321.018166 | 719.991283 | 716.977904 | 891.619367 | Non-sig. |
| PF3D7_1036000 | MSP11 | 9847.24914 | 0.632526069 | 0.47706734 | 1.32586327 | 0.18488499 | 0.80052251 | 10287.243 | 12358.9748 | 5158.17974 | 11584.599 | Non-sig. |
| PF3D7_0316100 | N/A | 1148.06 | 0.632179383 | 0.54105833 | 1.16841262 | 0.24264037 | 0.8422984 | 1301.27443 | 1192.29474 | 500.113394 | 1598.55744 | Non-sig. |
| PF3D7_0313700 | N/A | 616.312017 | 0.631567265 | 0.46786093 | 1.34990383 | 0.17704683 | 0.79022251 | 358.315513 | 767.560734 | 608.392459 | 730.97936 | Non-sig. |
| PF3D7_1413600 | N/A | 54.568597 | 0.631497297 | 0.63763068 | 0.99038098 | 0.32198794 | 0.88694295 | 54.6205953 | 65.4909001 | 31.1976163 | 66.9652761 | Non-sig. |
| PF3D7_1332400 | N/A | 1152.48622 | 0.631434667 | 0.41935131 | 1.50574148 | 0.13213355 | 0.72037933 | 962.558255 | 1292.76957 | 846.004351 | 1508.61272 | Non-sig. |
| PF3D7_1409400 | N/A | 5769.9336 | 0.629070833 | 0.56312681 | 1.11710332 | 0.26395015 | 0.85144857 | 2192.20727 | 6350.88148 | 6870.39611 | 7666.24955 | Non-sig. |
| PF3D7_1246100 | ALAS | 139.422355 | 0.628981927 | 0.50703021 | 1.24052161 | 0.21478253 | 0.83503497 | 91.5319964 | 172.77469 | 127.289456 | 166.09328 | Non-sig. |
| PF3D7_0602100 | MTR4 | 673.539061 | 0.628795813 | 0.42180494 | 1.49072651 | 0.13603331 | 0.72505481 | 552.141345 | 800.213459 | 506.008255 | 835.793185 | Non-sig. |
| PF3D7_1368900 | N/A | 1877.81768 | 0.628368162 | 0.55514457 | 1.13190004 | 0.25767646 | 0.85030738 | 738.105585 | 2326.0335 | 2211.86034 | 2235.27131 | Non-sig. |
| PF3D7_0100600 | RIF | 4.47493027 | 0.627382824 | 1.71348468 | 0.3661444 | 0.71425733 | 0.9875928 | 6.2875722 | 4.29731926 | 0.82064917 | 6.49418043 | Non-sig. |
| PF3D7_0518600 | WDR26 | 2480.4834 | 0.626631334 | 0.48260642 | 1.29843141 | 0.19413913 | 0.81267745 | 2610.97303 | 3058.22393 | 1289.65464 | 2963.082 | Non-sig. |
| PF3D7_0916400 | N/A | 1103.29174 | 0.624822796 | 0.45364234 | 1.37734675 | 0.16840507 | 0.77543678 | 962.389245 | 1048.96388 | 773.848278 | 1627.96555 | Non-sig. |
| PF3D7_0606000 | KIC1 | 5431.09432 | 0.623027902 | 0.41730606 | 1.49297593 | 0.13544349 | 0.72467461 | 4838.13921 | 6282.42887 | 3714.63818 | 6889.17103 | Non-sig. |
| PF3D7_0730500 | N/A | 7241.67589 | 0.62247891 | 0.51082731 | 1.21857015 | 0.22300738 | 0.83793839 | 7944.10999 | 7546.13214 | 3462.7151 | 10013.7463 | Non-sig. |
| PF3D7_0425500 | N/A | 14.2098121 | 0.62194289 | 0.9901442 | 0.62813365 | 0.52991639 | 0.93272327 | 8.36924806 | 19.3472014 | 13.9479896 | 15.1748092 | Non-sig. |
| PF3D7_1212900 | BDP2 | 3993.66024 | 0.621871218 | 0.47566769 | 1.30736486 | 0.19108883 | 0.80801527 | 3708.00024 | 3502.27131 | 2584.25801 | 6180.11141 | Non-sig. |
| PF3D7_1435700 | N/A | 1463.86002 | 0.621409766 | 0.4174672 | 1.48852356 | 0.13661287 | 0.72522545 | 1272.59138 | 1802.00181 | 1034.32809 | 1746.51881 | Non-sig. |
| PF3D7_0412000 | N/A | 2356.00721 | 0.621177353 | 0.42018966 | 1.47832613 | 0.13932051 | 0.73226195 | 1601.84344 | 2885.91497 | 2110.91256 | 2825.35785 | Non-sig. |
| PF3D7_1210200 | N/A | 139.823957 | 0.621138135 | 0.51521722 | 1.20558498 | 0.22797755 | 0.83828834 | 132.696595 | 163.070459 | 87.8861857 | 175.642587 | Non-sig. |
| PF3D7_0208100 | N/A | 4488.62687 | 0.619341558 | 0.53117109 | 1.16599259 | 0.24361744 | 0.84313671 | 5154.3418 | 5417.86852 | 1925.50366 | 5456.79348 | Non-sig. |
| PF3D7_0613300 | ROP14 | 7675.99971 | 0.618530867 | 0.53650716 | 1.15288466 | 0.24895773 | 0.8482655 | 8895.10241 | 9098.4845 | 3215.92807 | 9494.48388 | Non-sig. |
| PF3D7_0424100 | RH5 | 4410.40056 | 0.618278609 | 0.52910624 | 1.16853397 | 0.24259145 | 0.8422984 | 5051.11591 | 5346.66355 | 1908.45252 | 5335.37026 | Non-sig. |
| PF3D7_0419300 | N/A | 1210.34366 | 0.617817815 | 0.4791914 | 1.28929238 | 0.19729646 | 0.81471289 | 1254.36684 | 1551.65772 | 656.181625 | 1379.16846 | Non-sig. |
| PF3D7_1460500 | N/A | 2553.93318 | 0.616888096 | 0.4756427 | 1.29695694 | 0.194646 | 0.81327643 | 2649.99152 | 2872.31766 | 1382.57564 | 3310.8479 | Non-sig. |
| PF3D7_1216200 | N/A | 2653.12177 | 0.614776653 | 0.43033152 | 1.42861173 | 0.15311586 | 0.75391062 | 2230.78454 | 3761.67016 | 1961.79192 | 2658.24046 | Non-sig. |
| PF3D7_1246200 | ACT1 | 76814.0586 | 0.611778502 | 0.40229453 | 1.5207229 | 0.12832939 | 0.71411411 | 63734.2425 | 89617.8782 | 57800.3158 | 96103.798 | Non-sig. |
| PF3D7_0715100 | MYCBP | 50.2855121 | 0.611057542 | 0.73582475 | 0.83043896 | 0.40629065 | 0.90723586 | 60.1811827 | 73.2542682 | 19.6646767 | 48.0419206 | Non-sig. |
| PF3D7_0215200 | N/A | 169.66178 | 0.610848712 | 0.60366667 | 1.01189736 | 0.31158713 | 0.8804206 | 199.94503 | 228.5624 | 68.9855441 | 181.154145 | Non-sig. |
| PF3D7_0422000 | N/A | 505.393448 | 0.610431289 | 0.47489506 | 1.28540247 | 0.19865168 | 0.81528784 | 298.807827 | 668.554295 | 500.949625 | 553.262046 | Non-sig. |
| PF3D7_0414700 | N/A | 2844.02633 | 0.61027737 | 0.40749716 | 1.4976236 | 0.13423107 | 0.72178914 | 2152.78807 | 3597.62814 | 2349.75753 | 3275.93158 | Non-sig. |
| PF3D7_0403400 | N/A | 2034.31318 | 0.609780508 | 0.43320864 | 1.40759083 | 0.15925226 | 0.7667698 | 1465.60202 | 2049.50131 | 1755.62004 | 2866.52934 | Non-sig. |
| PF3D7_0523700 | N/A | 241.944412 | 0.609111194 | 0.50084004 | 1.21617813 | 0.22391707 | 0.83828834 | 142.249004 | 262.805288 | 240.682158 | 322.041197 | Non-sig. |
| PF3D7_0318700 | N/A | 1620.61129 | 0.609089526 | 0.55331689 | 1.10079692 | 0.27098505 | 0.8556331 | 1917.52219 | 1902.69018 | 649.942896 | 2012.28989 | Non-sig. |
| PF3D7_0815700 | Ub | 83.1862704 | 0.608985219 | 0.62787936 | 0.969908 | 0.33209235 | 0.88994703 | 37.8527804 | 94.4516535 | 93.6095847 | 106.831063 | Non-sig. |
| PF3D7_0813800 | GMD | 345.83489 | 0.60854296 | 0.47516924 | 1.28068677 | 0.20030371 | 0.81595849 | 215.187659 | 459.595945 | 332.461879 | 376.094076 | Non-sig. |
| PF3D7_1328800 | SIR2A | 2097.40011 | 0.607321024 | 0.42070848 | 1.44356736 | 0.14886071 | 0.74754886 | 1600.94646 | 2865.4395 | 1723.682 | 2199.53248 | Non-sig. |
| PF3D7_1237100 | N/A | 2381.79003 | 0.606975593 | 0.41952843 | 1.44680445 | 0.1479517 | 0.74652201 | 2041.96531 | 3184.35119 | 1734.20284 | 2566.64079 | Non-sig. |
| PF3D7_0507500 | SUB1 | 11328.3571 | 0.606505333 | 0.41327291 | 1.46756616 | 0.14222211 | 0.73871836 | 7911.06667 | 13120.1465 | 10051.961 | 14230.2542 | Non-sig. |
| PF3D7_1329800 | N/A | 300.90718 | 0.60588546 | 0.53036898 | 1.14238479 | 0.25329411 | 0.85030738 | 147.48818 | 348.638762 | 329.386363 | 378.115414 | Non-sig. |

| | | | | | | | | | | | | |
|---|---|---|---|---|---|---|---|---|---|---|---|---|
| PF3D7_1372800 | N/A | 2.2250329 | 0.605479518 | 2.20374448 | 0.27475033 | 0.78350806 | 0.99028237 | 1.04600195 | 3.22466364 | 2.46136264 | 2.16810339 | Non-sig. |
| PF3D7_1318800 | SEC63 | 6309.85908 | 0.605403561 | 0.48751049 | 1.24182674 | 0.21430049 | 0.83421309 | 3213.83509 | 7558.02494 | 6795.80243 | 7671.77388 | Non-sig. |
| PF3D7_0723100 | CPH1 | 48.6781825 | 0.60537663 | 0.62984629 | 0.96114978 | 0.33647686 | 0.8920875 | 41.9880088 | 57.9863373 | 35.2984474 | 59.4399365 | Non-sig. |
| PF3D7_1455100 | PTP1 | 1046.53376 | 0.605263983 | 0.63812091 | 0.94850988 | 0.34286995 | 0.8930363 | 1365.35917 | 1290.29445 | 295.430179 | 1235.05123 | Non-sig. |
| PF3D7_0916300 | N/A | 245.410763 | 0.605153734 | 0.48993431 | 1.23517322 | 0.21676604 | 0.83527883 | 149.534565 | 316.041696 | 239.561308 | 276.505482 | Non-sig. |
| PF3D7_0321100 | N/A | 3616.43538 | 0.604731455 | 0.53712067 | 1.12587635 | 0.26021786 | 0.85030738 | 4207.22187 | 4182.11593 | 1532.04204 | 4544.36167 | Non-sig. |
| PF3D7_1134900 | N/A | 152.646986 | 0.603993446 | 0.51942118 | 1.16282021 | 0.24490246 | 0.84331907 | 151.422781 | 188.899219 | 91.147405 | 179.11854 | Non-sig. |
| PF3D7_1205800 | HMGB3 | 14095.6928 | 0.601748699 | 0.54864608 | 1.09678848 | 0.27273387 | 0.85618956 | 16709.0927 | 15862.9985 | 5687.24675 | 18123.4332 | Non-sig. |
| PF3D7_1001100 | ACBP1 | 340.113526 | 0.601634887 | 0.47621808 | 1.26336004 | 0.20645982 | 0.82391338 | 332.807775 | 432.768814 | 207.850455 | 387.027063 | Non-sig. |
| PF3D7_1237500 | N/A | 3022.41925 | 0.601385748 | 0.41829101 | 1.43772095 | 0.15051325 | 0.75169515 | 2643.3778 | 3317.53258 | 2159.66855 | 3969.09807 | Non-sig. |
| PF3D7_0620200 | N/A | 79.1923673 | 0.597588685 | 0.5559471 | 1.07490206 | 0.28241858 | 0.86390383 | 66.1573917 | 86.9698069 | 59.9294644 | 103.712806 | Non-sig. |
| PF3D7_0804900 | N/A | 2749.34058 | 0.594967143 | 0.50111957 | 1.18727582 | 0.23511884 | 0.83828834 | 1358.74787 | 3064.55727 | 3021.47024 | 3552.58692 | Non-sig. |
| PF3D7_1435300 | N/A | 9554.58938 | 0.594853681 | 0.52987041 | 1.12263992 | 0.26159045 | 0.85030738 | 4250.96124 | 10047.1793 | 10973.0893 | 12947.1277 | Non-sig. |
| PF3D7_1318200 | G3PAT | 47.7869477 | 0.594724157 | 0.75450877 | 0.78822696 | 0.43056396 | 0.91355742 | 16.7802933 | 65.5194074 | 59.0951139 | 49.7529761 | Non-sig. |
| PF3D7_1247300 | N/A | 678.71176 | 0.59457856 | 0.4570301 | 1.30096149 | 0.19327164 | 0.81266121 | 416.851847 | 837.99182 | 664.431129 | 795.572243 | Non-sig. |
| PF3D7_0718900 | N/A | 1.81065148 | 0.594093709 | 2.44456333 | 0.24302652 | 0.80798486 | 0.99028237 | 2.08522085 | 1.07585439 | 0.82002618 | 3.26150449 | Non-sig. |
| PF3D7_0223500 | VAR | 435.392686 | 0.593751128 | 0.44869118 | 1.32329574 | 0.18573705 | 0.80052251 | 330.929704 | 602.807718 | 363.090035 | 444.743286 | Non-sig. |
| PF3D7_0405200 | CERLI2 | 4805.65825 | 0.593066457 | 0.50977913 | 1.16337924 | 0.24467567 | 0.84331907 | 5363.1546 | 6334.87464 | 2300.44968 | 5224.15406 | Non-sig. |
| PF3D7_1231000 | N/A | 1295.5706 | 0.592468664 | 0.67754351 | 0.87443633 | 0.38188068 | 0.90292925 | 1764.64357 | 1533.18245 | 302.249863 | 1582.20651 | Non-sig. |
| PF3D7_0505200 | ALP2b | 1430.55503 | 0.592433901 | 0.53239529 | 1.11277074 | 0.26580691 | 0.85416239 | 629.064566 | 1694.54516 | 1652.25953 | 1746.35086 | Non-sig. |
| PF3D7_0927300 | FH | 1975.29815 | 0.59034309 | 0.42787641 | 1.3797047 | 0.16767759 | 0.77517962 | 1790.30301 | 2607.43274 | 1363.32154 | 2140.13532 | Non-sig. |
| PF3D7_1228200 | N/A | 187.9537 | 0.589760399 | 0.4704157 | 1.25370049 | 0.20995089 | 0.82566351 | 147.346392 | 233.919464 | 152.757054 | 217.79189 | Non-sig. |
| PF3D7_0704400 | N/A | 3935.38909 | 0.589161537 | 0.46555454 | 1.26550486 | 0.20569041 | 0.82391338 | 2204.5769 | 4777.11395 | 4080.65685 | 4679.20864 | Non-sig. |
| PF3D7_0219600 | RFC1 | 2514.28927 | 0.588636634 | 0.41578796 | 1.41571353 | 0.1568594 | 0.75894163 | 1773.39507 | 3021.14539 | 2243.12788 | 3019.48873 | Non-sig. |
| PF3D7_1331100 | POLQ | 3710.29991 | 0.587540045 | 0.49084529 | 1.19699639 | 0.23130796 | 0.83828834 | 1890.8564 | 4335.01933 | 4038.82778 | 4576.49611 | Non-sig. |
| PF3D7_0319800 | N/A | 60.1529965 | 0.587441905 | 0.59989032 | 0.97924884 | 0.32745704 | 0.88852997 | 53.5448783 | 65.4995936 | 42.6879227 | 78.8795914 | Non-sig. |
| PF3D7_0216500 | N/A | 1028.75532 | 0.586675712 | 0.497915 | 1.17826479 | 0.23869106 | 0.84066259 | 1120.33827 | 1219.55163 | 524.921819 | 1250.20957 | Non-sig. |
| PF3D7_0104600 | N/A | 130.213531 | 0.586602431 | 0.50806963 | 1.15457095 | 0.24826617 | 0.84804265 | 90.5957292 | 164.13052 | 117.461031 | 148.666845 | Non-sig. |
| PF3D7_1252900 | N/A | 124.185809 | 0.585950205 | 0.54944492 | 1.06644031 | 0.28622465 | 0.86595658 | 116.675297 | 185.700108 | 82.1090492 | 112.258782 | Non-sig. |
| PF3D7_0409800 | N/A | 6068.83392 | 0.584313086 | 0.43849362 | 1.33254639 | 0.18268071 | 0.80052251 | 5873.84248 | 7816.09492 | 3839.23576 | 6746.16254 | Non-sig. |
| PF3D7_0406000 | N/A | 2678.97489 | 0.584076941 | 0.42163558 | 1.38526483 | 0.1659715 | 0.77284241 | 2423.13456 | 3439.33016 | 1865.01831 | 2988.41655 | Non-sig. |
| PF3D7_1432300 | N/A | 574.505157 | 0.582794054 | 0.54333856 | 1.07261678 | 0.28344309 | 0.86551387 | 259.18251 | 603.933398 | 660.426524 | 774.478196 | Non-sig. |
| PF3D7_1423100 | N/A | 2209.90484 | 0.581887318 | 0.58370246 | 0.99689029 | 0.31881776 | 0.88651753 | 2755.59213 | 2700.16411 | 785.253847 | 2598.60926 | Non-sig. |
| PF3D7_0820600 | N/A | 321.788233 | 0.581683671 | 0.47423907 | 1.22656209 | 0.21998723 | 0.83527883 | 208.207389 | 341.306052 | 307.10955 | 430.529942 | Non-sig. |
| PF3D7_1432200 | N/A | 2246.70265 | 0.581057954 | 0.51573225 | 1.12666592 | 0.25988376 | 0.85030738 | 2544.64372 | 2524.01263 | 1056.37051 | 2861.78373 | Non-sig. |
| PF3D7_1319700 | N/A | 346.606337 | 0.581015501 | 0.50535533 | 1.14971678 | 0.25026054 | 0.8494288 | 186.340435 | 412.008962 | 368.770399 | 419.305552 | Non-sig. |
| PF3D7_1107200 | N/A | 1354.14992 | 0.580788049 | 0.4659425 | 1.2464801 | 0.21258819 | 0.83121199 | 1367.50915 | 1815.42948 | 803.241872 | 1430.41917 | Non-sig. |
| PF3D7_1251400 | N/A | 70.4987336 | 0.576724314 | 0.63978628 | 0.90143277 | 0.36735827 | 0.89607003 | 35.7440651 | 68.6952349 | 77.1846607 | 100.370974 | Non-sig. |
| PF3D7_0113900 | CBP1 | 40.0471981 | 0.576448895 | 0.66907563 | 0.86156015 | 0.38892961 | 0.90444842 | 31.460493 | 52.631543 | 32.8301473 | 43.2666091 | Non-sig. |
| PF3D7_0800700 | SURF8.3 | 449.538495 | 0.57505543 | 0.44793511 | 1.2837918 | 0.19921481 | 0.81595849 | 418.499319 | 528.763857 | 303.9561 | 546.934704 | Non-sig. |
| PF3D7_1025100 | GFPT | 3059.29981 | 0.574597355 | 0.41363236 | 1.38914992 | 0.16478717 | 0.77284241 | 2237.28784 | 3895.25565 | 2678.55928 | 3426.09649 | Non-sig. |
| PF3D7_1130800 | N/A | 36.4232986 | 0.574151016 | 0.69675939 | 0.82403054 | 0.40992221 | 0.90748263 | 27.3061086 | 50.4623753 | 31.1967744 | 36.7279361 | Non-sig. |
| PF3D7_1420700 | P113 | 11623.4741 | 0.572644735 | 0.4780269 | 1.19793412 | 0.23094266 | 0.83828834 | 12195.5996 | 11825.9179 | 6497.7056 | 15974.6733 | Non-sig. |
| PF3D7_1029900 | N/A | 14323.3684 | 0.572637782 | 0.43054197 | 1.33003939 | 0.18350529 | 0.80052251 | 13550.733 | 15552.305 | 9484.50826 | 18705.9273 | Non-sig. |
| PF3D7_1368700 | N/A | 1917.85017 | 0.572480772 | 0.50361448 | 1.13674406 | 0.25564529 | 0.85030738 | 2124.87023 | 2509.83062 | 960.063928 | 2076.63589 | Non-sig. |
| PF3D7_1025900 | N/A | 13382.0042 | 0.570724512 | 0.54992675 | 1.03781915 | 0.29935426 | 0.87358937 | 16122.0585 | 16576.2002 | 5416.5486 | 15413.2093 | Non-sig. |
| PF3D7_1002600 | N/A | 379.766641 | 0.570482122 | 0.47760912 | 1.194454 | 0.23230042 | 0.83828834 | 228.287654 | 442.140377 | 382.684809 | 465.953725 | Non-sig. |
| PF3D7_0924400 | N/A | 14189.5593 | 0.570428465 | 0.48758594 | 1.16990343 | 0.24203984 | 0.8422984 | 15343.1372 | 14985.7714 | 7497.9549 | 18931.3735 | Non-sig. |
| PF3D7_0914200 | N/A | 606.343838 | 0.569019085 | 0.4538287 | 1.25381909 | 0.20990777 | 0.82566351 | 398.95943 | 645.928427 | 577.374097 | 803.113396 | Non-sig. |
| PF3D7_0706500 | N/A | 6365.85485 | 0.568941792 | 0.4796722 | 1.18610541 | 0.23558067 | 0.83828834 | 6639.15723 | 6283.21513 | 3614.52199 | 8926.52506 | Non-sig. |
| PF3D7_0702400 | SEMP1 | 539.352532 | 0.568150667 | 0.44216854 | 1.28491878 | 0.19882067 | 0.81528784 | 369.444085 | 620.196382 | 499.340941 | 668.42872 | Non-sig. |
| PF3D7_1021900 | N/A | 15798.5211 | 0.56762225 | 0.71555911 | 0.79325697 | 0.42762812 | 0.91236574 | 22458.8105 | 17068.2181 | 3001.79753 | 20665.2583 | Non-sig. |
| PF3D7_0730400 | IMP2 | 2384.93273 | 0.566910185 | 0.44883027 | 1.26308366 | 0.20655912 | 0.82391338 | 1515.98937 | 3205.58763 | 2328.31821 | 2489.83571 | Non-sig. |
| PF3D7_1136100 | N/A | 40.5908621 | 0.566404623 | 0.67697826 | 0.8366659 | 0.40278039 | 0.90716539 | 36.7868298 | 55.8185154 | 28.7378 | 41.0203032 | Non-sig. |

| | | | | | | | | | | | | |
|---|---|---|---|---|---|---|---|---|---|---|---|---|
| PF3D7_1232700 | ND9 | 1657.70451 | 0.566326444 | 0.44802925 | 1.26403901 | 0.20621603 | 0.82391338 | 1027.4915 | 2066.48971 | 1645.0902 | 1891.74665 | Non-sig. |
| PF3D7_0618000 | N/A | 14059.8921 | 0.566038169 | 0.40389745 | 1.40144031 | 0.16108244 | 0.7667698 | 11971.9939 | 16152.5022 | 10701.2224 | 17413.8499 | Non-sig. |
| PF3D7_1231200 | N/A | 4734.56567 | 0.565248283 | 0.62505323 | 0.90432023 | 0.36582562 | 0.89567376 | 6234.66675 | 5916.24875 | 1403.30182 | 5384.04535 | Non-sig. |
| PF3D7_1316200 | N/A | 1109.10468 | 0.565090579 | 0.43062478 | 1.31225746 | 0.18943328 | 0.80547152 | 754.491815 | 1309.66808 | 1034.52138 | 1337.73743 | Non-sig. |
| PF3D7_1146500 | LRR10 | 16.3056711 | 0.565074943 | 0.92898102 | 0.60827394 | 0.5430058 | 0.93953127 | 15.6989033 | 18.2701063 | 10.6669105 | 20.5867643 | Non-sig. |
| PF3D7_1227500 | SOC2 | 6260.37147 | 0.564389818 | 0.46489615 | 1.21401268 | 0.22474288 | 0.83828834 | 6553.90448 | 7178.46265 | 3548.91415 | 7760.2046 | Non-sig. |
| PF3D7_0515400 | N/A | 4979.90498 | 0.563754439 | 0.5487875 | 1.02727275 | 0.30429205 | 0.87568674 | 5982.10979 | 5467.63385 | 2056.58853 | 6413.28775 | Non-sig. |
| PF3D7_1410900 | N/A | 3013.5196 | 0.562860892 | 0.54774577 | 1.02759513 | 0.30414031 | 0.87568674 | 3618.04575 | 3756.15106 | 1248.44238 | 3431.43919 | Non-sig. |
| PF3D7_1127600 | N/A | 3276.83714 | 0.562804827 | 0.40787714 | 1.37983911 | 0.16763619 | 0.77517962 | 2680.21339 | 4190.36603 | 2611.14039 | 3625.62873 | Non-sig. |
| PF3D7_1325900 | N/A | 4152.91945 | 0.561440571 | 0.42457944 | 1.32234518 | 0.18605324 | 0.80073927 | 3662.14586 | 4253.70247 | 3047.77795 | 5648.05152 | Non-sig. |
| PF3D7_0703800 | N/A | 129.169665 | 0.561382026 | 0.63168544 | 0.88870503 | 0.37416163 | 0.89654416 | 158.974412 | 144.845358 | 50.1005413 | 162.758348 | Non-sig. |
| PF3D7_0514000 | TTL | 260.460997 | 0.560935356 | 0.47722981 | 1.17539882 | 0.23983519 | 0.84181962 | 252.884973 | 302.848709 | 168.272411 | 317.837894 | Non-sig. |
| PF3D7_1474600 | VMP1 | 1488.16491 | 0.560220814 | 0.49095728 | 1.14107853 | 0.25383725 | 0.85030738 | 784.905659 | 1654.747 | 1620.28787 | 1892.71909 | Non-sig. |
| PF3D7_1241400 | N/A | 392.307852 | 0.559584036 | 0.50922407 | 1.09889549 | 0.27181365 | 0.8556331 | 222.423964 | 379.681599 | 411.572144 | 555.553702 | Non-sig. |
| PF3D7_0811700 | N/A | 550.659441 | 0.55861823 | 0.49914301 | 1.11915466 | 0.26307416 | 0.85095134 | 293.972361 | 687.590317 | 596.362881 | 624.712204 | Non-sig. |
| PF3D7_1145400 | DYN1 | 7753.16042 | 0.558583266 | 0.43622373 | 1.28049721 | 0.20037033 | 0.81595849 | 4941.70897 | 8871.07548 | 7599.37163 | 9600.48561 | Non-sig. |
| PF3D7_1325200 | N/A | 272.973938 | 0.557964147 | 0.49736703 | 1.12183581 | 0.26193226 | 0.85030738 | 278.573802 | 359.583747 | 163.399547 | 290.338657 | Non-sig. |
| PF3D7_1425600 | N/A | 190.585776 | 0.5566456 | 0.48760888 | 1.14158217 | 0.25362774 | 0.85030738 | 178.106371 | 207.015831 | 130.613137 | 246.607765 | Non-sig. |
| PF3D7_1213900 | N/A | 1053.71614 | 0.556641795 | 0.52379516 | 1.06270892 | 0.28791397 | 0.86724628 | 553.043287 | 937.587932 | 1152.42799 | 1571.80537 | Non-sig. |
| PF3D7_0631400 | MC-2TM | 97.4925084 | 0.556576381 | 0.56599777 | 0.98335438 | 0.32543305 | 0.88694295 | 62.5439508 | 138.792839 | 95.1191546 | 93.5140894 | Non-sig. |
| PF3D7_1312100 | N/A | 2811.13698 | 0.554284885 | 0.56606879 | 0.97918291 | 0.32748961 | 0.88852997 | 3473.18518 | 3346.97592 | 1082.61274 | 3341.77406 | Non-sig. |
| PF3D7_0522100 | N/A | 2104.96406 | 0.553948028 | 0.41871869 | 1.32295989 | 0.18584872 | 0.80052251 | 1895.4461 | 2349.5786 | 1516.2114 | 2658.62014 | Non-sig. |
| PF3D7_1351600 | GK | 158.818364 | 0.553362496 | 0.48197021 | 1.14812594 | 0.25091658 | 0.8494288 | 133.513539 | 191.059047 | 123.987953 | 186.712918 | Non-sig. |
| PF3D7_1031800 | N/A | 1306.20222 | 0.552884581 | 0.42185617 | 1.31059973 | 0.18999303 | 0.80577283 | 1149.93987 | 1690.62032 | 968.066424 | 1416.18224 | Non-sig. |
| PF3D7_0704100 | BTP2 | 12981.1343 | 0.552777513 | 0.61435193 | 0.89977339 | 0.36824086 | 0.89615989 | 16945.3867 | 13559.4852 | 4103.53438 | 17316.1309 | Non-sig. |
| PF3D7_0918500 | TR | 5778.05968 | 0.552136457 | 0.43050928 | 1.28251929 | 0.19966054 | 0.81595849 | 5343.76906 | 7794.96066 | 4027.8433 | 5945.6657 | Non-sig. |
| PF3D7_0718400 | N/A | 1505.78862 | 0.550461726 | 0.42399756 | 1.29826625 | 0.19419586 | 0.81267745 | 1134.34227 | 2006.64069 | 1309.46154 | 1572.70997 | Non-sig. |
| PF3D7_1436900 | N/A | 312.888888 | 0.550270313 | 0.47761716 | 1.15211588 | 0.24927346 | 0.84847675 | 196.427684 | 391.869751 | 311.156182 | 352.101933 | Non-sig. |
| PF3D7_1121100 | N/A | 5557.03606 | 0.550255736 | 0.59138519 | 0.93045235 | 0.35213693 | 0.8930363 | 7106.36109 | 6560.14355 | 1914.01745 | 6647.62216 | Non-sig. |
| PF3D7_1319400 | N/A | 8157.96603 | 0.550061527 | 0.55029558 | 0.99957468 | 0.31751638 | 0.88651753 | 9913.85863 | 9975.97905 | 3329.27797 | 9412.74846 | Non-sig. |
| PF3D7_1366100 | DIP13 | 1376.26317 | 0.549613307 | 0.51099775 | 1.07556894 | 0.28212008 | 0.86390383 | 1517.59906 | 1946.61434 | 717.263978 | 1323.57531 | Non-sig. |
| PF3D7_0805300 | N/A | 5964.1923 | 0.549574965 | 0.68413231 | 0.80331679 | 0.42179166 | 0.91135898 | 8327.81711 | 6284.30289 | 1356.122 | 7888.52719 | Non-sig. |
| PF3D7_0605600 | N/A | 471.904863 | 0.549527916 | 0.55702397 | 0.98654268 | 0.32386688 | 0.88694295 | 563.741037 | 581.385735 | 202.896728 | 539.595952 | Non-sig. |
| PF3D7_0525100 | ACS10 | 6489.47836 | 0.549305333 | 0.51540672 | 1.06577062 | 0.28652735 | 0.86595658 | 3051.71727 | 7325.14061 | 7485.35049 | 8095.70507 | Non-sig. |
| PF3D7_0405100 | SEC24B | 6258.41669 | 0.549093981 | 0.58260203 | 0.94248553 | 0.34594412 | 0.8930363 | 2241.44355 | 7163.11294 | 7921.29624 | 7707.81405 | Non-sig. |
| PF3D7_1203800 | N/A | 212.770684 | 0.548264687 | 0.46243254 | 1.1856101 | 0.23577631 | 0.83828834 | 167.377133 | 250.000664 | 178.22284 | 255.482101 | Non-sig. |
| PF3D7_0935390 | N/A | 657.128233 | 0.547658084 | 0.46198065 | 1.18545675 | 0.23583691 | 0.83828834 | 410.436988 | 715.721432 | 657.011761 | 845.342749 | Non-sig. |
| PF3D7_1366900 | N/A | 12163.1416 | 0.546894536 | 0.45892145 | 1.19169531 | 0.23338074 | 0.83828834 | 12694.6674 | 14013.0865 | 7075.6607 | 14869.1521 | Non-sig. |
| PF3D7_0617900 | H3.3 | 14376.4943 | 0.545666678 | 0.40048353 | 1.36251963 | 0.17303395 | 0.78315911 | 11931.8081 | 16988.5912 | 11447.553 | 17138.0249 | Non-sig. |
| PF3D7_1446300 | N/A | 1653.88384 | 0.544418894 | 0.44348052 | 1.22760499 | 0.21959529 | 0.83527883 | 1290.40714 | 1538.3399 | 1400.48645 | 2386.30186 | Non-sig. |
| PF3D7_0920700 | N/A | 6742.48511 | 0.544289691 | 0.42127615 | 1.29200215 | 0.1963564 | 0.81437577 | 4860.4676 | 7107.65976 | 6110.31327 | 8891.4998 | Non-sig. |
| PF3D7_1207000 | N/A | 11467.7613 | 0.543749784 | 0.52427174 | 1.03715258 | 0.29966475 | 0.87376107 | 13414.8053 | 12445.6124 | 5249.45095 | 14761.1765 | Non-sig. |
| PF3D7_1017900 | RPN5 | 2798.18916 | 0.54202753 | 0.41512096 | 1.30570985 | 0.19165125 | 0.80841426 | 2529.78775 | 3442.4023 | 2027.67621 | 3192.89036 | Non-sig. |
| PF3D7_0707700 | N/A | 4874.25222 | 0.541579007 | 0.42285639 | 1.28076343 | 0.20027677 | 0.81595849 | 4372.86923 | 5042.24518 | 3567.22203 | 6514.67243 | Non-sig. |
| PF3D7_0531400 | N/A | 331.719354 | 0.540690957 | 0.47271478 | 1.14379955 | 0.25270678 | 0.85030738 | 310.748844 | 328.265672 | 229.991618 | 457.87128 | Non-sig. |
| PF3D7_0409600 | RPA1 | 6759.73965 | 0.539578623 | 0.42758575 | 1.26191909 | 0.2069779 | 0.82391338 | 4497.25296 | 7935.05948 | 6522.8147 | 8083.83147 | Non-sig. |
| PF3D7_1354800 | MCA1 | 333.42944 | 0.539362382 | 0.47663103 | 1.13161408 | 0.25779672 | 0.85030738 | 207.416371 | 396.97675 | 335.923606 | 393.401033 | Non-sig. |
| PF3D7_0805500 | N/A | 937.511427 | 0.53859975 | 0.43942753 | 1.22568505 | 0.22031722 | 0.83527883 | 671.254109 | 955.733239 | 857.595236 | 1265.46312 | Non-sig. |
| PF3D7_1208400 | N/A | 1957.95837 | 0.538376872 | 0.44115492 | 1.22038053 | 0.22232065 | 0.83675579 | 1942.05267 | 2322.31018 | 1251.88594 | 2315.58469 | Non-sig. |
| PF3D7_1107300 | PAIP1 | 25202.9561 | 0.537962312 | 0.50188084 | 1.0718925 | 0.28376832 | 0.86551387 | 28447.6717 | 26812.3468 | 12668.249 | 32883.557 | Non-sig. |
| PF3D7_1146800 | N/A | 347.341387 | 0.537457586 | 0.65228333 | 0.82396339 | 0.40996036 | 0.90748263 | 462.852596 | 357.139796 | 104.333966 | 465.039189 | Non-sig. |
| PF3D7_1372300 | N/A | 229.701989 | 0.537112888 | 0.47207641 | 1.13776685 | 0.25521785 | 0.85030738 | 203.298948 | 305.739555 | 171.671648 | 238.097806 | Non-sig. |
| PF3D7_0315700 | N/A | 61.8457598 | 0.536206157 | 0.62799436 | 0.85383913 | 0.39319413 | 0.90601068 | 64.2036885 | 84.7679547 | 36.9576295 | 61.4537664 | Non-sig. |
| PF3D7_1134600 | N/A | 475.724631 | 0.534624674 | 0.68551468 | 0.779888 | 0.4354568 | 0.91522571 | 660.922374 | 511.610244 | 116.652425 | 613.713478 | Non-sig. |

| | | | | | | | | | | | | |
|---|---|---|---|---|---|---|---|---|---|---|---|---|
| PF3D7_1401700 | N/A | 122.268676 | 0.533949772 | 0.52171119 | 1.02345855 | 0.30609109 | 0.87685698 | 81.4880332 | 148.384729 | 118.122717 | 141.079224 | Non-sig. |
| PF3D7_0512400 | N/A | 12.3773123 | 0.532966346 | 1.07852005 | 0.49416452 | 0.62118999 | 0.9639908 | 14.596546 | 17.2136702 | 5.74018323 | 11.9588498 | Non-sig. |
| PF3D7_0723300 | N/A | 6435.67772 | 0.532792876 | 0.58542195 | 0.91010061 | 0.36276945 | 0.89567376 | 8232.75243 | 7467.64084 | 2289.00879 | 7753.30884 | Non-sig. |
| PF3D7_1143800 | CAP93 | 1537.11238 | 0.531891106 | 0.4574724 | 1.16267365 | 0.24496193 | 0.84331907 | 1191.33822 | 1349.54981 | 1322.44382 | 2285.11766 | Non-sig. |
| PF3D7_1228500 | N/A | 937.72517 | 0.531603617 | 0.44223727 | 1.20207783 | 0.22933338 | 0.83828834 | 828.541873 | 1306.22939 | 705.354428 | 910.774993 | Non-sig. |
| PF3D7_1017200 | N/A | 227.15611 | 0.531283147 | 0.48955656 | 1.08523344 | 0.27781828 | 0.85972841 | 143.913446 | 273.768528 | 227.416756 | 263.52571 | Non-sig. |
| PF3D7_0901400 | RIF | 2.27919871 | 0.530942146 | 2.20624617 | 0.24065408 | 0.80982323 | 0.99028237 | 2.09542535 | 4.29756751 | 1.64123839 | 1.08256359 | Non-sig. |
| PF3D7_1026600 | N/A | 75114.8454 | 0.53084818 | 0.81749105 | 0.64936268 | 0.51610398 | 0.93247207 | 115013.915 | 91914.3281 | 7885.12123 | 85646.0168 | Non-sig. |
| PF3D7_0823300 | GCN5 | 6425.66189 | 0.529970413 | 0.41068595 | 1.29045177 | 0.19689385 | 0.81437577 | 5191.63078 | 6787.85974 | 5325.41312 | 8397.74389 | Non-sig. |
| PF3D7_1210400 | N/A | 640.127576 | 0.529304699 | 0.43468617 | 1.2176709 | 0.22334906 | 0.83828834 | 458.080793 | 794.987247 | 589.73232 | 717.709945 | Non-sig. |
| PF3D7_0826900 | N/A | 1346.6863 | 0.52904254 | 0.51411753 | 1.02903034 | 0.30346542 | 0.87520946 | 1549.23652 | 1654.60631 | 656.197901 | 1526.70447 | Non-sig. |
| PF3D7_1120700 | N/A | 654.570961 | 0.528353952 | 0.57827479 | 0.91367281 | 0.36088879 | 0.89560738 | 818.690327 | 753.025815 | 253.823498 | 792.744204 | Non-sig. |
| PF3D7_0630100 | N/A | 160.022361 | 0.528191011 | 0.53856036 | 0.98074618 | 0.32671793 | 0.88852997 | 166.274815 | 153.446767 | 96.0874866 | 224.280374 | Non-sig. |
| PF3D7_0108300 | ARP | 21312.9795 | 0.527950091 | 0.40766314 | 1.29506457 | 0.19529795 | 0.81344649 | 18008.9421 | 22841.2849 | 16903.5514 | 27498.1397 | Non-sig. |
| PF3D7_1348000 | N/A | 546.844655 | 0.52773927 | 0.52581114 | 1.00366696 | 0.31553917 | 0.88486757 | 630.964318 | 656.5354 | 265.310745 | 634.568157 | Non-sig. |
| PF3D7_1205000 | N/A | 2166.40581 | 0.527698502 | 0.46413582 | 1.13694845 | 0.25555984 | 0.85030738 | 2279.88317 | 2441.38149 | 1269.60967 | 2674.74893 | Non-sig. |
| PF3D7_1456400 | N/A | 82.1400232 | 0.527052838 | 0.71889781 | 0.73314013 | 0.46347296 | 0.92509266 | 109.482519 | 89.0553128 | 25.4605331 | 104.561728 | Non-sig. |
| PF3D7_1437700 | N/A | 1155.13049 | 0.526913299 | 0.56132798 | 0.93869059 | 0.34788962 | 0.8930363 | 467.518696 | 1376.50873 | 1425.02846 | 1351.46608 | Non-sig. |
| PF3D7_0704300 | N/A | 11945.8416 | 0.526881297 | 0.64213641 | 0.82051304 | 0.4119237 | 0.90748263 | 16257.6793 | 13076.8105 | 3319.63671 | 15129.2401 | Non-sig. |
| PF3D7_0209600 | N/A | 4162.48498 | 0.526318315 | 0.46670881 | 1.12772311 | 0.25943687 | 0.85030738 | 2380.56276 | 4903.65442 | 4442.09474 | 4923.62802 | Non-sig. |
| PF3D7_0307600 | N/A | 1467.44504 | 0.526264502 | 0.41270538 | 1.27515785 | 0.20225338 | 0.81948076 | 1278.77098 | 1784.43487 | 1126.79087 | 1679.78345 | Non-sig. |
| PF3D7_0518500 | DDX23 | 2393.62848 | 0.525555603 | 0.41220848 | 1.27497522 | 0.20231801 | 0.81948076 | 2146.65639 | 2837.51343 | 1778.30724 | 2812.03685 | Non-sig. |
| PF3D7_0103600 | PSH1 | 805.578466 | 0.52510156 | 0.59255367 | 0.88616708 | 0.3755275 | 0.89654416 | 1028.3433 | 879.608697 | 293.251176 | 1021.11069 | Non-sig. |
| PF3D7_0915000 | NDH2 | 2657.08071 | 0.523695331 | 0.49958562 | 1.04825941 | 0.29451911 | 0.87170077 | 1353.42055 | 3284.21631 | 3006.253 | 2984.43299 | Non-sig. |
| PF3D7_1469200 | SHLP1 | 3043.4326 | 0.52134545 | 0.40975571 | 1.27233236 | 0.20325506 | 0.82139228 | 2344.16167 | 3797.66247 | 2654.61427 | 3377.29201 | Non-sig. |
| PF3D7_0423400 | AARP | 3201.49665 | 0.521319779 | 0.4900635 | 1.06378007 | 0.28742834 | 0.86724628 | 3558.10997 | 4055.94171 | 1700.86114 | 3491.07378 | Non-sig. |
| PF3D7_1340500 | N/A | 376.047257 | 0.521240838 | 0.53523322 | 0.97385742 | 0.33012734 | 0.88975609 | 434.92885 | 417.337335 | 183.163117 | 468.759728 | Non-sig. |
| PF3D7_0312400 | GSK3 | 5718.34861 | 0.521150521 | 0.4049673 | 1.28689532 | 0.19813078 | 0.81528784 | 4516.56873 | 7083.61283 | 4876.57781 | 6396.63507 | Non-sig. |
| PF3D7_0932400 | RF1 | 1011.84246 | 0.521136854 | 0.42583191 | 1.22380882 | 0.22102435 | 0.83527883 | 921.676834 | 1274.20225 | 740.600616 | 1110.89015 | Non-sig. |
| PF3D7_0919900 | RCC-PIP | 20456.6283 | 0.521043205 | 0.60604278 | 0.85974657 | 0.38992876 | 0.90444842 | 26644.0595 | 19751.8655 | 6960.84805 | 28469.7402 | Non-sig. |
| PF3D7_1455000 | PPM3 | 116.521171 | 0.520389709 | 0.59924185 | 0.8684135 | 0.38516801 | 0.90292925 | 136.776161 | 145.945058 | 55.0221902 | 128.341275 | Non-sig. |
| PF3D7_1020200 | N/A | 330.607522 | 0.520307615 | 0.71525633 | 0.72744217 | 0.46695513 | 0.92538923 | 471.395356 | 404.557746 | 72.2691226 | 374.207862 | Non-sig. |
| PF3D7_1107700 | PES | 823.944457 | 0.520003352 | 0.43941251 | 1.18340589 | 0.23664832 | 0.83828834 | 793.421688 | 1014.0604 | 560.912687 | 927.383048 | Non-sig. |
| PF3D7_1453900 | N/A | 1129.11251 | 0.519435848 | 0.43216513 | 1.20193837 | 0.22938741 | 0.83828834 | 776.439806 | 1303.3632 | 1079.34068 | 1357.30635 | Non-sig. |
| PF3D7_0311900 | N/A | 253.318705 | 0.518343136 | 0.55553366 | 0.93305441 | 0.35079188 | 0.8930363 | 118.906533 | 287.5864 | 297.291284 | 309.490604 | Non-sig. |
| PF3D7_1340400 | N/A | 10.0816747 | 0.516248947 | 1.12644046 | 0.45830114 | 0.64673611 | 0.97317009 | 8.38788545 | 9.66752841 | 8.20726361 | 14.0640214 | Non-sig. |
| PF3D7_1239900 | VPS16 | 888.318627 | 0.515848368 | 0.43625137 | 1.18245673 | 0.23702453 | 0.83828834 | 608.707518 | 1068.54646 | 853.396622 | 1022.62391 | Non-sig. |
| PF3D7_0801700 | SENP2 | 2099.28076 | 0.515815808 | 0.52915016 | 0.97480044 | 0.32965926 | 0.88972733 | 2496.8546 | 2371.51027 | 959.486233 | 2569.27195 | Non-sig. |
| PF3D7_1107100 | N/A | 3152.53901 | 0.515439357 | 0.48801677 | 1.0561919 | 0.29088055 | 0.86724628 | 3453.16486 | 4218.06258 | 1737.81653 | 3201.11207 | Non-sig. |
| PF3D7_1352200 | UTP15 | 1505.96944 | 0.515341097 | 0.44238305 | 1.16492053 | 0.24405116 | 0.84331907 | 990.383551 | 1903.5436 | 1488.97203 | 1640.97857 | Non-sig. |
| PF3D7_1458000 | FP1 | 1558.74553 | 0.515302083 | 0.42387367 | 1.21569733 | 0.22410024 | 0.83828834 | 1102.01233 | 1854.15332 | 1464.34908 | 1814.46737 | Non-sig. |
| PF3D7_1366800 | N/A | 925.181173 | 0.515029586 | 0.48199584 | 1.06853533 | 0.2852791 | 0.86551387 | 519.653028 | 1108.59638 | 1003.50647 | 1068.96881 | Non-sig. |
| PF3D7_1202700 | N/A | 491.658457 | 0.515003974 | 0.53012572 | 0.97147517 | 0.33131171 | 0.88994703 | 564.461111 | 657.65452 | 245.586527 | 498.931668 | Non-sig. |
| PF3D7_1238800 | ACS11 | 8014.10297 | 0.51489612 | 0.42066042 | 1.22401845 | 0.22094527 | 0.83527883 | 5652.65483 | 8805.31075 | 7545.07981 | 10053.3665 | Non-sig. |
| PF3D7_1037200 | N/A | 51.2517042 | 0.514852311 | 0.62773744 | 0.82017143 | 0.41211839 | 0.90748263 | 48.3773543 | 64.3951452 | 36.1313129 | 56.1030044 | Non-sig. |
| PF3D7_1100800 | MC-2TM | 97.9828698 | 0.514215781 | 0.57774886 | 0.8900334 | 0.37344795 | 0.89654416 | 54.4658833 | 120.341281 | 106.675225 | 110.44909 | Non-sig. |
| PF3D7_1228700 | N/A | 3405.22344 | 0.513836762 | 0.4201351 | 1.22302745 | 0.22131933 | 0.83527883 | 2396.68874 | 3997.00659 | 3213.36658 | 4013.83186 | Non-sig. |
| PF3D7_0937700 | RIF | 9.18165171 | 0.5135466 | 1.18373437 | 0.43383601 | 0.66440753 | 0.97883902 | 9.43299098 | 12.8913324 | 5.74472032 | 8.6575631 | Non-sig. |
| PF3D7_1011200 | BET1 | 186.748228 | 0.512127845 | 0.51927117 | 0.98624355 | 0.32401361 | 0.88694295 | 113.366381 | 248.686139 | 194.269799 | 190.670594 | Non-sig. |
| PF3D7_1442700 | N/A | 2473.11515 | 0.511741179 | 0.4950122 | 1.03379508 | 0.30123197 | 0.87381251 | 2757.51561 | 2559.12822 | 1320.96408 | 3254.85269 | Non-sig. |
| PF3D7_0604600 | N/A | 308.274701 | 0.511533061 | 0.45549157 | 1.12303519 | 0.26142255 | 0.85030738 | 264.4633 | 316.443898 | 243.965595 | 408.226011 | Non-sig. |
| PF3D7_1005700 | N/A | 337.654022 | 0.511197222 | 0.4662177 | 1.0964775 | 0.27286986 | 0.85618956 | 242.161095 | 336.893842 | 314.569015 | 456.992139 | Non-sig. |
| PF3D7_0510700 | N/A | 12.4385591 | 0.511032791 | 1.09076422 | 0.46850894 | 0.63942068 | 0.96967357 | 15.7145649 | 15.041788 | 4.92365266 | 14.0742307 | Non-sig. |
| PF3D7_0113200 | N/A | 922.585282 | 0.510946999 | 0.46994184 | 1.08725581 | 0.27692377 | 0.85948573 | 928.294649 | 884.169335 | 593.782835 | 1284.09431 | Non-sig. |

| | | | | | | | | | | | |
|---|---|---|---|---|---|---|---|---|---|---|---|
| PF3D7_0208300 | N/A | 349.350161 | 0.510615944 | 0.45935503 | 1.11159324 | 0.26631308 | 0.85416239 | 243.430007 | 376.497407 | 332.689755 | 444.783475 | Non-sig. |
| PF3D7_1019000 | N/A | 9502.47296 | 0.510455198 | 0.5754172 | 0.88710452 | 0.37502263 | 0.89654416 | 11857.0614 | 8676.88946 | 3820.80652 | 13655.1344 | Non-sig. |
| PF3D7_1036500 | N/A | 10013.4714 | 0.509888082 | 0.51831773 | 0.98373653 | 0.32524507 | 0.88694295 | 11787.3695 | 12545.7235 | 4737.38288 | 10983.4096 | Non-sig. |
| PF3D7_1339000 | N/A | 12.8910591 | 0.508837944 | 1.03417382 | 0.49202362 | 0.62270265 | 0.96429926 | 7.27189992 | 13.9997938 | 13.9321884 | 16.3603542 | Non-sig. |
| PF3D7_0808300 | N/A | 4391.04256 | 0.507485218 | 0.4138081 | 1.22637817 | 0.2200564 | 0.83527883 | 4025.19156 | 5389.32477 | 3228.2091 | 4921.44481 | Non-sig. |
| PF3D7_1407000 | CCp3 | 243.14559 | 0.505221167 | 0.52577406 | 0.96090927 | 0.33659779 | 0.8920875 | 130.594309 | 281.079547 | 271.052305 | 289.856201 | Non-sig. |
| PF3D7_0214600 | STK2 | 19612.0941 | 0.504645877 | 0.70970231 | 0.71106698 | 0.47704273 | 0.92815732 | 28531.5871 | 22701.832 | 3901.98486 | 23312.9722 | Non-sig. |
| PF3D7_0214800 | N/A | 1462.31092 | 0.504365919 | 0.48934995 | 1.03068554 | 0.30268831 | 0.87475317 | 1420.66086 | 1190.60318 | 998.084628 | 2239.89501 | Non-sig. |
| PF3D7_0206000 | RAD2 | 2822.71573 | 0.503824563 | 0.47694873 | 1.05634953 | 0.29080855 | 0.86724628 | 3082.11725 | 3093.57798 | 1587.85007 | 3527.31762 | Non-sig. |
| PF3D7_0530100 | SYN6 | 2345.29627 | 0.503727647 | 0.45287198 | 1.11229589 | 0.26601095 | 0.85416239 | 2417.78162 | 2956.82225 | 1462.49336 | 2544.08785 | Non-sig. |
| PF3D7_0926500 | N/A | 5029.34948 | 0.502799919 | 0.43884773 | 1.14572751 | 0.25190792 | 0.85030738 | 3597.92175 | 4856.44095 | 4725.30625 | 6937.72895 | Non-sig. |
| PF3D7_0104800 | NPT1 | 696.093204 | 0.502342317 | 0.57992042 | 0.86622629 | 0.38636608 | 0.90388766 | 278.824826 | 691.084491 | 872.970513 | 941.492985 | Non-sig. |
| PF3D7_1031500 | DDX3X | 1825.63781 | 0.501190036 | 0.40766393 | 1.22941963 | 0.21891452 | 0.83527883 | 1465.99166 | 2139.22275 | 1557.31225 | 2140.02457 | Non-sig. |
| PF3D7_1329600 | N/A | 1889.27194 | 0.500933944 | 0.48224591 | 1.03875208 | 0.29892005 | 0.87351265 | 2070.20052 | 2417.1484 | 1059.25413 | 2010.48471 | Non-sig. |
| PF3D7_0826200 | N/A | 373.80395 | 0.500160535 | 0.47752522 | 1.04740129 | 0.29491454 | 0.87170077 | 387.198249 | 452.853502 | 232.408895 | 422.755155 | Non-sig. |
| PF3D7_0409200 | SOF1 | 1532.77176 | 0.500033621 | 0.42477266 | 1.17717938 | 0.23912392 | 0.84066259 | 1115.52587 | 1939.51778 | 1423.83322 | 1652.21016 | Non-sig. |
| PF3D7_0629300 | PL | 3218.08289 | 0.499458082 | 0.40688647 | 1.22751215 | 0.21963016 | 0.83527883 | 2498.54139 | 3730.47952 | 2834.42044 | 3808.89021 | Non-sig. |
| PF3D7_1315400 | N/A | 5011.82688 | 0.496254379 | 0.40221231 | 1.23381201 | 0.21727296 | 0.83527883 | 4105.46926 | 5798.95617 | 4211.00981 | 5931.87229 | Non-sig. |
| PF3D7_0724000 | N/A | 618.599512 | 0.496182769 | 0.42776533 | 1.15994154 | 0.24607261 | 0.8454557 | 490.033899 | 668.251297 | 536.413264 | 779.699587 | Non-sig. |
| PF3D7_1333100 | N/A | 73.6294997 | 0.495803509 | 0.79211371 | 0.62592467 | 0.53136436 | 0.93351492 | 105.27766 | 101.929143 | 17.2476299 | 70.0635658 | Non-sig. |
| PF3D7_0410600 | ATRP | 1284.90092 | 0.495686224 | 0.45521739 | 1.08890002 | 0.27619798 | 0.85889199 | 811.236325 | 1622.05612 | 1321.06739 | 1385.24385 | Non-sig. |
| PF3D7_1215400 | N/A | 2837.88937 | 0.495539344 | 0.40516908 | 1.22304334 | 0.22131333 | 0.83527883 | 2437.29886 | 3338.27014 | 2273.25003 | 3302.73844 | Non-sig. |
| PF3D7_1228600 | MSP9 | 86935.2927 | 0.49480582 | 0.58186263 | 0.85038254 | 0.39511244 | 0.90645858 | 112625.953 | 104249.664 | 31717.6464 | 99147.9077 | Non-sig. |
| PF3D7_0522900 | N/A | 372.377246 | 0.493830829 | 0.45785614 | 1.07857204 | 0.28077855 | 0.86335738 | 350.906576 | 384.009602 | 267.793121 | 486.799683 | Non-sig. |
| PF3D7_0303200 | N/A | 12806.8889 | 0.493063866 | 0.63704404 | 0.77398709 | 0.43893842 | 0.9169936 | 17567.3873 | 13554.7394 | 3712.01943 | 16393.4093 | Non-sig. |
| PF3D7_0920000 | ELO3 | 314.585756 | 0.49215597 | 0.47504627 | 1.03601691 | 0.30019425 | 0.87381251 | 203.130414 | 360.534979 | 319.476568 | 375.201063 | Non-sig. |
| PF3D7_1103400 | SufD | 2260.6275 | 0.491226645 | 0.4152709 | 1.18290649 | 0.23684621 | 0.83828834 | 2076.75304 | 2548.70343 | 1682.29916 | 2734.75439 | Non-sig. |
| PF3D7_0728700 | N/A | 169.89485 | 0.490844046 | 0.602623 | 0.81451264 | 0.41535134 | 0.90793155 | 210.637585 | 215.636901 | 72.2807105 | 181.024204 | Non-sig. |
| PF3D7_1227400 | N/A | 2212.87351 | 0.490674834 | 0.41817959 | 1.17335911 | 0.24065183 | 0.84181962 | 1612.65718 | 2524.24661 | 2067.44324 | 2647.147 | Non-sig. |
| PF3D7_1018400 | N/A | 75.2876688 | 0.489760306 | 0.5688345 | 0.8609891 | 0.38924405 | 0.90444842 | 71.2190908 | 94.5559187 | 54.1574456 | 81.2182202 | Non-sig. |
| PF3D7_1466200 | HDP1 | 1950.94094 | 0.487425503 | 0.53372949 | 0.91324447 | 0.36111398 | 0.89560738 | 945.770672 | 1823.2182 | 2302.75503 | 2732.01984 | Non-sig. |
| PF3D7_0933100 | N/A | 402.815182 | 0.487206903 | 0.51838097 | 0.93986264 | 0.34728802 | 0.8930363 | 461.017245 | 477.486382 | 210.248941 | 462.508162 | Non-sig. |
| PF3D7_0725400 | N/A | 9494.09364 | 0.487012057 | 0.46279305 | 1.05233226 | 0.29264713 | 0.87005865 | 9804.11883 | 13054.1271 | 6009.5487 | 9108.57991 | Non-sig. |
| PF3D7_0714200 | N/A | 4449.31371 | 0.486624162 | 0.42947478 | 1.13306807 | 0.25718567 | 0.85030738 | 4178.21563 | 4473.27752 | 3233.91216 | 5911.84952 | Non-sig. |
| PF3D7_1254100 | N/A | 5.0907493 | 0.485781767 | 1.54691574 | 0.31403247 | 0.75349638 | 0.99028237 | 5.22905991 | 8.59954122 | 3.28171868 | 3.25267737 | Non-sig. |
| PF3D7_1402900 | N/A | 314.022707 | 0.484881033 | 0.45795932 | 1.05878627 | 0.28969713 | 0.86724628 | 251.582507 | 313.308894 | 271.849629 | 419.349799 | Non-sig. |
| PF3D7_0528300 | N/A | 1657.83221 | 0.484481506 | 0.46827654 | 1.03460554 | 0.30085317 | 0.87381251 | 978.156733 | 2017.05024 | 1785.58092 | 1850.54094 | Non-sig. |
| PF3D7_1231500 | N/A | 472.053624 | 0.483637559 | 0.44292192 | 1.09192508 | 0.27486603 | 0.85802262 | 449.189243 | 541.951354 | 338.336465 | 558.737432 | Non-sig. |
| PF3D7_1326700 | N/A | 834.411714 | 0.483630582 | 0.47210698 | 1.02440888 | 0.30564219 | 0.87685698 | 882.953907 | 871.608172 | 509.076684 | 1074.00809 | Non-sig. |
| PF3D7_1320600 | RAB11a | 2926.88482 | 0.482872562 | 0.44140942 | 1.09393352 | 0.27398414 | 0.85618956 | 2934.77886 | 3748.65982 | 1948.6957 | 3075.40492 | Non-sig. |
| PF3D7_1301700 | CBP2 | 350.284578 | 0.482591852 | 0.46567375 | 1.03633038 | 0.30004804 | 0.87381251 | 351.499437 | 410.977464 | 233.238826 | 405.422583 | Non-sig. |
| PF3D7_0215400 | N/A | 3620.25797 | 0.482327709 | 0.43631586 | 1.10545537 | 0.26896232 | 0.8556331 | 3615.94314 | 3913.22151 | 2425.67468 | 4526.19253 | Non-sig. |
| PF3D7_1009700 | N/A | 7163.13986 | 0.482104379 | 0.40490911 | 1.19064839 | 0.23379165 | 0.83828834 | 6346.12596 | 8112.04814 | 5608.59143 | 8585.79389 | Non-sig. |
| PF3D7_1355300 | SET6 | 254.181956 | 0.482016973 | 0.57585131 | 0.83705111 | 0.40256384 | 0.90716539 | 112.579395 | 268.279953 | 311.24594 | 324.622534 | Non-sig. |
| PF3D7_0936800 | N/A | 545.076697 | 0.481954986 | 0.42521472 | 1.13343908 | 0.25702991 | 0.85030738 | 467.138253 | 636.365271 | 442.640382 | 634.162883 | Non-sig. |
| PF3D7_1023000 | N/A | 1279.46408 | 0.481930471 | 0.48871498 | 0.98611765 | 0.32407538 | 0.88694295 | 1435.4151 | 1482.19941 | 700.443628 | 1499.79819 | Non-sig. |
| PF3D7_1462700 | CYT1 | 1689.68989 | 0.481771466 | 0.51152043 | 0.94184208 | 0.3462735 | 0.8930363 | 845.149004 | 2086.68229 | 1974.73963 | 1852.18865 | Non-sig. |
| PF3D7_0400100 | VAR | 2517.46423 | 0.481629135 | 0.44980551 | 1.07074975 | 0.28428197 | 0.86551387 | 2575.15443 | 3246.64009 | 1627.38146 | 2620.68092 | Non-sig. |
| PF3D7_1243400 | N/A | 2649.04409 | 0.480525488 | 0.59079978 | 0.81334744 | 0.41601888 | 0.90793155 | 3464.28708 | 3372.93131 | 959.996659 | 2798.9613 | Non-sig. |
| PF3D7_0310900 | N/A | 1339.62175 | 0.47940792 | 0.52506137 | 0.91305121 | 0.36121561 | 0.89560738 | 690.453969 | 1225.37921 | 1547.29322 | 1895.36062 | Non-sig. |
| PF3D7_0823000 | VPS15 | 482.289322 | 0.479275463 | 0.43578537 | 1.09979704 | 0.27142056 | 0.8556331 | 361.290952 | 554.627622 | 444.372284 | 568.866431 | Non-sig. |
| PF3D7_1426300 | N/A | 930.476263 | 0.479090519 | 0.42103678 | 1.13788282 | 0.25516941 | 0.85030738 | 797.622489 | 1180.39124 | 757.189681 | 986.701641 | Non-sig. |
| PF3D7_1440000 | N/A | 2241.60673 | 0.477569139 | 0.43877147 | 1.0884234 | 0.27640824 | 0.85889199 | 2057.41627 | 2120.50327 | 1690.6112 | 3097.89617 | Non-sig. |
| PF3D7_1415700 | SPT | 171.811551 | 0.477521639 | 0.49746435 | 0.95991128 | 0.33709987 | 0.8920875 | 117.882182 | 185.631091 | 169.183815 | 214.549114 | Non-sig. |

| | | | | | | | | | | | | |
|---|---|---|---|---|---|---|---|---|---|---|---|---|
| PF3D7_0713500 | N/A | 522.832835 | 0.476957143 | 0.44638726 | 1.06848287 | 0.28530275 | 0.86551387 | 366.532625 | 576.094279 | 507.613328 | 641.09111 | Non-sig. |
| PF3D7_1315000 | N/A | 584.350343 | 0.476896195 | 0.42625936 | 1.11879349 | 0.26322825 | 0.85095134 | 460.669826 | 657.890372 | 516.520079 | 702.321095 | Non-sig. |
| PF3D7_0104100 | N/A | 1203.54061 | 0.475771811 | 0.46575627 | 1.02150383 | 0.3070158 | 0.87685698 | 727.889157 | 1334.51822 | 1285.48871 | 1466.26635 | Non-sig. |
| PF3D7_0516400 | PhLP1 | 1012.95223 | 0.4753706 | 0.4499262 | 1.05655239 | 0.29071592 | 0.86724628 | 1032.85034 | 1248.51598 | 662.573929 | 1107.86867 | Non-sig. |
| PF3D7_1308700 | N/A | 1493.52263 | 0.475156251 | 0.63022045 | 0.75395244 | 0.45087778 | 0.92081798 | 2042.51934 | 1832.54519 | 457.478041 | 1641.54795 | Non-sig. |
| PF3D7_1024200 | N/A | 138.124015 | 0.475132476 | 0.49497024 | 0.9599213 | 0.33709483 | 0.8920875 | 123.727463 | 157.900763 | 107.516099 | 163.351736 | Non-sig. |
| PF3D7_1007300 | N/A | 3454.85389 | 0.474966083 | 0.40783988 | 1.16458961 | 0.24418515 | 0.84331907 | 3108.30709 | 4027.62437 | 2674.28065 | 4009.20346 | Non-sig. |
| PF3D7_0815300 | N/A | 622.256723 | 0.474925517 | 0.47721771 | 0.99519676 | 0.31964058 | 0.88662607 | 375.004337 | 776.681483 | 666.15397 | 671.187104 | Non-sig. |
| PF3D7_0904600 | N/A | 5661.45632 | 0.474375611 | 0.50597877 | 0.93754054 | 0.34848058 | 0.8930363 | 6604.04526 | 6020.516 | 2874.34147 | 7146.92255 | Non-sig. |
| PF3D7_1412000 | N/A | 829.888125 | 0.473727865 | 0.60008958 | 0.78942859 | 0.42986155 | 0.91352046 | 299.629909 | 868.34212 | 1089.62422 | 1061.95625 | Non-sig. |
| PF3D7_1228100 | LRR13 | 127.02104 | 0.473626313 | 0.67044742 | 0.7064332 | 0.47991878 | 0.92880667 | 167.921965 | 176.115503 | 45.1477518 | 118.89894 | Non-sig. |
| PF3D7_1246300 | KIC4 | 5690.81315 | 0.472702273 | 0.44710919 | 1.05724124 | 0.2904015 | 0.86724628 | 5950.3572 | 6611.09694 | 3583.51786 | 6618.2806 | Non-sig. |
| PF3D7_1404700 | N/A | 697.296661 | 0.471945889 | 0.63183453 | 0.74694539 | 0.45509652 | 0.92229829 | 942.326518 | 673.136238 | 226.59834 | 947.125546 | Non-sig. |
| PF3D7_0405300 | LISP2 | 71.8031457 | 0.471907996 | 0.625375 | 0.75460004 | 0.450489 | 0.92081798 | 81.1430556 | 71.8670962 | 39.4298733 | 94.7725576 | Non-sig. |
| PF3D7_1473100 | N/A | 710.047419 | 0.471198311 | 0.43568954 | 1.08150017 | 0.27947469 | 0.86247429 | 515.753336 | 767.195777 | 674.275712 | 882.96485 | Non-sig. |
| PF3D7_1129100 | PV1 | 8331.52558 | 0.47114117 | 0.41715981 | 1.12940211 | 0.25872824 | 0.85030738 | 6138.61369 | 8943.63433 | 7827.3449 | 10416.5094 | Non-sig. |
| PF3D7_0411900 | N/A | 4211.42746 | 0.471096626 | 0.41837748 | 1.12600857 | 0.26016189 | 0.85030738 | 3349.81507 | 4249.51948 | 3709.85197 | 5536.52331 | Non-sig. |
| PF3D7_1243600 | N/A | 8494.64914 | 0.471038278 | 0.45313313 | 1.0395141 | 0.2985657 | 0.87330688 | 9022.23837 | 9631.66011 | 5218.31505 | 10106.383 | Non-sig. |
| PF3D7_1334600 | MSRP3 | 1178.18806 | 0.470662668 | 0.42520246 | 1.10691425 | 0.268331 | 0.8556331 | 910.302925 | 1518.86471 | 1064.95958 | 1218.62504 | Non-sig. |
| PF3D7_1121800 | N/A | 127.829399 | 0.470355228 | 0.56819502 | 0.82780597 | 0.40778039 | 0.90723586 | 146.483679 | 144.805539 | 68.1811999 | 151.847176 | Non-sig. |
| PF3D7_1356700 | N/A | 278.370398 | 0.469859973 | 0.47633225 | 0.98641227 | 0.32393085 | 0.88694295 | 188.258324 | 296.214432 | 278.377238 | 350.631596 | Non-sig. |
| PF3D7_1225200 | N/A | 1566.12239 | 0.469785334 | 0.43479875 | 1.08046615 | 0.27993465 | 0.86300615 | 1548.49549 | 1923.45883 | 1078.49862 | 1714.03662 | Non-sig. |
| PF3D7_0324600 | N/A | 5.5715916 | 0.46950269 | 1.78016321 | 0.26374137 | 0.79197921 | 0.99028237 | 5.24321715 | 12.9306449 | 4.11250436 | 0 | Non-sig. |
| PF3D7_1240900 | VAR | 2114.58685 | 0.469401742 | 0.42358261 | 1.10817048 | 0.2677882 | 0.8556331 | 1958.55689 | 2723.21181 | 1588.77799 | 2187.8007 | Non-sig. |
| PF3D7_1421100 | N/A | 560.01535 | 0.469217143 | 0.44849788 | 1.04619701 | 0.29547008 | 0.87191643 | 385.595624 | 618.959135 | 553.636793 | 681.86985 | Non-sig. |
| PF3D7_0508900 | AAP6 | 24808.866 | 0.468852318 | 0.71482191 | 0.65590088 | 0.5118879 | 0.93247207 | 36755.1654 | 29006.0051 | 4870.78744 | 28603.5061 | Non-sig. |
| PF3D7_1248700 | N/A | 26092.1579 | 0.468385809 | 0.42545863 | 1.10089624 | 0.27094182 | 0.8556331 | 23709.6821 | 25389.6721 | 20077.3243 | 35191.953 | Non-sig. |
| PF3D7_0304100 | IMC1e | 3516.8009 | 0.467507019 | 0.40719142 | 1.14812592 | 0.25091659 | 0.8494288 | 3128.74081 | 3932.35839 | 2775.20854 | 4230.89586 | Non-sig. |
| PF3D7_1002800 | N/A | 659.235465 | 0.467059197 | 0.45933026 | 1.01682654 | 0.30923596 | 0.87940531 | 681.699225 | 727.286115 | 425.501742 | 802.454779 | Non-sig. |
| PF3D7_0920500 | N/A | 4077.29507 | 0.466627567 | 0.4515937 | 1.03329068 | 0.30146789 | 0.87381251 | 4275.69917 | 5088.43634 | 2571.84635 | 4373.19841 | Non-sig. |
| PF3D7_0323800 | N/A | 1320.07151 | 0.466296554 | 0.45714878 | 1.0200105 | 0.30772348 | 0.87709318 | 829.371622 | 1443.76486 | 1387.44635 | 1619.70321 | Non-sig. |
| PF3D7_1012800 | N/A | 244.706346 | 0.465041943 | 0.45430066 | 1.02364355 | 0.30600367 | 0.87685698 | 207.51579 | 289.646667 | 203.706651 | 277.956277 | Non-sig. |
| PF3D7_1237300 | N/A | 8.37628352 | 0.464369066 | 1.25691512 | 0.36945141 | 0.71179128 | 0.9875928 | 8.37008158 | 13.9725582 | 5.74340121 | 5.41909307 | Non-sig. |
| PF3D7_0715200 | N/A | 4945.19213 | 0.463931357 | 0.4712148 | 0.98454326 | 0.32484847 | 0.88694295 | 5200.59367 | 4603.50664 | 3113.43331 | 6863.23489 | Non-sig. |
| PF3D7_0911300 | CRMP1 | 101.413502 | 0.463746458 | 0.55750139 | 0.83183014 | 0.40550483 | 0.90723586 | 64.3099042 | 105.106261 | 105.975742 | 130.262101 | Non-sig. |
| PF3D7_0607700 | N/A | 9533.48692 | 0.463088327 | 0.440949 | 1.05020837 | 0.29362232 | 0.87149079 | 7327.1038 | 8663.76165 | 8705.61793 | 13437.4643 | Non-sig. |
| PF3D7_1434400 | N/A | 2377.73841 | 0.462153756 | 0.42242602 | 1.09404661 | 0.27393454 | 0.85618956 | 2257.20761 | 2961.38098 | 1743.23394 | 2549.1311 | Non-sig. |
| PF3D7_0615800 | COG4 | 1538.48415 | 0.461430453 | 0.40937181 | 1.12716714 | 0.25967182 | 0.85030738 | 1249.29766 | 1771.13996 | 1339.69781 | 1793.80119 | Non-sig. |
| PF3D7_1209500 | PDEalpha | 1744.98437 | 0.461134595 | 0.49180518 | 0.93763672 | 0.34843113 | 0.8930363 | 950.645701 | 2148.70389 | 1986.06223 | 1894.52567 | Non-sig. |
| PF3D7_0213500 | N/A | 308.017911 | 0.46079057 | 0.45809873 | 1.00587612 | 0.31447517 | 0.88454055 | 283.782837 | 397.170708 | 234.828694 | 316.289402 | Non-sig. |
| PF3D7_0908100 | N/A | 142.231783 | 0.460199519 | 0.50583356 | 0.90978446 | 0.36293619 | 0.89567376 | 99.7393185 | 161.065092 | 139.556389 | 168.566333 | Non-sig. |
| PF3D7_1402200 | ApiCOX19 | 4437.33501 | 0.459968228 | 0.41824535 | 1.09975694 | 0.27143804 | 0.8556331 | 3913.57351 | 5821.89398 | 3558.30584 | 4455.56671 | Non-sig. |
| PF3D7_1343300 | N/A | 4749.13256 | 0.459547699 | 0.49139867 | 0.93518304 | 0.34969398 | 0.8930363 | 5457.8967 | 5324.82568 | 2540.69614 | 5673.11172 | Non-sig. |
| PF3D7_0819800 | N/A | 592.515218 | 0.458771981 | 0.68923657 | 0.66562339 | 0.50565186 | 0.93170321 | 853.216469 | 762.741518 | 145.386783 | 608.716103 | Non-sig. |
| PF3D7_1143100 | AP2-O | 17000.6068 | 0.458177418 | 0.59921423 | 0.76463041 | 0.44449166 | 0.91974025 | 22411.7343 | 15466.1516 | 6235.74272 | 23888.7984 | Non-sig. |
| PF3D7_1475600 | BDP4 | 1484.70215 | 0.458001448 | 0.41322169 | 1.10836741 | 0.26770317 | 0.8556331 | 1191.02347 | 1618.93651 | 1310.88389 | 1817.96472 | Non-sig. |
| PF3D7_1357500 | N/A | 2395.46422 | 0.457293698 | 0.48703452 | 0.93893488 | 0.34776418 | 0.8930363 | 1313.10741 | 2719.72675 | 2724.41107 | 2824.61167 | Non-sig. |
| PF3D7_1203900 | UBC | 1283.53033 | 0.457101689 | 0.42119492 | 1.08524977 | 0.27781105 | 0.85972841 | 969.979585 | 1398.70972 | 1193.61745 | 1571.81455 | Non-sig. |
| PF3D7_1301500 | N/A | 226.843375 | 0.456514408 | 0.48513665 | 0.9410017 | 0.34670399 | 0.8930363 | 159.789967 | 296.251863 | 222.520017 | 228.811653 | Non-sig. |
| PF3D7_1242800 | rabGDI | 6036.83732 | 0.456215797 | 0.4922765 | 0.92674705 | 0.35405788 | 0.8930363 | 3242.70198 | 7490.7692 | 6937.31636 | 6476.56173 | Non-sig. |
| PF3D7_1244000 | N/A | 1050.87892 | 0.455595094 | 0.45723746 | 0.99640807 | 0.31905192 | 0.88651753 | 674.112382 | 1121.06356 | 1098.19447 | 1310.14527 | Non-sig. |
| PF3D7_0821300 | DHX36 | 949.211567 | 0.455075049 | 0.45517908 | 0.99977144 | 0.31742113 | 0.88651753 | 990.720784 | 1050.38934 | 611.065134 | 1144.67101 | Non-sig. |
| PF3D7_1242100 | N/A | 17.9199718 | 0.455008561 | 1.0044289 | 0.45300226 | 0.65054712 | 0.97317009 | 7.20518945 | 11.8831415 | 22.8924944 | 29.6990618 | Non-sig. |
| PF3D7_1468300 | LRR14.1 | 45.7066543 | 0.45482256 | 0.6806174 | 0.66824998 | 0.50397404 | 0.93170321 | 26.0895638 | 57.0051694 | 50.8496215 | 48.8822625 | Non-sig. |

| | | | | | | | | | | | | |
|---|---|---|---|---|---|---|---|---|---|---|---|---|
| PF3D7_1478600 | PTP3 | 1339.25763 | 0.454315402 | 0.58290794 | 0.77939477 | 0.4357472 | 0.91522571 | 1751.37763 | 1595.03274 | 509.300109 | 1501.32004 | Non-sig. |
| PF3D7_1146600 | ORP1 | 11024.4605 | 0.453871458 | 0.44271831 | 1.02519243 | 0.3052724 | 0.87685698 | 11238.9086 | 11237.4702 | 7370.34892 | 14251.1143 | Non-sig. |
| PF3D7_0504100 | N/A | 691.779574 | 0.45310745 | 0.46443346 | 0.97561328 | 0.32925614 | 0.88909228 | 715.284428 | 899.003587 | 453.065274 | 699.765006 | Non-sig. |
| PF3D7_0805800 | N/A | 13.5159811 | 0.45232031 | 0.99442297 | 0.45485706 | 0.64921208 | 0.97317009 | 10.4935481 | 13.960963 | 12.3126974 | 17.296716 | Non-sig. |
| PF3D7_0423600 | N/A | 2892.80722 | 0.452314115 | 0.58217332 | 0.77694065 | 0.43719379 | 0.91524921 | 3789.03186 | 3050.70032 | 1097.44189 | 3634.0548 | Non-sig. |
| PF3D7_0821100 | PK1 | 1867.27974 | 0.452287671 | 0.47811519 | 0.94598055 | 0.34415851 | 0.8930363 | 1066.532 | 2129.74367 | 2086.9869 | 2185.85637 | Non-sig. |
| PF3D7_0220300 | N/A | 230.627341 | 0.452088501 | 0.59999327 | 0.75348929 | 0.45115595 | 0.92081798 | 294.695671 | 239.28393 | 95.2680031 | 293.26176 | Non-sig. |
| PF3D7_0300100 | VAR | 1426.73866 | 0.450718159 | 0.41750822 | 1.07954319 | 0.28034565 | 0.86335738 | 1253.12389 | 1805.19823 | 1158.27645 | 1490.35606 | Non-sig. |
| PF3D7_0703000 | N/A | 294.960014 | 0.450336407 | 0.57014196 | 0.78986715 | 0.42960536 | 0.91334237 | 135.937824 | 288.542904 | 362.260504 | 393.098823 | Non-sig. |
| PF3D7_0408100 | N/A | 1531.73423 | 0.448715625 | 0.67505355 | 0.66471116 | 0.50623525 | 0.93170321 | 2209.33471 | 1728.13754 | 381.961952 | 1807.50271 | Non-sig. |
| PF3D7_1476000 | N/A | 41.2626708 | 0.44837464 | 0.70320969 | 0.63761157 | 0.52372656 | 0.93247207 | 47.0209203 | 46.2333841 | 22.9689208 | 48.8274581 | Non-sig. |
| PF3D7_1400600 | RIF | 2.81377457 | 0.446409791 | 1.9697753 | 0.2266298 | 0.82071162 | 0.99096399 | 3.14433033 | 3.22283311 | 1.64134866 | 3.24658616 | Non-sig. |
| PF3D7_1013400 | N/A | 658.851628 | 0.446244865 | 0.52997108 | 0.84201738 | 0.39977822 | 0.90645858 | 788.246505 | 843.414763 | 327.684398 | 676.060845 | Non-sig. |
| PF3D7_0933000 | N/A | 2954.28553 | 0.446141309 | 0.42525437 | 1.04911635 | 0.29412457 | 0.87170077 | 2907.20714 | 3302.34729 | 2095.23815 | 3512.34953 | Non-sig. |
| PF3D7_1362700 | N/A | 3162.86878 | 0.445667429 | 0.67970174 | 0.65568087 | 0.51202948 | 0.93247207 | 4596.10836 | 3558.69985 | 760.711607 | 3735.9553 | Non-sig. |
| PF3D7_0308100 | N/A | 9863.92533 | 0.445571231 | 0.48700041 | 0.9149299 | 0.36022843 | 0.89560738 | 11320.3333 | 10826.0938 | 5385.49253 | 11923.7817 | Non-sig. |
| PF3D7_1113500 | N/A | 840.601597 | 0.445421786 | 0.48115734 | 0.92573 | 0.35458632 | 0.89325348 | 492.771765 | 913.032825 | 930.574544 | 1026.02725 | Non-sig. |
| PF3D7_0418000 | N/A | 1423.17573 | 0.445290854 | 0.43981145 | 1.01245854 | 0.31131886 | 0.8804206 | 1396.15342 | 1429.56956 | 1014.61341 | 1852.36656 | Non-sig. |
| PF3D7_0619200 | N/A | 2506.03099 | 0.445276311 | 0.4752588 | 0.93691335 | 0.34880313 | 0.8930363 | 2787.93972 | 3087.52995 | 1457.12721 | 2691.52708 | Non-sig. |
| PF3D7_1035700 | DBLMSP | 22497.0927 | 0.444512074 | 0.40175598 | 1.10642304 | 0.26854346 | 0.8556331 | 18156.0165 | 26271.6102 | 19960.7944 | 25599.9496 | Non-sig. |
| PF3D7_0104400 | LytB | 1274.17101 | 0.444170856 | 0.4386592 | 1.01256478 | 0.31126809 | 0.8804206 | 1290.48142 | 1463.55896 | 868.925823 | 1473.71784 | Non-sig. |
| PF3D7_0407700 | N/A | 10912.7247 | 0.444008849 | 0.56115281 | 0.79124409 | 0.42880157 | 0.91236574 | 14052.0046 | 12297.7174 | 4441.64156 | 12859.5353 | Non-sig. |
| PF3D7_1145000 | N/A | 622.786584 | 0.443564735 | 0.62805313 | 0.70625352 | 0.48003049 | 0.92880667 | 850.730357 | 798.520926 | 205.288165 | 636.606888 | Non-sig. |
| PF3D7_1122200 | JmjC3 | 948.586492 | 0.442439461 | 0.48364351 | 0.91480492 | 0.36029405 | 0.89560738 | 1062.02216 | 1143.06715 | 546.880783 | 1042.37587 | Non-sig. |
| PF3D7_1236700 | N/A | 1233.90501 | 0.442379564 | 0.47463671 | 0.93203824 | 0.35131677 | 0.8930363 | 728.866917 | 1486.42504 | 1363.15279 | 1357.17531 | Non-sig. |
| PF3D7_0725000 | N/A | 3670.12926 | 0.441931249 | 0.42498431 | 1.03987663 | 0.29839722 | 0.87330688 | 3626.76848 | 4346.49597 | 2598.21052 | 4109.04205 | Non-sig. |
| PF3D7_0407900 | N/A | 9773.87685 | 0.441365559 | 0.59191759 | 0.74565374 | 0.45587661 | 0.92247173 | 13079.0843 | 11046.8685 | 3502.11078 | 11467.4438 | Non-sig. |
| PF3D7_0221400 | N/A | 2.81397252 | 0.440669656 | 2.25760097 | 0.19519378 | 0.84524125 | 0.99165628 | 3.146402 | 6.46453181 | 1.64495629 | 0 | Non-sig. |
| PF3D7_1363400 | DOA1 | 2234.38766 | 0.440486657 | 0.4879911 | 0.90265306 | 0.36671006 | 0.89567376 | 1230.30719 | 2628.29951 | 2561.10369 | 2517.84026 | Non-sig. |
| PF3D7_1217200 | MRD1 | 1673.63598 | 0.439531537 | 0.54310515 | 0.80929363 | 0.41834627 | 0.9088998 | 2088.53108 | 1800.17821 | 753.201597 | 2052.63301 | Non-sig. |
| PF3D7_0419500 | N/A | 731.890396 | 0.439075173 | 0.48165238 | 0.91160179 | 0.36197838 | 0.89567376 | 431.711061 | 802.519873 | 810.662427 | 882.668224 | Non-sig. |
| PF3D7_0728200 | ALP3 | 365.338839 | 0.438906446 | 0.47188128 | 0.93012049 | 0.3523087 | 0.8930363 | 244.018229 | 458.263949 | 376.101707 | 382.971471 | Non-sig. |
| PF3D7_0322200 | N/A | 109.530652 | 0.438809414 | 0.64419495 | 0.68117487 | 0.49576085 | 0.93170321 | 142.005233 | 125.563518 | 44.3445184 | 126.209336 | Non-sig. |
| PF3D7_0420000 | N/A | 5754.41602 | 0.438634643 | 0.48826267 | 0.89835793 | 0.36899476 | 0.89654416 | 6603.56307 | 6092.75628 | 3169.4748 | 7151.86995 | Non-sig. |
| PF3D7_0515600 | GEP | 222.507875 | 0.436595638 | 0.4717724 | 0.925437 | 0.35473864 | 0.89325348 | 214.899035 | 264.969734 | 163.459431 | 246.703303 | Non-sig. |
| PF3D7_1441600 | N/A | 470.2048 | 0.435990563 | 0.52814355 | 0.82551527 | 0.40907912 | 0.90748263 | 247.170379 | 458.265648 | 551.833545 | 623.549629 | Non-sig. |
| PF3D7_0934900 | N/A | 161.082596 | 0.4358547 | 0.55559776 | 0.78447886 | 0.43275918 | 0.91381353 | 187.154187 | 200.711808 | 87.0399379 | 169.42445 | Non-sig. |
| PF3D7_1206100 | ISN1 | 265.890395 | 0.435289036 | 0.46188948 | 0.9424095 | 0.34598302 | 0.8930363 | 232.167403 | 345.693571 | 220.030273 | 265.670335 | Non-sig. |
| PF3D7_0202000 | KAHRP | 6697.55624 | 0.435113132 | 0.46873315 | 0.92827471 | 0.35326509 | 0.8930363 | 7333.03869 | 6759.64819 | 4057.64809 | 8639.88998 | Non-sig. |
| PF3D7_0503500 | N/A | 988.908346 | 0.435096419 | 0.49746167 | 0.87463305 | 0.3817736 | 0.90292925 | 538.095513 | 1129.72923 | 1143.31846 | 1144.49018 | Non-sig. |
| PF3D7_0929200 | N/A | 2167.30187 | 0.434937759 | 0.46133211 | 0.94278666 | 0.34579004 | 0.8930363 | 2358.68009 | 2450.30588 | 1327.8019 | 2532.41961 | Non-sig. |
| PF3D7_1017400 | HAD5 | 1890.38574 | 0.434091438 | 0.5146544 | 0.84346202 | 0.3989701 | 0.90645858 | 2231.26667 | 2508.31783 | 985.368352 | 1836.59011 | Non-sig. |
| PF3D7_1443000 | SRPK2 | 1683.75236 | 0.434056684 | 0.45603154 | 0.95181285 | 0.3411919 | 0.8930363 | 1092.57474 | 1721.69088 | 1771.81723 | 2148.92662 | Non-sig. |
| PF3D7_0211700 | TKL1 | 1402.28853 | 0.433922303 | 0.46817759 | 0.92683271 | 0.3540134 | 0.8930363 | 1540.26671 | 1588.70006 | 846.061328 | 1634.12603 | Non-sig. |
| PF3D7_0216000 | N/A | 881.1163 | 0.432694803 | 0.6017156 | 0.71910185 | 0.47207817 | 0.92682482 | 1178.34845 | 897.760869 | 322.019835 | 1126.33605 | Non-sig. |
| PF3D7_0932500 | DHHC6 | 542.429046 | 0.432127411 | 0.4266336 | 1.01287711 | 0.31111886 | 0.8804206 | 482.696352 | 636.44814 | 440.962167 | 609.609525 | Non-sig. |
| PF3D7_0814400 | N/A | 1192.51389 | 0.431483775 | 0.4138873 | 1.04251515 | 0.29717291 | 0.87301237 | 955.826332 | 1389.49394 | 1075.08363 | 1349.65167 | Non-sig. |
| PF3D7_1456600 | N/A | 79.9105513 | 0.431039419 | 0.58458602 | 0.7373413 | 0.46091482 | 0.92336312 | 52.2618311 | 78.4825594 | 83.676275 | 105.22154 | Non-sig. |
| PF3D7_1239100 | N/A | 464.210665 | 0.430845992 | 0.51347789 | 0.83907409 | 0.40142773 | 0.90667178 | 249.269159 | 517.295638 | 541.156369 | 549.121493 | Non-sig. |
| PF3D7_0717200 | N/A | 2616.61926 | 0.430293243 | 0.5178583 | 0.83090923 | 0.40602491 | 0.90723586 | 3145.06995 | 3343.01933 | 1313.90558 | 2664.48219 | Non-sig. |
| PF3D7_0933700 | N/A | 388.616709 | 0.429905888 | 0.4851455 | 0.88613806 | 0.37554314 | 0.89654416 | 404.907193 | 523.431899 | 257.633048 | 368.494697 | Non-sig. |
| PF3D7_0911200 | N/A | 288.006256 | 0.429640283 | 0.46368738 | 0.92657316 | 0.3541482 | 0.8930363 | 205.801757 | 325.340925 | 284.865018 | 336.017323 | Non-sig. |
| PF3D7_0818500 | N/A | 4619.19473 | 0.429605139 | 0.49843108 | 0.86191483 | 0.38873439 | 0.90444842 | 5399.74294 | 5827.75161 | 2473.63851 | 4775.64585 | Non-sig. |
| PF3D7_1007200 | N/A | 4398.45335 | 0.428312886 | 0.40470928 | 1.05832239 | 0.28990849 | 0.86724628 | 3936.03967 | 4945.97674 | 3564.63047 | 5147.1665 | Non-sig. |

| | | | | | | | | | | | | |
|---|---|---|---|---|---|---|---|---|---|---|---|---|
| PF3D7_1361800 | GAC | 52803.2001 | 0.428171723 | 0.59437634 | 0.72037141 | 0.47129635 | 0.92682482 | 18827.0567 | 57130.2139 | 71221.7498 | 64033.7802 | Non-sig. |
| PF3D7_0932100 | N/A | 10709.5575 | 0.426464447 | 0.60922828 | 0.70000763 | 0.48392254 | 0.93078439 | 14666.4282 | 11252.3507 | 3610.23362 | 13309.2175 | Non-sig. |
| PF3D7_0300400 | N/A | 3.76163041 | 0.425973904 | 1.72812554 | 0.24649477 | 0.80529925 | 0.99028237 | 3.13755008 | 5.37466367 | 3.28173772 | 3.25257019 | Non-sig. |
| PF3D7_0310200 | N/A | 9689.17337 | 0.425837948 | 0.46379677 | 0.91815635 | 0.35853702 | 0.89558186 | 9784.48016 | 8445.55357 | 6754.7553 | 13771.9044 | Non-sig. |
| PF3D7_0731100 | PTP2 | 2935.6986 | 0.425787655 | 0.40877121 | 1.04162829 | 0.29758405 | 0.87301237 | 2318.35772 | 3310.18315 | 2692.72867 | 3421.52488 | Non-sig. |
| PF3D7_0630600 | N/A | 3350.30878 | 0.424929544 | 0.47384938 | 0.89676079 | 0.36984657 | 0.89654416 | 1946.56518 | 3822.8897 | 3773.93932 | 3857.84093 | Non-sig. |
| PF3D7_1232500 | N/A | 5391.85137 | 0.42457037 | 0.53519931 | 0.79329394 | 0.42760659 | 0.91236574 | 6666.5867 | 5306.19051 | 2542.13927 | 7052.489 | Non-sig. |
| PF3D7_0204500 | AspAT | 406.010758 | 0.42454567 | 0.5417666 | 0.78363205 | 0.43325604 | 0.91449822 | 197.230401 | 469.335382 | 495.769855 | 461.707395 | Non-sig. |
| PF3D7_0820500 | YIF1 | 1087.34182 | 0.424383003 | 0.55219263 | 0.7685416 | 0.44216549 | 0.91830371 | 488.413442 | 1093.37135 | 1368.27531 | 1399.30719 | Non-sig. |
| PF3D7_0803400 | RAD54 | 5127.34468 | 0.424363761 | 0.45213426 | 0.93857909 | 0.34794689 | 0.8930363 | 5461.93125 | 5341.74807 | 3295.68604 | 6410.01337 | Non-sig. |
| PF3D7_1204200 | N/A | 99.0526699 | 0.423971921 | 0.53466957 | 0.79296063 | 0.42780076 | 0.91236574 | 74.6590753 | 104.109529 | 94.4318259 | 123.010249 | Non-sig. |
| PF3D7_1445800 | N/A | 210.539259 | 0.42351369 | 0.64302624 | 0.65862583 | 0.51013607 | 0.93247207 | 72.6626802 | 227.444344 | 286.653712 | 255.396299 | Non-sig. |
| PF3D7_1019600 | N/A | 3262.47049 | 0.423363975 | 0.40879609 | 1.03563606 | 0.30037196 | 0.87381251 | 2622.97787 | 3532.59285 | 2951.29451 | 3943.01672 | Non-sig. |
| PF3D7_1037500 | DYN2 | 2459.87583 | 0.423200221 | 0.5357009 | 0.78999349 | 0.42953157 | 0.91334237 | 1139.61222 | 2621.54087 | 3063.25166 | 3015.09858 | Non-sig. |
| PF3D7_1310700 | PIC5 | 17677.7228 | 0.419757661 | 0.50610334 | 0.8293912 | 0.40688308 | 0.90723586 | 21234.6792 | 20637.0241 | 9013.79802 | 19825.3899 | Non-sig. |
| PF3D7_1418300 | N/A | 291.113042 | 0.419736676 | 0.51750712 | 0.81107421 | 0.41732305 | 0.90869129 | 326.996851 | 384.940093 | 171.441999 | 281.073223 | Non-sig. |
| PF3D7_1105300 | N/A | 2457.3702 | 0.419261858 | 0.60093188 | 0.69768617 | 0.48537348 | 0.93170321 | 873.854044 | 2553.55259 | 3331.28944 | 3070.78471 | Non-sig. |
| PF3D7_1014600 | ADA2 | 23971.6024 | 0.418514752 | 0.43023876 | 0.97275 | 0.33067757 | 0.88975609 | 24489.1569 | 26171.2214 | 16548.7387 | 28677.2926 | Non-sig. |
| PF3D7_1227600 | SPC24 | 8778.32299 | 0.417787938 | 0.40559166 | 1.03007034 | 0.30297699 | 0.87475317 | 6942.7216 | 10087.8976 | 8089.31829 | 9993.3545 | Non-sig. |
| PF3D7_1227100 | DH60 | 1282.63447 | 0.417239821 | 0.50448345 | 0.82706344 | 0.4082011 | 0.90723586 | 680.260329 | 1538.48751 | 1516.21365 | 1395.57638 | Non-sig. |
| PF3D7_0207100 | N/A | 17960.5315 | 0.416930572 | 0.54546948 | 0.76435179 | 0.44465763 | 0.91974025 | 22878.358 | 19461.7308 | 7888.51619 | 21613.5211 | Non-sig. |
| PF3D7_1236200 | N/A | 1718.44763 | 0.416678378 | 0.51584605 | 0.80775723 | 0.41923035 | 0.90940098 | 2074.45863 | 1824.42526 | 869.963087 | 2104.94353 | Non-sig. |
| PF3D7_0931900 | AKLP2 | 676.344874 | 0.416392723 | 0.44855616 | 0.92829563 | 0.35325424 | 0.8930363 | 691.864708 | 812.498883 | 467.226787 | 733.78912 | Non-sig. |
| PF3D7_0726100 | N/A | 250.776083 | 0.415989234 | 0.47533128 | 0.87515644 | 0.38148879 | 0.90292925 | 221.147176 | 233.881481 | 208.624746 | 339.45093 | Non-sig. |
| PF3D7_0723500 | N/A | 53.6477162 | 0.415862076 | 0.61833426 | 0.67255221 | 0.50123221 | 0.93170321 | 40.9619664 | 67.6397563 | 50.9006133 | 55.0885289 | Non-sig. |
| PF3D7_0937800 | VAR | 1989.86622 | 0.415605329 | 0.45368364 | 0.91606859 | 0.35963092 | 0.89560738 | 2033.07956 | 2667.55729 | 1377.61517 | 1881.21285 | Non-sig. |
| PF3D7_1233000 | EF-G | 1034.66236 | 0.414512218 | 0.44974815 | 0.92165408 | 0.35670904 | 0.89552149 | 686.361159 | 1194.20446 | 1087.40838 | 1170.67544 | Non-sig. |
| PF3D7_0611400 | SWIB | 1146.43797 | 0.411667523 | 0.48761678 | 0.84424396 | 0.39853309 | 0.90645858 | 651.772753 | 1351.84159 | 1315.78444 | 1266.35311 | Non-sig. |
| PF3D7_1431400 | SRA | 10721.7748 | 0.411341585 | 0.61712503 | 0.66654497 | 0.50506283 | 0.93170321 | 14918.1718 | 12854.3862 | 3489.39223 | 11625.1487 | Non-sig. |
| PF3D7_1422600 | N/A | 940.732765 | 0.411313744 | 0.43312338 | 0.94964568 | 0.34229232 | 0.8930363 | 680.364826 | 1050.74589 | 934.469525 | 1097.35082 | Non-sig. |
| PF3D7_1348700 | WDR16 | 70.3254551 | 0.410898268 | 0.61414287 | 0.66905974 | 0.50345737 | 0.93170321 | 79.9304169 | 83.7129191 | 41.0585454 | 76.5999392 | Non-sig. |
| PF3D7_0707600 | MED10 | 368.761576 | 0.410882608 | 0.4372967 | 0.93959686 | 0.34742439 | 0.8930363 | 304.732066 | 421.88759 | 328.418168 | 420.008481 | Non-sig. |
| PF3D7_1478100 | N/A | 158.493922 | 0.410861387 | 0.48670096 | 0.84417624 | 0.39857093 | 0.90645858 | 125.953486 | 192.218484 | 146.116999 | 169.686717 | Non-sig. |
| PF3D7_1130700 | SMC1 | 2593.76494 | 0.410383934 | 0.48402751 | 0.8478525 | 0.39652012 | 0.90645858 | 2955.04358 | 2630.07813 | 1499.99328 | 3289.94479 | Non-sig. |
| PF3D7_1450400 | N/A | 3112.91747 | 0.409628218 | 0.42459811 | 0.9647434 | 0.33467335 | 0.89204385 | 3061.90052 | 3282.10882 | 2286.18845 | 3821.47207 | Non-sig. |
| PF3D7_0807700 | DegP | 1305.039 | 0.408308296 | 0.55171819 | 0.74006677 | 0.45925948 | 0.92288312 | 1668.69269 | 1523.56172 | 574.923509 | 1452.97807 | Non-sig. |
| PF3D7_1038400 | Pf11-1 | 381.999278 | 0.408234898 | 0.70332333 | 0.58043702 | 0.56161995 | 0.94472957 | 444.025571 | 142.627751 | 212.788145 | 728.555645 | Non-sig. |
| PF3D7_0627900 | POP4 | 986.382175 | 0.408164275 | 0.4282308 | 0.95314087 | 0.3405187 | 0.8930363 | 734.488453 | 1168.5863 | 960.859451 | 1081.5945 | Non-sig. |
| PF3D7_1019100 | N/A | 6749.40005 | 0.407120857 | 0.73993591 | 0.55021097 | 0.58217468 | 0.95129458 | 10416.8012 | 7634.87146 | 1190.30887 | 7755.61871 | Non-sig. |
| PF3D7_1108100 | N/A | 4710.23235 | 0.405958235 | 0.44692384 | 0.90833874 | 0.36369928 | 0.89567376 | 4626.04541 | 4275.87244 | 3477.88412 | 6461.12744 | Non-sig. |
| PF3D7_1334100 | N/A | 1478.63567 | 0.405746613 | 0.40893262 | 0.99220897 | 0.3210956 | 0.88694295 | 1254.52811 | 1670.31341 | 1289.58609 | 1700.11508 | Non-sig. |
| PF3D7_0220700 | N/A | 355.013943 | 0.405491795 | 0.48884305 | 0.82949281 | 0.4068256 | 0.90723586 | 338.418708 | 298.419357 | 272.595714 | 510.621992 | Non-sig. |
| PF3D7_0730200 | N/A | 475.200285 | 0.404992521 | 0.55248956 | 0.73303198 | 0.46353892 | 0.92509266 | 232.693066 | 440.961962 | 584.789703 | 642.356409 | Non-sig. |
| PF3D7_1315100 | PK9 | 105.613153 | 0.40444428 | 0.53131874 | 0.76120839 | 0.44653261 | 0.91974025 | 104.811011 | 112.799159 | 77.1433871 | 127.699056 | Non-sig. |
| PF3D7_1368400 | N/A | 1878.05264 | 0.403948044 | 0.47075985 | 0.85807667 | 0.39085013 | 0.90538866 | 2083.46417 | 2354.35518 | 1150.56357 | 1923.82766 | Non-sig. |
| PF3D7_0613500 | N/A | 1541.20502 | 0.403574547 | 0.41836956 | 0.9646365 | 0.33472691 | 0.89204385 | 1177.81159 | 1746.40654 | 1476.08091 | 1764.52105 | Non-sig. |
| PF3D7_1339900 | ABCB5 | 622.057701 | 0.403294322 | 0.55185302 | 0.73080024 | 0.46490118 | 0.92538923 | 283.254178 | 690.932978 | 787.669573 | 726.374076 | Non-sig. |
| PF3D7_0207400 | SERA7 | 9488.82269 | 0.40323355 | 0.40145593 | 1.00442794 | 0.31517239 | 0.88486757 | 7951.65483 | 10987.1298 | 8390.95532 | 10625.5508 | Non-sig. |
| PF3D7_0422200 | N/A | 3942.84742 | 0.402926807 | 0.44336357 | 0.90879547 | 0.3634581 | 0.89567376 | 3886.82022 | 3657.05272 | 2904.99137 | 5322.52537 | Non-sig. |
| PF3D7_0623600 | N/A | 947.932849 | 0.402597889 | 0.50347567 | 0.79963723 | 0.42392101 | 0.91236574 | 514.82476 | 1144.81458 | 1117.81712 | 1014.27493 | Non-sig. |
| PF3D7_1417300 | ATG4 | 1903.5576 | 0.402338302 | 0.54393198 | 0.73968496 | 0.45949118 | 0.92288312 | 2420.91313 | 2222.7062 | 859.197347 | 2111.41371 | Non-sig. |
| PF3D7_0205900 | RPN1 | 12682.2544 | 0.401503289 | 0.51033439 | 0.78674551 | 0.43143086 | 0.91355742 | 6418.5366 | 14854.0046 | 15438.6744 | 14017.8021 | Non-sig. |
| PF3D7_1209900 | ABCB7 | 928.246148 | 0.401072188 | 0.42858412 | 0.9358074 | 0.34937236 | 0.8930363 | 690.675139 | 1054.67532 | 909.208831 | 1058.4253 | Non-sig. |
| PF3D7_1361100 | SEC24A | 6946.71761 | 0.400587497 | 0.54856812 | 0.73024203 | 0.46524225 | 0.92538923 | 3067.86573 | 7195.4531 | 8908.56176 | 8614.98983 | Non-sig. |

| | | | | | | | | | | | | |
|---|---|---|---|---|---|---|---|---|---|---|---|---|
| PF3D7_1017700 | N/A | 250.632226 | 0.400364066 | 0.79540876 | 0.5033438 | 0.61472258 | 0.96217474 | 392.270411 | 342.369198 | 40.2370211 | 227.652274 | Non-sig. |
| PF3D7_0809400 | N/A | 2364.68564 | 0.399103381 | 0.47237525 | 0.8448863 | 0.39817432 | 0.90645858 | 2672.83561 | 2776.81243 | 1406.89871 | 2602.19582 | Non-sig. |
| PF3D7_0606500 | PTB | 1251.57992 | 0.398536206 | 0.41791865 | 0.95362149 | 0.34027527 | 0.8930363 | 980.740851 | 1376.34051 | 1178.71513 | 1470.52319 | Non-sig. |
| PF3D7_1312000 | MCAT | 132.059946 | 0.398197191 | 0.51991761 | 0.76588518 | 0.44374462 | 0.91974025 | 130.337136 | 172.820232 | 97.7093956 | 127.37302 | Non-sig. |
| PF3D7_1347700 | ECT | 3428.11324 | 0.398113412 | 0.42008033 | 0.94770782 | 0.34327822 | 0.8930363 | 2747.14236 | 3402.83697 | 3168.93822 | 4393.53539 | Non-sig. |
| PF3D7_0416100 | GATA | 2381.84391 | 0.397739458 | 0.41808816 | 0.95132916 | 0.34143731 | 0.8930363 | 1912.29655 | 2418.24136 | 2198.75335 | 2998.08439 | Non-sig. |
| PF3D7_1446000 | N/A | 3.87345037 | 0.397635028 | 1.73737556 | 0.22887108 | 0.81896912 | 0.99096399 | 5.09058593 | 4.43056425 | 1.63573007 | 4.33692124 | Non-sig. |
| PF3D7_0407800 | CINCH | 25874.3556 | 0.397384259 | 0.41943311 | 0.94743178 | 0.3434188 | 0.8930363 | 20647.8593 | 25345.2451 | 24018.4914 | 33485.8267 | Non-sig. |
| PF3D7_1322500 | DHHC5 | 3797.69865 | 0.397190478 | 0.41831555 | 0.94949966 | 0.34236654 | 0.8930363 | 3057.80945 | 4842.86529 | 3498.49313 | 3791.62674 | Non-sig. |
| PF3D7_0724900 | N/A | 3242.20194 | 0.397004848 | 0.70842986 | 0.56040107 | 0.5752059 | 0.94896603 | 4846.56311 | 2675.15516 | 751.634921 | 4695.45459 | Non-sig. |
| PF3D7_0914000 | N/A | 362.161585 | 0.396663479 | 0.47882739 | 0.82840599 | 0.40744062 | 0.90723586 | 361.089278 | 328.322485 | 264.461982 | 494.772594 | Non-sig. |
| PF3D7_0206800 | MSP2 | 44843.4814 | 0.395945497 | 0.44295321 | 0.89387657 | 0.37138794 | 0.89654416 | 31006.385 | 58011.5547 | 46449.7924 | 43906.1937 | Non-sig. |
| PF3D7_1445900 | DDX17 | 3170.5855 | 0.395906329 | 0.44647711 | 0.88673377 | 0.37522225 | 0.89654416 | 2078.35307 | 3568.50981 | 3397.83221 | 3637.6469 | Non-sig. |
| PF3D7_0914900 | N/A | 3038.31021 | 0.395222726 | 0.40717551 | 0.97064463 | 0.33172527 | 0.88994703 | 2469.81394 | 3364.33094 | 2779.53919 | 3539.55676 | Non-sig. |
| PF3D7_1244100 | N/A | 4490.18116 | 0.394337957 | 0.40529833 | 0.97295727 | 0.33057454 | 0.88975609 | 4076.57966 | 5269.04024 | 3684.13506 | 4930.96968 | Non-sig. |
| PF3D7_1428200 | MFS5 | 3848.85877 | 0.393603826 | 0.4311666 | 0.91288108 | 0.36130509 | 0.89560738 | 2984.00523 | 3659.27859 | 3669.95827 | 5082.193 | Non-sig. |
| PF3D7_1306900 | SNRPA | 464.302221 | 0.393491884 | 0.43265377 | 0.90948448 | 0.36309445 | 0.89567376 | 399.56283 | 564.561027 | 403.179133 | 489.905893 | Non-sig. |
| PF3D7_1471600 | APR2 | 121.29205 | 0.393433581 | 0.55683243 | 0.70655652 | 0.47984212 | 0.92880667 | 109.537077 | 94.406853 | 100.208232 | 181.016039 | Non-sig. |
| PF3D7_1432400 | LRR5 | 247.614738 | 0.393278818 | 0.45700724 | 0.86055271 | 0.38948444 | 0.90444842 | 228.626666 | 272.467895 | 199.606249 | 289.758145 | Non-sig. |
| PF3D7_0506800 | TCF25 | 1600.62212 | 0.392635748 | 0.41050184 | 0.95647745 | 0.33883107 | 0.8930363 | 1456.63916 | 1780.85425 | 1311.74323 | 1853.25186 | Non-sig. |
| PF3D7_1324700 | YKT6.2 | 922.18861 | 0.391646207 | 0.45651261 | 0.85790884 | 0.3909428 | 0.90538866 | 603.360896 | 1058.37689 | 991.881056 | 1035.1356 | Non-sig. |
| PF3D7_0619300 | N/A | 1515.28436 | 0.391498376 | 0.45491272 | 0.86060108 | 0.38945779 | 0.90444842 | 1621.57017 | 1555.11199 | 1000.62074 | 1883.83456 | Non-sig. |
| PF3D7_0523900 | N/A | 344.451415 | 0.391173214 | 0.44027866 | 0.88846734 | 0.37428942 | 0.89654416 | 301.216925 | 373.339693 | 294.873328 | 408.375714 | Non-sig. |
| PF3D7_1367400 | N/A | 457.063581 | 0.390825506 | 0.46239575 | 0.84521863 | 0.39798878 | 0.90645858 | 308.62405 | 514.935009 | 482.156091 | 522.539173 | Non-sig. |
| PF3D7_1422200 | N/A | 1084.36369 | 0.390353141 | 0.42251764 | 0.92387418 | 0.35555183 | 0.89402637 | 830.295722 | 1226.31076 | 1046.6362 | 1234.21206 | Non-sig. |
| PF3D7_1420300 | HEP1 | 444.114864 | 0.390274591 | 0.44329118 | 0.88040233 | 0.37864139 | 0.9000162 | 411.782769 | 561.493411 | 357.134804 | 446.04847 | Non-sig. |
| PF3D7_0106500 | N/A | 1464.14648 | 0.390110977 | 0.46159069 | 0.84514481 | 0.39802999 | 0.90645858 | 1611.85739 | 1715.29603 | 923.269737 | 1606.16275 | Non-sig. |
| PF3D7_0729500 | MT-A70 | 1739.93874 | 0.389479837 | 0.41078941 | 0.94812531 | 0.34306566 | 0.8930363 | 1554.31312 | 2087.78118 | 1458.72468 | 1858.93598 | Non-sig. |
| PF3D7_0528100 | N/A | 4428.82288 | 0.389007495 | 0.47786877 | 0.8140467 | 0.4156182 | 0.90793155 | 2568.71502 | 4839.4983 | 5101.53612 | 5205.54207 | Non-sig. |
| PF3D7_1472600 | PDI-14 | 2650.15143 | 0.38890595 | 0.40716273 | 0.95516097 | 0.3394963 | 0.8930363 | 2193.42076 | 3114.0791 | 2396.70848 | 2896.39738 | Non-sig. |
| PF3D7_0701600 | MC-2TM | 104.364595 | 0.388358501 | 0.57878494 | 0.67098929 | 0.50222735 | 0.93170321 | 64.013613 | 141.755626 | 116.561155 | 95.1279847 | Non-sig. |
| PF3D7_1011800 | PREBP | 15762.8637 | 0.387920893 | 0.62619557 | 0.6194884 | 0.53559466 | 0.93526055 | 22358.7679 | 18413.0495 | 4954.36836 | 17325.2691 | Non-sig. |
| PF3D7_0916000 | MFS2 | 8125.16617 | 0.387740994 | 0.42194491 | 0.91893747 | 0.35812828 | 0.89552149 | 7290.64983 | 7743.916 | 6789.05766 | 10677.0412 | Non-sig. |
| PF3D7_0305200 | N/A | 6157.22724 | 0.387207323 | 0.5510814 | 0.70263181 | 0.48228523 | 0.92880667 | 7996.82207 | 6997.2373 | 2675.39001 | 6959.45959 | Non-sig. |
| PF3D7_0924600 | N/A | 3725.82926 | 0.386605641 | 0.55740401 | 0.69358245 | 0.4879441 | 0.93170321 | 4864.29048 | 3942.81362 | 1595.32269 | 4500.89025 | Non-sig. |
| PF3D7_0713600 | N/A | 594.579904 | 0.385856908 | 0.43224345 | 0.89268421 | 0.37202631 | 0.89654416 | 464.23469 | 711.37528 | 566.699353 | 636.010293 | Non-sig. |
| PF3D7_0404900 | P41 | 5994.15321 | 0.385828786 | 0.40888958 | 0.9436014 | 0.34537337 | 0.8930363 | 5281.0911 | 7426.2545 | 5113.68169 | 6155.58554 | Non-sig. |
| PF3D7_1474900 | CITH | 559.241787 | 0.385583089 | 0.54091695 | 0.71283233 | 0.47594952 | 0.92705167 | 269.659126 | 639.372245 | 699.828169 | 628.107607 | Non-sig. |
| PF3D7_1109400 | ENP1 | 1484.74209 | 0.385248003 | 0.41768824 | 0.92233386 | 0.35635446 | 0.89552149 | 1159.40668 | 1747.06274 | 1415.78753 | 1616.71143 | Non-sig. |
| PF3D7_1468800 | U2AF2 | 1672.93945 | 0.38458529 | 0.44669598 | 0.86095535 | 0.38926264 | 0.90444842 | 1784.98614 | 1938.63763 | 1117.86861 | 1850.2654 | Non-sig. |
| PF3D7_0526200 | ARFGAP2 | 2759.86019 | 0.384550125 | 0.41246969 | 0.93231123 | 0.3511757 | 0.8930363 | 2539.81492 | 3376.65265 | 2248.69124 | 2874.28196 | Non-sig. |
| PF3D7_0802500 | N/A | 2705.49629 | 0.384171852 | 0.40983151 | 0.93738973 | 0.34855812 | 0.8930363 | 2221.31086 | 2874.72467 | 2473.38137 | 3252.56825 | Non-sig. |
| PF3D7_0924500 | N/A | 2346.55988 | 0.384092559 | 0.40622126 | 0.94552551 | 0.34439066 | 0.8930363 | 1993.0506 | 2581.0531 | 2078.96803 | 2733.16778 | Non-sig. |
| PF3D7_0613800 | ApiAP2 | 38966.972 | 0.384031611 | 0.44654793 | 0.86000087 | 0.38978856 | 0.90444842 | 35988.9667 | 33347.4252 | 31633.2815 | 54898.2144 | Non-sig. |
| PF3D7_0731800 | GEXP08 | 319.983068 | 0.383855879 | 0.45334723 | 0.84671496 | 0.39715402 | 0.90645858 | 249.635609 | 332.563047 | 305.556696 | 392.176921 | Non-sig. |
| PF3D7_0936400 | REX4 | 111.286947 | 0.382715307 | 0.57369598 | 0.66710474 | 0.50470523 | 0.93170321 | 129.445757 | 137.349767 | 64.0586679 | 114.293597 | Non-sig. |
| PF3D7_1110500 | VPS35 | 3059.30981 | 0.382134903 | 0.48688012 | 0.78486447 | 0.43253304 | 0.91370017 | 1722.26049 | 3414.62087 | 3590.32418 | 3510.03371 | Non-sig. |
| PF3D7_0409000 | N/A | 541.433212 | 0.382060586 | 0.45528253 | 0.83917251 | 0.40137251 | 0.90667178 | 437.461721 | 492.301309 | 502.757344 | 733.212475 | Non-sig. |
| PF3D7_0623100 | NAB2 | 3404.38898 | 0.381878931 | 0.45962892 | 0.83084183 | 0.40606299 | 0.90723586 | 3771.8709 | 4063.72086 | 2141.35615 | 3640.60801 | Non-sig. |
| PF3D7_0321500 | N/A | 2951.55281 | 0.381858645 | 0.49720901 | 0.76800427 | 0.44248465 | 0.91860646 | 1598.94403 | 3426.46483 | 3527.04891 | 3253.75347 | Non-sig. |
| PF3D7_0730000 | N/A | 175.555487 | 0.381632257 | 0.49413556 | 0.772323 | 0.43992314 | 0.91706217 | 124.955579 | 191.122435 | 179.78627 | 206.357665 | Non-sig. |
| PF3D7_1329300 | CAF1B | 1763.98017 | 0.381606522 | 0.57526393 | 0.66335903 | 0.50710064 | 0.93170321 | 732.742984 | 1681.28763 | 2330.89979 | 2310.99028 | Non-sig. |
| PF3D7_0400700 | RIF | 2.39646614 | 0.379607147 | 2.71074071 | 0.14003816 | 0.88862984 | 0.99457954 | 4.20289414 | 5.38297043 | 0 | 0 | Non-sig. |
| PF3D7_1430200 | PMIX | 4178.63407 | 0.379114953 | 0.42120802 | 0.90006584 | 0.36808522 | 0.89615989 | 4174.27066 | 4901.17859 | 3091.4338 | 4547.65325 | Non-sig. |

| | | | | | | | | | | | | |
|---|---|---|---|---|---|---|---|---|---|---|---|---|
| PF3D7_1457500 | VPS4 | 1274.09869 | 0.37833096 | 0.46468416 | 0.81416798 | 0.41554873 | 0.90793155 | 804.795928 | 1522.8071 | 1410.84321 | 1357.94851 | Non-sig. |
| PF3D7_0931000 | N/A | 743.458808 | 0.377717846 | 0.44667393 | 0.84562323 | 0.39776296 | 0.90645858 | 775.338391 | 852.832733 | 518.307943 | 827.356167 | Non-sig. |
| PF3D7_0521800 | N/A | 391.296996 | 0.37740418 | 0.44622719 | 0.84576688 | 0.39768281 | 0.90645858 | 322.740828 | 494.823403 | 358.004075 | 389.619676 | Non-sig. |
| PF3D7_1405800 | BOP1 | 1731.64587 | 0.377243696 | 0.41061955 | 0.9187183 | 0.35824294 | 0.89552149 | 1598.75062 | 1994.36914 | 1414.40384 | 1919.0599 | Non-sig. |
| PF3D7_0220500 | N/A | 1363.16862 | 0.376624579 | 0.50910299 | 0.73978073 | 0.45943305 | 0.92288312 | 1628.90967 | 1756.95822 | 743.964962 | 1322.84161 | Non-sig. |
| PF3D7_1451200 | N/A | 12611.6509 | 0.376184151 | 0.41452693 | 0.90750231 | 0.36414123 | 0.89567376 | 11428.4026 | 12449.2374 | 10524.9775 | 16043.9861 | Non-sig. |
| PF3D7_1118500 | NOP56 | 2349.52379 | 0.376180025 | 0.45846941 | 0.82051281 | 0.41192383 | 0.90748263 | 2600.34929 | 2734.92249 | 1489.8658 | 2572.9576 | Non-sig. |
| PF3D7_1406900 | RlmN | 450.293237 | 0.37567189 | 0.44291523 | 0.84818011 | 0.39633767 | 0.90645858 | 442.586841 | 529.170935 | 341.581989 | 487.833182 | Non-sig. |
| PF3D7_1113100 | PRL | 8030.77416 | 0.375429078 | 0.42883135 | 0.87547023 | 0.3813181 | 0.90292925 | 7941.06501 | 7856.155 | 6042.5729 | 10283.3037 | Non-sig. |
| PF3D7_0210600 | CERLI1 | 6398.02112 | 0.3743034 | 0.43015297 | 0.87016348 | 0.38421107 | 0.90292925 | 6586.97426 | 7697.57823 | 4558.63976 | 6748.89221 | Non-sig. |
| PF3D7_1410800 | N/A | 1233.40399 | 0.374040348 | 0.42181919 | 0.88673146 | 0.3752235 | 0.89654416 | 958.841463 | 1466.18836 | 1189.80628 | 1318.77984 | Non-sig. |
| PF3D7_0101300 | MC-2TM | 78.6793941 | 0.373859761 | 0.5797336 | 0.64488199 | 0.51900366 | 0.93247207 | 63.9851968 | 109.552082 | 73.0502103 | 68.1300876 | Non-sig. |
| PF3D7_0723400 | N/A | 2417.08461 | 0.373680965 | 0.57064158 | 0.65484356 | 0.51256848 | 0.93247207 | 3225.47413 | 2626.95505 | 986.556261 | 2829.35299 | Non-sig. |
| PF3D7_1456100 | SHMT | 32.3624116 | 0.372912266 | 0.77031727 | 0.48410217 | 0.62831336 | 0.96655174 | 16.8139021 | 33.2780541 | 39.4106324 | 39.9470576 | Non-sig. |
| PF3D7_0828900 | N/A | 1781.09 | 0.372471777 | 0.43501098 | 0.85623534 | 0.39186762 | 0.90601068 | 1678.75263 | 2360.39137 | 1426.24908 | 1658.96693 | Non-sig. |
| PF3D7_0711100 | N/A | 180.222314 | 0.371895459 | 0.4826546 | 0.7705209 | 0.44099096 | 0.91706217 | 144.149608 | 182.42827 | 169.995701 | 224.315676 | Non-sig. |
| PF3D7_1140800 | N/A | 1416.69399 | 0.370796216 | 0.44824378 | 0.82722 | 0.40811238 | 0.90723586 | 949.817356 | 1572.82841 | 1521.13622 | 1622.99398 | Non-sig. |
| PF3D7_0927700 | PPP4 | 1091.36352 | 0.36979003 | 0.539309 | 0.68567376 | 0.49291885 | 0.93170321 | 515.778928 | 1264.70218 | 1388.31024 | 1196.66272 | Non-sig. |
| PF3D7_0625800 | N/A | 276.644129 | 0.369748934 | 0.45071217 | 0.82036598 | 0.4120075 | 0.90748263 | 227.288446 | 319.779666 | 255.407328 | 304.101074 | Non-sig. |
| PF3D7_1302100 | G27/25 | 1972.78687 | 0.369202654 | 0.46622397 | 0.79189978 | 0.42841912 | 0.91236574 | 1233.15616 | 2060.36959 | 2209.94194 | 2387.67978 | Non-sig. |
| PF3D7_0924100 | N/A | 646.672532 | 0.369001992 | 0.46031959 | 0.80162131 | 0.42277205 | 0.91161836 | 698.800564 | 715.719911 | 430.343272 | 741.826382 | Non-sig. |
| PF3D7_1232000 | aFRS | 531.246969 | 0.368978642 | 0.4531194 | 0.81430776 | 0.41546867 | 0.90793155 | 388.386986 | 668.482995 | 538.761879 | 529.356014 | Non-sig. |
| PF3D7_1010300 | SDH4 | 10355.7233 | 0.368556362 | 0.41308247 | 0.89221013 | 0.37228031 | 0.89654416 | 8642.38562 | 13029.3425 | 9437.78367 | 10313.3814 | Non-sig. |
| PF3D7_0914400 | KIC3 | 1944.68199 | 0.368160563 | 0.40746354 | 0.90354235 | 0.36623812 | 0.89567376 | 1707.26686 | 2114.24098 | 1688.51863 | 2268.70151 | Non-sig. |
| PF3D7_0806400 | ALG13 | 461.863497 | 0.366738108 | 0.44110126 | 0.83141479 | 0.40573935 | 0.90723586 | 430.269196 | 577.756379 | 376.773748 | 462.654667 | Non-sig. |
| PF3D7_1113900 | MAPK2 | 127.840556 | 0.366053356 | 0.57372447 | 0.63802989 | 0.52345423 | 0.93247207 | 152.285576 | 125.618755 | 71.4243186 | 162.033575 | Non-sig. |
| PF3D7_1112700 | N/A | 177.176915 | 0.365270905 | 0.48026156 | 0.76056661 | 0.44691597 | 0.91974025 | 149.528111 | 179.169294 | 160.162891 | 219.847366 | Non-sig. |
| PF3D7_1112200 | COQ4 | 113.921595 | 0.364690044 | 0.53614972 | 0.68020188 | 0.49637664 | 0.93170321 | 77.5397392 | 123.551047 | 121.454139 | 133.141455 | Non-sig. |
| PF3D7_0417700 | N/A | 166.469946 | 0.364542747 | 0.51008835 | 0.7146659 | 0.47481552 | 0.92682482 | 172.248658 | 215.803617 | 119.041555 | 158.785955 | Non-sig. |
| PF3D7_0807400 | COQ10 | 537.859574 | 0.364286107 | 0.44119395 | 0.82568247 | 0.40898424 | 0.90748263 | 524.496105 | 663.374118 | 416.290488 | 547.277585 | Non-sig. |
| PF3D7_0810600 | DBP1 | 5608.87666 | 0.364084424 | 0.48347927 | 0.75305074 | 0.45141942 | 0.92081798 | 3232.57578 | 5858.49499 | 6576.77016 | 6767.6657 | Non-sig. |
| PF3D7_1207100 | N/A | 3483.16159 | 0.363977642 | 0.53982931 | 0.6742458 | 0.50015506 | 0.93170321 | 4479.44198 | 4136.7347 | 1613.18872 | 3703.28094 | Non-sig. |
| PF3D7_1136800 | N/A | 888.420778 | 0.363938499 | 0.43246478 | 0.84154484 | 0.40004278 | 0.90645858 | 669.894731 | 934.400178 | 883.811428 | 1065.57677 | Non-sig. |
| PF3D7_0311000 | N/A | 6.730108 | 0.363113894 | 1.3319445 | 0.27261939 | 0.78514581 | 0.99028237 | 5.200811 | 7.53571787 | 6.55774913 | 7.62615399 | Non-sig. |
| PF3D7_0908500 | N/A | 6217.83737 | 0.362942551 | 0.4015904 | 0.90376301 | 0.36612108 | 0.89567376 | 5519.53767 | 6991.21881 | 5360.12623 | 7000.46676 | Non-sig. |
| PF3D7_1475400 | CRMP4 | 420.28363 | 0.362717895 | 0.49314129 | 0.7355253 | 0.46201964 | 0.92403928 | 304.763652 | 349.612586 | 430.495316 | 596.262967 | Non-sig. |
| PF3D7_0422800 | SR12 | 309.628839 | 0.362276496 | 0.49726691 | 0.7285353 | 0.46628597 | 0.92538923 | 191.066197 | 352.202569 | 350.528502 | 344.71809 | Non-sig. |
| PF3D7_1466300 | RPN2 | 11925.8068 | 0.361694902 | 0.50077298 | 0.7222732 | 0.47012653 | 0.92682482 | 6404.63519 | 12474.0688 | 14472.2418 | 14352.2815 | Non-sig. |
| PF3D7_0817600 | N/A | 18693.6626 | 0.361455504 | 0.55148508 | 0.65542209 | 0.51219603 | 0.93247207 | 24568.9027 | 21377.2852 | 8159.69741 | 20668.7651 | Non-sig. |
| PF3D7_0108100 | N/A | 251.673604 | 0.361166992 | 0.45329478 | 0.79675966 | 0.42559063 | 0.91236574 | 227.200781 | 277.959459 | 213.509395 | 288.024782 | Non-sig. |
| PF3D7_1445400 | CLK1 | 4161.19803 | 0.36087341 | 0.40785334 | 0.88481171 | 0.37625819 | 0.89712687 | 3933.42066 | 4626.07977 | 3353.62556 | 4731.66613 | Non-sig. |
| PF3D7_0619700 | N/A | 1436.50187 | 0.360717817 | 0.51526951 | 0.70005659 | 0.48389197 | 0.93078439 | 1759.01966 | 1788.16529 | 757.081769 | 1441.74075 | Non-sig. |
| PF3D7_1312600 | BCKDHA | 2703.75386 | 0.360507824 | 0.42201939 | 0.85424469 | 0.39296942 | 0.90601068 | 2095.60033 | 3332.7603 | 2639.60352 | 2747.0513 | Non-sig. |
| PF3D7_0319400 | N/A | 4264.31237 | 0.360046503 | 0.40428534 | 0.89057522 | 0.3731571 | 0.89654416 | 3910.61302 | 4794.7362 | 3559.35789 | 4792.54236 | Non-sig. |
| PF3D7_1124600 | EK | 8835.59998 | 0.359140169 | 0.41528265 | 0.86480899 | 0.38714364 | 0.90421052 | 7201.61147 | 8790.35855 | 8281.26326 | 11069.1666 | Non-sig. |
| PF3D7_0311800 | N/A | 2061.64467 | 0.359107885 | 0.41113141 | 0.87346253 | 0.38241102 | 0.90292925 | 1729.62455 | 2472.50825 | 1883.05995 | 2161.38595 | Non-sig. |
| PF3D7_1436600 | PKG | 3961.59779 | 0.358897261 | 0.4501131 | 0.79734907 | 0.42524833 | 0.91236574 | 2715.12015 | 3850.77202 | 4227.31447 | 5053.1845 | Non-sig. |
| PF3D7_0617300 | N/A | 1059.69315 | 0.35858039 | 0.41268631 | 0.86889336 | 0.38490546 | 0.90292925 | 914.467819 | 1201.23831 | 942.857572 | 1180.20888 | Non-sig. |
| PF3D7_0519500 | CCR4-1 | 2356.49787 | 0.357853029 | 0.58140963 | 0.61549209 | 0.53822979 | 0.93646681 | 3158.39455 | 2145.05836 | 973.488947 | 3149.04964 | Non-sig. |
| PF3D7_0207800 | SERA3 | 8004.00385 | 0.357841215 | 0.4310792 | 0.83010549 | 0.40647914 | 0.90723586 | 5633.48761 | 9151.64618 | 8399.06933 | 8831.81225 | Non-sig. |
| PF3D7_1349600 | N/A | 1407.59482 | 0.357710929 | 0.50623955 | 0.70660408 | 0.47981255 | 0.92880667 | 1698.12993 | 1751.64532 | 770.227198 | 1410.37683 | Non-sig. |
| PF3D7_1344300 | N/A | 10726.4055 | 0.357391958 | 0.45726532 | 0.78158553 | 0.43445819 | 0.91522571 | 7259.4555 | 10022.0417 | 11549.3702 | 14074.7546 | Non-sig. |
| PF3D7_0418600 | N/A | 9202.1786 | 0.357074573 | 0.49909691 | 0.71544137 | 0.47433636 | 0.92682482 | 10906.7047 | 8545.32462 | 5232.01175 | 12124.6733 | Non-sig. |
| PF3D7_0523400 | N/A | 3317.11874 | 0.356767138 | 0.41590877 | 0.85780142 | 0.39100212 | 0.90538866 | 2562.83348 | 3586.85468 | 3255.07732 | 3863.70947 | Non-sig. |

| | | | | | | | | | | | | |
|---|---|---|---|---|---|---|---|---|---|---|---|---|
| PF3D7_0802800 | CNA | 8670.00909 | 0.35662036 | 0.40971667 | 0.87040724 | 0.38407789 | 0.90292925 | 8144.06014 | 10485.5956 | 7063.78404 | 8986.59655 | Non-sig. |
| PF3D7_1453100 | N/A | 432.803959 | 0.356565504 | 0.44250233 | 0.8057935 | 0.42036193 | 0.90969969 | 426.803946 | 498.047747 | 332.547892 | 473.816252 | Non-sig. |
| PF3D7_0104000 | TRSP | 36.3171612 | 0.354892951 | 0.74792356 | 0.47450431 | 0.63514033 | 0.9668494 | 43.4420407 | 49.601315 | 20.4710357 | 31.7542532 | Non-sig. |
| PF3D7_0411800 | N/A | 1920.13017 | 0.354310504 | 0.40996932 | 0.86423662 | 0.38745792 | 0.90421052 | 1793.91192 | 2181.64182 | 1577.24609 | 2127.72084 | Non-sig. |
| PF3D7_1346000 | N/A | 149.810136 | 0.352736564 | 0.52392962 | 0.67325181 | 0.50078711 | 0.93170321 | 164.052053 | 152.404528 | 99.3600196 | 183.423943 | Non-sig. |
| PF3D7_1206400 | TUM1 | 87.8707346 | 0.350970744 | 0.53817922 | 0.65214474 | 0.51430781 | 0.93247207 | 75.635145 | 101.991515 | 78.8162454 | 95.0400333 | Non-sig. |
| PF3D7_0827100 | IF2c | 1305.7111 | 0.350399327 | 0.43222555 | 0.81068628 | 0.41754585 | 0.90869129 | 956.673469 | 1405.23803 | 1338.92834 | 1522.00456 | Non-sig. |
| PF3D7_1238900 | PK2 | 2725.57639 | 0.349713204 | 0.50531613 | 0.69206815 | 0.48889453 | 0.93170321 | 3303.94293 | 2675.1126 | 1490.14115 | 3433.10888 | Non-sig. |
| PF3D7_1474700 | N/A | 125.873827 | 0.349063406 | 0.51066344 | 0.68354885 | 0.49426009 | 0.93170321 | 118.86364 | 124.490326 | 102.649348 | 157.491993 | Non-sig. |
| PF3D7_0620400 | MSP10 | 1728.00207 | 0.349037153 | 0.42447187 | 0.82228572 | 0.4109143 | 0.90748263 | 1382.76066 | 1707.61425 | 1657.07126 | 2164.56211 | Non-sig. |
| PF3D7_0905500 | N/A | 1956.8584 | 0.348618919 | 0.56597864 | 0.61595773 | 0.53792242 | 0.93646681 | 824.3403 | 2123.33323 | 2618.36347 | 2261.39658 | Non-sig. |
| PF3D7_1017300 | GRASP | 2087.96396 | 0.348526837 | 0.48457429 | 0.71924335 | 0.47199099 | 0.92682482 | 2457.86021 | 2470.80536 | 1216.59793 | 2206.59234 | Non-sig. |
| PF3D7_0606100 | N/A | 1778.07777 | 0.348085671 | 0.4433401 | 0.78514367 | 0.43236934 | 0.91370017 | 1909.11713 | 2038.76898 | 1220.40593 | 1944.01905 | Non-sig. |
| PF3D7_0603700 | mFRS | 225.103159 | 0.34797986 | 0.54673541 | 0.63646849 | 0.52447112 | 0.93247207 | 119.903752 | 262.939415 | 275.916701 | 241.652769 | Non-sig. |
| PF3D7_1136500 | CK1 | 13460.9885 | 0.34720612 | 0.41098934 | 0.84480566 | 0.39821935 | 0.90645858 | 10863.4113 | 16271.2364 | 12834.4079 | 13874.8985 | Non-sig. |
| PF3D7_0618800 | N/A | 336.891062 | 0.347135365 | 0.49157845 | 0.70616474 | 0.4800857 | 0.92880667 | 211.678667 | 381.930211 | 381.112671 | 372.842698 | Non-sig. |
| PF3D7_1033900 | N/A | 1884.23809 | 0.34705248 | 0.46325061 | 0.74916789 | 0.45375602 | 0.92203286 | 2128.67976 | 2103.55358 | 1189.05612 | 2115.66292 | Non-sig. |
| PF3D7_0513100 | N/A | 650.821346 | 0.346903667 | 0.45076679 | 0.76958569 | 0.4415457 | 0.91773602 | 643.770208 | 607.98012 | 502.284533 | 849.250522 | Non-sig. |
| PF3D7_1321500 | PDEbeta | 8603.93521 | 0.346668678 | 0.4055712 | 0.85476651 | 0.39268042 | 0.90601068 | 7198.18629 | 9078.65648 | 7952.03731 | 10186.8608 | Non-sig. |
| PF3D7_0820900 | N/A | 1811.6004 | 0.346639052 | 0.48511666 | 0.71454782 | 0.4748885 | 0.92682482 | 2135.6803 | 1939.43722 | 1054.71605 | 2116.56802 | Non-sig. |
| PF3D7_1454300 | KIN | 121.201935 | 0.345949744 | 0.56308546 | 0.6143823 | 0.53896273 | 0.93711173 | 138.990302 | 113.725986 | 74.7419634 | 157.349487 | Non-sig. |
| PF3D7_0826400 | CPT | 809.596588 | 0.345653342 | 0.42506184 | 0.81318365 | 0.41611277 | 0.90793155 | 745.174612 | 993.034011 | 681.041211 | 819.136516 | Non-sig. |
| PF3D7_0510200 | CYP87 | 1708.17139 | 0.345501808 | 0.42389826 | 0.81505832 | 0.41503893 | 0.90793155 | 1318.63276 | 2058.72412 | 1690.39226 | 1764.93641 | Non-sig. |
| PF3D7_1317100 | MCM4 | 11025.6607 | 0.344455189 | 0.49255541 | 0.69932271 | 0.48435038 | 0.93109154 | 6116.88657 | 12802.4351 | 13314.0029 | 11869.3182 | Non-sig. |
| PF3D7_0933800 | N/A | 146.235339 | 0.344059754 | 0.48904249 | 0.70353754 | 0.48172082 | 0.92880667 | 134.631726 | 157.762676 | 123.177572 | 169.36938 | Non-sig. |
| PF3D7_1228400 | N/A | 1405.53833 | 0.343964132 | 0.45235437 | 0.76038645 | 0.44702362 | 0.91974025 | 1537.32063 | 1502.82332 | 940.547354 | 1641.46201 | Non-sig. |
| PF3D7_0918900 | gammaGCS | 1541.45779 | 0.343003128 | 0.49581909 | 0.69179089 | 0.48906866 | 0.93170321 | 907.30178 | 1447.00699 | 1810.47962 | 2001.04275 | Non-sig. |
| PF3D7_0809100 | VAR | 857.696678 | 0.342832379 | 0.41729215 | 0.82156441 | 0.41132485 | 0.90748263 | 783.111671 | 990.002026 | 729.474366 | 928.198647 | Non-sig. |
| PF3D7_1373500 | VAR | 887.958689 | 0.342582077 | 0.43532849 | 0.78695073 | 0.43131071 | 0.91355742 | 873.713963 | 1110.14678 | 692.502171 | 875.471838 | Non-sig. |
| PF3D7_0606300 | N/A | 55.3873172 | 0.34253812 | 0.6071373 | 0.5641856 | 0.57262783 | 0.94850402 | 45.1021028 | 65.517344 | 52.5300324 | 58.3997895 | Non-sig. |
| PF3D7_1028500 | N/A | 1500.08948 | 0.342338448 | 0.42915007 | 0.79771267 | 0.42503725 | 0.91236574 | 1441.15423 | 1446.34826 | 1204.85599 | 1907.99943 | Non-sig. |
| PF3D7_0210800 | N/A | 244.929171 | 0.342265329 | 0.45734591 | 0.748373 | 0.45423521 | 0.92203286 | 213.563171 | 254.336776 | 218.443074 | 293.373663 | Non-sig. |
| PF3D7_0207600 | SERA5 | 104931.979 | 0.341241403 | 0.5399219 | 0.63201994 | 0.52737385 | 0.93270419 | 48623.639 | 109038.571 | 136535.5 | 125530.206 | Non-sig. |
| PF3D7_1324800 | DHFS-FPGS | 1764.57941 | 0.340584825 | 0.41423818 | 0.82219564 | 0.41096556 | 0.90748263 | 1439.98811 | 2083.08697 | 1674.40596 | 1860.83661 | Non-sig. |
| PF3D7_0105900 | N/A | 666.118648 | 0.340421055 | 0.43252338 | 0.78705815 | 0.43124783 | 0.91355742 | 517.779588 | 708.211059 | 657.83248 | 780.651463 | Non-sig. |
| PF3D7_1335100 | MSP7 | 28524.4295 | 0.340194044 | 0.41094609 | 0.82783132 | 0.40776603 | 0.90723586 | 22226.0595 | 32153.6687 | 28127.4403 | 31590.5497 | Non-sig. |
| PF3D7_0204400 | N/A | 22.9150302 | 0.33979129 | 0.90081338 | 0.37720498 | 0.70602128 | 0.98615891 | 9.24977202 | 21.623841 | 31.0427889 | 29.7437188 | Non-sig. |
| PF3D7_1305200 | ApiAP2 | 181.79739 | 0.338990635 | 0.48184968 | 0.70351948 | 0.48173207 | 0.92880667 | 139.526426 | 204.060335 | 181.39841 | 202.20439 | Non-sig. |
| PF3D7_1110200 | PRPF6 | 4612.05858 | 0.338562826 | 0.40814924 | 0.82950743 | 0.40681733 | 0.90723586 | 4358.46274 | 5400.081 | 3788.38222 | 4901.30835 | Non-sig. |
| PF3D7_0611600 | BTP1 | 4950.58716 | 0.337546566 | 0.56417457 | 0.59830164 | 0.54963869 | 0.94142878 | 6662.5339 | 5705.18464 | 2086.05755 | 5348.57257 | Non-sig. |
| PF3D7_1439800 | N/A | 4313.62602 | 0.336373379 | 0.49998535 | 0.67276647 | 0.50109587 | 0.93170321 | 5283.7395 | 4922.18712 | 2342.71253 | 4705.86492 | Non-sig. |
| PF3D7_0621100 | N/A | 10480.272 | 0.336220958 | 0.4467517 | 0.75259021 | 0.4516962 | 0.92081798 | 6951.59173 | 11494.0331 | 11577.1847 | 11898.2784 | Non-sig. |
| PF3D7_1455700 | N/A | 1628.87307 | 0.335987198 | 0.43772798 | 0.76757076 | 0.44274224 | 0.9187812 | 1718.09497 | 1722.85203 | 1162.28173 | 1912.26353 | Non-sig. |
| PF3D7_0308300 | PIC4 | 3250.43846 | 0.335749993 | 0.41688765 | 0.80537284 | 0.42060456 | 0.90969969 | 3192.82473 | 3385.68871 | 2555.16273 | 3868.07768 | Non-sig. |
| PF3D7_1031100 | N/A | 197.041402 | 0.335079201 | 0.49249857 | 0.68036584 | 0.49627285 | 0.93170321 | 152.190545 | 183.628814 | 196.190063 | 256.156187 | Non-sig. |
| PF3D7_0215900 | DHHC11 | 103.989901 | 0.334775739 | 0.55499209 | 0.60320812 | 0.54637024 | 0.94142878 | 68.9172873 | 126.896805 | 114.832502 | 105.313009 | Non-sig. |
| PF3D7_0924200 | N/A | 662.398743 | 0.334684291 | 0.4712168 | 0.71025543 | 0.47754575 | 0.92879493 | 425.634537 | 726.228971 | 745.850315 | 751.88115 | Non-sig. |
| PF3D7_1325400 | N/A | 11871.1474 | 0.334241937 | 0.51504882 | 0.64895195 | 0.51636944 | 0.93247207 | 14798.8338 | 11626.8396 | 6205.83197 | 14853.084 | Non-sig. |
| PF3D7_0323300 | N/A | 144.809854 | 0.334040136 | 0.52007395 | 0.64229353 | 0.52068261 | 0.93247207 | 97.5858537 | 164.311753 | 158.422186 | 158.919624 | Non-sig. |
| PF3D7_1004600 | N/A | 1391.66352 | 0.33403645 | 0.48122414 | 0.69413902 | 0.48759503 | 0.93170321 | 1625.95009 | 1655.41118 | 836.96232 | 1448.33049 | Non-sig. |
| PF3D7_0502300 | N/A | 30.4973638 | 0.334002627 | 1.11161928 | 0.30046495 | 0.76382253 | 0.99028237 | 52.5273214 | 36.5016573 | 1.64202113 | 31.3184553 | Non-sig. |
| PF3D7_0935300 | N/A | 216.486906 | 0.333655244 | 0.48072128 | 0.69407213 | 0.48763698 | 0.93170321 | 165.518711 | 269.68144 | 217.459453 | 213.288021 | Non-sig. |
| PF3D7_0928400 | N/A | 107.337225 | 0.333605747 | 0.59847023 | 0.55743081 | 0.57723312 | 0.9497432 | 57.5672718 | 104.244873 | 132.107424 | 135.429332 | Non-sig. |
| PF3D7_1435200 | N/A | 1505.85078 | 0.333216678 | 0.62609282 | 0.5322161 | 0.59457633 | 0.9542761 | 2153.42633 | 1948.86503 | 512.434139 | 1408.67763 | Non-sig. |

| | | | | | | | | | | | | |
|---|---|---|---|---|---|---|---|---|---|---|---|---|
| PF3D7_0616600 | N/A | 1407.66946 | 0.333038916 | 0.53581517 | 0.62155559 | 0.53423413 | 0.93470782 | 1806.58248 | 1695.79332 | 685.660043 | 1442.64201 | Non-sig. |
| PF3D7_1121600 | EXP1 | 14539.1623 | 0.332946296 | 0.40217192 | 0.82787056 | 0.40774381 | 0.90723586 | 13340.1871 | 16760.5813 | 12397.6635 | 15658.2171 | Non-sig. |
| PF3D7_1415100 | N/A | 753.969923 | 0.332887707 | 0.42791981 | 0.77792076 | 0.43661573 | 0.91522571 | 727.143582 | 902.454609 | 607.72615 | 778.555351 | Non-sig. |
| PF3D7_1448200 | CAPG2 | 63.2452792 | 0.332363024 | 0.59554755 | 0.55807974 | 0.57678994 | 0.9497432 | 65.2910055 | 66.5167704 | 46.8173561 | 74.3559847 | Non-sig. |
| PF3D7_1112000 | N/A | 1119.56286 | 0.332361236 | 0.42176117 | 0.78803185 | 0.43067808 | 0.91355742 | 1056.41067 | 1363.60334 | 926.032231 | 1132.2052 | Non-sig. |
| PF3D7_1418100 | LISP1 | 46.706066 | 0.332163442 | 0.6675999 | 0.49754867 | 0.61880219 | 0.96395246 | 31.6293682 | 42.8999235 | 50.9345104 | 61.3604619 | Non-sig. |
| PF3D7_1462400 | N/A | 1257.59783 | 0.332162468 | 0.46222594 | 0.71861494 | 0.4723782 | 0.92682482 | 1297.97032 | 1096.06642 | 929.15798 | 1707.19661 | Non-sig. |
| PF3D7_0202400 | PTEF | 2740.49036 | 0.332096793 | 0.63289111 | 0.52472975 | 0.59977107 | 0.95655566 | 3975.2018 | 3403.43767 | 878.145593 | 2705.17639 | Non-sig. |
| PF3D7_1133200 | N/A | 16501.6044 | 0.331551091 | 0.6179228 | 0.53655747 | 0.59157333 | 0.95321031 | 23729.8049 | 18755.2558 | 5498.18118 | 18023.1758 | Non-sig. |
| PF3D7_1473300 | N/A | 186.432977 | 0.33140293 | 0.52268486 | 0.63403965 | 0.52605494 | 0.93247207 | 214.719459 | 197.432805 | 115.801089 | 217.778553 | Non-sig. |
| PF3D7_1038800 | N/A | 415.721911 | 0.3313176 | 0.43542392 | 0.76090814 | 0.44671194 | 0.91974025 | 363.452242 | 493.458236 | 372.921081 | 433.056084 | Non-sig. |
| PF3D7_1142200 | N/A | 48.8009393 | 0.331132171 | 0.64382063 | 0.51432364 | 0.60702573 | 0.95715771 | 38.7877328 | 64.4530946 | 47.6007749 | 44.3621547 | Non-sig. |
| PF3D7_1341100 | N/A | 5.90286893 | 0.331043569 | 1.44319179 | 0.22938294 | 0.8185713 | 0.99096399 | 3.02378764 | 7.71929989 | 7.38973823 | 5.47864996 | Non-sig. |
| PF3D7_1250100 | G377 | 70.4024862 | 0.330664153 | 0.60442768 | 0.54706983 | 0.58433076 | 0.95151408 | 54.8194405 | 57.9163247 | 69.828508 | 99.0456714 | Non-sig. |
| PF3D7_0828700 | N/A | 1486.43445 | 0.330602867 | 0.63333097 | 0.52200647 | 0.60166582 | 0.95693578 | 2160.21999 | 1613.49422 | 473.936512 | 1698.08709 | Non-sig. |
| PF3D7_1105700 | RTCB | 944.075365 | 0.330339356 | 0.52700356 | 0.62682566 | 0.53077352 | 0.93351492 | 480.885965 | 1117.081 | 1191.62822 | 986.706276 | Non-sig. |
| PF3D7_0305300 | N/A | 2125.05432 | 0.330124152 | 0.4248577 | 0.77702288 | 0.43714527 | 0.91524921 | 1928.55233 | 2001.83459 | 1837.43598 | 2732.3944 | Non-sig. |
| PF3D7_0717300 | N/A | 1056.93743 | 0.329337361 | 0.44675837 | 0.73717111 | 0.4610183 | 0.92336312 | 1124.07914 | 1075.16125 | 749.827878 | 1278.68146 | Non-sig. |
| PF3D7_1327300 | N/A | 19932.5155 | 0.329310829 | 0.89751892 | 0.36691241 | 0.71368435 | 0.9875928 | 34021.6995 | 23517.4217 | 1313.55438 | 20877.3866 | Non-sig. |
| PF3D7_1356000 | N/A | 158.462888 | 0.328957415 | 0.6120478 | 0.53747013 | 0.59094291 | 0.95315309 | 210.668418 | 195.245268 | 70.6397704 | 157.298094 | Non-sig. |
| PF3D7_0811900 | N/A | 2371.13819 | 0.328801415 | 0.42995411 | 0.76473606 | 0.44442873 | 0.91974025 | 2474.57586 | 2719.40249 | 1729.88653 | 2560.6879 | Non-sig. |
| PF3D7_1139200 | N/A | 168.155868 | 0.328305003 | 0.53544156 | 0.61314815 | 0.53977838 | 0.93711173 | 100.86432 | 182.502258 | 197.04661 | 192.210286 | Non-sig. |
| PF3D7_1429200 | ApiAP2 | 195.251013 | 0.328289991 | 0.57711014 | 0.56885154 | 0.56945689 | 0.94850402 | 249.68988 | 192.016588 | 96.9275713 | 242.370011 | Non-sig. |
| PF3D7_0826500 | UBE4B | 3979.30488 | 0.328270419 | 0.49493721 | 0.6632567 | 0.50716617 | 0.93170321 | 2225.6558 | 4166.13696 | 4830.98762 | 4694.43915 | Non-sig. |
| PF3D7_1475500 | CCp1 | 700.7032 | 0.328161265 | 0.43980762 | 0.74614728 | 0.45557845 | 0.92229829 | 569.007837 | 893.53275 | 673.583905 | 666.688307 | Non-sig. |
| PF3D7_0924300 | TPK | 852.139626 | 0.328022322 | 0.44288485 | 0.74064922 | 0.45890616 | 0.92288312 | 895.036959 | 1003.80458 | 616.575472 | 893.141489 | Non-sig. |
| PF3D7_1348100 | N/A | 536.05023 | 0.327934229 | 0.46857137 | 0.69985973 | 0.48401491 | 0.93078439 | 567.792033 | 695.154947 | 383.220999 | 498.032942 | Non-sig. |
| PF3D7_1420900 | N/A | 717.06197 | 0.327725806 | 0.41934033 | 0.78152705 | 0.43449258 | 0.91522571 | 619.122864 | 786.807428 | 652.768422 | 809.549167 | Non-sig. |
| PF3D7_0930000 | N/A | 67.2290998 | 0.32755681 | 0.58894943 | 0.55617137 | 0.57809371 | 0.95057122 | 65.144172 | 86.9559712 | 54.1882024 | 62.6280536 | Non-sig. |
| PF3D7_1209600 | PBGD | 287.305624 | 0.327420784 | 0.48744489 | 0.67170831 | 0.50176941 | 0.93170321 | 190.056438 | 312.453914 | 319.345782 | 327.366364 | Non-sig. |
| PF3D7_1449100 | N/A | 5398.71259 | 0.326875389 | 0.4184867 | 0.78108907 | 0.43475011 | 0.91522571 | 5351.91902 | 5495.86489 | 4227.69267 | 6519.37379 | Non-sig. |
| PF3D7_0806800 | N/A | 18521.9544 | 0.326792597 | 0.4304001 | 0.75927631 | 0.44768729 | 0.92039086 | 14774.8062 | 17066.5917 | 18091.3083 | 24155.1115 | Non-sig. |
| PF3D7_1456000 | AP2-HC | 3877.79546 | 0.32660917 | 0.46270989 | 0.70586165 | 0.48027418 | 0.92880667 | 4437.27781 | 4210.48604 | 2444.52233 | 4418.89566 | Non-sig. |
| PF3D7_1032700 | N/A | 1861.14165 | 0.326178168 | 0.65890981 | 0.49502703 | 0.62058104 | 0.9639908 | 2774.78892 | 2251.10002 | 528.916321 | 1889.76135 | Non-sig. |
| PF3D7_0304800 | N/A | 420.315336 | 0.325839914 | 0.43357956 | 0.75151124 | 0.45234504 | 0.92103152 | 355.038325 | 465.561964 | 390.994826 | 469.66623 | Non-sig. |
| PF3D7_0408200 | N/A | 583.460717 | 0.32521341 | 0.7461596 | 0.43584967 | 0.66294581 | 0.97883902 | 925.472519 | 693.322601 | 110.857284 | 604.190464 | Non-sig. |
| PF3D7_1035400 | MSP3 | 32206.5901 | 0.324804772 | 0.5520022 | 0.5884121 | 0.55625572 | 0.94243456 | 42734.7332 | 39646.7368 | 14458.4909 | 31986.3993 | Non-sig. |
| PF3D7_1339600 | N/A | 524.481049 | 0.324287133 | 0.44506786 | 0.72862402 | 0.46623168 | 0.92538923 | 534.311872 | 626.760413 | 397.45195 | 539.399962 | Non-sig. |
| PF3D7_1020500 | N/A | 267.941798 | 0.324172956 | 0.47542107 | 0.68186493 | 0.49532436 | 0.93170321 | 270.007295 | 253.360001 | 206.075915 | 342.323981 | Non-sig. |
| PF3D7_0815500 | N/A | 2730.17617 | 0.324006745 | 0.48292437 | 0.67092647 | 0.50226737 | 0.93170321 | 3214.92327 | 3358.3724 | 1635.21679 | 2712.19221 | Non-sig. |
| PF3D7_1125000 | N/A | 979.397377 | 0.32354591 | 0.43403981 | 0.74542911 | 0.45601235 | 0.92247173 | 740.679418 | 1168.03346 | 999.180338 | 1009.69629 | Non-sig. |
| PF3D7_1358000 | N/A | 1491.4959 | 0.323519457 | 0.41961136 | 0.77099786 | 0.4407082 | 0.91706217 | 1388.18747 | 1479.07744 | 1261.80738 | 1836.9113 | Non-sig. |
| PF3D7_1311300 | N/A | 430.887586 | 0.323502205 | 0.49172364 | 0.65789435 | 0.51060602 | 0.93247207 | 266.249377 | 500.59791 | 498.97975 | 457.723305 | Non-sig. |
| PF3D7_1228300 | NEK1 | 5716.39503 | 0.323357475 | 0.43132369 | 0.74968633 | 0.45344365 | 0.92203286 | 4081.17632 | 6311.52594 | 6075.43083 | 6397.44702 | Non-sig. |
| PF3D7_1455800 | CCp2 | 93.2392785 | 0.322685961 | 0.59530824 | 0.54204854 | 0.58778507 | 0.95246063 | 53.7316386 | 92.2533424 | 111.713461 | 115.258672 | Non-sig. |
| PF3D7_0630400 | N/A | 330.672649 | 0.322358787 | 0.49585009 | 0.65011339 | 0.51561898 | 0.93247207 | 218.774645 | 310.160037 | 368.710968 | 425.044947 | Non-sig. |
| PF3D7_0216700 | ATG11 | 13497.9008 | 0.322289236 | 0.84331347 | 0.38217015 | 0.70233516 | 0.98512564 | 22677.9829 | 16453.5177 | 1315.37703 | 13544.7257 | Non-sig. |
| PF3D7_1015400 | N/A | 2384.49748 | 0.32152405 | 0.40634937 | 0.79125028 | 0.42879796 | 0.91236574 | 2043.50009 | 2690.90688 | 2196.20959 | 2607.37336 | Non-sig. |
| PF3D7_0110600 | PIP5K | 5034.89781 | 0.321178683 | 0.48438252 | 0.66306828 | 0.50728682 | 0.93170321 | 5859.42048 | 4635.90618 | 3094.44747 | 6549.8171 | Non-sig. |
| PF3D7_0620700 | N/A | 1342.80352 | 0.320824313 | 0.41006641 | 0.7823716 | 0.43399622 | 0.91522571 | 1174.54571 | 1514.42536 | 1213.65057 | 1468.59243 | Non-sig. |
| PF3D7_0204700 | HT1 | 1906.60403 | 0.320419923 | 0.64191551 | 0.49916215 | 0.61766516 | 0.96395246 | 598.790446 | 1803.81223 | 2792.2783 | 2431.53515 | Non-sig. |
| PF3D7_1129300 | N/A | 4147.39471 | 0.319810276 | 0.55109768 | 0.58031505 | 0.56170218 | 0.94472957 | 5523.94193 | 4623.11793 | 1855.66205 | 4586.85695 | Non-sig. |
| PF3D7_1239000 | N/A | 2950.47782 | 0.319695771 | 0.46429047 | 0.68856846 | 0.49109487 | 0.93170321 | 1870.1044 | 3095.83531 | 3379.34248 | 3456.62909 | Non-sig. |
| PF3D7_1128900 | N/A | 7527.24993 | 0.319009498 | 0.41249528 | 0.7733652 | 0.43930628 | 0.91706217 | 5990.08162 | 8808.91372 | 7406.57813 | 7903.42628 | Non-sig. |

| | | | | | | | | | | | | |
|---|---|---|---|---|---|---|---|---|---|---|---|---|
| PF3D7_0317800 | N/A | 905.368705 | 0.318458617 | 0.54357615 | 0.58585834 | 0.55797072 | 0.94272305 | 1159.72166 | 1172.37051 | 452.371227 | 837.011421 | Non-sig. |
| PF3D7_0212100 | N/A | 3150.33669 | 0.318276049 | 0.42092494 | 0.75613493 | 0.4495683 | 0.92081798 | 3090.01107 | 3120.24744 | 2518.61893 | 3872.46932 | Non-sig. |
| PF3D7_1249500 | N/A | 776.600536 | 0.317889713 | 0.44410883 | 0.71579238 | 0.47411956 | 0.92682482 | 564.596783 | 912.685006 | 817.935468 | 811.184886 | Non-sig. |
| PF3D7_0107600 | eIK2 | 175.993681 | 0.317730149 | 0.59353215 | 0.53532087 | 0.592428 | 0.95322751 | 230.775098 | 176.987946 | 82.9664992 | 213.24518 | Non-sig. |
| PF3D7_1467000 | N/A | 37.822582 | 0.317631379 | 0.69263435 | 0.4585845 | 0.64653257 | 0.97317009 | 33.6774515 | 33.2642559 | 33.6719525 | 50.6766682 | Non-sig. |
| PF3D7_1338400 | N/A | 215.893542 | 0.316839185 | 0.54076021 | 0.58591439 | 0.55793305 | 0.94272305 | 257.598918 | 280.13816 | 127.275299 | 198.56179 | Non-sig. |
| PF3D7_1322000 | N/A | 2657.05608 | 0.316198142 | 0.41511437 | 0.76171331 | 0.44623113 | 0.91974025 | 2102.2202 | 2868.69982 | 2631.68743 | 3025.61688 | Non-sig. |
| PF3D7_1451600 | LAP5 | 91.0401669 | 0.316098877 | 0.66435714 | 0.47579661 | 0.63421928 | 0.9668494 | 37.8831391 | 93.3572929 | 124.009051 | 108.911184 | Non-sig. |
| PF3D7_1444100 | N/A | 2054.00842 | 0.316034365 | 0.49311645 | 0.64089196 | 0.52159288 | 0.93247207 | 2361.28444 | 1759.82669 | 1298.89496 | 2796.0276 | Non-sig. |
| PF3D7_0504900 | N/A | 406.113892 | 0.315890134 | 0.50974289 | 0.61970483 | 0.53545213 | 0.93526055 | 489.373645 | 448.265908 | 234.637906 | 452.17811 | Non-sig. |
| PF3D7_1476700 | LPL1 | 36.9376043 | 0.315755338 | 0.69564261 | 0.45390454 | 0.64989753 | 0.97317009 | 34.653145 | 33.2849369 | 31.1960444 | 48.6162907 | Non-sig. |
| PF3D7_1035800 | M712 | 27.6407875 | 0.315393486 | 0.8096289 | 0.38955315 | 0.696867 | 0.98334178 | 22.1143247 | 18.2385554 | 27.1047058 | 43.1055639 | Non-sig. |
| PF3D7_0219800 | N/A | 1553.07955 | 0.313849594 | 0.416522 | 0.75350063 | 0.45114913 | 0.92081798 | 1526.20433 | 1714.3243 | 1243.57596 | 1728.2136 | Non-sig. |
| PF3D7_1432800 | N/A | 1748.20977 | 0.313764246 | 0.41241412 | 0.76079899 | 0.44677714 | 0.91974025 | 1678.06504 | 1890.53666 | 1439.75731 | 1984.48006 | Non-sig. |
| PF3D7_0627800 | ACAS | 3086.83379 | 0.31333116 | 0.56872799 | 0.55093325 | 0.58167944 | 0.95129458 | 1304.02735 | 3151.82338 | 4201.41785 | 3690.06656 | Non-sig. |
| PF3D7_0530500 | N/A | 26.2511264 | 0.312981399 | 0.80498133 | 0.38880579 | 0.69741982 | 0.98334178 | 30.5283786 | 34.3347052 | 16.4261127 | 23.7153092 | Non-sig. |
| PF3D7_0420600 | N/A | 3522.66417 | 0.31208382 | 0.41281154 | 0.75599587 | 0.44965167 | 0.92081798 | 3474.75606 | 3795.02056 | 2811.67308 | 4009.20701 | Non-sig. |
| PF3D7_0619600 | N/A | 190.160576 | 0.311873716 | 0.48829972 | 0.63869321 | 0.52302253 | 0.93247207 | 149.514536 | 239.256128 | 189.728418 | 182.143225 | Non-sig. |
| PF3D7_1034600 | IF3b | 1200.74035 | 0.310989306 | 0.44373337 | 0.70084723 | 0.48339836 | 0.9306121 | 1261.34467 | 1485.95177 | 882.541868 | 1173.12308 | Non-sig. |
| PF3D7_1116100 | N/A | 19968.659 | 0.310715631 | 0.44683995 | 0.69536224 | 0.48682832 | 0.93170321 | 21790.9282 | 19425.5003 | 13862.5681 | 24795.6393 | Non-sig. |
| PF3D7_1453500 | NTH | 31.951643 | 0.310623669 | 0.78956782 | 0.39340974 | 0.69401688 | 0.98334178 | 15.7954766 | 40.7688766 | 41.067523 | 30.1746959 | Non-sig. |
| PF3D7_1458800 | N/A | 224.770746 | 0.310583363 | 0.47536961 | 0.65335132 | 0.51352981 | 0.93247207 | 176.958299 | 220.993012 | 224.239501 | 276.892172 | Non-sig. |
| PF3D7_1003000 | N/A | 3411.25911 | 0.310418583 | 0.50296311 | 0.61717963 | 0.53711625 | 0.93646681 | 4202.4185 | 3384.22555 | 1889.33561 | 4169.0568 | Non-sig. |
| PF3D7_1031400 | OTU | 3659.92792 | 0.310359139 | 0.4096581 | 0.75760528 | 0.44868732 | 0.92081798 | 3411.53086 | 3732.64904 | 3124.06328 | 4371.46851 | Non-sig. |
| PF3D7_1211700 | MCM5 | 8814.22834 | 0.310010012 | 0.56186388 | 0.55175288 | 0.58111768 | 0.95129458 | 3770.52492 | 9860.20975 | 11970.7334 | 9655.44528 | Non-sig. |
| PF3D7_0911800 | N/A | 1635.74052 | 0.309936552 | 0.4580345 | 0.6766664 | 0.49861764 | 0.93170321 | 1786.53059 | 2078.20552 | 1135.17232 | 1543.05366 | Non-sig. |
| PF3D7_1405000 | N/A | 852.227873 | 0.30988699 | 0.41788148 | 0.74156669 | 0.45834991 | 0.92288312 | 737.743219 | 901.355218 | 784.309471 | 985.503584 | Non-sig. |
| PF3D7_1320000 | GP1 | 2093.19276 | 0.309800284 | 0.46103377 | 0.67196875 | 0.50160359 | 0.93170321 | 1464.60497 | 1900.7915 | 2273.73927 | 2733.6353 | Non-sig. |
| PF3D7_0934500 | N/A | 2469.91499 | 0.309430827 | 0.42992028 | 0.71974001 | 0.47168509 | 0.92682482 | 2591.54678 | 2857.35555 | 1820.80915 | 2609.94847 | Non-sig. |
| PF3D7_1429800 | N/A | 8794.93999 | 0.30930618 | 0.46443458 | 0.66598439 | 0.50542108 | 0.93170321 | 5510.55537 | 9437.52481 | 10200.5629 | 10031.1169 | Non-sig. |
| PF3D7_1335500 | N/A | 22.8961195 | 0.309128132 | 0.81844535 | 0.37770162 | 0.70565227 | 0.98615891 | 22.0517685 | 20.4005057 | 18.8817812 | 30.2504224 | Non-sig. |
| PF3D7_0217000 | N/A | 270.267889 | 0.308669492 | 0.46099077 | 0.66957847 | 0.50312654 | 0.93170321 | 269.529877 | 278.958417 | 213.544722 | 319.038539 | Non-sig. |
| PF3D7_1307700 | N/A | 2458.86926 | 0.308658504 | 0.44550012 | 0.69283596 | 0.4884125 | 0.93170321 | 2094.44975 | 2095.01766 | 2299.15574 | 3346.85387 | Non-sig. |
| PF3D7_1118300 | N/A | 3600.42933 | 0.308577976 | 0.48113135 | 0.64135911 | 0.52128939 | 0.93247207 | 2218.39438 | 3414.16578 | 4214.93313 | 4554.22402 | Non-sig. |
| PF3D7_1229100 | MRP2 | 3019.56573 | 0.308512597 | 0.52522767 | 0.58738832 | 0.55694294 | 0.94272305 | 1632.07609 | 2629.765 | 3763.40337 | 4053.01846 | Non-sig. |
| PF3D7_1008100 | PHD1 | 15410.6905 | 0.308338138 | 0.43493557 | 0.7089283 | 0.47836897 | 0.92880667 | 11970.1621 | 14142.4965 | 15569.8936 | 19960.2095 | Non-sig. |
| PF3D7_0519700 | N/A | 2661.12212 | 0.308139994 | 0.41366612 | 0.74490025 | 0.45633202 | 0.92276605 | 2612.27026 | 2820.85889 | 2143.89827 | 3067.46106 | Non-sig. |
| PF3D7_0817200 | N/A | 1781.626 | 0.307945733 | 0.43639323 | 0.70566111 | 0.48039891 | 0.92880667 | 1406.83393 | 1655.43644 | 1777.39875 | 2286.83487 | Non-sig. |
| PF3D7_1472300 | N/A | 1276.74254 | 0.307817108 | 0.5018887 | 0.61331747 | 0.53966645 | 0.93711173 | 716.75826 | 1321.42677 | 1564.95571 | 1503.82943 | Non-sig. |
| PF3D7_1132700 | mRPL2 | 312.918264 | 0.307809515 | 0.54090249 | 0.56906655 | 0.56931098 | 0.94850402 | 172.283099 | 290.94277 | 386.693406 | 401.753781 | Non-sig. |
| PF3D7_1340300 | N/A | 838.922204 | 0.307680732 | 0.63149794 | 0.48722365 | 0.62609985 | 0.96651427 | 1221.58587 | 967.709 | 278.440551 | 887.953393 | Non-sig. |
| PF3D7_0910600 | YKT6.1 | 577.911608 | 0.307663136 | 0.43097987 | 0.71386892 | 0.47530823 | 0.92682482 | 529.909219 | 702.349395 | 503.176392 | 576.211425 | Non-sig. |
| PF3D7_0803300 | MORG1 | 957.662712 | 0.307489638 | 0.42265845 | 0.72751328 | 0.46691158 | 0.92538923 | 843.374606 | 1171.35729 | 868.590391 | 947.328562 | Non-sig. |
| PF3D7_1218700 | N/A | 93.5537129 | 0.307218053 | 0.58138156 | 0.52842758 | 0.5972026 | 0.95631916 | 109.212202 | 114.886041 | 58.2875612 | 91.829047 | Non-sig. |
| PF3D7_1442400 | KIC9 | 8997.672 | 0.306729438 | 0.46652315 | 0.65747957 | 0.5108726 | 0.93247207 | 10333.7352 | 8791.924 | 5756.15552 | 11108.8733 | Non-sig. |
| PF3D7_0320900 | H2A.Z | 12494.6511 | 0.305940048 | 0.42120965 | 0.72633674 | 0.46763236 | 0.92538923 | 9372.37469 | 14079.5309 | 12976.9483 | 13549.7506 | Non-sig. |
| PF3D7_1122500 | KIC10 | 948.939383 | 0.305178691 | 0.48541293 | 0.62869914 | 0.52954604 | 0.93270419 | 717.701054 | 742.251862 | 980.010722 | 1355.79389 | Non-sig. |
| PF3D7_1025000 | EPS15 | 8678.83771 | 0.305097187 | 0.58522022 | 0.5213374 | 0.60213175 | 0.95693578 | 12126.4103 | 10339.9033 | 3403.14938 | 8845.88789 | Non-sig. |
| PF3D7_1349300 | TKL3 | 2591.84702 | 0.305048493 | 0.48996055 | 0.62259806 | 0.53354868 | 0.93467024 | 1550.25948 | 2455.23084 | 3087.06897 | 3274.8288 | Non-sig. |
| PF3D7_0913000 | N/A | 663.346225 | 0.30497296 | 0.44282462 | 0.68869919 | 0.49101259 | 0.93170321 | 548.502975 | 850.060635 | 638.390353 | 616.430936 | Non-sig. |
| PF3D7_0111300 | N/A | 2511.42069 | 0.304279125 | 0.42580101 | 0.71460404 | 0.47485375 | 0.92682482 | 2584.66785 | 2587.30512 | 1910.66499 | 2963.0448 | Non-sig. |
| PF3D7_1312900 | EIF4G | 11228.6601 | 0.30422013 | 0.55426161 | 0.54887462 | 0.5830915 | 0.95129458 | 15124.2335 | 11855.7808 | 4974.4948 | 12960.1313 | Non-sig. |
| PF3D7_1452500 | STX6 | 1022.92335 | 0.304209361 | 0.52939649 | 0.57463426 | 0.56553867 | 0.94698087 | 1304.99432 | 1248.23089 | 526.357459 | 1012.11074 | Non-sig. |
| PF3D7_0807100 | PSH3 | 634.671122 | 0.303681977 | 0.48061025 | 0.63186745 | 0.52747349 | 0.93270419 | 725.850114 | 620.041214 | 410.705949 | 782.087212 | Non-sig. |

| | | | | | | | | | | | | |
|---|---|---|---|---|---|---|---|---|---|---|---|---|
| PF3D7_0525000 | N/A | 526.123654 | 0.302744775 | 0.44357167 | 0.68251602 | 0.49491271 | 0.93170321 | 545.716558 | 584.637281 | 396.743482 | 577.397294 | Non-sig. |
| PF3D7_1024400 | N/A | 258.391661 | 0.302369778 | 0.55052969 | 0.54923428 | 0.58284468 | 0.95129458 | 326.886024 | 308.072947 | 136.299922 | 262.307753 | Non-sig. |
| PF3D7_1014900 | KIC8 | 15649.1477 | 0.302204421 | 0.50841482 | 0.59440522 | 0.55224112 | 0.94170064 | 19466.8718 | 14874.3004 | 8565.7156 | 19689.7032 | Non-sig. |
| PF3D7_0828600 | FT1 | 1499.70417 | 0.302204366 | 0.50320411 | 0.60056022 | 0.54813294 | 0.94142878 | 830.060005 | 1606.50473 | 1855.94653 | 1706.3054 | Non-sig. |
| PF3D7_0504600 | BCKDHB | 1107.6781 | 0.30188623 | 0.47600355 | 0.63421004 | 0.52594376 | 0.93247207 | 688.406279 | 1255.93349 | 1295.65335 | 1190.71928 | Non-sig. |
| PF3D7_0216400 | VPS45 | 988.450472 | 0.301113769 | 0.46329255 | 0.64994304 | 0.51572902 | 0.93247207 | 652.461233 | 1037.6478 | 1118.55403 | 1145.13883 | Non-sig. |
| PF3D7_0930900 | NFU1 | 135.254517 | 0.300274287 | 0.52311668 | 0.57401016 | 0.56596093 | 0.94738104 | 149.093592 | 152.47221 | 93.5859294 | 145.866338 | Non-sig. |
| PF3D7_0309300 | N/A | 1318.17542 | 0.299756448 | 0.56031315 | 0.53498022 | 0.59266354 | 0.95322751 | 585.572487 | 1391.44806 | 1777.44838 | 1518.23276 | Non-sig. |
| PF3D7_1130000 | PAGM | 709.791173 | 0.299679662 | 0.43401778 | 0.69047785 | 0.48989373 | 0.93170321 | 722.537736 | 818.545759 | 550.325798 | 747.7554 | Non-sig. |
| PF3D7_1368200 | ABCE1 | 1531.73455 | 0.299122679 | 0.52340273 | 0.57149621 | 0.56766333 | 0.94843277 | 802.457537 | 1488.0393 | 1944.15655 | 1892.28483 | Non-sig. |
| PF3D7_1433700 | N/A | 1505.54394 | 0.299068068 | 0.4360483 | 0.68585995 | 0.49280143 | 0.93170321 | 1129.99609 | 1493.0017 | 1569.87829 | 1829.2997 | Non-sig. |
| PF3D7_1133300 | CDC50B | 2959.52751 | 0.299002403 | 0.41309439 | 0.72381134 | 0.46918157 | 0.92681728 | 2509.95322 | 3565.08856 | 2797.83839 | 2965.22986 | Non-sig. |
| PF3D7_0625300 | MtDNAP | 2775.54768 | 0.29870642 | 0.43234599 | 0.69089671 | 0.48963045 | 0.93170321 | 2948.9897 | 2901.66978 | 2029.74352 | 3221.78772 | Non-sig. |
| PF3D7_1316500 | PRP40 | 889.496029 | 0.29833518 | 0.46778872 | 0.63775625 | 0.52363236 | 0.93247207 | 1018.60183 | 992.312491 | 577.441033 | 969.628762 | Non-sig. |
| PF3D7_0605100 | KH1 | 3888.36518 | 0.298166083 | 0.55563239 | 0.53662473 | 0.59152686 | 0.95321031 | 1724.88517 | 4171.64239 | 5250.64135 | 4406.2918 | Non-sig. |
| PF3D7_0917800 | N/A | 212.719117 | 0.297799528 | 0.54115361 | 0.55030498 | 0.58211021 | 0.95129458 | 121.903783 | 209.328252 | 259.454449 | 260.189984 | Non-sig. |
| PF3D7_0926100 | N/A | 7525.57897 | 0.297632601 | 0.53857421 | 0.55263062 | 0.58051637 | 0.95129458 | 3617.41368 | 7422.7198 | 9886.2566 | 9175.92582 | Non-sig. |
| PF3D7_0604300 | N/A | 845.234927 | 0.297545594 | 0.43106918 | 0.69025022 | 0.49003685 | 0.93170321 | 708.494967 | 814.367866 | 808.169936 | 1049.90694 | Non-sig. |
| PF3D7_0408500 | FEN1 | 2001.32774 | 0.297054981 | 0.49561009 | 0.59937234 | 0.54892462 | 0.94142878 | 2450.87821 | 2336.06518 | 1141.54431 | 2076.82326 | Non-sig. |
| PF3D7_0818800 | IMP4 | 162.079559 | 0.296940155 | 0.48309848 | 0.6146576 | 0.53878087 | 0.93711173 | 139.792146 | 166.38138 | 151.079136 | 191.065573 | Non-sig. |
| PF3D7_1416200 | MCA3 | 6036.55541 | 0.296846267 | 0.41269867 | 0.71928089 | 0.47196787 | 0.92682482 | 5669.7824 | 5937.44058 | 5165.69623 | 7373.30241 | Non-sig. |
| PF3D7_0917600 | PRP43 | 996.409674 | 0.296582336 | 0.45550163 | 0.65111147 | 0.51497453 | 0.93247207 | 677.975947 | 1065.43317 | 1110.41535 | 1131.81423 | Non-sig. |
| PF3D7_1004300 | N/A | 5562.04517 | 0.296045224 | 0.44206753 | 0.66968326 | 0.50305973 | 0.93170321 | 5535.43888 | 4863.41477 | 4451.43989 | 7397.88715 | Non-sig. |
| PF3D7_0704500 | N/A | 7982.76444 | 0.295869956 | 0.45382886 | 0.6519417 | 0.51443879 | 0.93247207 | 6536.75691 | 6545.26931 | 7797.25719 | 11051.7744 | Non-sig. |
| PF3D7_1232400 | N/A | 965.782267 | 0.29525405 | 0.59121346 | 0.49940346 | 0.61749518 | 0.96395246 | 1352.41423 | 1116.43757 | 382.608872 | 1011.6684 | Non-sig. |
| PF3D7_1226200 | N/A | 127.467488 | 0.295006537 | 0.57239003 | 0.51539426 | 0.60627754 | 0.95706907 | 156.975176 | 129.799586 | 72.2847397 | 150.810451 | Non-sig. |
| PF3D7_1366400 | RHOP148 | 2852.7085 | 0.294574468 | 0.41777776 | 0.7050985 | 0.48074894 | 0.92880667 | 2883.37404 | 3023.73369 | 2241.77439 | 3261.95189 | Non-sig. |
| PF3D7_1234600 | TOC75 | 1703.12964 | 0.294314997 | 0.50354024 | 0.58449152 | 0.55888967 | 0.94389464 | 947.869555 | 1746.55139 | 2111.74683 | 2006.35079 | Non-sig. |
| PF3D7_1314000 | N/A | 345.353934 | 0.294287136 | 0.53517387 | 0.54989071 | 0.58239434 | 0.95129458 | 186.439471 | 351.871698 | 433.694491 | 409.410074 | Non-sig. |
| PF3D7_0417800 | CRK1 | 1604.08947 | 0.294233517 | 0.49141063 | 0.59875285 | 0.54933771 | 0.94142878 | 1952.827 | 1784.31539 | 929.72326 | 1749.49224 | Non-sig. |
| PF3D7_1439500 | ORP2 | 2199.96444 | 0.29400233 | 0.41799937 | 0.70335591 | 0.48183398 | 0.92880667 | 1906.07531 | 2133.5692 | 2047.01903 | 2713.19423 | Non-sig. |
| PF3D7_1372000 | N/A | 37.1449209 | 0.293007394 | 0.69907993 | 0.4191329 | 0.67511901 | 0.97898416 | 38.9393345 | 47.2138572 | 27.9230551 | 34.5034367 | Non-sig. |
| PF3D7_1470800 | N/A | 1786.38922 | 0.29278569 | 0.47503396 | 0.61634686 | 0.53766562 | 0.93646681 | 2103.01277 | 1848.5199 | 1108.83309 | 2085.1911 | Non-sig. |
| PF3D7_1430700 | GDH2 | 689.76086 | 0.29252783 | 0.4314871 | 0.67795266 | 0.49780171 | 0.93170321 | 577.365906 | 840.359632 | 662.679657 | 678.638245 | Non-sig. |
| PF3D7_1464700 | N/A | 1841.04512 | 0.292147971 | 0.49452037 | 0.59077035 | 0.55467431 | 0.94243456 | 1084.67124 | 1768.54231 | 2225.49966 | 2285.46729 | Non-sig. |
| PF3D7_1139400 | N/A | 30.3353808 | 0.291888119 | 0.76556073 | 0.38127363 | 0.70300022 | 0.98512564 | 35.8041081 | 37.5499777 | 18.8911552 | 29.0962823 | Non-sig. |
| PF3D7_1405300 | N/A | 533.587677 | 0.290191032 | 0.48556139 | 0.59764025 | 0.55008 | 0.94162321 | 342.229462 | 526.779341 | 617.653699 | 647.688207 | Non-sig. |
| PF3D7_0918400 | N/A | 48.7324788 | 0.28971616 | 0.71706887 | 0.40402836 | 0.68619186 | 0.98210107 | 24.233829 | 64.3613659 | 63.2504032 | 43.0843171 | Non-sig. |
| PF3D7_0103900 | PIESP15 | 674.187376 | 0.289145737 | 0.44047348 | 0.65644302 | 0.51153912 | 0.93247207 | 507.206902 | 767.249372 | 706.275196 | 716.018033 | Non-sig. |
| PF3D7_1347800 | CEP72 | 26.0942788 | 0.28885504 | 0.81472564 | 0.35454271 | 0.7229322 | 0.98844303 | 32.3634664 | 31.1883429 | 14.7636783 | 26.0616274 | Non-sig. |
| PF3D7_1248900 | RPT6 | 8343.4182 | 0.28865792 | 0.45595419 | 0.63308535 | 0.52667791 | 0.93270419 | 5742.60507 | 10539.1203 | 9280.11914 | 7811.82831 | Non-sig. |
| PF3D7_1320700 | N/A | 5395.55737 | 0.288356199 | 0.44893103 | 0.64231737 | 0.52066714 | 0.93247207 | 5921.52435 | 5155.33538 | 3795.02876 | 6710.341 | Non-sig. |
| PF3D7_1012900 | ATG18 | 3276.09951 | 0.287985722 | 0.41535534 | 0.69334782 | 0.4880913 | 0.93170321 | 3231.09868 | 3328.98545 | 2669.44626 | 3874.86766 | Non-sig. |
| PF3D7_1140900 | N/A | 1032.78036 | 0.287908879 | 0.45506781 | 0.63267248 | 0.52694755 | 0.93270419 | 708.151594 | 1078.00901 | 1151.67919 | 1193.28165 | Non-sig. |
| PF3D7_0828500 | N/A | 693.117987 | 0.287650681 | 0.45578337 | 0.63111271 | 0.52796683 | 0.93270419 | 485.465975 | 799.763188 | 762.750166 | 724.492617 | Non-sig. |
| PF3D7_1314200 | TERT | 856.602922 | 0.287596565 | 0.47583674 | 0.60440176 | 0.54557656 | 0.94121391 | 992.94395 | 861.26928 | 550.400285 | 1021.79817 | Non-sig. |
| PF3D7_1112800 | N/A | 75.4960919 | 0.287464834 | 0.56105447 | 0.51236528 | 0.60839538 | 0.9580165 | 74.5133866 | 84.8367607 | 61.5648268 | 81.0693936 | Non-sig. |
| PF3D7_0404800 | N/A | 681.324153 | 0.287441462 | 0.52605154 | 0.54641312 | 0.58478199 | 0.95152795 | 868.882991 | 776.985977 | 358.863665 | 720.563981 | Non-sig. |
| PF3D7_0627100 | N/A | 6858.79253 | 0.287191594 | 0.50822958 | 0.56508241 | 0.57201771 | 0.94850402 | 4176.04268 | 5575.27471 | 8180.35031 | 9503.50244 | Non-sig. |
| PF3D7_0200300 | N/A | 3.46125661 | 0.287119796 | 2.10879363 | 0.13615358 | 0.89169987 | 0.99457954 | 6.2945629 | 2.14715346 | 0 | 5.4033101 | Non-sig. |
| PF3D7_1150300 | RIF | 1.95800816 | 0.286427311 | 2.35349318 | 0.12170306 | 0.90313419 | 0.99457954 | 1.04748284 | 1.07445791 | 2.46176181 | 3.24833009 | Non-sig. |
| PF3D7_1104500 | N/A | 37.2909162 | 0.286348686 | 0.72329509 | 0.39589469 | 0.69218272 | 0.98334178 | 45.2071445 | 42.9341441 | 22.1701839 | 38.8521925 | Non-sig. |
| PF3D7_1224100 | N/A | 3771.13826 | 0.286293644 | 0.40743933 | 0.70266571 | 0.4822641 | 0.92880667 | 3584.77369 | 4353.96557 | 3211.6819 | 3934.1319 | Non-sig. |
| PF3D7_1368800 | ERCC4 | 1572.25316 | 0.286037663 | 0.42464164 | 0.67359778 | 0.50056707 | 0.93170321 | 1568.08796 | 1555.32576 | 1265.8667 | 1899.73223 | Non-sig. |

| | | | | | | | | | | | | |
|---|---|---|---|---|---|---|---|---|---|---|---|---|
| PF3D7_1227800 | ELP3 | 928.628844 | 0.285886167 | 0.41557546 | 0.68792842 | 0.49149786 | 0.93170321 | 808.877918 | 996.639889 | 864.909931 | 1044.08764 | Non-sig. |
| PF3D7_0307500 | SKA2 | 1123.8578 | 0.285857077 | 0.41500638 | 0.68880164 | 0.4909481 | 0.93170321 | 1075.7798 | 1258.95486 | 950.075292 | 1210.62125 | Non-sig. |
| PF3D7_1329100 | MyoF | 7900.28744 | 0.285441204 | 0.47968502 | 0.59505966 | 0.5518036 | 0.94170064 | 4842.61629 | 7641.94642 | 9399.57985 | 9717.00721 | Non-sig. |
| PF3D7_1368500 | N/A | 190.797191 | 0.285125072 | 0.55163268 | 0.51687487 | 0.60524351 | 0.95693578 | 106.320688 | 193.133885 | 237.356438 | 226.377753 | Non-sig. |
| PF3D7_0800100 | VAR | 1194.60812 | 0.285059652 | 0.43314114 | 0.65812186 | 0.51045983 | 0.93247207 | 896.859507 | 1263.54568 | 1256.84445 | 1361.18282 | Non-sig. |
| PF3D7_0806700 | N/A | 1704.58218 | 0.283682063 | 0.41184618 | 0.68880586 | 0.49094545 | 0.93170321 | 1586.54191 | 1746.08829 | 1488.56983 | 1997.12867 | Non-sig. |
| PF3D7_1217100 | SPO11 | 77.0523235 | 0.283224921 | 0.56607826 | 0.50032821 | 0.61684399 | 0.96358724 | 80.0359449 | 88.9867824 | 59.1424282 | 80.0441383 | Non-sig. |
| PF3D7_1458500 | SAS4 | 2332.86428 | 0.282721844 | 0.56202415 | 0.50304216 | 0.61493463 | 0.96217474 | 3189.4107 | 2503.65093 | 1021.04379 | 2617.3517 | Non-sig. |
| PF3D7_0105000 | N/A | 205.594205 | 0.282418023 | 0.50216341 | 0.56240264 | 0.57384172 | 0.94850402 | 138.547567 | 226.601565 | 232.281224 | 224.946463 | Non-sig. |
| PF3D7_0509900 | N/A | 212.813197 | 0.282191476 | 0.46644981 | 0.60497715 | 0.54519418 | 0.94121391 | 175.483638 | 237.244675 | 208.545537 | 229.978937 | Non-sig. |
| PF3D7_1030400 | N/A | 105.411372 | 0.281681978 | 0.60376539 | 0.46654211 | 0.64082751 | 0.97002976 | 134.817651 | 123.640958 | 55.783188 | 107.403693 | Non-sig. |
| PF3D7_1318400 | SMC2 | 6205.56391 | 0.281271456 | 0.59340768 | 0.47399362 | 0.63550446 | 0.96694503 | 8843.10068 | 7109.81716 | 2362.43713 | 6506.90068 | Non-sig. |
| PF3D7_1007900 | EIF3D | 1703.00455 | 0.281174966 | 0.45506541 | 0.61787813 | 0.53665567 | 0.93618761 | 1145.52225 | 1797.127 | 1929.30039 | 1940.06855 | Non-sig. |
| PF3D7_1421500 | TIM8 | 42.966926 | 0.280310768 | 0.76222873 | 0.36775151 | 0.71305853 | 0.9875928 | 17.5968169 | 47.433626 | 59.7851304 | 47.0521306 | Non-sig. |
| PF3D7_1217800 | DPH3 | 67.1221378 | 0.280295636 | 0.68088995 | 0.4116607 | 0.68058813 | 0.97982498 | 33.2851904 | 55.9796185 | 87.7098947 | 91.5138478 | Non-sig. |
| PF3D7_0321700 | N/A | 3169.33687 | 0.279042103 | 0.65547797 | 0.42570783 | 0.67032076 | 0.97883902 | 4810.40476 | 3643.59099 | 917.593724 | 3305.758 | Non-sig. |
| PF3D7_0301700 | N/A | 2370.06828 | 0.278766607 | 0.41158565 | 0.67729915 | 0.49821617 | 0.93170321 | 2102.8854 | 2371.03808 | 2180.68202 | 2825.66761 | Non-sig. |
| PF3D7_0323700 | SART1 | 2428.51633 | 0.278635046 | 0.65627906 | 0.42456793 | 0.67115167 | 0.97898416 | 3666.27361 | 3081.35937 | 723.59656 | 2242.83576 | Non-sig. |
| PF3D7_1030100 | PRP22 | 2670.39624 | 0.278535452 | 0.40847402 | 0.68189269 | 0.49530681 | 0.93170321 | 2576.1998 | 2987.42752 | 2250.73448 | 2867.22314 | Non-sig. |
| PF3D7_1458300 | N/A | 6207.72325 | 0.278158107 | 0.48763696 | 0.57042048 | 0.56839255 | 0.94850402 | 3603.77813 | 6685.90963 | 7618.09244 | 6923.11278 | Non-sig. |
| PF3D7_1014000 | FS | 214.498185 | 0.278141034 | 0.46811948 | 0.59416676 | 0.55240058 | 0.94170064 | 199.770223 | 258.671768 | 188.03893 | 211.511819 | Non-sig. |
| PF3D7_0904300 | N/A | 139.726401 | 0.277932652 | 0.49103467 | 0.56601431 | 0.57138405 | 0.94850402 | 128.593153 | 155.523445 | 124.044651 | 150.744355 | Non-sig. |
| PF3D7_0403500 | N/A | 730.210043 | 0.277817334 | 0.42156342 | 0.6590167 | 0.50988504 | 0.93247207 | 630.49296 | 838.402864 | 689.680617 | 762.263731 | Non-sig. |
| PF3D7_1472100 | YIP1 | 1359.87079 | 0.277715051 | 0.41486946 | 0.66940345 | 0.50323815 | 0.93170321 | 1133.86955 | 1528.92609 | 1324.78818 | 1451.89933 | Non-sig. |
| PF3D7_1467600 | N/A | 848.077546 | 0.276925309 | 0.72325137 | 0.38288944 | 0.70180174 | 0.98512564 | 1353.52318 | 907.373754 | 180.730395 | 950.682856 | Non-sig. |
| PF3D7_1416800 | KRS2 | 588.942336 | 0.276616396 | 0.42877025 | 0.64513896 | 0.51883714 | 0.93247207 | 531.482135 | 702.837871 | 533.825367 | 587.623971 | Non-sig. |
| PF3D7_1423800 | VPS3 | 1454.26892 | 0.276207397 | 0.43866114 | 0.62966006 | 0.52891702 | 0.93270419 | 1577.33411 | 1620.82565 | 1053.90283 | 1565.01308 | Non-sig. |
| PF3D7_1245100 | KLP8 | 647.736843 | 0.276190805 | 0.51299908 | 0.5383846 | 0.59031156 | 0.95315309 | 391.967656 | 563.157172 | 779.544776 | 856.277766 | Non-sig. |
| PF3D7_1011000 | ISP1 | 679.904459 | 0.276177702 | 0.4558066 | 0.60590984 | 0.54457463 | 0.94101643 | 480.173026 | 787.972575 | 749.602763 | 701.869472 | Non-sig. |
| PF3D7_1472200 | HDA1 | 7699.01943 | 0.276117962 | 0.4312289 | 0.64030486 | 0.52197442 | 0.93247207 | 8052.13244 | 7391.83908 | 5877.09583 | 9475.01037 | Non-sig. |
| PF3D7_1409500 | N/A | 871.306422 | 0.275752723 | 0.63668494 | 0.43310703 | 0.66493702 | 0.97883902 | 1291.16074 | 1007.38882 | 285.835515 | 900.840608 | Non-sig. |
| PF3D7_0724500 | N/A | 31.6150582 | 0.275244521 | 0.75474865 | 0.36468369 | 0.71534754 | 0.9875928 | 37.6653671 | 31.169553 | 19.6916613 | 37.9336515 | Non-sig. |
| PF3D7_1041200 | RIF | 31.4927991 | 0.27450821 | 0.73248992 | 0.37476039 | 0.70783867 | 0.98661649 | 24.1085384 | 30.0790742 | 32.8274568 | 38.9561272 | Non-sig. |
| PF3D7_1117200 | N/A | 289.844412 | 0.274429423 | 0.55818772 | 0.4916436 | 0.62297131 | 0.96429926 | 377.23064 | 286.402557 | 147.859849 | 347.884602 | Non-sig. |
| PF3D7_0510500 | TopoI | 3940.53863 | 0.274313492 | 0.5098711 | 0.53800557 | 0.5905732 | 0.95315309 | 5017.09402 | 4498.67567 | 2117.39711 | 4128.98773 | Non-sig. |
| PF3D7_0301600 | GEXP21 | 583.023672 | 0.273545645 | 0.58619994 | 0.46664222 | 0.64075587 | 0.97002976 | 243.157555 | 626.620529 | 812.261487 | 650.055118 | Non-sig. |
| PF3D7_1028900 | IMC1m | 2009.7801 | 0.273520671 | 0.49087081 | 0.55721519 | 0.57738041 | 0.9497432 | 2466.73565 | 2282.14038 | 1173.3163 | 2116.92807 | Non-sig. |
| PF3D7_0105800 | N/A | 8908.25941 | 0.27288827 | 0.42980446 | 0.6349126 | 0.52548542 | 0.93247207 | 9594.33848 | 10170.8423 | 6542.45198 | 9325.40487 | Non-sig. |
| PF3D7_1445000 | N/A | 463.883433 | 0.272805056 | 0.43025527 | 0.63405395 | 0.52604561 | 0.93247207 | 402.774483 | 501.582827 | 437.477101 | 513.699319 | Non-sig. |
| PF3D7_1345600 | N/A | 6863.09747 | 0.270368531 | 0.46063796 | 0.58694367 | 0.55724154 | 0.94272305 | 4519.53812 | 8214.97342 | 7923.89758 | 6793.98076 | Non-sig. |
| PF3D7_1125100 | ICAP2 | 642.717473 | 0.269996758 | 0.51988607 | 0.51933832 | 0.60352484 | 0.95693578 | 355.457078 | 632.151328 | 809.665362 | 773.596122 | Non-sig. |
| PF3D7_0322700 | N/A | 1563.53549 | 0.269749363 | 0.43265801 | 0.62347017 | 0.5329756 | 0.93467024 | 1221.91352 | 1512.2343 | 1613.46594 | 1906.52821 | Non-sig. |
| PF3D7_0713100 | MC-2TM | 34.0706064 | 0.269735446 | 0.81404281 | 0.3313529 | 0.74037794 | 0.98954916 | 47.2229398 | 46.1780285 | 14.7753457 | 28.1061113 | Non-sig. |
| PF3D7_0811000 | CUL1 | 3739.44153 | 0.269596726 | 0.40447888 | 0.66652856 | 0.50507331 | 0.93170321 | 3275.43163 | 4197.55998 | 3506.63566 | 3978.13885 | Non-sig. |
| PF3D7_0930500 | DGK1 | 9179.70774 | 0.269563228 | 0.50154481 | 0.53746589 | 0.59094584 | 0.95315309 | 11581.3077 | 9627.22416 | 5068.29984 | 10441.9992 | Non-sig. |
| PF3D7_1347900 | N/A | 946.087738 | 0.269518091 | 0.54329755 | 0.49607824 | 0.61983921 | 0.9639908 | 459.502694 | 1046.872 | 1256.02637 | 1021.9499 | Non-sig. |
| PF3D7_1230000 | N/A | 1722.77627 | 0.268439549 | 0.41544982 | 0.64614193 | 0.51818745 | 0.93247207 | 1707.37735 | 1806.26809 | 1418.68462 | 1958.775 | Non-sig. |
| PF3D7_1038300 | N/A | 5.94048043 | 0.267664276 | 1.50133802 | 0.17828382 | 0.85850009 | 0.99402081 | 8.38470361 | 9.66855661 | 2.46201378 | 3.24664772 | Non-sig. |
| PF3D7_1237200 | N/A | 864.82045 | 0.267593269 | 0.4195831 | 0.63775989 | 0.52362999 | 0.93247207 | 837.861937 | 917.083374 | 731.930138 | 972.406349 | Non-sig. |
| PF3D7_1315500 | N/A | 1588.83226 | 0.267443485 | 0.43208845 | 0.61895541 | 0.53594574 | 0.93549683 | 1206.55102 | 1858.5916 | 1677.19578 | 1612.99065 | Non-sig. |
| PF3D7_0607000 | N/A | 8289.67626 | 0.266328738 | 0.64058547 | 0.41575832 | 0.67758686 | 0.97898416 | 12491.6913 | 9363.12892 | 2562.11587 | 8741.76896 | Non-sig. |
| PF3D7_1115700 | FP2A | 3546.62261 | 0.266212984 | 0.45680627 | 0.58276999 | 0.56004815 | 0.94472957 | 2351.58643 | 4045.69396 | 4088.7331 | 3700.47696 | Non-sig. |
| PF3D7_0215800 | ORC5 | 784.800601 | 0.265818401 | 0.44426226 | 0.59833668 | 0.54961531 | 0.94142878 | 575.904374 | 878.721232 | 849.249603 | 835.327193 | Non-sig. |
| PF3D7_1425500 | N/A | 84.1806937 | 0.264823873 | 0.61069233 | 0.43364533 | 0.66454601 | 0.97883902 | 105.628307 | 93.519689 | 47.5850188 | 89.9897599 | Non-sig. |

| | | | | | | | | | | | | |
|---|---|---|---|---|---|---|---|---|---|---|---|---|
| PF3D7_1132400 | N/A | 2787.5611 | 0.263992082 | 0.40542451 | 0.65114979 | 0.5149498 | 0.93247207 | 2476.38389 | 2947.64371 | 2590.03887 | 3136.17792 | Non-sig. |
| PF3D7_0106700 | AARP2 | 3204.79075 | 0.263909223 | 0.40986825 | 0.64388794 | 0.5196481 | 0.93247207 | 2657.36545 | 3470.12332 | 3167.46401 | 3524.21023 | Non-sig. |
| PF3D7_0816700 | TRAPPC2L | 172.209947 | 0.263777935 | 0.4888605 | 0.53957711 | 0.5894887 | 0.95315309 | 136.405106 | 202.975524 | 176.47948 | 172.979679 | Non-sig. |
| PF3D7_1316400 | SAS6L | 9.89758247 | 0.263654545 | 1.13326091 | 0.23265123 | 0.81603224 | 0.99096399 | 7.29523751 | 10.7597524 | 10.6595446 | 10.8757954 | Non-sig. |
| PF3D7_1234100 | BDP5 | 7580.62671 | 0.263379888 | 0.48255833 | 0.54579907 | 0.58520406 | 0.95192236 | 9185.87277 | 7432.2512 | 4595.66853 | 9108.71433 | Non-sig. |
| PF3D7_1018000 | N/A | 425.975648 | 0.263352285 | 0.48172417 | 0.54668688 | 0.58459387 | 0.95151408 | 489.78926 | 425.079732 | 284.914648 | 504.118952 | Non-sig. |
| PF3D7_1459000 | DBP5 | 3650.50833 | 0.26326375 | 0.4186199 | 0.62888495 | 0.52942438 | 0.93270419 | 3722.84082 | 4256.27476 | 2914.05858 | 3708.85914 | Non-sig. |
| PF3D7_1407600 | N/A | 644.443165 | 0.263257611 | 0.63994034 | 0.41137837 | 0.68079511 | 0.9798567 | 960.089997 | 710.07465 | 211.938367 | 695.669644 | Non-sig. |
| PF3D7_1246900 | PKB | 556.230621 | 0.261440614 | 0.43566232 | 0.60009921 | 0.54844012 | 0.94142878 | 561.180964 | 643.746533 | 450.909485 | 569.085502 | Non-sig. |
| PF3D7_1449300 | BRF1 | 1466.09544 | 0.260882565 | 0.45615719 | 0.57191374 | 0.56738042 | 0.94825913 | 996.936894 | 1681.31458 | 1670.54369 | 1515.5866 | Non-sig. |
| PF3D7_1030300 | N/A | 2457.70224 | 0.260571624 | 0.50347933 | 0.51754185 | 0.60477796 | 0.95693578 | 3061.87243 | 2314.86788 | 1411.22174 | 3042.84691 | Non-sig. |
| PF3D7_1330500 | N/A | 1606.82732 | 0.260297269 | 0.5802777 | 0.44857362 | 0.65373927 | 0.97317009 | 2261.33451 | 1815.41838 | 663.621573 | 1686.93483 | Non-sig. |
| PF3D7_0612100 | EIF3L | 2556.75593 | 0.260003119 | 0.48761127 | 0.53321803 | 0.59388266 | 0.9542761 | 1508.73367 | 2832.50187 | 3144.85599 | 2740.93219 | Non-sig. |
| PF3D7_0613700 | N/A | 2064.34854 | 0.259773026 | 0.47310183 | 0.5490848 | 0.58294725 | 0.95129458 | 2454.00401 | 2105.92805 | 1304.34534 | 2393.11676 | Non-sig. |
| PF3D7_0908700 | N/A | 468.582624 | 0.259726694 | 0.44046892 | 0.58965953 | 0.55541893 | 0.94243456 | 420.326511 | 575.396382 | 432.692097 | 445.915507 | Non-sig. |
| PF3D7_0908200 | N/A | 2788.42062 | 0.259568303 | 0.430961 | 0.60230115 | 0.5469737 | 0.94142878 | 2384.54493 | 2501.46836 | 2691.86747 | 3575.80169 | Non-sig. |
| PF3D7_1039700 | MC-2TM | 15.891986 | 0.259488862 | 0.9359528 | 0.27724567 | 0.78159147 | 0.99028237 | 16.6783869 | 19.3664426 | 12.2998907 | 15.2232238 | Non-sig. |
| PF3D7_1023900 | CHD1 | 104340.459 | 0.259182815 | 0.40294209 | 0.64322596 | 0.52007749 | 0.93247207 | 93613.0247 | 106286.96 | 96373.259 | 121088.59 | Non-sig. |
| PF3D7_1134800 | N/A | 1822.69871 | 0.258765969 | 0.40763725 | 0.6347947 | 0.52556232 | 0.93247207 | 1609.87493 | 1988.11486 | 1709.4257 | 1983.37937 | Non-sig. |
| PF3D7_1228800 | N/A | 13066.2275 | 0.257974318 | 0.54148874 | 0.47641677 | 0.63377749 | 0.9668494 | 17527.8764 | 12822.8006 | 6274.79901 | 15639.4342 | Non-sig. |
| PF3D7_1117000 | N/A | 262.492164 | 0.257081568 | 0.58713737 | 0.43785591 | 0.66149074 | 0.97783983 | 119.01108 | 278.932747 | 358.939882 | 293.084949 | Non-sig. |
| PF3D7_0913800 | N/A | 2968.08511 | 0.256440781 | 0.70374314 | 0.36439542 | 0.71556275 | 0.9875928 | 4672.06543 | 4068.47707 | 738.292871 | 2393.50507 | Non-sig. |
| PF3D7_1247800 | DPAP2 | 1130.13125 | 0.256437356 | 0.46730508 | 0.5487579 | 0.58317161 | 0.95129458 | 1320.33211 | 1266.31063 | 739.917045 | 1193.96522 | Non-sig. |
| PF3D7_0204900 | COQ5 | 467.363996 | 0.256217902 | 0.49541201 | 0.51718145 | 0.6050295 | 0.95693578 | 291.777193 | 547.669529 | 559.836445 | 470.172816 | Non-sig. |
| PF3D7_0218200 | N/A | 1237.71059 | 0.256131685 | 0.56091242 | 0.456634 | 0.64793413 | 0.97317009 | 1698.95991 | 1359.22566 | 557.720999 | 1334.93577 | Non-sig. |
| PF3D7_0515100 | ROM9 | 321.339786 | 0.255872125 | 0.44492577 | 0.57508946 | 0.56523079 | 0.94698087 | 275.388548 | 362.815593 | 310.370782 | 336.78422 | Non-sig. |
| PF3D7_1305700 | N/A | 30.3325351 | 0.255431943 | 0.73313113 | 0.34841235 | 0.72753053 | 0.98844303 | 24.9405324 | 35.5345696 | 30.324986 | 30.5300522 | Non-sig. |
| PF3D7_1476500 | N/A | 419.884918 | 0.255350378 | 0.491161 | 0.5198914 | 0.60313927 | 0.95693578 | 483.385151 | 528.909114 | 282.549002 | 384.696407 | Non-sig. |
| PF3D7_0916100 | TRX3 | 266.695941 | 0.254922692 | 0.45717788 | 0.55760067 | 0.5771171 | 0.9497432 | 223.602461 | 310.327932 | 262.689497 | 270.163875 | Non-sig. |
| PF3D7_0415700 | N/A | 444.848505 | 0.254116568 | 0.43533935 | 0.58372065 | 0.55940827 | 0.94416854 | 421.566373 | 447.593386 | 390.028329 | 520.205934 | Non-sig. |
| PF3D7_1027300 | nPrx | 46002.7296 | 0.253761361 | 0.74916573 | 0.33872527 | 0.7348167 | 0.98909017 | 75415.3058 | 60231.8865 | 8519.85405 | 39843.8722 | Non-sig. |
| PF3D7_1417800 | MCM2 | 8259.42996 | 0.253418781 | 0.53304138 | 0.47542046 | 0.63448731 | 0.9668494 | 4058.24159 | 8721.84321 | 11013.0979 | 9244.5371 | Non-sig. |
| PF3D7_0603200 | BCS1 | 707.692164 | 0.253102634 | 0.44920028 | 0.56345164 | 0.57312738 | 0.94850402 | 693.02156 | 917.644451 | 598.624618 | 621.478029 | Non-sig. |
| PF3D7_0302100 | SRPK1 | 1755.2825 | 0.251886522 | 0.42116086 | 0.59807676 | 0.54978872 | 0.94142878 | 1808.56492 | 1973.59162 | 1396.50613 | 1842.46732 | Non-sig. |
| PF3D7_0108900 | N/A | 260.406765 | 0.251012222 | 0.64254826 | 0.39065116 | 0.69605511 | 0.98334178 | 382.546301 | 271.331115 | 93.4387529 | 294.310892 | Non-sig. |
| PF3D7_1412700 | N/A | 523.114078 | 0.249845827 | 0.54930992 | 0.45483582 | 0.64922736 | 0.97317009 | 697.50929 | 561.017372 | 258.751422 | 575.178229 | Non-sig. |
| PF3D7_1348800 | N/A | 134.785991 | 0.248329092 | 0.57988201 | 0.42824072 | 0.66847587 | 0.97883902 | 172.810992 | 135.155949 | 73.9300305 | 157.246991 | Non-sig. |
| PF3D7_0309000 | YVH1 | 745.816036 | 0.247729036 | 0.44613555 | 0.55527751 | 0.57870486 | 0.9512809 | 558.810206 | 875.61208 | 804.835494 | 744.006366 | Non-sig. |
| PF3D7_1235200 | VP2 | 4623.13226 | 0.247584696 | 0.57944556 | 0.42727861 | 0.66917641 | 0.97883902 | 1904.54672 | 4653.63146 | 6549.84772 | 5384.50316 | Non-sig. |
| PF3D7_0319700 | ABCI3 | 2734.84579 | 0.247577484 | 0.41632127 | 0.59467893 | 0.55205811 | 0.94170064 | 2389.01442 | 2630.85556 | 2612.43659 | 3307.07658 | Non-sig. |
| PF3D7_0917500 | N/A | 3526.26924 | 0.247534681 | 0.40617039 | 0.60943556 | 0.54223577 | 0.93953127 | 3157.14905 | 4046.09666 | 3291.81005 | 3610.02121 | Non-sig. |
| PF3D7_1362500 | EXO | 371.967957 | 0.247413446 | 0.45654788 | 0.54192223 | 0.58787208 | 0.95246063 | 284.226377 | 381.974864 | 395.868501 | 425.802088 | Non-sig. |
| PF3D7_0913900 | RRSapi | 263.138961 | 0.246707203 | 0.48533312 | 0.5083255 | 0.61122509 | 0.9597882 | 205.266695 | 232.835517 | 275.956267 | 338.497363 | Non-sig. |
| PF3D7_1009300 | N/A | 398.161193 | 0.246170647 | 0.43429211 | 0.56683195 | 0.57082836 | 0.94850402 | 375.483067 | 429.290447 | 353.10686 | 434.764398 | Non-sig. |
| PF3D7_1032100 | DCP1 | 2564.06581 | 0.24590324 | 0.43313307 | 0.5677314 | 0.57021737 | 0.94850402 | 2511.27239 | 2280.42597 | 2180.97286 | 3283.592 | Non-sig. |
| PF3D7_1033000 | HPR1 | 418.54718 | 0.245265866 | 0.4901094 | 0.50043085 | 0.61677173 | 0.96358724 | 491.310832 | 502.231674 | 275.107891 | 405.538321 | Non-sig. |
| PF3D7_0110500 | BDP3 | 2539.96067 | 0.245160302 | 0.53131285 | 0.46142363 | 0.6444947 | 0.97190751 | 3266.337 | 2146.24944 | 1383.37507 | 3363.88118 | Non-sig. |
| PF3D7_1120400 | N/A | 326.545575 | 0.244878973 | 0.50001158 | 0.4897466 | 0.62431322 | 0.96488082 | 207.364531 | 366.961075 | 390.114293 | 341.742403 | Non-sig. |
| PF3D7_0113800 | N/A | 1048.40118 | 0.243504968 | 0.43120995 | 0.56470164 | 0.57227672 | 0.94850402 | 965.674331 | 958.83447 | 954.598292 | 1314.49762 | Non-sig. |
| PF3D7_0512600 | RAB1b | 2893.71233 | 0.243376305 | 0.46985645 | 0.51798013 | 0.60447213 | 0.95693578 | 1835.99269 | 3177.45676 | 3464.07172 | 3097.32812 | Non-sig. |
| PF3D7_0416500 | MAF1 | 1570.50799 | 0.24332671 | 0.43072832 | 0.56491922 | 0.57212871 | 0.94850402 | 1685.52725 | 1708.70857 | 1191.47272 | 1696.32344 | Non-sig. |
| PF3D7_1116700 | DPAP1 | 6444.60164 | 0.243242297 | 0.42525725 | 0.57198859 | 0.5673297 | 0.94825913 | 5220.19154 | 6065.87732 | 6584.83555 | 7907.50217 | Non-sig. |
| PF3D7_0612300 | TMEM234 | 95.635597 | 0.242438605 | 0.55320451 | 0.43824409 | 0.66120935 | 0.97783983 | 70.9336946 | 116.175505 | 104.152636 | 91.2805526 | Non-sig. |
| PF3D7_1111300 | BOS1 | 320.996655 | 0.241915905 | 0.55561123 | 0.435405 | 0.66326848 | 0.97883902 | 418.659341 | 291.907002 | 169.987809 | 403.43247 | Non-sig. |

| | | | | | | | | | | | |
|---|---|---|---|---|---|---|---|---|---|---|---|
| PF3D7_0623400 | Mei2 | 94.9082912 | 0.241886383 | 0.5390189 | 0.44875307 | 0.6536098 | 0.97317009 | 74.4945462 | 102.026128 | 99.3201564 | 103.792334 Non-sig. |
| PF3D7_0516700 | N/A | 8861.32913 | 0.241300381 | 0.55236119 | 0.43685253 | 0.6622183 | 0.97864228 | 4282.47009 | 7754.23842 | 11961.1204 | 11447.4876 Non-sig. |
| PF3D7_1248500 | N/A | 10787.0408 | 0.241288677 | 0.48707177 | 0.4953863 | 0.62032747 | 0.9639908 | 6344.61592 | 11847.6315 | 13429.1599 | 11526.7561 Non-sig. |
| PF3D7_1008400 | RPT2 | 10264.466 | 0.240290194 | 0.40342136 | 0.59563081 | 0.55142189 | 0.94170064 | 9620.1835 | 11729.4028 | 9203.12101 | 10505.1567 Non-sig. |
| PF3D7_1203300 | N/A | 1619.86459 | 0.240109913 | 0.41104793 | 0.58414092 | 0.55912551 | 0.94399202 | 1571.24306 | 1710.11043 | 1399.59978 | 1798.50509 Non-sig. |
| PF3D7_0715300 | N/A | 648.887997 | 0.24005724 | 0.45889084 | 0.52312494 | 0.60088731 | 0.95693578 | 465.36958 | 641.600712 | 724.411959 | 764.169737 Non-sig. |
| PF3D7_0925800 | UPF2 | 1501.24338 | 0.239953959 | 0.42610583 | 0.56313231 | 0.57334479 | 0.94850402 | 1579.7139 | 1575.69637 | 1173.8678 | 1675.69544 Non-sig. |
| PF3D7_0828300 | N/A | 1763.5601 | 0.238959636 | 0.55606442 | 0.42973373 | 0.66738935 | 0.97883902 | 2427.08353 | 1966.59714 | 809.021202 | 1851.53853 Non-sig. |
| PF3D7_1200200 | RIF | 33.3385613 | 0.238826581 | 0.70738062 | 0.33762104 | 0.73564878 | 0.98954916 | 28.3053061 | 37.5973614 | 32.8282734 | 34.6233044 Non-sig. |
| PF3D7_0202700 | OPP | 391.84556 | 0.238159459 | 0.48565253 | 0.49039065 | 0.62385749 | 0.96488082 | 258.925533 | 439.932211 | 459.918949 | 408.605546 Non-sig. |
| PF3D7_1448300 | N/A | 6536.89224 | 0.237862727 | 0.42118861 | 0.5647416 | 0.57224953 | 0.94850402 | 5335.94632 | 6268.0039 | 6662.34924 | 7881.2695 Non-sig. |
| PF3D7_1114900 | N/A | 4319.16301 | 0.237790882 | 0.40810049 | 0.58267728 | 0.56011057 | 0.94472957 | 4049.79508 | 4298.83327 | 3878.27116 | 5049.75254 Non-sig. |
| PF3D7_0612400 | N/A | 375.986619 | 0.237773212 | 0.46128243 | 0.51546124 | 0.60623075 | 0.95706907 | 287.034086 | 446.964114 | 402.914272 | 367.034006 Non-sig. |
| PF3D7_0503200 | CDC6 | 1800.92475 | 0.236456953 | 0.49613913 | 0.47659404 | 0.63365123 | 0.9668494 | 2258.85362 | 1857.10637 | 1048.87268 | 2038.86632 Non-sig. |
| PF3D7_0727700 | N/A | 460.78683 | 0.236448353 | 0.44917422 | 0.52640677 | 0.59860561 | 0.95655566 | 366.392877 | 447.406144 | 479.654459 | 549.693841 Non-sig. |
| PF3D7_1303400 | LisH | 5170.22427 | 0.235904566 | 0.52047391 | 0.45324955 | 0.65036905 | 0.97317009 | 6780.10655 | 6023.43904 | 2717.22266 | 5160.12883 Non-sig. |
| PF3D7_1012000 | N/A | 3669.20904 | 0.235830835 | 0.46123553 | 0.51130241 | 0.60913932 | 0.95866549 | 4285.23745 | 4337.46536 | 2455.06163 | 3599.07173 Non-sig. |
| PF3D7_0906400 | N/A | 134.944894 | 0.234510293 | 0.6287324 | 0.37298904 | 0.7091566 | 0.98741437 | 188.391705 | 156.661936 | 59.9529715 | 134.772965 Non-sig. |
| PF3D7_1417200 | NOT1 | 9162.58129 | 0.234234205 | 0.44163556 | 0.53037895 | 0.59584922 | 0.95516163 | 8576.06443 | 7596.16883 | 8264.76002 | 12213.3319 Non-sig. |
| PF3D7_0215100 | N/A | 1147.2004 | 0.233986834 | 0.47122959 | 0.49654529 | 0.61950974 | 0.96395246 | 744.482548 | 1269.7094 | 1363.92238 | 1210.68728 Non-sig. |
| PF3D7_1139300 | AP2-G5 | 3414.71154 | 0.233902119 | 0.45216037 | 0.51729903 | 0.60494743 | 0.95693578 | 2630.14326 | 2933.39722 | 3646.65842 | 4448.64727 Non-sig. |
| PF3D7_0937200 | N/A | 32.8590818 | 0.233048635 | 0.71072575 | 0.32790234 | 0.7429855 | 0.98977551 | 28.3709497 | 37.5729304 | 32.0206412 | 33.4718059 Non-sig. |
| PF3D7_1223400 | N/A | 18666.2365 | 0.232486003 | 0.40458545 | 0.57462768 | 0.56554313 | 0.94698087 | 17562.1949 | 18716.8357 | 16768.8118 | 21617.1036 Non-sig. |
| PF3D7_1423700 | N/A | 6199.85987 | 0.232456876 | 0.49071509 | 0.47371047 | 0.63570638 | 0.96697457 | 7764.95476 | 6329.30799 | 3638.35505 | 7066.82167 Non-sig. |
| PF3D7_0806200 | DPY19 | 1422.89586 | 0.231888215 | 0.57841159 | 0.4009052 | 0.68848992 | 0.98334178 | 599.12691 | 1512.77412 | 2018.02976 | 1561.65264 Non-sig. |
| PF3D7_1223100 | PKAr | 22966.5863 | 0.231196189 | 0.40114495 | 0.57634077 | 0.56438487 | 0.94661461 | 20456.13 | 25446.86 | 21804.4021 | 24158.9531 Non-sig. |
| PF3D7_1477300 | Pfg14-744 | 273.362639 | 0.231188837 | 0.46457963 | 0.49763016 | 0.61874474 | 0.96395246 | 271.51571 | 334.791673 | 231.596028 | 255.547148 Non-sig. |
| PF3D7_1200600 | VAR2CSA | 971.582086 | 0.231021955 | 0.41911616 | 0.55121223 | 0.5814882 | 0.95129458 | 961.56914 | 1087.50148 | 826.458938 | 1010.79879 Non-sig. |
| PF3D7_1238200 | N/A | 4.50723049 | 0.23077889 | 1.70520173 | 0.13533818 | 0.8923445 | 0.99457954 | 4.19075137 | 1.07439901 | 4.10307873 | 8.66069286 Non-sig. |
| PF3D7_0905200 | N/A | 450.872897 | 0.230648024 | 0.55172598 | 0.41804814 | 0.67591192 | 0.97898416 | 226.507282 | 452.68976 | 602.9202 | 521.374345 Non-sig. |
| PF3D7_0105700 | N/A | 3437.58018 | 0.230250631 | 0.47691245 | 0.48279434 | 0.62924177 | 0.96682409 | 3588.8822 | 2592.22723 | 2738.96265 | 4830.24864 Non-sig. |
| PF3D7_1302700 | N/A | 974.315543 | 0.230209108 | 0.47473737 | 0.48491887 | 0.6277339 | 0.96655174 | 1165.43114 | 1060.8642 | 628.406262 | 1042.56057 Non-sig. |
| PF3D7_0926200 | N/A | 946.200162 | 0.229854134 | 0.56308035 | 0.40820841 | 0.68312068 | 0.98107166 | 1311.3342 | 1012.3295 | 431.045122 | 1030.09183 Non-sig. |
| PF3D7_1252800 | N/A | 1867.53922 | 0.229517913 | 0.41571417 | 0.55210509 | 0.58087635 | 0.95129458 | 1856.75702 | 1874.22102 | 1581.95994 | 2157.21889 Non-sig. |
| PF3D7_1020800 | DLAT | 131.728299 | 0.229053328 | 0.65655728 | 0.34887029 | 0.72718669 | 0.98844303 | 190.373781 | 157.763584 | 52.5555757 | 126.220254 Non-sig. |
| PF3D7_1469600 | ACC | 1013.78572 | 0.228834228 | 0.51708848 | 0.44254366 | 0.65809585 | 0.97671961 | 592.55166 | 881.581623 | 1274.19481 | 1306.81478 Non-sig. |
| PF3D7_1324500 | N/A | 986.381963 | 0.228242779 | 0.41954666 | 0.54402239 | 0.58642606 | 0.95246063 | 832.077323 | 1094.25502 | 984.836834 | 1034.35868 Non-sig. |
| PF3D7_1034400 | SDHA | 1103.03181 | 0.227567883 | 0.64102538 | 0.35500604 | 0.72258507 | 0.98844303 | 355.773054 | 1199.60714 | 1676.2515 | 1180.49555 Non-sig. |
| PF3D7_0207200 | ISCA1 | 291.022838 | 0.226148343 | 0.48918596 | 0.46229525 | 0.64386961 | 0.97190751 | 303.512857 | 385.448452 | 233.154159 | 241.975884 Non-sig. |
| PF3D7_1426700 | PEPC | 6962.20068 | 0.225717359 | 0.46043856 | 0.49022254 | 0.62397643 | 0.96488082 | 4760.37835 | 6519.39554 | 8076.65193 | 8492.37689 Non-sig. |
| PF3D7_0818900 | HSP70 | 43727.7037 | 0.225326123 | 0.43805089 | 0.51438344 | 0.60698394 | 0.95715771 | 31220.4466 | 47293.3252 | 49418.9721 | 46978.071 Non-sig. |
| PF3D7_0617400 | VAR | 759.949742 | 0.223993772 | 0.449103 | 0.49875813 | 0.61794979 | 0.96395246 | 830.626398 | 900.950846 | 571.760403 | 736.461321 Non-sig. |
| PF3D7_1139100 | N/A | 1195.06594 | 0.223502148 | 0.4230617 | 0.52829682 | 0.59729334 | 0.95631916 | 987.741971 | 1199.10369 | 1217.46724 | 1375.95084 Non-sig. |
| PF3D7_0112800 | N/A | 167.874127 | 0.223294185 | 0.52309219 | 0.42687348 | 0.66947148 | 0.97883902 | 186.028204 | 217.909546 | 123.982138 | 143.576619 Non-sig. |
| PF3D7_0709200 | GLP3 | 297.152036 | 0.223241855 | 0.45775639 | 0.48768703 | 0.62577154 | 0.96637958 | 261.962555 | 362.056745 | 286.402761 | 278.186082 Non-sig. |
| PF3D7_0520400 | N/A | 1248.18929 | 0.222742178 | 0.42302779 | 0.52654267 | 0.59851122 | 0.95655566 | 1295.09469 | 1380.37543 | 1009.13444 | 1308.1526 Non-sig. |
| PF3D7_1321700 | SF1 | 4073.42399 | 0.222606415 | 0.53405206 | 0.41682531 | 0.67680619 | 0.97898416 | 5480.41162 | 4688.88168 | 2039.57811 | 4084.82457 Non-sig. |
| PF3D7_1025200 | ATP12 | 410.448587 | 0.222537861 | 0.46698963 | 0.47653705 | 0.63369182 | 0.9668494 | 299.351698 | 404.75948 | 458.109357 | 479.573814 Non-sig. |
| PF3D7_0511400 | N/A | 74.651097 | 0.221889754 | 0.71131666 | 0.3119423 | 0.75508438 | 0.99028237 | 106.937397 | 100.974119 | 31.1871817 | 59.5056907 Non-sig. |
| PF3D7_1233700 | N/A | 769.093817 | 0.220840514 | 0.48618857 | 0.45422811 | 0.64966466 | 0.97317009 | 933.225494 | 893.915843 | 487.807869 | 761.426061 Non-sig. |
| PF3D7_1361200 | N/A | 3734.19506 | 0.220719497 | 0.59599102 | 0.3703403 | 0.71112895 | 0.9875928 | 5457.79577 | 4056.70844 | 1440.83949 | 3981.43653 Non-sig. |
| PF3D7_0418800 | N/A | 11.0865795 | 0.220395876 | 1.13900637 | 0.19349837 | 0.84656868 | 0.99165628 | 15.6637964 | 12.9045608 | 4.92145302 | 10.8565079 Non-sig. |
| PF3D7_0100100 | VAR | 1839.47923 | 0.220366615 | 0.47377869 | 0.46512563 | 0.64184149 | 0.97073258 | 2151.02967 | 2314.69989 | 1247.81259 | 1644.37475 Non-sig. |
| PF3D7_1108000 | N/A | 2439.56224 | 0.220255835 | 0.52195747 | 0.42198043 | 0.6730393 | 0.97898416 | 3221.75653 | 2492.83731 | 1286.05165 | 2757.60347 Non-sig. |

| | | | | | | | | | | | | |
|---|---|---|---|---|---|---|---|---|---|---|---|---|
| PF3D7_0624600 | ISWI | 35200.3782 | 0.219145917 | 0.42346522 | 0.5175063 | 0.60480277 | 0.95693578 | 32225.2498 | 31285.7883 | 32838.8279 | 44451.6467 | Non-sig. |
| PF3D7_1463900 | RON11 | 9345.70106 | 0.218999305 | 0.42010719 | 0.52129387 | 0.60216207 | 0.95693578 | 9381.74687 | 8737.30703 | 7893.83822 | 11369.9121 | Non-sig. |
| PF3D7_1355100 | MCM6 | 9701.17919 | 0.218225387 | 0.6194699 | 0.35227763 | 0.72463007 | 0.98844303 | 3355.03179 | 10524.8861 | 14582.2882 | 10342.5107 | Non-sig. |
| PF3D7_0722600 | UTP7 | 2167.05582 | 0.218193063 | 0.45567694 | 0.4788328 | 0.63205758 | 0.9668494 | 2530.62008 | 2376.62578 | 1476.69966 | 2284.27774 | Non-sig. |
| PF3D7_1106200 | N/A | 271.680291 | 0.217468874 | 0.46584423 | 0.46682745 | 0.64062333 | 0.97002976 | 227.258319 | 254.330894 | 275.110838 | 330.021112 | Non-sig. |
| PF3D7_1446100 | N/A | 160.888069 | 0.216985634 | 0.62944298 | 0.34472643 | 0.73030004 | 0.98844303 | 228.171406 | 149.190081 | 69.7967949 | 196.393993 | Non-sig. |
| PF3D7_1360700 | PIAS | 2390.41529 | 0.216386824 | 0.52186226 | 0.41464356 | 0.67840286 | 0.97898416 | 1285.49347 | 2274.07824 | 3137.07565 | 2865.01379 | Non-sig. |
| PF3D7_0922400 | pBAS | 231.043341 | 0.21638496 | 0.55724957 | 0.38830889 | 0.69778746 | 0.98334178 | 127.38171 | 213.51674 | 299.778483 | 283.496433 | Non-sig. |
| PF3D7_1027600 | SPO11-2 | 44.0744927 | 0.216347412 | 0.64847407 | 0.33362539 | 0.73866226 | 0.98954916 | 38.8382668 | 51.5436498 | 42.6863703 | 43.229684 | Non-sig. |
| PF3D7_1231100 | RAB2 | 4412.59413 | 0.215209354 | 0.41680802 | 0.51632729 | 0.60562584 | 0.95693578 | 4154.77217 | 5371.47229 | 4013.43269 | 4110.69936 | Non-sig. |
| PF3D7_1200800 | FIKK12 | 311.5131 | 0.21359054 | 0.44999742 | 0.47464837 | 0.63503763 | 0.9668494 | 308.501642 | 308.995505 | 268.576028 | 359.979225 | Non-sig. |
| PF3D7_1218600 | RRS | 751.396882 | 0.213479826 | 0.4968802 | 0.42964044 | 0.66745721 | 0.97883902 | 452.684539 | 822.94852 | 938.760448 | 791.194019 | Non-sig. |
| PF3D7_1034700 | N/A | 1478.66627 | 0.213479691 | 0.47505307 | 0.44938072 | 0.65315703 | 0.97317009 | 945.491785 | 1644.72174 | 1793.08708 | 1531.36447 | Non-sig. |
| PF3D7_1147300 | N/A | 6622.1262 | 0.213376223 | 0.41073725 | 0.51949568 | 0.60341513 | 0.95693578 | 6774.44431 | 6930.18081 | 5492.33462 | 7291.54506 | Non-sig. |
| PF3D7_1476800 | N/A | 478.94653 | 0.213235123 | 0.44017737 | 0.48443 | 0.62808073 | 0.96655174 | 386.311675 | 518.647285 | 500.75044 | 510.076718 | Non-sig. |
| PF3D7_1145600 | N/A | 682.698586 | 0.21311931 | 0.48238531 | 0.44180307 | 0.65863172 | 0.97688434 | 805.967946 | 838.099049 | 459.069795 | 627.657554 | Non-sig. |
| PF3D7_1464300 | N/A | 22.5491441 | 0.212098843 | 0.81752835 | 0.25943913 | 0.79529644 | 0.99028237 | 22.1018598 | 27.8986868 | 19.7105722 | 20.4854578 | Non-sig. |
| PF3D7_1337700 | PH2 | 2398.69718 | 0.21148797 | 0.55000581 | 0.38451952 | 0.70059343 | 0.98491338 | 3279.7565 | 2940.02643 | 1167.04639 | 2207.95941 | Non-sig. |
| PF3D7_1332600 | APN1 | 4189.31 | 0.211288119 | 0.51016632 | 0.41415537 | 0.67876033 | 0.97898416 | 5459.44786 | 4756.13927 | 2307.13495 | 4234.51794 | Non-sig. |
| PF3D7_0512900 | AKAL | 57.7491516 | 0.211135879 | 0.66858779 | 0.3157938 | 0.75215902 | 0.99028237 | 31.3584012 | 64.5054641 | 75.4732442 | 59.659497 | Non-sig. |
| PF3D7_1236100 | CARP | 1657.43513 | 0.210997341 | 0.466008 | 0.45277622 | 0.65070989 | 0.97317009 | 1395.71363 | 1289.7416 | 1677.07039 | 2267.21488 | Non-sig. |
| PF3D7_1423000 | NOG2 | 992.403133 | 0.210674632 | 0.42994701 | 0.49000138 | 0.62413292 | 0.96488082 | 1056.10832 | 1068.89681 | 784.242339 | 1060.36507 | Non-sig. |
| PF3D7_0518700 | PUF1 | 281.252977 | 0.210567329 | 0.55795232 | 0.37739305 | 0.70588153 | 0.98615891 | 369.904536 | 253.140205 | 151.970638 | 349.996529 | Non-sig. |
| PF3D7_0526400 | N/A | 688.854625 | 0.210435563 | 0.44243306 | 0.47563254 | 0.63433619 | 0.9668494 | 527.552982 | 712.788071 | 749.627265 | 765.450181 | Non-sig. |
| PF3D7_1106600 | N/A | 514.832043 | 0.20983326 | 0.46428203 | 0.45195215 | 0.65130346 | 0.97317009 | 584.723294 | 510.612349 | 370.461179 | 593.531349 | Non-sig. |
| PF3D7_0820700 | KDH | 3961.74424 | 0.209628808 | 0.51907235 | 0.40385277 | 0.68632099 | 0.98210107 | 2098.14348 | 4295.46158 | 5250.29227 | 4203.07962 | Non-sig. |
| PF3D7_1007100 | N/A | 22.4829208 | 0.209506033 | 0.83068669 | 0.25220824 | 0.80088011 | 0.99028237 | 24.6319384 | 19.4934502 | 17.1308512 | 28.6754436 | Non-sig. |
| PF3D7_0815800 | VPS9 | 15647.6252 | 0.209387606 | 0.58411455 | 0.35847011 | 0.71999153 | 0.98797939 | 22690.1536 | 16114.8385 | 6338.48666 | 17447.0219 | Non-sig. |
| PF3D7_1244400 | N/A | 3264.18789 | 0.209342207 | 0.40511459 | 0.51674813 | 0.60533199 | 0.95693578 | 3100.74415 | 3369.03553 | 2954.85031 | 3632.12155 | Non-sig. |
| PF3D7_0507300 | SOPT | 1191.63278 | 0.209178766 | 0.58098922 | 0.36003898 | 0.71881799 | 0.98797939 | 1698.67209 | 1136.1394 | 512.529629 | 1419.19001 | Non-sig. |
| PF3D7_0508200 | N/A | 552.323829 | 0.209011162 | 0.50115846 | 0.41705604 | 0.67663742 | 0.97898416 | 356.771168 | 697.475967 | 667.690398 | 487.357783 | Non-sig. |
| PF3D7_0514200 | N/A | 1214.98791 | 0.208647943 | 0.42790428 | 0.48760425 | 0.62583019 | 0.96637958 | 959.129421 | 1314.33022 | 1295.22665 | 1291.26536 | Non-sig. |
| PF3D7_1468400 | D13 | 4746.73651 | 0.207927039 | 0.47356749 | 0.43906527 | 0.66061424 | 0.97783983 | 5820.52424 | 5048.24266 | 2990.38558 | 5127.79357 | Non-sig. |
| PF3D7_1354700 | N/A | 21.1285897 | 0.207898269 | 0.83620119 | 0.24862231 | 0.80365295 | 0.99028237 | 22.0267295 | 24.7032227 | 17.2364336 | 20.5479728 | Non-sig. |
| PF3D7_0623500 | SOD2 | 375.675605 | 0.207155844 | 0.44632929 | 0.46413231 | 0.64255296 | 0.97144917 | 313.314311 | 430.812563 | 384.060258 | 374.515289 | Non-sig. |
| PF3D7_0929500 | N/A | 8.11321436 | 0.206966989 | 1.39823029 | 0.14802067 | 0.88232645 | 0.99457954 | 13.527281 | 11.8406513 | 1.63953632 | 5.44538878 | Non-sig. |
| PF3D7_1308100 | N/A | 488.695879 | 0.206894617 | 0.44620946 | 0.46367151 | 0.64288311 | 0.97144917 | 501.291032 | 588.607238 | 406.276011 | 458.609237 | Non-sig. |
| PF3D7_0917300 | N/A | 1490.08075 | 0.206498841 | 0.45545539 | 0.45338983 | 0.65026806 | 0.97317009 | 1031.78192 | 1642.0394 | 1735.14842 | 1551.35327 | Non-sig. |
| PF3D7_0508700 | PRP5 | 3030.06792 | 0.206460378 | 0.52594463 | 0.39255155 | 0.69465073 | 0.98334178 | 4045.60592 | 3016.48453 | 1582.05495 | 3476.12629 | Non-sig. |
| PF3D7_1445300 | N/A | 517.556748 | 0.205543581 | 0.4590935 | 0.44771616 | 0.65435805 | 0.97327035 | 379.817765 | 589.145472 | 581.428755 | 519.835001 | Non-sig. |
| PF3D7_1012100 | N/A | 1528.46827 | 0.205412416 | 0.44585583 | 0.46071489 | 0.64500317 | 0.97190751 | 1735.477 | 1705.3422 | 1104.4911 | 1568.5628 | Non-sig. |
| PF3D7_0917400 | N/A | 1295.99099 | 0.204813787 | 0.49475878 | 0.41396695 | 0.67889831 | 0.97898416 | 1636.51905 | 1360.12133 | 772.164938 | 1415.15866 | Non-sig. |
| PF3D7_0909600 | N/A | 2144.91626 | 0.204759972 | 0.40661749 | 0.50356902 | 0.61456427 | 0.96217474 | 1944.14916 | 2246.83994 | 2041.73045 | 2346.94549 | Non-sig. |
| PF3D7_0514900 | N/A | 588.931015 | 0.204674622 | 0.42404998 | 0.48266627 | 0.62933271 | 0.96682409 | 565.687759 | 646.135501 | 528.811874 | 615.088925 | Non-sig. |
| PF3D7_1123100 | CDPK7 | 10780.4521 | 0.204247609 | 0.59161659 | 0.34523644 | 0.72991662 | 0.98844303 | 15692.7954 | 9968.47818 | 4344.85908 | 13115.6757 | Non-sig. |
| PF3D7_1345200 | ROM6 | 758.953885 | 0.20408552 | 0.4287675 | 0.47598179 | 0.63408735 | 0.9668494 | 773.554617 | 872.373799 | 637.306991 | 752.580134 | Non-sig. |
| PF3D7_0813100 | N/A | 4611.67911 | 0.203959949 | 0.49101287 | 0.41538616 | 0.67785924 | 0.97898416 | 5836.10545 | 5193.68938 | 2736.75638 | 4680.16523 | Non-sig. |
| PF3D7_1120200 | N/A | 1268.03464 | 0.203930997 | 0.46089631 | 0.44246611 | 0.65815195 | 0.97671961 | 866.923791 | 1297.22983 | 1489.88388 | 1418.10104 | Non-sig. |
| PF3D7_1365700 | N/A | 1844.3801 | 0.203363242 | 0.47611254 | 0.42713272 | 0.66928266 | 0.97883902 | 1198.14147 | 1761.73096 | 2230.72002 | 2186.92795 | Non-sig. |
| PF3D7_1407300 | PRP38B | 3443.93565 | 0.203067968 | 0.42368087 | 0.47929464 | 0.63172904 | 0.9668494 | 3690.45108 | 3880.67919 | 2713.68733 | 3490.92498 | Non-sig. |
| PF3D7_0510900 | N/A | 7.53748396 | 0.202744906 | 1.34223361 | 0.15105039 | 0.87993597 | 0.99421828 | 8.40103577 | 3.32087441 | 5.65918226 | 12.7688434 | Non-sig. |
| PF3D7_1110700 | ALP1 | 1504.66388 | 0.202455137 | 0.43118503 | 0.46953192 | 0.63868947 | 0.96956356 | 1158.52365 | 1598.23935 | 1639.76235 | 1622.13018 | Non-sig. |
| PF3D7_1218400 | N/A | 2981.79286 | 0.202408167 | 0.41308669 | 0.48998956 | 0.62414129 | 0.96488082 | 2477.30749 | 3172.63395 | 3068.46234 | 3208.76768 | Non-sig. |
| PF3D7_1337400 | N/A | 831.640144 | 0.20175329 | 0.46519598 | 0.43369525 | 0.66450975 | 0.97883902 | 977.540386 | 926.076095 | 569.943268 | 853.000828 | Non-sig. |

| | | | | | | | | | | | | |
|---|---|---|---|---|---|---|---|---|---|---|---|---|
| PF3D7_1404400 | N/A | 276.859444 | 0.201670904 | 0.45762338 | 0.44069187 | 0.65943608 | 0.97707593 | 258.159168 | 331.834901 | 256.928091 | 260.515616 | Non-sig. |
| PF3D7_1019800 | N/A | 2048.17171 | 0.201399316 | 0.59108616 | 0.34072751 | 0.73330873 | 0.98909017 | 2966.97009 | 1874.37963 | 844.324754 | 2507.01235 | Non-sig. |
| PF3D7_1251700 | aTrpRS | 218.172419 | 0.201176075 | 0.48219525 | 0.41720875 | 0.67652573 | 0.97898416 | 165.154431 | 238.245642 | 240.612012 | 228.677591 | Non-sig. |
| PF3D7_1106300 | XRN1 | 2542.09038 | 0.200653608 | 0.41738709 | 0.48073746 | 0.63070311 | 0.9668494 | 2549.88536 | 2988.11337 | 2181.42355 | 2448.93925 | Non-sig. |
| PF3D7_1414200 | N/A | 4487.92257 | 0.200208391 | 0.41987811 | 0.47682503 | 0.63348672 | 0.9668494 | 4717.19706 | 4443.63049 | 3637.02968 | 5153.83307 | Non-sig. |
| PF3D7_0824000 | N/A | 169.13213 | 0.200066045 | 0.69036889 | 0.28979586 | 0.77197242 | 0.99028237 | 260.999761 | 198.512599 | 54.2036131 | 162.812547 | Non-sig. |
| PF3D7_1236000 | N/A | 905.599507 | 0.199463672 | 0.428275 | 0.46573737 | 0.6414035 | 0.97034716 | 940.800469 | 1032.60902 | 745.555757 | 903.432788 | Non-sig. |
| PF3D7_1209300 | TRZ | 12707.5743 | 0.199410382 | 0.48688341 | 0.40956495 | 0.68212511 | 0.98088141 | 16070.2011 | 13470.8312 | 7591.6922 | 13697.5727 | Non-sig. |
| PF3D7_0515700 | GAP40 | 30236.9197 | 0.199406969 | 0.50116067 | 0.3978903 | 0.69071105 | 0.98334178 | 17098.5002 | 33338.3093 | 39202.3183 | 31308.5512 | Non-sig. |
| PF3D7_1208900 | PPM11 | 15702.4144 | 0.198790049 | 0.59606697 | 0.33350288 | 0.73875472 | 0.98954916 | 6270.07589 | 14107.1844 | 22974.1299 | 19458.2675 | Non-sig. |
| PF3D7_0809700 | RUVB1 | 152.756436 | 0.198461283 | 0.5329593 | 0.37237606 | 0.70961288 | 0.9875928 | 183.814174 | 172.854388 | 100.981262 | 153.375921 | Non-sig. |
| PF3D7_1429900 | WRN | 1767.47349 | 0.198254204 | 0.4730223 | 0.41912232 | 0.67512674 | 0.97898416 | 1318.26451 | 1433.06564 | 1973.94586 | 2344.61795 | Non-sig. |
| PF3D7_1142700 | N/A | 486.506156 | 0.197883049 | 0.52382087 | 0.37776855 | 0.70560254 | 0.98615891 | 273.63787 | 541.880681 | 632.379158 | 498.126915 | Non-sig. |
| PF3D7_1111000 | NSUN2 | 311.544224 | 0.196948374 | 0.51591095 | 0.38174878 | 0.70264771 | 0.98512564 | 189.385242 | 339.106192 | 390.904954 | 326.780507 | Non-sig. |
| PF3D7_1324600 | N/A | 585.137525 | 0.195197876 | 0.47682475 | 0.40937026 | 0.68226796 | 0.98088141 | 699.713229 | 606.046814 | 391.83301 | 642.957046 | Non-sig. |
| PF3D7_1250300 | VPS26 | 1384.42857 | 0.194801905 | 0.41557404 | 0.46875378 | 0.63924564 | 0.96967357 | 1312.82844 | 1349.28674 | 1269.40057 | 1606.19852 | Non-sig. |
| PF3D7_0526300 | WDR55 | 554.000044 | 0.194297292 | 0.49260156 | 0.39443093 | 0.69326291 | 0.98334178 | 350.019042 | 617.214736 | 683.144944 | 565.621456 | Non-sig. |
| PF3D7_1011700 | RAD23 | 1844.29566 | 0.193939286 | 0.40886332 | 0.4743377 | 0.63525911 | 0.9668494 | 1639.19813 | 2002.45601 | 1801.7692 | 1933.75928 | Non-sig. |
| PF3D7_0923500 | N/A | 679.869018 | 0.193470494 | 0.43032701 | 0.44958947 | 0.65300648 | 0.97317009 | 685.636529 | 661.927147 | 583.175575 | 788.736822 | Non-sig. |
| PF3D7_0405700 | N/A | 1443.88062 | 0.193446311 | 0.45732957 | 0.42299104 | 0.6723018 | 0.97898416 | 1010.8901 | 1421.13733 | 1683.24684 | 1660.2482 | Non-sig. |
| PF3D7_0936600 | GEXP5 | 452.91222 | 0.193062795 | 0.4624719 | 0.41745843 | 0.67634312 | 0.97898416 | 331.058693 | 499.155987 | 513.989873 | 467.444327 | Non-sig. |
| PF3D7_1414800 | SMB1 | 609.57805 | 0.192501155 | 0.49762124 | 0.38684272 | 0.69887265 | 0.98377748 | 374.033414 | 635.53971 | 763.553781 | 665.185295 | Non-sig. |
| PF3D7_0523600 | N/A | 447.708171 | 0.192352984 | 0.44586494 | 0.43141536 | 0.66616638 | 0.97883902 | 403.317617 | 546.885127 | 432.456475 | 408.173465 | Non-sig. |
| PF3D7_1204100 | N/A | 523.964607 | 0.19162579 | 0.64330093 | 0.29787892 | 0.76579557 | 0.99028237 | 801.419378 | 585.77159 | 177.414102 | 531.253358 | Non-sig. |
| PF3D7_0403600 | N/A | 1458.85979 | 0.191493407 | 0.45145078 | 0.42417339 | 0.67143937 | 0.97898416 | 1047.71169 | 1435.51478 | 1676.36095 | 1675.85176 | Non-sig. |
| PF3D7_1133800 | N/A | 5264.9797 | 0.191414736 | 0.4021004 | 0.47603717 | 0.6340479 | 0.9668494 | 4920.77025 | 5716.24109 | 4911.66752 | 5511.23992 | Non-sig. |
| PF3D7_1311800 | M1AAP | 7746.24853 | 0.190623679 | 0.58354401 | 0.32666547 | 0.74392092 | 0.98998709 | 3192.57392 | 7647.18248 | 11277.4834 | 8867.75428 | Non-sig. |
| PF3D7_1226800 | ATX3 | 1697.20868 | 0.189935159 | 0.41432512 | 0.45842057 | 0.64665032 | 0.97317009 | 1712.77598 | 1892.69171 | 1458.64308 | 1724.72397 | Non-sig. |
| PF3D7_0932000 | N/A | 1744.91514 | 0.18927709 | 0.41275521 | 0.45856984 | 0.6465431 | 0.97317009 | 1501.28973 | 1903.10648 | 1759.82517 | 1815.43918 | Non-sig. |
| PF3D7_0103400 | N/A | 7454.84748 | 0.189127543 | 0.44054189 | 0.42930661 | 0.66770011 | 0.97883902 | 8524.02948 | 7523.57094 | 5410.08436 | 8361.70515 | Non-sig. |
| PF3D7_1423600 | N/A | 506.581288 | 0.189060747 | 0.43089489 | 0.43876303 | 0.66083325 | 0.97783983 | 481.197112 | 580.433098 | 465.699223 | 498.995721 | Non-sig. |
| PF3D7_0512500 | N/A | 621.219262 | 0.188347944 | 0.42623031 | 0.44189242 | 0.65856705 | 0.97688434 | 601.334189 | 708.073434 | 560.169637 | 615.299786 | Non-sig. |
| PF3D7_1206900 | N/A | 230.983511 | 0.188303736 | 0.50168798 | 0.37534034 | 0.70740737 | 0.9862748 | 274.473501 | 242.558873 | 157.662361 | 249.239311 | Non-sig. |
| PF3D7_1436200 | BCP1 | 12348.3936 | 0.188278427 | 0.49538406 | 0.38006557 | 0.70389674 | 0.98582247 | 15496.5481 | 10875.9117 | 7591.36999 | 15429.7445 | Non-sig. |
| PF3D7_0718000 | N/A | 243.700687 | 0.188096165 | 0.48161443 | 0.39055343 | 0.69612736 | 0.98334178 | 233.053738 | 205.907003 | 222.641526 | 313.200482 | Non-sig. |
| PF3D7_0602300 | PALM | 19.7154098 | 0.187659323 | 0.87198249 | 0.21520997 | 0.82960362 | 0.99110542 | 14.6509591 | 24.7190584 | 22.1540062 | 17.3376155 | Non-sig. |
| PF3D7_1125900 | N/A | 629.59603 | 0.187097516 | 0.48412719 | 0.38646356 | 0.69915338 | 0.98382859 | 733.716148 | 798.609168 | 444.212957 | 541.845847 | Non-sig. |
| PF3D7_1340200 | N/A | 12.1672334 | 0.18692949 | 1.03543549 | 0.18053224 | 0.85673474 | 0.99352464 | 10.4390736 | 13.9812717 | 12.3029082 | 11.94568 | Non-sig. |
| PF3D7_1009100 | N/A | 3.5514415 | 0.186927664 | 1.77387557 | 0.10537812 | 0.91607578 | 0.99619209 | 4.20134786 | 3.22091483 | 2.46294533 | 4.32055798 | Non-sig. |
| PF3D7_0822600 | SEC23 | 5633.06408 | 0.186558751 | 0.50769606 | 0.36746149 | 0.71327481 | 0.9875928 | 3155.9975 | 5618.56422 | 7382.32295 | 6375.37165 | Non-sig. |
| PF3D7_1034900 | MRScyt | 6295.14594 | 0.185823751 | 0.47559352 | 0.39071969 | 0.69600445 | 0.98334178 | 4005.42243 | 7209.81123 | 7774.79828 | 6190.55183 | Non-sig. |
| PF3D7_1116200 | PDX2 | 15.6349714 | 0.185394255 | 0.9716659 | 0.19080041 | 0.84868196 | 0.99165628 | 12.396235 | 10.6677981 | 16.8289681 | 22.6468842 | Non-sig. |
| PF3D7_1243500 | SNF7 | 1836.60144 | 0.185274493 | 0.45592085 | 0.40637425 | 0.68446764 | 0.98115715 | 2143.32662 | 2129.13936 | 1294.64457 | 1779.29522 | Non-sig. |
| PF3D7_0626000 | N/A | 896.406692 | 0.184940789 | 0.54408807 | 0.33990966 | 0.73392456 | 0.98909017 | 455.30615 | 864.517285 | 1222.35505 | 1043.44828 | Non-sig. |
| PF3D7_0603800 | CEP76 | 4622.32237 | 0.184561477 | 0.40810761 | 0.45223728 | 0.65109805 | 0.97317009 | 4315.2133 | 4500.8468 | 4338.90762 | 5334.32177 | Non-sig. |
| PF3D7_0910200 | N/A | 11111.792 | 0.184399408 | 0.49943681 | 0.36921469 | 0.71196771 | 0.9875928 | 14442.9624 | 11711.6932 | 6362.70147 | 11929.8109 | Non-sig. |
| PF3D7_1438700 | N/A | 1588.39555 | 0.184258535 | 0.43716496 | 0.42148514 | 0.67340086 | 0.97898416 | 1305.76446 | 1952.86586 | 1668.27893 | 1426.67295 | Non-sig. |
| PF3D7_1211600 | LSD1 | 2569.48799 | 0.183117484 | 0.51253571 | 0.35727751 | 0.72088406 | 0.98816139 | 3362.88111 | 2416.41411 | 1450.78388 | 3047.87284 | Non-sig. |
| PF3D7_0910900 | N/A | 1458.97401 | 0.182906689 | 0.42942994 | 0.42592906 | 0.67015954 | 0.97883902 | 1162.79 | 1657.90153 | 1570.231 | 1444.97349 | Non-sig. |
| PF3D7_0903300 | N/A | 13456.607 | 0.181675492 | 0.42553848 | 0.42693082 | 0.66942972 | 0.97883902 | 12225.7883 | 11766.0952 | 12995.0641 | 16839.4807 | Non-sig. |
| PF3D7_1414500 | ABCk2 | 13741.3159 | 0.181017067 | 0.50381617 | 0.3592919 | 0.71937674 | 0.98797939 | 17724.922 | 12512.6605 | 8036.22053 | 16691.4607 | Non-sig. |
| PF3D7_0323500 | SMN | 2702.82909 | 0.180651933 | 0.5369872 | 0.33641758 | 0.736556 | 0.98954916 | 3709.18018 | 2990.58053 | 1358.88847 | 2752.66717 | Non-sig. |
| PF3D7_0804700 | N/A | 511.06948 | 0.18049729 | 0.72395663 | 0.24932058 | 0.80311281 | 0.99028237 | 842.0063 | 520.22929 | 116.647079 | 565.39525 | Non-sig. |
| PF3D7_0524400 | RBG1 | 903.656319 | 0.18015953 | 0.5202319 | 0.34630619 | 0.72911261 | 0.98844303 | 502.715592 | 1030.31554 | 1191.51705 | 890.077095 | Non-sig. |

| | | | | | | | | | | | | |
|---|---|---|---|---|---|---|---|---|---|---|---|---|
| PF3D7_0521300 | N/A | 786.067388 | 0.179910079 | 0.56383018 | 0.31908558 | 0.74966162 | 0.99028237 | 1104.17339 | 766.982 | 370.463941 | 902.650228 | Non-sig. |
| PF3D7_1319200 | N/A | 178.38062 | 0.17923907 | 0.57923463 | 0.30944122 | 0.75698592 | 0.99028237 | 242.158361 | 183.468248 | 92.8099963 | 195.085876 | Non-sig. |
| PF3D7_1207900 | N/A | 133.334762 | 0.179029671 | 0.51189549 | 0.34973872 | 0.7265348 | 0.98844303 | 103.875217 | 147.131406 | 146.107968 | 136.224454 | Non-sig. |
| PF3D7_1019200 | N/A | 139.778669 | 0.179002822 | 0.5771803 | 0.31013328 | 0.7564596 | 0.99028237 | 182.118299 | 167.652333 | 80.4129693 | 128.931074 | Non-sig. |
| PF3D7_0627700 | N/A | 6073.33524 | 0.178220639 | 0.60225671 | 0.29592138 | 0.76729012 | 0.99028237 | 2381.1119 | 5460.46211 | 9015.84822 | 7435.91875 | Non-sig. |
| PF3D7_1000600 | RIF | 41.9779896 | 0.178090965 | 0.70729005 | 0.2517934 | 0.80120076 | 0.99028237 | 25.229013 | 49.3758505 | 53.3715955 | 39.9354993 | Non-sig. |
| PF3D7_1344600 | LipA | 435.601612 | 0.177853515 | 0.52046745 | 0.3417188 | 0.73256252 | 0.98904466 | 546.147221 | 552.931672 | 271.742518 | 371.585038 | Non-sig. |
| PF3D7_1024100 | N/A | 382.013509 | 0.177251144 | 0.72674524 | 0.24389722 | 0.80731042 | 0.99028237 | 628.808322 | 428.044775 | 88.7095014 | 382.491436 | Non-sig. |
| PF3D7_0206100 | SufE | 698.262338 | 0.176827249 | 0.46386242 | 0.38120624 | 0.70305022 | 0.98512564 | 820.192161 | 770.348803 | 491.155074 | 711.353315 | Non-sig. |
| PF3D7_1224800 | N/A | 730.260066 | 0.176546826 | 0.41908731 | 0.42126503 | 0.67356156 | 0.97898416 | 663.412733 | 773.880024 | 707.80817 | 775.939337 | Non-sig. |
| PF3D7_1238500 | N/A | 3212.12107 | 0.176242793 | 0.42550554 | 0.41419624 | 0.6787304 | 0.97898416 | 3431.72586 | 3079.42378 | 2600.79916 | 3736.53549 | Non-sig. |
| PF3D7_1226700 | RRP9 | 796.313763 | 0.175719038 | 0.46940717 | 0.37434247 | 0.70814954 | 0.98679018 | 953.991932 | 870.107737 | 542.085863 | 819.069522 | Non-sig. |
| PF3D7_0620300 | N/A | 1092.47477 | 0.175133432 | 0.42264092 | 0.41437879 | 0.67859672 | 0.97898416 | 1073.58731 | 1034.37892 | 978.969321 | 1282.96352 | Non-sig. |
| PF3D7_1407800 | PM4 | 2993.64791 | 0.174368138 | 0.43463112 | 0.4011865 | 0.68828282 | 0.98334178 | 2317.1555 | 2862.23543 | 3308.52063 | 3486.68008 | Non-sig. |
| PF3D7_1362100 | 14-3-3II | 135.53631 | 0.173411827 | 0.62749523 | 0.27635561 | 0.78227495 | 0.99028237 | 193.591971 | 149.391306 | 61.5383612 | 137.623602 | Non-sig. |
| PF3D7_1344900 | N/A | 1138.94582 | 0.173049168 | 0.42380189 | 0.40832561 | 0.68303464 | 0.98107166 | 1023.35344 | 1065.1031 | 1118.02172 | 1349.30501 | Non-sig. |
| PF3D7_0114500 | N/A | 19.2910442 | 0.172920144 | 0.89110788 | 0.19405074 | 0.84613615 | 0.99165628 | 15.7179909 | 26.8586953 | 20.5160121 | 14.0714787 | Non-sig. |
| PF3D7_0930200 | LRR8 | 6513.13536 | 0.172845336 | 0.4574608 | 0.37783638 | 0.70555214 | 0.98615891 | 7800.18507 | 7295.90172 | 4447.03854 | 6509.41611 | Non-sig. |
| PF3D7_1127000 | N/A | 337.554484 | 0.172844126 | 0.44451517 | 0.38883741 | 0.69739643 | 0.98334178 | 325.998733 | 386.37839 | 308.754747 | 329.086067 | Non-sig. |
| PF3D7_0529800 | N/A | 5445.15937 | 0.17284062 | 0.60341366 | 0.28643803 | 0.77454265 | 0.99028237 | 8173.96009 | 5612.81411 | 2065.21418 | 5928.64912 | Non-sig. |
| PF3D7_0704200 | NSUN1 | 449.572396 | 0.172405524 | 0.44122843 | 0.39073984 | 0.69598955 | 0.98334178 | 452.947952 | 426.962047 | 392.627341 | 525.752245 | Non-sig. |
| PF3D7_1426600 | N/A | 1164.99611 | 0.171782654 | 0.46846598 | 0.36669185 | 0.71384889 | 0.9875928 | 1410.61222 | 1227.21013 | 781.162507 | 1240.99958 | Non-sig. |
| PF3D7_0913700 | N/A | 1823.98638 | 0.171738295 | 0.43209992 | 0.39745042 | 0.69103534 | 0.98334178 | 2026.24769 | 1903.76248 | 1405.10147 | 1960.83387 | Non-sig. |
| PF3D7_1022600 | Kelch10 | 1277.21234 | 0.171731971 | 0.43648588 | 0.39344221 | 0.6939929 | 0.98334178 | 1426.19042 | 1347.41615 | 976.631744 | 1358.61105 | Non-sig. |
| PF3D7_0827000 | DBP10 | 1650.73657 | 0.171630499 | 0.44909612 | 0.38216874 | 0.70233621 | 0.98512564 | 1920.34443 | 1699.25214 | 1185.28059 | 1798.06913 | Non-sig. |
| PF3D7_1033500 | WDR70 | 1362.73959 | 0.171570843 | 0.41328542 | 0.41513887 | 0.67804024 | 0.97898416 | 1196.18344 | 1469.137 | 1367.3261 | 1418.3118 | Non-sig. |
| PF3D7_0927800 | COX5B | 1666.31193 | 0.171427212 | 0.42663818 | 0.40180935 | 0.68782434 | 0.98303031 | 1482.13163 | 2024.84106 | 1652.65714 | 1505.61791 | Non-sig. |
| PF3D7_0406500 | N/A | 10009.2419 | 0.17101507 | 0.71969324 | 0.23762217 | 0.81217415 | 0.99055243 | 16623.8798 | 12188.8296 | 2209.91297 | 9014.34505 | Non-sig. |
| PF3D7_1365400 | UBA4 | 87.7418272 | 0.170458517 | 0.66571369 | 0.25605379 | 0.7979093 | 0.99028237 | 39.9967956 | 79.4023049 | 124.835235 | 106.732973 | Non-sig. |
| PF3D7_1140600 | N/A | 1898.971 | 0.170452823 | 0.58810386 | 0.28983456 | 0.77194281 | 0.99028237 | 2793.86701 | 2030.46706 | 780.393974 | 1991.15594 | Non-sig. |
| PF3D7_0630300 | N/A | 2440.58522 | 0.170382853 | 0.49197329 | 0.34632542 | 0.72909817 | 0.98844303 | 1494.95486 | 2318.76706 | 3097.94676 | 2850.67221 | Non-sig. |
| PF3D7_1122600 | N/A | 334.957423 | 0.17035274 | 0.44120946 | 0.386104 | 0.69941964 | 0.98394192 | 318.265231 | 338.950507 | 312.155869 | 370.458083 | Non-sig. |
| PF3D7_1022700 | PLSCR | 2962.44654 | 0.17012255 | 0.45257559 | 0.37589865 | 0.70699225 | 0.986213 | 2068.25464 | 3206.59545 | 3507.3935 | 3067.54257 | Non-sig. |
| PF3D7_0308600 | PRPF19 | 1109.46154 | 0.169185623 | 0.43233622 | 0.39132882 | 0.6955542 | 0.98334178 | 878.973274 | 1122.32771 | 1209.77515 | 1226.77002 | Non-sig. |
| PF3D7_0923300 | PLP3 | 44.6338735 | 0.169082247 | 0.67613418 | 0.25007203 | 0.80253164 | 0.99028237 | 30.5126279 | 52.5826502 | 53.3800499 | 42.0601661 | Non-sig. |
| PF3D7_0409400 | DnaJ | 742.805945 | 0.169034126 | 0.41937507 | 0.40306193 | 0.68690266 | 0.982243 | 729.39851 | 785.444086 | 669.344137 | 787.037046 | Non-sig. |
| PF3D7_1035000 | N/A | 611.146233 | 0.168941097 | 0.46439629 | 0.3637865 | 0.71601745 | 0.9875928 | 718.242614 | 641.554532 | 432.859574 | 651.92821 | Non-sig. |
| PF3D7_1147800 | MAEBL | 513.310287 | 0.168841878 | 0.71618504 | 0.23575175 | 0.81362531 | 0.99071176 | 840.581619 | 632.931436 | 126.406815 | 453.32128 | Non-sig. |
| PF3D7_0722200 | RALP1 | 25989.64 | 0.168153524 | 0.63721185 | 0.26388951 | 0.79186506 | 0.99028237 | 40462.2488 | 29712.2635 | 8491.69273 | 25292.3549 | Non-sig. |
| PF3D7_0817700 | RON5 | 12186.7342 | 0.16803381 | 0.48030269 | 0.34984982 | 0.72645141 | 0.98844303 | 8026.61144 | 10686.1524 | 14928.7205 | 15105.4526 | Non-sig. |
| PF3D7_0708700 | COX4 | 1470.20875 | 0.168008209 | 0.43924861 | 0.38249002 | 0.70209793 | 0.98512564 | 1149.11142 | 1728.09554 | 1620.06786 | 1383.56017 | Non-sig. |
| PF3D7_1235300 | NOT4 | 9564.80138 | 0.167899763 | 0.70588998 | 0.23785543 | 0.81199323 | 0.99055243 | 15782.035 | 10994.9987 | 2236.08624 | 9246.08562 | Non-sig. |
| PF3D7_1419000 | N/A | 3824.1911 | 0.167776931 | 0.62453803 | 0.26864166 | 0.78820545 | 0.99028237 | 5878.79898 | 4057.0161 | 1325.773 | 4035.17631 | Non-sig. |
| PF3D7_0714000 | H2B.Z | 23430.4454 | 0.167698952 | 0.40551927 | 0.41354126 | 0.6792101 | 0.97898416 | 20273.2421 | 25255.9082 | 23867.049 | 24325.5822 | Non-sig. |
| PF3D7_1464000 | N/A | 1167.96843 | 0.167538361 | 0.70695286 | 0.23698661 | 0.81266718 | 0.99055243 | 1919.9548 | 1341.2338 | 280.882056 | 1129.80306 | Non-sig. |
| PF3D7_1143700 | N/A | 149.403515 | 0.167155607 | 0.48922545 | 0.34167398 | 0.73259626 | 0.98904466 | 129.488198 | 156.65013 | 151.942724 | 159.533008 | Non-sig. |
| PF3D7_0404400 | P36 | 185.365774 | 0.167033469 | 0.53804174 | 0.31044705 | 0.75622102 | 0.99028237 | 114.271806 | 189.05998 | 234.722968 | 203.408343 | Non-sig. |
| PF3D7_0610700 | N/A | 193.220197 | 0.166903677 | 0.7272013 | 0.2295151 | 0.81846859 | 0.99096399 | 49.5336473 | 207.191728 | 314.255788 | 201.899623 | Non-sig. |
| PF3D7_1245600 | N/A | 96.7203186 | 0.166805734 | 0.69912453 | 0.23859231 | 0.81142173 | 0.9905328 | 146.443607 | 89.0354503 | 36.142528 | 115.259689 | Non-sig. |
| PF3D7_1141100 | N/A | 96.1802836 | 0.166570502 | 0.62088042 | 0.26828113 | 0.78848293 | 0.99028237 | 126.488204 | 79.3700318 | 55.0399171 | 123.822982 | Non-sig. |
| PF3D7_1453000 | N/A | 752.428573 | 0.166522761 | 0.42294702 | 0.39372014 | 0.69378767 | 0.98334178 | 673.89738 | 743.508903 | 744.135238 | 848.172771 | Non-sig. |
| PF3D7_0605300 | ARK1 | 606.800829 | 0.16639935 | 0.43829907 | 0.37964796 | 0.70420675 | 0.9859639 | 539.28676 | 729.897561 | 604.333946 | 553.68505 | Non-sig. |
| PF3D7_1100200 | VAR | 116.370662 | 0.166347182 | 0.52185811 | 0.3187594 | 0.74990896 | 0.99028237 | 123.340338 | 136.211674 | 96.1164262 | 109.814208 | Non-sig. |
| PF3D7_0933200 | N/A | 326.093727 | 0.166276675 | 0.4825679 | 0.34456638 | 0.73042038 | 0.98844303 | 385.165834 | 352.500776 | 229.750952 | 336.957347 | Non-sig. |

| | | | | | | | | | | | | |
|---|---|---|---|---|---|---|---|---|---|---|---|---|
| PF3D7_0605500 | N/A | 924.810328 | 0.166116942 | 0.42368633 | 0.3920753 | 0.69500258 | 0.98334178 | 777.369409 | 957.279975 | 965.729906 | 998.86202 | Non-sig. |
| PF3D7_0206700 | ADSL | 601.961827 | 0.165111384 | 0.50923382 | 0.32423491 | 0.74576019 | 0.98998709 | 356.938992 | 655.540664 | 777.804135 | 617.563518 | Non-sig. |
| PF3D7_0922300 | N/A | 116.381639 | 0.164680528 | 0.51130025 | 0.32208184 | 0.74739069 | 0.99028237 | 116.164862 | 131.117201 | 103.385804 | 114.85869 | Non-sig. |
| PF3D7_0218400 | DDX47 | 401.761049 | 0.164167527 | 0.43403449 | 0.37823613 | 0.70525519 | 0.98615891 | 385.901072 | 413.22207 | 371.977127 | 435.943927 | Non-sig. |
| PF3D7_0830900 | N/A | 839.332095 | 0.163821419 | 0.55191306 | 0.29682468 | 0.76660036 | 0.99028237 | 1160.18206 | 1003.70351 | 423.665762 | 769.777052 | Non-sig. |
| PF3D7_0909400 | N/A | 2287.51141 | 0.163530251 | 0.53557806 | 0.30533411 | 0.7601117 | 0.99028237 | 1243.4185 | 1934.00749 | 3072.2131 | 2900.40655 | Non-sig. |
| PF3D7_0908800 | N/A | 994.252994 | 0.163178742 | 0.47727677 | 0.34189542 | 0.73242959 | 0.98904466 | 653.729342 | 1107.29379 | 1222.1031 | 993.885748 | Non-sig. |
| PF3D7_1416300 | N/A | 783.685149 | 0.162439388 | 0.49461321 | 0.328417 | 0.74259639 | 0.98975488 | 486.910747 | 873.636468 | 991.962403 | 782.230976 | Non-sig. |
| PF3D7_0507600 | CAF40 | 1159.66146 | 0.161936134 | 0.55050662 | 0.29415838 | 0.76863688 | 0.99028237 | 703.235232 | 798.272371 | 1485.7529 | 1651.38534 | Non-sig. |
| PF3D7_1453200 | N/A | 2177.1045 | 0.161913643 | 0.43102227 | 0.37565029 | 0.7071769 | 0.986213 | 1956.31822 | 1895.52408 | 2153.74188 | 2702.83381 | Non-sig. |
| PF3D7_1004800 | AAC2 | 634.853879 | 0.161904869 | 0.45647474 | 0.35468527 | 0.72282538 | 0.98844303 | 478.43441 | 737.814079 | 719.863421 | 603.303605 | Non-sig. |
| PF3D7_0212900 | ATEL1 | 5857.49369 | 0.161511795 | 0.42367197 | 0.38121898 | 0.70304077 | 0.98512564 | 6425.7787 | 6454.42345 | 4634.3726 | 5915.40001 | Non-sig. |
| PF3D7_1444500 | eIK1 | 964.380491 | 0.161091587 | 0.43967275 | 0.36638974 | 0.71407427 | 0.9875928 | 1013.83563 | 865.605209 | 807.501255 | 1170.57987 | Non-sig. |
| PF3D7_0918200 | N/A | 819.299971 | 0.160890518 | 0.41898103 | 0.3840043 | 0.70097526 | 0.98508406 | 802.151131 | 905.506303 | 745.229325 | 824.313126 | Non-sig. |
| PF3D7_1315200 | N/A | 572.382852 | 0.160618931 | 0.50781798 | 0.31629233 | 0.75178063 | 0.99028237 | 346.459702 | 556.811485 | 734.291842 | 651.96838 | Non-sig. |
| PF3D7_0714300 | DHHC4 | 1176.03975 | 0.160356683 | 0.49614237 | 0.32320699 | 0.74653849 | 0.98998709 | 1509.69955 | 1195.01466 | 712.168863 | 1287.27591 | Non-sig. |
| PF3D7_0218300 | N/A | 594.808072 | 0.160067736 | 0.4503385 | 0.35543871 | 0.72226095 | 0.98844303 | 661.701772 | 682.836991 | 462.211449 | 572.482076 | Non-sig. |
| PF3D7_0527000 | MCM3 | 14976.5334 | 0.159946819 | 0.4251054 | 0.37625214 | 0.70672945 | 0.986213 | 11647.8008 | 16213.9614 | 16646.3301 | 15398.0412 | Non-sig. |
| PF3D7_1215600 | N/A | 323.367652 | 0.15977803 | 0.46794219 | 0.34144822 | 0.73276618 | 0.98904466 | 249.831807 | 370.083281 | 360.938963 | 312.616556 | Non-sig. |
| PF3D7_0103000 | N/A | 575.03234 | 0.159740686 | 0.44690823 | 0.35743509 | 0.72076611 | 0.98816139 | 484.486934 | 695.604005 | 601.841136 | 518.197283 | Non-sig. |
| PF3D7_0115100 | N/A | 66.0971841 | 0.159314942 | 0.6366572 | 0.25023661 | 0.80240437 | 0.99028237 | 85.7503248 | 70.947388 | 39.3801896 | 68.3108339 | Non-sig. |
| PF3D7_1223000 | N/A | 714.419985 | 0.159044106 | 0.42917984 | 0.37057683 | 0.71095274 | 0.9875928 | 661.556804 | 841.088394 | 688.582484 | 666.452256 | Non-sig. |
| PF3D7_0418700 | N/A | 2566.25838 | 0.158637145 | 0.43566343 | 0.36412775 | 0.71576262 | 0.9875928 | 2895.52661 | 2547.57984 | 1955.35033 | 2866.57674 | Non-sig. |
| PF3D7_0616500 | TLP | 103.040261 | 0.158395636 | 0.54450659 | 0.29089755 | 0.77112967 | 0.99028237 | 116.917406 | 98.6962578 | 78.0318895 | 118.51549 | Non-sig. |
| PF3D7_1005800 | N/A | 403.894992 | 0.158242869 | 0.4417168 | 0.35824508 | 0.72015992 | 0.98797939 | 418.87754 | 447.769235 | 344.782372 | 404.150821 | Non-sig. |
| PF3D7_0308900 | SF3B1 | 1986.14132 | 0.158114162 | 0.43199806 | 0.36600665 | 0.71436011 | 0.9875928 | 1587.70506 | 1884.76131 | 2166.90967 | 2305.18924 | Non-sig. |
| PF3D7_1230800 | WTAP | 1023.97448 | 0.157946675 | 0.55412933 | 0.28503576 | 0.77561675 | 0.99028237 | 1441.11579 | 1082.62341 | 495.245245 | 1076.91347 | Non-sig. |
| PF3D7_0719800 | N/A | 691.353196 | 0.15769789 | 0.42692498 | 0.3693808 | 0.71184391 | 0.9875928 | 604.101042 | 776.877079 | 703.007058 | 681.427605 | Non-sig. |
| PF3D7_0934800 | PKAc | 10678.2505 | 0.156396296 | 0.40390643 | 0.38720922 | 0.69860132 | 0.98377748 | 10741.2498 | 11118.6197 | 9458.90672 | 11394.2256 | Non-sig. |
| PF3D7_1410400 | RAP1 | 43517.9049 | 0.15611363 | 0.5649382 | 0.27633753 | 0.78228883 | 0.99028237 | 20032.3055 | 39125.8848 | 62298.6942 | 52614.735 | Non-sig. |
| PF3D7_0715600 | N/A | 1822.25816 | 0.155782138 | 0.50862151 | 0.30628303 | 0.75938916 | 0.99028237 | 1037.87692 | 1973.21963 | 2409.67506 | 1868.26102 | Non-sig. |
| PF3D7_0733000 | VAR | 471.578802 | 0.155306001 | 0.43290893 | 0.35874982 | 0.71978226 | 0.98797939 | 440.604979 | 535.209924 | 451.821 | 458.679304 | Non-sig. |
| PF3D7_1346900 | N/A | 1644.90435 | 0.155082925 | 0.46377567 | 0.33439211 | 0.73808369 | 0.98954916 | 1122.53392 | 1811.95031 | 1990.30106 | 1654.83209 | Non-sig. |
| PF3D7_1117100 | UCH54 | 2020.8352 | 0.155017012 | 0.45525145 | 0.34050855 | 0.73347358 | 0.98909017 | 1418.48058 | 2039.27574 | 2405.94625 | 2219.63823 | Non-sig. |
| PF3D7_1212300 | N/A | 508.696404 | 0.154833182 | 0.57612806 | 0.26874786 | 0.78812372 | 0.99028237 | 725.853425 | 587.858389 | 237.388227 | 483.685576 | Non-sig. |
| PF3D7_1018200 | PPP8 | 14071.949 | 0.154702808 | 0.41082372 | 0.37656737 | 0.70649514 | 0.986213 | 12101.8158 | 13805.3431 | 14534.4372 | 15846.1999 | Non-sig. |
| PF3D7_1318000 | N/A | 215.410034 | 0.154565346 | 0.46366062 | 0.3333588 | 0.73886346 | 0.98954916 | 210.815569 | 213.739906 | 196.985882 | 240.098781 | Non-sig. |
| PF3D7_1464900 | N/A | 2636.71174 | 0.154465045 | 0.50370735 | 0.30665633 | 0.75910498 | 0.99028237 | 1533.14505 | 2578.12777 | 3457.85819 | 2977.71594 | Non-sig. |
| PF3D7_1003400 | N/A | 7160.82231 | 0.154026652 | 0.53111325 | 0.29000718 | 0.77181075 | 0.99028237 | 9838.63272 | 6901.11892 | 3719.64053 | 8183.89706 | Non-sig. |
| PF3D7_0921300 | N/A | 163.848025 | 0.153956766 | 0.5088621 | 0.30255105 | 0.76223202 | 0.99028237 | 121.979045 | 186.752816 | 188.042732 | 158.617505 | Non-sig. |
| PF3D7_0626200 | N/A | 81.6948954 | 0.153534503 | 0.56927205 | 0.26970322 | 0.78738858 | 0.99028237 | 60.9426152 | 90.1760253 | 93.5981531 | 82.0627881 | Non-sig. |
| PF3D7_0809200 | pfa55-14 | 1003.48772 | 0.153347729 | 0.46491753 | 0.32983856 | 0.74152195 | 0.98954916 | 716.44487 | 928.961298 | 1183.67646 | 1184.86825 | Non-sig. |
| PF3D7_0300900 | N/A | 4.59246748 | 0.153018359 | 1.61786495 | 0.09458043 | 0.9246481 | 0.99619209 | 2.08857894 | 5.37684777 | 6.56197132 | 4.34247188 | Non-sig. |
| PF3D7_1143500 | N/A | 953.515453 | 0.152819762 | 0.46938927 | 0.32557148 | 0.74474859 | 0.98998709 | 1159.24035 | 1014.96683 | 647.209775 | 992.644854 | Non-sig. |
| PF3D7_1440100 | N/A | 6523.76786 | 0.152818411 | 0.46849586 | 0.32618946 | 0.74428102 | 0.98998709 | 4303.88838 | 6424.88498 | 8052.91184 | 7313.38624 | Non-sig. |
| PF3D7_0316200 | N/A | 2046.03297 | 0.152387265 | 0.46299693 | 0.32913234 | 0.74205566 | 0.98954916 | 2479.13016 | 2037.88814 | 1397.35291 | 2269.76066 | Non-sig. |
| PF3D7_0513600 | N/A | 5594.48055 | 0.152055517 | 0.4892883 | 0.31076876 | 0.75597642 | 0.99028237 | 7212.70583 | 5743.30164 | 3387.54834 | 6034.36638 | Non-sig. |
| PF3D7_1420200 | UNC | 3964.46587 | 0.151484081 | 0.43725926 | 0.34643996 | 0.7290121 | 0.98844303 | 2953.93403 | 4197.62811 | 4558.82699 | 4147.47434 | Non-sig. |
| PF3D7_1474200 | N/A | 159.746341 | 0.15101252 | 0.52070285 | 0.29001669 | 0.77180347 | 0.99028237 | 120.20831 | 139.41929 | 182.382816 | 196.974947 | Non-sig. |
| PF3D7_0910400 | N/A | 1680.26608 | 0.150989644 | 0.42660046 | 0.3539369 | 0.72338617 | 0.98844303 | 1841.25751 | 1744.32996 | 1343.8004 | 1791.67643 | Non-sig. |
| PF3D7_0705200 | N/A | 140.455912 | 0.150430871 | 0.49423274 | 0.30437253 | 0.76084409 | 0.99028237 | 123.273357 | 142.668697 | 142.928861 | 152.952733 | Non-sig. |
| PF3D7_0705300 | ORC2 | 3408.6731 | 0.150135782 | 0.5449024 | 0.27552784 | 0.78291074 | 0.99028237 | 4785.3731 | 3382.87445 | 1677.97882 | 3788.46601 | Non-sig. |
| PF3D7_1139800 | N/A | 1278.4179 | 0.149546558 | 0.55970768 | 0.2671869 | 0.78932526 | 0.99028237 | 617.614021 | 1165.91865 | 1806.42372 | 1523.71522 | Non-sig. |
| PF3D7_0309700 | SBP2 | 465.243287 | 0.149087279 | 0.63695847 | 0.23406122 | 0.81493745 | 0.99096399 | 713.693096 | 548.518081 | 169.14367 | 429.618302 | Non-sig. |

| | | | | | | | | | | | | |
|---|---|---|---|---|---|---|---|---|---|---|---|---|
| PF3D7_0728100 | N/A | 2447.76712 | 0.149011113 | 0.44942825 | 0.33155707 | 0.74022374 | 0.98954916 | 2072.15624 | 1999.01882 | 2570.64302 | 3149.25042 | Non-sig. |
| PF3D7_1365800 | N/A | 785.557857 | 0.148626881 | 0.42711439 | 0.3479791 | 0.72785588 | 0.98844303 | 669.122788 | 781.589593 | 820.994326 | 870.524722 | Non-sig. |
| PF3D7_1455600 | FER2 | 3339.30651 | 0.148590548 | 0.43105273 | 0.34471548 | 0.73030828 | 0.98844303 | 2624.18392 | 3222.1486 | 3710.56637 | 3800.32714 | Non-sig. |
| PF3D7_0925400 | N/A | 139.812725 | 0.148408396 | 0.51456271 | 0.28841654 | 0.77302791 | 0.99028237 | 131.402737 | 116.998891 | 133.83983 | 177.009442 | Non-sig. |
| PF3D7_1403800 | MISFIT | 821.624079 | 0.147944159 | 0.50773362 | 0.29138145 | 0.7707596 | 0.99028237 | 1041.90526 | 702.639942 | 517.49037 | 1024.46075 | Non-sig. |
| PF3D7_1426400 | N/A | 8.78574349 | 0.14750534 | 1.18718436 | 0.12424805 | 0.90111888 | 0.99457954 | 9.32259979 | 9.70034592 | 7.37175715 | 8.7482711 | Non-sig. |
| PF3D7_1428700 | N/A | 657.254347 | 0.147134133 | 0.51787706 | 0.28411016 | 0.77632596 | 0.99028237 | 868.463666 | 645.925004 | 379.441978 | 735.186739 | Non-sig. |
| PF3D7_1364300 | PRP16 | 1700.81752 | 0.147117147 | 0.42247306 | 0.34822847 | 0.72766861 | 0.98844303 | 1399.7662 | 1718.63082 | 1828.39153 | 1856.48152 | Non-sig. |
| PF3D7_1117500 | TyrRSapi | 652.417768 | 0.146389681 | 0.45092397 | 0.32464382 | 0.74545066 | 0.98998709 | 492.037067 | 671.913552 | 746.400213 | 699.320242 | Non-sig. |
| PF3D7_1123500 | GP2 | 1404.20874 | 0.145942489 | 0.50374315 | 0.28971608 | 0.77203346 | 0.99028237 | 878.715138 | 1201.42162 | 1787.43126 | 1749.26694 | Non-sig. |
| PF3D7_0807600 | N/A | 5406.43777 | 0.145647119 | 0.45207518 | 0.32217455 | 0.74732046 | 0.99028237 | 6481.69377 | 5619.3856 | 3786.16937 | 5738.50234 | Non-sig. |
| PF3D7_0702500 | N/A | 687.715377 | 0.145384861 | 0.52733292 | 0.27569843 | 0.7827797 | 0.99028237 | 381.926463 | 670.700756 | 923.882319 | 774.351969 | Non-sig. |
| PF3D7_1321100 | N/A | 20416.9475 | 0.14532836 | 0.43878798 | 0.33120406 | 0.74049036 | 0.98954916 | 16076.6232 | 18190.4903 | 22702.1195 | 24698.5569 | Non-sig. |
| PF3D7_0935200 | VPS33 | 3619.46039 | 0.144766425 | 0.55162886 | 0.26243447 | 0.7929865 | 0.99028237 | 5139.98331 | 4017.38682 | 1736.46696 | 3584.00446 | Non-sig. |
| PF3D7_0203100 | N/A | 5088.8827 | 0.144754436 | 0.46907545 | 0.30859521 | 0.75762947 | 0.99028237 | 6181.46917 | 4591.57733 | 3486.44953 | 6096.03479 | Non-sig. |
| PF3D7_1102400 | PPCS | 1331.66527 | 0.144451699 | 0.48720129 | 0.29649285 | 0.76685373 | 0.99028237 | 839.371818 | 1315.28617 | 1690.37632 | 1481.62679 | Non-sig. |
| PF3D7_1409800 | CELF2 | 316.364579 | 0.144433808 | 0.54505353 | 0.26499014 | 0.79101706 | 0.99028237 | 184.313527 | 271.421223 | 416.43768 | 393.285886 | Non-sig. |
| PF3D7_0517900 | N/A | 3167.10704 | 0.144331577 | 0.41574572 | 0.34716311 | 0.72846879 | 0.98844303 | 2639.94705 | 3303.39137 | 3377.50729 | 3347.58243 | Non-sig. |
| PF3D7_0716000 | N/A | 1629.08405 | 0.143784392 | 0.45405878 | 0.31666471 | 0.75149803 | 0.99028237 | 1938.09405 | 1660.67092 | 1158.1579 | 1759.41331 | Non-sig. |
| PF3D7_0912000 | N/A | 1692.21969 | 0.143311819 | 0.42826096 | 0.33463666 | 0.73789919 | 0.98954916 | 1780.74679 | 1554.30556 | 1435.87854 | 1997.94789 | Non-sig. |
| PF3D7_1229600 | N/A | 3103.97859 | 0.142995092 | 0.6123857 | 0.23350495 | 0.81536933 | 0.99096399 | 4744.44021 | 3476.82899 | 1156.52886 | 3038.11629 | Non-sig. |
| PF3D7_0406100 | N/A | 5293.06398 | 0.141877118 | 0.49545971 | 0.2863545 | 0.77460662 | 0.99028237 | 3138.85455 | 5619.22922 | 6926.78302 | 5487.38913 | Non-sig. |
| PF3D7_0814300 | N/A | 566.146239 | 0.141759611 | 0.45606281 | 0.31083352 | 0.75592718 | 0.99028237 | 430.841963 | 546.034148 | 645.62119 | 642.087656 | Non-sig. |
| PF3D7_1457900 | N/A | 6567.35758 | 0.141263162 | 0.42007678 | 0.33627938 | 0.73666019 | 0.98954916 | 6697.21714 | 5952.3079 | 5795.06678 | 7824.83851 | Non-sig. |
| PF3D7_1208000 | N/A | 20.8570646 | 0.141201778 | 0.86885359 | 0.16251504 | 0.87090028 | 0.99402081 | 26.7130979 | 22.7106694 | 13.0679211 | 20.9365699 | Non-sig. |
| PF3D7_0505600 | N/A | 392.503687 | 0.140992467 | 0.46398717 | 0.30387148 | 0.76122581 | 0.99028237 | 447.269063 | 382.257082 | 299.637778 | 440.850825 | Non-sig. |
| PF3D7_1446500 | NUP313 | 3839.41832 | 0.140986715 | 0.53283174 | 0.26459894 | 0.79131843 | 0.99028237 | 2095.92694 | 3270.06683 | 5207.62311 | 4784.05642 | Non-sig. |
| PF3D7_0204200 | SEC66 | 2941.34691 | 0.140900728 | 0.50842356 | 0.27713257 | 0.78167831 | 0.99028237 | 3922.18834 | 2983.50049 | 1673.85233 | 3185.84646 | Non-sig. |
| PF3D7_0908900 | N/A | 494.020319 | 0.14062169 | 0.43529904 | 0.32304617 | 0.74666028 | 0.98998709 | 448.682964 | 567.513124 | 491.185039 | 468.70015 | Non-sig. |
| PF3D7_0722800 | N/A | 97.5773768 | 0.140549066 | 0.53629183 | 0.26207572 | 0.79326307 | 0.99028237 | 106.139064 | 105.201226 | 79.642492 | 99.3267245 | Non-sig. |
| PF3D7_0217100 | N/A | 2397.91098 | 0.140352902 | 0.47657893 | 0.29450085 | 0.76837521 | 0.99028237 | 1569.81671 | 2738.59291 | 2992.56592 | 2290.66837 | Non-sig. |
| PF3D7_0501800 | CAF1A | 4990.73047 | 0.140066572 | 0.45129937 | 0.31036288 | 0.75628502 | 0.99028237 | 5983.64873 | 5206.01984 | 3513.98604 | 5259.26728 | Non-sig. |
| PF3D7_1338600 | N/A | 4735.6479 | 0.138911739 | 0.42123279 | 0.32977428 | 0.74157052 | 0.98954916 | 5179.90316 | 5181.3344 | 3835.98387 | 4745.37018 | Non-sig. |
| PF3D7_1338500 | N/A | 85.2103742 | 0.138851779 | 0.54611959 | 0.2542516 | 0.79930119 | 0.99028237 | 76.7773788 | 81.5658038 | 85.4021856 | 97.0961287 | Non-sig. |
| PF3D7_1136400 | SRP72 | 2977.20712 | 0.137918553 | 0.43621612 | 0.31617023 | 0.7518733 | 0.99028237 | 3411.38339 | 3236.38903 | 2258.90625 | 3002.1498 | Non-sig. |
| PF3D7_0609200 | N/A | 159.229584 | 0.136793945 | 0.48074369 | 0.28454652 | 0.77599159 | 0.99028237 | 151.453774 | 170.643453 | 151.91779 | 162.903317 | Non-sig. |
| PF3D7_1470500 | PDEdelta | 457.164153 | 0.136711119 | 0.44591912 | 0.30658277 | 0.75916097 | 0.99028237 | 373.922148 | 506.355815 | 496.939258 | 451.43939 | Non-sig. |
| PF3D7_0917900 | BIP | 50867.0981 | 0.136388377 | 0.46674602 | 0.29221112 | 0.77012521 | 0.99028237 | 33498.1992 | 52853.4262 | 63430.4413 | 53686.3259 | Non-sig. |
| PF3D7_0823200 | N/A | 2773.49941 | 0.136146326 | 0.60418308 | 0.22533952 | 0.82171516 | 0.99096399 | 1079.58426 | 2650.89315 | 4205.46844 | 3158.05178 | Non-sig. |
| PF3D7_1421300 | N/A | 264.530315 | 0.136105068 | 0.70965416 | 0.1917907 | 0.84790615 | 0.99165628 | 430.561994 | 322.959431 | 73.9177048 | 230.682129 | Non-sig. |
| PF3D7_0106300 | ATP6 | 11030.6599 | 0.135547632 | 0.50037054 | 0.27089451 | 0.78647217 | 0.99028237 | 6426.6233 | 11055.9949 | 14598.6973 | 12041.3239 | Non-sig. |
| PF3D7_1322600 | N/A | 1045.86801 | 0.135535589 | 0.42795081 | 0.31670834 | 0.75146492 | 0.99028237 | 900.414646 | 1208.41612 | 1093.00947 | 981.631801 | Non-sig. |
| PF3D7_1360000 | N/A | 85.9857438 | 0.134880811 | 0.58329196 | 0.23124065 | 0.81712785 | 0.99096399 | 58.0547268 | 84.0494317 | 105.678207 | 96.1606098 | Non-sig. |
| PF3D7_1337900 | N/A | 1022.17209 | 0.13460778 | 0.43205222 | 0.31155442 | 0.75537918 | 0.99028237 | 1081.33873 | 1193.71766 | 867.854214 | 945.777766 | Non-sig. |
| PF3D7_0500900 | FIKK5 | 764.645459 | 0.133819436 | 0.42126559 | 0.3176605 | 0.75074248 | 0.99028237 | 779.393735 | 786.66631 | 679.119357 | 813.402432 | Non-sig. |
| PF3D7_1359800 | N/A | 216.869819 | 0.133086262 | 0.47356464 | 0.28103083 | 0.77868676 | 0.99028237 | 231.91056 | 244.386597 | 181.971223 | 209.210898 | Non-sig. |
| PF3D7_1307600 | rpoA | 2286.38953 | 0.132736005 | 0.41363708 | 0.32089968 | 0.74828642 | 0.99028237 | 2093.16443 | 2168.33711 | 2269.3416 | 2614.71497 | Non-sig. |
| PF3D7_1251800 | N/A | 910.38886 | 0.131543666 | 0.47229099 | 0.27852249 | 0.7806113 | 0.99028237 | 1117.30546 | 900.363691 | 620.842483 | 1003.04381 | Non-sig. |
| PF3D7_1232200 | LPD1 | 464.349543 | 0.131391352 | 0.43872301 | 0.29948589 | 0.76456934 | 0.99028237 | 455.361454 | 430.378653 | 431.109488 | 540.548578 | Non-sig. |
| PF3D7_1360800 | FLN | 11854.5025 | 0.130660472 | 0.47733485 | 0.27372917 | 0.78429276 | 0.99028237 | 7532.4412 | 12257.062 | 15103.3126 | 12525.1942 | Non-sig. |
| PF3D7_1419100 | DDX55 | 1009.18798 | 0.130544486 | 0.54949971 | 0.23756971 | 0.81221485 | 0.99055243 | 1424.01371 | 1091.036 | 503.507251 | 1018.19497 | Non-sig. |
| PF3D7_0302600 | ABCB4 | 1061.33648 | 0.130444208 | 0.45748949 | 0.2851305 | 0.77554416 | 0.99028237 | 760.664544 | 1067.35244 | 1265.81008 | 1151.51885 | Non-sig. |
| PF3D7_1005100 | UTP25 | 479.073243 | 0.130196177 | 0.53830523 | 0.24186311 | 0.80888624 | 0.99028237 | 654.113714 | 483.824507 | 261.202056 | 517.152695 | Non-sig. |
| PF3D7_1474400 | N/A | 856.655102 | 0.130124817 | 0.46887931 | 0.27752305 | 0.78137851 | 0.99028237 | 1045.75328 | 903.073677 | 590.663787 | 887.129661 | Non-sig. |

| | | | | | | | | | | | | |
|---|---|---|---|---|---|---|---|---|---|---|---|---|
| PF3D7_1018800 | N/A | 320.153751 | 0.129990645 | 0.52359933 | 0.24826358 | 0.80393048 | 0.99028237 | 417.985331 | 351.915954 | 193.833181 | 316.880535 | Non-sig. |
| PF3D7_0405900 | ASP | 20005.8762 | 0.12998241 | 0.59841879 | 0.21720977 | 0.82804487 | 0.99110542 | 30295.0914 | 22838.4825 | 7915.85867 | 18974.0723 | Non-sig. |
| PF3D7_1239500 | GyrB | 1108.41502 | 0.129830984 | 0.42588879 | 0.30484715 | 0.76048257 | 0.99028237 | 1095.39516 | 1007.35772 | 1021.79938 | 1309.10782 | Non-sig. |
| PF3D7_0912100 | N/A | 236.31235 | 0.129765562 | 0.51504713 | 0.25194891 | 0.80108055 | 0.99028237 | 166.181658 | 202.839229 | 284.949468 | 291.279044 | Non-sig. |
| PF3D7_1446700 | N/A | 2130.29364 | 0.129401718 | 0.41321187 | 0.3131607 | 0.75415858 | 0.99028237 | 2178.66503 | 2370.94284 | 1891.08198 | 2080.48472 | Non-sig. |
| PF3D7_0603100 | N/A | 680.891492 | 0.12933143 | 0.434844 | 0.2974203 | 0.76614565 | 0.99028237 | 565.69282 | 669.746742 | 734.918276 | 753.208131 | Non-sig. |
| PF3D7_1468000 | N/A | 258.428057 | 0.12928377 | 0.56955118 | 0.22699237 | 0.82042968 | 0.99096399 | 359.34821 | 245.635223 | 134.718759 | 294.010034 | Non-sig. |
| PF3D7_1104900 | N/A | 10224.572 | 0.129228253 | 0.63745427 | 0.20272553 | 0.83934957 | 0.99128968 | 16144.8602 | 9883.76668 | 3389.46196 | 11480.199 | Non-sig. |
| PF3D7_1141200 | N/A | 410.546145 | 0.128907239 | 0.45631403 | 0.28249677 | 0.77756263 | 0.99028237 | 392.990608 | 508.151717 | 391.452716 | 349.58954 | Non-sig. |
| PF3D7_1321300 | N/A | 1746.56938 | 0.128531103 | 0.4590831 | 0.2799735 | 0.77949784 | 0.99028237 | 1285.19585 | 1559.41592 | 2052.16485 | 2089.50088 | Non-sig. |
| PF3D7_1311500 | RPT1 | 7200.49826 | 0.128309988 | 0.46044203 | 0.27866698 | 0.78050041 | 0.99028237 | 4899.67372 | 7450.73943 | 8861.0144 | 7590.5655 | Non-sig. |
| PF3D7_1022000 | N/A | 1630.58501 | 0.127259767 | 0.44052161 | 0.28888428 | 0.77266994 | 0.99028237 | 1511.39675 | 1349.32506 | 1605.99011 | 2055.6281 | Non-sig. |
| PF3D7_1016200 | N/A | 508.810716 | 0.126969658 | 0.59641731 | 0.21288728 | 0.83141488 | 0.99110542 | 217.780931 | 563.397252 | 754.702017 | 499.362662 | Non-sig. |
| PF3D7_1219100 | CHC | 5857.80514 | 0.126825822 | 0.63585023 | 0.19945864 | 0.84190399 | 0.99128968 | 1983.59321 | 5406.94076 | 9217.00022 | 6823.68636 | Non-sig. |
| PF3D7_0706300 | N/A | 851.266512 | 0.126766485 | 0.56841554 | 0.22301728 | 0.82352206 | 0.99096399 | 1149.66198 | 1187.11008 | 478.444675 | 589.849309 | Non-sig. |
| PF3D7_0219300 | N/A | 370.715947 | 0.126687233 | 0.48525835 | 0.26107172 | 0.7940372 | 0.99028237 | 259.764786 | 414.745654 | 448.857878 | 359.49547 | Non-sig. |
| PF3D7_1304000 | CAPH | 3881.41161 | 0.126245727 | 0.51656691 | 0.24439375 | 0.80692588 | 0.99028237 | 2140.54945 | 4013.17219 | 5282.44444 | 4089.48036 | Non-sig. |
| PF3D7_1220400 | DRN1 | 740.465708 | 0.126182746 | 0.46711559 | 0.27013174 | 0.78705891 | 0.99028237 | 895.811778 | 799.364453 | 520.703892 | 745.98271 | Non-sig. |
| PF3D7_1405900 | N/A | 879.750968 | 0.126094429 | 0.66326363 | 0.19011208 | 0.8492213 | 0.99165628 | 1413.632 | 974.071688 | 269.418358 | 861.881825 | Non-sig. |
| PF3D7_1236300 | N/A | 494.795952 | 0.125671277 | 0.4657353 | 0.26983412 | 0.78728787 | 0.99028237 | 364.88434 | 550.599351 | 581.371939 | 482.328179 | Non-sig. |
| PF3D7_1349100 | N/A | 692.172696 | 0.125631316 | 0.42708556 | 0.2941596 | 0.76863595 | 0.99028237 | 702.138083 | 781.029716 | 622.056131 | 663.466855 | Non-sig. |
| PF3D7_0908600 | N/A | 295.322812 | 0.125413907 | 0.48441089 | 0.25889985 | 0.79571252 | 0.99028237 | 212.444005 | 311.241423 | 352.281556 | 305.324263 | Non-sig. |
| PF3D7_1145100 | SEC21 | 1394.74726 | 0.124379878 | 0.52613652 | 0.23640229 | 0.81312052 | 0.99055243 | 787.942193 | 1225.0919 | 1881.00685 | 1684.94811 | Non-sig. |
| PF3D7_1473900 | N/A | 432.134935 | 0.124170968 | 0.46106333 | 0.26931434 | 0.78768779 | 0.99028237 | 425.479867 | 357.251097 | 401.64944 | 544.159335 | Non-sig. |
| PF3D7_1365600 | TOP6B | 861.260208 | 0.123637062 | 0.59072336 | 0.20929774 | 0.83421582 | 0.99128968 | 1277.19485 | 1000.40293 | 371.957742 | 795.485306 | Non-sig. |
| PF3D7_1419400 | N/A | 2372.72732 | 0.123467385 | 0.44548364 | 0.27715358 | 0.78166218 | 0.99028237 | 2071.338 | 1935.96057 | 2471.06527 | 3012.54543 | Non-sig. |
| PF3D7_0214900 | RON6 | 15765.4705 | 0.123359959 | 0.71299892 | 0.17301563 | 0.86263914 | 0.99402081 | 26524.7559 | 18826.6917 | 3659.36826 | 14051.0662 | Non-sig. |
| PF3D7_0525200 | SMC6 | 6318.56859 | 0.123138077 | 0.46556908 | 0.26448938 | 0.79140284 | 0.99028237 | 7856.38831 | 6302.55355 | 4242.12119 | 6873.21132 | Non-sig. |
| PF3D7_1317600 | N/A | 1070.42982 | 0.1230287 | 0.44841363 | 0.27436432 | 0.78380466 | 0.99028237 | 1209.65351 | 1264.99043 | 840.207569 | 966.86779 | Non-sig. |
| PF3D7_1125200 | N/A | 371.408131 | 0.122800274 | 0.43779326 | 0.28049832 | 0.77909521 | 0.99028237 | 344.084695 | 402.408863 | 367.095819 | 372.043146 | Non-sig. |
| PF3D7_1031200 | MORN1 | 2913.62905 | 0.121551963 | 0.40676238 | 0.29882794 | 0.76507133 | 0.99028237 | 2871.97318 | 3189.43063 | 2709.99431 | 2883.11809 | Non-sig. |
| PF3D7_1211900 | ATP4 | 4703.12876 | 0.121529444 | 0.67646097 | 0.17965478 | 0.8574236 | 0.99352464 | 1326.79777 | 4193.98506 | 7683.13168 | 5608.60055 | Non-sig. |
| PF3D7_1038100 | N/A | 327.142504 | 0.121210207 | 0.44404869 | 0.27296603 | 0.78487933 | 0.99028237 | 306.455286 | 318.634214 | 320.330882 | 363.149635 | Non-sig. |
| PF3D7_0318200 | RPB1 | 7173.58904 | 0.120884559 | 0.57026275 | 0.21198046 | 0.83212228 | 0.99110542 | 3214.28246 | 6866.32273 | 10531.7624 | 8081.98854 | Non-sig. |
| PF3D7_1371700 | FIKK13 | 1772.08851 | 0.120832313 | 0.42361138 | 0.28524331 | 0.77545774 | 0.99028237 | 1459.88058 | 1775.82266 | 1935.76686 | 1916.88395 | Non-sig. |
| PF3D7_1323300 | N/A | 526.764919 | 0.120818029 | 0.44499827 | 0.27150224 | 0.78600478 | 0.99028237 | 557.148748 | 619.342394 | 452.428055 | 478.140479 | Non-sig. |
| PF3D7_0803000 | CYP81 | 2391.8618 | 0.120752081 | 0.49501592 | 0.24393575 | 0.80728058 | 0.99028237 | 3127.98385 | 2644.17671 | 1456.03646 | 2339.25018 | Non-sig. |
| PF3D7_0217300 | AP2S | 635.000269 | 0.120664067 | 0.42287139 | 0.2853446 | 0.77538015 | 0.99028237 | 613.634539 | 693.306813 | 603.301992 | 629.757733 | Non-sig. |
| PF3D7_0529300 | TIC22 | 255.373813 | 0.120495899 | 0.45468144 | 0.26501169 | 0.79100045 | 0.99028237 | 237.35545 | 250.163845 | 252.037538 | 281.938421 | Non-sig. |
| PF3D7_1244700 | N/A | 1020.07447 | 0.12036992 | 0.48899143 | 0.24615957 | 0.80555871 | 0.99028237 | 1268.86044 | 1242.67965 | 686.545397 | 882.212375 | Non-sig. |
| PF3D7_1021700 | N/A | 23200.4012 | 0.120321983 | 0.5118401 | 0.23507729 | 0.81414875 | 0.99096399 | 30114.6055 | 18455.1138 | 14352.7528 | 29879.1326 | Non-sig. |
| PF3D7_1100100 | VAR | 641.705193 | 0.120289135 | 0.43390439 | 0.27722498 | 0.78160736 | 0.99028237 | 536.444556 | 645.711781 | 693.321509 | 691.342924 | Non-sig. |
| PF3D7_1306700 | N/A | 329.668546 | 0.120234263 | 0.45784368 | 0.26260986 | 0.7928513 | 0.99028237 | 359.447631 | 371.41137 | 272.59693 | 315.218254 | Non-sig. |
| PF3D7_0410400 | RRP4 | 347.797763 | 0.119663402 | 0.44930188 | 0.26633185 | 0.78998364 | 0.99028237 | 304.606345 | 395.095744 | 362.054767 | 329.434196 | Non-sig. |
| PF3D7_1254800 | RIF | 3.13204437 | 0.119032542 | 1.95501949 | 0.0608856 | 0.95145031 | 0.99639249 | 5.23296778 | 3.2241364 | 0.82053158 | 3.25054173 | Non-sig. |
| PF3D7_1404600 | ACalpha | 158.304194 | 0.118523033 | 0.50058759 | 0.23676782 | 0.81283692 | 0.99055243 | 179.073265 | 169.936114 | 124.751029 | 159.456367 | Non-sig. |
| PF3D7_1009200 | N/A | 196.166658 | 0.118388875 | 0.50544053 | 0.23422909 | 0.81480713 | 0.99096399 | 141.095648 | 193.119818 | 234.903148 | 215.548018 | Non-sig. |
| PF3D7_1411300 | N/A | 1436.94166 | 0.118370334 | 0.41198394 | 0.28731784 | 0.77386896 | 0.99028237 | 1455.99303 | 1532.15507 | 1300.14323 | 1459.47529 | Non-sig. |
| PF3D7_1472900 | N/A | 292.291648 | 0.118347553 | 0.57766531 | 0.20487218 | 0.83767199 | 0.99128968 | 143.076626 | 309.067583 | 417.176907 | 299.845476 | Non-sig. |
| PF3D7_1420600 | PANK1 | 1387.70023 | 0.118292975 | 0.43019843 | 0.27497305 | 0.78333694 | 0.99028237 | 1118.38704 | 1506.67609 | 1543.07991 | 1382.65789 | Non-sig. |
| PF3D7_1338100 | RPN3 | 9640.40299 | 0.118238572 | 0.43055461 | 0.27461922 | 0.7836088 | 0.99028237 | 7578.49871 | 10843.5172 | 10912.4192 | 9227.17686 | Non-sig. |
| PF3D7_1438400 | MCA2 | 7873.58067 | 0.117908006 | 0.41243819 | 0.28588043 | 0.77496971 | 0.99028237 | 7813.41361 | 7263.53396 | 7290.67112 | 9126.704 | Non-sig. |
| PF3D7_1249300 | PPM4 | 2878.44115 | 0.117604678 | 0.43423594 | 0.27083128 | 0.7865208 | 0.99028237 | 2212.78973 | 3038.21118 | 3309.31755 | 2953.44615 | Non-sig. |
| PF3D7_0709700 | PARE | 2035.88338 | 0.117571282 | 0.49005563 | 0.23991415 | 0.81039681 | 0.99028237 | 1260.10132 | 2161.94723 | 2645.47909 | 2076.00588 | Non-sig. |

| | | | | | | | | | | | |
|---|---|---|---|---|---|---|---|---|---|---|---|
| PF3D7_1225800 | UBA1 | 6616.28378 | 0.117101026 | 0.5811422 | 0.2015015 | 0.84030646 | 0.99128968 | 2812.93579 | 6754.92526 | 9882.48916 | 7014.78493 | Non-sig. |
| PF3D7_1335700 | COG3 | 2557.93568 | 0.116748226 | 0.41469689 | 0.28152665 | 0.77830649 | 0.99028237 | 2650.98029 | 2454.31078 | 2258.12304 | 2868.32862 | Non-sig. |
| PF3D7_0820800 | N/A | 595.319624 | 0.116548313 | 0.48330138 | 0.24115038 | 0.80943857 | 0.99028237 | 398.969025 | 599.786624 | 743.279876 | 639.242969 | Non-sig. |
| PF3D7_1138000 | N/A | 9520.63093 | 0.115876622 | 0.48308472 | 0.23986812 | 0.8104325 | 0.99028237 | 10021.678 | 6572.38181 | 8255.41029 | 13233.0536 | Non-sig. |
| PF3D7_1206800 | N/A | 550.275498 | 0.11579658 | 0.54769721 | 0.21142445 | 0.83255608 | 0.99110542 | 306.2424 | 458.193318 | 749.811788 | 686.854487 | Non-sig. |
| PF3D7_0625600 | PAP | 98.7094166 | 0.115685357 | 0.55413323 | 0.20876813 | 0.83462925 | 0.99128968 | 84.2746851 | 80.456248 | 105.139855 | 124.966879 | Non-sig. |
| PF3D7_0515300 | PI3K | 3644.01054 | 0.115647276 | 0.41987239 | 0.27543434 | 0.78298256 | 0.99028237 | 3402.08859 | 3247.8553 | 3593.9428 | 4332.15549 | Non-sig. |
| PF3D7_0922100 | N/A | 5053.51509 | 0.115325682 | 0.52451481 | 0.21987116 | 0.82597149 | 0.99096399 | 6996.10758 | 5336.19727 | 2707.58157 | 5174.17394 | Non-sig. |
| PF3D7_0525800 | IMC1g | 23448.4966 | 0.115194013 | 0.50466701 | 0.22825747 | 0.81944609 | 0.99096399 | 30886.6446 | 20414.5674 | 14139.4538 | 28353.3204 | Non-sig. |
| PF3D7_1021800 | SEA1 | 30868.1049 | 0.114589698 | 0.51806687 | 0.22118708 | 0.82494677 | 0.99096399 | 41675.1335 | 26921.7616 | 17611.0073 | 37264.5174 | Non-sig. |
| PF3D7_1125600 | C4AP1 | 473.437182 | 0.114532991 | 0.46591952 | 0.24582141 | 0.80582049 | 0.99028237 | 380.376936 | 576.477783 | 528.750138 | 408.143872 | Non-sig. |
| PF3D7_0613900 | MyoE | 47045.3122 | 0.114194644 | 0.53269236 | 0.21437259 | 0.83025651 | 0.99110542 | 66081.1897 | 47177.4207 | 24288.0473 | 50634.5911 | Non-sig. |
| PF3D7_0807200 | N/A | 321.180848 | 0.114181152 | 0.54975128 | 0.20769602 | 0.83546633 | 0.99128968 | 175.634127 | 355.244676 | 440.967701 | 312.876888 | Non-sig. |
| PF3D7_0816000 | RRB1 | 586.465154 | 0.113951232 | 0.42597934 | 0.26750413 | 0.78908104 | 0.99028237 | 599.522285 | 604.223044 | 527.198526 | 614.916762 | Non-sig. |
| PF3D7_1468500 | DER1-1 | 1490.46452 | 0.113472053 | 0.42277816 | 0.26839621 | 0.78839436 | 0.99028237 | 1300.64215 | 1701.98303 | 1562.99588 | 1396.23702 | Non-sig. |
| PF3D7_0726600 | N/A | 844.220455 | 0.113229472 | 0.43041749 | 0.26306894 | 0.79249745 | 0.99028237 | 845.322135 | 993.559917 | 776.955431 | 761.044337 | Non-sig. |
| PF3D7_0911700 | N/A | 1754.61497 | 0.113125036 | 0.40922775 | 0.27643539 | 0.78221367 | 0.99028237 | 1604.04601 | 1846.31069 | 1767.59976 | 1800.50341 | Non-sig. |
| PF3D7_1441300 | N/A | 3999.54064 | 0.112942834 | 0.44740586 | 0.25243933 | 0.80070151 | 0.99028237 | 4355.90785 | 3287.31517 | 3330.39189 | 5024.54766 | Non-sig. |
| PF3D7_0316500 | NUF2 | 2378.26671 | 0.112655237 | 0.42242564 | 0.26668655 | 0.78971051 | 0.99028237 | 2622.90794 | 2509.48394 | 1948.21811 | 2432.45688 | Non-sig. |
| PF3D7_1356400 | N/A | 886.572499 | 0.11264514 | 0.53455516 | 0.21072688 | 0.8331004 | 0.99110542 | 473.962646 | 933.338183 | 1229.60616 | 909.383008 | Non-sig. |
| PF3D7_0902400 | FIKK9.5 | 304.002552 | 0.112450726 | 0.49661448 | 0.22643465 | 0.82086338 | 0.99096399 | 208.009988 | 328.51412 | 376.02341 | 303.462689 | Non-sig. |
| PF3D7_0206900 | MSP5 | 6106.53135 | 0.112002767 | 0.60980698 | 0.18366921 | 0.85427297 | 0.99295786 | 9347.25176 | 7434.99512 | 2392.4066 | 5251.47191 | Non-sig. |
| PF3D7_1303200 | VAMP8 | 280.172481 | 0.111894755 | 0.46108121 | 0.24267906 | 0.80825403 | 0.99028237 | 234.432845 | 308.866029 | 304.012883 | 273.378167 | Non-sig. |
| PF3D7_0612900 | NOG1 | 1241.48392 | 0.111816095 | 0.42660592 | 0.26210629 | 0.7932395 | 0.99028237 | 1365.29715 | 1304.73446 | 1021.6966 | 1274.20746 | Non-sig. |
| PF3D7_0821000 | N/A | 2841.7958 | 0.111684733 | 0.44911348 | 0.2486782 | 0.80360971 | 0.99028237 | 2078.4664 | 2753.83773 | 3384.9451 | 3149.93395 | Non-sig. |
| PF3D7_0518400 | CYC3 | 20.854571 | 0.111358795 | 0.83933566 | 0.13267492 | 0.89445047 | 0.99457954 | 22.9295396 | 21.5190739 | 17.2194723 | 21.7501981 | Non-sig. |
| PF3D7_0719400 | N/A | 4090.89355 | 0.111340946 | 0.42265798 | 0.26343037 | 0.79221889 | 0.99028237 | 4416.31127 | 3813.29271 | 3450.08815 | 4683.88208 | Non-sig. |
| PF3D7_1331500 | N/A | 3197.53123 | 0.111191253 | 0.61185659 | 0.18172764 | 0.85579648 | 0.99321133 | 4948.42188 | 3373.44876 | 1200.7347 | 3267.51958 | Non-sig. |
| PF3D7_0320500 | Nico | 5536.14637 | 0.110707469 | 0.41442482 | 0.26713523 | 0.78936504 | 0.99028237 | 5915.82568 | 6083.32163 | 4732.01791 | 5413.42024 | Non-sig. |
| PF3D7_1348600 | N/A | 2488.20732 | 0.110161632 | 0.40821913 | 0.26985907 | 0.78726868 | 0.99028237 | 2264.23066 | 2675.25617 | 2522.20381 | 2491.13864 | Non-sig. |
| PF3D7_0614300 | MFR1 | 7672.85023 | 0.109943698 | 0.54962847 | 0.20003276 | 0.84145496 | 0.99128968 | 3741.02256 | 7449.50073 | 11019.834 | 8481.04357 | Non-sig. |
| PF3D7_0511700 | CGI121 | 524.056814 | 0.109887091 | 0.43285302 | 0.25386698 | 0.79959832 | 0.99028237 | 521.358756 | 594.670073 | 486.906477 | 493.29195 | Non-sig. |
| PF3D7_1366000 | N/A | 203.638403 | 0.109643066 | 0.48408788 | 0.22649414 | 0.82081712 | 0.99096399 | 160.878323 | 212.514733 | 230.740422 | 210.420136 | Non-sig. |
| PF3D7_0905400 | RhopH3 | 90673.1578 | 0.109306363 | 0.49521501 | 0.22072506 | 0.82530652 | 0.99096399 | 54189.0109 | 90768.5896 | 120290.438 | 97444.5924 | Non-sig. |
| PF3D7_1146900 | N/A | 248.348527 | 0.109058473 | 0.47753135 | 0.22837971 | 0.81935106 | 0.99096399 | 252.207908 | 305.943613 | 225.789984 | 209.452603 | Non-sig. |
| PF3D7_0932800 | CSE1 | 2783.03926 | 0.108939061 | 0.56976181 | 0.19120106 | 0.84836808 | 0.99165628 | 1273.89806 | 2586.23045 | 4081.7278 | 3190.30074 | Non-sig. |
| PF3D7_0108800 | N/A | 599.303442 | 0.108560699 | 0.42769133 | 0.25382955 | 0.79962724 | 0.99028237 | 545.801879 | 662.203864 | 607.658916 | 581.549109 | Non-sig. |
| PF3D7_0723900 | N/A | 1942.49084 | 0.108402715 | 0.45926157 | 0.23603698 | 0.81340397 | 0.99066994 | 2361.05629 | 1866.51557 | 1378.3495 | 2164.04201 | Non-sig. |
| PF3D7_0523100 | QCR2 | 1441.06437 | 0.10835484 | 0.49868467 | 0.21728127 | 0.82798915 | 0.99110542 | 883.015562 | 1615.98106 | 1890.57 | 1374.69088 | Non-sig. |
| PF3D7_0816200 | VPS2 | 1711.59241 | 0.108318917 | 0.49084588 | 0.22067806 | 0.82534312 | 0.99096399 | 2124.55242 | 2155.4692 | 1170.48908 | 1395.85893 | Non-sig. |
| PF3D7_0932700 | N/A | 174.070507 | 0.108290055 | 0.48783936 | 0.22197892 | 0.82433029 | 0.99096399 | 148.507123 | 163.066344 | 186.453048 | 198.255514 | Non-sig. |
| PF3D7_0624400 | N/A | 1820.40025 | 0.108106301 | 0.41517854 | 0.26038509 | 0.79456674 | 0.99028237 | 1885.14418 | 1759.99463 | 1619.41961 | 2017.04256 | Non-sig. |
| PF3D7_1021500 | ROK1 | 672.958406 | 0.107922075 | 0.52176513 | 0.20684034 | 0.83613455 | 0.99128968 | 908.325096 | 744.670206 | 387.644528 | 651.193796 | Non-sig. |
| PF3D7_0612200 | LRR6 | 9362.57976 | 0.107387804 | 0.53254736 | 0.2016493 | 0.8401909 | 0.99128968 | 13139.6331 | 9066.90143 | 4889.35488 | 10354.4296 | Non-sig. |
| PF3D7_1127800 | N/A | 1420.50576 | 0.10731842 | 0.49699114 | 0.21593628 | 0.82903741 | 0.99110542 | 1869.15405 | 1482.45941 | 866.611721 | 1463.79786 | Non-sig. |
| PF3D7_1315900 | N/A | 2782.40818 | 0.107318246 | 0.57093312 | 0.18796991 | 0.85090024 | 0.99165628 | 1253.71572 | 2725.77824 | 4103.814 | 3046.32478 | Non-sig. |
| PF3D7_1103700 | CK2beta1 | 1292.704 | 0.107154015 | 0.5319125 | 0.20145046 | 0.84034637 | 0.99128968 | 690.776757 | 1379.22928 | 1798.27274 | 1302.53721 | Non-sig. |
| PF3D7_1429300 | N/A | 169.830262 | 0.107105799 | 0.59899267 | 0.17880987 | 0.858087 | 0.99391013 | 242.821639 | 184.624371 | 84.5729861 | 167.302052 | Non-sig. |
| PF3D7_1459100 | N/A | 176.178392 | 0.10697128 | 0.49816207 | 0.21473188 | 0.82997636 | 0.99110542 | 137.895727 | 197.427909 | 201.217758 | 168.172174 | Non-sig. |
| PF3D7_1205300 | N/A | 539.727757 | 0.106417935 | 0.43658288 | 0.24375197 | 0.80742292 | 0.99028237 | 469.729058 | 605.449267 | 569.810072 | 513.922633 | Non-sig. |
| PF3D7_0613100 | N/A | 816.007025 | 0.106004272 | 0.43103651 | 0.24592875 | 0.80573738 | 0.99028237 | 696.888426 | 914.936995 | 875.048233 | 777.154447 | Non-sig. |
| PF3D7_1032900 | N/A | 289.122795 | 0.105984786 | 0.53106576 | 0.19956998 | 0.8418169 | 0.99128968 | 173.881364 | 279.958346 | 382.799652 | 319.851819 | Non-sig. |
| PF3D7_1367900 | N/A | 54.9859373 | 0.105806383 | 0.63340592 | 0.16704357 | 0.86733577 | 0.99402081 | 52.5673595 | 71.9143406 | 53.3728273 | 42.0892218 | Non-sig. |
| PF3D7_1333700 | CenH3 | 2723.43125 | 0.1057756 | 0.47855506 | 0.2210312 | 0.82506814 | 0.99096399 | 1760.65605 | 2889.71805 | 3486.26402 | 2757.08686 | Non-sig. |

| | | | | | | | | | | | | |
|---|---|---|---|---|---|---|---|---|---|---|---|---|
| PF3D7_0413600 | RPT3 | 7334.45274 | 0.105679202 | 0.40704942 | 0.25962253 | 0.79515495 | 0.99028237 | 6579.51301 | 7985.29332 | 7552.27888 | 7220.72576 | Non-sig. |
| PF3D7_1428900 | N/A | 2782.06908 | 0.105496049 | 0.41046812 | 0.25701399 | 0.79716797 | 0.99028237 | 2810.27897 | 2673.08623 | 2550.58493 | 3094.32619 | Non-sig. |
| PF3D7_0505100 | TRS85 | 5838.9718 | 0.104948131 | 0.42614976 | 0.24627054 | 0.80547281 | 0.99028237 | 4863.4067 | 5353.95258 | 6389.78545 | 6748.74245 | Non-sig. |
| PF3D7_1035900 | MAAP | 9575.78355 | 0.104777191 | 0.41404746 | 0.25305599 | 0.80022495 | 0.99028237 | 10243.7551 | 9258.31846 | 8212.83142 | 10588.2292 | Non-sig. |
| PF3D7_1232800 | N/A | 30.3735422 | 0.104124549 | 0.73374191 | 0.14190896 | 0.88715192 | 0.99457954 | 33.2137115 | 32.3150575 | 25.4016687 | 30.563731 | Non-sig. |
| PF3D7_1329700 | ACBP1 | 756.574786 | 0.103747608 | 0.47644757 | 0.21775241 | 0.82762202 | 0.99110542 | 940.021248 | 733.913609 | 519.061085 | 833.303203 | Non-sig. |
| PF3D7_0705500 | N/A | 29630.4763 | 0.103611508 | 0.47762253 | 0.21693178 | 0.82826151 | 0.99110542 | 20547.1156 | 24756.6011 | 36586.4709 | 36631.7177 | Non-sig. |
| PF3D7_0912900 | RPN8 | 3006.88934 | 0.103148643 | 0.40917324 | 0.2520904 | 0.80097119 | 0.99028237 | 2727.02407 | 3284.50373 | 3071.7778 | 2944.25177 | Non-sig. |
| PF3D7_0801000 | N/A | 1747.41656 | 0.102886306 | 0.4618131 | 0.22278776 | 0.8237007 | 0.99096399 | 1257.18632 | 1599.31253 | 2112.78283 | 2020.38455 | Non-sig. |
| PF3D7_1237000 | UBA2 | 1106.90637 | 0.102602878 | 0.41965995 | 0.24449052 | 0.80685094 | 0.99028237 | 1112.66775 | 1250.01019 | 1022.51925 | 1042.42829 | Non-sig. |
| PF3D7_1321800 | N/A | 1267.75516 | 0.102160743 | 0.48075667 | 0.2124999 | 0.83171706 | 0.99110542 | 829.209625 | 1353.14889 | 1616.22045 | 1272.44168 | Non-sig. |
| PF3D7_1221600 | N/A | 416.192154 | 0.10179821 | 0.46356019 | 0.21960084 | 0.82618203 | 0.99096953 | 317.50746 | 451.879419 | 485.282331 | 410.099407 | Non-sig. |
| PF3D7_1127500 | N/A | 1323.16165 | 0.100756276 | 0.45618208 | 0.22086855 | 0.82519478 | 0.99096399 | 961.37818 | 1457.41661 | 1592.28223 | 1281.56958 | Non-sig. |
| PF3D7_0207500 | SERA6 | 16162.2407 | 0.100519764 | 0.53558878 | 0.18768086 | 0.85112683 | 0.99170221 | 8296.08217 | 17174.7114 | 22902.3656 | 16275.8037 | Non-sig. |
| PF3D7_0708500 | N/A | 1124.36217 | 0.100467255 | 0.63884008 | 0.15726511 | 0.87503592 | 0.99402081 | 1784.82117 | 1234.881 | 386.032161 | 1091.71437 | Non-sig. |
| PF3D7_1104600 | N/A | 170.714219 | 0.100356295 | 0.54708253 | 0.18343904 | 0.85445355 | 0.99295786 | 223.93176 | 180.323701 | 105.924103 | 172.677311 | Non-sig. |
| PF3D7_0905700 | ATG3 | 742.333776 | 0.100193209 | 0.46378694 | 0.21603284 | 0.82896215 | 0.99110542 | 539.889096 | 825.250187 | 892.946739 | 711.249082 | Non-sig. |
| PF3D7_1127900 | N/A | 1612.25845 | 0.100181772 | 0.5932719 | 0.16886317 | 0.86590427 | 0.99402081 | 2441.28006 | 1785.82476 | 671.729718 | 1550.19926 | Non-sig. |
| PF3D7_0503400 | ADF1 | 15589.0698 | 0.100021532 | 0.44490001 | 0.224818 | 0.82212086 | 0.99096399 | 18405.1799 | 17831.1696 | 11692.9049 | 14427.0248 | Non-sig. |
| PF3D7_0208400 | N/A | 4489.44072 | 0.099823644 | 0.59265439 | 0.16843483 | 0.86624121 | 0.99402081 | 6834.35824 | 4527.15406 | 1834.42897 | 4761.82161 | Non-sig. |
| PF3D7_1133600 | N/A | 96.1217842 | 0.099323139 | 0.59992865 | 0.16555825 | 0.8685046 | 0.99402081 | 58.9097557 | 85.8538065 | 126.465456 | 113.258119 | Non-sig. |
| PF3D7_1037900 | N/A | 225.380484 | 0.099188575 | 0.48315592 | 0.2052931 | 0.83734314 | 0.99128968 | 253.183147 | 259.805494 | 182.26453 | 206.268764 | Non-sig. |
| PF3D7_1129200 | RPN7 | 5967.41365 | 0.09887024 | 0.41797058 | 0.23654832 | 0.81300721 | 0.99055243 | 6044.00473 | 7005.12164 | 5482.10405 | 5338.42417 | Non-sig. |
| PF3D7_0217500 | CDPK1 | 11914.9175 | 0.098561797 | 0.43705261 | 0.22551472 | 0.82157887 | 0.99096399 | 9721.37522 | 10377.5728 | 13294.5788 | 14266.1433 | Non-sig. |
| PF3D7_1342100 | IRP | 2824.53086 | 0.09790914 | 0.50115768 | 0.19536594 | 0.84510648 | 0.99165628 | 1746.74592 | 2484.42764 | 3710.34358 | 3356.6063 | Non-sig. |
| PF3D7_1412500 | ACT2 | 396.451624 | 0.097282769 | 0.45094815 | 0.21572939 | 0.82919869 | 0.99110542 | 325.018739 | 426.063106 | 440.977926 | 393.746726 | Non-sig. |
| PF3D7_1108500 | SCS-alpha | 3164.88152 | 0.09692392 | 0.45256394 | 0.21416625 | 0.83041742 | 0.99110542 | 2360.95391 | 3688.79989 | 3755.99433 | 2853.77797 | Non-sig. |
| PF3D7_0916900 | N/A | 1548.36645 | 0.096575176 | 0.43872878 | 0.220125 | 0.8257738 | 0.99096399 | 1781.12936 | 1710.30929 | 1212.24193 | 1489.78521 | Non-sig. |
| PF3D7_1471000 | RCL1 | 217.034455 | 0.096552582 | 0.47473338 | 0.20338275 | 0.83883588 | 0.99128968 | 178.785029 | 218.943785 | 240.601498 | 229.807509 | Non-sig. |
| PF3D7_1015700 | N/A | 175.663188 | 0.096549272 | 0.49976145 | 0.19319072 | 0.84680961 | 0.99165628 | 135.474408 | 191.554504 | 203.860069 | 171.76377 | Non-sig. |
| PF3D7_1412800 | NMT | 1822.27897 | 0.096495547 | 0.42441551 | 0.22736103 | 0.82014302 | 0.99096399 | 1539.08709 | 2036.4639 | 1983.46116 | 1730.10372 | Non-sig. |
| PF3D7_1416100 | SEY1 | 2632.64912 | 0.096415889 | 0.50877475 | 0.18950604 | 0.84969622 | 0.99165628 | 1540.74553 | 2479.31893 | 3548.29831 | 2962.23372 | Non-sig. |
| PF3D7_1243000 | STX16 | 1052.94546 | 0.096367942 | 0.48523845 | 0.19859915 | 0.84257632 | 0.99128968 | 1348.75396 | 1155.20561 | 687.172618 | 1020.64966 | Non-sig. |
| PF3D7_1132100 | N/A | 35.4659732 | 0.09629209 | 0.743196 | 0.12956487 | 0.8969107 | 0.99457954 | 47.3828121 | 35.4053115 | 21.3548274 | 37.7209419 | Non-sig. |
| PF3D7_1140500 | MyoK | 535.023379 | 0.095239631 | 0.51333066 | 0.18553271 | 0.85281122 | 0.99257334 | 712.272963 | 549.215009 | 322.824207 | 555.781339 | Non-sig. |
| PF3D7_1309400 | N/A | 2978.604 | 0.095005369 | 0.46060195 | 0.2062635 | 0.83658509 | 0.99128968 | 3591.25626 | 2642.16221 | 2170.16298 | 3510.83455 | Non-sig. |
| PF3D7_1223300 | GyrA | 1888.16192 | 0.094930393 | 0.40702602 | 0.2332293 | 0.81558336 | 0.99096399 | 1828.6319 | 1907.38726 | 1823.49697 | 1993.13156 | Non-sig. |
| PF3D7_0728600 | N/A | 5279.93468 | 0.094872458 | 0.60715742 | 0.15625677 | 0.87583063 | 0.99402081 | 8188.21511 | 5407.84839 | 2024.98518 | 5498.69005 | Non-sig. |
| PF3D7_0934700 | N/A | 3929.21848 | 0.09478781 | 0.46887195 | 0.2021614 | 0.83979055 | 0.99128968 | 4926.45697 | 3709.0387 | 2674.25279 | 4407.12548 | Non-sig. |
| PF3D7_1352100 | ABCB6 | 2895.19395 | 0.094773457 | 0.42384025 | 0.22360655 | 0.82306346 | 0.99096399 | 2359.62186 | 2923.26786 | 3240.42658 | 3057.45952 | Non-sig. |
| PF3D7_0509100 | SMC4 | 8400.39069 | 0.094659134 | 0.58095464 | 0.16293722 | 0.87056787 | 0.99402081 | 12650.5784 | 8856.67865 | 3599.65314 | 8494.65262 | Non-sig. |
| PF3D7_0928500 | N/A | 234.546795 | 0.094454921 | 0.55661781 | 0.16969439 | 0.86525048 | 0.99402081 | 318.582104 | 275.844713 | 135.486652 | 208.27371 | Non-sig. |
| PF3D7_0517300 | SR1 | 4531.76215 | 0.093742652 | 0.41684092 | 0.22488832 | 0.82206616 | 0.99096399 | 3795.39665 | 4716.92519 | 4973.57953 | 4641.14722 | Non-sig. |
| PF3D7_0824300 | Obg2 | 816.065357 | 0.093380142 | 0.43806519 | 0.21316494 | 0.83119831 | 0.99110542 | 922.582116 | 869.266277 | 656.96506 | 815.447975 | Non-sig. |
| PF3D7_1222700 | GAP45 | 13731.9575 | 0.093114915 | 0.45170961 | 0.20613888 | 0.83668243 | 0.99128968 | 16811.7515 | 14402.1379 | 9766.5139 | 13947.4267 | Non-sig. |
| PF3D7_1362000 | ApiCOX24 | 605.188341 | 0.092931458 | 0.52786957 | 0.17605004 | 0.86025463 | 0.99402081 | 342.592501 | 651.600829 | 828.448262 | 598.111772 | Non-sig. |
| PF3D7_1021000 | N/A | 1034.56095 | 0.092834384 | 0.42863243 | 0.21658274 | 0.82853354 | 0.99110542 | 967.288684 | 1217.53113 | 1035.24453 | 918.179456 | Non-sig. |
| PF3D7_1020400 | RCM1 | 228.834088 | 0.092629183 | 0.47647876 | 0.19440359 | 0.84585988 | 0.99165628 | 256.735613 | 252.179701 | 186.420661 | 220.000378 | Non-sig. |
| PF3D7_1356800 | ARK3 | 25950.6155 | 0.092339606 | 0.5404343 | 0.17086186 | 0.86433239 | 0.99402081 | 35662.0805 | 19945.4937 | 14579.1437 | 33615.7443 | Non-sig. |
| PF3D7_0209900 | N/A | 16.2939867 | 0.09229674 | 0.92990954 | 0.09925346 | 0.92093703 | 0.99619209 | 13.4802537 | 14.0071285 | 18.0236031 | 19.6649613 | Non-sig. |
| PF3D7_0717100 | N/A | 1609.04734 | 0.091981386 | 0.44278647 | 0.20773306 | 0.83543741 | 0.99128968 | 1221.29472 | 1625.47393 | 1893.97403 | 1695.44669 | Non-sig. |
| PF3D7_0619900 | SF3A2 | 886.385415 | 0.091596451 | 0.42814276 | 0.21393904 | 0.83059459 | 0.99110542 | 837.892831 | 1032.57182 | 878.59317 | 796.483834 | Non-sig. |
| PF3D7_1234700 | ULG8 | 132.943828 | 0.091390835 | 0.509219 | 0.17947255 | 0.85756667 | 0.99352464 | 140.092357 | 119.07867 | 117.45884 | 155.145443 | Non-sig. |
| PF3D7_0917000 | MOP | 4493.95947 | 0.090671374 | 0.43532369 | 0.20828495 | 0.83500648 | 0.99128968 | 3465.17283 | 4432.06852 | 5240.13405 | 4838.46249 | Non-sig. |

| | | | | | | | | | | | | |
|---|---|---|---|---|---|---|---|---|---|---|---|---|
| PF3D7_1138500 | PPM2 | 6102.97794 | 0.09040652 | 0.48836001 | 0.18512269 | 0.8531328 | 0.99272954 | 3980.28485 | 5291.43661 | 7843.01527 | 7297.17502 | Non-sig. |
| PF3D7_0719900 | N/A | 6316.19295 | 0.090102297 | 0.44810041 | 0.20107613 | 0.84063904 | 0.99128968 | 4932.00399 | 5446.16671 | 7305.79841 | 7580.80268 | Non-sig. |
| PF3D7_1249800 | THO2 | 4572.92674 | 0.08956627 | 0.43782792 | 0.20456957 | 0.83790843 | 0.99128968 | 4351.90567 | 3694.82657 | 4510.11551 | 5734.85921 | Non-sig. |
| PF3D7_1020100 | N/A | 200.979242 | 0.089209658 | 0.67642743 | 0.13188356 | 0.89507639 | 0.99457954 | 319.308357 | 239.640881 | 70.5691512 | 174.39858 | Non-sig. |
| PF3D7_0926300 | N/A | 966.727743 | 0.089173782 | 0.49604564 | 0.17976931 | 0.85733368 | 0.99352464 | 1266.23743 | 1054.51081 | 607.840646 | 938.322084 | Non-sig. |
| PF3D7_1422400 | N/A | 757.519752 | 0.089050857 | 0.43294323 | 0.20568715 | 0.8370353 | 0.99128968 | 649.148559 | 715.451755 | 818.992229 | 846.486465 | Non-sig. |
| PF3D7_0809800 | N/A | 246.785129 | 0.088940481 | 0.5356123 | 0.16605384 | 0.86811458 | 0.99402081 | 327.549777 | 274.640056 | 151.133065 | 233.817619 | Non-sig. |
| PF3D7_1311200 | N/A | 840.770268 | 0.08833169 | 0.42300093 | 0.2088215 | 0.83458758 | 0.99128968 | 739.723926 | 842.377665 | 890.226722 | 890.752757 | Non-sig. |
| PF3D7_0630700 | N/A | 365.916394 | 0.088221023 | 0.44258176 | 0.19933272 | 0.84200249 | 0.99128968 | 385.173079 | 363.764127 | 324.398637 | 390.329734 | Non-sig. |
| PF3D7_1220300 | N/A | 3308.99274 | 0.088165371 | 0.45992764 | 0.19169401 | 0.8479819 | 0.99165628 | 2727.10242 | 2589.11828 | 3688.54949 | 4231.20075 | Non-sig. |
| PF3D7_1330800 | N/A | 2761.76748 | 0.087828043 | 0.55513655 | 0.1582098 | 0.87429148 | 0.99402081 | 3981.78244 | 3254.1803 | 1374.07211 | 2437.03505 | Non-sig. |
| PF3D7_0513200 | N/A | 4630.99484 | 0.08773418 | 0.40913471 | 0.21443837 | 0.83020522 | 0.99110542 | 4688.12002 | 4385.55185 | 4292.40004 | 5157.90743 | Non-sig. |
| PF3D7_1332200 | N/A | 9705.63706 | 0.087124022 | 0.54953121 | 0.15854245 | 0.87402938 | 0.99402081 | 13849.3708 | 8224.05662 | 4976.3578 | 11772.763 | Non-sig. |
| PF3D7_0214200 | mRPL13 | 772.653835 | 0.08639215 | 0.44882934 | 0.1924833 | 0.84736365 | 0.99165628 | 609.816993 | 873.040159 | 889.01322 | 718.744969 | Non-sig. |
| PF3D7_1326800 | SYN18 | 403.391371 | 0.086234615 | 0.45515917 | 0.18946035 | 0.84973203 | 0.99165628 | 322.665743 | 403.613372 | 459.802972 | 427.483399 | Non-sig. |
| PF3D7_0317200 | CRK4 | 26096.3666 | 0.08616352 | 0.49481528 | 0.1741327 | 0.86176118 | 0.99402081 | 15980.8395 | 24371.296 | 34653.4324 | 29379.8986 | Non-sig. |
| PF3D7_1034000 | SLY1 | 1046.0515 | 0.085684272 | 0.57709618 | 0.14847486 | 0.88196803 | 0.99457954 | 504.746044 | 848.971565 | 1524.87327 | 1305.61513 | Non-sig. |
| PF3D7_1439300 | N/A | 3187.27659 | 0.0855947 | 0.42897858 | 0.19953141 | 0.84184708 | 0.99128968 | 3567.66858 | 2962.51533 | 2618.04772 | 3600.87472 | Non-sig. |
| PF3D7_1136700 | AIP | 3510.32 | 0.085433665 | 0.48892352 | 0.1747383 | 0.86128527 | 0.99402081 | 4507.3199 | 4201.89327 | 2306.04212 | 3026.02473 | Non-sig. |
| PF3D7_1471700 | N/A | 93.7116271 | 0.08515809 | 0.56914016 | 0.14962587 | 0.8810598 | 0.99421828 | 115.592703 | 89.0999286 | 66.5051518 | 103.648726 | Non-sig. |
| PF3D7_0108400 | N/A | 186.688845 | 0.084457584 | 0.52342421 | 0.1613559 | 0.8718131 | 0.99402081 | 127.502496 | 206.848489 | 234.729322 | 177.675074 | Non-sig. |
| PF3D7_0812200 | N/A | 51.1811495 | 0.084437927 | 0.68570467 | 0.12314037 | 0.90199594 | 0.99457954 | 29.3838363 | 55.842838 | 69.7727482 | 49.7251756 | Non-sig. |
| PF3D7_0907200 | N/A | 2808.38657 | 0.0841984 | 0.43647703 | 0.19290454 | 0.84703373 | 0.99165628 | 3247.58694 | 2659.13857 | 2205.57981 | 3121.24098 | Non-sig. |
| PF3D7_0403200 | N/A | 1187.50142 | 0.083370816 | 0.66670681 | 0.12504869 | 0.900485 | 0.99457954 | 1933.07485 | 1453.794 | 373.704623 | 989.432196 | Non-sig. |
| PF3D7_1143000 | N/A | 387.732469 | 0.083028145 | 0.47585882 | 0.17448063 | 0.86148775 | 0.99402081 | 282.481562 | 402.637995 | 470.405248 | 395.405073 | Non-sig. |
| PF3D7_0807800 | RPN10 | 4930.60439 | 0.08295319 | 0.40882184 | 0.20290792 | 0.839207 | 0.99128968 | 5193.91044 | 4898.69427 | 4383.99972 | 5245.81314 | Non-sig. |
| PF3D7_0802700 | N/A | 117.98903 | 0.082792825 | 0.67875626 | 0.12197725 | 0.90291703 | 0.99457954 | 183.565082 | 129.958645 | 45.9642212 | 112.468171 | Non-sig. |
| PF3D7_0623000 | CS | 784.315617 | 0.082496006 | 0.52170161 | 0.15812872 | 0.87435537 | 0.99402081 | 449.257607 | 780.161928 | 1074.16477 | 833.678165 | Non-sig. |
| PF3D7_0210200 | N/A | 8520.71002 | 0.081420339 | 0.4048424 | 0.20111614 | 0.84060777 | 0.99128968 | 8430.25676 | 8167.36729 | 8130.44572 | 9354.7703 | Non-sig. |
| PF3D7_1319300 | TRM1 | 626.948045 | 0.081413647 | 0.43299216 | 0.18802568 | 0.85085652 | 0.99165628 | 539.122486 | 613.688421 | 679.250281 | 675.730992 | Non-sig. |
| PF3D7_1358900 | N/A | 681.639486 | 0.081359704 | 0.45881713 | 0.17732491 | 0.85925319 | 0.99402081 | 525.663451 | 783.548558 | 798.947465 | 618.39847 | Non-sig. |
| PF3D7_0426000 | VAR | 108.83301 | 0.080754154 | 0.60405035 | 0.13368779 | 0.89364946 | 0.99457954 | 148.600328 | 128.713847 | 63.2531032 | 94.7647634 | Non-sig. |
| PF3D7_0111800 | N/A | 1274.99504 | 0.08047768 | 0.41160363 | 0.19552228 | 0.8449841 | 0.99165628 | 1290.38461 | 1299.12049 | 1188.56172 | 1321.91333 | Non-sig. |
| PF3D7_1404500 | RRP5 | 776.937821 | 0.08002874 | 0.49942626 | 0.16024135 | 0.87269095 | 0.99402081 | 1020.08389 | 858.613122 | 491.053401 | 738.000869 | Non-sig. |
| PF3D7_1118700 | MLC-B | 7278.53484 | 0.079172613 | 0.57910123 | 0.13671636 | 0.891255 | 0.99457954 | 10960.7049 | 8150.84046 | 3197.4483 | 6805.14569 | Non-sig. |
| PF3D7_1400700 | N/A | 5.74917349 | 0.079138774 | 1.49892862 | 0.05279689 | 0.95789374 | 0.99691622 | 2.09147134 | 5.37480212 | 9.02462679 | 6.5057937 | Non-sig. |
| PF3D7_1362800 | N/A | 3389.98439 | 0.078996009 | 0.52543754 | 0.15034329 | 0.88049378 | 0.99421828 | 1850.7827 | 3270.34013 | 4743.21881 | 3695.59594 | Non-sig. |
| PF3D7_1356300 | N/A | 4158.62349 | 0.078984333 | 0.41347131 | 0.19102736 | 0.84850416 | 0.99165628 | 4310.30698 | 4682.33253 | 3779.42225 | 3862.43222 | Non-sig. |
| PF3D7_0606200 | PEX4 | 1962.34525 | 0.078825092 | 0.41383592 | 0.19047426 | 0.84893751 | 0.99165628 | 1939.64118 | 2207.90778 | 1877.88637 | 1823.94565 | Non-sig. |
| PF3D7_0203600 | N/A | 250.348033 | 0.078520377 | 0.46867927 | 0.16753542 | 0.86694878 | 0.99402081 | 235.729086 | 217.830623 | 251.305306 | 296.527117 | Non-sig. |
| PF3D7_1116600 | STT3 | 1561.60475 | 0.078389297 | 0.55375013 | 0.14156077 | 0.88742695 | 0.99457954 | 767.732286 | 1596.38117 | 2270.24989 | 1612.05564 | Non-sig. |
| PF3D7_1464400 | N/A | 747.562859 | 0.077874879 | 0.50305694 | 0.15480331 | 0.87697638 | 0.99421828 | 917.267071 | 970.473036 | 537.776353 | 564.734975 | Non-sig. |
| PF3D7_1363600 | N/A | 145.071921 | 0.077867587 | 0.49882369 | 0.15610242 | 0.87595229 | 0.99402081 | 126.370187 | 162.377115 | 155.848524 | 135.691858 | Non-sig. |
| PF3D7_1225900 | N/A | 891.158024 | 0.077832421 | 0.43168031 | 0.18030107 | 0.85691622 | 0.99352464 | 741.099439 | 885.192717 | 992.95775 | 945.382188 | Non-sig. |
| PF3D7_0812400 | KARalpha | 4908.05608 | 0.07779709 | 0.53847013 | 0.144478 | 0.88512302 | 0.99457954 | 2524.69302 | 5167.19349 | 7026.42136 | 4913.91647 | Non-sig. |
| PF3D7_1439600 | NDUFA4 | 1643.11342 | 0.077664893 | 0.46482586 | 0.16708385 | 0.86730407 | 0.99402081 | 1211.05037 | 1948.04846 | 1986.47597 | 1426.87888 | Non-sig. |
| PF3D7_1010100 | N/A | 382.847671 | 0.077258266 | 0.44335576 | 0.17425795 | 0.86166275 | 0.99402081 | 412.958281 | 387.813488 | 332.373517 | 398.245397 | Non-sig. |
| PF3D7_0903600 | N/A | 10004.2312 | 0.077253009 | 0.81400749 | 0.09490454 | 0.92439065 | 0.99619209 | 18160.8128 | 11789.9473 | 1312.42789 | 8753.73672 | Non-sig. |
| PF3D7_1207400 | N/A | 77.267554 | 0.077173894 | 0.76388645 | 0.10102797 | 0.91952826 | 0.99619209 | 126.889907 | 89.1515365 | 23.801832 | 69.2269402 | Non-sig. |
| PF3D7_1108700 | Pfj2 | 4681.84704 | 0.077088469 | 0.52102913 | 0.14795424 | 0.88237888 | 0.99457954 | 2585.37757 | 4607.33488 | 6527.81718 | 5006.85851 | Non-sig. |
| PF3D7_1221300 | N/A | 3393.77425 | 0.076999029 | 0.45042101 | 0.17094902 | 0.86426385 | 0.99402081 | 4142.06693 | 3459.29632 | 2464.70513 | 3509.02863 | Non-sig. |
| PF3D7_0726800 | OST2 | 33.1769491 | 0.076452345 | 0.74317987 | 0.10287193 | 0.91806462 | 0.99619209 | 21.8354164 | 33.3759162 | 42.6227634 | 34.8737004 | Non-sig. |
| PF3D7_1437000 | N/A | 2430.89624 | 0.076281954 | 0.50373157 | 0.15143374 | 0.87963358 | 0.99421828 | 3290.79513 | 2473.60689 | 1442.87618 | 2516.30674 | Non-sig. |
| PF3D7_0217600 | N/A | 1606.68189 | 0.076082257 | 0.41861626 | 0.18174702 | 0.85578127 | 0.99321133 | 1637.75383 | 1473.77645 | 1490.96418 | 1824.23311 | Non-sig. |

| | | | | | | | | | | | | |
|---|---|---|---|---|---|---|---|---|---|---|---|---|
| PF3D7_0304700 | N/A | 70.439971 | 0.075805951 | 0.70466173 | 0.10757779 | 0.91433062 | 0.99619209 | 107.083572 | 78.3860303 | 30.3737898 | 65.9164923 | Non-sig. |
| PF3D7_1473500 | N/A | 342.623024 | 0.075756364 | 0.51729047 | 0.14644841 | 0.88356742 | 0.99457954 | 224.258564 | 297.197527 | 442.695078 | 406.340926 | Non-sig. |
| PF3D7_0411200 | N/A | 415.423904 | 0.07562782 | 0.43393763 | 0.1742827 | 0.8616433 | 0.99402081 | 406.597749 | 405.507241 | 402.483742 | 447.106883 | Non-sig. |
| PF3D7_1343900 | PRPF4 | 781.733679 | 0.075574908 | 0.47706287 | 0.15841708 | 0.87412816 | 0.99402081 | 535.818559 | 773.606719 | 986.38578 | 831.123658 | Non-sig. |
| PF3D7_1477700 | Pfg14-748 | 526.462883 | 0.075308179 | 0.54932106 | 0.13709319 | 0.89095713 | 0.99457954 | 742.514931 | 601.015049 | 283.304192 | 479.017361 | Non-sig. |
| PF3D7_1143600 | N/A | 156.506685 | 0.075006796 | 0.78581414 | 0.09545106 | 0.92395656 | 0.99619209 | 270.712976 | 199.75342 | 34.4750906 | 121.085252 | Non-sig. |
| PF3D7_0602700 | N/A | 51.5148772 | 0.074315992 | 0.67537094 | 0.1100373 | 0.9123798 | 0.99619209 | 69.3810722 | 53.6691231 | 31.2019573 | 51.8073561 | Non-sig. |
| PF3D7_1312400 | IF2a | 318.710431 | 0.074055291 | 0.44353937 | 0.16696441 | 0.86739805 | 0.99402081 | 299.050179 | 315.425581 | 321.967342 | 338.398621 | Non-sig. |
| PF3D7_0413900 | USP13 | 4323.10786 | 0.073745367 | 0.40758876 | 0.18093082 | 0.85642188 | 0.99350189 | 4545.55695 | 4463.51537 | 3879.84584 | 4403.51329 | Non-sig. |
| PF3D7_1029100 | N/A | 307.489327 | 0.073718202 | 0.45364073 | 0.16250349 | 0.87090938 | 0.99402081 | 321.711138 | 283.975089 | 277.646571 | 346.624511 | Non-sig. |
| PF3D7_0709100 | N/A | 3308.92674 | 0.0735087 | 0.42802494 | 0.17173929 | 0.8636425 | 0.99402081 | 2653.39402 | 3454.33009 | 3795.66188 | 3332.32098 | Non-sig. |
| PF3D7_0530200 | PPT | 1099.30641 | 0.07350547 | 0.56682948 | 0.12967828 | 0.89682097 | 0.99457954 | 552.352873 | 899.284781 | 1589.88869 | 1355.6993 | Non-sig. |
| PF3D7_1419900 | YTH1 | 292.224249 | 0.07349898 | 0.452722 | 0.16234903 | 0.871031 | 0.99402081 | 256.585735 | 290.859238 | 312.859071 | 308.592953 | Non-sig. |
| PF3D7_0812600 | N/A | 396.236931 | 0.072894809 | 0.47549927 | 0.15330162 | 0.87816042 | 0.99421828 | 295.199277 | 442.315997 | 477.005927 | 370.426525 | Non-sig. |
| PF3D7_1401900 | N/A | 1028.87209 | 0.072794492 | 0.60534416 | 0.12025307 | 0.90428268 | 0.99457954 | 1571.01671 | 1254.67599 | 435.20859 | 854.587051 | Non-sig. |
| PF3D7_0515200 | N/A | 58.364781 | 0.072672157 | 0.61982715 | 0.11724584 | 0.90666524 | 0.99546099 | 43.2917327 | 58.2985863 | 70.356151 | 61.5126538 | Non-sig. |
| PF3D7_1434600 | METAP2 | 1237.31046 | 0.072502822 | 0.47529991 | 0.15254121 | 0.87876009 | 0.99421828 | 1576.63212 | 1235.88694 | 836.157308 | 1300.56549 | Non-sig. |
| PF3D7_1029400 | CDC50C | 1312.34979 | 0.072274324 | 0.44193554 | 0.16354042 | 0.87009295 | 0.99402081 | 1016.42614 | 1301.67447 | 1542.28344 | 1389.01511 | Non-sig. |
| PF3D7_1323800 | VPS52 | 1123.38916 | 0.072015283 | 0.43483482 | 0.16561526 | 0.86845973 | 0.99402081 | 1287.38691 | 1147.8196 | 903.556771 | 1154.79337 | Non-sig. |
| PF3D7_0621500 | RPP1 | 214.384224 | 0.071819488 | 0.48348947 | 0.14854406 | 0.88191342 | 0.99457954 | 170.004926 | 213.706794 | 247.89807 | 225.927106 | Non-sig. |
| PF3D7_1461600 | SF3B2 | 1518.54567 | 0.071721521 | 0.50190415 | 0.14289884 | 0.88637008 | 0.99457954 | 2038.85659 | 1637.38789 | 923.129456 | 1474.80873 | Non-sig. |
| PF3D7_1121900 | PDK1 | 1049.29661 | 0.071658955 | 0.42395457 | 0.16902508 | 0.86577691 | 0.99402081 | 1122.00266 | 996.013584 | 924.612691 | 1154.5575 | Non-sig. |
| PF3D7_0317500 | EG5 | 2792.5729 | 0.071582668 | 0.43690796 | 0.16383924 | 0.8698577 | 0.99402081 | 2729.25641 | 2266.51874 | 2717.35793 | 3457.15852 | Non-sig. |
| PF3D7_1236900 | N/A | 3598.09548 | 0.071536439 | 0.41085107 | 0.17411769 | 0.86177297 | 0.99402081 | 3765.27495 | 3927.55787 | 3252.65128 | 3446.89782 | Non-sig. |
| PF3D7_1016300 | GBP130 | 34462.6055 | 0.071285533 | 0.40261076 | 0.17705819 | 0.85946268 | 0.99402081 | 34274.5555 | 33278.0234 | 32948.2149 | 37349.6282 | Non-sig. |
| PF3D7_1465100 | COG6 | 888.949674 | 0.070951972 | 0.43237076 | 0.16409984 | 0.86965255 | 0.99402081 | 740.184363 | 873.34445 | 993.811329 | 948.458555 | Non-sig. |
| PF3D7_0104900 | N/A | 132.806356 | 0.070907359 | 0.49962279 | 0.14192179 | 0.88714178 | 0.99457954 | 118.136802 | 137.859313 | 140.87068 | 134.35863 | Non-sig. |
| PF3D7_1026900 | HCS1 | 48.8749479 | 0.070656523 | 0.68637632 | 0.10294138 | 0.9180095 | 0.99619209 | 34.5105169 | 64.4972287 | 60.7114186 | 35.7806273 | Non-sig. |
| PF3D7_0317300 | N/A | 12488.4518 | 0.070490333 | 0.43054709 | 0.1637227 | 0.86994944 | 0.99402081 | 12676.5218 | 10311.9487 | 11690.3759 | 15274.9607 | Non-sig. |
| PF3D7_1361400 | ADF2 | 1042.33182 | 0.070307755 | 0.44639449 | 0.15750139 | 0.87484971 | 0.99402081 | 1211.30825 | 1175.11688 | 822.804384 | 960.097774 | Non-sig. |
| PF3D7_1432900 | N/A | 766.276229 | 0.07015041 | 0.45970922 | 0.15259735 | 0.87871581 | 0.99421828 | 595.971835 | 674.10791 | 899.089272 | 895.935901 | Non-sig. |
| PF3D7_1326100 | N/A | 2113.74123 | 0.069635858 | 0.41257319 | 0.16878425 | 0.86596634 | 0.99402081 | 2216.88882 | 2278.99587 | 1908.69879 | 2050.38145 | Non-sig. |
| PF3D7_0607800 | N/A | 562.876966 | 0.069576848 | 0.43033262 | 0.16168156 | 0.87155663 | 0.99402081 | 507.212105 | 607.359289 | 591.301225 | 545.635246 | Non-sig. |
| PF3D7_1350800 | N/A | 80.8711367 | 0.06817792 | 0.62539964 | 0.10901497 | 0.91319062 | 0.99619209 | 110.600071 | 90.3523618 | 47.5677046 | 74.96441 | Non-sig. |
| PF3D7_0720400 | AIF | 1345.81128 | 0.067860683 | 0.42890118 | 0.15821985 | 0.87428356 | 0.99402081 | 1125.18713 | 1470.01046 | 1502.95383 | 1285.09371 | Non-sig. |
| PF3D7_1307400 | N/A | 45.8983111 | 0.067778007 | 0.73094249 | 0.09272687 | 0.92612055 | 0.99619209 | 66.0570664 | 53.7079285 | 23.8012218 | 40.0270276 | Non-sig. |
| PF3D7_1451300 | NSE2 | 1401.78995 | 0.067683949 | 0.42610103 | 0.15884484 | 0.87379112 | 0.99402081 | 1472.19627 | 1617.33883 | 1265.73793 | 1251.88679 | Non-sig. |
| PF3D7_1119600 | N/A | 1247.78662 | 0.067601336 | 0.49089952 | 0.13770911 | 0.89047032 | 0.99457954 | 793.155964 | 1279.82683 | 1643.60291 | 1274.56077 | Non-sig. |
| PF3D7_1218200 | N/A | 2573.22053 | 0.067555374 | 0.43389253 | 0.1556961 | 0.87627257 | 0.99402081 | 2103.75142 | 2342.54152 | 2922.0037 | 2924.58549 | Non-sig. |
| PF3D7_1317000 | USP39 | 2546.6774 | 0.067168964 | 0.41108484 | 0.16339441 | 0.87020791 | 0.99402081 | 2682.7472 | 2498.00486 | 2292.17565 | 2713.78189 | Non-sig. |
| PF3D7_0514100 | UvrD | 1707.43938 | 0.067101762 | 0.4599953 | 0.14587489 | 0.88402016 | 0.99457954 | 1998.44897 | 1414.94017 | 1337.2575 | 2079.11089 | Non-sig. |
| PF3D7_0106200 | N/A | 544.802289 | 0.066632391 | 0.43713354 | 0.15243029 | 0.87884757 | 0.99421828 | 564.622563 | 621.778284 | 499.906394 | 492.901913 | Non-sig. |
| PF3D7_1354400 | N/A | 1654.73129 | 0.066087458 | 0.47063248 | 0.14042265 | 0.88832607 | 0.99457954 | 1136.023 | 1781.50781 | 2097.32538 | 1604.06898 | Non-sig. |
| PF3D7_1241800 | DBP9 | 2273.15882 | 0.065776919 | 0.4100737 | 0.16040268 | 0.87256388 | 0.99402081 | 2240.54322 | 2500.61474 | 2202.16827 | 2149.30903 | Non-sig. |
| PF3D7_0937400 | RIF | 2.66379875 | 0.065260555 | 2.61723434 | 0.02493493 | 0.98010687 | 0.99691622 | 5.24148055 | 0 | 0 | 5.41371444 | Non-sig. |
| PF3D7_0931100 | N/A | 1590.16196 | 0.065176301 | 0.72752135 | 0.08958679 | 0.92861558 | 0.99619209 | 2757.31374 | 1751.9036 | 351.548599 | 1499.88192 | Non-sig. |
| PF3D7_1129800 | N/A | 472.666898 | 0.065149329 | 0.44856498 | 0.14523945 | 0.88452182 | 0.99457954 | 384.198708 | 486.064118 | 539.588533 | 480.816232 | Non-sig. |
| PF3D7_1359000 | N/A | 986.448547 | 0.064356884 | 0.59121762 | 0.10885481 | 0.91331765 | 0.99619209 | 433.330501 | 879.443769 | 1495.17923 | 1137.84069 | Non-sig. |
| PF3D7_0523200 | N/A | 393.624596 | 0.06431357 | 0.51228402 | 0.1255428 | 0.90009384 | 0.99457954 | 248.604452 | 420.520784 | 520.768699 | 384.604449 | Non-sig. |
| PF3D7_0529900 | N/A | 2079.2999 | 0.064303269 | 0.41827903 | 0.15373295 | 0.8778203 | 0.99421828 | 2284.66046 | 2098.63245 | 1781.45218 | 2152.45451 | Non-sig. |
| PF3D7_1104100 | SYN13 | 1480.79087 | 0.06414896 | 0.41571728 | 0.1543091 | 0.87736602 | 0.99421828 | 1563.20843 | 1589.67032 | 1332.65678 | 1437.62793 | Non-sig. |
| PF3D7_1432500 | N/A | 493.316436 | 0.064080257 | 0.46623337 | 0.13744245 | 0.89068108 | 0.99457954 | 589.734268 | 549.573089 | 375.244051 | 458.714337 | Non-sig. |
| PF3D7_1323900 | N/A | 572.227947 | 0.06359042 | 0.52699979 | 0.12066498 | 0.9039564 | 0.99457954 | 782.190002 | 654.992453 | 337.399899 | 514.329433 | Non-sig. |
| PF3D7_0936500 | VAP1 | 14.8078818 | 0.063400149 | 1.07983187 | 0.05871298 | 0.95318072 | 0.99643037 | 24.1801889 | 17.1737328 | 4.92669406 | 12.9509113 | Non-sig. |

| | | | | | | | | | | | | |
|---|---|---|---|---|---|---|---|---|---|---|---|---|
| PF3D7_1248400 | N/A | 129.361959 | 0.063395551 | 0.63052147 | 0.10054464 | 0.91991195 | 0.99619209 | 191.805496 | 141.804378 | 61.5506513 | 122.287309 | Non-sig. |
| PF3D7_1355700 | NIF3 | 4919.08808 | 0.063239031 | 0.49743598 | 0.12712999 | 0.89883752 | 0.99457954 | 3137.54115 | 4196.04435 | 6484.70112 | 5858.0657 | Non-sig. |
| PF3D7_0514800 | N/A | 392.300264 | 0.062799827 | 0.57481717 | 0.10925183 | 0.91300275 | 0.99619209 | 247.374202 | 250.002681 | 519.882454 | 551.941718 | Non-sig. |
| PF3D7_0306400 | N/A | 3244.35245 | 0.062613979 | 0.49237887 | 0.12716626 | 0.89880881 | 0.99457954 | 2017.45251 | 3426.80611 | 4330.10105 | 3203.05015 | Non-sig. |
| PF3D7_1034500 | N/A | 2201.614 | 0.062453024 | 0.41702971 | 0.14975677 | 0.88095652 | 0.99421828 | 1907.46618 | 2328.68181 | 2400.31051 | 2169.99751 | Non-sig. |
| PF3D7_0827600 | N/A | 3021.34663 | 0.062233674 | 0.49120791 | 0.12669518 | 0.89918166 | 0.99457954 | 4026.68495 | 3048.43232 | 1886.06982 | 3124.19944 | Non-sig. |
| PF3D7_1363200 | PNKP | 274.822042 | 0.061903608 | 0.45733119 | 0.13535838 | 0.89232853 | 0.99457954 | 238.803266 | 289.753743 | 298.920183 | 271.810975 | Non-sig. |
| PF3D7_0417300 | N/A | 1622.63544 | 0.061306771 | 0.54747374 | 0.11198121 | 0.91083831 | 0.99619209 | 2342.27382 | 1814.37233 | 834.447168 | 1499.44844 | Non-sig. |
| PF3D7_1328000 | N/A | 2361.25834 | 0.061238528 | 0.44727204 | 0.13691562 | 0.89109749 | 0.99457954 | 1833.07109 | 2114.01651 | 2788.97519 | 2708.97057 | Non-sig. |
| PF3D7_1306100 | N/A | 221.905065 | 0.060345186 | 0.46260656 | 0.13044602 | 0.89621356 | 0.99457954 | 203.229369 | 234.56103 | 231.231056 | 218.598803 | Non-sig. |
| PF3D7_1403500 | N/A | 180.276677 | 0.060246602 | 0.4866446 | 0.1238 | 0.90147363 | 0.99457954 | 203.024624 | 191.302831 | 150.155408 | 176.623845 | Non-sig. |
| PF3D7_0918800 | N/A | 319.134373 | 0.060080951 | 0.48605828 | 0.12360853 | 0.90162524 | 0.99457954 | 230.49493 | 320.83329 | 394.212671 | 330.996602 | Non-sig. |
| PF3D7_1350100 | KRS1 | 2703.90326 | 0.059615446 | 0.54426851 | 0.10953315 | 0.91277963 | 0.99619209 | 1387.22426 | 2963.65723 | 3908.4764 | 2556.25516 | Non-sig. |
| PF3D7_0217200 | N/A | 348.874513 | 0.059338827 | 0.52012201 | 0.11408636 | 0.90916933 | 0.99597145 | 440.509283 | 271.375542 | 243.152654 | 440.460572 | Non-sig. |
| PF3D7_1011600 | N/A | 191.726877 | 0.05932848 | 0.55280878 | 0.10732188 | 0.91453362 | 0.99619209 | 115.748132 | 179.20565 | 259.513315 | 212.440409 | Non-sig. |
| PF3D7_0605800 | RAD50 | 4080.10572 | 0.059204196 | 0.48890247 | 0.12109613 | 0.90361489 | 0.99457954 | 5393.10161 | 3800.01158 | 2600.06088 | 4527.2488 | Non-sig. |
| PF3D7_1114500 | N/A | 143.337598 | 0.058538833 | 0.61043262 | 0.09589729 | 0.92360215 | 0.99619209 | 73.6139426 | 126.645609 | 206.932026 | 166.158814 | Non-sig. |
| PF3D7_1105800 | N/A | 4963.22054 | 0.058403216 | 0.43828574 | 0.13325374 | 0.89399271 | 0.99457954 | 5904.65582 | 5305.21045 | 3821.16918 | 4821.84672 | Non-sig. |
| PF3D7_0604100 | SIP2 | 19727.7718 | 0.057309619 | 0.45459201 | 0.12606825 | 0.8996779 | 0.99457954 | 15186.7386 | 16592.3882 | 23485.002 | 23646.9583 | Non-sig. |
| PF3D7_0509400 | RNAPI | 2809.48847 | 0.057121846 | 0.47915523 | 0.11921366 | 0.90510609 | 0.99486557 | 1882.88652 | 2565.45965 | 3624.52692 | 3165.08079 | Non-sig. |
| PF3D7_1405100 | N/A | 1655.46364 | 0.057117727 | 0.46223058 | 0.12356977 | 0.90165593 | 0.99457954 | 2068.66539 | 1615.62261 | 1177.04594 | 1760.52062 | Non-sig. |
| PF3D7_1025400 | N/A | 3975.83538 | 0.056896224 | 0.40566154 | 0.14025541 | 0.8884582 | 0.99457954 | 4013.09979 | 4266.15562 | 3781.84299 | 3842.24311 | Non-sig. |
| PF3D7_1426800 | N/A | 1146.97037 | 0.056512938 | 0.45401268 | 0.12447436 | 0.9009397 | 0.99457954 | 891.680544 | 1015.74156 | 1357.08739 | 1323.37199 | Non-sig. |
| PF3D7_1436100 | N/A | 1190.57141 | 0.056145835 | 0.43747807 | 0.12833977 | 0.89788009 | 0.99457954 | 1331.14937 | 1059.72834 | 1003.84413 | 1367.5638 | Non-sig. |
| PF3D7_1315300 | N/A | 1068.82781 | 0.055628921 | 0.48824826 | 0.11393573 | 0.90928874 | 0.99597145 | 719.108996 | 944.551171 | 1377.01475 | 1234.63634 | Non-sig. |
| PF3D7_0529500 | N/A | 243.694454 | 0.055133375 | 0.48011814 | 0.11483294 | 0.90857754 | 0.99597145 | 193.923105 | 266.412116 | 283.959212 | 230.483384 | Non-sig. |
| PF3D7_0313300 | N/A | 109.376323 | 0.054756862 | 0.62350072 | 0.08782165 | 0.93001843 | 0.99619209 | 56.128494 | 110.904404 | 158.187639 | 112.284754 | Non-sig. |
| PF3D7_0515900 | NIF2 | 804.227738 | 0.054632208 | 0.44836336 | 0.12184806 | 0.90301935 | 0.99457954 | 663.141744 | 706.340524 | 914.665627 | 932.763057 | Non-sig. |
| PF3D7_1213800 | PRS | 1519.24425 | 0.05442743 | 0.52465301 | 0.10373986 | 0.91737579 | 0.99619209 | 846.839025 | 1580.22456 | 2133.95708 | 1515.95632 | Non-sig. |
| PF3D7_0211300 | N/A | 239.247271 | 0.053378579 | 0.46331689 | 0.11520966 | 0.90827894 | 0.99597145 | 215.953323 | 224.511613 | 253.597232 | 262.926917 | Non-sig. |
| PF3D7_1411700 | N/A | 550.328057 | 0.053127135 | 0.42530301 | 0.12491596 | 0.90059008 | 0.99457954 | 526.035158 | 574.130389 | 554.323057 | 546.823624 | Non-sig. |
| PF3D7_1112100 | N/A | 11819.9801 | 0.052959882 | 0.55819983 | 0.09487621 | 0.92441316 | 0.99619209 | 17566.9449 | 11755.1764 | 5639.5878 | 12318.2112 | Non-sig. |
| PF3D7_1431600 | SCS-beta | 1677.70778 | 0.052911939 | 0.4884884 | 0.1083177 | 0.91374368 | 0.99619209 | 1084.24495 | 1834.7793 | 2209.29834 | 1582.50854 | Non-sig. |
| PF3D7_0103300 | N/A | 2697.45955 | 0.052628346 | 0.42332313 | 0.12432192 | 0.90106039 | 0.99457954 | 3052.80313 | 2823.76447 | 2243.93767 | 2669.33293 | Non-sig. |
| PF3D7_0913600 | N/A | 1783.19493 | 0.052061002 | 0.44117503 | 0.11800532 | 0.90606344 | 0.99520197 | 2125.66383 | 1826.93315 | 1376.65633 | 1803.52639 | Non-sig. |
| PF3D7_0405000 | DDX51 | 616.149008 | 0.05193777 | 0.42477489 | 0.12227128 | 0.90268418 | 0.99457954 | 575.288658 | 652.531472 | 634.765112 | 602.010789 | Non-sig. |
| PF3D7_0415300 | CRK3 | 1484.23845 | 0.051564648 | 0.42856821 | 0.12031841 | 0.90423092 | 0.99457954 | 1679.39569 | 1590.82884 | 1236.24326 | 1430.48602 | Non-sig. |
| PF3D7_1403100 | CAPD2 | 9523.3696 | 0.051422132 | 0.46541025 | 0.11048775 | 0.91202256 | 0.99619209 | 6626.34322 | 8864.99478 | 12080.6656 | 10521.4748 | Non-sig. |
| PF3D7_0914300 | N/A | 728.456689 | 0.051298564 | 0.47871804 | 0.1071582 | 0.91466347 | 0.99619209 | 927.080811 | 702.863185 | 504.248364 | 779.634395 | Non-sig. |
| PF3D7_1315600 | PIS | 719.613617 | 0.051295824 | 0.41944941 | 0.12229323 | 0.90266679 | 0.99457954 | 729.984143 | 724.998309 | 683.704486 | 739.767531 | Non-sig. |
| PF3D7_1247000 | FBXO9 | 611.156658 | 0.050899065 | 0.44730745 | 0.11378989 | 0.90940435 | 0.99597145 | 492.931555 | 657.917732 | 707.630755 | 586.146592 | Non-sig. |
| PF3D7_0815200 | N/A | 3047.12027 | 0.050456372 | 0.63794186 | 0.07909243 | 0.9369591 | 0.99639249 | 1050.48001 | 2812.59455 | 4936.81408 | 3388.59245 | Non-sig. |
| PF3D7_1323700 | GAPM1 | 10876.6133 | 0.049019976 | 0.49032412 | 0.09997464 | 0.92036446 | 0.99619209 | 6844.83886 | 10243.5435 | 14538.5019 | 11879.5688 | Non-sig. |
| PF3D7_0520500 | N/A | 749.057678 | 0.048750506 | 0.54185394 | 0.08996983 | 0.92831119 | 0.99619209 | 413.522353 | 865.53474 | 1058.92655 | 658.247071 | Non-sig. |
| PF3D7_1028600 | N/A | 1429.79004 | 0.048459838 | 0.48087611 | 0.10077406 | 0.91972982 | 0.99619209 | 1862.49287 | 1498.01621 | 949.404266 | 1409.24683 | Non-sig. |
| PF3D7_0511800 | INO1 | 6719.78406 | 0.048209166 | 0.50359033 | 0.09573092 | 0.92373428 | 0.99619209 | 3991.83416 | 6670.01808 | 9222.83142 | 6994.4526 | Non-sig. |
| PF3D7_0627300 | RNF5 | 6123.62285 | 0.047140834 | 0.40570328 | 0.11619535 | 0.90749772 | 0.99563532 | 5599.43065 | 6317.40918 | 6447.64084 | 6130.01074 | Non-sig. |
| PF3D7_1015500 | N/A | 379.416656 | 0.04690874 | 0.45785295 | 0.10245372 | 0.91839654 | 0.99619209 | 306.123144 | 393.853322 | 440.166189 | 377.523969 | Non-sig. |
| PF3D7_1350500 | N/A | 3451.68273 | 0.046680324 | 0.43109286 | 0.10828369 | 0.91377066 | 0.99619209 | 2912.81216 | 3059.46271 | 3878.69082 | 3955.76522 | Non-sig. |
| PF3D7_0619000 | N/A | 4247.04961 | 0.046426678 | 0.46582206 | 0.09966612 | 0.9206094 | 0.99619209 | 5427.07829 | 4434.73912 | 2930.71918 | 4195.66184 | Non-sig. |
| PF3D7_1463300 | N/A | 838.496303 | 0.046316595 | 0.52539742 | 0.08815535 | 0.9297532 | 0.99619209 | 479.552331 | 899.393165 | 1170.15564 | 804.88407 | Non-sig. |
| PF3D7_1210700 | N/A | 567.208247 | 0.046084487 | 0.49868567 | 0.09241189 | 0.92637079 | 0.99619209 | 369.427281 | 609.474139 | 746.541623 | 543.389947 | Non-sig. |
| PF3D7_1343700 | Kelch13 | 2785.48337 | 0.045635914 | 0.44623778 | 0.10226815 | 0.91854383 | 0.99619209 | 2111.02221 | 2643.6478 | 3371.55819 | 3015.70526 | Non-sig. |
| PF3D7_1242000 | QSOX | 490.581116 | 0.045348529 | 0.57411841 | 0.07898811 | 0.93704208 | 0.99639249 | 246.056088 | 420.732928 | 719.322427 | 576.213023 | Non-sig. |

| | | | | | | | | | | | | |
|---|---|---|---|---|---|---|---|---|---|---|---|---|
| PF3D7_0609700 | KIC6 | 9945.90666 | 0.045095803 | 0.49605439 | 0.09090899 | 0.92756491 | 0.99619209 | 13316.6611 | 8783.34082 | 6264.65675 | 11418.968 | Non-sig. |
| PF3D7_0606700 | N/A | 5987.70855 | 0.044900961 | 0.44840277 | 0.10013533 | 0.92023688 | 0.99619209 | 4618.43514 | 5271.09615 | 7170.37962 | 6890.9233 | Non-sig. |
| PF3D7_0621900 | SRP68 | 2400.7139 | 0.044782198 | 0.48044838 | 0.09320918 | 0.92573738 | 0.99619209 | 1589.8938 | 2278.83992 | 3136.67398 | 2597.4479 | Non-sig. |
| PF3D7_1108800 | N/A | 3.75691806 | 0.044706508 | 1.72293726 | 0.02594784 | 0.97929894 | 0.99691622 | 4.13126681 | 3.23642253 | 3.27421294 | 4.38576996 | Non-sig. |
| PF3D7_1475300 | COX11 | 724.607197 | 0.044471451 | 0.43121617 | 0.1031303 | 0.91785956 | 0.99619209 | 668.866344 | 819.965939 | 757.935777 | 651.66073 | Non-sig. |
| PF3D7_1020600 | N/A | 490.69833 | 0.044113963 | 0.43962441 | 0.10034466 | 0.9200707 | 0.99619209 | 545.961582 | 499.828505 | 420.597402 | 496.405831 | Non-sig. |
| PF3D7_1328200 | N/A | 4455.98053 | 0.044027274 | 0.41773606 | 0.10539496 | 0.91606242 | 0.99619209 | 4633.19694 | 3949.05934 | 4142.86875 | 5098.7971 | Non-sig. |
| PF3D7_1003900 | N/A | 14.9175249 | 0.043877134 | 0.96194738 | 0.04561282 | 0.96361885 | 0.99691622 | 14.6141757 | 11.8304259 | 14.7634011 | 18.4620968 | Non-sig. |
| PF3D7_1144200 | N/A | 97.665856 | 0.042493497 | 0.53038957 | 0.08011752 | 0.93614379 | 0.99639249 | 95.5726419 | 89.1140771 | 96.8744193 | 109.102286 | Non-sig. |
| PF3D7_1206300 | N/A | 11807.0575 | 0.04168805 | 0.75333256 | 0.05533818 | 0.95586905 | 0.99691622 | 21063.4936 | 12429.6325 | 2209.85091 | 11525.253 | Non-sig. |
| PF3D7_1216700 | PLP2 | 108.748936 | 0.041435148 | 0.53094493 | 0.07804039 | 0.93779593 | 0.99639249 | 120.023946 | 95.4898531 | 94.4523252 | 125.029619 | Non-sig. |
| PF3D7_1467900 | N/A | 12988.2436 | 0.041151221 | 0.4801099 | 0.08571209 | 0.93169529 | 0.99619209 | 16766.551 | 11163.7795 | 8839.82724 | 15182.8166 | Non-sig. |
| PF3D7_0107800 | MRE11 | 1756.52141 | 0.040930439 | 0.42402625 | 0.09652808 | 0.92310118 | 0.99619209 | 1985.70725 | 1772.18673 | 1477.70585 | 1790.48581 | Non-sig. |
| PF3D7_0201800 | KAHsp40 | 299.03749 | 0.040601578 | 0.46178823 | 0.0879225 | 0.92993827 | 0.99619209 | 257.748333 | 333.750063 | 331.770611 | 272.880954 | Non-sig. |
| PF3D7_1235100 | N/A | 202.360871 | 0.040368105 | 0.54416092 | 0.07418413 | 0.94086387 | 0.99639249 | 266.443531 | 241.920882 | 132.89116 | 168.187914 | Non-sig. |
| PF3D7_1110400 | N/A | 11316.4185 | 0.040340777 | 0.47217824 | 0.08543549 | 0.93191518 | 0.99619209 | 14681.7801 | 10759.8543 | 7635.00492 | 12189.0346 | Non-sig. |
| PF3D7_1350400 | N/A | 546.89827 | 0.039964809 | 0.43964792 | 0.09090185 | 0.92757058 | 0.99619209 | 611.510124 | 583.719566 | 467.308199 | 525.05519 | Non-sig. |
| PF3D7_0601600 | N/A | 429.572494 | 0.039889948 | 0.43969727 | 0.09072139 | 0.92771397 | 0.99619209 | 383.309993 | 458.464699 | 463.841292 | 412.673993 | Non-sig. |
| PF3D7_1313200 | MTFMT | 176.051133 | 0.039779432 | 0.50123194 | 0.07936332 | 0.93674364 | 0.99639249 | 136.801198 | 171.691229 | 210.240772 | 185.471332 | Non-sig. |
| PF3D7_1123200 | LRR11 | 1659.83871 | 0.039740445 | 0.49545628 | 0.08020979 | 0.9360704 | 0.99639249 | 1220.35648 | 1226.31525 | 2053.34462 | 2139.33847 | Non-sig. |
| PF3D7_1254700 | RIF | 11.6488279 | 0.039691386 | 1.17606367 | 0.03374935 | 0.97307702 | 0.99691622 | 7.32810427 | 19.3434047 | 15.5907691 | 4.33303343 | Non-sig. |
| PF3D7_0910300 | N/A | 1055.96012 | 0.039617524 | 0.43283147 | 0.09153106 | 0.92707063 | 0.99619209 | 905.064462 | 1177.13168 | 1177.69319 | 963.951147 | Non-sig. |
| PF3D7_0111400 | GEXP19 | 2000.81529 | 0.039541822 | 0.52715174 | 0.07501032 | 0.9402065 | 0.99639249 | 2804.71171 | 2321.81183 | 1142.45832 | 1734.27929 | Non-sig. |
| PF3D7_0616000 | PDXK | 490.718601 | 0.039351711 | 0.4554323 | 0.08640518 | 0.93114433 | 0.99619209 | 387.205221 | 503.287244 | 580.61345 | 491.768491 | Non-sig. |
| PF3D7_1011900 | HO | 2110.17464 | 0.039075561 | 0.40984851 | 0.09534148 | 0.9240436 | 0.99619209 | 2045.03833 | 2284.69213 | 2118.13887 | 1992.82922 | Non-sig. |
| PF3D7_1015800 | N/A | 3487.01458 | 0.039040589 | 0.50335256 | 0.07756112 | 0.93817717 | 0.99639249 | 2135.29103 | 3924.7019 | 4744.02579 | 3144.03958 | Non-sig. |
| PF3D7_1405400 | MutS | 549.341017 | 0.039023296 | 0.47136396 | 0.08278803 | 0.93402008 | 0.99639249 | 406.690754 | 508.460464 | 676.856785 | 605.356066 | Non-sig. |
| PF3D7_1037600 | XPB | 1765.00441 | 0.038951784 | 0.45392946 | 0.08581021 | 0.93161729 | 0.99619209 | 1327.72086 | 1959.11501 | 2154.35915 | 1618.82263 | Non-sig. |
| PF3D7_0312100 | N/A | 1318.6743 | 0.038889951 | 0.43740569 | 0.08891048 | 0.92915305 | 0.99619209 | 1057.38882 | 1279.87878 | 1544.17579 | 1393.25381 | Non-sig. |
| PF3D7_1038700 | N/A | 61.1285261 | 0.038767191 | 0.60304968 | 0.06428523 | 0.94874311 | 0.99639249 | 71.4737881 | 64.4054139 | 49.2650702 | 59.3698322 | Non-sig. |
| PF3D7_0817000 | UBA3 | 94.015246 | 0.038655252 | 0.60713053 | 0.06366877 | 0.94923398 | 0.99639249 | 55.5738692 | 93.4508284 | 129.676469 | 97.3598181 | Non-sig. |
| PF3D7_1313500 | N/A | 1312.44169 | 0.038262296 | 0.42999658 | 0.08898279 | 0.92909559 | 0.99619209 | 1147.47479 | 1175.65156 | 1442.44996 | 1484.19045 | Non-sig. |
| PF3D7_1144800 | N/A | 284.717485 | 0.038155406 | 0.52384186 | 0.07283764 | 0.94193532 | 0.99639249 | 380.741368 | 306.942798 | 181.480455 | 269.70532 | Non-sig. |
| PF3D7_1372400 | ACS4 | 1365.09637 | 0.038070366 | 0.50390269 | 0.07555103 | 0.9397763 | 0.99639249 | 840.107051 | 1488.16981 | 1853.70621 | 1278.4024 | Non-sig. |
| PF3D7_1459200 | N/A | 1316.15793 | 0.037832544 | 0.48256544 | 0.07839878 | 0.93751084 | 0.99639249 | 1724.19692 | 1290.57591 | 873.950842 | 1375.90804 | Non-sig. |
| PF3D7_1322200 | N/A | 7760.45604 | 0.037729868 | 0.44625894 | 0.08454703 | 0.93262151 | 0.99619209 | 6316.32827 | 6468.25229 | 9001.42841 | 9255.81519 | Non-sig. |
| PF3D7_1146300 | N/A | 10.1747224 | 0.037668651 | 1.28531333 | 0.02930698 | 0.97661976 | 0.99691622 | 17.7356208 | 13.9835876 | 2.4603485 | 6.51933277 | Non-sig. |
| PF3D7_0407100 | N/A | 259.027445 | 0.037595253 | 0.48221001 | 0.07796448 | 0.93785631 | 0.99639249 | 199.848285 | 271.521869 | 311.229842 | 253.509782 | Non-sig. |
| PF3D7_1312200 | N/A | 261.362328 | 0.037009555 | 0.45101499 | 0.08205837 | 0.9346003 | 0.99639249 | 264.104992 | 268.574771 | 251.943852 | 260.825699 | Non-sig. |
| PF3D7_1320200 | N/A | 515.761101 | 0.036585415 | 0.42891118 | 0.08529835 | 0.9320242 | 0.99619209 | 491.198282 | 549.485434 | 527.200542 | 495.160147 | Non-sig. |
| PF3D7_0824100 | N/A | 222.148212 | 0.03630857 | 0.47862625 | 0.07585996 | 0.93953051 | 0.99639249 | 249.173565 | 250.088962 | 189.679389 | 199.650933 | Non-sig. |
| PF3D7_1443400 | WLP1 | 3083.24774 | 0.035870365 | 0.4243653 | 0.08452709 | 0.93263736 | 0.99619209 | 2615.41374 | 3374.27613 | 3474.2389 | 2869.06218 | Non-sig. |
| PF3D7_0614100 | N/A | 2013.95472 | 0.035363871 | 0.49420118 | 0.07155764 | 0.94295395 | 0.99639249 | 2683.5853 | 1804.47209 | 1295.30845 | 2272.45304 | Non-sig. |
| PF3D7_1342800 | PEPCK | 1115.8869 | 0.03518512 | 0.54077948 | 0.06506371 | 0.94812327 | 0.99639249 | 592.741383 | 1131.93483 | 1611.43479 | 1127.43662 | Non-sig. |
| PF3D7_0935800 | CLAG9 | 27881.1442 | 0.035175039 | 0.64121878 | 0.05485653 | 0.95625276 | 0.99691622 | 9335.97307 | 26648.2764 | 45746.2298 | 29794.0975 | Non-sig. |
| PF3D7_0220800 | CLAG2 | 16930.3675 | 0.035160962 | 0.57559495 | 0.06108629 | 0.95129049 | 0.99639249 | 7554.56159 | 17270.3457 | 25893.194 | 17003.3687 | Non-sig. |
| PF3D7_1214900 | N/A | 1332.92175 | 0.034844021 | 0.42385107 | 0.08220817 | 0.93448118 | 0.99639249 | 1144.73192 | 1392.19719 | 1488.74829 | 1306.00961 | Non-sig. |
| PF3D7_0201900 | EMP3 | 2226.91185 | 0.03463669 | 0.57750562 | 0.05997637 | 0.95217446 | 0.99639249 | 3359.88961 | 1886.55635 | 1040.8604 | 2620.34103 | Non-sig. |
| PF3D7_0813700 | ABCF1 | 1160.85216 | 0.034137101 | 0.4833474 | 0.07062643 | 0.94369508 | 0.99639249 | 1522.43942 | 1146.72138 | 772.140955 | 1202.10688 | Non-sig. |
| PF3D7_0422300 | N/A | 1914.57297 | 0.033807047 | 0.49772061 | 0.06792374 | 0.94584634 | 0.99639249 | 1197.43881 | 2074.85214 | 2586.486 | 1799.51491 | Non-sig. |
| PF3D7_1227300 | N/A | 1261.3887 | 0.033478108 | 0.43002579 | 0.0778514 | 0.93794626 | 0.99639249 | 1441.56798 | 1228.43842 | 1052.1458 | 1323.40261 | Non-sig. |
| PF3D7_0905000 | ATPTG4 | 192.552445 | 0.033162742 | 0.48917357 | 0.06779341 | 0.94595009 | 0.99639249 | 159.670392 | 213.646812 | 220.848763 | 176.043814 | Non-sig. |
| PF3D7_1312800 | AAP2 | 13463.961 | 0.032947185 | 0.57242197 | 0.05755751 | 0.95410109 | 0.99673729 | 20110.5385 | 10749.2813 | 6510.31676 | 16485.7074 | Non-sig. |
| PF3D7_0827800 | SET3 | 7955.21938 | 0.032827352 | 0.46931467 | 0.06994742 | 0.94423551 | 0.99639249 | 10160.0608 | 7103.63836 | 5569.70108 | 8987.47723 | Non-sig. |

| | | | | | | | | | | | | |
|---|---|---|---|---|---|---|---|---|---|---|---|---|
| PF3D7_0113100 | SURF1.1 | 189.375855 | 0.032648906 | 0.64953684 | 0.0502649 | 0.95991129 | 0.99691622 | 299.240683 | 183.42893 | 75.5721557 | 199.261651 | Non-sig. |
| PF3D7_1349700 | N/A | 956.047424 | 0.03261561 | 0.423696 | 0.0769788 | 0.93864041 | 0.99639249 | 839.137458 | 958.383275 | 1051.19533 | 975.473628 | Non-sig. |
| PF3D7_0807300 | RAB18 | 683.175742 | 0.03246194 | 0.44875519 | 0.07233775 | 0.94233312 | 0.99639249 | 553.575682 | 628.063066 | 797.18549 | 753.878729 | Non-sig. |
| PF3D7_1460200 | N/A | 262.70721 | 0.032216737 | 0.47780413 | 0.06742666 | 0.94624205 | 0.99639249 | 306.330277 | 293.202056 | 213.410089 | 237.886416 | Non-sig. |
| PF3D7_0103700 | PSTK | 139.617655 | 0.032075801 | 0.54842718 | 0.05848689 | 0.9533608 | 0.99643037 | 182.79718 | 146.006158 | 93.5953938 | 136.071889 | Non-sig. |
| PF3D7_1433800 | N/A | 2216.85977 | 0.031663601 | 0.42991161 | 0.07365142 | 0.94128775 | 0.99639249 | 2109.43717 | 1851.19145 | 2275.57714 | 2631.2333 | Non-sig. |
| PF3D7_1464100 | N/A | 214.290777 | 0.031615653 | 0.46499491 | 0.06799139 | 0.94579248 | 0.99639249 | 216.932436 | 232.80936 | 206.98163 | 200.439683 | Non-sig. |
| PF3D7_0607500 | COQ2 | 306.775149 | 0.031316008 | 0.55907004 | 0.05601446 | 0.95533029 | 0.99691547 | 166.322328 | 307.941835 | 440.217648 | 312.618785 | Non-sig. |
| PF3D7_1018600 | N/A | 3204.07866 | 0.031052867 | 0.48869215 | 0.0635428 | 0.94933428 | 0.99639249 | 4299.78065 | 3212.29153 | 2039.78236 | 3264.46009 | Non-sig. |
| PF3D7_1244600 | ARFGAP1 | 1454.77337 | 0.030783178 | 0.45135364 | 0.0682019 | 0.94562491 | 0.99639249 | 1099.6858 | 1386.77437 | 1778.5402 | 1554.09309 | Non-sig. |
| PF3D7_0703200 | N/A | 5660.65956 | 0.030752278 | 0.41292504 | 0.07447424 | 0.94063304 | 0.99639249 | 6210.55795 | 5463.11417 | 4990.26299 | 5978.70311 | Non-sig. |
| PF3D7_1216000 | N/A | 1124.17276 | 0.03051939 | 0.44414605 | 0.06871476 | 0.94521667 | 0.99639249 | 1340.36349 | 1190.10681 | 884.456225 | 1081.76454 | Non-sig. |
| PF3D7_1356600 | N/A | 706.752885 | 0.030065456 | 0.52969205 | 0.05676026 | 0.95473617 | 0.99689429 | 406.296375 | 652.380572 | 992.117213 | 776.217383 | Non-sig. |
| PF3D7_0211200 | RAB5a | 659.406901 | 0.029953435 | 0.46513254 | 0.06439763 | 0.94865361 | 0.99639249 | 489.576965 | 701.037556 | 815.27003 | 631.743053 | Non-sig. |
| PF3D7_0515800 | N/A | 208.853714 | 0.02939994 | 0.46974817 | 0.0625866 | 0.9500957 | 0.99639249 | 185.926928 | 217.94288 | 227.415671 | 204.129377 | Non-sig. |
| PF3D7_0726400 | N/A | 5412.22658 | 0.029324926 | 0.51080123 | 0.05740966 | 0.95421887 | 0.99673729 | 3403.94789 | 4340.17416 | 7310.15816 | 6594.62611 | Non-sig. |
| PF3D7_1344000 | N/A | 431.786591 | 0.028885351 | 0.43961668 | 0.06570577 | 0.94761208 | 0.99639249 | 466.537126 | 409.823605 | 388.505169 | 462.280465 | Non-sig. |
| PF3D7_1246500 | N/A | 163.530589 | 0.028791158 | 0.47988202 | 0.05999633 | 0.95215856 | 0.99639249 | 154.591034 | 163.136196 | 169.159056 | 167.236071 | Non-sig. |
| PF3D7_1479500 | N/A | 6.3715405 | 0.028525373 | 1.52205258 | 0.01874138 | 0.98504741 | 0.99691622 | 1.0459648 | 6.44939147 | 11.4862906 | 6.50451511 | Non-sig. |
| PF3D7_1025500 | N/A | 677.863103 | 0.02846448 | 0.43424624 | 0.06554917 | 0.94773676 | 0.99639249 | 740.284837 | 619.866701 | 602.199645 | 749.101227 | Non-sig. |
| PF3D7_0303500 | N/A | 2299.3275 | 0.028135861 | 0.44713854 | 0.06292426 | 0.94982682 | 0.99639249 | 2586.52703 | 1862.00983 | 1967.45092 | 2781.32223 | Non-sig. |
| PF3D7_1433900 | N/A | 765.469794 | 0.028050226 | 0.42560003 | 0.06590748 | 0.94745148 | 0.99639249 | 795.694495 | 702.56365 | 720.42784 | 843.19319 | Non-sig. |
| PF3D7_1210600 | N/A | 2490.46622 | 0.027660393 | 0.43077787 | 0.06421034 | 0.94880275 | 0.99639249 | 2021.81884 | 2423.24775 | 2911.12995 | 2605.66835 | Non-sig. |
| PF3D7_1238600 | NSM | 3916.6643 | 0.027501451 | 0.40643974 | 0.06766428 | 0.94605289 | 0.99639249 | 3695.50258 | 3765.93998 | 4063.09809 | 4142.11656 | Non-sig. |
| PF3D7_1471900 | N/A | 522.869088 | 0.027368576 | 0.58626275 | 0.04668312 | 0.96276578 | 0.99691622 | 794.686187 | 549.170651 | 241.514139 | 506.105376 | Non-sig. |
| PF3D7_1463700 | CYC1 | 313.011342 | 0.027361033 | 0.44791381 | 0.06108549 | 0.95129113 | 0.99639249 | 305.638306 | 343.567326 | 314.434256 | 288.40548 | Non-sig. |
| PF3D7_1126600 | N/A | 597.670557 | 0.026735085 | 0.51212293 | 0.05220443 | 0.9583658 | 0.99691622 | 805.166243 | 526.850988 | 379.430744 | 679.234252 | Non-sig. |
| PF3D7_0317400 | N/A | 198.419792 | 0.026678494 | 0.59344728 | 0.04495512 | 0.96414308 | 0.99691622 | 121.940734 | 133.099964 | 270.963681 | 267.674788 | Non-sig. |
| PF3D7_0922000 | N/A | 154.166442 | 0.026566829 | 0.57111927 | 0.04651713 | 0.96289808 | 0.99691622 | 214.639819 | 162.040568 | 91.1566459 | 148.828736 | Non-sig. |
| PF3D7_0810900 | N/A | 2110.78712 | 0.026468999 | 0.4383245 | 0.06038677 | 0.9518476 | 0.99639249 | 2386.26294 | 2458.91352 | 1796.83248 | 1801.13954 | Non-sig. |
| PF3D7_0303800 | N/A | 773.252131 | 0.026029129 | 0.42938774 | 0.06061917 | 0.9516625 | 0.99639249 | 852.666386 | 825.112094 | 680.03789 | 735.192154 | Non-sig. |
| PF3D7_1368100 | RPN11 | 1482.65931 | 0.025910261 | 0.43912258 | 0.05900462 | 0.95294843 | 0.99639249 | 1178.30444 | 1436.58745 | 1760.1393 | 1555.60605 | Non-sig. |
| PF3D7_1005500 | UPF1 | 8151.09699 | 0.025904068 | 0.47636482 | 0.05437863 | 0.9566335 | 0.99691622 | 5427.84921 | 7838.57214 | 10727.6475 | 8610.31907 | Non-sig. |
| PF3D7_1406600 | ClpC | 3027.52046 | 0.025900372 | 0.41303072 | 0.0627081 | 0.94999895 | 0.99639249 | 3239.11251 | 2837.2121 | 2761.69144 | 3272.0658 | Non-sig. |
| PF3D7_0302800 | N/A | 686.603537 | 0.02584483 | 0.43769045 | 0.05904819 | 0.95291372 | 0.99639249 | 582.995293 | 648.130725 | 777.729342 | 737.558787 | Non-sig. |
| PF3D7_0522300 | N/A | 472.704878 | 0.024877353 | 0.51100232 | 0.04868344 | 0.96117157 | 0.99691622 | 611.588949 | 576.466209 | 325.957205 | 376.807149 | Non-sig. |
| PF3D7_1145700 | N/A | 422.344428 | 0.024129818 | 0.44914527 | 0.05372386 | 0.95715517 | 0.99691622 | 479.021166 | 453.057421 | 358.77793 | 398.521196 | Non-sig. |
| PF3D7_1444300 | LPAAT | 2724.87676 | 0.024119684 | 0.46721158 | 0.05162476 | 0.95882769 | 0.99691622 | 1910.48499 | 2927.67424 | 3493.39679 | 2567.951 | Non-sig. |
| PF3D7_1459900 | N/A | 469.568705 | 0.024061048 | 0.47242523 | 0.05093091 | 0.95938057 | 0.99691622 | 354.783955 | 428.459592 | 576.263462 | 518.767812 | Non-sig. |
| PF3D7_1347600 | N/A | 49.8089855 | 0.023952761 | 0.63754777 | 0.03757014 | 0.97003041 | 0.99691622 | 55.387705 | 58.0583478 | 43.4776755 | 42.3122135 | Non-sig. |
| PF3D7_1411900 | N/A | 307.628916 | 0.023820412 | 0.54059055 | 0.04406368 | 0.96485364 | 0.99691622 | 181.995907 | 332.679984 | 427.850234 | 287.989541 | Non-sig. |
| PF3D7_1017800 | N/A | 732.983806 | 0.023604485 | 0.46607895 | 0.05064482 | 0.95960855 | 0.99691622 | 536.177703 | 723.041067 | 917.503553 | 755.212901 | Non-sig. |
| PF3D7_1343800 | N/A | 13366.6655 | 0.023212426 | 0.45037052 | 0.05154073 | 0.95889465 | 0.99691622 | 13497.2612 | 9957.27716 | 13021.0393 | 16991.0845 | Non-sig. |
| PF3D7_1330000 | N/A | 173.037404 | 0.022754778 | 0.57139239 | 0.03982338 | 0.96823393 | 0.99691622 | 98.8210268 | 173.795064 | 244.1701 | 175.363426 | Non-sig. |
| PF3D7_1337300 | N/A | 993.306755 | 0.022674088 | 0.4299961 | 0.05273091 | 0.95794631 | 0.99691622 | 852.964734 | 938.754241 | 1117.86003 | 1063.64801 | Non-sig. |
| PF3D7_1111500 | N/A | 580.383336 | 0.022294439 | 0.49451041 | 0.04508386 | 0.96404046 | 0.99691622 | 395.747934 | 654.438348 | 755.751157 | 515.595905 | Non-sig. |
| PF3D7_0106800 | RAB5c | 713.189083 | 0.021794981 | 0.43729572 | 0.04984037 | 0.9602496 | 0.99691622 | 596.374652 | 707.949006 | 819.032982 | 729.399692 | Non-sig. |
| PF3D7_0615200 | N/A | 66.916778 | 0.021393622 | 0.73998369 | 0.02891094 | 0.97693562 | 0.99691622 | 106.016651 | 82.6848939 | 27.0893678 | 51.8761995 | Non-sig. |
| PF3D7_0409100 | PRPF31 | 2924.91001 | 0.020650594 | 0.47271271 | 0.0436853 | 0.96515526 | 0.99691622 | 3779.80199 | 3227.82402 | 2028.48365 | 2663.53038 | Non-sig. |
| PF3D7_1126700 | N/A | 16089.907 | 0.020339969 | 0.75374619 | 0.02698517 | 0.97847156 | 0.99691622 | 28923.9109 | 17114.7084 | 3029.45352 | 15291.5553 | Non-sig. |
| PF3D7_0811800 | N/A | 254.274693 | 0.019919985 | 0.45511935 | 0.04376871 | 0.96508877 | 0.99691622 | 269.414538 | 260.750226 | 235.702891 | 251.231115 | Non-sig. |
| PF3D7_1359500 | N/A | 43.4962937 | 0.01985811 | 0.68441308 | 0.0290148 | 0.97685279 | 0.99691622 | 55.3675214 | 49.4621592 | 31.1710434 | 37.9844506 | Non-sig. |
| PF3D7_0529400 | N/A | 17300.2066 | 0.01982254 | 0.43609703 | 0.04545443 | 0.9637451 | 0.99691622 | 20455.9566 | 15543.8026 | 13907.0136 | 19294.0538 | Non-sig. |
| PF3D7_0615100 | ENR | 53.5062342 | 0.019423484 | 0.65275543 | 0.02975614 | 0.97626154 | 0.99691622 | 66.2029274 | 65.4831261 | 40.2314939 | 42.1073892 | Non-sig. |

| | | | | | | | | | | | | |
|---|---|---|---|---|---|---|---|---|---|---|---|---|
| PF3D7_0522200 | TAF10 | 532.191243 | 0.018256307 | 0.44605191 | 0.04092866 | 0.96735277 | 0.99691622 | 569.264474 | 617.347539 | 488.481417 | 453.671542 | Non-sig. |
| PF3D7_1308500 | N/A | 694.241413 | 0.017479462 | 0.47522024 | 0.03678181 | 0.97065897 | 0.99691622 | 887.67405 | 717.993415 | 492.705047 | 678.593138 | Non-sig. |
| PF3D7_0311500 | N/A | 457.967715 | 0.017362168 | 0.51102248 | 0.03397535 | 0.97289681 | 0.99691622 | 291.923412 | 487.464715 | 618.173098 | 434.309636 | Non-sig. |
| PF3D7_1451900 | TSR1 | 2257.93651 | 0.017126254 | 0.4748004 | 0.03607043 | 0.9712262 | 0.99691622 | 1531.67625 | 2282.81872 | 2957.05298 | 2260.1981 | Non-sig. |
| PF3D7_1360900 | N/A | 3235.10472 | 0.017036399 | 0.46916581 | 0.03631211 | 0.9710335 | 0.99691622 | 2230.07249 | 3230.92754 | 4201.6006 | 3277.81825 | Non-sig. |
| PF3D7_0414400 | N/A | 13.9955987 | 0.016960289 | 1.10061696 | 0.0154098 | 0.98770524 | 0.99691622 | 23.0338031 | 17.1936183 | 4.92348962 | 10.8314838 | Non-sig. |
| PF3D7_1202200 | PIC | 2507.88548 | 0.016942551 | 0.50776229 | 0.03336709 | 0.97338185 | 0.99691622 | 1508.25631 | 2695.93727 | 3477.69739 | 2349.65095 | Non-sig. |
| PF3D7_1402800 | N/A | 3928.49745 | 0.016858984 | 0.48867683 | 0.03449925 | 0.97247904 | 0.99691622 | 5089.15968 | 3160.13335 | 2722.23682 | 4742.45996 | Non-sig. |
| PF3D7_0914100 | N/A | 10862.8503 | 0.016544846 | 0.66740165 | 0.02478994 | 0.98022252 | 0.99691622 | 18373.5199 | 10747.4067 | 3228.01094 | 11102.4636 | Non-sig. |
| PF3D7_0322500 | IscA2 | 664.624365 | 0.016500625 | 0.4911373 | 0.03359677 | 0.9731987 | 0.99691622 | 827.673375 | 814.889283 | 494.231854 | 521.702949 | Non-sig. |
| PF3D7_1446800 | HDP | 484.617255 | 0.01644327 | 0.4434913 | 0.03707687 | 0.97042371 | 0.99691622 | 456.212273 | 554.945855 | 507.438292 | 419.872601 | Non-sig. |
| PF3D7_0706000 | N/A | 4643.10992 | 0.016414477 | 0.42626855 | 0.03850736 | 0.96928316 | 0.99691622 | 3824.82291 | 4492.39274 | 5408.34298 | 4846.88103 | Non-sig. |
| PF3D7_1137300 | N/A | 1289.51419 | 0.015049577 | 0.45019939 | 0.03342869 | 0.97333273 | 0.99691622 | 986.670658 | 1235.94777 | 1578.62997 | 1356.80835 | Non-sig. |
| PF3D7_1435000 | COX15 | 1049.95006 | 0.014860222 | 0.54122924 | 0.02745643 | 0.97809569 | 0.99691622 | 568.806791 | 1147.39355 | 1519.907 | 963.692889 | Non-sig. |
| PF3D7_1236800 | ICMT | 310.367675 | 0.014847853 | 0.50876942 | 0.02918385 | 0.97671796 | 0.99691622 | 209.164548 | 330.615076 | 408.078522 | 293.612553 | Non-sig. |
| PF3D7_1459300 | N/A | 458.833439 | 0.014809146 | 0.47984897 | 0.0308621 | 0.97537952 | 0.99691622 | 570.821671 | 517.369157 | 342.400918 | 404.742009 | Non-sig. |
| PF3D7_0926000 | N/A | 625.297844 | 0.014316385 | 0.42551536 | 0.03364481 | 0.97316039 | 0.99691622 | 667.864936 | 630.78016 | 576.624593 | 625.921688 | Non-sig. |
| PF3D7_1365300 | N/A | 420.937527 | 0.013949013 | 0.56182871 | 0.02482788 | 0.98019226 | 0.99691622 | 222.23182 | 384.065244 | 615.211838 | 462.241207 | Non-sig. |
| PF3D7_0610900 | SPT5 | 3865.49171 | 0.01373505 | 0.44886411 | 0.03059957 | 0.97558888 | 0.99691622 | 4784.18228 | 3648.30661 | 2910.29563 | 4119.18232 | Non-sig. |
| PF3D7_1316700 | N/A | 93.6153404 | 0.013489211 | 0.64426837 | 0.02093725 | 0.98329571 | 0.99691622 | 47.3038095 | 89.0915207 | 138.765845 | 99.300187 | Non-sig. |
| PF3D7_0519000 | SLBP | 288.73748 | 0.013455355 | 0.55110495 | 0.02441523 | 0.9805214 | 0.99691622 | 402.668287 | 338.10419 | 172.435462 | 241.74198 | Non-sig. |
| PF3D7_1430300 | N/A | 1328.54925 | 0.013447527 | 0.45122311 | 0.02980239 | 0.97622465 | 0.99691622 | 1479.97123 | 1055.39501 | 1164.88278 | 1613.948 | Non-sig. |
| PF3D7_1440900 | N/A | 1119.33875 | 0.013339835 | 0.48810213 | 0.02733001 | 0.97819652 | 0.99691622 | 1452.13328 | 1303.78293 | 776.504385 | 944.934407 | Non-sig. |
| PF3D7_0525400 | N/A | 757.003779 | 0.013227446 | 0.44192335 | 0.02993154 | 0.97612165 | 0.99691622 | 644.865446 | 675.330664 | 862.021119 | 845.797886 | Non-sig. |
| PF3D7_1231800 | N/A | 1592.04785 | 0.013064558 | 0.5821432 | 0.02244217 | 0.98209524 | 0.99691622 | 819.032814 | 1117.60751 | 2350.29744 | 2081.25362 | Non-sig. |
| PF3D7_0907600 | N/A | 1034.08237 | 0.012976301 | 0.44722425 | 0.0290152 | 0.97685247 | 0.99691622 | 1253.35536 | 1029.89331 | 805.772571 | 1047.30826 | Non-sig. |
| PF3D7_0522600 | N/A | 1478.31618 | 0.012308347 | 0.56742179 | 0.02169171 | 0.98269388 | 0.99691622 | 736.828868 | 1228.56824 | 2206.81527 | 1741.05233 | Non-sig. |
| PF3D7_1019500 | N/A | 2686.77747 | 0.012001315 | 0.54605162 | 0.02197835 | 0.98246522 | 0.99691622 | 1387.15684 | 2544.60677 | 3963.74914 | 2851.59712 | Non-sig. |
| PF3D7_0811100 | N/A | 439.105581 | 0.011778053 | 0.53802983 | 0.02189108 | 0.98253484 | 0.99691622 | 252.576129 | 415.268879 | 621.697173 | 466.880141 | Non-sig. |
| PF3D7_1419600 | N/A | 852.545901 | 0.011770388 | 0.52301857 | 0.02250472 | 0.98204534 | 0.99691622 | 1195.23379 | 775.175882 | 503.25815 | 936.515784 | Non-sig. |
| PF3D7_0304300 | N/A | 454.945469 | 0.011766794 | 0.43311859 | 0.0271676 | 0.97832605 | 0.99691622 | 421.446426 | 452.678409 | 484.643649 | 461.013391 | Non-sig. |
| PF3D7_0920200 | N/A | 1933.31741 | 0.011506874 | 0.4089748 | 0.0281359 | 0.97755376 | 0.99691622 | 1873.36034 | 1846.3963 | 1977.81501 | 2035.69799 | Non-sig. |
| PF3D7_0726300 | PMS1 | 595.995013 | 0.011260439 | 0.4593361 | 0.0245146 | 0.98044214 | 0.99691622 | 493.043968 | 504.181817 | 694.096416 | 692.657853 | Non-sig. |
| PF3D7_0910100 | N/A | 4334.0046 | 0.011120712 | 0.58620222 | 0.01897078 | 0.98486442 | 0.99691622 | 1880.11899 | 4200.38956 | 6754.08954 | 4501.42033 | Non-sig. |
| PF3D7_0908000 | PMP1 | 2109.31522 | 0.010696984 | 0.51968431 | 0.02058362 | 0.98357781 | 0.99691622 | 1232.49565 | 1885.73684 | 2970.1264 | 2348.902 | Non-sig. |
| PF3D7_1227700 | KIC2 | 6283.76505 | 0.01059481 | 0.52541215 | 0.02016476 | 0.98391194 | 0.99691622 | 8991.42362 | 6924.74965 | 3530.35227 | 5688.53464 | Non-sig. |
| PF3D7_0711500 | N/A | 2250.47118 | 0.009534386 | 0.40831939 | 0.02335031 | 0.98137084 | 0.99691622 | 2187.94525 | 2380.76584 | 2298.09188 | 2135.08176 | Non-sig. |
| PF3D7_0106900 | IspD | 1142.83221 | 0.00913536 | 0.52664598 | 0.0173463 | 0.98616035 | 0.99691622 | 1624.22723 | 1237.14229 | 654.574041 | 1055.38527 | Non-sig. |
| PF3D7_0619100 | N/A | 348.181208 | 0.009096222 | 0.44054835 | 0.0206475 | 0.98352685 | 0.99691622 | 327.902681 | 353.167209 | 366.196655 | 345.458287 | Non-sig. |
| PF3D7_1017000 | N/A | 2508.55777 | 0.009004186 | 0.49086636 | 0.01834346 | 0.98536486 | 0.99691622 | 1601.28111 | 2516.64396 | 3399.81875 | 2516.48726 | Non-sig. |
| PF3D7_0902100 | FIKK9.2 | 1937.02211 | 0.00885296 | 0.47754577 | 0.01853845 | 0.9852093 | 0.99691622 | 1318.09548 | 2062.0314 | 2543.73156 | 1824.23001 | Non-sig. |
| PF3D7_0704900 | RF2 | 522.192445 | 0.008311392 | 0.4735821 | 0.01755006 | 0.9859978 | 0.99691622 | 387.840946 | 566.127899 | 653.268756 | 481.532179 | Non-sig. |
| PF3D7_0933300 | N/A | 1749.88552 | 0.008010699 | 0.42559838 | 0.0188222 | 0.98498294 | 0.99691622 | 2009.34349 | 1801.30054 | 1480.91531 | 1707.98272 | Non-sig. |
| PF3D7_1011300 | ARV1 | 432.169528 | 0.007884223 | 0.45460482 | 0.01734303 | 0.98616296 | 0.99691622 | 356.71318 | 461.428275 | 505.065094 | 405.471562 | Non-sig. |
| PF3D7_0212700 | N/A | 226.09175 | 0.007833948 | 0.46150291 | 0.01697486 | 0.98645667 | 0.99691622 | 222.7015 | 210.425124 | 228.238353 | 243.002021 | Non-sig. |
| PF3D7_0106100 | N/A | 1197.20983 | 0.007359257 | 0.51967109 | 0.01416137 | 0.98870124 | 0.99698118 | 709.603624 | 1074.44543 | 1678.34702 | 1326.44326 | Non-sig. |
| PF3D7_1227000 | N/A | 849.816662 | 0.006871405 | 0.42903967 | 0.01601578 | 0.9872218 | 0.99691622 | 766.039227 | 929.653608 | 929.424665 | 774.149149 | Non-sig. |
| PF3D7_1137600 | N/A | 25.3227697 | 0.006262372 | 0.97109068 | 0.0064488 | 0.99485464 | 0.99863448 | 44.1588349 | 28.9799904 | 6.56902214 | 21.5832315 | Non-sig. |
| PF3D7_1412400 | N/A | 3137.06334 | 0.005827555 | 0.42662073 | 0.0136598 | 0.98910139 | 0.9971949 | 3202.95634 | 2608.32553 | 3058.52783 | 3678.44365 | Non-sig. |
| PF3D7_0912800 | GCD10 | 648.414942 | 0.005235416 | 0.42140049 | 0.01242385 | 0.99008746 | 0.99779905 | 628.37545 | 657.75013 | 666.062561 | 641.471625 | Non-sig. |
| PF3D7_0405400 | PRPF8 | 9250.5364 | 0.004928987 | 0.61491337 | 0.00801574 | 0.99360443 | 0.99863448 | 3566.67991 | 8406.27887 | 14902.4094 | 10126.7774 | Non-sig. |
| PF3D7_1219600 | ATP2 | 4920.18201 | 0.004815108 | 0.58967269 | 0.00816573 | 0.99348476 | 0.99863448 | 2105.29035 | 4758.38148 | 7718.26114 | 5098.79509 | Non-sig. |
| PF3D7_1333900 | N/A | 759.207722 | 0.004456926 | 0.45366169 | 0.00982434 | 0.99216144 | 0.99863448 | 711.024215 | 595.386475 | 804.968439 | 925.45176 | Non-sig. |
| PF3D7_0407000 | N/A | 120.959929 | 0.004356761 | 0.5065225 | 0.00860132 | 0.99313723 | 0.99863448 | 113.425749 | 115.955989 | 128.071043 | 126.386935 | Non-sig. |

| | | | | | | | | | | | | |
|---|---|---|---|---|---|---|---|---|---|---|---|---|
| PF3D7_1337200 | DXS | 998.117176 | 0.003981606 | 0.60873621 | 0.00654077 | 0.99478125 | 0.99863448 | 412.807024 | 884.029712 | 1580.29069 | 1115.34128 | Non-sig. |
| PF3D7_1409200 | N/A | 1407.0263 | 0.003869239 | 0.52593838 | 0.00735683 | 0.99413015 | 0.99863448 | 1923.36294 | 1755.0654 | 887.295374 | 1062.38149 | Non-sig. |
| PF3D7_0309500 | AS | 6327.73817 | 0.00334612 | 0.48721959 | 0.00686779 | 0.99452034 | 0.99863448 | 4119.96347 | 6873.03244 | 8520.49117 | 5797.4656 | Non-sig. |
| PF3D7_1123300 | N/A | 2974.42567 | 0.003009314 | 0.40781439 | 0.00737913 | 0.99411236 | 0.99863448 | 2791.99021 | 3079.32478 | 3150.57231 | 2875.81537 | Non-sig. |
| PF3D7_1458600 | N/A | 1300.06948 | 0.002827033 | 0.44654818 | 0.00633086 | 0.99494874 | 0.99863448 | 1021.86734 | 1212.19519 | 1575.46489 | 1390.75049 | Non-sig. |
| PF3D7_0409300 | N/A | 556.884242 | 0.002611294 | 0.44001769 | 0.00593452 | 0.99526497 | 0.99863448 | 479.288747 | 528.873045 | 633.297925 | 586.07725 | Non-sig. |
| PF3D7_0907500 | N/A | 1729.64483 | 0.002028231 | 0.42263038 | 0.00479907 | 0.99617091 | 0.99863448 | 1942.03347 | 1843.9604 | 1514.99388 | 1617.59159 | Non-sig. |
| PF3D7_0208900 | P230p | 85.3057906 | 0.001865209 | 0.61188274 | 0.00304831 | 0.9975678 | 0.99888759 | 56.9186796 | 67.5721223 | 113.365449 | 103.366912 | Non-sig. |
| PF3D7_0707200 | N/A | 3713.49269 | 0.001735628 | 0.42914121 | 0.00404442 | 0.99677303 | 0.99863448 | 4207.43097 | 3241.38944 | 3215.26243 | 4189.88792 | Non-sig. |
| PF3D7_1012200 | RA | 6348.24192 | 0.00168919 | 0.48080613 | 0.00351325 | 0.99719684 | 0.99870489 | 8361.79785 | 7184.26465 | 4327.59208 | 5519.31308 | Non-sig. |
| PF3D7_0610000 | N/A | 685.848701 | 0.000803714 | 0.4461479 | 0.00180145 | 0.99856265 | 0.99920548 | 718.662186 | 813.744118 | 652.717518 | 558.270981 | Non-sig. |
| PF3D7_1237400 | N/A | 919.656064 | 0.000716375 | 0.42848892 | 0.00167186 | 0.99866605 | 0.99920548 | 798.456198 | 950.102996 | 1040.22976 | 889.835304 | Non-sig. |
| PF3D7_1428500 | N/A | 419.941356 | 0.000417915 | 0.51370052 | 0.00081354 | 0.99935089 | 0.99935089 | 500.659528 | 287.451934 | 339.273201 | 552.38076 | Non-sig. |
| PF3D7_0935100 | KSH1 | 230.421655 | -0.000638214 | 0.49243211 | -0.00129604 | 0.99896591 | 0.99920548 | 177.282662 | 244.600809 | 283.451211 | 216.351937 | Non-sig. |
| PF3D7_0529000 | TIP | 2116.70406 | -0.000707176 | 0.57393468 | -0.00123215 | 0.99901688 | 0.99920548 | 992.103468 | 1910.69289 | 3241.95439 | 2322.0655 | Non-sig. |
| PF3D7_0711400 | SAP18 | 1349.40456 | -0.000932483 | 0.51402109 | -0.00181409 | 0.99855256 | 0.99920548 | 1891.03509 | 1445.19455 | 809.014523 | 1252.37407 | Non-sig. |
| PF3D7_1123000 | N/A | 7671.80324 | -0.001281651 | 0.58021984 | -0.00220891 | 0.99823755 | 0.99920548 | 11815.7941 | 6454.27558 | 3535.02538 | 8882.11791 | Non-sig. |
| PF3D7_1422500 | HRD1 | 466.236222 | -0.001833162 | 0.47218414 | -0.0038823 | 0.99690238 | 0.99863448 | 352.253022 | 433.614847 | 580.547083 | 498.529936 | Non-sig. |
| PF3D7_0620600 | N/A | 105.15998 | -0.002074016 | 0.54044665 | -0.0038376 | 0.99693805 | 0.99863448 | 126.067382 | 97.6952182 | 84.5643019 | 112.313017 | Non-sig. |
| PF3D7_0607200 | N/A | 457.09756 | -0.002280935 | 0.44276729 | -0.00515154 | 0.99588968 | 0.99863448 | 492.123195 | 508.740186 | 422.890986 | 404.635874 | Non-sig. |
| PF3D7_0205600 | N/A | 527.965874 | -0.002299497 | 0.52836058 | -0.00435213 | 0.99652751 | 0.99863448 | 318.918042 | 462.466316 | 737.511904 | 592.967233 | Non-sig. |
| PF3D7_1248600 | N/A | 1720.78821 | -0.00247415 | 0.47511278 | -0.0052075 | 0.99584503 | 0.99863448 | 1181.4053 | 1695.16588 | 2262.7883 | 1743.79337 | Non-sig. |
| PF3D7_1354900 | N/A | 1176.06034 | -0.002672456 | 0.52050268 | -0.00513438 | 0.99590338 | 0.99863448 | 1659.86002 | 1282.24144 | 694.806127 | 1067.33375 | Non-sig. |
| PF3D7_0316700 | YOP1 | 1251.62035 | -0.002749101 | 0.47661852 | -0.00576793 | 0.99539788 | 0.99863448 | 867.83342 | 1320.01661 | 1637.47022 | 1181.16116 | Non-sig. |
| PF3D7_1401800 | CK | 4514.61278 | -0.002862322 | 0.44541472 | -0.00642619 | 0.99487267 | 0.99863448 | 3847.46546 | 3642.97198 | 5190.5356 | 5377.47808 | Non-sig. |
| PF3D7_1002300 | N/A | 26.4216822 | -0.003031555 | 0.77327254 | -0.00392042 | 0.99687196 | 0.99863448 | 24.1330945 | 22.5528665 | 28.7291932 | 30.2715747 | Non-sig. |
| PF3D7_1118400 | HAD4 | 1754.82814 | -0.003187033 | 0.43418848 | -0.00734021 | 0.99414342 | 0.99863448 | 1428.9739 | 1704.05976 | 2084.32167 | 1801.95721 | Non-sig. |
| PF3D7_1358300 | ROM7 | 203.66048 | -0.003397497 | 0.46704195 | -0.0072745 | 0.99419584 | 0.99863448 | 193.433468 | 194.280123 | 214.309628 | 212.618701 | Non-sig. |
| PF3D7_0628800 | GATB | 785.261911 | -0.004069493 | 0.57999745 | -0.0070164 | 0.99440177 | 0.99863448 | 366.5562 | 752.02007 | 1205.80053 | 816.670848 | Non-sig. |
| PF3D7_0314500 | N/A | 720.466016 | -0.004144974 | 0.48129124 | -0.0086122 | 0.99312855 | 0.99863448 | 941.469655 | 742.409551 | 501.850643 | 696.134215 | Non-sig. |
| PF3D7_0911600 | N/A | 760.026692 | -0.004311555 | 0.43091852 | -0.0100055 | 0.9920169 | 0.99863448 | 839.113702 | 694.179296 | 683.341886 | 823.471884 | Non-sig. |
| PF3D7_1409300 | DDI1 | 5216.68776 | -0.004374645 | 0.44009228 | -0.00994029 | 0.99206893 | 0.99863448 | 6381.54053 | 5438.43713 | 4067.93278 | 4978.84059 | Non-sig. |
| PF3D7_0812500 | N/A | 2987.68698 | -0.004933101 | 0.54440843 | -0.0090614 | 0.99277015 | 0.99863448 | 4424.42759 | 3168.68963 | 1561.55633 | 2796.0744 | Non-sig. |
| PF3D7_1021100 | N/A | 872.205205 | -0.006072166 | 0.66648418 | -0.00911074 | 0.99273078 | 0.99863448 | 274.004395 | 872.045626 | 1473.70543 | 869.065374 | Non-sig. |
| PF3D7_1251100 | N/A | 582.752688 | -0.006647499 | 0.43501272 | -0.01528116 | 0.98780787 | 0.99691622 | 510.103679 | 586.997464 | 657.931321 | 575.978286 | Non-sig. |
| PF3D7_1145800 | N/A | 319.643489 | -0.006677588 | 0.44794249 | -0.01490724 | 0.98810618 | 0.99691622 | 317.972182 | 288.611821 | 322.783038 | 349.206917 | Non-sig. |
| PF3D7_1325600 | FIS1 | 406.910087 | -0.006860293 | 0.47384297 | -0.01447799 | 0.98844864 | 0.99691622 | 314.348386 | 445.196063 | 501.173905 | 366.921996 | Non-sig. |
| PF3D7_1364100 | P92 | 5728.41685 | -0.007022023 | 0.57247393 | -0.0122661 | 0.99021331 | 0.99779905 | 2656.06092 | 5336.84873 | 8828.2677 | 6092.49004 | Non-sig. |
| PF3D7_0608900 | N/A | 1044.38278 | -0.007066866 | 0.41839453 | -0.01689044 | 0.98652402 | 0.99691622 | 992.308974 | 970.845734 | 1101.49934 | 1112.87708 | Non-sig. |
| PF3D7_0603900 | N/A | 126.910594 | -0.00743397 | 0.60980172 | -0.0121908 | 0.99027339 | 0.99779905 | 184.978049 | 143.841583 | 69.7913263 | 109.031419 | Non-sig. |
| PF3D7_1347500 | ALBA4 | 13383.7181 | -0.007438439 | 0.40955464 | -0.01816226 | 0.98550941 | 0.99691622 | 12203.6193 | 14107.2257 | 14632.7044 | 12591.3231 | Non-sig. |
| PF3D7_1209100 | N/A | 134.25491 | -0.00762943 | 0.51469825 | -0.01482311 | 0.9881733 | 0.99691622 | 122.11783 | 111.586682 | 147.014617 | 156.300512 | Non-sig. |
| PF3D7_1231600 | PRP2 | 2651.43148 | -0.007853999 | 0.50233566 | -0.01563496 | 0.98752561 | 0.99691622 | 3681.37479 | 2497.23244 | 1636.2915 | 2790.82719 | Non-sig. |
| PF3D7_0513900 | N/A | 182.054288 | -0.007954512 | 0.50285763 | -0.01581862 | 0.9873791 | 0.99691622 | 142.515126 | 194.454911 | 222.395048 | 168.852067 | Non-sig. |
| PF3D7_0617100 | N/A | 2347.24615 | -0.008249668 | 0.4700931 | -0.01754901 | 0.98599864 | 0.99691622 | 1787.34823 | 1886.75167 | 2920.30156 | 2794.58315 | Non-sig. |
| PF3D7_0628000 | PTPS | 114.637599 | -0.008452884 | 0.51284472 | -0.01648235 | 0.98684959 | 0.99691622 | 106.797142 | 120.354042 | 123.087966 | 108.311245 | Non-sig. |
| PF3D7_0204000 | CAPH2 | 129.489234 | -0.008545677 | 0.49995229 | -0.01709298 | 0.98636244 | 0.99691622 | 135.774779 | 122.323437 | 124.013342 | 135.845377 | Non-sig. |
| PF3D7_1219000 | FRM2 | 9690.31117 | -0.008745074 | 0.44347781 | -0.01971931 | 0.98426729 | 0.99691622 | 11469.1205 | 8104.80601 | 7970.467 | 11216.8512 | Non-sig. |
| PF3D7_1414000 | RPN13 | 3612.05027 | -0.00901159 | 0.43087689 | -0.02091453 | 0.98331383 | 0.99691622 | 4055.24613 | 4138.00204 | 3191.57753 | 3063.37539 | Non-sig. |
| PF3D7_1032800 | LRR1 | 379.234097 | -0.00906439 | 0.45604133 | -0.01987625 | 0.98414209 | 0.99691622 | 314.908769 | 371.542516 | 445.745186 | 384.739917 | Non-sig. |
| PF3D7_1349200 | N/A | 1902.01577 | -0.009159817 | 0.50804329 | -0.0180296 | 0.98561524 | 0.99691622 | 1150.60322 | 1841.19895 | 2665.13645 | 1951.12445 | Non-sig. |
| PF3D7_1465400 | N/A | 316.941319 | -0.009242173 | 0.5356585 | -0.01725385 | 0.9862341 | 0.99691622 | 192.241597 | 297.384799 | 443.342301 | 334.796579 | Non-sig. |
| PF3D7_0727500 | N/A | 582.113252 | -0.009783996 | 0.43803727 | -0.02233599 | 0.98217994 | 0.99691622 | 509.362344 | 617.001399 | 658.654182 | 543.435085 | Non-sig. |
| PF3D7_0929000 | TAF7 | 739.511285 | -0.009791879 | 0.43830426 | -0.02234037 | 0.98217645 | 0.99691622 | 649.651114 | 817.920272 | 834.24 | 656.233753 | Non-sig. |

| | | | | | | | | | | | | |
|---|---|---|---|---|---|---|---|---|---|---|---|---|
| PF3D7_0519600 | N/A | 1554.89706 | -0.009918506 | 0.41879992 | -0.02368316 | 0.98110534 | 0.99691622 | 1538.53773 | 1727.29451 | 1581.93145 | 1371.82457 | Non-sig. |
| PF3D7_0831200 | N/A | 1521.26092 | -0.010152377 | 0.41443106 | -0.02449714 | 0.98045606 | 0.99691622 | 1395.56129 | 1498.70371 | 1657.5462 | 1533.23247 | Non-sig. |
| PF3D7_1218800 | PSOP17 | 61.1058109 | -0.010564875 | 0.6119146 | -0.01726528 | 0.98622499 | 0.99691622 | 73.5286292 | 68.7115266 | 49.2595008 | 52.9235869 | Non-sig. |
| PF3D7_1106700 | DNA2 | 1628.73211 | -0.010632025 | 0.43793003 | -0.02427791 | 0.98063094 | 0.99691622 | 1957.23889 | 1680.19073 | 1312.4774 | 1565.02142 | Non-sig. |
| PF3D7_1358200 | N/A | 1452.62693 | -0.011045122 | 0.43063472 | -0.02564847 | 0.97953773 | 0.99691622 | 1671.24822 | 1323.66136 | 1245.31542 | 1570.28273 | Non-sig. |
| PF3D7_1457600 | N/A | 889.046832 | -0.011685764 | 0.43612498 | -0.02679453 | 0.97862362 | 0.99691622 | 1040.75616 | 859.770048 | 744.746486 | 910.914638 | Non-sig. |
| PF3D7_0622500 | N/A | 304.043346 | -0.011688789 | 0.46301412 | -0.02524499 | 0.97985955 | 0.99691622 | 263.889156 | 333.810983 | 346.51305 | 271.960193 | Non-sig. |
| PF3D7_1356100 | N/A | 5811.6086 | -0.011887682 | 0.47898668 | -0.0248184 | 0.98019982 | 0.99691622 | 7665.4759 | 5046.28262 | 4005.95993 | 6528.71595 | Non-sig. |
| PF3D7_0314900 | N/A | 2115.19867 | -0.011934855 | 0.40615 | -0.02938534 | 0.97655727 | 0.99691622 | 2162.02286 | 2123.7408 | 2085.90023 | 2089.13081 | Non-sig. |
| PF3D7_0925500 | TLP2 | 305.911992 | -0.012138017 | 0.45835318 | -0.0264818 | 0.97887305 | 0.99691622 | 279.347521 | 340.357739 | 334.951682 | 268.991028 | Non-sig. |
| PF3D7_0603600 | N/A | 504.612944 | -0.012257364 | 0.43235305 | -0.02835036 | 0.97738272 | 0.99691622 | 516.528268 | 459.050398 | 497.008057 | 545.865053 | Non-sig. |
| PF3D7_1326000 | C3AP2 | 1840.51049 | -0.01293868 | 0.44523576 | -0.02906029 | 0.97681651 | 0.99691622 | 1546.72722 | 2128.51831 | 2150.61246 | 1536.18397 | Non-sig. |
| PF3D7_1320100 | ClpS | 399.197536 | -0.013170531 | 0.54383456 | -0.0242179 | 0.9806788 | 0.99691622 | 574.948636 | 397.165734 | 227.44076 | 397.235013 | Non-sig. |
| PF3D7_0504700 | CEP120 | 812.527836 | -0.013709954 | 0.74868614 | -0.01831202 | 0.98538994 | 0.99691622 | 1463.07076 | 893.445037 | 170.049182 | 723.546371 | Non-sig. |
| PF3D7_0808400 | SEC28 | 908.140662 | -0.013722452 | 0.42257633 | -0.03247331 | 0.9740946 | 0.99691622 | 977.634348 | 838.313638 | 847.378251 | 969.23641 | Non-sig. |
| PF3D7_1033600 | CEF1 | 2339.40658 | -0.01400194 | 0.46678003 | -0.02999687 | 0.97606955 | 0.99691622 | 3045.19795 | 2383.94386 | 1656.64774 | 2271.83676 | Non-sig. |
| PF3D7_1010500 | N/A | 28.3914314 | -0.014111213 | 0.90923507 | -0.01551987 | 0.98761743 | 0.99691622 | 46.5927558 | 37.7284821 | 10.6472042 | 18.5972837 | Non-sig. |
| PF3D7_1312300 | N/A | 193.875087 | -0.01427462 | 0.48419965 | -0.02948086 | 0.97648109 | 0.99691622 | 214.959463 | 218.050815 | 174.817577 | 167.672493 | Non-sig. |
| PF3D7_1008700 | N/A | 6456.28894 | -0.014572769 | 0.58262446 | -0.02501229 | 0.98004516 | 0.99691622 | 2900.29497 | 5749.83248 | 10077.1145 | 7097.91384 | Non-sig. |
| PF3D7_1208500 | N/A | 135.551117 | -0.014802888 | 0.51969559 | -0.02848377 | 0.97727631 | 0.99691622 | 113.804659 | 115.847298 | 158.539401 | 154.013112 | Non-sig. |
| PF3D7_1301400 | HYP12 | 233.178921 | -0.015242983 | 0.55465661 | -0.02748184 | 0.97807543 | 0.99691622 | 330.355274 | 253.260136 | 138.786451 | 210.313822 | Non-sig. |
| PF3D7_0808100 | N/A | 772.708852 | -0.015748001 | 0.54535111 | -0.02887681 | 0.97696284 | 0.99691622 | 428.910651 | 666.106351 | 1124.57816 | 871.240245 | Non-sig. |
| PF3D7_0829600 | ETRAMP8 | 71.5419734 | -0.015992116 | 0.59019934 | -0.02709613 | 0.97838306 | 0.99691622 | 88.2083151 | 76.2331675 | 55.8245349 | 65.9018759 | Non-sig. |
| PF3D7_1116500 | FT2 | 1080.94718 | -0.016067874 | 0.48857759 | -0.03288705 | 0.97376466 | 0.99691622 | 751.86389 | 914.081401 | 1421.75738 | 1236.08604 | Non-sig. |
| PF3D7_1243800 | WDR82 | 234.608735 | -0.016386165 | 0.51838034 | -0.03161031 | 0.97478282 | 0.99691622 | 162.009156 | 253.265508 | 309.596457 | 213.563817 | Non-sig. |
| PF3D7_0313900 | N/A | 32.5316045 | -0.016428198 | 0.76668056 | -0.0214277 | 0.98290448 | 0.99691622 | 44.2212696 | 25.749758 | 21.3546007 | 38.8007899 | Non-sig. |
| PF3D7_0722700 | C3AP1 | 907.743399 | -0.016472282 | 0.47186507 | -0.03490888 | 0.9721524 | 0.99691622 | 683.881263 | 1036.45971 | 1141.70964 | 768.922986 | Non-sig. |
| PF3D7_1413000 | N/A | 32.0469838 | -0.016559626 | 0.91286533 | -0.01814027 | 0.98552695 | 0.99691622 | 55.6408115 | 37.5824669 | 9.03002698 | 25.9346298 | Non-sig. |
| PF3D7_0416600 | PHBL | 636.257292 | -0.016804168 | 0.45736777 | -0.03674104 | 0.97069149 | 0.99691622 | 777.576519 | 680.34758 | 502.597159 | 584.507911 | Non-sig. |
| PF3D7_1245200 | N/A | 265.270955 | -0.016825286 | 0.54046437 | -0.03113117 | 0.97516493 | 0.99691622 | 161.202853 | 266.029206 | 372.107588 | 261.744171 | Non-sig. |
| PF3D7_1252700 | N/A | 3332.38595 | -0.017431577 | 0.56586383 | -0.03080525 | 0.97542485 | 0.99691622 | 1591.521 | 3291.32331 | 5113.12523 | 3333.57428 | Non-sig. |
| PF3D7_0309200 | ARK2 | 4839.80054 | -0.017486446 | 0.60030704 | -0.02912917 | 0.97676157 | 0.99691622 | 7716.1768 | 4292.52018 | 2022.48158 | 5328.0236 | Non-sig. |
| PF3D7_1140700 | N/A | 2326.0122 | -0.017789432 | 0.45822907 | -0.03882214 | 0.96903219 | 0.99691622 | 2965.67982 | 2186.8445 | 1715.33275 | 2436.19171 | Non-sig. |
| PF3D7_0729800 | N/A | 282.825317 | -0.018163836 | 0.52116111 | -0.03485263 | 0.97219725 | 0.99691622 | 381.488594 | 308.085111 | 188.023566 | 253.704 | Non-sig. |
| PF3D7_0732200 | RIF | 8.13939473 | -0.018212365 | 1.24767578 | -0.01459703 | 0.98835367 | 0.99691622 | 7.33661566 | 5.37142034 | 9.02738385 | 10.8221591 | Non-sig. |
| PF3D7_1038600 | N/A | 92.5865875 | -0.018279411 | 0.53666731 | -0.03406097 | 0.97282853 | 0.99691622 | 101.009038 | 97.6530262 | 85.4083339 | 86.2759524 | Non-sig. |
| PF3D7_0607100 | N/A | 23.9235503 | -0.018319589 | 0.96444614 | -0.01899493 | 0.98484515 | 0.99691622 | 40.9437185 | 28.9920934 | 7.38821914 | 18.37017 | Non-sig. |
| PF3D7_1412100 | MCMBP | 2923.349 | -0.01843703 | 0.4377294 | -0.0421197 | 0.96640328 | 0.99691622 | 2320.79915 | 2939.68012 | 3562.99609 | 2869.92065 | Non-sig. |
| PF3D7_1435600 | N/A | 16596.4336 | -0.018502423 | 0.54987864 | -0.03364819 | 0.97315769 | 0.99691622 | 24765.955 | 14170.4454 | 8640.16306 | 18809.1712 | Non-sig. |
| PF3D7_0615500 | CRK5 | 3753.91055 | -0.01870201 | 0.40694229 | -0.0459574 | 0.9633442 | 0.99691622 | 4043.66767 | 3730.34555 | 3512.91981 | 3728.70916 | Non-sig. |
| PF3D7_1309300 | PRPF3 | 2079.72439 | -0.019041399 | 0.512062 | -0.03718573 | 0.97033692 | 0.99691622 | 2945.45786 | 2127.67964 | 1241.8138 | 2003.94626 | Non-sig. |
| PF3D7_1450000 | N/A | 887.53786 | -0.019085156 | 0.51198197 | -0.03727701 | 0.97026414 | 0.99691622 | 1237.80771 | 812.456059 | 549.358513 | 950.529163 | Non-sig. |
| PF3D7_0507700 | NPL4 | 2580.98505 | -0.019201045 | 0.40853729 | -0.04699949 | 0.96251363 | 0.99691622 | 2662.88272 | 2730.04375 | 2533.47712 | 2397.53664 | Non-sig. |
| PF3D7_1002200 | PArt | 4609.90189 | -0.019217693 | 0.57907206 | -0.03318705 | 0.97352543 | 0.99691622 | 7189.3487 | 4883.69014 | 2092.26208 | 4274.30665 | Non-sig. |
| PF3D7_0315100 | eIF4E | 799.213812 | -0.019416999 | 0.49083206 | -0.03955935 | 0.96844443 | 0.99691622 | 538.160332 | 744.954106 | 1070.69585 | 843.044959 | Non-sig. |
| PF3D7_1138350 | N/A | 66.9739693 | -0.019704961 | 0.60142275 | -0.03276391 | 0.97386286 | 0.99691622 | 56.6655218 | 79.3413644 | 78.1040279 | 53.784963 | Non-sig. |
| PF3D7_0934400 | ApiAP2 | 1024.16184 | -0.01975584 | 0.41769974 | -0.04729675 | 0.96227672 | 0.99691622 | 1041.58841 | 1096.14073 | 1020.77782 | 938.140414 | Non-sig. |
| PF3D7_1360500 | GCbeta | 401.028464 | -0.020105889 | 0.46897597 | -0.0428719 | 0.96580365 | 0.99691622 | 496.543811 | 403.267809 | 311.353669 | 392.948568 | Non-sig. |
| PF3D7_1117300 | N/A | 564.000672 | -0.020112683 | 0.50315973 | -0.03997276 | 0.96811484 | 0.99691622 | 380.376857 | 485.270401 | 755.181606 | 635.173824 | Non-sig. |
| PF3D7_0528400 | DHHC7 | 6190.20389 | -0.020235172 | 0.43044453 | -0.04700994 | 0.96250531 | 0.99691622 | 7439.34057 | 6232.59958 | 5028.14447 | 6060.73093 | Non-sig. |
| PF3D7_1415300 | N/A | 1826.76941 | -0.020258189 | 0.43464103 | -0.04660901 | 0.96282485 | 0.99691622 | 1496.78519 | 1733.95413 | 2182.16882 | 1894.16951 | Non-sig. |
| PF3D7_1355800 | SF3B5 | 405.162161 | -0.020810377 | 0.46757139 | -0.04450738 | 0.96449997 | 0.99691622 | 316.634497 | 414.246618 | 499.290687 | 390.476844 | Non-sig. |
| PF3D7_1449900 | N/A | 432.124553 | -0.020813579 | 0.46415987 | -0.0448414 | 0.96423372 | 0.99691622 | 479.827591 | 344.673454 | 390.762045 | 513.235121 | Non-sig. |
| PF3D7_1113600 | N/A | 918.573949 | -0.020898176 | 0.41512792 | -0.05034153 | 0.95985023 | 0.99691622 | 952.941567 | 913.306006 | 897.554134 | 910.494088 | Non-sig. |

| | | | | | | | | | | | | |
|---|---|---|---|---|---|---|---|---|---|---|---|---|
| PF3D7_0410500 | N/A | 119.899097 | -0.020912236 | 0.71280764 | -0.02933784 | 0.97659515 | 0.99691622 | 196.736819 | 151.759561 | 45.0899486 | 86.0100602 | Non-sig. |
| PF3D7_0404700 | DPAP3 | 22666.5851 | -0.021996945 | 0.42633016 | -0.05159603 | 0.95885058 | 0.99691622 | 27047.245 | 22609.036 | 18631.7819 | 22378.2776 | Non-sig. |
| PF3D7_1468100 | MORC | 23844.3039 | -0.022075957 | 0.41604141 | -0.05306192 | 0.95768257 | 0.99691622 | 23705.6659 | 20310.5915 | 24347.7966 | 27013.1617 | Non-sig. |
| PF3D7_0803700 | g-tub | 445.698919 | -0.022159709 | 0.50966038 | -0.04347936 | 0.96531941 | 0.99691622 | 319.83479 | 344.476594 | 578.14013 | 540.344163 | Non-sig. |
| PF3D7_0416400 | HAT1 | 13415.8512 | -0.022918525 | 0.41306943 | -0.05548347 | 0.9557533 | 0.99691622 | 14978.1797 | 12439.0081 | 12066.8017 | 14179.4154 | Non-sig. |
| PF3D7_1205200 | HADO | 153.022563 | -0.023821549 | 0.57002287 | -0.04179051 | 0.9666657 | 0.99691622 | 92.3810961 | 146.036347 | 215.898234 | 157.774577 | Non-sig. |
| PF3D7_1471300 | N/A | 829.412086 | -0.02399083 | 0.43780826 | -0.05479757 | 0.95629973 | 0.99691622 | 969.8709 | 873.571636 | 702.946258 | 771.259551 | Non-sig. |
| PF3D7_0216600 | N/A | 2481.51704 | -0.024012005 | 0.42854352 | -0.05603166 | 0.95531659 | 0.99691547 | 2476.12271 | 2030.83104 | 2528.196 | 2890.91842 | Non-sig. |
| PF3D7_0310800 | N/A | 572.970386 | -0.024039631 | 0.49474744 | -0.0485897 | 0.96124628 | 0.99691622 | 390.851249 | 522.605252 | 764.335202 | 614.089841 | Non-sig. |
| PF3D7_0924700 | SF3A3 | 1182.34864 | -0.024129627 | 0.45589123 | -0.05292847 | 0.9577889 | 0.99691622 | 1490.38419 | 1191.03968 | 894.376375 | 1153.59431 | Non-sig. |
| PF3D7_1317700 | N/A | 149.743934 | -0.024134847 | 0.49052219 | -0.04920236 | 0.96075803 | 0.99691622 | 137.690316 | 153.500144 | 164.216517 | 143.56876 | Non-sig. |
| PF3D7_1340700 | RAB11b | 1274.87519 | -0.024277963 | 0.42261656 | -0.05744679 | 0.95418929 | 0.99673729 | 1296.5288 | 1423.09014 | 1274.6924 | 1105.18944 | Non-sig. |
| PF3D7_1437900 | HSP40 | 4135.14348 | -0.024497814 | 0.43169057 | -0.05674855 | 0.9547455 | 0.99689429 | 4927.08663 | 4394.2037 | 3413.65389 | 3805.62973 | Non-sig. |
| PF3D7_1328900 | N/A | 740.395054 | -0.024540442 | 0.4371466 | -0.05613778 | 0.95523204 | 0.99691547 | 851.339802 | 791.921761 | 642.221894 | 676.096761 | Non-sig. |
| PF3D7_1452300 | DER1-1 | 288.734352 | -0.024775598 | 0.48876111 | -0.05069061 | 0.95957206 | 0.99691622 | 218.483055 | 307.05543 | 363.698724 | 265.700201 | Non-sig. |
| PF3D7_0906300 | N/A | 187.674735 | -0.024881653 | 0.48024488 | -0.05181034 | 0.95867981 | 0.99691622 | 164.112962 | 186.729721 | 214.335961 | 185.520295 | Non-sig. |
| PF3D7_0305600 | APE1 | 5330.21911 | -0.025471986 | 0.41304113 | -0.06166937 | 0.95082613 | 0.99639249 | 5829.10171 | 5719.57739 | 4925.56759 | 4846.62974 | Non-sig. |
| PF3D7_0823400 | N/A | 15.2319835 | -0.02559299 | 0.94965957 | -0.02694965 | 0.9784999 | 0.99691622 | 16.8015249 | 12.8844917 | 13.9561251 | 17.2857923 | Non-sig. |
| PF3D7_1306400 | RPT4 | 11064.5076 | -0.025716255 | 0.42697378 | -0.06022912 | 0.95197315 | 0.99639249 | 12676.8765 | 12350.774 | 9649.55058 | 9580.8293 | Non-sig. |
| PF3D7_1147600 | N/A | 308.903868 | -0.026091716 | 0.48206132 | -0.0541253 | 0.95683533 | 0.99691622 | 265.172926 | 245.735922 | 358.063733 | 366.642889 | Non-sig. |
| PF3D7_1006900 | N/A | 668.2878 | -0.026144846 | 0.42923329 | -0.06091057 | 0.95143043 | 0.99639249 | 733.073671 | 707.544299 | 615.727242 | 616.805991 | Non-sig. |
| PF3D7_0513700 | PSOP12 | 72.5737145 | -0.02628214 | 0.6911672 | -0.03802573 | 0.96966716 | 0.99691622 | 33.6836589 | 77.2549113 | 112.517222 | 66.8390658 | Non-sig. |
| PF3D7_0211500 | N/A | 1481.39019 | -0.026806798 | 0.4296209 | -0.0623964 | 0.95024716 | 0.99639249 | 1741.18025 | 1446.10097 | 1249.33899 | 1488.94056 | Non-sig. |
| PF3D7_1314800 | N/A | 2.76231914 | -0.026826562 | 2.24219078 | -0.01196444 | 0.99045398 | 0.99779905 | 3.11512068 | 0 | 2.46162018 | 5.4725357 | Non-sig. |
| PF3D7_0829100 | CCDC124 | 762.617642 | -0.027563563 | 0.52837981 | -0.05216619 | 0.95839627 | 0.99691622 | 1076.31343 | 879.374416 | 463.833324 | 630.949395 | Non-sig. |
| PF3D7_0302000 | PRP46 | 1261.43101 | -0.028365448 | 0.43283963 | -0.06553339 | 0.94774932 | 0.99639249 | 1099.63272 | 1381.05491 | 1447.86021 | 1117.17619 | Non-sig. |
| PF3D7_1433300 | RBAP46 | 994.400155 | -0.028584658 | 0.43041597 | -0.0664117 | 0.94705005 | 0.99639249 | 867.042195 | 923.175085 | 1141.28365 | 1046.09969 | Non-sig. |
| PF3D7_0524100 | N/A | 2052.01138 | -0.028612534 | 0.47896762 | -0.05973793 | 0.95236436 | 0.99639249 | 2750.24606 | 2010.17198 | 1394.81522 | 2052.81227 | Non-sig. |
| PF3D7_0408300 | ZRANB2 | 503.363971 | -0.028950362 | 0.47819962 | -0.06054033 | 0.9517253 | 0.99639249 | 622.135705 | 582.897076 | 394.929757 | 413.493346 | Non-sig. |
| PF3D7_1449600 | N/A | 1368.12842 | -0.029090745 | 0.42915335 | -0.06778636 | 0.9459557 | 0.99639249 | 1600.47916 | 1388.8326 | 1163.56954 | 1319.63239 | Non-sig. |
| PF3D7_0616200 | NDC80 | 3194.58459 | -0.029098065 | 0.53749159 | -0.05413678 | 0.95682619 | 0.99691622 | 4693.41465 | 3550.37012 | 1760.56454 | 2773.98904 | Non-sig. |
| PF3D7_1121500 | N/A | 221.848919 | -0.029169469 | 0.48967898 | -0.05956855 | 0.95249926 | 0.99639249 | 175.402926 | 210.43423 | 272.565035 | 228.993485 | Non-sig. |
| PF3D7_1224700 | N/A | 2594.36104 | -0.029170251 | 0.44110858 | -0.06612941 | 0.94727479 | 0.99639249 | 3012.75868 | 2145.30098 | 2228.59996 | 2990.78455 | Non-sig. |
| PF3D7_1221700 | N/A | 624.287719 | -0.029421255 | 0.42415225 | -0.06936484 | 0.94469921 | 0.99639249 | 668.411697 | 616.965893 | 592.974219 | 618.799068 | Non-sig. |
| PF3D7_0709000 | CRT | 479.902923 | -0.029678659 | 0.49885276 | -0.05949383 | 0.95255878 | 0.99639249 | 323.922506 | 458.276966 | 645.43823 | 491.973989 | Non-sig. |
| PF3D7_1212700 | EIF3A | 13315.0109 | -0.029788161 | 0.4398816 | -0.06771859 | 0.94600965 | 0.99639249 | 16511.0066 | 13603.606 | 10394.2227 | 12751.2084 | Non-sig. |
| PF3D7_1363500 | CCR4-2 | 1624.00839 | -0.029908006 | 0.44071402 | -0.06786262 | 0.945895 | 0.99639249 | 1304.49574 | 1532.12108 | 1976.93484 | 1682.4819 | Non-sig. |
| PF3D7_1117800 | MLH | 2001.01615 | -0.030510563 | 0.41546456 | -0.07343722 | 0.9414582 | 0.99639249 | 1805.1463 | 1980.43196 | 2239.05543 | 1979.4309 | Non-sig. |
| PF3D7_0724400 | PAM18 | 341.903749 | -0.030595337 | 0.44609803 | -0.06858433 | 0.94532049 | 0.99639249 | 358.818953 | 369.750328 | 332.288652 | 306.757061 | Non-sig. |
| PF3D7_0909000 | N/A | 142.598124 | -0.031189987 | 0.68922221 | -0.04525389 | 0.96390494 | 0.99691622 | 236.859669 | 136.265609 | 51.7423146 | 145.524905 | Non-sig. |
| PF3D7_1210500 | N/A | 11.387099 | -0.031995399 | 1.10303465 | -0.0290067 | 0.97685924 | 0.99691622 | 7.35723284 | 13.9546998 | 15.6004705 | 8.6359931 | Non-sig. |
| PF3D7_1333600 | UTP4 | 838.055369 | -0.032033285 | 0.41638715 | -0.07693149 | 0.93867804 | 0.99639249 | 828.088659 | 838.864843 | 866.59879 | 818.669183 | Non-sig. |
| PF3D7_0306600 | ATPTG17 | 743.430417 | -0.032192018 | 0.45424646 | -0.07086906 | 0.94350197 | 0.99639249 | 597.407921 | 806.111643 | 905.814727 | 664.387376 | Non-sig. |
| PF3D7_0727300 | DNMT | 514.699657 | -0.03231838 | 0.44169454 | -0.07316907 | 0.94167158 | 0.99639249 | 593.515113 | 487.17193 | 447.583999 | 530.527586 | Non-sig. |
| PF3D7_0418900 | N/A | 310.624204 | -0.03231858 | 0.4535826 | -0.07125181 | 0.94319735 | 0.99639249 | 311.240071 | 345.605283 | 316.959858 | 268.691601 | Non-sig. |
| PF3D7_0818700 | N/A | 3273.22978 | -0.032622017 | 0.42996153 | -0.07587194 | 0.93952098 | 0.99639249 | 3480.90283 | 2659.04343 | 3139.63222 | 3813.34064 | Non-sig. |
| PF3D7_1406100 | N/A | 1304.7734 | -0.033174358 | 0.42670768 | -0.07774493 | 0.93803095 | 0.99639249 | 1448.61622 | 1146.73739 | 1191.05821 | 1432.68181 | Non-sig. |
| PF3D7_1361700 | COX2A | 1812.85683 | -0.033336331 | 0.43788432 | -0.07613045 | 0.93931531 | 0.99639249 | 1504.53773 | 1964.20553 | 2162.84629 | 1619.83775 | Non-sig. |
| PF3D7_0708000 | N/A | 596.747839 | -0.033374182 | 0.44190455 | -0.07552351 | 0.93979819 | 0.99639249 | 518.026923 | 541.815959 | 689.098011 | 638.050462 | Non-sig. |
| PF3D7_1343200 | N/A | 3479.85476 | -0.033448159 | 0.43448993 | -0.07698259 | 0.9386374 | 0.99639249 | 2807.64379 | 3407.33061 | 4232.4876 | 3471.95701 | Non-sig. |
| PF3D7_0824800 | N/A | 2137.73398 | -0.033472353 | 0.44984402 | -0.0744088 | 0.94068511 | 0.99639249 | 2694.00534 | 2107.87092 | 1631.35021 | 2117.70944 | Non-sig. |
| PF3D7_1111700 | N/A | 297.565782 | -0.033713086 | 0.45097605 | -0.07475583 | 0.94040899 | 0.99639249 | 310.558935 | 320.81152 | 291.565409 | 267.327266 | Non-sig. |
| PF3D7_0415200 | N/A | 1809.2693 | -0.034699904 | 0.49433415 | -0.07019524 | 0.94403826 | 0.99639249 | 2368.48962 | 1384.68561 | 1293.84861 | 2190.05334 | Non-sig. |
| PF3D7_0209100 | PATPL1 | 1469.8681 | -0.034892144 | 0.43401107 | -0.0803946 | 0.93592343 | 0.99639249 | 1230.61397 | 1538.62239 | 1744.45429 | 1365.78175 | Non-sig. |

| | | | | | | | | | | | | |
|---|---|---|---|---|---|---|---|---|---|---|---|---|
| PF3D7_0620500 | N/A | 1818.54135 | -0.034906071 | 0.41143332 | -0.08484016 | 0.93238846 | 0.99619209 | 1873.31648 | 1666.16639 | 1807.78781 | 1926.89471 | Non-sig. |
| PF3D7_0315300 | N/A | 433.224134 | -0.035005297 | 0.48949702 | -0.07151279 | 0.94298965 | 0.99639249 | 570.040537 | 418.345007 | 307.218003 | 437.29299 | Non-sig. |
| PF3D7_1461400 | N/A | 2811.63061 | -0.035378697 | 0.41442965 | -0.08536719 | 0.93196947 | 0.99619209 | 3118.8118 | 2611.49699 | 2573.52525 | 2942.68839 | Non-sig. |
| PF3D7_1003100 | C4AP3 | 277.897341 | -0.035649853 | 0.51145806 | -0.0697024 | 0.94443053 | 0.99639249 | 211.9505 | 333.997667 | 350.495858 | 215.145338 | Non-sig. |
| PF3D7_1460600 | ISP3 | 1782.76211 | -0.035763814 | 0.42638059 | -0.08387768 | 0.93315369 | 0.99639249 | 2076.21886 | 1675.77862 | 1533.69177 | 1845.3592 | Non-sig. |
| PF3D7_1031900 | N/A | 103.028371 | -0.035847605 | 0.52054191 | -0.06886593 | 0.94509633 | 0.99639249 | 109.307956 | 100.903085 | 99.3504437 | 102.552001 | Non-sig. |
| PF3D7_0618500 | MDH | 1034.11076 | -0.035912294 | 0.41507368 | -0.08652029 | 0.93105283 | 0.99619209 | 1047.20905 | 1076.85252 | 1046.75583 | 965.625644 | Non-sig. |
| PF3D7_1131300 | N/A | 156.370303 | -0.035935793 | 0.5056904 | -0.07106283 | 0.94334775 | 0.99639249 | 186.226439 | 169.553706 | 130.575102 | 139.125965 | Non-sig. |
| PF3D7_1454900 | N/A | 30.7202946 | -0.036530143 | 0.80404927 | -0.04543272 | 0.9637624 | 0.99691622 | 45.9828909 | 29.0290458 | 16.4067285 | 31.4625131 | Non-sig. |
| PF3D7_0920100 | N/A | 300.532201 | -0.03664595 | 0.45392289 | -0.08073166 | 0.93565536 | 0.99639249 | 329.680909 | 322.125736 | 279.115196 | 271.206964 | Non-sig. |
| PF3D7_0927600 | N/A | 6144.92276 | -0.036885089 | 0.57610423 | -0.06402503 | 0.9489503 | 0.99639249 | 9629.27278 | 5968.28787 | 2818.07132 | 6164.05908 | Non-sig. |
| PF3D7_1425300 | N/A | 2231.23382 | -0.037190775 | 0.58989759 | -0.06304616 | 0.94972975 | 0.99639249 | 3504.08474 | 2573.91086 | 1016.2764 | 1830.66329 | Non-sig. |
| PF3D7_1217300 | EngA | 2168.31268 | -0.037908766 | 0.46022627 | -0.08236984 | 0.93435262 | 0.99639249 | 2804.56322 | 2115.61802 | 1589.34672 | 2163.72276 | Non-sig. |
| PF3D7_1018700 | N/A | 1640.47686 | -0.0379318 | 0.4250793 | -0.08923464 | 0.92889543 | 0.99619209 | 1669.76415 | 1377.39425 | 1654.32233 | 1860.4267 | Non-sig. |
| PF3D7_1431300 | LSG1 | 803.647988 | -0.038331322 | 0.41916786 | -0.09144623 | 0.92713803 | 0.99619209 | 850.02861 | 762.74854 | 778.677311 | 823.137491 | Non-sig. |
| PF3D7_1465500 | K2 | 22.3925901 | -0.03932749 | 0.86354781 | -0.04554176 | 0.96367549 | 0.99691622 | 27.3887184 | 13.9458291 | 18.0708488 | 30.1649639 | Non-sig. |
| PF3D7_0317900 | N/A | 319.889587 | -0.03945897 | 0.49853045 | -0.07915057 | 0.93691286 | 0.99639249 | 229.409929 | 339.088165 | 418.840585 | 292.219669 | Non-sig. |
| PF3D7_1226600 | PCNA2 | 963.252355 | -0.039602251 | 0.53641073 | -0.07382822 | 0.94114707 | 0.99639249 | 537.385441 | 948.214235 | 1415.19312 | 952.216619 | Non-sig. |
| PF3D7_0319000 | ATPase7 | 6329.96577 | -0.039989924 | 0.54305136 | -0.0736393 | 0.9412974 | 0.99639249 | 3534.15815 | 5017.52631 | 9300.86356 | 7467.31505 | Non-sig. |
| PF3D7_1443100 | N/A | 3522.595 | -0.040465112 | 0.49604558 | -0.08157539 | 0.93498437 | 0.99639249 | 4900.52511 | 3282.14667 | 2243.82252 | 3663.8857 | Non-sig. |
| PF3D7_1233800 | N/A | 1735.38919 | -0.040680084 | 0.45086652 | -0.09022645 | 0.92810727 | 0.99619209 | 2038.21204 | 2034.88757 | 1481.68678 | 1386.77038 | Non-sig. |
| PF3D7_1006500 | N/A | 185.565468 | -0.041007062 | 0.48454844 | -0.08462944 | 0.932556 | 0.99619209 | 181.787509 | 210.407345 | 194.58759 | 155.47943 | Non-sig. |
| PF3D7_1220700 | N/A | 45.3085485 | -0.041053254 | 0.64325223 | -0.06382139 | 0.94911244 | 0.99639249 | 48.4091234 | 40.7760639 | 43.5268102 | 48.5221964 | Non-sig. |
| PF3D7_0416800 | SAR1 | 5753.90447 | -0.041074521 | 0.40925163 | -0.10036495 | 0.92005459 | 0.99619209 | 6313.92157 | 5964.28709 | 5357.81517 | 5379.59407 | Non-sig. |
| PF3D7_0306500 | COX6C | 2071.35962 | -0.041312264 | 0.41267827 | -0.10010768 | 0.92025884 | 0.99619209 | 2049.19432 | 2221.04174 | 2152.8058 | 1862.3966 | Non-sig. |
| PF3D7_1336900 | cTrpRS | 590.980897 | -0.041474229 | 0.5054512 | -0.08205387 | 0.93460387 | 0.99639249 | 383.977342 | 558.04485 | 814.639071 | 607.262323 | Non-sig. |
| PF3D7_0215700 | RPB2 | 3393.18016 | -0.04160455 | 0.54295173 | -0.07662661 | 0.93892059 | 0.99639249 | 1794.2655 | 3433.72817 | 5089.55866 | 3255.1683 | Non-sig. |
| PF3D7_1314300 | N/A | 577.522986 | -0.041613152 | 0.4338667 | -0.0959123 | 0.92359023 | 0.99619209 | 531.945229 | 612.893719 | 639.64512 | 525.607875 | Non-sig. |
| PF3D7_1472000 | ISY1 | 184.216123 | -0.041626558 | 0.47827766 | -0.08703429 | 0.93064425 | 0.99619209 | 194.03964 | 200.849497 | 179.750502 | 162.224852 | Non-sig. |
| PF3D7_0418300 | N/A | 31882.022 | -0.041691423 | 0.47982505 | -0.0868888 | 0.9307599 | 0.99619209 | 43312.1834 | 29909.4359 | 21373.521 | 32932.9478 | Non-sig. |
| PF3D7_0626300 | FabB/FabF | 158.788482 | -0.041794101 | 0.57068576 | -0.07323488 | 0.94161921 | 0.99639249 | 106.034969 | 194.353768 | 215.900625 | 118.864564 | Non-sig. |
| PF3D7_1444200 | N/A | 352.696351 | -0.043036055 | 0.46390614 | -0.09276888 | 0.92608717 | 0.99619209 | 412.373031 | 390.91319 | 303.71016 | 303.789024 | Non-sig. |
| PF3D7_1318600 | N/A | 382.486744 | -0.043134897 | 0.44968768 | -0.0959219 | 0.9235826 | 0.99619209 | 438.277876 | 401.466749 | 338.28348 | 351.91887 | Non-sig. |
| PF3D7_0310700 | TRAPPC4 | 551.818875 | -0.043219449 | 0.53166928 | -0.0812901 | 0.93521125 | 0.99639249 | 324.68653 | 523.275966 | 795.134664 | 564.17834 | Non-sig. |
| PF3D7_0603000 | N/A | 1398.96158 | -0.043382517 | 0.49557262 | -0.08754018 | 0.93024215 | 0.99619209 | 1930.27548 | 1451.57214 | 910.061925 | 1303.93676 | Non-sig. |
| PF3D7_1118100 | AP1S | 315.7128 | -0.04340715 | 0.47849523 | -0.09071595 | 0.9277183 | 0.99619209 | 255.579389 | 272.59989 | 385.132429 | 349.539491 | Non-sig. |
| PF3D7_1226000 | N/A | 137.010883 | -0.043453627 | 0.50837152 | -0.08547612 | 0.93188288 | 0.99619209 | 148.44492 | 155.575052 | 129.768404 | 114.255157 | Non-sig. |
| PF3D7_0303700 | BCKDH-E2 | 2458.83929 | -0.044210555 | 0.41115281 | -0.10752828 | 0.91436989 | 0.99619209 | 2403.19576 | 2613.65948 | 2589.77904 | 2228.72287 | Non-sig. |
| PF3D7_0802400 | N/A | 604.368062 | -0.044265631 | 0.43236907 | -0.10237927 | 0.91845564 | 0.99619209 | 559.545885 | 549.491085 | 667.618839 | 640.816439 | Non-sig. |
| PF3D7_1359700 | N/A | 3897.6796 | -0.044304902 | 0.50156653 | -0.08833305 | 0.92961197 | 0.99619209 | 2726.73254 | 2892.44335 | 5188.01929 | 4783.52322 | Non-sig. |
| PF3D7_1018900 | N/A | 183.515205 | -0.044430234 | 0.49105208 | -0.09047968 | 0.92790604 | 0.99619209 | 155.714581 | 166.334386 | 216.803898 | 195.207953 | Non-sig. |
| PF3D7_1208800 | N/A | 1461.56994 | -0.044544094 | 0.44576584 | -0.09992712 | 0.92040219 | 0.99619209 | 1636.05282 | 1149.97402 | 1332.3291 | 1727.92381 | Non-sig. |
| PF3D7_1148800 | N/A | 114.329771 | -0.04473422 | 0.51958874 | -0.08609544 | 0.93139055 | 0.99619209 | 127.186034 | 100.90076 | 105.099503 | 124.132788 | Non-sig. |
| PF3D7_1030000 | SPT4 | 1033.25805 | -0.045137919 | 0.43270109 | -0.10431663 | 0.91691808 | 0.99619209 | 1191.45358 | 1108.99418 | 907.560417 | 925.024032 | Non-sig. |
| PF3D7_1129400 | NSUN3 | 786.071316 | -0.045141767 | 0.47852057 | -0.09433611 | 0.92484217 | 0.99619209 | 1039.35278 | 786.407954 | 557.697938 | 760.826594 | Non-sig. |
| PF3D7_0420100 | RIO2 | 1415.85695 | -0.04520355 | 0.42835816 | -0.10552746 | 0.91595728 | 0.99619209 | 1654.97274 | 1460.26712 | 1221.29748 | 1326.89047 | Non-sig. |
| PF3D7_0817800 | C3AP3 | 1478.1321 | -0.045287662 | 0.4195113 | -0.10795338 | 0.91403267 | 0.99619209 | 1475.63339 | 1623.36889 | 1527.00972 | 1286.5164 | Non-sig. |
| PF3D7_1129900 | MFR5 | 330.005906 | -0.045407939 | 0.58647616 | -0.07742504 | 0.93828542 | 0.99639249 | 171.548014 | 263.966453 | 498.510926 | 385.998232 | Non-sig. |
| PF3D7_0316300 | PPase | 8740.96332 | -0.045670715 | 0.4102244 | -0.11133105 | 0.91135383 | 0.99619209 | 9064.08892 | 9509.84295 | 8694.56113 | 7695.36027 | Non-sig. |
| PF3D7_1140400 | N/A | 5558.86396 | -0.046468761 | 0.41084698 | -0.11310479 | 0.90994748 | 0.99605273 | 5060.31111 | 5573.34176 | 6236.31199 | 5365.49097 | Non-sig. |
| PF3D7_0313000 | N/A | 800.55995 | -0.046800847 | 0.45763405 | -0.10226697 | 0.91854477 | 0.99619209 | 1004.32038 | 832.267759 | 623.032096 | 742.619566 | Non-sig. |
| PF3D7_1463500 | N/A | 625.36024 | -0.047325213 | 0.51593317 | -0.09172741 | 0.92691462 | 0.99619209 | 391.368249 | 657.836283 | 879.525166 | 572.711261 | Non-sig. |
| PF3D7_1431100 | PIP2 | 434.744481 | -0.047491599 | 0.48113044 | -0.09870836 | 0.92136983 | 0.99619209 | 562.128633 | 449.573335 | 321.950348 | 405.325607 | Non-sig. |
| PF3D7_1145300 | CRIPT | 413.754508 | -0.047809886 | 0.4860972 | -0.09835458 | 0.92165074 | 0.99619209 | 490.184569 | 503.931529 | 351.206598 | 309.695336 | Non-sig. |

| | | | | | | | | | | | | |
|---|---|---|---|---|---|---|---|---|---|---|---|---|
| PF3D7_0810200 | ABCK1 | 618.321382 | -0.048261305 | 0.42507337 | -0.11353641 | 0.9096053 | 0.99597145 | 588.660146 | 621.184367 | 668.582324 | 594.858693 | Non-sig. |
| PF3D7_0514300 | N/A | 1217.66516 | -0.04831388 | 0.41256132 | -0.11710715 | 0.90677515 | 0.99546099 | 1285.98401 | 1162.92674 | 1190.17432 | 1231.57556 | Non-sig. |
| PF3D7_0933500 | N/A | 1289.81788 | -0.048447692 | 0.49634546 | -0.09760881 | 0.92224293 | 0.99619209 | 1122.05846 | 841.003959 | 1500.74251 | 1695.46658 | Non-sig. |
| PF3D7_0607300 | UROD | 216.144112 | -0.048626252 | 0.49751509 | -0.09773825 | 0.92214015 | 0.99619209 | 256.616327 | 251.139412 | 183.120815 | 173.699894 | Non-sig. |
| PF3D7_1454400 | APP | 3259.896 | -0.048821004 | 0.49538378 | -0.09855188 | 0.92149408 | 0.99619209 | 2079.24422 | 3170.07644 | 4550.49519 | 3239.76813 | Non-sig. |
| PF3D7_1207500 | N/A | 536.627275 | -0.049485888 | 0.42749408 | -0.11575807 | 0.90784428 | 0.99580932 | 520.322911 | 545.597672 | 571.278723 | 509.309794 | Non-sig. |
| PF3D7_0717800 | N/A | 356.340946 | -0.049625187 | 0.57629231 | -0.08611114 | 0.93137807 | 0.99619209 | 533.061987 | 285.347785 | 192.210343 | 414.743671 | Non-sig. |
| PF3D7_1369700 | SNRPA1 | 313.014581 | -0.04963502 | 0.44502766 | -0.11153244 | 0.91119414 | 0.99619209 | 330.648739 | 293.154849 | 306.192394 | 322.062341 | Non-sig. |
| PF3D7_1250600 | N/A | 1038.48547 | -0.04992505 | 0.41516623 | -0.12025316 | 0.90428261 | 0.99457954 | 1110.93069 | 997.769121 | 1002.04475 | 1043.1973 | Non-sig. |
| PF3D7_0211900 | N/A | 794.345357 | -0.050328533 | 0.45507717 | -0.1105934 | 0.91193878 | 0.99619209 | 979.387494 | 856.388571 | 637.255964 | 704.349401 | Non-sig. |
| PF3D7_0526000 | N/A | 428.469626 | -0.050445972 | 0.4338797 | -0.11626719 | 0.90744079 | 0.99563532 | 450.623128 | 442.041008 | 421.340631 | 399.873735 | Non-sig. |
| PF3D7_0727100 | N/A | 787.297722 | -0.050481967 | 0.47800833 | -0.10560897 | 0.91589261 | 0.99619209 | 751.956087 | 544.996862 | 850.115758 | 1002.12218 | Non-sig. |
| PF3D7_0818300 | N/A | 121.77221 | -0.050530885 | 0.58462692 | -0.0864327 | 0.93112245 | 0.99619209 | 173.370317 | 121.308405 | 74.7142458 | 117.695871 | Non-sig. |
| PF3D7_1037000 | REV3 | 1706.85739 | -0.051872161 | 0.42597664 | -0.12177232 | 0.90307934 | 0.99457954 | 1530.46122 | 1510.18854 | 1944.45973 | 1842.32009 | Non-sig. |
| PF3D7_1145200 | N/A | 33497.7729 | -0.052413439 | 0.51571965 | -0.10163165 | 0.91904906 | 0.99619209 | 47303.6808 | 26473.1586 | 20909.1025 | 39305.1498 | Non-sig. |
| PF3D7_1362600 | N/A | 242.305057 | -0.052517673 | 0.51704357 | -0.10157301 | 0.9190956 | 0.99619209 | 322.980994 | 213.750998 | 170.715514 | 261.77272 | Non-sig. |
| PF3D7_0933400 | N/A | 138.412977 | -0.052580389 | 0.5544872 | -0.09482706 | 0.92445219 | 0.99619209 | 187.735942 | 147.148735 | 94.3885987 | 124.378633 | Non-sig. |
| PF3D7_1412200 | N/A | 414.732918 | -0.052708537 | 0.43275511 | -0.12179761 | 0.90305931 | 0.99457954 | 416.725045 | 398.10701 | 427.874177 | 416.225438 | Non-sig. |
| PF3D7_0503000 | N/A | 1705.19363 | -0.052818691 | 0.44755689 | -0.11801559 | 0.90605531 | 0.99520197 | 2129.43494 | 1776.44078 | 1343.64159 | 1571.25722 | Non-sig. |
| PF3D7_0719500 | CDC50A | 1997.26852 | -0.053157352 | 0.46340376 | -0.11471066 | 0.90867446 | 0.99597145 | 2608.37835 | 2032.69452 | 1460.0539 | 1887.9473 | Non-sig. |
| PF3D7_1235600 | SHMT | 4868.58308 | -0.053617441 | 0.49179429 | -0.10902412 | 0.91318336 | 0.99619209 | 3170.38186 | 5123.90612 | 6747.35925 | 4432.68511 | Non-sig. |
| PF3D7_1331600 | DEH | 997.405064 | -0.053833647 | 0.45024473 | -0.1195653 | 0.90482751 | 0.99483507 | 798.15795 | 1051.51607 | 1233.62286 | 906.323379 | Non-sig. |
| PF3D7_0315500 | N/A | 570.974066 | -0.054053172 | 0.43680796 | -0.12374585 | 0.90151651 | 0.99457954 | 552.471983 | 630.227353 | 610.80942 | 490.38751 | Non-sig. |
| PF3D7_1018100 | N/A | 19.8216813 | -0.054205125 | 0.87319278 | -0.06207693 | 0.95050157 | 0.99639249 | 24.0414409 | 23.6520035 | 16.407227 | 15.1860538 | Non-sig. |
| PF3D7_0302900 | N/A | 5032.63725 | -0.054373682 | 0.5780882 | -0.09405776 | 0.92506327 | 0.99619209 | 2311.0406 | 4719.60477 | 7943.50302 | 5156.40064 | Non-sig. |
| PF3D7_1459400 | N/A | 1272.86216 | -0.054550153 | 0.50915497 | -0.1071386 | 0.91467902 | 0.99619209 | 785.960513 | 1252.32677 | 1807.52953 | 1245.63181 | Non-sig. |
| PF3D7_0609600 | N/A | 108.507352 | -0.054618144 | 0.61763425 | -0.08843121 | 0.92953396 | 0.99619209 | 82.1463498 | 64.3697332 | 138.811139 | 148.702184 | Non-sig. |
| PF3D7_1140300 | N/A | 289.846305 | -0.05469771 | 0.45382095 | -0.12052707 | 0.90406564 | 0.99457954 | 265.319486 | 295.051855 | 325.245255 | 273.768623 | Non-sig. |
| PF3D7_1021400 | N/A | 1616.38041 | -0.055134371 | 0.58891562 | -0.09362015 | 0.9254109 | 0.99619209 | 721.035655 | 1653.47303 | 2573.1096 | 1517.90334 | Non-sig. |
| PF3D7_0915900 | N/A | 607.727378 | -0.055247012 | 0.44356656 | -0.1245518 | 0.90087839 | 0.99457954 | 724.849624 | 610.623535 | 514.079979 | 581.356375 | Non-sig. |
| PF3D7_1032400 | PIGA | 1961.28263 | -0.055303554 | 0.44061361 | -0.12551486 | 0.90011596 | 0.99457954 | 1701.59122 | 2223.7899 | 2295.9737 | 1623.7757 | Non-sig. |
| PF3D7_1461100 | N/A | 825.697571 | -0.05532777 | 0.42356357 | -0.13062448 | 0.89607238 | 0.99457954 | 781.157022 | 755.349471 | 901.803998 | 864.479792 | Non-sig. |
| PF3D7_1322700 | N/A | 110.540542 | -0.055590465 | 0.52791951 | -0.10530102 | 0.91613695 | 0.99619209 | 131.088462 | 103.114182 | 94.3870646 | 113.572462 | Non-sig. |
| PF3D7_1013200 | N/A | 5732.1577 | -0.055749414 | 0.4033762 | -0.138207 | 0.89007682 | 0.99457954 | 5775.32228 | 5369.53001 | 5910.45672 | 5873.3218 | Non-sig. |
| PF3D7_0411300 | N/A | 626.101729 | -0.0558392 | 0.4313968 | -0.12943814 | 0.89701097 | 0.99457954 | 606.215331 | 678.154932 | 670.158079 | 549.878576 | Non-sig. |
| PF3D7_0704700 | PPAT | 1036.67121 | -0.055863713 | 0.49246408 | -0.11343713 | 0.909684 | 0.99597145 | 1417.22964 | 939.661479 | 696.582502 | 1093.21121 | Non-sig. |
| PF3D7_1362400 | Pcalp | 2596.3091 | -0.055928094 | 0.44502315 | -0.12567457 | 0.89998953 | 0.99457954 | 2727.79357 | 1955.31879 | 2565.49518 | 3136.62885 | Non-sig. |
| PF3D7_0109800 | cPheRS | 909.268824 | -0.056478624 | 0.4210016 | -0.13415299 | 0.8932816 | 0.99457954 | 863.663627 | 838.14655 | 990.373246 | 944.891874 | Non-sig. |
| PF3D7_1023700 | N/A | 28.5346987 | -0.056646831 | 0.74392303 | -0.07614609 | 0.93930287 | 0.99639249 | 29.4534209 | 28.9760627 | 28.7419001 | 26.9674111 | Non-sig. |
| PF3D7_1119500 | N/A | 491.210591 | -0.056706507 | 0.43767287 | -0.12956368 | 0.89691164 | 0.99457954 | 445.031859 | 461.755004 | 556.557172 | 501.49833 | Non-sig. |
| PF3D7_1418200 | N/A | 253.841771 | -0.05724165 | 0.53535493 | -0.1069228 | 0.91485022 | 0.99619209 | 351.492074 | 281.613077 | 166.554237 | 215.707698 | Non-sig. |
| PF3D7_0506900 | ROM4 | 16638.1439 | -0.057321836 | 0.40516443 | -0.14147796 | 0.88749237 | 0.99457954 | 16803.1027 | 15028.7269 | 17134.1697 | 17586.5765 | Non-sig. |
| PF3D7_1225500 | N/A | 270.228856 | -0.057484108 | 0.48416605 | -0.11872809 | 0.90549078 | 0.99508197 | 339.606248 | 273.699053 | 211.851377 | 255.758747 | Non-sig. |
| PF3D7_1030500 | RPN9 | 2765.38809 | -0.057641388 | 0.46691502 | -0.12345156 | 0.90174953 | 0.99457954 | 1977.57335 | 2702.29132 | 3663.3561 | 2718.33161 | Non-sig. |
| PF3D7_1405200 | BET5 | 114.345646 | -0.058070567 | 0.51615745 | -0.11250553 | 0.91042259 | 0.99616251 | 105.980839 | 104.171365 | 127.23036 | 120.000021 | Non-sig. |
| PF3D7_1406200 | SPT6 | 7840.13516 | -0.058215163 | 0.48721446 | -0.1194857 | 0.90489057 | 0.99483507 | 10860.8422 | 7711.39785 | 5136.11415 | 7652.18647 | Non-sig. |
| PF3D7_1242200 | TGT | 1359.96094 | -0.058309298 | 0.41160476 | -0.14166332 | 0.88734595 | 0.99457954 | 1405.1733 | 1398.41246 | 1369.72686 | 1266.53114 | Non-sig. |
| PF3D7_0414000 | SMC3 | 4307.57747 | -0.058371894 | 0.51716405 | -0.1128692 | 0.91013425 | 0.99605273 | 6254.96393 | 4409.0668 | 2534.83325 | 4031.4459 | Non-sig. |
| PF3D7_1351000 | N/A | 4830.85277 | -0.05844009 | 0.4038676 | -0.14470111 | 0.88494686 | 0.99457954 | 5060.69925 | 4551.94893 | 4796.707 | 4914.05589 | Non-sig. |
| PF3D7_1330700 | N/A | 402.807448 | -0.058782851 | 0.53086243 | -0.11073086 | 0.91182978 | 0.99619209 | 245.175068 | 403.481227 | 576.513542 | 386.059954 | Non-sig. |
| PF3D7_1114600 | N/A | 142.631927 | -0.058970453 | 0.6152298 | -0.0958511 | 0.92363883 | 0.99619209 | 212.606218 | 169.771685 | 78.7742221 | 109.375585 | Non-sig. |
| PF3D7_1313400 | N/A | 1517.38155 | -0.059433299 | 0.41173134 | -0.14434971 | 0.88522433 | 0.99457954 | 1640.31986 | 1522.47499 | 1457.02863 | 1449.70272 | Non-sig. |
| PF3D7_0614900 | N/A | 2347.55608 | -0.059461461 | 0.45973007 | -0.12933994 | 0.89708867 | 0.99457954 | 3056.58373 | 2351.57199 | 1735.58144 | 2246.48716 | Non-sig. |
| PF3D7_1403700 | TRAM | 133.97878 | -0.059901579 | 0.50816149 | -0.11787902 | 0.90616352 | 0.99520197 | 115.709395 | 129.855838 | 157.669739 | 132.680147 | Non-sig. |

| | | | | | | | | | | | | |
|---|---|---|---|---|---|---|---|---|---|---|---|---|
| PF3D7_1358400 | N/A | 875.065502 | -0.059995395 | 0.42703391 | -0.14049328 | 0.88827026 | 0.99457954 | 865.527826 | 960.143774 | 920.954109 | 753.636299 | Non-sig. |
| PF3D7_1313800 | N/A | 4249.26716 | -0.060045588 | 0.49460416 | -0.1214013 | 0.90337319 | 0.99457954 | 2852.02928 | 3508.10154 | 5823.00534 | 4813.93247 | Non-sig. |
| PF3D7_0419200 | GOT1 | 199.925856 | -0.060057483 | 0.54945194 | -0.10930434 | 0.9129611 | 0.99619209 | 155.192129 | 134.267286 | 252.7884 | 257.455609 | Non-sig. |
| PF3D7_1306800 | N/A | 869.579766 | -0.060198178 | 0.44923161 | -0.13400254 | 0.89340056 | 0.99457954 | 1078.57132 | 875.877574 | 697.118664 | 826.751503 | Non-sig. |
| PF3D7_0415500 | CBP20 | 836.25907 | -0.060518209 | 0.43221182 | -0.14001979 | 0.88864435 | 0.99457954 | 892.616786 | 933.349327 | 815.038655 | 704.031513 | Non-sig. |
| PF3D7_0604400 | N/A | 2.22166985 | -0.060800537 | 2.2077386 | -0.02753974 | 0.97802925 | 0.99691622 | 2.07368371 | 1.07750346 | 2.45772315 | 3.27776909 | Non-sig. |
| PF3D7_1142800 | ATPTG6 | 595.632453 | -0.060964614 | 0.43146599 | -0.14129645 | 0.88763575 | 0.99457954 | 672.379922 | 600.385644 | 544.184583 | 565.579663 | Non-sig. |
| PF3D7_0904800 | RPA1 | 3730.51614 | -0.061815085 | 0.44583491 | -0.13865017 | 0.8897266 | 0.99457954 | 2895.91692 | 3726.79418 | 4724.65171 | 3574.70175 | Non-sig. |
| PF3D7_1128600 | NOT2 | 144.154255 | -0.062094328 | 0.64220693 | -0.09668897 | 0.9229734 | 0.99619209 | 228.30883 | 155.603598 | 66.5192549 | 126.185339 | Non-sig. |
| PF3D7_0726500 | N/A | 11492.2543 | -0.062244186 | 0.47336742 | -0.13149233 | 0.89538585 | 0.99457954 | 8473.01419 | 9294.09512 | 15006.9582 | 13194.9497 | Non-sig. |
| PF3D7_0910800 | NBP35 | 1039.73492 | -0.062298999 | 0.43089733 | -0.14457968 | 0.88504274 | 0.99457954 | 898.159927 | 1026.02166 | 1226.00684 | 1008.75125 | Non-sig. |
| PF3D7_1038500 | N/A | 133.156133 | -0.06246339 | 0.50802736 | -0.12295281 | 0.90214447 | 0.99457954 | 155.608479 | 136.313003 | 116.600295 | 124.102756 | Non-sig. |
| PF3D7_1023400 | N/A | 705.793185 | -0.062823769 | 0.42468287 | -0.14793102 | 0.88239721 | 0.99457954 | 749.381596 | 743.894271 | 692.988488 | 636.908383 | Non-sig. |
| PF3D7_1332900 | cyto-IRS | 2963.65156 | -0.06323589 | 0.59058829 | -0.10707271 | 0.91473129 | 0.99619209 | 1299.09116 | 2974.60184 | 4757.71092 | 2823.20231 | Non-sig. |
| PF3D7_0406900 | N/A | 147.427467 | -0.063753326 | 0.73202636 | -0.08709157 | 0.93059873 | 0.99619209 | 258.148609 | 154.466326 | 43.5361578 | 133.558774 | Non-sig. |
| PF3D7_0825400 | ATPTG15 | 251.476766 | -0.063789482 | 0.48675493 | -0.13105051 | 0.89573535 | 0.99457954 | 198.787224 | 259.030541 | 315.065992 | 233.023305 | Non-sig. |
| PF3D7_0720000 | CSL4 | 509.614878 | -0.064110289 | 0.42768652 | -0.14990019 | 0.88084336 | 0.99421828 | 521.469488 | 519.794128 | 520.406136 | 476.789757 | Non-sig. |
| PF3D7_1305900 | N/A | 901.457983 | -0.064230734 | 0.44771176 | -0.14346448 | 0.88592336 | 0.99457954 | 725.063821 | 879.911587 | 1117.73007 | 883.126457 | Non-sig. |
| PF3D7_0721500 | N/A | 330.190749 | -0.064913141 | 0.49119788 | -0.13215273 | 0.89486349 | 0.99457954 | 428.302023 | 351.964931 | 247.198071 | 293.29797 | Non-sig. |
| PF3D7_1478000 | GEXP17 | 87.1277965 | -0.065113904 | 0.55876907 | -0.11653097 | 0.90723174 | 0.99563532 | 106.082686 | 79.4495159 | 72.246235 | 90.7327493 | Non-sig. |
| PF3D7_1136600 | N/A | 1766.16318 | -0.065202199 | 0.49248183 | -0.13239514 | 0.89467176 | 0.99457954 | 2437.00845 | 1572.57576 | 1175.46574 | 1879.60278 | Non-sig. |
| PF3D7_0518300 | N/A | 3944.27185 | -0.065671518 | 0.43025955 | -0.15263233 | 0.87868823 | 0.99421828 | 3352.68797 | 4185.74296 | 4715.16117 | 3523.4953 | Non-sig. |
| PF3D7_1417100 | N/A | 342.26933 | -0.066005484 | 0.47132477 | -0.14004247 | 0.88862644 | 0.99457954 | 274.579848 | 349.847702 | 425.388771 | 319.261001 | Non-sig. |
| PF3D7_0313800 | N/A | 1108.19567 | -0.066181982 | 0.41267713 | -0.1603723 | 0.87258781 | 0.99402081 | 1125.55787 | 1116.88981 | 1141.65357 | 1048.68143 | Non-sig. |
| PF3D7_1439400 | N/A | 3264.74097 | -0.066261244 | 0.4412006 | -0.15018394 | 0.8806195 | 0.99421828 | 2624.81502 | 3411.55167 | 4054.33275 | 2968.26444 | Non-sig. |
| PF3D7_0824900 | N/A | 512.814338 | -0.066446681 | 0.58571017 | -0.11344635 | 0.90967669 | 0.99597145 | 794.150416 | 584.677436 | 255.449223 | 416.980277 | Non-sig. |
| PF3D7_0731700 | N/A | 152.703759 | -0.066506585 | 0.53712136 | -0.12382041 | 0.90145747 | 0.99457954 | 121.804126 | 115.962277 | 190.458153 | 182.59048 | Non-sig. |
| PF3D7_0915800 | GLTP | 154.032556 | -0.066867039 | 0.52058169 | -0.12844678 | 0.89779542 | 0.99457954 | 190.656114 | 172.984105 | 124.730917 | 127.75909 | Non-sig. |
| PF3D7_1418900 | DBP4 | 807.999742 | -0.06687823 | 0.50404881 | -0.13268205 | 0.89444484 | 0.99457954 | 1125.66066 | 855.080023 | 528.125864 | 723.132422 | Non-sig. |
| PF3D7_1119000 | N/A | 2160.06331 | -0.067631595 | 0.44864579 | -0.15074608 | 0.88017602 | 0.99421828 | 2612.42467 | 2453.16472 | 1809.16246 | 1765.5014 | Non-sig. |
| PF3D7_1334800 | MSRP2 | 1861.51372 | -0.06779464 | 0.45300312 | -0.14965601 | 0.88103601 | 0.99421828 | 1572.09985 | 2155.55912 | 2238.1955 | 1480.20043 | Non-sig. |
| PF3D7_0729000 | SRP9 | 180.122414 | -0.067797058 | 0.49329284 | -0.13743775 | 0.89068479 | 0.99457954 | 218.830898 | 174.313192 | 150.050522 | 177.295042 | Non-sig. |
| PF3D7_1251300 | TMK | 725.189314 | -0.067873637 | 0.43117809 | -0.15741439 | 0.87491827 | 0.99402081 | 748.203937 | 800.066385 | 736.301725 | 616.185211 | Non-sig. |
| PF3D7_0514700 | N/A | 40.8492355 | -0.068127715 | 0.80241781 | -0.08490304 | 0.93233847 | 0.99619209 | 67.4246778 | 40.7636598 | 16.4283117 | 38.780293 | Non-sig. |
| PF3D7_1352400 | N/A | 2244.30192 | -0.068568297 | 0.54661006 | -0.1254428 | 0.90017301 | 0.99457954 | 1231.86241 | 1851.39775 | 3363.02216 | 2530.92538 | Non-sig. |
| PF3D7_0828100 | N/A | 365.683175 | -0.069302962 | 0.44238188 | -0.15665868 | 0.87551385 | 0.99402081 | 342.254321 | 366.914158 | 406.572416 | 346.991804 | Non-sig. |
| PF3D7_0115700 | VAR | 206.036329 | -0.0697101 | 0.5545946 | -0.1256956 | 0.89997289 | 0.99457954 | 290.884189 | 231.683364 | 131.435566 | 170.142196 | Non-sig. |
| PF3D7_0619800 | N/A | 1591.60464 | -0.069827894 | 0.42653466 | -0.16370978 | 0.86995962 | 0.99402081 | 1490.59473 | 1352.47193 | 1769.5216 | 1753.8303 | Non-sig. |
| PF3D7_1367700 | AlaRS | 2777.21017 | -0.069834877 | 0.50086258 | -0.13942922 | 0.88911099 | 0.99457954 | 1756.31216 | 2837.31523 | 3932.15932 | 2583.05396 | Non-sig. |
| PF3D7_1118200 | N/A | 1316.69046 | -0.070086943 | 0.4316277 | -0.16237823 | 0.87100801 | 0.99402081 | 1121.71391 | 1265.90623 | 1575.40779 | 1303.7339 | Non-sig. |
| PF3D7_0616300 | N/A | 157.389931 | -0.070557855 | 0.54965005 | -0.12836869 | 0.89785721 | 0.99457954 | 207.783799 | 185.785303 | 114.919635 | 121.070985 | Non-sig. |
| PF3D7_1414400 | PP1 | 9822.47127 | -0.070690645 | 0.43034389 | -0.16426548 | 0.86952216 | 0.99402081 | 11242.5598 | 8062.70425 | 8883.73024 | 11100.8907 | Non-sig. |
| PF3D7_1205100 | SEPSECS | 857.213143 | -0.070753422 | 0.44366815 | -0.15947375 | 0.87329564 | 0.99402081 | 987.943573 | 963.66695 | 768.668052 | 708.573998 | Non-sig. |
| PF3D7_0925200 | RRP8 | 372.053637 | -0.070990208 | 0.53185001 | -0.13347787 | 0.89381546 | 0.99457954 | 512.321401 | 442.39787 | 250.378538 | 283.116738 | Non-sig. |
| PF3D7_0406400 | cGLO2 | 807.032304 | -0.071041466 | 0.44864754 | -0.15834583 | 0.8741843 | 0.99402081 | 1001.32094 | 784.752569 | 652.72061 | 789.335096 | Non-sig. |
| PF3D7_1450800 | N/A | 225.574289 | -0.0713539 | 0.4952062 | -0.14408927 | 0.88542998 | 0.99457954 | 175.51975 | 229.716859 | 286.555206 | 210.505341 | Non-sig. |
| PF3D7_1358500 | N/A | 277.233654 | -0.071436864 | 0.4701405 | -0.15194791 | 0.87922802 | 0.99421828 | 249.416072 | 307.965251 | 318.649961 | 232.90333 | Non-sig. |
| PF3D7_1134700 | RPA2 | 774.810716 | -0.071809941 | 0.46424042 | -0.15468265 | 0.8770715 | 0.99421828 | 589.022929 | 731.561769 | 998.874004 | 779.784162 | Non-sig. |
| PF3D7_1452900 | N/A | 432.000097 | -0.071876381 | 0.46205686 | -0.15555744 | 0.87638188 | 0.99402081 | 423.439455 | 509.206629 | 462.040222 | 333.314082 | Non-sig. |
| PF3D7_1330400 | N/A | 3887.12035 | -0.071971686 | 0.50279739 | -0.14314252 | 0.88617762 | 0.99457954 | 2422.89427 | 3724.88235 | 5544.86352 | 3855.84127 | Non-sig. |
| PF3D7_1311400 | AP1M1 | 2659.78725 | -0.072439005 | 0.45739894 | -0.15837161 | 0.87416399 | 0.99402081 | 1992.03591 | 2500.25353 | 3460.76021 | 2686.09936 | Non-sig. |
| PF3D7_0501900 | TRAPPC13 | 736.686575 | -0.072582441 | 0.42833169 | -0.16945382 | 0.86543969 | 0.99402081 | 702.350046 | 784.601919 | 807.992606 | 651.801729 | Non-sig. |
| PF3D7_1462600 | N/A | 8.3085934 | -0.072853887 | 1.2313941 | -0.05916375 | 0.95282169 | 0.99639249 | 10.5039606 | 6.44172788 | 6.56791922 | 9.72076588 | Non-sig. |
| PF3D7_0908400 | N/A | 883.399917 | -0.073114133 | 0.52872731 | -0.13828326 | 0.89001656 | 0.99457954 | 520.406817 | 798.575516 | 1290.79693 | 923.820402 | Non-sig. |

| | | | | | | | | | | | | |
|---|---|---|---|---|---|---|---|---|---|---|---|---|
| PF3D7_0918000 | GAP50 | 33346.9648 | -0.073332564 | 0.50878777 | -0.14413193 | 0.88539629 | 0.99457954 | 20134.2114 | 32904.1784 | 48254.0606 | 32095.4088 | Non-sig. |
| PF3D7_0509600 | AsnRS | 1461.18078 | -0.073641114 | 0.41413986 | -0.17781701 | 0.85886669 | 0.99402081 | 1406.05059 | 1500.10554 | 1590.7978 | 1347.76918 | Non-sig. |
| PF3D7_1027800 | RPL3 | 8947.14391 | -0.07379218 | 0.43606263 | -0.16922381 | 0.8656206 | 0.99402081 | 7247.8351 | 9147.40961 | 11103.7396 | 8289.59129 | Non-sig. |
| PF3D7_1130600 | N/A | 438.7795 | -0.074592843 | 0.44300958 | -0.16837749 | 0.86628631 | 0.99402081 | 501.207165 | 455.976906 | 399.171325 | 398.762603 | Non-sig. |
| PF3D7_1138200 | N/A | 198.328773 | -0.074874857 | 0.51534701 | -0.14529018 | 0.88448177 | 0.99457954 | 192.03003 | 248.452888 | 214.882599 | 137.949575 | Non-sig. |
| PF3D7_1029500 | N/A | 3.90362486 | -0.074936641 | 1.70001928 | -0.04407988 | 0.96484073 | 0.99691622 | 3.08812144 | 4.3209842 | 4.90565186 | 3.29974195 | Non-sig. |
| PF3D7_0612800 | P12p | 103.337219 | -0.074942609 | 0.55761424 | -0.13439866 | 0.89308734 | 0.99457954 | 82.9797462 | 83.7428997 | 128.894993 | 117.731235 | Non-sig. |
| PF3D7_1016600 | N/A | 345.850451 | -0.075158123 | 0.47889194 | -0.15694171 | 0.87529078 | 0.99402081 | 266.915393 | 336.002873 | 442.547016 | 337.936521 | Non-sig. |
| PF3D7_1117700 | RAN | 17457.0033 | -0.075443588 | 0.48118863 | -0.15678589 | 0.87541359 | 0.99402081 | 23562.7369 | 19512.3567 | 12264.2917 | 14488.6277 | Non-sig. |
| PF3D7_0518000 | N/A | 833.88689 | -0.076015566 | 0.44646842 | -0.17025967 | 0.86480593 | 0.99402081 | 1017.52947 | 877.476195 | 694.394618 | 746.147275 | Non-sig. |
| PF3D7_0410300 | PPM1 | 1231.94625 | -0.076323589 | 0.41289646 | -0.18484922 | 0.8533473 | 0.99276109 | 1311.03907 | 1246.66545 | 1218.06509 | 1152.01539 | Non-sig. |
| PF3D7_1012500 | N/A | 864.737941 | -0.076368597 | 0.6007593 | -0.12712012 | 0.89884532 | 0.99457954 | 376.95045 | 812.180114 | 1397.91588 | 871.905323 | Non-sig. |
| PF3D7_0622800 | N/A | 3633.73459 | -0.076454775 | 0.45506345 | -0.16800905 | 0.86657616 | 0.99402081 | 2872.71163 | 3034.49677 | 4587.01879 | 4040.71115 | Non-sig. |
| PF3D7_0516800 | ApiAP2 | 10125.1668 | -0.077030428 | 0.42282806 | -0.18217908 | 0.85544219 | 0.99321133 | 10385.6484 | 8199.5673 | 10405.1773 | 11510.2742 | Non-sig. |
| PF3D7_1249400 | N/A | 692.820398 | -0.077195994 | 0.42252067 | -0.18270348 | 0.85503069 | 0.99321117 | 706.385527 | 717.927436 | 716.312866 | 630.655764 | Non-sig. |
| PF3D7_1249900 | KsgA2 | 1304.01144 | -0.077295468 | 0.60739164 | -0.12725804 | 0.89873617 | 0.99457954 | 2111.59614 | 1490.57466 | 566.645621 | 1047.22936 | Non-sig. |
| PF3D7_0614800 | NTHL1 | 1574.99621 | -0.077409355 | 0.43644998 | -0.17736135 | 0.85922457 | 0.99402081 | 1883.10412 | 1692.63468 | 1351.582 | 1372.66403 | Non-sig. |
| PF3D7_1339400 | ApiCOX14 | 1725.82038 | -0.077675944 | 0.44237913 | -0.17558682 | 0.86061856 | 0.99402081 | 1480.72192 | 1921.21334 | 2063.62831 | 1437.71794 | Non-sig. |
| PF3D7_1344500 | USB1 | 918.395039 | -0.0777965 | 0.42284151 | -0.18398501 | 0.85402521 | 0.99295786 | 1008.27484 | 963.628998 | 878.120441 | 823.555884 | Non-sig. |
| PF3D7_0704600 | UT | 22728.0694 | -0.078331654 | 0.41307738 | -0.18962949 | 0.84959947 | 0.99165628 | 24524.6769 | 19509.0322 | 22165.2743 | 24713.2942 | Non-sig. |
| PF3D7_0609900 | N/A | 2311.11244 | -0.078713605 | 0.41598907 | -0.18922037 | 0.84992011 | 0.99165628 | 2106.18527 | 2192.28304 | 2641.95021 | 2304.03124 | Non-sig. |
| PF3D7_0303000 | NSF | 482.315797 | -0.078982301 | 0.58410103 | -0.13522027 | 0.89243772 | 0.99457954 | 248.298124 | 375.595266 | 742.387273 | 562.982524 | Non-sig. |
| PF3D7_1347100 | TOP3 | 1503.06081 | -0.079067025 | 0.45839355 | -0.17248721 | 0.86305451 | 0.99402081 | 1945.59803 | 1536.6667 | 1143.17023 | 1386.80829 | Non-sig. |
| PF3D7_0911000 | N/A | 479.095578 | -0.079378617 | 0.47254409 | -0.1679814 | 0.86659791 | 0.99402081 | 370.961283 | 501.316049 | 613.329294 | 430.775686 | Non-sig. |
| PF3D7_0602500 | N/A | 344.355073 | -0.079391927 | 0.50969084 | -0.15576487 | 0.87621836 | 0.99402081 | 240.919972 | 375.601131 | 466.451187 | 294.448001 | Non-sig. |
| PF3D7_1024000 | N/A | 1196.63254 | -0.079459772 | 0.41756199 | -0.19029455 | 0.84907832 | 0.99165628 | 1191.12153 | 1264.10689 | 1267.99581 | 1063.30593 | Non-sig. |
| PF3D7_0508500 | ND6 | 8918.76125 | -0.079543993 | 0.42781474 | -0.18593093 | 0.85249892 | 0.99253574 | 7558.2134 | 8109.52243 | 10770.7071 | 9236.60205 | Non-sig. |
| PF3D7_0601900 | N/A | 596.060532 | -0.079659232 | 0.50929731 | -0.15641008 | 0.87570979 | 0.99402081 | 392.700459 | 520.369585 | 831.998283 | 639.173801 | Non-sig. |
| PF3D7_1241100 | PFS2 | 1020.64701 | -0.080323771 | 0.42678784 | -0.18820539 | 0.85071565 | 0.99165628 | 942.546149 | 1079.61482 | 1155.43226 | 904.994833 | Non-sig. |
| PF3D7_0603500 | CAX | 837.101296 | -0.080423975 | 0.61678391 | -0.13039247 | 0.89625593 | 0.99457954 | 340.850865 | 789.844691 | 1379.63439 | 838.07524 | Non-sig. |
| PF3D7_1033700 | BDP1 | 1114.54186 | -0.080434045 | 0.44729509 | -0.17982323 | 0.85729135 | 0.99352464 | 899.74995 | 1140.66683 | 1391.20775 | 1026.54289 | Non-sig. |
| PF3D7_1229800 | MyoJ | 12427.4357 | -0.080879321 | 0.43815266 | -0.18459165 | 0.85354934 | 0.99277813 | 13021.0152 | 9385.80643 | 12530.4015 | 14772.5196 | Non-sig. |
| PF3D7_1403300 | N/A | 89.6525636 | -0.08104012 | 0.5707209 | -0.14199606 | 0.88708312 | 0.99457954 | 110.565661 | 102.995758 | 73.9212293 | 71.1276058 | Non-sig. |
| PF3D7_1449700 | RRP6 | 813.226503 | -0.081413022 | 0.43814542 | -0.18581279 | 0.85259157 | 0.99253574 | 822.939001 | 655.40372 | 849.370192 | 925.193098 | Non-sig. |
| PF3D7_1361000 | PRMT5 | 342.028912 | -0.081629453 | 0.48990788 | -0.16662204 | 0.86766745 | 0.99402081 | 258.852983 | 298.319014 | 444.292785 | 366.650866 | Non-sig. |
| PF3D7_1021200 | N/A | 2571.00755 | -0.082224523 | 0.46849342 | -0.17550838 | 0.86068018 | 0.99402081 | 3438.58134 | 2408.04637 | 1850.24631 | 2587.15617 | Non-sig. |
| PF3D7_0918100 | N/A | 4271.34862 | -0.082413715 | 0.54321803 | -0.15171388 | 0.87941261 | 0.99421828 | 6459.92627 | 4622.82108 | 2327.08365 | 3675.56348 | Non-sig. |
| PF3D7_0716300 | N/A | 14903.5254 | -0.082691124 | 0.43475309 | -0.1902025 | 0.84915045 | 0.99165628 | 12093.7429 | 14054.945 | 18567.0531 | 14898.3609 | Non-sig. |
| PF3D7_1309200 | PPM6 | 7434.77039 | -0.082729461 | 0.41876045 | -0.19755796 | 0.84339093 | 0.99164151 | 8744.32645 | 6927.97093 | 6551.63068 | 7515.15351 | Non-sig. |
| PF3D7_0523300 | ApiCOX18 | 773.08581 | -0.083035068 | 0.4433707 | -0.18728136 | 0.85144004 | 0.99184901 | 860.102139 | 879.345293 | 730.655274 | 622.240534 | Non-sig. |
| PF3D7_1354100 | N/A | 101.903571 | -0.083219365 | 0.60759347 | -0.13696554 | 0.89105804 | 0.99457954 | 64.107358 | 81.5833494 | 145.328441 | 116.595133 | Non-sig. |
| PF3D7_0529200 | N/A | 99.6294872 | -0.083338516 | 0.61675183 | -0.13512488 | 0.89251314 | 0.99457954 | 57.7844674 | 103.061292 | 146.962865 | 90.7093246 | Non-sig. |
| PF3D7_0708900 | SCO1 | 322.01152 | -0.083365792 | 0.44342291 | -0.18800515 | 0.85087261 | 0.99165628 | 349.024049 | 309.130109 | 313.664341 | 316.227582 | Non-sig. |
| PF3D7_0624100 | N/A | 573.825073 | -0.08354686 | 0.50206266 | -0.16640724 | 0.86783648 | 0.99402081 | 384.86095 | 586.019156 | 795.690296 | 528.729891 | Non-sig. |
| PF3D7_0304200 | EHD | 1566.39185 | -0.084457146 | 0.50044676 | -0.1687635 | 0.86598267 | 0.99402081 | 2215.53625 | 1451.98248 | 1009.2687 | 1588.77996 | Non-sig. |
| PF3D7_1451800 | SORTLR | 4412.7162 | -0.084622901 | 0.53023891 | -0.15959391 | 0.87320098 | 0.99402081 | 2490.14514 | 4071.64988 | 6593.67438 | 4495.39539 | Non-sig. |
| PF3D7_1406400 | PPR1 | 627.161019 | -0.084753145 | 0.5105687 | -0.16599753 | 0.86815889 | 0.99402081 | 403.929876 | 636.401412 | 886.891859 | 581.420927 | Non-sig. |
| PF3D7_0505500 | MSH6 | 3888.74414 | -0.086313316 | 0.51891996 | -0.16633262 | 0.8678952 | 0.99402081 | 2308.25654 | 3503.52295 | 5701.45195 | 4041.74512 | Non-sig. |
| PF3D7_1369100 | N/A | 330.031391 | -0.086433601 | 0.49759615 | -0.17370231 | 0.86209942 | 0.99402081 | 441.95209 | 328.363867 | 238.161394 | 311.648212 | Non-sig. |
| PF3D7_0812100 | N/A | 4163.4998 | -0.08646926 | 0.52145628 | -0.16582265 | 0.86829652 | 0.99402081 | 2458.34544 | 3672.36424 | 6117.75742 | 4405.53211 | Non-sig. |
| PF3D7_0819500 | N/A | 272.920801 | -0.08658415 | 0.45438145 | -0.19055389 | 0.84887512 | 0.99165628 | 269.048985 | 286.612713 | 293.121659 | 242.899848 | Non-sig. |
| PF3D7_1402300 | RPN6 | 6781.90272 | -0.086593887 | 0.45837631 | -0.18891441 | 0.8501599 | 0.99165628 | 5033.36736 | 6320.29275 | 8937.07577 | 6836.87502 | Non-sig. |
| PF3D7_1310100 | N/A | 134.68394 | -0.086596036 | 0.50756804 | -0.17060971 | 0.86453066 | 0.99402081 | 122.112435 | 118.025843 | 155.226526 | 143.370956 | Non-sig. |
| PF3D7_1465000 | COX6A | 2478.25147 | -0.086683808 | 0.41024586 | -0.21129722 | 0.83265535 | 0.99110542 | 2439.46629 | 2567.97013 | 2665.84004 | 2239.72942 | Non-sig. |

| | | | | | | | | | | | | |
|---|---|---|---|---|---|---|---|---|---|---|---|---|
| PF3D7_1448700 | N/A | 181.496232 | -0.087301438 | 0.47498786 | -0.1837972 | 0.85417255 | 0.99295786 | 200.794356 | 179.252428 | 173.254448 | 172.683696 | Non-sig. |
| PF3D7_0210000 | Sec61-gamma | 3506.34386 | -0.087645326 | 0.43826668 | -0.19998172 | 0.84149487 | 0.99128968 | 2835.04494 | 3354.30447 | 4390.33584 | 3445.6902 | Non-sig. |
| PF3D7_1460100 | FCP | 183.476938 | -0.08772824 | 0.58519823 | -0.14991201 | 0.88083404 | 0.99421828 | 274.144853 | 155.676449 | 104.265148 | 199.821301 | Non-sig. |
| PF3D7_1446600 | CEN2 | 6522.79879 | -0.08793123 | 0.40475871 | -0.21724357 | 0.82801853 | 0.99110542 | 6624.02784 | 6723.25039 | 6818.99362 | 5924.92332 | Non-sig. |
| PF3D7_1406300 | GDPD | 1906.23606 | -0.088626595 | 0.4459599 | -0.19873221 | 0.84247223 | 0.99128968 | 1516.42219 | 1919.71062 | 2412.85069 | 1775.96076 | Non-sig. |
| PF3D7_0305400 | N/A | 208.513898 | -0.089228131 | 0.4884305 | -0.18268337 | 0.85504646 | 0.99321117 | 230.574442 | 240.641066 | 199.422364 | 163.417721 | Non-sig. |
| PF3D7_0216100 | N/A | 259.65554 | -0.089722315 | 0.45221508 | -0.19840629 | 0.8427272 | 0.99128968 | 256.965112 | 246.752136 | 278.442601 | 256.462311 | Non-sig. |
| PF3D7_1132300 | N/A | 1760.49343 | -0.090189864 | 0.4361615 | -0.20678089 | 0.83618098 | 0.99128968 | 1480.27688 | 1819.33238 | 2150.50929 | 1591.85515 | Non-sig. |
| PF3D7_0705400 | MCM7 | 7734.41481 | -0.090541486 | 0.44823172 | -0.20199705 | 0.83991903 | 0.99128968 | 5989.51101 | 7848.44633 | 9964.27913 | 7135.42278 | Non-sig. |
| PF3D7_0814100 | N/A | 203.304702 | -0.090633676 | 0.4660669 | -0.19446495 | 0.84581184 | 0.99165628 | 219.874009 | 192.094358 | 199.55492 | 201.69552 | Non-sig. |
| PF3D7_0524000 | KASbeta | 13275.0541 | -0.090671684 | 0.59120877 | -0.15336661 | 0.87810918 | 0.99421828 | 5954.80274 | 11045.6623 | 21428.994 | 14670.7574 | Non-sig. |
| PF3D7_1307900 | N/A | 3495.57394 | -0.090678445 | 0.41047398 | -0.22091156 | 0.8251613 | 0.99096399 | 3945.15647 | 3411.58968 | 3265.76158 | 3359.78804 | Non-sig. |
| PF3D7_0423300 | N/A | 5981.75078 | -0.090680115 | 0.7372698 | -0.12299448 | 0.90211147 | 0.99457954 | 11040.583 | 6344.72368 | 1299.1289 | 5242.56752 | Non-sig. |
| PF3D7_1438000 | N/A | 3761.52984 | -0.09083165 | 0.41638714 | -0.21814231 | 0.82731824 | 0.99110542 | 4379.20961 | 3693.70777 | 3380.77326 | 3592.42871 | Non-sig. |
| PF3D7_0318900 | N/A | 2468.50913 | -0.091313833 | 0.43271473 | -0.21102548 | 0.83286739 | 0.99110542 | 2956.56842 | 2638.79055 | 2136.84446 | 2141.83307 | Non-sig. |
| PF3D7_0829000 | N/A | 2481.44237 | -0.0915495 | 0.47450905 | -0.19293521 | 0.84700971 | 0.99165628 | 1905.58451 | 1932.81641 | 3214.44523 | 2872.92334 | Non-sig. |
| PF3D7_1115500 | ApiAP2 | 1365.18408 | -0.09176368 | 0.43735275 | -0.20981617 | 0.83381115 | 0.9911446 | 1241.17365 | 1523.95307 | 1575.86006 | 1119.74954 | Non-sig. |
| PF3D7_1216900 | N/A | 6143.21034 | -0.092464549 | 0.48697618 | -0.18987489 | 0.84940717 | 0.99165628 | 8576.33617 | 5622.71019 | 4104.03127 | 6269.76375 | Non-sig. |
| PF3D7_1230400 | ClpQ | 327.308641 | -0.092566478 | 0.47236712 | -0.195963 | 0.84463913 | 0.99165628 | 264.206718 | 321.180756 | 411.180694 | 312.666399 | Non-sig. |
| PF3D7_1437200 | N/A | 11718.127 | -0.092780646 | 0.46656788 | -0.19885777 | 0.84237401 | 0.99128968 | 8582.53007 | 10179.2554 | 15606.7627 | 12503.96 | Non-sig. |
| PF3D7_1356500 | N/A | 13.9906214 | -0.092818699 | 0.99875651 | -0.09293426 | 0.92595579 | 0.99619209 | 16.6252207 | 17.2294221 | 12.293631 | 9.81421189 | Non-sig. |
| PF3D7_0527400 | N/A | 405.894369 | -0.092952264 | 0.51337636 | -0.18106066 | 0.85631996 | 0.99350189 | 558.981024 | 335.837031 | 279.243557 | 449.515863 | Non-sig. |
| PF3D7_0907000 | N/A | 454.143621 | -0.094408966 | 0.46189805 | -0.20439351 | 0.838046 | 0.99128968 | 370.277511 | 462.563569 | 567.499568 | 416.233836 | Non-sig. |
| PF3D7_0302500 | CLAG3.1 | 71642.4641 | -0.094439721 | 0.54831616 | -0.17223589 | 0.86325208 | 0.99402081 | 39342.4812 | 56478.4347 | 108630.288 | 82118.6519 | Non-sig. |
| PF3D7_1126000 | ThrRS | 4205.56898 | -0.094725437 | 0.46837573 | -0.20224241 | 0.83972722 | 0.99128968 | 3036.56671 | 4320.15727 | 5650.28933 | 3815.26262 | Non-sig. |
| PF3D7_0619400 | N/A | 9723.12433 | -0.095341801 | 0.57174022 | -0.1667572 | 0.86756109 | 0.99402081 | 4680.5797 | 8492.87555 | 15407.6291 | 10311.413 | Non-sig. |
| PF3D7_1453800 | GluPho | 3256.09643 | -0.095779247 | 0.5919811 | -0.16179443 | 0.87146774 | 0.99402081 | 1431.53638 | 3176.46254 | 5296.36869 | 3120.01811 | Non-sig. |
| PF3D7_0607900 | N/A | 487.329286 | -0.096037194 | 0.43530651 | -0.22061971 | 0.82538856 | 0.99096399 | 508.548342 | 422.724318 | 498.548434 | 519.49605 | Non-sig. |
| PF3D7_1107800 | ApiAP2 | 24206.0977 | -0.096175211 | 0.50511515 | -0.19040255 | 0.8489937 | 0.99165628 | 15239.7108 | 20921.5151 | 34785.2238 | 25877.9411 | Non-sig. |
| PF3D7_1253400 | ACS3 | 2080.38029 | -0.096319678 | 0.54859034 | -0.17557669 | 0.86062652 | 0.99402081 | 1127.38611 | 2252.17494 | 3171.85362 | 1770.10648 | Non-sig. |
| PF3D7_0521500 | N/A | 336.130824 | -0.096470476 | 0.45070853 | -0.21404182 | 0.83051444 | 0.99110542 | 337.509011 | 364.930373 | 357.192313 | 284.891602 | Non-sig. |
| PF3D7_0520800 | N/A | 3164.80621 | -0.09692633 | 0.42685974 | -0.22706834 | 0.82037061 | 0.99096399 | 3179.81317 | 2536.69321 | 3362.30665 | 3580.4118 | Non-sig. |
| PF3D7_0707420 | N/A | 5.05680396 | -0.096932321 | 1.55180091 | -0.06246441 | 0.95019301 | 0.99639249 | 3.07885058 | 6.4987729 | 7.34075299 | 3.30883937 | Non-sig. |
| PF3D7_0521700 | DDX1 | 135.640355 | -0.097347236 | 0.5711892 | -0.17042906 | 0.86467272 | 0.99402081 | 86.3255864 | 130.899372 | 193.829685 | 131.506774 | Non-sig. |
| PF3D7_1142900 | N/A | 881.120554 | -0.098538159 | 0.47487735 | -0.20750234 | 0.83561757 | 0.99128968 | 1177.22753 | 896.081045 | 645.474646 | 805.698994 | Non-sig. |
| PF3D7_1122400 | N/A | 1004.25822 | -0.098938897 | 0.48421683 | -0.20432767 | 0.83809745 | 0.99128968 | 1364.74214 | 1059.98796 | 712.933381 | 879.369393 | Non-sig. |
| PF3D7_1218100 | N/A | 136.252393 | -0.098994617 | 0.71138413 | -0.13915775 | 0.88932549 | 0.99457954 | 236.188351 | 131.005266 | 45.97027 | 131.845686 | Non-sig. |
| PF3D7_0804500 | N/A | 16044.8947 | -0.099364371 | 0.42754478 | -0.23240693 | 0.81622197 | 0.99096399 | 15764.9603 | 12682.4451 | 17429.4169 | 18302.7564 | Non-sig. |
| PF3D7_0412100 | mtRPS12 | 415.621262 | -0.099568456 | 0.44977178 | -0.2213755 | 0.82480006 | 0.99096399 | 375.315641 | 439.004514 | 484.457583 | 363.70731 | Non-sig. |
| PF3D7_0925600 | N/A | 461.748484 | -0.100561307 | 0.43742239 | -0.2298952 | 0.81817321 | 0.99096399 | 454.813039 | 488.500248 | 500.802203 | 402.878446 | Non-sig. |
| PF3D7_1437800 | TRAPPC5 | 396.190989 | -0.10121117 | 0.52759524 | -0.19183488 | 0.84787155 | 0.99165628 | 248.274087 | 386.337077 | 571.562817 | 378.589973 | Non-sig. |
| PF3D7_0409700 | RF2 | 470.431681 | -0.102043597 | 0.4863443 | -0.2098176 | 0.83381003 | 0.9911446 | 625.691904 | 494.407306 | 348.708973 | 412.91854 | Non-sig. |
| PF3D7_0718100 | EST | 2507.48031 | -0.102105519 | 0.41478418 | -0.24616542 | 0.80555419 | 0.99028237 | 2737.75562 | 2183.67202 | 2454.66905 | 2653.82452 | Non-sig. |
| PF3D7_1305000 | N/A | 3763.71188 | -0.102232799 | 0.44679991 | -0.22881114 | 0.8190157 | 0.99096399 | 3193.74084 | 3024.94589 | 4600.06537 | 4236.09541 | Non-sig. |
| PF3D7_1035100 | N/A | 3398.73641 | -0.102505191 | 0.46023701 | -0.22272262 | 0.8237514 | 0.99096399 | 4510.87806 | 3344.42876 | 2528.28861 | 3211.35022 | Non-sig. |
| PF3D7_0703100 | CIA2 | 102.728598 | -0.102549049 | 0.52683554 | -0.19465097 | 0.8456662 | 0.99165628 | 94.4573538 | 96.6483932 | 118.205298 | 101.603345 | Non-sig. |
| PF3D7_0602000 | N/A | 622.496409 | -0.102580211 | 0.45552163 | -0.22519284 | 0.82182926 | 0.99096399 | 581.822953 | 481.588343 | 707.29919 | 719.275149 | Non-sig. |
| PF3D7_1250200 | N/A | 4848.90746 | -0.102590592 | 0.46294075 | -0.22160631 | 0.82462037 | 0.99096399 | 6232.12808 | 3876.73123 | 3810.6122 | 5476.15831 | Non-sig. |
| PF3D7_0720700 | PX1 | 16452.5857 | -0.102810982 | 0.42960601 | -0.23931458 | 0.81086166 | 0.9905328 | 19809.5126 | 14122.2096 | 14267.8438 | 17610.7768 | Non-sig. |
| PF3D7_0709400 | N/A | 1523.00396 | -0.103369013 | 0.43007874 | -0.24034904 | 0.81005968 | 0.99028237 | 1667.58805 | 1235.8074 | 1487.56509 | 1701.05529 | Non-sig. |
| PF3D7_1364500 | RRP45 | 430.923729 | -0.103535355 | 0.48715998 | -0.21252845 | 0.83169479 | 0.99110542 | 340.710491 | 341.303942 | 551.813938 | 489.866542 | Non-sig. |
| PF3D7_0813200 | N/A | 1875.00144 | -0.1036696 | 0.4545202 | -0.22808579 | 0.81957955 | 0.99096399 | 2308.27755 | 2131.52177 | 1576.61953 | 1483.58691 | Non-sig. |
| PF3D7_0925700 | HDAC1 | 1619.22399 | -0.104245058 | 0.57341884 | -0.18179566 | 0.85574309 | 0.99321133 | 786.643748 | 1458.00864 | 2568.37942 | 1663.86414 | Non-sig. |
| PF3D7_1328700 | N/A | 25.4155033 | -0.104376033 | 0.78027078 | -0.13376899 | 0.89358525 | 0.99457954 | 29.2028778 | 26.1838102 | 23.5073767 | 22.7679486 | Non-sig. |

| | | | | | | | | | | | | |
|---|---|---|---|---|---|---|---|---|---|---|---|---|
| PF3D7_1007400 | N/A | 1904.7518 | -0.104660467 | 0.46127651 | -0.22689312 | 0.82050685 | 0.99096399 | 2518.51051 | 1894.79133 | 1429.41439 | 1776.29098 | Non-sig. |
| PF3D7_1401300 | EH2 | 97.0382255 | -0.104877941 | 0.56373328 | -0.18604178 | 0.852412 | 0.99253574 | 73.6429309 | 89.0713732 | 127.288396 | 98.1502019 | Non-sig. |
| PF3D7_1008000 | HDA2 | 9061.81535 | -0.105087262 | 0.43542678 | -0.24134313 | 0.80928919 | 0.99028237 | 9405.14125 | 6867.49999 | 9378.27363 | 10596.3465 | Non-sig. |
| PF3D7_1130500 | COG2 | 2271.87517 | -0.105951075 | 0.40589094 | -0.26103336 | 0.79406678 | 0.99028237 | 2413.98886 | 2183.16861 | 2296.56893 | 2193.77429 | Non-sig. |
| PF3D7_0205200 | N/A | 43.3262494 | -0.106012876 | 0.71948203 | -0.14734611 | 0.88285884 | 0.99457954 | 61.3121422 | 51.6866467 | 28.6857976 | 31.6204113 | Non-sig. |
| PF3D7_0511500 | N/A | 27838.7738 | -0.106071803 | 0.44229548 | -0.23982113 | 0.81046892 | 0.99028237 | 30431.5139 | 20575.8403 | 27292.0012 | 33055.74 | Non-sig. |
| PF3D7_1024700 | N/A | 759.35962 | -0.106086478 | 0.42919489 | -0.24717554 | 0.80477238 | 0.99028237 | 773.045306 | 821.020389 | 801.468988 | 641.903798 | Non-sig. |
| PF3D7_1407400 | N/A | 461.27923 | -0.106247118 | 0.44217883 | -0.24028088 | 0.81011252 | 0.99028237 | 516.455192 | 492.601535 | 440.156957 | 395.903237 | Non-sig. |
| PF3D7_0414600 | APH | 1801.2229 | -0.106538901 | 0.5188753 | -0.2053266 | 0.83731696 | 0.99128968 | 2642.24052 | 1878.9483 | 1093.51807 | 1590.18472 | Non-sig. |
| PF3D7_1354200 | SAC1 | 3484.70943 | -0.106782005 | 0.49668086 | -0.21499118 | 0.8297742 | 0.99110542 | 2287.25302 | 3047.98446 | 4939.6273 | 3663.97293 | Non-sig. |
| PF3D7_1200900 | N/A | 887.795649 | -0.10680352 | 0.53350438 | -0.20019239 | 0.84133012 | 0.99128968 | 514.597411 | 868.106926 | 1326.33013 | 842.148128 | Non-sig. |
| PF3D7_0626400 | N/A | 1953.48841 | -0.107258497 | 0.49711216 | -0.21576317 | 0.82917236 | 0.99110542 | 2682.40414 | 1518.0513 | 1370.04218 | 2243.45602 | Non-sig. |
| PF3D7_0928300 | N/A | 114.647122 | -0.107397608 | 0.50915206 | -0.21093425 | 0.83293858 | 0.99110542 | 122.913578 | 110.57864 | 114.941421 | 110.154851 | Non-sig. |
| PF3D7_1308000 | N/A | 9137.84891 | -0.10747896 | 0.44978781 | -0.23895481 | 0.81114062 | 0.9905328 | 11918.0899 | 9039.53873 | 7038.35637 | 8555.41067 | Non-sig. |
| PF3D7_0223100 | RIF | 9.47230064 | -0.107501086 | 1.25762643 | -0.08547935 | 0.93188031 | 0.99619209 | 3.14097315 | 9.67141032 | 16.4116321 | 8.66518704 | Non-sig. |
| PF3D7_0305500 | N/A | 18281.0997 | -0.107925713 | 0.48917349 | -0.22062871 | 0.82538155 | 0.99096399 | 12610.7028 | 14913.7825 | 25318.1315 | 20281.7818 | Non-sig. |
| PF3D7_1138400 | GCalpha | 7133.69829 | -0.108216059 | 0.57409245 | -0.18849936 | 0.85048522 | 0.99165628 | 3839.24302 | 4879.69008 | 10962.6653 | 8853.19472 | Non-sig. |
| PF3D7_1411100 | N/A | 510.959647 | -0.108341019 | 0.53508283 | -0.20247523 | 0.83954522 | 0.99128968 | 304.297631 | 493.531012 | 755.630048 | 490.379899 | Non-sig. |
| PF3D7_1324000 | N/A | 656.417951 | -0.10855765 | 0.54953229 | -0.19754553 | 0.84340065 | 0.99164151 | 997.844671 | 597.480746 | 364.719355 | 665.627032 | Non-sig. |
| PF3D7_0205800 | PH1 | 689.843112 | -0.108610513 | 0.45485956 | -0.23877813 | 0.81127763 | 0.9905328 | 611.829716 | 778.248643 | 819.615668 | 549.678423 | Non-sig. |
| PF3D7_0616900 | N/A | 1343.97079 | -0.10895767 | 0.43251093 | -0.25191888 | 0.80110376 | 0.99028237 | 1632.19448 | 1250.73456 | 1157.41244 | 1335.54168 | Non-sig. |
| PF3D7_1345500 | UBC | 476.148332 | -0.109041596 | 0.49023866 | -0.22242554 | 0.82398263 | 0.99096399 | 344.110758 | 430.300043 | 643.863389 | 486.319136 | Non-sig. |
| PF3D7_1138900 | N/A | 165.930379 | -0.109090873 | 0.47868229 | -0.22789829 | 0.81972531 | 0.99096399 | 165.310186 | 162.007 | 179.051413 | 157.352919 | Non-sig. |
| PF3D7_0109700 | RRP36 | 128.76272 | -0.109100362 | 0.65934671 | -0.16546736 | 0.86857614 | 0.99402081 | 199.393042 | 166.453975 | 68.1312007 | 81.0726636 | Non-sig. |
| PF3D7_0310300 | N/A | 1355.41469 | -0.109452917 | 0.44214757 | -0.24754838 | 0.80448385 | 0.99028237 | 1697.51779 | 1256.0993 | 1116.33956 | 1351.70213 | Non-sig. |
| PF3D7_0826600 | VAMP7 | 547.954707 | -0.109925849 | 0.52679819 | -0.20866786 | 0.83470753 | 0.99128968 | 337.5182 | 558.124534 | 799.786146 | 496.389947 | Non-sig. |
| PF3D7_0110400 | RPB9 | 820.479465 | -0.110149075 | 0.42104595 | -0.2616082 | 0.79362352 | 0.99028237 | 784.939323 | 792.54692 | 918.531749 | 785.899868 | Non-sig. |
| PF3D7_1338700 | N/A | 4186.37206 | -0.111157187 | 0.40653349 | -0.27342688 | 0.78452509 | 0.99028237 | 4436.91224 | 3755.6637 | 4258.25465 | 4294.65765 | Non-sig. |
| PF3D7_0811500 | CARM1 | 579.681581 | -0.111300587 | 0.43227611 | -0.25747569 | 0.79681158 | 0.99028237 | 547.672833 | 593.265074 | 656.279156 | 521.509261 | Non-sig. |
| PF3D7_0905800 | N/A | 785.270815 | -0.111777806 | 0.44735743 | -0.24986241 | 0.80269375 | 0.99028237 | 939.876427 | 650.044864 | 691.650512 | 859.511456 | Non-sig. |
| PF3D7_0828200 | N/A | 2314.59801 | -0.111779004 | 0.41735529 | -0.26782697 | 0.78883251 | 0.99028237 | 2694.22481 | 2278.3792 | 2114.37817 | 2171.40986 | Non-sig. |
| PF3D7_1411200 | ROM8 | 5187.60186 | -0.112389791 | 0.520356 | -0.21598635 | 0.82899839 | 0.99110542 | 7528.38665 | 4042.19793 | 3251.08357 | 5928.7393 | Non-sig. |
| PF3D7_1428600 | RF1 | 481.669599 | -0.112422974 | 0.44317802 | -0.25367453 | 0.79974702 | 0.99028237 | 500.902475 | 530.40949 | 499.958891 | 395.40754 | Non-sig. |
| PF3D7_1015100 | N/A | 1042.9483 | -0.112633352 | 0.42313515 | -0.26618765 | 0.79009469 | 0.99028237 | 1174.61781 | 1082.94034 | 992.773565 | 921.461496 | Non-sig. |
| PF3D7_0321600 | DDX42 | 4162.31754 | -0.112988724 | 0.46327628 | -0.24389059 | 0.80731556 | 0.99028237 | 5501.07931 | 4448.56875 | 3149.6726 | 3549.94952 | Non-sig. |
| PF3D7_1249000 | N/A | 44.7988281 | -0.113471632 | 0.64785738 | -0.17514909 | 0.86096248 | 0.99402081 | 46.3030597 | 37.5572019 | 46.8116192 | 48.5234317 | Non-sig. |
| PF3D7_1020000 | RBM34 | 1555.94052 | -0.113658026 | 0.50773116 | -0.22385474 | 0.82287033 | 0.99096399 | 2192.66875 | 1765.52002 | 1042.05263 | 1223.52067 | Non-sig. |
| PF3D7_0807000 | N/A | 525.893466 | -0.113721516 | 0.46247525 | -0.24589752 | 0.80576157 | 0.99028237 | 420.256135 | 506.359645 | 672.712967 | 504.245116 | Non-sig. |
| PF3D7_1336400 | N/A | 586.576695 | -0.113962606 | 0.46778369 | -0.24362245 | 0.80752324 | 0.99028237 | 754.008455 | 620.107343 | 465.709775 | 506.481205 | Non-sig. |
| PF3D7_0306100 | N/A | 1356.48576 | -0.113999959 | 0.41316073 | -0.27592158 | 0.7826083 | 0.99028237 | 1504.44332 | 1292.88453 | 1315.75167 | 1312.86351 | Non-sig. |
| PF3D7_0528500 | CPalpha | 457.918277 | -0.114105865 | 0.43363327 | -0.26313909 | 0.79244337 | 0.99028237 | 479.987137 | 473.479149 | 472.061606 | 406.145217 | Non-sig. |
| PF3D7_0608310 | SAM50 | 482.944394 | -0.114247327 | 0.51200877 | -0.22313549 | 0.82343006 | 0.99096399 | 317.381695 | 459.443998 | 686.409176 | 468.542706 | Non-sig. |
| PF3D7_1005000 | MRSapi | 495.632899 | -0.114355393 | 0.42965998 | -0.26615323 | 0.7901212 | 0.99028237 | 526.286325 | 449.585674 | 504.270774 | 502.38882 | Non-sig. |
| PF3D7_1014500 | N/A | 253.953595 | -0.114474936 | 0.58193187 | -0.19671536 | 0.84405029 | 0.99165628 | 135.900276 | 230.637734 | 391.813932 | 257.462437 | Non-sig. |
| PF3D7_0923700 | N/A | 36.6244065 | -0.114904918 | 0.78647604 | -0.14610098 | 0.88384167 | 0.99457954 | 57.4856055 | 40.8544137 | 18.8680471 | 29.2895597 | Non-sig. |
| PF3D7_0614700 | N/A | 2532.38674 | -0.115220432 | 0.4751576 | -0.24248887 | 0.80840138 | 0.99028237 | 3474.55931 | 2383.52871 | 1792.68309 | 2478.77584 | Non-sig. |
| PF3D7_0528200 | EIF3E | 2612.55627 | -0.115733144 | 0.45897615 | -0.25215503 | 0.80092124 | 0.99028237 | 1976.92533 | 2620.57835 | 3457.35951 | 2395.3619 | Non-sig. |
| PF3D7_0809900 | JmjC1 | 950.650372 | -0.116178599 | 0.44166007 | -0.26304981 | 0.79251219 | 0.99028237 | 934.533526 | 742.312126 | 1043.21134 | 1082.5445 | Non-sig. |
| PF3D7_1241900 | N/A | 3818.59945 | -0.116332268 | 0.50089456 | -0.23224901 | 0.81634461 | 0.99096399 | 5500.13436 | 3854.7751 | 2445.17191 | 3474.31642 | Non-sig. |
| PF3D7_0408900 | KAE1 | 1762.14685 | -0.116349177 | 0.45109002 | -0.25792895 | 0.79646174 | 0.99028237 | 2232.82261 | 1888.06785 | 1433.76126 | 1493.93566 | Non-sig. |
| PF3D7_1013300 | N/A | 389.968746 | -0.11635482 | 0.5498091 | -0.21162767 | 0.83239752 | 0.99110542 | 233.67721 | 317.604738 | 577.368024 | 431.225011 | Non-sig. |
| PF3D7_0922200 | SAMS | 23589.0415 | -0.116991238 | 0.42392411 | -0.27597213 | 0.78256947 | 0.99028237 | 20666.7832 | 20895.1748 | 28422.9541 | 24371.2538 | Non-sig. |
| PF3D7_1462100 | N/A | 563.033645 | -0.117003426 | 0.47039101 | -0.24873653 | 0.80356459 | 0.99028237 | 444.515893 | 483.820371 | 726.934098 | 596.864219 | Non-sig. |
| PF3D7_1453400 | N/A | 918.113632 | -0.117110242 | 0.51737205 | -0.22635595 | 0.82092458 | 0.99096399 | 594.394022 | 740.107174 | 1315.99032 | 1021.96301 | Non-sig. |

| | | | | | | | | | | | | |
|---|---|---|---|---|---|---|---|---|---|---|---|---|
| PF3D7_1125300 | mtRNAP | 909.51858 | -0.11726168 | 0.42803227 | -0.27395523 | 0.78411903 | 0.99028237 | 986.123308 | 762.701386 | 906.848797 | 982.400829 | Non-sig. |
| PF3D7_1435800 | ERA | 135.155349 | -0.117418584 | 0.51710668 | -0.22706839 | 0.82037056 | 0.99096399 | 117.757112 | 113.773236 | 163.404479 | 145.68657 | Non-sig. |
| PF3D7_1402100 | N/A | 193.420763 | -0.117523781 | 0.47796286 | -0.24588476 | 0.80577144 | 0.99028237 | 194.760985 | 162.014319 | 207.792373 | 209.115374 | Non-sig. |
| PF3D7_0318300 | N/A | 1318.38292 | -0.117608944 | 0.47595352 | -0.24710174 | 0.80482949 | 0.99028237 | 1013.25481 | 1043.82872 | 1730.65642 | 1485.79174 | Non-sig. |
| PF3D7_1438800 | N/A | 401.90292 | -0.117666034 | 0.43408458 | -0.27106707 | 0.78633945 | 0.99028237 | 431.689627 | 388.390713 | 404.917034 | 382.614305 | Non-sig. |
| PF3D7_1144100 | N/A | 253.007651 | -0.117966923 | 0.52211997 | -0.22593835 | 0.82124937 | 0.99096399 | 190.645728 | 294.221428 | 335.839612 | 191.323836 | Non-sig. |
| PF3D7_1326300 | N/A | 1516.7775 | -0.118126905 | 0.4182848 | -0.28240784 | 0.77763081 | 0.99028237 | 1451.89552 | 1566.77406 | 1705.68049 | 1342.75993 | Non-sig. |
| PF3D7_1137900 | N/A | 18.0253117 | -0.118263633 | 0.97685535 | -0.12106566 | 0.90363903 | 0.99457954 | 29.4540178 | 16.0977209 | 8.2120006 | 18.3375074 | Non-sig. |
| PF3D7_1470700 | N/A | 478.326141 | -0.118298287 | 0.51482433 | -0.22978379 | 0.81825978 | 0.99096399 | 672.575428 | 522.553502 | 323.58588 | 394.589755 | Non-sig. |
| PF3D7_1310900 | N/A | 252.666758 | -0.118590591 | 0.46618364 | -0.254386 | 0.79919737 | 0.99028237 | 228.80071 | 257.727782 | 297.155904 | 226.982635 | Non-sig. |
| PF3D7_1119300 | U2AF1 | 1504.0785 | -0.118920032 | 0.46276407 | -0.25697767 | 0.79719601 | 0.99028237 | 1995.14031 | 1378.24077 | 1137.22012 | 1505.7128 | Non-sig. |
| PF3D7_1406700 | VPS29 | 848.932279 | -0.119123762 | 0.4534415 | -0.26271032 | 0.79277386 | 0.99028237 | 692.836468 | 879.072788 | 1074.84627 | 748.973592 | Non-sig. |
| PF3D7_1338800 | N/A | 192.535306 | -0.119294731 | 0.4849067 | -0.24601584 | 0.80566997 | 0.99028237 | 230.326516 | 198.5497 | 170.803641 | 170.461367 | Non-sig. |
| PF3D7_0616800 | MQO | 1010.11013 | -0.119416904 | 0.42885369 | -0.27845605 | 0.7806623 | 0.99028237 | 901.193293 | 960.274867 | 1202.41005 | 976.562325 | Non-sig. |
| PF3D7_1135500 | N/A | 181.032904 | -0.119422173 | 0.49564069 | -0.24094505 | 0.8095977 | 0.99028237 | 206.441954 | 204.122665 | 170.695028 | 142.871969 | Non-sig. |
| PF3D7_1447800 | N/A | 1117.13447 | -0.119537475 | 0.5255754 | -0.22744115 | 0.82008072 | 0.99096399 | 1642.01543 | 928.879429 | 685.107546 | 1212.53548 | Non-sig. |
| PF3D7_1468600 | N/A | 146.879591 | -0.119896994 | 0.54386346 | -0.22045422 | 0.82551743 | 0.99096399 | 120.136698 | 106.190977 | 185.649746 | 175.540941 | Non-sig. |
| PF3D7_1442800 | N/A | 30.3830206 | -0.120238868 | 0.73094772 | -0.16449722 | 0.86933974 | 0.99402081 | 34.6105368 | 30.0741941 | 28.7269648 | 28.1203866 | Non-sig. |
| PF3D7_1350900 | ApiAP2 | 133.02667 | -0.120517095 | 0.50611464 | -0.23812213 | 0.81178637 | 0.99055243 | 147.439277 | 111.57482 | 129.77565 | 143.316932 | Non-sig. |
| PF3D7_0818200 | 14-3-3I | 20816.8938 | -0.120727705 | 0.47323574 | -0.25511113 | 0.79863727 | 0.99028237 | 14674.6975 | 19198.9618 | 28699.7639 | 20694.152 | Non-sig. |
| PF3D7_1416400 | N/A | 127.619075 | -0.120863458 | 0.74682585 | -0.1618362 | 0.87143485 | 0.99402081 | 229.27054 | 124.496678 | 36.9525906 | 119.756493 | Non-sig. |
| PF3D7_1224400 | N/A | 389.96002 | -0.12096385 | 0.43923579 | -0.27539616 | 0.78301189 | 0.99028237 | 408.512938 | 404.487833 | 404.093481 | 342.745827 | Non-sig. |
| PF3D7_0606800 | VFT1 | 1963.68998 | -0.121609581 | 0.45340348 | -0.26821493 | 0.78853388 | 0.99028237 | 1530.18456 | 1956.16531 | 2562.34332 | 1806.06673 | Non-sig. |
| PF3D7_0212800 | MATE | 370.216345 | -0.121748069 | 0.48373282 | -0.25168453 | 0.80128491 | 0.99028237 | 283.454068 | 335.743909 | 487.938769 | 373.728635 | Non-sig. |
| PF3D7_1310800 | N/A | 767.5525 | -0.122007438 | 0.50480277 | -0.24169328 | 0.80901784 | 0.99028237 | 504.677891 | 715.491012 | 1094.96449 | 755.076614 | Non-sig. |
| PF3D7_1442500 | N/A | 324.397336 | -0.122091793 | 0.46054851 | -0.26510083 | 0.79093179 | 0.99028237 | 288.421777 | 336.915462 | 387.651854 | 284.600252 | Non-sig. |
| PF3D7_0420200 | ACPS | 145.426953 | -0.122665093 | 0.58185923 | -0.21081575 | 0.83303104 | 0.99110542 | 96.8231246 | 106.230871 | 206.138219 | 172.515599 | Non-sig. |
| PF3D7_1019700 | N/A | 3454.70158 | -0.122817227 | 0.6302071 | -0.19488392 | 0.84548383 | 0.99165628 | 5901.04729 | 3546.91642 | 1302.65633 | 3068.18627 | Non-sig. |
| PF3D7_1135900 | N/A | 6206.50746 | -0.123617984 | 0.41155186 | -0.30037037 | 0.76389466 | 0.99028237 | 7035.7707 | 6357.66354 | 5908.85162 | 5523.74398 | Non-sig. |
| PF3D7_1122700 | N/A | 753.815255 | -0.123670965 | 0.42562676 | -0.29056201 | 0.77138632 | 0.99028237 | 706.78692 | 712.921263 | 865.29544 | 730.257396 | Non-sig. |
| PF3D7_1429700 | N/A | 1360.87109 | -0.123774345 | 0.42672275 | -0.29005799 | 0.77177187 | 0.99028237 | 1362.09223 | 1489.94697 | 1476.28823 | 1115.15694 | Non-sig. |
| PF3D7_1333800 | N/A | 1348.82062 | -0.12413362 | 0.43565509 | -0.28493554 | 0.77569352 | 0.99028237 | 1628.66095 | 1419.49362 | 1185.16054 | 1161.96738 | Non-sig. |
| PF3D7_1234900 | N/A | 4375.012 | -0.124337115 | 0.44279969 | -0.28079766 | 0.7788656 | 0.99028237 | 3750.16325 | 4824.13299 | 5376.48612 | 3549.26563 | Non-sig. |
| PF3D7_1437300 | N/A | 1881.2505 | -0.124676654 | 0.432229 | -0.28845046 | 0.77300195 | 0.99028237 | 2238.54318 | 2012.47133 | 1686.61107 | 1587.3764 | Non-sig. |
| PF3D7_1363100 | N/A | 1181.64753 | -0.124736347 | 0.41124776 | -0.30331192 | 0.76165217 | 0.99028237 | 1237.45875 | 1114.66979 | 1227.94429 | 1146.51728 | Non-sig. |
| PF3D7_1013900 | N/A | 1124.54986 | -0.124990225 | 0.4214227 | -0.2965911 | 0.7667787 | 0.99028237 | 1263.97487 | 989.107507 | 1082.59023 | 1162.52683 | Non-sig. |
| PF3D7_1143300 | RPC40 | 1423.25674 | -0.125417284 | 0.42824771 | -0.29286154 | 0.76962799 | 0.99028237 | 1712.91727 | 1411.97533 | 1257.44072 | 1310.69363 | Non-sig. |
| PF3D7_1231900 | N/A | 426.554741 | -0.125947332 | 0.44537395 | -0.28279007 | 0.77733776 | 0.99028237 | 383.373121 | 403.376704 | 506.789278 | 412.67986 | Non-sig. |
| PF3D7_0929400 | RhopH2 | 47953.4368 | -0.126138746 | 0.59901343 | -0.21057749 | 0.83321698 | 0.99110542 | 21328.1154 | 37734.9951 | 78768.5095 | 53982.1269 | Non-sig. |
| PF3D7_1465300 | N/A | 414.296299 | -0.126159985 | 0.43441208 | -0.29041546 | 0.77149842 | 0.99028237 | 440.181721 | 416.267845 | 424.640619 | 376.095013 | Non-sig. |
| PF3D7_0710200 | N/A | 7246.66666 | -0.126334661 | 0.71142751 | -0.17757911 | 0.85905353 | 0.99402081 | 13297.5284 | 7597.65893 | 1830.34401 | 6261.13532 | Non-sig. |
| PF3D7_0532200 | N/A | 132.476024 | -0.126900983 | 0.53528745 | -0.23707072 | 0.81260192 | 0.99055243 | 101.602411 | 121.41489 | 174.784044 | 132.102752 | Non-sig. |
| PF3D7_0804300 | N/A | 274.038197 | -0.127170532 | 0.46159086 | -0.27550487 | 0.78292838 | 0.99028237 | 316.899981 | 238.189305 | 255.431187 | 285.632316 | Non-sig. |
| PF3D7_1003300 | ARD1 | 1940.17579 | -0.127216214 | 0.44117939 | -0.28835485 | 0.77307513 | 0.99028237 | 2357.97489 | 2118.56114 | 1693.54865 | 1590.61847 | Non-sig. |
| PF3D7_1027700 | CEN3 | 1821.8336 | -0.127457008 | 0.43691819 | -0.29171825 | 0.77050206 | 0.99028237 | 1676.44943 | 2022.96124 | 2127.96516 | 1459.95856 | Non-sig. |
| PF3D7_1130400 | RPT5 | 5667.9626 | -0.127498618 | 0.43556616 | -0.2927193 | 0.76973672 | 0.99028237 | 4672.20591 | 5329.62794 | 7164.05044 | 5505.96611 | Non-sig. |
| PF3D7_1335600 | N/A | 347.492002 | -0.127648016 | 0.43896308 | -0.29079442 | 0.77120855 | 0.99028237 | 367.755218 | 340.307648 | 357.971552 | 323.93359 | Non-sig. |
| PF3D7_0526700 | N/A | 1088.07614 | -0.128258219 | 0.42241949 | -0.30362761 | 0.76141161 | 0.99028237 | 1254.57044 | 988.251395 | 1018.38336 | 1091.09936 | Non-sig. |
| PF3D7_0907100 | N/A | 726.968789 | -0.128780652 | 0.45054862 | -0.28583075 | 0.77500775 | 0.99028237 | 918.580714 | 715.527161 | 600.445617 | 673.321663 | Non-sig. |
| PF3D7_1447400 | N/A | 107.166334 | -0.129006252 | 0.53159156 | -0.24267927 | 0.80825387 | 0.99028237 | 93.2857057 | 97.7581237 | 130.488186 | 107.133321 | Non-sig. |
| PF3D7_0916500 | UFD1 | 976.563376 | -0.129145979 | 0.43211299 | -0.29887086 | 0.76503858 | 0.99028237 | 1010.03145 | 1078.80909 | 1030.45007 | 786.962897 | Non-sig. |
| PF3D7_1030800 | N/A | 1131.23076 | -0.129289039 | 0.49602255 | -0.26065154 | 0.79436124 | 0.99028237 | 1455.22661 | 1386.21151 | 908.767437 | 774.717496 | Non-sig. |
| PF3D7_1233900 | SENP1 | 2545.65984 | -0.129371635 | 0.44080822 | -0.29348735 | 0.76914967 | 0.99028237 | 3166.01753 | 2159.56162 | 2153.6526 | 2703.40763 | Non-sig. |
| PF3D7_0923800 | TRXR | 3712.73743 | -0.129577762 | 0.48393801 | -0.26775694 | 0.78888642 | 0.99028237 | 2604.52371 | 3954.0869 | 5153.97528 | 3138.36384 | Non-sig. |

| | | | | | | | | | | | | |
|---|---|---|---|---|---|---|---|---|---|---|---|---|
| PF3D7_0802300 | PWP2 | 967.86965 | -0.12966823 | 0.49270615 | -0.26317558 | 0.79241525 | 0.99028237 | 660.466477 | 915.069912 | 1361.87661 | 934.065606 | Non-sig. |
| PF3D7_1119100 | N/A | 490.685597 | -0.129778972 | 0.50629586 | -0.25633031 | 0.7976958 | 0.99028237 | 694.132378 | 476.722456 | 331.664839 | 460.222714 | Non-sig. |
| PF3D7_0510300 | SIP | 46.9539804 | -0.129821664 | 0.6752191 | -0.19226598 | 0.84753386 | 0.99165628 | 63.821872 | 50.516172 | 34.4604072 | 39.0174701 | Non-sig. |
| PF3D7_1203700 | NAPL | 11443.2108 | -0.130175186 | 0.4005842 | -0.32496336 | 0.7452088 | 0.98998709 | 11764.3588 | 10908.7234 | 12153.8757 | 10945.8855 | Non-sig. |
| PF3D7_1333200 | UBA1 | 2350.13507 | -0.130287319 | 0.45075484 | -0.28904253 | 0.77254883 | 0.99028237 | 1860.28783 | 2103.35133 | 3051.7994 | 2385.10171 | Non-sig. |
| PF3D7_1211500 | mtRPS18 | 303.095942 | -0.130595502 | 0.46155944 | -0.28294406 | 0.77721972 | 0.99028237 | 347.120548 | 327.324591 | 286.596495 | 251.342133 | Non-sig. |
| PF3D7_0302700 | N/A | 355.121018 | -0.130847042 | 0.46435993 | -0.28177936 | 0.7781127 | 0.99028237 | 363.305895 | 406.199233 | 379.108701 | 271.870244 | Non-sig. |
| PF3D7_1452400 | N/A | 3261.25823 | -0.130867376 | 0.40477734 | -0.32330707 | 0.7464627 | 0.98998709 | 3526.83368 | 3053.06151 | 3291.36023 | 3173.77752 | Non-sig. |
| PF3D7_1121700 | GCN20 | 1198.40616 | -0.131678839 | 0.44559199 | -0.29551438 | 0.76760097 | 0.99028237 | 1387.47141 | 925.764552 | 1118.76774 | 1361.62096 | Non-sig. |
| PF3D7_0111600 | N/A | 51.8933082 | -0.131704393 | 0.63800867 | -0.20643041 | 0.83645472 | 0.99128968 | 41.9214273 | 52.640655 | 66.4737849 | 46.5373654 | Non-sig. |
| PF3D7_0506300 | N/A | 559.004542 | -0.131721768 | 0.50023212 | -0.26332129 | 0.79230295 | 0.99028237 | 778.512207 | 587.235111 | 390.806087 | 479.464765 | Non-sig. |
| PF3D7_1022200 | N/A | 693.822194 | -0.132053382 | 0.57419176 | -0.22998133 | 0.81810628 | 0.99096399 | 353.563531 | 594.487953 | 1097.18592 | 730.051378 | Non-sig. |
| PF3D7_1128300 | PFK11 | 195.056049 | -0.132200209 | 0.525919 | -0.2513699 | 0.80152813 | 0.99028237 | 264.468153 | 163.048217 | 143.751231 | 208.956594 | Non-sig. |
| PF3D7_0928700 | N/A | 470.342581 | -0.132604443 | 0.44928626 | -0.29514466 | 0.76788337 | 0.99028237 | 419.382949 | 409.758464 | 564.332607 | 487.896303 | Non-sig. |
| PF3D7_0320800 | DOZI | 920.742419 | -0.133030505 | 0.45931903 | -0.28962551 | 0.77210275 | 0.99028237 | 730.31531 | 813.384889 | 1195.7465 | 943.522974 | Non-sig. |
| PF3D7_0322100 | Prt1 | 1079.78167 | -0.133287286 | 0.41765954 | -0.31912903 | 0.74962867 | 0.99028237 | 1222.92043 | 1040.84739 | 1036.4403 | 1018.91856 | Non-sig. |
| PF3D7_1207200 | N/A | 528.980755 | -0.133465057 | 0.53943426 | -0.24741673 | 0.80458573 | 0.99028237 | 338.428669 | 394.681127 | 768.123079 | 614.690146 | Non-sig. |
| PF3D7_1201400 | N/A | 119.678484 | -0.133811922 | 0.67064398 | -0.19952751 | 0.84185013 | 0.99128968 | 199.007466 | 114.799096 | 51.7443738 | 113.163001 | Non-sig. |
| PF3D7_0918600 | RECQ1 | 663.665275 | -0.13387352 | 0.47992485 | -0.27894684 | 0.78028562 | 0.99028237 | 495.824067 | 576.178915 | 892.759571 | 689.898549 | Non-sig. |
| PF3D7_0519200 | N/A | 4901.49276 | -0.13443989 | 0.47633477 | -0.28223824 | 0.77776084 | 0.99028237 | 3506.84477 | 4236.64887 | 6752.2496 | 5110.22778 | Non-sig. |
| PF3D7_1344200 | GRP170 | 7539.76275 | -0.134506691 | 0.42719057 | -0.31486343 | 0.75286535 | 0.99028237 | 9291.4401 | 7522.08724 | 6490.76222 | 6854.76142 | Non-sig. |
| PF3D7_1239200 | ApiAP2 | 11843.9936 | -0.134512538 | 0.44656423 | -0.30121655 | 0.76324937 | 0.99028237 | 10294.1522 | 9188.39466 | 14497.1448 | 13396.2826 | Non-sig. |
| PF3D7_0814200 | ALBA1 | 18191.3466 | -0.134677016 | 0.49082491 | -0.27438912 | 0.7837856 | 0.99028237 | 25628.6977 | 19685.9352 | 12451.2988 | 14999.4545 | Non-sig. |
| PF3D7_1313300 | N/A | 93.9367226 | -0.134699627 | 0.53723527 | -0.25072745 | 0.80202484 | 0.99028237 | 108.923315 | 94.5559187 | 87.8007072 | 84.466949 | Non-sig. |
| PF3D7_1449500 | ApiAP2 | 1317.83834 | -0.134843318 | 0.50181766 | -0.26870979 | 0.78815302 | 0.99028237 | 1892.47792 | 1175.64453 | 866.623195 | 1336.60774 | Non-sig. |
| PF3D7_1440500 | N/A | 506.087473 | -0.134918536 | 0.43219763 | -0.31216862 | 0.75491238 | 0.99028237 | 561.875941 | 512.943284 | 497.669833 | 451.860835 | Non-sig. |
| PF3D7_0211000 | N/A | 197.686304 | -0.135902523 | 0.49099755 | -0.2767886 | 0.78194244 | 0.99028237 | 170.21045 | 206.474031 | 243.610936 | 170.4498 | Non-sig. |
| PF3D7_0927500 | N/A | 155.417177 | -0.136041882 | 0.50180037 | -0.27110758 | 0.78630829 | 0.99028237 | 190.16501 | 157.812235 | 135.46876 | 138.222704 | Non-sig. |
| PF3D7_1476100 | ETRAMP14.2 | 16.5242181 | -0.136351341 | 0.99774916 | -0.13665894 | 0.89130039 | 0.99457954 | 24.8805972 | 21.5511363 | 9.83053089 | 9.83460782 | Non-sig. |
| PF3D7_0905300 | N/A | 556.163036 | -0.136366556 | 0.48449522 | -0.2814611 | 0.77835676 | 0.99028237 | 542.042818 | 373.204826 | 622.74069 | 686.663808 | Non-sig. |
| PF3D7_1326900 | N/A | 537.493 | -0.136546399 | 0.53458694 | -0.25542412 | 0.79839554 | 0.99028237 | 791.019136 | 436.626249 | 335.125201 | 587.201416 | Non-sig. |
| PF3D7_1366300 | N/A | 4345.05672 | -0.136562705 | 0.43277197 | -0.31555349 | 0.75234145 | 0.99028237 | 3867.69432 | 3591.67131 | 5233.22236 | 4687.63891 | Non-sig. |
| PF3D7_1211200 | N/A | 1430.50203 | -0.136687679 | 0.46082819 | -0.2966131 | 0.76676191 | 0.99028237 | 1863.09291 | 1197.07635 | 1133.60505 | 1528.23381 | Non-sig. |
| PF3D7_1360100 | RRM1 | 665.882914 | -0.137192344 | 0.4293949 | -0.31950157 | 0.7493462 | 0.99028237 | 705.491766 | 704.400491 | 689.56712 | 564.072279 | Non-sig. |
| PF3D7_0919100 | N/A | 4194.66633 | -0.137316636 | 0.40397002 | -0.33991789 | 0.73391837 | 0.98909017 | 4345.0909 | 4156.77064 | 4443.18026 | 3833.62353 | Non-sig. |
| PF3D7_1479900 | N/A | 4.54846346 | -0.137622064 | 1.59492001 | -0.08628775 | 0.93123767 | 0.99619209 | 6.27693604 | 4.29937809 | 3.28187991 | 4.33565978 | Non-sig. |
| PF3D7_1331400 | N/A | 201.857122 | -0.137858516 | 0.46722897 | -0.29505558 | 0.76795142 | 0.99028237 | 207.889516 | 204.012538 | 215.083169 | 180.443266 | Non-sig. |
| PF3D7_0417200 | DHFR-TS | 1645.45837 | -0.137914846 | 0.48460882 | -0.28459005 | 0.77595824 | 0.99028237 | 1155.8729 | 1440.27854 | 2291.90086 | 1693.78116 | Non-sig. |
| PF3D7_1468900 | N/A | 451.203018 | -0.13819799 | 0.48065712 | -0.28751887 | 0.77371506 | 0.99028237 | 354.810588 | 375.521013 | 590.533223 | 483.947248 | Non-sig. |
| PF3D7_0624200 | N/A | 1395.35329 | -0.138312976 | 0.46177525 | -0.29952444 | 0.76453992 | 0.99028237 | 1826.30999 | 1480.83756 | 1098.33849 | 1175.92712 | Non-sig. |
| PF3D7_0318500 | N/A | 11668.5088 | -0.138467093 | 0.43526783 | -0.31811929 | 0.75039445 | 0.99028237 | 11726.1087 | 8773.65222 | 12729.9269 | 13444.3474 | Non-sig. |
| PF3D7_1119900 | SEC16 | 3388.15971 | -0.138543482 | 0.44587521 | -0.31072255 | 0.75601155 | 0.99028237 | 2757.20032 | 2936.76394 | 4343.9795 | 3514.69509 | Non-sig. |
| PF3D7_0921200 | N/A | 496.070991 | -0.13856748 | 0.46161976 | -0.30017667 | 0.7640424 | 0.99028237 | 411.083866 | 434.390871 | 628.4401 | 510.369128 | Non-sig. |
| PF3D7_0522500 | RPL17 | 509.038246 | -0.139515778 | 0.42715854 | -0.32661357 | 0.74396018 | 0.98998709 | 550.552016 | 488.385887 | 516.751 | 480.464081 | Non-sig. |
| PF3D7_0319200 | CCR4-4 | 683.838225 | -0.139710347 | 0.4377266 | -0.31917262 | 0.74959561 | 0.99028237 | 825.426778 | 624.43949 | 608.601367 | 676.885266 | Non-sig. |
| PF3D7_1017100 | RON12 | 8945.68249 | -0.139937497 | 0.42951432 | -0.32580403 | 0.74457263 | 0.98998709 | 10532.7307 | 9786.10131 | 8225.82633 | 7238.07156 | Non-sig. |
| PF3D7_0628700 | N/A | 393.262368 | -0.14000575 | 0.44376228 | -0.31549719 | 0.75238419 | 0.99028237 | 404.147204 | 417.407592 | 420.492008 | 331.002667 | Non-sig. |
| PF3D7_1455500 | N/A | 1739.19802 | -0.140157026 | 0.48539612 | -0.28874772 | 0.77277444 | 0.99028237 | 1227.09853 | 1484.73759 | 2419.80736 | 1825.14861 | Non-sig. |
| PF3D7_1330300 | N/A | 742.171595 | -0.140471623 | 0.47794959 | -0.29390468 | 0.76883074 | 0.99028237 | 992.886411 | 789.287477 | 563.94125 | 622.57124 | Non-sig. |
| PF3D7_1415500 | N/A | 456.950408 | -0.140568431 | 0.55655929 | -0.25256686 | 0.80060294 | 0.99028237 | 254.922474 | 422.667433 | 703.118716 | 447.09301 | Non-sig. |
| PF3D7_1005900 | N/A | 1475.59953 | -0.140572221 | 0.42172441 | -0.33332721 | 0.7388873 | 0.98954916 | 1709.60175 | 1298.50318 | 1385.39526 | 1508.89794 | Non-sig. |
| PF3D7_0109850 | N/A | 387.373278 | -0.140762895 | 0.53865717 | -0.26132186 | 0.79384431 | 0.99028237 | 236.570435 | 350.977695 | 575.612804 | 386.332178 | Non-sig. |
| PF3D7_1137200 | AEP | 3269.73668 | -0.14099862 | 0.54928201 | -0.25669623 | 0.79741328 | 0.99028237 | 5086.9492 | 3437.23439 | 1772.20288 | 2782.56024 | Non-sig. |
| PF3D7_0610800 | TK | 3181.50553 | -0.141656054 | 0.54167601 | -0.26151436 | 0.79369588 | 0.99028237 | 1766.0359 | 3269.51562 | 4908.74988 | 2781.7207 | Non-sig. |

| | | | | | | | | | | | | |
|---|---|---|---|---|---|---|---|---|---|---|---|---|
| PF3D7_0610200 | RBM25 | 1742.84933 | -0.142567755 | 0.61491646 | -0.23184898 | 0.81665531 | 0.99096399 | 2940.29006 | 1785.20914 | 717.869903 | 1528.02823 | Non-sig. |
| PF3D7_0905100 | NUP221 | 2422.03842 | -0.142692835 | 0.50706289 | -0.28141053 | 0.77839555 | 0.99028237 | 1686.33173 | 1781.58831 | 3396.80425 | 2823.42938 | Non-sig. |
| PF3D7_0912500 | N/A | 1533.49241 | -0.142828388 | 0.41337383 | -0.3455187 | 0.72970446 | 0.98844303 | 1728.5796 | 1497.5367 | 1490.19944 | 1417.65388 | Non-sig. |
| PF3D7_1460900 | ARPS10 | 623.597574 | -0.142844111 | 0.4332755 | -0.32968426 | 0.74163855 | 0.98954916 | 680.827969 | 663.335438 | 628.116186 | 522.110705 | Non-sig. |
| PF3D7_0312500 | MFR3 | 265.297877 | -0.142903633 | 0.5290655 | -0.27010575 | 0.7870789 | 0.99028237 | 182.069402 | 219.973074 | 374.498057 | 284.650974 | Non-sig. |
| PF3D7_0726050 | N/A | 20.6488937 | -0.14317784 | 0.93003052 | -0.15394961 | 0.87764946 | 0.99421828 | 24.6605815 | 30.3994467 | 18.7290816 | 8.8064652 | Non-sig. |
| PF3D7_1206600 | RPC2 | 1173.62411 | -0.143714737 | 0.52236482 | -0.27512331 | 0.78322151 | 0.99028237 | 723.899151 | 1002.94698 | 1739.81574 | 1227.83458 | Non-sig. |
| PF3D7_1007800 | N/A | 162.533137 | -0.143801823 | 0.65409976 | -0.21984693 | 0.82599037 | 0.99096399 | 271.754179 | 152.335495 | 69.8180157 | 156.224857 | Non-sig. |
| PF3D7_1221500 | N/A | 213.774622 | -0.144075546 | 0.50489132 | -0.28535952 | 0.77536872 | 0.99028237 | 165.289401 | 213.513468 | 283.354959 | 192.940658 | Non-sig. |
| PF3D7_0208700 | N/A | 551.606014 | -0.144597407 | 0.44246184 | -0.32680198 | 0.74381766 | 0.98998709 | 541.304923 | 599.624135 | 617.079095 | 448.415905 | Non-sig. |
| PF3D7_1035600 | H101 | 2028.29635 | -0.144655162 | 0.4494631 | -0.32183991 | 0.74757398 | 0.99028237 | 2608.10612 | 2113.72743 | 1651.94561 | 1739.40625 | Non-sig. |
| PF3D7_1107400 | RAD51 | 697.804315 | -0.144700333 | 0.52491257 | -0.27566559 | 0.78280493 | 0.99028237 | 428.944698 | 654.74463 | 1036.25187 | 671.276063 | Non-sig. |
| PF3D7_1236400 | N/A | 1195.30712 | -0.14477974 | 0.41375846 | -0.34991367 | 0.72640349 | 0.98844303 | 1313.16417 | 1178.2632 | 1197.36659 | 1092.43454 | Non-sig. |
| PF3D7_1019900 | ATG8 | 1050.88928 | -0.144977609 | 0.43773491 | -0.33119956 | 0.74049375 | 0.98954916 | 1072.45264 | 816.801303 | 1134.80299 | 1179.5002 | Non-sig. |
| PF3D7_0611000 | N/A | 393.64659 | -0.145125664 | 0.59960176 | -0.24203675 | 0.80875168 | 0.99028237 | 186.056172 | 375.690244 | 640.454435 | 372.385509 | Non-sig. |
| PF3D7_0713400 | SR25 | 1064.13209 | -0.145352166 | 0.50096704 | -0.29014317 | 0.77170671 | 0.99028237 | 704.751354 | 988.830603 | 1530.30058 | 1032.64582 | Non-sig. |
| PF3D7_1110800 | N/A | 67.0712976 | -0.145584353 | 0.66177163 | -0.21999183 | 0.82587752 | 0.99096399 | 101.739138 | 67.6659787 | 39.3973317 | 59.482742 | Non-sig. |
| PF3D7_1031600 | GEXP15 | 4346.52752 | -0.146212573 | 0.48789412 | -0.29968095 | 0.76442053 | 0.99028237 | 6196.03643 | 4123.30472 | 2937.49729 | 4129.27165 | Non-sig. |
| PF3D7_0827300 | N/A | 657.319148 | -0.146789242 | 0.43700464 | -0.33589859 | 0.73694734 | 0.98954916 | 790.327714 | 649.198747 | 591.307107 | 598.443023 | Non-sig. |
| PF3D7_1327800 | N/A | 963.873099 | -0.147404375 | 0.51263988 | -0.28753981 | 0.77369902 | 0.99028237 | 612.430851 | 932.505523 | 1413.36934 | 897.186682 | Non-sig. |
| PF3D7_1343400 | RAD5 | 993.981152 | -0.147442865 | 0.44014303 | -0.33498853 | 0.73763374 | 0.98954916 | 848.076523 | 917.182194 | 1241.15907 | 969.506816 | Non-sig. |
| PF3D7_1112300 | N/A | 238.664911 | -0.147895599 | 0.45750662 | -0.32326439 | 0.74649502 | 0.98998709 | 257.881359 | 214.599119 | 243.922683 | 238.256485 | Non-sig. |
| PF3D7_0520200 | MED17 | 1080.76335 | -0.148138284 | 0.43410601 | -0.34124909 | 0.73291607 | 0.98904466 | 943.305129 | 1011.77564 | 1328.89307 | 1039.07956 | Non-sig. |
| PF3D7_0920300 | N/A | 45.5415542 | -0.148524667 | 0.70376242 | -0.21104376 | 0.83285313 | 0.99110542 | 60.6195235 | 56.9814737 | 35.274905 | 29.2903145 | Non-sig. |
| PF3D7_0201500 | N/A | 1373.56083 | -0.148839277 | 0.41267492 | -0.36066955 | 0.71834649 | 0.98797939 | 1446.32671 | 1386.0031 | 1442.38085 | 1219.53265 | Non-sig. |
| PF3D7_1341600 | GPI10 | 314.025245 | -0.148976103 | 0.65533409 | -0.22732848 | 0.82016833 | 0.99096399 | 118.906907 | 288.659228 | 541.201834 | 307.33301 | Non-sig. |
| PF3D7_1440700 | AP3M | 1599.21198 | -0.149545796 | 0.46631714 | -0.32069548 | 0.74844118 | 0.99028237 | 1201.84541 | 1449.32673 | 2161.89991 | 1583.77586 | Non-sig. |
| PF3D7_1336000 | N/A | 838.12086 | -0.149618504 | 0.4317502 | -0.34653951 | 0.72893729 | 0.98844303 | 769.414586 | 850.145168 | 993.516845 | 739.40684 | Non-sig. |
| PF3D7_0616400 | N/A | 655.24138 | -0.149671043 | 0.46782391 | -0.3199303 | 0.74902117 | 0.99028237 | 535.345709 | 533.100233 | 842.816914 | 709.702664 | Non-sig. |
| PF3D7_1322100 | SET2 | 1090.15329 | -0.150132613 | 0.42965326 | -0.34942738 | 0.72676848 | 0.98844303 | 1094.19845 | 883.778855 | 1199.39169 | 1183.24418 | Non-sig. |
| PF3D7_0502200 | MPV17 | 489.181843 | -0.150277072 | 0.4378205 | -0.343239 | 0.73141866 | 0.98878696 | 464.984949 | 431.410509 | 564.17129 | 496.160623 | Non-sig. |
| PF3D7_0615700 | ALYREF | 1253.12149 | -0.150281301 | 0.44312135 | -0.33914254 | 0.73450235 | 0.98909017 | 1447.59996 | 1410.48295 | 1189.18036 | 965.222668 | Non-sig. |
| PF3D7_1437100 | N/A | 317.76443 | -0.150290126 | 0.5299562 | -0.28358971 | 0.77672482 | 0.99028237 | 456.312343 | 340.465566 | 212.586333 | 261.693477 | Non-sig. |
| PF3D7_1121200 | N/A | 101.43296 | -0.15037722 | 0.53627878 | -0.28040867 | 0.77916399 | 0.99028237 | 92.7903255 | 106.51015 | 120.543278 | 85.8880884 | Non-sig. |
| PF3D7_1469900 | MGET | 532.671182 | -0.150552258 | 0.74370762 | -0.20243474 | 0.83957687 | 0.99128968 | 985.74436 | 651.795307 | 135.449367 | 357.695693 | Non-sig. |
| PF3D7_0708800 | HSP110c | 6421.09289 | -0.151145071 | 0.41947993 | -0.36031538 | 0.7186113 | 0.98797939 | 7775.09299 | 6044.55055 | 5739.38997 | 6125.33805 | Non-sig. |
| PF3D7_0703300 | N/A | 299.910951 | -0.151426535 | 0.49734966 | -0.30446695 | 0.76077217 | 0.99028237 | 224.013469 | 282.634679 | 406.99624 | 285.999418 | Non-sig. |
| PF3D7_1403400 | N/A | 466.190062 | -0.151462829 | 0.48222286 | -0.31409301 | 0.7534504 | 0.99028237 | 608.730023 | 510.913823 | 372.788671 | 372.327732 | Non-sig. |
| PF3D7_0303600 | Plrx | 58.423014 | -0.151491204 | 0.72039948 | -0.21028777 | 0.83344308 | 0.9911446 | 32.4982584 | 73.0465708 | 90.2775422 | 37.8696847 | Non-sig. |
| PF3D7_0923400 | N/A | 1888.08014 | -0.151725759 | 0.50710329 | -0.2992009 | 0.76478677 | 0.99028237 | 2768.26412 | 1874.12456 | 1206.63201 | 1703.29987 | Non-sig. |
| PF3D7_1201500 | N/A | 429.977469 | -0.151777934 | 0.46260732 | -0.32809237 | 0.74284182 | 0.98977551 | 548.988964 | 415.346313 | 356.387401 | 399.187199 | Non-sig. |
| PF3D7_1115100 | N/A | 129.729793 | -0.151922551 | 0.56364006 | -0.26953824 | 0.78751552 | 0.99028237 | 174.005499 | 151.468466 | 99.3035359 | 94.1416727 | Non-sig. |
| PF3D7_0506700 | N/A | 1153.7801 | -0.152302826 | 0.42504427 | -0.35832226 | 0.72010216 | 0.98797939 | 1198.07446 | 948.474227 | 1231.15488 | 1237.41681 | Non-sig. |
| PF3D7_0516100 | ATPase1 | 740.920037 | -0.152521358 | 0.58984342 | -0.2585794 | 0.79595978 | 0.99028237 | 363.652972 | 584.549837 | 1196.09098 | 819.386358 | Non-sig. |
| PF3D7_1403600 | Sel1 | 1599.97077 | -0.154036573 | 0.48090095 | -0.32030831 | 0.74873462 | 0.99028237 | 2205.34891 | 1692.5833 | 1165.56519 | 1336.38567 | Non-sig. |
| PF3D7_0704800 | PPM12 | 1420.23691 | -0.154473011 | 0.45525976 | -0.33930741 | 0.73437816 | 0.98909017 | 1245.09518 | 1084.56574 | 1747.1132 | 1604.17352 | Non-sig. |
| PF3D7_0803100 | UTP14 | 2881.84795 | -0.154566583 | 0.58623734 | -0.26365872 | 0.79204291 | 0.99028237 | 4738.55998 | 2899.606 | 1333.9725 | 2555.25331 | Non-sig. |
| PF3D7_0524900 | TYW1 | 650.866365 | -0.154652168 | 0.42111189 | -0.36724722 | 0.71343462 | 0.9875928 | 669.935845 | 608.261478 | 701.471224 | 623.796912 | Non-sig. |
| PF3D7_1239700 | FTSH1 | 4883.76882 | -0.154847649 | 0.44837284 | -0.34535466 | 0.72982776 | 0.98844303 | 3854.43282 | 4640.7437 | 6436.50702 | 4603.39173 | Non-sig. |
| PF3D7_0928800 | N/A | 565.899227 | -0.154926508 | 0.54149465 | -0.28610903 | 0.77479462 | 0.99028237 | 859.380361 | 567.525893 | 333.475494 | 503.215159 | Non-sig. |
| PF3D7_1304400 | N/A | 1300.8122 | -0.155131063 | 0.46982978 | -0.33018567 | 0.74125967 | 0.98954916 | 1760.37572 | 1135.52632 | 981.280004 | 1326.06675 | Non-sig. |
| PF3D7_1235000 | PIH1 | 224.18193 | -0.155360269 | 0.47056697 | -0.33015549 | 0.74128247 | 0.98954916 | 220.409143 | 237.322885 | 252.008873 | 186.986817 | Non-sig. |
| PF3D7_0314000 | N/A | 1392.99254 | -0.155500109 | 0.42524806 | -0.36566918 | 0.71461195 | 0.9875928 | 1635.91341 | 1432.61714 | 1300.21653 | 1203.22306 | Non-sig. |
| PF3D7_1422700 | N/A | 1708.68187 | -0.155545105 | 0.54343177 | -0.28622748 | 0.7747039 | 0.99028237 | 2584.02545 | 1277.65177 | 1017.7177 | 1955.33254 | Non-sig. |

| | | | | | | | | | | | | |
|---|---|---|---|---|---|---|---|---|---|---|---|---|
| PF3D7_1249200 | N/A | 18.9667631 | -0.155802009 | 0.89990475 | -0.17313167 | 0.86254793 | 0.99402081 | 26.9654005 | 18.3160199 | 13.1086318 | 17.4770001 | Non-sig. |
| PF3D7_1302300 | N/A | 529.849157 | -0.156290292 | 0.47091722 | -0.33188485 | 0.73997621 | 0.98954916 | 414.607368 | 487.175679 | 702.169702 | 515.443877 | Non-sig. |
| PF3D7_1408900 | N/A | 244.528132 | -0.156315513 | 0.49293377 | -0.31711261 | 0.75115816 | 0.99028237 | 197.561883 | 252.269646 | 317.74781 | 210.533189 | Non-sig. |
| PF3D7_0423000 | N/A | 544.398919 | -0.157250276 | 0.53513575 | -0.29385119 | 0.76887162 | 0.99028237 | 794.58344 | 620.67726 | 353.802258 | 408.532718 | Non-sig. |
| PF3D7_1439000 | CTR1 | 3851.38754 | -0.157357253 | 0.47765132 | -0.32943959 | 0.74182345 | 0.98954916 | 2757.11679 | 3939.41609 | 5365.00731 | 3344.00996 | Non-sig. |
| PF3D7_0218100 | N/A | 718.60352 | -0.157499561 | 0.43163572 | -0.36489001 | 0.71519352 | 0.9875928 | 661.62595 | 702.185159 | 853.801447 | 656.801526 | Non-sig. |
| PF3D7_0309900 | N/A | 774.906063 | -0.157618238 | 0.44325565 | -0.35559217 | 0.72214601 | 0.98844303 | 698.59695 | 643.605749 | 935.620221 | 821.801333 | Non-sig. |
| PF3D7_0417500 | N/A | 1763.67247 | -0.157618319 | 0.47374805 | -0.33270495 | 0.73935701 | 0.98954916 | 1361.6046 | 1940.9907 | 2357.97171 | 1394.12287 | Non-sig. |
| PF3D7_0607400 | N/A | 551.928326 | -0.157637995 | 0.43908712 | -0.35901302 | 0.71958535 | 0.98797939 | 517.661923 | 475.390235 | 646.30909 | 568.352055 | Non-sig. |
| PF3D7_0206600 | N/A | 64.3378462 | -0.157814384 | 0.59527898 | -0.26510996 | 0.79092476 | 0.99028237 | 68.3707016 | 49.3672976 | 67.3375949 | 72.2757907 | Non-sig. |
| PF3D7_0631100 | N/A | 104.749688 | -0.157866421 | 0.53740837 | -0.29375505 | 0.76894509 | 0.99028237 | 131.579994 | 97.6409556 | 89.5213821 | 100.256419 | Non-sig. |
| PF3D7_1209700 | N/A | 398.818736 | -0.158676136 | 0.43491614 | -0.36484306 | 0.71522856 | 0.9875928 | 413.727324 | 391.906009 | 427.712326 | 361.929284 | Non-sig. |
| PF3D7_1340600 | DBR1 | 1308.89207 | -0.158790558 | 0.51083985 | -0.31084215 | 0.75592062 | 0.99028237 | 845.275118 | 1114.82172 | 1916.08817 | 1359.38328 | Non-sig. |
| PF3D7_1122100 | GPI16 | 486.140458 | -0.158862916 | 0.46081758 | -0.34474144 | 0.73028876 | 0.98844303 | 400.633249 | 460.443845 | 624.891777 | 458.59296 | Non-sig. |
| PF3D7_0703600 | N/A | 3051.13965 | -0.158892546 | 0.56775405 | -0.27986158 | 0.7795837 | 0.99028237 | 4721.43247 | 3690.43942 | 1716.88502 | 2075.8017 | Non-sig. |
| PF3D7_0204600 | N/A | 831.170747 | -0.159311141 | 0.47271683 | -0.33701178 | 0.73610802 | 0.98954916 | 641.565325 | 700.498267 | 1112.18986 | 870.429533 | Non-sig. |
| PF3D7_1102500 | GEXP02 | 1229.75665 | -0.15978843 | 0.41777532 | -0.38247456 | 0.7021094 | 0.98512564 | 1382.54495 | 1242.39956 | 1213.12149 | 1080.9606 | Non-sig. |
| PF3D7_0706700 | MSH2-2 | 42.1831857 | -0.160100356 | 0.66530474 | -0.24064214 | 0.80983249 | 0.99028237 | 50.5120762 | 35.4111456 | 38.5990073 | 44.2105135 | Non-sig. |
| PF3D7_0313400 | N/A | 308.74755 | -0.160553399 | 0.60143977 | -0.26694842 | 0.78950887 | 0.99028237 | 501.041235 | 312.23916 | 151.117853 | 270.591951 | Non-sig. |
| PF3D7_0927900 | PSD | 4683.45622 | -0.160755158 | 0.40382696 | -0.39807931 | 0.69057172 | 0.98334178 | 5156.52666 | 4365.52889 | 4731.77278 | 4479.99654 | Non-sig. |
| PF3D7_1477400 | N/A | 150.34497 | -0.160885103 | 0.53241315 | -0.30218094 | 0.76251414 | 0.99028237 | 133.412781 | 177.142863 | 183.904764 | 106.919472 | Non-sig. |
| PF3D7_0806000 | N/A | 1333.90395 | -0.161149039 | 0.41007101 | -0.39297838 | 0.69433545 | 0.98334178 | 1436.74151 | 1255.15236 | 1379.93684 | 1263.7851 | Non-sig. |
| PF3D7_1241700 | RFC4 | 1661.23124 | -0.161410958 | 0.47431414 | -0.34030391 | 0.73362767 | 0.98909017 | 1223.90416 | 1692.2085 | 2283.92409 | 1444.8882 | Non-sig. |
| PF3D7_1425800 | N/A | 2404.42744 | -0.161600682 | 0.66112732 | -0.24443201 | 0.80689625 | 0.99028237 | 4280.69418 | 2421.2699 | 797.56943 | 2118.17626 | Non-sig. |
| PF3D7_1431500 | MAPK1 | 1821.34739 | -0.161615711 | 0.45768626 | -0.35311463 | 0.72400251 | 0.98844303 | 2434.87721 | 1691.85601 | 1411.92853 | 1746.72781 | Non-sig. |
| PF3D7_1427300 | N/A | 2147.92737 | -0.161715534 | 0.4081025 | -0.39626205 | 0.69191172 | 0.98334178 | 2202.72019 | 1935.24327 | 2333.6102 | 2120.13581 | Non-sig. |
| PF3D7_0915500 | N/A | 56.9463097 | -0.161771562 | 0.60826188 | -0.26595709 | 0.79027225 | 0.99028237 | 53.679593 | 48.2852656 | 66.5239712 | 59.2964089 | Non-sig. |
| PF3D7_0812700 | SNRPC | 673.626533 | -0.161869876 | 0.45127395 | -0.35869537 | 0.71982299 | 0.98797939 | 571.874074 | 683.742665 | 850.665414 | 588.22398 | Non-sig. |
| PF3D7_0921800 | RPF1 | 228.585361 | -0.16188581 | 0.48267035 | -0.33539622 | 0.73732622 | 0.98954916 | 202.692653 | 241.576102 | 279.949348 | 190.123341 | Non-sig. |
| PF3D7_1447200 | N/A | 1587.56782 | -0.161949666 | 0.62144081 | -0.26060352 | 0.79439827 | 0.99028237 | 2706.42793 | 1672.0489 | 647.097864 | 1324.69658 | Non-sig. |
| PF3D7_0612600 | NCS6 | 1878.46162 | -0.162218252 | 0.40857284 | -0.39703631 | 0.69134068 | 0.98334178 | 2050.27953 | 1834.85782 | 1917.68568 | 1711.02342 | Non-sig. |
| PF3D7_1428800 | N/A | 625.274839 | -0.162418192 | 0.461419 | -0.35199719 | 0.72484037 | 0.98844303 | 808.879066 | 640.22504 | 512.256997 | 539.738254 | Non-sig. |
| PF3D7_1407900 | PMI | 742.730956 | -0.162621502 | 0.58606656 | -0.27747958 | 0.78141188 | 0.99028237 | 361.291631 | 636.158619 | 1207.4445 | 766.029079 | Non-sig. |
| PF3D7_0821600 | CLP1 | 535.300473 | -0.16287074 | 0.45919758 | -0.35468554 | 0.72282519 | 0.98844303 | 459.81335 | 444.264409 | 670.951172 | 566.172959 | Non-sig. |
| PF3D7_1215700 | N/A | 19.4421263 | -0.163198425 | 0.92683954 | -0.17608056 | 0.86023065 | 0.99402081 | 30.5346769 | 19.3121506 | 10.6773579 | 17.2443198 | Non-sig. |
| PF3D7_0210900 | N/A | 4122.94363 | -0.163630571 | 0.41057055 | -0.39854435 | 0.69022897 | 0.98334178 | 4484.8012 | 4257.47737 | 4228.25593 | 3521.24 | Non-sig. |
| PF3D7_1349500 | N/A | 12745.9085 | -0.163633957 | 0.45681981 | -0.35820241 | 0.72019184 | 0.98797939 | 11312.8304 | 9226.68733 | 15622.9431 | 14821.173 | Non-sig. |
| PF3D7_1220600 | N/A | 242.425596 | -0.164169261 | 0.45491879 | -0.36087597 | 0.71819216 | 0.98797939 | 255.566314 | 235.311311 | 256.840339 | 221.984419 | Non-sig. |
| PF3D7_1126500 | N/A | 939.563066 | -0.164331991 | 0.52068809 | -0.31560543 | 0.75230201 | 0.99028237 | 582.67341 | 866.729764 | 1403.01225 | 905.836845 | Non-sig. |
| PF3D7_1327500 | N/A | 687.817929 | -0.164465374 | 0.46405347 | -0.3544104 | 0.72303135 | 0.98844303 | 894.039475 | 577.198648 | 560.162709 | 719.870886 | Non-sig. |
| PF3D7_1430600 | N/A | 585.128074 | -0.164637757 | 0.42363786 | -0.38862852 | 0.69755097 | 0.98334178 | 617.729196 | 568.722233 | 619.228135 | 534.832731 | Non-sig. |
| PF3D7_1457800 | N/A | 259.649355 | -0.164845111 | 0.56341451 | -0.29258229 | 0.76984145 | 0.99028237 | 392.393788 | 279.022324 | 156.850905 | 210.330403 | Non-sig. |
| PF3D7_0105500 | N/A | 35.4463235 | -0.165083816 | 0.75338929 | -0.21912153 | 0.82655538 | 0.99110542 | 27.2365877 | 22.5631441 | 47.594692 | 44.3908701 | Non-sig. |
| PF3D7_0321900 | CARL | 3941.84342 | -0.16524124 | 0.42287091 | -0.39076048 | 0.69597429 | 0.98334178 | 3517.49255 | 3585.87835 | 4816.98957 | 3847.0132 | Non-sig. |
| PF3D7_1416000 | N/A | 3.43311875 | -0.16540228 | 2.06435029 | -0.08012317 | 0.9361393 | 0.99639249 | 3.13641242 | 6.48115512 | 4.11490747 | 0 | Non-sig. |
| PF3D7_1354000 | N/A | 52.0660191 | -0.166170355 | 0.65739657 | -0.25277034 | 0.80044569 | 0.99028237 | 72.5191347 | 46.1582395 | 37.7693625 | 51.8173395 | Non-sig. |
| PF3D7_0903700 | N/A | 4348.82791 | -0.166205835 | 0.6051472 | -0.27465356 | 0.78358241 | 0.99028237 | 1889.21309 | 3543.11723 | 7308.5364 | 4654.44491 | Non-sig. |
| PF3D7_1218900 | N/A | 328.3349 | -0.166413171 | 0.47879012 | -0.34757018 | 0.728163 | 0.98844303 | 422.048167 | 274.693317 | 272.663806 | 343.934308 | Non-sig. |
| PF3D7_1222900 | N/A | 238.994578 | -0.166783578 | 0.51181538 | -0.32586668 | 0.74452523 | 0.98998709 | 180.961107 | 252.186151 | 324.383213 | 198.447842 | Non-sig. |
| PF3D7_1147900 | PEX22 | 1208.36833 | -0.167048285 | 0.45157096 | -0.36992699 | 0.71143689 | 0.9875928 | 974.721277 | 1135.56248 | 1581.51277 | 1141.67678 | Non-sig. |
| PF3D7_0109600 | CoSP | 108.359556 | -0.167430139 | 0.54270056 | -0.30851293 | 0.75769207 | 0.99028237 | 140.976155 | 103.448468 | 88.4726167 | 100.540983 | Non-sig. |
| PF3D7_0406800 | N/A | 307.743551 | -0.168195211 | 0.51093526 | -0.32919085 | 0.74201144 | 0.98954916 | 236.708995 | 347.058992 | 414.3897 | 232.816516 | Non-sig. |
| PF3D7_0203300 | ERCC1 | 370.912871 | -0.168417476 | 0.44499158 | -0.3784734 | 0.70507895 | 0.98615891 | 354.140745 | 368.512373 | 430.828065 | 330.170302 | Non-sig. |
| PF3D7_0423200 | N/A | 891.202663 | -0.168533188 | 0.4547535 | -0.37060339 | 0.71093296 | 0.9875928 | 1131.05625 | 948.21544 | 755.559464 | 729.979502 | Non-sig. |

| | | | | | | | | | | | | |
|---|---|---|---|---|---|---|---|---|---|---|---|---|
| PF3D7_0404000 | N/A | 2136.40892 | -0.168842232 | 0.52256865 | -0.32310058 | 0.74661907 | 0.98998709 | 3212.93074 | 2225.40418 | 1309.98155 | 1797.31919 | Non-sig. |
| PF3D7_1208100 | N/A | 1032.26503 | -0.168910419 | 0.48965496 | -0.34495805 | 0.73012591 | 0.98844303 | 749.713537 | 835.445392 | 1435.23551 | 1108.66566 | Non-sig. |
| PF3D7_0713700 | N/A | 1157.72173 | -0.169676404 | 0.44834857 | -0.37844752 | 0.70509817 | 0.98615891 | 953.308494 | 1047.24318 | 1497.8909 | 1132.44434 | Non-sig. |
| PF3D7_1006200 | ALBA3 | 3816.07599 | -0.170045678 | 0.44744779 | -0.38003468 | 0.70391967 | 0.98582247 | 3080.51188 | 3356.34056 | 5000.64175 | 3826.80977 | Non-sig. |
| PF3D7_1328400 | N/A | 305.222056 | -0.170312907 | 0.47157662 | -0.36115639 | 0.71798254 | 0.98797939 | 389.567847 | 285.39325 | 257.076484 | 288.850643 | Non-sig. |
| PF3D7_1462900 | N/A | 150.873151 | -0.171587818 | 0.50076829 | -0.34264913 | 0.73186244 | 0.98904466 | 134.734956 | 135.196004 | 184.791135 | 148.770508 | Non-sig. |
| PF3D7_1110600 | N/A | 1639.24852 | -0.172004323 | 0.42627036 | -0.40350993 | 0.68657313 | 0.98210107 | 1889.35386 | 1738.27264 | 1584.46498 | 1344.90258 | Non-sig. |
| PF3D7_1441000 | NCS2 | 309.989906 | -0.172159133 | 0.49429614 | -0.34829148 | 0.72762129 | 0.98844303 | 419.004226 | 313.298179 | 238.181469 | 269.475748 | Non-sig. |
| PF3D7_1212800 | SDHB | 1378.44687 | -0.172335163 | 0.43043676 | -0.40037278 | 0.68888197 | 0.98334178 | 1251.50888 | 1405.62586 | 1669.67831 | 1186.97444 | Non-sig. |
| PF3D7_1427000 | N/A | 1491.29863 | -0.172364073 | 0.51345738 | -0.33569305 | 0.73710235 | 0.98954916 | 2218.74009 | 1470.7012 | 942.167137 | 1333.58608 | Non-sig. |
| PF3D7_0612000 | N/A | 504.747012 | -0.172444412 | 0.44858448 | -0.38441903 | 0.7006679 | 0.98491338 | 449.70012 | 509.928524 | 619.879029 | 439.480375 | Non-sig. |
| PF3D7_0209400 | N/A | 740.195352 | -0.172451905 | 0.46428383 | -0.37143638 | 0.71031254 | 0.9875928 | 588.874384 | 743.787828 | 979.631654 | 648.487543 | Non-sig. |
| PF3D7_1348300 | N/A | 422.366213 | -0.172694922 | 0.49272548 | -0.35048913 | 0.72597165 | 0.98844303 | 315.02437 | 362.574021 | 579.927327 | 431.939135 | Non-sig. |
| PF3D7_1205400 | N/A | 987.829392 | -0.17281299 | 0.52422493 | -0.32965428 | 0.74166121 | 0.98954916 | 700.481713 | 669.409867 | 1393.08518 | 1188.34081 | Non-sig. |
| PF3D7_1037100 | PyKII | 746.816246 | -0.172949834 | 0.45213158 | -0.38252102 | 0.70207494 | 0.98512564 | 621.073294 | 729.619527 | 961.743138 | 674.829023 | Non-sig. |
| PF3D7_1346200 | N/A | 56.5895206 | -0.172967564 | 0.61673642 | -0.28045622 | 0.77912751 | 0.99028237 | 48.393926 | 51.5111556 | 71.4457448 | 55.007256 | Non-sig. |
| PF3D7_1108300 | PCD | 159.78436 | -0.173379627 | 0.49039841 | -0.35354851 | 0.72367727 | 0.98844303 | 167.991528 | 169.652532 | 170.753051 | 130.740331 | Non-sig. |
| PF3D7_0805400 | N/A | 349.72527 | -0.173518286 | 0.44608919 | -0.38897666 | 0.69729341 | 0.98334178 | 398.478278 | 353.388514 | 343.071797 | 303.962492 | Non-sig. |
| PF3D7_1212500 | GAT | 2217.52549 | -0.173820635 | 0.5905646 | -0.29432959 | 0.76850607 | 0.99028237 | 1029.50559 | 1855.16613 | 3672.03404 | 2313.39621 | Non-sig. |
| PF3D7_1407100 | NOP1 | 1018.8172 | -0.174173144 | 0.41931227 | -0.41537813 | 0.67786512 | 0.97898416 | 1014.02116 | 1020.60643 | 1146.38764 | 894.253579 | Non-sig. |
| PF3D7_1226500 | N/A | 424.502305 | -0.174359838 | 0.43360306 | -0.40211856 | 0.68759677 | 0.98297024 | 462.467894 | 380.823792 | 437.810669 | 416.906864 | Non-sig. |
| PF3D7_1132600 | PRP38A | 1344.94308 | -0.174415167 | 0.4797341 | -0.36356633 | 0.71618187 | 0.9875928 | 1837.67633 | 1451.18443 | 1014.89723 | 1076.01432 | Non-sig. |
| PF3D7_1200400 | VAR | 451.276188 | -0.174482466 | 0.46689377 | -0.37370913 | 0.70862073 | 0.98718713 | 589.273686 | 437.597394 | 368.033478 | 410.200193 | Non-sig. |
| PF3D7_0602600 | N/A | 1348.49729 | -0.174962048 | 0.43104896 | -0.40589832 | 0.68481731 | 0.98115715 | 1200.0342 | 1207.86733 | 1660.10154 | 1325.98609 | Non-sig. |
| PF3D7_0615900 | N/A | 1807.63201 | -0.175395213 | 0.52604231 | -0.33342415 | 0.73881413 | 0.98954916 | 2750.76955 | 1603.54644 | 1084.34458 | 1791.86749 | Non-sig. |
| PF3D7_1142300 | N/A | 6541.99194 | -0.175549812 | 0.42297053 | -0.4150403 | 0.6781124 | 0.97898416 | 6145.54247 | 5403.7784 | 7733.3592 | 6885.2877 | Non-sig. |
| PF3D7_1115400 | FP3 | 4559.45391 | -0.17565911 | 0.41284117 | -0.42548835 | 0.67048071 | 0.97883902 | 4348.40938 | 4053.20008 | 5324.82968 | 4511.3765 | Non-sig. |
| PF3D7_0528700 | CYP23 | 493.819693 | -0.17581027 | 0.43318175 | -0.40585798 | 0.68484695 | 0.98115715 | 490.955248 | 493.747001 | 556.714893 | 433.861629 | Non-sig. |
| PF3D7_0824200 | N/A | 690.671037 | -0.175838183 | 0.5443783 | -0.32300733 | 0.74668969 | 0.98998709 | 398.107334 | 673.730117 | 1066.96293 | 623.883773 | Non-sig. |
| PF3D7_1441700 | ATP23 | 178.844015 | -0.175876293 | 0.47263512 | -0.37211854 | 0.70980459 | 0.9875928 | 192.281914 | 170.690968 | 187.195871 | 165.207306 | Non-sig. |
| PF3D7_0629800 | N/A | 1596.31974 | -0.175901395 | 0.40826873 | -0.43084709 | 0.66657956 | 0.97883902 | 1699.06992 | 1525.48039 | 1687.967 | 1472.76167 | Non-sig. |
| PF3D7_0923600 | LipL2 | 252.883936 | -0.17614841 | 0.52134525 | -0.33787286 | 0.735459 | 0.98954916 | 178.426449 | 222.288856 | 357.901852 | 252.918585 | Non-sig. |
| PF3D7_0709600 | POP1 | 574.367644 | -0.176177206 | 0.48082624 | -0.36640514 | 0.71406279 | 0.9875928 | 787.849228 | 526.742647 | 431.226322 | 551.652381 | Non-sig. |
| PF3D7_1438500 | CPSF3 | 528.322716 | -0.17621528 | 0.52828451 | -0.33356132 | 0.73871061 | 0.98954916 | 333.883237 | 453.794358 | 786.883416 | 538.729854 | Non-sig. |
| PF3D7_0631000 | N/A | 1317.19364 | -0.176232957 | 0.41669016 | -0.42293526 | 0.6723425 | 0.97898416 | 1431.04178 | 1347.80508 | 1364.08305 | 1125.84464 | Non-sig. |
| PF3D7_0108600 | N/A | 399.950689 | -0.176698885 | 0.45834383 | -0.38551601 | 0.69985515 | 0.98429322 | 482.735415 | 425.004604 | 366.241928 | 325.820808 | Non-sig. |
| PF3D7_1148640 | N/A | 48.7423622 | -0.176774259 | 4.56682677 | -0.03870833 | 0.96912293 | 0.99691622 | 0 | 0 | 103.44062 | 91.5288289 | Non-sig. |
| PF3D7_0920600 | N/A | 2046.02914 | -0.176907583 | 0.42180506 | -0.41940603 | 0.67491942 | 0.97898416 | 2460.94862 | 1830.08469 | 1881.85616 | 2011.22711 | Non-sig. |
| PF3D7_1442600 | TREP | 493.536933 | -0.177021716 | 0.69943356 | -0.25309297 | 0.80019637 | 0.99028237 | 902.483471 | 453.667801 | 145.409592 | 472.586867 | Non-sig. |
| PF3D7_0416300 | MCM9 | 2447.85253 | -0.177361748 | 0.42412921 | -0.41817858 | 0.67581656 | 0.97898416 | 2517.46587 | 1944.71881 | 2678.75284 | 2650.47258 | Non-sig. |
| PF3D7_1306300 | N/A | 204.605283 | -0.177579206 | 0.48817057 | -0.36376467 | 0.71603375 | 0.9875928 | 201.042531 | 156.657285 | 233.246 | 227.475315 | Non-sig. |
| PF3D7_0308000 | N/A | 572.373241 | -0.177804939 | 0.44866884 | -0.39629438 | 0.69188787 | 0.98334178 | 650.270929 | 630.981894 | 565.012817 | 443.227324 | Non-sig. |
| PF3D7_1202600 | NEMF | 19642.6722 | -0.178024149 | 0.42237013 | -0.4214885 | 0.67339841 | 0.97898416 | 19863.8555 | 15444.382 | 21842.2168 | 21420.2345 | Non-sig. |
| PF3D7_0516600 | MB2 | 3205.56736 | -0.178028825 | 0.43673855 | -0.40763249 | 0.68354351 | 0.98115715 | 3826.01183 | 2496.98542 | 2980.33569 | 3518.93649 | Non-sig. |
| PF3D7_1443700 | DPCK | 1357.10509 | -0.178635817 | 0.45238439 | -0.39487617 | 0.69293428 | 0.98334178 | 1090.11934 | 1269.81292 | 1791.64794 | 1276.84015 | Non-sig. |
| PF3D7_1345300 | ApiCOX16 | 894.162273 | -0.178870893 | 0.43980359 | -0.40670631 | 0.68422371 | 0.98115715 | 1073.68787 | 947.667436 | 825.513244 | 729.780537 | Non-sig. |
| PF3D7_1034800 | N/A | 372.151952 | -0.179501524 | 0.5227341 | -0.34338973 | 0.73130528 | 0.98878696 | 542.927545 | 371.497413 | 247.92873 | 326.254118 | Non-sig. |
| PF3D7_1442900 | ARFGEF | 10672.5778 | -0.179997737 | 0.4444774 | -0.40496488 | 0.68550333 | 0.98183202 | 8800.73465 | 9029.24864 | 13874.0103 | 10986.3177 | Non-sig. |
| PF3D7_1009800 | N/A | 432.104402 | -0.180007503 | 0.4611839 | -0.3903161 | 0.69630282 | 0.98334178 | 366.658207 | 381.880418 | 551.179424 | 428.69956 | Non-sig. |
| PF3D7_1436800 | ClpR | 124.641012 | -0.180144274 | 0.58195521 | -0.30955006 | 0.75690314 | 0.99028237 | 80.7267317 | 108.493663 | 183.84002 | 125.503635 | Non-sig. |
| PF3D7_0717400 | N/A | 1557.74955 | -0.18023145 | 0.43935322 | -0.41021994 | 0.68164462 | 0.98067094 | 1975.39401 | 1364.7027 | 1334.68346 | 1556.21802 | Non-sig. |
| PF3D7_0926900 | N/A | 1079.92821 | -0.180578522 | 0.49414272 | -0.36543799 | 0.71478449 | 0.9875928 | 1497.01881 | 1203.24601 | 798.119736 | 821.328294 | Non-sig. |
| PF3D7_1439700 | N/A | 1313.60863 | -0.180590212 | 0.45542953 | -0.39652723 | 0.69171612 | 0.98334178 | 1089.47472 | 1388.43567 | 1701.72547 | 1074.79866 | Non-sig. |
| PF3D7_1361900 | PCNA1 | 7416.66048 | -0.180965023 | 0.43459691 | -0.4163974 | 0.67711923 | 0.97898416 | 6595.91778 | 7880.26291 | 9166.31334 | 6024.1479 | Non-sig. |

| | | | | | | | | | | | | |
|---|---|---|---|---|---|---|---|---|---|---|---|---|
| PF3D7_1014400 | MRK | 643.22719 | -0.18213506 | 0.52498299 | -0.34693517 | 0.72864003 | 0.98844303 | 403.057788 | 627.737116 | 964.15958 | 577.954276 | Non-sig. |
| PF3D7_0322300 | DGAT | 2488.74983 | -0.18251641 | 0.43621343 | -0.4184108 | 0.67564679 | 0.97898416 | 2178.57835 | 2091.84143 | 3113.14475 | 2571.4348 | Non-sig. |
| PF3D7_0203000 | ROPE | 10367.9168 | -0.182626522 | 0.70505087 | -0.25902602 | 0.79561516 | 0.99028237 | 19299.0295 | 8865.97159 | 2747.86456 | 10558.8016 | Non-sig. |
| PF3D7_0507000 | N/A | 99.4739937 | -0.18273264 | 0.52812166 | -0.34600482 | 0.72933909 | 0.98844303 | 100.689721 | 100.962094 | 110.804886 | 85.4392738 | Non-sig. |
| PF3D7_1127300 | N/A | 330.227895 | -0.182862245 | 0.45992664 | -0.39759003 | 0.69093241 | 0.98334178 | 300.598919 | 337.058599 | 401.507644 | 281.746419 | Non-sig. |
| PF3D7_1009900 | N/A | 151.536728 | -0.182924614 | 0.48999124 | -0.37332221 | 0.70890865 | 0.9873286 | 154.690586 | 154.522106 | 167.535615 | 129.398604 | Non-sig. |
| PF3D7_0912200 | N/A | 5.17167021 | -0.182976443 | 1.49473992 | -0.12241357 | 0.9025715 | 0.99457954 | 5.24797231 | 5.36925712 | 5.74615683 | 4.32329458 | Non-sig. |
| PF3D7_1447700 | N/A | 664.915355 | -0.183049834 | 0.57449769 | -0.31862588 | 0.75001022 | 0.99028237 | 341.468678 | 586.153862 | 1072.26248 | 659.776399 | Non-sig. |
| PF3D7_0606400 | N/A | 9.32380653 | -0.183114945 | 1.20731978 | -0.15167062 | 0.87944673 | 0.99421828 | 13.3521827 | 11.8994489 | 6.53252935 | 5.51106512 | Non-sig. |
| PF3D7_0724300 | COQ3 | 281.763206 | -0.183116847 | 0.46792514 | -0.39133791 | 0.69554748 | 0.98334178 | 255.121702 | 238.376592 | 343.965221 | 289.589308 | Non-sig. |
| PF3D7_0105400 | N/A | 2028.91608 | -0.183416509 | 0.45109028 | -0.4066071 | 0.68429658 | 0.98115715 | 1677.88641 | 2117.17788 | 2637.31874 | 1683.28129 | Non-sig. |
| PF3D7_0630900 | HAS1 | 518.675519 | -0.18412419 | 0.42866944 | -0.42952488 | 0.66754129 | 0.97883902 | 574.615044 | 507.549771 | 528.894992 | 463.64227 | Non-sig. |
| PF3D7_1326600 | N/A | 4967.35808 | -0.184696846 | 0.42857834 | -0.43095236 | 0.66650301 | 0.97883902 | 5760.56417 | 3894.07746 | 4809.32788 | 5405.46282 | Non-sig. |
| PF3D7_1131600 | N/A | 251.291252 | -0.18486 | 0.48915767 | -0.37791496 | 0.70549377 | 0.98615891 | 203.447109 | 241.323818 | 331.064367 | 229.329713 | Non-sig. |
| PF3D7_1304300 | N/A | 43.8198893 | -0.185104924 | 0.66959362 | -0.27644368 | 0.78220731 | 0.99028237 | 43.9874934 | 31.1606257 | 49.2338888 | 50.8975493 | Non-sig. |
| PF3D7_1134000 | HSP70-3 | 4288.50475 | -0.185163236 | 0.4735925 | -0.39097587 | 0.69581507 | 0.98334178 | 3125.52468 | 4265.66339 | 6000.82881 | 3762.00213 | Non-sig. |
| PF3D7_0305000 | EF-Ts | 1249.54797 | -0.185253562 | 0.4353038 | -0.42557304 | 0.67041899 | 0.97883902 | 1095.01078 | 1219.47797 | 1564.11083 | 1119.59232 | Non-sig. |
| PF3D7_1128700 | GPI8 | 1153.88805 | -0.185363594 | 0.52302189 | -0.35440886 | 0.72303249 | 0.98844303 | 720.065371 | 1188.8314 | 1735.42585 | 971.229566 | Non-sig. |
| PF3D7_0920400 | N/A | 1274.42904 | -0.185807709 | 0.43456646 | -0.4275703 | 0.66896399 | 0.97883902 | 1136.16059 | 1100.48889 | 1576.4116 | 1284.65506 | Non-sig. |
| PF3D7_1224500 | ASF1 | 2192.7248 | -0.186268958 | 0.58072009 | -0.32075515 | 0.74839595 | 0.99028237 | 1048.37523 | 2060.14681 | 3619.42357 | 2042.95359 | Non-sig. |
| PF3D7_0722000 | N/A | 1095.90493 | -0.186363312 | 0.51233938 | -0.36374973 | 0.7160449 | 0.9875928 | 1587.88713 | 1209.58805 | 745.606396 | 840.538149 | Non-sig. |
| PF3D7_0217900 | N/A | 2163.21987 | -0.18672864 | 0.60206894 | -0.31014495 | 0.75645073 | 0.99028237 | 947.518859 | 1920.54828 | 3658.15359 | 2126.65877 | Non-sig. |
| PF3D7_0320100 | SEC22 | 1174.2969 | -0.186750811 | 0.41931094 | -0.44537548 | 0.65604843 | 0.97496341 | 1300.89732 | 1201.25073 | 1199.54843 | 995.491133 | Non-sig. |
| PF3D7_1423400 | N/A | 131.797118 | -0.187207522 | 0.58827795 | -0.31822971 | 0.7503107 | 0.99028237 | 193.84965 | 98.6904353 | 87.0703416 | 147.578044 | Non-sig. |
| PF3D7_0416200 | N/A | 587.961975 | -0.187689555 | 0.71734909 | -0.26164326 | 0.79359649 | 0.99028237 | 1079.43756 | 708.851546 | 173.188741 | 390.370056 | Non-sig. |
| PF3D7_1115300 | FP2B | 198.933401 | -0.189181867 | 0.52699657 | -0.35898121 | 0.71960915 | 0.98797939 | 145.247939 | 201.726174 | 278.414226 | 170.345266 | Non-sig. |
| PF3D7_1006400 | N/A | 217.376379 | -0.189241547 | 0.52358551 | -0.36143389 | 0.71777512 | 0.98797939 | 155.791076 | 200.646668 | 307.165234 | 205.90254 | Non-sig. |
| PF3D7_1438100 | SEC62 | 5413.88572 | -0.189314854 | 0.43973289 | -0.43052239 | 0.66681568 | 0.97883902 | 4482.05638 | 5198.56044 | 7054.87075 | 4920.05531 | Non-sig. |
| PF3D7_1338000 | N/A | 1463.66909 | -0.189321719 | 0.47385244 | -0.39953729 | 0.68949736 | 0.98334178 | 1087.05106 | 1290.90086 | 2031.78248 | 1444.94194 | Non-sig. |
| PF3D7_1477600 | SURF14.1 | 146.786339 | -0.189497809 | 0.57542183 | -0.32931981 | 0.74191397 | 0.98954916 | 220.278884 | 139.437656 | 92.8269473 | 134.601867 | Non-sig. |
| PF3D7_MIT04100 | LSUD | 2.98759995 | -0.189577716 | 1.96462088 | -0.09649583 | 0.92312679 | 0.99619209 | 3.12767994 | 4.55481496 | 3.2222479 | 1.04565699 | Non-sig. |
| PF3D7_0801500 | NOL10 | 588.323448 | -0.18960703 | 0.42316946 | -0.44806406 | 0.65410696 | 0.97317009 | 635.489754 | 559.096697 | 618.384855 | 540.322484 | Non-sig. |
| PF3D7_1456300 | N/A | 103.550851 | -0.189635445 | 0.5431076 | -0.34916736 | 0.72696367 | 0.98844303 | 88.3362991 | 104.107127 | 132.206459 | 89.5535188 | Non-sig. |
| PF3D7_0202800 | N/A | 520.704675 | -0.189816604 | 0.45036869 | -0.42146936 | 0.67341238 | 0.97898416 | 636.062998 | 542.862138 | 473.923351 | 429.970215 | Non-sig. |
| PF3D7_1220900 | HP1 | 11397.4526 | -0.190012359 | 0.43799346 | -0.43382465 | 0.66441578 | 0.97883902 | 14842.071 | 10414.7656 | 9452.05687 | 10880.917 | Non-sig. |
| PF3D7_0107200 | CCR4-3 | 123.286888 | -0.190594802 | 0.53927898 | -0.35342524 | 0.72376967 | 0.98844303 | 101.783578 | 131.019654 | 160.884842 | 99.459477 | Non-sig. |
| PF3D7_0609800 | DHHC2 | 755.056049 | -0.190793962 | 0.44418638 | -0.42953583 | 0.66753333 | 0.97883902 | 874.975481 | 584.927008 | 734.940274 | 825.381431 | Non-sig. |
| PF3D7_1353400 | N/A | 1110.53452 | -0.191037325 | 0.42626531 | -0.44816531 | 0.65403389 | 0.97317009 | 1110.2672 | 910.776684 | 1257.56101 | 1163.53317 | Non-sig. |
| PF3D7_0419800 | N/A | 585.709536 | -0.191044043 | 0.49889004 | -0.38293818 | 0.7017656 | 0.98512564 | 838.644346 | 561.42339 | 410.531804 | 532.238604 | Non-sig. |
| PF3D7_0705000 | N/A | 2496.41173 | -0.191409673 | 0.42235971 | -0.45319112 | 0.65041112 | 0.97317009 | 3057.43559 | 2361.04194 | 2266.30469 | 2300.86471 | Non-sig. |
| PF3D7_1310500 | EMC7 | 388.487172 | -0.191447128 | 0.45479159 | -0.42095573 | 0.67378741 | 0.97898416 | 344.671226 | 352.107233 | 483.597909 | 373.572321 | Non-sig. |
| PF3D7_0417400 | N/A | 10848.5518 | -0.191634224 | 0.42489309 | -0.45101752 | 0.65197693 | 0.97317009 | 11934.0127 | 8369.10012 | 11202.0419 | 11889.0525 | Non-sig. |
| PF3D7_0908300 | N/A | 782.959192 | -0.191741746 | 0.48668667 | -0.39397369 | 0.69360046 | 0.98334178 | 570.582901 | 784.302946 | 1099.18956 | 677.76136 | Non-sig. |
| PF3D7_1226300 | HAD2 | 5596.10921 | -0.19235838 | 0.51769562 | -0.37156656 | 0.71021559 | 0.9875928 | 3416.54919 | 5253.89167 | 8520.3588 | 5193.63717 | Non-sig. |
| PF3D7_1422100 | N/A | 1446.98989 | -0.193194257 | 0.45367948 | -0.42583865 | 0.67022542 | 0.97883902 | 1927.4527 | 1324.44751 | 1160.27834 | 1375.781 | Non-sig. |
| PF3D7_1026300 | N/A | 943.655774 | -0.193353409 | 0.43261273 | -0.4469434 | 0.65491592 | 0.97355347 | 1155.42936 | 926.898262 | 858.34409 | 833.951381 | Non-sig. |
| PF3D7_0310500 | DHX57 | 3206.56605 | -0.193412832 | 0.53475981 | -0.36168169 | 0.71758991 | 0.98797939 | 1916.34383 | 2524.80742 | 4925.6831 | 3459.42985 | Non-sig. |
| PF3D7_1369000 | GPAA1 | 927.749666 | -0.194178601 | 0.55269015 | -0.35133357 | 0.72533812 | 0.98844303 | 513.771949 | 911.98912 | 1466.05699 | 819.180604 | Non-sig. |
| PF3D7_0511200 | SCD | 3703.36292 | -0.194187497 | 0.54291405 | -0.35767631 | 0.72058556 | 0.98816139 | 2056.78664 | 3418.04761 | 5847.30088 | 3491.31655 | Non-sig. |
| PF3D7_1450300 | CPR | 405.158492 | -0.194554879 | 0.53011223 | -0.36700696 | 0.71361383 | 0.9875928 | 260.055706 | 356.207866 | 604.496433 | 399.873961 | Non-sig. |
| PF3D7_0417600 | N/A | 289.009752 | -0.194622982 | 0.7194573 | -0.2705136 | 0.78676516 | 0.99028237 | 533.49987 | 302.609749 | 83.7671265 | 236.162264 | Non-sig. |
| PF3D7_0714600 | N/A | 1423.47797 | -0.194656139 | 0.5923236 | -0.32863141 | 0.7424343 | 0.98975488 | 2358.26601 | 1539.85134 | 680.819953 | 1114.97458 | Non-sig. |
| PF3D7_1367500 | N/A | 1815.3101 | -0.195283603 | 0.43147548 | -0.4525949 | 0.65084047 | 0.97317009 | 1666.87569 | 1883.5706 | 2208.92787 | 1501.86623 | Non-sig. |
| PF3D7_1325100 | N/A | 3506.56488 | -0.195747446 | 0.41549269 | -0.47112128 | 0.63755413 | 0.96841129 | 3998.49455 | 3607.20628 | 3489.77791 | 2930.78076 | Non-sig. |

| | | | | | | | | | | | | |
|---|---|---|---|---|---|---|---|---|---|---|---|---|
| PF3D7_1327400 | N/A | 2952.67153 | -0.195921188 | 0.53022673 | -0.36950455 | 0.71175168 | 0.9875928 | 4501.20445 | 3192.06088 | 1804.84991 | 2312.57089 | Non-sig. |
| PF3D7_1206000 | SHLP2 | 6026.38613 | -0.195992223 | 0.41560412 | -0.47158392 | 0.6372238 | 0.96841129 | 6025.77147 | 6269.88913 | 6844.36333 | 4965.52058 | Non-sig. |
| PF3D7_1113200 | N/A | 520.255718 | -0.196144191 | 0.44557602 | -0.44020365 | 0.65978962 | 0.97707593 | 632.619937 | 529.229144 | 478.674557 | 440.499233 | Non-sig. |
| PF3D7_1311900 | vapA | 3505.59194 | -0.196383348 | 0.54606569 | -0.35963319 | 0.71912146 | 0.98797939 | 2085.19582 | 2540.47858 | 5402.10919 | 3994.58415 | Non-sig. |
| PF3D7_1021300 | N/A | 574.945961 | -0.196448685 | 0.5567814 | -0.35282911 | 0.72421656 | 0.98844303 | 322.816615 | 498.010648 | 904.900769 | 574.055812 | Non-sig. |
| PF3D7_0708400 | HSP90 | 27853.7373 | -0.196451334 | 0.41015045 | -0.47897384 | 0.63195724 | 0.9668494 | 33112.1327 | 26019.882 | 26382.5028 | 25900.4316 | Non-sig. |
| PF3D7_0106000 | N/A | 4982.35987 | -0.196818308 | 0.4246896 | -0.46344038 | 0.64304874 | 0.97144917 | 4480.60479 | 4348.33495 | 6162.5816 | 4937.91815 | Non-sig. |
| PF3D7_1110900 | N/A | 1464.60918 | -0.196826568 | 0.47806889 | -0.41171173 | 0.68055073 | 0.97982498 | 2062.25621 | 1333.87807 | 1066.78189 | 1395.52057 | Non-sig. |
| PF3D7_1434300 | HOP | 3441.85992 | -0.198058689 | 0.50288584 | -0.39384423 | 0.69369605 | 0.98334178 | 5108.45028 | 3358.7152 | 2247.39131 | 3052.88288 | Non-sig. |
| PF3D7_1017600 | N/A | 2609.48936 | -0.198140123 | 0.71229059 | -0.27817316 | 0.78087944 | 0.99028237 | 4882.246 | 2838.56494 | 694.898341 | 2022.24815 | Non-sig. |
| PF3D7_0517700 | EIF3B | 3124.44147 | -0.198254712 | 0.45703414 | -0.43378534 | 0.66444432 | 0.97883902 | 2434.52948 | 2997.97778 | 4242.75168 | 2822.50693 | Non-sig. |
| PF3D7_1319000 | N/A | 382.139699 | -0.198311317 | 0.45229535 | -0.43845535 | 0.66105623 | 0.97783983 | 388.716858 | 302.51552 | 427.95204 | 409.374378 | Non-sig. |
| PF3D7_0934300 | ATP8 | 418.045042 | -0.198469866 | 0.4703349 | -0.42197563 | 0.6730428 | 0.97898416 | 344.147195 | 414.547126 | 549.134708 | 364.351139 | Non-sig. |
| PF3D7_1032600 | N/A | 151.327284 | -0.198485636 | 0.52793194 | -0.37596823 | 0.70694052 | 0.986213 | 118.77549 | 129.883369 | 204.450909 | 152.199369 | Non-sig. |
| PF3D7_1352000 | N/A | 1690.78514 | -0.198547752 | 0.45060531 | -0.44062453 | 0.65948484 | 0.97707593 | 2114.08367 | 1295.7273 | 1499.9965 | 1853.33309 | Non-sig. |
| PF3D7_0404200 | N/A | 142.322815 | -0.198620169 | 0.50936183 | -0.38993925 | 0.69658147 | 0.98334178 | 168.878047 | 152.513914 | 135.430036 | 112.469262 | Non-sig. |
| PF3D7_0510000 | N/A | 586.009818 | -0.19866869 | 0.59035053 | -0.33652666 | 0.73647375 | 0.98954916 | 961.361099 | 617.980636 | 291.576401 | 473.121138 | Non-sig. |
| PF3D7_0827900 | PDI8 | 15349.8776 | -0.198711522 | 0.47207912 | -0.42092843 | 0.67380734 | 0.97898416 | 11129.2006 | 14417.5022 | 21681.1485 | 14171.659 | Non-sig. |
| PF3D7_1106100 | N/A | 554.05118 | -0.199143095 | 0.50210675 | -0.39661506 | 0.69165135 | 0.98334178 | 789.193433 | 574.745404 | 395.562056 | 456.703828 | Non-sig. |
| PF3D7_1433200 | N/A | 108.135363 | -0.199272569 | 0.74570875 | -0.26722573 | 0.78929536 | 0.99028237 | 196.970169 | 93.3321698 | 34.4983459 | 107.740765 | Non-sig. |
| PF3D7_0715900 | CDF | 2418.42193 | -0.199305217 | 0.42293778 | -0.47124004 | 0.63746933 | 0.96841129 | 2281.63761 | 2036.36899 | 2888.6272 | 2467.05393 | Non-sig. |
| PF3D7_1351900 | N/A | 1602.04073 | -0.199446168 | 0.56484651 | -0.35309799 | 0.72401498 | 0.98844303 | 2575.10252 | 1693.38114 | 850.457248 | 1289.22201 | Non-sig. |
| PF3D7_0318800 | N/A | 119.860598 | -0.199696018 | 0.60676598 | -0.32911538 | 0.74206848 | 0.98954916 | 185.952569 | 111.651142 | 70.6072993 | 111.231381 | Non-sig. |
| PF3D7_1331700 | N/A | 1283.07302 | -0.199741855 | 0.44364689 | -0.45022711 | 0.65254669 | 0.97317009 | 1088.14954 | 1131.76508 | 1655.11876 | 1257.25871 | Non-sig. |
| PF3D7_0806100 | N/A | 606.355235 | -0.200058137 | 0.50581028 | -0.39552011 | 0.69245909 | 0.98334178 | 883.892007 | 563.371562 | 413.085837 | 565.071532 | Non-sig. |
| PF3D7_0710500 | N/A | 269.158899 | -0.200104343 | 0.46281719 | -0.43236152 | 0.66547868 | 0.97883902 | 257.877451 | 275.002428 | 317.610202 | 226.145517 | Non-sig. |
| PF3D7_0604200 | N/A | 59.1456777 | -0.200481886 | 0.628516 | -0.31897658 | 0.74974427 | 0.99028237 | 45.9572341 | 50.5399453 | 80.3863702 | 59.6991614 | Non-sig. |
| PF3D7_1109100 | N/A | 114.577034 | -0.201089395 | 0.62256636 | -0.32300074 | 0.74669468 | 0.98998709 | 166.447613 | 69.7281456 | 78.855313 | 143.277064 | Non-sig. |
| PF3D7_1245300 | UBC12 | 183.689488 | -0.201139985 | 0.49385503 | -0.40728549 | 0.68379832 | 0.98115715 | 200.968027 | 204.189401 | 192.007905 | 137.592619 | Non-sig. |
| PF3D7_1225300 | N/A | 669.013229 | -0.201450168 | 0.43237269 | -0.46591788 | 0.64127428 | 0.97034716 | 786.78122 | 672.8996 | 644.633177 | 571.738919 | Non-sig. |
| PF3D7_1028200 | N/A | 892.961993 | -0.201562737 | 0.49959921 | -0.40344887 | 0.68661804 | 0.98210107 | 612.830385 | 874.821047 | 1297.33126 | 786.86528 | Non-sig. |
| PF3D7_1112600 | N/A | 942.336478 | -0.201652287 | 0.43691158 | -0.46154027 | 0.64441104 | 0.97190751 | 1174.56334 | 927.093188 | 841.806608 | 825.882778 | Non-sig. |
| PF3D7_0628600 | N/A | 593.860662 | -0.201869916 | 0.4309718 | -0.46840633 | 0.63949404 | 0.96967357 | 701.837979 | 560.391434 | 568.970721 | 544.242514 | Non-sig. |
| PF3D7_0803200 | N/A | 5320.20965 | -0.20190701 | 0.57566739 | -0.35073553 | 0.72578677 | 0.98844303 | 8797.84056 | 4821.81786 | 2586.3122 | 5074.86798 | Non-sig. |
| PF3D7_0830200 | N/A | 33.4395077 | -0.202124475 | 0.75693328 | -0.26703077 | 0.78944547 | 0.99028237 | 22.1215451 | 31.1099708 | 49.2840895 | 31.2424254 | Non-sig. |
| PF3D7_0816400 | N/A | 1699.79768 | -0.202197967 | 0.44310377 | -0.45632193 | 0.6481585 | 0.97317009 | 1572.68925 | 1856.55765 | 2064.60385 | 1305.33998 | Non-sig. |
| PF3D7_1247900 | UTP11 | 214.765684 | -0.202613884 | 0.52387036 | -0.3867634 | 0.69893137 | 0.98377748 | 293.198184 | 236.165531 | 166.674709 | 163.024313 | Non-sig. |
| PF3D7_1469300 | PNO1 | 413.730203 | -0.202847907 | 0.45128116 | -0.44949341 | 0.65307576 | 0.97317009 | 403.493766 | 437.994293 | 481.950824 | 331.481928 | Non-sig. |
| PF3D7_0930600 | CYP72 | 1109.16307 | -0.202918409 | 0.51211178 | -0.39623851 | 0.69192908 | 0.98334178 | 1644.27145 | 1142.71172 | 730.136818 | 919.532287 | Non-sig. |
| PF3D7_0621300 | PUF3 | 481.695808 | -0.20320375 | 0.45181085 | -0.44975402 | 0.65288781 | 0.97317009 | 609.947109 | 454.844083 | 421.381917 | 440.610124 | Non-sig. |
| PF3D7_0208500 | ACP | 635.892743 | -0.203391172 | 0.45029976 | -0.45167951 | 0.65149988 | 0.97317009 | 553.081039 | 636.472158 | 808.001046 | 546.01673 | Non-sig. |
| PF3D7_0921100 | N/A | 48.0444495 | -0.203492075 | 0.64740145 | -0.31432132 | 0.75327701 | 0.99028237 | 42.0044004 | 39.7270266 | 60.7503414 | 49.6960297 | Non-sig. |
| PF3D7_0519900 | N/A | 383.998509 | -0.203568825 | 0.44789832 | -0.45449785 | 0.64947054 | 0.97317009 | 357.911496 | 371.246167 | 464.036003 | 342.800368 | Non-sig. |
| PF3D7_1221800 | N/A | 1728.3186 | -0.203711751 | 0.51518962 | -0.39541121 | 0.69253943 | 0.98334178 | 2611.51465 | 1658.6503 | 1089.11041 | 1553.99905 | Non-sig. |
| PF3D7_1434100 | N/A | 174.775049 | -0.203928711 | 0.49321717 | -0.41346637 | 0.67926496 | 0.97898416 | 155.631877 | 153.479809 | 218.426028 | 171.562482 | Non-sig. |
| PF3D7_0907900 | PDF | 1743.58265 | -0.205289953 | 0.42394932 | -0.48423229 | 0.62822102 | 0.96655174 | 1837.9374 | 1855.228 | 1896.90547 | 1384.25973 | Non-sig. |
| PF3D7_1113000 | N/A | 3640.15168 | -0.205543483 | 0.4764756 | -0.43138302 | 0.66618989 | 0.97883902 | 3036.07708 | 2547.01462 | 4761.74932 | 4215.76572 | Non-sig. |
| PF3D7_1027200 | N/A | 291.628155 | -0.206061629 | 0.49994341 | -0.41216991 | 0.68021489 | 0.97982498 | 222.136623 | 287.5472 | 402.444937 | 254.383862 | Non-sig. |
| PF3D7_0525900 | NEK2 | 48.2625612 | -0.206301 | 0.65508823 | -0.31492094 | 0.75282168 | 0.99028237 | 64.976338 | 44.0466184 | 38.5710115 | 45.456277 | Non-sig. |
| PF3D7_0109500 | N/A | 487.718345 | -0.206308937 | 0.44175645 | -0.46701964 | 0.64048583 | 0.97002976 | 463.786278 | 490.790021 | 581.153004 | 415.144076 | Non-sig. |
| PF3D7_0726200 | FIKK7.1 | 222.942012 | -0.206429547 | 0.46905551 | -0.4400962 | 0.65986743 | 0.97707593 | 248.598554 | 182.369926 | 229.170136 | 231.629434 | Non-sig. |
| PF3D7_0601500 | N/A | 68.3771922 | -0.206610986 | 0.5854707 | -0.35289722 | 0.72416549 | 0.98844303 | 84.2114575 | 69.7434789 | 62.4186121 | 57.1352204 | Non-sig. |
| PF3D7_1313700 | N/A | 108.478827 | -0.206628335 | 0.600103 | -0.34432145 | 0.73060455 | 0.98844303 | 68.0936949 | 98.8471549 | 164.120579 | 102.853878 | Non-sig. |
| PF3D7_1435100 | N/A | 293.417003 | -0.206694218 | 0.45692173 | -0.45236241 | 0.65100792 | 0.97317009 | 342.379985 | 298.478751 | 286.518244 | 246.291034 | Non-sig. |

| | | | | | | | | | | | | |
|---|---|---|---|---|---|---|---|---|---|---|---|---|
| PF3D7_1031700 | I3 | 931.855107 | -0.207520815 | 0.45025889 | -0.46089221 | 0.64487594 | 0.97190751 | 776.645938 | 882.723081 | 1220.63293 | 847.418475 | Non-sig. |
| PF3D7_1032500 | Der1-2 | 1145.89141 | -0.207532255 | 0.5072332 | -0.40914565 | 0.68243278 | 0.98088141 | 752.931911 | 1053.96226 | 1703.07011 | 1073.60136 | Non-sig. |
| PF3D7_1203500 | SUA5 | 857.371642 | -0.207945538 | 0.4446665 | -0.46764382 | 0.64003932 | 0.96994518 | 1085.3096 | 863.973978 | 753.006268 | 727.196722 | Non-sig. |
| PF3D7_0928200 | N/A | 782.029688 | -0.208210701 | 0.46422573 | -0.44851176 | 0.6537839 | 0.97317009 | 620.208407 | 720.974331 | 1056.24857 | 730.687441 | Non-sig. |
| PF3D7_1206200 | EIF3C | 4777.96772 | -0.208244696 | 0.42980721 | -0.48450722 | 0.62802594 | 0.96655174 | 6035.68602 | 4719.97545 | 4208.9485 | 4147.26092 | Non-sig. |
| PF3D7_0707400 | N/A | 961.489027 | -0.20867492 | 0.42012661 | -0.49669532 | 0.61940393 | 0.96395246 | 1119.05457 | 917.373118 | 942.863521 | 866.664901 | Non-sig. |
| PF3D7_1362200 | RUVB3 | 661.999927 | -0.20871539 | 0.42188079 | -0.49472598 | 0.62079356 | 0.9639908 | 678.774147 | 605.204125 | 740.775897 | 623.24554 | Non-sig. |
| PF3D7_0311300 | N/A | 6969.85424 | -0.208824739 | 0.45207461 | -0.46192539 | 0.64413483 | 0.97190751 | 9168.12149 | 5564.87135 | 5778.9419 | 7367.48221 | Non-sig. |
| PF3D7_1247200 | N/A | 277.015269 | -0.208840019 | 0.48148828 | -0.43373853 | 0.66447832 | 0.97883902 | 245.143262 | 295.465027 | 348.757734 | 218.695054 | Non-sig. |
| PF3D7_1427200 | Sel4 | 384.663855 | -0.209222648 | 0.43644453 | -0.4793797 | 0.63166853 | 0.9668494 | 393.075638 | 358.531209 | 431.884625 | 355.163948 | Non-sig. |
| PF3D7_1311600 | N/A | 2652.73533 | -0.209277706 | 0.40614612 | -0.51527688 | 0.60635955 | 0.95706907 | 2971.79532 | 2473.99437 | 2717.87096 | 2447.28066 | Non-sig. |
| PF3D7_0618100 | N/A | 1277.55603 | -0.20939578 | 0.43461431 | -0.48179679 | 0.62995031 | 0.96682409 | 1598.82465 | 1105.1932 | 1141.58605 | 1264.62022 | Non-sig. |
| PF3D7_0911400 | N/A | 769.089809 | -0.209476728 | 0.43242067 | -0.48442811 | 0.62808208 | 0.96655174 | 742.696009 | 641.49832 | 906.836642 | 785.328266 | Non-sig. |
| PF3D7_0406600 | N/A | 6.41959553 | -0.209931039 | 1.42859877 | -0.14694891 | 0.88317234 | 0.99457954 | 3.08869627 | 7.56101983 | 10.6297787 | 4.39888728 | Non-sig. |
| PF3D7_1121000 | DHHC3 | 681.732894 | -0.209973276 | 0.46971081 | -0.44702671 | 0.65485578 | 0.97355347 | 905.522268 | 715.478884 | 557.232982 | 548.697442 | Non-sig. |
| PF3D7_1307100 | UTP6 | 869.00306 | -0.211254861 | 0.43817465 | -0.4821248 | 0.6297173 | 0.96682409 | 1065.97825 | 723.051911 | 799.214561 | 887.767516 | Non-sig. |
| PF3D7_1447900 | MDR2 | 1406.36653 | -0.211579586 | 0.61311376 | -0.34509026 | 0.73002652 | 0.98844303 | 612.235595 | 1116.63477 | 2406.02618 | 1490.56958 | Non-sig. |
| PF3D7_0910500 | N/A | 2460.23158 | -0.211630753 | 0.48919161 | -0.43261321 | 0.66529579 | 0.97883902 | 2638.87913 | 1469.49783 | 2641.8275 | 3090.72185 | Non-sig. |
| PF3D7_1324100 | N/A | 1473.42509 | -0.211951645 | 0.42133291 | -0.50305029 | 0.61492891 | 0.96217474 | 1388.04083 | 1408.93121 | 1774.73556 | 1321.99277 | Non-sig. |
| PF3D7_1013600 | N/A | 3295.68989 | -0.211999864 | 0.41042831 | -0.51653324 | 0.60548203 | 0.95693578 | 3749.46376 | 2837.98452 | 3325.41879 | 3269.89248 | Non-sig. |
| PF3D7_1136300 | TSN | 7590.59802 | -0.212011476 | 0.48722085 | -0.43514451 | 0.66345754 | 0.97883902 | 11040.367 | 7374.88584 | 5254.6477 | 6692.49155 | Non-sig. |
| PF3D7_1450700 | N/A | 2270.45018 | -0.212114992 | 0.61684822 | -0.34386902 | 0.73094479 | 0.98865088 | 3923.49412 | 2335.10041 | 950.985739 | 1872.22045 | Non-sig. |
| PF3D7_1223200 | N/A | 1284.90735 | -0.21220244 | 0.50814319 | -0.41760363 | 0.67623695 | 0.97898416 | 852.113007 | 1080.67571 | 1906.02506 | 1300.81564 | Non-sig. |
| PF3D7_0108700 | PSOP24 | 4257.8144 | -0.212245676 | 0.43806058 | -0.48451216 | 0.62802244 | 0.96655174 | 4982.16089 | 3162.38228 | 4158.843 | 4727.87145 | Non-sig. |
| PF3D7_1144600 | TFG2 | 1137.79627 | -0.212429048 | 0.42130731 | -0.50421401 | 0.614111 | 0.96217474 | 1288.47725 | 1157.18923 | 1154.42073 | 951.097858 | Non-sig. |
| PF3D7_1202000 | PSH2 | 336.563162 | -0.212725848 | 0.54305355 | -0.39172168 | 0.69526387 | 0.98334178 | 214.776348 | 273.595722 | 507.577202 | 350.303374 | Non-sig. |
| PF3D7_0314200 | N/A | 626.600641 | -0.212776856 | 0.44341604 | -0.47985828 | 0.63132817 | 0.9668494 | 565.741282 | 539.580492 | 779.524852 | 621.55594 | Non-sig. |
| PF3D7_1220100 | PRP17 | 991.526777 | -0.213519608 | 0.41967271 | -0.50877649 | 0.6109089 | 0.9597882 | 1034.71661 | 996.803845 | 1094.78375 | 839.802906 | Non-sig. |
| PF3D7_1420400 | GlyRS | 2117.60282 | -0.213849043 | 0.43069596 | -0.49651973 | 0.61952777 | 0.96395246 | 1883.86081 | 2067.41522 | 2664.45767 | 1854.6776 | Non-sig. |
| PF3D7_0629100 | NAPRT | 1101.1298 | -0.213949798 | 0.50249377 | -0.42577602 | 0.67027106 | 0.97883902 | 749.451836 | 934.57319 | 1615.48089 | 1105.0133 | Non-sig. |
| PF3D7_1446400 | pdhB | 205.402676 | -0.213999476 | 0.51044068 | -0.41924455 | 0.67503742 | 0.97898416 | 164.221019 | 212.446208 | 276.779822 | 168.163654 | Non-sig. |
| PF3D7_1474800 | N/A | 2320.2532 | -0.214627427 | 0.42552901 | -0.5043779 | 0.61399584 | 0.96217474 | 2154.62797 | 1959.02266 | 2830.25603 | 2337.10614 | Non-sig. |
| PF3D7_0911500 | N/A | 33.2334633 | -0.214743019 | 0.74114771 | -0.28974389 | 0.77201218 | 0.99028237 | 46.5711582 | 33.2241152 | 25.0023852 | 28.1361945 | Non-sig. |
| PF3D7_0517800 | DCX | 25.1305141 | -0.215289656 | 0.82836023 | -0.25989859 | 0.794942 | 0.99028237 | 34.4150804 | 30.1216933 | 19.6815604 | 16.3037224 | Non-sig. |
| PF3D7_1148100 | N/A | 27.2959277 | -0.215630357 | 0.80088029 | -0.26924168 | 0.7877437 | 0.99028237 | 19.9625667 | 20.397358 | 38.5882389 | 30.2355472 | Non-sig. |
| PF3D7_0716700 | N/A | 344.526544 | -0.215664284 | 0.48907958 | -0.44095949 | 0.65924232 | 0.97707593 | 273.710235 | 285.405028 | 466.503844 | 352.487069 | Non-sig. |
| PF3D7_1359200 | HMGB4 | 998.768982 | -0.215680609 | 0.44869497 | -0.48068425 | 0.63074092 | 0.9668494 | 1241.243 | 1070.47917 | 905.510132 | 777.843624 | Non-sig. |
| PF3D7_0306000 | QCR8 | 1097.21288 | -0.215864981 | 0.45834828 | -0.47096278 | 0.63766731 | 0.96841129 | 889.851727 | 1090.38972 | 1468.1882 | 940.421875 | Non-sig. |
| PF3D7_0930400 | N/A | 3191.75526 | -0.216319577 | 0.63776864 | -0.33918189 | 0.73447271 | 0.98909017 | 5650.47051 | 3285.42313 | 1211.20123 | 2619.92617 | Non-sig. |
| PF3D7_0728000 | eIF2alpha | 2758.60261 | -0.21634441 | 0.44892502 | -0.48191658 | 0.62986521 | 0.96682409 | 2235.51578 | 2498.64452 | 3694.32268 | 2605.92747 | Non-sig. |
| PF3D7_1126100 | ATG7 | 654.396606 | -0.216597436 | 0.5572867 | -0.38866428 | 0.69752451 | 0.98334178 | 392.924482 | 472.012098 | 1013.61111 | 739.038739 | Non-sig. |
| PF3D7_0831000 | GEXP09 | 31.9785452 | -0.216607586 | 0.71636102 | -0.3023721 | 0.76236842 | 0.99028237 | 32.5967197 | 30.0524698 | 36.1302619 | 29.1347294 | Non-sig. |
| PF3D7_1126300 | N/A | 619.928174 | -0.216614593 | 0.50730121 | -0.42699404 | 0.66938367 | 0.97883902 | 561.553544 | 379.939808 | 771.064371 | 767.154973 | Non-sig. |
| PF3D7_0605200 | N/A | 617.46469 | -0.217177331 | 0.44618458 | -0.48674325 | 0.62644029 | 0.96655174 | 607.891012 | 664.981838 | 719.728694 | 477.257215 | Non-sig. |
| PF3D7_1226400 | N/A | 1262.82213 | -0.218870504 | 0.42727719 | -0.51224477 | 0.60847971 | 0.9580165 | 1266.35696 | 1012.88379 | 1450.41687 | 1321.6309 | Non-sig. |
| PF3D7_1013500 | PI-PLC | 2122.51439 | -0.219404015 | 0.41989906 | -0.52251609 | 0.60131105 | 0.95693578 | 2346.98422 | 1705.68591 | 2220.24718 | 2217.14026 | Non-sig. |
| PF3D7_1006100 | NOT5 | 195.13238 | -0.22006602 | 0.46738001 | -0.4708503 | 0.63774764 | 0.96841129 | 208.123052 | 185.700221 | 211.841267 | 174.86498 | Non-sig. |
| PF3D7_0508000 | P38 | 7547.0996 | -0.220204231 | 0.46277537 | -0.47583395 | 0.63419268 | 0.9668494 | 5861.73485 | 7679.24571 | 10381.8992 | 6265.5186 | Non-sig. |
| PF3D7_0321800 | N/A | 6580.88455 | -0.220325897 | 0.43177103 | -0.51028411 | 0.60985243 | 0.95903775 | 6878.81402 | 4881.05346 | 7285.99391 | 7277.67682 | Non-sig. |
| PF3D7_1251500 | DRS1 | 630.560757 | -0.220334327 | 0.45650067 | -0.48265937 | 0.62933762 | 0.96682409 | 829.391049 | 596.593566 | 528.084095 | 568.174319 | Non-sig. |
| PF3D7_0716900 | DMT2 | 813.065336 | -0.220417475 | 0.48905362 | -0.45070206 | 0.6522043 | 0.97317009 | 585.221176 | 768.269139 | 1164.58305 | 734.187984 | Non-sig. |
| PF3D7_1201600 | NEK3 | 121.997074 | -0.220793976 | 0.57010767 | -0.3872847 | 0.69854545 | 0.98377748 | 173.352438 | 93.3994417 | 89.491266 | 131.745151 | Non-sig. |
| PF3D7_1330200 | N/A | 924.276355 | -0.221859459 | 0.42672174 | -0.519916 | 0.60312212 | 0.95693578 | 1056.77061 | 947.972269 | 933.719557 | 758.642987 | Non-sig. |
| PF3D7_0723800 | N/A | 12624.2752 | -0.221925582 | 0.50108521 | -0.44288991 | 0.65784537 | 0.97671961 | 18084.0097 | 9042.76264 | 9102.98125 | 14267.347 | Non-sig. |

| | | | | | | | | | | | | |
|---|---|---|---|---|---|---|---|---|---|---|---|---|
| PF3D7_1318700 | AAP4 | 2962.83471 | -0.221979885 | 0.571841 | -0.38818463 | 0.69787941 | 0.98334178 | 4778.77097 | 2108.68869 | 1602.21464 | 3361.66456 | Non-sig. |
| PF3D7_0904900 | CuTP | 10831.929 | -0.222318147 | 0.42263525 | -0.52602841 | 0.59886847 | 0.95655566 | 10884.6307 | 8460.11531 | 12445.0452 | 11537.9247 | Non-sig. |
| PF3D7_1352800 | MON1 | 1021.66506 | -0.222366239 | 0.41441173 | -0.53658288 | 0.59155578 | 0.95321031 | 1064.53569 | 916.2461 | 1135.91289 | 969.96558 | Non-sig. |
| PF3D7_1471400 | N/A | 3986.97659 | -0.222495306 | 0.40855318 | -0.54459325 | 0.5860333 | 0.95246063 | 4116.32538 | 3408.73238 | 4471.2368 | 3951.61181 | Non-sig. |
| PF3D7_0933900 | N/A | 823.229118 | -0.222611269 | 0.41747988 | -0.53322634 | 0.59387691 | 0.9542761 | 892.397595 | 790.294415 | 880.845903 | 729.378558 | Non-sig. |
| PF3D7_1127700 | N/A | 28.6746087 | -0.222929446 | 0.79541276 | -0.28026888 | 0.77927122 | 0.99028237 | 43.0300035 | 26.8468755 | 18.8799689 | 25.9415868 | Non-sig. |
| PF3D7_1406800 | GAPM3 | 12192.5297 | -0.223274511 | 0.46898112 | -0.47608422 | 0.63401438 | 0.9668494 | 9045.07699 | 10860.4102 | 17222.8641 | 11641.7673 | Non-sig. |
| PF3D7_0811600 | N/A | 3320.91777 | -0.223563082 | 0.63897515 | -0.34987759 | 0.72643057 | 0.98844303 | 1301.87802 | 2445.95766 | 5853.20283 | 3682.63258 | Non-sig. |
| PF3D7_1241200 | N/A | 2774.03851 | -0.223876729 | 0.45484292 | -0.49220669 | 0.62257323 | 0.96429926 | 2287.30342 | 2237.38371 | 3690.13192 | 2881.33497 | Non-sig. |
| PF3D7_1004500 | N/A | 2686.97987 | -0.224101463 | 0.40855545 | -0.54852153 | 0.58333385 | 0.95129458 | 2688.89172 | 2473.89718 | 3101.52029 | 2483.61027 | Non-sig. |
| PF3D7_1211400 | PfJ4 | 3525.10786 | -0.224861569 | 0.41404583 | -0.54308377 | 0.58707212 | 0.95246063 | 3656.69843 | 3607.98243 | 3941.79662 | 2893.95394 | Non-sig. |
| PF3D7_0112200 | MRP1 | 4012.39257 | -0.225064724 | 0.63561395 | -0.35409029 | 0.72327122 | 0.98844303 | 1516.87026 | 3525.48787 | 7132.24136 | 3874.97078 | Non-sig. |
| PF3D7_0318100 | STOML | 270.884656 | -0.225200113 | 0.53942721 | -0.41748008 | 0.67632729 | 0.97898416 | 177.350005 | 259.884978 | 406.322667 | 239.980974 | Non-sig. |
| PF3D7_1322300 | N/A | 1177.04595 | -0.22525613 | 0.42944453 | -0.52452904 | 0.59991063 | 0.95655566 | 1459.1553 | 1044.82827 | 1078.52422 | 1125.67602 | Non-sig. |
| PF3D7_0620100 | ATPTG12 | 257.145936 | -0.225535136 | 0.4532821 | -0.49756021 | 0.61879406 | 0.96395246 | 273.701442 | 249.195482 | 280.69688 | 224.989939 | Non-sig. |
| PF3D7_1466800 | N/A | 1027.32047 | -0.225622539 | 0.41631849 | -0.54194695 | 0.58785505 | 0.95246063 | 1035.38354 | 935.474004 | 1179.50961 | 958.914722 | Non-sig. |
| PF3D7_0621000 | N/A | 620.30596 | -0.225811776 | 0.45089432 | -0.50080865 | 0.6165058 | 0.96358724 | 768.496862 | 492.484121 | 569.167897 | 651.074959 | Non-sig. |
| PF3D7_0727600 | N/A | 142.295582 | -0.22681633 | 0.50056694 | -0.45311888 | 0.65046314 | 0.97317009 | 132.671567 | 128.746671 | 174.123907 | 133.640184 | Non-sig. |
| PF3D7_1464500 | N/A | 7725.56175 | -0.226940474 | 0.47814132 | -0.47463055 | 0.63505033 | 0.9668494 | 6051.05528 | 5685.21782 | 10612.5606 | 8553.4133 | Non-sig. |
| PF3D7_1351300 | N/A | 12.8813031 | -0.227580194 | 1.01039834 | -0.22523809 | 0.82179406 | 0.99096399 | 14.6719699 | 11.8174094 | 13.1305402 | 11.9052928 | Non-sig. |
| PF3D7_0510100 | N/A | 12443.8972 | -0.228138319 | 0.45608692 | -0.50020799 | 0.61692864 | 0.96358724 | 11875.1345 | 8546.86682 | 14976.249 | 14377.3386 | Non-sig. |
| PF3D7_0416900 | N/A | 1468.77522 | -0.228145819 | 0.44403678 | -0.51379937 | 0.60739227 | 0.95715771 | 1824.71016 | 1136.9014 | 1344.79637 | 1568.69296 | Non-sig. |
| PF3D7_1460800 | SNAPC3 | 279.389568 | -0.22890257 | 0.45961476 | -0.49803137 | 0.61846193 | 0.96395246 | 339.571294 | 268.340167 | 263.57777 | 246.06904 | Non-sig. |
| PF3D7_1348400 | N/A | 1811.80097 | -0.230758868 | 0.44792581 | -0.51517207 | 0.60643278 | 0.95706907 | 2101.38023 | 1291.26417 | 1811.48158 | 2043.07788 | Non-sig. |
| PF3D7_0732000 | N/A | 5.84756483 | -0.230961624 | 1.44613254 | -0.15970986 | 0.87310963 | 0.99402081 | 5.22925014 | 7.52452913 | 7.38390986 | 3.25257019 | Non-sig. |
| PF3D7_1353300 | N/A | 4070.55697 | -0.230969911 | 0.4642381 | -0.49752468 | 0.6188191 | 0.96395246 | 5569.03868 | 3255.3523 | 3222.64405 | 4235.19285 | Non-sig. |
| PF3D7_1142400 | CPO | 1722.38503 | -0.231001808 | 0.49265117 | -0.46889528 | 0.63914449 | 0.96967357 | 1192.83078 | 1557.2071 | 2526.81146 | 1612.69079 | Non-sig. |
| PF3D7_0811400 | N/A | 4476.58065 | -0.231099273 | 0.42179376 | -0.54789637 | 0.58376306 | 0.95151408 | 4279.47774 | 4562.59256 | 5389.10221 | 3675.15011 | Non-sig. |
| PF3D7_1411800 | N/A | 266.087927 | -0.231752033 | 0.56033061 | -0.41359874 | 0.679168 | 0.97898416 | 405.123176 | 289.732738 | 169.997992 | 199.497803 | Non-sig. |
| PF3D7_0906890 | N/A | 53.319072 | -0.231786563 | 0.64160134 | -0.36126259 | 0.71790315 | 0.98797939 | 42.8606687 | 53.7530786 | 72.1929008 | 44.4696399 | Non-sig. |
| PF3D7_1022100 | N/A | 280.562867 | -0.23199523 | 0.47465172 | -0.48876939 | 0.62500497 | 0.96566822 | 356.783969 | 284.588115 | 249.537744 | 231.34164 | Non-sig. |
| PF3D7_0906100 | VPS46 | 971.551567 | -0.23221867 | 0.43359028 | -0.53557167 | 0.59225461 | 0.95322751 | 1138.57419 | 1018.04397 | 960.680638 | 768.90747 | Non-sig. |
| PF3D7_0935000 | SNRPB2 | 1139.66581 | -0.232804236 | 0.41783296 | -0.55717059 | 0.57741088 | 0.9497432 | 1167.12744 | 1124.10352 | 1295.65129 | 971.781006 | Non-sig. |
| PF3D7_1303600 | N/A | 495.700763 | -0.23316626 | 0.43918285 | -0.5309093 | 0.59548163 | 0.95502327 | 467.902441 | 465.038071 | 603.297188 | 446.565354 | Non-sig. |
| PF3D7_1111800 | N/A | 465.681358 | -0.233495363 | 0.48217143 | -0.48425798 | 0.6282028 | 0.96655174 | 630.000786 | 486.401785 | 376.80944 | 369.513423 | Non-sig. |
| PF3D7_1414700 | N/A | 309.500088 | -0.233692537 | 0.48728115 | -0.4795846 | 0.6315228 | 0.9668494 | 408.833813 | 243.497205 | 260.396698 | 325.272635 | Non-sig. |
| PF3D7_0713800 | NELFA | 3020.55128 | -0.234303808 | 0.52644038 | -0.44507187 | 0.65626781 | 0.97501595 | 4594.21744 | 3310.01799 | 1936.69924 | 2241.27043 | Non-sig. |
| PF3D7_1366200 | N/A | 606.453842 | -0.234599888 | 0.42566782 | -0.55113371 | 0.58154202 | 0.95129458 | 634.19807 | 590.828854 | 677.073607 | 523.714836 | Non-sig. |
| PF3D7_1131200 | N/A | 609.091421 | -0.234806452 | 0.55240541 | -0.42506182 | 0.67079161 | 0.97898416 | 937.267715 | 687.052894 | 380.140746 | 431.90433 | Non-sig. |
| PF3D7_0805700 | FIKK8 | 5606.9217 | -0.234901451 | 0.4509487 | -0.52090505 | 0.60243292 | 0.95693578 | 4557.98295 | 4744.22115 | 7566.50711 | 5558.97559 | Non-sig. |
| PF3D7_0213300 | N/A | 60.5699217 | -0.235207067 | 0.59043594 | -0.3983617 | 0.69036359 | 0.98334178 | 63.7124339 | 52.6907544 | 67.2619113 | 58.6145871 | Non-sig. |
| PF3D7_0115400 | N/A | 14.6120525 | -0.235629817 | 1.09126063 | -0.21592442 | 0.82904666 | 0.99110542 | 5.2301975 | 16.1231704 | 26.2546715 | 10.8401707 | Non-sig. |
| PF3D7_1342400 | CK2beta2 | 3222.86587 | -0.235744044 | 0.43838763 | -0.53775249 | 0.59074793 | 0.95315309 | 2738.84323 | 3014.03288 | 4232.10986 | 2906.47752 | Non-sig. |
| PF3D7_0727800 | N/A | 2900.23985 | -0.236789128 | 0.52202151 | -0.45360033 | 0.65011652 | 0.97317009 | 1850.66126 | 2254.63496 | 4424.43448 | 3071.22868 | Non-sig. |
| PF3D7_0107700 | N/A | 161.866934 | -0.236923162 | 0.59575283 | -0.39768701 | 0.69086091 | 0.98334178 | 95.5043485 | 127.791604 | 254.46855 | 169.703234 | Non-sig. |
| PF3D7_1419500 | N/A | 156.695657 | -0.236961221 | 0.58367514 | -0.40598135 | 0.6847563 | 0.98115715 | 235.076034 | 176.118597 | 104.249112 | 111.338887 | Non-sig. |
| PF3D7_0710100 | N/A | 1040.07441 | -0.237223329 | 0.4573178 | -0.51872753 | 0.60395076 | 0.95693578 | 1382.2032 | 1056.01732 | 868.826167 | 853.250964 | Non-sig. |
| PF3D7_1245500 | N/A | 1460.6184 | -0.237386331 | 0.57350578 | -0.41392143 | 0.67893165 | 0.97898416 | 2411.87059 | 1474.23462 | 749.515163 | 1206.85321 | Non-sig. |
| PF3D7_1320300 | N/A | 735.550868 | -0.237567452 | 0.44631492 | -0.5322866 | 0.59452751 | 0.9542761 | 641.621839 | 652.510006 | 950.114214 | 697.957414 | Non-sig. |
| PF3D7_1448400 | HRD3 | 1773.01201 | -0.237596964 | 0.51533028 | -0.46105765 | 0.64475725 | 0.97190751 | 1135.78343 | 1510.75801 | 2701.23984 | 1744.26678 | Non-sig. |
| PF3D7_0921600 | N/A | 776.542737 | -0.237628947 | 0.42444213 | -0.55986183 | 0.57557369 | 0.94896603 | 834.596125 | 643.58988 | 846.096157 | 781.888786 | Non-sig. |
| PF3D7_0902800 | SERA9 | 3478.99319 | -0.237706633 | 0.54507518 | -0.4360988 | 0.66276505 | 0.97883902 | 1945.11287 | 3265.72448 | 5584.44353 | 3120.69187 | Non-sig. |
| PF3D7_1428300 | N/A | 2114.36969 | -0.237807046 | 0.41156701 | -0.57780881 | 0.56339321 | 0.94603921 | 2342.81774 | 2096.9533 | 2233.69277 | 1784.01493 | Non-sig. |
| PF3D7_1234500 | N/A | 349.545075 | -0.238065175 | 0.44228514 | -0.53826176 | 0.59039635 | 0.95315309 | 403.645574 | 331.681211 | 353.070771 | 309.782745 | Non-sig. |

| | | | | | | | | | | | | |
|---|---|---|---|---|---|---|---|---|---|---|---|---|
| PF3D7_1239600 | ThzK | 429.748033 | -0.238674626 | 0.51477529 | -0.46364818 | 0.64289983 | 0.97144917 | 303.686674 | 445.502479 | 626.459615 | 343.343365 | Non-sig. |
| PF3D7_0506000 | N/A | 109.789521 | -0.239000863 | 0.52199643 | -0.45785919 | 0.64705361 | 0.97317009 | 103.837429 | 102.035841 | 133.786904 | 99.4979094 | Non-sig. |
| PF3D7_1225600 | N/A | 1129.23996 | -0.239659383 | 0.56313696 | -0.42557921 | 0.67041449 | 0.97883902 | 1839.04856 | 1116.95482 | 606.943536 | 954.012905 | Non-sig. |
| PF3D7_1457000 | SPP | 6348.50014 | -0.239968239 | 0.48638809 | -0.49336783 | 0.62175271 | 0.96429926 | 4444.66728 | 6024.06647 | 9305.55106 | 5619.71577 | Non-sig. |
| PF3D7_1250800 | N/A | 3096.70058 | -0.240085515 | 0.40767379 | -0.58891574 | 0.5559178 | 0.94243456 | 3363.45923 | 2641.98425 | 3344.09604 | 3037.2628 | Non-sig. |
| PF3D7_1324400 | PRELID | 368.553416 | -0.240144147 | 0.46806756 | -0.51305446 | 0.60791323 | 0.95769381 | 477.544849 | 306.051366 | 320.961458 | 369.655991 | Non-sig. |
| PF3D7_1314100 | N/A | 389.082397 | -0.240215144 | 0.57537722 | -0.41749158 | 0.67631888 | 0.97898416 | 211.666978 | 378.717584 | 630.801101 | 335.143924 | Non-sig. |
| PF3D7_1448000 | UTP12 | 615.44977 | -0.240251231 | 0.55835881 | -0.43028108 | 0.66699119 | 0.97883902 | 356.13472 | 480.563802 | 976.686134 | 648.414425 | Non-sig. |
| PF3D7_0622300 | N/A | 168.349188 | -0.24050246 | 0.48004116 | -0.50100383 | 0.61636843 | 0.96358724 | 194.753542 | 158.797199 | 170.010807 | 149.835202 | Non-sig. |
| PF3D7_0729300 | NMD3 | 649.45438 | -0.241641499 | 0.44478921 | -0.54327194 | 0.58694258 | 0.95246063 | 824.475538 | 631.928015 | 583.152859 | 558.261107 | Non-sig. |
| PF3D7_1408000 | PMII | 1158.7841 | -0.241894524 | 0.52318777 | -0.46234743 | 0.64383219 | 0.97190751 | 786.361972 | 832.618672 | 1724.73917 | 1291.41658 | Non-sig. |
| PF3D7_0912600 | N/A | 75.2312321 | -0.242401966 | 0.58488515 | -0.4144437 | 0.67854919 | 0.97898416 | 99.9813272 | 63.3091422 | 63.2377233 | 74.3967357 | Non-sig. |
| PF3D7_1430000 | N/A | 497.492135 | -0.242994746 | 0.59648059 | -0.40738081 | 0.68372832 | 0.98115715 | 827.639096 | 531.266143 | 251.275031 | 379.788271 | Non-sig. |
| PF3D7_1310600 | RAB5b | 2537.49751 | -0.243303091 | 0.41998411 | -0.57931499 | 0.56237665 | 0.94519828 | 2995.30692 | 2571.99178 | 2506.72902 | 2075.96233 | Non-sig. |
| PF3D7_0725100 | N/A | 3835.29182 | -0.243436402 | 0.4306629 | -0.56525975 | 0.5718971 | 0.94850402 | 3492.40761 | 3148.1838 | 4823.61063 | 3876.96524 | Non-sig. |
| PF3D7_0313500 | PfG | 365.399043 | -0.243905024 | 0.49507704 | -0.49266075 | 0.62225232 | 0.96429926 | 515.910986 | 328.310099 | 276.78812 | 340.586967 | Non-sig. |
| PF3D7_1225100 | api-IRS | 278.204737 | -0.243921208 | 0.4801762 | -0.50798271 | 0.61146547 | 0.9597882 | 250.772877 | 216.681944 | 352.396433 | 292.967694 | Non-sig. |
| PF3D7_0815900 | aLipDH | 204.573833 | -0.244491178 | 0.48001378 | -0.509342 | 0.61051253 | 0.95950619 | 256.653456 | 192.103104 | 187.231408 | 182.307362 | Non-sig. |
| PF3D7_1237700 | N/A | 8327.90352 | -0.244524504 | 0.49476466 | -0.49422387 | 0.62114808 | 0.9639908 | 5654.66271 | 7983.59831 | 12408.9586 | 7264.39443 | Non-sig. |
| PF3D7_0920900 | PRP24 | 838.994505 | -0.244624888 | 0.47858791 | -0.51113888 | 0.60925381 | 0.95866549 | 1147.60792 | 892.897167 | 672.558042 | 642.914894 | Non-sig. |
| PF3D7_1007500 | N/A | 233.761981 | -0.244865498 | 0.4981345 | -0.49156502 | 0.62302687 | 0.96429926 | 188.716726 | 223.776312 | 318.171351 | 204.383536 | Non-sig. |
| PF3D7_0932300 | M18AAP | 934.584676 | -0.245120807 | 0.54587161 | -0.4490448 | 0.65339934 | 0.97317009 | 539.911018 | 893.833725 | 1487.32411 | 817.269849 | Non-sig. |
| PF3D7_0517400 | FACT-L | 4690.07148 | -0.245457118 | 0.40692286 | -0.60320307 | 0.54637361 | 0.94142878 | 5303.00873 | 4020.88657 | 4873.23163 | 4563.15901 | Non-sig. |
| PF3D7_1305100 | AMR3 | 1280.95901 | -0.246082505 | 0.44488281 | -0.55314006 | 0.58016751 | 0.95129458 | 1645.19733 | 1310.35895 | 1134.89607 | 1033.38367 | Non-sig. |
| PF3D7_1230600 | NSUN4 | 405.672812 | -0.246091464 | 0.44982575 | -0.54708177 | 0.58432256 | 0.95151408 | 378.515471 | 394.983259 | 501.711662 | 347.480855 | Non-sig. |
| PF3D7_0419700 | Pf34 | 5010.40072 | -0.246376239 | 0.42377788 | -0.58138061 | 0.56098396 | 0.94472957 | 6264.94892 | 4895.46616 | 4609.62292 | 4271.56489 | Non-sig. |
| PF3D7_1308800 | INT | 2380.33718 | -0.246905315 | 0.42885445 | -0.5757322 | 0.5647962 | 0.94662774 | 2899.91208 | 2452.28052 | 2267.29822 | 1901.85789 | Non-sig. |
| PF3D7_0804400 | METAP1c | 1457.58411 | -0.247060971 | 0.41909349 | -0.58951278 | 0.55551734 | 0.94243456 | 1483.62636 | 1205.92228 | 1680.46418 | 1460.32362 | Non-sig. |
| PF3D7_0531000 | N/A | 986.380313 | -0.247148937 | 0.43944355 | -0.56241338 | 0.5738344 | 0.94850402 | 916.651261 | 792.978612 | 1224.50769 | 1011.38369 | Non-sig. |
| PF3D7_1224000 | GCH1 | 2551.49598 | -0.247414839 | 0.48723459 | -0.50779408 | 0.61159777 | 0.9597882 | 3652.90529 | 2693.14007 | 1886.88705 | 1973.05154 | Non-sig. |
| PF3D7_0525700 | N/A | 356.839652 | -0.247628136 | 0.4566269 | -0.54229862 | 0.58761281 | 0.95246063 | 325.546497 | 311.370664 | 449.062965 | 341.378481 | Non-sig. |
| PF3D7_0517200 | N/A | 404.485122 | -0.247641711 | 0.49025278 | -0.50513066 | 0.61346707 | 0.96215172 | 553.274416 | 314.448595 | 325.191071 | 425.026406 | Non-sig. |
| PF3D7_0710000 | N/A | 3524.08342 | -0.248312494 | 0.4376001 | -0.56744159 | 0.5704142 | 0.94850402 | 3191.35054 | 2783.14549 | 4461.68599 | 3660.15166 | Non-sig. |
| PF3D7_1214000 | N/A | 736.292813 | -0.248441575 | 0.43309236 | -0.57364572 | 0.56620757 | 0.94749454 | 728.398061 | 746.800668 | 870.568896 | 599.403627 | Non-sig. |
| PF3D7_0717700 | N/A | 4472.56874 | -0.248803933 | 0.43736046 | -0.56887615 | 0.56944019 | 0.94850402 | 3855.12044 | 4343.22848 | 5859.20918 | 3832.71687 | Non-sig. |
| PF3D7_0324800 | RIF | 13.5564217 | -0.248911266 | 0.99947139 | -0.24904291 | 0.80332759 | 0.99028237 | 14.6747625 | 15.0395371 | 14.7723555 | 9.7390318 | Non-sig. |
| PF3D7_0403900 | SET8 | 2444.58795 | -0.248918374 | 0.47222684 | -0.52711611 | 0.59811296 | 0.95655566 | 3181.06121 | 1685.33617 | 2129.0461 | 2782.90832 | Non-sig. |
| PF3D7_0305100 | N/A | 3516.35264 | -0.248995518 | 0.42993661 | -0.57914472 | 0.56249153 | 0.94519828 | 3605.94436 | 2641.52226 | 4032.07974 | 3785.86421 | Non-sig. |
| PF3D7_0525500 | N/A | 392.255566 | -0.249516565 | 0.45836578 | -0.54436124 | 0.58619291 | 0.95246063 | 358.314318 | 322.860458 | 493.699743 | 394.147747 | Non-sig. |
| PF3D7_1001700 | N/A | 3127.9994 | -0.249699658 | 0.42599512 | -0.58615615 | 0.55777058 | 0.94272305 | 3787.5219 | 3215.61585 | 3008.66453 | 2500.19532 | Non-sig. |
| PF3D7_1212100 | PPP1 | 974.974555 | -0.250343592 | 0.61272544 | -0.40857385 | 0.68285242 | 0.98107166 | 1671.4975 | 1064.59917 | 447.553189 | 716.248361 | Non-sig. |
| PF3D7_1224600 | CCHL | 244.049195 | -0.250373708 | 0.48224957 | -0.5191787 | 0.60363613 | 0.95693578 | 305.557043 | 255.547277 | 224.937217 | 190.155243 | Non-sig. |
| PF3D7_1010800 | N/A | 299.292411 | -0.25047229 | 0.47409715 | -0.52831427 | 0.59728123 | 0.95631916 | 259.434051 | 290.955714 | 390.787544 | 255.992335 | Non-sig. |
| PF3D7_1139500 | N/A | 15.2715896 | -0.250749053 | 1.02946283 | -0.24357271 | 0.80756176 | 0.99028237 | 12.6160205 | 21.467037 | 20.5279689 | 6.47533179 | Non-sig. |
| PF3D7_1340000 | PSOP7 | 153.357719 | -0.250816731 | 0.59850799 | -0.41906998 | 0.67516499 | 0.97898416 | 244.920881 | 144.909181 | 88.6782733 | 134.922542 | Non-sig. |
| PF3D7_1215200 | CYP32 | 371.128165 | -0.251062727 | 0.4591509 | -0.54679786 | 0.58451762 | 0.95151408 | 342.621072 | 374.634863 | 463.904978 | 303.351749 | Non-sig. |
| PF3D7_0812800 | N/A | 65.4032908 | -0.251186153 | 0.6012746 | -0.41775614 | 0.67612543 | 0.97898416 | 55.3111973 | 56.7615337 | 86.7181034 | 62.8223289 | Non-sig. |
| PF3D7_1466900 | N/A | 2122.16412 | -0.25129126 | 0.48044273 | -0.52304102 | 0.6009457 | 0.95693578 | 2910.42802 | 1529.61646 | 1702.85891 | 2345.7531 | Non-sig. |
| PF3D7_0420400 | RRF2 | 489.966957 | -0.251440874 | 0.52427114 | -0.47960083 | 0.63151126 | 0.9668494 | 724.020233 | 529.664788 | 341.381828 | 364.800978 | Non-sig. |
| PF3D7_1010900 | N/A | 329.628043 | -0.251525813 | 0.50718829 | -0.49592196 | 0.61994947 | 0.9639908 | 466.433157 | 341.399798 | 250.40733 | 260.271886 | Non-sig. |
| PF3D7_1305400 | N/A | 447.695054 | -0.251663656 | 0.45440214 | -0.55383466 | 0.579692 | 0.95129458 | 404.872055 | 442.187785 | 568.240773 | 375.479601 | Non-sig. |
| PF3D7_0520100 | PPM9 | 506.867108 | -0.251916501 | 0.49412466 | -0.50982378 | 0.61017493 | 0.95926017 | 374.8459 | 487.093008 | 726.883848 | 438.645676 | Non-sig. |
| PF3D7_0624000 | HK | 7215.8813 | -0.253412022 | 0.59498013 | -0.42591678 | 0.67016849 | 0.97883902 | 3275.00822 | 6565.41115 | 12420.6383 | 6602.46747 | Non-sig. |
| PF3D7_1443500 | PARN | 553.853763 | -0.254015507 | 0.43049519 | -0.59005423 | 0.55515429 | 0.94243456 | 547.749247 | 507.42018 | 657.116519 | 503.129107 | Non-sig. |

| | | | | | | | | | | | | |
|---|---|---|---|---|---|---|---|---|---|---|---|---|
| PF3D7_1013800 | N/A | 4582.83914 | -0.254516882 | 0.74339888 | -0.3423692 | 0.73207306 | 0.98904466 | 8900.81938 | 5146.51602 | 1071.58675 | 3212.4344 | Non-sig. |
| PF3D7_1133100 | N/A | 1241.97338 | -0.25482474 | 0.4332057 | -0.58823034 | 0.55637769 | 0.94243456 | 1580.09581 | 1172.52711 | 1122.85404 | 1092.41655 | Non-sig. |
| PF3D7_1451400 | Sir2B | 383.115361 | -0.255286226 | 0.51270818 | -0.49791721 | 0.6185424 | 0.96395246 | 282.387599 | 294.986414 | 551.185761 | 403.901672 | Non-sig. |
| PF3D7_0904100 | N/A | 1866.335 | -0.255328255 | 0.44944753 | -0.56809358 | 0.56997143 | 0.94850402 | 1922.32788 | 1299.13606 | 2139.72377 | 2104.1523 | Non-sig. |
| PF3D7_1033200 | ETRAMP10.2 | 1517.03164 | -0.256831611 | 0.45682976 | -0.56220421 | 0.5739769 | 0.94850402 | 1231.14705 | 1307.01851 | 2072.0026 | 1457.95838 | Non-sig. |
| PF3D7_1410200 | CTPS | 1247.59941 | -0.257104293 | 0.41313566 | -0.62232413 | 0.53372876 | 0.93467024 | 1292.56972 | 1172.514 | 1424.31776 | 1100.99617 | Non-sig. |
| PF3D7_1003700 | N/A | 697.820502 | -0.257498306 | 0.43447326 | -0.59266778 | 0.55340351 | 0.94213746 | 652.251299 | 626.438402 | 867.444574 | 645.147735 | Non-sig. |
| PF3D7_0209800 | UAP56 | 6494.24871 | -0.257870287 | 0.44289419 | -0.58223904 | 0.56040568 | 0.94472957 | 5464.08188 | 6296.28797 | 8681.88367 | 5534.74131 | Non-sig. |
| PF3D7_1144900 | RAB6 | 2481.19172 | -0.258132689 | 0.41499177 | -0.62201881 | 0.5339295 | 0.93467024 | 2661.96147 | 2008.46815 | 2743.16424 | 2511.17303 | Non-sig. |
| PF3D7_1110300 | N/A | 478.259377 | -0.258279718 | 0.57393515 | -0.45001551 | 0.65269926 | 0.97317009 | 749.03599 | 559.114409 | 293.172697 | 311.714413 | Non-sig. |
| PF3D7_0826100 | HEUL | 13572.2831 | -0.258459439 | 0.46758151 | -0.55275804 | 0.58042911 | 0.95129458 | 11761.7571 | 9423.12243 | 17807.6106 | 15296.6424 | Non-sig. |
| PF3D7_0731300 | PfG174 | 976.300117 | -0.258601218 | 0.469893 | -0.55034065 | 0.58208575 | 0.95129458 | 820.172561 | 732.801909 | 1306.72987 | 1045.49613 | Non-sig. |
| PF3D7_1442300 | tRIP | 1825.07666 | -0.258859303 | 0.42546357 | -0.60841709 | 0.54291088 | 0.93953127 | 1703.41504 | 1757.77197 | 2273.16158 | 1565.95807 | Non-sig. |
| PF3D7_0609400 | CLS | 421.251129 | -0.258911787 | 0.52385307 | -0.49424505 | 0.62113313 | 0.9639908 | 635.691016 | 405.604087 | 282.518157 | 361.191255 | Non-sig. |
| PF3D7_1458200 | N/A | 211.171819 | -0.259242876 | 0.53549494 | -0.48411825 | 0.62830195 | 0.96655174 | 148.245556 | 187.130058 | 311.677654 | 197.634008 | Non-sig. |
| PF3D7_1467100 | MAG | 468.692174 | -0.259868537 | 0.49466898 | -0.52533825 | 0.59934807 | 0.95655566 | 355.994605 | 473.107405 | 665.316656 | 380.350031 | Non-sig. |
| PF3D7_1238000 | N/A | 615.001267 | -0.25991404 | 0.43234118 | -0.60117808 | 0.54772139 | 0.94142878 | 747.161495 | 575.616733 | 593.474937 | 543.751903 | Non-sig. |
| PF3D7_1348200 | N/A | 247.005327 | -0.260036079 | 0.50027288 | -0.51978848 | 0.60321101 | 0.95693578 | 337.562555 | 249.217712 | 201.072708 | 200.168333 | Non-sig. |
| PF3D7_0219500 | N/A | 513.278097 | -0.260167726 | 0.52877258 | -0.49202197 | 0.62270381 | 0.96429926 | 776.780818 | 535.632756 | 342.394475 | 398.304338 | Non-sig. |
| PF3D7_1315800 | MYB1 | 841.475234 | -0.260966224 | 0.44223155 | -0.59011218 | 0.55511544 | 0.94243456 | 1084.3959 | 805.989131 | 750.559509 | 724.956402 | Non-sig. |
| PF3D7_1007600 | N/A | 972.604512 | -0.26112165 | 0.43429679 | -0.60125162 | 0.54767241 | 0.94142878 | 931.620536 | 985.296069 | 1189.01078 | 784.490667 | Non-sig. |
| PF3D7_1302600 | DOHH | 573.20835 | -0.261338063 | 0.4612257 | -0.56661644 | 0.5709748 | 0.94850402 | 480.599997 | 505.81776 | 769.131889 | 537.283755 | Non-sig. |
| PF3D7_1350700 | N/A | 622.131478 | -0.261413019 | 0.42884484 | -0.60957483 | 0.54214349 | 0.93953127 | 731.033353 | 599.921653 | 625.745836 | 531.82507 | Non-sig. |
| PF3D7_0606600 | N/A | 2930.78989 | -0.261548755 | 0.46542927 | -0.56195167 | 0.57414895 | 0.94850402 | 2530.13584 | 2094.63859 | 3861.10781 | 3237.27731 | Non-sig. |
| PF3D7_0308200 | CCT7 | 2870.23816 | -0.26157723 | 0.50217017 | -0.5208936 | 0.60244089 | 0.95693578 | 1915.13612 | 2647.95913 | 4343.99244 | 2573.86497 | Non-sig. |
| PF3D7_1429400 | N/A | 884.344337 | -0.261774067 | 0.42215955 | -0.62008326 | 0.53520297 | 0.93519306 | 884.471498 | 765.787173 | 1044.13765 | 842.981024 | Non-sig. |
| PF3D7_1458900 | TVP23 | 347.286008 | -0.262118486 | 0.45113551 | -0.58101941 | 0.56122737 | 0.94472957 | 408.794154 | 350.207037 | 348.791305 | 281.351536 | Non-sig. |
| PF3D7_1343000 | PMT | 59015.404 | -0.262458935 | 0.41629831 | -0.63045881 | 0.52839445 | 0.93270419 | 61965.9738 | 61181.5109 | 66771.5767 | 46142.5546 | Non-sig. |
| PF3D7_1435900 | N/A | 147.399145 | -0.262480308 | 0.50019369 | -0.52475733 | 0.5997519 | 0.95655566 | 137.532101 | 139.590649 | 183.885319 | 128.58851 | Non-sig. |
| PF3D7_0932200 | PFN | 14421.9548 | -0.262707864 | 0.42359735 | -0.62018297 | 0.53513733 | 0.93519306 | 13465.1576 | 14369.7597 | 17997.5291 | 11855.3728 | Non-sig. |
| PF3D7_1467400 | RPL22 | 1017.63604 | -0.262814482 | 0.43776008 | -0.60036192 | 0.54826506 | 0.94142878 | 927.470856 | 998.15873 | 1292.48817 | 852.426406 | Non-sig. |
| PF3D7_1369400 | N/A | 1124.57792 | -0.263893044 | 0.66729167 | -0.39546881 | 0.69249693 | 0.98334178 | 391.125949 | 863.375827 | 2062.82575 | 1180.98416 | Non-sig. |
| PF3D7_1418000 | UFD1 | 1996.76695 | -0.263945883 | 0.47987943 | -0.55002542 | 0.58230194 | 0.95129458 | 2867.07338 | 1991.25668 | 1491.06327 | 1637.67447 | Non-sig. |
| PF3D7_0213900 | N/A | 427.3203 | -0.264305182 | 0.48127202 | -0.54918045 | 0.58288162 | 0.95129458 | 584.670086 | 345.423086 | 348.27727 | 430.910758 | Non-sig. |
| PF3D7_1146000 | RSA4 | 448.739759 | -0.26449962 | 0.43470846 | -0.60845289 | 0.54288714 | 0.93953127 | 473.58114 | 377.71206 | 505.899056 | 437.766781 | Non-sig. |
| PF3D7_API05900 | N/A | 5.28754765 | -0.264818472 | 1.80452468 | -0.14675248 | 0.88332739 | 0.99457954 | 0 | 5.36030592 | 11.4744569 | 4.31542782 | Non-sig. |
| PF3D7_1114700 | CLK3 | 7355.95875 | -0.264829052 | 0.42135573 | -0.62851655 | 0.52966561 | 0.93270419 | 9165.81866 | 7184.92731 | 6892.80326 | 6180.28577 | Non-sig. |
| PF3D7_0608100 | N/A | 2691.78155 | -0.264849319 | 0.55027618 | -0.48130253 | 0.6303015 | 0.9668494 | 4385.76517 | 2448.87201 | 1490.93172 | 2441.5573 | Non-sig. |
| PF3D7_1440300 | PBGS | 386.800525 | -0.264986757 | 0.44240591 | -0.59896749 | 0.54919456 | 0.94142878 | 394.373738 | 323.05056 | 449.997706 | 379.780097 | Non-sig. |
| PF3D7_0927100 | N/A | 188.01301 | -0.265019442 | 0.47887212 | -0.55342425 | 0.57997294 | 0.95129458 | 218.422119 | 152.466035 | 192.102313 | 189.061574 | Non-sig. |
| PF3D7_1469400 | NT3 | 231.344719 | -0.265142818 | 0.64005095 | -0.41425268 | 0.67868907 | 0.97898416 | 103.157021 | 192.063866 | 401.613121 | 228.544868 | Non-sig. |
| PF3D7_0505400 | N/A | 2530.5261 | -0.265599947 | 0.43460296 | -0.6111324 | 0.54111193 | 0.93839968 | 3309.93191 | 2310.03581 | 2215.91004 | 2286.22665 | Non-sig. |
| PF3D7_1302050 | N/A | 3.00823458 | -0.265615646 | 1.96257147 | -0.13534062 | 0.89234257 | 0.99457954 | 4.14816575 | 4.3594032 | 2.43552038 | 1.089849 | Non-sig. |
| PF3D7_1229300 | PIC1 | 4082.87403 | -0.265786754 | 0.46386038 | -0.57298869 | 0.56665235 | 0.9476402 | 5371.2256 | 2908.74205 | 3544.79391 | 4506.73458 | Non-sig. |
| PF3D7_0415400 | N/A | 2346.32948 | -0.266051221 | 0.41125213 | -0.64692971 | 0.51767744 | 0.93247207 | 2396.08547 | 2252.69987 | 2727.96548 | 2008.56711 | Non-sig. |
| PF3D7_1314400 | N/A | 45.0905478 | -0.266068591 | 0.64237232 | -0.41419685 | 0.67872995 | 0.97898416 | 46.2039723 | 41.1201566 | 52.2395595 | 40.7985029 | Non-sig. |
| PF3D7_0628900 | N/A | 409.00123 | -0.266290527 | 0.44329081 | -0.60071295 | 0.54803119 | 0.94142878 | 392.915128 | 371.175637 | 500.232254 | 371.681902 | Non-sig. |
| PF3D7_0626500 | CEP135 | 62.3401652 | -0.2672158 | 0.7662553 | -0.34872946 | 0.72729242 | 0.98844303 | 112.622745 | 59.0151987 | 23.8172558 | 53.905461 | Non-sig. |
| PF3D7_1241300 | N/A | 165.39908 | -0.267399809 | 0.49493124 | -0.54027669 | 0.58900623 | 0.95315309 | 160.902427 | 130.937907 | 200.363493 | 169.392492 | Non-sig. |
| PF3D7_1408400 | FANCJ | 1947.97178 | -0.267715162 | 0.53165231 | -0.50355309 | 0.61457547 | 0.96217474 | 1220.95351 | 1482.53226 | 3035.1023 | 2053.29905 | Non-sig. |
| PF3D7_1215000 | Trx-Px2 | 940.875221 | -0.267818576 | 0.4278843 | -0.62591354 | 0.53137165 | 0.93351492 | 991.13517 | 964.231734 | 1064.74097 | 743.393006 | Non-sig. |
| PF3D7_1023100 | N/A | 280.094217 | -0.267973993 | 0.49865811 | -0.53739022 | 0.5909981 | 0.95315309 | 271.014622 | 186.60373 | 340.944937 | 321.813581 | Non-sig. |
| PF3D7_0313100 | HRD3 | 2279.99979 | -0.268166512 | 0.41855547 | -0.64069527 | 0.52172069 | 0.93247207 | 2185.80358 | 2149.02928 | 2796.63346 | 1988.53285 | Non-sig. |
| PF3D7_1015600 | HSP60 | 2161.5599 | -0.268624301 | 0.47504747 | -0.56546834 | 0.57175525 | 0.94850402 | 1651.0361 | 2168.69299 | 3073.11279 | 1753.39771 | Non-sig. |

| | | | | | | | | | | | | |
|---|---|---|---|---|---|---|---|---|---|---|---|---|
| PF3D7_1359900 | N/A | 5837.94669 | -0.268927178 | 0.45420804 | -0.59207929 | 0.55379749 | 0.94243456 | 6239.97925 | 3873.06148 | 6520.97876 | 6717.76726 | Non-sig. |
| PF3D7_1214400 | EMC6 | 248.530859 | -0.26916183 | 0.59709189 | -0.45078795 | 0.65214238 | 0.97317009 | 132.791628 | 246.174405 | 410.201738 | 204.955664 | Non-sig. |
| PF3D7_1031300 | N/A | 1350.82514 | -0.269590327 | 0.51952772 | -0.51891423 | 0.60382055 | 0.95693578 | 2045.9832 | 1008.35511 | 907.669294 | 1441.29294 | Non-sig. |
| PF3D7_0909900 | N/A | 1317.40662 | -0.269620679 | 0.45983066 | -0.58634777 | 0.55764184 | 0.94272305 | 1058.91245 | 1169.23164 | 1821.1186 | 1220.3638 | Non-sig. |
| PF3D7_0211100 | N/A | 109.116747 | -0.269621478 | 0.52210118 | -0.51641614 | 0.6055638 | 0.95693578 | 132.702033 | 104.063501 | 105.957 | 93.7444535 | Non-sig. |
| PF3D7_1431000 | N/A | 586.314082 | -0.269659078 | 0.42482991 | -0.63474597 | 0.5255941 | 0.93247207 | 615.952114 | 548.51538 | 665.902613 | 514.886223 | Non-sig. |
| PF3D7_1210100 | SYN2 | 1771.28263 | -0.270316295 | 0.41171331 | -0.65656438 | 0.51146106 | 0.93247207 | 1854.00261 | 1705.15502 | 2019.42692 | 1506.54596 | Non-sig. |
| PF3D7_0107100 | N/A | 180.44939 | -0.271013497 | 0.48052255 | -0.56399745 | 0.57275587 | 0.94850402 | 178.668262 | 174.972501 | 215.941786 | 152.215013 | Non-sig. |
| PF3D7_1217700 | N/A | 1085.13729 | -0.271359388 | 0.44591573 | -0.6085441 | 0.54282666 | 0.93953127 | 1401.74082 | 1099.04371 | 972.242458 | 867.522183 | Non-sig. |
| PF3D7_0305700 | N/A | 2802.57685 | -0.271365449 | 0.42781276 | -0.63430892 | 0.52587924 | 0.93247207 | 2664.0063 | 2834.92728 | 3466.60542 | 2244.7684 | Non-sig. |
| PF3D7_0703400 | N/A | 335.853969 | -0.271867979 | 0.46719677 | -0.58191323 | 0.56062513 | 0.94472957 | 296.86151 | 285.407414 | 437.755736 | 323.391217 | Non-sig. |
| PF3D7_1436500 | N/A | 393.294285 | -0.272067526 | 0.48333564 | -0.56289564 | 0.57350595 | 0.94850402 | 318.401356 | 324.196064 | 541.892634 | 388.687088 | Non-sig. |
| PF3D7_1215100 | N/A | 619.81819 | -0.27236488 | 0.47203068 | -0.57700673 | 0.56393491 | 0.94638173 | 853.058248 | 510.66655 | 503.49839 | 612.049571 | Non-sig. |
| PF3D7_0605400 | N/A | 357.290325 | -0.272586933 | 0.45464114 | -0.59956504 | 0.54879616 | 0.94142878 | 450.828706 | 334.711277 | 331.217029 | 312.404286 | Non-sig. |
| PF3D7_1421000 | DIX | 129.313922 | -0.272928539 | 0.50479834 | -0.54066846 | 0.58873612 | 0.95315309 | 137.111888 | 104.463023 | 145.874892 | 129.805885 | Non-sig. |
| PF3D7_1012700 | NIF4 | 1875.00785 | -0.273130078 | 0.48801396 | -0.55967678 | 0.57569993 | 0.94896603 | 2533.50654 | 1246.38966 | 1570.64024 | 2149.49494 | Non-sig. |
| PF3D7_0721000 | N/A | 6653.93375 | -0.273148463 | 0.42679468 | -0.63999969 | 0.5221728 | 0.93247207 | 6759.32787 | 5022.48094 | 7804.5079 | 7029.41828 | Non-sig. |
| PF3D7_0934000 | N/A | 598.034505 | -0.273252043 | 0.42269424 | -0.6464532 | 0.5179859 | 0.93247207 | 657.746581 | 550.473584 | 651.262042 | 532.655812 | Non-sig. |
| PF3D7_1425700 | N/A | 1024.84758 | -0.273612349 | 0.45816384 | -0.59719324 | 0.55037837 | 0.94170064 | 1379.85141 | 1027.09312 | 863.871795 | 828.57399 | Non-sig. |
| PF3D7_0513400 | YihA1 | 190.943484 | -0.274173963 | 0.49008929 | -0.55943675 | 0.57586368 | 0.94896603 | 170.000228 | 170.751979 | 247.896862 | 175.124867 | Non-sig. |
| PF3D7_0929100 | N/A | 768.516117 | -0.27435427 | 0.44157423 | -0.62130951 | 0.534396 | 0.93470782 | 937.950516 | 793.991867 | 744.923159 | 597.198927 | Non-sig. |
| PF3D7_0707800 | N/A | 2353.26161 | -0.274367678 | 0.40626377 | -0.6753437 | 0.49945743 | 0.93170321 | 2643.31681 | 2075.71548 | 2509.43433 | 2184.57983 | Non-sig. |
| PF3D7_1235700 | N/A | 1992.70496 | -0.274536349 | 0.58119908 | -0.47236198 | 0.63666844 | 0.96815998 | 984.271241 | 1905.57929 | 3378.84792 | 1702.12137 | Non-sig. |
| PF3D7_1352300 | N/A | 328.635057 | -0.275135472 | 0.44332841 | -0.62061322 | 0.53485414 | 0.93519306 | 337.802203 | 296.165281 | 381.884328 | 298.688415 | Non-sig. |
| PF3D7_0931800 | N/A | 5283.4249 | -0.275803693 | 0.46426983 | -0.59405905 | 0.55247262 | 0.94170064 | 4182.18395 | 5309.18282 | 7391.37672 | 4250.95612 | Non-sig. |
| PF3D7_0710400 | RAD14 | 236.773763 | -0.275828698 | 0.46025547 | -0.59929478 | 0.54897633 | 0.94142878 | 246.877698 | 228.672333 | 271.75436 | 199.790662 | Non-sig. |
| PF3D7_0311100 | N/A | 1178.30238 | -0.27598905 | 0.54083779 | -0.51029913 | 0.60984191 | 0.95903775 | 1882.7765 | 1140.64267 | 698.898102 | 990.892253 | Non-sig. |
| PF3D7_1450200 | N/A | 40.503998 | -0.276222279 | 0.70407546 | -0.39231914 | 0.69482242 | 0.98334178 | 30.3448947 | 33.318131 | 58.255609 | 40.0973574 | Non-sig. |
| PF3D7_1420100 | N/A | 279.830017 | -0.276254263 | 0.47961572 | -0.57599085 | 0.56462137 | 0.94662774 | 253.943311 | 216.679678 | 358.97025 | 289.726828 | Non-sig. |
| PF3D7_1450100 | SRP54 | 1451.73608 | -0.276336599 | 0.4381696 | -0.63066128 | 0.52826202 | 0.93270419 | 1283.47186 | 1256.66286 | 1897.00273 | 1369.80687 | Non-sig. |
| PF3D7_0208800 | N/A | 163.500722 | -0.276360649 | 0.50900274 | -0.54294531 | 0.58716746 | 0.95246063 | 138.79303 | 145.970884 | 219.243363 | 149.99561 | Non-sig. |
| PF3D7_0903200 | RAB7 | 7500.11359 | -0.276361229 | 0.41440577 | -0.66688558 | 0.50484523 | 0.93170321 | 8006.54005 | 7638.31389 | 8426.00261 | 5929.59779 | Non-sig. |
| PF3D7_1028300 | EBP2 | 347.800243 | -0.276521864 | 0.50894512 | -0.54332355 | 0.58690706 | 0.95246063 | 509.474606 | 313.491805 | 252.867728 | 315.366833 | Non-sig. |
| PF3D7_0907400 | ClpY | 9532.54488 | -0.276931943 | 0.43532301 | -0.63615279 | 0.52467684 | 0.93247207 | 11920.5641 | 7228.20471 | 8968.91063 | 10012.5001 | Non-sig. |
| PF3D7_1454800 | N/A | 9.58690242 | -0.277007677 | 1.18296483 | -0.23416392 | 0.81485773 | 0.99096399 | 6.20710706 | 7.54765749 | 14.7397437 | 9.8531014 | Non-sig. |
| PF3D7_0314700 | RNF1 | 610.594538 | -0.277115841 | 0.45011273 | -0.61565875 | 0.53811977 | 0.93646681 | 775.588235 | 485.909661 | 562.695517 | 618.184737 | Non-sig. |
| PF3D7_1413500 | SufC | 858.586478 | -0.277202938 | 0.45163509 | -0.61377635 | 0.53936312 | 0.93711173 | 748.337383 | 849.089268 | 1133.08725 | 703.832007 | Non-sig. |
| PF3D7_0609100 | ZIP1 | 776.106181 | -0.277241103 | 0.4386615 | -0.63201603 | 0.5273764 | 0.93270419 | 709.419231 | 675.991957 | 991.290904 | 727.722632 | Non-sig. |
| PF3D7_1038900 | N/A | 218.802542 | -0.277256964 | 0.51495396 | -0.53841117 | 0.59029321 | 0.95315309 | 170.490017 | 179.198381 | 308.795783 | 216.725988 | Non-sig. |
| PF3D7_0802000 | GDH3 | 10225.623 | -0.277271528 | 0.46675482 | -0.59404106 | 0.55248465 | 0.94170064 | 7867.90127 | 8543.54633 | 14542.2901 | 9948.75438 | Non-sig. |
| PF3D7_1362900 | N/A | 798.571578 | -0.27729063 | 0.45250548 | -0.61278955 | 0.5400155 | 0.93711173 | 1054.63128 | 656.497693 | 695.746723 | 787.410613 | Non-sig. |
| PF3D7_0527600 | N/A | 2116.47911 | -0.277365152 | 0.4303802 | -0.64446541 | 0.51927368 | 0.93247207 | 2623.53599 | 1679.52446 | 2015.22825 | 2147.62775 | Non-sig. |
| PF3D7_1342900 | AP2-HS | 1557.3814 | -0.277634398 | 0.44378058 | -0.62561186 | 0.53156956 | 0.93351492 | 1536.22817 | 1131.46573 | 1877.20339 | 1684.62833 | Non-sig. |
| PF3D7_0934100 | XPD | 1340.3193 | -0.27812484 | 0.45841723 | -0.60670677 | 0.54404553 | 0.94060399 | 1084.90759 | 1199.37461 | 1853.05153 | 1223.94347 | Non-sig. |
| PF3D7_0619500 | ACS12 | 1196.36584 | -0.278142009 | 0.49505519 | -0.5618404 | 0.57422476 | 0.94850402 | 848.092461 | 1013.72232 | 1774.26365 | 1149.38491 | Non-sig. |
| PF3D7_1124000 | ERO1 | 1325.22057 | -0.280043419 | 0.43583407 | -0.64254595 | 0.52051877 | 0.93247207 | 1185.2362 | 1171.70159 | 1721.42945 | 1222.51503 | Non-sig. |
| PF3D7_1020700 | N/A | 583.2227 | -0.280053272 | 0.42617103 | -0.65713823 | 0.51109204 | 0.93247207 | 620.496623 | 496.677771 | 658.771391 | 556.945014 | Non-sig. |
| PF3D7_0926700 | NADSYN | 2002.16203 | -0.280118987 | 0.52701312 | -0.53152185 | 0.59505721 | 0.95475866 | 1231.86142 | 1888.32283 | 3159.65205 | 1728.81181 | Non-sig. |
| PF3D7_0823600 | LipB | 59.0663946 | -0.280266843 | 0.61716455 | -0.45412012 | 0.64974237 | 0.97317009 | 78.7941309 | 57.9725698 | 50.9030128 | 48.595865 | Non-sig. |
| PF3D7_1457400 | N/A | 1586.19614 | -0.280534599 | 0.45101105 | -0.6220127 | 0.53393352 | 0.93467024 | 1360.08709 | 1275.19382 | 2119.53942 | 1589.96422 | Non-sig. |
| PF3D7_1147500 | FTB | 123.845526 | -0.280629389 | 0.50192914 | -0.55910161 | 0.57609238 | 0.94904771 | 133.709154 | 109.438208 | 137.982215 | 114.252526 | Non-sig. |
| PF3D7_1015200 | CysRS | 787.179091 | -0.280858361 | 0.512093 | -0.54845187 | 0.58338167 | 0.95129458 | 529.773032 | 681.674657 | 1197.07842 | 740.190262 | Non-sig. |
| PF3D7_0317000 | N/A | 4987.30177 | -0.281756788 | 0.47052477 | -0.59881393 | 0.54929697 | 0.94142878 | 3826.68358 | 4917.78196 | 7118.56403 | 4086.17752 | Non-sig. |
| PF3D7_1409000 | N/A | 325.825934 | -0.282007479 | 0.5185905 | -0.54379607 | 0.58658181 | 0.95246063 | 236.862587 | 257.508919 | 478.001459 | 330.930772 | Non-sig. |

| | | | | | | | | | | | | |
|---|---|---|---|---|---|---|---|---|---|---|---|---|
| PF3D7_0310400 | PIESP1 | 7843.7607 | -0.282166778 | 0.41708335 | -0.67652372 | 0.49870819 | 0.93170321 | 7921.84909 | 6246.85056 | 9294.81065 | 7911.53251 | Non-sig. |
| PF3D7_0521200 | N/A | 84.3588541 | -0.282538431 | 0.70014466 | -0.40354293 | 0.68654886 | 0.98210107 | 146.865338 | 69.8107258 | 38.5777299 | 82.1816229 | Non-sig. |
| PF3D7_1332300 | TRAPPC2 | 878.047238 | -0.28283281 | 0.4159699 | -0.67993577 | 0.49654513 | 0.93170321 | 971.293365 | 767.710903 | 956.39728 | 816.787405 | Non-sig. |
| PF3D7_0323200 | N/A | 689.079199 | -0.282926387 | 0.41988559 | -0.67381781 | 0.50042715 | 0.93170321 | 746.001857 | 633.38395 | 766.844448 | 610.086541 | Non-sig. |
| PF3D7_1003800 | EFTUD2 | 2385.71642 | -0.284494026 | 0.53157069 | -0.53519509 | 0.59251496 | 0.95322751 | 1435.11649 | 2118.53987 | 3804.90558 | 2184.30375 | Non-sig. |
| PF3D7_0618600 | ROM10 | 270.699171 | -0.285278996 | 0.48063994 | -0.59353993 | 0.55281987 | 0.94170064 | 231.969874 | 239.76848 | 362.580821 | 248.477509 | Non-sig. |
| PF3D7_0901800 | N/A | 677.385565 | -0.285332301 | 0.45216693 | -0.6310331 | 0.52801888 | 0.93270419 | 593.920641 | 652.623758 | 894.176231 | 568.821631 | Non-sig. |
| PF3D7_1410300 | N/A | 5089.02939 | -0.285603402 | 0.44846761 | -0.63684288 | 0.52422719 | 0.93247207 | 6257.19823 | 3562.69277 | 4925.16296 | 5611.0636 | Non-sig. |
| PF3D7_1106000 | RUVB2 | 1101.6527 | -0.286454516 | 0.54582666 | -0.52480858 | 0.59971627 | 0.95655566 | 638.830414 | 985.046309 | 1782.15517 | 1000.57889 | Non-sig. |
| PF3D7_1367100 | SNP1 | 1123.79246 | -0.286746626 | 0.41198865 | -0.69600613 | 0.48642499 | 0.93170321 | 1217.05079 | 1001.28548 | 1253.14565 | 1023.68791 | Non-sig. |
| PF3D7_0521400 | N/A | 854.640245 | -0.286853256 | 0.41740125 | -0.68723621 | 0.49193389 | 0.93170321 | 929.42094 | 806.077306 | 949.220584 | 733.842153 | Non-sig. |
| PF3D7_0923000 | RPB3 | 314.098254 | -0.286939629 | 0.4657523 | -0.61607775 | 0.53784321 | 0.93646681 | 287.974535 | 307.005965 | 402.325818 | 259.086697 | Non-sig. |
| PF3D7_1123600 | N/A | 670.002384 | -0.287107816 | 0.44215953 | -0.64933084 | 0.51612456 | 0.93247207 | 611.914836 | 627.596282 | 860.800657 | 579.69776 | Non-sig. |
| PF3D7_1303500 | NHE | 18078.4745 | -0.287920596 | 0.41871733 | -0.68762522 | 0.49168882 | 0.93170321 | 22489.268 | 14985.5871 | 17263.8765 | 17575.1663 | Non-sig. |
| PF3D7_0526800 | N/A | 1511.45138 | -0.287972095 | 0.44034941 | -0.65396271 | 0.51313583 | 0.93247207 | 1339.31461 | 1459.46273 | 1984.07348 | 1262.95469 | Non-sig. |
| PF3D7_1364600 | N/A | 1482.05627 | -0.288154376 | 0.41603705 | -0.6926171 | 0.48854988 | 0.93170321 | 1785.78897 | 1329.31199 | 1473.4807 | 1339.64341 | Non-sig. |
| PF3D7_1333300 | N/A | 730.37165 | -0.28851278 | 0.48129427 | -0.59945193 | 0.54887155 | 0.94142878 | 565.668272 | 602.135425 | 1040.3736 | 713.309307 | Non-sig. |
| PF3D7_1365100 | mtRPS17 | 125.244071 | -0.288735289 | 0.51307494 | -0.56275462 | 0.57360199 | 0.94850402 | 128.576634 | 127.933829 | 146.845976 | 97.6198443 | Non-sig. |
| PF3D7_0510800 | N/A | 66.4606606 | -0.289140884 | 0.67126466 | -0.43074051 | 0.66665706 | 0.97883902 | 106.182984 | 67.6214988 | 40.2346957 | 51.803464 | Non-sig. |
| PF3D7_0903100 | RER1 | 194.752706 | -0.289295842 | 0.49385762 | -0.58578795 | 0.55801802 | 0.94272305 | 170.599931 | 167.653599 | 257.648489 | 183.108803 | Non-sig. |
| PF3D7_1119400 | N/A | 3729.3732 | -0.28982025 | 0.48066611 | -0.60295546 | 0.54653832 | 0.94142878 | 2788.15913 | 2978.4767 | 5416.96376 | 3733.89322 | Non-sig. |
| PF3D7_0623200 | FNR | 477.383958 | -0.289919189 | 0.49951704 | -0.58039899 | 0.56164558 | 0.94472957 | 350.394033 | 444.260862 | 699.697966 | 415.18297 | Non-sig. |
| PF3D7_0829700 | N/A | 90.5737658 | -0.290954417 | 0.58529967 | -0.49710334 | 0.61911618 | 0.96395246 | 67.9066944 | 78.493371 | 131.247464 | 84.6475343 | Non-sig. |
| PF3D7_1441800 | VPS60 | 523.200129 | -0.291025165 | 0.47981459 | -0.60653672 | 0.54415841 | 0.94060399 | 701.132102 | 559.47592 | 450.663211 | 381.529284 | Non-sig. |
| PF3D7_0311600 | OST1 | 495.46236 | -0.291594483 | 0.51914162 | -0.56168581 | 0.5743301 | 0.94850402 | 338.941279 | 482.843412 | 751.483427 | 408.581321 | Non-sig. |
| PF3D7_1441500 | N/A | 1529.84427 | -0.291995693 | 0.49634667 | -0.58828982 | 0.55633778 | 0.94243456 | 2295.2848 | 1446.6529 | 1073.30645 | 1304.13293 | Non-sig. |
| PF3D7_0213200 | N/A | 298.676624 | -0.29215661 | 0.48166957 | -0.60654986 | 0.54414969 | 0.94060399 | 403.443556 | 288.996525 | 254.400239 | 247.866176 | Non-sig. |
| PF3D7_0804600 | N/A | 808.826028 | -0.292262983 | 0.42556181 | -0.68676976 | 0.49222783 | 0.93170321 | 957.360477 | 784.471426 | 823.679461 | 669.79275 | Non-sig. |
| PF3D7_1126900 | SNRPF | 631.636261 | -0.292273843 | 0.4401236 | -0.66407219 | 0.50664411 | 0.93170321 | 782.073298 | 625.42384 | 608.867924 | 510.179982 | Non-sig. |
| PF3D7_1471200 | SulP | 4781.48042 | -0.292380471 | 0.57520687 | -0.5083049 | 0.61123953 | 0.9597882 | 2411.63694 | 4009.2063 | 8116.69929 | 4588.37917 | Non-sig. |
| PF3D7_1228900 | N/A | 1080.76664 | -0.292418816 | 0.50383348 | -0.58038782 | 0.56165312 | 0.94472957 | 1625.38131 | 1066.52024 | 754.780556 | 876.384461 | Non-sig. |
| PF3D7_1015300 | METAP1b | 545.96945 | -0.292609655 | 0.43657974 | -0.67023186 | 0.50270999 | 0.93170321 | 536.911264 | 524.668172 | 665.264683 | 457.033681 | Non-sig. |
| PF3D7_0611100 | SDH3 | 267.047779 | -0.29336429 | 0.46473137 | -0.63125562 | 0.5278734 | 0.93270419 | 252.363064 | 251.245937 | 335.720119 | 228.861996 | Non-sig. |
| PF3D7_1365900 | N/A | 7580.22715 | -0.29466575 | 0.41314385 | -0.71322796 | 0.47570471 | 0.9269156 | 7967.32937 | 7579.97775 | 8735.95448 | 6037.64701 | Non-sig. |
| PF3D7_1120000 | N/A | 2433.78738 | -0.295042173 | 0.6812116 | -0.43311384 | 0.66493207 | 0.97883902 | 4574.58731 | 2574.842 | 789.324335 | 1796.39587 | Non-sig. |
| PF3D7_1355500 | PP5 | 1834.65545 | -0.295207493 | 0.49972201 | -0.59074342 | 0.55469235 | 0.94243456 | 1259.50527 | 1653.25595 | 2783.58728 | 1642.27329 | Non-sig. |
| PF3D7_0218700 | PRP45 | 1067.69009 | -0.295240577 | 0.59674603 | -0.4947508 | 0.62077604 | 0.9639908 | 1825.4099 | 1133.22495 | 528.047475 | 784.078022 | Non-sig. |
| PF3D7_0510400 | N/A | 93.9504069 | -0.295786927 | 0.56750082 | -0.52120969 | 0.60222071 | 0.95693578 | 86.2683932 | 103.021989 | 120.718907 | 65.7923381 | Non-sig. |
| PF3D7_1309700 | VPS18 | 459.390767 | -0.296099193 | 0.46774004 | -0.63304222 | 0.52670607 | 0.93270419 | 495.207999 | 314.298902 | 517.502384 | 510.553785 | Non-sig. |
| PF3D7_1135100 | PPM8 | 1292.61755 | -0.296301614 | 0.49433597 | -0.59939319 | 0.54891072 | 0.94142878 | 930.822345 | 1050.35334 | 1918.6539 | 1270.64063 | Non-sig. |
| PF3D7_1134200 | N/A | 582.546893 | -0.296379304 | 0.43063875 | -0.68823184 | 0.49130679 | 0.93170321 | 612.329553 | 476.238628 | 671.96381 | 569.655581 | Non-sig. |
| PF3D7_0308800 | N/A | 1048.87417 | -0.296674305 | 0.46611556 | -0.6364823 | 0.52446211 | 0.93247207 | 1462.89716 | 1019.95174 | 850.046294 | 862.6015 | Non-sig. |
| PF3D7_1365500 | GCVT | 1021.02796 | -0.296816446 | 0.43017077 | -0.68999678 | 0.49019621 | 0.93170321 | 971.163543 | 972.532753 | 1280.06169 | 860.353831 | Non-sig. |
| PF3D7_1239400 | N/A | 27.8638221 | -0.29701458 | 0.81154185 | -0.365988 | 0.71437403 | 0.9875928 | 43.5535972 | 26.9364455 | 18.0237645 | 22.9414813 | Non-sig. |
| PF3D7_0913100 | N/A | 124.018403 | -0.297320263 | 0.58750437 | -0.50607328 | 0.61280521 | 0.96139828 | 85.2096459 | 92.2925703 | 188.05612 | 130.515277 | Non-sig. |
| PF3D7_1122300 | N/A | 414.116829 | -0.297543792 | 0.43752512 | -0.68006106 | 0.4964658 | 0.93170321 | 442.245431 | 342.706972 | 471.056308 | 400.458606 | Non-sig. |
| PF3D7_1244200 | TFB2 | 1036.43182 | -0.297719083 | 0.4156375 | -0.71629505 | 0.47380918 | 0.92682482 | 1080.90377 | 905.381891 | 1205.01703 | 954.4246 | Non-sig. |
| PF3D7_1212400 | N/A | 361.951374 | -0.298651538 | 0.44036237 | -0.67819495 | 0.4976481 | 0.93170321 | 401.218496 | 346.787645 | 397.353399 | 302.445957 | Non-sig. |
| PF3D7_0718800 | N/A | 15.6680797 | -0.299780695 | 0.9512921 | -0.31513002 | 0.75266294 | 0.99028237 | 19.8523427 | 17.2033481 | 14.7656591 | 10.8509687 | Non-sig. |
| PF3D7_1345400 | N/A | 4471.71892 | -0.300213251 | 0.58412494 | -0.51395383 | 0.60728427 | 0.95715771 | 7679.47515 | 3634.48431 | 2191.50608 | 4381.41015 | Non-sig. |
| PF3D7_0411400 | N/A | 284.289311 | -0.300247231 | 0.45345031 | -0.66213921 | 0.50788201 | 0.93170321 | 343.424891 | 261.749803 | 284.20697 | 247.775582 | Non-sig. |
| PF3D7_0611900 | N/A | 540.093664 | -0.300482107 | 0.45366641 | -0.66234154 | 0.50775236 | 0.93170321 | 501.537571 | 545.247232 | 690.57313 | 423.016723 | Non-sig. |
| PF3D7_1470400 | MPC2 | 384.52245 | -0.300655995 | 0.48143498 | -0.62449969 | 0.53229947 | 0.93443426 | 491.147651 | 419.890912 | 357.895445 | 269.155791 | Non-sig. |
| PF3D7_0203900 | N/A | 486.07325 | -0.300713713 | 0.45135082 | -0.66625272 | 0.50524958 | 0.93170321 | 439.751314 | 447.50379 | 633.156566 | 423.881328 | Non-sig. |

| | | | | | | | | | | | | |
|---|---|---|---|---|---|---|---|---|---|---|---|---|
| PF3D7_1102600 | GEXP14 | 100.387973 | -0.300917751 | 0.52967941 | -0.56811299 | 0.56995825 | 0.94850402 | 120.73707 | 96.639055 | 100.973152 | 83.2026132 | Non-sig. |
| PF3D7_1244900 | N/A | 479.384714 | -0.301672783 | 0.46149224 | -0.65368982 | 0.51331166 | 0.93247207 | 619.377327 | 490.256009 | 439.44659 | 368.45893 | Non-sig. |
| PF3D7_1412900 | N/A | 923.067988 | -0.301747449 | 0.44887539 | -0.67222988 | 0.50143736 | 0.93170321 | 1236.78734 | 785.631882 | 802.529785 | 867.322947 | Non-sig. |
| PF3D7_1306500 | N/A | 490.821544 | -0.302144317 | 0.4540225 | -0.66548314 | 0.50574152 | 0.93170321 | 586.287669 | 361.416214 | 497.857055 | 517.725235 | Non-sig. |
| PF3D7_0921700 | N/A | 1757.46318 | -0.302187381 | 0.53158884 | -0.56846073 | 0.56972216 | 0.94850402 | 2779.3737 | 1788.45858 | 1102.66449 | 1359.35596 | Non-sig. |
| PF3D7_1302500 | N/A | 1864.51651 | -0.302360896 | 0.50086282 | -0.60368006 | 0.54605637 | 0.94142878 | 1283.00529 | 1578.74934 | 2835.04493 | 1761.26649 | Non-sig. |
| PF3D7_1245400 | N/A | 587.797125 | -0.302422965 | 0.45997496 | -0.65747702 | 0.51087425 | 0.93247207 | 728.755873 | 630.268879 | 569.736871 | 422.426877 | Non-sig. |
| PF3D7_0914800 | N/A | 719.387272 | -0.302427301 | 0.44570904 | -0.67853077 | 0.49743522 | 0.93170321 | 641.913966 | 633.111228 | 946.904896 | 655.619 | Non-sig. |
| PF3D7_1447100 | N/A | 552.659784 | -0.30255955 | 0.47254828 | -0.64027224 | 0.52199562 | 0.93247207 | 666.898628 | 619.779902 | 553.999363 | 369.961243 | Non-sig. |
| PF3D7_1108400 | CK2alpha | 4905.66746 | -0.302711998 | 0.47017891 | -0.643823 | 0.51969022 | 0.93247207 | 3776.41572 | 4743.09373 | 7060.19784 | 4042.96257 | Non-sig. |
| PF3D7_0931500 | N/A | 2246.61871 | -0.302742444 | 0.47615467 | -0.63580694 | 0.52490226 | 0.93247207 | 3255.03136 | 2172.92672 | 1708.22602 | 1850.29074 | Non-sig. |
| PF3D7_1363300 | N/A | 167.796654 | -0.302949365 | 0.49516652 | -0.6118131 | 0.54066142 | 0.93792541 | 190.968987 | 176.126434 | 179.763125 | 124.328071 | Non-sig. |
| PF3D7_0711300 | N/A | 272.036492 | -0.303139839 | 0.47091453 | -0.64372582 | 0.51975324 | 0.93247207 | 246.317895 | 250.303281 | 354.534374 | 236.990419 | Non-sig. |
| PF3D7_0907800 | N/A | 1630.93944 | -0.303162867 | 0.42237327 | -0.71776054 | 0.47290495 | 0.92682482 | 1992.35546 | 1585.42275 | 1611.11912 | 1334.86042 | Non-sig. |
| PF3D7_0906700 | LRR9 | 347.972749 | -0.303960988 | 0.44149502 | -0.68848112 | 0.49114985 | 0.93170321 | 409.352522 | 310.117018 | 359.709012 | 312.712444 | Non-sig. |
| PF3D7_0906000 | RNaseII | 1140.49303 | -0.30411695 | 0.43914412 | -0.69252197 | 0.48860959 | 0.93170321 | 1480.72949 | 937.466611 | 1039.97907 | 1103.79696 | Non-sig. |
| PF3D7_0407300 | N/A | 164.370241 | -0.304974162 | 0.48011443 | -0.6352114 | 0.52529055 | 0.93247207 | 186.884959 | 140.662463 | 176.498613 | 153.434929 | Non-sig. |
| PF3D7_1335800 | N/A | 1215.41591 | -0.305110448 | 0.48124299 | -0.63400497 | 0.52607758 | 0.93247207 | 1776.52823 | 1111.84763 | 910.694691 | 1062.59309 | Non-sig. |
| PF3D7_0422700 | EIF4A3 | 520.750298 | -0.305296119 | 0.53937887 | -0.56601424 | 0.5713841 | 0.94850402 | 328.150825 | 428.218118 | 822.820028 | 503.812222 | Non-sig. |
| PF3D7_0819100 | N/A | 223.671345 | -0.305798685 | 0.46954846 | -0.65126119 | 0.5148779 | 0.93247207 | 217.722207 | 209.277705 | 276.73838 | 190.947087 | Non-sig. |
| PF3D7_1422000 | COX14 | 149.05821 | -0.305817059 | 0.51241708 | -0.59681277 | 0.55063239 | 0.94170064 | 137.803705 | 151.960518 | 191.674315 | 114.794302 | Non-sig. |
| PF3D7_0409500 | N/A | 921.638592 | -0.305861187 | 0.5336526 | -0.57314662 | 0.56654542 | 0.9476402 | 568.674115 | 839.401854 | 1468.93698 | 809.541418 | Non-sig. |
| PF3D7_1024800 | EXP3 | 5706.74307 | -0.306430029 | 0.46918428 | -0.6531123 | 0.51368388 | 0.93247207 | 4967.84193 | 3920.23209 | 7653.0177 | 6285.88057 | Non-sig. |
| PF3D7_0726700 | N/A | 701.206171 | -0.306572778 | 0.45222238 | -0.67792483 | 0.49781936 | 0.93170321 | 686.373661 | 742.236655 | 864.381899 | 511.832467 | Non-sig. |
| PF3D7_1238700 | N/A | 2028.2762 | -0.30657559 | 0.46254001 | -0.6628088 | 0.50745301 | 0.93170321 | 2862.49173 | 1922.707 | 1623.74037 | 1704.16568 | Non-sig. |
| PF3D7_1314600 | LipL1 | 215.874099 | -0.306977714 | 0.54815565 | -0.56001925 | 0.57546631 | 0.94896603 | 158.699456 | 235.085752 | 318.575714 | 151.135473 | Non-sig. |
| PF3D7_1454500 | ISCU | 950.248108 | -0.307398065 | 0.43410751 | -0.70811507 | 0.4788738 | 0.92880667 | 1200.88622 | 917.885353 | 901.484336 | 780.736521 | Non-sig. |
| PF3D7_0820300 | N/A | 3140.26187 | -0.307815731 | 0.41520238 | -0.74136311 | 0.4584733 | 0.92288312 | 3601.65138 | 2486.64182 | 3346.39829 | 3126.356 | Non-sig. |
| PF3D7_1409700 | N/A | 293.462158 | -0.307863365 | 0.51674448 | -0.59577485 | 0.55132565 | 0.94170064 | 213.730805 | 254.280645 | 435.303427 | 270.533757 | Non-sig. |
| PF3D7_0411100 | MED8 | 153.69349 | -0.307984592 | 0.50250863 | -0.61289414 | 0.53994634 | 0.93711173 | 140.680321 | 131.272135 | 199.248436 | 143.57307 | Non-sig. |
| PF3D7_1353600 | ERD2 | 1932.41978 | -0.308282393 | 0.45536053 | -0.67700728 | 0.49840134 | 0.93170321 | 1665.34055 | 1482.02931 | 2610.63249 | 1971.67678 | Non-sig. |
| PF3D7_1448900 | SDHC | 169.125333 | -0.308749012 | 0.49351042 | -0.62561802 | 0.53156552 | 0.93351492 | 156.77678 | 156.967395 | 217.392287 | 145.36487 | Non-sig. |
| PF3D7_1467500 | N/A | 1596.81368 | -0.308789144 | 0.47713055 | -0.64717957 | 0.51751573 | 0.93247207 | 2320.65997 | 1508.874 | 1213.74323 | 1343.97751 | Non-sig. |
| PF3D7_1342200 | N/A | 108.335318 | -0.309064178 | 0.52271542 | -0.59126662 | 0.55434179 | 0.94243456 | 111.606061 | 86.9054332 | 128.127604 | 106.702172 | Non-sig. |
| PF3D7_1443600 | N/A | 744.371556 | -0.309188049 | 0.57502927 | -0.53769098 | 0.59079041 | 0.95315309 | 422.560383 | 519.164785 | 1224.78247 | 810.97859 | Non-sig. |
| PF3D7_0532300 | N/A | 940.821788 | -0.309881046 | 0.46677328 | -0.66387914 | 0.50676767 | 0.93170321 | 757.048951 | 870.119821 | 1325.63492 | 810.483461 | Non-sig. |
| PF3D7_1340100 | RRP42 | 299.437851 | -0.310001385 | 0.47001319 | -0.65955891 | 0.50953693 | 0.93247207 | 267.010361 | 262.979158 | 395.771768 | 271.990118 | Non-sig. |
| PF3D7_1404200 | N/A | 49.5234949 | -0.310401984 | 0.75381329 | -0.41177568 | 0.68050385 | 0.97982498 | 78.6432276 | 62.3003391 | 31.1880787 | 25.9623341 | Non-sig. |
| PF3D7_1452700 | N/A | 1153.33543 | -0.310549276 | 0.43196654 | -0.71891975 | 0.47219037 | 0.92682482 | 1219.0869 | 1192.90945 | 1334.8481 | 866.497264 | Non-sig. |
| PF3D7_1405500 | CBWD1 | 210.927093 | -0.310672561 | 0.46284066 | -0.67123006 | 0.50207398 | 0.93170321 | 231.409179 | 182.444216 | 235.681698 | 194.173278 | Non-sig. |
| PF3D7_1305500 | MKP1 | 105.922386 | -0.310770482 | 0.53927698 | -0.57627248 | 0.56443102 | 0.94661461 | 98.9462965 | 107.298321 | 135.513574 | 81.9313523 | Non-sig. |
| PF3D7_0721300 | DDX31 | 799.961047 | -0.311085251 | 0.45659319 | -0.6813182 | 0.49567017 | 0.93170321 | 1090.2305 | 673.699724 | 681.758261 | 754.155704 | Non-sig. |
| PF3D7_1458700 | N/A | 969.402611 | -0.311440965 | 0.58149609 | -0.53558566 | 0.59224494 | 0.95322751 | 518.159123 | 702.700927 | 1628.77498 | 1027.97541 | Non-sig. |
| PF3D7_0627000 | N/A | 56.7334938 | -0.311549932 | 0.68476273 | -0.45497502 | 0.64912721 | 0.97317009 | 90.5592297 | 54.7163795 | 35.3179675 | 46.3403984 | Non-sig. |
| PF3D7_0621700 | N/A | 602.877316 | -0.311924898 | 0.42253623 | -0.73822048 | 0.46038048 | 0.92336312 | 670.222147 | 528.812759 | 665.379385 | 547.094974 | Non-sig. |
| PF3D7_0622200 | MIAB | 1042.31045 | -0.311933209 | 0.46051174 | -0.67736212 | 0.49817623 | 0.93170321 | 1437.95668 | 846.570909 | 871.401552 | 1013.31266 | Non-sig. |
| PF3D7_1137700 | N/A | 28.4376357 | -0.312066361 | 0.84667765 | -0.36857753 | 0.71244264 | 0.9875928 | 48.3832765 | 22.5375252 | 14.7813086 | 28.0484324 | Non-sig. |
| PF3D7_0912400 | N/A | 2148.87404 | -0.312373386 | 0.50484842 | -0.61874688 | 0.53608312 | 0.93549683 | 1453.27315 | 1834.56772 | 3307.61377 | 2000.04151 | Non-sig. |
| PF3D7_0820200 | PGPS | 2126.16107 | -0.312667883 | 0.42399829 | -0.73742723 | 0.46086258 | 0.92336312 | 2003.09416 | 1843.12339 | 2708.04619 | 1950.38052 | Non-sig. |
| PF3D7_1037800 | N/A | 57.9660561 | -0.313311286 | 0.6695209 | -0.46796342 | 0.63981075 | 0.96987621 | 88.4134472 | 61.1613364 | 40.2428962 | 42.0465445 | Non-sig. |
| PF3D7_1429100 | N/A | 2630.0929 | -0.313338553 | 0.60221028 | -0.52031419 | 0.60284461 | 0.95693578 | 4591.03834 | 2741.55455 | 1238.48879 | 1949.28994 | Non-sig. |
| PF3D7_0904400 | SPC3 | 910.467486 | -0.313808798 | 0.49130184 | -0.63872913 | 0.52299916 | 0.93247207 | 665.788223 | 835.334753 | 1352.11218 | 788.634789 | Non-sig. |
| PF3D7_1225700 | N/A | 1397.79241 | -0.313882762 | 0.41442338 | -0.75739636 | 0.44881244 | 0.92081798 | 1430.88974 | 1216.4202 | 1667.51879 | 1276.3409 | Non-sig. |
| PF3D7_0609300 | N/A | 443.078388 | -0.313895543 | 0.55148067 | -0.56918685 | 0.56922935 | 0.94850402 | 676.916548 | 286.476778 | 305.52796 | 503.392266 | Non-sig. |

| | | | | | | | | | | | | |
|---|---|---|---|---|---|---|---|---|---|---|---|---|
| PF3D7_0401600 | RIF | 933.232268 | -0.3139025 | 0.41725882 | -0.75229686 | 0.45187255 | 0.92081798 | 1097.0676 | 820.790752 | 971.737351 | 843.33337 | Non-sig. |
| PF3D7_1355600 | PIP1 | 157.594632 | -0.314489953 | 0.55724048 | -0.56437026 | 0.57250218 | 0.94850402 | 237.956898 | 127.675226 | 111.701345 | 153.04506 | Non-sig. |
| PF3D7_0816100 | PPCDC | 1650.4875 | -0.314728372 | 0.42552221 | -0.73962854 | 0.45952542 | 0.92288312 | 1815.20202 | 1693.61979 | 1844.40947 | 1248.71872 | Non-sig. |
| PF3D7_0527900 | DDX41 | 923.147973 | -0.314835482 | 0.4245064 | -0.74165073 | 0.45829898 | 0.92288312 | 914.348389 | 856.340298 | 1132.48299 | 789.420209 | Non-sig. |
| PF3D7_1129000 | SpdSyn | 1696.58104 | -0.314878538 | 0.46820428 | -0.67252384 | 0.50125027 | 0.93170321 | 1410.64462 | 1267.42532 | 2351.09798 | 1757.15627 | Non-sig. |
| PF3D7_1110100 | N/A | 1690.19423 | -0.315375706 | 0.43999968 | -0.71676349 | 0.47352005 | 0.92682482 | 1799.34417 | 1197.3713 | 1949.01035 | 1815.05107 | Non-sig. |
| PF3D7_1363800 | GEX1 | 139.360319 | -0.315525374 | 0.5288904 | -0.59657989 | 0.5507879 | 0.94170064 | 139.855045 | 154.554129 | 169.156293 | 93.8758103 | Non-sig. |
| PF3D7_1334000 | N/A | 482.704206 | -0.315625385 | 0.50651584 | -0.62313034 | 0.53319887 | 0.93467024 | 724.37543 | 432.559859 | 346.512584 | 427.368952 | Non-sig. |
| PF3D7_1009000 | DPH5 | 346.162756 | -0.315920192 | 0.4883936 | -0.64685571 | 0.51772533 | 0.93247207 | 286.56224 | 344.698559 | 481.042206 | 272.34802 | Non-sig. |
| PF3D7_1469000 | IF1 | 361.220159 | -0.315950917 | 0.50177808 | -0.62966265 | 0.52891532 | 0.93270419 | 273.077141 | 333.905098 | 527.887723 | 310.010673 | Non-sig. |
| PF3D7_0915600 | N/A | 112.500152 | -0.316219455 | 0.59676397 | -0.52989032 | 0.59618798 | 0.95541558 | 166.862006 | 125.645584 | 82.9067744 | 74.5862446 | Non-sig. |
| PF3D7_1032300 | N/A | 338.084168 | -0.316360517 | 0.49235624 | -0.64254394 | 0.52052007 | 0.93247207 | 472.644429 | 339.059066 | 277.599347 | 263.033828 | Non-sig. |
| PF3D7_0805900 | N/A | 66.7806274 | -0.316913583 | 0.64341107 | -0.49255227 | 0.62232898 | 0.96429926 | 99.1403424 | 72.0631386 | 49.2093111 | 46.7097175 | Non-sig. |
| PF3D7_0214400 | N/A | 748.668407 | -0.3171221 | 0.65490232 | -0.48422809 | 0.628224 | 0.96655174 | 1381.53375 | 620.254561 | 280.041664 | 712.843655 | Non-sig. |
| PF3D7_0617000 | TOM40 | 233.798419 | -0.317199745 | 0.51760242 | -0.61282509 | 0.539992 | 0.93711173 | 181.891327 | 183.540228 | 336.662054 | 233.100066 | Non-sig. |
| PF3D7_1001800 | N/A | 1531.63469 | -0.317748738 | 0.42490797 | -0.74780602 | 0.45457717 | 0.92203286 | 1458.8704 | 1409.41484 | 1940.21434 | 1318.03916 | Non-sig. |
| PF3D7_1359300 | DIS3 | 1457.81449 | -0.318324928 | 0.4160157 | -0.76517528 | 0.44416718 | 0.91974025 | 1471.52594 | 1261.58311 | 1764.35295 | 1333.79595 | Non-sig. |
| PF3D7_0936100 | ETRAMP9 | 702.76568 | -0.319008961 | 0.46616985 | -0.68431916 | 0.49377365 | 0.93170321 | 862.412434 | 775.953023 | 697.996024 | 474.701238 | Non-sig. |
| PF3D7_0801800 | N/A | 5276.49314 | -0.319070892 | 0.44658892 | -0.71446218 | 0.47494143 | 0.92682482 | 5093.66031 | 3706.28785 | 6621.41665 | 5684.60777 | Non-sig. |
| PF3D7_0929300 | N/A | 770.195542 | -0.319152079 | 0.51132022 | -0.62417262 | 0.53251423 | 0.93450161 | 1181.52728 | 724.361098 | 528.869355 | 646.024433 | Non-sig. |
| PF3D7_1458400 | ADCL | 229.63261 | -0.319554242 | 0.55180391 | -0.57910834 | 0.56251608 | 0.94519828 | 152.451127 | 220.034069 | 357.18976 | 188.855484 | Non-sig. |
| PF3D7_0102800 | N/A | 394.12752 | -0.319979701 | 0.43997979 | -0.72726 | 0.4670667 | 0.92538923 | 421.950053 | 379.874195 | 453.32621 | 321.359622 | Non-sig. |
| PF3D7_1129700 | N/A | 744.317534 | -0.320290533 | 0.41996805 | -0.76265453 | 0.44566945 | 0.91974025 | 785.07122 | 663.147203 | 868.072716 | 660.978996 | Non-sig. |
| PF3D7_0729100 | N/A | 3447.53889 | -0.321720741 | 0.48238574 | -0.66693668 | 0.50481258 | 0.93170321 | 3126.65296 | 2166.53149 | 4533.88269 | 3963.08844 | Non-sig. |
| PF3D7_1329400 | AMPD | 395.547186 | -0.32380792 | 0.54966717 | -0.58909816 | 0.55579543 | 0.94243456 | 251.685407 | 400.173871 | 627.516457 | 302.813008 | Non-sig. |
| PF3D7_1006000 | IMP4 | 106.793388 | -0.323928206 | 0.7013809 | -0.46184349 | 0.64419356 | 0.97190751 | 193.400517 | 97.6814231 | 44.3385976 | 91.7530142 | Non-sig. |
| PF3D7_0805000 | N/A | 382.097695 | -0.324213391 | 0.5573582 | -0.58169664 | 0.56077104 | 0.94472957 | 230.259345 | 358.492093 | 619.131545 | 320.507797 | Non-sig. |
| PF3D7_0817500 | HINT1 | 6506.88959 | -0.324297399 | 0.4606889 | -0.70394012 | 0.48147006 | 0.92880667 | 8791.86288 | 6914.34264 | 5678.67325 | 4642.67959 | Non-sig. |
| PF3D7_1367600 | N/A | 227.247121 | -0.324539481 | 0.48480534 | -0.66942224 | 0.50322617 | 0.93170321 | 216.304203 | 231.925495 | 288.970256 | 171.788531 | Non-sig. |
| PF3D7_1345800 | N/A | 1237.97894 | -0.325109767 | 0.50759784 | -0.6404869 | 0.5218561 | 0.93247207 | 842.820465 | 1057.70546 | 1910.61738 | 1140.77246 | Non-sig. |
| PF3D7_1419800 | GR | 941.801303 | -0.325644782 | 0.5112185 | -0.63699725 | 0.52412663 | 0.93247207 | 642.103741 | 783.474664 | 1453.04053 | 888.586278 | Non-sig. |
| PF3D7_1247500 | N/A | 1611.76824 | -0.325902151 | 0.41094735 | -0.79305085 | 0.42774819 | 0.91236574 | 1846.58026 | 1354.04611 | 1739.54833 | 1506.89826 | Non-sig. |
| PF3D7_0715800 | DMT1 | 734.646191 | -0.325974458 | 0.56362059 | -0.57835796 | 0.56302247 | 0.94574922 | 409.079858 | 634.302768 | 1225.16723 | 670.034912 | Non-sig. |
| PF3D7_1134100 | PDI-11 | 1803.92014 | -0.326100118 | 0.58798283 | -0.55460823 | 0.57916266 | 0.95129458 | 880.169351 | 1670.53931 | 3133.34933 | 1531.62255 | Non-sig. |
| PF3D7_0826300 | N/A | 1333.09186 | -0.3267895 | 0.43108694 | -0.75805939 | 0.44841543 | 0.92081798 | 1646.34542 | 1332.63433 | 1320.64803 | 1032.73965 | Non-sig. |
| PF3D7_1119800 | ASF1 | 1104.32132 | -0.327170685 | 0.41678406 | -0.78498848 | 0.43246032 | 0.91370017 | 1180.6961 | 1045.95557 | 1277.26995 | 913.363654 | Non-sig. |
| PF3D7_0929700 | N/A | 245.132348 | -0.327549438 | 0.55881283 | -0.58615233 | 0.55777316 | 0.94272305 | 155.430631 | 209.36674 | 389.957852 | 225.774168 | Non-sig. |
| PF3D7_1419700 | EIF3H | 2350.74847 | -0.327699121 | 0.48919383 | -0.66987582 | 0.50293696 | 0.93170321 | 3521.16165 | 2262.0122 | 1712.32183 | 1907.49819 | Non-sig. |
| PF3D7_1463200 | RFC3 | 1285.23654 | -0.327909657 | 0.47706617 | -0.68734629 | 0.49186454 | 0.93170321 | 997.888932 | 1237.51474 | 1863.17477 | 1042.36774 | Non-sig. |
| PF3D7_1202100 | SRAP1 | 1151.20311 | -0.328032413 | 0.54609115 | -0.60069168 | 0.54804536 | 0.94142878 | 1847.2993 | 1216.07472 | 716.066435 | 825.371988 | Non-sig. |
| PF3D7_0930100 | N/A | 2324.77217 | -0.328041392 | 0.44275511 | -0.74090933 | 0.45874842 | 0.92288312 | 3134.19775 | 1899.9802 | 2041.9199 | 2222.99082 | Non-sig. |
| PF3D7_1454700 | 6PGD | 3180.53546 | -0.328280998 | 0.52788595 | -0.62187864 | 0.53402167 | 0.93467024 | 1988.80111 | 3092.62232 | 5092.55509 | 2548.16331 | Non-sig. |
| PF3D7_0205700 | N/A | 373.353544 | -0.328319698 | 0.46017899 | -0.71346086 | 0.47556063 | 0.9269156 | 482.230199 | 363.835666 | 349.239465 | 298.108847 | Non-sig. |
| PF3D7_1026000 | N/A | 1227.95237 | -0.328615088 | 0.58463835 | -0.56208268 | 0.57405969 | 0.94850402 | 2114.82514 | 1182.90619 | 619.920955 | 994.157214 | Non-sig. |
| PF3D7_0502900 | N/A | 949.90501 | -0.329178801 | 0.43230656 | -0.76144761 | 0.44638976 | 0.91974025 | 1122.79665 | 962.739494 | 992.894133 | 721.189766 | Non-sig. |
| PF3D7_1029200 | N/A | 110.457713 | -0.329259557 | 0.51410931 | -0.6404466 | 0.5218823 | 0.93247207 | 129.439763 | 97.6483719 | 116.618682 | 98.1240348 | Non-sig. |
| PF3D7_1134500 | N/A | 1556.97473 | -0.329457035 | 0.46268709 | -0.7120515 | 0.47643289 | 0.92731133 | 2153.04364 | 1185.25245 | 1315.16313 | 1574.4397 | Non-sig. |
| PF3D7_1446900 | N/A | 1644.60965 | -0.329567715 | 0.44265459 | -0.74452569 | 0.4565585 | 0.92287178 | 1446.14571 | 1535.17053 | 2216.91398 | 1380.2084 | Non-sig. |
| PF3D7_1206700 | EIF5 | 5939.71424 | -0.329927139 | 0.40787881 | -0.80888522 | 0.41858117 | 0.9088998 | 6674.14628 | 4803.21663 | 6557.7481 | 5723.74596 | Non-sig. |
| PF3D7_0629400 | PABP3 | 1643.68219 | -0.330010969 | 0.51702003 | -0.63829436 | 0.52328208 | 0.93247207 | 1071.45164 | 1406.33467 | 2589.93655 | 1507.00588 | Non-sig. |
| PF3D7_0516000 | N/A | 464.250441 | -0.33045099 | 0.44837488 | -0.73699711 | 0.46112411 | 0.92336312 | 598.373099 | 390.476795 | 436.169017 | 431.982854 | Non-sig. |
| PF3D7_0218800 | N/A | 296.219966 | -0.33065422 | 0.45385523 | -0.72854558 | 0.46627968 | 0.92538923 | 301.72488 | 282.032931 | 358.231066 | 242.890987 | Non-sig. |
| PF3D7_0928000 | COX6B | 2396.62297 | -0.33079316 | 0.48110136 | -0.68757477 | 0.4917206 | 0.93170321 | 3390.8197 | 2559.46056 | 1950.09591 | 1686.11571 | Non-sig. |
| PF3D7_1208700 | N/A | 375.805474 | -0.330822435 | 0.45314488 | -0.73005887 | 0.4653542 | 0.92538923 | 479.660553 | 353.297969 | 357.902941 | 312.360434 | Non-sig. |

| | | | | | | | | | | | | |
|---|---|---|---|---|---|---|---|---|---|---|---|---|
| PF3D7_0405600 | N/A | 885.598136 | -0.330978016 | 0.50296474 | -0.65805411 | 0.51050336 | 0.93247207 | 631.576871 | 855.566258 | 1341.60536 | 713.644062 | Non-sig. |
| PF3D7_0509800 | PI4KB | 6173.08915 | -0.331794 | 0.45641015 | -0.72696456 | 0.46724767 | 0.92538923 | 7877.34027 | 4168.27056 | 5882.46256 | 6764.28321 | Non-sig. |
| PF3D7_0524500 | N/A | 376.214307 | -0.331805531 | 0.44326919 | -0.74854183 | 0.45413341 | 0.92203286 | 376.21013 | 332.780772 | 462.253676 | 333.612651 | Non-sig. |
| PF3D7_1115900 | DHHC9 | 749.770618 | -0.331928756 | 0.45794797 | -0.72481761 | 0.46856393 | 0.92594245 | 645.260739 | 711.386448 | 1025.79796 | 616.637327 | Non-sig. |
| PF3D7_1465200 | MED4 | 554.062438 | -0.33214947 | 0.44982092 | -0.73840379 | 0.46026911 | 0.92336312 | 679.31669 | 569.913456 | 555.919533 | 411.100073 | Non-sig. |
| PF3D7_1346100 | SEC61A | 5366.5163 | -0.332191246 | 0.51805921 | -0.64122255 | 0.5213781 | 0.93247207 | 3443.3076 | 4450.74072 | 8519.64356 | 5052.3733 | Non-sig. |
| PF3D7_1449400 | MUS81 | 2482.15598 | -0.332302203 | 0.46918615 | -0.70825237 | 0.47878855 | 0.92880667 | 3606.81714 | 2192.7248 | 1927.00342 | 2202.07856 | Non-sig. |
| PF3D7_1330600 | N/A | 1293.50088 | -0.332852333 | 0.44801797 | -0.74294416 | 0.45751549 | 0.92288312 | 1113.50462 | 1149.51898 | 1770.3713 | 1140.60861 | Non-sig. |
| PF3D7_1318300 | N/A | 1216.85496 | -0.333179176 | 0.50641166 | -0.65792161 | 0.5105885 | 0.93247207 | 1139.38823 | 692.889147 | 1573.96299 | 1461.17947 | Non-sig. |
| PF3D7_1410600 | eIF2gamma | 6234.88152 | -0.333518982 | 0.49642519 | -0.67184138 | 0.50168468 | 0.93170321 | 4318.45342 | 5595.00278 | 9585.96964 | 5440.10023 | Non-sig. |
| PF3D7_1470600 | N/A | 607.612488 | -0.334040231 | 0.50286583 | -0.66427308 | 0.50651555 | 0.93170321 | 917.080732 | 560.235296 | 438.503955 | 514.629968 | Non-sig. |
| PF3D7_1234300 | N/A | 325.364694 | -0.334576622 | 0.54279177 | -0.61639959 | 0.53763083 | 0.93646681 | 212.355455 | 273.709874 | 513.193193 | 302.200254 | Non-sig. |
| PF3D7_0930700 | N/A | 194.057819 | -0.335320735 | 0.47540911 | -0.7053309 | 0.48060434 | 0.92880667 | 233.541182 | 185.657498 | 199.553544 | 157.479053 | Non-sig. |
| PF3D7_0519300 | COX10 | 618.910927 | -0.335493274 | 0.46980487 | -0.71411196 | 0.47515795 | 0.92682482 | 513.594893 | 585.875705 | 867.262155 | 508.910955 | Non-sig. |
| PF3D7_1472400 | N/A | 1115.73954 | -0.335544138 | 0.42685953 | -0.78607625 | 0.43182281 | 0.91370017 | 1127.02327 | 885.052029 | 1362.67933 | 1088.20354 | Non-sig. |
| PF3D7_1215900 | SR10 | 371.64611 | -0.335762263 | 0.54463446 | -0.61649104 | 0.53757048 | 0.93646681 | 245.190585 | 280.071079 | 583.911205 | 377.411571 | Non-sig. |
| PF3D7_1216100 | N/A | 153.113145 | -0.33589866 | 0.5056951 | -0.66423159 | 0.5065421 | 0.93170321 | 142.220051 | 148.344539 | 199.36221 | 122.525781 | Non-sig. |
| PF3D7_0528800 | N/A | 335.914625 | -0.336215459 | 0.44243533 | -0.75992001 | 0.4473024 | 0.91995656 | 396.690198 | 291.881976 | 353.133616 | 301.952711 | Non-sig. |
| PF3D7_1014300 | N/A | 3935.45063 | -0.336264326 | 0.46492845 | -0.72326038 | 0.46951993 | 0.92682482 | 5690.54475 | 3571.53054 | 3093.80091 | 3385.92634 | Non-sig. |
| PF3D7_1432600 | N/A | 573.655488 | -0.336281983 | 0.43425727 | -0.77438423 | 0.43870361 | 0.9169936 | 686.65542 | 458.09865 | 593.84826 | 556.01962 | Non-sig. |
| PF3D7_0210300 | N/A | 663.090952 | -0.336380403 | 0.61262255 | -0.54908263 | 0.58294874 | 0.95129458 | 302.969993 | 609.558894 | 1176.78947 | 563.045451 | Non-sig. |
| PF3D7_0410900 | N/A | 710.178905 | -0.336504059 | 0.49648716 | -0.67776991 | 0.49791759 | 0.93170321 | 1031.08781 | 514.951427 | 554.427483 | 740.248903 | Non-sig. |
| PF3D7_0909300 | ARP | 735.824791 | -0.336913957 | 0.42817789 | -0.78685511 | 0.43136669 | 0.91355742 | 853.815402 | 720.436999 | 788.945842 | 580.10092 | Non-sig. |
| PF3D7_0625100 | SMS2 | 34.7565493 | -0.336959361 | 0.69889375 | -0.48213246 | 0.62971185 | 0.96682409 | 39.85251 | 33.2969088 | 37.7550967 | 28.1216816 | Non-sig. |
| PF3D7_0915300 | N/A | 214.714866 | -0.336995893 | 0.4719757 | -0.71401111 | 0.47522031 | 0.92682482 | 218.930013 | 208.16275 | 260.347207 | 171.419495 | Non-sig. |
| PF3D7_0423700 | ETRAMP4 | 1971.05325 | -0.337076681 | 0.4917035 | -0.68552833 | 0.49301059 | 0.93170321 | 1567.20424 | 1352.24986 | 2833.08631 | 2131.67258 | Non-sig. |
| PF3D7_0907700 | PA28 | 3475.5236 | -0.337584942 | 0.44611349 | -0.75672435 | 0.44921503 | 0.92081798 | 2947.58614 | 3022.35877 | 4812.77573 | 3119.37375 | Non-sig. |
| PF3D7_0727200 | NFS1 | 299.411821 | -0.337919563 | 0.47298945 | -0.71443362 | 0.47495909 | 0.92682482 | 265.199348 | 267.190087 | 403.239349 | 262.0185 | Non-sig. |
| PF3D7_1364800 | RPB5 | 445.601863 | -0.338411908 | 0.43093669 | -0.7852938 | 0.43228133 | 0.91370017 | 482.103139 | 395.099237 | 513.113325 | 392.091749 | Non-sig. |
| PF3D7_0504400 | N/A | 1721.09039 | -0.338845235 | 0.44846648 | -0.75556423 | 0.44991051 | 0.92081798 | 1466.81941 | 1517.35335 | 2377.49619 | 1522.69262 | Non-sig. |
| PF3D7_1433400 | PHD2 | 33729.1098 | -0.338997716 | 0.42544398 | -0.79680928 | 0.4255618 | 0.91236574 | 39116.8271 | 24489.9393 | 36230.7091 | 35078.964 | Non-sig. |
| PF3D7_1362300 | TMCO1 | 220.105288 | -0.339099145 | 0.46084111 | -0.7358266 | 0.46183623 | 0.92402128 | 236.341297 | 192.177717 | 255.328275 | 196.573863 | Non-sig. |
| PF3D7_1417500 | CBF5 | 541.251117 | -0.339104676 | 0.48512633 | -0.69900283 | 0.48455026 | 0.93113794 | 422.076041 | 482.850376 | 786.791992 | 473.286057 | Non-sig. |
| PF3D7_1475200 | N/A | 673.664761 | -0.339230042 | 0.42287621 | -0.80219703 | 0.42243899 | 0.91161836 | 730.494388 | 562.2484 | 774.476157 | 627.440099 | Non-sig. |
| PF3D7_1128400 | FPPS/GGPPS | 1329.62725 | -0.33929651 | 0.42630744 | -0.79589629 | 0.42609232 | 0.91236574 | 1312.84964 | 1278.72276 | 1657.54631 | 1069.39028 | Non-sig. |
| PF3D7_0915400 | PFK9 | 9394.76193 | -0.339661328 | 0.61334428 | -0.55378576 | 0.57972548 | 0.95129458 | 4037.45175 | 8166.92249 | 16953.4375 | 8421.23597 | Non-sig. |
| PF3D7_1303900 | N/A | 22.1440817 | -0.340020418 | 1.05168571 | -0.32330992 | 0.74646054 | 0.98998709 | 44.7181796 | 23.6850446 | 4.91814116 | 15.2549613 | Non-sig. |
| PF3D7_0825700 | CryPH | 21.4645642 | -0.340313211 | 0.91678789 | -0.37120169 | 0.71048732 | 0.9875928 | 36.6166941 | 21.4966779 | 11.4866572 | 16.2582274 | Non-sig. |
| PF3D7_0608000 | DPH7 | 398.607404 | -0.340889229 | 0.45578076 | -0.74792369 | 0.45450619 | 0.92203286 | 409.947583 | 408.187413 | 480.943507 | 295.351114 | Non-sig. |
| PF3D7_1140100 | N/A | 1586.6915 | -0.341117959 | 0.52262718 | -0.65269846 | 0.51395069 | 0.93247207 | 1016.94355 | 1455.76611 | 2529.53994 | 1344.5164 | Non-sig. |
| PF3D7_0519800 | N/A | 1413.32202 | -0.34118118 | 0.43096234 | -0.79167284 | 0.42855146 | 0.91236574 | 1634.85094 | 1040.58144 | 1524.51522 | 1453.34048 | Non-sig. |
| PF3D7_1028000 | N/A | 404.098908 | -0.341316958 | 0.4638541 | -0.73582827 | 0.46183521 | 0.92402128 | 509.291241 | 417.43125 | 394.197367 | 295.475774 | Non-sig. |
| PF3D7_0107900 | N/A | 204.164701 | -0.341665274 | 0.46795865 | -0.73011851 | 0.46531775 | 0.92538923 | 212.668734 | 174.055368 | 243.720459 | 186.214243 | Non-sig. |
| PF3D7_1239800 | N/A | 10254.2452 | -0.34260854 | 0.48710559 | -0.7033558 | 0.48183405 | 0.92880667 | 13179.7179 | 6062.45278 | 9752.69169 | 12022.1185 | Non-sig. |
| PF3D7_1405700 | N/A | 1916.19171 | -0.343508683 | 0.49537023 | -0.69343829 | 0.48803454 | 0.93170321 | 2789.43035 | 1317.48519 | 1497.31713 | 2060.53419 | Non-sig. |
| PF3D7_0623900 | N/A | 595.695701 | -0.343530174 | 0.42961923 | -0.79961546 | 0.42393363 | 0.91236574 | 603.521978 | 535.850574 | 728.942137 | 514.468114 | Non-sig. |
| PF3D7_0904500 | N/A | 299.157159 | -0.343595868 | 0.46358829 | -0.74116597 | 0.45859281 | 0.92288312 | 285.688934 | 285.588545 | 383.397848 | 241.953309 | Non-sig. |
| PF3D7_1323200 | N/A | 588.29211 | -0.343859685 | 0.44540779 | -0.77201093 | 0.44010795 | 0.91706217 | 542.890261 | 528.553026 | 773.06746 | 508.65769 | Non-sig. |
| PF3D7_1402700 | SR140 | 1390.2325 | -0.343987865 | 0.41765366 | -0.8236199 | 0.41015557 | 0.90748263 | 1415.91598 | 1286.4706 | 1694.35998 | 1164.18344 | Non-sig. |
| PF3D7_0211400 | KASIII | 106.330837 | -0.344449438 | 0.52868577 | -0.65152017 | 0.51471076 | 0.93247207 | 123.860522 | 107.383105 | 114.103617 | 79.976106 | Non-sig. |
| PF3D7_1369600 | N/A | 1049.72412 | -0.344760344 | 0.50560559 | -0.68187605 | 0.49531733 | 0.93170321 | 1619.27252 | 963.48154 | 730.157704 | 885.984727 | Non-sig. |
| PF3D7_1420800 | N/A | 8.59978925 | -0.344844663 | 1.23436252 | -0.27937065 | 0.77996039 | 0.99028237 | 9.39635806 | 4.30177696 | 9.84253907 | 10.8584829 | Non-sig. |
| PF3D7_0722400 | OLA1 | 2975.12583 | -0.345542964 | 0.47980339 | -0.72017617 | 0.47141653 | 0.92682482 | 2261.28834 | 2873.41052 | 4397.86657 | 2367.93786 | Non-sig. |
| PF3D7_0905600 | WDR66 | 71.4164971 | -0.345626555 | 0.57131821 | -0.60496331 | 0.54520338 | 0.94121391 | 76.8764981 | 55.7879313 | 82.9573717 | 70.0441873 | Non-sig. |

| | | | | | | | | | | | | |
|---|---|---|---|---|---|---|---|---|---|---|---|---|
| PF3D7_1464200 | ZFP | 74.5664803 | -0.345698861 | 0.5675883 | -0.60906622 | 0.54248054 | 0.93953127 | 72.426007 | 64.4300895 | 94.4020644 | 67.0077603 | Non-sig. |
| PF3D7_1434200 | CAM | 12265.05 | -0.34600102 | 0.4289983 | -0.80653239 | 0.41993594 | 0.90957507 | 12231.9915 | 12523.3997 | 15225.5028 | 9079.30599 | Non-sig. |
| PF3D7_1252100 | RON3 | 18031.3104 | -0.346335542 | 0.47656371 | -0.72673503 | 0.46738829 | 0.92538923 | 14811.152 | 12573.61 | 25559.1901 | 19181.2895 | Non-sig. |
| PF3D7_1422800 | ARP4 | 372.584763 | -0.346511743 | 0.52193797 | -0.66389449 | 0.50675784 | 0.93170321 | 260.780547 | 339.161105 | 573.159953 | 317.237446 | Non-sig. |
| PF3D7_1352500 | N/A | 7130.45357 | -0.346617036 | 0.41628919 | -0.83263521 | 0.40505049 | 0.90723586 | 6896.96694 | 6381.82229 | 9068.72616 | 6174.29887 | Non-sig. |
| PF3D7_1451000 | N/A | 484.549861 | -0.346731153 | 0.49135457 | -0.70566385 | 0.48039721 | 0.92880667 | 711.581289 | 414.169872 | 373.689868 | 438.758416 | Non-sig. |
| PF3D7_0925300 | aPRS | 1743.69063 | -0.346929846 | 0.40831756 | -0.84965692 | 0.39551585 | 0.90645858 | 1902.0961 | 1572.65099 | 2002.55709 | 1497.45834 | Non-sig. |
| PF3D7_1467300 | DXR | 1416.59836 | -0.347052457 | 0.4648494 | -0.74659117 | 0.45531038 | 0.92229829 | 1166.68373 | 1372.56783 | 2005.39891 | 1121.74298 | Non-sig. |
| PF3D7_1413300 | N/A | 33.7538972 | -0.347195106 | 0.79274737 | -0.43796437 | 0.66141211 | 0.97783983 | 19.721541 | 34.4688945 | 55.7201957 | 25.1049576 | Non-sig. |
| PF3D7_0303900 | N/A | 95.0563697 | -0.347358155 | 0.54713777 | -0.63486415 | 0.52551702 | 0.93247207 | 89.7597743 | 92.4960471 | 123.025761 | 74.9438959 | Non-sig. |
| PF3D7_1141900 | IMC1b | 14.1453338 | -0.347389967 | 1.11533068 | -0.31146814 | 0.75544476 | 0.99028237 | 25.2117896 | 6.44156399 | 6.56802386 | 18.3599578 | Non-sig. |
| PF3D7_1005600 | N/A | 1352.70356 | -0.347403596 | 0.52317101 | -0.66403449 | 0.50666824 | 0.93170321 | 2157.63328 | 1131.87701 | 872.272155 | 1249.03178 | Non-sig. |
| PF3D7_1317300 | N/A | 1773.82396 | -0.347491742 | 0.41777242 | -0.83177282 | 0.40553719 | 0.90723586 | 1917.56018 | 1733.48228 | 2055.24616 | 1389.00722 | Non-sig. |
| PF3D7_1013700 | N/A | 98.1833113 | -0.348055042 | 0.53189834 | -0.65436384 | 0.51287743 | 0.93247207 | 122.388134 | 85.9900205 | 97.6346833 | 86.7204075 | Non-sig. |
| PF3D7_1235400 | N/A | 389.258832 | -0.348130084 | 0.44510475 | -0.78213068 | 0.43413778 | 0.91522571 | 412.797966 | 304.691734 | 459.136753 | 380.408875 | Non-sig. |
| PF3D7_1112900 | N/A | 213.847619 | -0.348335851 | 0.64760029 | -0.53788711 | 0.59065499 | 0.95315309 | 381.709359 | 212.676162 | 97.6691775 | 163.33578 | Non-sig. |
| PF3D7_0614200 | NAR1 | 385.093759 | -0.348388228 | 0.54643427 | -0.63756658 | 0.52375585 | 0.93247207 | 262.088859 | 268.249708 | 600.360767 | 409.675704 | Non-sig. |
| PF3D7_0923900 | PABP2 | 1624.01066 | -0.348617672 | 0.42897422 | -0.81267745 | 0.41640301 | 0.90793155 | 1525.6024 | 1377.08119 | 2112.76655 | 1480.59249 | Non-sig. |
| PF3D7_1443900 | HSP90 | 2045.6827 | -0.349164884 | 0.4532113 | -0.77042404 | 0.4410484 | 0.91706217 | 1765.8076 | 1593.23695 | 2818.01814 | 2005.66812 | Non-sig. |
| PF3D7_0418200 | EIF3M | 2363.04211 | -0.34924235 | 0.44078679 | -0.79231583 | 0.42817654 | 0.91236574 | 2135.64872 | 2293.00064 | 3159.4891 | 1864.02998 | Non-sig. |
| PF3D7_0400500 | RIF | 1.85725601 | -0.349603367 | 2.68219432 | -0.1303423 | 0.89629562 | 0.99457954 | 4.19219566 | 1.07380062 | 0 | 2.16302778 | Non-sig. |
| PF3D7_1214700 | N/A | 174.57668 | -0.349942339 | 0.49439328 | -0.7078218 | 0.47905593 | 0.92880667 | 221.485834 | 172.885391 | 169.924961 | 134.010533 | Non-sig. |
| PF3D7_1316000 | N/A | 40.4875781 | -0.350006816 | 0.73470341 | -0.47639198 | 0.63379514 | 0.9668494 | 62.9946917 | 42.9504159 | 27.91139 | 28.0938146 | Non-sig. |
| PF3D7_1415400 | N/A | 2212.62581 | -0.350013844 | 0.5894189 | -0.59382868 | 0.55262671 | 0.94170064 | 3863.1463 | 1706.58859 | 1096.6381 | 2184.13025 | Non-sig. |
| PF3D7_1340800 | MPC1 | 383.655245 | -0.350315703 | 0.45160085 | -0.77571977 | 0.43791447 | 0.91593796 | 470.066816 | 378.866003 | 390.048876 | 295.639283 | Non-sig. |
| PF3D7_0913500 | AMR4 | 574.001324 | -0.350880545 | 0.45973587 | -0.76322203 | 0.44533099 | 0.91974025 | 548.634362 | 593.612981 | 738.147395 | 415.610557 | Non-sig. |
| PF3D7_0212400 | N/A | 3184.78455 | -0.351076903 | 0.52296393 | -0.67132145 | 0.50201577 | 0.93170321 | 2144.68931 | 2263.97858 | 4995.77798 | 3334.69231 | Non-sig. |
| PF3D7_1132800 | AQP | 5705.29913 | -0.351181644 | 0.58931621 | -0.59591377 | 0.55123283 | 0.94170064 | 2741.88237 | 5162.26559 | 10050.3485 | 4866.70013 | Non-sig. |
| PF3D7_0205000 | N/A | 235.797286 | -0.351227321 | 0.70080356 | -0.50117799 | 0.61624586 | 0.96358724 | 446.902915 | 235.051412 | 82.1146312 | 179.120187 | Non-sig. |
| PF3D7_1233400 | N/A | 618.56397 | -0.351670787 | 0.42268041 | -0.83200163 | 0.40540803 | 0.90723586 | 698.077095 | 562.200155 | 689.098739 | 524.879888 | Non-sig. |
| PF3D7_1428000 | N/A | 529.987704 | -0.351672306 | 0.44284489 | -0.79412073 | 0.42712515 | 0.91236574 | 508.354521 | 439.937971 | 680.017486 | 491.640836 | Non-sig. |
| PF3D7_0903000 | N/A | 573.058656 | -0.352183251 | 0.47006812 | -0.74921748 | 0.45372614 | 0.92203286 | 713.334699 | 624.003242 | 572.097936 | 382.798747 | Non-sig. |
| PF3D7_1329900 | N/A | 145.125978 | -0.352433095 | 0.5207595 | -0.67676748 | 0.49855349 | 0.93170321 | 135.697955 | 105.171975 | 189.696991 | 149.936991 | Non-sig. |
| PF3D7_1210000 | RPL1 | 1135.03145 | -0.352939095 | 0.423965 | -0.83247224 | 0.40514244 | 0.90723586 | 1401.78993 | 1072.20103 | 1144.69114 | 921.443703 | Non-sig. |
| PF3D7_1456500 | N/A | 4117.3266 | -0.353287367 | 0.43444163 | -0.8131987 | 0.41610414 | 0.90793155 | 3823.22445 | 3212.31311 | 5414.47835 | 4019.29051 | Non-sig. |
| PF3D7_1002900 | N/A | 1396.75292 | -0.353472039 | 0.42362774 | -0.83439304 | 0.40405954 | 0.90723586 | 1425.61395 | 1350.8347 | 1708.30624 | 1102.25681 | Non-sig. |
| PF3D7_1427500 | MSH2-1 | 1822.16816 | -0.353807798 | 0.50916792 | -0.69487449 | 0.48713396 | 0.93170321 | 1227.87533 | 1577.09725 | 2860.78003 | 1622.92003 | Non-sig. |
| PF3D7_1356900 | PK5 | 783.403625 | -0.354214387 | 0.52462097 | -0.67518153 | 0.49956044 | 0.93170321 | 522.801062 | 763.035152 | 1235.08168 | 612.696605 | Non-sig. |
| PF3D7_1307300 | DBP6 | 232.168681 | -0.354225616 | 0.47709003 | -0.74247121 | 0.45780189 | 0.92288312 | 214.587159 | 203.906174 | 306.306023 | 203.875369 | Non-sig. |
| PF3D7_1436700 | N/A | 401.050694 | -0.354328814 | 0.53774869 | -0.65891153 | 0.50995258 | 0.93247207 | 263.256684 | 329.370052 | 636.546075 | 375.029966 | Non-sig. |
| PF3D7_1214100 | PIGO | 2033.33852 | -0.354354467 | 0.54773703 | -0.64694269 | 0.51766904 | 0.93247207 | 1175.73993 | 1752.82809 | 3387.52897 | 1817.2571 | Non-sig. |
| PF3D7_0627400 | TIM22 | 198.802248 | -0.354533819 | 0.5092884 | -0.69613567 | 0.48634387 | 0.93170321 | 163.643915 | 168.621496 | 282.354313 | 180.589268 | Non-sig. |
| PF3D7_1024900 | N/A | 854.186596 | -0.35459271 | 0.42776356 | -0.82894557 | 0.40713521 | 0.90723586 | 849.265377 | 710.113878 | 1067.87002 | 789.49711 | Non-sig. |
| PF3D7_1002400 | TRA2B | 5457.47532 | -0.355086614 | 0.42877876 | -0.8281348 | 0.40759416 | 0.90723586 | 6940.83587 | 5422.41088 | 5310.77 | 4155.88452 | Non-sig. |
| PF3D7_0614600 | N/A | 335.521322 | -0.355473162 | 0.51055224 | -0.69625228 | 0.48627085 | 0.93170321 | 503.923033 | 290.824214 | 249.649063 | 297.688979 | Non-sig. |
| PF3D7_0916200 | N/A | 993.310858 | -0.355553549 | 0.42096072 | -0.84462404 | 0.39832078 | 0.90645858 | 1206.0207 | 822.830098 | 1024.26838 | 920.124251 | Non-sig. |
| PF3D7_0608400 | C4AP4 | 530.928169 | -0.355715557 | 0.4370789 | -0.81384747 | 0.41573234 | 0.90793155 | 615.483464 | 522.693784 | 576.666862 | 408.868567 | Non-sig. |
| PF3D7_1430900 | COX2B | 1369.84978 | -0.355927979 | 0.42794112 | -0.83172184 | 0.40556597 | 0.90723586 | 1470.50104 | 1380.46668 | 1605.41194 | 1023.01944 | Non-sig. |
| PF3D7_1465600 | N/A | 27.6916851 | -0.356152581 | 0.78688126 | -0.45261287 | 0.65082753 | 0.97317009 | 40.9497516 | 23.6222102 | 21.3442981 | 24.8504806 | Non-sig. |
| PF3D7_0214000 | CCT8 | 1599.33253 | -0.356378014 | 0.52559708 | -0.67804413 | 0.49774371 | 0.93170321 | 1020.8468 | 1485.08846 | 2570.56571 | 1320.82916 | Non-sig. |
| PF3D7_1421900 | CTR2 | 2408.84878 | -0.357006743 | 0.41148581 | -0.86760401 | 0.38561115 | 0.90356827 | 2770.76443 | 2293.68466 | 2640.0353 | 1930.91072 | Non-sig. |
| PF3D7_0505700 | N/A | 1866.72174 | -0.357286776 | 0.47146175 | -0.75782771 | 0.44855413 | 0.92081798 | 2730.82693 | 1560.96806 | 1462.85467 | 1712.23729 | Non-sig. |
| PF3D7_0803800 | N/A | 6208.72113 | -0.358222831 | 0.43130393 | -0.83055777 | 0.4062235 | 0.90723586 | 7801.05402 | 6304.95499 | 6150.29365 | 4578.58188 | Non-sig. |
| PF3D7_1002500 | N/A | 761.996772 | -0.358333601 | 0.43941113 | -0.81548595 | 0.4147942 | 0.90793155 | 956.237956 | 752.525073 | 756.184186 | 583.039873 | Non-sig. |

| | | | | | | | | | | | | |
|---|---|---|---|---|---|---|---|---|---|---|---|---|
| PF3D7_1202400 | CYP26 | 813.196207 | -0.358871371 | 0.49164476 | -0.7299404 | 0.46542661 | 0.92538923 | 663.388557 | 860.138668 | 1164.02107 | 565.236533 | Non-sig. |
| PF3D7_0820400 | N/A | 114.745029 | -0.358895738 | 0.51488266 | -0.69704375 | 0.48577542 | 0.93170321 | 115.734215 | 100.873657 | 142.072339 | 100.299905 | Non-sig. |
| PF3D7_1454200 | N/A | 197.002303 | -0.358957846 | 0.62149314 | -0.5775733 | 0.56355224 | 0.94603921 | 155.924874 | 89.0355617 | 286.675739 | 256.373036 | Non-sig. |
| PF3D7_0303100 | N/A | 1552.6533 | -0.359246874 | 0.49051564 | -0.73238618 | 0.46393289 | 0.92538923 | 1184.61344 | 1153.0543 | 2305.05061 | 1567.89486 | Non-sig. |
| PF3D7_0813500 | N/A | 276.772121 | -0.359325835 | 0.48527361 | -0.74046028 | 0.45902075 | 0.92288312 | 376.013457 | 212.581518 | 246.293707 | 272.199803 | Non-sig. |
| PF3D7_1347000 | WDR92 | 103.346253 | -0.359607475 | 0.53873284 | -0.66750613 | 0.5044489 | 0.93170321 | 121.57039 | 107.430341 | 110.789359 | 73.594923 | Non-sig. |
| PF3D7_1441400 | FACT-S | 2033.22658 | -0.359714691 | 0.49415858 | -0.72793372 | 0.46665416 | 0.92538923 | 1456.32659 | 1858.58628 | 3114.16151 | 1703.83194 | Non-sig. |
| PF3D7_0807900 | TyrRS | 2698.42264 | -0.360860758 | 0.41657602 | -0.86625428 | 0.38635073 | 0.90388766 | 2661.26076 | 2374.17111 | 3406.8962 | 2351.36249 | Non-sig. |
| PF3D7_1326400 | N/A | 531.281209 | -0.360928151 | 0.42837187 | -0.84255801 | 0.39947568 | 0.90645858 | 559.878148 | 468.920938 | 634.837614 | 461.488134 | Non-sig. |
| PF3D7_1012300 | QCR7 | 1839.36712 | -0.361097922 | 0.43277124 | -0.83438521 | 0.40406395 | 0.90723586 | 2267.49464 | 1864.95191 | 1869.34899 | 1355.67292 | Non-sig. |
| PF3D7_0830400 | N/A | 2462.05836 | -0.361142007 | 0.41697018 | -0.86610991 | 0.38642989 | 0.90388766 | 2809.45155 | 2418.19294 | 2727.80492 | 1892.78401 | Non-sig. |
| PF3D7_1106900 | N/A | 63.3317821 | -0.36115623 | 0.80449184 | -0.44892466 | 0.653486 | 0.97317009 | 122.183339 | 61.4426488 | 20.4756272 | 49.225513 | Non-sig. |
| PF3D7_0212500 | N/A | 6861.87168 | -0.361763212 | 0.48046701 | -0.75294079 | 0.4514855 | 0.92081798 | 5844.58549 | 4524.75326 | 9590.62732 | 7487.52066 | Non-sig. |
| PF3D7_1118000 | N/A | 468.580512 | -0.362196962 | 0.47566759 | -0.76144974 | 0.44638849 | 0.91974025 | 389.750475 | 429.431959 | 664.196875 | 390.942738 | Non-sig. |
| PF3D7_0324900 | VAR | 1763.94844 | -0.362238284 | 0.45014097 | -0.80472186 | 0.4209802 | 0.90997678 | 2421.89095 | 1704.38153 | 1546.82831 | 1382.69295 | Non-sig. |
| PF3D7_0822400 | N/A | 110.188067 | -0.362415493 | 0.75639784 | -0.47913343 | 0.63184371 | 0.9668494 | 211.226208 | 118.083172 | 36.9475159 | 74.495371 | Non-sig. |
| PF3D7_0313200 | N/A | 378.534553 | -0.362579609 | 0.44011561 | -0.8238281 | 0.41003724 | 0.90748263 | 425.870286 | 357.9023 | 425.839523 | 304.526103 | Non-sig. |
| PF3D7_1217500 | N/A | 2821.99698 | -0.362580649 | 0.45411498 | -0.79843358 | 0.42461892 | 0.91236574 | 4008.18613 | 2448.39965 | 2341.59818 | 2489.80396 | Non-sig. |
| PF3D7_0110100 | SELB | 462.144781 | -0.363021309 | 0.43896351 | -0.82699655 | 0.40823901 | 0.90723586 | 463.272719 | 388.391646 | 576.574714 | 420.340046 | Non-sig. |
| PF3D7_1127400 | N/A | 185.74625 | -0.363306485 | 0.48044031 | -0.75619484 | 0.44953238 | 0.92081798 | 183.911469 | 169.608288 | 234.002734 | 155.462509 | Non-sig. |
| PF3D7_0627600 | N/A | 143.354225 | -0.363373346 | 0.539768 | -0.67320284 | 0.50081826 | 0.93170321 | 206.216607 | 140.579385 | 116.614218 | 110.006692 | Non-sig. |
| PF3D7_1004400 | N/A | 3598.35696 | -0.363604923 | 0.40569611 | -0.89624947 | 0.37011954 | 0.89654416 | 4275.50132 | 3125.21669 | 3823.41641 | 3169.29342 | Non-sig. |
| PF3D7_1357400 | N/A | 2908.64203 | -0.363646772 | 0.41391996 | -0.87854369 | 0.37964874 | 0.90130671 | 3599.38173 | 2422.10786 | 2947.32323 | 2665.75531 | Non-sig. |
| PF3D7_1316600 | CCT | 8065.7271 | -0.363761415 | 0.41725411 | -0.87179827 | 0.38331844 | 0.90292925 | 10199.4206 | 6547.16956 | 7955.17574 | 7561.14248 | Non-sig. |
| PF3D7_1008500 | N/A | 4749.09469 | -0.363964277 | 0.4272346 | -0.8519073 | 0.39426555 | 0.90621209 | 5052.21371 | 3442.06066 | 5637.70252 | 4864.40187 | Non-sig. |
| PF3D7_0823800 | N/A | 2201.45961 | -0.364174678 | 0.54977177 | -0.66241066 | 0.50770807 | 0.93170321 | 1288.87125 | 1708.01614 | 3666.48877 | 2142.46226 | Non-sig. |
| PF3D7_0319900 | N/A | 224.152895 | -0.364543084 | 0.56195235 | -0.64870817 | 0.51652702 | 0.93247207 | 170.250292 | 137.403997 | 334.172607 | 254.784683 | Non-sig. |
| PF3D7_0708300 | BUD32 | 138.398751 | -0.364579505 | 0.57439975 | -0.6347139 | 0.52561502 | 0.93247207 | 102.869774 | 147.113826 | 208.506805 | 95.1046004 | Non-sig. |
| PF3D7_1432000 | SYN11 | 529.278299 | -0.364666411 | 0.44570801 | -0.81817333 | 0.41325822 | 0.90793155 | 694.174494 | 475.438377 | 497.634836 | 449.86549 | Non-sig. |
| PF3D7_1358100 | N/A | 1745.80388 | -0.364773075 | 0.60352842 | -0.60440083 | 0.54557719 | 0.94121391 | 3105.60469 | 1732.80898 | 825.41828 | 1319.38358 | Non-sig. |
| PF3D7_0419900 | N/A | 13086.1105 | -0.364945299 | 0.42461906 | -0.85946518 | 0.39008392 | 0.90444842 | 16569.9802 | 10005.7994 | 12895.1295 | 12873.5329 | Non-sig. |
| PF3D7_1113700 | GLO1 | 463.599914 | -0.365009421 | 0.46776718 | -0.78032285 | 0.43520087 | 0.91522571 | 399.166365 | 420.854859 | 644.481577 | 389.896853 | Non-sig. |
| PF3D7_1343500 | N/A | 678.14515 | -0.365054385 | 0.43175297 | -0.84551678 | 0.39782237 | 0.90645858 | 763.329726 | 667.609984 | 763.651056 | 517.989834 | Non-sig. |
| PF3D7_0109950 | C2AP3 | 352.917178 | -0.365646073 | 0.4583809 | -0.79769046 | 0.42505014 | 0.91236574 | 332.902834 | 325.897478 | 461.747369 | 291.12103 | Non-sig. |
| PF3D7_0905900 | SEC27 | 1334.57971 | -0.365832754 | 0.46364378 | -0.78903842 | 0.43008955 | 0.91355742 | 1099.2049 | 1085.78411 | 1906.3059 | 1247.02392 | Non-sig. |
| PF3D7_1475100 | N/A | 935.321283 | -0.365931157 | 0.46403311 | -0.78858847 | 0.43035257 | 0.91355742 | 772.341515 | 807.729275 | 1334.0145 | 827.199842 | Non-sig. |
| PF3D7_0717600 | IMC32 | 5361.96348 | -0.366614933 | 0.49268427 | -0.74411738 | 0.45680546 | 0.92288312 | 8229.75194 | 4327.73655 | 3849.78075 | 5040.58467 | Non-sig. |
| PF3D7_1455200 | N/A | 536.037979 | -0.366965809 | 0.42931241 | -0.85477569 | 0.39267534 | 0.90601068 | 606.482941 | 503.188561 | 601.217573 | 433.262843 | Non-sig. |
| PF3D7_1209200 | LSM7 | 497.166079 | -0.366970174 | 0.4473666 | -0.82028961 | 0.41205103 | 0.90748263 | 548.102877 | 507.074296 | 571.999159 | 361.487982 | Non-sig. |
| PF3D7_1323600 | N/A | 529.255993 | -0.368603518 | 0.43952988 | -0.83863132 | 0.40167623 | 0.90667178 | 526.23372 | 428.378266 | 666.640524 | 495.771462 | Non-sig. |
| PF3D7_0928900 | GK | 104.218775 | -0.368746021 | 0.56482772 | -0.65284689 | 0.51385499 | 0.93247207 | 82.8850164 | 86.9916645 | 151.852071 | 95.1463482 | Non-sig. |
| PF3D7_1314700 | N/A | 1394.36931 | -0.369089985 | 0.56335083 | -0.65516898 | 0.51235897 | 0.93247207 | 2343.99551 | 1418.69521 | 799.866616 | 1014.91991 | Non-sig. |
| PF3D7_1466000 | N/A | 13.6019356 | -0.369693991 | 1.0071837 | -0.36705716 | 0.71357638 | 0.9875928 | 16.7434285 | 15.0465218 | 13.9487494 | 8.66904273 | Non-sig. |
| PF3D7_1315700 | GCD14 | 952.284502 | -0.369697248 | 0.41947799 | -0.88132692 | 0.3781409 | 0.9000162 | 1032.02199 | 892.867593 | 1115.20471 | 769.043717 | Non-sig. |
| PF3D7_1014800 | N/A | 145.610715 | -0.37041857 | 0.56978919 | -0.65009758 | 0.5156292 | 0.93247207 | 225.172003 | 143.795682 | 103.475966 | 109.999209 | Non-sig. |
| PF3D7_0817300 | N/A | 2195.57565 | -0.370839071 | 0.58936889 | -0.62921386 | 0.52920906 | 0.93270419 | 1310.38375 | 1229.11621 | 3641.7564 | 2601.04621 | Non-sig. |
| PF3D7_0924000 | N/A | 6766.77497 | -0.371648225 | 0.4629716 | -0.80274519 | 0.42212203 | 0.91161836 | 9469.64349 | 4863.49604 | 5797.73722 | 6936.22314 | Non-sig. |
| PF3D7_1409900 | CDS | 2063.0683 | -0.371665739 | 0.46907563 | -0.79233649 | 0.4281645 | 0.91236574 | 1668.6267 | 1608.05548 | 2985.80383 | 1989.7872 | Non-sig. |
| PF3D7_1406500 | WDR65 | 91.3865846 | -0.372008867 | 0.59721112 | -0.62291015 | 0.53334357 | 0.93467024 | 139.059265 | 71.87337 | 67.355729 | 87.257974 | Non-sig. |
| PF3D7_1408100 | PMIII | 2281.69918 | -0.372193457 | 0.53613509 | -0.69421581 | 0.48754688 | 0.93170321 | 1385.61828 | 1998.17148 | 3762.84684 | 1980.16012 | Non-sig. |
| PF3D7_1423200 | CYP52 | 552.284036 | -0.372279222 | 0.46440349 | -0.80162882 | 0.4227677 | 0.91161836 | 752.138422 | 541.954772 | 494.363477 | 420.679473 | Non-sig. |
| PF3D7_0318000 | N/A | 14.9100297 | -0.37248984 | 0.95598015 | -0.38964181 | 0.69680143 | 0.98334178 | 15.5977762 | 11.8428659 | 18.0329017 | 14.1665751 | Non-sig. |
| PF3D7_1328100 | N/A | 5352.33028 | -0.372597046 | 0.43550958 | -0.85554271 | 0.39225077 | 0.90601068 | 4910.36414 | 5123.83574 | 7168.78229 | 4206.33894 | Non-sig. |
| PF3D7_1358700 | YOP1L | 3694.39939 | -0.372988925 | 0.41064049 | -0.90831014 | 0.36371439 | 0.89567376 | 4074.98486 | 2915.87184 | 4263.63611 | 3523.10474 | Non-sig. |

| | | | | | | | | | | | | |
|---|---|---|---|---|---|---|---|---|---|---|---|---|
| PF3D7_1237800 | HUB1 | 226.387833 | -0.373605621 | 0.48947988 | -0.76327064 | 0.44530201 | 0.91974025 | 305.232556 | 219.105835 | 206.017019 | 175.195923 | Non-sig. |
| PF3D7_1467200 | WDR79 | 129.161032 | -0.374085919 | 0.50121318 | -0.7463609 | 0.45544943 | 0.92229829 | 154.548629 | 107.334721 | 137.127202 | 117.633576 | Non-sig. |
| PF3D7_0719300 | ARP6 | 1632.33773 | -0.374595162 | 0.41078893 | -0.91189205 | 0.36182555 | 0.89567376 | 1944.81101 | 1456.75805 | 1741.40579 | 1386.37609 | Non-sig. |
| PF3D7_0207000 | MSP4 | 21551.3662 | -0.376499902 | 0.43264617 | -0.8702259 | 0.38417696 | 0.90292925 | 27825.0201 | 21588.4054 | 20870.4506 | 15921.5888 | Non-sig. |
| PF3D7_0419400 | N/A | 2910.2779 | -0.377290481 | 0.47346796 | -0.79686593 | 0.4255289 | 0.91236574 | 2361.1628 | 2150.48609 | 4215.90878 | 2913.55391 | Non-sig. |
| PF3D7_1241500 | N/A | 540.612148 | -0.377721153 | 0.47645738 | -0.79277007 | 0.4279118 | 0.91236574 | 779.581255 | 480.767624 | 442.636943 | 459.462771 | Non-sig. |
| PF3D7_0518200 | MDM2 | 3757.42585 | -0.377874424 | 0.45108787 | -0.83769583 | 0.40220156 | 0.90713659 | 3288.49528 | 3680.79648 | 5204.70136 | 2855.71027 | Non-sig. |
| PF3D7_1329000 | N/A | 1694.41642 | -0.378418982 | 0.44331787 | -0.85360642 | 0.3933231 | 0.90601068 | 1536.91046 | 1328.89256 | 2293.62417 | 1618.23852 | Non-sig. |
| PF3D7_1474300 | N/A | 335.237905 | -0.378662541 | 0.47460266 | -0.79785171 | 0.42495655 | 0.91236574 | 460.031759 | 315.449974 | 298.134023 | 267.335863 | Non-sig. |
| PF3D7_1410700 | N/A | 2161.17635 | -0.378777403 | 0.43383642 | -0.87308808 | 0.38261507 | 0.90292925 | 1974.36864 | 1864.14328 | 2911.94895 | 1894.24453 | Non-sig. |
| PF3D7_0809500 | N/A | 183.060991 | -0.379379815 | 0.47735778 | -0.79474941 | 0.42675929 | 0.91236574 | 199.815797 | 174.124813 | 214.142664 | 144.160688 | Non-sig. |
| PF3D7_1124100 | BDCP | 4723.75296 | -0.379683293 | 0.40765297 | -0.93138852 | 0.35165263 | 0.8930363 | 5269.28406 | 3774.58018 | 5414.26 | 4436.88759 | Non-sig. |
| PF3D7_1457700 | N/A | 875.887346 | -0.379782177 | 0.45251598 | -0.83926799 | 0.40131894 | 0.90667178 | 1184.41907 | 670.411386 | 796.805136 | 851.91379 | Non-sig. |
| PF3D7_1144500 | AOS1 | 343.294955 | -0.380214483 | 0.503811 | -0.75467683 | 0.45044291 | 0.92081798 | 270.41415 | 331.58948 | 505.881617 | 265.294574 | Non-sig. |
| PF3D7_0108000 | N/A | 4430.60441 | -0.380904462 | 0.42211327 | -0.90237501 | 0.36685769 | 0.89567376 | 4749.25108 | 4424.3107 | 5274.93988 | 3273.916 | Non-sig. |
| PF3D7_1128500 | mHCF101 | 309.525732 | -0.380996223 | 0.46328551 | -0.82237888 | 0.41086129 | 0.90748263 | 290.366787 | 263.993846 | 409.787311 | 273.954982 | Non-sig. |
| PF3D7_1104000 | N/A | 1390.24129 | -0.381244899 | 0.47506394 | -0.80251281 | 0.42225638 | 0.91161836 | 1087.31023 | 1238.23749 | 2058.14838 | 1177.26906 | Non-sig. |
| PF3D7_0915700 | ApiCOX30 | 442.135495 | -0.381551097 | 0.44343708 | -0.86044022 | 0.38954642 | 0.90444842 | 542.57854 | 424.184868 | 458.041589 | 343.736983 | Non-sig. |
| PF3D7_1010200 | N/A | 1241.65267 | -0.381853009 | 0.43848953 | -0.87083724 | 0.38384303 | 0.90292925 | 1291.04912 | 888.122072 | 1518.89961 | 1268.53988 | Non-sig. |
| PF3D7_1022800 | ISPG | 2512.61081 | -0.382507638 | 0.40743359 | -0.93882204 | 0.34782212 | 0.8930363 | 2742.11647 | 2277.68769 | 2945.36682 | 2085.27227 | Non-sig. |
| PF3D7_0508400 | N/A | 185.707857 | -0.382874775 | 0.57256819 | -0.66869725 | 0.50368862 | 0.93170321 | 293.464626 | 192.296848 | 127.204045 | 129.865909 | Non-sig. |
| PF3D7_0602900 | N/A | 1603.8116 | -0.382898847 | 0.42119346 | -0.90908071 | 0.36330752 | 0.89567376 | 2036.09096 | 1330.63318 | 1594.85387 | 1453.66841 | Non-sig. |
| PF3D7_1341200 | N/A | 4077.13114 | -0.384062248 | 0.43634067 | -0.88018898 | 0.37875694 | 0.9000162 | 3826.64489 | 3979.74407 | 5406.45317 | 3095.68241 | Non-sig. |
| PF3D7_0516900 | RPL2 | 5353.99884 | -0.384086015 | 0.41969667 | -0.91515144 | 0.36011213 | 0.89560738 | 5154.17511 | 4631.10194 | 6970.67562 | 4660.04269 | Non-sig. |
| PF3D7_1120500 | N/A | 1058.31145 | -0.384536513 | 0.4312523 | -0.89167412 | 0.37256763 | 0.89654416 | 1012.05475 | 935.5147 | 1384.81866 | 900.857692 | Non-sig. |
| PF3D7_0800200 | VAR | 207.565128 | -0.384570178 | 0.53214188 | -0.72268355 | 0.46987433 | 0.92682482 | 311.003697 | 202.705522 | 159.374084 | 157.177209 | Non-sig. |
| PF3D7_0423500 | GAPM2 | 7246.67172 | -0.385016917 | 0.48573584 | -0.79264672 | 0.42798368 | 0.91236574 | 5746.61959 | 5043.96974 | 10669.0085 | 7527.08901 | Non-sig. |
| PF3D7_0725200 | N/A | 948.124962 | -0.385503019 | 0.43138893 | -0.89363215 | 0.37151874 | 0.89654416 | 1113.25347 | 943.177698 | 1034.89675 | 701.171929 | Non-sig. |
| PF3D7_0824400 | NT2 | 1314.38311 | -0.385590883 | 0.53492739 | -0.72082845 | 0.47101507 | 0.92682482 | 815.937956 | 1127.64417 | 2161.71964 | 1152.23066 | Non-sig. |
| PF3D7_0801400 | N/A | 108.908457 | -0.385835658 | 0.55854878 | -0.69078238 | 0.48970231 | 0.93170321 | 151.748471 | 111.771568 | 95.1807471 | 76.9330416 | Non-sig. |
| PF3D7_1220500 | TSR3 | 190.046759 | -0.386410459 | 0.46940814 | -0.82318653 | 0.41040193 | 0.90748263 | 209.003622 | 167.490594 | 221.662574 | 162.030246 | Non-sig. |
| PF3D7_1212200 | N/A | 129.0198 | -0.387199855 | 0.50913526 | -0.76050489 | 0.44695285 | 0.91974025 | 130.567245 | 101.924102 | 161.80416 | 121.783694 | Non-sig. |
| PF3D7_1223600 | N/A | 547.540828 | -0.387303694 | 0.53972791 | -0.71759064 | 0.47300973 | 0.92682482 | 356.182532 | 419.404785 | 884.706108 | 529.86989 | Non-sig. |
| PF3D7_0218000 | RFC2 | 1895.08198 | -0.38736831 | 0.47038807 | -0.82350793 | 0.41021921 | 0.90748263 | 1550.96125 | 1848.58282 | 2744.74866 | 1436.03519 | Non-sig. |
| PF3D7_0509000 | alphaSNAP | 2058.51905 | -0.387672906 | 0.4141273 | -0.93612015 | 0.34921133 | 0.8930363 | 2380.97328 | 1958.93662 | 2285.94399 | 1608.2223 | Non-sig. |
| PF3D7_1444900 | N/A | 61.035807 | -0.388522109 | 0.58954044 | -0.65902537 | 0.50987948 | 0.93247207 | 67.0004522 | 50.5076434 | 71.3894256 | 55.2457067 | Non-sig. |
| PF3D7_0312300 | RPN12 | 1885.74578 | -0.388572774 | 0.42923722 | -0.90526346 | 0.36532583 | 0.89567376 | 1779.46651 | 1687.47126 | 2496.69582 | 1579.34953 | Non-sig. |
| PF3D7_0111500 | N/A | 3397.93616 | -0.388735198 | 0.41826204 | -0.92940587 | 0.35267879 | 0.8930363 | 4248.94577 | 3208.99245 | 3457.13139 | 2676.67501 | Non-sig. |
| PF3D7_1325500 | N/A | 341.625707 | -0.388894507 | 0.44071853 | -0.88241015 | 0.37755505 | 0.89940947 | 391.301692 | 309.079788 | 383.497687 | 282.623661 | Non-sig. |
| PF3D7_1352700 | AQR | 2640.24936 | -0.389849663 | 0.51011541 | -0.76423816 | 0.44472533 | 0.91974025 | 1871.14169 | 1904.83191 | 4118.20111 | 2666.82271 | Non-sig. |
| PF3D7_0602400 | EF-G | 1824.34711 | -0.389967838 | 0.43080874 | -0.90519947 | 0.36535972 | 0.89567376 | 1857.97058 | 1350.86622 | 2280.7551 | 1807.79654 | Non-sig. |
| PF3D7_1415600 | N/A | 1667.22881 | -0.390055735 | 0.44133496 | -0.88380883 | 0.37679942 | 0.89801319 | 1908.01082 | 1136.00154 | 1874.48802 | 1750.41486 | Non-sig. |
| PF3D7_0713900 | N/A | 6833.54605 | -0.390117741 | 0.48927247 | -0.79734251 | 0.42525214 | 0.91236574 | 5248.12346 | 4888.23335 | 10255.3642 | 6942.46319 | Non-sig. |
| PF3D7_1004200 | N/A | 10441.601 | -0.390679896 | 0.48229644 | -0.81004101 | 0.41791661 | 0.90869129 | 8198.77949 | 7521.58599 | 15494.5715 | 10551.4669 | Non-sig. |
| PF3D7_1226900 | PV2 | 270.632874 | -0.391303392 | 0.54222554 | -0.72166167 | 0.47050251 | 0.92682482 | 201.795624 | 182.485238 | 412.179861 | 286.070775 | Non-sig. |
| PF3D7_1454600 | N/A | 434.730632 | -0.391371262 | 0.43729586 | -0.89498049 | 0.37079752 | 0.89654416 | 456.350541 | 355.493554 | 530.242474 | 396.83596 | Non-sig. |
| PF3D7_0310100 | CDPK3 | 55.4430664 | -0.391720009 | 0.62618483 | -0.62556612 | 0.53159957 | 0.93351492 | 76.7044437 | 50.455484 | 49.2622769 | 45.3500611 | Non-sig. |
| PF3D7_0214500 | N/A | 27.9709574 | -0.391771022 | 0.758304 | -0.51664111 | 0.60540671 | 0.95693578 | 27.3159894 | 25.7654036 | 36.1248793 | 22.6775572 | Non-sig. |
| PF3D7_1331200 | N/A | 1966.54451 | -0.392372074 | 0.54821113 | -0.71573168 | 0.47415704 | 0.92682482 | 3312.83438 | 1625.82712 | 1152.16252 | 1775.354 | Non-sig. |
| PF3D7_1242900 | TIM13 | 409.457035 | -0.392541355 | 0.46351946 | -0.84687137 | 0.39706682 | 0.90645858 | 556.217568 | 382.101418 | 373.616259 | 325.892894 | Non-sig. |
| PF3D7_0901700 | N/A | 1080.33148 | -0.392587462 | 0.47558912 | -0.82547612 | 0.40910133 | 0.90748263 | 895.754664 | 1072.27271 | 1556.84641 | 796.452128 | Non-sig. |
| PF3D7_1336100 | N/A | 123.105405 | -0.392666654 | 0.5208058 | -0.75395983 | 0.45087334 | 0.92081798 | 163.986343 | 111.832704 | 115.658204 | 100.94437 | Non-sig. |
| PF3D7_0710900 | RPL1 | 654.333354 | -0.392689982 | 0.44293276 | -0.88656793 | 0.37531157 | 0.89654416 | 784.480651 | 660.067701 | 701.264371 | 471.520695 | Non-sig. |
| PF3D7_1138100 | URM1 | 118.883442 | -0.392914395 | 0.53607866 | -0.73294168 | 0.463594 | 0.92509266 | 103.949369 | 97.7100514 | 165.828777 | 108.045568 | Non-sig. |

| | | | | | | | | | | | | |
|---|---|---|---|---|---|---|---|---|---|---|---|---|
| PF3D7_0105300 | CAP | 6360.35923 | -0.393185209 | 0.44269736 | -0.88815802 | 0.37445576 | 0.89654416 | 8664.72473 | 6195.38325 | 5778.9862 | 4802.34274 | Non-sig. |
| PF3D7_0413500 | PGM2 | 5118.7231 | -0.39348359 | 0.43429388 | -0.9060307 | 0.3649196 | 0.89567376 | 6864.35832 | 4866.93052 | 4760.79474 | 3982.80881 | Non-sig. |
| PF3D7_1304900 | RPB11 | 356.709223 | -0.393810026 | 0.45450976 | -0.8664501 | 0.38624338 | 0.90388766 | 347.80322 | 328.49572 | 462.24585 | 288.2921 | Non-sig. |
| PF3D7_1364900 | FC | 140.622602 | -0.394374351 | 0.52471906 | -0.75159144 | 0.45229679 | 0.92103152 | 127.167386 | 138.475896 | 192.130084 | 104.717043 | Non-sig. |
| PF3D7_1138600 | N/A | 657.454057 | -0.395797697 | 0.43090551 | -0.91852549 | 0.35834382 | 0.89552149 | 681.851522 | 612.735671 | 812.20885 | 523.020184 | Non-sig. |
| PF3D7_1123800 | SMC5 | 1796.38889 | -0.396135492 | 0.53079467 | -0.74630647 | 0.45548231 | 0.92229829 | 2916.04492 | 1328.04641 | 1167.22649 | 1774.23772 | Non-sig. |
| PF3D7_0502100 | N/A | 577.313535 | -0.396180395 | 0.45854604 | -0.86399263 | 0.38759194 | 0.90421052 | 519.543697 | 540.915421 | 792.428854 | 456.366169 | Non-sig. |
| PF3D7_0723700 | N/A | 605.607118 | -0.397384742 | 0.44583762 | -0.89132169 | 0.37275661 | 0.89654416 | 636.036231 | 436.655152 | 740.859139 | 608.877949 | Non-sig. |
| PF3D7_0518100 | AMR2 | 1041.84787 | -0.39748913 | 0.55014917 | -0.72251156 | 0.46998002 | 0.92682482 | 624.744243 | 812.050451 | 1743.87135 | 986.725436 | Non-sig. |
| PF3D7_1234800 | SF3B3 | 2160.04848 | -0.397714285 | 0.61015695 | -0.65182292 | 0.51451541 | 0.93247207 | 972.631249 | 1824.58283 | 3938.85775 | 1904.12207 | Non-sig. |
| PF3D7_0531200 | N/A | 142.370037 | -0.397962661 | 0.49382803 | -0.80587297 | 0.4203161 | 0.90969969 | 175.259952 | 123.497327 | 148.575861 | 122.147006 | Non-sig. |
| PF3D7_1320800 | N/A | 1139.26919 | -0.39824797 | 0.48628053 | -0.81896754 | 0.41280494 | 0.90786241 | 910.702203 | 1141.95964 | 1680.08857 | 824.326366 | Non-sig. |
| PF3D7_1317500 | N/A | 35.9674616 | -0.398330937 | 0.72771009 | -0.54737586 | 0.58412054 | 0.95151408 | 27.7576117 | 31.3469294 | 53.9286815 | 30.836624 | Non-sig. |
| PF3D7_1209000 | N/A | 699.432351 | -0.39837058 | 0.4601962 | -0.86565379 | 0.38668005 | 0.90407366 | 906.498336 | 724.089339 | 684.426762 | 482.714969 | Non-sig. |
| PF3D7_1245000 | N/A | 187.363902 | -0.399280138 | 0.47780399 | -0.83565676 | 0.40334803 | 0.90723586 | 234.231776 | 152.455423 | 192.110693 | 170.657715 | Non-sig. |
| PF3D7_1450900 | N/A | 180.887288 | -0.399310289 | 0.51780692 | -0.77115673 | 0.44061404 | 0.91706217 | 153.492833 | 137.389561 | 257.830676 | 174.836081 | Non-sig. |
| PF3D7_1363000 | N/A | 175.262919 | -0.39943836 | 0.51769444 | -0.77157166 | 0.44036817 | 0.91706217 | 252.751129 | 158.77989 | 146.202754 | 143.317903 | Non-sig. |
| PF3D7_0311700 | PM6 | 42.5385668 | -0.4001827 | 0.73998154 | -0.54080092 | 0.58864481 | 0.95315309 | 68.2305142 | 46.1743056 | 28.7312944 | 27.0181529 | Non-sig. |
| PF3D7_0402400 | GEXP18 | 9861.28526 | -0.400268016 | 0.44315235 | -0.90322891 | 0.36640442 | 0.89567376 | 13741.444 | 9207.68066 | 8699.92032 | 7796.09612 | Non-sig. |
| PF3D7_0307400 | ClpP | 1680.23866 | -0.401092689 | 0.41555263 | -0.96520312 | 0.33444309 | 0.89204385 | 2080.93615 | 1515.93252 | 1743.79752 | 1380.28846 | Non-sig. |
| PF3D7_1137000 | N/A | 154.96235 | -0.401367201 | 0.48788953 | -0.82266 | 0.41070137 | 0.90748263 | 190.969995 | 137.924656 | 161.862916 | 129.091833 | Non-sig. |
| PF3D7_0730100 | N/A | 271.467188 | -0.401618324 | 0.45254324 | -0.88746951 | 0.37482618 | 0.89654416 | 330.624051 | 230.671496 | 287.468423 | 237.104782 | Non-sig. |
| PF3D7_1011100 | N/A | 508.481707 | -0.401723665 | 0.51448206 | -0.78083124 | 0.43490176 | 0.91522571 | 384.452165 | 359.384684 | 772.924306 | 517.165674 | Non-sig. |
| PF3D7_0918300 | EIF3F | 1012.46277 | -0.402379079 | 0.44785392 | -0.89846055 | 0.36894007 | 0.89654416 | 906.123172 | 911.311793 | 1399.15179 | 833.264324 | Non-sig. |
| PF3D7_1411400 | PREX | 1598.96585 | -0.402758213 | 0.49300821 | -0.81694017 | 0.41396262 | 0.90793155 | 1323.80637 | 1043.63164 | 2317.39543 | 1711.02995 | Non-sig. |
| PF3D7_0629900 | N/A | 1208.1471 | -0.402885967 | 0.43313127 | -0.93017058 | 0.35228277 | 0.8930363 | 1168.64591 | 954.900441 | 1582.69048 | 1126.35156 | Non-sig. |
| PF3D7_0417000 | N/A | 17.5261885 | -0.403099613 | 0.94401746 | -0.4270044 | 0.66937612 | 0.97883902 | 25.2054716 | 20.3997216 | 14.7774022 | 9.72215844 | Non-sig. |
| PF3D7_1105500 | CEN4 | 959.69984 | -0.403938102 | 0.46272331 | -0.87295818 | 0.38268587 | 0.90292925 | 823.337268 | 906.189921 | 1362.79279 | 746.479378 | Non-sig. |
| PF3D7_1108600 | ERC | 29038.4419 | -0.404047522 | 0.46453169 | -0.86979539 | 0.38441223 | 0.90292925 | 23452.755 | 27167.7236 | 42703.7756 | 22829.5134 | Non-sig. |
| PF3D7_1427700 | N/A | 180.660091 | -0.404543259 | 0.49321547 | -0.82021608 | 0.41209294 | 0.90748263 | 171.197724 | 167.776104 | 240.316086 | 143.350452 | Non-sig. |
| PF3D7_0921400 | NifU | 83.3101614 | -0.404576985 | 0.55775429 | -0.72536777 | 0.46822644 | 0.92594245 | 84.7433444 | 80.6119781 | 105.021478 | 62.8638455 | Non-sig. |
| PF3D7_0927400 | N/A | 157.044469 | -0.405083585 | 0.6245781 | -0.64857155 | 0.51661535 | 0.93247207 | 273.591114 | 136.281494 | 84.5884506 | 133.716818 | Non-sig. |
| PF3D7_1018500 | RDS3 | 540.850085 | -0.405337144 | 0.4476198 | -0.90553891 | 0.36517996 | 0.89567376 | 630.262249 | 547.79679 | 602.433829 | 382.907472 | Non-sig. |
| PF3D7_1216400 | N/A | 1440.5514 | -0.405475955 | 0.45336548 | -0.89436884 | 0.37112458 | 0.89654416 | 1462.35014 | 961.344723 | 1820.86637 | 1517.64436 | Non-sig. |
| PF3D7_1135400 | N/A | 868.570752 | -0.405532922 | 0.64532252 | -0.62841899 | 0.5297295 | 0.93270419 | 345.594037 | 699.993286 | 1633.78132 | 794.91436 | Non-sig. |
| PF3D7_1008900 | AK1 | 3061.60616 | -0.405629218 | 0.48903651 | -0.82944567 | 0.40685227 | 0.90723586 | 2323.19604 | 2984.76828 | 4654.91404 | 2283.54627 | Non-sig. |
| PF3D7_0823900 | DTC | 1272.16843 | -0.405912482 | 0.51102904 | -0.79430414 | 0.4270184 | 0.91236574 | 888.025387 | 1202.28823 | 2011.60075 | 986.759344 | Non-sig. |
| PF3D7_0931400 | N/A | 1831.98398 | -0.406143203 | 0.54201287 | -0.74932391 | 0.453662 | 0.92203286 | 3008.93627 | 1870.66583 | 1167.70345 | 1280.63036 | Non-sig. |
| PF3D7_0514600 | N/A | 291.397855 | -0.406642219 | 0.48575589 | -0.83713286 | 0.40251789 | 0.90716539 | 254.44371 | 272.625158 | 409.744086 | 228.778467 | Non-sig. |
| PF3D7_0921900 | N/A | 997.008229 | -0.407474232 | 0.56427734 | -0.72211695 | 0.47022258 | 0.92682482 | 1677.16178 | 1032.69748 | 596.90531 | 681.26835 | Non-sig. |
| PF3D7_1345700 | IDH | 2617.62334 | -0.40750256 | 0.53915025 | -0.75582374 | 0.44975488 | 0.92081798 | 1604.22329 | 2452.82664 | 4365.194 | 2048.24942 | Non-sig. |
| PF3D7_1425900 | N/A | 270.607992 | -0.407841768 | 0.4529706 | -0.90037138 | 0.36792265 | 0.89615989 | 318.380971 | 216.850274 | 298.858637 | 248.342088 | Non-sig. |
| PF3D7_1113800 | N/A | 1715.15558 | -0.407930491 | 0.42912632 | -0.95060702 | 0.3418039 | 0.8930363 | 1628.63421 | 1488.16837 | 2283.25492 | 1460.56484 | Non-sig. |
| PF3D7_0216900 | N/A | 309.068998 | -0.408235871 | 0.59175066 | -0.68987818 | 0.4902708 | 0.93170321 | 533.722791 | 240.475786 | 171.593763 | 290.483652 | Non-sig. |
| PF3D7_0321200 | ALG7 | 138.830551 | -0.408406038 | 0.55419155 | -0.73694022 | 0.46115871 | 0.92336312 | 110.397498 | 138.459809 | 206.103204 | 100.361693 | Non-sig. |
| PF3D7_0816300 | N/A | 81.4472985 | -0.408458467 | 0.63977294 | -0.63844287 | 0.52318543 | 0.93247207 | 132.637591 | 79.3995027 | 53.384659 | 60.3674412 | Non-sig. |
| PF3D7_1331300 | SEC11 | 2268.692 | -0.409180118 | 0.44774749 | -0.91386357 | 0.36078854 | 0.89560738 | 2013.40717 | 2131.5104 | 3162.92781 | 1766.92263 | Non-sig. |
| PF3D7_0628100 | HECT1 | 69329.8416 | -0.4095246 | 0.5307655 | -0.77157352 | 0.44036707 | 0.91706217 | 55618.911 | 37673.9578 | 102589.596 | 81436.902 | Non-sig. |
| PF3D7_0815400 | ATPTG3 | 634.818636 | -0.40964615 | 0.43287828 | -0.94633104 | 0.34397977 | 0.8930363 | 719.092703 | 613.391248 | 729.595496 | 477.195098 | Non-sig. |
| PF3D7_0610600 | CDPK2 | 1415.70752 | -0.409665628 | 0.41296038 | -0.99202162 | 0.32118698 | 0.88694295 | 1515.11153 | 1252.32517 | 1715.55302 | 1179.84038 | Non-sig. |
| PF3D7_0813600 | N/A | 812.748039 | -0.409727294 | 0.57112568 | -0.71740303 | 0.47312545 | 0.92682482 | 1371.41857 | 854.304417 | 483.668484 | 541.600682 | Non-sig. |
| PF3D7_0613400 | N/A | 2932.32299 | -0.410184475 | 0.41626538 | -0.98539176 | 0.32443168 | 0.88694295 | 3533.25339 | 2796.3284 | 3159.60792 | 2240.10226 | Non-sig. |
| PF3D7_1357300 | N/A | 5.11597919 | -0.410573255 | 1.61274885 | -0.25457978 | 0.79904767 | 0.99028237 | 5.13330519 | 1.08226291 | 6.5290826 | 7.71926606 | Non-sig. |
| PF3D7_1415200 | N/A | 81.5682885 | -0.410810763 | 0.55456882 | -0.74077508 | 0.45882983 | 0.92288312 | 90.1657846 | 78.4153684 | 96.0242338 | 61.6677673 | Non-sig. |

| | | | | | | | | | | | | |
|---|---|---|---|---|---|---|---|---|---|---|---|---|
| PF3D7_1452000 | RON2 | 10140.7158 | -0.410898217 | 0.47709709 | -0.86124653 | 0.38910228 | 0.90444842 | 9286.55429 | 6319.53 | 13863.5422 | 11093.2365 | Non-sig. |
| PF3D7_1353700 | ROMO1 | 198.206852 | -0.41104252 | 0.50590825 | -0.81248432 | 0.41651378 | 0.90793155 | 178.224396 | 195.504495 | 274.117662 | 144.980855 | Non-sig. |
| PF3D7_0716200 | N/A | 1380.62782 | -0.41117694 | 0.42921341 | -0.95797784 | 0.33807393 | 0.8930363 | 1358.17007 | 1087.93959 | 1793.77386 | 1282.62775 | Non-sig. |
| PF3D7_0209200 | MTR3 | 679.845215 | -0.411338023 | 0.42745865 | -0.96228728 | 0.33590532 | 0.8920875 | 759.175716 | 641.0137 | 793.028343 | 526.1631 | Non-sig. |
| PF3D7_0301500 | N/A | 390.03445 | -0.411423772 | 0.50733809 | -0.81094595 | 0.41739671 | 0.90869129 | 294.812786 | 351.911907 | 595.466352 | 317.946754 | Non-sig. |
| PF3D7_0513000 | N/A | 613.034663 | -0.411432521 | 0.43330507 | -0.9495216 | 0.34235539 | 0.8930363 | 776.784543 | 555.164209 | 623.059543 | 497.130357 | Non-sig. |
| PF3D7_0204100 | N/A | 12115.1349 | -0.412079925 | 0.44773048 | -0.92037497 | 0.35737684 | 0.89552149 | 10981.7622 | 8828.19894 | 16685.4362 | 11965.1422 | Non-sig. |
| PF3D7_0823700 | TOM7 | 174.053673 | -0.412277193 | 0.52339882 | -0.78769225 | 0.43087674 | 0.91355742 | 141.29013 | 148.310274 | 256.005853 | 150.608433 | Non-sig. |
| PF3D7_1462300 | N/A | 3757.02538 | -0.412475562 | 0.48526261 | -0.85000483 | 0.3953224 | 0.90645858 | 2872.44438 | 2836.13279 | 5708.21582 | 3611.30852 | Non-sig. |
| PF3D7_1437400 | PANK2 | 1266.24739 | -0.412920154 | 0.41226235 | -1.0015956 | 0.31653895 | 0.88585961 | 1378.78621 | 1101.98663 | 1513.6175 | 1070.59923 | Non-sig. |
| PF3D7_1459800 | N/A | 1587.22629 | -0.414998569 | 0.52180113 | -0.79531942 | 0.42642772 | 0.91236574 | 1050.21611 | 1459.53091 | 2577.37655 | 1261.78161 | Non-sig. |
| PF3D7_0322400 | N/A | 1082.12409 | -0.415027643 | 0.48130203 | -0.86230188 | 0.38852142 | 0.90444842 | 1631.01225 | 954.872078 | 842.69349 | 899.918553 | Non-sig. |
| PF3D7_1313600 | N/A | 2943.6785 | -0.415853906 | 0.44896433 | -0.92625154 | 0.35431527 | 0.8930363 | 3048.07641 | 1941.39998 | 3681.86319 | 3103.37442 | Non-sig. |
| PF3D7_1452600 | N/A | 3196.20615 | -0.415926576 | 0.46677709 | -0.89106039 | 0.37289677 | 0.89654416 | 2782.35253 | 2225.42848 | 4524.96252 | 3252.08109 | Non-sig. |
| PF3D7_1233600 | AARP1 | 10099.5886 | -0.415999556 | 0.49837293 | -0.8347154 | 0.40387797 | 0.90723586 | 7642.48928 | 6947.15889 | 15448.6076 | 10360.0987 | Non-sig. |
| PF3D7_0305800 | N/A | 263.081203 | -0.416013571 | 0.73224285 | -0.56813606 | 0.56994258 | 0.94850402 | 525.407964 | 231.745691 | 76.3841999 | 218.786958 | Non-sig. |
| PF3D7_0520300 | LSM2 | 338.33144 | -0.416400695 | 0.44849401 | -0.92844203 | 0.35317832 | 0.8930363 | 349.215575 | 306.121232 | 424.330535 | 273.658421 | Non-sig. |
| PF3D7_1205700 | tGLO2 | 250.00397 | -0.4164586 | 0.46891963 | -0.88812362 | 0.37447426 | 0.89654416 | 244.782931 | 226.523532 | 326.763523 | 201.945895 | Non-sig. |
| PF3D7_0216300 | N/A | 486.88736 | -0.416521843 | 0.42847597 | -0.97210082 | 0.33100039 | 0.88994703 | 545.749604 | 406.557343 | 567.603808 | 427.638684 | Non-sig. |
| PF3D7_0414100 | N/A | 429.653558 | -0.416610063 | 0.47614564 | -0.87496352 | 0.38159375 | 0.90292925 | 364.567451 | 360.421875 | 617.720104 | 375.904802 | Non-sig. |
| PF3D7_1342700 | RPB12 | 142.439117 | -0.416731377 | 0.50924323 | -0.81833465 | 0.41316613 | 0.90793155 | 133.664396 | 125.819468 | 191.936668 | 118.335936 | Non-sig. |
| PF3D7_0912700 | N/A | 117.446264 | -0.416939392 | 0.53523228 | -0.77898776 | 0.43598692 | 0.91522571 | 163.783437 | 106.492392 | 104.973808 | 94.5354193 | Non-sig. |
| PF3D7_0819600 | N/A | 7429.34931 | -0.417628086 | 0.42067573 | -0.99275536 | 0.32082919 | 0.88694295 | 9804.90895 | 6564.35968 | 7189.71415 | 6158.41447 | Non-sig. |
| PF3D7_1468700 | eIF4A | 11293.193 | -0.418511076 | 0.55845101 | -0.74941412 | 0.45360764 | 0.92203286 | 6306.32564 | 10277.1525 | 19533.0001 | 9056.29381 | Non-sig. |
| PF3D7_1028400 | N/A | 584.200908 | -0.419356567 | 0.49198696 | -0.85237334 | 0.39400691 | 0.90601068 | 882.376069 | 530.184129 | 454.927449 | 469.315984 | Non-sig. |
| PF3D7_0501500 | RAP3 | 10931.7186 | -0.419808967 | 0.45674202 | -0.91913804 | 0.35802337 | 0.89552149 | 9133.73435 | 10044.2815 | 15888.2075 | 8660.65125 | Non-sig. |
| PF3D7_0311200 | N/A | 699.436309 | -0.420022931 | 0.45122772 | -0.93084471 | 0.3519339 | 0.8930363 | 715.454247 | 488.06803 | 885.511999 | 708.710958 | Non-sig. |
| PF3D7_0914600 | ELF1 | 789.290299 | -0.420791637 | 0.43508978 | -0.96713748 | 0.33347531 | 0.89204385 | 1043.73444 | 678.40243 | 763.609476 | 671.41485 | Non-sig. |
| PF3D7_1454000 | N/A | 456.738677 | -0.421007029 | 0.43610628 | -0.96537713 | 0.33435595 | 0.89204385 | 571.262979 | 398.137475 | 474.663151 | 382.891103 | Non-sig. |
| PF3D7_0703700 | N/A | 746.617438 | -0.421508231 | 0.42671811 | -0.98779082 | 0.3232551 | 0.88694295 | 775.400776 | 672.036187 | 934.330679 | 604.70211 | Non-sig. |
| PF3D7_1363700 | AMR1 | 1030.85004 | -0.421563835 | 0.44565283 | -0.94594674 | 0.34417575 | 0.8930363 | 1428.80118 | 900.162769 | 932.214216 | 862.221978 | Non-sig. |
| PF3D7_0200600 | RIF | 2.53871545 | -0.421947913 | 2.11371565 | -0.19962378 | 0.84177483 | 0.99128968 | 4.19312928 | 1.07422823 | 1.64139639 | 3.2461079 | Non-sig. |
| PF3D7_1148600 | N/A | 66.1706137 | -0.422281412 | 0.71013242 | -0.59465164 | 0.55207635 | 0.94170064 | 69.4134705 | 24.6845573 | 82.1171351 | 88.4672919 | Non-sig. |
| PF3D7_1201900 | N/A | 1514.21448 | -0.422499769 | 0.40997646 | -1.03054641 | 0.30275358 | 0.87475317 | 1674.7497 | 1325.82929 | 1793.94174 | 1262.33718 | Non-sig. |
| PF3D7_1474500 | SF3A1 | 1326.11835 | -0.42287742 | 0.45771541 | -0.92388722 | 0.35554504 | 0.89402637 | 1275.36825 | 899.031894 | 1762.65349 | 1367.41978 | Non-sig. |
| PF3D7_0312200 | N/A | 567.958715 | -0.422965 | 0.51736878 | -0.81753098 | 0.41362505 | 0.90793155 | 803.663019 | 330.419351 | 497.759968 | 639.992522 | Non-sig. |
| PF3D7_1230500 | N/A | 114.465698 | -0.423179828 | 0.59121633 | -0.71577832 | 0.47412824 | 0.92682482 | 78.9015616 | 93.3644948 | 183.130406 | 102.466329 | Non-sig. |
| PF3D7_0731600 | ACS5 | 6762.0466 | -0.42328776 | 0.4088851 | -1.03522421 | 0.30056421 | 0.87381251 | 7357.09022 | 5300.47583 | 8136.82176 | 6253.79858 | Non-sig. |
| PF3D7_1344700 | N/A | 462.876648 | -0.423312142 | 0.56271783 | -0.7522636 | 0.45189256 | 0.92081798 | 451.854159 | 211.340808 | 608.645023 | 579.666602 | Non-sig. |
| PF3D7_0317700 | N/A | 3254.95392 | -0.423574001 | 0.42317652 | -1.00093928 | 0.31685616 | 0.88585961 | 3435.71507 | 2398.16248 | 4022.94603 | 3162.99208 | Non-sig. |
| PF3D7_1242500 | N/A | 965.688818 | -0.423845383 | 0.43561975 | -0.97297099 | 0.33056772 | 0.88975609 | 1228.7076 | 932.807403 | 984.478787 | 716.76148 | Non-sig. |
| PF3D7_1020900 | ARF1 | 6414.68676 | -0.423953262 | 0.41690486 | -1.01690649 | 0.30919792 | 0.87940531 | 6394.23075 | 5210.15703 | 8306.57621 | 5747.78303 | Non-sig. |
| PF3D7_0802900 | N/A | 249.820149 | -0.424419472 | 0.45424698 | -0.93433636 | 0.35013041 | 0.8930363 | 296.001942 | 208.323956 | 276.639093 | 218.315606 | Non-sig. |
| PF3D7_1304100 | LigI | 855.189378 | -0.424612981 | 0.50824529 | -0.83544893 | 0.40346499 | 0.90723586 | 668.810068 | 567.513456 | 1291.20836 | 893.225624 | Non-sig. |
| PF3D7_1480000 | RIF | 2.53429763 | -0.424640181 | 2.11454184 | -0.200819 | 0.8408401 | 0.99128968 | 4.18982556 | 3.22339653 | 1.64116719 | 1.08280123 | Non-sig. |
| PF3D7_1238300 | CWC22 | 1824.56577 | -0.424746698 | 0.53190829 | -0.79853371 | 0.42456084 | 0.91236574 | 3027.93811 | 1637.12046 | 1154.85014 | 1478.35438 | Non-sig. |
| PF3D7_1338900 | N/A | 1983.25129 | -0.424761456 | 0.40851618 | -1.03976654 | 0.29844838 | 0.87330688 | 2313.07663 | 1610.45569 | 2233.17873 | 1776.29413 | Non-sig. |
| PF3D7_1322900 | N/A | 31.5901716 | -0.425020534 | 0.75249848 | -0.56481248 | 0.57220131 | 0.94850402 | 35.5083639 | 35.486523 | 36.9114214 | 18.4543779 | Non-sig. |
| PF3D7_0110700 | CAF1C | 2264.34433 | -0.425418944 | 0.40758622 | -1.04375204 | 0.29660013 | 0.87299304 | 2573.03879 | 2030.82665 | 2618.55089 | 1834.96099 | Non-sig. |
| PF3D7_0821800 | SEC61B | 2841.81287 | -0.425792681 | 0.41296696 | -1.03105751 | 0.30251385 | 0.87475317 | 3172.20158 | 2670.92009 | 3344.08757 | 2180.04222 | Non-sig. |
| PF3D7_1229000 | N/A | 735.98206 | -0.426152879 | 0.50640001 | -0.8415341 | 0.40004879 | 0.90645858 | 530.065725 | 629.641123 | 1157.43158 | 626.78981 | Non-sig. |
| PF3D7_1024500 | N/A | 559.084591 | -0.426366152 | 0.43666493 | -0.97641492 | 0.32885889 | 0.88909228 | 710.941754 | 511.079342 | 571.384034 | 442.933236 | Non-sig. |
| PF3D7_1204400 | G37 | 10.1281297 | -0.426490088 | 1.13250923 | -0.37658862 | 0.70647935 | 0.986213 | 12.5722541 | 6.44639406 | 10.6680047 | 10.8258661 | Non-sig. |
| PF3D7_0415000 | GET3 | 264.511675 | -0.426831331 | 0.50006671 | -0.85354878 | 0.39335505 | 0.90601068 | 236.27886 | 184.676087 | 370.249835 | 266.841919 | Non-sig. |

| | | | | | | | | | | | | |
|---|---|---|---|---|---|---|---|---|---|---|---|---|
| PF3D7_1368000 | N/A | 180.091369 | -0.426992169 | 0.52720076 | -0.80992328 | 0.41798427 | 0.90869129 | 268.99349 | 162.454012 | 144.324607 | 144.593366 | Non-sig. |
| PF3D7_1456700 | N/A | 1556.62206 | -0.427305931 | 0.47549937 | -0.89864667 | 0.36884089 | 0.89654416 | 1300.49079 | 1112.56202 | 2270.21162 | 1543.22382 | Non-sig. |
| PF3D7_1118900 | N/A | 22.2079559 | -0.428231705 | 0.83285709 | -0.51417189 | 0.60713182 | 0.95715771 | 31.341067 | 19.3546134 | 19.6870688 | 18.4490745 | Non-sig. |
| PF3D7_1010600 | eIF2beta | 4656.88934 | -0.428360735 | 0.41826104 | -1.02414688 | 0.30576591 | 0.87685698 | 6041.42009 | 4190.12033 | 4645.15778 | 3750.85918 | Non-sig. |
| PF3D7_0707300 | RAMA | 85119.5002 | -0.428453106 | 0.60653293 | -0.7063971 | 0.47994122 | 0.92880667 | 156731.722 | 74709.8648 | 38602.5114 | 70433.9028 | Non-sig. |
| PF3D7_1349800 | N/A | 171.477031 | -0.428737954 | 0.55230058 | -0.77627649 | 0.43758575 | 0.91561189 | 127.249632 | 164.209015 | 266.056441 | 128.393035 | Non-sig. |
| PF3D7_1248100 | N/A | 984.815767 | -0.429414277 | 0.42688128 | -1.00593374 | 0.31444745 | 0.88454055 | 982.174163 | 830.970104 | 1278.2936 | 847.825199 | Non-sig. |
| PF3D7_0910700 | ALP5a | 70.5995876 | -0.429562491 | 0.61062984 | -0.70347445 | 0.48176013 | 0.92880667 | 56.7917851 | 64.3948335 | 105.109591 | 56.1021407 | Non-sig. |
| PF3D7_1313000 | NEDD8 | 680.296132 | -0.429693776 | 0.45654137 | -0.94119351 | 0.3466057 | 0.8930363 | 871.449535 | 694.717838 | 690.399646 | 464.617508 | Non-sig. |
| PF3D7_1006700 | N/A | 474.591916 | -0.429891286 | 0.66055075 | -0.65080735 | 0.51517086 | 0.93247207 | 905.955291 | 395.029682 | 183.920075 | 413.462615 | Non-sig. |
| PF3D7_0505900 | MED11 | 652.077417 | -0.430108715 | 0.48422083 | -0.88824909 | 0.37440678 | 0.89654416 | 537.309334 | 629.304219 | 959.582045 | 482.114069 | Non-sig. |
| PF3D7_1316800 | SEC20 | 1870.14871 | -0.430482013 | 0.42745262 | -1.00708709 | 0.31389292 | 0.88410671 | 2141.4591 | 1853.96114 | 2152.76842 | 1332.40619 | Non-sig. |
| PF3D7_0422500 | BRR2 | 6653.9636 | -0.430546287 | 0.50734509 | -0.8486261 | 0.39608938 | 0.90645858 | 4806.15993 | 4676.21494 | 10472.6221 | 6660.85738 | Non-sig. |
| PF3D7_0935400 | GDV1 | 324.933472 | -0.430568888 | 0.53936845 | -0.79828342 | 0.42470604 | 0.91236574 | 224.289852 | 249.979441 | 521.555141 | 303.909454 | Non-sig. |
| PF3D7_1141800 | N/A | 1340.93109 | -0.431722027 | 0.46128183 | -0.93591813 | 0.34931534 | 0.8930363 | 1545.32913 | 832.781923 | 1534.83076 | 1450.78257 | Non-sig. |
| PF3D7_0804000 | N/A | 2694.68474 | -0.432298356 | 0.4826995 | -0.89558485 | 0.37047453 | 0.89654416 | 4135.28854 | 2167.18282 | 2055.84304 | 2420.42455 | Non-sig. |
| PF3D7_1115000 | N/A | 639.993677 | -0.432350484 | 0.43178233 | -1.00131584 | 0.31667414 | 0.88585961 | 766.258323 | 605.872714 | 704.15319 | 483.690482 | Non-sig. |
| PF3D7_0218600 | N/A | 3396.62987 | -0.432502569 | 0.41502572 | -1.04211028 | 0.29736055 | 0.87301237 | 3743.24278 | 2563.39005 | 4060.67793 | 3219.20873 | Non-sig. |
| PF3D7_1402400 | N/A | 827.513483 | -0.432599091 | 0.47611149 | -0.90860881 | 0.36355666 | 0.89567376 | 1231.51529 | 725.519279 | 670.071251 | 682.948112 | Non-sig. |
| PF3D7_1345000 | N/A | 65.884279 | -0.433044732 | 0.64788975 | -0.66839262 | 0.503883 | 0.93170321 | 103.968039 | 64.4209618 | 47.6157086 | 47.532407 | Non-sig. |
| PF3D7_1326200 | N/A | 75.8532618 | -0.433107346 | 0.5688268 | -0.76140461 | 0.44641544 | 0.91974025 | 75.6899844 | 59.0356974 | 98.532849 | 70.1545163 | Non-sig. |
| PF3D7_1410000 | EMC2 | 587.536517 | -0.433517865 | 0.44418303 | -0.97598926 | 0.32906979 | 0.88909228 | 560.243134 | 494.844599 | 789.886947 | 505.171389 | Non-sig. |
| PF3D7_1407500 | TRM112 | 224.721222 | -0.434998775 | 0.48782533 | -0.8917101 | 0.37254834 | 0.89654416 | 203.725589 | 184.675908 | 312.783311 | 197.700081 | Non-sig. |
| PF3D7_1368300 | N/A | 201.20662 | -0.435085912 | 0.51717015 | -0.84128195 | 0.40019 | 0.90645858 | 163.000749 | 169.882219 | 299.415267 | 172.528247 | Non-sig. |
| PF3D7_0709500 | N/A | 646.019841 | -0.435171661 | 0.42518994 | -1.0234759 | 0.30608289 | 0.87685698 | 768.628604 | 586.028105 | 716.85347 | 512.569186 | Non-sig. |
| PF3D7_1103500 | N/A | 44.4030356 | -0.436433372 | 0.66613544 | -0.65517213 | 0.51235694 | 0.93247207 | 62.0038401 | 34.3511388 | 40.2319808 | 41.0251828 | Non-sig. |
| PF3D7_1307000 | RRP40 | 502.668428 | -0.436518805 | 0.44265676 | -0.98613384 | 0.32406744 | 0.88694295 | 579.077723 | 490.571272 | 577.210126 | 363.814592 | Non-sig. |
| PF3D7_0827200 | N/A | 113.400745 | -0.436959317 | 0.64497619 | -0.67748131 | 0.49810062 | 0.93170321 | 197.90592 | 107.296288 | 63.2404118 | 85.1603611 | Non-sig. |
| PF3D7_0215600 | GET2 | 386.933619 | -0.437229581 | 0.56133753 | -0.77890673 | 0.43603466 | 0.91522571 | 642.545601 | 385.30299 | 247.990038 | 271.895846 | Non-sig. |
| PF3D7_1140200 | N/A | 3927.84998 | -0.437501321 | 0.64318012 | -0.68021587 | 0.49636779 | 0.93170321 | 7349.60211 | 4255.76038 | 1688.50507 | 2417.53234 | Non-sig. |
| PF3D7_0503300 | SRSF12 | 2765.65899 | -0.437970354 | 0.52021148 | -0.84190829 | 0.39983929 | 0.90645858 | 4539.86481 | 2462.34892 | 1824.98401 | 2235.43822 | Non-sig. |
| PF3D7_0521100 | N/A | 66.3236972 | -0.438328874 | 0.66197062 | -0.66215759 | 0.50787023 | 0.93170321 | 43.1926108 | 61.1460172 | 109.247689 | 51.7084716 | Non-sig. |
| PF3D7_1409600 | N/A | 691.900092 | -0.438458217 | 0.47637846 | -0.9203989 | 0.35736434 | 0.89552149 | 866.496993 | 423.802134 | 726.052136 | 751.249104 | Non-sig. |
| PF3D7_1113300 | UGT | 2465.74237 | -0.438566405 | 0.56748045 | -0.77283086 | 0.43962249 | 0.91706217 | 1350.07766 | 2207.80963 | 4324.92432 | 1980.15788 | Non-sig. |
| PF3D7_0811200 | EMC1 | 1419.65504 | -0.439551635 | 0.50206949 | -0.87547968 | 0.38131296 | 0.90292925 | 1017.58345 | 1168.23551 | 2250.63715 | 1242.16405 | Non-sig. |
| PF3D7_1445700 | N/A | 2297.04268 | -0.439612343 | 0.44709905 | -0.98325493 | 0.32548199 | 0.88694295 | 3217.19597 | 2147.75628 | 2071.69513 | 1751.52333 | Non-sig. |
| PF3D7_1304600 | N/A | 277.46545 | -0.439865897 | 0.48195995 | -0.91266069 | 0.36142102 | 0.89560738 | 289.69404 | 185.58578 | 349.097134 | 285.484847 | Non-sig. |
| PF3D7_0716500 | N/A | 518.269028 | -0.440449601 | 0.42659684 | -1.03247273 | 0.30185071 | 0.87436035 | 588.425345 | 449.80362 | 605.106359 | 429.740788 | Non-sig. |
| PF3D7_0114000 | EPF1 | 927.324156 | -0.440635434 | 0.66815548 | -0.65948039 | 0.50958734 | 0.93247207 | 1800.86242 | 852.76362 | 335.150944 | 720.519644 | Non-sig. |
| PF3D7_0624500 | APC11 | 2952.67366 | -0.4406715 | 0.44237538 | -0.99614834 | 0.31917808 | 0.88651753 | 3926.81981 | 2926.13129 | 2873.64434 | 2084.0992 | Non-sig. |
| PF3D7_1201800 | COX19 | 324.168542 | -0.440828855 | 0.45430373 | -0.97033951 | 0.33187729 | 0.88994703 | 323.653871 | 267.301107 | 422.84009 | 282.879102 | Non-sig. |
| PF3D7_0601200 | MC-2TM | 189.869396 | -0.441944716 | 0.47654887 | -0.92738594 | 0.35372619 | 0.8930363 | 195.201462 | 164.304725 | 242.154412 | 157.816984 | Non-sig. |
| PF3D7_1352600 | N/A | 10.1826062 | -0.442150609 | 1.12171816 | -0.39417264 | 0.69345358 | 0.98334178 | 13.6388275 | 8.59185398 | 9.84980298 | 8.64994021 | Non-sig. |
| PF3D7_1435500 | CLC | 1085.19104 | -0.443097182 | 0.42270257 | -1.04824813 | 0.2945243 | 0.87170077 | 1360.619 | 850.061046 | 1140.5621 | 989.522006 | Non-sig. |
| PF3D7_1466700 | NIP7 | 133.449597 | -0.443325149 | 0.52302049 | -0.84762482 | 0.39664694 | 0.90645858 | 154.121564 | 138.569377 | 153.474803 | 87.632644 | Non-sig. |
| PF3D7_0609000 | NUP637 | 13916.2889 | -0.443388477 | 0.5491038 | -0.80747661 | 0.41939195 | 0.90940098 | 9215.97302 | 8460.33253 | 22859.9234 | 15128.9265 | Non-sig. |
| PF3D7_1357700 | UTP21 | 661.366031 | -0.44342115 | 0.42347218 | -1.04710809 | 0.29504973 | 0.87170077 | 778.594752 | 594.280414 | 745.853768 | 526.73519 | Non-sig. |
| PF3D7_1429600 | GEXP03 | 223.62115 | -0.443946185 | 0.46626331 | -0.95213622 | 0.3410279 | 0.8930363 | 271.045499 | 171.958105 | 244.514935 | 206.966062 | Non-sig. |
| PF3D7_1114400 | N/A | 250.193333 | -0.444090633 | 0.45806045 | -0.96950225 | 0.33229466 | 0.89003898 | 263.872692 | 213.59984 | 312.843667 | 210.457131 | Non-sig. |
| PF3D7_0204800 | N/A | 546.464781 | -0.445175379 | 0.45058759 | -0.98798856 | 0.32315825 | 0.88694295 | 739.004423 | 500.197961 | 521.4 | 425.256741 | Non-sig. |
| PF3D7_1319900 | N/A | 147.759097 | -0.445270926 | 0.52702196 | -0.84488116 | 0.39817719 | 0.90645858 | 125.359532 | 127.656978 | 215.208421 | 122.811457 | Non-sig. |
| PF3D7_1365000 | N/A | 639.134855 | -0.445457475 | 0.6106812 | -0.72944357 | 0.46573037 | 0.92538923 | 1156.59608 | 634.331286 | 317.779104 | 447.832951 | Non-sig. |
| PF3D7_0629000 | GNA1 | 20.3891638 | -0.44558085 | 0.84736259 | -0.52584438 | 0.59899634 | 0.95655566 | 24.0613137 | 19.3472633 | 22.9731153 | 15.1749629 | Non-sig. |
| PF3D7_1472500 | N/A | 140.898422 | -0.445781972 | 0.53305817 | -0.83627266 | 0.40300153 | 0.90716539 | 206.203626 | 125.737332 | 118.979456 | 112.673273 | Non-sig. |

| | | | | | | | | | | | | |
|---|---|---|---|---|---|---|---|---|---|---|---|---|
| PF3D7_0802100 | AP2-LT | 16560.6931 | -0.446046834 | 0.44771258 | -0.99627944 | 0.31911439 | 0.88651753 | 20189.55 | 10539.6717 | 18011.6758 | 17501.8748 | Non-sig. |
| PF3D7_1357600 | N/A | 131.267948 | -0.446047077 | 0.55253924 | -0.8072677 | 0.41951227 | 0.90940098 | 118.959116 | 138.751458 | 183.712493 | 83.6487258 | Non-sig. |
| PF3D7_1120600 | N/A | 1718.10824 | -0.44614951 | 0.52135757 | -0.85574572 | 0.39213845 | 0.90601068 | 2780.69389 | 1234.51807 | 1182.95203 | 1674.26899 | Non-sig. |
| PF3D7_1444700 | N/A | 375.339331 | -0.446329465 | 0.45385917 | -0.9834096 | 0.32540589 | 0.88694295 | 444.835604 | 368.375739 | 421.079077 | 267.066904 | Non-sig. |
| PF3D7_1419200 | ATrx1 | 836.309802 | -0.446358532 | 0.46323905 | -0.96355981 | 0.33526667 | 0.8920875 | 724.239 | 761.807328 | 1204.84942 | 654.343458 | Non-sig. |
| PF3D7_0721700 | PSOP1 | 168.991467 | -0.446916802 | 0.51118968 | -0.87426805 | 0.3819723 | 0.90292925 | 148.302881 | 139.518643 | 241.427362 | 146.716982 | Non-sig. |
| PF3D7_0102900 | N/A | 1841.00521 | -0.447033169 | 0.53166122 | -0.84082335 | 0.40044691 | 0.90646318 | 1172.67237 | 1630.91354 | 3074.95069 | 1485.48424 | Non-sig. |
| PF3D7_1123700 | N/A | 452.802443 | -0.447106324 | 0.44515633 | -1.00438047 | 0.31519526 | 0.88486757 | 519.632452 | 439.434104 | 525.190365 | 326.95285 | Non-sig. |
| PF3D7_0323100 | N/A | 2105.54916 | -0.447176933 | 0.47935981 | -0.93286279 | 0.35089082 | 0.8930363 | 1717.82987 | 1504.8873 | 3140.46529 | 2059.01417 | Non-sig. |
| PF3D7_0508600 | N/A | 966.005081 | -0.447336143 | 0.41838249 | -1.06920379 | 0.28497785 | 0.86551387 | 1192.73251 | 792.820072 | 1036.5091 | 841.958642 | Non-sig. |
| PF3D7_0403700 | CLF1 | 3401.64253 | -0.447628556 | 0.40598068 | -1.10258585 | 0.27020706 | 0.8556331 | 4095.33174 | 2983.83524 | 3755.06034 | 2772.34281 | Non-sig. |
| PF3D7_1111400 | N/A | 595.492281 | -0.447816083 | 0.46021012 | -0.97306873 | 0.33051914 | 0.88975609 | 696.576087 | 621.036673 | 677.805085 | 386.551281 | Non-sig. |
| PF3D7_1222000 | N/A | 92.2357447 | -0.447879078 | 0.57851308 | -0.77419007 | 0.4388184 | 0.9169936 | 83.0134299 | 61.1897593 | 129.725235 | 95.014555 | Non-sig. |
| PF3D7_0511600 | ARNP | 906.13231 | -0.448054163 | 0.47048197 | -0.95233014 | 0.34092958 | 0.8930363 | 1334.86571 | 819.386258 | 756.829104 | 713.448171 | Non-sig. |
| PF3D7_0520900 | SAHH | 16132.0341 | -0.448105623 | 0.55315976 | -0.81008355 | 0.41789216 | 0.90869129 | 9291.03581 | 14620.6651 | 27943.4877 | 12672.9477 | Non-sig. |
| PF3D7_0616700 | N/A | 258.895868 | -0.448231821 | 0.48338994 | -0.92726758 | 0.35378762 | 0.8930363 | 338.987501 | 179.18168 | 258.713573 | 258.700719 | Non-sig. |
| PF3D7_1013100 | UTP13 | 2154.44882 | -0.448360239 | 0.40745979 | -1.1003791 | 0.27116698 | 0.8556331 | 2398.60676 | 1769.8313 | 2574.46915 | 1874.88809 | Non-sig. |
| PF3D7_1417700 | N/A | 1137.34792 | -0.448617189 | 0.41476134 | -1.08162731 | 0.27941817 | 0.86247429 | 1318.87895 | 1022.5648 | 1306.67 | 901.277923 | Non-sig. |
| PF3D7_1241600 | SAMC | 202.114582 | -0.448824008 | 0.52602221 | -0.85324155 | 0.39352537 | 0.90601068 | 172.448367 | 205.000373 | 293.969309 | 137.040279 | Non-sig. |
| PF3D7_0718700 | N/A | 221.883451 | -0.449040762 | 0.56435378 | -0.79567247 | 0.42622243 | 0.91236574 | 152.392808 | 220.057727 | 359.628992 | 155.454277 | Non-sig. |
| PF3D7_1112400 | N/A | 453.56785 | -0.449361256 | 0.45351478 | -0.99084149 | 0.32176299 | 0.88694295 | 460.336194 | 437.173543 | 586.830841 | 329.93082 | Non-sig. |
| PF3D7_1143900 | N/A | 59.23509 | -0.449696402 | 0.59509799 | -0.75566783 | 0.44984838 | 0.92081798 | 66.163039 | 47.241726 | 70.6028954 | 52.9326997 | Non-sig. |
| PF3D7_1118800 | ARC40 | 122.360262 | -0.450044359 | 0.55669887 | -0.80841615 | 0.41885106 | 0.90908353 | 178.502821 | 122.405247 | 104.259088 | 84.2738928 | Non-sig. |
| PF3D7_0105200 | N/A | 1149.72943 | -0.450224818 | 0.42438273 | -1.06089336 | 0.28873837 | 0.86724628 | 1315.12275 | 846.701871 | 1340.23619 | 1096.8569 | Non-sig. |
| PF3D7_0625200 | N/A | 386.360834 | -0.450315082 | 0.71810512 | -0.62708797 | 0.53060157 | 0.93351492 | 765.361521 | 417.535464 | 127.273104 | 235.273245 | Non-sig. |
| PF3D7_1132200 | TCP1 | 3024.68369 | -0.450701215 | 0.4521217 | -0.99685817 | 0.31883336 | 0.88651753 | 2615.79758 | 2707.43242 | 4370.63191 | 2404.87284 | Non-sig. |
| PF3D7_1022900 | ApiCOX13 | 786.727901 | -0.452264217 | 0.43643951 | -1.03625866 | 0.30008149 | 0.87381251 | 773.728279 | 705.475045 | 1044.20092 | 623.507359 | Non-sig. |
| PF3D7_0817900 | HMGB2 | 5469.22814 | -0.452427789 | 0.40284446 | -1.12308305 | 0.26140222 | 0.85030738 | 6517.59306 | 4532.49528 | 6122.1301 | 4704.69414 | Non-sig. |
| PF3D7_0626700 | N/A | 1546.08508 | -0.45284596 | 0.52996794 | -0.85447802 | 0.39284018 | 0.90601068 | 1017.5117 | 1461.55341 | 2555.70479 | 1149.57044 | Non-sig. |
| PF3D7_1415800 | KsgA1 | 357.903321 | -0.453502519 | 0.48885235 | -0.92768811 | 0.35356938 | 0.8930363 | 315.586732 | 349.85465 | 511.6099 | 254.562003 | Non-sig. |
| PF3D7_0618700 | N/A | 652.133405 | -0.453527174 | 0.46064009 | -0.98455863 | 0.32484092 | 0.88694295 | 886.427925 | 641.186551 | 621.346542 | 459.572603 | Non-sig. |
| PF3D7_0315000 | N/A | 180.269074 | -0.45509637 | 0.51667575 | -0.8808162 | 0.37841731 | 0.9000162 | 251.346033 | 180.518955 | 165.752796 | 123.458511 | Non-sig. |
| PF3D7_0503900 | N/A | 212.935136 | -0.455538265 | 0.46478968 | -0.98009548 | 0.32703899 | 0.88852997 | 256.877315 | 188.837185 | 235.71758 | 170.308464 | Non-sig. |
| PF3D7_1103600 | ALP2a | 258.912532 | -0.455573751 | 0.53749863 | -0.84758123 | 0.39667123 | 0.90645858 | 194.624586 | 181.363022 | 404.036857 | 255.625664 | Non-sig. |
| PF3D7_1443200 | N/A | 219.437257 | -0.45619551 | 0.46037344 | -0.99092491 | 0.32172225 | 0.88694295 | 258.86996 | 184.568072 | 248.839664 | 185.471332 | Non-sig. |
| PF3D7_0922600 | N/A | 1668.20374 | -0.456253458 | 0.4984494 | -0.91534559 | 0.36001023 | 0.89560738 | 1214.15146 | 1354.28429 | 2645.17126 | 1459.20794 | Non-sig. |
| PF3D7_1444400 | N/A | 129.478821 | -0.456821681 | 0.5051995 | -0.90424016 | 0.36586807 | 0.89567376 | 133.638175 | 106.235306 | 165.891791 | 112.150011 | Non-sig. |
| PF3D7_0515500 | GEP1 | 42.7140185 | -0.45687031 | 0.66814582 | -0.68378832 | 0.49410884 | 0.93170321 | 42.10332 | 31.1168222 | 56.6689629 | 40.9669688 | Non-sig. |
| PF3D7_1321200 | N/A | 362.422318 | -0.456884904 | 0.47270532 | -0.96653218 | 0.33377795 | 0.89204385 | 515.903095 | 303.584852 | 322.967015 | 307.234311 | Non-sig. |
| PF3D7_1004700 | N/A | 239.671665 | -0.456972946 | 0.50020364 | -0.9135738 | 0.36094084 | 0.89560738 | 349.613894 | 215.826991 | 205.230122 | 188.015652 | Non-sig. |
| PF3D7_0526900 | N/A | 1907.28085 | -0.457177431 | 0.41601068 | -1.098956 | 0.27178726 | 0.8556331 | 2073.97148 | 1753.10735 | 2339.91836 | 1462.12621 | Non-sig. |
| PF3D7_0821500 | NEP1 | 179.591124 | -0.457215692 | 0.4767366 | -0.95905306 | 0.33753202 | 0.89233763 | 223.603353 | 155.700691 | 192.09183 | 146.968623 | Non-sig. |
| PF3D7_1357200 | GluRS | 722.241816 | -0.457353995 | 0.43085438 | -1.0615048 | 0.28846055 | 0.86724628 | 798.541582 | 681.43963 | 872.960029 | 536.026021 | Non-sig. |
| PF3D7_0318400 | N/A | 76.797246 | -0.457951935 | 0.7137408 | -0.64162219 | 0.52111852 | 0.93247207 | 141.881301 | 64.4073512 | 36.1270392 | 64.7732928 | Non-sig. |
| PF3D7_1251600 | N/A | 832.389787 | -0.458315517 | 0.51425117 | -0.89122892 | 0.37280637 | 0.89654416 | 1343.35941 | 719.939147 | 583.956718 | 682.303869 | Non-sig. |
| PF3D7_0720600 | N/A | 870.091921 | -0.458605635 | 0.44511733 | -1.0303028 | 0.30286788 | 0.87475317 | 1123.57092 | 855.301909 | 891.017411 | 610.477441 | Non-sig. |
| PF3D7_1430400 | ATG5 | 174.203468 | -0.458675919 | 0.50636834 | -0.90581477 | 0.3650339 | 0.89567376 | 167.456955 | 123.374331 | 235.733699 | 170.248886 | Non-sig. |
| PF3D7_1222100 | N/A | 649.380339 | -0.458736025 | 0.64733002 | -0.70865866 | 0.47853632 | 0.92880667 | 264.952516 | 589.260113 | 1238.26027 | 505.048457 | Non-sig. |
| PF3D7_1028800 | N/A | 1074.63324 | -0.458782469 | 0.42424455 | -1.08141039 | 0.2795146 | 0.86247429 | 1313.76078 | 1005.42119 | 1174.46072 | 804.890245 | Non-sig. |
| PF3D7_1034100 | N/A | 6.46279316 | -0.458885829 | 1.39384667 | -0.3292226 | 0.74198744 | 0.98954916 | 9.26804442 | 7.56046558 | 5.72425686 | 3.29840579 | Non-sig. |
| PF3D7_0500800 | MESA | 26257.0023 | -0.458927207 | 0.67332243 | -0.6815861 | 0.49550071 | 0.93170321 | 51944.2597 | 23652.7132 | 8852.86101 | 20578.1753 | Non-sig. |
| PF3D7_0919400 | PDI-Trans | 2377.74784 | -0.459612439 | 0.48237883 | -0.95280391 | 0.34068943 | 0.8930363 | 1823.81186 | 1996.93844 | 3682.55031 | 2007.69073 | Non-sig. |
| PF3D7_1415000 | UDG | 580.041139 | -0.459637056 | 0.44564708 | -1.03139251 | 0.30235679 | 0.87475317 | 789.519593 | 488.745556 | 553.991519 | 487.907886 | Non-sig. |
| PF3D7_1364400 | N/A | 2747.6468 | -0.459745643 | 0.43917515 | -1.04683893 | 0.29517387 | 0.87170077 | 2789.77641 | 1901.52641 | 3573.6457 | 2725.63866 | Non-sig. |

| | | | | | | | | | | | | |
|---|---|---|---|---|---|---|---|---|---|---|---|---|
| PF3D7_0530700 | N/A | 395.75568 | -0.460094345 | 0.4861793 | -0.94634704 | 0.34397161 | 0.8930363 | 330.765128 | 316.464747 | 585.657831 | 350.135017 | Non-sig. |
| PF3D7_1355900 | N/A | 787.28533 | -0.460428474 | 0.42408497 | -1.08569863 | 0.27761235 | 0.85972841 | 838.222817 | 696.533212 | 985.406151 | 628.97914 | Non-sig. |
| PF3D7_0517100 | N/A | 204.056852 | -0.461360984 | 0.4738267 | -0.9736914 | 0.33020979 | 0.88975609 | 261.874452 | 162.060118 | 211.040523 | 181.252314 | Non-sig. |
| PF3D7_0831100 | SURF8.1 | 97.2275376 | -0.461982034 | 0.62120665 | -0.74368495 | 0.4570671 | 0.92288312 | 162.308144 | 83.6629728 | 63.2535833 | 79.6854504 | Non-sig. |
| PF3D7_0817400 | N/A | 433.218082 | -0.462310547 | 0.47805356 | -0.96706851 | 0.33350979 | 0.89204385 | 370.609211 | 380.883903 | 633.243913 | 348.135303 | Non-sig. |
| PF3D7_1214600 | N/A | 204.128596 | -0.462459048 | 0.5321749 | -0.86899823 | 0.3848481 | 0.90292925 | 166.779278 | 202.994181 | 306.153666 | 140.587261 | Non-sig. |
| PF3D7_0903400 | DDX60 | 5750.32143 | -0.463243362 | 0.50827233 | -0.91140779 | 0.36208055 | 0.89567376 | 4019.94096 | 4373.69372 | 9311.09599 | 5296.55507 | Non-sig. |
| PF3D7_0617200 | N/A | 4762.2827 | -0.463610855 | 0.63627956 | -0.72862761 | 0.46622949 | 0.92538923 | 8985.39446 | 4909.87515 | 2056.82473 | 3097.03647 | Non-sig. |
| PF3D7_0810100 | N/A | 235.80598 | -0.464369598 | 0.49554225 | -0.93709386 | 0.34871028 | 0.8930363 | 303.181481 | 243.7984 | 243.779764 | 152.464273 | Non-sig. |
| PF3D7_1235900 | XAB2 | 2612.54596 | -0.46466808 | 0.46777375 | -0.99336075 | 0.32053418 | 0.88694295 | 3939.60645 | 2191.66501 | 2120.02455 | 2198.88782 | Non-sig. |
| PF3D7_0603400 | TEX1 | 1713.78169 | -0.464933455 | 0.4937509 | -0.94163566 | 0.34637921 | 0.8930363 | 2584.68016 | 1135.96702 | 1390.69584 | 1743.78376 | Non-sig. |
| PF3D7_0628500 | N/A | 72.4107945 | -0.465607127 | 0.57288845 | -0.8127361 | 0.41636938 | 0.90793155 | 73.5030181 | 56.907299 | 94.4082556 | 64.8246054 | Non-sig. |
| PF3D7_0816800 | DMC1 | 46.2293291 | -0.465878343 | 0.65235154 | -0.71415229 | 0.47513302 | 0.92682482 | 59.7205607 | 45.1242602 | 47.5960457 | 32.47645 | Non-sig. |
| PF3D7_1021600 | DPA | 818.868819 | -0.466728404 | 0.55154227 | -0.84622417 | 0.3974277 | 0.90645858 | 507.214174 | 768.31935 | 1392.84388 | 607.097873 | Non-sig. |
| PF3D7_1450500 | N/A | 1947.61053 | -0.466731179 | 0.45951583 | -1.01570208 | 0.30977128 | 0.87951138 | 2578.13924 | 1248.62815 | 1941.85254 | 2021.82219 | Non-sig. |
| PF3D7_0904000 | N/A | 3215.88602 | -0.467002261 | 0.40817786 | -1.14411463 | 0.2525761 | 0.85030738 | 3859.5213 | 2518.86457 | 3604.28343 | 2880.87479 | Non-sig. |
| PF3D7_1351200 | N/A | 1205.71394 | -0.467401625 | 0.42240105 | -1.10653518 | 0.26849495 | 0.8556331 | 1303.54473 | 910.676694 | 1495.04673 | 1113.58761 | Non-sig. |
| PF3D7_1249100 | N/A | 682.010462 | -0.467836126 | 0.42675354 | -1.09626771 | 0.27296164 | 0.85618956 | 780.182257 | 626.816784 | 803.070689 | 517.972118 | Non-sig. |
| PF3D7_1015000 | N/A | 1723.10982 | -0.468123511 | 0.42104178 | -1.11182198 | 0.2662147 | 0.85416239 | 2117.67924 | 1616.40557 | 1882.8706 | 1275.48389 | Non-sig. |
| PF3D7_0110300 | N/A | 6.52493644 | -0.468466183 | 1.3827311 | -0.33879775 | 0.7347621 | 0.98909017 | 10.3080897 | 6.52327772 | 4.88054612 | 4.38783224 | Non-sig. |
| PF3D7_0715700 | N/A | 152.316632 | -0.468485157 | 0.49240711 | -0.95141833 | 0.34139206 | 0.8930363 | 169.796218 | 113.872989 | 183.831506 | 141.765814 | Non-sig. |
| PF3D7_1332700 | DBP8 | 580.592368 | -0.468679983 | 0.43687824 | -1.07279316 | 0.28336393 | 0.86551387 | 584.211081 | 497.862557 | 763.805352 | 476.49048 | Non-sig. |
| PF3D7_1345900 | SPC25 | 998.101497 | -0.468818801 | 0.45096044 | -1.03960073 | 0.29852544 | 0.87330688 | 1366.51295 | 957.887082 | 951.392148 | 716.613807 | Non-sig. |
| PF3D7_1232100 | CPN60 | 4117.20752 | -0.468842096 | 0.41079994 | -1.14129057 | 0.25374903 | 0.85030738 | 5044.47684 | 3164.17083 | 4516.34804 | 3743.83437 | Non-sig. |
| PF3D7_0825000 | N/A | 4054.03024 | -0.469277685 | 0.55663249 | -0.84306557 | 0.39919178 | 0.90645858 | 6871.52049 | 2572.23513 | 2544.02585 | 4228.33948 | Non-sig. |
| PF3D7_1029300 | N/A | 541.267275 | -0.469670676 | 0.45970852 | -1.02167059 | 0.30693684 | 0.87685698 | 766.284788 | 473.189538 | 491.131501 | 434.463274 | Non-sig. |
| PF3D7_0822500 | DLC1 | 50.2387369 | -0.469750765 | 0.67438074 | -0.6965661 | 0.48607438 | 0.93170321 | 77.454876 | 48.3609187 | 39.386105 | 35.7530479 | Non-sig. |
| PF3D7_0418500 | BET3 | 266.013981 | -0.470075876 | 0.48889612 | -0.96150461 | 0.33629851 | 0.8920875 | 381.842587 | 206.370809 | 236.2949 | 239.547629 | Non-sig. |
| PF3D7_1472700 | N/A | 1087.66873 | -0.470276 | 0.43296474 | -1.08617622 | 0.27740104 | 0.85972841 | 1251.33481 | 760.784527 | 1275.42911 | 1063.12649 | Non-sig. |
| PF3D7_0514500 | N/A | 19.2706018 | -0.470289549 | 0.87327845 | -0.53853333 | 0.5902089 | 0.95315309 | 20.9552366 | 19.3387996 | 23.7981545 | 12.9902163 | Non-sig. |
| PF3D7_0408400 | N/A | 512.745917 | -0.470421722 | 0.43966031 | -1.06996631 | 0.28463447 | 0.86551387 | 672.333036 | 413.148865 | 519.018745 | 446.483022 | Non-sig. |
| PF3D7_1445100 | N/A | 755.455148 | -0.470521127 | 0.47174056 | -0.99741503 | 0.3185631 | 0.88651753 | 662.476896 | 543.91493 | 1092.43061 | 722.99815 | Non-sig. |
| PF3D7_1128000 | N/A | 3401.65903 | -0.470598851 | 0.45528518 | -1.03363535 | 0.30130667 | 0.87381251 | 3437.67955 | 2176.68663 | 4465.3617 | 3526.90825 | Non-sig. |
| PF3D7_1410100 | N/A | 64.303568 | -0.470925832 | 0.61435537 | -0.76653653 | 0.44335712 | 0.91969696 | 94.5497228 | 56.8995503 | 55.0077637 | 50.7572351 | Non-sig. |
| PF3D7_1107000 | LSM4 | 551.996898 | -0.471508457 | 0.43219294 | -1.0909675 | 0.27528718 | 0.85802262 | 594.026573 | 495.05577 | 688.708968 | 430.196281 | Non-sig. |
| PF3D7_1248000 | N/A | 248.583769 | -0.472143674 | 0.64307544 | -0.7341964 | 0.46282904 | 0.92495974 | 114.479744 | 198.626484 | 463.028752 | 218.200095 | Non-sig. |
| PF3D7_0809250 | C4AP5 | 182.328877 | -0.472628299 | 0.49192191 | -0.96077911 | 0.33666324 | 0.8920875 | 218.979911 | 180.413059 | 204.895402 | 125.027135 | Non-sig. |
| PF3D7_0313600 | N/A | 487.971784 | -0.472866756 | 0.61220714 | -0.77239667 | 0.43987952 | 0.91706217 | 881.825356 | 493.614837 | 252.945116 | 323.501826 | Non-sig. |
| PF3D7_0527100 | UBC13 | 4082.0711 | -0.473452036 | 0.42623447 | -1.11077837 | 0.26666376 | 0.85416239 | 4464.26393 | 3978.00956 | 5027.55004 | 2858.46087 | Non-sig. |
| PF3D7_1329200 | N/A | 313.747692 | -0.473668977 | 0.45634187 | -1.03796957 | 0.29928422 | 0.87358937 | 411.11211 | 246.864157 | 318.599529 | 278.414974 | Non-sig. |
| PF3D7_0608500 | N/A | 5348.20574 | -0.474174606 | 0.46852927 | -1.01204905 | 0.3115146 | 0.8804206 | 4482.59876 | 5089.32241 | 7955.72036 | 3865.18142 | Non-sig. |
| PF3D7_0504800 | N/A | 10080.8267 | -0.47424161 | 0.49396797 | -0.96006551 | 0.33702225 | 0.8920875 | 8158.97276 | 6491.4366 | 15286.6766 | 10386.221 | Non-sig. |
| PF3D7_0524600 | N/A | 2040.42714 | -0.475315328 | 0.42427173 | -1.12030874 | 0.26258222 | 0.85086275 | 2466.2658 | 1959.1388 | 2280.8703 | 1455.43366 | Non-sig. |
| PF3D7_1205900 | N/A | 3178.85969 | -0.475424142 | 0.43121088 | -1.10253282 | 0.2702301 | 0.8556331 | 4265.04288 | 2360.26223 | 3131.0218 | 2959.11183 | Non-sig. |
| PF3D7_1457200 | TRX1 | 4728.99504 | -0.475470512 | 0.45048196 | -1.0554707 | 0.2912101 | 0.86724628 | 4163.27482 | 4296.57548 | 6839.07346 | 3617.05639 | Non-sig. |
| PF3D7_1137500 | N/A | 1076.97963 | -0.475879256 | 0.44990807 | -1.05772554 | 0.29018059 | 0.86724628 | 1028.12141 | 1029.66367 | 1477.72712 | 772.406303 | Non-sig. |
| PF3D7_1320500 | SNAP23 | 479.096005 | -0.477001631 | 0.43396518 | -1.09917029 | 0.2716938 | 0.8556331 | 562.709277 | 435.956137 | 552.473849 | 365.244757 | Non-sig. |
| PF3D7_1427400 | N/A | 461.788974 | -0.477310133 | 0.53779291 | -0.8875352 | 0.37479083 | 0.89654416 | 327.601743 | 314.1977 | 747.14612 | 458.210333 | Non-sig. |
| PF3D7_1473200 | N/A | 4172.33427 | -0.477336966 | 0.42044172 | -1.13532255 | 0.2562402 | 0.85030738 | 4197.01504 | 3713.01618 | 5515.52506 | 3263.78083 | Non-sig. |
| PF3D7_1353900 | N/A | 3285.10013 | -0.47783024 | 0.43262134 | -1.10449992 | 0.26937634 | 0.8556331 | 3219.55237 | 3063.81078 | 4428.69494 | 2428.34242 | Non-sig. |
| PF3D7_0305900 | N/A | 181.532724 | -0.478488458 | 0.47326476 | -1.01103758 | 0.31199844 | 0.88111287 | 202.825922 | 148.353963 | 219.864618 | 155.086391 | Non-sig. |
| PF3D7_0913300 | N/A | 878.895406 | -0.478503864 | 0.42434483 | -1.12762978 | 0.25947631 | 0.85030738 | 918.910193 | 761.824426 | 1127.63136 | 707.215652 | Non-sig. |
| PF3D7_0320300 | CCT5 | 4018.5548 | -0.478581258 | 0.51076086 | -0.93699674 | 0.34876023 | 0.8930363 | 2795.3656 | 3648.97379 | 6562.41242 | 3067.46737 | Non-sig. |
| PF3D7_0904700 | HU | 648.006026 | -0.478787534 | 0.49893971 | -0.95961001 | 0.33725153 | 0.8920875 | 527.04557 | 442.391644 | 981.820904 | 640.765985 | Non-sig. |

| | | | | | | | | | | | | |
|---|---|---|---|---|---|---|---|---|---|---|---|---|
| PF3D7_0810800 | PPPK-DHPS | 1013.35433 | -0.478790237 | 0.45731515 | -1.04695906 | 0.29511846 | 0.87170077 | 897.840769 | 907.756377 | 1461.89142 | 785.928753 | Non-sig. |
| PF3D7_1223500 | N/A | 3002.13142 | -0.478828857 | 0.43093389 | -1.11114227 | 0.26650711 | 0.85416239 | 3274.06646 | 2064.48298 | 3717.48199 | 2952.49424 | Non-sig. |
| PF3D7_1306200 | N/A | 1001.01077 | -0.480163177 | 0.4783715 | -1.00374536 | 0.31550137 | 0.88486757 | 890.547127 | 663.097241 | 1441.38026 | 1009.01845 | Non-sig. |
| PF3D7_0309100 | OMD | 97.8162158 | -0.481414028 | 0.56762788 | -0.84811554 | 0.39637363 | 0.90645858 | 145.284003 | 75.2578072 | 82.8555007 | 87.8675525 | Non-sig. |
| PF3D7_1135800 | N/A | 89.8115302 | -0.481470763 | 0.55117641 | -0.87353297 | 0.38237265 | 0.90292925 | 121.613513 | 79.4905505 | 87.816338 | 70.3257189 | Non-sig. |
| PF3D7_1303300 | N/A | 253.411387 | -0.481538885 | 0.48222728 | -0.99857246 | 0.31800184 | 0.88651753 | 357.599189 | 220.012 | 233.214337 | 202.820023 | Non-sig. |
| PF3D7_1314900 | P44 | 661.881983 | -0.482091883 | 0.53540732 | -0.90042079 | 0.36789636 | 0.89615989 | 440.617535 | 601.035566 | 1101.9888 | 503.88603 | Non-sig. |
| PF3D7_0827500 | N/A | 1567.91009 | -0.482155939 | 0.40938235 | -1.17776434 | 0.23889057 | 0.84066259 | 1806.04228 | 1261.42433 | 1848.94333 | 1355.23042 | Non-sig. |
| PF3D7_0405500 | N/A | 101.067052 | -0.482504283 | 0.53445134 | -0.90280301 | 0.36663045 | 0.89567376 | 101.061068 | 88.2367957 | 134.467971 | 80.5023745 | Non-sig. |
| PF3D7_1244300 | N/A | 119.648899 | -0.482945013 | 0.57605408 | -0.8383675 | 0.40182434 | 0.90667178 | 182.560751 | 120.979214 | 96.6041418 | 78.4514879 | Non-sig. |
| PF3D7_1447000 | N/A | 9630.20993 | -0.483089525 | 0.40799574 | -1.18405532 | 0.23639116 | 0.83828834 | 11853.2006 | 8649.18156 | 10602.1966 | 7416.26095 | Non-sig. |
| PF3D7_1027100 | MPP10 | 792.35553 | -0.483255071 | 0.55013734 | -0.87842623 | 0.37971245 | 0.90130671 | 1359.29257 | 695.24131 | 488.692764 | 626.195475 | Non-sig. |
| PF3D7_0824600 | DRE2 | 529.421173 | -0.483381698 | 0.43989332 | -1.0988612 | 0.27182861 | 0.8556331 | 681.39516 | 483.191666 | 553.30263 | 399.795239 | Non-sig. |
| PF3D7_1425200 | N/A | 247.524256 | -0.483606358 | 0.49475286 | -0.97747056 | 0.32833625 | 0.88909228 | 215.701438 | 214.620621 | 361.344433 | 198.430534 | Non-sig. |
| PF3D7_0933600 | QCR1 | 2350.80178 | -0.483618458 | 0.42190712 | -1.14626759 | 0.25168445 | 0.85030738 | 2445.7793 | 2153.34285 | 3036.44303 | 1767.64193 | Non-sig. |
| PF3D7_1009500 | N/A | 874.118012 | -0.484007979 | 0.43078196 | -1.12355675 | 0.26120111 | 0.85030738 | 1152.0015 | 771.591237 | 886.914803 | 685.964509 | Non-sig. |
| PF3D7_1470300 | N/A | 157.018622 | -0.484112503 | 0.49462601 | -0.97874454 | 0.32770622 | 0.88852997 | 203.772413 | 119.172692 | 162.55538 | 142.574003 | Non-sig. |
| PF3D7_1237600 | PWP1 | 552.293025 | -0.484419234 | 0.44891068 | -1.07909937 | 0.28054343 | 0.86335738 | 755.498524 | 490.404944 | 532.985185 | 430.283448 | Non-sig. |
| PF3D7_0321400 | pPK1 | 2529.93625 | -0.484455497 | 0.41539586 | -1.16625018 | 0.24351331 | 0.84313671 | 3215.20577 | 1957.04173 | 2686.42495 | 2261.07254 | Non-sig. |
| PF3D7_1459700 | N/A | 360.609988 | -0.48449239 | 0.4906002 | -0.98755034 | 0.32337292 | 0.88694295 | 323.728458 | 354.231547 | 517.281622 | 247.198325 | Non-sig. |
| PF3D7_0721900 | N/A | 1131.78868 | -0.484542543 | 0.42506657 | -1.13992154 | 0.25431899 | 0.85030738 | 1463.53803 | 1013.17415 | 1176.75131 | 873.691243 | Non-sig. |
| PF3D7_1430800 | PIP3 | 181.363439 | -0.484556865 | 0.47410303 | -1.02204971 | 0.30675737 | 0.87685698 | 202.96646 | 144.890701 | 220.071005 | 157.525589 | Non-sig. |
| PF3D7_1346600 | C2AP4 | 454.680395 | -0.484902833 | 0.45455945 | -1.06675339 | 0.28608321 | 0.86595658 | 471.267914 | 437.057529 | 589.391515 | 321.004623 | Non-sig. |
| PF3D7_0109200 | CFIM25 | 386.20192 | -0.484976676 | 0.45328084 | -1.06992537 | 0.2846529 | 0.86551387 | 380.337348 | 308.107934 | 520.530212 | 335.832186 | Non-sig. |
| PF3D7_1354500 | ADSS | 1651.26198 | -0.485272695 | 0.45525829 | -1.06592829 | 0.28645606 | 0.86595658 | 1452.93021 | 1483.07469 | 2399.60902 | 1269.43403 | Non-sig. |
| PF3D7_0926400 | N/A | 782.334869 | -0.485636047 | 0.53685853 | -0.90458849 | 0.36568344 | 0.89567376 | 525.18323 | 556.900428 | 1300.0769 | 747.178919 | Non-sig. |
| PF3D7_1147400 | N/A | 788.319575 | -0.485805963 | 0.49065378 | -0.99011968 | 0.32211563 | 0.88694295 | 678.552839 | 798.736669 | 1160.85873 | 515.130058 | Non-sig. |
| PF3D7_1246800 | SRPRB | 3082.13829 | -0.485934146 | 0.45510296 | -1.06774553 | 0.28563532 | 0.86572992 | 2778.73521 | 2935.15084 | 4413.76395 | 2200.90316 | Non-sig. |
| PF3D7_1027400 | RPB7 | 428.955714 | -0.486257736 | 0.44674525 | -1.08844522 | 0.27639861 | 0.85889199 | 432.856989 | 376.818143 | 568.146664 | 338.001062 | Non-sig. |
| PF3D7_1357800 | CCT4 | 2037.82475 | -0.487014932 | 0.48009895 | -1.01440532 | 0.31038939 | 0.88032279 | 1600.43346 | 1772.42776 | 3156.38963 | 1622.04815 | Non-sig. |
| PF3D7_0716400 | N/A | 1275.87525 | -0.487066118 | 0.43423255 | -1.12167113 | 0.2620023 | 0.85030738 | 1685.9544 | 1190.18226 | 1292.64258 | 934.721755 | Non-sig. |
| PF3D7_1210300 | N/A | 123.583199 | -0.487096698 | 0.51536854 | -0.94514247 | 0.34458614 | 0.8930363 | 122.614858 | 104.208277 | 165.772857 | 101.736802 | Non-sig. |
| PF3D7_1126200 | RPS18 | 4390.6113 | -0.487348153 | 0.43862539 | -1.11108057 | 0.26653367 | 0.85416239 | 4101.61316 | 3992.77172 | 6148.63062 | 3319.42969 | Non-sig. |
| PF3D7_0810300 | PPM5 | 18471.6902 | -0.487621138 | 0.51762081 | -0.94204314 | 0.34617055 | 0.8930363 | 30567.3413 | 13618.5785 | 12560.917 | 17139.9241 | Non-sig. |
| PF3D7_1405600 | RNR | 3973.05911 | -0.488028233 | 0.47887127 | -1.01912198 | 0.30814506 | 0.87724478 | 3132.72041 | 3575.01816 | 6144.43997 | 3040.05791 | Non-sig. |
| PF3D7_1203600 | CC1HL | 166.194011 | -0.488264605 | 0.49285502 | -0.99068608 | 0.32183889 | 0.88694295 | 198.48578 | 159.978418 | 189.6421 | 116.669747 | Non-sig. |
| PF3D7_0825900 | N/A | 67.958956 | -0.48865424 | 0.68316522 | -0.71527973 | 0.47443621 | 0.92682482 | 117.868748 | 64.3827891 | 41.0632146 | 48.521072 | Non-sig. |
| PF3D7_0727400 | N/A | 3206.79215 | -0.488706579 | 0.5666765 | -0.8624084 | 0.38846282 | 0.90444842 | 1791.59593 | 2892.98374 | 5697.69028 | 2444.89864 | Non-sig. |
| PF3D7_0622600 | QCR9 | 1010.67769 | -0.489179379 | 0.4517824 | -1.08277653 | 0.27890762 | 0.8621077 | 939.03472 | 934.204851 | 1421.58509 | 747.886108 | Non-sig. |
| PF3D7_0816600 | ClpB1 | 1568.00513 | -0.48920368 | 0.48446929 | -1.00977231 | 0.31260439 | 0.88188394 | 1233.51318 | 1190.7681 | 2428.87518 | 1418.86406 | Non-sig. |
| PF3D7_0211800 | AsnRS | 3439.87011 | -0.489627329 | 0.50470735 | -0.97012126 | 0.33198606 | 0.88994703 | 2431.66058 | 2941.36831 | 5604.13231 | 2782.31926 | Non-sig. |
| PF3D7_0215000 | ACS9 | 1305.33583 | -0.490901112 | 0.48305594 | -1.01624072 | 0.30951477 | 0.87951138 | 2004.33336 | 971.815513 | 1046.53405 | 1198.66039 | Non-sig. |
| PF3D7_0211600 | ALG14 | 285.623091 | -0.492037825 | 0.47881096 | -1.02762441 | 0.30412654 | 0.87568674 | 273.815694 | 268.78469 | 393.76368 | 206.128299 | Non-sig. |
| PF3D7_0812220 | YihA3 | 397.40096 | -0.492451305 | 0.44111748 | -1.11637222 | 0.26426284 | 0.85162075 | 501.575064 | 350.101234 | 427.658534 | 310.269007 | Non-sig. |
| PF3D7_1369200 | N/A | 1490.57012 | -0.492944393 | 0.51264771 | -0.96156559 | 0.33626786 | 0.8920875 | 1126.77793 | 978.187187 | 2358.49382 | 1498.82155 | Non-sig. |
| PF3D7_1307800 | CEP170 | 988.741062 | -0.493253295 | 0.55217178 | -0.89329681 | 0.37169824 | 0.89654416 | 1703.24732 | 912.177989 | 609.341338 | 730.197606 | Non-sig. |
| PF3D7_1011400 | N/A | 1889.47529 | -0.494394919 | 0.45543217 | -1.08555115 | 0.27767762 | 0.85972841 | 1658.23771 | 1449.47164 | 2761.7111 | 1688.48073 | Non-sig. |
| PF3D7_0109300 | N/A | 62.8477888 | -0.494451068 | 0.60580354 | -0.81619046 | 0.41439121 | 0.90793155 | 59.8925108 | 46.1614003 | 87.0300226 | 58.3072214 | Non-sig. |
| PF3D7_0615600 | N/A | 2179.36742 | -0.494811999 | 0.4592352 | -1.07746967 | 0.28127049 | 0.86386728 | 1878.44229 | 1667.82011 | 3220.27348 | 1950.93381 | Non-sig. |
| PF3D7_0523800 | N/A | 548.44062 | -0.495014693 | 0.53417808 | -0.92668477 | 0.35409023 | 0.8930363 | 390.887009 | 371.154604 | 892.071076 | 539.649789 | Non-sig. |
| PF3D7_1016900 | ETRAMP10.3 | 3266.41655 | -0.49518153 | 0.42880799 | -1.15478616 | 0.24817801 | 0.84804265 | 4497.13179 | 2814.74594 | 3146.16271 | 2607.62575 | Non-sig. |
| PF3D7_0507900 | N/A | 154.647718 | -0.495512969 | 0.54054722 | -0.91668767 | 0.35930633 | 0.89560738 | 234.013407 | 145.106013 | 128.077356 | 111.394094 | Non-sig. |
| PF3D7_1470900 | N/A | 4248.60846 | -0.495940425 | 0.42302677 | -1.17236179 | 0.24105183 | 0.84185407 | 4437.09733 | 3952.22867 | 5506.2944 | 3098.81344 | Non-sig. |
| PF3D7_0937300 | RIF | 1.94923396 | -0.495977746 | 2.32884182 | -0.21297185 | 0.83134892 | 0.99110542 | 2.09639464 | 1.07425265 | 2.46205926 | 2.16422929 | Non-sig. |

| | | | | | | | | | | | | |
|---|---|---|---|---|---|---|---|---|---|---|---|---|
| PF3D7_0413800 | N/A | 249.880031 | -0.496270181 | 0.46115223 | -1.07615263 | 0.281859 | 0.86390383 | 261.352371 | 211.547969 | 323.427272 | 203.192512 | Non-sig. |
| PF3D7_0815600 | EIF3G | 2222.29559 | -0.496361838 | 0.46015045 | -1.07869466 | 0.28072387 | 0.86335738 | 3250.71842 | 2070.36332 | 1951.24384 | 1616.85677 | Non-sig. |
| PF3D7_0205400 | N/A | 451.788759 | -0.4963708 | 0.46279371 | -1.07255303 | 0.28347171 | 0.86551387 | 413.464137 | 375.599192 | 643.836102 | 374.255604 | Non-sig. |
| PF3D7_0213100 | SIS1 | 935.853817 | -0.496555693 | 0.41841366 | -1.18675785 | 0.23532315 | 0.83828834 | 1021.24277 | 756.630475 | 1169.35269 | 796.189333 | Non-sig. |
| PF3D7_1445200 | MAK5 | 983.993026 | -0.497024725 | 0.51403045 | -0.9669169 | 0.33358558 | 0.89204385 | 1611.54426 | 817.465336 | 692.428848 | 814.533659 | Non-sig. |
| PF3D7_0728900 | N/A | 1141.86116 | -0.497069585 | 0.49916404 | -0.99580407 | 0.31934535 | 0.88651753 | 1821.99698 | 893.95243 | 851.589012 | 999.906216 | Non-sig. |
| PF3D7_1124700 | MGE1 | 672.186985 | -0.497193245 | 0.49937981 | -0.99562143 | 0.31943412 | 0.88651753 | 1056.72345 | 601.055977 | 517.316126 | 513.652384 | Non-sig. |
| PF3D7_0103200 | NT4 | 1031.20353 | -0.497416093 | 0.47023336 | -1.0578069 | 0.29014348 | 0.86724628 | 860.494141 | 873.565072 | 1553.71559 | 837.039321 | Non-sig. |
| PF3D7_0710700 | N/A | 381.163189 | -0.498133331 | 0.60443528 | -0.82413014 | 0.40986562 | 0.90748263 | 686.04197 | 370.379134 | 206.896526 | 261.335126 | Non-sig. |
| PF3D7_0608600 | N/A | 393.648597 | -0.499081409 | 0.49624039 | -1.00572509 | 0.31454783 | 0.88454055 | 580.82154 | 384.365253 | 341.526889 | 267.880705 | Non-sig. |
| PF3D7_1104200 | SNF2L | 4859.45414 | -0.499109912 | 0.44128758 | -1.13103096 | 0.25804206 | 0.85030738 | 5117.50677 | 3195.54986 | 6265.88733 | 4858.87261 | Non-sig. |
| PF3D7_1341900 | N/A | 1555.30403 | -0.499209594 | 0.41750023 | -1.19571096 | 0.23180937 | 0.83828834 | 1651.20936 | 1357.64831 | 1992.16384 | 1220.19461 | Non-sig. |
| PF3D7_0504000 | ATPase3 | 3204.68682 | -0.499300821 | 0.51806167 | -0.96378646 | 0.335153 | 0.8920875 | 2287.44645 | 2170.84605 | 5219.80261 | 3140.65216 | Non-sig. |
| PF3D7_0217400 | N/A | 787.793051 | -0.500372406 | 0.44801197 | -1.11687284 | 0.2640487 | 0.85144857 | 785.134286 | 747.109832 | 1060.83494 | 558.093144 | Non-sig. |
| PF3D7_1424400 | N/A | 5488.28323 | -0.501821272 | 0.40800132 | -1.22995012 | 0.21871578 | 0.83527883 | 6685.36549 | 4897.9153 | 6181.25981 | 4188.59234 | Non-sig. |
| PF3D7_0806600 | N/A | 271.08959 | -0.502059582 | 0.5329341 | -0.94206692 | 0.34615838 | 0.8930363 | 433.786918 | 201.706788 | 202.047158 | 246.817497 | Non-sig. |
| PF3D7_1010000 | N/A | 592.813519 | -0.502323116 | 0.49779344 | -1.0090995 | 0.31292692 | 0.88232401 | 460.551606 | 516.591861 | 929.157231 | 464.953378 | Non-sig. |
| PF3D7_1242600 | N/A | 378.9029 | -0.502577424 | 0.44210092 | -1.13679344 | 0.25562465 | 0.85030738 | 463.96018 | 340.186253 | 424.569947 | 286.89522 | Non-sig. |
| PF3D7_1003200 | N/A | 103.888866 | -0.50326968 | 0.55557342 | -0.90585629 | 0.36501192 | 0.89567376 | 153.587621 | 87.0931185 | 90.2339748 | 84.640749 | Non-sig. |
| PF3D7_1461200 | IMP3 | 225.144494 | -0.503828308 | 0.47769272 | -1.05471213 | 0.291557 | 0.86779157 | 245.674628 | 215.826991 | 282.396648 | 156.67971 | Non-sig. |
| PF3D7_0704000 | N/A | 1211.00627 | -0.503957502 | 0.53800534 | -0.93671468 | 0.34890534 | 0.8930363 | 809.68959 | 829.047437 | 2030.79852 | 1174.48955 | Non-sig. |
| PF3D7_0708600 | IMC1d | 45.8398193 | -0.504080327 | 0.68075711 | -0.74047016 | 0.45901476 | 0.92288312 | 69.9316583 | 33.3412827 | 37.7281001 | 42.3582363 | Non-sig. |
| PF3D7_1235800 | N/A | 156.755879 | -0.504570442 | 0.56976351 | -0.88557873 | 0.37584458 | 0.89654416 | 257.135491 | 137.297303 | 110.897033 | 121.693688 | Non-sig. |
| PF3D7_0604700 | GILP | 746.887396 | -0.504900262 | 0.42438011 | -1.18973591 | 0.23415021 | 0.83828834 | 924.231084 | 662.447987 | 828.360295 | 572.510218 | Non-sig. |
| PF3D7_0916600 | N/A | 75.5244673 | -0.50505437 | 0.60077779 | -0.84066752 | 0.40053422 | 0.90646318 | 71.2570313 | 77.3523916 | 105.863185 | 47.6252609 | Non-sig. |
| PF3D7_1220000 | N/A | 17.5955965 | -0.505084513 | 0.89693261 | -0.56312426 | 0.57335027 | 0.94850402 | 19.9374607 | 12.8870937 | 21.3419557 | 16.2158758 | Non-sig. |
| PF3D7_0213600 | N/A | 196.446674 | -0.505244361 | 0.54916293 | -0.92002635 | 0.35755899 | 0.89552149 | 318.313153 | 166.250319 | 142.938276 | 158.284947 | Non-sig. |
| PF3D7_1346400 | N/A | 27644.9113 | -0.505868684 | 0.5694669 | -0.88831973 | 0.37436879 | 0.89654416 | 17479.3698 | 15760.3789 | 47405.5523 | 29934.344 | Non-sig. |
| PF3D7_0109400 | N/A | 728.891734 | -0.506083802 | 0.42183863 | -1.19970947 | 0.2302522 | 0.83828834 | 897.29643 | 629.091242 | 813.642157 | 575.537106 | Non-sig. |
| PF3D7_1108200 | DTD | 575.971439 | -0.506323625 | 0.48719727 | -1.03925792 | 0.2986848 | 0.87330688 | 875.346732 | 426.321072 | 476.929571 | 525.28838 | Non-sig. |
| PF3D7_0212300 | ERF1 | 1882.34179 | -0.508883512 | 0.5697031 | -0.89324336 | 0.37172686 | 0.89654416 | 1044.30308 | 1595.48138 | 3377.22121 | 1512.36149 | Non-sig. |
| PF3D7_0512800 | N/A | 470.227908 | -0.509324822 | 0.45040105 | -1.13082513 | 0.2581287 | 0.85030738 | 498.430597 | 331.450428 | 606.23015 | 444.800456 | Non-sig. |
| PF3D7_0831600 | CLAG8 | 5106.54487 | -0.509538027 | 0.56495775 | -0.90190466 | 0.36710752 | 0.89587085 | 2989.52892 | 3454.29249 | 9008.2892 | 4974.06884 | Non-sig. |
| PF3D7_1461900 | N/A | 1385.5096 | -0.509925952 | 0.57135806 | -0.89248055 | 0.37213541 | 0.89654416 | 768.991717 | 1127.46632 | 2486.38495 | 1159.19543 | Non-sig. |
| PF3D7_1438600 | GET4 | 226.110312 | -0.509956303 | 0.64303301 | -0.7930484 | 0.42774963 | 0.91236574 | 107.296491 | 174.928699 | 423.739032 | 198.477027 | Non-sig. |
| PF3D7_0532600 | N/A | 305.612368 | -0.510182549 | 0.49888521 | -1.02264517 | 0.30647565 | 0.87685698 | 453.271466 | 287.819495 | 265.128276 | 216.230237 | Non-sig. |
| PF3D7_1253100 | N/A | 877.549328 | -0.510735041 | 0.4221878 | -1.20973423 | 0.22638089 | 0.83828834 | 1103.61728 | 769.649026 | 959.02333 | 677.90768 | Non-sig. |
| PF3D7_1456200 | N/A | 153.008772 | -0.510875419 | 0.60905645 | -0.83879814 | 0.40158259 | 0.90667178 | 91.615543 | 131.961695 | 267.761157 | 120.696691 | Non-sig. |
| PF3D7_1320900 | N/A | 253.926403 | -0.511522121 | 0.48920877 | -1.0456111 | 0.29574062 | 0.87191643 | 230.914092 | 226.869153 | 365.86022 | 192.062145 | Non-sig. |
| PF3D7_0930800 | N/A | 5104.54271 | -0.512225378 | 0.49064063 | -1.04399299 | 0.29648863 | 0.87299304 | 4539.51078 | 3031.82102 | 7462.92634 | 5383.91271 | Non-sig. |
| PF3D7_1445500 | N/A | 453.160853 | -0.513559017 | 0.44309299 | -1.15903213 | 0.24644309 | 0.84582594 | 523.068838 | 322.982429 | 542.860464 | 423.731681 | Non-sig. |
| PF3D7_1107500 | N/A | 263.976627 | -0.513834195 | 0.48165729 | -1.06680456 | 0.2860601 | 0.86595658 | 261.322761 | 251.288319 | 359.538871 | 183.756558 | Non-sig. |
| PF3D7_1334300 | MSRP5 | 656.984209 | -0.514167301 | 0.58049125 | -0.88574514 | 0.37575488 | 0.89654416 | 1156.73703 | 656.848477 | 389.226132 | 425.125195 | Non-sig. |
| PF3D7_0810400 | AQP2 | 266.157642 | -0.514435865 | 0.48147862 | -1.06845007 | 0.28531754 | 0.86551387 | 321.624166 | 170.675522 | 304.621041 | 267.709838 | Non-sig. |
| PF3D7_0936200 | N/A | 265.65337 | -0.514744198 | 0.55467624 | -0.92800838 | 0.35340322 | 0.8930363 | 178.884992 | 224.274801 | 445.936515 | 213.51717 | Non-sig. |
| PF3D7_0418400 | N/A | 162.786838 | -0.514984809 | 0.50877778 | -1.01219987 | 0.3114425 | 0.8804206 | 182.190329 | 163.592453 | 200.85918 | 104.505388 | Non-sig. |
| PF3D7_1106400 | N/A | 276.953349 | -0.515199361 | 0.45539479 | -1.13132467 | 0.25791846 | 0.85030738 | 315.302861 | 250.346352 | 336.442773 | 205.72141 | Non-sig. |
| PF3D7_1006800 | GBP2 | 5443.89332 | -0.515263954 | 0.43074766 | -1.19620836 | 0.23161526 | 0.83828834 | 5164.3588 | 4494.67664 | 7647.12487 | 4469.41298 | Non-sig. |
| PF3D7_0906200 | N/A | 483.318272 | -0.515513345 | 0.4354532 | -1.18385477 | 0.23647056 | 0.83828834 | 553.578261 | 436.726346 | 583.921882 | 359.0466 | Non-sig. |
| PF3D7_1233300 | PPR2 | 1174.09669 | -0.515532499 | 0.4199637 | -1.22756444 | 0.21961052 | 0.83527883 | 1526.42023 | 956.893859 | 1237.0354 | 976.037271 | Non-sig. |
| PF3D7_0922800 | N/A | 2215.67264 | -0.516161469 | 0.44203269 | -1.16769978 | 0.24292789 | 0.8422984 | 2580.19401 | 1419.94221 | 2635.50198 | 2227.05235 | Non-sig. |
| PF3D7_1440600 | SOC3 | 115.238623 | -0.516513128 | 0.59303281 | -0.87096888 | 0.38377114 | 0.90292925 | 190.242171 | 99.829016 | 81.2870482 | 89.5962582 | Non-sig. |
| PF3D7_1306600 | N/A | 768.619826 | -0.516513753 | 0.42949157 | -1.20261674 | 0.22912467 | 0.83828834 | 853.674127 | 561.364854 | 955.780283 | 703.660041 | Non-sig. |
| PF3D7_0513500 | PAM16 | 114.743575 | -0.516749765 | 0.56749645 | -0.91057797 | 0.36251778 | 0.89567376 | 150.466945 | 124.722618 | 119.766515 | 64.0182229 | Non-sig. |

| | | | | | | | | | | | | |
|---|---|---|---|---|---|---|---|---|---|---|---|---|
| PF3D7_0909800 | SNRPD3 | 476.959102 | -0.517173928 | 0.43099022 | -1.19996673 | 0.23015226 | 0.83828834 | 594.589263 | 395.276249 | 528.563163 | 389.407733 | Non-sig. |
| PF3D7_1353800 | N/A | 2233.189 | -0.517433916 | 0.41338481 | -1.25170037 | 0.21067906 | 0.82618629 | 2370.72709 | 1847.93577 | 2888.01371 | 1826.07942 | Non-sig. |
| PF3D7_0528600 | N/A | 280.574719 | -0.517646861 | 0.45824292 | -1.12963416 | 0.25863041 | 0.85030738 | 322.085812 | 259.298651 | 338.654496 | 202.259915 | Non-sig. |
| PF3D7_1339800 | MZT1 | 288.125434 | -0.517838014 | 0.46332388 | -1.11765882 | 0.26371274 | 0.85144857 | 292.67549 | 255.616163 | 385.775868 | 218.434214 | Non-sig. |
| PF3D7_0216200 | N/A | 230.191552 | -0.518192312 | 0.49763714 | -1.04130555 | 0.29773376 | 0.87301237 | 339.672529 | 192.282243 | 202.715523 | 186.095911 | Non-sig. |
| PF3D7_0807500 | N/A | 4328.41638 | -0.518537817 | 0.41081853 | -1.26220648 | 0.2068745 | 0.82391338 | 5193.31855 | 3913.90439 | 5002.74133 | 3203.70125 | Non-sig. |
| PF3D7_0303300 | RPB6 | 892.249995 | -0.518557071 | 0.43288729 | -1.19790319 | 0.23095471 | 0.83828834 | 1047.43144 | 846.806035 | 1054.36974 | 620.392771 | Non-sig. |
| PF3D7_1221200 | N/A | 204.124825 | -0.519775755 | 0.47524281 | -1.09370567 | 0.27408409 | 0.85618956 | 223.575403 | 186.254464 | 257.379834 | 149.289598 | Non-sig. |
| PF3D7_0212000 | N/A | 450.762996 | -0.520531742 | 0.43142863 | -1.20653037 | 0.22761306 | 0.83828834 | 551.759319 | 377.85671 | 510.704752 | 362.731201 | Non-sig. |
| PF3D7_0814900 | FeSOD | 2897.07112 | -0.520853112 | 0.45166252 | -1.1531909 | 0.24883204 | 0.8482655 | 2733.91766 | 2769.24284 | 4094.75139 | 1990.37259 | Non-sig. |
| PF3D7_1212000 | TPx(Gl) | 1111.77815 | -0.521213738 | 0.50977115 | -1.02244652 | 0.30656961 | 0.87685698 | 811.384606 | 1005.10822 | 1809.23742 | 821.382369 | Non-sig. |
| PF3D7_0710800 | USE1 | 566.988856 | -0.521985884 | 0.53102053 | -0.98298626 | 0.3256142 | 0.88694295 | 936.508532 | 529.199137 | 400.688614 | 401.559143 | Non-sig. |
| PF3D7_0508300 | TPT | 947.234576 | -0.522833732 | 0.69460205 | -0.75270975 | 0.45162435 | 0.92081798 | 311.738718 | 666.224421 | 1922.00452 | 888.970642 | Non-sig. |
| PF3D7_1341700 | N/A | 11.2782325 | -0.522863394 | 1.17921959 | -0.44339782 | 0.65747801 | 0.97654009 | 21.7747702 | 10.774874 | 4.9154804 | 7.64780553 | Non-sig. |
| PF3D7_1348500 | N/A | 1109.9053 | -0.523214743 | 0.42625846 | -1.22745891 | 0.21965016 | 0.83527883 | 1135.66697 | 938.057249 | 1482.17088 | 883.726101 | Non-sig. |
| PF3D7_1467700 | N/A | 155.568691 | -0.52343986 | 0.54549119 | -0.95957527 | 0.33726902 | 0.8920875 | 128.81712 | 148.282693 | 237.962682 | 107.212268 | Non-sig. |
| PF3D7_0423100 | N/A | 1928.25014 | -0.523534811 | 0.61412326 | -0.85249143 | 0.39394139 | 0.90601068 | 3646.10728 | 1763.57371 | 902.867768 | 1400.4518 | Non-sig. |
| PF3D7_0524300 | N/A | 661.482074 | -0.523743488 | 0.47851881 | -1.09450972 | 0.27373149 | 0.85618956 | 979.318229 | 624.596567 | 581.406746 | 460.606752 | Non-sig. |
| PF3D7_0323600 | N/A | 785.146879 | -0.523917033 | 0.42761263 | -1.22521413 | 0.22049455 | 0.83527883 | 944.030723 | 718.935074 | 908.323075 | 569.298642 | Non-sig. |
| PF3D7_0418100 | SOC1 | 825.167843 | -0.524703286 | 0.44724255 | -1.17319626 | 0.24071711 | 0.84181962 | 835.61904 | 573.873613 | 1111.41819 | 779.76053 | Non-sig. |
| PF3D7_1109200 | N/A | 63.1747478 | -0.525027393 | 0.59940581 | -0.87591308 | 0.38107729 | 0.90292925 | 85.9758259 | 46.1893024 | 63.1959298 | 57.3379333 | Non-sig. |
| PF3D7_1310200 | N/A | 1792.33535 | -0.525131512 | 0.43482873 | -1.20767436 | 0.22717254 | 0.83828834 | 1970.02938 | 1733.21087 | 2259.86937 | 1206.23178 | Non-sig. |
| PF3D7_0506100 | N/A | 766.290765 | -0.525592343 | 0.42643553 | -1.23252473 | 0.21775313 | 0.83527883 | 982.651489 | 668.563382 | 826.149281 | 587.798907 | Non-sig. |
| PF3D7_1437500 | N/A | 471.871885 | -0.52565935 | 0.56611501 | -0.92853808 | 0.35312852 | 0.8930363 | 295.668918 | 325.79706 | 817.84204 | 448.179523 | Non-sig. |
| PF3D7_0707900 | N/A | 1031.14873 | -0.525848169 | 0.41969742 | -1.25292208 | 0.21023406 | 0.82566351 | 1305.88314 | 894.078035 | 1128.23611 | 796.397619 | Non-sig. |
| PF3D7_1427600 | MIT3 | 667.122816 | -0.526048192 | 0.43530514 | -1.20845849 | 0.22687095 | 0.83828834 | 716.554398 | 607.375226 | 858.194022 | 486.367616 | Non-sig. |
| PF3D7_1213200 | MED18 | 297.525278 | -0.526085586 | 0.53378324 | -0.98557906 | 0.32433972 | 0.88694295 | 220.295697 | 271.725624 | 481.81033 | 216.26946 | Non-sig. |
| PF3D7_0822300 | SNRPG | 182.099031 | -0.52704538 | 0.51009532 | -1.0332292 | 0.30149665 | 0.87381251 | 161.437627 | 155.762513 | 268.368943 | 142.827042 | Non-sig. |
| PF3D7_1334700 | MSRP7 | 339.643903 | -0.527561169 | 0.44966284 | -1.1732372 | 0.2407007 | 0.84181962 | 356.193418 | 289.852804 | 445.821537 | 266.707854 | Non-sig. |
| PF3D7_0604000 | N/A | 312.395385 | -0.528093892 | 0.53919441 | -0.97941276 | 0.32737607 | 0.88852997 | 271.919957 | 173.761358 | 465.77672 | 338.123508 | Non-sig. |
| PF3D7_1404000 | RPB4 | 130.374732 | -0.528717358 | 0.51143834 | -1.03378513 | 0.30123662 | 0.87381251 | 174.519074 | 115.230153 | 133.584065 | 98.165635 | Non-sig. |
| PF3D7_1147700 | N/A | 1580.84666 | -0.529615882 | 0.43735429 | -1.21095389 | 0.22591308 | 0.83828834 | 2139.73532 | 1467.62521 | 1596.01467 | 1120.01145 | Non-sig. |
| PF3D7_0808450 | MPODD | 18.6137132 | -0.530342178 | 0.94513669 | -0.56112749 | 0.57471063 | 0.94883668 | 15.4658951 | 21.7464196 | 28.4698909 | 8.77264713 | Non-sig. |
| PF3D7_1403200 | N/A | 42.131423 | -0.530542178 | 0.66819815 | -0.7939893 | 0.42720166 | 0.91236574 | 50.3459778 | 40.8142095 | 49.2468093 | 28.1186952 | Non-sig. |
| PF3D7_1346500 | N/A | 462.961202 | -0.530607531 | 0.49669995 | -1.06826572 | 0.28540067 | 0.86551387 | 718.446224 | 412.10511 | 376.107664 | 345.18581 | Non-sig. |
| PF3D7_1133900 | N/A | 2477.56078 | -0.530727013 | 0.41274029 | -1.28586191 | 0.19849126 | 0.81528784 | 2652.76978 | 1968.44939 | 3203.52711 | 2085.49684 | Non-sig. |
| PF3D7_1343600 | N/A | 237.969893 | -0.530743555 | 0.46619193 | -1.13846578 | 0.25492604 | 0.85030738 | 288.252614 | 218.923876 | 274.28482 | 170.41826 | Non-sig. |
| PF3D7_0507200 | SUB3 | 1301.23737 | -0.531156484 | 0.46364912 | -1.14560012 | 0.25196065 | 0.85030738 | 1684.19759 | 786.495992 | 1392.09475 | 1342.16113 | Non-sig. |
| PF3D7_0319300 | N/A | 336.509975 | -0.531157153 | 0.45657648 | -1.16334759 | 0.2446885 | 0.84331907 | 431.177016 | 311.45871 | 364.439762 | 238.964411 | Non-sig. |
| PF3D7_1222300 | GRP94 | 14194.9866 | -0.531679365 | 0.45319558 | -1.17317862 | 0.24072418 | 0.84181962 | 12332.2881 | 11004.3196 | 21230.3367 | 12213.0021 | Non-sig. |
| PF3D7_0830100 | N/A | 696.587095 | -0.531908168 | 0.43904282 | -1.21151775 | 0.22569704 | 0.83828834 | 730.458524 | 502.167721 | 916.551405 | 637.170729 | Non-sig. |
| PF3D7_0420500 | N/A | 122.735225 | -0.533122505 | 0.56474996 | -0.94399742 | 0.34517096 | 0.8930363 | 97.2990061 | 91.3599544 | 192.813232 | 109.468709 | Non-sig. |
| PF3D7_0820000 | SRCAP | 2823.0999 | -0.53335922 | 0.47675088 | -1.11873777 | 0.26325203 | 0.85095134 | 2323.17108 | 2029.29757 | 4354.72682 | 2585.20415 | Non-sig. |
| PF3D7_0501600 | RAP2 | 8966.97452 | -0.533537595 | 0.42262377 | -1.26244104 | 0.20679013 | 0.82391338 | 8963.25474 | 6877.7374 | 12249.4118 | 7777.49415 | Non-sig. |
| PF3D7_0902600 | FIKK9.7 | 146.155039 | -0.533591083 | 0.5031785 | -1.06044094 | 0.28894405 | 0.86724628 | 146.110666 | 114.854149 | 199.525892 | 124.129447 | Non-sig. |
| PF3D7_0522400 | N/A | 23064.8786 | -0.533891047 | 0.47175781 | -1.1317058 | 0.25775814 | 0.85030738 | 22414.2218 | 13724.4538 | 32154.8095 | 23966.0292 | Non-sig. |
| PF3D7_1332800 | N/A | 394.156565 | -0.533921112 | 0.54980527 | -0.97110949 | 0.33149376 | 0.88994703 | 258.839427 | 324.077947 | 673.413579 | 320.295305 | Non-sig. |
| PF3D7_1303000 | ATPTG13 | 345.900316 | -0.534038808 | 0.56000948 | -0.95362458 | 0.34027371 | 0.8930363 | 224.893247 | 311.900897 | 593.25232 | 253.554802 | Non-sig. |
| PF3D7_1317900 | NOC4 | 261.991566 | -0.53403934 | 0.45813261 | -1.16568725 | 0.24374092 | 0.84313671 | 339.059977 | 204.920013 | 280.899507 | 223.086767 | Non-sig. |
| PF3D7_1316900 | N/A | 3520.89443 | -0.534562173 | 0.43711354 | -1.22293666 | 0.22135362 | 0.83527883 | 4257.12831 | 2269.592 | 4074.55457 | 3482.30285 | Non-sig. |
| PF3D7_0626900 | N/A | 543.504172 | -0.534584125 | 0.45025162 | -1.18730083 | 0.23510898 | 0.83828834 | 741.072763 | 493.764027 | 545.207568 | 393.972328 | Non-sig. |
| PF3D7_1463800 | N/A | 64.3262393 | -0.535368017 | 0.61925848 | -0.86453077 | 0.38729638 | 0.90421052 | 94.9622528 | 44.0988276 | 57.4105224 | 60.8333544 | Non-sig. |
| PF3D7_0706100 | N/A | 1230.74745 | -0.535832238 | 0.47764676 | -1.12181696 | 0.26194027 | 0.85030738 | 1123.28888 | 783.042949 | 1789.93302 | 1226.72494 | Non-sig. |
| PF3D7_0618900 | GPI1 | 1501.77081 | -0.535894979 | 0.42487227 | -1.26130844 | 0.20719774 | 0.82391338 | 2013.65644 | 1295.89492 | 1541.6715 | 1155.8604 | Non-sig. |

| | | | | | | | | | | | | |
|---|---|---|---|---|---|---|---|---|---|---|---|---|
| PF3D7_0802200 | 1-CysPxn | 2694.57189 | -0.536564633 | 0.41871107 | -1.28146751 | 0.20002951 | 0.81595849 | 2903.97989 | 2418.21667 | 3475.83611 | 1980.2549 | Non-sig. |
| PF3D7_1439100 | N/A | 1002.76916 | -0.537498765 | 0.58801076 | -0.91409682 | 0.36066597 | 0.89560738 | 612.467548 | 565.22934 | 1762.12578 | 1071.25396 | Non-sig. |
| PF3D7_0320600 | N/A | 235.151762 | -0.537929622 | 0.46538572 | -1.1558791 | 0.24773061 | 0.84730586 | 250.727938 | 180.432296 | 306.161118 | 203.285697 | Non-sig. |
| PF3D7_0509300 | N/A | 205.324713 | -0.53795857 | 0.47047238 | -1.14344347 | 0.25285451 | 0.85030738 | 261.467268 | 178.236705 | 224.93892 | 156.655959 | Non-sig. |
| PF3D7_1308400 | N/A | 11473.6674 | -0.538097234 | 0.50574962 | -1.06395973 | 0.28734694 | 0.86724628 | 8969.31808 | 7207.71788 | 18208.2885 | 11509.345 | Non-sig. |
| PF3D7_0628200 | PK4 | 14258.3286 | -0.538261305 | 0.44595396 | -1.20698851 | 0.22743657 | 0.83828834 | 14141.6871 | 9412.20637 | 19633.6581 | 13845.7627 | Non-sig. |
| PF3D7_1338200 | RPL6 | 5421.59193 | -0.538879452 | 0.42003114 | -1.28295118 | 0.19950918 | 0.81595849 | 6206.62676 | 5074.29269 | 6638.39647 | 3767.05179 | Non-sig. |
| PF3D7_1354300 | N/A | 1513.24338 | -0.539485576 | 0.5619523 | -0.96002023 | 0.33704503 | 0.8920875 | 2699.87733 | 1329.14732 | 886.291947 | 1137.65694 | Non-sig. |
| PF3D7_1441100 | N/A | 8705.70296 | -0.539988484 | 0.50094073 | -1.07794886 | 0.28105658 | 0.863711 | 14148.4287 | 8034.76683 | 6484.23716 | 6155.37914 | Non-sig. |
| PF3D7_0815000 | Sel3 | 413.090396 | -0.540079264 | 0.53222726 | -1.0147531 | 0.31022354 | 0.88032279 | 292.298592 | 358.490056 | 686.466586 | 315.106347 | Non-sig. |
| PF3D7_1430100 | PTPA | 353.777147 | -0.540268701 | 0.46658822 | -1.1579133 | 0.24689942 | 0.84582594 | 348.991143 | 323.092459 | 489.373965 | 253.651021 | Non-sig. |
| PF3D7_0419000 | N/A | 117.059402 | -0.540718133 | 0.591624 | -0.91395571 | 0.36074012 | 0.89560738 | 185.746123 | 118.129384 | 91.9357081 | 72.4263927 | Non-sig. |
| PF3D7_0602200 | N/A | 386.388223 | -0.541045238 | 0.45946314 | -1.17755961 | 0.23897222 | 0.84066259 | 392.897541 | 354.28252 | 522.982134 | 275.390698 | Non-sig. |
| PF3D7_1129500 | N/A | 695.329715 | -0.541566541 | 0.41968474 | -1.29041274 | 0.19690739 | 0.81437577 | 813.427688 | 573.02211 | 835.211932 | 559.65713 | Non-sig. |
| PF3D7_0816900 | AK2 | 18.5710016 | -0.541684213 | 0.91792895 | -0.59011562 | 0.55511314 | 0.94243456 | 29.3432987 | 18.263381 | 14.7718006 | 11.9055261 | Non-sig. |
| PF3D7_MIT02100 | COX1 | 5916.09968 | -0.542013571 | 0.53045181 | -1.02179606 | 0.30687743 | 0.87685698 | 4065.49976 | 3886.66575 | 9963.28196 | 5748.95124 | Non-sig. |
| PF3D7_0610100 | SLU7 | 2305.65153 | -0.542290949 | 0.59027971 | -0.91870166 | 0.35825164 | 0.89552149 | 4267.68356 | 2091.77454 | 1200.54299 | 1662.60504 | Non-sig. |
| PF3D7_0511900 | N/A | 86.5581586 | -0.543394604 | 0.56921994 | -0.9546303 | 0.33976469 | 0.8930363 | 89.4567289 | 55.7979315 | 115.798892 | 85.1790824 | Non-sig. |
| PF3D7_1349900 | N/A | 111.803654 | -0.544216213 | 0.51305253 | -1.06074169 | 0.28880731 | 0.86724628 | 134.149902 | 94.7411648 | 131.149852 | 87.1736979 | Non-sig. |
| PF3D7_1330100 | N/A | 431.396701 | -0.544761291 | 0.49330545 | -1.10430829 | 0.26945943 | 0.8556331 | 629.013361 | 426.55257 | 394.957858 | 275.063014 | Non-sig. |
| PF3D7_0724200 | TAP42 | 313.803169 | -0.545083688 | 0.45911377 | -1.18725188 | 0.23512828 | 0.83828834 | 422.256608 | 274.847533 | 322.648681 | 235.459854 | Non-sig. |
| PF3D7_0529100 | ATrx2 | 127.914383 | -0.545249329 | 0.55134983 | -0.98893534 | 0.32269478 | 0.88694295 | 109.812912 | 119.317942 | 193.622129 | 88.9045483 | Non-sig. |
| PF3D7_0528900 | N/A | 42.4603713 | -0.54562447 | 0.6716845 | -0.81232256 | 0.41660657 | 0.90793155 | 59.8271404 | 31.1416539 | 41.0440083 | 37.8286826 | Non-sig. |
| PF3D7_0509500 | N/A | 225.252079 | -0.545852844 | 0.51373806 | -1.06251199 | 0.28800332 | 0.86724628 | 347.758396 | 186.798446 | 187.193014 | 179.258458 | Non-sig. |
| PF3D7_0827400 | N/A | 60.0352215 | -0.545918502 | 0.63453042 | -0.8603504 | 0.38959592 | 0.90444842 | 88.5170624 | 38.7426582 | 54.1113799 | 58.7697854 | Non-sig. |
| PF3D7_0203200 | N/A | 480.092326 | -0.546364316 | 0.46756306 | -1.16853611 | 0.24259058 | 0.8422984 | 706.52013 | 388.549819 | 433.553177 | 391.746178 | Non-sig. |
| PF3D7_0504500 | N/A | 39.8149048 | -0.546929309 | 0.69580893 | -0.78603376 | 0.43184771 | 0.91370017 | 57.7021478 | 37.5894678 | 36.9366952 | 27.0313082 | Non-sig. |
| PF3D7_1034200 | RPL27 | 131.89504 | -0.547621365 | 0.50010666 | -1.09500913 | 0.27351264 | 0.85618956 | 166.564577 | 111.933265 | 146.748853 | 102.333464 | Non-sig. |
| PF3D7_1141600 | DPM1 | 202.099283 | -0.547902257 | 0.52101719 | -1.05160111 | 0.2929826 | 0.87056747 | 168.133623 | 152.701702 | 311.694376 | 175.86743 | Non-sig. |
| PF3D7_0711000 | CDC48 | 2420.48527 | -0.548346889 | 0.45296058 | -1.21058413 | 0.22605483 | 0.83828834 | 2695.09213 | 1473.89547 | 3054.88644 | 2458.06702 | Non-sig. |
| PF3D7_0416700 | N/A | 911.793095 | -0.548703517 | 0.41624311 | -1.31822847 | 0.18742718 | 0.80209142 | 1109.75228 | 767.739501 | 1056.5275 | 713.153099 | Non-sig. |
| PF3D7_1336700 | PV3 | 1199.09322 | -0.548842576 | 0.4349519 | -1.26184659 | 0.20700399 | 0.82391338 | 1387.21556 | 801.711839 | 1461.68461 | 1145.76089 | Non-sig. |
| PF3D7_1111100 | RFC5 | 2283.73212 | -0.548986889 | 0.44441307 | -1.2353077 | 0.21671601 | 0.83527883 | 2119.75715 | 1986.42719 | 3306.18416 | 1722.55999 | Non-sig. |
| PF3D7_1145900 | N/A | 107.581818 | -0.549315418 | 0.61989546 | -0.88614203 | 0.375541 | 0.89654416 | 183.630167 | 100.94957 | 72.2340106 | 73.5135265 | Non-sig. |
| PF3D7_1342300 | N/A | 647.065586 | -0.549484259 | 0.43389821 | -1.26638978 | 0.20537357 | 0.82391338 | 766.745984 | 594.795605 | 770.90189 | 455.818867 | Non-sig. |
| PF3D7_1342000 | N/A | 8485.07114 | -0.549543621 | 0.5541195 | -0.99174208 | 0.32132336 | 0.88694295 | 15102.7846 | 7577.25507 | 5061.255 | 6198.98986 | Non-sig. |
| PF3D7_0722500 | CWC15 | 1236.48566 | -0.549917225 | 0.59464152 | -0.92478781 | 0.35507629 | 0.89367896 | 2276.52142 | 1186.61299 | 662.447282 | 820.360951 | Non-sig. |
| PF3D7_1133000 | N/A | 818.053293 | -0.550788334 | 0.4448037 | -1.23827282 | 0.21561492 | 0.83503497 | 783.396312 | 684.459261 | 1161.09921 | 643.258389 | Non-sig. |
| PF3D7_0724100 | N/A | 4350.76983 | -0.551426966 | 0.64520075 | -0.85465952 | 0.39273967 | 0.90601068 | 8586.1674 | 3936.76809 | 1758.69037 | 3121.45344 | Non-sig. |
| PF3D7_1022400 | SRSF4 | 5229.32442 | -0.552606461 | 0.42028532 | -1.31483645 | 0.18856487 | 0.80500943 | 7156.29963 | 4326.70531 | 5281.41262 | 4152.88013 | Non-sig. |
| PF3D7_1338300 | EF-1gamma | 5879.4704 | -0.552767806 | 0.43725567 | -1.26417529 | 0.20616712 | 0.82391338 | 5805.08926 | 5420.74032 | 8179.24346 | 4112.80854 | Non-sig. |
| PF3D7_0630500 | YTM1 | 344.18473 | -0.553236894 | 0.48173952 | -1.14841501 | 0.25079728 | 0.8494288 | 305.24304 | 282.198956 | 513.304051 | 275.992871 | Non-sig. |
| PF3D7_1124300 | N/A | 3178.80736 | -0.55328662 | 0.40861161 | -1.35406484 | 0.17571569 | 0.78744529 | 4072.29055 | 2603.99426 | 3489.79299 | 2549.15166 | Non-sig. |
| PF3D7_1104400 | Trx-mero | 2708.60284 | -0.553399785 | 0.45677712 | -1.2115313 | 0.22569184 | 0.83828834 | 2666.74633 | 1732.09545 | 3776.72531 | 2658.84428 | Non-sig. |
| PF3D7_1448600 | VTI1 | 447.262463 | -0.553572802 | 0.4352711 | -1.27178855 | 0.20344826 | 0.82152939 | 540.392209 | 391.986078 | 523.695698 | 332.975869 | Non-sig. |
| PF3D7_0714500 | N/A | 1480.30326 | -0.554839911 | 0.42269754 | -1.31261685 | 0.18931209 | 0.80547152 | 1543.46683 | 1140.51343 | 1979.39243 | 1257.84035 | Non-sig. |
| PF3D7_0921000 | N/A | 770.413943 | -0.555034879 | 0.46108308 | -1.20376327 | 0.22868109 | 0.83828834 | 840.093101 | 771.794894 | 993.371242 | 476.396535 | Non-sig. |
| PF3D7_1026800 | RPS2 | 8441.18758 | -0.555459191 | 0.44230225 | -1.25583623 | 0.20917535 | 0.82566351 | 7867.13161 | 7476.86892 | 12225.4612 | 6195.28859 | Non-sig. |
| PF3D7_0902200 | FIKK9.3 | 504.841106 | -0.556944686 | 0.48105868 | -1.15774792 | 0.24696692 | 0.84582594 | 434.833009 | 378.738346 | 767.120046 | 438.673024 | Non-sig. |
| PF3D7_1214200 | SET5 | 59.1532766 | -0.55696189 | 0.63856186 | -0.8722129 | 0.38309224 | 0.90292925 | 66.1380384 | 63.3535808 | 74.7027436 | 32.4187435 | Non-sig. |
| PF3D7_1472800 | N/A | 1346.1411 | -0.557169601 | 0.4636751 | -1.20163796 | 0.22950383 | 0.83828834 | 2023.74172 | 1188.3868 | 1182.29519 | 990.140676 | Non-sig. |
| PF3D7_1041000 | RIF | 2.00413659 | -0.557603088 | 2.31387101 | -0.24098279 | 0.80956846 | 0.99028237 | 3.14452718 | 2.14851771 | 1.64136685 | 1.08213463 | Non-sig. |
| PF3D7_0720200 | N/A | 326.280724 | -0.558493981 | 0.51238888 | -1.08998067 | 0.27572166 | 0.85877328 | 512.546987 | 296.668076 | 265.007233 | 230.900599 | Non-sig. |
| PF3D7_1234000 | N/A | 459.195408 | -0.558786394 | 0.43138703 | -1.29532497 | 0.19520815 | 0.81344649 | 561.212766 | 387.518844 | 532.872727 | 355.177293 | Non-sig. |

| | | | | | | | | | | | | |
|---|---|---|---|---|---|---|---|---|---|---|---|---|
| PF3D7_1106500 | N/A | 1401.02395 | -0.558916683 | 0.42329203 | -1.32040446 | 0.18670002 | 0.80156944 | 1529.17262 | 1014.70176 | 1808.86608 | 1251.35534 | Non-sig. |
| PF3D7_0729200 | AOP | 204.463958 | -0.559116673 | 0.48290942 | -1.15780858 | 0.24694216 | 0.84582594 | 204.672671 | 155.708083 | 282.38299 | 175.092089 | Non-sig. |
| PF3D7_1137400 | N/A | 616.656166 | -0.559528038 | 0.46706002 | -1.19797889 | 0.23092524 | 0.83828834 | 547.065836 | 494.72069 | 922.226157 | 502.611982 | Non-sig. |
| PF3D7_1311700 | N/A | 365.769524 | -0.559632226 | 0.45226063 | -1.23741088 | 0.21593458 | 0.83527883 | 424.069771 | 339.093314 | 447.58032 | 252.334692 | Non-sig. |
| PF3D7_1429500 | DPH2 | 84.4842858 | -0.561082238 | 0.57213888 | -0.98067489 | 0.32675309 | 0.88852997 | 78.9195159 | 61.1659274 | 122.365546 | 75.4861536 | Non-sig. |
| PF3D7_0626600 | N/A | 106.671042 | -0.562515098 | 0.59471691 | -0.94585355 | 0.34422329 | 0.8930363 | 78.5911368 | 94.5422088 | 175.617531 | 77.933292 | Non-sig. |
| PF3D7_0720500 | N/A | 339.266779 | -0.562880241 | 0.53201893 | -1.05800791 | 0.29005183 | 0.86724628 | 245.784271 | 280.217855 | 563.195685 | 267.869306 | Non-sig. |
| PF3D7_1400400 | RIF | 62.1670492 | -0.563753934 | 0.59612432 | -0.9456986 | 0.34430234 | 0.8930363 | 82.7426964 | 54.7989204 | 65.6457927 | 45.4807875 | Non-sig. |
| PF3D7_1318100 | FD | 863.889727 | -0.564636089 | 0.45411788 | -1.24336899 | 0.21373189 | 0.83322411 | 1140.87187 | 842.12442 | 920.872027 | 551.690591 | Non-sig. |
| PF3D7_1207800 | N/A | 296.725692 | -0.564898939 | 0.58151895 | -0.97141966 | 0.33133934 | 0.88994703 | 528.587354 | 254.27373 | 179.873696 | 224.167987 | Non-sig. |
| PF3D7_1451700 | CNB | 3391.57125 | -0.56652359 | 0.41794497 | -1.35549806 | 0.17525893 | 0.78744529 | 4269.60609 | 3072.33083 | 3828.56705 | 2395.78104 | Non-sig. |
| PF3D7_0111200 | N/A | 110.13557 | -0.567337468 | 0.51809836 | -1.09503814 | 0.27349993 | 0.85618956 | 131.060437 | 96.8878012 | 131.974701 | 80.6193421 | Non-sig. |
| PF3D7_0316800 | N/A | 6200.58228 | -0.567936911 | 0.42777933 | -1.32763991 | 0.18429709 | 0.80052251 | 6340.32876 | 5575.05695 | 8470.59714 | 4416.34628 | Non-sig. |
| PF3D7_0407500 | N/A | 131.012021 | -0.568288014 | 0.5702694 | -0.99652552 | 0.31899487 | 0.88651753 | 98.5482451 | 106.345659 | 214.205963 | 104.948219 | Non-sig. |
| PF3D7_0923200 | NOS | 518.424058 | -0.568391759 | 0.55212588 | -1.02946046 | 0.30326335 | 0.87510307 | 333.866603 | 394.7959 | 904.333103 | 440.700625 | Non-sig. |
| PF3D7_1361600 | YfhJ | 360.249127 | -0.568947366 | 0.45371504 | -1.2539751 | 0.20985106 | 0.82566351 | 416.8217 | 333.831464 | 443.923562 | 246.41978 | Non-sig. |
| PF3D7_1443800 | CWC24 | 1052.65523 | -0.56912951 | 0.4682909 | -1.21533328 | 0.224239 | 0.83828834 | 1584.25788 | 946.537169 | 931.246271 | 748.579598 | Non-sig. |
| PF3D7_1122000 | N/A | 159.716256 | -0.569615037 | 0.5157663 | -1.10440531 | 0.26941736 | 0.8556331 | 152.442992 | 110.555361 | 229.091978 | 146.774695 | Non-sig. |
| PF3D7_0603300 | DHODH | 686.029617 | -0.570232675 | 0.43765742 | -1.30292017 | 0.19260202 | 0.81094926 | 712.416378 | 598.773676 | 927.194162 | 505.734254 | Non-sig. |
| PF3D7_1038000 | N/A | 20547.4313 | -0.57116657 | 0.43517097 | -1.31251074 | 0.18934786 | 0.80547152 | 29756.8489 | 17205.6243 | 19368.5005 | 15858.7516 | Non-sig. |
| PF3D7_0919000 | NAPS | 8080.35917 | -0.571228596 | 0.41281943 | -1.38372507 | 0.16644266 | 0.77284241 | 8658.83903 | 6908.92031 | 10660.0001 | 6093.67724 | Non-sig. |
| PF3D7_0520600 | Ap4AH | 598.891065 | -0.571465474 | 0.44677569 | -1.27908812 | 0.20086603 | 0.81683598 | 618.313595 | 541.195947 | 813.515391 | 422.539329 | Non-sig. |
| PF3D7_0721400 | N/A | 1274.90708 | -0.571508581 | 0.44473667 | -1.28504938 | 0.19877503 | 0.81528784 | 1241.91213 | 1142.24421 | 1806.26575 | 909.206214 | Non-sig. |
| PF3D7_API02400 | ORF91 | 2.05438956 | -0.572267338 | 2.34718141 | -0.24381044 | 0.80737763 | 0.99028237 | 4.12266776 | 1.08714236 | 0.81344624 | 2.19430188 | Non-sig. |
| PF3D7_1248300 | N/A | 233.641275 | -0.572296353 | 0.51873788 | -1.10324767 | 0.26991962 | 0.8556331 | 189.966732 | 183.631269 | 368.571542 | 192.395558 | Non-sig. |
| PF3D7_1104800 | N/A | 632.595601 | -0.572407984 | 0.56637655 | -1.01064916 | 0.31218438 | 0.88116827 | 377.706399 | 504.37129 | 1134.93205 | 513.372667 | Non-sig. |
| PF3D7_0402600 | N/A | 2.68443566 | -0.572888737 | 2.00842002 | -0.28524349 | 0.7754576 | 0.99028237 | 3.1381185 | 2.14975605 | 3.28183394 | 2.16803414 | Non-sig. |
| PF3D7_0716800 | EIF3I | 1901.34621 | -0.573752822 | 0.4463923 | -1.28531074 | 0.19868372 | 0.81528784 | 1746.321 | 1529.51322 | 2802.49337 | 1527.05725 | Non-sig. |
| PF3D7_1444600 | N/A | 33.8806481 | -0.57407321 | 0.74241133 | -0.77325491 | 0.43937153 | 0.91706217 | 28.7465091 | 31.3856242 | 52.2326608 | 23.1577982 | Non-sig. |
| PF3D7_0702900 | N/A | 555.572848 | -0.574257253 | 0.43357963 | -1.32445626 | 0.18535156 | 0.80052251 | 691.643804 | 492.905592 | 637.817076 | 399.924919 | Non-sig. |
| PF3D7_0301900 | N/A | 177.388337 | -0.574791196 | 0.48586945 | -1.18301572 | 0.23680292 | 0.83828834 | 194.542463 | 158.845635 | 229.922677 | 126.242572 | Non-sig. |
| PF3D7_1247400 | FKBP35 | 2900.44606 | -0.575222615 | 0.42450529 | -1.35504227 | 0.17540409 | 0.78744529 | 3195.13239 | 2658.5051 | 3747.04869 | 2001.09806 | Non-sig. |
| PF3D7_0611200 | AP2-EXP2 | 666.464088 | -0.575249875 | 0.45941356 | -1.25213953 | 0.21051903 | 0.82617023 | 800.483757 | 662.405304 | 794.732535 | 408.234754 | Non-sig. |
| PF3D7_API01600 | RPS19 | 2.06639293 | -0.575416855 | 2.56585206 | -0.22425956 | 0.82255534 | 0.99096399 | 4.14337 | 0 | 0.81719873 | 3.30500297 | Non-sig. |
| PF3D7_1250700 | N/A | 1237.87326 | -0.575851886 | 0.46075689 | -1.24979549 | 0.21137426 | 0.82768725 | 1110.36527 | 1085.9 | 1852.79668 | 902.431113 | Non-sig. |
| PF3D7_0314400 | PPP6 | 273.762073 | -0.575961986 | 0.57101712 | -1.00865974 | 0.31313784 | 0.88244909 | 174.680314 | 225.346296 | 480.430949 | 214.590731 | Non-sig. |
| PF3D7_1402000 | N/A | 369.830609 | -0.576994592 | 0.4814315 | -1.19849779 | 0.23072328 | 0.83828834 | 338.564146 | 331.656289 | 546.874798 | 262.227203 | Non-sig. |
| PF3D7_1227900 | RPUSP | 121.451746 | -0.577683491 | 0.5128773 | -1.12635808 | 0.26001398 | 0.85030738 | 158.760044 | 89.0868707 | 132.201354 | 105.758717 | Non-sig. |
| PF3D7_0220200 | N/A | 3468.54581 | -0.577988309 | 0.41438976 | -1.39479389 | 0.163078 | 0.76935642 | 4637.15755 | 2823.84873 | 3671.38206 | 2741.7949 | Non-sig. |
| PF3D7_1471800 | N/A | 91.4244227 | -0.578168955 | 0.59145115 | -0.97754303 | 0.32830038 | 0.88909228 | 143.672114 | 83.78041 | 75.5095737 | 62.7355932 | Non-sig. |
| PF3D7_1124800 | RRS1 | 1063.00661 | -0.578738472 | 0.55233762 | -1.0477984 | 0.2947315 | 0.87170077 | 1835.34835 | 1043.67867 | 711.74748 | 661.251947 | Non-sig. |
| PF3D7_0932600 | RPS6 | 268.210354 | -0.578771905 | 0.45022464 | -1.28551806 | 0.19861131 | 0.81528784 | 316.168114 | 211.894145 | 326.415768 | 218.36339 | Non-sig. |
| PF3D7_1131400 | N/A | 335.998456 | -0.57998069 | 0.46826193 | -1.23858178 | 0.21550042 | 0.83503497 | 482.627704 | 256.364036 | 322.830293 | 282.17179 | Non-sig. |
| PF3D7_0213000 | N/A | 1446.85443 | -0.581411867 | 0.48829878 | -1.19068876 | 0.23377579 | 0.83828834 | 2316.34189 | 1257.76954 | 1152.96434 | 1060.34194 | Non-sig. |
| PF3D7_1302900 | N/A | 201.971284 | -0.581657312 | 0.53176789 | -1.09381803 | 0.27403479 | 0.85618956 | 169.509132 | 135.187516 | 314.567837 | 188.62065 | Non-sig. |
| PF3D7_0622700 | N/A | 79.1196161 | -0.581666228 | 0.68211024 | -0.85274519 | 0.39380063 | 0.90601068 | 44.1542589 | 66.5485828 | 145.336973 | 60.4386495 | Non-sig. |
| PF3D7_0312900 | N/A | 65.4239314 | -0.582117435 | 0.66974234 | -0.86916624 | 0.38475621 | 0.90292925 | 111.953561 | 61.2637235 | 45.1269753 | 43.3514661 | Non-sig. |
| PF3D7_1408700 | N/A | 14752.66 | -0.582564195 | 0.53698581 | -1.0848782 | 0.27797561 | 0.85972841 | 10161.2976 | 9164.80636 | 25221.2702 | 14463.2659 | Non-sig. |
| PF3D7_0105600 | N/A | 543.235779 | -0.582626159 | 0.48437746 | -1.20283499 | 0.22904018 | 0.83828834 | 522.435059 | 334.710253 | 780.317154 | 535.480648 | Non-sig. |
| PF3D7_0936700 | N/A | 6.05314114 | -0.582741035 | 1.42096569 | -0.41010212 | 0.68173104 | 0.98067094 | 6.29577146 | 3.22181201 | 8.20838269 | 6.48659838 | Non-sig. |
| PF3D7_1412300 | NTF2 | 3483.43304 | -0.583338758 | 0.46346885 | -1.25863638 | 0.2081617 | 0.82398515 | 2966.53992 | 2870.58501 | 5389.6825 | 2706.92472 | Non-sig. |
| PF3D7_0931700 | N/A | 215.105649 | -0.583348132 | 0.62195634 | -0.93792457 | 0.34828317 | 0.8930363 | 392.290072 | 210.35554 | 123.996807 | 133.780175 | Non-sig. |
| PF3D7_0720100 | ESF2 | 424.598818 | -0.584358447 | 0.48109477 | -1.21464312 | 0.22450224 | 0.83828834 | 633.225149 | 383.310624 | 385.847708 | 296.011792 | Non-sig. |
| PF3D7_0532100 | ETRAMP5 | 10955.2564 | -0.585951228 | 0.47765525 | -1.22672416 | 0.21992628 | 0.83527883 | 8783.21599 | 9181.32763 | 17516.3405 | 8340.1416 | Non-sig. |

| | | | | | | | | | | | | |
|---|---|---|---|---|---|---|---|---|---|---|---|---|
| PF3D7_1135300 | N/A | 112.949034 | -0.586320564 | 0.64022575 | -0.91580285 | 0.35977031 | 0.89560738 | 67.2965713 | 79.4243453 | 203.643357 | 101.431862 | Non-sig. |
| PF3D7_0826000 | N/A | 208.016256 | -0.586741706 | 0.54586192 | -1.07489034 | 0.28242383 | 0.86390383 | 329.664747 | 129.809392 | 170.025467 | 202.565416 | Non-sig. |
| PF3D7_0824700 | LMF1 | 396.880144 | -0.586959717 | 0.4724359 | -1.24241133 | 0.21408484 | 0.83398638 | 364.021178 | 328.37989 | 588.815642 | 306.303867 | Non-sig. |
| PF3D7_1344800 | ATCase | 1180.01712 | -0.587463516 | 0.50743175 | -1.15771926 | 0.24697862 | 0.84582594 | 880.786571 | 1029.07519 | 1952.93863 | 857.268085 | Non-sig. |
| PF3D7_1033300 | N/A | 398.707984 | -0.587776037 | 0.51025791 | -1.15191951 | 0.24935415 | 0.84847675 | 348.385797 | 391.980885 | 609.068096 | 245.397158 | Non-sig. |
| PF3D7_1216300 | SRP19 | 992.041199 | -0.587834115 | 0.42579117 | -1.38056906 | 0.1674115 | 0.77517962 | 1341.12327 | 798.813138 | 1041.80442 | 786.423972 | Non-sig. |
| PF3D7_1031000 | P25 | 161.486255 | -0.588043911 | 0.51368579 | -1.1447541 | 0.25231103 | 0.85030738 | 230.47284 | 109.591161 | 157.555983 | 148.325038 | Non-sig. |
| PF3D7_0724700 | BDP6 | 2769.56047 | -0.588096042 | 0.4176492 | -1.40811007 | 0.15909848 | 0.7667698 | 3106.10364 | 1966.94371 | 3546.54045 | 2458.65408 | Non-sig. |
| PF3D7_0503100 | IspE | 624.736752 | -0.588127567 | 0.4668581 | -1.2597566 | 0.20775718 | 0.82398515 | 558.549292 | 497.962462 | 941.909053 | 500.526202 | Non-sig. |
| PF3D7_1309600 | N/A | 143.674247 | -0.588761683 | 0.49913748 | -1.17955816 | 0.23817599 | 0.84066259 | 194.229941 | 117.040508 | 151.04937 | 112.377171 | Non-sig. |
| PF3D7_1442100 | RPA3 | 463.73111 | -0.588776621 | 0.43632435 | -1.34940125 | 0.17720812 | 0.79027662 | 510.102624 | 388.958359 | 603.947712 | 351.915746 | Non-sig. |
| PF3D7_1142000 | N/A | 120.529971 | -0.588918403 | 0.50763071 | -1.16013156 | 0.24599525 | 0.8454557 | 136.79455 | 92.2852972 | 152.751763 | 100.288273 | Non-sig. |
| PF3D7_1026200 | N/A | 179.739906 | -0.588994687 | 0.52443669 | -1.12309969 | 0.26139516 | 0.85030738 | 152.465305 | 150.263241 | 279.187353 | 137.043725 | Non-sig. |
| PF3D7_1414600 | Pgt1 | 806.148045 | -0.589085864 | 0.46098929 | -1.27787322 | 0.20129413 | 0.81783459 | 1172.64519 | 553.795568 | 764.508041 | 733.643378 | Non-sig. |
| PF3D7_1137100 | N/A | 801.468111 | -0.58910008 | 0.43532539 | -1.35324081 | 0.17597871 | 0.78744529 | 1099.93791 | 682.792954 | 825.938716 | 597.202869 | Non-sig. |
| PF3D7_0531300 | N/A | 575.262775 | -0.589309151 | 0.4753162 | -1.2398255 | 0.21503994 | 0.83503497 | 841.550823 | 540.187591 | 540.934238 | 378.378445 | Non-sig. |
| PF3D7_0618200 | N/A | 1701.20951 | -0.58938868 | 0.43523701 | -1.35417868 | 0.17567938 | 0.78744529 | 1821.97885 | 1574.04 | 2265.83186 | 1142.98735 | Non-sig. |
| PF3D7_0627200 | N/A | 249.598113 | -0.589775899 | 0.46846694 | -1.25894883 | 0.20804882 | 0.82398515 | 269.23364 | 221.569719 | 330.520579 | 177.068514 | Non-sig. |
| PF3D7_1116400 | SEC12 | 2396.24958 | -0.589811619 | 0.48026086 | -1.22810679 | 0.21940688 | 0.83527883 | 2239.21246 | 1428.08872 | 3519.3259 | 2398.37125 | Non-sig. |
| PF3D7_0322600 | N/A | 113.968521 | -0.590848126 | 0.5519996 | -1.07037781 | 0.28444928 | 0.86551387 | 170.631404 | 78.3139564 | 103.494418 | 103.434304 | Non-sig. |
| PF3D7_0822000 | N/A | 790.27891 | -0.591327999 | 0.4180567 | -1.41446843 | 0.15722441 | 0.75912848 | 920.312531 | 620.322631 | 979.668273 | 640.812206 | Non-sig. |
| PF3D7_0722100 | PELO | 773.619167 | -0.591346862 | 0.53204191 | -1.1114667 | 0.26636752 | 0.85416239 | 529.270432 | 649.222977 | 1330.41011 | 585.573146 | Non-sig. |
| PF3D7_0524700 | TOM22 | 316.276456 | -0.591681304 | 0.52583189 | -1.12522901 | 0.260492 | 0.85030738 | 256.202187 | 300.64508 | 504.055104 | 204.203453 | Non-sig. |
| PF3D7_0112700 | N/A | 62.6469239 | -0.593013352 | 0.67298462 | -0.88116925 | 0.37822622 | 0.9000162 | 49.5053392 | 34.3291778 | 101.030681 | 65.7224978 | Non-sig. |
| PF3D7_1333400 | N/A | 1121.00738 | -0.594630048 | 0.46922998 | -1.26724649 | 0.20506717 | 0.82391338 | 976.283459 | 967.787183 | 1721.10785 | 818.85103 | Non-sig. |
| PF3D7_0829300 | LSM8 | 987.240134 | -0.594720365 | 0.45986172 | -1.29325912 | 0.19592145 | 0.81434658 | 1366.04029 | 952.199218 | 1009.88587 | 620.835159 | Non-sig. |
| PF3D7_1024600 | N/A | 258.701227 | -0.595447814 | 0.49775873 | -1.19625789 | 0.23159593 | 0.83828834 | 343.618663 | 258.713304 | 279.15644 | 153.316501 | Non-sig. |
| PF3D7_0425800 | VAR | 21.0143589 | -0.596520905 | 1.13180901 | -0.52705086 | 0.59815827 | 0.95655566 | 47.4437177 | 11.7975024 | 3.28609341 | 21.5301221 | Non-sig. |
| PF3D7_1122900 | N/A | 356.471664 | -0.596928481 | 0.47336807 | -1.26102396 | 0.20730021 | 0.82391338 | 434.454175 | 217.706166 | 423.919719 | 349.806596 | Non-sig. |
| PF3D7_1250900 | N/A | 132.191458 | -0.597443232 | 0.53393262 | -1.11894874 | 0.26316201 | 0.85095134 | 158.305199 | 77.3424376 | 160.038755 | 133.079442 | Non-sig. |
| PF3D7_1143400 | N/A | 2758.99941 | -0.597817938 | 0.41997685 | -1.42345451 | 0.15460444 | 0.7570531 | 3296.70595 | 2503.11642 | 3348.47227 | 1887.70299 | Non-sig. |
| PF3D7_0717900 | N/A | 448.276423 | -0.597819505 | 0.43897781 | -1.36184447 | 0.17324698 | 0.78315911 | 598.576908 | 359.510837 | 481.227344 | 353.790602 | Non-sig. |
| PF3D7_0830800 | SURF8.2 | 1709.72391 | -0.598194662 | 0.55005164 | -1.08752454 | 0.27680506 | 0.85948573 | 3054.84587 | 1262.3944 | 1063.84067 | 1457.8147 | Non-sig. |
| PF3D7_1437600 | N/A | 53.3509355 | -0.600131277 | 0.68192403 | -0.88005592 | 0.37882902 | 0.9000162 | 38.9104796 | 47.2237002 | 89.5064146 | 37.7631476 | Non-sig. |
| PF3D7_1439200 | N/A | 363.297132 | -0.60095104 | 0.45355176 | -1.32498889 | 0.18517484 | 0.80052251 | 388.598318 | 258.554965 | 487.067296 | 318.967948 | Non-sig. |
| PF3D7_0519100 | N/A | 547.234776 | -0.601481938 | 0.44036007 | -1.36588665 | 0.17197454 | 0.78315911 | 656.246507 | 498.437265 | 663.128037 | 371.127294 | Non-sig. |
| PF3D7_1317200 | ApiAP2 | 1457.66588 | -0.601678604 | 0.4508197 | -1.33463244 | 0.18199667 | 0.79970932 | 1395.73167 | 1011.40199 | 2118.6644 | 1304.86546 | Non-sig. |
| PF3D7_1125400 | TIM44 | 731.7154 | -0.601771733 | 0.47748907 | -1.26028379 | 0.207567 | 0.82398515 | 1137.04753 | 618.005024 | 627.489904 | 544.319143 | Non-sig. |
| PF3D7_1426200 | PRMT1 | 2437.48657 | -0.602441769 | 0.41196962 | -1.46234513 | 0.14364666 | 0.7412338 | 3090.39899 | 2085.22351 | 2787.94751 | 1786.37628 | Non-sig. |
| PF3D7_1144700 | TIC20 | 685.17367 | -0.602843615 | 0.50783048 | -1.18709616 | 0.23518969 | 0.83828834 | 536.626645 | 617.372868 | 1115.67929 | 471.015879 | Non-sig. |
| PF3D7_0217700 | N/A | 256.139791 | -0.603132329 | 0.47663082 | -1.26540773 | 0.20572521 | 0.82391338 | 291.020349 | 241.550659 | 326.765954 | 165.222202 | Non-sig. |
| PF3D7_0817100 | N/A | 273.81299 | -0.603259229 | 0.5158329 | -1.16948576 | 0.24220798 | 0.8422984 | 438.966607 | 208.155541 | 221.751602 | 226.378208 | Non-sig. |
| PF3D7_0922500 | PGK | 4657.35419 | -0.604251592 | 0.59184926 | -1.02095522 | 0.30727566 | 0.87709318 | 2388.40835 | 3832.69007 | 8848.63425 | 3559.68411 | Non-sig. |
| PF3D7_0204300 | N/A | 888.786088 | -0.60538537 | 0.49612299 | -1.22023244 | 0.22237677 | 0.83675579 | 1439.93002 | 767.191145 | 705.487222 | 642.535969 | Non-sig. |
| PF3D7_0720900 | N/A | 359.714389 | -0.606032976 | 0.45204861 | -1.34063675 | 0.18003842 | 0.79619661 | 373.978188 | 295.237616 | 494.244565 | 275.397186 | Non-sig. |
| PF3D7_0613000 | N/A | 625.444727 | -0.606458766 | 0.50252912 | -1.20681318 | 0.2275041 | 0.83828834 | 495.696379 | 544.242439 | 1013.98193 | 447.85816 | Non-sig. |
| PF3D7_1138300 | N/A | 61.8623653 | -0.606723292 | 0.61512546 | -0.98634073 | 0.32396594 | 0.88694295 | 56.519472 | 47.2865555 | 92.7210769 | 50.9223568 | Non-sig. |
| PF3D7_0322900 | RPS3A | 4890.26914 | -0.607065463 | 0.44088547 | -1.37692326 | 0.16853598 | 0.77543678 | 4585.55044 | 4047.15019 | 7222.69734 | 3705.67861 | Non-sig. |
| PF3D7_1210900 | PIGM | 236.882626 | -0.607509666 | 0.58344342 | -1.04124865 | 0.29776016 | 0.87301237 | 148.146479 | 192.16341 | 423.65122 | 183.569394 | Non-sig. |
| PF3D7_1141700 | N/A | 523.956653 | -0.607716843 | 0.45833023 | -1.32593665 | 0.18486068 | 0.80052251 | 741.03264 | 470.343548 | 524.544395 | 359.906031 | Non-sig. |
| PF3D7_0922700 | PRP18 | 1002.01958 | -0.60804137 | 0.51396214 | -1.18304701 | 0.23679052 | 0.83828834 | 1655.32581 | 926.550657 | 765.154591 | 661.047246 | Non-sig. |
| PF3D7_1032000 | RimM | 711.224739 | -0.608537803 | 0.44421256 | -1.36992479 | 0.17071038 | 0.78169715 | 701.119795 | 577.247646 | 1016.78139 | 549.750124 | Non-sig. |
| PF3D7_0530750 | DPM3 | 80.9550194 | -0.60898072 | 0.7153711 | -0.85127945 | 0.39461414 | 0.90645858 | 38.6753614 | 69.8809194 | 156.689018 | 58.5747789 | Non-sig. |
| PF3D7_1421150 | N/A | 3793.88107 | -0.609059035 | 0.57358043 | -1.06185462 | 0.28830169 | 0.86724628 | 6095.97877 | 4368.36867 | 3070.19213 | 1640.98471 | Non-sig. |

| | | | | | | | | | | | | |
|---|---|---|---|---|---|---|---|---|---|---|---|---|
| PF3D7_0728800 | N/A | 1093.46352 | -0.609526493 | 0.42662703 | -1.42871044 | 0.15308748 | 0.75391062 | 1398.09113 | 973.360407 | 1244.13012 | 758.272428 | Non-sig. |
| PF3D7_0208200 | KRR1 | 549.229116 | -0.609609883 | 0.43407229 | -1.40439713 | 0.16020063 | 0.7667698 | 621.84973 | 475.610964 | 705.23337 | 394.222402 | Non-sig. |
| PF3D7_1307200 | N/A | 862.814018 | -0.611311558 | 0.51087615 | -1.19659444 | 0.23146467 | 0.83828834 | 1401.30743 | 813.620639 | 684.802569 | 551.525431 | Non-sig. |
| PF3D7_1432700 | N/A | 1186.72017 | -0.611949425 | 0.4289075 | -1.42676317 | 0.15364817 | 0.75442818 | 1241.77257 | 1005.69748 | 1627.49091 | 871.919733 | Non-sig. |
| PF3D7_0818400 | FCF1 | 1403.89059 | -0.612524316 | 0.48773333 | -1.25585906 | 0.20916708 | 0.82566351 | 2240.27713 | 1255.26271 | 1155.01774 | 965.004757 | Non-sig. |
| PF3D7_1353500 | TFB4 | 319.895609 | -0.613114634 | 0.49274908 | -1.24427353 | 0.21339891 | 0.83292034 | 278.429818 | 246.914254 | 495.080316 | 259.158047 | Non-sig. |
| PF3D7_0934200 | N/A | 304.338517 | -0.613585306 | 0.4625872 | -1.32642085 | 0.18470034 | 0.80052251 | 328.361 | 270.890027 | 407.749047 | 210.353996 | Non-sig. |
| PF3D7_1367800 | PSOP2 | 70.1622035 | -0.613628165 | 0.94049585 | -0.65245175 | 0.51410979 | 0.93247207 | 160.06235 | 60.0802892 | 9.85622186 | 50.6499528 | Non-sig. |
| PF3D7_1468200 | TLS1 | 368.179516 | -0.615065809 | 0.43835814 | -1.40311254 | 0.16058328 | 0.7667698 | 456.740517 | 279.172727 | 434.273248 | 302.531571 | Non-sig. |
| PF3D7_1223800 | YHM2 | 284.328144 | -0.615170272 | 0.46950733 | -1.31024638 | 0.1901125 | 0.80577283 | 295.669249 | 251.356624 | 392.306329 | 197.980376 | Non-sig. |
| PF3D7_1456900 | N/A | 321.924963 | -0.615394951 | 0.47104848 | -1.30643654 | 0.19140415 | 0.80801527 | 324.030502 | 286.530404 | 454.953758 | 222.185187 | Non-sig. |
| PF3D7_1004900 | N/A | 964.775938 | -0.615495442 | 0.43121053 | -1.42736646 | 0.15347429 | 0.75442818 | 1295.09815 | 841.918015 | 1040.04003 | 682.047554 | Non-sig. |
| PF3D7_0405800 | N/A | 293.329149 | -0.615521664 | 0.48012877 | -1.28199287 | 0.19984515 | 0.81595849 | 312.444701 | 277.902007 | 397.418719 | 185.551169 | Non-sig. |
| PF3D7_0205100 | N/A | 278.689513 | -0.616484452 | 0.65957803 | -0.93466493 | 0.34996101 | 0.8930363 | 549.313868 | 253.243988 | 125.651889 | 186.548306 | Non-sig. |
| PF3D7_0516200 | RPS11 | 2601.7581 | -0.617128678 | 0.4346699 | -1.41976401 | 0.1556764 | 0.75876132 | 2537.15121 | 2136.97731 | 3762.4389 | 1970.46498 | Non-sig. |
| PF3D7_1410500 | N/A | 177.341158 | -0.617237508 | 0.51202625 | -1.20548021 | 0.22801797 | 0.83828834 | 211.905701 | 176.137831 | 217.512732 | 103.808366 | Non-sig. |
| PF3D7_0321000 | N/A | 983.080113 | -0.617257002 | 0.62309449 | -0.99063146 | 0.32186557 | 0.88694295 | 1923.38936 | 815.688089 | 457.383894 | 735.859109 | Non-sig. |
| PF3D7_0916800 | N/A | 490.365587 | -0.617319125 | 0.46138748 | -1.33796246 | 0.18090868 | 0.79699299 | 463.96164 | 396.990268 | 723.263087 | 377.247353 | Non-sig. |
| PF3D7_0203800 | N/A | 227.714652 | -0.617739432 | 0.4640362 | -1.33123112 | 0.18311297 | 0.80052251 | 294.503038 | 191.033061 | 257.029943 | 168.292568 | Non-sig. |
| PF3D7_1224900 | SF3B6 | 470.297741 | -0.618065914 | 0.46468305 | -1.3300806 | 0.18349172 | 0.80052251 | 629.655144 | 447.160951 | 509.497175 | 294.877692 | Non-sig. |
| PF3D7_0630200 | PSOP6 | 31.1378944 | -0.619416474 | 0.7973305 | -0.77686289 | 0.43723967 | 0.91524921 | 53.4986549 | 29.1739282 | 22.069627 | 19.8093676 | Non-sig. |
| PF3D7_0404300 | N/A | 3250.67799 | -0.619686214 | 0.64948131 | -0.95412479 | 0.34002048 | 0.8930363 | 6542.32029 | 2958.4461 | 1334.53851 | 2167.40706 | Non-sig. |
| PF3D7_0917100 | OGG1 | 556.598916 | -0.619693895 | 0.45701495 | -1.35595978 | 0.17511197 | 0.78744529 | 819.111319 | 433.459479 | 529.754405 | 444.070459 | Non-sig. |
| PF3D7_0822800 | SNRNP40 | 401.988766 | -0.619796687 | 0.55612881 | -1.11448404 | 0.26507159 | 0.85370776 | 273.370783 | 268.331433 | 700.424936 | 365.827915 | Non-sig. |
| PF3D7_1361300 | N/A | 30.8417716 | -0.619939162 | 0.79708229 | -0.77776055 | 0.43671019 | 0.91522571 | 53.5239389 | 21.4776341 | 21.3424901 | 27.0230233 | Non-sig. |
| PF3D7_1336300 | N/A | 26.0616283 | -0.620217186 | 0.8106431 | -0.76509279 | 0.44421629 | 0.91974025 | 41.9155872 | 18.2637974 | 21.3367378 | 22.7303907 | Non-sig. |
| PF3D7_1116300 | CYP19C | 469.747661 | -0.620364236 | 0.43420854 | -1.42872418 | 0.15308352 | 0.75391062 | 550.54808 | 399.358493 | 587.855156 | 341.228913 | Non-sig. |
| PF3D7_1252300 | N/A | 1547.30498 | -0.62045438 | 0.44157904 | -1.40508113 | 0.15999715 | 0.7667698 | 1489.55454 | 1159.8736 | 2260.22787 | 1279.56392 | Non-sig. |
| PF3D7_0726900 | TIM50 | 690.155093 | -0.621045032 | 0.44267205 | -1.40294612 | 0.16063291 | 0.7667698 | 739.829818 | 469.995622 | 932.954681 | 617.84025 | Non-sig. |
| PF3D7_1433000 | NOP10 | 127.960219 | -0.621416431 | 0.53316404 | -1.16552577 | 0.24380624 | 0.84313671 | 117.215005 | 99.9489623 | 192.823672 | 101.853237 | Non-sig. |
| PF3D7_0719100 | N/A | 707.876557 | -0.621640757 | 0.4621655 | -1.34506094 | 0.17860556 | 0.79309819 | 999.847504 | 662.838546 | 716.442406 | 452.377773 | Non-sig. |
| PF3D7_0402000 | N/A | 3516.09385 | -0.62191927 | 0.51583366 | -1.20565857 | 0.22794917 | 0.83828834 | 6010.29751 | 3056.80194 | 2514.86839 | 2482.40755 | Non-sig. |
| PF3D7_0929900 | N/A | 223.478888 | -0.622063848 | 0.50530016 | -1.23107789 | 0.21829373 | 0.83527883 | 199.368948 | 189.012212 | 342.290422 | 163.243967 | Non-sig. |
| PF3D7_0705900 | N/A | 1456.55465 | -0.622516501 | 0.50952808 | -1.22175113 | 0.22180176 | 0.83637416 | 1087.82507 | 1271.29401 | 2443.93466 | 1023.16486 | Non-sig. |
| PF3D7_0504300 | N/A | 52.1301359 | -0.622671795 | 0.71625607 | -0.86934243 | 0.38465986 | 0.90292925 | 92.126559 | 49.4329116 | 34.4638942 | 32.4971787 | Non-sig. |
| PF3D7_0324200 | EPF3 | 6.88870945 | -0.623074714 | 1.72453927 | -0.36129923 | 0.71787577 | 0.98797939 | 16.7715454 | 4.29505453 | 0 | 6.48823789 | Non-sig. |
| PF3D7_0419600 | RANBP1 | 5458.22732 | -0.623089598 | 0.43057255 | -1.44711872 | 0.14786368 | 0.74652201 | 7637.01554 | 4806.26278 | 5601.0014 | 3788.62957 | Non-sig. |
| PF3D7_0530000 | N/A | 943.153047 | -0.623332715 | 0.43208355 | -1.4426208 | 0.14912732 | 0.74817855 | 1018.6407 | 823.045301 | 1268.83546 | 662.090725 | Non-sig. |
| PF3D7_0530300 | PIC6 | 4554.99532 | -0.623468913 | 0.42853747 | -1.45487609 | 0.14570357 | 0.7422476 | 5755.80068 | 2942.27293 | 5292.62506 | 4229.28259 | Non-sig. |
| PF3D7_1324300 | N/A | 1490.86628 | -0.623476057 | 0.47993501 | -1.29908433 | 0.19391499 | 0.81267745 | 1425.73723 | 878.32263 | 2190.26594 | 1469.1393 | Non-sig. |
| PF3D7_0919700 | N/A | 199.489986 | -0.623562849 | 0.52499162 | -1.18775771 | 0.23492888 | 0.83828834 | 173.350016 | 180.35825 | 310.345096 | 133.906582 | Non-sig. |
| PF3D7_1007000 | TMEM147 | 677.0436 | -0.62385821 | 0.56507669 | -1.10402397 | 0.26958274 | 0.8556331 | 414.416878 | 575.242097 | 1227.67166 | 490.843759 | Non-sig. |
| PF3D7_0624700 | PIGL | 693.105245 | -0.624251195 | 0.48403105 | -1.28969246 | 0.19715746 | 0.81471289 | 592.857472 | 602.477232 | 1088.43338 | 488.652899 | Non-sig. |
| PF3D7_0705600 | N/A | 459.744299 | -0.624435837 | 0.44105657 | -1.41577267 | 0.15684207 | 0.75894163 | 506.679681 | 330.411005 | 608.656612 | 393.229896 | Non-sig. |
| PF3D7_1333000 | CPN20 | 975.440643 | -0.626512577 | 0.42776232 | -1.46462779 | 0.1430225 | 0.74065302 | 1097.78273 | 854.576584 | 1270.09556 | 679.307702 | Non-sig. |
| PF3D7_1372100 | GEXP04 | 378.984907 | -0.627034565 | 0.472969 | -1.32574136 | 0.18492538 | 0.80052251 | 565.512255 | 314.344147 | 354.827143 | 281.256085 | Non-sig. |
| PF3D7_1124500 | pdhA | 245.830571 | -0.628317146 | 0.46201802 | -1.35994079 | 0.17384866 | 0.78387252 | 329.420585 | 187.780003 | 267.734935 | 198.386763 | Non-sig. |
| PF3D7_1453600 | N/A | 128.543963 | -0.628713806 | 0.52024596 | -1.20849339 | 0.22685754 | 0.83828834 | 126.452576 | 98.6840855 | 185.64793 | 103.391262 | Non-sig. |
| PF3D7_1408200 | AP2-G2 | 1156.89702 | -0.628777547 | 0.4530923 | -1.38774715 | 0.16521405 | 0.77284241 | 1457.2895 | 691.87147 | 1352.91746 | 1125.50965 | Non-sig. |
| PF3D7_1025800 | C4AP2 | 684.727536 | -0.62882412 | 0.45351397 | -1.38655953 | 0.16557612 | 0.77284241 | 905.122345 | 645.729784 | 758.237075 | 429.820942 | Non-sig. |
| PF3D7_0624300 | N/A | 38.7185983 | -0.629273346 | 0.68493541 | -0.91873385 | 0.3582348 | 0.89552149 | 53.9674594 | 28.0071269 | 40.150327 | 32.7494799 | Non-sig. |
| PF3D7_1467800 | N/A | 192.513793 | -0.629343329 | 0.52373738 | -1.20163914 | 0.22950337 | 0.83828834 | 163.198755 | 159.160141 | 304.315194 | 143.381081 | Non-sig. |
| PF3D7_1242400 | N/A | 214.83805 | -0.630080032 | 0.59801545 | -1.05361833 | 0.2920577 | 0.86879375 | 390.932821 | 152.433906 | 131.369341 | 184.61613 | Non-sig. |
| PF3D7_1202500 | TGS1 | 688.52996 | -0.630358809 | 0.44429853 | -1.41877313 | 0.15596517 | 0.75894163 | 824.698746 | 440.921268 | 848.486417 | 640.01341 | Non-sig. |

| | | | | | | | | | | | | |
|---|---|---|---|---|---|---|---|---|---|---|---|---|
| PF3D7_1354600 | N/A | 120.901867 | -0.630508225 | 0.53585985 | -1.17662897 | 0.23934363 | 0.84087702 | 181.519319 | 96.6556685 | 112.453886 | 92.9785958 | Non-sig. |
| PF3D7_0414500 | N/A | 178.933902 | -0.63062934 | 0.50324517 | -1.25312547 | 0.21016005 | 0.82566351 | 260.915969 | 154.53187 | 174.098796 | 126.188973 | Non-sig. |
| PF3D7_1250500 | N/A | 898.265153 | -0.630678329 | 0.43525812 | -1.44897545 | 0.14734444 | 0.74558819 | 964.071077 | 787.249031 | 1218.89323 | 622.847277 | Non-sig. |
| PF3D7_0822200 | N/A | 69.8443857 | -0.631417191 | 0.57374135 | -1.10052585 | 0.27110307 | 0.8556331 | 76.9033465 | 53.6369692 | 92.8193453 | 56.0178818 | Non-sig. |
| PF3D7_1413900 | N/A | 522.167801 | -0.633271902 | 0.51120135 | -1.23879152 | 0.21542271 | 0.83503497 | 422.988524 | 341.246046 | 846.696849 | 477.739783 | Non-sig. |
| PF3D7_1221100 | N/A | 49.5120534 | -0.633767649 | 0.65610305 | -0.96595748 | 0.33406546 | 0.89204385 | 74.5661868 | 44.0204519 | 45.9741403 | 33.4874346 | Non-sig. |
| PF3D7_0904200 | PH | 108.832423 | -0.635958697 | 0.65100381 | -0.97688935 | 0.32862393 | 0.88909228 | 200.270723 | 98.8222461 | 64.8379527 | 71.3987719 | Non-sig. |
| PF3D7_0218500 | SNRPD2 | 662.256931 | -0.636127372 | 0.42053908 | -1.51264746 | 0.13036923 | 0.71872652 | 807.92312 | 521.558625 | 803.962088 | 515.583891 | Non-sig. |
| PF3D7_0935500 | GEXP22 | 1952.40502 | -0.636725974 | 0.43201653 | -1.47384634 | 0.14052297 | 0.73498918 | 2789.37241 | 1441.17971 | 1963.57659 | 1615.49137 | Non-sig. |
| PF3D7_1003500 | N/A | 4623.73208 | -0.63677783 | 0.41661573 | -1.52845363 | 0.12639995 | 0.71393062 | 5710.98964 | 4100.62298 | 5544.826 | 3138.48969 | Non-sig. |
| PF3D7_1346300 | ALBA2 | 3265.04206 | -0.637596407 | 0.43399235 | -1.46914203 | 0.14179427 | 0.73825055 | 3623.50693 | 3015.23831 | 4326.44019 | 2094.98282 | Non-sig. |
| PF3D7_0527300 | METAP1a | 897.38618 | -0.639223711 | 0.46084017 | -1.38708333 | 0.16541635 | 0.77284241 | 829.987625 | 763.027968 | 1355.80573 | 640.723398 | Non-sig. |
| PF3D7_0203400 | N/A | 1247.34883 | -0.63930523 | 0.49265085 | -1.2976842 | 0.19439588 | 0.81287241 | 999.561934 | 1064.84217 | 2038.74566 | 886.24555 | Non-sig. |
| PF3D7_0421800 | N/A | 2.10385492 | -0.640019548 | 2.57562948 | -0.24849054 | 0.80375489 | 0.99028237 | 5.15304201 | 1.08455266 | 0 | 2.177825 | Non-sig. |
| PF3D7_0914700 | MFR4 | 456.218931 | -0.640451334 | 0.59740387 | -1.07205757 | 0.28369418 | 0.86551387 | 273.700892 | 268.240384 | 837.730781 | 445.203666 | Non-sig. |
| PF3D7_0608200 | N/A | 72.7469971 | -0.64048603 | 0.56811098 | -1.12739598 | 0.2595751 | 0.85030738 | 90.40943 | 62.4546981 | 86.876987 | 51.2468734 | Non-sig. |
| PF3D7_0205500 | N/A | 340.915237 | -0.640643559 | 0.45175907 | -1.41810889 | 0.15615897 | 0.75894163 | 366.262499 | 281.584908 | 464.411303 | 251.402239 | Non-sig. |
| PF3D7_0611300 | ATPTG2 | 695.17322 | -0.642942925 | 0.42040406 | -1.52934518 | 0.12617889 | 0.71393062 | 819.128204 | 542.212349 | 875.962227 | 543.390101 | Non-sig. |
| PF3D7_1449800 | N/A | 404.072418 | -0.643840648 | 0.44078386 | -1.460672 | 0.14410548 | 0.7412338 | 451.79178 | 332.797473 | 533.668665 | 298.031754 | Non-sig. |
| PF3D7_0612500 | N/A | 128.828833 | -0.64471779 | 0.54360739 | -1.18599895 | 0.23562271 | 0.83828834 | 200.198902 | 108.226291 | 114.258946 | 92.6311927 | Non-sig. |
| PF3D7_1457300 | N/A | 2192.02256 | -0.644834837 | 0.50765204 | -1.27022998 | 0.20400272 | 0.82253153 | 1614.43159 | 1636.0812 | 3733.09187 | 1784.48556 | Non-sig. |
| PF3D7_1427900 | N/A | 7774.72145 | -0.645238014 | 0.40756775 | -1.58314296 | 0.11338888 | 0.68498781 | 10099.2202 | 6439.22093 | 8870.68392 | 5689.76071 | Non-sig. |
| PF3D7_1202900 | HMGB1 | 5465.24443 | -0.645585927 | 0.42400253 | -1.52259924 | 0.127859 | 0.71411411 | 7787.08176 | 4299.80207 | 5549.19873 | 4224.89515 | Non-sig. |
| PF3D7_0219400 | N/A | 4637.79004 | -0.646020996 | 0.4293821 | -1.50453641 | 0.13244331 | 0.72037933 | 5165.25441 | 4218.63502 | 6152.9748 | 3014.29591 | Non-sig. |
| PF3D7_1470000 | ATG12 | 112.659132 | -0.646498298 | 0.69173158 | -0.93460862 | 0.34999003 | 0.8930363 | 52.7992104 | 90.5512564 | 221.948818 | 85.3372434 | Non-sig. |
| PF3D7_0314300 | DER1-2 | 128.347011 | -0.646503292 | 0.62055337 | -1.04181739 | 0.29749635 | 0.87301237 | 87.244067 | 76.2120908 | 225.800057 | 124.131828 | Non-sig. |
| PF3D7_1429000 | N/A | 214.810753 | -0.646707281 | 0.48810047 | -1.32494705 | 0.18518872 | 0.80052251 | 215.862406 | 185.036603 | 308.341112 | 150.002891 | Non-sig. |
| PF3D7_1476300 | N/A | 8170.25759 | -0.647114618 | 0.40616154 | -1.59324443 | 0.11110537 | 0.68366578 | 10619.6811 | 6071.76206 | 9325.41492 | 6664.17228 | Non-sig. |
| PF3D7_1242300 | N/A | 319.073027 | -0.647190176 | 0.47412143 | -1.36503041 | 0.17224349 | 0.78315911 | 406.279719 | 302.807972 | 372.693326 | 194.51109 | Non-sig. |
| PF3D7_0319100 | RBX1 | 1294.23268 | -0.648383752 | 0.45366348 | -1.42921744 | 0.15294175 | 0.75391062 | 1888.00228 | 1162.47388 | 1272.71096 | 853.743621 | Non-sig. |
| PF3D7_0202600 | N/A | 2851.98962 | -0.648723434 | 0.40470652 | -1.60294782 | 0.10894617 | 0.6779986 | 3520.60498 | 2170.90983 | 3444.64054 | 2271.80314 | Non-sig. |
| PF3D7_1141400 | PIGH | 2175.77495 | -0.649007086 | 0.46483533 | -1.3962086 | 0.16265169 | 0.76900226 | 3183.63736 | 2059.9179 | 2130.7026 | 1328.84193 | Non-sig. |
| PF3D7_0625700 | N/A | 463.266285 | -0.649241477 | 0.50761322 | -1.27900821 | 0.20089416 | 0.81683598 | 374.533685 | 399.524053 | 756.789807 | 322.217596 | Non-sig. |
| PF3D7_0821900 | N/A | 298.276086 | -0.649668104 | 0.4719018 | -1.3767019 | 0.16860443 | 0.77543678 | 359.365447 | 279.341888 | 369.278637 | 185.118374 | Non-sig. |
| PF3D7_1016100 | N/A | 27.1091471 | -0.650016358 | 0.92128345 | -0.70555523 | 0.48046478 | 0.92880667 | 53.246029 | 29.0369945 | 13.1233854 | 13.0301795 | Non-sig. |
| PF3D7_1477500 | N/A | 365.442817 | -0.650274656 | 0.44095879 | -1.47468351 | 0.14029765 | 0.73498918 | 470.742422 | 270.356008 | 422.179962 | 298.492879 | Non-sig. |
| PF3D7_1308200 | cpsSII | 7258.48807 | -0.650506674 | 0.54972315 | -1.18333506 | 0.23667638 | 0.83828834 | 4555.52853 | 5009.68653 | 13179.6374 | 6289.09975 | Non-sig. |
| PF3D7_0419100 | N/A | 92.6340397 | -0.651721026 | 0.57692829 | -1.12963957 | 0.25862813 | 0.85030738 | 80.924129 | 66.5544388 | 145.328986 | 77.7286053 | Non-sig. |
| PF3D7_0616100 | N/A | 159.599248 | -0.652478801 | 0.56597012 | -1.15285026 | 0.24897185 | 0.8482655 | 261.562137 | 150.474173 | 128.816044 | 97.5446392 | Non-sig. |
| PF3D7_1408800 | N/A | 25.6713024 | -0.653761639 | 0.90173837 | -0.72500147 | 0.46845113 | 0.92594245 | 47.3030164 | 27.9083187 | 15.6008479 | 11.8730265 | Non-sig. |
| PF3D7_0512200 | GS | 220.688565 | -0.654005023 | 0.49396727 | -1.32398452 | 0.18550819 | 0.80052251 | 209.420241 | 160.958544 | 330.146242 | 182.229232 | Non-sig. |
| PF3D7_0422100 | N/A | 1394.00191 | -0.654405316 | 0.43448907 | -1.50614908 | 0.13202891 | 0.72037933 | 1422.22916 | 1165.47018 | 1987.30925 | 1000.99903 | Non-sig. |
| PF3D7_1341300 | N/A | 4387.69015 | -0.655114736 | 0.4579796 | -1.43044523 | 0.15258927 | 0.75391062 | 4082.1986 | 3904.89043 | 6651.84435 | 2911.82723 | Non-sig. |
| PF3D7_1025300 | N/A | 635.250369 | -0.655306825 | 0.58926178 | -1.112081 | 0.26610333 | 0.85416239 | 448.467204 | 304.717478 | 1105.48421 | 682.332582 | Non-sig. |
| PF3D7_0626800 | PyrK | 21482.5804 | -0.655913386 | 0.53813887 | -1.21885524 | 0.22289914 | 0.83793839 | 13846.2897 | 17314.6481 | 38720.7057 | 16048.6782 | Non-sig. |
| PF3D7_1107600 | N/A | 32.6489438 | -0.6565232 | 0.86267824 | -0.76102905 | 0.44663972 | 0.91974025 | 65.2846574 | 27.8948929 | 14.7841773 | 22.6320475 | Non-sig. |
| PF3D7_1351400 | N/A | 4505.88004 | -0.657038791 | 0.44875191 | -1.46414705 | 0.14315377 | 0.74065302 | 4279.30894 | 3906.37375 | 6749.59461 | 3088.24284 | Non-sig. |
| PF3D7_0301400 | N/A | 294.309322 | -0.658068065 | 0.5757187 | -1.1430375 | 0.25302302 | 0.85030738 | 186.770968 | 220.158339 | 533.543068 | 236.764912 | Non-sig. |
| PF3D7_1301900 | N/A | 195.138611 | -0.659168461 | 0.55363381 | -1.19062176 | 0.23380211 | 0.83828834 | 325.591108 | 175.318794 | 152.548446 | 127.096094 | Non-sig. |
| PF3D7_1117600 | ICT1 | 235.122192 | -0.659434012 | 0.49276987 | -1.33821902 | 0.18082505 | 0.79699299 | 253.810864 | 221.238115 | 321.991831 | 143.447959 | Non-sig. |
| PF3D7_1422300 | N/A | 233.360642 | -0.660981168 | 0.46435794 | -1.42343032 | 0.15461145 | 0.7570531 | 300.852156 | 164.196691 | 270.996084 | 197.397637 | Non-sig. |
| PF3D7_0105100 | N/A | 323.460421 | -0.662226297 | 0.45569849 | -1.45321151 | 0.14616504 | 0.7426703 | 342.945278 | 261.00396 | 449.779022 | 240.113422 | Non-sig. |
| PF3D7_0729400 | N/A | 531.973184 | -0.662538297 | 0.44583285 | -1.48606881 | 0.13726092 | 0.7277391 | 564.849563 | 453.963959 | 739.07171 | 370.007504 | Non-sig. |
| PF3D7_0512300 | MAT1 | 329.265117 | -0.662862059 | 0.45644885 | -1.45221543 | 0.14644171 | 0.7426703 | 366.994929 | 285.494318 | 440.136267 | 224.434957 | Non-sig. |

| | | | | | | | | | | | | |
|---|---|---|---|---|---|---|---|---|---|---|---|---|
| PF3D7_0316400 | N/A | 1630.012 | -0.662935157 | 0.47873372 | -1.38476803 | 0.16612341 | 0.77284241 | 2484.20366 | 985.951735 | 1512.117 | 1537.7756 | Non-sig. |
| PF3D7_0626100 | N/A | 459.61424 | -0.663878805 | 0.47291418 | -1.40380397 | 0.16037723 | 0.7667698 | 430.676887 | 389.720852 | 696.20356 | 321.855659 | Non-sig. |
| PF3D7_1421600 | N/A | 628.488963 | -0.664802934 | 0.42690799 | -1.5572511 | 0.11941088 | 0.70181652 | 827.153279 | 505.410524 | 714.494987 | 466.89706 | Non-sig. |
| PF3D7_1203200 | SRP14 | 178.562002 | -0.665053255 | 0.51672038 | -1.28706603 | 0.19807127 | 0.81528784 | 232.989933 | 175.262203 | 205.076697 | 100.919174 | Non-sig. |
| PF3D7_1448800 | N/A | 317.616339 | -0.666382222 | 0.46106189 | -1.44532053 | 0.14836787 | 0.74754886 | 366.729626 | 283.859053 | 412.615358 | 207.261321 | Non-sig. |
| PF3D7_0502700 | N/A | 108.796794 | -0.666534771 | 0.56581057 | -1.17801753 | 0.23878962 | 0.84066259 | 143.876421 | 57.9787007 | 123.144294 | 110.187762 | Non-sig. |
| PF3D7_0312800 | N/A | 3108.72009 | -0.666590076 | 0.44518022 | -1.49734881 | 0.13430252 | 0.72178914 | 3346.53607 | 2893.36075 | 4282.15162 | 1912.83191 | Non-sig. |
| PF3D7_1228000 | N/A | 354.671603 | -0.666631735 | 0.45416022 | -1.46783381 | 0.14214937 | 0.73871836 | 370.35963 | 280.306475 | 499.8806 | 268.139708 | Non-sig. |
| PF3D7_0629200 | N/A | 3258.38308 | -0.666864416 | 0.40635876 | -1.64107307 | 0.10078225 | 0.65652696 | 4146.14588 | 2632.74683 | 3850.54927 | 2404.09034 | Non-sig. |
| PF3D7_0511300 | N/A | 571.636868 | -0.667233069 | 0.46969441 | -1.42056848 | 0.15544225 | 0.75842353 | 521.450295 | 426.954332 | 881.367889 | 456.774955 | Non-sig. |
| PF3D7_0927000 | P23 | 1526.29343 | -0.667238118 | 0.47696869 | -1.39891388 | 0.16183881 | 0.76806182 | 2417.4831 | 1333.84459 | 1328.92812 | 1024.91792 | Non-sig. |
| PF3D7_0316600 | FNT | 4961.95346 | -0.667689479 | 0.54330946 | -1.22893034 | 0.21909792 | 0.83527883 | 3156.69678 | 3877.43584 | 9023.21023 | 3790.47098 | Non-sig. |
| PF3D7_1120300 | MIT1 | 875.191748 | -0.668369579 | 0.43455423 | -1.53805792 | 0.12403445 | 0.708883 | 1232.35869 | 717.86829 | 916.530798 | 634.009213 | Non-sig. |
| PF3D7_1327200 | RPR2 | 282.787757 | -0.669274778 | 0.50448884 | -1.32663941 | 0.18462799 | 0.80052251 | 447.64024 | 238.47513 | 247.032471 | 198.003186 | Non-sig. |
| PF3D7_1367000 | SKP1 | 1755.91471 | -0.670914257 | 0.45900231 | -1.46167947 | 0.14382906 | 0.7412338 | 1631.06667 | 1514.38305 | 2682.72361 | 1195.48553 | Non-sig. |
| PF3D7_0531500 | N/A | 13.2991045 | -0.67092842 | 1.00574926 | -0.66709313 | 0.50471265 | 0.93170321 | 14.6101557 | 9.6801231 | 18.0434141 | 10.862725 | Non-sig. |
| PF3D7_1414300 | N/A | 6455.46788 | -0.671035151 | 0.44450231 | -1.50963252 | 0.13113722 | 0.72037933 | 6157.65092 | 5439.13672 | 9702.70741 | 4522.37649 | Non-sig. |
| PF3D7_1219900 | N/A | 575.326805 | -0.671224482 | 0.43809458 | -1.53214514 | 0.12548662 | 0.71290716 | 623.072334 | 476.873045 | 790.420446 | 410.941394 | Non-sig. |
| PF3D7_0520700 | N/A | 362.370348 | -0.671303649 | 0.48311746 | -1.38952471 | 0.16467325 | 0.77284241 | 339.062057 | 246.761893 | 551.122137 | 312.535305 | Non-sig. |
| PF3D7_0820100 | N/A | 319.272428 | -0.672476996 | 0.49197773 | -1.36688503 | 0.17166134 | 0.78315911 | 497.617824 | 265.224838 | 287.31904 | 226.928011 | Non-sig. |
| PF3D7_0528000 | UMP1 | 1185.6855 | -0.672557551 | 0.46997413 | -1.43105228 | 0.15241523 | 0.75391062 | 1059.81491 | 1011.41162 | 1854.28546 | 817.229991 | Non-sig. |
| PF3D7_1359600 | N/A | 11315.7705 | -0.673166399 | 0.40594748 | -1.65825983 | 0.09726503 | 0.65042594 | 14739.1856 | 8233.58356 | 13078.6524 | 9211.66053 | Non-sig. |
| PF3D7_0103100 | VPS51 | 1440.41243 | -0.673709263 | 0.45620185 | -1.47677889 | 0.13973492 | 0.73371218 | 1490.16431 | 882.77157 | 2051.1947 | 1337.51913 | Non-sig. |
| PF3D7_1357000 | N/A | 18123.3315 | -0.674219294 | 0.5910872 | -1.14064269 | 0.25401865 | 0.85030738 | 9428.79059 | 13723.6828 | 35136.3784 | 14204.4741 | Non-sig. |
| PF3D7_0920800 | IMPDH | 643.361887 | -0.674553655 | 0.56624956 | -1.19126566 | 0.23354931 | 0.83828834 | 396.786319 | 523.634919 | 1185.10092 | 467.925386 | Non-sig. |
| PF3D7_0623300 | N/A | 1177.84479 | -0.675318032 | 0.47157451 | -1.43204947 | 0.15212966 | 0.75391062 | 1071.7942 | 1039.15915 | 1825.17738 | 775.24842 | Non-sig. |
| PF3D7_1030600 | KAE1 | 566.846012 | -0.676355525 | 0.44981923 | -1.50361628 | 0.1326802 | 0.72037933 | 829.028379 | 430.329949 | 565.819766 | 442.205953 | Non-sig. |
| PF3D7_0709300 | MED14 | 1716.76468 | -0.676460421 | 0.48854358 | -1.38464703 | 0.16616042 | 0.77284241 | 1461.09637 | 1095.05865 | 2762.72709 | 1548.1766 | Non-sig. |
| PF3D7_0608300 | N/A | 460.19884 | -0.676537385 | 0.43569183 | -1.55278877 | 0.12047361 | 0.70293962 | 613.262696 | 341.108745 | 519.15228 | 367.27164 | Non-sig. |
| PF3D7_0610400 | H3 | 67202.4888 | -0.676760729 | 0.49813446 | -1.35859046 | 0.1742764 | 0.78513297 | 53926.0845 | 42324.2697 | 111437.43 | 61122.1705 | Non-sig. |
| PF3D7_1339500 | N/A | 49.6621274 | -0.676767588 | 0.68452717 | -0.98866431 | 0.32282741 | 0.88694295 | 44.9820054 | 49.4437112 | 77.1235401 | 27.099253 | Non-sig. |
| PF3D7_1036700 | PhLP2 | 247.73151 | -0.677004846 | 0.52384664 | -1.29237221 | 0.19622827 | 0.81437577 | 394.751668 | 225.492099 | 215.080442 | 155.601829 | Non-sig. |
| PF3D7_1145500 | ABCB3 | 642.626604 | -0.67722646 | 0.46627372 | -1.45242255 | 0.14638415 | 0.7426703 | 599.003037 | 535.402988 | 982.285895 | 453.814497 | Non-sig. |
| PF3D7_1027900 | DHHC10 | 66.5935619 | -0.677827383 | 0.61193279 | -1.10768272 | 0.26799886 | 0.8556331 | 59.5883861 | 49.4524606 | 104.188107 | 53.1452938 | Non-sig. |
| PF3D7_1434000 | CAF16 | 435.726376 | -0.678537841 | 0.44876477 | -1.51201226 | 0.13053074 | 0.7188689 | 460.299563 | 357.695425 | 612.265241 | 312.645276 | Non-sig. |
| PF3D7_1214500 | N/A | 32.1931494 | -0.678708008 | 0.72335795 | -0.93827407 | 0.34810358 | 0.8930363 | 35.7092779 | 22.5467901 | 43.5115672 | 27.0049623 | Non-sig. |
| PF3D7_1414900 | N/A | 148.397366 | -0.67877583 | 0.56478592 | -1.20182853 | 0.22942997 | 0.83828834 | 220.089166 | 76.1728892 | 145.379668 | 151.947741 | Non-sig. |
| PF3D7_1455400 | HlyIII | 488.580278 | -0.678846102 | 0.54962437 | -1.23510917 | 0.21678988 | 0.83527883 | 329.888482 | 352.128568 | 872.742094 | 399.561969 | Non-sig. |
| PF3D7_0729600 | CMC1 | 825.54383 | -0.68014351 | 0.49244065 | -1.38116849 | 0.16722716 | 0.77512642 | 1267.68222 | 788.429659 | 765.744428 | 480.319011 | Non-sig. |
| PF3D7_0413700 | N/A | 2586.98042 | -0.680714774 | 0.42797062 | -1.59056428 | 0.11170767 | 0.68406239 | 2635.87102 | 2009.05205 | 3736.39485 | 1966.60377 | Non-sig. |
| PF3D7_1334200 | N/A | 756.549547 | -0.682260891 | 0.53893556 | -1.26594151 | 0.20553402 | 0.82391338 | 1341.51263 | 637.42927 | 523.125114 | 524.131178 | Non-sig. |
| PF3D7_1360200 | EMC3 | 541.514053 | -0.68293557 | 0.44253469 | -1.54323624 | 0.12277347 | 0.7079393 | 760.28628 | 387.75612 | 574.526379 | 443.487432 | Non-sig. |
| PF3D7_1470100 | N/A | 1163.09343 | -0.6840023 | 0.57528585 | -1.18897814 | 0.23444827 | 0.83828834 | 674.643567 | 832.298929 | 2192.58279 | 952.848422 | Non-sig. |
| PF3D7_0531600 | N/A | 3280.29746 | -0.685094761 | 0.42741688 | -1.60287248 | 0.10896281 | 0.6779986 | 3624.94313 | 2158.34466 | 4464.63674 | 2873.26531 | Non-sig. |
| PF3D7_1319500 | N/A | 28.2652741 | -0.6860108 | 0.75315458 | -0.91084993 | 0.36237445 | 0.89567376 | 35.3112539 | 23.6935846 | 34.4189591 | 19.637299 | Non-sig. |
| PF3D7_1449200 | N/A | 12918.1157 | -0.687046508 | 0.42753418 | -1.60699786 | 0.10805484 | 0.67668385 | 13197.6117 | 9156.26798 | 18676.6893 | 10641.8939 | Non-sig. |
| PF3D7_1336800 | NUDC | 1125.5077 | -0.687279957 | 0.50016061 | -1.37411851 | 0.16940489 | 0.77841034 | 1882.47431 | 951.979983 | 895.068179 | 772.508343 | Non-sig. |
| PF3D7_1023300 | N/A | 309.482804 | -0.689757644 | 0.46488592 | -1.48371377 | 0.13788487 | 0.72905594 | 398.91678 | 278.113648 | 365.302839 | 195.597947 | Non-sig. |
| PF3D7_1206500 | N/A | 265.698465 | -0.689892136 | 0.63240878 | -1.09089589 | 0.2753187 | 0.85802262 | 516.739844 | 233.922648 | 139.615481 | 172.515886 | Non-sig. |
| PF3D7_1115200 | SET7 | 4295.74634 | -0.69063038 | 0.41810586 | -1.65180748 | 0.0985738 | 0.65401834 | 5913.571 | 3022.9161 | 4696.06279 | 3550.43546 | Non-sig. |
| PF3D7_0718500 | N/A | 1430.98802 | -0.691838908 | 0.44978565 | -1.53815246 | 0.12401134 | 0.708883 | 2093.08348 | 1259.41792 | 1442.43612 | 929.014553 | Non-sig. |
| PF3D7_1311000 | ISD11 | 64.3906076 | -0.691860541 | 0.59218861 | -1.16831112 | 0.24268129 | 0.8422984 | 85.3440686 | 55.9705255 | 73.7816587 | 42.4661774 | Non-sig. |
| PF3D7_0217800 | RPS26 | 3982.54168 | -0.693242303 | 0.47115504 | -1.47136768 | 0.14119171 | 0.73697898 | 3589.07537 | 3543.74146 | 6253.49518 | 2543.85473 | Non-sig. |
| PF3D7_1436400 | N/A | 165.388559 | -0.694751079 | 0.5464611 | -1.27136421 | 0.20359911 | 0.82152939 | 159.067318 | 91.1839333 | 249.70684 | 161.596145 | Non-sig. |

| | | | | | | | | | | | | |
|---|---|---|---|---|---|---|---|---|---|---|---|---|
| PF3D7_0720800 | N/A | 669.334303 | -0.694895837 | 0.42223318 | -1.64576323 | 0.09981251 | 0.65652696 | 859.028151 | 527.542968 | 795.984808 | 494.781286 | Non-sig. |
| PF3D7_1432100 | N/A | 439.994306 | -0.695452975 | 0.45429884 | -1.53082712 | 0.12581213 | 0.71290716 | 634.353612 | 310.432366 | 453.86418 | 361.327067 | Non-sig. |
| PF3D7_0813300 | N/A | 2986.59687 | -0.695731987 | 0.44230076 | -1.57298393 | 0.11572254 | 0.69120407 | 4318.32943 | 2629.26701 | 3068.02333 | 1930.76769 | Non-sig. |
| PF3D7_1246700 | N/A | 772.705655 | -0.697426194 | 0.4465636 | -1.56176228 | 0.118344 | 0.697844 | 1057.94267 | 690.110297 | 854.005915 | 488.763733 | Non-sig. |
| PF3D7_1204700 | N/A | 396.335474 | -0.697840017 | 0.4662725 | -1.49663558 | 0.13448811 | 0.72190273 | 414.068499 | 348.876633 | 566.534127 | 255.862635 | Non-sig. |
| PF3D7_1433600 | SPC1 | 636.619933 | -0.698458129 | 0.43389847 | -1.60972711 | 0.10745744 | 0.67533753 | 765.75059 | 547.103268 | 809.798187 | 423.827687 | Non-sig. |
| PF3D7_0928600 | N/A | 220.774945 | -0.698477564 | 0.51111701 | -1.36657076 | 0.17175989 | 0.78315911 | 195.570589 | 158.85099 | 350.624711 | 178.05349 | Non-sig. |
| PF3D7_1244800 | N/A | 1167.24441 | -0.698628182 | 0.52442645 | -1.33217571 | 0.18280246 | 0.80052251 | 2014.61479 | 1046.32698 | 874.591013 | 733.44485 | Non-sig. |
| PF3D7_0821200 | N/A | 589.322935 | -0.698932954 | 0.4375361 | -1.59742922 | 0.11017006 | 0.68028088 | 726.609665 | 512.377209 | 732.089534 | 386.215333 | Non-sig. |
| PF3D7_1207600 | MiaA | 647.523601 | -0.69898651 | 0.43246742 | -1.61627554 | 0.10603475 | 0.66957345 | 888.366185 | 522.550309 | 714.519757 | 464.658152 | Non-sig. |
| PF3D7_0629700 | SET1 | 22734.4076 | -0.700766693 | 0.49922838 | -1.40369964 | 0.16040831 | 0.7667698 | 20724.0469 | 12182.6246 | 35575.3655 | 22455.5933 | Non-sig. |
| PF3D7_1465800 | N/A | 216.402499 | -0.700956322 | 0.52984311 | -1.32295071 | 0.18585177 | 0.80052251 | 182.441943 | 143.70455 | 353.270931 | 186.192571 | Non-sig. |
| PF3D7_0213400 | PK7 | 619.242218 | -0.701605423 | 0.45565481 | -1.53977398 | 0.12361546 | 0.70887094 | 608.129986 | 504.368453 | 925.523081 | 438.947354 | Non-sig. |
| PF3D7_0415100 | N/A | 240.211382 | -0.702112987 | 0.47156855 | -1.4888885 | 0.13651673 | 0.72522545 | 253.366328 | 191.057778 | 341.584121 | 174.837302 | Non-sig. |
| PF3D7_0714100 | N/A | 34.7580007 | -0.702803727 | 0.86463914 | -0.81282895 | 0.41631614 | 0.90793155 | 71.5052019 | 23.6124023 | 14.7806603 | 29.1337384 | Non-sig. |
| PF3D7_0618300 | N/A | 5120.76069 | -0.703398379 | 0.47183429 | -1.4907742 | 0.13602079 | 0.72505481 | 4581.63748 | 4515.32233 | 8108.02783 | 3278.05513 | Non-sig. |
| PF3D7_0203700 | MAK16 | 710.911219 | -0.704302714 | 0.49805162 | -1.41411591 | 0.15732787 | 0.75912848 | 1177.74077 | 593.528793 | 584.657927 | 487.717387 | Non-sig. |
| PF3D7_1447300 | N/A | 239.952115 | -0.704438799 | 0.49003214 | -1.4375359 | 0.15056578 | 0.75169515 | 254.887699 | 216.957075 | 339.805619 | 148.158067 | Non-sig. |
| PF3D7_1469700 | MED6 | 477.650192 | -0.704781334 | 0.43978565 | -1.60255644 | 0.10903262 | 0.6779986 | 533.760501 | 390.765398 | 650.258124 | 335.816745 | Non-sig. |
| PF3D7_0810500 | PPM7 | 644.434501 | -0.705485917 | 0.43426809 | -1.62454008 | 0.10426059 | 0.6615241 | 818.855129 | 554.074772 | 779.043438 | 425.764663 | Non-sig. |
| PF3D7_1441900 | TFB5 | 346.221453 | -0.705780592 | 0.4519999 | -1.56146181 | 0.11841483 | 0.697844 | 445.835645 | 297.697447 | 412.72471 | 228.628012 | Non-sig. |
| PF3D7_0210400 | N/A | 552.366556 | -0.706092891 | 0.46121517 | -1.53094031 | 0.12578415 | 0.71290716 | 537.510147 | 453.385444 | 832.117066 | 386.453565 | Non-sig. |
| PF3D7_0521000 | N/A | 907.062514 | -0.706504706 | 0.59398473 | -1.18943244 | 0.23426954 | 0.83828834 | 1736.64214 | 810.126094 | 513.299634 | 568.182193 | Non-sig. |
| PF3D7_0530800 | N/A | 114.166386 | -0.707335011 | 0.51116156 | -1.38377974 | 0.16642591 | 0.77284241 | 139.556267 | 89.1367988 | 143.646697 | 84.325779 | Non-sig. |
| PF3D7_1317400 | N/A | 557.05066 | -0.707457534 | 0.44896196 | -1.57576275 | 0.11508049 | 0.68970185 | 619.239367 | 488.607128 | 762.59246 | 357.763683 | Non-sig. |
| PF3D7_1022300 | ZIPCO | 130.163614 | -0.708138485 | 0.59442493 | -1.1913001 | 0.23353579 | 0.83828834 | 93.629262 | 109.462016 | 229.118057 | 88.4451199 | Non-sig. |
| PF3D7_1201700 | N/A | 192.296116 | -0.708342638 | 0.51864287 | -1.36576184 | 0.17201373 | 0.78315911 | 305.749941 | 135.265761 | 171.595246 | 156.573517 | Non-sig. |
| PF3D7_0415600 | GAK | 133.285493 | -0.709930654 | 0.55649255 | -1.27572356 | 0.20205326 | 0.81948076 | 112.320682 | 110.602807 | 218.358782 | 91.8596993 | Non-sig. |
| PF3D7_1323000 | FabZ | 99.9035045 | -0.710714411 | 0.53636463 | -1.32505831 | 0.18515181 | 0.80052251 | 140.713792 | 71.0117139 | 107.41783 | 80.4706821 | Non-sig. |
| PF3D7_0913400 | N/A | 100.172002 | -0.71071776 | 0.57519089 | -1.23562067 | 0.2165996 | 0.83527883 | 156.318721 | 90.5522967 | 92.5461355 | 61.2708568 | Non-sig. |
| PF3D7_1413400 | N/A | 2169.66526 | -0.710773583 | 0.44306739 | -1.60421101 | 0.10866755 | 0.6779986 | 3296.60246 | 1651.6464 | 2090.7594 | 1639.65277 | Non-sig. |
| PF3D7_0212200 | N/A | 207.206041 | -0.711097656 | 0.48667334 | -1.46113952 | 0.14397716 | 0.7412338 | 295.407138 | 176.03272 | 219.236363 | 138.147944 | Non-sig. |
| PF3D7_1311100 | N/A | 78.1856265 | -0.711482049 | 0.99062703 | -0.71821385 | 0.47262544 | 0.92682482 | 186.951605 | 68.7137139 | 7.38877778 | 49.688409 | Non-sig. |
| PF3D7_0307200 | N/A | 5265.51462 | -0.712089333 | 0.4277121 | -1.66488003 | 0.09593669 | 0.64772846 | 5539.76835 | 4393.02903 | 7538.57573 | 3590.68539 | Non-sig. |
| PF3D7_0111700 | N/A | 22.0183878 | -0.712106721 | 0.86181188 | -0.82629021 | 0.40863948 | 0.90748263 | 20.9758451 | 11.8148995 | 33.6515199 | 21.6312865 | Non-sig. |
| PF3D7_1132900 | GCVH | 310.243181 | -0.7131094 | 0.45974962 | -1.55108207 | 0.12088202 | 0.70312626 | 433.387817 | 256.49152 | 337.512268 | 213.581118 | Non-sig. |
| PF3D7_0306800 | CCT2 | 1964.86355 | -0.713640057 | 0.63656015 | -1.12108818 | 0.26225033 | 0.85030738 | 863.779814 | 1575.35543 | 4018.25074 | 1402.06821 | Non-sig. |
| PF3D7_1312700 | N/A | 1572.48905 | -0.714065165 | 0.42793772 | -1.66861934 | 0.09519284 | 0.64740909 | 2244.00799 | 1260.78344 | 1663.91933 | 1121.24543 | Non-sig. |
| PF3D7_1023200 | OMPDC | 372.221227 | -0.714110433 | 0.45165267 | -1.58110531 | 0.11385396 | 0.68623241 | 433.979384 | 319.904815 | 490.984941 | 244.015769 | Non-sig. |
| PF3D7_1451100 | eEF2 | 13573.1927 | -0.714304381 | 0.68007695 | -1.05032876 | 0.29356699 | 0.87149079 | 4731.03902 | 10110.6271 | 29001.424 | 10449.6808 | Non-sig. |
| PF3D7_0719600 | N/A | 3624.75297 | -0.714469121 | 0.45320535 | -1.57647988 | 0.11491525 | 0.68949152 | 3343.48285 | 2887.56578 | 5665.08391 | 2602.87933 | Non-sig. |
| PF3D7_1131100 | SR1 | 3031.16619 | -0.71526021 | 0.46783068 | -1.52888692 | 0.12629248 | 0.71393062 | 3129.49859 | 1711.39917 | 4405.43755 | 2878.32945 | Non-sig. |
| PF3D7_1108900 | N/A | 113.654688 | -0.715411882 | 0.54483047 | -1.31309081 | 0.18915235 | 0.80547152 | 108.310824 | 78.3499443 | 174.083747 | 93.8742367 | Non-sig. |
| PF3D7_1465700 | PM8 | 12.1086072 | -0.715522049 | 1.08556924 | -0.65912152 | 0.50981774 | 0.93247207 | 17.8242566 | 12.8900379 | 12.3108954 | 5.40923899 | Non-sig. |
| PF3D7_0422600 | N/A | 59.8741928 | -0.715745971 | 0.59556999 | -1.20178313 | 0.22944756 | 0.83828834 | 70.06854 | 47.2976399 | 78.7610858 | 43.3695055 | Non-sig. |
| PF3D7_0615300 | GWT1 | 638.909238 | -0.715770515 | 0.42589934 | -1.6806096 | 0.09283877 | 0.64025761 | 846.103172 | 463.560097 | 742.418923 | 503.55476 | Non-sig. |
| PF3D7_1006300 | N/A | 5.77222522 | -0.717018728 | 1.47685387 | -0.48550418 | 0.62731875 | 0.96655174 | 10.3036619 | 5.39876583 | 4.08990898 | 3.29656413 | Non-sig. |
| PF3D7_0614400 | CWF7 | 901.342642 | -0.719288375 | 0.422923 | -1.70075494 | 0.08898901 | 0.62981332 | 1043.71109 | 630.81808 | 1199.20307 | 731.638327 | Non-sig. |
| PF3D7_1009600 | PPM10 | 213.832893 | -0.720568669 | 0.5015666 | -1.43663606 | 0.15082143 | 0.75169515 | 248.133358 | 121.286247 | 284.110762 | 201.801206 | Non-sig. |
| PF3D7_1119200 | N/A | 95.541303 | -0.720587684 | 0.65395296 | -1.10189529 | 0.2705072 | 0.8556331 | 161.653834 | 100.93918 | 76.3431536 | 43.2290447 | Non-sig. |
| PF3D7_1212600 | N/A | 233.51796 | -0.721530337 | 0.47964488 | -1.50430113 | 0.13250385 | 0.72037933 | 249.040104 | 197.75863 | 332.303724 | 154.969381 | Non-sig. |
| PF3D7_0706200 | N/A | 96.202884 | -0.722109389 | 0.64194109 | -1.1248842 | 0.26063811 | 0.85030738 | 78.5899544 | 99.9143464 | 160.845535 | 45.4617005 | Non-sig. |
| PF3D7_1366500 | NDK | 1571.56704 | -0.722152484 | 0.55532098 | -1.30042357 | 0.19345584 | 0.81267745 | 1005.62804 | 1328.60461 | 2907.86117 | 1044.17435 | Non-sig. |
| PF3D7_1135000 | N/A | 172.420568 | -0.722205138 | 0.53624371 | -1.34678528 | 0.1780494 | 0.79202829 | 153.364383 | 151.378787 | 275.854682 | 109.08442 | Non-sig. |

| | | | | | | | | | | | | |
|---|---|---|---|---|---|---|---|---|---|---|---|---|
| PF3D7_1324200 | N/A | 1452.59208 | -0.722605275 | 0.62759314 | -1.1513913 | 0.24957129 | 0.84867055 | 2930.05818 | 1254.62327 | 688.14029 | 937.546574 | Non-sig. |
| PF3D7_1314500 | N/A | 935.403718 | -0.722729748 | 0.41773201 | -1.73012778 | 0.08360745 | 0.61468103 | 1140.29308 | 749.468306 | 1189.53415 | 662.319332 | Non-sig. |
| PF3D7_1238100 | N/A | 1272.54984 | -0.722904656 | 0.41841432 | -1.72772446 | 0.08403764 | 0.61549563 | 1494.83574 | 1040.31518 | 1674.84347 | 880.204979 | Non-sig. |
| PF3D7_1331800 | N/A | 4007.37539 | -0.723362537 | 0.48809507 | -1.48201156 | 0.13833722 | 0.72945925 | 3469.31995 | 3610.47312 | 6513.43186 | 2436.27664 | Non-sig. |
| PF3D7_0801600 | N/A | 58.1738778 | -0.723882052 | 0.63294688 | -1.14366951 | 0.25276072 | 0.85030738 | 91.7537426 | 39.8060767 | 53.3021516 | 47.8335403 | Non-sig. |
| PF3D7_0114100 | MC-2TM | 110.818438 | -0.725261668 | 0.5178708 | -1.40046837 | 0.16137311 | 0.76690962 | 149.022088 | 80.541939 | 127.23327 | 86.476455 | Non-sig. |
| PF3D7_0202300 | N/A | 61.4247556 | -0.726998197 | 0.67523232 | -1.07666381 | 0.28163049 | 0.86390383 | 43.0026617 | 42.9626604 | 109.984001 | 49.7496992 | Non-sig. |
| PF3D7_0719700 | RPS10 | 4839.45036 | -0.728194811 | 0.41359411 | -1.76065084 | 0.07829752 | 0.60381404 | 6228.37323 | 4061.23783 | 5842.69149 | 3225.49888 | Non-sig. |
| PF3D7_1371900 | N/A | 75.8917834 | -0.72864821 | 0.56068195 | -1.29957494 | 0.19374669 | 0.81267745 | 93.2921046 | 61.2319332 | 96.0207743 | 53.0223214 | Non-sig. |
| PF3D7_0306900 | N/A | 5614.3018 | -0.728732937 | 0.50014288 | -1.4570495 | 0.14510272 | 0.74131203 | 4495.35214 | 4899.25484 | 9510.09811 | 3552.50213 | Non-sig. |
| PF3D7_0111900 | N/A | 641.95094 | -0.728910861 | 0.42199023 | -1.7273169 | 0.08411077 | 0.61549563 | 803.796274 | 497.81453 | 797.726516 | 468.46644 | Non-sig. |
| PF3D7_1350300 | N/A | 354.225603 | -0.729446326 | 0.515176 | -1.41591675 | 0.15679988 | 0.75894163 | 586.046528 | 304.919333 | 298.012673 | 227.923879 | Non-sig. |
| PF3D7_1305600 | S2P | 800.528375 | -0.73033774 | 0.44410359 | -1.64452112 | 0.10006861 | 0.65652696 | 818.664824 | 643.009507 | 1179.04436 | 561.394813 | Non-sig. |
| PF3D7_1323500 | PMV | 1257.57518 | -0.730818526 | 0.56516987 | -1.29309535 | 0.19597808 | 0.81434658 | 780.448071 | 829.279456 | 2358.17706 | 1062.39613 | Non-sig. |
| PF3D7_1435400 | EMC4 | 820.768811 | -0.730981361 | 0.59678919 | -1.22485692 | 0.22062914 | 0.83527883 | 446.587811 | 658.059706 | 1601.85817 | 576.569553 | Non-sig. |
| PF3D7_0630800 | N/A | 53.7955197 | -0.731711455 | 0.8933359 | -0.81907763 | 0.41274212 | 0.90786241 | 119.710444 | 53.6854351 | 14.7770645 | 27.0091354 | Non-sig. |
| PF3D7_1434500 | N/A | 3460.54486 | -0.732262042 | 0.49139496 | -1.49017003 | 0.13617953 | 0.7251047 | 3621.68077 | 1741.57799 | 5018.97472 | 3459.94594 | Non-sig. |
| PF3D7_1462000 | TLP1 | 5.08158933 | -0.732674413 | 1.51981216 | -0.48208222 | 0.62974754 | 0.96682409 | 6.15996623 | 4.32905162 | 6.5290826 | 3.30825688 | Non-sig. |
| PF3D7_0829200 | PHB1 | 337.739362 | -0.734214475 | 0.44532786 | -1.64870546 | 0.09920799 | 0.65421395 | 386.406107 | 255.547796 | 457.2623 | 251.741244 | Non-sig. |
| PF3D7_1139900 | EMC8 | 237.481985 | -0.735330867 | 0.58400464 | -1.2591182 | 0.20798764 | 0.82398515 | 178.615355 | 124.530667 | 414.619502 | 232.162417 | Non-sig. |
| PF3D7_1465900 | N/A | 5854.84829 | -0.736798504 | 0.45674331 | -1.61315663 | 0.10671048 | 0.67223797 | 5469.89328 | 4901.75288 | 9166.38583 | 3881.36117 | Non-sig. |
| PF3D7_1343100 | N/A | 5489.12517 | -0.736820245 | 0.54102645 | -1.3618932 | 0.17323159 | 0.78315911 | 9872.3517 | 3255.44709 | 3850.2163 | 4978.4856 | Non-sig. |
| PF3D7_0213800 | N/A | 75.4236912 | -0.73810564 | 0.56664037 | -1.30259981 | 0.19271142 | 0.81094926 | 104.096076 | 52.5922659 | 84.5772135 | 60.4292093 | Non-sig. |
| PF3D7_0501700 | APC3 | 48.8011768 | -0.739862776 | 0.63340364 | -1.16807471 | 0.24277663 | 0.8422984 | 54.7068533 | 36.4869417 | 67.3398131 | 36.671099 | Non-sig. |
| PF3D7_1325700 | N/A | 294.294212 | -0.740403992 | 0.5079172 | -1.45772578 | 0.14491614 | 0.74131203 | 264.014605 | 253.276102 | 472.183888 | 187.702254 | Non-sig. |
| PF3D7_1204500 | N/A | 672.802105 | -0.740698299 | 0.44087599 | -1.68006041 | 0.09294556 | 0.64025761 | 732.018597 | 557.232429 | 951.502727 | 450.454668 | Non-sig. |
| PF3D7_1320400 | SPB | 358.808854 | -0.740708468 | 0.47271966 | -1.56690852 | 0.11713608 | 0.69494617 | 362.436425 | 235.113998 | 535.298346 | 302.386648 | Non-sig. |
| PF3D7_0516300 | N/A | 313.000182 | -0.741413332 | 0.53011206 | -1.39859736 | 0.16193376 | 0.76806182 | 244.340142 | 218.855225 | 538.820051 | 249.985309 | Non-sig. |
| PF3D7_1008200 | N/A | 279.940628 | -0.74182677 | 0.47840514 | -1.55062459 | 0.12099168 | 0.70312626 | 324.642742 | 166.401174 | 376.027705 | 252.69089 | Non-sig. |
| PF3D7_1316100 | IPK2 | 2122.54174 | -0.741901658 | 0.48959994 | -1.51532219 | 0.12969083 | 0.71812182 | 3325.45973 | 1178.87002 | 1987.88616 | 1997.95104 | Non-sig. |
| PF3D7_1223700 | VIT | 1346.02044 | -0.742097382 | 0.51395688 | -1.44389035 | 0.14876982 | 0.74754886 | 1002.85119 | 1029.37974 | 2366.41357 | 985.437255 | Non-sig. |
| PF3D7_1448100 | N/A | 90.1098336 | -0.742125034 | 0.54835974 | -1.35335434 | 0.17594246 | 0.78744529 | 120.56278 | 58.0069336 | 105.050959 | 76.8186608 | Non-sig. |
| PF3D7_0522800 | BUD31 | 280.531091 | -0.743194958 | 0.46240392 | -1.60724189 | 0.10800132 | 0.67668385 | 343.646738 | 240.458298 | 358.803279 | 179.216048 | Non-sig. |
| PF3D7_1309000 | N/A | 488.393628 | -0.743763524 | 0.44804803 | -1.66000848 | 0.09691275 | 0.64910712 | 510.188397 | 358.442046 | 712.802321 | 372.141751 | Non-sig. |
| PF3D7_0825600 | COA5 | 99.7485201 | -0.7441687 | 0.58975243 | -1.26183236 | 0.20700912 | 0.82391338 | 167.165123 | 83.868159 | 82.8507755 | 65.1100232 | Non-sig. |
| PF3D7_0501000 | N/A | 1989.52919 | -0.745828671 | 0.48681121 | -1.53206964 | 0.12550525 | 0.71290716 | 1630.10779 | 1509.26611 | 3354.90604 | 1463.83682 | Non-sig. |
| PF3D7_1347300 | N/A | 190.328521 | -0.746224095 | 0.62237534 | -1.19899368 | 0.2305304 | 0.83828834 | 109.034909 | 142.867379 | 367.680876 | 141.730921 | Non-sig. |
| PF3D7_1247100 | N/A | 489.718839 | -0.746295548 | 0.43840133 | -1.70231133 | 0.08869702 | 0.62981332 | 617.175728 | 410.012555 | 610.098528 | 321.588546 | Non-sig. |
| PF3D7_1434700 | TIM17 | 217.637616 | -0.747537539 | 0.49685754 | -1.50453095 | 0.13244471 | 0.72037933 | 221.906563 | 137.376007 | 323.539929 | 187.727966 | Non-sig. |
| PF3D7_0915100 | UBC9 | 714.660371 | -0.748008691 | 0.42419827 | -1.76334689 | 0.077842 | 0.60344518 | 850.864897 | 567.937411 | 940.863002 | 498.976174 | Non-sig. |
| PF3D7_0806900 | N/A | 322.504928 | -0.748940781 | 0.47089042 | -1.59047784 | 0.11172714 | 0.68406239 | 374.321268 | 193.196661 | 434.381182 | 288.120602 | Non-sig. |
| PF3D7_0406200 | Pfs16 | 9015.32219 | -0.749046578 | 0.41744486 | -1.79436053 | 0.07275564 | 0.58363337 | 12921.7576 | 6600.38789 | 9687.40241 | 6851.74089 | Non-sig. |
| PF3D7_1339300 | N/A | 6401.32295 | -0.750546587 | 0.47337581 | -1.58551953 | 0.11284834 | 0.68406239 | 10332.0107 | 4039.63564 | 5727.95684 | 5505.6886 | Non-sig. |
| PF3D7_0220600 | N/A | 504.861784 | -0.751280822 | 0.50749097 | -1.48038263 | 0.13877116 | 0.73009893 | 408.364362 | 389.506928 | 858.233437 | 363.342408 | Non-sig. |
| PF3D7_0304900 | N/A | 250.149579 | -0.752366897 | 0.45643586 | -1.64835187 | 0.09928049 | 0.65421395 | 301.147735 | 195.456595 | 326.693631 | 177.300356 | Non-sig. |
| PF3D7_0715500 | N/A | 249.003878 | -0.75348639 | 0.5580405 | -1.35023604 | 0.17694028 | 0.79022251 | 185.17788 | 212.816366 | 439.776158 | 158.245108 | Non-sig. |
| PF3D7_1471100 | EXP2 | 7093.06399 | -0.755444792 | 0.43140426 | -1.75112965 | 0.07992358 | 0.60514927 | 7270.45903 | 4882.07142 | 10547.0654 | 5672.6601 | Non-sig. |
| PF3D7_0608700 | CCT6 | 2510.90603 | -0.755926154 | 0.50595832 | -1.49404827 | 0.135163 | 0.72405824 | 1925.09522 | 2007.2858 | 4382.7869 | 1728.45621 | Non-sig. |
| PF3D7_1128100 | N/A | 1578.05427 | -0.756346581 | 0.45034042 | -1.67949966 | 0.0930547 | 0.64025761 | 2123.33486 | 1444.80079 | 1841.7143 | 902.367113 | Non-sig. |
| PF3D7_1204600 | N/A | 167.246858 | -0.757524165 | 0.50559727 | -1.49827582 | 0.1340616 | 0.72178914 | 192.087171 | 149.262865 | 228.206898 | 99.4304973 | Non-sig. |
| PF3D7_1233100 | N/A | 538.156604 | -0.758094282 | 0.47922134 | -1.58192931 | 0.11366571 | 0.68587805 | 866.272702 | 415.673643 | 486.712937 | 383.967134 | Non-sig. |
| PF3D7_0417900 | N/A | 79.8022688 | -0.759255595 | 0.69039179 | -1.09974597 | 0.27144282 | 0.8556331 | 155.703353 | 60.0962226 | 45.1669089 | 58.2425902 | Non-sig. |
| PF3D7_0112000 | TatD | 364.090834 | -0.760169378 | 0.4416747 | -1.7211069 | 0.08523143 | 0.61879091 | 423.088384 | 272.739838 | 492.548618 | 267.986495 | Non-sig. |
| PF3D7_1225400 | N/A | 117.741508 | -0.760676595 | 0.55734717 | -1.36481647 | 0.17231074 | 0.78315911 | 192.927379 | 88.0811557 | 103.412092 | 86.5454052 | Non-sig. |

| | | | | | | | | | | | | |
|---|---|---|---|---|---|---|---|---|---|---|---|---|
| PF3D7_1010400 | N/A | 149.679386 | -0.761179371 | 0.7186306 | -1.05920813 | 0.28950501 | 0.86724628 | 314.412959 | 136.435709 | 62.3705441 | 85.4983307 | Non-sig. |
| PF3D7_1357100 | N/A | 19270.2753 | -0.761589408 | 0.56909751 | -1.33824062 | 0.18081801 | 0.79699299 | 11279.5575 | 14945.7799 | 37203.5516 | 13652.2124 | Non-sig. |
| PF3D7_1142500 | RPL28 | 1054.15871 | -0.762513043 | 0.43462304 | -1.75442387 | 0.0793579 | 0.60514927 | 1122.34295 | 839.270087 | 1530.37751 | 724.644285 | Non-sig. |
| PF3D7_0727900 | N/A | 634.50017 | -0.763416967 | 0.59145193 | -1.29075066 | 0.19679015 | 0.81437577 | 454.238574 | 291.750234 | 1142.66599 | 649.345882 | Non-sig. |
| PF3D7_0310000 | N/A | 486.828117 | -0.763641968 | 0.43473865 | -1.75655411 | 0.07899384 | 0.60514927 | 581.123993 | 389.874169 | 644.321725 | 331.992581 | Non-sig. |
| PF3D7_1250400 | N/A | 442.865725 | -0.764691151 | 0.61469973 | -1.24400763 | 0.21349676 | 0.83292034 | 879.701581 | 353.044081 | 235.697959 | 303.019281 | Non-sig. |
| PF3D7_1470950 | N/A | 29.5482278 | -0.7653038 | 0.80229051 | -0.95389861 | 0.34013497 | 0.8930363 | 52.5640139 | 23.9171318 | 21.9630486 | 19.7487169 | Non-sig. |
| PF3D7_0715400 | PSOP20 | 432.707895 | -0.765540948 | 0.46367876 | -1.65101579 | 0.09873535 | 0.65401834 | 626.174634 | 371.624695 | 463.740335 | 269.291917 | Non-sig. |
| PF3D7_1337000 | OCT1 | 219.126591 | -0.765648768 | 0.47880543 | -1.59908121 | 0.10980256 | 0.67975706 | 230.612732 | 158.784284 | 321.147732 | 165.961616 | Non-sig. |
| PF3D7_1105200 | WDR8 | 54.5769857 | -0.765683239 | 0.64805756 | -1.18150498 | 0.23740218 | 0.83906386 | 85.0278063 | 48.3216146 | 52.5374514 | 32.4210703 | Non-sig. |
| PF3D7_0319500 | N/A | 1589.20389 | -0.765811351 | 0.41891548 | -1.82808083 | 0.06753742 | 0.56795755 | 1940.85326 | 1309.47243 | 2061.84574 | 1044.64413 | Non-sig. |
| PF3D7_1364000 | N/A | 624.634546 | -0.765894155 | 0.5451726 | -1.40486545 | 0.16006129 | 0.7667698 | 586.660431 | 281.074258 | 986.477619 | 644.325875 | Non-sig. |
| PF3D7_1455900 | PPRD | 252.728399 | -0.765896393 | 0.51684363 | -1.48187257 | 0.13837421 | 0.72945925 | 225.866178 | 215.789559 | 410.504867 | 158.75299 | Non-sig. |
| PF3D7_0522700 | SufA | 174.563546 | -0.766931655 | 0.51955396 | -1.47613475 | 0.13990772 | 0.73389218 | 184.471254 | 158.992164 | 255.230666 | 99.5601004 | Non-sig. |
| PF3D7_1213700 | RPB8 | 341.348668 | -0.76699762 | 0.46903794 | -1.63525708 | 0.10199517 | 0.65895255 | 378.224276 | 296.532158 | 481.703466 | 208.934773 | Non-sig. |
| PF3D7_0323000 | TMA7 | 940.645901 | -0.768210009 | 0.43972657 | -1.74701749 | 0.0806343 | 0.6065657 | 1006.08814 | 770.901254 | 1364.43958 | 621.154629 | Non-sig. |
| PF3D7_0813900 | RPS16 | 5432.71779 | -0.769032823 | 0.48936671 | -1.57148577 | 0.11606986 | 0.6917189 | 4432.8344 | 4359.35064 | 9261.59749 | 3677.08861 | Non-sig. |
| PF3D7_1130200 | PfP0 | 11845.4127 | -0.769610182 | 0.43495756 | -1.76939143 | 0.07682857 | 0.59858498 | 11981.2489 | 9466.72104 | 17882.6718 | 8051.00887 | Non-sig. |
| PF3D7_1450600 | N/A | 236.919741 | -0.771443261 | 0.68005115 | -1.13439005 | 0.25663097 | 0.85030738 | 133.469735 | 97.6852608 | 463.901716 | 252.62225 | Non-sig. |
| PF3D7_1128200 | MBF1 | 528.404736 | -0.771636691 | 0.44180582 | -1.74655165 | 0.08071514 | 0.6065657 | 568.783181 | 395.400511 | 763.966749 | 385.468504 | Non-sig. |
| PF3D7_0728500 | N/A | 11.0621278 | -0.772211519 | 1.09879173 | -0.70278242 | 0.48219135 | 0.92880667 | 16.4858591 | 9.7177785 | 11.4517451 | 6.59312826 | Non-sig. |
| PF3D7_0219100 | N/A | 123.824528 | -0.773011676 | 0.52592052 | -1.46982603 | 0.14160888 | 0.73825055 | 125.107658 | 90.1618011 | 187.214758 | 92.8138931 | Non-sig. |
| PF3D7_1143200 | N/A | 4121.87102 | -0.773168945 | 0.41681365 | -1.85495113 | 0.06360321 | 0.54881073 | 4708.09509 | 3327.37559 | 5693.15458 | 2758.85881 | Non-sig. |
| PF3D7_1309800 | N/A | 527.013883 | -0.77411966 | 0.4363378 | -1.77412928 | 0.07604177 | 0.5968434 | 595.190784 | 409.994238 | 734.93055 | 367.93996 | Non-sig. |
| PF3D7_1111200 | N/A | 370.539804 | -0.775057219 | 0.50763227 | -1.5268084 | 0.12680866 | 0.71411411 | 622.836918 | 289.74657 | 312.885563 | 256.690165 | Non-sig. |
| PF3D7_0716600 | SufS | 178.721677 | -0.775966145 | 0.53735408 | -1.44404997 | 0.14872492 | 0.74754886 | 158.797336 | 110.546288 | 292.333043 | 153.210041 | Non-sig. |
| PF3D7_0906910 | N/A | 336.469317 | -0.777709853 | 0.45575198 | -1.70643219 | 0.08792764 | 0.62914105 | 479.402176 | 266.680944 | 370.759778 | 229.034369 | Non-sig. |
| PF3D7_0320700 | SPC25 | 3546.22512 | -0.778149553 | 0.43505529 | -1.78862222 | 0.07367568 | 0.58628445 | 5224.0173 | 2928.85873 | 3736.26986 | 2295.75459 | Non-sig. |
| PF3D7_1112500 | N/A | 335.525545 | -0.77952941 | 0.44780973 | -1.74076034 | 0.0817256 | 0.60888892 | 412.898575 | 271.609023 | 435.141868 | 222.452716 | Non-sig. |
| PF3D7_0717000 | N/A | 437.019099 | -0.779815247 | 0.52083502 | -1.49724043 | 0.13433071 | 0.72178914 | 341.625659 | 326.314592 | 762.796515 | 317.339631 | Non-sig. |
| PF3D7_1313900 | N/A | 111.168983 | -0.779880387 | 0.63549314 | -1.22720504 | 0.21974553 | 0.83527883 | 82.969929 | 55.8304546 | 197.853801 | 108.021747 | Non-sig. |
| PF3D7_0818000 | SNRNP27 | 1764.40673 | -0.780125628 | 0.43207594 | -1.80552897 | 0.07099195 | 0.58066864 | 2596.82002 | 1202.67617 | 1863.64966 | 1394.48108 | Non-sig. |
| PF3D7_0507100 | RPL4 | 9572.8967 | -0.780638669 | 0.40084789 | -1.94746858 | 0.05147858 | 0.49951196 | 12112.3628 | 7156.27262 | 12090.521 | 6932.43037 | Non-sig. |
| PF3D7_0320000 | I2 | 1394.83505 | -0.780693281 | 0.42589282 | -1.83307454 | 0.06679149 | 0.56527363 | 1769.40886 | 1176.19201 | 1757.17546 | 876.563844 | Non-sig. |
| PF3D7_1452200 | N/A | 92.1346455 | -0.782008867 | 0.556692 | -1.40474241 | 0.16009789 | 0.7667698 | 92.4967138 | 66.5520137 | 140.405776 | 69.0840782 | Non-sig. |
| PF3D7_1253000 | GECO | 3816.65782 | -0.782121106 | 0.41358335 | -1.89108459 | 0.05861305 | 0.52454723 | 4668.34196 | 3115.55311 | 4984.82292 | 2497.91328 | Non-sig. |
| PF3D7_0306200 | AHA1 | 615.294392 | -0.784073585 | 0.45577832 | -1.72029593 | 0.08537866 | 0.61879091 | 743.932882 | 549.423395 | 813.02994 | 354.791353 | Non-sig. |
| PF3D7_1037700 | ERH | 143.690475 | -0.784752592 | 0.50422117 | -1.55636581 | 0.11962113 | 0.70181652 | 154.248607 | 104.171503 | 209.314287 | 107.027502 | Non-sig. |
| PF3D7_0721800 | N/A | 3091.50946 | -0.785561956 | 0.43226692 | -1.81730758 | 0.06917003 | 0.57496879 | 3486.05405 | 2633.23177 | 4339.83678 | 1906.91522 | Non-sig. |
| PF3D7_0715000 | PNPase | 667.595853 | -0.786883541 | 0.51077564 | -1.5405659 | 0.12342247 | 0.70887094 | 527.05159 | 507.889749 | 1163.24667 | 472.195399 | Non-sig. |
| PF3D7_0819000 | N/A | 1803.45594 | -0.787208098 | 0.41540003 | -1.89506029 | 0.05808441 | 0.52246389 | 2510.53624 | 1350.56131 | 2056.82755 | 1295.89867 | Non-sig. |
| PF3D7_0812000 | N/A | 126.418187 | -0.789778641 | 0.50610275 | -1.56051049 | 0.11863929 | 0.69838998 | 174.299581 | 94.4887439 | 146.126442 | 90.7579797 | Non-sig. |
| PF3D7_0527500 | HIP | 5158.73325 | -0.793301152 | 0.45668997 | -1.73706716 | 0.08237532 | 0.60888892 | 8266.7126 | 4088.43761 | 4818.26198 | 3461.52081 | Non-sig. |
| PF3D7_0821700 | N/A | 3126.76433 | -0.793767361 | 0.43817188 | -1.81154336 | 0.07005678 | 0.57813215 | 3173.56889 | 2463.66151 | 4757.99829 | 2111.82863 | Non-sig. |
| PF3D7_0307900 | N/A | 520.125649 | -0.793771949 | 0.47163022 | -1.68303879 | 0.09236758 | 0.63969079 | 804.48954 | 327.106272 | 515.101259 | 433.805526 | Non-sig. |
| PF3D7_1451500 | N/A | 182.022957 | -0.794376074 | 0.48057571 | -1.6529676 | 0.09833746 | 0.65401834 | 247.62773 | 121.355737 | 214.23788 | 144.870481 | Non-sig. |
| PF3D7_1324900 | LDH | 11375.3556 | -0.796031914 | 0.60240186 | -1.32143003 | 0.18635802 | 0.80139998 | 5822.88225 | 8418.06773 | 23049.5729 | 8210.89958 | Non-sig. |
| PF3D7_1413200 | N/A | 246.606211 | -0.796144717 | 0.89437139 | -0.89017239 | 0.37337333 | 0.89654416 | 594.125119 | 224.317714 | 32.0247735 | 135.957238 | Non-sig. |
| PF3D7_1309500 | GAR1 | 390.995976 | -0.796161333 | 0.44232907 | -1.79992995 | 0.0718717 | 0.58133781 | 468.591139 | 310.255266 | 523.814062 | 261.323436 | Non-sig. |
| PF3D7_1010700 | N/A | 1769.70368 | -0.796867662 | 0.42297492 | -1.8839596 | 0.05957044 | 0.52776621 | 2257.17417 | 1475.94884 | 2235.62085 | 1110.07085 | Non-sig. |
| PF3D7_1209800 | ATP11 | 98.8326409 | -0.797393349 | 0.53245413 | -1.49758131 | 0.13424206 | 0.72178914 | 112.175804 | 73.0457683 | 138.700095 | 71.408896 | Non-sig. |
| PF3D7_1303100 | N/A | 715.530479 | -0.798494359 | 0.44982984 | -1.77510314 | 0.07588086 | 0.59646408 | 831.510911 | 621.931375 | 985.705308 | 422.974321 | Non-sig. |
| PF3D7_1242700 | N/A | 4292.68999 | -0.799401344 | 0.43685782 | -1.82988906 | 0.06726653 | 0.56748101 | 4543.94387 | 3554.24032 | 6360.84103 | 2711.73473 | Non-sig. |
| PF3D7_0931200 | Sel2 | 236.77723 | -0.799514835 | 0.57208056 | -1.39755638 | 0.16224632 | 0.76885602 | 163.739532 | 169.667712 | 437.530116 | 176.17156 | Non-sig. |

| | | | | | | | | | | | | |
|---|---|---|---|---|---|---|---|---|---|---|---|---|
| PF3D7_1356200 | TIM23 | 623.520103 | -0.800020985 | 0.43411486 | -1.84287858 | 0.06534674 | 0.5620244 | 786.28597 | 513.275382 | 797.919909 | 396.599152 | Non-sig. |
| PF3D7_0822100 | MED7 | 107.656421 | -0.800542551 | 0.55721755 | -1.43667864 | 0.15080933 | 0.75169515 | 101.846379 | 86.9734029 | 171.574428 | 70.2314732 | Non-sig. |
| PF3D7_0522000 | N/A | 604.167013 | -0.801640311 | 0.57180028 | -1.40195858 | 0.16092762 | 0.7667698 | 1127.9653 | 540.194119 | 407.954891 | 340.553742 | Non-sig. |
| PF3D7_1105400 | RPS4 | 7321.44934 | -0.802096015 | 0.49176616 | -1.63105167 | 0.10287942 | 0.66002937 | 5910.21971 | 5632.15389 | 12701.2266 | 5042.19719 | Non-sig. |
| PF3D7_1234400 | MiGS | 22.1483962 | -0.802160964 | 0.82176883 | -0.97613944 | 0.32899537 | 0.88909228 | 28.3878225 | 16.0999947 | 27.9183441 | 16.1874236 | Non-sig. |
| PF3D7_MIT02200 | SSUA | 37.2636868 | -0.803176693 | 0.6873084 | -1.16858268 | 0.24257181 | 0.8422984 | 45.1118389 | 25.1966429 | 49.6417192 | 29.1045461 | Non-sig. |
| PF3D7_0913200 | EF-1beta | 4494.10752 | -0.803317168 | 0.42296496 | -1.8992523 | 0.05753131 | 0.51925194 | 6442.40241 | 3602.27851 | 4985.62968 | 2946.11949 | Non-sig. |
| PF3D7_1360400 | N/A | 341.929745 | -0.805004376 | 0.44248243 | -1.81929115 | 0.06886702 | 0.57367529 | 437.318746 | 264.228797 | 432.540318 | 233.631121 | Non-sig. |
| PF3D7_1461000 | N/A | 196.601159 | -0.805213442 | 0.47394878 | -1.6989461 | 0.08932934 | 0.62981332 | 254.63094 | 128.772654 | 245.549633 | 157.451408 | Non-sig. |
| PF3D7_1327700 | UPF3B | 272.000312 | -0.805476017 | 0.45432644 | -1.77290147 | 0.07624503 | 0.59693801 | 373.473936 | 201.752167 | 318.631459 | 194.143688 | Non-sig. |
| PF3D7_1347400 | N/A | 222.695737 | -0.806217333 | 0.5043944 | -1.59838677 | 0.10995693 | 0.67975706 | 207.798822 | 166.451664 | 358.718006 | 157.814456 | Non-sig. |
| PF3D7_0107000 | CEN1 | 3731.72514 | -0.806262204 | 0.42695234 | -1.88841268 | 0.05897057 | 0.52596984 | 5300.58588 | 3073.80145 | 4195.85504 | 2356.65818 | Non-sig. |
| PF3D7_1305300 | GCN1 | 2966.37716 | -0.806615394 | 0.53096371 | -1.51915354 | 0.12872385 | 0.71411411 | 2158.23524 | 1807.14559 | 5390.86805 | 2509.25976 | Non-sig. |
| PF3D7_0622400 | N/A | 356.785295 | -0.808458159 | 0.4756098 | -1.69983496 | 0.08916197 | 0.62981332 | 442.416018 | 321.400517 | 466.010544 | 197.314101 | Non-sig. |
| PF3D7_1469800 | N/A | 1322.08778 | -0.808536488 | 0.48445306 | -1.66896765 | 0.09512379 | 0.64740909 | 2172.09192 | 1124.54526 | 1194.44613 | 797.267824 | Non-sig. |
| PF3D7_0728300 | N/A | 293.5061 | -0.808848137 | 0.47802941 | -1.69204682 | 0.09063705 | 0.63196333 | 392.070597 | 259.960815 | 355.364406 | 166.628581 | Non-sig. |
| PF3D7_0526500 | N/A | 1293.87738 | -0.809984025 | 0.44652346 | -1.81397866 | 0.06968101 | 0.57592823 | 1532.8624 | 740.181634 | 1762.75874 | 1139.70674 | Non-sig. |
| PF3D7_1005400 | N/A | 5.18963165 | -0.812910557 | 1.53158705 | -0.53076353 | 0.59558265 | 0.95502327 | 4.17807322 | 4.30123509 | 9.02299733 | 3.25622096 | Non-sig. |
| PF3D7_0825300 | N/A | 308.985576 | -0.813059374 | 0.44768377 | -1.81614666 | 0.06934788 | 0.57496879 | 395.330025 | 240.60535 | 392.316137 | 207.690791 | Non-sig. |
| PF3D7_1012600 | GMPS | 977.106795 | -0.813824542 | 0.52401206 | -1.55306452 | 0.12040772 | 0.70293962 | 721.067826 | 715.673375 | 1769.89927 | 701.786711 | Non-sig. |
| PF3D7_1454100 | N/A | 274.172467 | -0.814008024 | 0.52697792 | -1.54467198 | 0.12242563 | 0.7079393 | 241.825195 | 238.569375 | 457.055115 | 159.240184 | Non-sig. |
| PF3D7_0306700 | EMC5 | 328.80126 | -0.815646004 | 0.4908653 | -1.66164935 | 0.0965831 | 0.64895933 | 312.193294 | 265.130606 | 526.321385 | 211.559754 | Non-sig. |
| PF3D7_0819900 | LSM3 | 435.605124 | -0.816220007 | 0.48017778 | -1.69982877 | 0.08916314 | 0.62981332 | 609.400005 | 396.77797 | 501.981889 | 234.260631 | Non-sig. |
| PF3D7_1115600 | CYP19B | 9128.16681 | -0.8167989 | 0.41579151 | -1.96444345 | 0.04947869 | 0.48997774 | 11143.9471 | 7468.84476 | 12146.6175 | 5753.2579 | Non-sig. |
| PF3D7_0809600 | N/A | 1727.95612 | -0.818117866 | 0.47709325 | -1.71479656 | 0.08638253 | 0.62521128 | 1561.79001 | 1082.08154 | 2848.33704 | 1419.6159 | Non-sig. |
| PF3D7_1213100 | N/A | 164.118783 | -0.818589728 | 0.55272535 | -1.48100631 | 0.13860489 | 0.72994902 | 262.318574 | 151.184981 | 156.729932 | 86.2416438 | Non-sig. |
| PF3D7_0615000 | N/A | 234.520572 | -0.822187601 | 0.54762892 | -1.50135898 | 0.13326274 | 0.72178914 | 409.056294 | 197.770132 | 190.349762 | 140.906101 | Non-sig. |
| PF3D7_1411600 | Obg1 | 262.459912 | -0.823788828 | 0.50651749 | -1.62637784 | 0.1038693 | 0.6615241 | 295.488252 | 136.307502 | 375.265325 | 242.778566 | Non-sig. |
| PF3D7_0710300 | N/A | 222.260465 | -0.824313573 | 0.48525293 | -1.69872973 | 0.08937012 | 0.62981332 | 229.83398 | 168.600274 | 338.195315 | 152.412293 | Non-sig. |
| PF3D7_1123400 | ERF3 | 3288.92079 | -0.82583248 | 0.48953966 | -1.68695725 | 0.09161157 | 0.6361181 | 2721.27222 | 2533.24811 | 5689.19719 | 2211.96564 | Non-sig. |
| PF3D7_1367200 | N/A | 17.2489635 | -0.825902611 | 0.98774444 | -0.8361501 | 0.40307047 | 0.90716539 | 33.5395235 | 13.9656101 | 10.6687663 | 10.8219539 | Non-sig. |
| PF3D7_0304500 | GLP1 | 405.2122 | -0.826108317 | 0.47969728 | -1.72214511 | 0.08504324 | 0.61879091 | 427.116998 | 351.16509 | 609.075539 | 233.491174 | Non-sig. |
| PF3D7_0501200 | PIESP2 | 2891.49905 | -0.827016739 | 0.49475897 | -1.67155483 | 0.09461214 | 0.6459473 | 2331.4465 | 2124.53423 | 5064.88179 | 2045.13368 | Non-sig. |
| PF3D7_1463400 | N/A | 306.141945 | -0.828150559 | 0.46076647 | -1.7973325 | 0.07228285 | 0.58363337 | 435.283984 | 244.71928 | 348.15874 | 196.405776 | Non-sig. |
| PF3D7_1036600 | N/A | 438.614881 | -0.828344732 | 0.48098288 | -1.72219172 | 0.08503479 | 0.61879091 | 689.907762 | 365.061319 | 432.639343 | 266.851098 | Non-sig. |
| PF3D7_1223900 | N/A | 369.320408 | -0.830337439 | 0.44664587 | -1.85905095 | 0.06301992 | 0.54644766 | 442.205765 | 292.027033 | 503.264613 | 239.78422 | Non-sig. |
| PF3D7_0424000 | N/A | 296.696463 | -0.830534212 | 0.46869444 | -1.77201636 | 0.07639184 | 0.59693801 | 413.191236 | 249.014824 | 346.514552 | 178.06524 | Non-sig. |
| PF3D7_0209300 | IspF | 422.931613 | -0.83072234 | 0.44112388 | -1.88319512 | 0.05967393 | 0.52780047 | 500.105573 | 328.700038 | 582.711163 | 280.209679 | Non-sig. |
| PF3D7_1436000 | GPI | 1657.6119 | -0.832280963 | 0.5885264 | -1.41417778 | 0.15730971 | 0.75912848 | 935.344262 | 1110.33122 | 3310.19799 | 1274.57412 | Non-sig. |
| PF3D7_0613200 | N/A | 322.196832 | -0.832690592 | 0.47937323 | -1.73704026 | 0.08238006 | 0.60888892 | 382.533049 | 285.122791 | 442.791056 | 178.340431 | Non-sig. |
| PF3D7_1462200 | N/A | 323.703162 | -0.833144315 | 0.50724504 | -1.64248883 | 0.10048874 | 0.65652696 | 535.856092 | 266.673871 | 293.658758 | 198.623928 | Non-sig. |
| PF3D7_1411500 | N/A | 424.976523 | -0.833920933 | 0.54806211 | -1.52158107 | 0.12811408 | 0.71411411 | 784.51941 | 322.931174 | 304.721749 | 287.733762 | Non-sig. |
| PF3D7_1246600 | BUD13 | 442.720988 | -0.834744217 | 0.50922071 | -1.63925817 | 0.1011595 | 0.65679295 | 765.414095 | 335.953926 | 369.507012 | 300.008919 | Non-sig. |
| PF3D7_1350000 | N/A | 38.9803435 | -0.835230061 | 0.83766523 | -0.9970929 | 0.31871942 | 0.88651753 | 67.176899 | 45.1012222 | 32.8354277 | 10.8078253 | Non-sig. |
| PF3D7_MIT04000 | LSUE | 16200.7709 | -0.835523681 | 0.44443122 | -1.87998421 | 0.06011023 | 0.53077333 | 20739.2284 | 14478.4604 | 20791.0862 | 8794.3086 | Non-sig. |
| PF3D7_1001600 | XL2 | 10352.3662 | -0.835544318 | 0.4634873 | -1.80273402 | 0.07143 | 0.58131508 | 9780.80558 | 6357.86294 | 16757.1964 | 8513.59976 | Non-sig. |
| PF3D7_1309100 | N/A | 4657.23104 | -0.837069831 | 0.43552309 | -1.92198726 | 0.05460736 | 0.50314747 | 6444.94931 | 3992.48099 | 5498.44236 | 2693.0515 | Non-sig. |
| PF3D7_1433100 | N/A | 342.463367 | -0.837374918 | 0.48103745 | -1.74076865 | 0.08172414 | 0.60888892 | 392.795139 | 188.822338 | 485.421203 | 302.81479 | Non-sig. |
| PF3D7_0511100 | N/A | 30.0893316 | -0.838170522 | 0.74102687 | -1.1310933 | 0.25801582 | 0.85030738 | 34.4908916 | 20.4272637 | 42.6581091 | 22.7810618 | Non-sig. |
| PF3D7_1351100 | N/A | 647.396877 | -0.838413441 | 0.43208087 | -1.94040859 | 0.05233005 | 0.50193661 | 753.440991 | 500.250265 | 907.258833 | 428.637418 | Non-sig. |
| PF3D7_1310000 | OSCP | 122.577776 | -0.839165524 | 0.5073745 | -1.65393713 | 0.09814029 | 0.65401834 | 167.661092 | 90.2448357 | 146.895038 | 85.5101399 | Non-sig. |
| PF3D7_1431800 | E-NTPDase | 254.877506 | -0.839829711 | 0.50921867 | -1.64925162 | 0.0990961 | 0.65421395 | 228.435144 | 186.695269 | 425.435883 | 178.943729 | Non-sig. |
| PF3D7_1027500 | N/A | 17.4068457 | -0.839832285 | 0.9477011 | -0.88617845 | 0.37552138 | 0.89654416 | 14.6105049 | 15.2819514 | 29.9866339 | 9.74829242 | Non-sig. |
| PF3D7_0310600 | EIF3K | 1166.95577 | -0.840115846 | 0.42280749 | -1.98699376 | 0.04692309 | 0.48052729 | 1402.13704 | 922.266879 | 1592.6921 | 750.727049 | Non-sig. |

| | | | | | | | | | | | | |
|---|---|---|---|---|---|---|---|---|---|---|---|---|
| PF3D7_1005200 | N/A | 8.33762788 | -0.841257165 | 1.24189824 | -0.67739621 | 0.4981546 | 0.93170321 | 12.4170771 | 7.5471056 | 9.00809005 | 4.37823873 | Non-sig. |
| PF3D7_0208600 | RRF1 | 2291.34769 | -0.842991188 | 0.48762163 | -1.72878135 | 0.08384823 | 0.61527417 | 3955.25508 | 1752.63608 | 1929.73036 | 1527.76923 | Non-sig. |
| PF3D7_1408500 | N/A | 360.487683 | -0.844555362 | 0.51767135 | -1.63145085 | 0.10279522 | 0.66002937 | 617.699754 | 295.236885 | 308.69808 | 220.316011 | Non-sig. |
| PF3D7_1214300 | N/A | 750.79045 | -0.844976755 | 0.51182297 | -1.65091606 | 0.09875572 | 0.65401834 | 593.83649 | 546.379472 | 1335.0729 | 527.872934 | Non-sig. |
| PF3D7_0502600 | N/A | 186.081977 | -0.848147867 | 0.49515418 | -1.71289652 | 0.08673158 | 0.62688118 | 283.173266 | 123.390808 | 195.470288 | 142.293545 | Non-sig. |
| PF3D7_0906800 | N/A | 130.360342 | -0.848672976 | 0.53485621 | -1.5867311 | 0.11257356 | 0.68406239 | 165.248103 | 68.672476 | 170.005553 | 117.515235 | Non-sig. |
| PF3D7_0526100 | N/A | 186.078762 | -0.850787901 | 0.5590883 | -1.52174156 | 0.12807385 | 0.71411411 | 149.329678 | 151.331187 | 329.284126 | 114.370056 | Non-sig. |
| PF3D7_0725600 | N/A | 4522.46411 | -0.852393136 | 0.43387158 | -1.96462079 | 0.04945814 | 0.48997774 | 4913.14062 | 2799.0929 | 6728.56005 | 3649.06286 | Non-sig. |
| PF3D7_1459500 | N/A | 58.7867137 | -0.852608392 | 0.60884331 | -1.40037408 | 0.16140133 | 0.76690962 | 64.2095767 | 43.9923604 | 87.0571448 | 39.887773 | Non-sig. |
| PF3D7_0703900 | N/A | 1710.80298 | -0.854043529 | 0.51790468 | -1.64903611 | 0.09914024 | 0.65421395 | 1378.16116 | 973.793417 | 3027.38411 | 1463.87323 | Non-sig. |
| PF3D7_1369500 | N/A | 1094.99846 | -0.854202705 | 0.49228968 | -1.73516272 | 0.08271199 | 0.6103177 | 932.224738 | 713.392512 | 1887.56637 | 846.810207 | Non-sig. |
| PF3D7_1208300 | mACP | 819.075293 | -0.855886182 | 0.44223823 | -1.93535096 | 0.05294724 | 0.50202873 | 1205.72111 | 653.855719 | 904.713243 | 512.011104 | Non-sig. |
| PF3D7_0312000 | N/A | 151.875142 | -0.857249855 | 0.54672723 | -1.56796627 | 0.116889 | 0.69425779 | 135.446297 | 121.593694 | 255.818071 | 94.6425066 | Non-sig. |
| PF3D7_0819200 | PLP5 | 83.1903383 | -0.857921618 | 0.55162519 | -1.55526185 | 0.11988372 | 0.70181652 | 97.790842 | 61.1781219 | 116.602352 | 57.1900376 | Non-sig. |
| PF3D7_1227200 | K1 | 2346.00567 | -0.857992855 | 0.50361615 | -1.7036643 | 0.08844382 | 0.62981332 | 2365.40249 | 1115.46101 | 3681.9438 | 2221.21539 | Non-sig. |
| PF3D7_1441200 | N/A | 2024.29942 | -0.858340353 | 0.55713992 | -1.54061902 | 0.12340954 | 0.70887094 | 1294.94157 | 1446.31261 | 3923.43908 | 1432.50441 | Non-sig. |
| PF3D7_1317800 | RPS19 | 6197.95156 | -0.860417797 | 0.4140175 | -2.078216 | 0.03768947 | 0.43374033 | 8057.53201 | 4949.23556 | 7929.0221 | 3856.01659 | Non-sig. |
| PF3D7_1325800 | N/A | 416.093864 | -0.862651211 | 0.52775739 | -1.63456017 | 0.10214128 | 0.65895255 | 736.453355 | 252.086288 | 337.605755 | 338.230057 | Non-sig. |
| PF3D7_1025700 | N/A | 736.205994 | -0.86294118 | 0.4696306 | -1.83748925 | 0.0661377 | 0.56240843 | 726.515684 | 596.424479 | 1173.40748 | 448.476328 | Non-sig. |
| PF3D7_1424100 | RPL5 | 7598.68449 | -0.864210596 | 0.41771569 | -2.06889665 | 0.03855579 | 0.43554063 | 8395.31643 | 5401.27608 | 11222.3231 | 5375.82237 | Non-sig. |
| PF3D7_1130900 | N/A | 176.029211 | -0.865017677 | 0.49232476 | -1.75700624 | 0.07891675 | 0.60514927 | 251.741757 | 108.351245 | 202.883008 | 141.140836 | Non-sig. |
| PF3D7_1421800 | N/A | 183.593581 | -0.870744369 | 0.62112826 | -1.40187531 | 0.16095249 | 0.7667698 | 355.941297 | 158.913871 | 119.035123 | 100.484032 | Non-sig. |
| PF3D7_1217000 | N/A | 37.4856637 | -0.871932049 | 0.70664165 | -1.23390978 | 0.21723652 | 0.83527883 | 58.4843641 | 30.1097771 | 38.5519291 | 22.7965845 | Non-sig. |
| PF3D7_1104300 | N/A | 2004.34456 | -0.872070493 | 0.44909297 | -1.94184845 | 0.05215545 | 0.50193661 | 2262.81186 | 1122.09137 | 2921.74622 | 1710.7288 | Non-sig. |
| PF3D7_1426900 | QCR6 | 2409.87588 | -0.872439617 | 0.43061374 | -2.02603757 | 0.04276094 | 0.46423655 | 3532.03252 | 1915.61137 | 2702.32519 | 1489.53444 | Non-sig. |
| PF3D7_1371300 | N/A | 39.5106901 | -0.872698794 | 4.56856357 | -0.19102258 | 0.8485079 | 0.99165628 | 102.228963 | 55.8137977 | 0 | 0 | Non-sig. |
| PF3D7_0814000 | N/A | 5639.74136 | -0.875098115 | 0.43670218 | -2.00387853 | 0.04508307 | 0.47730745 | 8542.8835 | 4508.25422 | 6056.50166 | 3451.32607 | Non-sig. |
| PF3D7_1204300 | EIF5A | 7621.92639 | -0.875294559 | 0.45263634 | -1.93376994 | 0.05314142 | 0.50202873 | 7415.3195 | 5907.46091 | 12315.8071 | 4849.11804 | Non-sig. |
| PF3D7_0530600 | N/A | 495.423113 | -0.877213149 | 0.51363312 | -1.70785939 | 0.08766244 | 0.62914105 | 859.685473 | 404.694194 | 423.670766 | 293.642018 | Non-sig. |
| PF3D7_0210100 | N/A | 2461.60733 | -0.878187437 | 0.44028905 | -1.99457024 | 0.04608977 | 0.47851132 | 3472.92764 | 2079.78833 | 2904.28772 | 1389.42561 | Non-sig. |
| PF3D7_0624900 | N/A | 158.255212 | -0.87933133 | 0.64081649 | -1.3722046 | 0.16999975 | 0.77948916 | 317.531842 | 125.578981 | 92.7878882 | 97.122137 | Non-sig. |
| PF3D7_1245900 | N/A | 194.852585 | -0.880313392 | 0.49556756 | -1.77637414 | 0.07567127 | 0.59646408 | 202.773755 | 147.07648 | 302.13668 | 127.423426 | Non-sig. |
| PF3D7_0915200 | N/A | 160.844545 | -0.883560271 | 0.49403192 | -1.78846798 | 0.07370054 | 0.58628445 | 237.513638 | 115.934078 | 179.813307 | 110.117157 | Non-sig. |
| PF3D7_0315400 | N/A | 1373.11247 | -0.883860565 | 0.42270425 | -2.09096685 | 0.03653104 | 0.42994016 | 1768.32932 | 1089.81865 | 1793.76698 | 840.534913 | Non-sig. |
| PF3D7_1123900 | SNU13 | 941.86537 | -0.883902178 | 0.44750036 | -1.9751988 | 0.04824559 | 0.48501926 | 1062.49837 | 775.257485 | 1380.79011 | 548.915513 | Non-sig. |
| PF3D7_1110000 | N/A | 214.375109 | -0.884007552 | 0.48660409 | -1.81668748 | 0.06926498 | 0.57496879 | 280.611113 | 180.716419 | 275.554372 | 120.618531 | Non-sig. |
| PF3D7_1442000 | N/A | 133.63848 | -0.884080281 | 0.50210869 | -1.76073487 | 0.07828329 | 0.60381404 | 176.861657 | 84.9040654 | 169.847921 | 102.940278 | Non-sig. |
| PF3D7_0107400 | ATPk | 354.83303 | -0.885804299 | 0.44102038 | -2.00853369 | 0.04458661 | 0.47433706 | 434.001308 | 250.123369 | 486.884036 | 248.323407 | Non-sig. |
| PF3D7_1124400 | LSM1 | 233.790866 | -0.887760298 | 0.46494186 | -1.90940068 | 0.05621042 | 0.5116887 | 330.636812 | 173.042024 | 276.506213 | 154.978414 | Non-sig. |
| PF3D7_1135700 | N/A | 227.090621 | -0.88829397 | 0.46309463 | -1.91816944 | 0.05508953 | 0.50446754 | 278.835624 | 167.755883 | 310.874734 | 150.896242 | Non-sig. |
| PF3D7_1125500 | SNRPD1 | 743.138478 | -0.888938726 | 0.43168014 | -2.05925323 | 0.03946999 | 0.44209722 | 858.817852 | 559.430955 | 1071.30267 | 483.00243 | Non-sig. |
| PF3D7_0503800 | RPL31 | 2051.2613 | -0.890557051 | 0.44262706 | -2.01198057 | 0.04422199 | 0.472355 | 2226.10726 | 1653.32975 | 3103.77091 | 1221.83729 | Non-sig. |
| PF3D7_1308300 | RPS27 | 3645.29148 | -0.890602144 | 0.41762947 | -2.13251747 | 0.03296433 | 0.40615121 | 4267.46361 | 2788.35761 | 5204.49574 | 2320.84896 | Non-sig. |
| PF3D7_0825500 | KRI1 | 853.320565 | -0.89067639 | 0.59259569 | -1.50300855 | 0.13283684 | 0.72037933 | 1719.90513 | 665.257885 | 497.700543 | 530.418708 | Non-sig. |
| PF3D7_0922900 | FabG | 96.9814865 | -0.893081885 | 0.54758849 | -1.63093618 | 0.10290379 | 0.66002937 | 140.590475 | 78.3997856 | 111.631891 | 57.3037949 | Non-sig. |
| PF3D7_0513800 | RAB1a | 814.409173 | -0.894227499 | 0.45442177 | -1.96783596 | 0.04908691 | 0.48792209 | 829.336534 | 526.345996 | 1288.53609 | 613.418073 | Non-sig. |
| PF3D7_0728400 | N/A | 171.388851 | -0.895969996 | 0.51930011 | -1.72534144 | 0.08446596 | 0.61653204 | 228.603319 | 152.720816 | 217.349414 | 86.8818541 | Non-sig. |
| PF3D7_1135600 | CAPD3 | 348.160919 | -0.896211799 | 0.48168934 | -1.86055974 | 0.06280638 | 0.54548881 | 344.655544 | 222.026077 | 561.071529 | 264.890524 | Non-sig. |
| PF3D7_MIT02300 | CYTB | 3218.09315 | -0.897985976 | 0.56546205 | -1.58805702 | 0.11227345 | 0.68406239 | 2044.19131 | 1956.15868 | 6332.52965 | 2539.49298 | Non-sig. |
| PF3D7_0314600 | N/A | 2200.29193 | -0.898189529 | 0.58192213 | -1.5434875 | 0.12271254 | 0.7079393 | 4446.23675 | 1515.35781 | 1281.87976 | 1557.69338 | Non-sig. |
| PF3D7_1120800 | N/A | 100.511248 | -0.898533903 | 0.60200738 | -1.49256294 | 0.13555163 | 0.72467461 | 80.2774923 | 75.3107179 | 181.22398 | 65.232803 | Non-sig. |
| PF3D7_1462500 | N/A | 22.4024104 | -0.898844847 | 0.85086766 | -1.05638619 | 0.29079181 | 0.86724628 | 37.8694514 | 13.9513581 | 20.5300015 | 17.2588305 | Non-sig. |
| PF3D7_1030700 | N/A | 145.044564 | -0.899414788 | 0.4996503 | -1.80008857 | 0.07184665 | 0.58133781 | 165.82586 | 97.7221438 | 211.784657 | 104.845593 | Non-sig. |
| PF3D7_0709800 | N/A | 283.618859 | -0.900022602 | 0.52909467 | -1.70106155 | 0.08893143 | 0.62981332 | 500.879976 | 218.092606 | 237.98574 | 177.517116 | Non-sig. |

| | | | | | | | | | | | | |
|---|---|---|---|---|---|---|---|---|---|---|---|---|
| PF3D7_0521600 | N/A | 39.3591435 | -0.901768501 | 0.75100404 | -1.20075053 | 0.22984799 | 0.83828834 | 59.9109199 | 39.7168455 | 42.6962776 | 15.1125312 | Non-sig. |
| PF3D7_0713200 | EPF1 | 10.8712205 | -0.902489069 | 1.69562016 | -0.53224719 | 0.5945548 | 0.9542761 | 28.387377 | 1.07308532 | 0 | 14.0244197 | Non-sig. |
| PF3D7_1114000 | N/A | 23.4398057 | -0.903377467 | 0.93860649 | -0.96246667 | 0.33581524 | 0.8920875 | 49.7460301 | 17.2424336 | 11.4683031 | 15.3024559 | Non-sig. |
| PF3D7_1126400 | MED21 | 271.756668 | -0.905201581 | 0.4630544 | -1.95484932 | 0.05060086 | 0.49645064 | 358.26294 | 215.96202 | 350.394894 | 162.40682 | Non-sig. |
| PF3D7_1401600 | N/A | 12211.9119 | -0.905757037 | 0.40947458 | -2.21199819 | 0.02696679 | 0.36452847 | 16086.1136 | 7606.7542 | 15762.3566 | 9392.42311 | Non-sig. |
| PF3D7_1239300 | N/A | 61.8065614 | -0.906710004 | 0.61922914 | -1.46425602 | 0.14312401 | 0.74065302 | 60.9508461 | 46.1597856 | 100.168821 | 39.9467932 | Non-sig. |
| PF3D7_1417900 | ATPTG9 | 642.962577 | -0.908636077 | 0.43845656 | -2.07235143 | 0.03823268 | 0.43374033 | 761.856398 | 501.516167 | 916.069945 | 392.407799 | Non-sig. |
| PF3D7_0425900 | RIF | 24.1014255 | -0.914268569 | 0.80958642 | -1.12930325 | 0.25876993 | 0.85030738 | 36.7596704 | 18.2521467 | 26.2711499 | 15.1227348 | Non-sig. |
| PF3D7_0306300 | GRX1 | 2492.45609 | -0.914387456 | 0.45165599 | -2.02452193 | 0.04291648 | 0.46497239 | 2560.15722 | 1968.03879 | 3953.47696 | 1488.15138 | Non-sig. |
| PF3D7_1463000 | MED20 | 296.128389 | -0.91489442 | 0.4691268 | -1.9502071 | 0.05115144 | 0.49816237 | 420.203914 | 170.700599 | 353.853384 | 239.755659 | Non-sig. |
| PF3D7_1419300 | GST | 696.489003 | -0.914988365 | 0.53730248 | -1.70292974 | 0.08858122 | 0.62981332 | 513.584486 | 519.350655 | 1306.6375 | 446.38337 | Non-sig. |
| PF3D7_1203100 | N/A | 412.861579 | -0.91568355 | 0.48936963 | -1.87114912 | 0.06132441 | 0.53711809 | 636.273118 | 354.500666 | 443.172805 | 217.499727 | Non-sig. |
| PF3D7_1308900 | DCP2 | 2393.55478 | -0.916045608 | 0.44168098 | -2.07399834 | 0.03807947 | 0.43374033 | 2726.45229 | 1357.00542 | 3531.25812 | 1959.50327 | Non-sig. |
| PF3D7_0505800 | SUMO | 4636.32345 | -0.916145929 | 0.41867808 | -2.18818697 | 0.02865599 | 0.37778095 | 5670.42402 | 3624.22814 | 6451.24282 | 2799.39883 | Non-sig. |
| PF3D7_1415900 | N/A | 428.779239 | -0.916438151 | 0.45966008 | -1.99373013 | 0.04618155 | 0.47851132 | 665.775852 | 316.881773 | 455.501611 | 276.957719 | Non-sig. |
| PF3D7_0921500 | N/A | 20.1729245 | -0.917450453 | 0.90314206 | -1.01584291 | 0.3097042 | 0.87951138 | 15.7368696 | 13.9618222 | 36.9366621 | 14.0563442 | Non-sig. |
| PF3D7_1205600 | N/A | 3165.72415 | -0.917933615 | 0.45580421 | -2.01387702 | 0.04402245 | 0.47123681 | 5039.22403 | 2546.59079 | 3241.33104 | 1835.75075 | Non-sig. |
| PF3D7_0113000 | GARP | 2966.97493 | -0.918491571 | 0.77013978 | -1.1926297 | 0.23301443 | 0.83828834 | 7064.47518 | 2387.15497 | 697.321114 | 1718.94845 | Non-sig. |
| PF3D7_0317600 | RPS11 | 4967.49098 | -0.91906995 | 0.44171285 | -2.08069551 | 0.03746178 | 0.43374033 | 5116.4368 | 3745.87839 | 7880.06728 | 3127.58143 | Non-sig. |
| PF3D7_1469100 | N/A | 365.941115 | -0.919631814 | 0.44152133 | -2.08287066 | 0.03726301 | 0.43374033 | 506.095075 | 265.21327 | 451.513706 | 240.942407 | Non-sig. |
| PF3D7_0716100 | SDA1 | 455.092672 | -0.921177712 | 0.60614752 | -1.51972528 | 0.12858004 | 0.71411411 | 931.15852 | 334.705758 | 260.38168 | 294.124728 | Non-sig. |
| PF3D7_1016400 | FIKK10.1 | 1921.34617 | -0.921953286 | 0.60164374 | -1.53239073 | 0.12542604 | 0.71290716 | 1036.91975 | 1332.613 | 3993.19311 | 1322.65883 | Non-sig. |
| PF3D7_1246000 | N/A | 20.3282518 | -0.923520998 | 0.87349488 | -1.05727122 | 0.29038782 | 0.86724628 | 33.5990544 | 12.8849218 | 19.7023524 | 15.1266785 | Non-sig. |
| PF3D7_0403300 | N/A | 107.037826 | -0.92398282 | 0.57360715 | -1.61082864 | 0.10721708 | 0.67462717 | 136.831898 | 98.928407 | 143.547685 | 48.8433144 | Non-sig. |
| PF3D7_1231700 | N/A | 71.8071021 | -0.92509286 | 0.57208955 | -1.61704206 | 0.1058692 | 0.66932581 | 87.1318231 | 46.1733084 | 100.971347 | 52.9519304 | Non-sig. |
| PF3D7_0605000 | N/A | 227.469936 | -0.927332972 | 0.47990548 | -1.9323242 | 0.0533195 | 0.50202873 | 286.09087 | 184.783582 | 310.206604 | 128.798687 | Non-sig. |
| PF3D7_1336200 | N/A | 1137.154 | -0.928064355 | 0.48016404 | -1.93280687 | 0.05325999 | 0.50202873 | 1904.35391 | 910.009165 | 1077.45034 | 656.802603 | Non-sig. |
| PF3D7_1358800 | RPS15 | 4622.46706 | -0.928215234 | 0.42367469 | -2.19086781 | 0.02846136 | 0.37697071 | 5685.37561 | 3672.74383 | 6435.04468 | 2696.70411 | Non-sig. |
| PF3D7_0919600 | WBP1 | 244.709648 | -0.931287985 | 0.47780086 | -1.94911325 | 0.0512819 | 0.49851654 | 261.800752 | 169.586256 | 380.182652 | 167.268933 | Non-sig. |
| PF3D7_1460700 | RPL27 | 4037.64385 | -0.931548487 | 0.42327972 | -2.20078698 | 0.02775111 | 0.37221613 | 4658.90675 | 3078.07832 | 5936.43218 | 2477.15817 | Non-sig. |
| PF3D7_0606900 | GLP2 | 491.00891 | -0.932139174 | 0.4731264 | -1.97016945 | 0.04881895 | 0.48633045 | 478.741829 | 343.395039 | 809.737023 | 332.161749 | Non-sig. |
| PF3D7_0823100 | N/A | 234.553018 | -0.932167762 | 0.4833298 | -1.92863706 | 0.05377593 | 0.50202873 | 338.903908 | 189.9243 | 276.770529 | 132.613334 | Non-sig. |
| PF3D7_0509200 | LRR2 | 28.3002378 | -0.933617917 | 0.76899985 | -1.21406775 | 0.22472185 | 0.83828834 | 43.9909229 | 17.191694 | 30.3613181 | 21.6570165 | Non-sig. |
| PF3D7_1461700 | N/A | 80.1042797 | -0.935273395 | 0.7229669 | -1.29366005 | 0.19578287 | 0.81434658 | 167.903012 | 65.5084508 | 42.6842974 | 44.3213586 | Non-sig. |
| PF3D7_1131000 | N/A | 403.179704 | -0.935491721 | 0.43629452 | -2.14417483 | 0.03201888 | 0.40103081 | 546.810317 | 288.870243 | 512.224147 | 264.814111 | Non-sig. |
| PF3D7_0710600 | RPL34 | 3223.20715 | -0.935652393 | 0.4100083 | -2.28203282 | 0.0224874 | 0.33002288 | 4232.33891 | 2408.589 | 4234.16266 | 2017.73802 | Non-sig. |
| PF3D7_0415900 | N/A | 7437.83723 | -0.937071891 | 0.40748412 | -2.29965256 | 0.02146791 | 0.3235449 | 9608.86405 | 5545.43016 | 9934.91663 | 4662.13808 | Non-sig. |
| PF3D7_1350200 | SNRPE | 292.326143 | -0.93720765 | 0.48867915 | -1.91783841 | 0.0551315 | 0.50446754 | 472.28187 | 223.780419 | 296.026501 | 177.215782 | Non-sig. |
| PF3D7_0517000 | N/A | 4866.03138 | -0.937219832 | 0.42665377 | -2.19667536 | 0.02804364 | 0.37518991 | 5644.67215 | 3786.37854 | 7141.74828 | 2891.32655 | Non-sig. |
| PF3D7_1006600 | PhLP3 | 678.494571 | -0.93818117 | 0.46558635 | -2.01505298 | 0.0438991 | 0.47123681 | 804.982269 | 576.880356 | 978.251054 | 353.864604 | Non-sig. |
| PF3D7_0902000 | FIKK9.1 | 395.359067 | -0.938492961 | 0.46045532 | -2.03818466 | 0.04153147 | 0.45538799 | 446.15072 | 235.007059 | 592.937321 | 307.341168 | Non-sig. |
| PF3D7_0708100 | RPB10 | 314.366332 | -0.939955396 | 0.46070812 | -2.04024055 | 0.04132637 | 0.45424714 | 350.411181 | 224.462503 | 476.073386 | 206.518258 | Non-sig. |
| PF3D7_1318500 | N/A | 14.2210886 | -0.940921258 | 1.02231343 | -0.92038432 | 0.35737196 | 0.89552149 | 25.1468677 | 7.52059536 | 12.3094409 | 11.9074503 | Non-sig. |
| PF3D7_1308600 | N/A | 195.071208 | -0.94617983 | 0.51532786 | -1.83607351 | 0.06634679 | 0.56240843 | 209.639255 | 104.055912 | 303.921403 | 162.668263 | Non-sig. |
| PF3D7_1250000 | FCF2 | 540.080997 | -0.948302998 | 0.47021397 | -2.01674783 | 0.04372184 | 0.47123681 | 698.915535 | 463.925116 | 723.99487 | 273.488466 | Non-sig. |
| PF3D7_0610500 | N/A | 473.915216 | -0.949677533 | 0.51864517 | -1.83107372 | 0.06708954 | 0.56689057 | 389.036159 | 305.895953 | 859.72769 | 341.001064 | Non-sig. |
| PF3D7_0819400 | PLP4 | 8.66520672 | -0.949990779 | 1.22127232 | -0.77786974 | 0.43664581 | 0.91522571 | 10.5138905 | 7.5133545 | 12.3168923 | 4.31668955 | Non-sig. |
| PF3D7_0706400 | RPL37 | 2307.71268 | -0.950232254 | 0.41714737 | -2.27792941 | 0.02273078 | 0.33267319 | 3055.26742 | 1764.5762 | 3027.48011 | 1383.52701 | Non-sig. |
| PF3D7_0929800 | N/A | 9.62356981 | -0.950714578 | 1.16054942 | -0.81919353 | 0.412676 | 0.90786241 | 12.3363908 | 5.42980257 | 13.0269681 | 7.70111783 | Non-sig. |
| PF3D7_1403000 | N/A | 32.456591 | -0.951640781 | 0.88879176 | -1.07071288 | 0.28429855 | 0.86551387 | 69.2962464 | 30.0650371 | 16.4185517 | 14.0465288 | Non-sig. |
| PF3D7_0810700 | N/A | 111.466663 | -0.951795297 | 0.53097318 | -1.79254872 | 0.07304511 | 0.58427436 | 170.055613 | 81.609838 | 123.955549 | 70.2456524 | Non-sig. |
| PF3D7_1039000 | FIKK10.2 | 731.263754 | -0.952488606 | 0.41949319 | -2.27056991 | 0.02317303 | 0.3380532 | 991.312653 | 500.969731 | 937.228455 | 495.544178 | Non-sig. |
| PF3D7_0630000 | N/A | 8.7266709 | -0.953621539 | 1.30658007 | -0.72986077 | 0.46547529 | 0.92538923 | 15.6778924 | 9.67604534 | 7.38314886 | 2.16959702 | Non-sig. |
| PF3D7_0826700 | RACK1 | 5514.05259 | -0.953819335 | 0.54887985 | -1.73775615 | 0.08225379 | 0.60888892 | 3693.58665 | 3913.05425 | 10852.5741 | 3596.99533 | Non-sig. |

| | | | | | | | | | | | | |
|---|---|---|---|---|---|---|---|---|---|---|---|---|
| PF3D7_1109900 | RPL36 | 2404.97312 | -0.954714739 | 0.42681178 | -2.23685192 | 0.02529602 | 0.35869122 | 3011.3522 | 1897.81404 | 3334.42287 | 1376.30338 | Non-sig. |
| PF3D7_1102300 | N/A | 54.7408446 | -0.956620939 | 0.86817508 | -1.1018756 | 0.27051576 | 0.8556331 | 128.25351 | 33.2745902 | 16.4221648 | 41.013113 | Non-sig. |
| PF3D7_1412600 | DHS | 263.565428 | -0.957232255 | 0.45239587 | -2.11591731 | 0.03435184 | 0.41634064 | 342.763015 | 173.879038 | 353.086086 | 184.533575 | Non-sig. |
| PF3D7_1303800 | N/A | 12322.4525 | -0.958023121 | 0.51409866 | -1.86350052 | 0.06239188 | 0.54367134 | 10374.9712 | 6434.34152 | 22164.4586 | 10316.0386 | Non-sig. |
| PF3D7_1403900 | CPPED1 | 374.658857 | -0.958893208 | 0.58556674 | -1.63754726 | 0.10151617 | 0.65830194 | 247.037145 | 216.823632 | 742.27734 | 292.497309 | Non-sig. |
| PF3D7_0509700 | N/A | 76.2147839 | -0.959019774 | 0.57577621 | -1.66561201 | 0.09579072 | 0.64772846 | 118.500492 | 53.7057449 | 82.8960355 | 49.7568632 | Non-sig. |
| PF3D7_1406000 | RBM8A | 510.319498 | -0.959044842 | 0.45342409 | -2.11511663 | 0.03442001 | 0.41634064 | 684.455169 | 414.117454 | 663.48559 | 279.219776 | Non-sig. |
| PF3D7_1253200 | N/A | 118.899677 | -0.960585603 | 0.52303146 | -1.83657328 | 0.06627292 | 0.56240843 | 156.534114 | 94.4744208 | 157.634506 | 66.9556656 | Non-sig. |
| PF3D7_0907300 | N/A | 234.819437 | -0.960802176 | 0.48635204 | -1.97552821 | 0.04820824 | 0.48501926 | 375.142362 | 164.245492 | 245.493349 | 154.396544 | Non-sig. |
| PF3D7_0110200 | ERV1 | 390.367687 | -0.961826814 | 0.49232632 | -1.95363678 | 0.05074419 | 0.49693661 | 379.84306 | 302.850735 | 651.745036 | 227.031917 | Non-sig. |
| PF3D7_1462800 | GAPDH | 56578.7446 | -0.962773569 | 0.56432101 | -1.7060743 | 0.08799425 | 0.62914105 | 35570.187 | 40965.6519 | 114003.057 | 35776.0825 | Non-sig. |
| PF3D7_1333500 | N/A | 215.998623 | -0.963482787 | 0.52698526 | -1.82829173 | 0.06750578 | 0.56795755 | 342.21207 | 186.843363 | 229.030617 | 105.908444 | Non-sig. |
| PF3D7_0723600 | N/A | 189.912256 | -0.963490446 | 0.48307644 | -1.99448859 | 0.04609869 | 0.47851132 | 218.082283 | 135.345874 | 283.97069 | 122.250178 | Non-sig. |
| PF3D7_1035200 | N/A | 137068.309 | -0.963952697 | 0.50208678 | -1.9198926 | 0.05487147 | 0.50446754 | 239805.077 | 115442.472 | 122653.999 | 70371.6867 | Non-sig. |
| PF3D7_0705800 | CRISP | 61.5140194 | -0.964703741 | 0.64412995 | -1.49768497 | 0.13421512 | 0.72178914 | 95.5735419 | 54.9222678 | 67.2023205 | 28.3579474 | Non-sig. |
| PF3D7_1202800 | N/A | 190.036965 | -0.964973225 | 0.48938775 | -1.97179685 | 0.0486328 | 0.48633045 | 284.394856 | 143.902514 | 218.347835 | 113.502654 | Non-sig. |
| PF3D7_0621800 | N/A | 2716.45834 | -0.971188112 | 0.41120062 | -2.36183524 | 0.01818472 | 0.29462587 | 3335.62859 | 1934.39326 | 3859.81352 | 1735.99802 | Non-sig. |
| PF3D7_1348900 | N/A | 202.349162 | -0.971489728 | 0.47265529 | -2.0553874 | 0.0398416 | 0.44531815 | 281.819097 | 148.42185 | 254.255146 | 124.900557 | Non-sig. |
| PF3D7_1104700 | RPC25 | 721.615898 | -0.971732112 | 0.43665797 | -2.22538504 | 0.0260554 | 0.36318813 | 838.036358 | 532.081832 | 1073.6419 | 442.703498 | Non-sig. |
| PF3D7_1004000 | N/A | 6589.62418 | -0.972253555 | 0.4364658 | -2.22755951 | 0.0259099 | 0.36314991 | 7256.14405 | 5051.13389 | 10203.0457 | 3848.17305 | Non-sig. |
| PF3D7_1440200 | SPP | 1046.10561 | -0.97334932 | 0.5342151 | -1.82201762 | 0.06845231 | 0.57292315 | 949.411094 | 470.89988 | 1822.78743 | 941.324015 | Non-sig. |
| PF3D7_0901900 | N/A | 106.014041 | -0.975168726 | 0.62283701 | -1.56568848 | 0.11742157 | 0.69586069 | 78.8796692 | 61.1746143 | 202.008478 | 81.9934011 | Non-sig. |
| PF3D7_1225000 | N/A | 102.036567 | -0.975403311 | 0.53030101 | -1.83933895 | 0.06586535 | 0.56240843 | 131.706346 | 61.1431991 | 138.822452 | 76.4742721 | Non-sig. |
| PF3D7_1124900 | RPL35 | 1927.29302 | -0.976661442 | 0.42428066 | -2.30192309 | 0.02133951 | 0.3235449 | 2496.99779 | 1491.76953 | 2614.64926 | 1105.75551 | Non-sig. |
| PF3D7_1431700 | N/A | 2780.44649 | -0.977541312 | 0.41865824 | -2.33493867 | 0.01954662 | 0.30912841 | 3580.34821 | 2117.77906 | 3795.58474 | 1628.07393 | Non-sig. |
| PF3D7_0804800 | CYP24 | 752.997301 | -0.978291448 | 0.43234086 | -2.26277816 | 0.02364937 | 0.34047378 | 1045.11649 | 572.566871 | 952.827221 | 441.478622 | Non-sig. |
| PF3D7_0219200 | N/A | 5643.39566 | -0.978574916 | 0.45299147 | -2.1602502 | 0.03075331 | 0.39012163 | 6166.43538 | 4565.8 | 8807.81758 | 3033.52966 | Non-sig. |
| PF3D7_1032200 | N/A | 7.32832532 | -0.979695858 | 1.37544145 | -0.7122774 | 0.47629302 | 0.92731133 | 15.4162867 | 5.40596025 | 4.08457749 | 4.40647685 | Non-sig. |
| PF3D7_1139600 | N/A | 41.5698406 | -0.980288375 | 0.71369074 | -1.37354785 | 0.1695821 | 0.77854935 | 57.817915 | 38.641702 | 52.5507215 | 17.2690239 | Non-sig. |
| PF3D7_1302800 | N/A | 4091.54824 | -0.980740454 | 0.42100235 | -2.32953677 | 0.01983065 | 0.31175896 | 5085.13262 | 3132.78017 | 5776.95439 | 2371.32576 | Non-sig. |
| PF3D7_1208600 | TIM10 | 188.339385 | -0.981657261 | 0.52029126 | -1.88674562 | 0.05919455 | 0.52683388 | 198.856786 | 153.682904 | 301.136415 | 99.6814345 | Non-sig. |
| PF3D7_1325000 | LSM6 | 470.818814 | -0.982117114 | 0.46740073 | -2.10123143 | 0.03562066 | 0.42313505 | 491.977859 | 349.26638 | 758.191114 | 283.839904 | Non-sig. |
| PF3D7_1476200 | N/A | 382.18696 | -0.984297648 | 0.51439426 | -1.91350823 | 0.05568302 | 0.50776014 | 686.335495 | 242.491507 | 329.341325 | 270.579512 | Non-sig. |
| PF3D7_1427800 | RRP41 | 269.349013 | -0.98525224 | 0.49659606 | -1.98401138 | 0.04725457 | 0.48052729 | 261.609663 | 182.503662 | 454.026044 | 179.256684 | Non-sig. |
| PF3D7_0614500 | RPL19 | 7255.14637 | -0.987039462 | 0.43538291 | -2.26706066 | 0.02338652 | 0.33880673 | 10348.1765 | 5837.55066 | 8940.93505 | 3893.92321 | Non-sig. |
| PF3D7_1026100 | N/A | 126.930803 | -0.987291706 | 0.68007103 | -1.45174793 | 0.14657171 | 0.7426703 | 267.910547 | 100.914103 | 69.7868693 | 69.1116947 | Non-sig. |
| PF3D7_1346700 | P48/45 | 484.396426 | -0.987608557 | 0.44649186 | -2.21192961 | 0.02697153 | 0.36452847 | 543.012377 | 327.283549 | 744.880529 | 322.409247 | Non-sig. |
| PF3D7_1421400 | RPC34 | 286.742092 | -0.98842679 | 0.45071964 | -2.19299694 | 0.0283076 | 0.37681824 | 371.797809 | 203.994581 | 390.783355 | 180.392623 | Non-sig. |
| PF3D7_0512000 | PFD6 | 872.90841 | -0.989033837 | 0.45783151 | -2.16025723 | 0.03075276 | 0.39012163 | 1273.58928 | 717.180283 | 1048.34942 | 452.514661 | Non-sig. |
| PF3D7_0923100 | N/A | 13.7587899 | -0.989758589 | 1.00595643 | -0.98389807 | 0.32516563 | 0.88694295 | 21.8758244 | 8.60877736 | 14.7583523 | 9.7922055 | Non-sig. |
| PF3D7_1421200 | RPS25 | 2872.75902 | -0.991342253 | 0.44728349 | -2.21636228 | 0.0266667 | 0.36318813 | 3901.50119 | 2387.32467 | 3743.88084 | 1458.32937 | Non-sig. |
| PF3D7_1327000 | N/A | 582.392175 | -0.992145712 | 0.42964498 | -2.30922218 | 0.02093126 | 0.31957865 | 716.414568 | 372.232574 | 833.742367 | 407.17919 | Non-sig. |
| PF3D7_1446200 | LAP | 4134.14643 | -0.99224659 | 0.69020745 | -1.43760631 | 0.15054579 | 0.75169515 | 1493.17508 | 3022.27485 | 9511.21167 | 2509.92409 | Non-sig. |
| PF3D7_1023500 | N/A | 8.98079364 | -0.994627202 | 1.22966327 | -0.80886144 | 0.41859485 | 0.9088998 | 13.3484404 | 8.67934612 | 10.5893139 | 3.30607414 | Non-sig. |
| PF3D7_1475900 | N/A | 27.4512333 | -0.994707161 | 0.87177511 | -1.14101349 | 0.25386431 | 0.85030738 | 56.8130313 | 20.3895528 | 16.4244493 | 16.1778996 | Non-sig. |
| PF3D7_0422400 | RPS19 | 5752.82245 | -0.994832573 | 0.42371952 | -2.34785639 | 0.0188818 | 0.30332278 | 6560.93482 | 4177.37729 | 8761.47862 | 3511.49907 | Non-sig. |
| PF3D7_1037300 | AAC1 | 1966.76934 | -0.995967 | 0.42879808 | -2.32269465 | 0.02019556 | 0.3154509 | 2206.37988 | 1395.74803 | 3033.31402 | 1231.63544 | Non-sig. |
| PF3D7_1360300 | N/A | 34.5889138 | -0.997022433 | 0.74711351 | -1.33449927 | 0.18204029 | 0.79970932 | 30.4881835 | 22.5404996 | 61.583718 | 23.7432543 | Non-sig. |
| PF3D7_0309600 | PfP2 | 6308.09017 | -0.997503893 | 0.41849257 | -2.38356419 | 0.01714589 | 0.28476159 | 8850.0524 | 4784.42127 | 7961.87712 | 3636.00988 | Non-sig. |
| PF3D7_0520000 | RPS9 | 6018.30883 | -0.998436287 | 0.43550138 | -2.2926134 | 0.02187027 | 0.32547388 | 6611.74434 | 4492.3271 | 9431.14323 | 3538.02065 | Non-sig. |
| PF3D7_1232300 | N/A | 140.484975 | -0.99866113 | 0.537716 | -1.85722784 | 0.06327875 | 0.54779542 | 234.264368 | 106.285191 | 140.392018 | 80.9983213 | Non-sig. |
| PF3D7_0532400 | LyMP | 1837.18417 | -0.998961613 | 0.41031057 | -2.43464754 | 0.0149063 | 0.26055806 | 2298.41891 | 1209.17078 | 2599.50522 | 1241.64179 | Non-sig. |
| PF3D7_0518800 | PSOP13 | 17.941031 | -0.999151372 | 1.04667774 | -0.95459312 | 0.3397835 | 0.8930363 | 40.558349 | 11.8425223 | 7.37721173 | 11.986041 | Non-sig. |
| PF3D7_1213300 | N/A | 23.0128093 | -0.999573956 | 0.88700042 | -1.12691486 | 0.25977848 | 0.85030738 | 45.1511178 | 16.2581541 | 16.3051369 | 14.3368286 | Non-sig. |

| | | | | | | | | | | | | |
|---|---|---|---|---|---|---|---|---|---|---|---|---|
| PF3D7_1319800 | Pfg17 | 82.4892928 | -1.000208671 | 0.55435158 | -1.80428577 | 0.07118652 | 0.58066864 | 113.29134 | 49.405098 | 106.703565 | 60.5571682 | Non-sig. |
| PF3D7_MIT01700 | RNA2 | 11.7505211 | -1.000815931 | 1.06354699 | -0.94101712 | 0.34669608 | 0.8930363 | 15.9958376 | 8.63881323 | 15.3404033 | 7.02703012 | Non-sig. |
| PF3D7_1105100 | H2B | 73839.4359 | -1.001091226 | 0.46326392 | -2.16095226 | 0.03069903 | 0.39012163 | 112974.881 | 36203.76 | 83980.1098 | 62198.993 | Non-sig. |
| PF3D7_1014700 | PHB2 | 497.561076 | -1.001913832 | 0.50673934 | -1.97717791 | 0.04802152 | 0.48501926 | 449.579176 | 382.206881 | 877.641523 | 280.816724 | Non-sig. |
| PF3D7_1347200 | NT1 | 7549.22825 | -1.002989205 | 0.52698 | -1.90327754 | 0.05700434 | 0.51625469 | 5608.30959 | 5420.796 | 14536.6216 | 4631.18581 | Non-sig. |
| PF3D7_0304400 | RPL44 | 3239.4627 | -1.003144124 | 0.44359875 | -2.26137725 | 0.02373591 | 0.34079358 | 3567.38423 | 2490.54853 | 5077.32424 | 1822.59382 | Non-sig. |
| PF3D7_1440800 | MFS6 | 321.893396 | -1.004265334 | 0.50346552 | -1.99470529 | 0.04607503 | 0.47851132 | 389.395169 | 149.152671 | 469.764724 | 279.261022 | Non-sig. |
| PF3D7_1149000 | Pf332 | 13346.184 | -1.005589924 | 0.59247782 | -1.69726172 | 0.08964719 | 0.62981332 | 10504.8119 | 4896.65748 | 25130.7761 | 12852.4904 | Non-sig. |
| PF3D7_0721100 | N/A | 6268.83345 | -1.011306438 | 0.46381009 | -2.18043215 | 0.02922544 | 0.38231209 | 6160.32802 | 4732.28827 | 10599.9898 | 3582.72772 | Non-sig. |
| PF3D7_1229700 | N/A | 203.410146 | -1.012653314 | 0.54121679 | -1.87106781 | 0.06133568 | 0.53711809 | 182.892162 | 156.970259 | 360.95004 | 112.828121 | Non-sig. |
| PF3D7_0617600 | N/A | 54.5940937 | -1.017165925 | 0.63746401 | -1.59564447 | 0.11056819 | 0.6811515 | 87.1362919 | 41.8775141 | 59.1057125 | 30.2568563 | Non-sig. |
| PF3D7_1452800 | N/A | 136.04803 | -1.017370705 | 0.5815384 | -1.74944716 | 0.08021375 | 0.60623748 | 244.567954 | 110.605117 | 119.849489 | 69.1695608 | Non-sig. |
| PF3D7_1229500 | CCT3 | 3309.02594 | -1.017494809 | 0.48306775 | -2.10631908 | 0.03517665 | 0.42141755 | 2917.22391 | 2249.95607 | 5942.21201 | 2126.71176 | Non-sig. |
| PF3D7_1438900 | Trx-Px1 | 3120.76044 | -1.017559368 | 0.53528068 | -1.90098279 | 0.05730427 | 0.51808534 | 2334.23274 | 2380.14609 | 6021.25457 | 1747.40837 | Non-sig. |
| PF3D7_1323400 | RPL23 | 7093.88981 | -1.018669097 | 0.42423785 | -2.4011745 | 0.01634254 | 0.27399615 | 9755.33916 | 5481.77372 | 9243.15932 | 3895.28704 | Non-sig. |
| PF3D7_1127100 | dUTPase | 1832.432 | -1.019421619 | 0.42298241 | -2.41008042 | 0.015949 | 0.27257363 | 2393.8524 | 1380.84711 | 2514.49788 | 1040.53061 | Non-sig. |
| PF3D7_1029600 | ADA | 9639.15276 | -1.020956066 | 0.58864594 | -1.73441452 | 0.08284456 | 0.61044573 | 5591.17652 | 6930.15064 | 20237.1431 | 5798.14072 | Non-sig. |
| PF3D7_1416700 | N/A | 48.2575752 | -1.023877837 | 0.68123252 | -1.50297852 | 0.13284458 | 0.72037933 | 78.5963929 | 40.8242797 | 50.8804483 | 22.7291801 | Non-sig. |
| PF3D7_1120900 | HSBP | 446.902542 | -1.023962104 | 0.45627532 | -2.24417594 | 0.02482108 | 0.35350027 | 529.696613 | 339.775151 | 668.546903 | 249.591502 | Non-sig. |
| PF3D7_1023600 | N/A | 55.7513513 | -1.03092679 | 0.80893256 | -1.2744286 | 0.20251156 | 0.81963808 | 118.682377 | 53.6826378 | 31.197044 | 19.4433468 | Non-sig. |
| PF3D7_1461300 | N/A | 4587.4385 | -1.033409612 | 0.40764493 | -2.53507287 | 0.01124239 | 0.21636979 | 6143.57565 | 3253.10097 | 6183.66215 | 2769.41522 | Non-sig. |
| PF3D7_1323100 | N/A | 2829.03798 | -1.033610174 | 0.42511232 | -2.43138138 | 0.01504137 | 0.26127604 | 3182.88413 | 1869.14435 | 4419.43359 | 1844.68987 | Non-sig. |
| PF3D7_1401200 | N/A | 972.725508 | -1.034356956 | 0.44571121 | -2.32068866 | 0.02030365 | 0.3154509 | 1290.67511 | 771.624486 | 1323.76695 | 504.835485 | Non-sig. |
| PF3D7_1337100 | PK6 | 246.84513 | -1.035146858 | 0.51713212 | -2.00170675 | 0.04531628 | 0.47730745 | 244.727737 | 193.255062 | 418.700473 | 130.69725 | Non-sig. |
| PF3D7_1416900 | N/A | 675.36696 | -1.036394077 | 0.51110861 | -2.02773748 | 0.04258705 | 0.46329811 | 611.103216 | 532.828294 | 1204.76912 | 352.767211 | Non-sig. |
| PF3D7_1214800 | OBC13 | 14.7867017 | -1.038936312 | 0.9842517 | -1.05555958 | 0.29116948 | 0.86724628 | 16.7787317 | 11.8152834 | 22.9810552 | 7.57173665 | Non-sig. |
| PF3D7_1222800 | N/A | 80.6139544 | -1.039075395 | 0.60388885 | -1.72064015 | 0.08531614 | 0.61879091 | 139.847735 | 61.1785749 | 77.1871158 | 44.2423922 | Non-sig. |
| PF3D7_0511000 | TCTP | 4321.76893 | -1.043827536 | 0.48938306 | -2.1329458 | 0.03292917 | 0.40615121 | 3879.50098 | 3230.34359 | 7761.11636 | 2416.11479 | Non-sig. |
| PF3D7_0104700 | MFR2 | 11.5754362 | -1.049180526 | 1.09649252 | -0.95685152 | 0.33864221 | 0.8930363 | 18.9305361 | 9.65923348 | 12.3175235 | 5.39445163 | Non-sig. |
| PF3D7_1475000 | MED31 | 17.6250556 | -1.05184588 | 1.03100943 | -1.02020975 | 0.30762899 | 0.87709318 | 12.3389951 | 5.403925 | 35.1392338 | 17.6180685 | Non-sig. |
| PF3D7_1019400 | N/A | 3368.42776 | -1.052490796 | 0.51246908 | -2.05376449 | 0.03999849 | 0.44613054 | 2737.31331 | 2454.56745 | 6353.20421 | 1928.62606 | Non-sig. |
| PF3D7_1001200 | ACBP2 | 2546.35435 | -1.054658252 | 0.43496022 | -2.42472348 | 0.01532005 | 0.26352477 | 2806.75517 | 1783.45508 | 4068.58277 | 1526.62437 | Non-sig. |
| PF3D7_1470200 | N/A | 212.398476 | -1.058761533 | 0.48460039 | -2.18481364 | 0.02890251 | 0.37902356 | 279.305427 | 163.20683 | 294.716823 | 112.364826 | Non-sig. |
| PF3D7_1040300 | RIF | 4.12776301 | -1.059388935 | 1.75167002 | -0.60478796 | 0.5453199 | 0.94121391 | 2.09668934 | 3.22263098 | 9.02777518 | 2.16395653 | Non-sig. |
| PF3D7_1001000 | CSA-L | 406.613267 | -1.060261434 | 0.47507804 | -2.23176267 | 0.02563065 | 0.3611468 | 449.729762 | 309.171991 | 649.443696 | 218.107617 | Non-sig. |
| PF3D7_1336600 | N/A | 362.95822 | -1.060403985 | 0.48522179 | -2.18540057 | 0.02885949 | 0.37902356 | 515.12281 | 298.567773 | 466.223175 | 171.91912 | Non-sig. |
| PF3D7_1144000 | RPS21 | 3474.73552 | -1.064715685 | 0.44466905 | -2.39440027 | 0.01664757 | 0.2773548 | 3970.26744 | 2652.10101 | 5433.08602 | 1843.48761 | Non-sig. |
| PF3D7_0824500 | N/A | 490.960246 | -1.064735211 | 0.46083603 | -2.31044262 | 0.02086366 | 0.31957865 | 797.591642 | 343.517556 | 531.220396 | 291.511389 | Non-sig. |
| PF3D7_1120100 | PGM1 | 6250.89768 | -1.067322765 | 0.5173414 | -2.06309172 | 0.03910391 | 0.43985677 | 4941.56897 | 4465.07331 | 11984.4981 | 3612.45036 | Non-sig. |
| PF3D7_1144400 | N/A | 170.770288 | -1.078106412 | 0.54667794 | -1.97210519 | 0.0485976 | 0.48633045 | 300.223398 | 87.9659241 | 163.459127 | 131.432703 | Non-sig. |
| PF3D7_0604800 | N/A | 207.359654 | -1.079669287 | 0.54618303 | -1.97675362 | 0.04806949 | 0.48501926 | 175.951384 | 129.796935 | 386.892532 | 136.797764 | Non-sig. |
| PF3D7_1142600 | N/A | 5298.43297 | -1.080244389 | 0.41814076 | -2.58344677 | 0.00978186 | 0.19689529 | 7196.97201 | 3887.71289 | 7191.68379 | 2917.36319 | Non-sig. |
| PF3D7_0726000 | N/A | 45019.7328 | -1.081286108 | 0.61018345 | -1.77206724 | 0.0763834 | 0.59693801 | 28385.959 | 18818.4574 | 93899.6803 | 38974.8343 | Non-sig. |
| PF3D7_0825200 | IF3a | 500.743825 | -1.082772133 | 0.44308866 | -2.44369183 | 0.01453784 | 0.25674317 | 769.764891 | 340.489953 | 590.946924 | 301.773531 | Non-sig. |
| PF3D7_0505000 | N/A | 742.44733 | -1.085070751 | 0.50826836 | -2.13483829 | 0.03277422 | 0.40615121 | 813.911405 | 318.533881 | 1204.35017 | 632.993867 | Non-sig. |
| PF3D7_0201600 | RLP1 | 386.608512 | -1.088187343 | 0.44136358 | -2.46551233 | 0.01368175 | 0.24576103 | 507.226446 | 267.193039 | 544.49257 | 227.521993 | Non-sig. |
| PF3D7_1038200 | N/A | 36.4181599 | -1.088682655 | 0.69382501 | -1.56910264 | 0.11662402 | 0.69346137 | 49.9060412 | 22.6099468 | 49.1794885 | 23.9771629 | Non-sig. |
| PF3D7_1463600 | N/A | 15.3408037 | -1.089101539 | 0.96976983 | -1.12305158 | 0.26141559 | 0.85030738 | 16.4216614 | 10.8291374 | 25.2892402 | 8.82317568 | Non-sig. |
| PF3D7_0903900 | RPL32 | 4962.55161 | -1.092284738 | 0.44303755 | -2.46544507 | 0.01368432 | 0.24576103 | 7203.61478 | 3883.4706 | 6309.02512 | 2454.09593 | Non-sig. |
| PF3D7_1460400 | UCHL3 | 483.801863 | -1.094398934 | 0.47280242 | -2.3147067 | 0.02062899 | 0.31863668 | 492.049412 | 315.750187 | 825.742552 | 301.665301 | Non-sig. |
| PF3D7_0917700 | N/A | 47.2701779 | -1.095552186 | 0.65810933 | -1.66469632 | 0.09597336 | 0.64772846 | 55.6813432 | 24.6920009 | 73.0704901 | 35.6368776 | Non-sig. |
| PF3D7_1458100 | PET117 | 75.522864 | -1.09631216 | 0.57473141 | -1.90752088 | 0.05645317 | 0.51301697 | 108.29431 | 55.962892 | 97.5631136 | 40.2711405 | Non-sig. |
| PF3D7_1325300 | N/A | 12.8596926 | -1.097318136 | 1.13659324 | -0.96544489 | 0.33432203 | 0.89204385 | 27.7733114 | 10.8058404 | 7.35599896 | 5.50361959 | Non-sig. |
| PF3D7_0928100 | N/A | 75.3435432 | -1.10080323 | 0.79166872 | -1.39048468 | 0.16438175 | 0.77284241 | 170.397688 | 66.5388759 | 35.311064 | 29.1265443 | Non-sig. |

| | | | | | | | | | | | | |
|---|---|---|---|---|---|---|---|---|---|---|---|---|
| PF3D7_0721600 | N/A | 8645.18645 | -1.101778335 | 0.41667817 | -2.644195 | 0.00818855 | 0.17215445 | 10944.7865 | 6159.58526 | 12644.6038 | 4831.7702 | Non-sig. |
| PF3D7_0825800 | N/A | 69.7015616 | -1.104255285 | 0.63470518 | -1.73979246 | 0.08189547 | 0.60888892 | 128.038797 | 38.6596448 | 62.3858381 | 49.7219666 | Non-sig. |
| PF3D7_1134400 | N/A | 255.518721 | -1.105602784 | 0.47405365 | -2.33223135 | 0.01968852 | 0.31044583 | 381.861797 | 186.876307 | 316.010806 | 137.325976 | Non-sig. |
| PF3D7_0104500 | N/A | 541.26944 | -1.111218549 | 0.44402535 | -2.50260159 | 0.01232842 | 0.22917891 | 645.141493 | 374.83266 | 834.752537 | 310.351069 | Non-sig. |
| PF3D7_1243100 | N/A | 601.225932 | -1.113622001 | 0.45924707 | -2.42488644 | 0.01531318 | 0.26352477 | 985.974368 | 422.2268 | 658.966303 | 337.736257 | Non-sig. |
| PF3D7_1368600 | TIM9 | 257.618821 | -1.115113198 | 0.47157338 | -2.36466527 | 0.01804637 | 0.29328125 | 410.419186 | 160.000826 | 294.69927 | 165.356003 | Non-sig. |
| PF3D7_1304500 | N/A | 1103.50216 | -1.116089262 | 0.44873931 | -2.48716622 | 0.01287652 | 0.2379425 | 1191.40975 | 748.453279 | 1828.95211 | 645.193492 | Non-sig. |
| PF3D7_0320200 | N/A | 159.914577 | -1.12147669 | 0.51181606 | -2.19117135 | 0.0284394 | 0.37697071 | 261.804134 | 90.1594545 | 176.5431 | 111.151619 | Non-sig. |
| PF3D7_1408600 | N/A | 6566.08171 | -1.12345885 | 0.42027889 | -2.67312703 | 0.00751478 | 0.16451776 | 8303.3373 | 4683.80824 | 9698.30215 | 3578.87916 | Non-sig. |
| PF3D7_1216500 | MDV1 | 2198.83491 | -1.129552747 | 0.61455329 | -1.83800619 | 0.06606149 | 0.56240843 | 4831.2452 | 1575.74382 | 1205.37211 | 1182.97851 | Non-sig. |
| PF3D7_1357900 | N/A | 559.933949 | -1.129876943 | 0.51067252 | -2.21252741 | 0.02693024 | 0.36452847 | 526.405533 | 422.971338 | 1010.6916 | 279.667321 | Non-sig. |
| PF3D7_0831300 | GEXP13 | 1402.25113 | -1.13298764 | 0.44779716 | -2.53013586 | 0.01140184 | 0.2165123 | 2354.35243 | 924.718392 | 1498.23499 | 831.698699 | Non-sig. |
| PF3D7_0924800 | TRF1 | 2.61022738 | -1.133613867 | 2.08211588 | -0.54445282 | 0.5861299 | 0.95246063 | 3.10303093 | 1.08848341 | 4.06367706 | 2.18571811 | Non-sig. |
| PF3D7_1025600 | COX17 | 424.276326 | -1.136275106 | 0.46074498 | -2.46616925 | 0.01365668 | 0.24576103 | 523.457478 | 312.041091 | 642.94701 | 218.659727 | Non-sig. |
| PF3D7_0532000 | N/A | 26032.1127 | -1.136365607 | 0.56807851 | -2.00036717 | 0.04546063 | 0.47763064 | 19337.1821 | 11282.9181 | 52233.3021 | 21275.0484 | Non-sig. |
| PF3D7_0112300 | N/A | 28.3357499 | -1.138519141 | 0.8603552 | -1.32331291 | 0.18573134 | 0.80052251 | 17.8791997 | 16.0986313 | 59.9454924 | 19.4196761 | Non-sig. |
| PF3D7_0819300 | N/A | 118.756951 | -1.139012807 | 0.65266164 | -1.74518117 | 0.08095334 | 0.60749403 | 91.6168632 | 47.2056183 | 234.907618 | 101.297705 | Non-sig. |
| PF3D7_0600400 | VAR | 14.7246474 | -1.142759012 | 1.28539531 | -0.88903312 | 0.37398528 | 0.89654416 | 38.9845827 | 9.6542239 | 1.64286561 | 8.6169172 | Non-sig. |
| PF3D7_0308500 | N/A | 1079.03442 | -1.145801024 | 0.45274904 | -2.53076414 | 0.01138144 | 0.2165123 | 1223.16079 | 778.473812 | 1749.39091 | 565.112153 | Non-sig. |
| PF3D7_1322800 | C2AP2 | 128.293415 | -1.149511183 | 0.52364046 | -2.1952299 | 0.02814711 | 0.37562567 | 155.069954 | 92.3962861 | 198.591769 | 67.1156508 | Non-sig. |
| PF3D7_1236650 | N/A | 53.267647 | -1.149511974 | 0.62570131 | -1.83715769 | 0.06618662 | 0.56240843 | 68.4597928 | 37.3989538 | 78.3637032 | 28.848138 | Non-sig. |
| PF3D7_0110900 | AKLP1 | 149.159944 | -1.149744049 | 0.495258 | -2.32150526 | 0.02025959 | 0.3154509 | 188.792173 | 86.9993873 | 222.430331 | 98.4178832 | Non-sig. |
| PF3D7_1033400 | HAD1 | 3081.2359 | -1.151825144 | 0.49870361 | -2.30963867 | 0.02090817 | 0.31957865 | 2640.79263 | 1981.88452 | 5858.62177 | 1843.64469 | Non-sig. |
| PF3D7_0521900 | N/A | 386.108061 | -1.155839723 | 0.69733931 | -1.65749974 | 0.09741848 | 0.65042594 | 909.330217 | 266.024112 | 156.894904 | 212.183012 | Non-sig. |
| PF3D7_0414300 | N/A | 538.237969 | -1.159218053 | 0.45681558 | -2.53760621 | 0.01116135 | 0.21636979 | 595.627288 | 354.461454 | 891.331961 | 311.531171 | Non-sig. |
| PF3D7_1321000 | N/A | 13.9857296 | -1.166777774 | 1.01549613 | -1.14897314 | 0.25056706 | 0.8494288 | 23.1340359 | 10.7329102 | 15.6018081 | 6.47416409 | Non-sig. |
| PF3D7_0700800 | MC-2TM | 99.7211432 | -1.17054084 | 0.5523403 | -2.11923851 | 0.03407032 | 0.41400122 | 133.336738 | 75.1635484 | 142.840261 | 47.5440254 | Non-sig. |
| PF3D7_1349000 | N/A | 17.5459977 | -1.172658583 | 0.91141175 | -1.28663974 | 0.19821989 | 0.81528784 | 27.2967629 | 10.7377074 | 21.3438751 | 10.8056454 | Non-sig. |
| PF3D7_1349400 | N/A | 241.223881 | -1.177551586 | 0.47659446 | -2.47076222 | 0.01348254 | 0.24576103 | 295.411847 | 167.445008 | 373.606084 | 128.432585 | Non-sig. |
| PF3D7_1213600 | DLC8 | 835.880483 | -1.177869934 | 0.45537579 | -2.58658882 | 0.00969312 | 0.19600817 | 889.912641 | 508.860015 | 1428.58655 | 516.162723 | Non-sig. |
| PF3D7_0828000 | ROM3 | 44.8896751 | -1.180992988 | 0.73724118 | -1.60190861 | 0.10917582 | 0.6780932 | 33.4262528 | 29.0329135 | 91.050862 | 26.048672 | Non-sig. |
| PF3D7_0222100 | MC-2TM | 134.609546 | -1.181450478 | 0.51410411 | -2.29807631 | 0.02155745 | 0.3235449 | 166.784684 | 93.4409164 | 206.839586 | 71.3729979 | Non-sig. |
| PF3D7_0414200 | N/A | 241.509423 | -1.181602169 | 0.60743365 | -1.94523661 | 0.05174651 | 0.50119377 | 158.401716 | 152.708466 | 512.127041 | 142.800469 | Non-sig. |
| PF3D7_0810000 | N/A | 551.932205 | -1.182598662 | 0.46418915 | -2.54766546 | 0.01084464 | 0.21201073 | 656.673661 | 402.758351 | 875.790848 | 272.505961 | Non-sig. |
| PF3D7_0407400 | N/A | 223.798435 | -1.183934365 | 0.53299937 | -2.22126785 | 0.02633283 | 0.36318813 | 375.484558 | 177.221252 | 246.232595 | 96.2553348 | Non-sig. |
| PF3D7_1144300 | RPL41 | 1166.88675 | -1.184940494 | 0.48289608 | -2.45382093 | 0.01413474 | 0.25214083 | 1183.90223 | 848.121927 | 2057.64955 | 577.873303 | Non-sig. |
| PF3D7_0512700 | OPRT | 516.362083 | -1.185083514 | 0.43107348 | -2.74914504 | 0.00597509 | 0.14324004 | 700.568085 | 337.12332 | 733.95648 | 293.800446 | Non-sig. |
| PF3D7_1447600 | N/A | 5.37186321 | -1.188293575 | 1.52412881 | -0.7796543 | 0.43559439 | 0.91522571 | 9.23994934 | 4.32905162 | 5.71294728 | 2.20550459 | Non-sig. |
| PF3D7_0830700 | N/A | 284.2649 | -1.191266632 | 0.47031918 | -2.53288976 | 0.01131265 | 0.21636979 | 336.561419 | 191.035212 | 454.110418 | 155.352549 | Non-sig. |
| PF3D7_0512100 | N/A | 150.238749 | -1.194218977 | 0.53694075 | -2.22411685 | 0.02614058 | 0.36318813 | 159.075506 | 107.666413 | 259.011973 | 75.2011018 | Non-sig. |
| PF3D7_0808000 | N/A | 159.160106 | -1.196259385 | 0.54980027 | -2.17580719 | 0.02956968 | 0.38502203 | 158.036866 | 77.2349938 | 285.034438 | 116.334125 | Non-sig. |
| PF3D7_0221200 | N/A | 600.113945 | -1.200210826 | 0.45896704 | -2.6150262 | 0.00892206 | 0.18464485 | 678.710707 | 409.918535 | 993.70752 | 318.119017 | Non-sig. |
| PF3D7_1126800 | SR-MG | 231.80075 | -1.202456713 | 0.84274243 | -1.42683774 | 0.15362667 | 0.75442818 | 601.346989 | 127.667021 | 45.1751385 | 153.013851 | Non-sig. |
| PF3D7_1229200 | N/A | 5.38201214 | -1.203100738 | 1.65687022 | -0.72612853 | 0.46775998 | 0.92538923 | 12.5828397 | 1.07414561 | 2.46221409 | 5.40884916 | Non-sig. |
| PF3D7_0622000 | N/A | 80.0767954 | -1.203815193 | 0.59083566 | -2.0374789 | 0.04160207 | 0.45538799 | 83.7482057 | 53.731935 | 139.495995 | 43.3310461 | Non-sig. |
| PF3D7_1130100 | RPL38 | 3394.22863 | -1.207005803 | 0.44567573 | -2.7082601 | 0.0067637 | 0.15295744 | 5311.78424 | 2477.54049 | 4161.68482 | 1625.90496 | Non-sig. |
| PF3D7_API01300 | RPL4 | 21.5090961 | -1.207353801 | 1.06128769 | -1.13763102 | 0.25527459 | 0.85030738 | 8.30789047 | 6.46200023 | 51.6277746 | 19.638719 | Non-sig. |
| PF3D7_0513300 | PNP | 7936.15583 | -1.215247895 | 0.53846724 | -2.25686505 | 0.02401651 | 0.34389043 | 6066.15623 | 5508.4537 | 16121.8238 | 4048.18963 | Non-sig. |
| PF3D7_0611700 | RPL39 | 2110.85484 | -1.217942832 | 0.45305528 | -2.68828749 | 0.00718195 | 0.15922925 | 2338.77614 | 1460.08949 | 3566.02508 | 1078.52866 | Non-sig. |
| PF3D7_0519400 | RPS24 | 6326.61164 | -1.2215869 | 0.43285995 | -2.82212968 | 0.00477059 | 0.1245053 | 9349.79258 | 4537.80945 | 8361.81074 | 3057.03378 | Non-sig. |
| PF3D7_1466600 | N/A | 37.8835092 | -1.222785259 | 0.83950673 | -1.45655206 | 0.14524007 | 0.74131203 | 85.7058118 | 26.8779037 | 20.5087468 | 18.4415746 | Non-sig. |
| PF3D7_1426100 | BTF3 | 2263.35765 | -1.223393847 | 0.41557795 | -2.94383727 | 0.0032417 | 0.094887 | 3212.76068 | 1507.00172 | 3125.97876 | 1207.68946 | Non-sig. |
| PF3D7_0110000 | N/A | 834.00237 | -1.224360791 | 0.4231945 | -2.89313968 | 0.00381412 | 0.10470046 | 1273.19064 | 495.041954 | 1063.04381 | 504.733082 | Non-sig. |
| PF3D7_0706600 | N/A | 51.1405295 | -1.224402983 | 0.62580461 | -1.95652598 | 0.05040322 | 0.4954291 | 72.628227 | 31.1168222 | 70.6308813 | 30.1861875 | Non-sig. |

| | | | | | | | | | | | | |
|---|---|---|---|---|---|---|---|---|---|---|---|---|
| PF3D7_1473600 | N/A | 115.739042 | -1.225291178 | 0.55198963 | -2.21977211 | 0.02643424 | 0.36318813 | 122.983395 | 78.358768 | 201.167551 | 60.4464539 | Non-sig. |
| PF3D7_0801100 | N/A | 122.943164 | -1.226254021 | 0.50734797 | -2.41698815 | 0.01564953 | 0.26832101 | 172.832574 | 70.7934835 | 171.685693 | 76.4609043 | Non-sig. |
| PF3D7_1434800 | N/A | 1648.52366 | -1.231694524 | 0.45630716 | -2.69926628 | 0.00694925 | 0.15600488 | 1796.48165 | 1118.29236 | 2828.15417 | 851.16647 | Non-sig. |
| PF3D7_0621200 | PDX1 | 870.065154 | -1.239260084 | 0.42481283 | -2.91719083 | 0.003532 | 0.10006693 | 1155.48951 | 470.119844 | 1289.17059 | 565.480664 | Non-sig. |
| PF3D7_1040200 | N/A | 21.6380664 | -1.239328774 | 1.03366142 | -1.19896975 | 0.23053971 | 0.83828834 | 7.34315131 | 13.9622068 | 53.3520314 | 11.8948763 | Non-sig. |
| PF3D7_1309900 | N/A | 46.2667158 | -1.242824343 | 0.69194552 | -1.79613035 | 0.07247379 | 0.58363337 | 47.1286298 | 21.4901923 | 82.8767879 | 33.5712529 | Non-sig. |
| PF3D7_0925000 | N/A | 98.9263663 | -1.243096091 | 0.64367921 | -1.93123542 | 0.05345395 | 0.50202873 | 203.648501 | 68.7239173 | 74.7017958 | 48.6312513 | Non-sig. |
| PF3D7_1149400 | N/A | 741.54448 | -1.243360165 | 0.44894197 | -2.76953426 | 0.00561365 | 0.13705584 | 1099.15453 | 528.020303 | 986.243755 | 352.759331 | Non-sig. |
| PF3D7_0213700 | N/A | 140.575992 | -1.243557978 | 0.51015245 | -2.4376203 | 0.01478429 | 0.25936156 | 167.882396 | 78.3980599 | 227.371043 | 88.6524688 | Non-sig. |
| PF3D7_1460300 | RPL29 | 2605.24692 | -1.243830195 | 0.43961537 | -2.8293601 | 0.00466412 | 0.12247164 | 3493.41832 | 1873.41471 | 3833.68016 | 1220.4745 | Non-sig. |
| PF3D7_1207300 | LIMP | 9.23098544 | -1.245563837 | 1.19422801 | -1.04298662 | 0.29695449 | 0.87301237 | 15.3941741 | 5.42109007 | 10.5952839 | 5.5133937 | Non-sig. |
| PF3D7_0322000 | CYP19A | 8483.90566 | -1.247669988 | 0.45096403 | -2.76667296 | 0.00566315 | 0.13763022 | 9096.92321 | 5588.47619 | 14782.2847 | 4467.93857 | Non-sig. |
| PF3D7_1105000 | H4 | 44699.6448 | -1.25938629 | 0.43354539 | -2.9048545 | 0.00367424 | 0.1021834 | 57044.5925 | 21364.9145 | 69072.2198 | 31316.8523 | Non-sig. |
| PF3D7_1024300 | ATPTG10 | 769.331827 | -1.265649882 | 0.43118718 | -2.93526788 | 0.0033326 | 0.0959571 | 1018.09034 | 504.075682 | 1155.25985 | 399.901432 | Non-sig. |
| PF3D7_1453700 | P23 | 5406.18592 | -1.268276813 | 0.41809333 | -3.03347772 | 0.00241753 | 0.0762384 | 7321.01074 | 3604.29119 | 7959.79229 | 2739.64946 | Non-sig. |
| PF3D7_1130310 | N/A | 1.87265659 | -1.270001039 | 2.42160751 | -0.52444545 | 0.59996875 | 0.95655566 | 2.02086993 | 1.10175948 | 3.26879631 | 1.09920064 | Non-sig. |
| PF3D7_MIT01400 | COX3 | 614.202951 | -1.27549633 | 0.59316138 | -2.15033612 | 0.03152864 | 0.39723726 | 412.215108 | 289.328124 | 1326.19093 | 429.077638 | Non-sig. |
| PF3D7_1001300 | N/A | 268.374546 | -1.279173763 | 0.5401982 | -2.36797115 | 0.01788593 | 0.29246808 | 241.317986 | 179.351688 | 518.76223 | 134.066281 | Non-sig. |
| PF3D7_1103100 | RPP1 | 6747.87429 | -1.284046773 | 0.42287609 | -3.03646103 | 0.00239373 | 0.07610251 | 9384.81835 | 4573.83929 | 9749.37593 | 3283.46358 | Non-sig. |
| PF3D7_0714400 | N/A | 96.5147884 | -1.289842968 | 0.62299057 | -2.07040527 | 0.03841441 | 0.43487081 | 191.222979 | 66.5595453 | 82.9238608 | 45.3527683 | Non-sig. |
| PF3D7_0102300 | N/A | 1287.0818 | -1.291070324 | 0.52012506 | -2.48223058 | 0.01305628 | 0.23935 | 1148.12155 | 900.759765 | 2506.49702 | 592.948849 | Non-sig. |
| PF3D7_1439900 | TIM | 4718.87594 | -1.292672121 | 0.53874599 | -2.39940925 | 0.01642155 | 0.27445228 | 3614.13017 | 3049.22859 | 9789.70633 | 2422.43868 | Non-sig. |
| PF3D7_1149100 | N/A | 4504.96587 | -1.295322001 | 0.47263008 | -2.74066769 | 0.00613145 | 0.1443399 | 8116.5586 | 3072.54447 | 4686.79588 | 2143.96455 | Non-sig. |
| PF3D7_1334500 | MSRP6 | 1048.33537 | -1.298268888 | 0.42511262 | -3.05394105 | 0.00225856 | 0.07432221 | 1665.50169 | 583.677563 | 1315.74902 | 628.413221 | Non-sig. |
| PF3D7_1477100 | N/A | 5.61135719 | -1.304026241 | 1.5500241 | -0.84129417 | 0.40018316 | 0.90645858 | 9.42539258 | 5.37260131 | 6.56445685 | 1.08297802 | Non-sig. |
| PF3D7_0705700 | N/A | 5231.24913 | -1.305129647 | 0.42478407 | -3.07245432 | 0.00212306 | 0.07164324 | 6679.15322 | 3425.35093 | 8217.3584 | 2603.13396 | Non-sig. |
| PF3D7_1015900 | ENO | 15049.8039 | -1.305154142 | 0.59118488 | -2.20769202 | 0.02726575 | 0.36756728 | 9221.28541 | 9635.00081 | 33634.7593 | 7708.16987 | Non-sig. |
| PF3D7_API02900 | TUFA | 888.631357 | -1.318507926 | 0.8114054 | -1.62496815 | 0.10416935 | 0.6615241 | 328.31926 | 180.43845 | 2208.74498 | 837.022734 | Non-sig. |
| PF3D7_0928150 | N/A | 7.66107066 | -1.318710488 | 1.41608722 | -0.93123536 | 0.35173183 | 0.8930363 | 18.6800396 | 5.45026427 | 3.24675669 | 3.2672221 | Non-sig. |
| PF3D7_1452100 | LRR3 | 31.074529 | -1.322833602 | 0.78401347 | -1.68725877 | 0.0915536 | 0.6361181 | 30.4397883 | 13.9598692 | 58.2829706 | 21.6154877 | Non-sig. |
| PF3D7_0421100 | VAR | 337.946705 | -1.323047916 | 0.4878127 | -2.71220475 | 0.00668373 | 0.15241957 | 467.939643 | 140.51205 | 497.811377 | 245.523752 | Non-sig. |
| PF3D7_0719000 | N/A | 380.736037 | -1.323391867 | 0.4611457 | -2.86979118 | 0.00410743 | 0.11102634 | 497.809716 | 254.459448 | 590.275467 | 180.399515 | Non-sig. |
| PF3D7_0106400 | N/A | 396.00565 | -1.328160501 | 0.48751027 | -2.7243744 | 0.00644234 | 0.14904599 | 721.935548 | 239.681625 | 411.064959 | 211.340469 | Non-sig. |
| PF3D7_1302400 | N/A | 7.69804259 | -1.332638486 | 1.36117085 | -0.97903837 | 0.32756102 | 0.88852997 | 15.5150145 | 2.15657891 | 6.55073143 | 6.5698455 | Non-sig. |
| PF3D7_0625900 | N/A | 50.6966082 | -1.343265206 | 0.6932736 | -1.93756866 | 0.05267587 | 0.50193661 | 96.3122275 | 21.4926718 | 49.2340077 | 35.7475258 | Non-sig. |
| PF3D7_API02600 | RPS11 | 5.78700397 | -1.344893893 | 1.5021543 | -0.89531009 | 0.37062135 | 0.89654416 | 5.14124796 | 2.16157 | 11.4406808 | 4.40451714 | Non-sig. |
| PF3D7_MIT03200 | LSUA | 7324.63787 | -1.347241159 | 0.41179958 | -3.27159434 | 0.00106943 | 0.04292299 | 9895.67961 | 4537.51761 | 11136.3001 | 3729.05414 | Sig. genes |
| PF3D7_0532500 | N/A | 135.471074 | -1.358261238 | 0.51330298 | -2.64611993 | 0.00814209 | 0.17185982 | 220.390143 | 83.0197797 | 169.513385 | 68.9609896 | Non-sig. |
| PF3D7_0604900 | N/A | 270.657012 | -1.361772669 | 0.47057231 | -2.89386483 | 0.00380532 | 0.10470046 | 434.043571 | 170.912559 | 345.405003 | 132.266913 | Non-sig. |
| PF3D7_MIT03300 | RNA7 | 255.938276 | -1.363400033 | 0.45969143 | -2.96590263 | 0.00301796 | 0.09084746 | 358.841023 | 155.260763 | 378.304678 | 131.34664 | Non-sig. |
| PF3D7_0617800 | H2A | 137532.71 | -1.366491763 | 0.54175845 | -2.52232664 | 0.01165814 | 0.21980365 | 145956.887 | 44642.3947 | 250438.71 | 109092.847 | Non-sig. |
| PF3D7_0319600 | EF-1delta | 2345.94852 | -1.372753565 | 0.44774816 | -3.06590553 | 0.00217012 | 0.07230999 | 2721.09279 | 1491.17273 | 4048.44954 | 1123.07902 | Non-sig. |
| PF3D7_1141500 | N/A | 8.79424986 | -1.380628891 | 1.40871669 | -0.98006143 | 0.3270558 | 0.88852997 | 23.0146267 | 6.44913363 | 2.46139742 | 3.25184163 | Non-sig. |
| PF3D7_1404100 | N/A | 830.012023 | -1.388978911 | 0.42869925 | -3.23998444 | 0.00119536 | 0.0463197 | 1259.94585 | 511.104026 | 1142.73345 | 406.264763 | Sig. genes |
| PF3D7_MIT01000 | SSUB | 314.323528 | -1.394489331 | 0.51944931 | -2.68455324 | 0.00726268 | 0.1603237 | 297.959871 | 181.698469 | 612.718138 | 164.917633 | Non-sig. |
| PF3D7_0307100 | RPS12 | 4751.3944 | -1.4010608 | 0.48176915 | -2.90815798 | 0.00363565 | 0.10191351 | 4727.85702 | 2980.77971 | 9057.61577 | 2239.32512 | Non-sig. |
| PF3D7_1426000 | RPL21 | 4400.57053 | -1.40449634 | 0.47694302 | -2.94478856 | 0.00323175 | 0.094887 | 4330.57699 | 2582.81539 | 8445.35173 | 2243.538 | Non-sig. |
| PF3D7_MIT00900 | LSUG | 187.72591 | -1.408362204 | 0.49698915 | -2.83378863 | 0.00459997 | 0.12185333 | 240.939447 | 85.9425166 | 304.435201 | 119.586476 | Non-sig. |
| PF3D7_1215800 | RSPH9 | 22.8571088 | -1.420448253 | 0.87391936 | -1.6253768 | 0.1040823 | 0.6615241 | 46.1158698 | 12.8914088 | 20.5167815 | 11.904375 | Non-sig. |
| PF3D7_0709050 | N/A | 95.563858 | -1.421282403 | 0.58151348 | -2.44410912 | 0.01452103 | 0.25674317 | 101.971637 | 44.1337617 | 176.249048 | 59.9009848 | Non-sig. |
| PF3D7_MIT01300 | RNA10 | 172.034704 | -1.42663711 | 0.51045541 | -2.79483199 | 0.00519267 | 0.12795703 | 194.59155 | 93.5317924 | 306.865953 | 93.1495216 | Non-sig. |
| PF3D7_0803900 | N/A | 7.1456882 | -1.441699012 | 1.3650044 | -1.05618635 | 0.29088309 | 0.86724628 | 10.3266851 | 5.55748458 | 10.5518656 | 2.14671752 | Non-sig. |
| PF3D7_1466500 | N/A | 84.7123681 | -1.451318158 | 0.62648928 | -2.31658897 | 0.02052613 | 0.317975 | 172.75689 | 48.3567851 | 75.493719 | 42.242078 | Non-sig. |
| PF3D7_0627500 | DJ1 | 2252.8283 | -1.452211518 | 0.46969353 | -3.0918278 | 0.00198928 | 0.06799495 | 2353.70751 | 1327.93955 | 4245.59893 | 1084.0672 | Non-sig. |

| | | | | | | | | | | | | |
|---|---|---|---|---|---|---|---|---|---|---|---|---|
| PF3D7_MIT04200 | RNA8 | 163.095651 | -1.488677298 | 0.53007285 | -2.80843906 | 0.00497823 | 0.12701704 | 246.969576 | 111.106341 | 233.976923 | 60.3297659 | Non-sig. |
| PF3D7_1443300 | LSM5 | 798.292996 | -1.490353357 | 0.43067238 | -3.46052686 | 0.00053912 | 0.02550227 | 1206.32458 | 472.918831 | 1148.6781 | 365.250482 | Sig. genes |
| PF3D7_0523500 | N/A | 28.9775691 | -1.498053603 | 0.77801629 | -1.92547845 | 0.05416953 | 0.50202873 | 51.1912004 | 12.9030199 | 34.4524592 | 17.3635968 | Non-sig. |
| PF3D7_1201000 | N/A | 3967.78319 | -1.508072073 | 0.4036358 | -3.73621977 | 0.00018681 | 0.010642 | 5853.91436 | 2121.25787 | 5888.73235 | 2007.2282 | Sig. genes |
| PF3D7_1248800 | N/A | 17.8477371 | -1.509105698 | 0.95485698 | -1.58045208 | 0.11400337 | 0.68635213 | 30.8863637 | 13.0369229 | 21.9752927 | 5.4923692 | Non-sig. |
| PF3D7_1474100 | N/A | 573.069592 | -1.519622034 | 0.5323806 | -2.85439033 | 0.00431195 | 0.11537737 | 536.821461 | 361.797308 | 1162.54801 | 231.111593 | Non-sig. |
| PF3D7_0209000 | P230 | 370.974895 | -1.524134269 | 0.5597986 | -2.72264753 | 0.00647611 | 0.14917581 | 529.265097 | 101.8873 | 571.781018 | 280.966167 | Non-sig. |
| PF3D7_1221000 | SET10 | 4515.67102 | -1.527947747 | 0.48856751 | -3.12740352 | 0.00176358 | 0.06146994 | 4315.91318 | 2240.51304 | 9095.756 | 2410.50186 | Non-sig. |
| PF3D7_0502800 | N/A | 62.7432374 | -1.528263397 | 0.66573913 | -2.29558898 | 0.02169939 | 0.32475533 | 76.3769944 | 45.1421112 | 109.945109 | 19.508735 | Non-sig. |
| PF3D7_1365200 | N/A | 333.957584 | -1.535499554 | 0.49073886 | -3.12895451 | 0.00175429 | 0.06146994 | 366.891445 | 184.768296 | 626.190499 | 157.980095 | Non-sig. |
| PF3D7_1034300 | TLAP2 | 30.6700143 | -1.545774707 | 0.77794882 | -1.98698766 | 0.04692376 | 0.48052729 | 57.7754708 | 12.8832249 | 33.6609638 | 18.3603978 | Non-sig. |
| PF3D7_MIT02000 | RNA3 | 26.9963431 | -1.572640401 | 0.84381124 | -1.86373484 | 0.06235895 | 0.54367134 | 23.0804757 | 13.7436234 | 57.6484482 | 13.512825 | Non-sig. |
| PF3D7_1402500 | N/A | 4011.35636 | -1.600051439 | 0.44213976 | -3.61888156 | 0.00029588 | 0.01507276 | 4701.31282 | 2113.32949 | 7364.01106 | 1866.77206 | Sig. genes |
| PF3D7_API02200 | RPL6 | 4.33874146 | -1.602959713 | 2.05123649 | -0.78146022 | 0.43453187 | 0.91522571 | 0 | 1.07125792 | 13.02188 | 3.2618279 | Non-sig. |
| PF3D7_API02800 | RPS7 | 7.79637782 | -1.607552784 | 1.53209171 | -1.04925363 | 0.2940614 | 0.87170077 | 14.4562664 | 0 | 9.01801177 | 7.71123306 | Non-sig. |
| PF3D7_1215300 | CPN10 | 438.315435 | -1.620017886 | 0.44014973 | -3.68060636 | 0.00023268 | 0.01255588 | 654.1475 | 236.433889 | 668.738459 | 193.941891 | Sig. genes |
| PF3D7_1229400 | MIF | 4456.00383 | -1.629108586 | 0.54637541 | -2.98166527 | 0.00286685 | 0.08850955 | 3532.34238 | 2439.70185 | 9936.97352 | 1914.99757 | Non-sig. |
| PF3D7_1371800 | N/A | 380.228371 | -1.647470792 | 0.44475832 | -3.70419333 | 0.00021206 | 0.01158266 | 620.790284 | 173.84341 | 532.160455 | 194.119335 | Sig. genes |
| PF3D7_API02700 | RPS12 | 398.65108 | -1.677252176 | 0.59766394 | -2.80634661 | 0.00501068 | 0.12701704 | 573.956012 | 85.4866413 | 640.750847 | 294.410819 | Non-sig. |
| PF3D7_1012400 | HGPRT | 17803.5278 | -1.73363895 | 0.41393801 | -4.18816076 | 2.8122E-05 | 0.00229219 | 26196.6999 | 9250.65131 | 28554.2115 | 7212.54854 | Sig. genes |
| PF3D7_API02300 | RPS5 | 32.9893167 | -1.746821462 | 0.87231299 | -2.00251685 | 0.04522917 | 0.47730745 | 30.2454677 | 6.45452714 | 71.3464857 | 23.9107864 | Non-sig. |
| PF3D7_MIT02700 | RNA5 | 21.3680118 | -1.760682836 | 0.90868229 | -1.93762205 | 0.05266935 | 0.50193661 | 20.7749782 | 7.95139014 | 45.194028 | 11.551651 | Non-sig. |
| PF3D7_0531800 | N/A | 157.860367 | -1.761658126 | 0.52311897 | -3.36760514 | 0.00075824 | 0.03347634 | 224.815571 | 89.6272455 | 262.725716 | 54.2729349 | Sig. genes |
| PF3D7_0725800 | N/A | 157.381643 | -1.770385407 | 0.52190033 | -3.39219061 | 0.00069336 | 0.03113077 | 224.815571 | 88.5207363 | 261.91733 | 54.2729349 | Sig. genes |
| PF3D7_1149900 | N/A | 28.0545748 | -1.820077262 | 0.80421472 | -2.26317327 | 0.02362501 | 0.34047378 | 32.4351437 | 13.9724282 | 54.9728608 | 10.8378665 | Non-sig. |
| PF3D7_1477200 | N/A | 3.70794525 | -1.837475032 | 2.00309158 | -0.91731953 | 0.35897522 | 0.89560738 | 4.18650345 | 0 | 7.38984124 | 3.25543631 | Non-sig. |
| PF3D7_0831400 | N/A | 1170.36336 | -1.844193704 | 0.42957284 | -4.29308735 | 1.7621E-05 | 0.00169734 | 1595.49945 | 548.422474 | 2066.0765 | 471.45501 | Sig. genes |
| PF3D7_1418800 | N/A | 151.529805 | -1.889788335 | 0.52074212 | -3.62902918 | 0.00028449 | 0.01463323 | 232.220476 | 79.2557002 | 245.106676 | 49.5363668 | Sig. genes |
| PF3D7_API02000 | RPL14 | 9.11780011 | -1.916807842 | 1.35334868 | -1.41634441 | 0.15667469 | 0.75894163 | 5.13196018 | 2.16555729 | 23.6590312 | 5.51465173 | Non-sig. |
| PF3D7_0825100 | N/A | 3.91526818 | -1.922062175 | 1.82018124 | -1.05597296 | 0.29098057 | 0.86724628 | 8.30380675 | 1.07715295 | 4.09707321 | 2.18303979 | Non-sig. |
| PF3D7_API01500 | RPL2 | 81.8360911 | -1.924378679 | 0.70066851 | -2.74648944 | 0.00602368 | 0.14347438 | 110.593552 | 13.9833412 | 148.442644 | 54.3248273 | Non-sig. |
| PF3D7_1109700 | N/A | 22.0635677 | -2.151712304 | 0.92015273 | -2.33842952 | 0.01936498 | 0.30809506 | 28.2963966 | 4.37417561 | 43.7318906 | 11.8518078 | Non-sig. |
| PF3D7_1424500 | N/A | 4.45834247 | -2.166194195 | 1.7268589 | -1.25441297 | 0.20969194 | 0.82566351 | 7.31233625 | 1.11028924 | 7.27850421 | 2.13224018 | Non-sig. |
| PF3D7_API03800 | ORF79 | 10.4418434 | -2.16742436 | 1.27822911 | -1.69564622 | 0.0899529 | 0.62981332 | 11.4155798 | 1.09005285 | 22.7272968 | 6.5344442 | Non-sig. |
| PF3D7_1300500 | RIF | 19.4624085 | -2.191586266 | 0.94783446 | -2.31220361 | 0.02076647 | 0.31957865 | 21.9874439 | 8.59673951 | 41.8498439 | 5.41560658 | Non-sig. |
| PF3D7_API01800 | RPL16 | 35.8613001 | -2.231975193 | 0.9553959 | -2.33617832 | 0.01948195 | 0.30902805 | 22.6105538 | 4.3247682 | 95.5791132 | 20.9307651 | Non-sig. |
| PF3D7_0901100 | RIF | 32.1405176 | -2.310694266 | 0.76114422 | -3.03581662 | 0.00239885 | 0.07610251 | 54.4810591 | 11.8183106 | 52.5196284 | 9.7430723 | Non-sig. |
| PF3D7_MIT01100 | RNA1 | 10.0194019 | -2.381260402 | 1.42245035 | -1.6740552 | 0.09411975 | 0.64424605 | 10.4515042 | 0 | 23.1512891 | 6.47481424 | Non-sig. |
| PF3D7_API03600 | ClpM | 59.5902738 | -2.413416829 | 1.00341709 | -2.40519804 | 0.0161637 | 0.27349786 | 27.3507057 | 3.21952609 | 173.273819 | 34.5170446 | Non-sig. |
| PF3D7_MIT03600 | SSUD | 3.23408777 | -2.433732326 | 2.1466198 | -1.13375099 | 0.25689901 | 0.85030738 | 6.8265509 | 0 | 4.12075074 | 1.98904946 | Non-sig. |
| PF3D7_1424900 | N/A | 3.4407426 | -2.442554699 | 1.98035171 | -1.2333944 | 0.21742866 | 0.83527883 | 4.24790249 | 1.1149989 | 7.36856588 | 1.03150313 | Non-sig. |
| PF3D7_1300600 | RIF | 2.2899462 | -2.912476187 | 2.58270867 | -1.12768282 | 0.2594539 | 0.85030738 | 3.14928809 | 1.07665972 | 4.93383698 | 0 | Non-sig. |
| PF3D7_0302400 | N/A | 9.50273413 | -2.955207547 | 1.50002044 | -1.97011152 | 0.04882559 | 0.48633045 | 30.4085293 | 1.07414561 | 3.28295212 | 3.2453095 | Non-sig. |
| PF3D7_0322800 | N/A | 7.54088185 | -3.040777869 | 1.57847387 | -1.92640368 | 0.05405399 | 0.50202873 | 22.8200403 | 2.15473165 | 4.09658258 | 1.09217283 | Non-sig. |
| PF3D7_API04900 | N/A | 47.815831 | -3.14672135 | 0.98778203 | -3.18564344 | 0.00144432 | 0.05388756 | 90.5273993 | 0 | 81.3233669 | 19.412558 | Non-sig. |
| PF3D7_1300700 | RIF | 13.1587232 | -3.491210586 | 1.23868426 | -2.81848305 | 0.00482512 | 0.1253111 | 14.674156 | 2.14853019 | 33.647897 | 2.16430951 | Non-sig. |
| PF3D7_1300900 | N/A | 36.4857526 | -3.503478456 | 0.93590327 | -3.74341942 | 0.00018153 | 0.01054778 | 28.2499639 | 8.59841733 | 105.843269 | 3.25135996 | Sig. genes |
| PF3D7_0700300 | RIF | 4.04366943 | -3.809462896 | 2.20808353 | -1.72523496 | 0.08448514 | 0.61653204 | 5.23669938 | 0 | 9.85362846 | 1.08434988 | Non-sig. |
| PF3D7_1351500 | N/A | 2.02165748 | -4.358076192 | 2.84005646 | -1.53450337 | 0.12490587 | 0.71232646 | 3.17869516 | 0 | 4.90793475 | 0 | Non-sig. |
| PF3D7_0833400 | RIF | 2.2878374 | -4.53454414 | 2.71589235 | -1.66963324 | 0.09499195 | 0.64740909 | 4.20730629 | 0 | 4.94404333 | 0 | Non-sig. |
| PF3D7_1418600 | N/A | 89.7126537 | -4.847124036 | 2.85890321 | -1.69544881 | 0.08999031 | 0.62981332 | 332.753268 | 0 | 14.0449128 | 12.052434 | Non-sig. |
| PF3D7_API05700 | N/A | 3.42843422 | -5.108392261 | 2.81416339 | -1.81524367 | 0.06948647 | 0.5752177 | 13.7137369 | 0 | 0 | 0 | Non-sig. |
| PF3D7_1418700 | N/A | 3.49181821 | -5.152062388 | 2.77600762 | -1.85592516 | 0.06346423 | 0.54850484 | 0 | 0 | 13.9672728 | 0 | Non-sig. |
| PF3D7_0832300 | N/A | 3.52578783 | -5.162540711 | 2.38910273 | -2.16087012 | 0.03070537 | 0.39012163 | 4.20157388 | 0 | 9.90157745 | 0 | Non-sig. |

| | | | | | | | | | | | | |
|---|---|---|---|---|---|---|---|---|---|---|---|---|
| PF3D7_0831700 | HSP70x | 2945.8736 | -5.349920842 | 0.63334428 | -8.44709749 | 2.9863E-17 | 4.2116E-14 | 4003.77977 | 55.7842585 | 7497.68102 | 226.249347 | Sig. genes |
| PF3D7_0601300 | EPF1 | 4.55934819 | -5.520034879 | 2.29864455 | -2.40143039 | 0.01633112 | 0.27399615 | 15.7631341 | 0 | 2.47425866 | 0 | Non-sig. |
| PF3D7_0601100 | EPF3 | 4.51882357 | -5.523408038 | 2.60376338 | -2.1213172 | 0.03389512 | 0.41377036 | 0 | 0 | 18.0752943 | 0 | Non-sig. |
| PF3D7_1253800 | N/A | 4.51882357 | -5.523408038 | 2.60376338 | -2.1213172 | 0.03389512 | 0.41377036 | 0 | 0 | 18.0752943 | 0 | Non-sig. |
| PF3D7_0833500 | VAR | 174.274271 | -6.323076102 | 0.7492005 | -8.43976488 | 3.1798E-17 | 4.2116E-14 | 290.892224 | 1.07259982 | 397.5945 | 7.5377596 | Sig. genes |
| PF3D7_0832200 | N/A | 568.705507 | -7.184742289 | 0.70639762 | -10.1709605 | 2.6726E-24 | 1.4159E-20 | 553.742048 | 5.68699238 | 1705.25665 | 10.1363337 | Sig. genes |
| PF3D7_0832000 | N/A | 16.4691074 | -7.379516641 | 1.72688827 | -4.2733029 | 1.926E-05 | 0.00170065 | 37.8442988 | 0 | 28.0321308 | 0 | Sig. genes |
| PF3D7_0832900 | N/A | 33.2932526 | -8.400279964 | 1.66772937 | -5.0369563 | 4.7299E-07 | 8.0836E-05 | 32.5886336 | 0 | 100.584377 | 0 | Sig. genes |

| | | | | | |
|---|---|---|---|---|---|
| PF3D7_1466000 | N/A | 16.7434285 | 15.04652181 | 13.9487494 | 8.66904273 |
| PF3D7_1466100 | PPKL | 169.557392 | 481.7010486 | 421.361146 | 589.424388 |
| PF3D7_1466200 | HDP1 | 945.770672 | 1823.2182 | 2302.75503 | 2732.01984 |
| PF3D7_1466300 | RPN2 | 6404.63519 | 12474.06876 | 14472.2418 | 14352.2815 |
| PF3D7_1466400 | ApiAP2 | 1265.52599 | 2534.709481 | 1980.50381 | 3059.85068 |
| PF3D7_1466500 | N/A | 172.75689 | 48.35678509 | 75.493719 | 42.242078 |
| PF3D7_1466600 | N/A | 85.7058118 | 26.87790373 | 20.5087468 | 18.4415746 |
| PF3D7_1466700 | NIP7 | 154.121564 | 138.5693769 | 153.474803 | 87.632644 |
| PF3D7_1466800 | N/A | 1035.38354 | 935.4740037 | 1179.50961 | 958.914722 |
| PF3D7_1466900 | N/A | 2910.42802 | 1529.616462 | 1702.85891 | 2345.7531 |
| PF3D7_1467000 | N/A | 33.6774515 | 33.2642559 | 33.6719525 | 50.6766682 |
| PF3D7_1467100 | MAG | 355.994605 | 473.1074049 | 665.316656 | 380.350031 |
| PF3D7_1467200 | WDR79 | 154.548629 | 107.3347212 | 137.127202 | 117.633576 |
| PF3D7_1467300 | DXR | 1166.68373 | 1372.567826 | 2005.39891 | 1121.74298 |
| PF3D7_1467400 | RPL22 | 927.470856 | 998.1587305 | 1292.48817 | 852.426406 |
| PF3D7_1467500 | N/A | 2320.65997 | 1508.873998 | 1213.74323 | 1343.97751 |
| PF3D7_1467600 | N/A | 1353.52318 | 907.3737541 | 180.730395 | 950.682856 |
| PF3D7_1467700 | N/A | 128.81712 | 148.282693 | 237.962682 | 107.212268 |
| PF3D7_1467800 | N/A | 163.198755 | 159.160141 | 304.315194 | 143.381081 |
| PF3D7_1467900 | N/A | 16766.551 | 11163.77953 | 8839.82724 | 15182.8166 |
| PF3D7_1468000 | N/A | 359.34821 | 245.6352234 | 134.718759 | 294.010034 |
| PF3D7_1468100 | MORC | 23705.6659 | 20310.59146 | 24347.7966 | 27013.1617 |

| | | | | | |
|---|---|---|---|---|---|
| PF3D7_1468200 | TLS1 | 456.740517 | 279.1727271 | 434.273248 | 302.531571 |
| PF3D7_1468300 | LRR14.1 | 26.0895638 | 57.00516936 | 50.8496215 | 48.8822625 |
| PF3D7_1468400 | D13 | 5820.52424 | 5048.242665 | 2990.38558 | 5127.79357 |
| PF3D7_1468500 | DER1-1 | 1300.64215 | 1701.983032 | 1562.99588 | 1396.23702 |
| PF3D7_1468600 | N/A | 120.136698 | 106.1909765 | 185.649746 | 175.540941 |
| PF3D7_1468700 | eIF4A | 6306.32564 | 10277.15251 | 19533.0001 | 9056.29381 |
| PF3D7_1468800 | U2AF2 | 1784.98614 | 1938.637632 | 1117.86861 | 1850.2654 |
| PF3D7_1468900 | N/A | 354.810588 | 375.5210135 | 590.533223 | 483.947248 |
| PF3D7_1469000 | IF1 | 273.077141 | 333.9050984 | 527.887723 | 310.010673 |
| PF3D7_1469100 | N/A | 506.095075 | 265.2132696 | 451.513706 | 240.942407 |
| PF3D7_1469200 | SHLP1 | 2344.16167 | 3797.662471 | 2654.61427 | 3377.29201 |
| PF3D7_1469300 | PNO1 | 403.493766 | 437.9942929 | 481.950824 | 331.481928 |
| PF3D7_1469400 | NT3 | 103.157021 | 192.0638661 | 401.613121 | 228.544868 |
| PF3D7_1469500 | N/A | 38.9348492 | 90.13846616 | 42.7049887 | 73.3273507 |
| PF3D7_1469600 | ACC | 592.55166 | 881.5816232 | 1274.19481 | 1306.81478 |
| PF3D7_1469700 | MED6 | 533.760501 | 390.7653981 | 650.258124 | 335.816745 |
| PF3D7_1469800 | N/A | 2172.09192 | 1124.545257 | 1194.44613 | 797.267824 |
| PF3D7_1469900 | MGET | 985.74436 | 651.7953067 | 135.449367 | 357.695693 |
| PF3D7_1470000 | ATG12 | 52.7992104 | 90.5512564 | 221.948818 | 85.3372434 |
| PF3D7_1470100 | N/A | 674.643567 | 832.2989294 | 2192.58279 | 952.848422 |
| PF3D7_1470200 | N/A | 279.305427 | 163.2068299 | 294.716823 | 112.364826 |
| PF3D7_1470300 | N/A | 203.772413 | 119.1726922 | 162.55538 | 142.574003 |
| PF3D7_1470400 | MPC2 | 491.147651 | 419.8909124 | 357.895445 | 269.155791 |
| PF3D7_1470500 | PDEdelta | 373.922148 | 506.355815 | 496.939258 | 451.43939 |
| PF3D7_1470600 | N/A | 917.080732 | 560.2352964 | 438.503955 | 514.629968 |
| PF3D7_1470700 | N/A | 672.575428 | 522.553502 | 323.58588 | 394.589755 |
| PF3D7_1470800 | N/A | 2103.01277 | 1848.519898 | 1108.83309 | 2085.1911 |
| PF3D7_1470900 | N/A | 4437.09733 | 3952.228665 | 5506.2944 | 3098.81344 |
| PF3D7_1470950 | N/A | 52.5640139 | 23.91713182 | 21.9630486 | 19.7487169 |
| PF3D7_1471000 | RCL1 | 178.785029 | 218.9437847 | 240.601498 | 229.807509 |
| PF3D7_1471100 | EXP2 | 7270.45903 | 4882.071422 | 10547.0654 | 5672.6601 |
| PF3D7_1471200 | SulP | 2411.63694 | 4009.2063 | 8116.69929 | 4588.37917 |
| PF3D7_1471300 | N/A | 969.8709 | 873.5716361 | 702.946258 | 771.259551 |
| PF3D7_1471400 | N/A | 4116.32538 | 3408.732383 | 4471.2368 | 3951.61181 |
| PF3D7_1471500 | TMEM43 | 25.1869455 | 70.87681652 | 37.7595812 | 65.9380303 |
| PF3D7_1471600 | APR2 | 109.537077 | 94.406853 | 100.208232 | 181.016039 |
| PF3D7_1471700 | N/A | 115.592703 | 89.09992859 | 66.5051518 | 103.648726 |
| PF3D7_1471800 | N/A | 143.672114 | 83.78041001 | 75.5095737 | 62.7355932 |
| PF3D7_1471900 | N/A | 794.686187 | 549.1706506 | 241.514139 | 506.105376 |
| PF3D7_1472000 | ISY1 | 194.03964 | 200.8494969 | 179.750502 | 162.224852 |
| PF3D7_1472100 | YIP1 | 1133.86955 | 1528.926085 | 1324.78818 | 1451.89933 |
| PF3D7_1472200 | HDA1 | 8052.13244 | 7391.839076 | 5877.09583 | 9475.01037 |
| PF3D7_1472300 | N/A | 716.75826 | 1321.426772 | 1564.95571 | 1503.82943 |
| PF3D7_1472400 | N/A | 1127.02327 | 885.0520288 | 1362.67933 | 1088.20354 |
| PF3D7_1472500 | N/A | 206.203626 | 125.7373323 | 118.979456 | 112.673273 |
| PF3D7_1472600 | PDI-14 | 2193.42076 | 3114.079099 | 2396.70848 | 2896.39738 |
| PF3D7_1472700 | N/A | 1251.33481 | 760.7845269 | 1275.42911 | 1063.12649 |
| PF3D7_1472800 | N/A | 2023.74172 | 1188.386802 | 1182.29519 | 990.140676 |
| PF3D7_1472900 | N/A | 143.076626 | 309.0675826 | 417.176907 | 299.845476 |
| PF3D7_1473000 | N/A | 1.03009585 | 0 | 0.81895513 | 1.10543768 |
| PF3D7_1473100 | N/A | 515.753336 | 767.1957773 | 674.275712 | 882.96485 |
| PF3D7_1473200 | N/A | 4197.01504 | 3713.016178 | 5515.52506 | 3263.78083 |
| PF3D7_1473300 | N/A | 214.719459 | 197.4328049 | 115.801089 | 217.778553 |
| PF3D7_1473400 | N/A | 3037.60762 | 4384.530169 | 2405.00588 | 4683.7071 |

| | | | | | |
|---|---|---|---|---|---|
| PF3D7_1473500 | N/A | 224.258564 | 297.1975271 | 442.695078 | 406.340926 |
| PF3D7_1473600 | N/A | 122.983395 | 78.358768 | 201.167551 | 60.4464539 |
| PF3D7_1473700 | NUP116 | 515.36986 | 646.7830931 | 100.219584 | 632.111703 |
| PF3D7_1473900 | N/A | 425.479867 | 357.2510971 | 401.64944 | 544.159335 |
| PF3D7_1474000 | N/A | 690.521062 | 784.0132082 | 252.204582 | 920.470522 |
| PF3D7_1474100 | N/A | 536.821461 | 361.7973084 | 1162.54801 | 231.111593 |
| PF3D7_1474200 | N/A | 120.20831 | 139.41929 | 182.382816 | 196.974947 |
| PF3D7_1474300 | N/A | 460.031759 | 315.4499738 | 298.134023 | 267.335863 |
| PF3D7_1474400 | N/A | 1045.75328 | 903.0736765 | 590.663787 | 887.129661 |
| PF3D7_1474500 | SF3A1 | 1275.36825 | 899.0318942 | 1762.65349 | 1367.41978 |
| PF3D7_1474600 | VMP1 | 784.905659 | 1654.747003 | 1620.28787 | 1892.71909 |
| PF3D7_1474700 | N/A | 118.86364 | 124.4903256 | 102.649348 | 157.491993 |
| PF3D7_1474800 | N/A | 2154.62797 | 1959.022664 | 2830.25603 | 2337.10614 |
| PF3D7_1474900 | CITH | 269.659126 | 639.3722446 | 699.828169 | 628.107607 |
| PF3D7_1475000 | MED31 | 12.3389951 | 5.403925004 | 35.1392338 | 17.6180685 |
| PF3D7_1475100 | N/A | 772.341515 | 807.7292751 | 1334.0145 | 827.199842 |
| PF3D7_1475200 | N/A | 730.494388 | 562.2484002 | 774.476157 | 627.440099 |
| PF3D7_1475300 | COX11 | 668.866344 | 819.9659389 | 757.935777 | 651.66073 |
| PF3D7_1475400 | CRMP4 | 304.763652 | 349.612586 | 430.495316 | 596.262967 |
| PF3D7_1475500 | CCp1 | 569.007837 | 893.5327503 | 673.583905 | 666.688307 |
| PF3D7_1475600 | BDP4 | 1191.02347 | 1618.936512 | 1310.88389 | 1817.96472 |
| PF3D7_1475700 | N/A | 21.0071367 | 81.59607554 | 47.6171016 | 73.4482418 |
| PF3D7_1475800 | N/A | 259.489874 | 582.5370059 | 211.687851 | 368.476438 |
| PF3D7_1475900 | N/A | 56.8130313 | 20.38955281 | 16.4244493 | 16.1778996 |
| PF3D7_1476000 | N/A | 47.0209203 | 46.23338412 | 22.9689208 | 48.8274581 |
| PF3D7_1476100 | ETRAMP14.2 | 24.8805972 | 21.55113633 | 9.83053089 | 9.83460782 |
| PF3D7_1476200 | N/A | 686.335495 | 242.4915067 | 329.341325 | 270.579512 |
| PF3D7_1476300 | N/A | 10619.6811 | 6071.762057 | 9325.41492 | 6664.17228 |
| PF3D7_1476500 | N/A | 483.385151 | 528.9091136 | 282.549002 | 384.696407 |
| PF3D7_1476600 | N/A | 307.690363 | 449.4461357 | 210.290162 | 459.890262 |
| PF3D7_1476700 | LPL1 | 34.653145 | 33.28493692 | 31.1960444 | 48.6162907 |
| PF3D7_1476800 | N/A | 386.311675 | 518.6472847 | 500.75044 | 510.076718 |
| PF3D7_1477100 | N/A | 9.42539258 | 5.372601305 | 6.56445685 | 1.08297802 |
| PF3D7_1477200 | N/A | 4.18650345 | 0 | 7.38984124 | 3.25543631 |
| PF3D7_1477300 | Pfg14-744 | 271.51571 | 334.7916725 | 231.596028 | 255.547148 |
| PF3D7_1477400 | N/A | 133.412781 | 177.1428628 | 183.904764 | 106.919472 |
| PF3D7_1477500 | N/A | 470.742422 | 270.3560076 | 422.179962 | 298.492879 |
| PF3D7_1477600 | SURF14.1 | 220.278884 | 139.4376558 | 92.8269473 | 134.601867 |
| PF3D7_1477700 | Pfg14-748 | 742.514931 | 601.0150493 | 283.304192 | 479.017361 |
| PF3D7_1477800 | ACBP | 1102.16594 | 1650.91187 | 127.216821 | 1381.2865 |
| PF3D7_1478000 | GEXP17 | 106.082686 | 79.44951587 | 72.246235 | 90.7327493 |
| PF3D7_1478100 | N/A | 125.953486 | 192.2184837 | 146.116999 | 169.686717 |
| PF3D7_1478600 | PTP3 | 1751.37763 | 1595.03274 | 509.300109 | 1501.32004 |
| PF3D7_1478800 | N/A | 120.783731 | 551.8565146 | 126.430269 | 516.325888 |
| PF3D7_1478900 | N/A | 17747.3704 | 34018.51232 | 13527.3241 | 38819.8952 |
| PF3D7_1479000 | ACS1a | 67.3819947 | 973.1376373 | 58.3140639 | 838.559272 |
| PF3D7_1479100 | N/A | 46.8294351 | 179.7578634 | 9.01759753 | 121.966912 |
| PF3D7_1479200 | N/A | 107.672908 | 742.8703534 | 18.0481843 | 625.836345 |
| PF3D7_1479400 | RIF | 1.05000286 | 6.458737404 | 0.82227348 | 0 |
| PF3D7_1479500 | N/A | 1.0459648 | 6.449391467 | 11.4862906 | 6.50451511 |
| PF3D7_1479600 | RIF | 1.05029178 | 2.154073311 | 0 | 0 |
| PF3D7_1479700 | RIF | 1.04724102 | 1.073973605 | 0 | 5.41086947 |
| PF3D7_1479800 | RIF | 1.04756107 | 4.297741739 | 0.82059814 | 6.49622489 |
| PF3D7_1479900 | N/A | 6.27693604 | 4.299378091 | 3.28187991 | 4.33565978 |

| | | | | | |
|---|---|---|---|---|---|
| PF3D7_1480000 | RIF | 4.18982556 | 3.22339653 | 1.64116719 | 1.08280123 |
| PF3D7_API00100 | RPS4 | 0 | 0 | 0 | 0 |
| PF3D7_API00200 | N/A | 0 | 0 | 0 | 0 |
| PF3D7_API00300 | N/A | 0 | 0 | 0 | 0 |
| PF3D7_API00450 | N/A | 0 | 0 | 0 | 0 |
| PF3D7_API00600 | N/A | 0 | 0 | 0 | 0 |
| PF3D7_API00700 | N/A | 0 | 0 | 0 | 0 |
| PF3D7_API00800 | N/A | 0 | 0 | 0 | 0 |
| PF3D7_API00900 | N/A | 0 | 0 | 0 | 0 |
| PF3D7_API01000 | N/A | 0 | 0 | 0 | 0 |
| PF3D7_API01100 | N/A | 0 | 0 | 0 | 0 |
| PF3D7_API01200 | N/A | 0 | 0 | 0 | 0 |
| PF3D7_API01300 | RPL4 | 8.30789047 | 6.462000225 | 51.6277746 | 19.638719 |
| PF3D7_API01400 | RPL23 | 0 | 0 | 0 | 0 |
| PF3D7_API01500 | RPL2 | 110.593552 | 13.98334121 | 148.442644 | 54.3248273 |
| PF3D7_API01600 | RPS19 | 4.14337 | 0 | 0.81719873 | 3.30500297 |
| PF3D7_API01700 | RPS3 | 2.08461219 | 0 | 1.64166407 | 1.0899898 |
| PF3D7_API01800 | RPL16 | 22.6105538 | 4.324768201 | 95.5791132 | 20.9307651 |
| PF3D7_API01900 | RPS17 | 0 | 0 | 0.82160429 | 0 |
| PF3D7_API02000 | RPL14 | 5.13196018 | 2.165557285 | 23.6590312 | 5.51465173 |
| PF3D7_API02100 | RPS8 | 0 | 0 | 0 | 0 |
| PF3D7_API02200 | RPL6 | 0 | 1.071257919 | 13.02188 | 3.2618279 |
| PF3D7_API02300 | RPS5 | 30.2454677 | 6.454527142 | 71.3464857 | 23.9107864 |
| PF3D7_API02400 | ORF91 | 4.12266776 | 1.087142355 | 0.81344624 | 2.19430188 |
| PF3D7_API02500 | RPL36 | 0 | 0 | 0 | 0 |
| PF3D7_API02600 | RPS11 | 5.14124796 | 2.161570002 | 11.4406808 | 4.40451714 |
| PF3D7_API02700 | RPS12 | 573.956012 | 85.48664133 | 640.750847 | 294.410819 |
| PF3D7_API02800 | RPS7 | 14.4562664 | 0 | 9.01801177 | 7.71123306 |
| PF3D7_API02900 | TUFA | 328.31926 | 180.4384496 | 2208.74498 | 837.022734 |
| PF3D7_API03000 | ORF78 | 0 | 0 | 2.43557752 | 3.26663117 |
| PF3D7_API03100 | N/A | 0 | 0 | 0 | 0 |
| PF3D7_API03200 | N/A | 0 | 0 | 0 | 0 |
| PF3D7_API03300 | N/A | 0 | 0 | 0 | 0 |
| PF3D7_API03400 | N/A | 0 | 0 | 0 | 0 |
| PF3D7_API03500 | RPL11 | 1.04422304 | 0 | 1.6600145 | 0 |
| PF3D7_API03600 | ClpM | 27.3507057 | 3.219526095 | 173.273819 | 34.5170446 |
| PF3D7_API03700 | N/A | 0 | 0 | 0 | 0 |
| PF3D7_API03800 | ORF79 | 11.4155798 | 1.090052854 | 22.7272968 | 6.5344442 |
| PF3D7_API03900 | N/A | 0 | 0 | 0 | 0 |
| PF3D7_API04000 | YCF93 | 0 | 0 | 0 | 0 |
| PF3D7_API04100 | RPS2 | 0 | 0 | 0 | 0 |
| PF3D7_API04200 | rpoC2 | 0 | 0 | 0 | 0 |
| PF3D7_API04300 | rpoC1 | 1.05490284 | 0 | 0 | 0 |
| PF3D7_API04400 | rpoB | 0 | 0 | 0 | 0 |
| PF3D7_API04500 | ORF101 | 0 | 0 | 0 | 0 |
| PF3D7_API04600 | ORF51 | 0 | 0 | 0 | 0 |
| PF3D7_API04700 | SUFB | 0 | 0 | 0 | 0 |
| PF3D7_API04800 | N/A | 0 | 0 | 0 | 0 |
| PF3D7_API04900 | N/A | 90.5273993 | 0 | 81.3233669 | 19.412558 |
| PF3D7_API05000 | N/A | 0 | 0 | 0 | 0 |
| PF3D7_API05100 | N/A | 0 | 0 | 0 | 0 |
| PF3D7_API05200 | N/A | 0 | 0 | 0 | 0 |
| PF3D7_API05300 | N/A | 0 | 0 | 0 | 0 |
| PF3D7_API05400 | N/A | 0 | 0 | 0 | 0 |

| | | | | | |
|---|---|---|---|---|---|
| PF3D7_API05500 | N/A | 0 | 0 | 0 | 0 |
| PF3D7_API05600 | N/A | 0 | 0 | 0 | 0 |
| PF3D7_API05700 | N/A | 13.7137369 | 0 | 0 | 0 |
| PF3D7_API05800 | N/A | 0 | 0 | 0 | 0 |
| PF3D7_API05900 | N/A | 0 | 5.360305915 | 11.4744569 | 4.31542782 |
| PF3D7_API06000 | N/A | 0 | 0 | 0 | 0 |
| PF3D7_API06100 | N/A | 0 | 0 | 0 | 0 |
| PF3D7_API06200 | N/A | 0 | 0 | 0 | 0 |
| PF3D7_API06300 | N/A | 0 | 0 | 0 | 0 |
| PF3D7_API06400 | N/A | 0 | 0 | 0 | 0 |
| PF3D7_API06500 | N/A | 0 | 0 | 0 | 0 |
| PF3D7_API06600 | N/A | 0 | 0 | 0 | 0 |
| PF3D7_API06700 | N/A | 0 | 10.72328849 | 0 | 0 |
| PF3D7_API06800 | N/A | 0 | 0 | 0 | 0 |
| PF3D7_MIT00100 | N/A | 0 | 0 | 0 | 0 |
| PF3D7_MIT00200 | RNA20 | 0 | 0 | 0 | 0 |
| PF3D7_MIT00300 | RNA9 | 0 | 0 | 0 | 0 |
| PF3D7_MIT00400 | RNA17 | 0 | 0 | 0 | 0 |
| PF3D7_MIT00500 | RNA23t | 0 | 0 | 0 | 0 |
| PF3D7_MIT00600 | LSUC | 0 | 0 | 0 | 0 |
| PF3D7_MIT00700 | RNA24t | 0 | 0 | 0 | 0 |
| PF3D7_MIT00800 | RNA25t | 0 | 0 | 0 | 0 |
| PF3D7_MIT00900 | LSUG | 240.939447 | 85.94251657 | 304.435201 | 119.586476 |
| PF3D7_MIT01000 | SSUB | 297.959871 | 181.6984688 | 612.718138 | 164.917633 |
| PF3D7_MIT01100 | RNA1 | 10.4515042 | 0 | 23.1512891 | 6.47481424 |
| PF3D7_MIT01200 | RNA15 | 0 | 0 | 0 | 0 |
| PF3D7_MIT01300 | RNA10 | 194.59155 | 93.53179242 | 306.865953 | 93.1495216 |
| PF3D7_MIT01400 | COX3 | 412.215108 | 289.3281237 | 1326.19093 | 429.077638 |
| PF3D7_MIT01500 | LSUF | 0 | 2.158559334 | 1.6900777 | 1.03938601 |
| PF3D7_MIT01600 | SSUE | 0 | 0 | 0 | 0 |
| PF3D7_MIT01700 | RNA2 | 15.9958376 | 8.638813229 | 15.3404033 | 7.02703012 |
| PF3D7_MIT01800 | RNA26t | 0 | 0 | 0 | 0 |
| PF3D7_MIT01900 | RNA21 | 0 | 0 | 0 | 0 |
| PF3D7_MIT02000 | RNA3 | 23.0804757 | 13.74362341 | 57.6484482 | 13.512825 |
| PF3D7_MIT02100 | COX1 | 4065.49976 | 3886.66575 | 9963.28196 | 5748.95124 |
| PF3D7_MIT02200 | SSUA | 45.1118389 | 25.19664292 | 49.6417192 | 29.1045461 |
| PF3D7_MIT02300 | CYTB | 2044.19131 | 1956.158676 | 6332.52965 | 2539.49298 |
| PF3D7_MIT02400 | RNA27t | 0 | 0 | 0.82160429 | 0 |
| PF3D7_MIT02500 | LSUB | 0 | 0 | 0 | 0 |
| PF3D7_MIT02600 | RNA4 | 2.09428497 | 1.099425162 | 0.80729394 | 0 |
| PF3D7_MIT02700 | RNA5 | 20.7749782 | 7.951390137 | 45.194028 | 11.551651 |
| PF3D7_MIT02800 | RNA6 | 0 | 0 | 0 | 0 |
| PF3D7_MIT02900 | RNA12 | 1.05490284 | 0 | 0 | 0 |
| PF3D7_MIT03000 | RNA18 | 0 | 0 | 0 | 0 |
| PF3D7_MIT03100 | RNA13 | 0 | 0 | 0 | 0 |
| PF3D7_MIT03200 | LSUA | 9895.67961 | 4537.517606 | 11136.3001 | 3729.05414 |
| PF3D7_MIT03300 | RNA7 | 358.841023 | 155.260763 | 378.304678 | 131.34664 |
| PF3D7_MIT03400 | RNA11 | 0 | 0 | 0 | 0 |
| PF3D7_MIT03500 | RNA22 | 0 | 0 | 0 | 0 |
| PF3D7_MIT03600 | SSUD | 6.8265509 | 0 | 4.12075074 | 1.98904946 |
| PF3D7_MIT03700 | SSUF | 0 | 0 | 1.66356418 | 1.06279618 |
| PF3D7_MIT03800 | RNA14 | 0 | 0 | 0 | 0 |
| PF3D7_MIT03900 | RNA19 | 0 | 0 | 0 | 0 |
| PF3D7_MIT04000 | LSUE | 20739.2284 | 14478.46039 | 20791.0862 | 8794.3086 |

| | | | | | |
|---|---|---|---|---|---|
| PF3D7_MIT04100 | LSUD | 3.12767994 | 4.554814957 | 3.2222479 | 1.04565699 |
| PF3D7_MIT04200 | RNA8 | 246.969576 | 111.1063407 | 233.976923 | 60.3297659 |

### Significantly deregulated genes shown

| gene_id | gene_name | baseMean | log2FoldChange | lfcSE | stat | pvalue | padj | 01_JP_S1 | 02_JP_S2 | 03_JP_S3 | 04_JP_S4 | sig |
|---|---|---|---|---|---|---|---|---|---|---|---|---|
| PF3D7_1255100 | RIF | 147.464972 | 5.921610151 | 0.72648659 | 8.15102472 | 3.6085E-16 | 3.1863E-13 | 8.3817014 | 339.507833 | 1.64123839 | 240.329117 | Sig. genes |
| PF3D7_0101600 | RIF | 37.1331716 | 5.661533899 | 1.1104928 | 5.09821755 | 3.4287E-07 | 6.2379E-05 | 3.14328293 | 88.0567073 | 0 | 57.3326961 | Sig. genes |
| PF3D7_0412400 | VAR | 324.642616 | 4.676104275 | 0.6733819 | 6.94420842 | 3.8059E-12 | 1.8331E-09 | 42.1632811 | 549.15281 | 7.39343837 | 699.860934 | Sig. genes |
| PF3D7_0631900 | N/A | 163.528712 | 4.120673442 | 0.6666414 | 6.18124445 | 6.3598E-10 | 1.982E-07 | 6.27671638 | 362.226339 | 28.7163033 | 256.895489 | Sig. genes |
| PF3D7_0413100 | VAR | 71.2906139 | 4.064505801 | 0.90283001 | 4.50196132 | 6.7329E-06 | 0.00075896 | 0 | 162.986639 | 15.6035626 | 106.572254 | Sig. genes |
| PF3D7_0425200 | N/A | 143.660193 | 4.02169277 | 0.58422322 | 6.88382907 | 5.8265E-12 | 2.3745E-09 | 23.0781103 | 285.6895 | 10.6703764 | 255.202784 | Sig. genes |
| PF3D7_0712900 | VAR | 197.609312 | 3.965676312 | 0.51356184 | 7.72190605 | 1.146E-14 | 8.6738E-12 | 26.3480798 | 347.525513 | 21.3582618 | 395.205392 | Sig. genes |
| PF3D7_0425100 | N/A | 64.4411399 | 3.856353739 | 0.78292742 | 4.92555711 | 8.412E-07 | 0.00012045 | 14.631373 | 138.6929 | 2.461023 | 101.979264 | Sig. genes |
| PF3D7_1479000 | ACS1a | 484.348242 | 3.851159185 | 0.45233968 | 8.51386556 | 1.6823E-17 | 4.2116E-14 | 67.3819947 | 973.137637 | 58.3140639 | 838.559272 | Sig. genes |
| PF3D7_1148200 | N/A | 747.804978 | 3.718372609 | 0.45093968 | 8.24583148 | 1.6402E-16 | 1.7379E-13 | 117.201143 | 1188.36933 | 94.2194419 | 1591.43 | Sig. genes |
| PF3D7_1370500 | N/A | 959.806226 | 3.636647033 | 0.5012759 | 7.25478128 | 4.0231E-13 | 2.6643E-10 | 194.490947 | 1633.14664 | 91.8068248 | 1919.78049 | Sig. genes |
| PF3D7_0935600 | GIG | 3296.12516 | 3.499472447 | 0.51301391 | 6.82139878 | 9.0158E-12 | 3.4118E-09 | 758.026194 | 6160.75015 | 313.744555 | 5951.97973 | Sig. genes |
| PF3D7_1479200 | N/A | 373.606948 | 3.452431013 | 0.68866087 | 5.0132528 | 5.3518E-07 | 8.4674E-05 | 107.672908 | 742.870353 | 18.0481843 | 625.836345 | Sig. genes |
| PF3D7_0202200 | PTP1 | 368.896774 | 3.445386779 | 0.48718199 | 7.07207333 | 1.5264E-12 | 8.9852E-10 | 76.8446618 | 610.519869 | 47.6354415 | 740.587123 | Sig. genes |
| PF3D7_0102200 | RESA | 27786.8823 | 3.382108993 | 0.75800095 | 4.46187962 | 8.1244E-06 | 0.00086224 | 8832.8789 | 49243.0973 | 895.384546 | 52176.1686 | Sig. genes |
| PF3D7_0712600 | VAR | 154.696898 | 3.350530969 | 0.53838204 | 6.22333352 | 4.867E-10 | 1.6116E-07 | 31.6180094 | 232.755635 | 23.8227178 | 330.59123 | Sig. genes |
| PF3D7_0702000 | N/A | 827.790246 | 3.325777239 | 0.72998409 | 4.5559585 | 5.2147E-06 | 0.00061395 | 267.246392 | 1683.72172 | 33.6704458 | 1326.52243 | Sig. genes |
| PF3D7_0829800 | N/A | 2864.00317 | 3.228717914 | 0.46900598 | 6.88417216 | 5.8125E-12 | 2.3745E-09 | 388.354932 | 4811.47247 | 715.32748 | 5540.85779 | Sig. genes |
| PF3D7_0712800 | VAR | 67.6344179 | 3.193437918 | 0.62550554 | 5.10537112 | 3.3015E-07 | 6.2379E-05 | 12.6473318 | 130.857634 | 13.9650661 | 113.06764 | Sig. genes |
| PF3D7_0831800 | HRP2 | 966.038595 | 3.192153975 | 0.4595747 | 6.94588704 | 3.7609E-12 | 1.8331E-09 | 143.219698 | 1782.00322 | 237.372417 | 1701.55904 | Sig. genes |
| PF3D7_0702300 | STARP | 312.402092 | 3.18043644 | 0.66581758 | 4.77673848 | 1.7816E-06 | 0.00024203 | 104.251081 | 538.576957 | 20.5337937 | 586.246537 | Sig. genes |
| PF3D7_0711700 | VAR | 120.406062 | 3.110292221 | 0.65708317 | 4.73348334 | 2.207E-06 | 0.00028519 | 38.9955449 | 173.762271 | 11.5006224 | 257.365811 | Sig. genes |
| PF3D7_0412900 | VAR | 131.211613 | 3.09499195 | 0.55789041 | 5.54767013 | 2.895E-08 | 6.9717E-06 | 34.7811876 | 263.858284 | 20.5369672 | 205.670012 | Sig. genes |
| PF3D7_0221700 | N/A | 269.10043 | 2.92073452 | 0.83567451 | 3.49506237 | 0.00047395 | 0.02262155 | 117.876903 | 582.652903 | 8.21274226 | 367.659174 | Sig. genes |
| PF3D7_1200700 | ACS7 | 2200.68667 | 2.909515877 | 0.59711539 | 4.8726191 | 1.1013E-06 | 0.00015354 | 818.615753 | 3753.4258 | 216.034047 | 4014.67107 | Sig. genes |
| PF3D7_0412700 | VAR | 128.012664 | 2.902326347 | 0.57917125 | 5.0111713 | 5.41E-07 | 8.4674E-05 | 41.1111922 | 254.194216 | 19.716002 | 197.029247 | Sig. genes |
| PF3D7_0935900 | REX1 | 1819.53833 | 2.862709247 | 0.7125015 | 4.01782909 | 5.8737E-05 | 0.00426284 | 774.333682 | 3031.52527 | 105.962691 | 3366.33168 | Sig. genes |
| PF3D7_0207300 | SERA8 | 85.6024445 | 2.830283474 | 0.57284061 | 4.94078706 | 7.7808E-07 | 0.00011778 | 19.9790859 | 148.114803 | 22.1709303 | 152.144958 | Sig. genes |
| PF3D7_1352900 | N/A | 225.334622 | 2.82814138 | 0.55534556 | 5.09257942 | 3.5322E-07 | 6.2379E-05 | 78.9441105 | 372.328361 | 32.8516007 | 417.214414 | Sig. genes |
| PF3D7_0315200 | CTRP | 472.749512 | 2.773315653 | 0.58559553 | 4.73588934 | 2.181E-06 | 0.00028519 | 184.43005 | 724.018682 | 57.5026593 | 925.046657 | Sig. genes |
| PF3D7_1401100 | N/A | 1616.07888 | 2.756782246 | 0.41446593 | 6.65140864 | 2.903E-11 | 1.0253E-08 | 446.471004 | 2873.15914 | 386.853879 | 2757.83149 | Sig. genes |
| PF3D7_0424800 | N/A | 67.3481893 | 2.729785986 | 0.72733117 | 3.75315415 | 0.00017462 | 0.01051312 | 28.3311069 | 138.537731 | 7.38755065 | 95.1363691 | Sig. genes |
| PF3D7_1407700 | N/A | 44.5078722 | 2.705923137 | 0.82046792 | 3.2980243 | 0.00097368 | 0.03968109 | 19.9286162 | 94.5171167 | 4.10390078 | 59.4818553 | Sig. genes |
| PF3D7_1221400 | IMC1h | 230.614839 | 2.69702061 | 0.47172626 | 5.71734254 | 1.082E-08 | 2.8663E-06 | 59.9672014 | 418.52654 | 63.2337406 | 380.731874 | Sig. genes |
| PF3D7_0801300 | WARP | 148.670449 | 2.678988781 | 0.68466481 | 3.9128472 | 9.1214E-05 | 0.00623873 | 66.9194756 | 269.823378 | 13.9468891 | 243.992053 | Sig. genes |

| | | | | | | | | | | | | |
|---|---|---|---|---|---|---|---|---|---|---|---|---|
| PF3D7_1115800 | N/A | 3664.66186 | 2.63131269 | 0.60042556 | 4.38241282 | 1.1737E-05 | 0.00119584 | 1620.52272 | 6523.12582 | 417.241846 | 6097.75704 | Sig. genes |
| PF3D7_0902500 | FIKK9.6 | 666.287426 | 2.614373756 | 0.45092069 | 5.79785718 | 6.7168E-09 | 1.8729E-06 | 224.226334 | 1166.30606 | 150.299009 | 1124.31831 | Sig. genes |
| PF3D7_0301800 | N/A | 1186.03502 | 2.594848708 | 0.5034771 | 5.15385649 | 2.5518E-07 | 5.4078E-05 | 460.836355 | 2241.75094 | 213.519426 | 1828.03335 | Sig. genes |
| PF3D7_1401400 | ETRAMP14 | 3110.5344 | 2.568849662 | 0.50235174 | 5.11364737 | 3.16E-07 | 6.2006E-05 | 1247.43619 | 5330.6744 | 547.728765 | 5316.29822 | Sig. genes |
| PF3D7_1002100 | PTP5 | 2209.43167 | 2.514083058 | 0.42930503 | 5.85616956 | 4.7366E-09 | 1.3942E-06 | 771.666522 | 3710.24614 | 545.351318 | 3810.46268 | Sig. genes |
| PF3D7_1148900 | N/A | 120.348396 | 2.483204816 | 0.58070709 | 4.27617442 | 1.9013E-05 | 0.00170065 | 50.4690058 | 195.344681 | 22.9917186 | 212.588178 | Sig. genes |
| PF3D7_1149600 | N/A | 2033.23576 | 2.481025169 | 0.54753283 | 4.53128112 | 5.8627E-06 | 0.00067523 | 918.254836 | 3414.9182 | 317.8633 | 3481.90669 | Sig. genes |
| PF3D7_0501300 | SBP1 | 1674.19272 | 2.476218915 | 0.53938723 | 4.59080007 | 4.4155E-06 | 0.00053167 | 747.329038 | 2989.36642 | 273.48903 | 2686.5864 | Sig. genes |
| PF3D7_1149200 | RESA3 | 21673.4427 | 2.452373106 | 0.73178906 | 3.35120221 | 0.00080462 | 0.03523018 | 11953.5 | 33672.3987 | 1439.92714 | 39627.9451 | Sig. genes |
| PF3D7_1479100 | N/A | 89.392952 | 2.447873942 | 0.74105882 | 3.3032114 | 0.00095584 | 0.03925625 | 46.8294351 | 179.757863 | 9.01759753 | 121.966912 | Sig. genes |
| PF3D7_1404800 | N/A | 2671.08521 | 2.442105647 | 0.46027026 | 5.30580807 | 1.1217E-07 | 2.5839E-05 | 1036.71186 | 3920.75927 | 624.281871 | 5102.58783 | Sig. genes |
| PF3D7_0108500 | N/A | 3058.68094 | 2.440882895 | 0.66413853 | 3.67526171 | 0.00023761 | 0.01258835 | 1607.79036 | 5764.69544 | 295.680094 | 4566.55788 | Sig. genes |
| PF3D7_0424600 | N/A | 5168.90345 | 2.436398106 | 0.43699565 | 5.57533714 | 2.4705E-08 | 6.2327E-06 | 1954.68867 | 8994.56266 | 1269.83748 | 8456.52499 | Sig. genes |
| PF3D7_0605900 | N/A | 942.656415 | 2.360429394 | 0.47914723 | 4.92631337 | 8.3796E-07 | 0.00012045 | 223.170009 | 1343.35036 | 390.940576 | 1813.16471 | Sig. genes |
| PF3D7_0324300 | EPF4 | 188.982343 | 2.352752651 | 0.60154697 | 3.91117033 | 9.185E-05 | 0.00623873 | 88.9380615 | 404.121613 | 35.2813086 | 227.588388 | Sig. genes |
| PF3D7_1029700 | N/A | 41.4947026 | 2.345737688 | 0.69582316 | 3.37116934 | 0.0007485 | 0.0333239 | 11.5718603 | 66.5362131 | 15.602952 | 72.2677848 | Sig. genes |
| PF3D7_0805200 | GAMER | 2003.94472 | 2.266209458 | 0.66792068 | 3.39293202 | 0.00069149 | 0.03113077 | 1170.8524 | 3524.60945 | 209.275543 | 3111.0415 | Sig. genes |
| PF3D7_0730800 | N/A | 3903.99137 | 2.202953809 | 0.58825352 | 3.74490544 | 0.00018046 | 0.01054778 | 2185.11733 | 6690.15551 | 602.002904 | 6138.68975 | Sig. genes |
| PF3D7_1102700 | ETRAMP11.1 | 5302.52957 | 2.194062438 | 0.57199968 | 3.83577565 | 0.00012517 | 0.00780169 | 2921.70795 | 9348.03272 | 882.816541 | 8057.56108 | Sig. genes |
| PF3D7_0402100 | N/A | 4354.06502 | 2.182185039 | 0.48957644 | 4.4572918 | 8.3002E-06 | 0.00086224 | 2138.36585 | 6896.56955 | 1006.86097 | 7374.4637 | Sig. genes |
| PF3D7_0731500 | EBA175 | 60363.9877 | 2.17062345 | 0.42282718 | 5.13359485 | 2.8426E-07 | 5.7923E-05 | 24181.2409 | 85027.8241 | 19702.7014 | 112544.185 | Sig. genes |
| PF3D7_0202500 | ETRAMP2 | 10880.5003 | 2.15942263 | 0.5254957 | 4.10930598 | 3.9685E-05 | 0.00300359 | 5702.91389 | 19971.3541 | 2258.03638 | 15589.6969 | Sig. genes |
| PF3D7_1201200 | N/A | 2396.51096 | 2.151157889 | 0.41065202 | 5.23839604 | 1.6198E-07 | 3.5757E-05 | 838.231737 | 4145.52536 | 923.201462 | 3679.08527 | Sig. genes |
| PF3D7_0402900 | N/A | 89.8922552 | 2.14527275 | 0.57247622 | 3.74735698 | 0.00017871 | 0.01054778 | 37.7243107 | 171.894204 | 28.7225697 | 121.227937 | Sig. genes |
| PF3D7_0302200 | CLAG3.2 | 2772.12371 | 2.141431293 | 0.51797564 | 4.13423165 | 3.5614E-05 | 0.00279136 | 617.377873 | 3783.47561 | 1430.91429 | 5256.72706 | Sig. genes |
| PF3D7_0219700 | GEXP20 | 194.73734 | 2.136944463 | 0.49910188 | 4.28157969 | 1.8557E-05 | 0.00170065 | 87.1400005 | 332.868571 | 57.464318 | 301.476471 | Sig. genes |
| PF3D7_0814500 | N/A | 525.611109 | 2.135293946 | 0.45910328 | 4.65100996 | 3.3031E-06 | 0.00040698 | 231.651564 | 769.258746 | 158.518796 | 943.01533 | Sig. genes |
| PF3D7_0102700 | MaTrA | 340.975532 | 2.130169047 | 0.50427782 | 4.22419735 | 2.3979E-05 | 0.00202067 | 167.065942 | 591.512663 | 87.0297809 | 518.293743 | Sig. genes |
| PF3D7_1478800 | N/A | 328.849101 | 2.111019499 | 0.4468021 | 4.72473046 | 2.3042E-06 | 0.00029066 | 120.783731 | 551.856515 | 126.430269 | 516.325888 | Sig. genes |
| PF3D7_1353200 | MAHRP2 | 155.248386 | 2.110064882 | 0.52099204 | 4.05009044 | 5.1198E-05 | 0.00376731 | 44.1922939 | 252.179544 | 72.2688306 | 252.352874 | Sig. genes |
| PF3D7_1249700 | N/A | 90.710989 | 2.094626486 | 0.64259961 | 3.25961371 | 0.00111564 | 0.04444108 | 49.5120349 | 113.711072 | 19.7137684 | 179.907081 | Sig. genes |
| PF3D7_1218500 | DRPC | 2143.9741 | 2.076481931 | 0.41444108 | 5.01031879 | 5.434E-07 | 8.4674E-05 | 801.606784 | 3134.63724 | 841.92996 | 3797.72242 | Sig. genes |
| PF3D7_0712300 | VAR | 121.741369 | 2.071024695 | 0.51628393 | 4.0114065 | 6.0358E-05 | 0.0043191 | 48.4819979 | 207.011813 | 45.1811868 | 186.29048 | Sig. genes |
| PF3D7_0215300 | ACS8 | 354.165023 | 2.049524335 | 0.47898478 | 4.27889243 | 1.8783E-05 | 0.00170065 | 171.717492 | 556.750097 | 104.319278 | 583.873227 | Sig. genes |
| PF3D7_0201700 | N/A | 392.7173 | 2.035311804 | 0.49247013 | 4.13286347 | 3.5827E-05 | 0.00279136 | 200.1126 | 605.064838 | 108.421717 | 657.270043 | Sig. genes |
| PF3D7_0702200 | LPL20 | 2826.06953 | 2.01253598 | 0.5664898 | 3.55264295 | 0.00038138 | 0.01870889 | 1713.56397 | 4469.34981 | 532.250343 | 4589.11398 | Sig. genes |
| PF3D7_1255200 | VAR | 334.087504 | 2.006021682 | 0.46264515 | 4.33598339 | 1.4511E-05 | 0.00145055 | 123.312958 | 613.527169 | 142.936856 | 456.573033 | Sig. genes |
| PF3D7_1350600 | N/A | 99.8833485 | 2.004972262 | 0.5359206 | 3.74117407 | 0.00018316 | 0.01054778 | 38.6373644 | 153.77935 | 41.0121085 | 166.104571 | Sig. genes |

| | | | | | | | | | | | | |
|---|---|---|---|---|---|---|---|---|---|---|---|---|
| PF3D7_0814600 | PIMMS22 | 125.132769 | 1.991141988 | 0.51053193 | 3.90013215 | 9.614E-05 | 0.00644748 | 51.3833399 | 199.787365 | 49.2448597 | 200.115509 | Sig. genes |
| PF3D7_0623800 | TKL4 | 5944.58975 | 1.968293526 | 0.53793843 | 3.65895691 | 0.00025324 | 0.01328402 | 3478.32349 | 7608.21066 | 1362.09742 | 11329.7274 | Sig. genes |
| PF3D7_1312500 | N/A | 1202.12762 | 1.966880499 | 0.55479006 | 3.54526992 | 0.00039221 | 0.01906362 | 727.141349 | 2131.12533 | 252.945343 | 1697.29846 | Sig. genes |
| PF3D7_0900100 | VAR | 524.613941 | 1.944129271 | 0.45403043 | 4.28193607 | 1.8527E-05 | 0.00170065 | 246.626116 | 944.960337 | 186.47512 | 720.394193 | Sig. genes |
| PF3D7_1016700 | N/A | 755.814635 | 1.942750759 | 0.48467501 | 4.00835764 | 6.1142E-05 | 0.0043191 | 407.715219 | 1093.10472 | 216.854328 | 1305.58427 | Sig. genes |
| PF3D7_1416500 | GDH1 | 909.765071 | 1.93621152 | 0.43417922 | 4.45947528 | 8.2161E-06 | 0.00086224 | 433.138229 | 1370.68082 | 321.055898 | 1514.18534 | Sig. genes |
| PF3D7_1416600 | N/A | 801.369583 | 1.90410352 | 0.57523842 | 3.31011189 | 0.00093259 | 0.03921306 | 511.329475 | 1076.79519 | 165.125724 | 1452.22795 | Sig. genes |
| PF3D7_0707500 | N/A | 1532.83349 | 1.90240123 | 0.45093726 | 4.21877139 | 2.4564E-05 | 0.00203342 | 766.126145 | 2036.94257 | 528.19712 | 2800.06814 | Sig. genes |
| PF3D7_0828800 | GAMA | 4593.01103 | 1.874758937 | 0.41975212 | 4.46634777 | 7.9566E-06 | 0.00086224 | 2225.58299 | 6722.12521 | 1710.94368 | 7713.39223 | Sig. genes |
| PF3D7_0219900 | N/A | 601.524935 | 1.863898364 | 0.45644133 | 4.08354427 | 4.4354E-05 | 0.00330968 | 315.322037 | 915.633156 | 203.62856 | 971.515988 | Sig. genes |
| PF3D7_0320400 | Cap380 | 323.460567 | 1.860492725 | 0.45173038 | 4.11859107 | 3.812E-05 | 0.00292692 | 133.884967 | 462.259021 | 145.407415 | 552.290863 | Sig. genes |
| PF3D7_0407600 | N/A | 83.7814706 | 1.855959114 | 0.57002975 | 3.25589868 | 0.00113034 | 0.04469064 | 43.176699 | 125.523265 | 29.5689029 | 136.857016 | Sig. genes |
| PF3D7_1147000 | SLARP | 764.523505 | 1.846348325 | 0.43713616 | 4.22373739 | 2.4028E-05 | 0.00202067 | 373.178863 | 1076.82684 | 292.456174 | 1315.63214 | Sig. genes |
| PF3D7_0731200 | N/A | 619.239074 | 1.840748832 | 0.42649621 | 4.31597937 | 1.589E-05 | 0.00155895 | 259.68524 | 923.077115 | 280.823655 | 1013.37029 | Sig. genes |
| PF3D7_1125800 | N/A | 11658.8946 | 1.838727504 | 0.49985991 | 3.67848564 | 0.00023462 | 0.01255588 | 7069.63304 | 19009.0685 | 3120.19673 | 17436.68 | Sig. genes |
| PF3D7_1240400 | VAR | 814.502693 | 1.820426623 | 0.4377688 | 4.15842024 | 3.2046E-05 | 0.00257239 | 403.701937 | 1409.36049 | 315.45181 | 1129.49653 | Sig. genes |
| PF3D7_0408600 | SIAP1 | 207.439476 | 1.793879089 | 0.48367736 | 3.70883414 | 0.00020822 | 0.01158266 | 77.8955982 | 311.163213 | 107.589976 | 333.109115 | Sig. genes |
| PF3D7_0502400 | MSP8 | 709.076547 | 1.784147081 | 0.53953899 | 3.30679916 | 0.00094369 | 0.03925625 | 463.035181 | 1112.76314 | 175.74919 | 1084.75868 | Sig. genes |
| PF3D7_0625400 | N/A | 1119.99268 | 1.764454151 | 0.4556105 | 3.87272492 | 0.00010763 | 0.00695364 | 613.434524 | 1486.24734 | 405.691328 | 1974.59755 | Sig. genes |
| PF3D7_0424500 | FIKK4.1 | 1109.00757 | 1.758921684 | 0.54297593 | 3.23941003 | 0.00119777 | 0.0463197 | 740.468011 | 1582.36383 | 271.878961 | 1841.3195 | Sig. genes |
| PF3D7_0927200 | N/A | 1086.50218 | 1.754079356 | 0.45474147 | 3.85731115 | 0.00011464 | 0.00729089 | 612.242759 | 1548.90999 | 381.974889 | 1802.88108 | Sig. genes |
| PF3D7_1105600 | PTEX88 | 3623.11959 | 1.751719293 | 0.45900712 | 3.81632274 | 0.00013546 | 0.00834468 | 1230.5072 | 4971.01327 | 2087.21902 | 6203.73888 | Sig. genes |
| PF3D7_0501100 | HSP40 | 2784.02104 | 1.741910913 | 0.40935306 | 4.25527757 | 2.0879E-05 | 0.00181339 | 1194.89426 | 4436.15333 | 1368.05572 | 4136.98086 | Sig. genes |
| PF3D7_1026500 | N/A | 315.578242 | 1.727110229 | 0.48082514 | 3.59197158 | 0.00032819 | 0.01655931 | 181.742062 | 479.261264 | 111.45126 | 489.858383 | Sig. genes |
| PF3D7_1114200 | N/A | 830.16875 | 1.721901106 | 0.436721 | 3.94279441 | 8.0538E-05 | 0.00561432 | 410.424379 | 1084.8585 | 362.175788 | 1463.21633 | Sig. genes |
| PF3D7_0202100 | LSAP2 | 7033.35463 | 1.70502735 | 0.46021201 | 3.70487364 | 0.0002115 | 0.01158266 | 4229.55899 | 11437.1001 | 2374.47816 | 10092.2812 | Sig. genes |
| PF3D7_1455300 | N/A | 4332.106 | 1.681283943 | 0.43609909 | 3.85527967 | 0.0001156 | 0.00729089 | 2462.47775 | 6084.9508 | 1656.72134 | 7124.27411 | Sig. genes |
| PF3D7_1226100 | HAD3 | 404.71533 | 1.669024682 | 0.44436071 | 3.75601316 | 0.00017264 | 0.01051312 | 189.341092 | 682.549615 | 197.908086 | 549.062526 | Sig. genes |
| PF3D7_0911100 | N/A | 20608.1451 | 1.65337583 | 0.46856778 | 3.52857345 | 0.00041781 | 0.02012306 | 13053.5237 | 31258.935 | 6830.9401 | 31289.1815 | Sig. genes |
| PF3D7_1345100 | TRX2 | 396.804341 | 1.652792553 | 0.460919 | 3.58586333 | 0.00033596 | 0.01679191 | 224.913083 | 550.327846 | 158.349595 | 653.626842 | Sig. genes |
| PF3D7_0424900 | N/A | 357.040227 | 1.640805647 | 0.44967178 | 3.64889621 | 0.00026337 | 0.01367972 | 177.671127 | 602.154671 | 169.149713 | 479.185396 | Sig. genes |
| PF3D7_0818600 | PBLP | 888.088709 | 1.632913715 | 0.4214782 | 3.87425428 | 0.00010695 | 0.00695364 | 440.081382 | 1452.64168 | 426.087873 | 1233.54391 | Sig. genes |
| PF3D7_0323400 | RIPR | 15444.2683 | 1.62889597 | 0.41887884 | 3.88870439 | 0.00010078 | 0.00667421 | 6405.2536 | 23923.035 | 8688.66346 | 22760.1209 | Sig. genes |
| PF3D7_0424700 | FIKK4.2 | 688.753346 | 1.612673504 | 0.4751541 | 3.394001 | 0.00068879 | 0.03113077 | 427.921594 | 909.555719 | 251.373255 | 1166.16281 | Sig. genes |
| PF3D7_0926600 | N/A | 231.46143 | 1.579646572 | 0.47579143 | 3.32004 | 0.00090005 | 0.03845517 | 133.771439 | 329.344377 | 98.5667352 | 364.163168 | Sig. genes |
| PF3D7_0613600 | N/A | 3709.1296 | 1.578712473 | 0.47685094 | 3.31070436 | 0.00093061 | 0.03921306 | 1255.65592 | 5826.15329 | 2465.02412 | 5289.68508 | Sig. genes |
| PF3D7_1351700 | IMC1f | 23806.516 | 1.522817589 | 0.40804107 | 3.73202039 | 0.00018995 | 0.01070591 | 11083.2052 | 34607.3263 | 13500.4704 | 36035.0622 | Sig. genes |
| PF3D7_1205500 | N/A | 7989.33905 | 1.517681285 | 0.4433238 | 3.42341487 | 0.0006184 | 0.02848924 | 3345.28081 | 10126.3977 | 4926.3982 | 13559.2795 | Sig. genes |

| | | | | | | | | | | | | |
|---|---|---|---|---|---|---|---|---|---|---|---|---|
| PF3D7_0805100 | N/A | 513.896912 | 1.48799776 | 0.43146884 | 3.44867958 | 0.00056333 | 0.02641192 | 279.959394 | 804.767787 | 260.344214 | 710.516253 | Sig. genes |
| PF3D7_0808200 | PMX | 10218.1918 | 1.468265335 | 0.42716535 | 3.43722949 | 0.0005877 | 0.02731246 | 6380.27537 | 15866.634 | 4470.60515 | 14155.2529 | Sig. genes |
| PF3D7_0410000 | EVP1 | 4045.54152 | 1.457792137 | 0.40729594 | 3.57919637 | 0.00034465 | 0.01706513 | 2261.37734 | 5593.55824 | 2057.56516 | 6269.66535 | Sig. genes |
| PF3D7_0414900 | ARO | 6107.41942 | 1.44844614 | 0.43536338 | 3.32698201 | 0.00087792 | 0.03781482 | 3949.14932 | 9367.89703 | 2602.25494 | 8510.37637 | Sig. genes |
| PF3D7_0505300 | N/A | 548.238968 | 1.442038456 | 0.43180903 | 3.33952824 | 0.00083921 | 0.03644365 | 319.933391 | 773.663842 | 270.196533 | 829.162105 | Sig. genes |
| PF3D7_1000100 | VAR | 1096.99964 | 1.414599286 | 0.43211943 | 3.2736304 | 0.00106175 | 0.04292299 | 694.549885 | 1626.06661 | 502.742451 | 1564.63961 | Sig. genes |
| PF3D7_1342600 | MyoA | 31953.5556 | 1.406548463 | 0.43234987 | 3.25326445 | 0.00114087 | 0.04477292 | 21025.3396 | 44217.2146 | 13982.8513 | 48588.8168 | Sig. genes |
| PF3D7_1133400 | AMA1 | 21789.1908 | 1.400775761 | 0.43515011 | 3.21906331 | 0.0012861 | 0.04901987 | 9482.44733 | 33414.5172 | 14458.4847 | 29801.3138 | Sig. genes |
| PF3D7_0611800 | N/A | 7992.45088 | 1.345976948 | 0.40724756 | 3.30505838 | 0.00094957 | 0.03925625 | 4366.93535 | 10543.7465 | 4658.85981 | 12400.2619 | Sig. genes |
| PF3D7_1224300 | PABP1 | 24772.3412 | 1.317519026 | 0.40892152 | 3.22193612 | 0.00127328 | 0.0488827 | 12950.2511 | 33237.8334 | 15422.6709 | 37478.6096 | Sig. genes |
| PF3D7_MIT03200 | LSUA | 7324.63787 | -1.347241159 | 0.41179958 | -3.2715943 | 0.00106943 | 0.04292299 | 9895.67961 | 4537.51761 | 11136.3001 | 3729.05414 | Sig. genes |
| PF3D7_1404100 | N/A | 830.012023 | -1.388978911 | 0.42869925 | -3.2399844 | 0.00119536 | 0.0463197 | 1259.94585 | 511.104026 | 1142.73345 | 406.264763 | Sig. genes |
| PF3D7_1443300 | LSM5 | 798.292996 | -1.490353357 | 0.43067238 | -3.4605269 | 0.00053912 | 0.02550227 | 1206.32458 | 472.918831 | 1148.6781 | 365.250482 | Sig. genes |
| PF3D7_1201000 | N/A | 3967.78319 | -1.508072073 | 0.4036358 | -3.7362198 | 0.00018681 | 0.010642 | 5853.91436 | 2121.25787 | 5888.73235 | 2007.2282 | Sig. genes |
| PF3D7_1402500 | N/A | 4011.35636 | -1.600051439 | 0.44213976 | -3.6188816 | 0.00029588 | 0.01507276 | 4701.31282 | 2113.32949 | 7364.01106 | 1866.77206 | Sig. genes |
| PF3D7_1215300 | CPN10 | 438.315435 | -1.620017886 | 0.44014973 | -3.6806064 | 0.00023268 | 0.01255588 | 654.1475 | 236.433889 | 668.738459 | 193.941891 | Sig. genes |
| PF3D7_1371800 | N/A | 380.228371 | -1.647470792 | 0.44475832 | -3.7041933 | 0.00021206 | 0.01158266 | 620.790284 | 173.84341 | 532.160455 | 194.119335 | Sig. genes |
| PF3D7_1012400 | HGPRT | 17803.5278 | -1.73363895 | 0.41393801 | -4.1881608 | 2.8122E-05 | 0.00229219 | 26196.6999 | 9250.65131 | 28554.2115 | 7212.54854 | Sig. genes |
| PF3D7_0531800 | N/A | 157.860367 | -1.761658126 | 0.52311897 | -3.3676051 | 0.00075824 | 0.03347634 | 224.815571 | 89.6272455 | 262.725716 | 54.2729349 | Sig. genes |
| PF3D7_0725800 | N/A | 157.381643 | -1.770385407 | 0.52190033 | -3.3921906 | 0.00069336 | 0.03113077 | 224.815571 | 88.5207363 | 261.91733 | 54.2729349 | Sig. genes |
| PF3D7_0831400 | N/A | 1170.36336 | -1.844193704 | 0.42957284 | -4.2930873 | 1.7621E-05 | 0.00169734 | 1595.49945 | 548.422474 | 2066.0765 | 471.45501 | Sig. genes |
| PF3D7_1418800 | N/A | 151.529805 | -1.889788335 | 0.52074212 | -3.6290292 | 0.00028449 | 0.01463323 | 232.220476 | 79.2557002 | 245.106676 | 49.5363668 | Sig. genes |
| PF3D7_1300900 | N/A | 36.4857526 | -3.503478456 | 0.93590327 | -3.7434194 | 0.00018153 | 0.01054778 | 28.2499639 | 8.59841733 | 105.843269 | 3.25135996 | Sig. genes |
| PF3D7_0831700 | HSP70x | 2945.8736 | -5.349920842 | 0.63334428 | -8.4470975 | 2.9863E-17 | 4.2116E-14 | 4003.77977 | 55.7842585 | 7497.68102 | 226.249347 | Sig. genes |
| PF3D7_0833500 | VAR | 174.274271 | -6.323076102 | 0.7492005 | -8.4397649 | 3.1798E-17 | 4.2116E-14 | 290.892224 | 1.07259982 | 397.5945 | 7.5377596 | Sig. genes |
| PF3D7_0832200 | N/A | 568.705507 | -7.184742289 | 0.70639762 | -10.17096 | 2.6726E-24 | 1.4159E-20 | 553.742048 | 5.68699238 | 1705.25665 | 10.1363337 | Sig. genes |
| PF3D7_0832000 | N/A | 16.4691074 | -7.379516641 | 1.72688827 | -4.2733029 | 1.926E-05 | 0.00170065 | 37.8442988 | 0 | 28.0321308 | 0 | Sig. genes |
| PF3D7_0832900 | N/A | 33.2932526 | -8.400279964 | 1.66772937 | -5.0369563 | 4.7299E-07 | 8.0836E-05 | 32.5886336 | 0 | 100.584377 | 0 | Sig. genes |

### Upregulated genes

| Gene ID | Short name | Product description | Biological process | Cellular compartment | Molecular function | Stage peak expression | Sex | Molecular weight | baseMean | log2FoldChange | lfcSE | stat | pvalue | padj | 01_JP_S1 | 02_JP_S2 | 03_JP_S3 | 04_JP_S4 | sig |
|---|---|---|---|---|---|---|---|---|---|---|---|---|---|---|---|---|---|---|---|
| PF3D7_1255100 | RIF | rifin | modulation by symbiont of host erythrocyte aggregation; antigenic variation; cytoadherence to microvasculature, mediated by symbiont protein | host cell plasma membrane; symbiont-containing vacuole; Maurer's cleft | N/A | RG | no | 39.7 | 147.465 | 5.92161 | 0.726487 | 8.151025 | 3.61E-16 | 3.19E-13 | 8.381701 | 339.5078 | 1.641238 | 240.3291 | Sig. genes |
| PF3D7_0101600 | RIF | rifin | modulation by symbiont of host erythrocyte aggregation; antigenic variation; cytoadherence to microvasculature, mediated by symbiont protein | integral component of membrane; host cell plasma membrane; symbiont-containing vacuole; Maurer's cleft | N/A | RG | male | 41.4 | 37.13317 | 5.661534 | 1.110493 | 5.098218 | 3.43E-07 | 6.24E-05 | 3.143283 | 88.05671 | 0 | 57.3327 | Sig. genes |
| PF3D7_0412400 | VAR | erythrocyte membrane protein 1, PfEMP1 | obsolete pathogenesis; modulation by symbiont of host erythrocyte aggregation; antigenic variation; cytoadherence to microvasculature, mediated by symbiont protein; cell-cell adhesion | integral component of membrane; host cell plasma membrane; infected host cell surface knob | host cell surface receptor binding; cell adhesion molecule binding | RG | no | 250.8 | 324.6426 | 4.676104 | 0.673382 | 6.944208 | 3.81E-12 | 1.83E-09 | 42.16328 | 549.1528 | 7.393438 | 699.8609 | Sig. genes |
| PF3D7_0631900 | N/A | stevor | modulation by symbiont of host erythrocyte aggregation; entry into host | integral component of membrane; rhoptry; Maurer's cleft; host cell membrane | N/A | RG | no | 34.1 | 163.5287 | 4.120673 | 0.666641 | 6.181244 | 6.36E-10 | 1.98E-07 | 6.276716 | 362.2263 | 28.7163 | 256.8955 | Sig. genes |
| PF3D7_0413100 | VAR | erythrocyte membrane protein 1, PfEMP1 | obsolete pathogenesis; modulation by symbiont of host erythrocyte aggregation; antigenic variation; cytoadherence to microvasculature, mediated by symbiont protein; cell-cell adhesion | integral component of membrane; host cell plasma membrane; infected host cell surface knob | host cell surface receptor binding; cell adhesion molecule binding | RG | (male) | 263.7 | 71.29061 | 4.064506 | 0.90283 | 4.501961 | 6.73E-06 | 0.000759 | 0 | 162.9866 | 15.60356 | 106.5723 | Sig. genes |
| PF3D7_0425200 | N/A | Plasmodium exported protein (hyp15), unknown function | N/A | integral component of membrane; host cell | N/A | RG | male | 27.0 | 143.6602 | 4.021693 | 0.584223 | 6.883829 | 5.83E-12 | 2.37E-09 | 23.07811 | 285.6895 | 10.67038 | 255.2028 | Sig. genes |
| PF3D7_0712900 | VAR | erythrocyte membrane protein 1, PfEMP1 | obsolete pathogenesis; modulation by symbiont of host erythrocyte aggregation; antigenic variation; cytoadherence to microvasculature, mediated by symbiont protein; cell-cell adhesion | integral component of membrane; host cell plasma membrane; infected host cell surface knob | host cell surface receptor binding; cell adhesion molecule binding | RG | no | 248.9 | 197.6093 | 3.965676 | 0.513562 | 7.721906 | 1.15E-14 | 8.67E-12 | 26.34808 | 347.5255 | 21.35826 | 395.2054 | Sig. genes |
| PF3D7_0425100 | N/A | Plasmodium exported protein (hyp6), unknown function | N/A | integral component of membrane; host cell | N/A | RG | no | 24.1 | 64.44114 | 3.856354 | 0.782927 | 4.925557 | 8.41E-07 | 0.00012 | 14.63137 | 138.6929 | 2.461023 | 101.9793 | Sig. genes |
| PF3D7_1479000 | ACS1a | acyl-CoA synthetase | fatty acid metabolic process | endoplasmic reticulum; membrane; host cell cytoplasm | long-chain fatty acid-CoA ligase activity; decanoate-CoA ligase activity | OK | no | 94.9 | 484.3482 | 3.851159 | 0.45234 | 8.513866 | 1.68E-17 | 4.21E-14 | 67.38199 | 973.1376 | 58.31406 | 838.5593 | Sig. genes |
| PF3D7_1148200 | N/A | non-coding RNA | N/A | N/A | N/A | OK | no | N/A | 747.805 | 3.718373 | 0.45094 | 8.245831 | 1.64E-16 | 1.74E-13 | 117.2011 | 1188.369 | 94.21944 | 1591.43 | Sig. genes |
| PF3D7_1370500 | N/A | non-coding RNA | N/A | N/A | N/A | OK | female | N/A | 959.8062 | 3.636647 | 0.501276 | 7.254781 | 4.02E-13 | 2.66E-10 | 194.4909 | 1633.147 | 91.80682 | 1919.78 | Sig. genes |
| PF3D7_0935600 | GIG | gametocytogenesis-implicated protein | N/A | Maurer's cleft | N/A | RG | male | 58.3 | 3296.125 | 3.499472 | 0.513014 | 6.821399 | 9.02E-12 | 3.41E-09 | 758.0262 | 6160.75 | 313.7446 | 5951.98 | Sig. genes |
| PF3D7_1479200 | N/A | Plasmodium exported protein (PHISTa), unknown function | N/A | host cell | N/A | RG | male | 34.0 | 373.6069 | 3.452431 | 0.688661 | 5.013253 | 5.35E-07 | 8.47E-05 | 107.6729 | 742.8704 | 18.04818 | 625.8363 | Sig. genes |
| PF3D7_0202200 | PTP1 | EMP1-trafficking protein | N/A | membrane; integral component of membrane; Maurer's cleft; host cell cytoplasm; protein-containing complex | protein-containing complex binding | RG | male | 33.7 | 368.8968 | 3.445387 | 0.487182 | 7.072073 | 1.53E-12 | 8.99E-10 | 76.84466 | 610.5199 | 47.63544 | 740.5871 | Sig. genes |

| | | | | | | | | | | | | | | | | | | | |
|---|---|---|---|---|---|---|---|---|---|---|---|---|---|---|---|---|---|---|---|
| PF3D7_0102200 | RESA | ring-infected erythrocyte surface antigen | regulation of immune response | host cell plasma membrane; symbiont-containing vacuole; food vacuole; merozoite dense granule; host cell surface; extracellular vesicle | protein binding | RG | no | 126.5 | 27786.88 | 3.382109 | 0.758001 | 4.46188 | 8.12E-06 | 0.000862 | 8832.879 | 49243.1 | 895.3845 | 52176.17 | Sig. genes |
| PF3D7_0712600 | VAR | erythrocyte membrane protein 1, PfEMP1 | obsolete pathogenesis; modulation by symbiont of host erythrocyte aggregation; antigenic variation; cytoadherence to microvasculature, mediated by symbiont protein; cell-cell adhesion | integral component of membrane; host cell plasma membrane; infected host cell surface knob | host cell surface receptor binding; cell adhesion molecule binding | RG | female | 251.7 | 154.6969 | 3.350531 | 0.538382 | 6.223334 | 4.87E-10 | 1.61E-07 | 31.61801 | 232.7556 | 23.82272 | 330.5912 | Sig. genes |
| PF3D7_0702000 | N/A | Plasmodium exported protein (hyp12), unknown function | N/A | Maurer's cleft; host cell | N/A | RG | (male) | 28.8 | 827.7902 | 3.325777 | 0.729984 | 4.555958 | 5.21E-06 | 0.000614 | 267.2464 | 1683.722 | 33.67045 | 1326.522 | Sig. genes |
| PF3D7_0829800 | N/A | non-coding RNA | N/A | N/A | N/A | OK | no | N/A | 2864.003 | 3.228718 | 0.469006 | 6.884172 | 5.81E-12 | 2.37E-09 | 388.3549 | 4811.472 | 715.3275 | 5540.858 | Sig. genes |
| PF3D7_0712800 | VAR | erythrocyte membrane protein 1, PfEMP1 | obsolete pathogenesis; modulation by symbiont of host erythrocyte aggregation; antigenic variation; cytoadherence to microvasculature, mediated by symbiont protein; cell-cell adhesion | integral component of membrane; host cell plasma membrane; infected host cell surface knob | host cell surface receptor binding; cell adhesion molecule binding | RG | female | 256.5 | 67.63442 | 3.193438 | 0.625506 | 5.105371 | 3.3E-07 | 6.24E-05 | 12.64733 | 130.8576 | 13.96507 | 113.0676 | Sig. genes |
| PF3D7_0831800 | HRP2 | histidine-rich protein II | heme catabolic process | cytoplasm; food vacuole; Maurer's cleft; host cell cytoplasm; host cell membrane | N/A | RG | no | 32.4 | 966.0386 | 3.192154 | 0.459575 | 6.945887 | 3.76E-12 | 1.83E-09 | 143.2197 | 1782.003 | 237.3724 | 1701.559 | Sig. genes |
| PF3D7_0702300 | STARP | sporozoite threonine and asparagine-rich protein | N/A | cell surface | N/A | RG | male | 66.0 | 312.4021 | 3.180436 | 0.665818 | 4.776738 | 1.78E-06 | 0.000242 | 104.2511 | 538.577 | 20.53379 | 586.2465 | Sig. genes |
| PF3D7_0711700 | VAR | erythrocyte membrane protein 1, PfEMP1 | obsolete pathogenesis; modulation by symbiont of host erythrocyte aggregation; antigenic variation; cytoadherence to microvasculature, mediated by symbiont protein; cell-cell adhesion | integral component of membrane; host cell plasma membrane; infected host cell surface knob | host cell surface receptor binding; cell adhesion molecule binding | RG | female | 251.0 | 120.4061 | 3.110292 | 0.657083 | 4.733483 | 2.21E-06 | 0.000285 | 38.99554 | 173.7623 | 11.50062 | 257.3658 | Sig. genes |
| PF3D7_0412900 | VAR | erythrocyte membrane protein 1, PfEMP1 | obsolete pathogenesis; modulation by symbiont of host erythrocyte aggregation; antigenic variation; cytoadherence to microvasculature, mediated by symbiont protein; cell-cell adhesion | integral component of membrane; host cell plasma membrane; infected host cell surface knob | host cell surface receptor binding; cell adhesion molecule binding | RG | no | 261.8 | 131.2116 | 3.094992 | 0.55789 | 5.54767 | 2.9E-08 | 6.97E-06 | 34.78119 | 263.8583 | 20.53697 | 205.67 | Sig. genes |
| PF3D7_0221700 | N/A | Plasmodium exported protein, unknown function | N/A | Maurer's cleft; host cell | N/A | RG | no | 58.3 | 269.1004 | 2.920735 | 0.835675 | 3.495062 | 0.000474 | 0.022622 | 117.8769 | 582.6529 | 8.212742 | 367.6592 | Sig. genes |
| PF3D7_1200700 | ACS7 | acyl-CoA synthetase | N/A | endoplasmic reticulum; membrane; Maurer's cleft | long-chain fatty acid-CoA ligase activity; decanoate-CoA ligase activity | RG | no | 104.6 | 2200.687 | 2.909516 | 0.597115 | 4.872619 | 1.1E-06 | 0.000154 | 818.6158 | 3753.426 | 216.034 | 4014.671 | Sig. genes |
| PF3D7_0412700 | VAR | erythrocyte membrane protein 1, PfEMP1 | obsolete pathogenesis; modulation by symbiont of host erythrocyte aggregation; antigenic variation; cytoadherence to microvasculature, mediated by symbiont protein; cell-cell adhesion | integral component of membrane; host cell plasma membrane; infected host cell surface knob | host cell surface receptor binding; cell adhesion molecule binding | RG | female | 256.1 | 128.0127 | 2.902326 | 0.579171 | 5.011171 | 5.41E-07 | 8.47E-05 | 41.11119 | 254.1942 | 19.716 | 197.0292 | Sig. genes |
| PF3D7_0935900 | REX1 | ring-exported protein 1 | N/A | cytoplasm; integral component of membrane; Maurer's cleft; host cell cytoplasm; host cell membrane; extracellular vesicle | protein binding | RG | male | 83.0 | 1819.538 | 2.862709 | 0.712501 | 4.017829 | 5.87E-05 | 0.004263 | 774.3337 | 3031.525 | 105.9627 | 3366.332 | Sig. genes |

| | | | | | | | | | | | | | | | | | | | |
|---|---|---|---|---|---|---|---|---|---|---|---|---|---|---|---|---|---|---|---|
| PF3D7_0207300 | SERA8 | serine repeat antigen 8 | proteolysis; proteolysis involved in cellular protein catabolic process | extracellular space; lysosome | cysteine-type endopeptidase activity; cysteine-type peptidase activity | OK | male | 79.4 | 85.60244 | 2.830283 | 0.572841 | 4.940787 | 7.78E-07 | 0.000118 | 19.97909 | 148.1148 | 22.17093 | 152.145 | Sig. genes |
| PF3D7_1352900 | N/A | Plasmodium exported protein, unknown function | N/A | Maurer's cleft; host cell | N/A | RG | male | 68.9 | 225.3346 | 2.828141 | 0.555346 | 5.092579 | 3.53E-07 | 6.24E-05 | 78.94411 | 372.3284 | 32.8516 | 417.2144 | Sig. genes |
| PF3D7_0315200 | CTRP | circumsporozoite- and TRAP-related protein | B cell activation involved in immune response; entry into host; cell-cell adhesion | microneme | GPI anchor binding; heparan sulfate proteoglycan binding | OK | no | 237.9 | 472.7495 | 2.773316 | 0.585596 | 4.735889 | 2.18E-06 | 0.000285 | 184.4301 | 724.0187 | 57.50266 | 925.0467 | Sig. genes |
| PF3D7_1401100 | N/A | DnaJ protein, putative | N/A | integral component of membrane; Maurer's cleft; host cell | N/A | RG | no | 57.1 | 1616.079 | 2.756782 | 0.414466 | 6.651409 | 2.9E-11 | 1.03E-08 | 446.471 | 2873.159 | 386.8539 | 2757.831 | Sig. genes |
| PF3D7_0424800 | N/A | Plasmodium exported protein (PHISTb), unknown function | N/A | host cell | N/A | RG | male | 36.7 | 67.34819 | 2.729786 | 0.727331 | 3.753154 | 0.000175 | 0.010513 | 28.33111 | 138.5377 | 7.387551 | 95.13637 | Sig. genes |
| PF3D7_1407700 | N/A | conserved protein, unknown function | N/A | inner membrane pellicle complex | N/A | OK | female | 47.3 | 44.50787 | 2.705923 | 0.820468 | 3.298024 | 0.000974 | 0.039681 | 19.92862 | 94.51712 | 4.103901 | 59.48186 | Sig. genes |
| PF3D7_1221400 | IMC1h | inner membrane complex protein 1h, putative | N/A | inner membrane pellicle complex | N/A | OK | female | 60.1 | 230.6148 | 2.697021 | 0.471726 | 5.717343 | 1.08E-08 | 2.87E-06 | 59.9672 | 418.5265 | 63.23374 | 380.7319 | Sig. genes |
| PF3D7_0801300 | WARP | von Willebrand factor A domain-related protein | cell adhesion | microneme lumen | heparin binding; LBD domain binding; cell adhesion molecule binding | OK | no | 32.0 | 148.6704 | 2.678989 | 0.684665 | 3.912847 | 9.12E-05 | 0.006239 | 66.91948 | 269.8234 | 13.94689 | 243.9921 | Sig. genes |
| PF3D7_1115800 | N/A | conserved Plasmodium protein, unknown function | N/A | nucleus; cell surface; Maurer's cleft | N/A | RG | no | 33.9 | 3664.662 | 2.631313 | 0.600426 | 4.382413 | 1.17E-05 | 0.001196 | 1620.523 | 6523.126 | 417.2418 | 6097.757 | Sig. genes |
| PF3D7_0902500 | FIKK9.6 | serine/threonine protein kinase, FIKK family | protein phosphorylation; intracellular signal transduction | nucleus; cytoplasm; Maurer's cleft | protein kinase activity; protein serine/threonine kinase activity; ATP binding | RG | male | 70.1 | 666.2874 | 2.614374 | 0.450921 | 5.797857 | 6.72E-09 | 1.87E-06 | 224.2263 | 1166.306 | 150.299 | 1124.318 | Sig. genes |
| PF3D7_0301800 | N/A | Plasmodium exported protein, unknown function | N/A | integral component of membrane; Maurer's cleft; host cell | N/A | RG | male | 30.8 | 1186.035 | 2.594849 | 0.503477 | 5.153856 | 2.55E-07 | 5.41E-05 | 460.8364 | 2241.751 | 213.5194 | 1828.033 | Sig. genes |
| PF3D7_1401400 | ETRAMP14 | early transcribed membrane protein 14.1 | N/A | symbiont-containing vacuole membrane | N/A | RG | male | 11.4 | 3110.534 | 2.56885 | 0.502352 | 5.113647 | 3.16E-07 | 6.2E-05 | 1247.436 | 5330.674 | 547.7288 | 5316.298 | Sig. genes |
| PF3D7_1002100 | PTP5 | EMP1-trafficking protein | N/A | Maurer's cleft; host cell cytosol | protein binding | RG | no | 70.0 | 2209.432 | 2.514083 | 0.429305 | 5.85617 | 4.74E-09 | 1.39E-06 | 771.6665 | 3710.246 | 545.3513 | 3810.463 | Sig. genes |
| PF3D7_1148900 | N/A | Plasmodium exported protein, unknown function | N/A | Maurer's cleft | N/A | RG | male | 10.3 | 120.3484 | 2.483205 | 0.580707 | 4.276174 | 1.9E-05 | 0.001701 | 50.46901 | 195.3447 | 22.99172 | 212.5882 | Sig. genes |
| PF3D7_1149600 | N/A | DnaJ protein, putative | N/A | Maurer's cleft; host cell | N/A | RG | male | 83.1 | 2033.236 | 2.481025 | 0.547533 | 4.531281 | 5.86E-06 | 0.000675 | 918.2548 | 3414.918 | 317.8633 | 3481.907 | Sig. genes |
| PF3D7_0501300 | SBP1 | skeleton-binding protein 1 | protein transport | nucleus; cytoplasm; membrane; integral component of membrane; symbiont-containing vacuolar space; symbiont-containing vacuole membrane; Maurer's cleft; extracellular vesicle | phosphoprotein phosphatase activity; protein binding; metal ion binding | RG | no | 36.3 | 1674.193 | 2.476219 | 0.539387 | 4.5908 | 4.42E-06 | 0.000532 | 747.329 | 2989.366 | 273.489 | 2686.586 | Sig. genes |
| PF3D7_1149200 | RESA3 | ring-infected erythrocyte surface antigen | N/A | Maurer's cleft | N/A | (RG); OK | male | 126.9 | 21673.44 | 2.452373 | 0.731789 | 3.351202 | 0.000805 | 0.03523 | 11953.5 | 33672.4 | 1439.927 | 39627.95 | Sig. genes |
| PF3D7_1479100 | N/A | Plasmodium exported protein, unknown function | N/A | host cell | N/A | ITZ; mGC | no | 24.9 | 89.39295 | 2.447874 | 0.741059 | 3.303211 | 0.000956 | 0.039256 | 46.82944 | 179.7579 | 9.017598 | 121.9669 | Sig. genes |
| PF3D7_1404800 | N/A | conserved Plasmodium protein, unknown function | N/A | Maurer's cleft; host cell | N/A | RG | male | 122.8 | 2671.085 | 2.442106 | 0.46027 | 5.305808 | 1.12E-07 | 2.58E-05 | 1036.712 | 3920.759 | 624.2819 | 5102.588 | Sig. genes |
| PF3D7_0108500 | N/A | ELM2 domain-containing protein, putative | N/A | nucleus; Maurer's cleft | N/A | RG | no | 19.7 | 3058.681 | 2.440883 | 0.664139 | 3.675262 | 0.000238 | 0.012588 | 1607.79 | 5764.695 | 295.6801 | 4566.558 | Sig. genes |
| PF3D7_0424600 | N/A | Plasmodium exported protein (PHISTb) | cytoadherence to microvasculature, mediated by symbiont protein | nucleus; host cell surface; host cell periphery; extracellular vesicle | N/A | RG; (OK) | male | 36.0 | 5168.903 | 2.436398 | 0.436996 | 5.575337 | 2.47E-08 | 6.23E-06 | 1954.689 | 8994.563 | 1269.837 | 8456.525 | Sig. genes |

| | | | | | | | | | | | | | | | | | | | |
|---|---|---|---|---|---|---|---|---|---|---|---|---|---|---|---|---|---|---|---|
| PF3D7_0605900 | N/A | elongation of fatty acids protein, putative | fatty acid elongation, saturated fatty acid; fatty acid elongation, unsaturated fatty acid; sphingolipid biosynthetic process; fatty acid elongation, monounsaturated fatty acid; fatty acid elongation, polyunsaturated fatty acid; very long-chain fatty acid biosynthetic process | endoplasmic reticulum; integral component of membrane; integral component of endoplasmic reticulum membrane | fatty acid elongase activity; ligase activity; 3-oxo-arachidoyl-CoA synthase activity; 3-oxo-cerotoyl-CoA synthase activity; 3-oxo-lignoceronyl-CoA synthase activity; very-long-chain 3-ketoacyl-CoA synthase activity | OK | female | 34.7 | 942.6564 | 2.360429 | 0.479147 | 4.926313 | 8.38E-07 | 0.00012 | 223.17 | 1343.35 | 390.9406 | 1813.165 | Sig. genes |
| PF3D7_0324300 | EPF4 | exported protein family 4 | N/A | integral component of membrane; Maurer's cleft | N/A | nonunique (RG) | no | 35.7 | 188.9823 | 2.352753 | 0.601547 | 3.91117 | 9.18E-05 | 0.006239 | 88.93806 | 404.1216 | 35.28131 | 227.5884 | Sig. genes |
| PF3D7_1029700 | N/A | conserved Plasmodium protein, unknown function | N/A | N/A | N/A | OK | no | 91.3 | 41.4947 | 2.345738 | 0.695823 | 3.371169 | 0.000748 | 0.033324 | 11.57186 | 66.53621 | 15.60295 | 72.26778 | Sig. genes |
| PF3D7_0805200 | GAMER | gamete release protein, putative | cell motility | cytoplasm; inner membrane pellicle complex | N/A | OK | female | 10.3 | 2003.945 | 2.266209 | 0.667921 | 3.392932 | 0.000691 | 0.031131 | 1170.852 | 3524.609 | 209.2755 | 3111.041 | Sig. genes |
| PF3D7_0730800 | N/A | Plasmodium exported protein, unknown function | N/A | host cell | N/A | RG | no | 30.1 | 3903.991 | 2.202954 | 0.588254 | 3.744905 | 0.00018 | 0.010548 | 2185.117 | 6690.156 | 602.0029 | 6138.69 | Sig. genes |
| PF3D7_1102700 | ETRAMP11.1 | early transcribed membrane protein 11.1 | N/A | symbiont-containing vacuole membrane | N/A | RG | male | 10.2 | 5302.53 | 2.194062 | 0.572 | 3.835776 | 0.000125 | 0.007802 | 2921.708 | 9348.033 | 882.8165 | 8057.561 | Sig. genes |
| PF3D7_0402100 | N/A | Plasmodium exported protein (PHISTb), unknown function | N/A | integral component of membrane; Maurer's cleft; host cell | protein binding | RG | no | 67.8 | 4354.065 | 2.182185 | 0.489576 | 4.457292 | 8.3E-06 | 0.000862 | 2138.366 | 6896.57 | 1006.861 | 7374.464 | Sig. genes |
| PF3D7_0731500 | EBA175 | erythrocyte binding antigen-175 | obsolete pathogenesis; entry into host; modulation by symbiont of host receptor-mediated signal transduction | integral component of membrane; microneme; extracellular vesicle | heparin binding; host cell surface receptor binding | OK | (female) | 174.6 | 60363.99 | 2.170623 | 0.422827 | 5.133595 | 2.84E-07 | 5.79E-05 | 24181.24 | 85027.82 | 19702.7 | 112544.2 | Sig. genes |
| PF3D7_0202500 | ETRAMP2 | early transcribed membrane protein 2 | N/A | nucleus; symbiont-containing vacuole membrane; extracellular vesicle | N/A | RG | male | 11.5 | 10880.5 | 2.159423 | 0.525496 | 4.109306 | 3.97E-05 | 0.003004 | 5702.914 | 19971.35 | 2258.036 | 15589.7 | Sig. genes |
| PF3D7_1201200 | N/A | Plasmodium exported protein (PHISTa-like), unknown function | N/A | Maurer's cleft; host cell | N/A | RG | male | 25.9 | 2396.511 | 2.151158 | 0.410652 | 5.238396 | 1.62E-07 | 3.58E-05 | 838.2317 | 4145.525 | 923.2015 | 3679.085 | Sig. genes |
| PF3D7_0402900 | N/A | probable protein, unknown function | N/A | N/A | N/A | RG; mGC | (male) | 11.6 | 89.89226 | 2.145273 | 0.572476 | 3.747357 | 0.000179 | 0.010548 | 37.72431 | 171.8942 | 28.72257 | 121.2279 | Sig. genes |
| PF3D7_0302200 | CLAG3.2 | cytoadherence linked asexual protein 3.2 | cytoadherence to microvasculature, mediated by symbiont protein | nucleus; host cell plasma membrane; rhoptry | anion channel activity; protein binding; cell adhesion molecule binding | SZ | male | 167.5 | 2772.124 | 2.141431 | 0.517976 | 4.134232 | 3.56E-05 | 0.002791 | 617.3779 | 3783.476 | 1430.914 | 5256.727 | Sig. genes |
| PF3D7_0219700 | GEXP20 | Plasmodium exported protein (PHISTc), unknown function | N/A | Maurer's cleft; host cell | N/A | RG | female | 35.3 | 194.7373 | 2.136944 | 0.499102 | 4.28158 | 1.86E-05 | 0.001701 | 87.14 | 332.8686 | 57.46432 | 301.4765 | Sig. genes |
| PF3D7_0814500 | N/A | conserved protein, unknown function | N/A | N/A | N/A | RG | no | 90.3 | 525.6111 | 2.135294 | 0.459103 | 4.65101 | 3.3E-06 | 0.000407 | 231.6516 | 769.2587 | 158.5188 | 943.0153 | Sig. genes |
| PF3D7_0102700 | MaTrA | merozoite-associated tryptophan-rich antigen | N/A | cell surface | N/A | OK | male | 34.5 | 340.9755 | 2.130169 | 0.504278 | 4.224197 | 2.4E-05 | 0.002021 | 167.0659 | 591.5127 | 87.02978 | 518.2937 | Sig. genes |
| PF3D7_1478800 | N/A | Plasmodium exported protein, unknown function | N/A | Maurer's cleft; host cell | N/A | RG | male | 24.1 | 328.8491 | 2.111019 | 0.446802 | 4.72473 | 2.3E-06 | 0.000291 | 120.7837 | 551.8565 | 126.4303 | 516.3259 | Sig. genes |
| PF3D7_1353200 | MAHRP2 | membrane associated histidine-rich protein 2 | N/A | nucleus; integral component of membrane; Maurer's cleft; extracellular vesicle | protein binding | RG | male | 15.8 | 155.2484 | 2.110065 | 0.520992 | 4.05009 | 5.12E-05 | 0.003767 | 44.19229 | 252.1795 | 72.26883 | 252.3529 | Sig. genes |
| PF3D7_1249700 | N/A | conserved Plasmodium protein, unknown function | N/A | N/A | N/A | OK | male | 105.8 | 90.71099 | 2.094626 | 0.6426 | 3.259614 | 0.001116 | 0.044441 | 49.51203 | 113.7111 | 19.71377 | 179.9071 | Sig. genes |
| PF3D7_1218500 | DRPC | dynamin-like protein, putative | mitochondrial fission; mitochondrion organization | nucleus; cytoplasm; mitochondrion; membrane | GTPase activity; GTP binding; microtubule binding | RG | male | 122.8 | 2143.974 | 2.076482 | 0.414441 | 5.010319 | 5.43E-07 | 8.47E-05 | 801.6068 | 3134.637 | 841.93 | 3797.722 | Sig. genes |
| PF3D7_0712300 | VAR | erythrocyte membrane protein 1, PfEMP1 | obsolete pathogenesis; modulation by symbiont of host erythrocyte aggregation; antigenic variation; cytoadherence to microvasculature, mediated by symbiont protein; cell-cell adhesion | integral component of membrane; host cell plasma membrane; infected host cell surface knob | host cell surface receptor binding; cell adhesion molecule binding | RG | female | 257.0 | 121.7414 | 2.071025 | 0.516284 | 4.011406 | 6.04E-05 | 0.004319 | 48.482 | 207.0118 | 45.18119 | 186.2905 | Sig. genes |

| | | | | | | | | | | | | | | | | | | | |
|---|---|---|---|---|---|---|---|---|---|---|---|---|---|---|---|---|---|---|---|
| PF3D7_0215300 | ACS8 | acyl-CoA synthetase | fatty acid metabolic process; long-chain fatty acid transport | endoplasmic reticulum; membrane | long-chain fatty acid-CoA ligase activity; decanoate-CoA ligase activity | RG | male | 102.7 | 354.165 | 2.049524 | 0.478985 | 4.278892 | 1.88E-05 | 0.001701 | 171.7175 | 556.7501 | 104.3193 | 583.8732 | Sig. genes |
| PF3D7_0201700 | N/A | DnaJ protein, putative | N/A | integral component of membrane; Maurer's cleft | N/A | RG | no | 105.9 | 392.7173 | 2.035312 | 0.49247 | 4.132863 | 3.58E-05 | 0.002791 | 200.1126 | 605.0648 | 108.4217 | 657.27 | Sig. genes |
| PF3D7_0702200 | LPL20 | lysophospholipase LPL20 | phospholipid metabolic process; glycerophospholipid catabolic process | cytosol; membrane; vesicle; cell periphery | lysophospholipase activity; lipase activity; hydrolase activity; phosphatidyl phospholipase B activity | RG | male | 49.2 | 2826.07 | 2.012536 | 0.56649 | 3.552643 | 0.000381 | 0.018709 | 1713.564 | 4469.35 | 532.2503 | 4589.114 | Sig. genes |
| PF3D7_1255200 | VAR | erythrocyte membrane protein 1, PfEMP1 | obsolete pathogenesis; modulation by symbiont of host erythrocyte aggregation; antigenic variation; cytoadherence to microvasculature, mediated by symbiont protein; cell-cell adhesion | integral component of membrane; host cell plasma membrane; infected host cell surface knob | integrin binding; host cell surface receptor binding; cell adhesion molecule binding | RG | no | 255.4 | 334.0875 | 2.006022 | 0.462645 | 4.335983 | 1.45E-05 | 0.001451 | 123.313 | 613.5272 | 142.9369 | 456.573 | Sig. genes |
| PF3D7_1350600 | N/A | conserved Plasmodium protein, unknown function | N/A | integral component of membrane | N/A | mGC | no | 31.4 | 99.88335 | 2.004972 | 0.535921 | 3.741174 | 0.000183 | 0.010548 | 38.63736 | 153.7793 | 41.01211 | 166.1046 | Sig. genes |
| PF3D7_0814600 | PIMMS22 | structural integrity maintenance protein SIMP, putative | regulation of cell shape; cell gliding | cytoplasm; cell surface; membrane; cell periphery | N/A | OK | female | 44.8 | 125.1328 | 1.991142 | 0.510532 | 3.900132 | 9.61E-05 | 0.006447 | 51.38334 | 199.7874 | 49.24486 | 200.1155 | Sig. genes |
| PF3D7_0623800 | TKL4 | tyrosine kinase-like protein, putative | protein phosphorylation | N/A | protein kinase activity; protein serine/threonine kinase activity; non-membrane spanning protein tyrosine kinase activity; ATP binding | RG | male | 279.8 | 5944.59 | 1.968294 | 0.537938 | 3.658957 | 0.000253 | 0.013284 | 3478.323 | 7608.211 | 1362.097 | 11329.73 | Sig. genes |
| PF3D7_1312500 | N/A | conserved Plasmodium protein, unknown function | N/A | N/A | N/A | OK | male | 7.2 | 1202.128 | 1.96688 | 0.55479 | 3.54527 | 0.000392 | 0.019064 | 727.1413 | 2131.125 | 252.9453 | 1697.298 | Sig. genes |
| PF3D7_0900100 | VAR | erythrocyte membrane protein 1, PfEMP1 | obsolete pathogenesis; cell-cell adhesion | integral component of membrane; host cell plasma membrane; infected host cell surface knob | host cell surface receptor binding; cell adhesion molecule binding | RG | male | 258.4 | 524.6139 | 1.944129 | 0.45403 | 4.281936 | 1.85E-05 | 0.001701 | 246.6261 | 944.9603 | 186.4751 | 720.3942 | Sig. genes |
| PF3D7_1016700 | N/A | Plasmodium exported protein (PHISTc), unknown function | N/A | Maurer's cleft | N/A | RG | no | 98.8 | 755.8146 | 1.942751 | 0.484675 | 4.008358 | 6.11E-05 | 0.004319 | 407.7152 | 1093.105 | 216.8543 | 1305.584 | Sig. genes |
| PF3D7_1416500 | GDH1 | NADP-specific glutamate dehydrogenase | cellular amino acid metabolic process; glutamate metabolic process; glutamate biosynthetic process; oxidation-reduction process | nucleus; cytoplasm; cytosol | nucleotide binding; glutamate dehydrogenase (NADP+) activity; oxidoreductase activity; oxidoreductase activity, acting on the CH-NH2 group of donors, NAD or NADP as acceptor | RG | female | 52.5 | 909.7651 | 1.936212 | 0.434179 | 4.459475 | 8.22E-06 | 0.000862 | 433.1382 | 1370.681 | 321.0559 | 1514.185 | Sig. genes |
| PF3D7_1416600 | N/A | conserved Plasmodium protein, unknown function | N/A | N/A | N/A | mGC | no | 357.6 | 801.3696 | 1.904104 | 0.575238 | 3.310112 | 0.000933 | 0.039213 | 511.3295 | 1076.795 | 165.1257 | 1452.228 | Sig. genes |
| PF3D7_0707500 | N/A | conserved Plasmodium protein, unknown function | N/A | N/A | N/A | RG; (OK) | male | 194.4 | 1532.833 | 1.902401 | 0.450937 | 4.218771 | 2.46E-05 | 0.002033 | 766.1261 | 2036.943 | 528.1971 | 2800.068 | Sig. genes |
| PF3D7_0828800 | GAMA | GPI-anchored micronemal antigen | entry into host | cytoplasm; cell surface; integral component of membrane; microneme; extrinsic component of cell outer membrane; osmiophilic body; anchored component of plasma membrane | host cell surface receptor binding; host cell surface binding | OK | no | 85.3 | 4593.011 | 1.874759 | 0.419752 | 4.466348 | 7.96E-06 | 0.000862 | 2225.583 | 6722.125 | 1710.944 | 7713.392 | Sig. genes |
| PF3D7_0219900 | N/A | Plasmodium exported protein, unknown function | N/A | host cell | N/A | RG | no | 27.0 | 601.5249 | 1.863898 | 0.456441 | 4.083544 | 4.44E-05 | 0.00331 | 315.322 | 915.6332 | 203.6286 | 971.516 | Sig. genes |
| PF3D7_0320400 | Cap380 | oocyst capsule protein Cap380 | N/A | cell surface | N/A | OK | no | 360.5 | 323.4606 | 1.860493 | 0.45173 | 4.118591 | 3.81E-05 | 0.002927 | 133.885 | 462.259 | 145.4074 | 552.2909 | Sig. genes |
| PF3D7_0407600 | N/A | conserved Plasmodium protein, unknown function | N/A | N/A | N/A | OK | no | 142.9 | 83.78147 | 1.855959 | 0.57003 | 3.255899 | 0.00113 | 0.044691 | 43.1767 | 125.5233 | 29.5689 | 136.857 | Sig. genes |
| PF3D7_1147000 | SLARP | sporozoite and liver stage asparagine-rich protein | regulation of gene expression | nucleus | N/A | OK | no | 348.0 | 764.5235 | 1.846348 | 0.437136 | 4.223737 | 2.4E-05 | 0.002021 | 373.1789 | 1076.827 | 292.4562 | 1315.632 | Sig. genes |

| | | | | | | | | | | | | | | | | | | | |
|---|---|---|---|---|---|---|---|---|---|---|---|---|---|---|---|---|---|---|---|
| PF3D7_0731200 | N/A | Plasmodium exported protein, unknown function | N/A | integral component of membrane; Maurer's cleft; host cell | N/A | RG | male | 37.0 | 619.2391 | 1.840749 | 0.426496 | 4.315979 | 1.59E-05 | 0.001559 | 259.6852 | 923.0771 | 280.8237 | 1013.37 | Sig. genes |
| PF3D7_1125800 | N/A | kelch domain-containing protein, putative | N/A | N/A | protein binding | OK | female | 67.6 | 11658.89 | 1.838728 | 0.49986 | 3.678486 | 0.000235 | 0.012556 | 7069.633 | 19009.07 | 3120.197 | 17436.68 | Sig. genes |
| PF3D7_1240400 | VAR | erythrocyte membrane protein 1, PfEMP1 | obsolete pathogenesis; modulation by symbiont of host erythrocyte aggregation; antigenic variation; cytoadherence to microvasculature, mediated by symbiont protein; cell-cell adhesion | integral component of membrane; host cell plasma membrane; infected host cell surface knob | host cell surface receptor binding; cell adhesion molecule binding | RG | no | 260.6 | 814.5027 | 1.820427 | 0.437769 | 4.15842 | 3.2E-05 | 0.002572 | 403.7019 | 1409.36 | 315.4518 | 1129.497 | Sig. genes |
| PF3D7_0408600 | SIAP1 | sporozoite invasion-associated protein 1 | N/A | cell surface; apical part of cell | N/A | OK | male | 112.9 | 207.4395 | 1.793879 | 0.483677 | 3.708834 | 0.000208 | 0.011583 | 77.8956 | 311.1632 | 107.59 | 333.1091 | Sig. genes |
| PF3D7_0502400 | MSP8 | merozoite surface protein 8 | N/A | plasma membrane; membrane; integral component of membrane; symbiont-containing vacuole; food vacuole; Maurer's cleft | N/A | RG | male | 69.4 | 709.0765 | 1.784147 | 0.539539 | 3.306799 | 0.000944 | 0.039256 | 463.0352 | 1112.763 | 175.7492 | 1084.759 | Sig. genes |
| PF3D7_0625400 | N/A | conserved Plasmodium protein, unknown function | N/A | N/A | N/A | RG | no | 48.5 | 1119.993 | 1.764454 | 0.455611 | 3.872725 | 0.000108 | 0.006954 | 613.4345 | 1486.247 | 405.6913 | 1974.598 | Sig. genes |
| PF3D7_0424500 | FIKK4.1 | serine/threonine protein kinase, FIKK family | protein phosphorylation; cytoadherence to microvasculature, mediated by symbiont protein; intracellular signal transduction; modulation by symbiont of host process | nucleus; cytoplasm; integral component of membrane; Maurer's cleft; host cell periphery | protein kinase activity; protein serine/threonine kinase activity; ATP binding | RG | male | 74.4 | 1109.008 | 1.758922 | 0.542976 | 3.23941 | 0.001198 | 0.04632 | 740.468 | 1582.364 | 271.879 | 1841.319 | Sig. genes |
| PF3D7_0927200 | N/A | gametocyte development protein GD1, putative | N/A | cytosol | metal ion binding | OK | no | 189.7 | 1086.502 | 1.754079 | 0.454741 | 3.857311 | 0.000115 | 0.007291 | 612.2428 | 1548.91 | 381.9749 | 1802.881 | Sig. genes |
| PF3D7_1105600 | PTEX88 | translocon component PTEX88 | translocation of peptides or proteins into host cell cytoplasm | symbiont-containing vacuole; symbiont-containing vacuole membrane; merozoite dense granule; PTEX complex | RNA binding; protein binding | RG | male | 90.8 | 3623.12 | 1.751719 | 0.459007 | 3.816323 | 0.000135 | 0.008345 | 1230.507 | 4971.013 | 2087.219 | 6203.739 | Sig. genes |
| PF3D7_0501100 | HSP40 | heat shock protein 40, type II, co-chaperone J domain protein JDP | protein folding; positive regulation of ATP-dependent activity; chaperone cofactor-dependent protein refolding | cytosol; integral component of membrane; host cell cytosol | ATPase activator activity; protein binding; unfolded protein binding; chaperone binding | RG | male | 46.4 | 2784.021 | 1.741911 | 0.409353 | 4.255278 | 2.09E-05 | 0.001813 | 1194.894 | 4436.153 | 1368.056 | 4136.981 | Sig. genes |
| PF3D7_1026500 | N/A | conserved Plasmodium protein, unknown function | N/A | N/A | N/A | OK | female | 8.4 | 315.5782 | 1.72711 | 0.480825 | 3.591972 | 0.000328 | 0.016559 | 181.7421 | 479.2613 | 111.4513 | 489.8584 | Sig. genes |
| PF3D7_1114200 | N/A | GTPase-activating protein, putative | intracellular protein transport; activation of GTPase activity | Maurer's cleft | GTPase activator activity | RG | no | 44.6 | 830.1687 | 1.721901 | 0.436721 | 3.942794 | 8.05E-05 | 0.005614 | 410.4244 | 1084.859 | 362.1758 | 1463.216 | Sig. genes |
| PF3D7_0202100 | LSAP2 | liver stage associated protein 2 | N/A | Maurer's cleft; host cell | N/A | (RG); OK | male | 35.3 | 7033.355 | 1.705027 | 0.460212 | 3.704874 | 0.000211 | 0.011583 | 4229.559 | 11437.1 | 2374.478 | 10092.28 | Sig. genes |
| PF3D7_1455300 | N/A | conserved protein, unknown function | N/A | nucleus | N/A | OK | female | 69.5 | 4332.106 | 1.681284 | 0.436099 | 3.85528 | 0.000116 | 0.007291 | 2462.478 | 6084.951 | 1656.721 | 7124.274 | Sig. genes |
| PF3D7_1226100 | HAD3 | haloacid dehalogenase-like hydrolase, putative | N/A | cytosol | magnesium ion binding; catalytic activity; alkaline phosphatase activity; hydrolase activity; phosphatase activity | mGC | no | 36.2 | 404.7153 | 1.669025 | 0.444361 | 3.756013 | 0.000173 | 0.010513 | 189.3411 | 682.5496 | 197.9081 | 549.0625 | Sig. genes |
| PF3D7_0911100 | N/A | START domain-containing protein, putative | N/A | N/A | N/A | OK | female | 137.1 | 20608.15 | 1.653376 | 0.468568 | 3.528573 | 0.000418 | 0.020123 | 13053.52 | 31258.94 | 6830.94 | 31289.18 | Sig. genes |
| PF3D7_1345100 | TRX2 | thioredoxin 2 | cell redox homeostasis | cytoplasm; mitochondrion; symbiont-containing vacuole; PTEX complex | protein binding; protein-disulfide reductase activity; oxidoreductase activity, acting on a sulfur group of donors, disulfide as acceptor; protein-disulfide reductase (NAD(P)) activity | RG | no | 18.6 | 396.8043 | 1.652793 | 0.460919 | 3.585863 | 0.000336 | 0.016792 | 224.9131 | 550.3278 | 158.3496 | 653.6268 | Sig. genes |

| | | | | | | | | | | | | | | | | | | | |
|---|---|---|---|---|---|---|---|---|---|---|---|---|---|---|---|---|---|---|---|
| PF3D7_0424900 | N/A | Plasmodium exported protein (PHISTa), unknown function | N/A | integral component of membrane; host cell | N/A | RG | male | 33.2 | 357.0402 | 1.640806 | 0.449672 | 3.648896 | 0.000263 | 0.01368 | 177.6711 | 602.1547 | 169.1497 | 479.1854 | Sig. genes |
| PF3D7_0818600 | PBLP | BEM46-like protein, putative | lipid metabolic process; entry into host; cell maturation | plasma membrane; cell surface; membrane; integral component of membrane | palmitoyl-(protein) hydrolase activity; lipase activity | OK | no | 34.9 | 888.0887 | 1.632914 | 0.421478 | 3.874254 | 0.000107 | 0.006954 | 440.0814 | 1452.642 | 426.0879 | 1233.544 | Sig. genes |
| PF3D7_0323400 | RIPR | Rh5 interacting protein | entry into host; protein insertion into membrane | extracellular region; plasma membrane; membrane; host cell plasma membrane; microneme; cytoplasmic vesicle; protein-containing complex; host cell membrane; microneme lumen; host extracellular space | protein binding; host cell surface binding | OK | female | 125.9 | 15444.27 | 1.628896 | 0.418879 | 3.888704 | 0.000101 | 0.006674 | 6405.254 | 23923.04 | 8688.663 | 22760.12 | Sig. genes |
| PF3D7_0424700 | FIKK4.2 | serine/threonine protein kinase, FIKK family | protein phosphorylation; intracellular signal transduction; modulation by symbiont of host process | nucleus; cytoplasm; integral component of membrane; Maurer's cleft; host cell cytoplasm; host cell periphery | protein kinase activity; protein serine/threonine kinase activity; ATP binding; kinase activity | RG | no | 143.0 | 688.7533 | 1.612674 | 0.475154 | 3.394001 | 0.000689 | 0.031131 | 427.9216 | 909.5557 | 251.3733 | 1166.163 | Sig. genes |
| PF3D7_0926600 | N/A | conserved Plasmodium membrane protein, unknown function | N/A | membrane; integral component of membrane | N/A | OK | no | 163.2 | 231.4614 | 1.579647 | 0.475791 | 3.32004 | 0.0009 | 0.038455 | 133.7714 | 329.3444 | 98.56674 | 364.1632 | Sig. genes |
| PF3D7_0613600 | N/A | conserved Plasmodium protein, unknown function | N/A | integral component of membrane | N/A | mGC | female | 123.7 | 3709.13 | 1.578712 | 0.476851 | 3.310704 | 0.000931 | 0.039213 | 1255.656 | 5826.153 | 2465.024 | 5289.685 | Sig. genes |
| PF3D7_1351700 | IMC1f | inner membrane complex protein 1f | N/A | basal part of cell; inner membrane pellicle complex | N/A | OK | no | 151.5 | 23806.52 | 1.522818 | 0.408041 | 3.73202 | 0.00019 | 0.010706 | 11083.21 | 34607.33 | 13500.47 | 36035.06 | Sig. genes |
| PF3D7_1205500 | N/A | zinc finger protein, putative | protein ubiquitination | nucleus; cytoplasm | protein binding; ubiquitin protein ligase activity | RG | no | 204.9 | 7989.339 | 1.517681 | 0.443324 | 3.423415 | 0.000618 | 0.028489 | 3345.281 | 10126.4 | 4926.398 | 13559.28 | Sig. genes |
| PF3D7_0805100 | N/A | conserved protein, unknown function | N/A | N/A | N/A | OK | female | 64.5 | 513.8969 | 1.487998 | 0.431469 | 3.44868 | 0.000563 | 0.026412 | 279.9594 | 804.7678 | 260.3442 | 710.5163 | Sig. genes |
| PF3D7_0808200 | PMX | plasmepsin X | proteolysis; response to xenobiotic stimulus; exit from host cell; entry into host | exoneme; apical part of cell | aspartic-type endopeptidase activity | OK | female | 65.1 | 10218.19 | 1.468265 | 0.427165 | 3.437229 | 0.000588 | 0.027312 | 6380.275 | 15866.63 | 4470.605 | 14155.25 | Sig. genes |
| PF3D7_0410000 | EVP1 | erythrocyte vesicle protein 1 | N/A | integral component of membrane; host cell cytoplasmic vesicle; host cell cytosol | N/A | OK | no | 100.5 | 4045.542 | 1.457792 | 0.407296 | 3.579196 | 0.000345 | 0.017065 | 2261.377 | 5593.558 | 2057.565 | 6269.665 | Sig. genes |
| PF3D7_0414900 | ARO | armadillo-domain containing rhoptry protein | N/A | nucleus; cytosol; rhoptry; anchored component of membrane; apical part of cell | DNA binding | SZ | male | 30.8 | 6107.419 | 1.448446 | 0.435363 | 3.326982 | 0.000878 | 0.037815 | 3949.149 | 9367.897 | 2602.255 | 8510.376 | Sig. genes |
| PF3D7_0505300 | N/A | UDP-N-acetylglucosamine transporter, putative | pyrimidine nucleotide-sugar transmembrane transport | Golgi membrane; Golgi apparatus; integral component of membrane; integral component of Golgi membrane | sialic acid transmembrane transporter activity; pyrimidine nucleotide-sugar transmembrane transporter activity; | OK | male | 74.0 | 548.239 | 1.442038 | 0.431809 | 3.339528 | 0.000839 | 0.036444 | 319.9334 | 773.6638 | 270.1965 | 829.1621 | Sig. genes |
| PF3D7_1000100 | VAR | erythrocyte membrane protein 1, PfEMP1 | obsolete pathogenesis; modulation by symbiont of host erythrocyte aggregation; antigenic variation; cytoadherence to microvasculature, mediated by symbiont protein; cell-cell adhesion | integral component of membrane; host cell plasma membrane; infected host cell surface knob | host cell surface receptor binding; cell adhesion molecule binding | RG | no | 252.3 | 1097 | 1.414599 | 0.432119 | 3.27363 | 0.001062 | 0.042923 | 694.5499 | 1626.067 | 502.7425 | 1564.64 | Sig. genes |
| PF3D7_1342600 | MyoA | myosin A | actin filament organization; response to xenobiotic stimulus; vesicle transport along actin filament; entry into host; cell motility | nucleus; cytoplasm; cytosol; plasma membrane; cell surface; actin cytoskeleton; myosin complex; pellicle; vesicle; perinuclear region of cytoplasm; inner membrane pellicle complex; cell periphery | microfilament motor activity; cytoskeletal motor activity; actin binding; protein binding; ATP binding; ATP hydrolysis activity; actin-dependent ATPase activity; actin filament binding | OK | female | 92.3 | 31953.56 | 1.406548 | 0.43235 | 3.253264 | 0.001141 | 0.044773 | 21025.34 | 44217.21 | 13982.85 | 48588.82 | Sig. genes |

| | | | | | | | | | | | | | | | | | | |
|---|---|---|---|---|---|---|---|---|---|---|---|---|---|---|---|---|---|---|
| PF3D7_1133400 | AMA1 | apical membrane antigen 1 | obsolete pathogenesis; entry into host | plasma membrane; membrane; integral component of membrane; apical complex; microneme | protein binding; host cell surface binding | OK | no | 72.0 | 21789.19 | 1.400776 | 0.43515 | 3.219063 | 0.001286 | 0.04902 | 9482.447 | 33414.52 | 14458.48 | 29801.31 | Sig. genes |
| PF3D7_0611800 | N/A | conserved Plasmodium protein, unknown function | N/A | nucleus | N/A | OK | female | 374.9 | 7992.451 | 1.345977 | 0.407248 | 3.305058 | 0.00095 | 0.039256 | 4366.935 | 10543.75 | 4658.86 | 12400.26 | Sig. genes |
| PF3D7_1224300 | PABP1 | polyadenylate-binding protein 1, putative | RNA processing | nucleus; cytoplasm; cytosol; cell surface; nuclear periphery; P granule; ribonucleoprotein complex | nucleic acid binding; RNA binding; mRNA binding; mRNA 3'-UTR binding; poly(A) binding; poly(U) RNA binding | RG | no | 97.2 | 24772.34 | 1.317519 | 0.408922 | 3.221936 | 0.001273 | 0.048883 | 12950.25 | 33237.83 | 15422.67 | 37478.61 | Sig. genes |

log-fold change ≥2

### Downregulated genes

| Gene ID | Short name | Product description | Biological process | Cellular compartment | Molecular function | Stage peak expression | Sex | Molecular weight | baseMean | log2FoldChange | lfcSE | stat | pvalue | padj | 01_JP_S1 | 02_JP_S2 | 03_JP_S3 | 04_JP_S4 | sig |
|---|---|---|---|---|---|---|---|---|---|---|---|---|---|---|---|---|---|---|---|
| PF3D7_MIT03200 | LSUA | large subunit ribosomal RNA fragment A | N/A | N/A | N/A | mGC | no | N/A | 7324.638 | -1.34724 | 0.4118 | -3.27159 | 0.001069 | 0.042923 | 9895.68 | 4537.518 | 11136.3 | 3729.054 | Sig. genes |
| PF3D7_1404100 | N/A | cytochrome c, putative | mitochondrial electron transport, ubiquinol to cytochrome c; mitochondrial electron transport, cytochrome c to oxygen; oxidation-reduction process | mitochondrion; mitochondrial respirasome; mitochondrial intermembrane space; Maurer's cleft | electron transfer activity; heme binding; metal ion binding | mGC | no | 12.9 | 830.012 | -1.38898 | 0.428699 | -3.23998 | 0.001195 | 0.04632 | 1259.946 | 511.104 | 1142.733 | 406.2648 | Sig. genes |
| PF3D7_1443300 | LSM5 | U6 snRNA-associated Sm-like protein LSm5, putative | mRNA splicing, via spliceosome; RNA splicing | nucleus; spliceosomal complex; U6 snRNP; sno(s)RNA-containing ribonucleoprotein complex; U4/U6 x U5 tri-snRNP complex; Lsm1-7-Pat1 complex | RNA binding | RG | no | 11.4 | 798.293 | -1.49035 | 0.430672 | -3.46053 | 0.000539 | 0.025502 | 1206.325 | 472.9188 | 1148.678 | 365.2505 | Sig. genes |
| PF3D7_1201000 | N/A | Plasmodium exported protein (PHISTb), unknown function | N/A | nucleus; symbiont-containing vacuole; infected host cell surface knob; host cell surface; host cell periphery | protein binding | eTZ | male | 77.1 | 3967.783 | -1.50807 | 0.403636 | -3.73622 | 0.000187 | 0.010642 | 5853.914 | 2121.258 | 5888.732 | 2007.228 | Sig. genes |
| PF3D7_1402500 | N/A | ribosomal protein S27a, putative | translation; protein ubiquitination; modification-dependent protein catabolic process | nucleus; cytoplasm; ribosome | structural constituent of ribosome; protein binding; protein tag; ubiquitin protein ligase binding; metal ion binding | RG | no | 17.4 | 4011.356 | -1.60005 | 0.44214 | -3.61888 | 0.000296 | 0.015073 | 4701.313 | 2113.329 | 7364.011 | 1866.772 | Sig. genes |
| PF3D7_1215300 | CPN10 | 10 kDa chaperonin | protein folding; response to unfolded protein; response to heat; chaperone cofactor-dependent protein refolding | mitochondrion; mitochondrial matrix; food vacuole | ATP binding; metal ion binding; unfolded protein binding; chaperone binding | OK | no | 11.2 | 438.3154 | -1.62002 | 0.44015 | -3.68061 | 0.000233 | 0.012556 | 654.1475 | 236.4339 | 668.7385 | 193.9419 | Sig. genes |
| PF3D7_1371800 | N/A | Plasmodium exported protein, unknown function | N/A | integral component of membrane; host cell | N/A | eTZ | male | 53.4 | 380.2284 | -1.64747 | 0.444758 | -3.70419 | 0.000212 | 0.011583 | 620.7903 | 173.8434 | 532.1605 | 194.1193 | Sig. genes |
| PF3D7_1012400 | HGPRT | hypoxanthine-guanine phosphoribosyltransferase | purine ribonucleoside salvage; adenine salvage; guanine salvage; nucleoside metabolic process; response to xenobiotic stimulus; GMP salvage; IMP salvage; hypoxanthine metabolic process | cytoplasm | nucleotide binding; magnesium ion binding; RNA binding; hypoxanthine phosphoribosyltransferase activity; guanine phosphoribosyltransferase activity | eTZ | female | 26.4 | 17803.53 | -1.73364 | 0.413938 | -4.18816 | 2.81E-05 | 0.002292 | 26196.7 | 9250.651 | 28554.21 | 7212.549 | Sig. genes |
| PF3D7_0531800 | N/A | 5.8S ribosomal RNA | N/A | N/A | N/A | eTZ | no | N/A | 157.8604 | -1.76166 | 0.523119 | -3.36761 | 0.000758 | 0.033476 | 224.8156 | 89.62725 | 262.7257 | 54.27293 | Sig. genes |
| PF3D7_0725800 | N/A | 5.8S ribosomal RNA | N/A | N/A | N/A | eTZ | no | N/A | 157.3816 | -1.77039 | 0.5219 | -3.39219 | 0.000693 | 0.031131 | 224.8156 | 88.52074 | 261.9173 | 54.27293 | Sig. genes |
| PF3D7_0831400 | N/A | Plasmodium exported protein, unknown function | N/A | host cell | N/A | eTZ | male | 36.4 | 1170.363 | -1.84419 | 0.429573 | -4.29309 | 1.76E-05 | 0.001697 | 1595.499 | 548.4225 | 2066.076 | 471.455 | Sig. genes |
| PF3D7_1418800 | N/A | signal recognition particle RNA | N/A | N/A | N/A | OK | no | N/A | 151.5298 | -1.88979 | 0.520742 | -3.62903 | 0.000284 | 0.014633 | 232.2205 | 79.2557 | 245.1067 | 49.53637 | Sig. genes |
| PF3D7_1300900 | N/A | stevor | modulation by symbiont of host erythrocyte aggregation | integral component of membrane; Maurer's cleft | N/A | RG | male | 34.7 | 36.48575 | -3.50348 | 0.935903 | -3.74342 | 0.000182 | 0.010548 | 28.24996 | 8.598417 | 105.8433 | 3.25136 | Sig. genes |
| PF3D7_0831700 | HSP70x | heat shock protein 70 | response to unfolded protein; cytoadherence to microvasculature, mediated by symbiont protein; cellular response to unfolded protein; protein refolding; chaperone cofactor-dependent protein refolding | nucleus; cytoplasm; symbiont-containing vacuole; Maurer's cleft; host cell cytoplasm; protein-containing complex; host cell cytosol; extracellular vesicle | RNA binding; protein binding; ATP binding; ATP hydrolysis activity; heat shock protein binding; protein folding chaperone; unfolded protein binding; misfolded protein binding | RG | male | 75.1 | 2945.874 | -5.34992 | 0.633344 | -8.4471 | 2.99E-17 | 4.21E-14 | 4003.78 | 55.78426 | 7497.681 | 226.2493 | Sig. genes |
| PF3D7_0833500 | VAR | erythrocyte membrane protein 1, PfEMP1 | obsolete pathogenesis | integral component of membrane | host cell surface receptor binding | RG | male | 255.8 | 174.2743 | -6.32308 | 0.749201 | -8.43976 | 3.18E-17 | 4.21E-14 | 290.8922 | 1.0726 | 397.5945 | 7.53776 | Sig. genes |
| PF3D7_0832200 | N/A | Plasmodium exported protein (PHISTa-like), unknown function | N/A | integral component of membrane; host cell; host cell cytosol | N/A | eTZ | male | 33.5 | 568.7055 | -7.18474 | 0.706398 | -10.171 | 2.67E-24 | 1.42E-20 | 553.742 | 5.686992 | 1705.257 | 10.13633 | Sig. genes |
| PF3D7_0832000 | N/A | stevor | modulation by symbiont of host erythrocyte aggregation | integral component of membrane; Maurer's cleft | N/A | RG | male | 34.4 | 16.46911 | -7.37952 | 1.726888 | -4.2733 | 1.93E-05 | 0.001701 | 37.8443 | 0 | 28.03213 | 0 | Sig. genes |
| PF3D7_0832900 | N/A | stevor | modulation by symbiont of host erythrocyte aggregation | integral component of membrane; Maurer's cleft | N/A | RG | no | 33.9 | 33.29325 | -8.40028 | 1.667729 | -5.03696 | 4.73E-07 | 8.08E-05 | 32.58863 | 0 | 100.5844 | 0 | Sig. genes |

log-fold change ≤-2
