## Supplementary material for "*Plasmodium falciparum* SET10 is a histone H3 lysine K18 methyltransferase that participates in a chromatin modulation network crucial for intraerythrocytic development": Table S2

Table S2. List of peptides following *Pf* SET10 TurboID

| Protein IDs | Majority protein IDs | Peptide counts (all) | Peptide counts (razor+unique) | Peptide counts (unique) | Protein names | Gene names | Fasta headers |
|---|---|---|---|---|---|---|---|
| Q8I586 | Q8I586 | 178 | 178 | 178 | | PFL1620w | [illegible] |
| C6KSY0 | C6KSY0 | 98 | 98 | 98 | | ApiAP2 | [illegible] |
| Q8IKF6 | Q8IKF6 | 68 | 68 | 68 | | PF14_064 | [illegible] |
| C0H4K6 | C0H4K6 | 90 | 90 | 90 | | MAL7P1.1 | [illegible] |
| Q8I350 | Q8I350 | 79 | 79 | 79 | | PFI0495w | [illegible] |
| Q8IJW2 | Q8IJW2 | 78 | 78 | 78 | | PF10_007 | [illegible] |
| O77384 | O77384 | 82 | 82 | 82 | Protein Pl | PFC0760c | [illegible] |
| Q8I5K3 | Q8I5K3 | 70 | 70 | 70 | | PFL1010c | [illegible] |
| Q8I2Z8 | Q8I2Z8 | 53 | 53 | 53 | Probable | PFI0755c | [illegible] |
| Q8I1X5 | Q8I1X5 | 64 | 64 | 64 | | PFD0265w | [illegible] |
| Q8I2X4 | Q8I2X4 | 30 | 30 | 30 | | PFI0875w | [illegible] |
| O77380 | O77380 | 63 | 63 | 63 | | PFC0780w | [illegible] |
| C6KST7 | C6KST7 | 72 | 72 | 72 | | PFE0445w | [illegible] |
| Q8I531;BF | Q8I531 | 69;1 | 69;1 | 69;1 | | ApiAP2 | [illegible] |
| Q8IB24 | Q8IB24 | 31 | 31 | 26 | | PF08_005 | [illegible] |
| C6KSQ6 | C6KSQ6 | 61 | 61 | 61 | Probable | PFF0285c | [illegible] |
| C6KTD2 | C6KTD2 | 70 | 70 | 70 | Putative h | SET1 | [illegible] |
| C0H552 | C0H552 | 75 | 75 | 75 | | PFI1120c | [illegible] |
| Q8I4R5 | Q8I4R5 | 61 | 61 | 61 | | PFL2505c | [illegible] |
| C6KTB7 | C6KTB7 | 83 | 83 | 83 | Putative E | PFF1365c | [illegible] |
| Q8I5L6 | Q8I5L6 | 50 | 50 | 50 | Clathrin h | PFL0930w | [illegible] |
| A0A144A4 | A0A144A4 | 58 | 58 | 58 | | | [illegible] |
| Q8IES7 | Q8IES7 | 76 | 76 | 75 | | MAL13P1 | [illegible] |
| Q8I3U0 | Q8I3U0 | 55 | 55 | 55 | | PFE0840c | [illegible] |
| C0H571 | C0H571 | 43 | 43 | 43 | | RhopH2 | [illegible] |
| C6KTA4 | C6KTA4 | 21 | 21 | 21 | Pyruvate k | PFF1300w | [illegible] |
| A0A5K1K8 | A0A5K1K8 | 41 | 41 | 41 | | | [illegible] |
| Q8ILB9 | Q8ILB9 | 71 | 71 | 71 | | PF14_032 | [illegible] |
| C0H5A6 | C0H5A6 | 61 | 61 | 61 | | MAL13P1 | [illegible] |
| Q8I3M5 | Q8I3M5 | 43 | 43 | 43 | | PFE1195w | [illegible] |
| A0A144A1 | A0A144A1 | 34;34 | 34;34 | 34;34 | Protein SE | PF14_015 | [illegible] |
| Q8IBJ1 | Q8IBJ1 | 53 | 53 | 53 | Ubiquitin | MAL7P1.1 | [illegible] |
| Q8ILC9 | Q8ILC9 | 56 | 56 | 56 | | PF14_031 | [illegible] |
| A0A144A0 | A0A144A0 | 49;49 | 49;49 | 49;49 | | PF11_029 | [illegible] |
| Q8IFP1 | Q8IFP1 | 51 | 51 | 51 | | PFD1060w | [illegible] |
| Q8IJZ9 | Q8IJZ9 | 41 | 41 | 41 | | PF10_004 | [illegible] |
| O77375 | O77375 | 45 | 45 | 45 | DNA-direc | PFC0805w | [illegible] |
| Q8IKM7 | Q8IKM7 | 41 | 41 | 41 | | PF14_057 | [illegible] |
| Q8ILZ2 | Q8ILZ2 | 55 | 55 | 55 | | PF14_010 | [illegible] |
| A0A144A1 | A0A144A1 | 18;18 | 18;18 | 16;16 | Actin-1 | PFL2215w | [illegible] |
| O77382 | O77382 | 42 | 42 | 41 | Kinesin-li | PFC0770c | [illegible] |
| Q8IEN3 | Q8IEN3 | 39 | 39 | 39 | | cpsSII | [illegible] |
| Q8I0V4 | Q8I0V4 | 28 | 28 | 28 | | PFL1070c | [illegible] |
| Q8IJR8 | Q8IJR8 | 39 | 39 | 39 | | PF10_012 | [illegible] |
| Q8IBU8 | Q8IBU8 | 41 | 41 | 41 | | PF07_006 | [illegible] |
| Q8IAQ8 | Q8IAQ8 | 25 | 25 | 25 | | PF08_011 | [illegible] |
| C6KT34 | C6KT34 | 34 | 34 | 33 | | PFF0940c | [illegible] |
| C0H5L6 | C0H5L6 | 46 | 46 | 46 | | MAL13P1 | [illegible] |
| Q8IB94 | Q8IB94 | 51 | 51 | 51 | | MAL8P1.2 | [illegible] |
| O97292 | O97292 | 42 | 42 | 42 | | PFC0965w | [illegible] |
| Q8I0P6 | Q8I0P6 | 21 | 21 | 21 | Elongation factor 1-al | | [illegible] |
| Q9U0H9 | Q9U0H9 | 35 | 35 | 35 | | PFD0540c | [illegible] |
| Q8IKW5 | Q8IKW5 | 25 | 25 | 25 | | PF14_048 | [illegible] |
| Q8IBB0 | Q8IBB0 | 38 | 38 | 38 | Histone-ly | PF08_001 | [illegible] |
| Q8IBY8 | Q8IBY8 | 52 | 52 | 52 | | PF07_004 | [illegible] |
| Q8IEE5 | Q8IEE5 | 31 | 31 | 30 | | PF13_009 | [illegible] |
| Q8IKK7 | Q8IKK7 | 19 | 19 | 19 | Glyceralde | GAPDH | [illegible] |
| Q8ILR7;A0 | Q8ILR7 | 31;1 | 31;1 | 31;1 | DNA helic | PF14_017 | [illegible] |
| Q8I1N9 | Q8I1N9 | 31 | 31 | 31 | | PFD0970c | [illegible] |
| C6KTD3 | C6KTD3 | 29 | 29 | 29 | | PFF1445c | [illegible] |
| O96153 | O96153 | 30 | 30 | 30 | | PFB0260w | [illegible] |
| Q8IIS4 | Q8IIS4 | 40 | 40 | 39 | | PF11_009 | [illegible] |
| Q8II57 | Q8II57 | 40 | 40 | 40 | | PF11_031 | [illegible] |
| Q8I5D0 | Q8I5D0 | 34 | 34 | 34 | | PFL1395c | [illegible] |
| Q8ILH5 | Q8ILH5 | 33 | 33 | 33 | | PF14_026 | [illegible] |
| Q8I5M6 | Q8I5M6 | 31 | 31 | 31 | | PFL0880c | [illegible] |
| C6KT82 | C6KT82 | 39 | 39 | 39 | | PFF1185w | [illegible] |
| Q8IJG6 | Q8IJG6 | 36 | 36 | 36 | | PF10_023 | [illegible] |
| C6KSR5 | C6KSR5 | 33 | 33 | 33 | | PFF0330w | [illegible] |
| C0H5I7 | C0H5I7 | 19 | 19 | 19 | T-comple | MAL13P1 | [illegible] |
| K7NTP5 | K7NTP5 | 23 | 18 | 18 | | | [illegible] |
| O77312 | O77312 | 28 | 28 | 28 | | PFC0135c | [illegible] |
| Q8IHP1 | Q8IHP1 | 35 | 35 | 35 | | PF11_048 | [illegible] |
| Q8I574 | Q8I574 | 32 | 32 | 32 | | PFL1680w | [illegible] |
| Q8I2G2 | Q8I2G2 | 30 | 30 | 30 | | CLAG9 | [illegible] |
| Q8ILG6 | Q8ILG6 | 26 | 26 | 26 | | PF14_027 | [illegible] |
| Q8I3W6 | Q8I3W6 | 28 | 28 | 28 | | PFE0705c | [illegible] |
| Q8IKF0 | Q8IKF0 | 25 | 25 | 25 | | H45 | [illegible] |
| Q8IIW0 | Q8IIW0 | 33 | 33 | 33 | | PF11_005 | [illegible] |
| Q8IHR4 | Q8IHR4 | 28 | 28 | 28 | | dyn1 | [illegible] |
| Q8IC16 | Q8IC16 | 26 | 25 | 25 | DNA helic | PF07_002 | [illegible] |
| Q76NM6 | Q76NM6 | 18 | 18 | 18 | V-type pr | vapA | [illegible] |
| C0H4L1 | C0H4L1 | 27 | 27 | 27 | | MAL7P1.2 | [illegible] |
| Q8IKP5 | Q8IKP5 | 25 | 25 | 25 | | PF14_055 | [illegible] |
| C6KST5 | C6KST5 | 23 | 23 | 23 | | PFF0430w | [illegible] |
| Q8I5V9 | Q8I5V9 | 35 | 35 | 35 | | PFL0445w | [illegible] |
| C6KTB4 | C6KTB4 | 26 | 26 | 26 | | PFF1350c | [illegible] |
| O77385 | O77385 | 36 | 36 | 36 | | Pfcrk-4 | [illegible] |
| Q8I447 | Q8I447 | 36 | 36 | 36 | | PFE0270c | [illegible] |
| Q8IEK1 | Q8IEK1 | 34 | 34 | 34 | | MAL13P1 | [illegible] |
| Q7K6A4 | Q7K6A4 | 16 | 16 | 16 | S-adenosy | PfSAMS | [illegible] |
| Q8I492 | Q8I492 | 25 | 25 | 25 | | MESA | [illegible] |
| Q8I4U5 | Q8I4U5 | 17 | 16 | 16 | | PFL2345c | [illegible] |
| Q8I3X5 | Q8I3X5 | 34 | 34 | 34 | | PFE0655w | [illegible] |
| O77313 | O77313 | 23 | 23 | 23 | | PFC0140c | [illegible] |
| O77310;O | O77310;O | 32;24;6 | 32;24;6 | 28;20;3 | | RhopH1/3 | [illegible] |
| Q8IE50 | Q8IE50 | 31 | 31 | 31 | | MAL13P1 | [illegible] |
| Q8I3K0 | Q8I3K0 | 33 | 33 | 33 | | PFE1320w | [illegible] |
| Q8I283 | Q8I283 | 28 | 28 | 28 | Cyclin-de | PFA_0285 | [illegible] |
| Q8IKI5 | Q8IKI5 | 26 | 26 | 26 | | PF14_062 | [illegible] |
| Q8IB67 | Q8IB67 | 27 | 27 | 27 | | gcn5 | [illegible] |
| Q8I3J5 | Q8I3J5 | 26 | 26 | 26 | DNA helic | PFE1345c | [illegible] |
| Q8IEK3 | Q8IEK3 | 21 | 21 | 20 | | PF13_006 | [illegible] |
| O96236 | O96236 | 26 | 26 | 26 | DNA-direc | PFB0715w | [illegible] |
| Q8IL88 | Q8IL88 | 22 | 22 | 22 | | PF14_035 | [illegible] |
| Q8IKH3 | Q8IKH3 | 22 | 22 | 22 | | PF14_063 | [illegible] |
| Q8IDH3 | Q8IDH3 | 38 | 38 | 38 | | PF13_027 | [illegible] |
| C0H4V6 | C0H4V6 | 14 | 14 | 14 | | MAL8P1.6 | [illegible] |
| O97239 | O97239 | 38 | 38 | 38 | Protein do | PFC0245c | [illegible] |
| A0A143ZU | A0A143ZU | 25 | 25 | 25 | | | [illegible] |
| O97313 | O97313 | 13 | 13 | 13 | 40S ribosomal protei | | [illegible] |
| C6KSN9 | C6KSN9 | 28 | 28 | 28 | | ApiAP2 | [illegible] |
| Q8II43 | Q8II43 | 20 | 20 | 20 | T-comple | PF11_033 | [illegible] |
| Q8I3Y8 | Q8I3Y8 | 26 | 26 | 26 | | PFE0585c | [illegible] |
| A0A143ZZ | A0A143ZZ | 39 | 39 | 39 | | | [illegible] |
| A0A5K1K9 | A0A5K1K9 | 42 | 42 | 42 | | | [illegible] |
| Q8I2I8 | Q8I2I8 | 25 | 25 | 25 | | PFI1590c | [illegible] |
| Q8IKJ0 | Q8IKJ0 | 20 | 20 | 20 | | PF14_061 | [illegible] |
| O96271 | O96271 | 23 | 23 | 23 | | PFB0895c | [illegible] |
| C0H4W3 | C0H4W3 | 26 | 26 | 26 | Probable | PF08_004 | [illegible] |
| Q7KQL1 | Q7KQL1 | 25 | 25 | 25 | | PF10_032 | [illegible] |
| Q8IBH8 | Q8IBH8 | 31 | 31 | 31 | | PF07_011 | [illegible] |
| O77314 | O77314 | 30 | 30 | 30 | | PFC0145c | [illegible] |
| Q8IL69 | Q8IL69 | 24 | 24 | 24 | | PF14_038 | [illegible] |
| A0A5K1K8 | A0A5K1K8 | 28 | 28 | 28 | | | [illegible] |
| Q8IHY0 | Q8IHY0 | 22 | 22 | 22 | | PF11_039 | [illegible] |
| Q8IBZ6 | Q8IBZ6 | 31 | 31 | 31 | | PF07_003 | [illegible] |
| Q8I0W8 | Q8I0W8 | 23 | 23 | 23 | | PFE0675c | [illegible] |
| Q8II72 | Q8II72 | 14 | 14 | 14 | | PF11_030 | [illegible] |
| Q6LFN2 | Q6LFN2 | 26 | 26 | 26 | RING fing | PFF0165c | [illegible] |
| Q8ILZ1 | Q8ILZ1 | 24 | 24 | 24 | | PF14_010 | [illegible] |
| A0A143ZV | A0A143ZV | 23;23 | 23;23 | 23;23 | Serine-re | SERA | [illegible] |
| C0H598 | C0H598 | 23 | 23 | 23 | | MAL13P1 | [illegible] |
| Q8IAM0 | Q8IAM0 | 21 | 21 | 21 | | PF08_013 | [illegible] |
| Q8IDF0 | Q8IDF0 | 29 | 29 | 28 | DNA helic | PF13_029 | [illegible] |
| Q8I4U7 | Q8I4U7 | 22 | 22 | 22 | | PFL2335w | [illegible] |
| Q9U0H1 | Q9U0H1 | 26 | 26 | 26 | DNA poly | PFD0590c | [illegible] |
| A0A143ZV | A0A143ZV | 17;17 | 17;17 | 17;17 | Tubulin b | PF10_008 | [illegible] |
| C6KSV9 | C6KSV9 | 20 | 20 | 20 | | PFF0560c | [illegible] |
| Q8I1Z2 | Q8I1Z2 | 24 | 24 | 24 | | PFD0180c | [illegible] |
| Q8I1V1 | Q8I1V1 | 17 | 17 | 17 | | PFD0665c | [illegible] |
| Q8I4T6 | Q8I4T6 | 24 | 24 | 24 | | PFL2390c | [illegible] |
| A0A5K1K9 | A0A5K1K9 | 29 | 29 | 29 | | | [illegible] |
| Q8IL02 | Q8IL02 | 11 | 11 | 11 | | PF14_044 | [illegible] |
| O97247 | O97247 | 23 | 23 | 23 | | PFC0285c | [illegible] |
| A0A144A0 | A0A144A0 | 20;20 | 20;20 | 20;20 | Proline--t | proRS | [illegible] |
| Q8I3Z1 | Q8I3Z1 | 36 | 36 | 36 | MATH and | PFE0570w | [illegible] |
| C0H5J0 | C0H5J0 | 28 | 28 | 28 | | MAL13P1 | [illegible] |
| C0H4P2 | C0H4P2 | 28 | 28 | 28 | | MAL7P1.1 | [illegible] |
| Q8ILC8 | Q8ILC8 | 27 | 27 | 27 | DNA topo | PF14_031 | [illegible] |
| Q8I5U3 | Q8I5U3 | 23 | 23 | 23 | | PFL0530c | [illegible] |
| Q8I5C4 | Q8I5C4 | 20 | 20 | 20 | T-comple | PFL1425w | [illegible] |
| Q8IEQ1 | Q8IEQ1 | 19 | 18 | 18 | | PF13_003 | [illegible] |
| Q8IJW0 | Q8IJW0 | 16 | 16 | 16 | | PF10_008 | [illegible] |
| A0A144A2 | A0A144A2 | 21 | 21 | 21 | | | [illegible] |
| C0H4M3 | C0H4M3 | 23 | 23 | 23 | | MAL7P1.2 | [illegible] |
| Q9U0J0 | Q9U0J0 | 21 | 21 | 21 | | PFD0470c | [illegible] |
| Q8II61 | Q8II61 | 10 | 10 | 10 | | PF11_031 | [illegible] |
| A0A5K1K9 | A0A5K1K9 | 24 | 24 | 24 | | | [illegible] |
| Q8I5H4 | Q8I5H4 | 20 | 20 | 20 | Polyaden | PFL1170w | [illegible] |
| Q8I3A1 | Q8I3A1 | 17 | 17 | 17 | | PFI0235w | [illegible] |
| Q8IKV6 | Q8IKV6 | 25 | 25 | 25 | | PfRON2 | [illegible] |
| Q8I517 | Q8I517 | 28 | 28 | 28 | | PFL1980c | [illegible] |
| Q8IM66 | Q8IM66 | 17 | 17 | 17 | | PF14_002 | [illegible] |
| Q8I3U1 | Q8I3U1 | 22 | 22 | 22 | Ubiquitin | PFE0835w | [illegible] |
| Q6ZLZ9 | Q6ZLZ9 | 15 | 15 | 4 | | PFI0180w | [illegible] |
| Q8IAW0 | Q8IAW0 | 15 | 15 | 15 | Importin | PF08_008 | [illegible] |
| O96220 | O96220 | 14 | 14 | 14 | | PFB0635w | [illegible] |
| Q8I240 | Q8I240 | 27 | 27 | 27 | | PFA_0510 | [illegible] |
| Q8ID85 | Q8ID85 | 17 | 17 | 17 | | PF13_033 | [illegible] |
| Q8IL94 | Q8IL94 | 13 | 13 | 13 | Ribonucle | PF14_035 | [illegible] |
| Q8IBU0 | Q8IBU0 | 18 | 18 | 18 | | PF07_006 | [illegible] |
| C6KTD8 | C6KTD8 | 28 | 28 | 28 | | PFF1470c | [illegible] |
| Q8IJA4 | Q8IJA4 | 23 | 23 | 22 | | PF10_029 | [illegible] |
| P27362 | P27362 | 14 | 14 | 14 | Phosphog | PGK | [illegible] |
| Q8ILE3 | Q8ILE3 | 10 | 10 | 10 | | PF14_030 | [illegible] |
| Q8ILL5;BF | Q8ILL5 | 20;1 | 20;1 | 20;1 | | PF14_022 | [illegible] |
| Q8IKY8 | Q8IKY8 | 30 | 30 | 30 | | PF14_046 | [illegible] |
| Q8IIR6 | Q8IIR6 | 17 | 17 | 17 | | PF11_009 | [illegible] |
| C0H4L6 | C0H4L6 | 18 | 18 | 18 | | MAL7P1.2 | [illegible] |
| A0A144A3 | A0A144A3 | 16;16 | 16;16 | 16;16 | Fructose-l | PF14_042 | [illegible] |

| | | | | | | | | | | | | | | | | | | | | | |
|---|---|---|---|---|---|---|---|---|---|---|---|---|---|---|---|---|---|---|---|---|---|
| Q8IBL5 | Q8IBL5 | 29 | 29 | 29 | | PF07_010[illegible] | tr\|Q8IBL5\|Q8IBL5_PLAF7 Uncharacterized protein OS=Plasmo[illegible] | 1 | 29 | 29 | 29 | 8 | 12 | 12 | 27 | 27 | 28 | 13 | 11 | [illegible] | [illegible] |
| Q8IER6 | Q8IER6 | 24 | 24 | 24 | | PF13_002[illegible] | tr\|Q8IER6\|Q8IER6_PLAF7 MCU1 domain-containing protein, pu[illegible] | 1 | 24 | 24 | 24 | 1 | 1 | 0 | 22 | 23 | 22 | 1 | 3 | [illegible] | [illegible] |
| Q8IBA0 | Q8IBA0 | 11 | 11 | 11 | | PfRACK | tr\|Q8IBA0\|Q8IBA0_PLAF7 Receptor for activated c kinase OS=[illegible] | 1 | 11 | 11 | 11 | 5 | 6 | 5 | 11 | 11 | 11 | 7 | 8 | [illegible] | [illegible] |
| Q8ILQ6 | Q8ILQ6 | 33 | 33 | 33 | | PF14_018[illegible] | tr\|Q8ILQ6\|Q8ILQ6_PLAF7 Uncharacterized protein OS=Plasmo[illegible] | 1 | 33 | 33 | 33 | 15 | 17 | 14 | 30 | 30 | 32 | 17 | 15 | [illegible] | [illegible] |
| O96205 | O96205 | 33 | 33 | 33 | | PFB0560w | tr\|O96205\|O96205_PLAF7 Uncharacterized protein OS=Plasmo[illegible] | 1 | 33 | 33 | 33 | 4 | 7 | 5 | 32 | 31 | 29 | 8 | 7 | [illegible] | [illegible] |
| Q8I1R6 | Q8I1R6 | 17 | 17 | 17 | Bifunctior | DHFR-TS | tr\|Q8I1R6\|Q8I1R6_PLAF7 Bifunctional dihydrofolate reductas[illegible] | 1 | 17 | 17 | 17 | 3 | 3 | 2 | 15 | 17 | 15 | 6 | 6 | [illegible] | [illegible] |
| Q8II60 | Q8II60 | 18 | 18 | 18 | | PF11_031[illegible] | tr\|Q8II60\|Q8II60_PLAF7 26S protease regulatory subunit 6A, p[illegible] | 1 | 18 | 18 | 18 | 9 | 9 | 8 | 17 | 16 | 17 | 12 | 9 | [illegible] | [illegible] |
| Q8IC35 | Q8IC35 | 22 | 22 | 22 | | MAL7P1.1[illegible] | tr\|Q8IC35\|Q8IC35_PLAF7 Erythrocyte membrane-associated a[illegible] | 1 | 22 | 22 | 22 | 10 | 15 | 14 | 19 | 19 | 16 | 16 | 15 | [illegible] | [illegible] |
| Q8IJI4 | Q8IJI4 | 23 | 23 | 23 | | PF10_021[illegible] | tr\|Q8IJI4\|Q8IJI4_PLAF7 PHAX domain-containing protein, puta[illegible] | 1 | 23 | 23 | 23 | 11 | 11 | 9 | 21 | 21 | 23 | 14 | 11 | [illegible] | [illegible] |
| Q8IL56 | Q8IL56 | 17 | 17 | 17 | | PF14_039[illegible] | tr\|Q8IL56\|Q8IL56_PLAF7 FACT complex subunit SSRP1 OS=Pla[illegible] | 1 | 17 | 17 | 17 | 13 | 16 | 14 | 17 | 17 | 17 | 15 | 14 | [illegible] | [illegible] |
| O97282 | O97282 | 20 | 20 | 20 | | PFC0900w | tr\|O97282\|O97282_PLAF7 T-complex protein 1 subunit epsilo[illegible] | 1 | 20 | 20 | 20 | 13 | 11 | 13 | 16 | 16 | 15 | 15 | 15 | [illegible] | [illegible] |
| Q8IJP9 | Q8IJP9 | 20 | 20 | 20 | | PF10_014[illegible] | tr\|Q8IJP9\|Q8IJP9_PLAF7 Transcriptional coactivator ADA2 OS=[illegible] | 1 | 20 | 20 | 20 | 7 | 4 | 7 | 19 | 20 | 18 | 10 | 8 | [illegible] | [illegible] |
| C6KTB1 | C6KTB1 | 9 | 9 | 9 | | PFF1335c | tr\|C6KTB1\|C6KTB1_PLAF7 Protein DJ-1 OS=Plasmodium falcip[illegible] | 1 | 9 | 9 | 9 | 8 | 7 | 7 | 8 | 9 | 7 | 7 | 7 | [illegible] | [illegible] |
| Q8IAX9 | Q8IAX9 | 20 | 20 | 20 | | MAL8P1.9[illegible] | tr\|Q8IAX9\|Q8IAX9_PLAF7 AAA family ATPase, putative OS=Pl[illegible] | 1 | 20 | 20 | 20 | 1 | 2 | 1 | 18 | 18 | 18 | 1 | 2 | [illegible] | [illegible] |
| Q8I548 | Q8I548 | 23 | 23 | 23 | | PFL1815c | tr\|Q8I548\|Q8I548_PLAF7 Uncharacterized protein OS=Plasmo[illegible] | 1 | 23 | 23 | 23 | 8 | 8 | 9 | 23 | 23 | 22 | 10 | 9 | [illegible] | [illegible] |
| Q8I5G5 | Q8I5G5 | 19 | 19 | 19 | | PFL1215c | tr\|Q8I5G5\|Q8I5G5_PLAF7 Uncharacterized protein OS=Plasmo[illegible] | 1 | 19 | 19 | 19 | 7 | 8 | 9 | 19 | 19 | 19 | 11 | 10 | [illegible] | [illegible] |
| C6KT76 | C6KT76 | 12 | 12 | 12 | Hexokina[illegible] | PFF1155w | tr\|C6KT76\|C6KT76_PLAF7 Phosphotransferase OS=Plasmodiu[illegible] | 1 | 12 | 12 | 12 | 9 | 11 | 11 | 10 | 12 | 12 | 11 | 12 | [illegible] | [illegible] |
| Q8IIU3 | Q8IIU3 | 16 | 16 | 16 | | PF11_007[illegible] | tr\|Q8IIU3\|Q8IIU3_PLAF7 RuvB-like helicase OS=Plasmodium f[illegible] | 1 | 16 | 16 | 16 | 2 | 4 | 2 | 16 | 16 | 16 | 5 | 5 | [illegible] | [illegible] |
| Q8I538 | Q8I538 | 19 | 19 | 19 | | PFL1865w | tr\|Q8I538\|Q8I538_PLAF7 Uncharacterized protein OS=Plasmo[illegible] | 1 | 19 | 19 | 19 | 3 | 4 | 4 | 18 | 18 | 19 | 3 | 5 | [illegible] | [illegible] |
| Q8IDQ9 | Q8IDQ9 | 15 | 15 | 15 | | PfPMT | tr\|Q8IDQ9\|Q8IDQ9_PLAF7 Phosphoethanolamine N-methyltr[illegible] | 1 | 15 | 15 | 15 | 9 | 11 | 10 | 15 | 15 | 15 | 14 | 12 | [illegible] | [illegible] |
| [illegible] | [illegible] | [illegible] | [illegible] | [illegible] | [illegible] | [illegible] | [illegible] | [illegible] | [illegible] | [illegible] | [illegible] | [illegible] | [illegible] | [illegible] | [illegible] | [illegible] | [illegible] | [illegible] | [illegible] | [illegible] | [illegible] |

| | | | | | | | | | | | | | |
|---|---|---|---|---|---|---|---|---|---|---|---|---|---|
| A0A5K1K9 | A0A5K1K9 | 11 | 11 | 11 | | | tr\|A0A5K1K976\|A0A5K1K976_PLAF7 Uncharacterized protein ( | 1 | 11 | 11 | 11 | 2 | [illegible] |
| A0A1437V | A0A1437V | 15;15 | 15;15 | 15;15 | Uncharact | PFB0460c | tr\|A0A1437WL6\|A0A1437WL6_PLAF7 Uncharacterized protein | 2 | 15 | 15 | 15 | 0 | [illegible] |
| Q8IKL1 | Q8IKL1 | 15 | 15 | 15 | | PF14_059: | tr\|Q8IKL1\|Q8IKL1_PLAF7 GTP-binding protein, putative OS=Pl | 1 | 15 | 15 | 15 | 6 | [illegible] |
| Q8IKV8 | Q8IKV8 | 12 | 12 | 12 | | PF14_049: | tr\|Q8IKV8\|Q8IKV8_PLAF7 Sortilin OS=Plasmodium falciparum | 1 | 12 | 12 | 12 | 6 | [illegible] |
| C6KTC7 | C6KTC7 | 11 | 11 | 11 | | PFF1415c | tr\|C6KTC7\|C6KTC7_PLAF7 DnaJ protein, putative OS=Plasmod | 1 | 11 | 11 | 11 | 7 | [illegible] |
| Q8I5X9 | Q8I5X9 | 14 | 14 | 14 | DNA-direc | PFL0330c | tr\|Q8I5X9\|Q8I5X9_PLAF7 DNA-directed RNA polymerase subu | 1 | 14 | 14 | 14 | 1 | [illegible] |
| Q8IM38 | Q8IM38 | 11 | 11 | 11 | | PF14_005: | tr\|Q8IM38\|Q8IM38_PLAF7 Ribonucleoside-diphosphate redu | 1 | 11 | 11 | 11 | 1 | [illegible] |
| Q8ID37 | Q8ID37 | 9 | 9 | 9 | | MAL13P1. | tr\|Q8ID37\|Q8ID37_PLAF7 U1 small nuclear ribonucleoprotein | 1 | 9 | 9 | 9 | 1 | [illegible] |
| Q8IKG9 | Q8IKG9 | 13 | 13 | 13 | | PF14_063( | tr\|Q8IKG9\|Q8IKG9_PLAF7 NOC3p domain-containing protein | 1 | 13 | 13 | 13 | 3 | [illegible] |
| A0A144A1 | A0A144A1 | 17;17 | 17;17 | 17;17 | Protein Pl | PF14_017! | tr\|A0A144A1P7\|A0A144A1P7_PLAF7 HSA domain-containing r | 2 | 17 | 17 | 17 | 0 | [illegible] |
| Q8I413 | Q8I413 | 18 | 18 | 18 | | PFE0450w | tr\|Q8I413\|Q8I413_PLAF7 Structural maintenance of chromoso | 1 | 18 | 18 | 18 | 1 | [illegible] |
| A0A1437V | A0A1437V | 11;11 | 11;11 | 11;11 | | PFE1305c | tr\|A0A1437VA6\|A0A1437VA6_PLAF7 ADP-ribosylation factor ( | 2 | 11 | 11 | 11 | 1 | [illegible] |
| Q7KQK3 | Q7KQK3 | 7 | 7 | 7 | | PFL0565w | tr\|Q7KQK3\|Q7KQK3_PLAF7 Heat shock protein DNAJ homolog | 1 | 7 | 7 | 7 | 5 | [illegible] |
| Q8IET7 | Q8IET7 | 8 | 8 | 8 | | PF13_001 | tr\|Q8IET7\|Q8IET7_PLAF7 40S ribosomal protein S7 OS=Plasmo | 1 | 8 | 8 | 8 | 5 | [illegible] |
| Q8IL13 | Q8IL13 | 10 | 10 | 10 | | PF14_043: | tr\|Q8IL13\|Q8IL13_PLAF7 ATP-dependent RNA helicase DDX5 | 1 | 10 | 10 | 10 | 2 | [illegible] |
| Q8I6U4 | Q8I6U4 | 6 | 6 | 1 | | PF11_016! | tr\|Q8I6U4\|Q8I6U4_PLAF7 Cysteine proteinase falcipain 2a OS: | 1 | 6 | 6 | 1 | 5 | [illegible] |
| C0H4M6 | C0H4M6 | 8 | 8 | 8 | | MAL7P1.3 | tr\|C0H4M6\|C0H4M6_PLAF7 Regulator of chromosome conden | 1 | 8 | 8 | 8 | 2 | [illegible] |
| Q8I3P6 | Q8I3P6 | 12 | 12 | 12 | | PFE1085w | tr\|Q8I3P6\|Q8I3P6_PLAF7 ATP-dependent RNA helicase DDX1 | 1 | 12 | 12 | 12 | 4 | [illegible] |
| Q8IAR1 | Q8IAR1 | 15 | 15 | 15 | | PF08_011 | tr\|Q8IAR1\|Q8IAR1_PLAF7 DNA helicase PSH3 OS=Plasmodium | 1 | 15 | 15 | 15 | 0 | [illegible] |
| Q8IJZ5 | Q8IJZ5 | 21 | 21 | 21 | | PF10_004! | tr\|Q8IJZ5\|Q8IJZ5_PLAF7 WD repeat-containing protein, putati | 1 | 21 | 21 | 21 | 2 | [illegible] |
| Q8IK82 | Q8IK82 | 15 | 15 | 15 | | PF14_072 | tr\|Q8IK82\|Q8IK82_PLAF7 Bromodomain protein, putative OS= | 1 | 15 | 15 | 15 | 3 | [illegible] |
| O96210 | O96210 | 8 | 8 | 8 | | PFB0585w | tr\|O96210\|O96210_PLAF7 Arginyl-tRNA--protein transferase ( | 1 | 8 | 8 | 8 | 5 | [illegible] |
| Q8IDD0 | Q8IDD0 | 12 | 12 | 12 | | PF13_030( | tr\|Q8IDD0\|Q8IDD0_PLAF7 DNA helicase, putative OS=Plasmo | 1 | 12 | 12 | 12 | 3 | [illegible] |

[illegible]

| | | | | | | | | | | | | | | | | | | | | |
|---|---|---|---|---|---|---|---|---|---|---|---|---|---|---|---|---|---|---|---|---|
| O96152;A0 | O96152;A0 | 6;6 | 6;6 | 6;6 | | PFB0255w | tr\|O96152\|O96152_PLAF7 RNA-binding protein, putative OS=P | 2 | 6 | 6 | 6 | 0 | 1 | 1 | 5 | 5 | 5 | 1 | 1 | [illegible] |
| Q8IKI6 | Q8IKI6 | 8 | 8 | 8 | | PF14_061 | tr\|Q8IKI6\|Q8IKI6_PLAF7 Mediator of RNA polymerase II trans | 1 | 8 | 8 | 8 | 0 | 0 | 0 | 8 | 8 | 8 | 1 | 0 | [illegible] |
| A0A144A0 | A0A144A0 | 7;7 | 7;7 | 7;7 | Pre-mRN | PFL2310w | tr\|A0A144A0M5\|A0A144A0M5_PLAF7 Pre-mRNA-splicing fact | 2 | 7 | 7 | 7 | 0 | 1 | 1 | 5 | 6 | 6 | 1 | 0 | [illegible] |
| Q8I533 | Q8I533 | 9 | 9 | 9 | | PFL1890c | tr\|Q8I533\|Q8I533_PLAF7 HD superfamily phosphohydrolase | 1 | 9 | 9 | 9 | 1 | 1 | 1 | 9 | 9 | 9 | 2 | 3 | [illegible] |
| Q8I0V9 | Q8I0V9 | 6 | 6 | 6 | | PFA_0450 | tr\|Q8I0V9\|Q8I0V9_PLAF7 Cleavage and polyadenylation speci | 1 | 6 | 6 | 6 | 0 | 0 | 0 | 6 | 6 | 6 | 0 | 2 | [illegible] |
| O96240 | O96240 | 5 | 5 | 5 | | PFB0735c | tr\|O96240\|O96240_PLAF7 Alba domain-containing protein OS | 1 | 5 | 5 | 5 | 0 | 0 | 0 | 5 | 5 | 4 | 0 | 0 | [illegible] |
| Q8IKZ8 | Q8IKZ8 | 5 | 5 | 5 | | PF14_045 | tr\|Q8IKZ8\|Q8IKZ8_PLAF7 Uncharacterized protein OS=Plasmo | 1 | 5 | 5 | 5 | 5 | 5 | 5 | 3 | 4 | 4 | 5 | 5 | [illegible] |
| Q8I608 | Q8I608 | 7 | 7 | 7 | | PFL0185c | tr\|Q8I608\|Q8I608_PLAF7 Nucleosome assembly protein OS=P | 1 | 7 | 7 | 7 | 1 | 3 | 1 | 7 | 7 | 7 | 4 | 4 | [illegible] |
| C0H4C9 | C0H4C9 | 12 | 12 | 12 | | PFE0320w | tr\|C0H4C9\|C0H4C9_PLAF7 Uncharacterized protein OS=Plasm | 1 | 12 | 12 | 12 | 0 | 0 | 0 | 12 | 11 | 11 | 0 | 0 | [illegible] |
| Q8IJU4 | Q8IJU4 | 7 | 7 | 7 | | PF10_009 | tr\|Q8IJU4\|Q8IJU4_PLAF7 PI31 domain-containing protein, put | 1 | 7 | 7 | 7 | 0 | 2 | 2 | 7 | 7 | 7 | 3 | 3 | [illegible] |
| Q8I546 | Q8I546 | 4 | 4 | 4 | | PFL1825w | tr\|Q8I546\|Q8I546_PLAF7 DUF4149 domain-containing protein | 1 | 4 | 4 | 4 | 4 | 4 | 3 | 4 | 4 | 4 | 4 | 4 | [illegible] |
| Q8IEM3 | Q8IEM3 | 4 | 4 | 4 | | PF13_004 | tr\|Q8IEM3\|Q8IEM3_PLAF7 60S ribosomal protein L24, putative | 1 | 4 | 4 | 4 | 3 | 4 | 4 | 4 | 3 | 4 | 3 | 4 | [illegible] |
| Q8IHU4 | Q8IHU4 | 8 | 8 | 8 | | PF11_043 | tr\|Q8IHU4\|Q8IHU4_PLAF7 J domain-containing protein OS=Pl | 1 | 8 | 8 | 8 | 0 | 1 | 0 | 8 | 8 | 8 | 0 | 1 | [illegible] |
| Q8IJN3-O | Q8IJN3 | 8;1 | 8;1 | 8;1 | | FIKK-10.1 | tr\|Q8IJN3\|Q8IJN3_PLAF7 Serine/threonine protein kinase, FII | 2 | 8 | 8 | 8 | 2 | 3 | 3 | 8 | 7 | 6 | 6 | 3 | [illegible] |
| Q8I488 | Q8I488 | 6 | 6 | 6 | | PFE0060w | tr\|Q8I488\|Q8I488_PLAF7 Parasite-infected erythrocyte surfac | 1 | 6 | 6 | 6 | 5 | 6 | 5 | 6 | 6 | 6 | 6 | 5 | [illegible] |
| Q8IIV8 | Q8IIV8 | 8 | 8 | 8 | | PF11_005 | tr\|Q8IIV8\|Q8IIV8_PLAF7 Thioredoxin-like mero protein OS=Pl | 1 | 8 | 8 | 8 | 4 | 5 | 6 | 8 | 8 | 8 | 7 | 7 | [illegible] |
| C6KSP3 | C6KSP3 | 10 | 10 | 10 | | PFF0220w | tr\|C6KSP3\|C6KSP3_PLAF7 GYF domain-containing protein OS= | 1 | 10 | 10 | 10 | 7 | 7 | 7 | 9 | 9 | 10 | 7 | 5 | [illegible] |
| A0A143ZY | A0A143ZY | 10 | 10 | 10 | | | tr\|A0A143ZY29\|A0A143ZY29_PLAF7 Regulator of nonsense tra | 1 | 10 | 10 | 10 | 0 | 0 | 1 | 10 | 7 | 10 | 1 | 2 | [illegible] |
| Q8ID62 | Q8ID62 | 9 | 9 | 9 | | PF13_034 | tr\|Q8ID62\|Q8ID62_PLAF7 Exosome complex component RRP4 | 1 | 9 | 9 | 9 | 0 | 0 | 0 | 8 | 8 | 7 | 0 | 0 | [illegible] |
| Q8IIG8 | Q8IIG8 | 14 | 14 | 14 | | PF11_020 | tr\|Q8IIG8\|Q8IIG8_PLAF7 Protein transport protein SEC16, put | 1 | 14 | 14 | 14 | 4 | 7 | 9 | 13 | 14 | 12 | 7 | 7 | [illegible] |

[illegible]

| | | | | | | | | | | | | |
|---|---|---|---|---|---|---|---|---|---|---|---|---|
| Q8IL96 | Q8IL96 | 4 | 4 | 4 | | PF14_035[illegible] | tr\|Q8IL96\|Q8IL96_PLAF7 N-acetyltransferase, GNAT family, pu[illegible] | 1 | 4 | 4 | 4 | 4 |
| C6KTB5 | C6KTB5 | 4 | 4 | 4 | | PFF1355w | tr\|C6KTB5\|C6KTB5_PLAF7 Ribonuclease P protein subunit p29 | 1 | 4 | 4 | 4 | 0 |
| Q8IIP2 | Q8IIP2 | 5 | 5 | 5 | | PF11_012[illegible] | tr\|Q8IIP2\|Q8IIP2_PLAF7 Protein kinase domain-containing pro[illegible] | 1 | 5 | 5 | 5 | 0 |
| Q8I1T8 | Q8I1T8 | 4 | 4 | 4 | ATPase AS[illegible] | PFD0725c | sp\|Q8I1T8\|ASNA_PLAF7 ATPase ASNA1 homolog OS=Plasmod[illegible] | 1 | 4 | 4 | 4 | 1 |
| O96154 | O96154 | 7 | 7 | 7 | | PFB0265c | tr\|O96154\|O96154_PLAF7 DNA repair protein RAD2, putative [illegible] | 1 | 7 | 7 | 7 | 0 |
| Q8IJ60 | Q8IJ60 | 7 | 7 | 7 | | PF10_034[illegible] | tr\|Q8IJ60\|Q8IJ60_PLAF7 Methionine--tRNA ligase OS=Plasmo[illegible] | 1 | 7 | 7 | 7 | 4 |
| C6KT23 | C6KT23 | 5 | 5 | 5 | | PFF0885w | tr\|C6KT23\|C6KT23_PLAF7 60S ribosomal protein L27a, putative | 1 | 5 | 5 | 5 | 3 |
| Q8IEP4 | Q8IEP4 | 7 | 7 | 7 | | PF13_003[illegible] | tr\|Q8IEP4\|Q8IEP4_PLAF7 U3 small nucleolar RNA-associated r[illegible] | 1 | 7 | 7 | 7 | 1 |
| Q8IKF5 | Q8IKF5 | 4 | 4 | 4 | | PF14_065[illegible] | tr\|Q8IKF5\|Q8IKF5_PLAF7 Telomere length and silencing prote[illegible] | 1 | 4 | 4 | 4 | 0 |
| C0H5C1 | C0H5C1 | 9 | 9 | 9 | | MAL13P1.[illegible] | tr\|C0H5C1\|C0H5C1_PLAF7 Uncharacterized protein OS=Plasm[illegible] | 1 | 9 | 9 | 9 | 0 |
| Q8ILD0 | Q8ILD0 | 5 | 5 | 5 | | PF14_031[illegible] | tr\|Q8ILD0\|Q8ILD0_PLAF7 Chromatin assembly factor 1 P55 sub[illegible] | 1 | 5 | 5 | 5 | 3 |
| Q8II17 | Q8II17 | 8 | 8 | 8 | DNA-dire[illegible] | PF11_035[illegible] | tr\|Q8II17\|Q8II17_PLAF7 DNA-directed RNA polymerase subun[illegible] | 1 | 8 | 8 | 8 | 1 |
| Q8I446 | Q8I446 | 5 | 5 | 5 | | PFE0275w | tr\|Q8I446\|Q8I446_PLAF7 BCNT-C domain-containing protein [illegible] | 1 | 5 | 5 | 5 | 0 |
| Q8IAW7 | Q8IAW7 | 6 | 6 | 6 | | MAL8P1.9[illegible] | tr\|Q8IAW7\|Q8IAW7_PLAF7 CS domain protein, putative OS=P[illegible] | 1 | 6 | 6 | 6 | 2 |
| Q8I3N7 | Q8I3N7 | 5 | 5 | 5 | | PFE1130w | tr\|Q8I3N7\|Q8I3N7_PLAF7 Inner membrane complex protein [illegible] | 1 | 5 | 5 | 5 | 4 |
| Q8IJ89 | Q8IJ89 | 8 | 8 | 8 | | PF10_031[illegible] | tr\|Q8IJ89\|Q8IJ89_PLAF7 Uncharacterized protein OS=Plasmod[illegible] | 1 | 8 | 8 | 8 | 4 |
| C0H4V8 | C0H4V8 | 8 | 8 | 8 | | MAL8P1.6[illegible] | tr\|C0H4V8\|C0H4V8_PLAF7 DNA helicase, putative OS=Plasmo[illegible] | 1 | 8 | 8 | 8 | 2 |
| Q8IHS2 | Q8IHS2 | 5 | 5 | 5 | | Aos1 | tr\|Q8IHS2\|Q8IHS2_PLAF7 SUMO-activating enzyme subunit 1 [illegible] | 1 | 5 | 5 | 5 | 1 |
| O96176 | O96176 | 7 | 7 | 7 | | PFB0410c | tr\|O96176\|O96176_PLAF7 Patatin-like phospholipase, putativ[illegible] | 1 | 7 | 7 | 7 | 0 |
| Q8IJZ3 | Q8IJZ3 | 6 | 6 | 6 | | PF10_004[illegible] | tr\|Q8IJZ3\|Q8IJZ3_PLAF7 RNA-binding protein, putative OS=Pl[illegible] | 1 | 6 | 6 | 6 | 0 |

| | | | | | | | | | | | | | | | | | | | |
|---|---|---|---|---|---|---|---|---|---|---|---|---|---|---|---|---|---|---|---|
| Q8IKP1 | Q8IKP1 | 4 | 4 | 4 | | PF14_056[illegible] | tr\|Q8IKP1\|Q8IKP1_PLAF7 ATP-dependent RNA helicase DBP5 | 1 | 4 | 4 | 4 | 1 | 2 | 2 | 3 | 4 | 2 | 2 | 1 |
| A0A144A0[illegible] | A0A144A0[illegible] | 5 | 5 | 5 | | | tr\|A0A144A0I9\|A0A144A0I9_PLAF7 Uncharacterized protein C[illegible] | 1 | 5 | 5 | 5 | 1 | 0 | 0 | 5 | 5 | 5 | 1 | 1 |
| Q8IIW5 | Q8IIW5 | 4 | 4 | 4 | Casein kir[illegible] | PF11_004[illegible] | tr\|Q8IIW5\|Q8IIW5_PLAF7 Casein kinase II subunit beta OS=Pl[illegible] | 1 | 4 | 4 | 4 | 1 | 1 | 1 | 3 | 4 | 3 | 1 | 1 |
| O77389 | O77389 | 5 | 5 | 5 | | PFC0725c | tr\|O77389\|O77389_PLAF7 Formate-nitrite transporter OS=Pla[illegible] | 1 | 5 | 5 | 5 | 3 | 3 | 2 | 3 | 3 | 3 | 2 | 3 |
| O96149 | O96149 | 3 | 3 | 3 | | PFB0240w | tr\|O96149\|O96149_PLAF7 PCI domain-containing protein, put[illegible] | 1 | 3 | 3 | 3 | 2 | 2 | 2 | 3 | 3 | 3 | 2 | 3 |
| Q8I3F4 | Q8I3F4 | 4 | 4 | 4 | | PFE1570c | tr\|Q8I3F4\|Q8I3F4_PLAF7 G-patch domain-containing protein [illegible] | 1 | 4 | 4 | 4 | 0 | 0 | 0 | 4 | 4 | 3 | 0 | 0 |
| Q8ILV5 | Q8ILV5 | 2 | 2 | 2 | | PF14_013[illegible] | tr\|Q8ILV5\|Q8ILV5_PLAF7 26S proteasome regulatory subunit [illegible] | 1 | 2 | 2 | 2 | 2 | 2 | 1 | 2 | 2 | 2 | 2 | 1 |
| Q8IEL1 | Q8IEL1 | 4 | 4 | 4 | | PF13_005[illegible] | tr\|Q8IEL1\|Q8IEL1_PLAF7 Uncharacterized protein OS=Plasmo[illegible] | 1 | 4 | 4 | 4 | 1 | 0 | 0 | 3 | 4 | 3 | 1 | 1 |
| C6KSS5 | C6KSS5 | 6 | 6 | 6 | Protein Pl[illegible] | PFF0380w | sp\|C6KSS5\|LRR2_PLAF7 Protein PFF0380w OS=Plasmodium fa[illegible] | 1 | 6 | 6 | 6 | 0 | 0 | 0 | 6 | 4 | 5 | 0 | 0 |
| Q8IBH7 | Q8IBH7 | 4 | 4 | 4 | | PF07_011[illegible] | tr\|Q8IBH7\|Q8IBH7_PLAF7 Eukaryotic translation initiation fac[illegible] | 1 | 4 | 4 | 4 | 1 | 1 | 2 | 3 | 3 | 3 | 2 | 2 |
| Q8I3I0 | Q8I3I0 | 2 | 2 | 2 | | PFE1430c | tr\|Q8I3I0\|Q8I3I0_PLAF7 Peptidyl-prolyl cis-trans isomerase [illegible] | 1 | 2 | 2 | 2 | 0 | 0 | 0 | 2 | 2 | 2 | 1 | 1 |
| Q8IK88 | Q8IK88 | 2 | 2 | 2 | | PF14_071[illegible] | tr\|Q8IK88\|Q8IK88_PLAF7 Mediator of RNA polymerase II trans[illegible] | 1 | 2 | 2 | 2 | 1 | 1 | 1 | 2 | 2 | 2 | 0 | 0 |
| A0A144A1[illegible] | A0A144A1[illegible] | 4;4 | 4;4 | 4;4 | Glutathio[illegible] | GST | tr\|A0A144A1J6\|A0A144A1J6_PLAF7 Glutathione S-transferase | 2 | 4 | 4 | 4 | 1 | 1 | 1 | 3 | 3 | 4 | 1 | 2 |
| Q8IIW2 | Q8IIW2 | 4 | 4 | 4 | | PF11_005[illegible] | tr\|Q8IIW2\|Q8IIW2_PLAF7 Phenylalanine--tRNA ligase beta su[illegible] | 1 | 4 | 4 | 4 | 1 | 1 | 1 | 4 | 3 | 4 | 3 | 2 |
| Q8IL48 | Q8IL48 | 4 | 4 | 4 | | PF14_040[illegible] | tr\|Q8IL48\|Q8IL48_PLAF7 tRNA import protein tRIP OS=Plasmo[illegible] | 1 | 4 | 4 | 4 | 4 | 4 | 4 | 4 | 3 | 4 | 4 | 4 |
| B9ZSH7;B[illegible] | B9ZSH7;B[illegible] | 4;2 | 4;2 | 0;0 | | PfMC-2TM[illegible] | tr\|B9ZSH7\|B9ZSH7_PLAF7 Pfmc-2TM Maurers cleft two transm[illegible] | 2 | 4 | 4 | 0 | 1 | 1 | 1 | 4 | 4 | 4 | 1 | 1 |
| Q8IEQ6 | Q8IEQ6 | 5 | 5 | 5 | | PF13_003[illegible] | tr\|Q8IEQ6\|Q8IEQ6_PLAF7 Uncharacterized protein OS=Plasmo[illegible] | 1 | 5 | 5 | 5 | 0 | 0 | 0 | 4 | 5 | 5 | 0 | 2 |
| C6S3I6 | C6S3I6 | 3 | 3 | 3 | | PF14_078[illegible] | tr\|C6S3I6\|C6S3I6_PLAF7 Replication factor A protein 3, putativ[illegible] | 1 | 3 | 3 | 3 | 0 | 1 | 1 | 3 | 3 | 3 | 1 | 1 |
| A0A5K1K9[illegible] | A0A5K1K9[illegible] | 3 | 3 | 3 | | | tr\|A0A5K1K903\|A0A5K1K903_PLAF7 U3 small nucleolar RNA-a[illegible] | 1 | 3 | 3 | 3 | 1 | 0 | 1 | 3 | 3 | 3 | 0 | 0 |
| Q8IBC0 | Q8IBC0 | 2 | 2 | 2 | | PF08_000[illegible] | tr\|Q8IBC0\|Q8IBC0_PLAF7 Uncharacterized protein OS=Plasmo[illegible] | 1 | 2 | 2 | 2 | 0 | 0 | 0 | 2 | 2 | 2 | 0 | 0 |

[illegible]

| | | | | | | | |
|---|---|---|---|---|---|---|---|
| Q8IJV6 | Q8IJV6 | 4 | 4 | 4 | | PF10_008 | [illegible] |
| Q8I4Y9 | Q8I4Y9 | 4 | 4 | 4 | | PFL2120w | [illegible] |
| A0A144A1 | A0A144A1 | 3;3 | 3;3 | 3;3 | | PFL0105w | [illegible] |
| Q8I577 | Q8I577 | 3 | 3 | 3 | | PFL1665c | [illegible] |
| C6KT35 | C6KT35 | 2 | 2 | 2 | | PfACS12 | [illegible] |
| Q8I3G2 | Q8I3G2 | 2 | 2 | 2 | | PFE1530c | [illegible] |
| A0A5K1K9 | A0A5K1K9 | 2 | 2 | 2 | | | [illegible] |
| Q8IKG2 | Q8IKG2 | 2 | 2 | 2 | | PF14_064 | [illegible] |
| Q8I450 | Q8I450 | 3 | 3 | 3 | | PFE0255w | [illegible] |
| Q8IJI7 | Q8IJI7 | 2 | 2 | 2 | | PF10_0210 | [illegible] |
| Q8ID74 | Q8ID74 | 2 | 2 | 2 | | PF13_0334 | [illegible] |
| Q8IEF7 | Q8IEF7 | 2 | 2 | 2 | | PF13_008 | [illegible] |
| Q8IKZ4 | Q8IKZ4 | 3 | 3 | 3 | | PF14_045 | [illegible] |
| Q8IC05 | Q8IC05 | 30 | 30 | 30 | | PF07_002 | [illegible] |
| Q8I0U8;O | Q8I0U8 | 28;1 | 28;1 | 28;1 | | MSP1 | [illegible] |
| Q8IJ52 | Q8IJ52 | 23 | 23 | 23 | | PF10_034 | [illegible] |
| A0A14377 | A0A14377 | 15;15 | 15;15 | 15;15 | Enolase | ENO | [illegible] |
| A0A144A3 | A0A144A3 | 11;11 | 9;9 | 9;9 | Actin-2 | PF14_012 | [illegible] |
| Q8IIV2 | Q8IIV2 | 5 | 5 | 5 | Histone H | PF11_006 | [illegible] |
| Q8IM10 | Q8IM10 | 5 | 5 | 5 | 40S ribos | PF14_008 | [illegible] |
| Q8IDN6 | Q8IDN6 | 5 | 5 | 5 | | Sec61 | [illegible] |
| Q8I0V2 | Q8I0V2 | 6 | 6 | 6 | ATP synth | PFL1725w | [illegible] |
| Q8I485 | Q8I485 | 11 | 11 | 11 | | RAP3 | [illegible] |
| O97320;C | O97320 | 4;1 | 4;1 | 4;1 | Histone H | PFC0920w | [illegible] |
| Q8IJ45 | Q8IJ45 | 11 | 11 | 11 | | PF10_035 | [illegible] |
| A0A1437Y | A0A1437Y | 9;9 | 9;9 | 9;9 | ADP-ribos | ARF1 | [illegible] |
| Q8IDJ8 | Q8IDJ8 | 9 | 9 | 9 | Lysine--tR | PF13_026 | [illegible] |
| Q8IEQ3 | Q8IEQ3 | 7 | 7 | 7 | | PF13_003 | [illegible] |
| Q9NFA0 | Q9NFA0 | 3 | 3 | 3 | | PFC0912w | [illegible] |
| A0A144A1 | A0A144A1 | 5;5 | 5;5 | 5;5 | Calmodul | PF14_032 | [illegible] |
| Q8I2W2 | Q8I2W2 | 4 | 4 | 4 | | PFI0935w | [illegible] |
| Q8IAX6 | Q8IAX6 | 4 | 4 | 4 | 60S ribos | PF08_007 | [illegible] |
| Q8I207 | Q8I207 | 3 | 3 | 3 | | PFD0080c | [illegible] |
| Q8I2Q0 | Q8I2Q0 | 3 | 3 | 3 | | PFI1270w | [illegible] |
| Q8ILK3 | Q8ILK3 | 3 | 3 | 3 | | PF14_024 | [illegible] |
| C6KSV0;C | C6KSV0;C | 3;2 | 3;2 | 3;2 | Histone H | PFE0510w | [illegible] |
| Q8I305 | Q8I305 | 4 | 4 | 4 | | PFI0720w | [illegible] |
| Q6ZMA7 | Q6ZMA7 | 2 | 2 | 2 | | PFD0310w | [illegible] |
| Q8IL28 | Q8IL28 | 4 | 4 | 4 | | PF14_042 | [illegible] |
| Q8I0U7 | Q8I0U7 | 3 | 3 | 3 | | PFI1545c | [illegible] |
| Q8IKE9 | Q8IKE9 | 4 | 4 | 4 | | PF14_065 | [illegible] |
| Q8IBJ7 | Q8IBJ7 | 2 | 2 | 2 | | MAL7P1.1 | [illegible] |
| Q8I527 | Q8I527 | 2 | 2 | 2 | | PFL1920c | [illegible] |
| Q8I3R6 | Q8I3R6 | 3 | 3 | 3 | 40S ribos | PFE0975c | [illegible] |
| Q8IFP3 | Q8IFP3 | 13 | 2 | 2 | | PFD1050w | [illegible] |
| O97250 | O97250 | 3 | 3 | 3 | | PFC0300c | [illegible] |
| A0A1437Y | A0A1437Y | 4;4 | 4;4 | 4;4 | | PFE0055c | [illegible] |
| Q8IKL9 | Q8IKL9 | 2 | 2 | 2 | | PF14_058 | [illegible] |
| Q8ILL2 | Q8ILL2 | 2 | 2 | 2 | | PF14_023 | [illegible] |
| A0A144A2 | A0A144A2 | 4;4 | 4;4 | 4;4 | STI1-like | PF14_032 | [illegible] |
| Q8IE00 | Q8IE00 | 2 | 2 | 2 | Eukaryoti | EIF6 | [illegible] |
| Q8I1T4 | Q8I1T4 | 4 | 4 | 4 | | Pfcrk-3 | [illegible] |
| Q8I6S4 | Q8I6S4 | 3 | 3 | 3 | Peptidyl-p | PfCyP24 | [illegible] |
| Q8IDI5 | Q8IDI5 | 2 | 2 | 2 | | PF13_026 | [illegible] |
| C6KSY6 | C6KSY6 | 3 | 3 | 3 | | PFF0700c | [illegible] |
| Q8IIH2 | Q8IIH2 | 4 | 4 | 4 | | PF11_020 | [illegible] |
| Q8ILV2 | Q8ILV2 | 2 | 2 | 2 | | PF14_014 | [illegible] |
| Q8I257 | Q8I257 | 2 | 2 | 2 | | PFA_0420 | [illegible] |
| Q8I420 | Q8I420 | 2 | 2 | 2 | | PfoTPT | [illegible] |
| Q8I2A9 | Q8I2A9 | 3 | 3 | 3 | | PFA_0155 | [illegible] |
| Q76NM4 | Q76NM4 | 3 | 3 | 2 | | Rab11a | [illegible] |
| Q8IDZ5 | Q8IDZ5 | 2 | 2 | 2 | | MAL13P1 | [illegible] |
| Q8I274 | Q8I274 | 4 | 4 | 4 | | Rab5c | [illegible] |
| Q8I1T6 | Q8I1T6 | 3 | 3 | 3 | | PFD0735c | [illegible] |
| Q8IJU2 | Q8IJU2 | 2 | 2 | 2 | | PF10_010 | [illegible] |
| Q8IM01 | Q8IM01 | 2 | 2 | 2 | | PF14_009 | [illegible] |
| Q8I441 | Q8I441 | 2 | 2 | 2 | | PFE0300c | [illegible] |
| Q8IJH1 | Q8IJH1 | 4 | 4 | 4 | | PF10_022 | [illegible] |
| Q8IDR3 | Q8IDR3 | 4 | 4 | 4 | Myosin-A | PF13_023 | [illegible] |
| A0A144A3 | A0A144A3 | 3;3 | 3;3 | 3;3 | Queuine t | PF14_032 | [illegible] |
| Q8I542 | Q8I542 | 2 | 2 | 2 | | PFL1845c | [illegible] |
| Q8ILP1 | Q8ILP1 | 4 | 4 | 4 | | PF14_077 | [illegible] |
| Q8IKM1 | Q8IKM1 | 3 | 3 | 3 | | PF14_058 | [illegible] |
| Q8IB31 | Q8IB31 | 2 | 2 | 2 | | MAL8P1.6 | [illegible] |
| A0A5K1K8 | A0A5K1K8 | 4 | 4 | 4 | | | [illegible] |
| C6KSW1 | C6KSW1 | 2 | 2 | 2 | | PFF0570c | [illegible] |
| Q8IC42 | Q8IC42 | 4 | 4 | 4 | | PF07_000 | [illegible] |
| Q8IKX8 | Q8IKX8 | 4 | 4 | 4 | | PF14_047 | [illegible] |
| Q8IKH5 | Q8IKH5 | 2 | 2 | 2 | Serine/th | PF14_063 | [illegible] |
| Q8IJK2 | Q8IJK2 | 2 | 2 | 2 | Autophag | PF10_019 | [illegible] |
| Q8ILX1 | Q8ILX1 | 3 | 3 | 3 | | PF14_012 | [illegible] |
| A0A5K1K9 | A0A5K1K9 | 2 | 2 | 2 | | | [illegible] |
| A0A1437X | A0A1437X | 4;4 | 4;4 | 4;4 | Uncharact | PFB0315w | [illegible] |
| O97225 | O97225 | 3 | 3 | 3 | | PFC0165w | [illegible] |
| Q8IJT4 | Q8IJT4 | 2 | 2 | 2 | | PF10_010 | [illegible] |
| Q8I5K4 | Q8I5K4 | 5 | 5 | 5 | | PFL1005c | [illegible] |
| C0H5F0 | C0H5F0 | 3 | 3 | 3 | | MAL13P1 | [illegible] |
| Q8I275 | Q8I275 | 4 | 4 | 4 | | AARP2 | [illegible] |
| Q8II92 | Q8II92 | 5 | 5 | 5 | | PF11_028 | [illegible] |
| A0A144A2 | A0A144A2 | 4;4 | 4;4 | 4;4 | NAD-depe | Sir2B | [illegible] |
| Q8IKS3 | Q8IKS3 | 5 | 5 | 5 | | PF14_052 | [illegible] |
| Q8IJP4 | Q8IJP4 | 2 | 2 | 2 | | PF10_014 | [illegible] |
| Q8I6T3 | Q8I6T3 | 2 | 2 | 2 | Proteasor | PF13_015 | [illegible] |
| Q8I5E2 | Q8I5E2 | 4 | 4 | 4 | | PFL1335w | [illegible] |
| C0H4M0 | C0H4M0 | 2 | 2 | 2 | | RAMA | [illegible] |
| Q8IIQ7 | Q8IIQ7 | 2 | 2 | 2 | | PF11_011 | [illegible] |
| Q8IKE8 | Q8IKE8 | 4 | 4 | 4 | | PF14_065 | [illegible] |
| Q8I398 | Q8I398 | 6 | 6 | 6 | | PFI0250c | [illegible] |
| Q8IDG3 | Q8IDG3 | 3 | 3 | 3 | Proteasor | PF13_028 | [illegible] |
| Q8I2G0 | Q8I2G0 | 3 | 3 | 3 | | REX2 | [illegible] |
| Q7KQL4 | Q7KQL4 | 6 | 6 | 6 | DNA poly | PF10_016 | [illegible] |
| Q8I5I4 | Q8I5I4 | 3 | 3 | 3 | | PFL1115w | [illegible] |
| A0A1437V | A0A1437V | 6 | 2 | 2 | | | [illegible] |
| Q8ILC5 | Q8ILC5 | 2 | 2 | 2 | | PF14_031 | [illegible] |
| Q8IHS5 | Q8IHS5 | 2 | 2 | 2 | 40S ribos | PF11_045 | [illegible] |
| Q8IKT8 | Q8IKT8 | 2 | 2 | 2 | | PF14_051 | [illegible] |
| Q8IK92 | Q8IK92 | 2 | 2 | 2 | | PF14_071 | [illegible] |
| Q8I603 | Q8I603 | 2 | 2 | 2 | Eukaryoti | PFL0210c | [illegible] |
| Q8IJN4 | Q8IJN4 | 2 | 2 | 2 | | PF10_015 | [illegible] |
| Q8IM73 | Q8IM73 | 2 | 2 | 2 | | PF14_001 | [illegible] |
| C6KT04 | C6KT04 | 2 | 2 | 2 | | PFF0790c | [illegible] |
| O77327 | O77327 | 2 | 2 | 2 | | PFC0375c | [illegible] |
| A0A1437Y | A0A1437Y | 2 | 2 | 2 | | | [illegible] |
| A0A5K1K9 | A0A5K1K9 | 2 | 2 | 2 | | | [illegible] |
| Q8IBF2 | Q8IBF2 | 3 | 3 | 3 | | MAL7P1.1 | [illegible] |
| Q8ILV6 | Q8ILV6 | 2 | 2 | 2 | | PF14_013 | [illegible] |
| A0A5K1K9 | A0A5K1K9 | 6 | 6 | 6 | | | [illegible] |
| Q8IKK3 | Q8IKK3 | 4 | 4 | 4 | | PF14_060 | [illegible] |
| Q8I261 | Q8I261 | 2 | 2 | 2 | | PFA_0400 | [illegible] |
| C6KT66 | C6KT66 | 4 | 4 | 4 | | PfCS | [illegible] |
| Q8I249 | Q8I249 | 2 | 2 | 2 | | PFA_0465 | [illegible] |
| Q8IIU9 | Q8IIU9 | 3 | 3 | 3 | | PF11_006 | [illegible] |
| Q8I4T9 | Q8I4T9 | 2 | 2 | 2 | | PFL2375c | [illegible] |
| Q8ILL9 | Q8ILL9 | 4 | 4 | 4 | Serine/th | PP7 | [illegible] |
| Q8IBN7 | Q8IBN7 | 3 | 3 | 3 | | MAL7P1.1 | [illegible] |
| Q8I3P4 | Q8I3P4 | 2 | 2 | 2 | | PFE1095w | [illegible] |
| Q8I3B0 | Q8I3B0 | 3 | 3 | 3 | | PFI0190w | [illegible] |
| Q8I2V5 | Q8I2V5 | 2 | 2 | 2 | | PFI0970c | [illegible] |
| O97319 | O97319 | 2 | 2 | 2 | | PFC0870w | [illegible] |
| Q8I6V0 | Q8I6V0 | 2 | 2 | 2 | | PF14_055 | [illegible] |
| Q8IIQ0 | Q8IIQ0 | 4 | 4 | 4 | | PF11_011 | [illegible] |
| Q8IL35 | Q8IL35 | 3 | 3 | 3 | | PF14_041 | [illegible] |
| Q8IHX4 | Q8IHX4 | 2 | 2 | 2 | | PF11_040 | [illegible] |
| Q8I3W5 | Q8I3W5 | 5 | 5 | 5 | | PFE0710w | [illegible] |
| Q8ILY2 | Q8ILY2 | 2 | 2 | 2 | | PF14_011 | [illegible] |
| Q8I239 | Q8I239 | 2 | 2 | 2 | | NCS | [illegible] |
| Q8IDX1 | Q8IDX1 | 2 | 2 | 2 | | MAL13P1 | [illegible] |
| Q8IJQ6 | Q8IJQ6 | 5 | 5 | 5 | | PF10_013 | [illegible] |
| O77317 | O77317 | 4 | 4 | 4 | | PFC0150w | [illegible] |
| Q8I2H3 | Q8I2H3 | 3 | 3 | 3 | | PFI1670c | [illegible] |
| Q8I418 | Q8I418 | 2 | 2 | 2 | | RCC1 | [illegible] |
| A0A5K1K9 | A0A5K1K9 | 2 | 2 | 2 | | | [illegible] |
| Q8I3R4 | Q8I3R4 | 3 | 3 | 3 | | PFE0985w | [illegible] |
| Q8I427 | Q8I427 | 4 | 4 | 4 | | PFE0375w | [illegible] |
| C0H5H0 | C0H5H0 | 3 | 3 | 3 | | MAL13P1 | [illegible] |
| Q8IID2 | Q8IID2 | 4 | 4 | 4 | | PF11_024 | [illegible] |
| Q8I3V5 | Q8I3V5 | 2 | 2 | 2 | | PFE0765w | [illegible] |
| Q8IIH7 | Q8IIH7 | 2 | 2 | 2 | | PF11_019 | [illegible] |
| Q8IIK1 | Q8IIK1 | 2 | 2 | 2 | | PF11_017 | [illegible] |
| Q8IK08 | Q8IK08 | 2 | 2 | 2 | | PF10_003 | [illegible] |
| A0A14377 | A0A14377 | 6;6 | 6;6 | 6;6 | Uncharact | PFL1135c | [illegible] |
| Q8I5T1 | Q8I5T1 | 2 | 2 | 2 | | PFL0600w | [illegible] |
| Q8I422 | Q8I422 | 3 | 3 | 3 | | PFE0400w | [illegible] |
| Q8IIL9 | Q8IIL9 | 2 | 2 | 2 | | PF11_015 | [illegible] |
| Q8IKI7 | Q8IKI7 | 3 | 3 | 3 | | PF14_061 | [illegible] |
| Q8I433 | Q8I433 | 3 | 3 | 3 | | ROM4 | [illegible] |
| Q8I5E7 | Q8I5E7 | 3 | 2 | 2 | | PFL1310c | [illegible] |
| A0A144A3 | A0A144A3 | 2 | 2 | 2 | | | [illegible] |
| Q7KWJ5 | Q7KWJ5 | 2 | 2 | 2 | | HT1 | [illegible] |
| Q8I5M5 | Q8I5M5 | 2 | 2 | 2 | | PFL0885w | [illegible] |
| C6KSY1 | C6KSY1 | 4 | 4 | 4 | | MyoE | [illegible] |
| Q8ILJ1 | Q8ILJ1 | 5 | 5 | 5 | | PF14_025 | [illegible] |
| C6KT11 | C6KT11 | 2 | 2 | 2 | | PFF0825c | [illegible] |
| O77318 | O77318 | 2 | 2 | 2 | | PFC0330w | [illegible] |
| P46468 | P46468 | 2 | 1 | 1 | Putative c | PF07_004 | [illegible] |
| Q8I2I3 | Q8I2I3 | 2 | 2 | 2 | | PFI1620c | [illegible] |
| Q8IAN4 | Q8IAN4 | 3 | 3 | 3 | | PF08_012 | [illegible] |
| Q8IDB5 | Q8IDB5 | 2 | 2 | 2 | | MAL13P1 | [illegible] |
| Q8IIX6 | Q8IIX6 | 2 | 2 | 2 | | PF11_003 | [illegible] |
| Q8IE14 | Q8IE14 | 2 | 2 | 2 | Signal pep | SP21 | [illegible] |
| C0H4A0 | C0H4A0 | 2 | 2 | 2 | | PFD0450c | [illegible] |
| Q76NM1 | Q76NM1 | 2 | 2 | 2 | ER lumen | ERD2 | [illegible] |
| C0H4C6 | C0H4C6 | 3 | 3 | 3 | | PFE0245c | [illegible] |
| A0A144A4 | A0A144A4 | 3 | 3 | 3 | | | [illegible] |
| A0A144A6 | A0A144A6 | 4;4 | 4;4 | 4;4 | Bifunctior | PF14_051 | [illegible] |
| Q8ILK9 | Q8ILK9 | 2 | 2 | 2 | | PF14_023 | [illegible] |
| C0H4W0 | C0H4W0 | 2 | 2 | 2 | | MAL8P1.6 | [illegible] |
| A0A1437Y | A0A1437Y | 3 | 3 | 3 | | | [illegible] |
| Q8I2Z3 | Q8I2Z3 | 2 | 2 | 2 | | PFI0780w | [illegible] |
| Q8IEL8 | Q8IEL8 | 2 | 2 | 2 | | PF13_005 | [illegible] |
| Q8ILZ3 | Q8ILZ3 | 4 | 4 | 4 | CTP synth | PF14_010 | [illegible] |
| C0H4R1 | C0H4R1 | 2 | 2 | 2 | | MAL8P1.1 | [illegible] |
| Q8IJ37 | Q8IJ37 | 2 | 2 | 2 | Pyruvate k | PF10_036 | [illegible] |
| Q8ILA9 | Q8ILA9 | 2 | 2 | 2 | | PF14_033 | [illegible] |
| O97245 | O97245 | 3 | 3 | 3 | Glycerol-3 | PFC0275w | [illegible] |
| A0A1437Y | A0A1437Y | 2;2 | 2;2 | 2;2 | Uncharact | PF11_020 | [illegible] |
| C0H5G3 | C0H5G3 | 2 | 2 | 2 | 60S ribos | MAL13P1 | [illegible] |
| C6KSL5 | C6KSL5 | 3 | 1 | 0 | | PfMC-2TM | [illegible] |
| O96217 | O96217 | 2 | 2 | 2 | | PFB0620w | [illegible] |
| Q8I480 | Q8I480 | 2 | 2 | 2 | RING fing | PFE0100w | [illegible] |
| Q8IIN9 | Q8IIN9 | 2 | 2 | 2 | | PF11_013 | [illegible] |
| Q8IKA7 | Q8IKA7 | 2 | 2 | 2 | | PF14_069 | [illegible] |
| Q8IKW9 | Q8IKW9 | 2 | 2 | 2 | | PF14_048 | [illegible] |
| Q8IBS3 | Q8IBS3 | 3 | 3 | 3 | | PF07_007 | [illegible] |
| A0A5K1K8 | A0A5K1K8 | 3 | 3 | 3 | | | [illegible] |
| Q8IJV1 | Q8IJV1 | 2 | 2 | 2 | | PF10_009 | [illegible] |
| Q8I3L3 | Q8I3L3 | 2 | 2 | 2 | | PFE1255w | [illegible] |
| Q8IBI7 | Q8IBI7 | 2 | 2 | 2 | | PF07_011 | [illegible] |
| C0H524 | C0H524 | 2 | 2 | 2 | | PFI0385c | [illegible] |
| Q8IJT1 | Q8IJT1 | 2 | 2 | 2 | Proteasor | PF10_011 | [illegible] |
| Q8IBZ4 | Q8IBZ4 | 2 | 2 | 2 | | PF07_003 | [illegible] |
| Q8IKP8 | Q8IKP8 | 3 | 3 | 3 | | PF14_055 | [illegible] |

| | | | | | | | | | | | | | | |
|---|---|---|---|---|---|---|---|---|---|---|---|---|---|---|
| Q8IBI5 | Q8IBI5 | 3 | 3 | 3 | | MAL7P1.1 | tr\|Q8IBI5\|Q8IBI5_PLAF7 Cysteine desulfurase, putative OS=Pl | 1 | 3 | 3 | 3 | 0 | 1 | [illegible] |
| Q8IDX6;C( | Q8IDX6;C( | 2;1 | 2;1 | 2;1 | Reticuloc | PF13_019 | sp\|Q8IDX6\|RBP2A_PLAF7 Reticulocyte-binding protein 2 hom | 2 | 2 | 2 | 2 | 1 | 2 | [illegible] |
| Q8IL78 | Q8IL78 | 2 | 2 | 2 | | PF14_037 | tr\|Q8IL78\|Q8IL78_PLAF7 DEAD/DEAH box helicase, putative O | 1 | 2 | 2 | 2 | 0 | 0 | [illegible] |
| Q8I665 | Q8I665 | 5 | 5 | 5 | | PFB0194w | tr\|Q8I665\|Q8I665_PLAF7 Uncharacterized protein OS=Plasmo | 1 | 5 | 5 | 5 | 4 | 4 | [illegible] |
| Q8IBV7 | Q8IBV7 | 3 | 3 | 3 | Histone H | PF07_005 | tr\|Q8IBV7\|Q8IBV7_PLAF7 Histone H2B OS=Plasmodium falcip | 1 | 3 | 3 | 3 | 1 | 1 | [illegible] |
| C6KSX0 | C6KSX0 | 2 | 2 | 2 | Merozoite | PF12 | sp\|C6KSX0\|PF12_PLAF7 Merozoite surface protein P12 OS=Pla | 1 | 2 | 2 | 2 | 1 | 1 | [illegible] |
| Q8IIK3 | Q8IIK3 | 3 | 3 | 2 | | PF11_017 | tr\|Q8IIK3\|Q8IIK3_PLAF7 Peptidyl-prolyl cis-trans isomerase O | 1 | 3 | 3 | 2 | 1 | 1 | [illegible] |
| Q8IE43 | Q8IE43 | 2 | 2 | 2 | | PF13_015 | tr\|Q8IE43\|Q8IE43_PLAF7 Uncharacterized protein OS=Plasmo | 1 | 2 | 2 | 2 | 1 | 1 | [illegible] |
| Q8IDX8 | Q8IDX8 | 2 | 2 | 2 | | MSP7 | tr\|Q8IDX8\|Q8IDX8_PLAF7 Merozoite surface protein 7 OS=Pla | 1 | 2 | 2 | 2 | 1 | 1 | [illegible] |
| Q8IJ69 | Q8IJ69 | 2 | 2 | 2 | | PF10_033 | tr\|Q8IJ69\|Q8IJ69_PLAF7 Sec1 family protein, putative OS=Plas | 1 | 2 | 2 | 2 | 1 | 0 | [illegible] |
| Q8I4S7 | Q8I4S7 | 2 | 2 | 2 | | PFL2435w | tr\|Q8I4S7\|Q8I4S7_PLAF7 Transcription initiation factor IIA sub | 1 | 2 | 2 | 2 | 0 | 0 | [illegible] |
| Q8I468 | Q8I468 | 2 | 2 | 2 | | PFE0160c | tr\|Q8I468\|Q8I468_PLAF7 Serine/arginine-rich splicing factor 1 | 1 | 2 | 2 | 2 | 1 | 1 | [illegible] |
| C6KSR7 | C6KSR7 | 2 | 2 | 2 | | PFF0340c | tr\|C6KSR7\|C6KSR7_PLAF7 Glutaredoxin-like protein OS=Plasm | 1 | 2 | 2 | 2 | 1 | 1 | [illegible] |
| C0H4S5 | C0H4S5 | 2 | 2 | 2 | | MAL8P1.1 | tr\|C0H4S5\|C0H4S5_PLAF7 YEATS domain-containing protein, | 1 | 2 | 2 | 2 | 0 | 0 | [illegible] |
| Q8IHN4 | Q8IHN4 | 3 | 3 | 3 | | PF11_050 | tr\|Q8IHN4\|Q8IHN4_PLAF7 Antigen 332, DBL-like protein OS=P | 1 | 3 | 3 | 3 | 2 | 2 | [illegible] |
| C6KT80 | C6KT80 | 4 | 4 | 4 | | PFF1175c | tr\|C6KT80\|C6KT80_PLAF7 Uncharacterized protein OS=Plasmo | 1 | 4 | 4 | 4 | 0 | 0 | [illegible] |
| C6KSQ9 | C6KSQ9 | 2 | 2 | 2 | | PFF0300w | tr\|C6KSQ9\|C6KSQ9_PLAF7 RNA-binding protein, putative OS= | 1 | 2 | 2 | 2 | 0 | 0 | [illegible] |
| Q8I1T0 | Q8I1T0 | 2 | 2 | 2 | | PFD0760c | tr\|Q8I1T0\|Q8I1T0_PLAF7 Uncharacterized protein OS=Plasmo | 1 | 2 | 2 | 2 | 0 | 0 | [illegible] |
| Q8I2N8 | Q8I2N8 | 2 | 2 | 2 | | PFI1330c | tr\|Q8I2N8\|Q8I2N8_PLAF7 Uncharacterized protein OS=Plasmo | 1 | 2 | 2 | 2 | 0 | 0 | [illegible] |
| Q8I416 | Q8I416 | 2 | 2 | 2 | | PFE0430w | tr\|Q8I416\|Q8I416_PLAF7 Pre-mRNA-processing ATP-depende | 1 | 2 | 2 | 2 | 0 | 0 | [illegible] |
| Q8IBL4 | Q8IBL4 | 2 | 2 | 2 | | MAL7P1.1 | tr\|Q8IBL4\|Q8IBL4_PLAF7 RNA-binding protein, putative OS=Pl | 1 | 2 | 2 | 2 | 1 | 1 | [illegible] |
| Q8IED9 | Q8IED9 | 2 | 2 | 2 | | MAL13P1 | tr\|Q8IED9\|Q8IED9_PLAF7 Uncharacterized protein OS=Plasmo | 1 | 2 | 2 | 2 | 0 | 1 | [illegible] |
| Q8I2C5 | Q8I2C5 | 3 | 3 | 3 | | RIF | tr\|Q8I2C5\|Q8I2C5_PLAF7 Rifin OS=Plasmodium falciparum (is | 1 | 3 | 3 | 3 | 0 | 0 | [illegible] |
| Q8I717 | Q8I717 | 2 | 2 | 2 | | PF14_067 | tr\|Q8I717\|Q8I717_PLAF7 RNA 3-terminal phosphate cyclase-li | 1 | 2 | 2 | 2 | 0 | 0 | [illegible] |
| Q8IEP8 | Q8IEP8 | 3 | 3 | 3 | | MAL13P1 | tr\|Q8IEP8\|Q8IEP8_PLAF7 Uncharacterized protein OS=Plasmo | 1 | 3 | 3 | 3 | 2 | 2 | [illegible] |
| C6S3B8 | C6S3B8 | 3 | 3 | 3 | | PFB0850c | tr\|C6S3B8\|C6S3B8_PLAF7 SUZ domain-containing protein, put | 1 | 3 | 3 | 3 | 1 | 1 | [illegible] |
| O96243 | O96243 | 2 | 2 | 2 | | PFB0750w | tr\|O96243\|O96243_PLAF7 Vacuolar protein sorting-associated | 1 | 2 | 2 | 2 | 1 | 1 | [illegible] |
| Q8IKR1 | Q8IKR1 | 3 | 3 | 3 | | PF14_054 | tr\|Q8IKR1\|Q8IKR1_PLAF7 V-type H(+)-translocating pyrophos | 1 | 3 | 3 | 3 | 1 | 1 | [illegible] |
| O96252 | O96252 | 4 | 4 | 4 | ATP synth | PFB0795w | tr\|O96252\|O96252_PLAF7 ATP synthase subunit alpha OS=Pla | 1 | 4 | 4 | 4 | 1 | 3 | [illegible] |
| C0H4A1 | C0H4A1 | 2 | 2 | 2 | | PFD0465c | tr\|C0H4A1\|C0H4A1_PLAF7 NAD(P)-binding protein, putative ( | 1 | 2 | 2 | 2 | 0 | 0 | [illegible] |
| Q8IJC0 | Q8IJC0 | 2 | 2 | 2 | | PF10_027 | tr\|Q8IJC0\|Q8IJC0_PLAF7 Nucleolar preribosomal assembly pro | 1 | 2 | 2 | 2 | 0 | 0 | [illegible] |
| Q9NLB2 | Q9NLB2 | 2 | 2 | 2 | | GRX1 | tr\|Q9NLB2\|Q9NLB2_PLAF7 Glutaredoxin OS=Plasmodium falc | 1 | 2 | 2 | 2 | 2 | 1 | [illegible] |
| A0A144A4 | A0A144A4 | 2;2 | 2;2 | 2;2 | Triosepho | TPI | tr\|A0A144A4G4\|A0A144A4G4_PLAF7 Triosephosphate isomer | 2 | 2 | 2 | 2 | 1 | 1 | [illegible] |
| Q8I600 | Q8I600 | 5 | 5 | 5 | | PFL0225c | tr\|Q8I600\|Q8I600_PLAF7 Uncharacterized protein OS=Plasmo | 1 | 5 | 5 | 5 | 0 | 0 | [illegible] |
| O77366 | O77366 | 2 | 2 | 2 | | PFC0850c | tr\|O77366\|O77366_PLAF7 CCR4 domain-containing protein 4 | 1 | 2 | 2 | 2 | 0 | 0 | [illegible] |
| Q8I4T0 | Q8I4T0 | 2 | 2 | 2 | | PFL2420w | tr\|Q8I4T0\|Q8I4T0_PLAF7 Uncharacterized protein OS=Plasmo | 1 | 2 | 2 | 2 | 1 | 1 | [illegible] |
| B3FEM7 | B3FEM7 | 2 | 2 | 2 | | PDEgamm | tr\|B3FEM7\|B3FEM7_PLAF7 Phosphodiesterase OS=Plasmodiu | 1 | 2 | 2 | 2 | 1 | 1 | [illegible] |
| C6KSX6 | C6KSX6 | 3 | 3 | 3 | | PFF0645c | tr\|C6KSX6\|C6KSX6_PLAF7 Rhoptry protein ROP14 OS=Plasmo | 1 | 3 | 3 | 3 | 0 | 2 | [illegible] |
| O97253 | O97253 | 2 | 2 | 2 | | PFC0315c | tr\|O97253\|O97253_PLAF7 Spindle and kinetochore-associated | 1 | 2 | 2 | 2 | 0 | 0 | [illegible] |
| Q8IDJ7 | Q8IDJ7 | 4 | 4 | 4 | | MAL13P1 | tr\|Q8IDJ7\|Q8IDJ7_PLAF7 Small nuclear ribonucleoprotein F O | 1 | 4 | 4 | 4 | 0 | 2 | [illegible] |
| Q8ILF4 | Q8ILF4 | 2 | 2 | 2 | | PF14_029 | tr\|Q8ILF4\|Q8ILF4_PLAF7 PhIL1 interacting protein PIP2 OS=Pla | 1 | 2 | 2 | 2 | 2 | 1 | [illegible] |
| Q8IKG8 | Q8IKG8 | 3 | 3 | 3 | | PF14_063 | tr\|Q8IKG8\|Q8IKG8_PLAF7 Uncharacterized protein OS=Plasmo | 1 | 3 | 3 | 3 | 1 | 2 | [illegible] |
| A0A144A2 | A0A144A2 | 2 | 2 | 2 | | | tr\|A0A144A2I0\|A0A144A2I0_PLAF7 DNA-directed RNA polyme | 1 | 2 | 2 | 2 | 0 | 0 | [illegible] |
| C6KTB8 | C6KTB8 | 5 | 5 | 5 | | PfPK4 | tr\|C6KTB8\|C6KTB8_PLAF7 Protein kinase PK4 OS=Plasmodium | 1 | 5 | 5 | 5 | 0 | 0 | [illegible] |
| Q8IIC3 | Q8IIC3 | 2 | 2 | 2 | | PF11_025 | tr\|Q8IIC3\|Q8IIC3_PLAF7 Endoplasmic reticulum oxidoreducti | 1 | 2 | 2 | 2 | 0 | 0 | [illegible] |
| Q8IEA1 | Q8IEA1 | 2 | 2 | 2 | | MAL13P1 | tr\|Q8IEA1\|Q8IEA1_PLAF7 Uncharacterized protein OS=Plasmo | 1 | 2 | 2 | 2 | 0 | 0 | [illegible] |
| C6KT64 | C6KT64 | 3 | 3 | 3 | | PFF1095w | tr\|C6KT64\|C6KT64_PLAF7 Leucine--tRNA ligase, putative OS=P | 1 | 3 | 3 | 3 | 0 | 1 | [illegible] |
| Q8I1T2 | Q8I1T2 | 2 | 2 | 2 | | PFD0750w | tr\|Q8I1T2\|Q8I1T2_PLAF7 Nuclear cap-binding protein subunit | 1 | 2 | 2 | 2 | 1 | 0 | [illegible] |
| Q9U0J3 | Q9U0J3 | 3 | 3 | 3 | | PFD0460c | tr\|Q9U0J3\|Q9U0J3_PLAF7 Methyltransferase, putative OS=Pla | 1 | 3 | 3 | 3 | 1 | 1 | [illegible] |
| O77316 | O77316 | 2 | 2 | 2 | | PFC0130c | tr\|O77316\|O77316_PLAF7 RNA-binding protein, putative OS=P | 1 | 2 | 2 | 2 | 0 | 0 | [illegible] |
| O97287 | O97287 | 2 | 2 | 2 | | PFC0930c | tr\|O97287\|O97287_PLAF7 Uncharacterized protein OS=Plasmo | 1 | 2 | 2 | 2 | 0 | 0 | [illegible] |
| Q8IDA3 | Q8IDA3 | 3 | 3 | 3 | | MAL13P1 | tr\|Q8IDA3\|Q8IDA3_PLAF7 ER membrane protein complex sub | 1 | 3 | 3 | 3 | 1 | 1 | [illegible] |
| Q8ILI2 | Q8ILI2 | 3 | 3 | 3 | | PF14_026 | tr\|Q8ILI2\|Q8ILI2_PLAF7 Proliferation-associated protein 2g4, | 1 | 3 | 3 | 3 | 1 | 2 | [illegible] |
| Q8IK86 | Q8IK86 | 4 | 4 | 4 | | PF14_072 | tr\|Q8IK86\|Q8IK86_PLAF7 HA2 domain-containing protein OS= | 1 | 4 | 4 | 4 | 0 | 0 | [illegible] |
| Q8IIQ6 | Q8IIQ6 | 2 | 2 | 2 | Vacuolar | PF11_011 | tr\|Q8IIQ6\|Q8IIQ6_PLAF7 Vacuolar protein sorting-associated | 1 | 2 | 2 | 2 | 0 | 0 | [illegible] |
| Q7KQJ9 | Q7KQJ9 | 3 | 3 | 3 | Proliferat | PFL1285c | tr\|Q7KQJ9\|Q7KQJ9_PLAF7 Proliferating cell nuclear antigen O | 1 | 3 | 3 | 3 | 1 | 1 | [illegible] |
| C0H5H9 | C0H5H9 | 2 | 2 | 2 | | PF13_026 | tr\|C0H5H9\|C0H5H9_PLAF7 Ubiquitin-activating enzyme E1, pu | 1 | 2 | 2 | 2 | 0 | 0 | [illegible] |
| Q8IJR0 | Q8IJR0 | 2 | 2 | 2 | Phosphoi | PI PLC | tr\|Q8IJR0\|Q8IJR0_PLAF7 Phosphoinositide phospholipase C O | 1 | 2 | 2 | 2 | 0 | 0 | [illegible] |
| Q8IM06 | Q8IM06 | 2 | 2 | 2 | | PF14_008 | tr\|Q8IM06\|Q8IM06_PLAF7 WD repeat-containing protein, put | 1 | 2 | 2 | 2 | 0 | 0 | [illegible] |
| Q8I6Z3 | Q8I6Z3 | 2 | 2 | 2 | Protein pe | MAL7P1.1 | tr\|Q8I6Z3\|Q8I6Z3_PLAF7 Protein pelota homolog OS=Plasmo | 1 | 2 | 2 | 2 | 0 | 0 | [illegible] |
| Q8ILD9 | Q8ILD9 | 2 | 2 | 2 | | PF14_030 | tr\|Q8ILD9\|Q8ILD9_PLAF7 Leucine-rich repeat protein OS=Plas | 1 | 2 | 2 | 2 | 1 | 1 | [illegible] |
| A0A1437V | A0A1437V | 2 | 2 | 2 | | | tr\|A0A1437VW7\|A0A1437VW7_PLAF7 Uncharacterized protei | 1 | 2 | 2 | 2 | 1 | 1 | [illegible] |
| A0A144A3 | A0A144A3 | 2;1;1;1 | 2;1;1;1 | 2;1;1;1 | | 3D7Surf8. | tr\|A0A144A325\|A0A144A325_PLAF7 Surface-associated inters | 4 | 2 | 2 | 2 | 0 | 0 | [illegible] |
| Q8I2M7 | Q8I2M7 | 3 | 3 | 3 | | PFI1385c | tr\|Q8I2M7\|Q8I2M7_PLAF7 Uncharacterized protein OS=Plasm | 1 | 3 | 3 | 3 | 1 | 1 | [illegible] |
| Q8IM62 | Q8IM62 | 2 | 2 | 2 | | PF14_002 | tr\|Q8IM62\|Q8IM62_PLAF7 Uncharacterized protein OS=Plasm | 1 | 2 | 2 | 2 | 0 | 0 | [illegible] |
| Q8ILY8 | Q8ILY8 | 3 | 3 | 3 | | PF14_010 | tr\|Q8ILY8\|Q8ILY8_PLAF7 Uncharacterized protein OS=Plasmo | 1 | 3 | 3 | 3 | 2 | 3 | [illegible] |
| Q8I3B4 | Q8I3B4 | 2 | 2 | 2 | | PFI0165c | tr\|Q8I3B4\|Q8I3B4_PLAF7 ATP-dependent RNA helicase DDX6 | 1 | 2 | 2 | 2 | 0 | 0 | [illegible] |
| C0H4L9 | C0H4L9 | 3 | 3 | 3 | | MAL7P1.2 | tr\|C0H4L9\|C0H4L9_PLAF7 Uncharacterized protein OS=Plasmo | 1 | 3 | 3 | 3 | 0 | 1 | [illegible] |
| Q8IJR3 | Q8IJR3 | 3 | 3 | 3 | | PF10_012 | tr\|Q8IJR3\|Q8IJR3_PLAF7 Uncharacterized protein OS=Plasmo | 1 | 3 | 3 | 3 | 0 | 0 | [illegible] |
| Q8IJE1 | Q8IJE1 | 2 | 2 | 2 | | PF10_025 | tr\|Q8IJE1\|Q8IJE1_PLAF7 Uncharacterized protein OS=Plasmod | 1 | 2 | 2 | 2 | 1 | 1 | [illegible] |
| Q8I657 | Q8I657 | 2 | 2 | 2 | | PFB0826c | tr\|Q8I657\|Q8I657_PLAF7 E2F-associated phosphoprotein, put | 1 | 2 | 2 | 2 | 1 | 1 | [illegible] |
| Q8ID94 | Q8ID94 | 2 | 2 | 2 | Uncharact | MAL13P1 | sp\|Q8ID94\|YPF12_PLAF7 Uncharacterized protein MAL13P1.30 | 1 | 2 | 2 | 2 | 0 | 0 | [illegible] |
| O77363 | O77363 | 2 | 2 | 2 | | PFC0545c | tr\|O77363\|O77363_PLAF7 Uncharacterized protein OS=Plasmo | 1 | 2 | 2 | 2 | 0 | 0 | [illegible] |
| Q8I3R5 | Q8I3R5 | 4 | 4 | 4 | | PFE0980c | tr\|Q8I3R5\|Q8I3R5_PLAF7 CCR4 domain-containing protein 1, | 1 | 4 | 4 | 4 | 0 | 2 | [illegible] |
| Q8IE02 | Q8IE02 | 3 | 3 | 3 | | apn1 | tr\|Q8IE02\|Q8IE02_PLAF7 DNA-(Apurinic or apyrimidinic site) l | 1 | 3 | 3 | 3 | 0 | 0 | [illegible] |
| A0A144A5 | A0A144A5 | 3 | 3 | 3 | | | tr\|A0A144A5C9\|A0A144A5C9_PLAF7 Uncharacterized protein | 1 | 3 | 3 | 3 | 0 | 0 | [illegible] |
| A0A5K1K8 | A0A5K1K8 | 2 | 2 | 2 | | | tr\|A0A5K1K8Z2\|A0A5K1K8Z2_PLAF7 Uncharacterized protein | 1 | 2 | 2 | 2 | 0 | 0 | [illegible] |
| O77393 | O77393 | 3 | 3 | 3 | | PFC0705c | tr\|O77393\|O77393_PLAF7 Uncharacterized protein OS=Plasmo | 1 | 3 | 3 | 3 | 0 | 0 | [illegible] |
| Q76NN7 | Q76NN7 | 2 | 2 | 2 | Peptidyl-p | PfCyP19 | tr\|Q76NN7\|Q76NN7_PLAF7 Peptidyl-prolyl cis-trans isomeras | 1 | 2 | 2 | 2 | 1 | 1 | [illegible] |
| Q8I1U8 | Q8I1U8 | 2 | 2 | 2 | Ubiquitin | PFD0680c | tr\|Q8I1U8\|Q8I1U8_PLAF7 Ubiquitin carboxyl-terminal hydrola | 1 | 2 | 2 | 2 | 0 | 1 | [illegible] |
| Q8I562 | Q8I562 | 3 | 3 | 3 | | PFL1745c | tr\|Q8I562\|Q8I562_PLAF7 Clustered-asparagine-rich protein O | 1 | 3 | 3 | 3 | 0 | 0 | [illegible] |
| Q8I5B6 | Q8I5B6 | 2 | 2 | 2 | | HslV | tr\|Q8I5B6\|Q8I5B6_PLAF7 ATP-dependent protease subunit Cl | 1 | 2 | 2 | 2 | 1 | 1 | [illegible] |
| Q8I5X8 | Q8I5X8 | 3 | 3 | 3 | | PFL0335c | tr\|Q8I5X8\|Q8I5X8_PLAF7 Eukaryotic translation initiation fact | 1 | 3 | 3 | 3 | 0 | 0 | [illegible] |
| Q8I617 | Q8I617 | 2 | 2 | 2 | | PFL0140w | tr\|Q8I617\|Q8I617_PLAF7 DNA/RNA-binding protein, putative | 1 | 2 | 2 | 2 | 0 | 0 | [illegible] |
| Q8IDD4 | Q8IDD4 | 2 | 2 | 2 | | MAL13P1 | tr\|Q8IDD4\|Q8IDD4_PLAF7 Serine/threonine protein kinase, p | 1 | 2 | 2 | 2 | 1 | 1 | [illegible] |
| Q8I406 | Q8I406 | 2 | 2 | 2 | | PFE0485w | tr\|Q8I406\|Q8I406_PLAF7 Phosphatidylinositol 4-kinase OS=Pl | 1 | 2 | 2 | 2 | 1 | 1 | [illegible] |
| Q8IBQ0 | Q8IBQ0 | 2 | 2 | 2 | | PF07_008 | tr\|Q8IBQ0\|Q8IBQ0_PLAF7 Uncharacterized protein OS=Plasm | 1 | 2 | 2 | 2 | 1 | 1 | [illegible] |
| Q8I604 | Q8I604 | 2 | 2 | 2 | | PFL0205w | tr\|Q8I604\|Q8I604_PLAF7 Uncharacterized protein OS=Plasmo | 1 | 2 | 2 | 2 | 1 | 1 | [illegible] |
| Q8I5N4 | Q8I5N4 | 2 | 2 | 2 | | PFL0840c | tr\|Q8I5N4\|Q8I5N4_PLAF7 Uncharacterized protein OS=Plasmo | 1 | 2 | 2 | 2 | 0 | 0 | [illegible] |
| Q8I5L0 | Q8I5L0 | 2 | 2 | 2 | | PFL0975w | tr\|Q8I5L0\|Q8I5L0_PLAF7 Cell cycle associated protein, putativ | 1 | 2 | 2 | 2 | 0 | 0 | [illegible] |
| Q8ID32 | Q8ID32 | 2 | 2 | 2 | | MAL13P1 | tr\|Q8ID32\|Q8ID32_PLAF7 Ribosome biogenesis protein MRT4 | 1 | 2 | 2 | 2 | 0 | 0 | [illegible] |
| Q8III9 | Q8III9 | 2 | 2 | 2 | | PF11_018 | tr\|Q8III9\|Q8III9_PLAF7 Uncharacterized protein OS=Plasmodi | 1 | 2 | 2 | 2 | 0 | 0 | [illegible] |
| C6KT98 | C6KT98 | 2 | 2 | 2 | | PFF1270c | tr\|C6KT98\|C6KT98_PLAF7 Uncharacterized protein OS=Plasmo | 1 | 2 | 2 | 2 | 0 | 0 | [illegible] |
| Q8IHU2 | Q8IHU2 | 3 | 3 | 3 | | PF11_043 | tr\|Q8IHU2\|Q8IHU2_PLAF7 Uncharacterized protein OS=Plasm | 1 | 3 | 3 | 3 | 0 | 0 | [illegible] |
| Q8I540 | Q8I540 | 2 | 2 | 2 | | PFL1855w | tr\|Q8I540\|Q8I540_PLAF7 Pre-mRNA-splicing factor CWC22, pu | 1 | 2 | 2 | 2 | 1 | 0 | [illegible] |
| Q8I5T0 | Q8I5T0 | 2 | 2 | 2 | | PFL0605c | tr\|Q8I5T0\|Q8I5T0_PLAF7 Uncharacterized protein OS=Plasmo | 1 | 2 | 2 | 2 | 1 | 0 | [illegible] |
| A0A144A1 | A0A144A1 | 2 | 2 | 2 | | | tr\|A0A144A1B4\|A0A144A1B4_PLAF7 Dynein heavy chain, puta | 1 | 2 | 2 | 2 | 1 | 1 | [illegible] |
| C0H4N5 | C0H4N5 | 2 | 2 | 2 | | MAL7P1.8 | tr\|C0H4N5\|C0H4N5_PLAF7 Dynein heavy chain, putative OS=P | 1 | 2 | 2 | 2 | 1 | 1 | [illegible] |
| C0H520 | C0H520 | 3 | 3 | 3 | | PFI0295c | tr\|C0H520\|C0H520_PLAF7 Exoribonuclease II OS=Plasmodium | 1 | 3 | 3 | 3 | 1 | 0 | [illegible] |
| C0H5G2 | C0H5G2 | 2 | 1 | 1 | | Rab11b | tr\|C0H5G2\|C0H5G2_PLAF7 Ras-related protein Rab-11B OS=Pl | 1 | 2 | 1 | 1 | 2 | 2 | [illegible] |
| C6KSQ7 | C6KSQ7 | 2 | 2 | 2 | Elongatio | PFF0290w | tr\|C6KSQ7\|C6KSQ7_PLAF7 Elongation of fatty acids protein O | 1 | 2 | 2 | 2 | 2 | 2 | [illegible] |
| O96287 | O96287 | 3 | 1 | 0 | | MC-2TM | tr\|O96287\|O96287_PLAF7 Pfmc-2TM Maurers cleft two transm | 1 | 3 | 1 | 0 | 1 | 1 | [illegible] |
| O97305 | O97305 | 2 | 2 | 2 | | PFC1055w | tr\|O97305\|O97305_PLAF7 BSD domain-containing protein OS= | 1 | 2 | 2 | 2 | 0 | 0 | [illegible] |
| Q8I1X9 | Q8I1X9 | 2 | 2 | 2 | | PFD0245c | tr\|Q8I1X9\|Q8I1X9_PLAF7 ATP-dependent RNA helicase DDX5 | 1 | 2 | 2 | 2 | 0 | 0 | [illegible] |
| Q8I434 | Q8I434 | 2 | 2 | 2 | | PFE0335w | tr\|Q8I434\|Q8I434_PLAF7 Transcription factor 25, putative OS= | 1 | 2 | 2 | 2 | 1 | 1 | [illegible] |
| Q8IC13 | Q8IC13 | 2 | 2 | 2 | | PF07_002 | tr\|Q8IC13\|Q8IC13_PLAF7 Mediator of RNA polymerase II trans | 1 | 2 | 2 | 2 | 0 | 0 | [illegible] |
| Q8IDF6 | Q8IDF6 | 2 | 2 | 2 | Adenylos | Adss | sp\|Q8IDF6\|PURA_PLAF7 Adenylosuccinate synthetase OS=Pla | 1 | 2 | 2 | 2 | 1 | 0 | [illegible] |
| Q8IHN1 | Q8IHN1 | 4 | 1 | 1 | | PF11_050 | tr\|Q8IHN1\|Q8IHN1_PLAF7 Ring-infected erythrocyte surface a | 1 | 4 | 1 | 1 | 2 | 3 | [illegible] |
| Q8IKL5 | Q8IKL5 | 2 | 2 | 2 | | PF14_058 | tr\|Q8IKL5\|Q8IKL5_PLAF7 Valine--tRNA ligase, putative OS=Pla | 1 | 2 | 2 | 2 | 0 | 0 | [illegible] |
| Q8IKN5 | Q8IKN5 | 2 | 2 | 2 | | PF14_056 | tr\|Q8IKN5\|Q8IKN5_PLAF7 Uncharacterized protein OS=Plasm | 1 | 2 | 2 | 2 | 0 | 0 | [illegible] |
| Q8IK20 | Q8IK20 | 2 | 2 | 2 | | PF10_002 | tr\|Q8IK20\|Q8IK20_PLAF7 Exported lipase 2 OS=Plasmodium f | 1 | 2 | 2 | 2 | 1 | 1 | [illegible] |
| O96233 | O96233 | 2 | 2 | 2 | | PFB0700c | tr\|O96233\|O96233_PLAF7 Uncharacterized protein OS=Plasmo | 1 | 2 | 2 | 2 | 0 | 0 | [illegible] |
| O96163 | O96163 | 3 | 3 | 3 | | SERA-7 | tr\|O96163\|O96163_PLAF7 Serine repeat antigen 7 OS=Plasmo | 1 | 3 | 3 | 3 | 2 | 2 | [illegible] |
| Q8IB46 | Q8IB46 | 2 | 2 | 2 | | PF08_004 | tr\|Q8IB46\|Q8IB46_PLAF7 Uncharacterized protein OS=Plasmo | 1 | 2 | 2 | 2 | 0 | 0 | [illegible] |
| Q8I5G2 | Q8I5G2 | 2 | 2 | 2 | | PFL1230w | tr\|Q8I5G2\|Q8I5G2_PLAF7 Small subunit rRNA processing facto | 1 | 2 | 2 | 2 | 0 | 0 | [illegible] |
| Q8I5H0 | Q8I5H0 | 4 | 4 | 4 | | PFL1190c | tr\|Q8I5H0\|Q8I5H0_PLAF7 Uncharacterized protein OS=Plasmo | 1 | 4 | 4 | 4 | 0 | 0 | [illegible] |
| Q8IBW2 | Q8IBW2 | 2 | 2 | 2 | | MAL7P1.6 | tr\|Q8IBW2\|Q8IBW2_PLAF7 Uncharacterized protein OS=Plasm | 1 | 2 | 2 | 2 | 1 | 1 | [illegible] |
| C0H5M6 | C0H5M6 | 2 | 2 | 2 | | MAL13P1. | tr\|C0H5M6\|C0H5M6_PLAF7 Uncharacterized protein OS=Plasm | 1 | 2 | 2 | 2 | 0 | 0 | [illegible] |
| O97224 | O97224 | 2 | 2 | 2 | Palmitoyl | PFC0160w | tr\|O97224\|O97224_PLAF7 Palmitoyltransferase OS=Plasmodiu | 1 | 2 | 2 | 2 | 1 | 1 | [illegible] |
| Q8I2T0 | Q8I2T0 | 2 | 2 | 2 | | PFI1100w | tr\|Q8I2T0\|Q8I2T0_PLAF7 Para-aminobenzoic acid synthetase | 1 | 2 | 2 | 2 | 0 | 1 | [illegible] |
| Q8I518 | Q8I518 | 2 | 2 | 2 | | PFL1975c | tr\|Q8I518\|Q8I518_PLAF7 Polyadenylation factor subunit 2, pu | 1 | 2 | 2 | 2 | 0 | 0 | [illegible] |
| Q8I6U5 | Q8I6U5 | 6 | 1 | 1 | | PF11_016 | tr\|Q8I6U5\|Q8I6U5_PLAF7 Cysteine proteinase falcipain 2b OS | 1 | 6 | 1 | 1 | 5 | 5 | [illegible] |
| Q8IAL3 | Q8IAL3 | 2 | 2 | 2 | | PF08_013 | tr\|Q8IAL3\|Q8IAL3_PLAF7 Nucleolar protein 10, putative OS=P | 1 | 2 | 2 | 2 | 0 | 0 | [illegible] |
| Q8IIK2 | Q8IIK2 | 2 | 2 | 2 | | PF11_017 | tr\|Q8IIK2\|Q8IIK2_PLAF7 Guanine nucleotide-exchange factor | 1 | 2 | 2 | 2 | 0 | 1 | [illegible] |
| Q8IIK5 | Q8IIK5 | 2 | 2 | 2 | | RON4 | tr\|Q8IIK5\|Q8IIK5_PLAF7 Rhoptry neck protein 4 OS=Plasmodiu | 1 | 2 | 2 | 2 | 1 | 1 | [illegible] |
| Q8IJR9 | Q8IJR9 | 2 | 2 | 2 | | PF10_012 | tr\|Q8IJR9\|Q8IJR9_PLAF7 GMP synthase [glutamine-hydrolyzin | 1 | 2 | 2 | 2 | 2 | 2 | [illegible] |
| Q8IK84 | Q8IK84 | 2 | 2 | 2 | | PCRMP 4 | tr\|Q8IK84\|Q8IK84_PLAF7 Cysteine repeat modular protein 4 C | 1 | 2 | 2 | 2 | 0 | 1 | [illegible] |
| Q8IL45 | Q8IL45 | 2 | 2 | 2 | | TREP | tr\|Q8IL45\|Q8IL45_PLAF7 TRAP-like protein OS=Plasmodium f | 1 | 2 | 2 | 2 | 0 | 0 | [illegible] |
| Q8ILB6 | Q8ILB6 | 2 | 2 | 2 | | PF14_032 | tr\|Q8ILB6\|Q8ILB6_PLAF7 Mitochondrial acidic protein MAM33 | 1 | 2 | 2 | 2 | 1 | 1 | [illegible] |
| Q8ILF2 | Q8ILF2 | 3 | 3 | 3 | | PF14_029 | tr\|Q8ILF2\|Q8ILF2_PLAF7 Large subunit GTPase 1, putative OS= | 1 | 3 | 3 | 3 | 0 | 1 | [illegible] |
| A0A5K1K8 | A0A5K1K8 | 2 | 2 | 2 | | | tr\|A0A5K1K8G8\|A0A5K1K8G8_PLAF7 Vacuolar protein sorting- | 1 | 2 | 2 | 2 | 2 | 2 | [illegible] |
| A0A5K1K9 | A0A5K1K9 | 3 | 3 | 3 | | | tr\|A0A5K1K999\|A0A5K1K999_PLAF7 Kinetochore protein SPC | 1 | 3 | 3 | 3 | 1 | 1 | [illegible] |
| Q8I1Q1 | Q8I1Q1 | 2 | 2 | 2 | | PFD0900w | tr\|Q8I1Q1\|Q8I1Q1_PLAF7 Regulator of chromosome condens | 1 | 2 | 2 | 2 | 0 | 0 | [illegible] |
| Q8I379 | Q8I379 | 2 | 2 | 2 | | PFI0345w | tr\|Q8I379\|Q8I379_PLAF7 GTPase-activating protein, putative | 1 | 2 | 2 | 2 | 0 | 0 | [illegible] |
| Q8I3R7 | Q8I3R7 | 3 | 3 | 3 | | PFE0965c | tr\|Q8I3R7\|Q8I3R7_PLAF7 V-type proton ATPase proteolipid su | 1 | 3 | 3 | 3 | 3 | 2 | [illegible] |
| Q8IDT3 | Q8IDT3 | 2 | 2 | 2 | | PF13_021 | tr\|Q8IDT3\|Q8IDT3_PLAF7 Nucleolar complex protein 2, putati | 1 | 2 | 2 | 2 | 0 | 0 | [illegible] |
| Q8IIA5 | Q8IIA5 | 2 | 2 | 2 | | PF11_026 | tr\|Q8IIA5\|Q8IIA5_PLAF7 YTBD domain-containing protein, pu | 1 | 2 | 2 | 2 | 0 | 0 | [illegible] |
| Q9NFE6 | Q9NFE6 | 2 | 2 | 2 | Eukaryoti | PFC0441c | tr\|Q9NFE6\|Q9NFE6_PLAF7 Eukaryotic translation initiation fa | 1 | 2 | 2 | 2 | 0 | 0 | [illegible] |
| Q8IEN2 | Q8IEN2 | 2 | 2 | 2 | | PF13_004 | tr\|Q8IEN2\|Q8IEN2_PLAF7 40S ribosomal protein S27 OS=Plasn | 1 | 2 | 2 | 2 | 2 | 2 | [illegible] |
