## Supplementary material for "*Plasmodium falciparum* SET10 is a histone H3 lysine K18 methyltransferase that participates in a chromatin modulation network crucial for intraerythrocytic development": Table S3

Table S3. List of interactors of PfSET10

Curation steps:

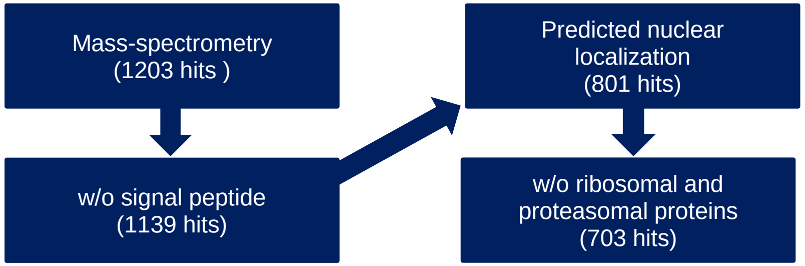

### PfSET10 interactors

| | | | | | | Expression | | | | | | | | Sex | | | GO Terms | | |
|---|---|---|---|---|---|---|---|---|---|---|---|---|---|---|---|---|---|---|---|
| Gene ID | Short name | Product Description | Mol. weight [kDa] | Signal Peptide | # TM Domains | Ring | Early Trophozoite | Late Trophozoite | Schizont | Gametocyte II | Gametocyte V | Ookinete | Peak expression | male gametocyte | female gametocyte | Sex specificity | Curated GO Functions (Molecular) | Curated GO Processes (Biological) | Curated GO Components (Cellular) |
| PF3D7_1233600 | AARP1 | asparagine and aspartate rich protein 1 | 646.33 | no | 0 | 50.78 | 15.99 | 24.42 | 3.49 | 10.84 | 46.00 | 181.42 | O | 125.79 | 43.18 | M | zinc ion binding | N/A | plasma membrane |
| PF3D7_0613800 | ApiAP2 | AP2 domain transcription factor, putative | 485.58 | no | 0 | 9.78 | 1.34 | 89.95 | 46.14 | 11.78 | 17.07 | 97.31 | O | 85.41 | 6.69 | M | sequence-specific DNA binding, DNA-binding transcription factor activity, DNA binding | response to xenobiotic stimulus, regulation of transcription, DNA-templated | nucleus |
| PF3D7_1468100 | MORC | MORC family protein | 295.83 | no | 0 | 170.77 | 129.57 | 91.30 | 24.26 | 7.88 | 27.78 | 110.92 | R | 155.68 | 84.96 | M | protein binding | regulation of cell cycle | nucleus |
| PF3D7_0704600 | UT | HECT-type E3 ubiquitin ligase UT | 460.41 | no | 0 | 134.30 | 57.58 | 41.68 | 13.08 | 14.52 | 61.20 | 373.27 | O | 561.82 | 158.74 | M | ubiquitin-protein transferase activity | response to drug | endoplasmic reticulum, Golgi apparatus |
| PF3D7_0910200 | N/A | conserved Plasmodium protein, unknown function | 368.34 | no | 0 | 18.62 | 6.72 | 13.52 | 4.05 | 3.60 | 11.87 | 40.91 | O | 55.00 | 33.69 | M | N/A | N/A | nucleus |
| PF3D7_1008100 | PHD1 | PHD finger protein PHD1 | 453.00 | no | 0 | 86.17 | 53.30 | 21.26 | 8.32 | 3.22 | 17.25 | 68.53 | R | 35.99 | 17.39 | M | methylated histone binding, metal ion binding | entry into host, histone H3-K9 acetylation, histone H3-K4 trimethylation | histone acetyltransferase complex, nucleus |
| PF3D7_0317300 | NUP335 | nucleoporin NUP335, putative | 402.95 | no | 0 | 53.47 | 39.23 | 25.60 | 8.19 | 6.84 | 23.42 | 78.67 | O | 166.65 | 27.89 | M | DNA-binding transcription factor activity, protein binding, sequence-specific DNA binding | protein autoprocessing, positive regulation of transcription, DNA-templated | nucleus, cytoplasm, endoplasmic reticulum membrane, nuclear periphery, membrane |
| PF3D7_1221000 | SET10 | histone-lysine N-methyltransferase, H3 lysine-4 specific | 271.07 | no | 0 | 5.25 | 3.05 | 21.93 | 5.72 | 2.16 | 12.63 | 50.60 | O | 9.49 | 12.36 | 0 | histone methyltransferase activity (H3-K4 specific), metal ion binding | chromatin remodelling;regulation of transcription | nuclear periphery, nucleus |
| PF3D7_0915400 | PFK9 | ATP-dependent 6-phosphofructokinase | 159.45 | no | 0 | 431.40 | 123.27 | 115.68 | 12.05 | 4.03 | 6.74 | 60.44 | R | 112.38 | 18.75 | M | 6-phosphofructokinase activity; ATP binding, phosphofructokinase activity, metal ion binding, diphosphate-fructose-6-phosphate 1-phosphotransferase activity | response to glucose, response to xenobiotic stimulus, glycolytic process, fructose 6-phosphate metabolic process | cytoplasm, cytosol, 6-phosphofructokinase complex |
| PF3D7_0405400 | PRPF8 | pre-mRNA-processing-splicing factor 8, putative | 366.39 | no | 0 | 156.78 | 102.89 | 99.78 | 16.79 | 6.56 | 15.84 | 402.18 | O | 85.17 | 40.86 | M | U1 snRNA binding, U6 snRNA binding, U2 snRNA binding, U5 snRNA binding, U1 snRNA binding, U6 snRNA binding, pre-mRNA intronic binding | spliceosomal tri-snRNP complex assembly, RNA splicing, via transesterification reactions, RNA splicing | nucleus, cytoplasm, spliceosomal complex, U5 snRNP, apicoplast |
| PF3D7_0917900 | BIP | heat shock protein 70 | 72.39 | yes | 0 | 653.49 | 329.31 | 1163.63 | 248.63 | 22.32 | 24.05 | 457.83 | T | 195.73 | 166.21 | M | ATP binding, ATP hydrolysis activity; heat shock protein binding; protein folding chaperone; unfolded protein binding; misfolded protein binding | response to unfolded protein; ubiquitin-dependent ERAD pathway; endoplasmic reticulum unfolded protein response; cellular response to unfolded protein; chaperone cofactor-dependent protein refolding | food vacuole; nucleus; membrane; endoplasmic reticulum; extracellular vesicle |
| PF3D7_0317700 | N/A | CPSF (cleavage and polyadenylation specific factor), subunit A, putative | 338.53 | no | 0 | 35.07 | 13.47 | 13.36 | 2.84 | 5.65 | 38.85 | 85.22 | O | 166.90 | 117.96 | M | damaged DNA binding; nucleic acid binding; protein binding | nucleotide-excision repair; DNA repair; proteasome-mediated ubiquitin-dependent protein catabolic process | nucleus; cullin-RING ubiquitin ligase complex |
| PF3D7_0609000 | NUP637 | nucleoporin NUP637, putative | 720.51 | no | 0 | 86.14 | 62.26 | 38.45 | 12.58 | 8.74 | 47.35 | 51.50 | R | 79.23 | 33.93 | M | protein binding | nucleocytoplasmic transport | nuclear periphery; nucleus |
| PF3D7_1239200 | ApiAP2 | AP2 domain transcription factor, putative | 299.38 | no | 0 | 16650.00 | 28095.00 | 61.7 | 45480.00 | 34029.00 | 28.91 | 223.22 | S | 69.00 | 39.57 | M | DNA binding, DNA-binding transcription factor activity | regulation of transcription, DNA-templated, | heterochromatin, nucleus |
| PF3D7_0818900 | HSP70 | heat shock protein 70 | 73.91 | no | 0 | 3426.39 | 3329.13 | 3055.78 | 1342.83 | 388.88 | 170.78 | 1845.88 | R | 1425.79 | 301.35 | M | ATP binding; ATP hydrolysis activity; heat shock protein binding; phosphatidylinositol-3-phosphate binding; protein folding chaperone; unfolded protein binding; misfolded protein binding | response to unfolded protein; response to heat; cellular response to unfolded protein; | nucleus; cytoplasm; cytosol; cell surface; food vacuole; extracellular vesicle |
| PF3D7_0605800 | RAD50 | DNA repair protein RAD50, putative | 267.95 | no | 0 | 20.12 | 6.26 | 12.65 | 2.62 | 1.82 | 9.21 | 35.80 | O | 19.13 | 14.92 | M | G-quadruplex DNA binding; metal ion binding; single-stranded telomeric DNA binding adenylate kinase activity; double-stranded telomeric DNA binding; single-stranded DNA endodeoxyribonuclease activity | telomere maintenance via recombination; double-strand break repair; telomere maintenance via telomerase; DNA duplex unwinding; chromosome organization involved in meiotic cell cycle | condensed nuclear chromosome; nucleus; Mre11 complex; site of double-strand break |
| PF3D7_0629700 | SET1 | SET domain protein, putative | 796.04 | no | 0 | 45.67 | 31.36 | 25.30 | 4.30 | 3.13 | 23.45 | 102.36 | O | 82.65 | 38.44 | M | histone methyltransferase activity (H3-K4 specific) | positive regulation of transcription, DNA-templated, histone H3-K4 methylation | nucleus;histone methyltransferase complex |
| PF3D7_0922800 | N/A | conserved Plasmodium protein, unknown function | 516.45 | no | 2 | 10.86 | 3.03 | 7.97 | 0.99 | 3.65 | 34.85 | 35.96 | O | 61.71 | 21.16 | M | protein kinase binding | signal transduction | integral component of membrane |
| PF3D7_1252100 | RON3 | rhoptry neck protein 3 | 263.15 | yes | 3 | 5.87 | 0.67 | 56.25 | 101.19 | 3.59 | 1.98 | 98.22 | S | 1.39 | 0.38 | 0 | RNA binding, host cell surface binding; protein binding | entry into host;symbiont intracellular protein transport in host | rhoptry; extracellular vesicle |
| PF3D7_0628100 | HECT1 | HECT domain-containing protein 1, putative | 1206.00 | no | 2 | 8.24 | 0.65 | 21.71 | 36.80 | 2.47 | 28.22 | 948.15 | O | 8.36 | 58.58 | F | ubiquitin protein ligase activity, protein binding, ubiquitin-protein transferase activity | N/A | integral component of membrane |
| PF3D7_1219100 | CHC | clathrin heavy chain, putative | 232.95 | no | 0 | 39.47 | 9.25 | 27.53 | 6.29 | 1.92 | 28.74 | 51.03 | O | 14.72 | 23.77 | F | clathrin light chain binding, structural molecule activity, protein binding | receptor-mediated endocytosis, intracellular protein transport, vesicle-mediated transport | clathrin complex; extracellular vesicle, clathrin coat of trans-Golgi network vesicle |
| PF3D7_1472200 | HDA1 | histone deacetylase, putative | 268.92 | no | 3 | 127.77 | 27.82 | 14.49 | 4.13 | 5.76 | 46.66 | 129.46 | O | 122.03 | 38.33 | M | NAD-dependent histone deacetylase activity (H3-K14 specific) | chromatin remodelling;regulation of transcription | nucleus, integral component of membrane |
| PF3D7_1303800 | N/A | conserved Plasmodium protein, unknown function | 1111.80 | no | 2 | 5.85 | 3.71 | 11.23 | 1.84 | 2.07 | 33.23 | 81.14 | O | 109.15 | 24.81 | M | N/A | N/A | nucleus, integral component of membrane |
| PF3D7_0516800 | ApiAP2 | AP2 domain transcription factor AP2-O2, putative | 276.42 | no | 0 | 12.31 | 1.70 | 43.02 | 9.63 | 6.21 | 80.08 | 288.62 | O | 122.19 | 204.96 | F | sequence-specific DNA binding, DNA-binding transcription factor activity, DNA binding | regulation of transcription, DNA-templated | heterochromatin, nucleus |
| PF3D7_0929400 | RhopH2 | high molecular weight rhoptry protein 2 | 162.66 | yes | 0 | 14.32 | 1.05 | 231.19 | 161.35 | 4.05 | 13.19 | 291.69 | O | 15.10 | 33.85 | F | RNA binding; protein binding; protein-containing complex binding | entry into host | extracellular vesicle; host cell membrane; host cell cytoplasm; rhoptry; symbiont-containing vacuole membrane |
| PF3D7_0626800 | PyrK | pyruvate kinase | 55.66 | no | 0 | 1031.61 | 455.47 | 887.62 | 126.79 | 23.95 | 11.73 | 120.11 | R | 319.67 | 32.71 | M | RNA binding; magnesium ion binding; catalytic activity; pyruvate kinase activity; potassium ion binding | glycolytic process; protein homotetramerization; | cytoplasm |
| PF3D7_1322300 | N/A | BRCT domain-containing protein, putative | 178.35 | no | 0 | 5.93 | 3.97 | 8.03 | 1.79 | 4.15 | 27.02 | 75.21 | O | 508.88 | 10.67 | M | N/A | N/A | nucleus |
| PF3D7_1434500 | N/A | dynein-related AAA-type ATPase, putative | 971.00 | no | 9 | 46.73 | 45.83 | 7.08 | 0.54 | 1.98 | 20.47 | 18.71 | R | 64.58 | 16.90 | M | ATP binding; ATP hydrolysis activity | rRNA processing; ribosomal large subunit assembly | nucleus; membrane; integral component of membrane; preribosome, large subunit precursor |
| PF3D7_1308400 | N/A | conserved Plasmodium protein, unknown function | 737.69 | no | 3 | 15.92 | 18.24 | 15.04 | 3.71 | 2.80 | 16.91 | 31.38 | O | 30.75 | 13.91 | M | N/A | N/A | N/A |
| PF3D7_0524000 | KASbeta | karyopherin beta | 127.35 | no | 0 | 347.01 | 93.65 | 99.26 | 14.61 | 7.36 | 26.71 | 97.28 | R | 73.58 | 16.93 | M | RNA binding; nuclear localization sequence binding; protein domain specific binding; protein heterodimerization activity; nuclear import signal receptor activity | protein import into nucleus; NLS-bearing protein import into nucleus; nucleocytoplasmic transport | nucleus; nuclear pore; cytoplasm; food vacuole; RNA N6-methyladenosine methyltransferase complex |
| PF3D7_1416100 | SEY1 | Protein SEY1 homolog | 110.61 | no | 2 | 31.54 | 41334.00 | 45306.00 | 11810.00 | 13881.00 | 45337.00 | 19603.00 | GC V | 25.87 | 24.32 | 0 | GTPase activity;GTP binding | endoplasmic reticulum organiziation;endoplasmic reticulum membrane fusion | cytoplasm;endoplasmic reticulum membrane |
| PF3D7_0726500 | N/A | ubiquitin carboxyl-terminal hydrolase, putative | 373.16 | no | 0 | 76.66 | 27.76 | 41.70 | 8.34 | 12.86 | 90.14 | 137.99 | O | 112.68 | 49.59 | M | cysteine-type endopeptidase activity; thiol-dependent deubiquitinase; protein binding | protein deubiquitination; ubiquitin-dependent protein catabolic process | nucleus; cytosol |
| PF3D7_1433400 | PHD2 | PHD finger protein PHD2, putative | 682.45 | no | 12 | 16.22 | 42.77 | 44.79 | 13.13 | 3.44 | 35.46 | 76.44 | O | 157.85 | 59.14 | M | metal ion binding | N/A | histone acetyltransferase complex; nucleus; membrane; integral component of membrane |
| PF3D7_1128000 | N/A | Uncharacterized protein | 211.53 | no | 0 | 55.96 | 35.29 | 33.4 | 21276.00 | 45506.00 | 22.75 | 105.73 | GC II | 17.89 | 16.75 | 0 | N/A | N/A | N/A |
| PF3D7_0422500 | BRR2 | pre-mRNA-splicing helicase BRR2, putative | 337.90 | no | 0 | 105.44 | 72.81 | 27.09 | 4.35 | 8.08 | 18.99 | 136.41 | O | 92.14 | 36.50 | M | RNA helicase activity; nucleic acid binding; mRNA binding | regulation of translation; RNA splicing, via transesterification reactions | spliceosomal complex; cytoplasmic stress granule;nucleus |
| PF3D7_1003800 | EFTUD2 | U5 small nuclear ribonucleoprotein component, putative | 142.94 | no | 0 | 77.87 | 48.98 | 25.68 | 2.72 | 8.20 | 15.49 | 24.12 | R | 81.59 | 18.30 | M | GTPase activity; U5 snRNA binding, RNA binding GTP binding; | RNA splicing, via transesterification reactions; mRNA splicing, via spliceosome | U4/U6 x U5 tri-snRNP complex; U2-type catalytic step 2 spliceosome; ribonucleoprotein complex |
| PF3D7_0318200 | RPB1 | DNA-directed RNA polymerase II subunit RPB1 | 278.67 | no | 0 | 227.11 | 99.19 | 104.71 | 27.79 | 14.63 | 10.43 | 78.93 | R | 39.53 | 34.43 | 0 | DNA-directed 5'-3' RNA polymerase activity; protein binding; metal ion binding | transcription, DNA-templated; transcription by RNA polymerase II | transcription by RNA polymerase II; transcription, DNA-templated |
| PF3D7_1460500 | N/A | conserved Plasmodium protein, unknown function | 192.01 | no | 0 | 4.68 | 2.11 | 7.58 | 1.74 | 2.35 | 14.39 | 14.84 | O | 36.53 | 2.71 | M | N/A | N/A | nucleus |
| PF3D7_1410300 | N/A | WD repeat-containing protein, putative | 526.73 | no | 0 | 26.36 | 4.05 | 9.03 | 1.22 | 3.16 | 22.31 | 15.23 | R | 128.48 | 24.03 | M | U3 snoRNA binding | endonucleolytic cleavage in 5'-ETS of tricistronic rRNA transcript (SSU-rRNA, 5.8S rRNA, LSU-rRNA) | nucleus;cytoplasm |
| PF3D7_1246200 | ActI | Actin-1 | 41.87 | no | 0 | 55.25 | 77.34 | 1088.75 | 970.19 | 191.01 | 748.05 | 2474.74 | O | 240.73 | 2466.7 | F | protein binding | structural complex | cytoskeleton;cytoplasm;nucleus |
| PF3D7_0317500 | EG5 | kinesin-5 | 191.78 | no | 0 | 42.54 | 25.61 | 21.30 | 5.23 | 5.77 | 17.14 | 44.40 | O | 110.89 | 21.40 | M | ATP hydrolysis activity, microtubule motor activity; microtubule binding, plus-end-directed microtubule motor activity | microtubule-based movement, response to xenobiotic stimulus | nucleus, kinesin complex |
| PF3D7_1308200 | cpsSII | carbamoyl phosphate synthetase | 273.43 | no | 0 | 131.81 | 92.56 | 20.97 | 1.88 | 1.73 | 7.79 | 30.79 | R | 21.64 | 9.93 | M | carbamoyl-phosphate synthase (glutamine-hydrolyzing) activity, ATP binding. | de novo' pyrimidine nucleobase biosynthetic process, nitrogen compound metabolic process, glutamine metabolic process, glutamate catabolic process | nucleus, cytoplasm |
| PF3D7_1222300 | GRP94 | endoplasmin, putative | 95.02 | yes | 0 | 60.07 | 86.81 | 168.73 | 31.49 | 18.41 | 167.27 | 960.16 | O | 172.76 | 728.94 | F | RNA binding ATP binding, unfolded protein binding | protein folding, response to heat | endoplasmic reticulum;cell surface;food vacuole;perinuclear region of cytoplasm;extracellular vesicle |
| PF3D7_1012700 | NIF4 | NLI interacting factor-like phosphatase, putative | 171.01 | no | 0 | 16.21 | 6.12 | 11.28 | 2.63 | 2.17 | 7.39 | 29.05 | O | 4.24 | 13.92 | F | RNA polymerase II CTD heptapeptide repeat phosphatase activity | dephosphorylation of RNA polymerase II C-terminal domain | nucleus |
| PF3D7_0715200 | N/A | conserved Plasmodium protein, unknown function | 258.40 | yes | 1 | 5.20 | 0.51 | 19.11 | 5.35 | 11.36 | 157.89 | 1393.58 | O | 117.85 | 635.92 | F | N/A | N/A | apicoplast |
| PF3D7_0806800 | N/A | V-type proton ATPase subunit a, putative | 123.00 | no | 6 | 301.21 | 111.23 | 95.72 | 38.98 | 9.05 | 28.61 | 411.22 | O | 137.31 | 54.45 | M | ATPase binding; proton-transporting ATPase activity, rotational mechanism, proton-transporting ATP synthase activity, rotational mechanism, hydrolase activity, proton transmembrane transporter activity | vacuolar acidification, proton transmembrane transport | membrane, integral component of membrane, proton-transporting two-sector ATPase complex, vacuolar proton-transporting V-type ATPase complex, proton-transporting V-type ATPase, V0 domain |
| PF3D7_0619400 | N/A | cell division cycle protein 48 homologue, putative | 92.39 | no | 3 | 60.44 | 96.67 | 256.32 | 91.56 | 19.79 | 18.16 | 284.25 | O | 199.42 | 68.50 | M | ATP hydrolysis activity; polyubiquitin modification-dependent protein binding, hydrolase activity, RNA binding | regulation of cell cycle process, macroautophagy, ubiquitin-dependent ERAD pathway, retrograde protein transport, ER to cytosol; mitotic spindle disassembly | nucleus; cytoplasm;endoplasmic reticulum, cytosol |
| PF3D7_1369400 | N/A | conserved Plasmodium protein, unknown function | 346.81 | no | 2 | 3.46 | 1.66 | 3.61 | 0.65 | 2.47 | 15.03 | 8.04 | G V | 10.66 | 6.96 | 0 | N/A | N/A | integral component of membrane |
| PF3D7_0826100 | HEUL | HECT-like E3 ubiquitin ligase, putative | 1005.10 | no | 2 | 99.48 | 75.67 | 24.99 | 3.25 | 6.02 | 43.36 | 68.66 | R | 119.77 | 52.32 | M | ubiquitin protein ligase activity; ubiquitin-protein transferase activity | proteasome-mediated ubiquitin-dependent protein catabolic process; positive regulation of protein catabolic process; protein ubiquitination; ubiquitin-dependent protein catabolic process | cytoplasm; integral component of membrane |
| PF3D7_0321800 | N/A | WD repeat-containing protein, putative | 322.24 | no | 0 | 77.86 | 29.97 | 17.73 | 10.61 | 2.66 | 15.76 | 43.51 | R | 184.29 | 15.87 | M | protein binding | RNA splicing | nucleus; precatalytic spliceosome |
| PF3D7_1357100 | N/A | elongation factor 1-alpha | 48.96 | no | 0 | 119.25 | 251.23 | 99.01 | 20.40 | 18.26 | 13.44 | 5.82 | T | 14.96 | 20.46 | F | GTPase activity; translation elongation factor activity; RNA binding GTP binding | translation; translational elongation | nucleus; mitochondrion; eukaryotic translation elongation factor 1 complex; extracellular vesicle |
| PF3D7_0410900 | AKIT7 | apicomplexan kinetochore protein 7, putative | 117.53 | no | 0 | 22.49 | 12.17 | 8.33 | 1.00 | 3.33 | 22.24 | 189.56 | O | 133.30 | 46.13 | M | N/A | N/A | kinetochore; nucleus |
| PF3D7_1451100 | eEF2 | elongation factor 2 | 93.52 | no | 0 | 1381.80 | 1093.50 | 653.01 | 116.96 | 38.30 | 31.65 | 275.30 | R | 123.94 | 185.42 | F | GTP binding; GTPase activity; translation elongation factor activity; RNA binding | translational elongation; response to xenobiotic stimulus | nucleus; ribonucleoprotein complex |
| PF3D7_0827800 | SET3 | SET domain protein, putative | 283.55 | no | 0 | 64.54 | 10.91 | 23.63 | 8.84 | 6.71 | 49.68 | 167.97 | O | 119.89 | 325.00 | F | protein binding; histone-lysine N-methyltransferase activity | chromatin remodelling;regulation of transcription | nuclear periphery; nucleus |
| PF3D7_0710200 | N/A | conserved Plasmodium protein, unknown function | 350.70 | no | 0 | 8.14 | 11.06 | 10.76 | 10.36 | 4.34 | 56.98 | 330.97 | O | 3852.79 | 1172.20 | M | N/A | N/A | nucleus |
| PF3D7_1317100 | MCM4 | DNA replication licensing factor MCM4 | 115.22 | no | 0 | 10.44 | 6.05 | 148.92 | 45.16 | 5.69 | 21.55 | 67.84 | T | 785.94 | 69.19 | M | DNA replication origin binding; single-stranded DNA binding; ATP binding | double-strand break repair via break-induced replication; DNA replication; pre-replicative complex assembly involved in nuclear cell cycle DNA replication; DNA unwinding involved in DNA replication; DNA duplex unwinding; mitotic DNA replication initiation | MCM complex; nucleus |
| PF3D7_1462800 | GAPDH | glyceraldehyde-3-phosphate dehydrogenase | 36.64 | no | 0 | 5476.96 | 3601.64 | 8667.43 | 1684.88 | 256.77 | 93.03 | 434.72 | T | 511.84 | 25.12 | M | glyceraldehyde-3-phosphate dehydrogenase (NAD+) (phosphorylating) activity; oxidoreductase activity:acting on the aldehyde or oxo group of donors:NAD or NADP as acceptor:NADP binding | gluconeogenesis; glucose metabolic process; glycolytic process; response to xenobiotic stimulus; entry into host; oxidation-reduction process | cytosol; cell surface; membrane; food vacuole; extracellular vesicle |
| PF3D7_1417800 | MCM2 | DNA replication licensing factor MCM2 | 111.50 | no | 0 | 9.07 | 9.88 | 103.25 | 21.20 | 6.25 | 26.17 | 111.84 | O | 1583.73 | 87.28 | F | DNA binding;DNA helicase activity;protein binding;ATP binding | DNA replication intiation;DNA duplex unwinding;regulation of cell cycle | nucleus;cytoplasm;MCM complex |
| PF3D7_0420000 | N/A | zinc finger protein, putative | 401.14 | no | 0 | 64.52 | 36.23 | 11.67 | 1.96 | 2.55 | 21.96 | 96.31 | O | 83.95 | 39.48 | M | zinc ion binding | N/A | cytoplasm; membrane; nucleus |
| PF3D7_0629800 | N/A | cullin-2, putative | 136.34 | no | 0 | 69.76 | 7.61 | 10.68 | 1.65 | 2.34 | 30.56 | 11.76 | R | 48.74 | 24.90 | M | protein binding; ubiquitin-protein transferase activity; ubiquitin protein ligase binding | G1/S transition of mitotic cell cycle; ubiquitin-dependent protein catabolic process; protein ubiquitination; regulation of cell cycle | cullin-RING ubiquitin ligase complex;nucleus |
| PF3D7_0205900 | RPN1 | 26S proteasome regulatory subunit RPN1, putative | 108.36 | no | 0 | 95.37 | 81.94 | 202.97 | 133.40 | 48.64 | 59.85 | 270.09 | O | 37.03 | 121.60 | F | enzyme regulator activity | proteasome-mediated ubiquitin-dependent protein catabolic process; ubiquitin-dependent protein catabolic process | nucleus;cytoplasm;proteasome complex; proteasome regulatory particle; proteasome storage granule |
| PF3D7_1107800 | ApiAP2 | AP2 domain transcription factor, putative | 206.87 | no | 0 | 39.31 | 93.68 | 44.62 | 35.76 | 10.53 | 72.68 | 200.31 | O | 29.60 | 146.15 | F | sequence-specific DNA binding; DNA-binding transcription factor activity; DNA binding | regulation of transcription, DNA-templated | heterochromatin; nucleus |
| PF3D7_1130700 | SMC1 | structural maintenance of chromosomes protein 1 | 214.67 | no | 0 | 6.89 | 8.26 | 26.27 | 5.38 | 5.10 | 45.26 | 97.20 | O | 196.05 | 131.32 | M | ATP hydrolysis activity;ATP binding; double-stranded DNA binding; protein binding | mitotic sister chromatid segregation; chromosome organization | chromosome;cohesin complex; nuclear mitotic cohesin complex |
| PF3D7_1228800 | N/A | WD repeat-containing protein, putative | 378.91 | no | 1 | 37.43 | 30.41 | 51.02 | 14.24 | 7.36 | 39.98 | 253.23 | O | 225.37 | 160.00 | M | protein binding | nucleosome assembly | CAF-1 complex; nucleus; integral component of membrane |
| PF3D7_1428900 | N/A | conserved Plasmodium protein, unknown function | 172.15 | no | 0 | 8.17 | 4.99 | 23.26 | 5.97 | 4.02 | 15.46 | 55.15 | O | 11.28 | 47.39 | F | N/A | N/A | N/A |
| PF3D7_1218200 | N/A | symplekin domain-containing protein, putative | 256.15 | no | 2 | 30.51 | 9.13 | 8.59 | 2.49 | 2.42 | 17.87 | 31.06 | O | 40.69 | 10.02 | M | N/A | N/A | integral component of membrane |

| | | | | | | | | | | | | | | | | | | | |
|---|---|---|---|---|---|---|---|---|---|---|---|---|---|---|---|---|---|---|---|
| PF3D7_0624600 | ISWI | ISWI chromatin-remodeling complex ATPase | 315.62 | no | 0 | 236.36 | 81.38 | 73.90 | 18.07 | 8.80 | 44.49 | 197.63 | R | 451.10 | 62.63 | M | ATP-dependent activity, acting on DNA; transcription cis-regulatory region binding; DNA binding; DNA helicase activity; helicase activity | positive regulation of transcription, DNA-templated; chromatin remodeling | nucleus |
| PF3D7_1023900 | CHD1 | chromodomain-helicase-DNA-binding protein 1 homolog, putative | 381.25 | no | 0 | 626.71 | 717.52 | 387.05 | 195.77 | 20.21 | 57.73 | 783.10 | O | 575.01 | 126.36 | M | protein binding; DNA binding; DNA helicase activity; ATP binding | N/A | nucleus |
| PF3D7_0606700 | N/A | coatomer alpha subunit, putative | 176.94 | no | 0 | 124.45 | 30.55 | 56.95 | 26.60 | 7.86 | 26.86 | 92.69 | R | 74.66 | 40.43 | M | structural molecule activity; protein binding | endoplasmic reticulum to Golgi vesicle-mediated transport; ? intracellular protein transport; retrograde vesicle-mediated transport, Golgi to endoplasmic reticulum; vesicle-mediated transport | Golgi membrane;membrane coat;COPI vesicle coat |
| PF3D7_1357800 | CCT4 | T-complex protein 1 subunit delta | 57.98 | no | 0 | 328.28 | 188.56 | 112.06 | 31.14 | 9.69 | 11.12 | 36.31 | R | 43.45 | 26.43 | M | unfolded protein binding; ATP binding | protein folding; response to xenobiotic stimulus | nucleus; cytosol; chaperonin-containing T-complex |
| PF3D7_0831700 | HSP70x | heat shock protein 70 | 75.05 | yes | 1 | 2948.21 | 393.22 | 249.55 | 22.30 | 7.27 | 5.50 | 104.84 | R | 56.87 | 2.05 | M | ATP binding, ATP hydrolysis activity, heat shock protein binding, protein folding chaperone, unfolded protein binding, misfolded protein binding | response to unfolded protein; cytoadherence to microvasculature, mediated by symbiont protein; cellular response to unfolded protein;? chaperone cofactor-dependent protein refolding | nucleus, cytoplasm, symbiont-containing vacuole, Maurer's cleft, protein-containing complex, host cell cytosol |
| PF3D7_0302900 | N/A | exportin-1, putative | 147.94 | no | 0 | 70.88 | 14.79 | 42.43 | 10.66 | 8.60 | 14.64 | 23.16 | R | 57.21 | 18.62 | M | nuclear export signal receptor activity | protein export from nucleus, regulation of protein export from nucleus, ribosomal small subunit export from nucleus | ? cytoplasm; nucleus |
| PF3D7_1148000 | N/A | serine/threonine protein kinase, putative | 170.70 | no | 0 | 6.11 | 1.34 | 12.99 | 2.78 | 4.63 | 22.21 | 81.72 | O | 59.36 | 41.96 | M | ATP binding, protein serine/threonine kinase activity, protein kinase activity ? | regulation of chromatin assembly or disassembly, protein phosphorylation, chromosome segregation, peptidyl-serine phosphorylation, intracellular signal transduction | nucleus |
| PF3D7_1234800 | SF3B3 | splicing factor 3B subunit 3, putative | 151.81 | no | 0 | 77.19 | 39.63 | 36.93 | 3.75 | 9.01 | 18.18 | 91.56 | O | 46.12 | 37.97 | M | nucleic acid binding, protein binding | RNA splicing, ? mRNA splicing, via spliceosome, RNA splicing, via transesterification reactions, spliceosomal complex assembly | spliceosomal complex, U2 snRNP nucleus, cytoplasm |
| PF3D7_0935800 | CLAG9 | cytoadherence linked asexual protein 9 | 160.42 | yes | 2 | 10.88 | 0.76 | 119.53 | 73.47 | 0.93 | 3.04 | 68.85 | T | 2.32 | 4.58 | 0 | RNA binding, protein binding, host cell surface receptor binding | cell-cell adhesion, ? cytoadherence to microvasculature, mediated by symbiont protein | rhoptry, apical part of cell |
| PF3D7_1429800 | N/A | coatomer subunit beta, putative | 159.97 | no | 0 | 168.39 | 18.50 | 49.33 | 19.22 | 7.65 | 19.78 | 32.47 | R | 64.54 | 38.52 | M | protein binding | endoplasmic reticulum to Golgi vesicle-mediated transport; | Golgi membrane; membrane coat;COPI vesicle coat |
| PF3D7_0514100 | UvrD | ATP-dependent DNA helicase UvrD | 170.29 | no | 1 | 6.20 | 3.20 | 17.97 | 6.27 | 6.44 | 29.73 | 89.45 | O | 27.77 | 33.96 | F | 3'-5' DNA helicase activity; single-stranded DNA helicase activity; ATP binding; DNA helicase activity; protein binding | recombinational repair, DNA repair | nucleus |
| PF3D7_1468700 | eIF4A | eukaryotic initiation factor 4A | 45.31 | no | 0 | 1715.49 | 1152.94 | 678.76 | 92.91 | 79.47 | 83.48 | 368.49 | R | 299.10 | 214.37 | M | 3'-5' DNA/RNA helicase activity; ATP hydrolysis activity; helicase activity; translation initiation factor activity; nucleic acid binding | cytoplasmic translational initiation; regulation of translational initiation | nucleus; cytoplasm; cytosol; cytoplasmic stress granule; eukaryotic translation initiation factor 4F complex; food vacuole |
| PF3D7_1104200 | SNF2L | chromatin remodeling protein | 167.44 | no | 0 | 185.35 | 54.00 | 35.85 | 6.97 | 3.54 | 23.34 | 164.78 | R | 22.56 | 21.17 | M | ATP binding; DNA binding; helicase activity | regulation of transcription, DNA-templated; nucleosome positioning; chromatin remodeling | nucleus |
| PF3D7_1145400 | DYN1 | dynamin-like protein | 96.37 | no | 0 | 104.32 | 24.52 | 141.95 | 195.44 | 14.85 | 47.22 | 125.00 | S | 70.70 | 413.90 | F | GTPase activity; RNA binding; microtubule binding | mitochondrial fission; organelle fission; | nucleus; cytoplasm; plasma membrane; symbiont-containing vacuole membrane; food vacuole |
| PF3D7_0705400 | MCM7 | DNA replication licensing factor MCM7 | 94.24 | no | 0 | 11.77 | 5.78 | 133.58 | 36.61 | 6.29 | 12.61 | 117.12 | T | 730.03 | 91.54 | M | ATP-dependent activity, acting on DNA; ATP binding; single-stranded DNA binding; DNA replication origin binding; DNA binding | DNA duplex unwinding; DNA strand elongation involved in DNA replication; pre-replicative complex assembly involved in nuclear cell cycle DNA replication; double-strand break repair via break-induced replication; DNA replication | nucleus; cytoplasm; MCM complex |
| PF3D7_1311900 | vapA | V-type proton ATPase catalytic subunit A | 68.58 | no | 0 | 215.18 | 98.82 | 149.85 | 52.77 | 11.05 | 17.98 | 44.13 | R | 125.94 | 38.91 | M | proton-transporting ATPase activity, rotational mechanism; ? ATP binding | proton transmembrane transport; ATP metabolic process; vacuolar acidification | vacuolar proton-transporting V-type ATPase, V1 domain; vacuolar membrane; plasma membrane; food vacuole; proton-transporting V-type ATPase, V1 domain |
| PF3D7_0706000 | N/A | importin-7, putative | 145.51 | no | 0 | 87.79 | 27.74 | 41.24 | 10.56 | 14.46 | 84.34 | 38.58 | R | 77.58 | 44.48 | M | N/A | protein import into nucleus; intracellular protein transport | cytoplasm; cytosol; nuclear envelope; nucleus; nuclear pore |
| PF3D7_1458600 | N/A | zinc finger protein, putative | 139.59 | no | 0 | 67.96 | 8.21 | 7.98 | 1.50 | 3.18 | 13.13 | 7.82 | R | 31.50 | 10.95 | M | protein binding | N/A | nucleus; cytoplasm |
| PF3D7_0608700 | CCT6 | T-complex protein 1 subunit zeta | 61.53 | no | 0 | 86.46 | 57.11 | 45.00 | 7.41 | 8.79 | 20.33 | 190.37 | O | 59.35 | 35.85 | M | unfolded protein binding | protein folding | chaperonin-containing T-complex; nucleus; cytosol |
| PF3D7_1208900 | PPM11 | protein phosphatase PPM11, putative | 166.86 | no | 0 | 10.69 | 16.35 | 63.35 | 21.44 | 14.28 | 51.73 | 81.43 | O | 199.06 | 27.11 | M | phosphoprotein phosphatase activity;phosphatase activity | N/A | nucleus |
| PF3D7_0627800 | ACAS | acetyl-CoA synthetase | 113.88 | no | 0 | 121.57 | 49.14 | 54.06 | 13.24 | 3.01 | 7.58 | 34.99 | R | 23.00 | 17.90 | M | acetate-CoA ligase activity | acetyl-CoA biosynthetic process; generation of precursor metabolites and energy; chromatin organization; response to xenobiotic stimulus; acetyl-CoA biosynthetic process from acetate; histone acetylation | nucleus;cytoplasm |
| PF3D7_0317200 | CRK4 | cdc2-related protein kinase 4 | 182.22 | no | 0 | 10.39 | 5.75 | 73.63 | 29.01 | 3.48 | 8.00 | 243.56 | O | 43.80 | 24.98 | M | protein kinase activity; cyclin-dependent protein serine/threonine kinase activity; ? ATP binding | mitotic cell cycle; DNA replication; protein phosphorylation; peptidyl-serine phosphorylation; regulation of cell cycle | nucleus;cytosol;protein kinase CK2 complex |
| PF3D7_0505500 | MSH6 | DNA mismatch repair protein MSH6, putative | 156.44 | no | 0 | 9.34 | 6.55 | 55.50 | 9.61 | 2.49 | 7.86 | 43.93 | T | 219.05 | 23.24 | M | ATP-dependent activity, acting on DNA; mismatched DNA binding; ? four-way junction DNA binding | meiotic mismatch repair; DNA repair; pyrimidine dimer repair; maintenance of DNA repeat elements; negative regulation of DNA recombination | nucleus; mismatch repair complex; MutSalpha complex |
| PF3D7_1311800 | M1AAP | M1-family alanyl aminopeptidase | 126.06 | no | 1 | 231.49 | 204.48 | 77.95 | 6.48 | 2.65 | 8.23 | 26.03 | R | 18.74 | 10.06 | M | aminopeptidase activity; metallopeptidase activity;zinc ion binding | proteolysis; response to xenobiotic stimulus | cytoplasm; cytosol; cell surface; food vacuole; extracellular vesicle; integral component of membrane; microneme; food vacuole |
| PF3D7_0922200 | SAMS | S-adenosylmethionine synthetase | 44.84 | no | 0 | 315.15 | 102.02 | 890.99 | 258.09 | 11.44 | 83.26 | 194.91 | T | 44.53 | 171.00 | F | methionine adenosyltransferase activity | protein modification | nucleus;cytoplasm |
| PF3D7_0500800 | MESA | mature parasite-infected erythrocyte surface antigen | 168.29 | no | 0 | 37.84 | 309.37 | 162.54 | 1.16 | 12.27 | 10.68 | 182.49 | T | 582.57 | 11.74 | M | protein binding | N/A | Maurer's cleft; infected host cell surface knob ?host cell plasma membrane; nucleus |
| PF3D7_1248900 | RPT6 | 26S protease regulatory subunit 8, putative | 49.55 | no | 0 | 80.34 | 66.05 | 178.09 | 57.21 | 26.44 | 35.09 | 140.92 | T | 79.96 | 175.91 | F | ATP hydrolysis activity; hydrolase activity; peptidase activity; transcription coregulator activity | proteasome regulatory particle assembly; proteasome-mediated ubiquitin-dependent protein catabolic process; | proteasome complex; nucleus; cytoplasm; ? proteasome regulatory particle, base subcomplex |
| PF3D7_0513200 | N/A | CPSF domain-containing protein, putative | 332.13 | no | 0 | 76.27 | 13.90 | 11.28 | 4.06 | 2.27 | 18.49 | 76.44 | O | 64.45 | 16.96 | M | polynucleotide adenylyltransferase activity;nucleic acid binding; protein binding | histone mRNA catabolic process; sno(s)RNA polyadenylation | nucleus; nucleolus; TRAMP complex |
| PF3D7_0303000 | NSF | N-ethylmaleimide-sensitive fusion protein | 89.13 | no | 0 | 13.10 | 3.04 | 8.97 | 1.09 | 2.59 | 4.81 | 20.57 | O | 11.59 | 33.59 | F | ATP hydrolysis activity; metal ion binding; ATP binding | Golgi to plasma membrane protein transport; endoplasmic reticulum to Golgi vesicle-mediated transport;SNARE complex disassembly | Golgi stack; food vacuole; host cell cytoplasm |
| PF3D7_0302500 | CLAG3.1 | Cytoadherence linked asexual protein 3.1 | 167.24 | yes | 0 | 10.50 | 0.66 | 104.73 | 181.65 | 3.53 | 3.09 | 76.13 | S | 2.61 | 0.64 | M | RNA binding; anion channel activity; protein binding; cell adhesion molecule binding | cytoadherence to microvasculature, mediated by symbiont protein | host cell plasma membrane; rhoptry; host cell cytoplasm; extracellular vesicle |
| PF3D7_1329100 | MyoF | myosin F, putative | 250.29 | no | 0 | 19.44 | 66.45 | 51.14 | 22.61 | 5.62 | 19.54 | 106.53 | O | 79.52 | 39.69 | M | actin filament binding; microfilament motor activity; cytoskeletal motor activity; protein binding; ATP binding; actin-dependent ATPase activity | cytoskeleton organization; actin filament organization; actin filament-based movement; vesicle transport along actin filament | cytoplasm; cytosol; nuclear envelope; nucleus; nuclear pore; actin cytoskeleton; myosin complex; vesicle; apical part of cell |
| PF3D7_0526500 | N/A | Suf domain-containing protein, putative | 165.38 | no | 0 | 88.79 | 24.83 | 16.01 | 1.62 | 3.76 | 15.92 | 41.01 | R | 80.94 | 45.64 | M | mRNA binding | RNA 3'-end processing | nucleus |
| PF3D7_0105800 | N/A | cyclin-dependent kinases regulatory subunit, putative | 99.31 | no | 0 | 5.42 | 1.25 | 24.78 | 12.70 | 4.33 | 26.24 | 56.62 | O | 28.40 | 36.31 | F | kinase activity | signal transduction | nucleus |
| PF3D7_1465300 | N/A | ribonuclease Z, putative | 100.04 | no | 0 | 34.03 | 9.81 | 5.09 | 0.43 | 1.59 | 14.89 | 17.85 | R | 10.16 | 13.76 | 0 | 3'-tRNA processing endoribonuclease activity | mitochondrial tRNA 3'-trailer cleavage, endonucleolytic | mitochondrion |
| PF3D7_0823300 | GCN5 | histone acetyltransferase GCN5 | 170.92 | no | 0 | 138.06 | 181.90 | 71.62 | 12.74 | 7.97 | 24.22 | 59.65 | T | 59.27 | 31.03 | M | H3 histone acetyltransferase activity | chromatin remodeling | nucleus |
| PF3D7_0527000 | MCM3 | DNA replication licensing factor MCM3, putative | 109.80 | no | 0 | 53.07 | 30.87 | 511.11 | 273.79 | 16.69 | 48.61 | 269.17 | T | 2853.14 | 372.70 | M | ATP hydrolysis activity | DNA replication | nucleus |
| PF3D7_1311500 | RPT1 | 26S protease regulatory subunit 7, putative | 46.84 | no | 0 | 81.96 | 50.75 | 189.27 | 46.12 | 7.85 | 12.95 | 126.39 | T | 378.22 | 148.07 | M | TBP-class protein binding | proteasome regulatory particle assembly; proteasome-mediated ubiquitin-dependent protein catabolic process; | proteasome complex; nucleus; cytoplasm; ? proteasome regulatory particle, base subcomplex |
| PF3D7_0215700 | RPB2 | DNA-directed RNA polymerase II subunit RPB2, putative | 151.72 | no | 0 | 99.34 | 58.36 | 67.24 | 8.90 | 1.43 | 4.46 | 56.77 | R | 202.29 | 68.53 | M | DNA-directed 5'-3' RNA polymerase activity | transcription by RNA polymerase II | RNA polymerase II, core complex |
| PF3D7_1437900 | HSP40 | HSP40, subfamily A | 48.47 | no | 0 | 217.89 | 220.27 | 89.47 | 8.74 | 27.79 | 75.89 | 222.01 | O | 178.64 | 320.86 | F | ATP hydrolysis activity;heat shock protein binding | protein folding;repsonse to heat;response to xenobiotic stimulus | nucleus;cytosol;host cell plasma membrane |
| PF3D7_1466300 | RPN2 | 26S proteasome regulatory subunit RPN2, putative | 132.77 | no | 0 | 225.33 | 118.29 | 193.13 | 100.95 | 9.41 | 13.39 | 180.04 | R | 74.16 | 107.13 | F | endopeptidase activity | proteasome regulatory particle assembly; proteasome-mediated ubiquitin-dependent protein catabolic process; | proteasome complex; nucleus; cytoplasm; ? proteasome regulatory particle, base subcomplex |
| PF3D7_1352700 | AQR | intron-binding protein aquarius, putative | 304.30 | no | 0 | 54.96 | 21.31 | 9.91 | 1.34 | 1.54 | 8.94 | 5.34 | R | 36.03 | 6.85 | M | RNA binding;RNA helicase activity | nuclear-transcribed mRNA catabolic process, nonsense-mediated decay;mRNA splicing, via spliceosome | spliceosomal complex;catalytic step 2 spliceosome;nucleus |
| PF3D7_0818200 | 14-3-3I | 14-3-3 protein | 30.19 | no | 0 | 316.38 | 298.08 | 1144.07 | 511.61 | 242.14 | 88.32 | 323.33 | T | 267.35 | 91.82 | M | RNA binding;protein binding;histone binding | entry into host | nucleus;cytoplasm;food vacuole |
| PF3D7_0305500 | N/A | protein dopey homolog, putative | 467.88 | no | 0 | 87.86 | 100.77 | 31.51 | 17.36 | 6.90 | 30.10 | 68.16 | T | 25.46 | 11.08 | M | N/A | Golgi to endosome transport;protein transport | cytosol;trans-Golgi network;endosome;peroxisome |
| PF3D7_0107800 | MRE11 | double-strand break repair protein MRE11 | 146.15 | no | 0 | 6.20 | 4.86 | 7.78 | 1.16 | 1.93 | 18.27 | 39.47 | O | 15.12 | 12.23 | 0 | nuclease activity;protein binding;single-stranded DNA endodeoxyribonuclease activity | DNA repair;double-strand break repair via homologous recombination;double-strand break repair via nonhomologous end joining;meiotic DNA double-strand break formation;mitochondrial double-strand break repair via homologous recombination;mitotic G2 DNA damage checkpoint signaling;telomere maintenance | Mre11 complex;nucleus;site of double-strand break |
| PF3D7_0322900 | RPS3A | 40S ribosomal protein S3A, putative | 30.05 | no | 0 | 414.72 | 482.85 | 209.93 | 44.12 | 40.69 | 29.20 | 132.31 | T | 93.24 | 60.94 | M | RNA binding;structural constituent of ribosome | translation | cytosol;cytosolic small ribosomal subunit;extracellular vesicle;nucleus |
| PF3D7_0604100 | SIP2 | AP2 domain transcription factor | 229.62 | no | 0 | 5.73 | 0.57 | 29.83 | 15.16 | 6.58 | 5.73 | 40.37 | O | 45.16 | 1.87 | M | sequence-specific DNA binding;telomeric DNA binding | N/A | nuclear periphery;nucleus |
| PF3D7_1132200 | TCP1 | T-complex protein 1 subunit alpha | 60.26 | no | 0 | 295.57 | 252.35 | 125.91 | 21.29 | 30.46 | 59.02 | 174.64 | R | 328.72 | 202.50 | M | ATP binding;unfolded protein binding | protein folding | chaperonin-containing T-complex;cytosol;intracellular;nucleus |
| PF3D7_0511800 | INO1 | inositol-3-phosphate synthase | 69.11 | no | 0 | 132.59 | 17.25 | 80.78 | 17.50 | 2.47 | 36.51 | 298.86 | O | 82.39 | 480.43 | F | inositol-3-phosphate synthase activity | inositol biosynthetic process | cytoplasm |
| PF3D7_0804500 | N/A | conserved Plasmodium membrane protein | 720.95 | no | 8 | 20.68 | 21.82 | 19.27 | 5.83 | 4.01 | 55.55 | 74.26 | O | 375.68 | 50.13 | M | N/A | vacuolar acidification | RAVE complex |
| PF3D7_1366300 | N/A | conserved Plasmodium protein, unknown function | 494.68 | no | 0 | 75.37 | 41.91 | 14.07 | 1.33 | 3.05 | 27.38 | 94.33 | O | 58.40 | 32.96 | M | N/A | N/A | RNA N6-methyladenosine methyltransferase complex;nucleus |
| PF3D7_0932800 | CSE1 | importin alpha re-exporter, putative | 159.11 | no | 0 | 48.05 | 33.94 | 46.16 | 12.39 | 15.18 | 32.16 | 16.80 | R | 58.79 | 27.77 | M | nuclear export signal receptor activity | protein export from nucleus | cytosol;nuclear envelope |
| PF3D7_1464700 | N/A | ATP synthase (C/AC39) subunit, putative | 44.53 | no | 0 | 51.00 | 19.16 | 23.83 | 6.78 | 4.64 | 36.85 | 83.72 | O | 99.58 | 32.97 | M | P-type proton-exporting transporter activity;proton-transporting ATPase activity, rotational mechanism | vacuolar acidification;vacuolar transport | food vacuole;lysosomal membrane;nucleus;vacuolar proton-transporting V-type ATPase complex;vacuolar proton-transporting V-type ATPase, V0 domain |
| PF3D7_0219600 | RFC1 | replication factor C subunit 1 | 104.18 | no | 0 | 6.02 | 2.72 | 24.02 | 5.31 | 3.69 | 11.36 | 106.06 | O | 282.45 | 17.91 | M | DNA binding | DNA repair;DNA replication | vacuolar acidification;vacuolar transport |
| PF3D7_0820000 | SRCAP | Snf2-related CBP activator, putative | 246.75 | no | 0 | 41.42 | 33.94 | 15.68 | 1.21 | 2.94 | 11.26 | 93.84 | O | 39.79 | 33.45 | M | ATP binding;ATP hydrolysis activity;helicase activity;histone binding;transcription coactivator activity | ATP-dependent chromatin remodeling;chromatin remodeling;gene silencing;histone exchange | Swr1 complex;cytoplasm;membrane;nucleus |
| PF3D7_1033600 | CEF1 | pre-mRNA-splicing factor CEF1, putative | 108.41 | no | 0 | 194.63 | 116.32 | 60.04 | 17.35 | 7.74 | 14.41 | 248.40 | O | 176.84 | 91.41 | M | DNA binding;protein binding | mRNA splicing, via spliceosome;regulation of transcription, DNA-templated | ATP-dependent chromatin remodeling;chromatin remodeling;gene silencing;histone exchange |
| PF3D7_0727900 | N/A | conserved Plasmodium protein, unknown function | 230.43 | no | 2 | 38.79 | 4.71 | 4.81 | 1.50 | 1.94 | 24.82 | 8.71 | R | 7.54 | 7.14 | 0 | DNA-binding transcription factor activity;sequence-specific DNA binding | positive regulation of transcription, DNA-templated;protein autoprocessing | endoplasmic reticulum membrane;nucleus |
| PF3D7_0303100 | N/A | CLP1 P-loop domain-containing protein, putative | 170.17 | no | 0 | 3.82 | 2.53 | 4.30 | 1.18 | 1.16 | 8.17 | 11.88 | O | 18.64 | 7.25 | M | polynucleotide 5'-hydroxyl-kinase activity | RNA processing;cleavage in ITS2 between 5.8S rRNA and LSU-rRNA of tricistronic rRNA transcript (SSU-rRNA, 5.8S rRNA, LSU-rRNA) | cytosol;nucleus |
| PF3D7_1440100 | RAD21 | cohesin complex subunit RAD21 | 118.50 | no | 0 | 3.85 | 1.40 | 26.43 | 10.09 | 1.34 | 5.96 | 78.05 | O | 163.27 | 1.93 | M | N/A | N/A | cohesin complex;mitochondrion;nucleus |
| PF3D7_1361800 | GAC | glideosome-associated connector | 291.02 | no | 0 | 8.40 | 0.95 | 123.41 | 90.35 | 3.55 | 35.83 | 646.45 | O | 23.58 | 351.38 | F | RNA binding;actin filament binding | cell gliding | apical part of cell;cell surface;cytoplasm;cytosol |
| PF3D7_1138500 | PPM2 | protein phosphatase PPM2, protein phosphatase 2C | 105.40 | no | 0 | 138.71 | 30.49 | 78.17 | 16.89 | 10.96 | 57.72 | 293.37 | O | 65.16 | 61.85 | M | magnesium-dependent protein serine/threonine phosphatase activity;protein serine/threonine phosphatase activity | protein dephosphorylation | cytoplasm;nucleus |
| PF3D7_0709300 | MED14 | mediator of RNA polymerase II transcription subunit 14 | 325.49 | no | 0 | 19.66 | 9.05 | 8.28 | 1.61 | 1.74 | 11.70 | 17.26 | R | 30.96 | 12.10 | M | protein binding | regulation of transcription | mediator complex;nucleolus;nucleus |
| PF3D7_0513600 | N/A | deoxyribodipyrimidine photo-lyase, putative | 129.20 | no | 0 | 13.47 | 26.26 | 47.79 | 5.71 | 17.26 | 94.76 | 1193.82 | O | 100.80 | 190.88 | F | deoxyribodipyrimidine photo-lyase activity | DNA repair | cytostome;food vacuole;mediator complex;nucleolus;symbiont-containing vacuole membrane |
| PF3D7_1129100 | PV1 | parasitophorous vacuolar protein 1 | 51.95 | yes | 0 | 335.87 | 787.46 | 190.70 | 43.51 | 29.39 | 56.59 | 291.08 | T | 304.17 | 129.23 | M | protein binding | N/A | Maurer's cleft;food vacuole;merozoite dense granule;protein-containing complex;symbiont-containing vacuole |
| PF3D7_0603400 | TEX1 | trophozoite exported protein 1 | 132.47 | no | 0 | 22.98 | 22.01 | 12.65 | 0.91 | 3.28 | 17.64 | 65.08 | O | 124.79 | 33.72 | M | ubiquitin protein ligase activity | proteasome-mediated ubiquitin-dependent protein catabolic process | Maurer's cleft |
| PF3D7_1410400 | RAP1 | rhoptry-associated protein 1 | 90.05 | yes | 0 | 12.07 | 1.05 | 263.12 | 566.49 | 1.72 | 3.71 | 215.24 | S | 11.61 | 2.06 | M | RNA binding | N/A | extracellular vesicle;food vacuole;rhoptry;symbiont-containing vacuole |
| PF3D7_0207600 | SERA5 | Serine-repeat antigen protein | 111.77 | yes | 0 | 80.16 | 12328.00 | 2965.83 | 4124.57 | 113.8 | 13.75 | 877.08 | T | 14154.00 | 34547.00 | F | cysteine-type endopeptidase activity;cysteine-type peptidase activity;kinase binding;peptidase activity;protein binding | exit from host cell;immunoglobulin production;proteolysis;proteolysis involved in cellular protein catabolic process;regulation of immune response | extracellular space;extracellular vesicle;lysosome;symbiont-containing vacuole space;symbiont-containing vacuole |
| PF3D7_1304000 | CAPH | condensin complex subunit 2, putative | 119.22 | no | 0 | 4.84 | 0.86 | 29.56 | 18.96 | 11.09 | 12.11 | 74.38 | O | 140.41 | 26.73 | M | DNA topoisomerase binding;DNA topoisomerase type II (double strand cut, ATP-hydrolyzing) activator activity;chromatin binding | mitotic chromosome condensation | condensin complex;nucleus |
| PF3D7_0802000 | GDH3 | glutamate dehydrogenase, putative | 160.42 | no | 0 | 170.50 | 430.05 | 200.66 | 20.77 | 11.31 | 26.03 | 501.36 | O | 115.71 | 37.02 | M | glutamate dehydrogenase (NAD+) activity | glutamate catabolic process;glutamate metabolic process | cytosol;mitochondrion;nucleus |

| | | | | | | | | | | | | | | | | | | | |
|---|---|---|---|---|---|---|---|---|---|---|---|---|---|---|---|---|---|---|---|
| PF3D7_1343000 | PMT | phosphoethanolamine N-methyltransferase | 31.04 | no | 0 | 260.59 | 26.67 | 445.83 | 56.14 | 11.86 | 21.48 | 95.75 | T | 11.00 | 29.68 | F | RNA binding;phosphoethanolamine N-methyltransferase activity;protein binding | phosphatidylcholine biosynthetic process | Golgi apparatus;extracellular vesicle;food vacuole |
| PF3D7_1408700 | N/A | conserved protein, unknown function | 851.88 | no | 0 | 37.27 | 10.13 | 14.03 | 3.19 | 3.36 | 46.01 | 87.60 | O | 88.85 | 43.04 | M | K63-linked polyubiquitin modification-dependent protein binding;thiol-dependent deubiquitinase | plastid organization;protein deubiquitination involved in ubiquitin-dependent protein catabolic process | cytoplasm;nucleus |
| PF3D7_1213900 | N/A | W2 domain-containing protein, putative | 134.55 | no | 0 | 101.63 | 31.40 | 16.90 | 1.13 | 4.11 | 21.58 | 48.64 | R | 37.46 | 48.80 | F | guanyl-nucleotide exchange factor activity;translation initiation factor activity;translation initiation factor binding | | eukaryotic translation initiation factor 2B complex |
| PF3D7_1222600 | AP2-G | AP2 domain transcription factor AP2-G | 284.06 | no | 0 | 6.31 | 0.44 | 0.95 | 0.04 | 0.84 | 2.23 | 2.59 | R | 0.16 | 0.17 | 0 | DNA-binding transcription activator activity;protein binding;sequence-specific DNA binding | response to xenobiotic stimulus | nucleus |
| PF3D7_0608800 | OAT | ornithine aminotransferase | 46.06 | no | 0 | 191.69 | 1715.73 | 1569.11 | 142.75 | 141.12 | 301.74 | 854.11 | T | 114.62 | 833.80 | F | RNA binding;identical protein binding;ornithine-oxo-acid transaminase activity;protein binding;pyridoxal phosphate binding | arginine catabolic process to glutamate;arginine catabolic process to proline via ornithine;ornithine metabolic process | cytoplasm;extracellular vesicle;food vacuole;intracellular;mitochondrion |
| PF3D7_1211700 | MCM5 | DNA replication licensing factor MCM5, putative | 85.86 | no | 0 | 11.75 | 4.56 | 194.84 | 67.29 | 5.70 | 18.80 | 37.43 | T | 1362.90 | 99.36 | M | ATP binding;ATP-dependent activity, acting on DNA;DNA helicase activity;DNA replication origin binding;RNA binding;single-stranded DNA binding | DNA replication initiation;DNA strand elongation involved in DNA replication;double-strand break repair via break-induced replication;pre-replicative complex assembly involved in nuclear cell cycle DNA replication | MCM complex;nuclear pre-replicative complex;nucleus |
| PF3D7_1231100 | RAB2 | ras-related protein Rab-2 | 24.42 | no | 0 | 54.86 | 25.66 | 43.65 | 13.51 | 27.12 | 51.75 | 75.03 | O | 71.44 | 40.71 | M | N/A | endoplasmic reticulum to Golgi vesicle-mediated transport | Golgi apparatus;food vacuole;vacuole |
| PF3D7_0102900 | N/A | aspartate--tRNA ligase | 73.45 | no | 0 | 59.33 | 45.30 | 24.36 | 3.75 | 7.32 | 20.19 | 47.74 | R | 26.44 | 71.25 | F | RNA binding;aspartate-tRNA ligase activity;protein homodimerization activity | aspartyl-tRNA aminoacylation | aminoacyl-tRNA synthetase multienzyme complex;cytosol;nucleus |
| PF3D7_1234100 | BDP5 | bromodomain protein 5 | 466.01 | no | 0 | 68.35 | 28.88 | 11.75 | 2.61 | 2.92 | 31.91 | 83.41 | O | 100.42 | 56.57 | M | protein binding | N/A | nucleus;transcription regulator complex |
| PF3D7_1231900 | N/A | WD repeat-containing protein, putative | 130.78 | no | 0 | 7.94 | 3.14 | 2.37 | 0.44 | 0.78 | 12.02 | 12.75 | O | 9.88 | 16.12 | F | protein binding | N/A | N/A |
| PF3D7_1019000 | eIF2A | eukaryotic translation initiation factor subunit eIF2A | 280.75 | no | 2 | 225.70 | 93.84 | 47.82 | 21.01 | 11.12 | 21.07 | 293.04 | O | 54.48 | 31.32 | M | mRNA binding;ribosome binding;tRNA binding;translation initiation factor activity | | cytosolic small ribosomal subunit;nucleus |
| PF3D7_0317400 | N/A | DNA replication complex GINS protein, putative | 61.49 | no | 0 | 8.65 | 8.98 | 12.81 | 2.64 | 5.32 | 25.80 | 45.50 | O | 8.44 | 45.64 | F | N/A | double-strand break repair via break-induced replication | GINS complex;replication fork protection complex |
| PF3D7_1205600 | N/A | tetratricopeptide repeat protein, putative | 39.02 | no | 0 | 18.28 | 50.17 | 85.34 | 19.59 | 20.31 | 47.55 | 177.97 | O | 2047.55 | 140.64 | M | histone binding | CENP-A containing chromatin assembly;DNA replication-dependent chromatin assembly | nucleoplasm;nucleus |
| PF3D7_0725000 | N/A | exonuclease I, putative | 157.93 | no | 0 | 10.51 | 6.57 | 15.78 | 4.83 | 1.75 | 18.82 | 115.46 | O | 9.15 | 25.87 | F | 5'-3' exonuclease activity;5'-flap endonuclease activity;flap endonuclease activity | DNA recombination;mismatch repair | nucleus |
| PF3D7_0703200 | N/A | conserved Plasmodium protein, unknown function | 182.54 | no | 0 | 11.53 | 17.90 | 48.64 | 42.06 | 12.08 | 44.49 | 162.80 | O | 42.79 | 60.42 | F | N/A | N/A | nucleus |
| PF3D7_1012400 | HGPRT | hypoxanthine-guanine phosphoribosyltransferase | 26.36 | no | 0 | 752.59 | 1396.42 | 877.39 | 110.33 | 95.30 | 127.31 | 309.40 | T | 37.91 | 118.91 | F | RNA binding;hypoxanthine phosphoribosyltransferase activity;magnesium ion binding | GMP salvage;IMP salvage;adenine salvage;guanine salvage;hypoxanthine metabolic process;purine ribonucleoside salvage;response to xenobiotic stimulus | |
| PF3D7_1143300 | RPC40 | DNA-directed RNA polymerases I and III subunit RPAC1 | 38.96 | no | 0 | 105.20 | 31.73 | 12.05 | 2.45 | 5.62 | 56.14 | 102.24 | R | 55.67 | 50.72 | M | | tRNA transcription by RNA polymerase III;transcription by RNA polymerase I | RNA polymerase I complex;RNA polymerase III complex |
| PF3D7_1408000 | PMII | plasmepsin II | 51.48 | no | 0 | 49.61 | 121.67 | 65.81 | 3.60 | 11.25 | 7.38 | 55.50 | T | 13.97 | 2.81 | M | aspartic-type endopeptidase activity | hemoglobin catabolic process;response to xenobiotic stimulus | food vacuole;nucleus |
| PF3D7_1016300 | GBP130 | GBP130 protein | 95.84 | no | 1 | 592.71 | 139.11 | 491.06 | 60.9 | 26.99 | 34608.00 | 136.75 | GC V | 449.39 | 45108.00 | F | mRNA binding;protein binding | N/A | host cell cytoplasm;infected host cell surface knob |
| PF3D7_0708800 | HSP110c | heat shock protein 110 | 99.96 | no | 1 | 303.12 | 163.20 | 71.91 | 8.00 | 18.46 | 42.18 | 256.20 | R | 96.20 | 105.75 | F | ATP hydrolysis activity;protein binding | protein folding;response to heat | cytoplasm;cytosol;inner membrane pellicle complex;nucleus |
| PF3D7_0823800 | N/A | DnaJ protein, putative | 76.68 | no | 0 | 25.29 | 15.54 | 28.59 | 7.15 | 3.60 | 22.20 | 12.31 | T | 20.43 | 17.83 | M | RNA binding;unfolded protein binding | chaperone cofactor-dependent protein refolding;protein refolding | cytoplasm;food vacuole;nucleus |
| PF3D7_0917000 | MOP | merozoite organizing protein | 215.35 | no | 3 | 9.08 | 4.80 | 13.43 | 2.87 | 0.67 | 7.49 | 17.43 | O | 66.93 | 12.66 | M | N/A | cell division;mitotic chromosome condensation | apical part of cell;condensed chromosome;condensin complex |
| PF3D7_1249300 | PPM4 | protein phosphatase PPM4, putative | 119.60 | no | 0 | 29.40 | 13.49 | 25.15 | 4.69 | 2.70 | 11.17 | 124.95 | O | 48.51 | 30.66 | M | magnesium-dependent protein serine/threonine phosphatase activity;protein serine/threonine phosphatase activity | | nucleus |
| PF3D7_0309300 | N/A | N2227-like protein, putative | 112.25 | no | 0 | 37.83 | 11.42 | 24.53 | 9.60 | 4.12 | 7.50 | 29.73 | R | 24.17 | 15.36 | M | S-adenosylmethionine-dependent methyltransferase activity | N/A | cytoplasm;nucleus |
| PF3D7_1309400 | N/A | HORMA domain protein, putative | 158.12 | no | 0 | 24.72 | 22.75 | 59.20 | 28.26 | 10.48 | 158.85 | 346.56 | O | 139.32 | 2078.18 | F | N/A | N/A | nucleus |
| PF3D7_1421100 | N/A | conserved Plasmodium protein, unknown function | 150.71 | no | 0 | 28.74 | 2.45 | 4.84 | 2.11 | 1.47 | 10.45 | 5.96 | R | 6.39 | 6.66 | 0 | N/A | N/A | N/A |
| PF3D7_0805700 | FIKK8 | serine/threonine protein kinase, FIKK family | 171.80 | no | 0 | 7.04 | 1.11 | 16.65 | 6.97 | 2.04 | 10.72 | 87.91 | O | 13.35 | 12.90 | 0 | protein serine/threonine kinase activity | intracellular signal transduction;protein phosphorylation | cytoplasm;cytosol;nucleus |
| PF3D7_0501600 | RAP2 | rhoptry-associated protein 2 | 46.74 | yes | 0 | 11.35 | 1.00 | 74.17 | 50.69 | 0.74 | 4.76 | 16.26 | T | 1.80 | 1.19 | 0 | RNA binding;protein binding | entry into host | extracellular vesicle;food vacuole;rhoptry |
| PF3D7_0416900 | N/A | conserved Plasmodium protein, unknown function | 289.70 | no | 0 | 29.85 | 18.11 | 6.73 | 1.49 | 1.74 | 23.86 | 86.98 | O | 15.11 | 17.45 | F | N/A | N/A | N/A |
| PF3D7_0729100 | AKIT8 | apicomplexan kinetochore protein 8, putative | 304.52 | no | 0 | 7.69 | 3.83 | 13.36 | 1.61 | 3.38 | 21.43 | 63.27 | O | 53.85 | 20.87 | M | allantoicase activity | allantoin catabolic process | kinetochore;nucleus |
| PF3D7_0306100 | N/A | conserved Plasmodium protein, unknown function | 74.33 | no | 0 | 33.32 | 10.03 | 10.27 | 1.63 | 2.25 | 27.52 | 54.24 | O | 22.87 | 32.48 | F | N/A | N/A | nucleus |
| PF3D7_0108300 | ARP | conserved Plasmodium protein, unknown function | 254.99 | no | 0 | 407.70 | 553.88 | 117.16 | 81.20 | 26.45 | 68.73 | 191.81 | T | 188.07 | 116.00 | M | mRNA binding | N/A | host cell cytoplasm;host cell surface |
| PF3D7_1443400 | WLP1 | WD repeat-containing protein | 95.82 | no | 0 | 10.02 | 11.57 | 26.19 | 6.93 | 1.99 | 7.03 | 8.60 | T | 23.14 | 4.20 | M | protein binding | complex formation | cytosol;inner pelicle |
| PF3D7_1206800 | N/A | conserved Plasmodium protein, unknown function | 86.31 | no | 0 | 9.83 | 4.78 | 9.14 | 0.64 | 5.18 | 15.16 | 8.02 | GV | 9.90 | 5.83 | M | N/A | N/A | N/A |
| PF3D7_0817700 | RON5 | rhoptry neck protein 5 | 133.69 | no | 0 | 9.26 | 0.70 | 87.16 | 138.20 | 16.98 | 17.42 | 99.94 | S | 86.88 | 59.67 | M | RNA binding;host cell surface binding | N/A | protein-containing complex;rhoptry neck |
| PF3D7_1011800 | PREBP | PRE-binding protein | 131.62 | no | 0 | 393.91 | 383.43 | 88.77 | 14.46 | 23.65 | 184.56 | 573.99 | O | 377.24 | 2674.04 | F | DNA-binding transcription factor activity;RNA binding;double-stranded DNA binding;mRNA binding | regulation of gene expression;regulation of transcription, DNA-templated | cytoplasm;nucleus |
| PF3D7_1015800 | N/A | ribonucleoside-diphosphate reductase small chain, putative | 40.39 | no | 0 | 7.61 | 3.65 | 56.77 | 10.65 | 17.43 | 55.43 | 26.92 | T | 206.75 | 49.49 | M | ribonucleoside-diphosphate reductase activity, thioredoxin disulfide as acceptor | DNA replication;deoxyribonucleoside diphosphate metabolic process;deoxyribonucleotide biosynthetic process | ribonucleoside-diphosphate reductase complex |
| PF3D7_1224500 | ASF1 | histone chaperone ASF1 | 31.55 | no | 0 | 4.62 | 2.91 | 31.66 | 6.35 | 19.20 | 52.48 | 29.64 | GV | 35.66 | 23.46 | M | chromatin binding;histone binding;unfolded protein binding | DNA replication-dependent chromatin assembly;DNA replication-independent chromatin assembly;chromatin assembly;gene silencing;histone exchange | nucleus |
| PF3D7_1212700 | EIF3A | eukaryotic translation initiation factor 3 subunit A, putative | 166.06 | no | 0 | 1252.39 | 541.64 | 142.06 | 22.49 | 14.34 | 34.28 | 377.21 | R | 205.89 | 277.96 | F | mRNA binding;translation initiation factor activity | formation of cytoplasmic translation initiation complex;regulation of translational initiation;translation reinitiation;translational initiation | eukaryotic translation initiation factor 3 complex;eukaryotic translation initiation factor 3 complex, eIF3e;eukaryotic translation initiation factor 3 complex, eIF3m;multi-eIF complex;nucleus |
| PF3D7_1138800 | N/A | WD repeat-containing protein, putative | 222.08 | no | 0 | 78.02 | 18.27 | 13.61 | 4.09 | 2.74 | 18.55 | 102.40 | O | 48.83 | 28.15 | M | protein binding | N/A | nucleus |
| PF3D7_1325100 | N/A | phosphoribosylpyrophosphate synthetase | 49.38 | no | 0 | 651.23 | 226.86 | 147.30 | 57.81 | 16.22 | 77.72 | 47.56 | R | 12.65 | 47.31 | F | RNA binding;ribose phosphate diphosphokinase activity | 5-phosphoribose 1-diphosphate biosynthetic process;nucleotide biosynthetic process;pentose-phosphate shunt;purine nucleotide biosynthetic process;purine ribonucleoside salvage | cytoplasm;mitochondrion;nucleus;ribose phosphate diphosphokinase complex |
| PF3D7_1108400 | CK2alpha | casein kinase 2, alpha subunit | 39.89 | no | 0 | 101.82 | 47.36 | 63.15 | 17.65 | 6.41 | 27.58 | 137.34 | O | 19.73 | 33.65 | F | RNA binding;protein binding;protein serine/threonine kinase activity | peptidyl-serine phosphorylation;peptidyl-threonine phosphorylation;protein autophosphorylation;protein phosphorylation;regulation of cell cycle | cytoplasm;cytosol;extracellular vesicle;nucleus;protein kinase CK2 complex |
| PF3D7_0308600 | PRPF19 | pre-mRNA-processing factor 19, putative | 60.75 | no | 0 | 87.33 | 38.22 | 41.68 | 4.64 | 3.19 | 6.89 | 87.33 | O | 17.59 | 7.95 | M | protein binding;ubiquitin-protein transferase activity | generation of catalytic spliceosome for first transesterification step | Prp19 complex;U2-type catalytic step 1 spliceosome;catalytic step 2 spliceosome;nuclear matrix;nucleus |
| PF3D7_0209800 | UAP56 | ATP-dependent RNA helicase UAP56 | 52.22 | no | 0 | 435.76 | 272.99 | 202.38 | 49.10 | 24.66 | 75.83 | 356.27 | R | 115.16 | 216.34 | F | RNA binding;RNA helicase activity | N/A | RNA N6-methyladenosine methyltransferase complex;cytoplasm;nucleus |
| PF3D7_0206700 | ADSL | adenylosuccinate lyase | 55.18 | no | 0 | 1.31 | 0.30 | 0.14 | 0.00 | 0.14 | 0.16 | 0.00 | R | 4.22 | 21.68 | F | N6-(1,2-dicarboxyethyl)AMP AMP-lyase (fumarate-forming) activity | purine nucleotide metabolic process | Maurer's cleft;nucleus |
| PF3D7_1223100 | PKAr | cAMP-dependent protein kinase regulatory subunit | 50.83 | no | 0 | 10.63 | 1.61 | 151.01 | 346.12 | 7.79 | 21.53 | 315.60 | S | 306.72 | 87.24 | M | 14-3-3 protein binding;RNA binding;cAMP-dependent protein kinase regulator activity;protein binding | | cAMP-dependent protein kinase complex;extracellular vesicle;nucleus |
| PF3D7_0205600 | N/A | conserved Plasmodium protein, unknown function | 76.45 | no | 0 | 14.59 | 11.45 | 23.75 | 5.60 | 6.45 | 11.15 | 61.75 | O | 192.46 | 20.89 | M | N/A | N/A | nucleus |
| PF3D7_1438700 | N/A | DNA primase small subunit | 53.49 | no | 0 | 21002.00 | 22282.00 | 42.4 | 45505.00 | 34455.00 | 27.69 | 55.58 | S | 326.36 | 70.91 | M | DNA primase activity;RNA binding | DNA replication, synthesis of RNA primer | alpha DNA polymerase:primase complex |
| PF3D7_0520900 | SAHH | adenosylhomocysteinase | 53.84 | no | 0 | 158.74 | 85.38 | 357.91 | 77.62 | 7.03 | 7.53 | 52.19 | T | 23.21 | 17.86 | M | adenosylhomocysteinase activity;protein binding | S-adenosylmethionine cycle | Maurer's cleft;cytosol;intracellular |
| PF3D7_0106300 | ATP6 | calcium-transporting ATPase | 139.41 | no | 8 | 140.62 | 63.55 | 152.06 | 117.60 | 12.14 | 15.80 | 92.53 | T | 76.75 | 44.14 | M | ATP hydrolysis activity;P-type calcium transporter activity;P-type proton-exporting transporter activity;RNA binding | calcium ion transmembrane transport;calcium ion transport;cellular calcium ion homeostasis;response to drug;response to xenobiotic stimulus | |
| PF3D7_0520200 | MED17 | mediator of RNA polymerase II transcription subunit 17, putative | 82.39 | no | 0 | 9.15 | 11.46 | 10.71 | 1.28 | 1.51 | 6.68 | 40.65 | O | 36.17 | 13.04 | M | N/A | N/A | mediator complex;nucleus |
| PF3D7_0501800 | CAF1A | chromatin assembly factor 1 subunit A | 125.35 | no | 0 | 4.88 | 4.06 | 27.53 | 3.78 | 2.10 | 9.17 | 63.82 | O | 236.91 | 34.05 | M | histone binding;protein binding;transcription corepressor activity | DNA replication-independent chromatin assembly;mitotic sister chromatid segregation;obsolete regulation of chromatin silencing | HIR complex;nucleus |
| PF3D7_1037300 | AAC1 | ADP/ATP carrier protein 1 | 33.73 | no | 3 | 212.24 | 50.77 | 25.66 | 4.96 | 37.64 | 57.17 | 45.17 | R | 23.99 | 56.60 | F | ATP:ADP antiporter activity;RNA binding | N/A | cell surface;food vacuole;mitochondrial inner membrane;mitochondrion |
| PF3D7_0517000 | N/A | 60S ribosomal protein L12, putative | 18.11 | no | 0 | 305.23 | 372.68 | 146.88 | 18.76 | 112.87 | 133.48 | 270.47 | T | 72.26 | 111.64 | F | large ribosomal subunit rRNA binding;rRNA binding;structural constituent of ribosome | ribosomal large subunit assembly;translation | cytosolic large ribosomal subunit;large ribosomal subunit;nucleus;ribosome |
| PF3D7_1442900 | ARFGEF | Sec7 domain-containing protein ARFGEF, putative | 404.58 | no | 2 | 148.08 | 18.68 | 27.70 | 6.99 | 3.76 | 27.09 | 38.97 | T | 51.30 | 23.23 | M | N/A | response to brefeldin A | integral component of membrane |
| PF3D7_0422700 | EIF4A3 | eukaryotic initiation factor 4A-III, putative | 44.80 | no | 0 | 63.93 | 31.93 | 38.19 | 2.07 | 5.63 | 14.41 | 86.68 | O | 68.36 | 27.70 | M | RNA binding;RNA helicase activity;mRNA binding;translation initiation factor activity | regulation of translation;regulation of translational initiation | catalytic step 2 spliceosome;eukaryotic translation initiation factor 4F complex;nucleolus;nucleus |
| PF3D7_0815200 | N/A | importin subunit beta, putative | 99.96 | no | 0 | 80.94 | 34.55 | 40.57 | 4.74 | 7.58 | 25.51 | 61.61 | R | 79.78 | 22.95 | M | nuclear import signal receptor activity;nuclear localization sequence binding | NLS-bearing protein import into nucleus;protein import into nucleus | cytoplasm;nuclear pore;nucleus |
| PF3D7_1465900 | N/A | 40S ribosomal protein S3 | 24.67 | no | 0 | 256.94 | 318.02 | 161.44 | 15.43 | 79.36 | 127.59 | 573.21 | O | 208.09 | 546.87 | F | RNA binding;structural constituent of ribosome | translation | cytosolic small ribosomal subunit;extracellular vesicle;nucleus |
| PF3D7_1206200 | EIF3C | eukaryotic translation initiation factor 3 subunit C, putative | 116.01 | no | 0 | 352.53 | 186.86 | 66.18 | 21.86 | 15.58 | 19.57 | 102.70 | R | 156.82 | 56.50 | M | mRNA binding;translation initiation factor activity;translation initiation factor binding | regulation of translational initiation;translational initiation | eukaryotic translation initiation factor 3 complex;nucleus |
| PF3D7_0821000 | N/A | conserved Plasmodium protein, unknown function | 60.54 | no | 2 | 3.14 | 3.38 | 48.00 | 19.71 | 4.07 | 22.79 | 46.18 | T | 16.24 | 18.38 | 0 | RNA binding | N/A | integral component of membrane |
| PF3D7_1429900 | WRN | ADP-dependent DNA helicase RecQ | 169.54 | no | 0 | 11.82 | 5.98 | 11.13 | 2.72 | 0.79 | 25.42 | 66.19 | O | 165.82 | 101.35 | M | 3'-5' DNA helicase activity;DNA helicase activity;four-way junction helicase activity;protein binding | DNA duplex unwinding;DNA recombination;DNA repair;DNA unwinding involved in DNA replication;double-strand break repair via homologous recombination | chromosome;chromosome, telomeric region;cytoplasm;nucleus |
| PF3D7_1338100 | RPN3 | 26S proteasome regulatory subunit RPN3, putative | 59.07 | no | 0 | 98.98 | 64.68 | 135.80 | 130.06 | 12.21 | 33.34 | 96.02 | T | 103.08 | 90.99 | M | enzyme regulator activity | regulation of protein catabolic process | proteasome complex; nucleus; cytoplasm |
| PF3D7_1329000 | N/A | DNA-directed RNA polymerase III subunit RPC1, putative | 274.83 | no | 0 | 71.87 | 38.90 | 13.44 | 3.58 | 2.22 | 14.13 | 53.81 | R | 19.77 | 19.71 | 0 | DNA-directed 5'-3' RNA polymerase activity | transcription by RNA polymerase III | RNA polymerase III complex |
| PF3D7_0320800 | DOZI | ATP-dependent RNA helicase DDX6 | 49.41 | no | 0 | 6.50 | 20.89 | 54.61 | 11.23 | 47.25 | 163.76 | 237.14 | O | 110.82 | 263.97 | F | RNA helicase activity;mRNA binding;protein binding | P-body assembly;stress granule assembly | P-body;cytoplasm;cytoplasmic stress granule;nucleus |
| PF3D7_1133800 | USMTase | RNA (uracil-5-)methyltransferase | 101.56 | no | 1 | 33.85 | 17.73 | 28.92 | 9.73 | 2.34 | 40.58 | 89.27 | O | 17.42 | 224.16 | F | S-adenosylmethionine-dependent tRNA (m5U54) methyltransferase activity;RNA methyltransferase activity | RNA processing | apicoplast |
| PF3D7_0905400 | RhopH3 | high molecular weight rhoptry protein 3 | 104.85 | yes | 0 | 33.34 | 3.03 | 558.08 | 530.00 | 15.27 | 5.58 | 583.41 | O | 15.74 | 5.58 | M | RNA binding;host cell surface receptor binding;protein binding;protein-containing complex binding | entry into host | cell surface;extracellular vesicle;infected host cell surface knob;rhoptry;symbiont-containing vacuole membrane |
| PF3D7_0510100 | N/A | KH domain-containing protein, putative | 297.57 | no | 0 | 6.52 | 7.19 | 38.61 | 10.19 | 2.95 | 17.42 | 30.71 | T | 91.56 | 10.18 | M | RNA binding | N/A | nucleus |
| PF3D7_0410800 | AKIT3 | apicomplexan kinetochore protein 3, putative | 150.27 | no | 0 | 10.68 | 11.15 | 55.02 | 17.66 | 11.24 | 38.06 | 107.14 | O | 322.61 | 77.92 | M | protein binding | cell division | kinetochore;nucleus |
| PF3D7_0814200 | ALBA1 | DNA/RNA-binding protein Alba 1 | 27.26 | no | 0 | 3045.36 | 2028.17 | 1052.00 | 492.03 | 221.15 | 158.23 | 1025.18 | R | 161.56 | 545.58 | F | DNA binding;RNA binding;mRNA binding;translation repressor activity, mRNA regulatory element binding | | cytoplasm;extracellular vesicle;food vacuole;nucleus |
| PF3D7_0620500 | N/A | cleavage stimulation factor subunit 1, putative | 76.49 | no | 0 | 31.44 | 15.11 | 9.28 | 2.22 | 2.95 | 12.55 | 20.34 | R | 5.21 | 17.95 | F | RNA binding | RNA processing | mRNA cleavage stimulating factor complex |
| PF3D7_0630600 | N/A | deubiquitinating enzyme MINDY, putative | 110.12 | no | 0 | 16.52 | 7.76 | 38.11 | 9.58 | 5.62 | 33.99 | 105.26 | O | 74.44 | 39.73 | M | thiol-dependent deubiquitinase;Lys48-specific deubiquitinase activity;Lys48-specific deubiquitinase activity [illegible] | protein K48-linked deubiquitination | cytoplasm;nucleus |
| PF3D7_0515000 | CWC2 | pre-mRNA-splicing factor CWC2, putative | 98.38 | no | 0 | 80.85 | 64.26 | 86.26 | 54.79 | 6.78 | 29.88 | 138.15 | O | 133.81 | 57.80 | M | U6 snRNA binding;mRNA binding;pre-mRNA binding;protein binding | mRNA splicing, via spliceosome;positive regulation of cell cycle;spliceosomal snRNP assembly | Prp19 complex;U2-type catalytic step 1 spliceosome;U2-type catalytic step 2 spliceosome;nucleus;spliceosomal complex |
| PF3D7_1449500 | ApiAP2 | AP2 domain transcription factor AP2-O5, putative | 84.43 | no | 0 | 12.33 | 6.91 | 15.43 | 2.38 | 4.29 | 27.87 | 41.68 | O | 77.68 | 62.28 | M | DNA-binding transcription factor activity | regulation of transcription, DNA-templated;cell motility | heterochromatin;nucleus |
| PF3D7_1223500 | N/A | conserved Plasmodium protein, unknown function | 544.35 | no | 0 | 21.96 | 8.56 | 6.90 | 1.21 | 2.10 | 34.53 | 11.70 | GV | 30.88 | 33.09 | 0 | signaling receptor activity;transmembrane signaling receptor activity | toll-like receptor signaling pathway;immune response | integral component of plasma membrane |
| PF3D7_0418200 | EIF3M | eukaryotic translation initiation factor 3 subunit M, putative | 51.03 | no | 0 | 94.39 | 40.37 | 29.60 | 4.56 | 10.95 | 32.90 | 33.66 | R | 140.78 | 21.84 | M | translation initiation factor activity | cytoplasmic translational initiation | eukaryotic translation initiation factor 3 complex;nucleus |
| PF3D7_1229000 | N/A | conserved Plasmodium membrane protein, unknown function | 223.74 | no | 7 | 33.96 | 6.20 | 5.49 | 0.47 | 1.24 | 9.64 | 26.54 | R | 13.57 | 13.57 | 0 | N/A | N/A | integral component of membrane |
| PF3D7_1118300 | N/A | insulinase, putative | 173.62 | no | 0 | 149.71 | 39.73 | 51.13 | 10.20 | 14.30 | 18.53 | 61.07 | R | 50.92 | 24.59 | M | metalloendopeptidase activity; metal ion binding | N/A | N/A |
| PF3D7_1134000 | HSP70-3 | heat shock protein 70 | 73.30 | no | 0 | 95.08 | 272.42 | 123.14 | 7.53 | 20.29 | 21.48 | 163.92 | T | 371.73 | 193.18 | M | ATP binding;ATP hydrolysis activity;heat shock protein binding;misfolded protein binding;protein folding chaperone;unfolded protein binding | cellular response to unfolded protein;chaperone cofactor-dependent protein refolding;protein refolding;response to heat;response to unfolded protein | cytoplasm;food vacuole;mitochondrion;nucleus |
| PF3D7_1117700 | RAN | GTP-binding nuclear protein RAN/TC4 | 24.88 | no | 0 | 645.89 | 348.61 | 516.00 | 167.18 | 256.89 | 470.32 | 436.49 | R | 180.16 | 308.94 | F | GTP binding;GTPase activity | RNA export from nucleus;intracellular protein transport;protein import into nucleus;regulation of cell cycle;ribosomal subunit export from nucleus | cytoplasm;extracellular vesicle;food vacuole;nucleus |
| PF3D7_1205800 | HMGB3 | high mobility group protein B3, putative | 267.02 | no | 0 | 111.87 | 55.52 | 45.64 | 13.44 | 4.28 | 30.02 | 208.46 | O | 60.29 | 63.24 | 0 | DNA binding | N/A | nucleus |

| | | | | | | | | | | | | | | | | | | | |
|---|---|---|---|---|---|---|---|---|---|---|---|---|---|---|---|---|---|---|---|
| PF3D7_1007700 | AP2-I | AP2 domain transcription factor AP2-I | 182.66 | no | 0 | 35.10 | 204.32 | 285.73 | 99.61 | 2.49 | 14.17 | 189.67 | T | 299.60 | 65.11 | M | protein binding;sequence-specific DNA binding | response to xenobiotic stimulus | Maurer's cleft;nucleus;protein-containing complex |
| PF3D7_0218000 | RFC2 | replication factor C subunit 2, putative | 37.92 | no | 0 | 8.69 | 6.06 | 58.69 | 10.39 | 13.35 | 40.20 | 886.79 | O | 241.59 | 33.82 | M | RNA binding | DNA repair;DNA replication;DNA-dependent DNA replication | DNA replication factor C complex;nucleus |
| PF3D7_0507700 | NPL4 | nuclear protein localization protein 4, putative | 62.57 | no | 0 | 56.39 | 19.95 | 28.39 | 4.74 | 5.56 | 32.48 | 166.21 | O | 48.98 | 38.85 | M | ubiquitin binding;ubiquitin protein ligase binding | ubiquitin-dependent ERAD pathway;ubiquitin-dependent protein catabolic process | nucleus |
| PF3D7_1017900 | RPN5 | 26S proteasome regulatory subunit p55, putative | 55.21 | no | 0 | 81.76 | 28.08 | 53.35 | 20.16 | 6.70 | 38.10 | 62.76 | 81.76 | 26.05 | 48.36 | F | endopeptidase activity | proteasome-mediated ubiquitin-dependent protein catabolic process;ubiquitin-dependent protein catabolic process | cytoplasm;nuclear proteasome complex;nucleus;proteasome regulatory particle;proteasome regulatory particle, lid subcomplex |
| PF3D7_0517400 | FACT-L | FACT complex subunit SPT16, putative | 132.68 | no | 0 | 139.58 | 175.55 | 174.46 | 143.00 | 27.06 | 23.39 | 1056.64 | O | 484.10 | 74.42 | M | nucleosome binding | DNA replication-independent chromatin organization;positive regulation of transcription elongation from RNA polymerase II promoter;transcription elongation from RNA polymerase II promoter;transcription, DNA-templated | FACT complex;nucleus |
| PF3D7_1456500 | N/A | STAG domain-containing protein, putative | 283.20 | no | 0 | 5.28 | 2.08 | 13.94 | 2.25 | 2.51 | 88.83 | 66.84 | GV | 134.03 | 118.31 | M | protein binding | N/A | cohesin complex;nucleus |
| PF3D7_1306600 | N/A | V-type proton ATPase subunit H, putative | 53.83 | no | 0 | 30.02 | 12.80 | 22.26 | 5.39 | 3.84 | 22.84 | 47.96 | O | 41.04 | 15.08 | M | proton-transporting ATP synthase activity, rotational mechanism;proton-transporting ATPase activity, rotational mechanism | vacuolar acidification | vacuolar proton-transporting V-type ATPase, V1 domain |
| PF3D7_0922600 | N/A | glutamine synthetase | 63.23 | no | 0 | 183.13 | 30.36 | 31.55 | 3.95 | 10.59 | 22.89 | 144.78 | R | 19.79 | 43.68 | F | glutamate-ammonia ligase activity;catalytic activity | glutamine biosynthetic process;nitrogen compound metabolic process | cytosol |
| PF3D7_0406100 | N/A | V-type proton ATPase subunit B | 55.79 | no | 0 | 327.22 | 176.18 | 222.98 | 35.39 | 39.39 | 60.60 | 320.99 | R | 257.44 | 218.04 | 0 | protein binding;proton-transporting ATPase activity, rotational mechanism | proton-transporting V-type ATPase complex assembly;vacuolar acidification | cell periphery;food vacuole;nucleus |
| PF3D7_0309600 | PfP2 | 60S acidic ribosomal protein P2 | 11.95 | no | 0 | 317.26 | 634.57 | 246.28 | 172.14 | 305.70 | 170.07 | 584.90 | T | 301.99 | 315.73 | F | RNA binding | entry into host;protein tetramerization | Maurer's cleft;cell surface;host cell surface;nucleus |
| PF3D7_1103800 | NOT1-G | CCR4-NOT transcription complex subunit NOT1-G, putative | 391.94 | no | 4 | 7.06 | 8.95 | 17.42 | 2.72 | 5.79 | 50.75 | 124.88 | O | 380.35 | 147.15 | M | mRNA binding;molecular adaptor activity | negative regulation of translation;nuclear-transcribed mRNA catabolic process, deadenylation-dependent decay | CCR4-NOT complex;CCR4-NOT core complex;P-body;cytosol;nucleus |
| PF3D7_1328200 | N/A | BRCA2 protein, putative | 320.13 | no | 0 | 11.49 | 1.29 | 7.23 | 2.38 | 1.15 | 10.80 | 100.26 | O | 29.68 | 11.97 | M | N/A | DNA repair | nucleus |
| PF3D7_0610900 | SPT5 | transcription elongation factor SPT5, putative | 149.36 | no | 0 | 144.61 | 74.80 | 24.35 | 3.25 | 15.75 | 32.14 | 146.31 | O | 213.18 | 73.26 | M | mRNA binding | regulation of transcription by RNA polymerase II;transcription elongation from RNA polymerase II promoter;transcription, DNA-templated | DSIF complex;nucleus |
| PF3D7_1012900 | ATG18 | autophagy-related protein 18 | 43.52 | no | 0 | 181.88 | 53.41 | 27.07 | 4.97 | 6.49 | 27.66 | 32.06 | R | 29.26 | 13.13 | M | phosphatidylinositol-3,5-bisphosphate binding;phosphatidylinositol-3-phosphate binding;protein binding | autophagosome assembly;autophagy of mitochondrion;autophagy of nucleus;plastid fission;protein lipidation;protein localization to phagophore assembly site;response to xenobiotic stimulus | cytoplasmic vesicle;cytosol;extrinsic component of membrane;food vacuole;phagophore assembly site membrane |
| PF3D7_0605100 | KH1 | KH domain-containing protein, putative | 85.63 | no | 0 | 94.39 | 30.37 | 24.80 | 21.69 | 8.18 | 31.37 | 150.30 | O | 110.44 | 21.84 | M | N6-methyladenosine-containing RNA binding;RNA binding;mRNA binding | regulation of gene expression | cytoplasm;nucleus |
| PF3D7_1322200 | STU2 | protein STU2, putative | 402.41 | no | 0 | 7.42 | 6.79 | 22.70 | 9.88 | 9.18 | 46.69 | 172.04 | O | 302.30 | 31.79 | M | N/A | N/A | kinetochore;nucleus |
| PF3D7_1417200 | NOT1 | CCR4-NOT transcription complex subunit NOT1, putative | 519.82 | no | 0 | 89.22 | 42.65 | 19.22 | 3.54 | 6.25 | 51.71 | 134.31 | O | 53.47 | 50.80 | 0 | mRNA binding;molecular adaptor activity | negative regulation of translation;nuclear-transcribed mRNA catabolic process, deadenylation-dependent decay | CCR4-NOT complex;CCR4-NOT core complex;P-body;cytosol;nucleus |
| PF3D7_0501000 | N/A | Plasmodium exported protein, unknown function | 30.72 | no | 1 | 97.26 | 295.98 | 76.18 | 2.87 | 30.52 | 7.85 | 15.94 | T | 7.29 | 0.63 | M | N/A | N/A | Maurer's cleft;extracellular vesicle;nucleus |
| PF3D7_1304100 | LigI | DNA ligase I | 104.50 | yes | 0 | 5.94 | 4.61 | 25.48 | 9.42 | 5.54 | 4.85 | 39.60 | O | 107.28 | 5.26 | M | DNA ligase (ATP) activity;calcium ion binding;magnesium ion binding;manganese ion binding | DNA ligation;DNA ligation involved in DNA repair;lagging strand elongation | apicoplast;cytoplasm;mitochondrion;nucleus |
| PF3D7_0302000 | PRP46 | pre-mRNA-splicing factor PRP46, putative | 69.57 | no | 0 | 81.27 | 35.17 | 22.89 | 2.60 | 7.81 | 22.93 | 47.13 | R | 95.92 | 40.16 | M | RNA binding | mRNA splicing, via spliceosome | Cul4-RING E3 ubiquitin ligase complex;Prp19 complex;catalytic step 2 spliceosome;spliceosomal complex |
| PF3D7_1230800 | WTAP | pre-mRNA-splicing regulator WTAP, putative | 71.31 | no | 0 | 108.15 | 43.58 | 19.74 | 2.49 | 6.32 | 15.78 | 168.80 | O | 242.04 | 120.20 | M | protein binding | mRNA methylation;regulation of alternative mRNA splicing, via spliceosome | RNA N6-methyladenosine methyltransferase complex;nucleus |
| PF3D7_1223400 | N/A | phospholipid-transporting ATPase, putative | 190.44 | no | 10 | 352.87 | 150.90 | 74.98 | 34.36 | 14.67 | 32.90 | 115.95 | R | 88.29 | 51.43 | M | ATP binding;ATPase-coupled intramembrane lipid transporter activity | aminophospholipid transport;endocytosis;phospholipid translocation;retrograde vesicle-mediated transport, Golgi to endoplasmic reticulum | endosome;integral component of membrane;membrane;plasma membrane;trans-Golgi network |
| PF3D7_1305500 | MKP1 | mitogen-activated protein kinase phosphatase 1, putative | 92.28 | no | 2 | 5.80 | 4.16 | 3.67 | 0.21 | 3.47 | 23.77 | 4.87 | GV | 26.40 | 14.11 | M | MAP kinase activity | cell cycle;response to drug;response to xenobiotic stimulus | integral component of membrane |
| PF3D7_0810800 | PPPK-DHPS | hydroxymethyldihydropterin pyrophosphokinase-dihydropteroate synthase | 83.37 | no | 0 | 20.91 | 15.29 | 10.16 | 0.87 | 0.96 | 1.90 | 7.46 | R | 14.14 | 2.96 | M | dihydropteroate synthase activity | dihydrofolate biosynthetic process;response to xenobiotic stimulus;tetrahydrofolate biosynthetic process | mitochondrial envelope;nucleus |
| PF3D7_1421200 | RPS25 | 40S ribosomal protein S25 | 11.66 | no | 0 | 273.22 | 496.15 | 161.45 | 100.96 | 318.88 | 181.82 | 299.03 | T | 163.83 | 244.34 | F | structural constituent of ribosome | translation | cytosolic small ribosomal subunit;nucleus |
| PF3D7_1030500 | RPN9 | 26S proteasome regulatory subunit RPN9, putative | 46.80 | no | 0 | 18.61 | 9.68 | 15.02 | 4.31 | 2.13 | 17.23 | 10.99 | R | 8.72 | 9.74 | 0 | endopeptidase activity;structural molecule activity | proteasome assembly;ubiquitin-dependent protein catabolic process | cytosol;nucleus;proteasome regulatory particle;proteasome regulatory particle, lid subcomplex |
| PF3D7_0520100 | PPM9 | protein phosphatase PPM9, putative | 83.40 | no | 0 | 29.88 | 11.77 | 6.90 | 1.29 | 3.37 | 8.77 | 10.57 | R | 112.10 | 17.24 | M | catalytic activity;magnesium-dependent protein serine/threonine phosphatase activity;protein serine/threonine phosphatase activity | | cytoplasm |
| PF3D7_1407800 | PM4 | plasmepsin IV | 51.05 | no | 1 | 294.46 | 136.02 | 68.28 | 5.97 | 24.35 | 23.31 | 37.62 | R | 34.19 | 3.62 | M | RNA binding;aspartic-type endopeptidase activity | hemoglobin catabolic process | food vacuole;nucleus |
| PF3D7_1111700 | N/A | conserved Plasmodium protein, unknown function | 95.65 | no | 0 | 24.88 | 4.79 | 4.11 | 0.20 | 3.06 | 23.30 | 6.94 | R | 11.89 | 17.19 | F | N/A | N/A | N/A |
| PF3D7_0507100 | RPL4 | 60S ribosomal protein L4 | 46.21 | no | 0 | 1015.63 | 1201.12 | 449.42 | 114.39 | 19.70 | 33.01 | 1682.00 | O | 103.13 | 153.31 | F | RNA binding;structural constituent of ribosome | translation | cytosolic large ribosomal subunit;nucleus |
| PF3D7_0629400 | PABP3 | polyadenylate-binding protein 3, putative | 90.78 | no | 0 | 7.40 | 22.89 | 26.01 | 1.72 | 4.84 | 27.03 | 8.94 | GV | 74.20 | 42.02 | M | RNA binding;mRNA binding | N/A | cytosol;nucleus |
| PF3D7_0317600 | RPS11 | 40S ribosomal protein S11, putative | 18.76 | no | 0 | 719.09 | 557.12 | 229.51 | 44.62 | 128.79 | 161.81 | 689.78 | R | 72.78 | 405.01 | F | structural constituent of ribosome | translation | cytosolic small ribosomal subunit;extracellular vesicle;nucleus |
| PF3D7_1126500 | N/A | WD repeat-containing protein, putative | 163.53 | no | 0 | 63.27 | 17.69 | 6.45 | 0.42 | 3.43 | 37.47 | 17.75 | R | 5.73 | 14.72 | F | protein binding | N/A | N/A |
| PF3D7_0305600 | APE1 | DNA-(apurinic or apyrimidinic site) endonuclease | 72.44 | no | 0 | 7.32 | 1.45 | 22.35 | 7.53 | 3.58 | 102.14 | 172.82 | O | 368.56 | 508.80 | F | 3'-5' exonuclease activity;DNA-(apurinic or apyrimidinic site) endonuclease activity;double-stranded DNA 3'-5' exodeoxyribonuclease activity;phosphatase activity;phosphodiesterase I activity;ribonuclease activity | DNA repair;base-excision repair;entry into host | mitochondrion;nucleus |
| PF3D7_1427500 | MSH2-1 | DNA mismatch repair protein MSH2, putative | 94.74 | no | 0 | 3.85 | 2.31 | 17.95 | 2.51 | 0.90 | 3.35 | 9.92 | T | 3.36 | 2.14 | 0 | ATP-dependent activity, acting on DNA;DNA binding;damaged DNA binding;double-strand/single-strand DNA junction binding | DNA recombination;maintenance of DNA repeat elements;mismatch repair;negative regulation of DNA recombination;postreplication repair | MutSalpha complex;MutSbeta complex;mismatch repair complex;nucleus |
| PF3D7_1333600 | UTP4 | U3 small nucleolar RNA-associated protein 4, putative | 159.34 | no | 1 | 118.79 | 83.82 | 11.71 | 1.78 | 4.58 | 20.87 | 22.24 | R | 41.72 | 24.81 | F | protein binding | maturation of SSU-rRNA from tricistronic rRNA transcript (SSU-rRNA, 5.8S rRNA, LSU-rRNA) | 90S preribosome;membrane;nucleolus;small-subunit processome;t-UTP complex |
| PF3D7_1345400 | N/A | conserved Plasmodium protein, unknown function | 174.49 | no | 0 | 9.27 | 8.90 | 19.93 | 5.55 | 2.79 | 19.76 | 213.43 | O | 1725.45 | 142.65 | F | N/A | N/A | N/A |
| PF3D7_1474800 | N/A | proteasome subunit alpha type-1, putative | 28.84 | no | 0 | 102.83 | 39.66 | 90.00 | 14.65 | 29.03 | 67.83 | 201.62 | O | 150.12 | 52.39 | F | endopeptidase activity | proteasomal protein catabolic process;proteasomal ubiquitin-independent protein catabolic process;proteasome-mediated ubiquitin-dependent protein catabolic process;ubiquitin-dependent protein catabolic process | cytoplasm;nucleus;proteasome core complex;proteasome core complex, alpha-subunit complex |
| PF3D7_0618000 | N/A | conserved Plasmodium membrane protein, unknown function | 92.75 | no | 4 | 4.88 | 0.46 | 15.89 | 13.20 | 1.46 | 3.97 | 52.92 | O | 8.67 | 1.39 | F | N/A | N/A | integral component of membrane;food vacuole |
| PF3D7_1369200 | N/A | conserved Plasmodium protein, unknown function | 259.08 | no | 0 | 20.37 | 7.01 | 7.00 | 1.03 | 1.20 | 25.91 | 23.23 | GV | 13.10 | 34.75 | M | N/A | N/A | N/A |
| PF3D7_0704100 | BTP2 | basal complex transmembrane protein 2 | 425.26 | no | 7 | 7.24 | 0.77 | 15.24 | 16.12 | 4.86 | 220.48 | 148.40 | GV | 59.05 | 457.23 | M | N/A | N/A | basal part of cell;inner membrane pellicle complex |
| PF3D7_1245100 | KLP8 | kinesin-13, putative | 153.12 | no | 0 | 3.53 | 0.56 | 11.61 | 2.52 | 2.73 | 30.72 | 63.98 | O | 112.10 | 28.28 | 0 | cytoskeletal motor activity | axoneme assembly;microtubule-based movement | cytoplasm;kinesin complex;kinetochore;nucleus |
| PF3D7_1027800 | RPL3 | 60S ribosomal protein L3 | 44.22 | no | 0 | 770.37 | 705.27 | 466.74 | 64.19 | 52.59 | 45.18 | 446.97 | R | 211.93 | 219.59 | F | RNA binding;structural constituent of ribosome | ribosomal large subunit assembly;translation | cytosolic large ribosomal subunit;intracellular;nucleus |
| PF3D7_1424100 | RPL5 | 60S ribosomal protein L5, putative | 34.00 | no | 0 | 504.89 | 589.18 | 262.21 | 53.34 | 13.25 | 26.40 | 437.26 | T | 146.76 | 106.81 | M | 5S rRNA binding;RNA binding;structural constituent of ribosome | ribosomal large subunit assembly;translation | cytosolic large ribosomal subunit;extracellular vesicle;nucleus |
| PF3D7_0424600 | N/A | Plasmodium exported protein (PHISTb) | 35.96 | no | 0 | 55.89 | 0.78 | 5.51 | 0.04 | 0.55 | 0.70 | 44.03 | R | 0.34 | 0.04 | 0 | N/A | cytoadherence to microvasculature, mediated by symbiont protein | extracellular vesicle;host cell periphery;host cell surface |
| PF3D7_0212400 | NUP390 | nucleoporin NUP390, putative | 494.21 | no | 7 | 18.97 | 13.56 | 9.58 | 1.65 | 3.32 | 21.83 | 26.29 | O | 18.83 | 7.73 | M | protein binding | nucleocytoplasmic transport | nuclear periphery |
| PF3D7_1328000 | N/A | conserved Plasmodium protein, unknown function | 261.45 | no | 0 | 3.61 | 0.46 | 7.83 | 1.76 | 0.88 | 12.63 | 85.74 | O | 300.50 | 12.12 | M | N/A | N/A | N/A |
| PF3D7_0608500 | N/A | proteasome subunit alpha type-2, putative | 26.53 | no | 0 | 27.23 | 14.55 | 40.38 | 9.35 | 14.22 | 57.75 | 75.37 | O | 61.22 | 53.59 | M | endopeptidase activity | proteasomal protein catabolic process;proteasomal ubiquitin-independent protein catabolic process;proteasome-mediated ubiquitin-dependent protein catabolic process;ubiquitin-dependent protein catabolic process | cytoplasm;extracellular vesicle;nucleus;proteasome core complex;proteasome core complex, alpha-subunit complex |
| PF3D7_0802100 | AP2-LT | AP2 domain transcription factor AP2-LT | 299.29 | no | 0 | 7.72 | 1.64 | 39.37 | 26.49 | 7.16 | 32.48 | 220.82 | O | 142.23 | 54.00 | M | sequence-specific DNA binding | regulation of transcription | histone acetyltransferase complex;nucleus |
| PF3D7_1326600 | N/A | conserved Plasmodium protein, unknown function | 575.46 | no | 0 | 5.23 | 1.44 | 8.33 | 1.36 | 1.62 | 15.97 | 24.12 | O | 247.48 | 16.86 | M | N/A | N/A | N/A |
| PF3D7_0821600 | CLP1 | polyribonucleotide 5'-hydroxyl-kinase Clp1, putative | 79.06 | no | 0 | 51.37 | 17.43 | 11.49 | 0.87 | 3.16 | 21.91 | 81.13 | O | 12.29 | 27.11 | F | polydeoxyribonucleotide kinase activity;polynucleotide 5'-hydroxyl-kinase activity | RNA processing;mRNA polyadenylation;tRNA splicing, via endonucleolytic cleavage and ligation | nucleus;tRNA-intron endonuclease complex |
| PF3D7_0322700 | N/A | conserved Plasmodium protein, unknown function | 372.68 | no | 7 | 16.18 | 14.85 | 11.66 | 1.74 | 3.09 | 17.13 | 15.59 | GV | 59.66 | 16.42 | M | N/A | N/A | N/A |
| PF3D7_0416800 | SAR1 | small GTP-binding protein sar1 | 22.02 | no | 0 | 225.73 | 204.89 | 284.79 | 151.66 | 104.27 | 164.20 | 455.41 | O | 144.46 | 143.91 | 0 | GTPase activity | endoplasmic reticulum to Golgi vesicle-mediated transport;intracellular protein transport;membrane organization;positive regulation of protein exit from endoplasmic reticulum;regulation of COPII vesicle coating;vesicle organization | COPII vesicle coat;Maurer's cleft;endoplasmic reticulum;endoplasmic reticulum exit site;food vacuole |
| PF3D7_0621200 | PDX1 | pyridoxine biosynthesis protein PDX1 | 33.01 | no | 0 | 223.47 | 1032.83 | 102.45 | 15.53 | 83.20 | 106.48 | 160.93 | T | 43.88 | 257.79 | F | amine-lyase activity;catalytic activity;glutaminase activity;identical protein binding | cellular amino acid metabolic process;pyridoxal phosphate biosynthetic process;pyridoxine biosynthetic process;response to singlet oxygen;vitamin B6 biosynthetic process | cytosol;extracellular vesicle;nucleus |
| PF3D7_1129200 | RPN7 | 26S proteasome regulatory subunit RPN7, putative | 46.62 | no | 0 | 73.85 | 21.92 | 67.53 | 14.76 | 9.22 | 42.44 | 107.29 | O | 23.62 | 65.43 | F | endopeptidase activity | proteasome-mediated ubiquitin-dependent protein catabolic process;proteolysis | nucleus;proteasome complex |
| PF3D7_0214100 | SEC31 | protein transport protein SEC31 | 166.71 | no | 0 | 412.56 | 94.90 | 268.41 | 535.68 | 14.17 | 17.78 | 89.82 | S | 41.62 | 26.85 | M | structural molecule activity | COPII-coated vesicle budding;COPII-coated vesicle cargo loading;endoplasmic reticulum organization;endoplasmic reticulum to Golgi vesicle-mediated transport;intracellular protein transport;translocation of peptides or proteins into host cell cytoplasm | COPII vesicle coat;COPII-coated ER to Golgi transport vesicle;Maurer's cleft;endoplasmic reticulum exit site;host cell cytoplasm |
| PF3D7_0811500 | CARM1 | histone-arginine methyltransferase CARM1, putative | 108.43 | no | 0 | 48.98 | 10.74 | 3.63 | 0.68 | 1.74 | 7.54 | 5.48 | R | 3.06 | 3.00 | 0 | histone methyltransferase activity | histone methylation | nucleus |
| PF3D7_0722200 | RALP1 | rhoptry-associated leucine zipper-like protein 1 | 87.88 | yes | 1 | 10.93 | 0.94 | 75.57 | 83.40 | 9.61 | 9.65 | 212.23 | O | 17.21 | 26.33 | F | host cell surface receptor binding | entry into host | rhoptry neck |
| PF3D7_1338300 | EF-1gamma | elongation factor 1-gamma, putative | 47.78 | no | 0 | 488.79 | 165.11 | 106.59 | 26.11 | 13.81 | 67.05 | 138.72 | R | 10.21 | 114.70 | F | glutathione transferase activity;translation elongation factor activity | translational elongation | eukaryotic translation elongation factor 1 complex;nucleus |
| PF3D7_1134800 | N/A | coatomer subunit delta | 56.84 | no | 0 | 184.16 | 46.05 | 48.07 | 37.73 | 28.07 | 35.57 | 16.57 | R | 54.67 | 67.53 | F | protein binding | Golgi localization;endoplasmic reticulum to Golgi vesicle-mediated transport;retrograde vesicle-mediated transport, Golgi to endoplasmic reticulum | COPI vesicle coat;COPI-coated vesicle;cytoplasm;cytosol |
| PF3D7_0713700 | N/A | conserved Plasmodium protein, unknown function | 24.38 | yes | 2 | 87.64 | 11.77 | 32.63 | 6.74 | 56.84 | 330.75 | 35.67 | GV | 81.11 | 135.52 | F | N/A | N/A | nucleus |
| PF3D7_0316800 | N/A | 40S ribosomal protein S15A, putative | 14.87 | no | 0 | 237.36 | 212.36 | 135.61 | 19.70 | 225.81 | 167.35 | 275.29 | O | 159.49 | 273.05 | F | RNA binding;structural constituent of ribosome | translation | cytosolic small ribosomal subunit;nucleus |
| PF3D7_0819000 | N/A | ribonuclease, putative | 157.67 | no | 0 | 77.37 | 25.93 | 9.81 | 0.73 | 2.69 | 22.02 | 93.05 | O | 54.18 | 27.29 | M | RNA binding | mRNA 3'-end processing by stem-loop binding and cleavage;mRNA polyadenylation;pre-mRNA cleavage required for polyadenylation | mRNA cleavage and polyadenylation specificity factor complex |
| PF3D7_1006700 | N/A | conserved Plasmodium protein, unknown function | 45.62 | no | 0 | 39.04 | 24.62 | 12.39 | 0.44 | 14.69 | 100.71 | 181.79 | O | 265.79 | 363.06 | F | N/A | N/A | nucleus |
| PF3D7_1454400 | APP | aminopeptidase P | 90.13 | yes | 1 | 166.57 | 173.38 | 68.35 | 10.01 | 7.25 | 31.06 | 75.96 | T | 39.23 | 69.75 | F | aminopeptidase activity;metalloaminopeptidase activity;protein homodimerization activity | hemoglobin catabolic process;proteolysis | cytosol;food vacuole |
| PF3D7_1110200 | PRPF6 | pre-mRNA-processing factor 6, putative | 152.15 | no | 0 | 172.13 | 70.74 | 43.29 | 7.94 | 3.00 | 12.88 | 157.43 | R | 72.23 | 49.06 | M | RNA bindding | mRNA splicing, via spliceosome;spliceosomal tri-snRNP complex assembly | U4/U6 x U5 tri-snRNP complex;catalytic step 2 spliceosome |
| PF3D7_1136500 | CK1 | Casein kinase 1 | 37.79 | no | 0 | 756.48 | 111.09 | 331.05 | 88.97 | 93.86 | 156.19 | 928.77 | O | 895.88 | 546.88 | M | protein serine/threonine kinase activity | endocytosis;peptidyl-serine phosphorylation | cytoplasm;nucleus |
| PF3D7_1217900 | N/A | PPPDE peptidase domain-containing protein, putative | 56.09 | no | 0 | 10.83 | 5.84 | 23.09 | 29.59 | 8.96 | 83.20 | 70.28 | GV | 67.55 | 92.55 | F | peptidase activity | protein modification by small protein removal | N/A |
| PF3D7_0305100 | N/A | conserved Plasmodium protein, unknown function | 475.13 | no | 2 | 8.82 | 14.44 | 12.02 | 2.88 | 4.34 | 40.65 | 33.74 | GV | 268.59 | 57.20 | M | N/A | N/A | N/A |
| PF3D7_0811400 | N/A | conserved protein, unknown function | 71.37 | no | 0 | 179.07 | 76.29 | 53.67 | 13.10 | 10.31 | 12.55 | 63.45 | R | 149.38 | 25.93 | M | N/A | N/A | nucleus |
| PF3D7_1355700 | NIF3 | NLI interacting factor-like phosphatase, putative | 151.54 | no | 0 | 63.86 | 32.67 | 22.22 | 5.36 | 8.66 | 42.63 | 25.36 | R | 19.83 | 18.81 | 0 | RNA polymerase II CTD heptapeptide repeat phosphatase activity | dephosphorylation of RNA polymerase II C-terminal domain | nucleus |
| PF3D7_1013600 | N/A | Uncharacterized protein | 269.27 | no | 0 | 27485.00 | 28856.00 | 45363.00 | 27454.00 | 28856.00 | 15.38 | 29921.00 | T | 39.93 | 19.54 | M | | | nucleus |
| PF3D7_1241700 | RFC4 | replication factor C subunit 4, putative | 37.64 | no | 0 | 8.96 | 5.46 | 35.56 | 11.77 | 8.63 | 24.56 | 49.68 | O | 130.71 | 54.65 | M | ATP binding;DNA clamp loader activity | DNA repair;DNA replication;DNA-dependent DNA replication;leading strand elongation;response to xenobiotic stimulus | DNA replication factor C complex;nucleus |
| PF3D7_0529400 | AKIT5 | apicomplexan kinetochore protein 5, putative | 185.37 | no | 0 | 11.01 | 18.82 | 62.00 | 31.27 | 9.27 | 38.11 | 461.55 | O | 288.64 | 93.96 | M | N/A | N/A | nucleus |
| PF3D7_1420200 | UNC | myosin-specific chaperone UNC, putative | 148.02 | no | 0 | 6.49 | 2.02 | 12.58 | 8.86 | 3.59 | 19.00 | 25.79 | O | 15.46 | 15.52 | 0 | protein binding | protein folding | cytosol |

| | | | | | | | | | | | | | | | | | | |
|---|---|---|---|---|---|---|---|---|---|---|---|---|---|---|---|---|---|---|
| PF3D7_1422400 | N/A | nucleolar RNA-associated protein, putative | 195.36 | no | 0 | 36.18 | 14.53 | 3.87 | 0.89 | 2.11 | 14.64 | 30.65 | O | 36.45 | 18.82 | M | RNA binding | rRNA processing;tRNA export from nucleus | CURI complex;UTP-C complex;nucleolus;small-subunit processome |
| PF3D7_0933200 | N/A | calcyclin-binding protein, putative | 24.00 | no | 0 | 10.12 | 20.78 | 48.46 | 14.26 | 175.70 | 139.55 | 37.16 | G II | 332.92 | 221.68 | M | protein binding | protein stabilization;regulation of protein stability | N/A |
| PF3D7_0730500 | N/A | conserved Plasmodium protein, unknown function | 329.46 | no | 0 | 5.58 | 0.92 | 10.46 | 3.78 | 2.68 | 26.83 | 67.63 | O | 19.23 | 47.57 | F | N/A | N/A | nucleus |
| PF3D7_1004300 | RFUL | RING finger E3 ubiquitin-protein ligase | 131.70 | no | 0 | 166.81 | 29.31 | 59.63 | 14.23 | 9.65 | 88.75 | 244.47 | O | 180.51 | 739.51 | F | ubiquitin protein ligase activity | protein ubiquitination;ubiquitin-dependent protein catabolic process | cytoplasm;nucleus |
| PF3D7_0607000 | N/A | translation initiation factor IF-2, putative | 112.36 | no | 0 | 262.17 | 174.02 | 64.16 | 15.05 | 9.03 | 23.82 | 207.45 | R | 175.02 | 100.50 | M | translation initiation factor activity | translational initiation | nucleus |
| PF3D7_0910100 | N/A | exportin-7, putative | 139.18 | no | 0 | 37.56 | 8.20 | 37.48 | 8.39 | 4.83 | 7.55 | 13.01 | R | 40.83 | 8.46 | M | nuclear export signal receptor activity | plastid organization;protein export from nucleus | cytoplasm;nuclear pore;nucleus |
| PF3D7_1343300 | N/A | CDT1-like protein, putative | 114.58 | no | 0 | 8.77 | 13.07 | 34.37 | 6.99 | 7.79 | 46.36 | 129.45 | O | 342.31 | 59.51 | M | N/A | N/A | nucleus |
| PF3D7_1456700 | N/A | conserved Plasmodium protein, unknown function | 77.81 | no | 0 | 52.58 | 20.59 | 54.59 | 10.84 | 16.52 | 31.83 | 34.37 | T | 97.21 | 21.47 | M | N/A | N/A | N/A |
| PF3D7_0922100 | N/A | ubiquitin-like protein, putative | 182.09 | no | 2 | 235.85 | 27.78 | 21.69 | 3.04 | 5.73 | 27.64 | 59.17 | R | 171.26 | 59.16 | M | polyubiquitin modification-dependent protein binding | ubiquitin-dependent protein catabolic process | cytosol |
| PF3D7_1422800 | ARP4 | actin-related protein ARP4 | 60.66 | no | 0 | 15.09 | 5.60 | 8.36 | 0.84 | 5.33 | 13.50 | 43.63 | O | 38.95 | 9.25 | M | chromatin binding;protein binding;structural constituent of cytoskeleton | ATP-dependent chromatin remodeling;chromatin remodeling;cytoskeleton organization;endocytosis;exocytosis;histone H4 acetylation;regulation of gene expression;regulation of transcription by RNA polymerase II;schizogony | NuA4 histone acetyltransferase complex;SWI/SNF complex;actin filament;nucleus |
| PF3D7_1360700 | PIAS | E3 SUMO-protein ligase PIAS, putative | 130.64 | no | 0 | 33.57 | 18.75 | 30.26 | 7.74 | 2.71 | 16.61 | 52.92 | O | 78.64 | 18.78 | M | SUMO transferase activity | protein sumoylation | N/A |
| PF3D7_1118500 | NOP56 | nucleolar protein 56, putative | 69.39 | no | 0 | 411.95 | 138.07 | 42.30 | 11.69 | 21.43 | 15.69 | 130.05 | R | 401.74 | 99.98 | M | snoRNA binding | rRNA processing | box C/D RNP complex;nucleus;small-subunit processome |
| PF3D7_1457300 | N/A | MA3 domain-containing protein, putative | 75.99 | no | 0 | 182.45 | 36.58 | 26.15 | 4.41 | 4.98 | 44.93 | 31.83 | R | 9.68 | 17.11 | F | translation initiation factor binding | translation | apicoplast;nucleus |
| PF3D7_0212100 | N/A | conserved Plasmodium protein, unknown function | 214.68 | no | 0 | 49.63 | 26.06 | 45.75 | 11.10 | 8.00 | 14.18 | 180.40 | O | 75.13 | 26.74 | M | N/A | N/A | nuclear periphery;nucleus |
| PF3D7_1304500 | N/A | small heat shock protein, putative | 25.20 | no | 0 | 5.08 | 14.65 | 19.44 | 1.12 | 64.54 | 169.50 | 77.14 | GV | 45.09 | 90.32 | F | protein self-association;unfolded protein binding | protein complex oligomerization;protein folding;response to heat;response to hydrogen peroxide;response to reactive oxygen species;response to salt stress;response to unfolded protein | mitochondrion |
| PF3D7_1406200 | SPT6 | transcription elongation factor SPT6, putative | 338.38 | no | 0 | 102.71 | 61.35 | 21.10 | 2.44 | 5.82 | 29.94 | 60.85 | O | 224.56 | 96.85 | M | histone binding;nucleosome binding | mRNA transcription by RNA polymerase II;nucleosome organization;obsolete chromatin maintenance;positive regulation of transcription elongation from RNA polymerase II promoter;regulation of mRNA processing;transcription elongation from RNA polymerase II promoter | nucleus;transcription elongation factor complex;transcriptionally active chromatin |
| PF3D7_0407800 | CINCH | protein CINCH | 230.12 | no | 0 | 5.14 | 0.48 | 36.03 | 22.44 | 4.14 | 43.47 | 225.65 | O | 46.25 | 176.58 | F | protein binding | cell division | basal part of cell;inner membrane pellicle complex;nucleus |
| PF3D7_0921600 | N/A | tetratricopeptide repeat protein, putative | 198.03 | no | 0 | 33.35 | 11.83 | 5.09 | 0.75 | 1.27 | 13.07 | 26.52 | R | 30.94 | 13.19 | M | protein binding | N/A | N/A |
| PF3D7_0808100 | N/A | AP-3 complex subunit delta, putative | 193.08 | no | 2 | 6.66 | 5.57 | 8.17 | 1.88 | 1.28 | 10.71 | 4.12 | GV | 2.91 | 11.01 | F | protein binding | Golgi to vacuole transport;intracellular protein transport;protein targeting to vacuole | AP-3 adaptor complex;endosome membrane |
| PF3D7_1105400 | RPS4 | 40S ribosomal protein S4, putative | 29.77 | no | 0 | 288.16 | 391.25 | 236.48 | 50.29 | 86.13 | 104.03 | 334.58 | T | 86.41 | 114.11 | F | RNA binding;structural constituent of ribosome | translation | cytosolic small ribosomal subunit;extracellular vesicle;nucleus;small ribosomal subunit |
| PF3D7_0612900 | NOG1 | nucleolar GTP-binding protein 1, putative | 79.53 | no | 0 | 424.38 | 254.88 | 72.02 | 12.96 | 26.81 | 55.44 | 284.10 | R | 52.09 | 385.59 | F | GTP binding | ribosomal large subunit biogenesis | nucleolus;preribosome, large subunit precursor |
| PF3D7_0525800 | IMC1g | inner membrane complex protein 1g | 34.34 | no | 0 | 12.68 | 1.27 | 89.65 | 27.95 | 7.02 | 87.12 | 816.21 | O | 3.10 | 27.46 | F | N/A | component of protein complex | inner membrane pellicle complex |
| PF3D7_1446500 | NUP313 | nucleoporin NUP313, putative | 346.92 | no | 0 | 69.27 | 31.12 | 23.23 | 1.91 | 3.37 | 27.38 | 52.11 | R | 63.87 | 19.05 | M | protein binding | nucleocytoplasmic transport | nuclear pore;nucleus |
| PF3D7_1463200 | RFC3 | replication factor C subunit 3, putative | 39.23 | no | 0 | 4.54 | 3.54 | 45.00 | 6.50 | 6.03 | 11.15 | 8.77 | T | 141.14 | 15.91 | M | DNA binding | DNA repair;DNA replication;DNA strand elongation involved in DNA replication;DNA-dependent DNA replication;leading strand elongation | DNA replication factor C complex;PCNA complex;nucleus |
| PF3D7_1441200 | N/A | 60S ribosomal protein L1, putative | 24.81 | no | 0 | 66.45 | 52.63 | 24.83 | 3.54 | 14.53 | 36.07 | 21.44 | R | 30.17 | 15.71 | M | RNA binding;structural constituent of ribosome | maturation of LSU-rRNA;translation | cytosolic large ribosomal subunit;intracellular;nucleus |
| PF3D7_1466400 | ApiAP2 | AP2 domain transcription factor AP2-EXP | 92.09 | no | 0 | 200.36 | 29.73 | 20.00 | 2.82 | 3.59 | 14.47 | 15.52 | R | 64.32 | 9.47 | M | sequence-specific DNA binding | positive regulation of gene expression;response to xenobiotic stimulus | heterochromatin;nucleus |
| PF3D7_0704400 | N/A | phosphoinositide-binding protein, putative | 118.84 | no | 0 | 7.37 | 2.67 | 15.19 | 4.91 | 1.25 | 23.51 | 7.79 | GV | 22.36 | 56.25 | F | transcription coactivator activity | histone H3 acetylation;regulation of transcription, DNA-templated;transcription initiation from RNA polymerase II promoter | nucleus;transcription factor TFIID complex |
| PF3D7_1412100 | MCMBP | mini-chromosome maintenance complex-binding protein | 112.77 | no | 0 | 8.12 | 5.38 | 15.08 | 6.86 | 1.21 | 7.49 | 16.46 | O | 27.05 | 5.82 | M | chromatin binding | DNA-dependent DNA replication;schizogony | MCM complex;cytoplasm;nucleus |
| PF3D7_1345800 | N/A | conserved Plasmodium protein, unknown function | 118.92 | no | 0 | 33.18 | 23.32 | 21.71 | 4.63 | 6.31 | 24.30 | 17.32 | R | 109.60 | 17.73 | M | N/A | N/A | nucleus |
| PF3D7_0210200 | N/A | Uncharacterized protein | 310.02 | no | 0 | 22586.00 | 22859.00 | 29.31 | 32721.00 | 19603.00 | 45.48 | 53.73 | S | 548.18 | 52.03 | M | N/A | N/A | nucleolus;nucleus |
| PF3D7_1462300 | N/A | GTP-binding protein, putative | 161.22 | yes | 3 | 96.52 | 30.63 | 31.09 | 4.30 | 7.51 | 37.90 | 55.87 | R | 32.46 | 21.36 | M | RNA binding;GTPase activity;GTP binding | N/A | nucleus;integral component of membrane |
| PF3D7_1451800 | SORTLR | sortilin | 102.27 | no | 2 | 15.72 | 39.26 | 125.00 | 31.58 | 4.45 | 55.11 | 212.66 | O | 49.53 | 480.74 | F | lipid transporter activity;protein binding | endocytosis;post-Golgi vesicle-mediated transport;protein transport | Golgi apparatus;cis-Golgi network;endoplasmic reticulum;integral component of membrane |
| PF3D7_0629200 | N/A | DnaJ protein, putative | 44.67 | no | 1 | 36.24 | 32.17 | 27.98 | 3.44 | 2.96 | 12.81 | 215.58 | O | 15.63 | 18.89 | 0 | RNA binding | response to heat;response to unfolded protein | apicoplast;nucleus;symbiont-containing vacuole |
| PF3D7_1206600 | RPC2 | DNA-directed RNA polymerase III subunit RPC2, putative | 167.29 | no | 0 | 95.46 | 39.26 | 14.03 | 1.06 | 2.24 | 23.32 | 71.43 | R | 38.00 | 44.63 | F | DNA-directed 5'-3' RNA polymerase activity | transcription by RNA polymerase III | RNA polymerase III complex |
| PF3D7_1405600 | RNR | ribonucleoside-diphosphate reductase small chain, putative | 40.60 | no | 0 | 6.45 | 6.44 | 43.54 | 10.75 | 27.90 | 32.33 | 118.87 | O | 494.07 | 46.18 | M | ribonucleoside-diphosphate reductase activity, thioredoxin disulfide as acceptor | deoxyribonucleotide biosynthetic process | intracellular;ribonucleoside-diphosphate reductase complex;cytoplasm;nucleus |
| PF3D7_1367100 | SNP1 | U1 small nuclear ribonucleoprotein 70 kDa homolog, putative | 51.84 | no | 0 | 411.70 | 108.40 | 99.27 | 49.07 | 15.74 | 12.26 | 268.47 | R | 262.26 | 101.69 | M | RNA binding;U1 snRNA binding;mRNA binding;snRNA binding | RNA processing;mRNA splicing, via spliceosome | U1 snRNP;U2-type prespliceosome;spliceosomal complex |
| PF3D7_1466800 | N/A | NOC3 domain-containing protein, putative | 105.45 | no | 0 | 25.90 | 12.41 | 4.08 | 0.23 | 47.43 | 116.99 | 628.46 | O | 19.05 | 307.57 | F | chromatin binding | N/A | nucleus |
| PF3D7_1417600 | OSCP | ookinete surface and oocyst capsule protein OSCP, putative | 548.86 | no | 1 | 66.92 | 45.79 | 17.73 | 46844.00 | 30407.00 | 45350.00 | 71.3 | S | 67.25 | 45624.00 | F | signaling receptor activity | cell motility | capsule;cell surface;integral component of plasma membrane |
| PF3D7_0509100 | SMC4 | structural maintenance of chromosomes protein 4, putative | 201.67 | no | 0 | 12.70 | 5.30 | 49.19 | 17.16 | 9.97 | 12.86 | 132.53 | O | 749.06 | 46.70 | M | chromatin binding | mitotic chromosome condensation | chromosome, centromeric region;condensin complex;nucleus |
| PF3D7_0526200 | ARFGAP2 | ADP-ribosylation factor GTPase-activating protein, putative | 58.51 | no | 0 | 45512.00 | 29707.00 | 45310.00 | 13820.00 | 12086.00 | 45468.00 | 33.86 | R | 40.92 | 76.72 | F | GTPase activator activity | N/A | N/A |
| PF3D7_1211400 | PfJ4 | heat shock protein DNAJ homologue Pfj4 | 28.06 | no | 0 | 170.37 | 50.43 | 83.14 | 8.07 | 162.81 | 95.63 | 127.01 | O | 213.38 | 92.31 | M | Hsp70 protein binding;unfolded protein binding | response to heat;response to unfolded protein | Maurer's cleft;cytoplasm;host cell cytoplasm;nucleus |
| PF3D7_1302800 | N/A | 40S ribosomal protein S7, putative | 22.48 | no | 0 | 99.48 | 117.31 | 46.10 | 4.25 | 26.45 | 145.33 | 204.69 | O | 90.68 | 82.38 | M | RNA binding;structural constituent of ribosome | rRNA processing;ribosomal small subunit biogenesis;translation | cytosolic small ribosomal subunit;nucleus;small-subunit processome |
| PF3D7_1445900 | DDX17 | ATP-dependent RNA helicase DDX17 | 60.04 | no | 0 | 425.42 | 127.30 | 77.55 | 7.59 | 21.68 | 59.39 | 267.15 | R | 240.31 | 119.07 | M | ATP hydrolysis activity;RNA binding;RNA helicase activity;helicase activity;nucleic acid binding | | cytoplasm;host cell membrane;nucleus;ribonucleoprotein complex |
| PF3D7_1115700 | FP2A | cysteine proteinase falcipain 2a | 55.93 | no | 1 | 86.49 | 113.43 | 19.58 | 0.85 | 1.25 | 7.11 | 6.80 | T | 3.60 | 9.55 | F | cysteine-type endopeptidase activity;protein binding | proteolysis;proteolysis involved in cellular protein catabolic process | extracellular space;food vacuole;lysosome;membrane |
| PF3D7_0711500 | N/A | regulator of chromosome condensation, putative | 78.90 | no | 0 | 27.37 | 30.63 | 28.12 | 2.71 | 4.71 | 18.73 | 50.38 | O | 71.90 | 35.29 | M | chromatin binding | Ran protein signal transduction;chromosome condensation;chromosome segregation;regulation of mitotic spindle assembly | Maurer's cleft;cytoplasm;nucleus |
| PF3D7_0521700 | DDX1 | ATP-dependent RNA helicase DDX1, putative | 97.34 | no | 0 | 15.61 | 3.91 | 4.11 | 0.65 | 1.15 | 4.03 | 6.91 | R | 11.86 | 6.42 | M | RNA binding | rRNA processing | nucleus |
| PF3D7_0807100 | PSH3 | DNA helicase PSH3 | 157.20 | no | 0 | 16.63 | 13.93 | 9.47 | 2.40 | 4.95 | 45.14 | 34.89 | GV | 66.75 | 31.31 | M | 3'-5' DNA helicase activity;ATP hydrolysis activity;RNA binding;RNA helicase activity;single-stranded DNA helicase activity | DNA duplex unwinding | cytoplasm;nucleus |
| PF3D7_1004200 | N/A | WD repeat-containing protein, putative | 358.07 | no | 4 | 12.85 | 12.49 | 23.40 | 8.26 | 2.93 | 21.24 | 37.30 | O | 25.21 | 43.36 | F | protein binding | RNA splicing | nucleus;precatalytic spliceosome;integral component of membrane |
| PF3D7_1475600 | BDP4 | bromodomain protein 4, putative | 85.80 | no | 0 | 54.56 | 18.05 | 13.16 | 2.03 | 2.69 | 13.61 | 21.76 | R | 33.04 | 29.24 | M | protein binding | regulation of transcription | nucleus |
| PF3D7_0212900 | ATEL1 | arginyl-tRNA--protein transferase | 42.94 | no | 0 | 2.92 | 0.42 | 20.02 | 8.07 | 3.89 | 17.55 | 32.19 | O | 138.14 | 18.62 | M | arginyltransferase activity;catalytic activity;leucyltransferase activity | protein arginylation | cytoplasm |
| PF3D7_1357500 | N/A | DNA helicase, putative | 106.29 | no | 0 | 10.13 | 5.57 | 23.90 | 8.13 | 2.70 | 5.64 | 23.43 | O | 131.45 | 10.02 | M | ATP-dependent DNA/DNA annealing activity;ATP-dependent activity, acting on DNA;DNA helicase activity | DNA repair;DNA rewinding;cellular response to DNA damage stimulus;replication fork processing;replication fork protection | nuclear replication fork;nucleus |
| PF3D7_1120100 | PGM1 | phosphoglycerate mutase, putative | 28.77 | no | 0 | 1236.80 | 235.64 | 358.89 | 60.81 | 38.32 | 33.55 | 82.78 | R | 197.09 | 52.49 | M | phosphoglycerate mutase activity;protein binding | glycolytic process | Maurer's cleft;cytosol;nucleus |
| PF3D7_1242700 | N/A | 40S ribosomal protein S17, putative | 16.07 | no | 0 | 273.91 | 363.31 | 204.11 | 14.89 | 277.19 | 324.95 | 425.63 | O | 243.60 | 196.71 | M | RNA binding;structural constituent of ribosome | translation | cytosolic small ribosomal subunit;mitochondrion;nucleus |
| PF3D7_0811300 | CAF1 | CCR4-associated factor 1 | 200.21 | no | 0 | 402.20 | 70.06 | 117.56 | 38.84 | 12.49 | 69.82 | 206.03 | R | 129.59 | 48.34 | M | poly(A)-specific ribonuclease activity | exonucleolytic catabolism of deadenylated mRNA;negative regulation of translation | CCR4-NOT complex;CCR4-NOT core complex;P-body;cytosol;nucleus |
| PF3D7_0517700 | EIF3B | eukaryotic translation initiation factor 3 subunit B, putative | 84.06 | no | 0 | 162.41 | 84.94 | 55.68 | 3.89 | 11.43 | 23.64 | 136.98 | R | 62.62 | 55.88 | M | protein binding;RNA binding; translation initiation factor activity | translational initiation | eukaryotic translation initiation factor 3 complex;nucleus |
| PF3D7_0612100 | EIF3L | eukaryotic translation initiation factor 3 subunit L, putative | 78.13 | no | 2 | 117.96 | 23.59 | 30.02 | 4.76 | 1.47 | 9.55 | 19.84 | R | 31.75 | 21.04 | M | protein binding;RNA binding; translation initiation factor activity | translational initiation | eukaryotic translation initiation factor 3 complex;nucleus |
| PF3D7_1330600 | N/A | elongation factor Tu, putative | 57.63 | no | 0 | 7.98 | 6.81 | 48.77 | 8.67 | 5.87 | 31.84 | 200.76 | O | 45.31 | 199.91 | F | GTP binding;mRNA binding;translation elongation factor activity | mitochondrial translational elongation;translational elongation | mitochondrion;nucleus |
| PF3D7_1211800 | PfpUB | polyubiquitin | 14.62 | no | 0 | 69.15 | 43.1 | 111.26 | 64.84 | 45615.00 | 46.64 | 72.64 | GC II | 1283.13 | 1053.03 | M | protein tag;ubiquitin protein ligase binding | modification-dependent protein catabolic process;protein polyubiquitination;protein ubiquitination;ubiquitin-dependent protein catabolic process | cell surface;cytoplasm;nucleus;ubiquitin ligase complex |
| PF3D7_1316500 | PRP40 | pre-mRNA-processing factor 40, putative | 106.12 | no | 0 | 30.88 | 26.83 | 22.97 | 2.56 | 2.73 | 10.38 | 58.69 | O | 47.37 | 27.64 | M | protein binding;RNA binding | mRNA cis splicing, via spliceosome | spliceosomal complex;nucleus |
| PF3D7_0516200 | RPS11 | 40S ribosomal protein S11 | 16.08 | no | 0 | 390.82 | 422.05 | 175.46 | 68.46 | 193.51 | 161.36 | 117.78 | T | 110.70 | 172.59 | F | RNA binding | translation | cytosolic small ribosomal subunit;extracellular vesicle;nucleus |
| PF3D7_0807800 | RPN10 | 26S proteasome regulatory subunit RPN10, putative | 55.07 | no | 0 | 204.37 | 39.57 | 80.09 | 26.26 | 11.01 | 103.98 | 331.06 | O | 149.45 | 159.35 | F | polyubiquitin modification-dependent protein binding | proteasome assembly;proteasome-mediated ubiquitin-dependent protein catabolic process | cytosol;nucleus;proteasome regulatory particle, base subcomplex |
| PF3D7_0111400 | GEXP19 | conserved Plasmodium protein, unknown function | 85.04 | no | 0 | 3.88 | 1.36 | 6.44 | 1.12 | 3.06 | 30.24 | 186.57 | O | 127.40 | 91.28 | M | N/A | N/A | N/A |
| PF3D7_0823900 | DTC | dicarboxylate/tricarboxylate carrier | 35.39 | no | 1 | 9.82 | 7.89 | 16.76 | 3.54 | 9.66 | 10.80 | 34.54 | O | 57.97 | 24.92 | M | dicarboxylic acid transmembrane transporter activity;oxoglutarate:malate antiporter activity | alpha-ketoglutarate transport;dicarboxylic acid transport;malate transport;mitochondrial transport | mitochondrial inner membrane;mitochondrion |
| PF3D7_1342000 | N/A | 40S ribosomal protein S6 | 35.38 | no | 0 | 373.69 | 306.10 | 149.68 | 19.30 | 14.43 | 36.14 | 266.82 | R | 391.35 | 213.34 | M | structural constituent of ribosome | translation | cytosolic small ribosomal subunit;nucleus |
| PF3D7_0414000 | SMC3 | structural maintenance of chromosomes protein 3 | 141.22 | no | 0 | 4.75 | 6.70 | 16.43 | 2.39 | 4.70 | 42.00 | 98.77 | O | 167.87 | 111.69 | M | ATP binding;centromeric DNA binding;chromatin binding;minor groove of adenine-thymine-rich DNA binding;protein binding | chromosome organization;mitotic sister chromatid segregation;regulation of DNA replication | chromatin;chromosome, centromeric region;cohesin complex;nuclear chromosome;nuclear mitotic cohesin complex;nuclear periphery;nucleus |
| PF3D7_1122400 | N/A | conserved Plasmodium protein, unknown function | 101.94 | no | 0 | 47.66 | 9.44 | 11.77 | 1.14 | 2.32 | 19.68 | 29.12 | R | 48.56 | 27.06 | M | N/A | N/A | N/A |
| PF3D7_0613500 | N/A | AP-3 complex subunit beta, putative | 159.77 | no | 0 | 4.65 | 4.31 | 7.60 | 2.11 | 3.13 | 33.52 | 61.38 | O | 38.51 | 42.08 | 0 | N/A | vesicle-mediated transport | AP-3 adaptor complex;coated vesicle |
| PF3D7_0422400 | RPS19 | 40S ribosomal protein S19 | 19.72 | no | 0 | 265.03 | 189.45 | 98.34 | 8.91 | 81.63 | 161.57 | 229.58 | R | 80.80 | 83.79 | 0 | RNA binding;structural constituent of ribosome | ribosomal small subunit assembly;translation | cytosolic small ribosomal subunit;extracellular vesicle;nucleus |
| PF3D7_1329300 | CAF1B | chromatin assembly factor 1 subunit B, putative | 66.51 | no | 0 | 7.86 | 1.35 | 36.46 | 6.21 | 8.40 | 23.98 | 40.73 | O | 143.86 | 25.16 | M | chromatin binding;histone binding;protein binding;unfolded protein binding | chromatin assembly;chromatin assembly or disassembly | nucleus |
| PF3D7_1359600 | N/A | conserved Plasmodium protein, unknown function | 235.14 | no | 0 | 7.69 | 49.44 | 30.86 | 4.06 | 2.98 | 18.14 | 35.41 | T | 20.51 | 32.19 | F | N/A | N/A | nucleus |
| PF3D7_1403100 | CAPD2 | condensin complex subunit 1, putative | 229.53 | no | 1 | 8.74 | 5.50 | 41.88 | 13.10 | 15.82 | 33.23 | 615.29 | O | 256.00 | 109.38 | M | chromatin binding;histone binding | chromosome separation;meiotic chromosome condensation;mitotic chromosome condensation | condensed chromosome, centromeric region;condensin complex |
| PF3D7_0315100 | eIF4E | eukaryotic translation initiation factor 4E | 26.95 | no | 0 | 32.14 | 15.40 | 12.51 | 2.04 | 13.42 | 19.80 | 48.77 | O | 15.76 | 22.43 | F | RNA 7-methylguanosine cap binding;RNA cap binding;mRNA cap binding;protein binding;translation initiation factor activity | translational initiation | P granule;eukaryotic translation initiation factor 4F complex;nucleus |
| PF3D7_0302100 | SRPK1 | serine/threonine protein kinase | 157.37 | no | 0 | 38.29 | 33.71 | 24.41 | 2.20 | 3.78 | 26.91 | 68.03 | O | 259.41 | 38.10 | M | MAP kinase activity;protein serine/threonine kinase activity | intracellular signal transduction;protein phosphorylation;regulation of gene expression;regulation of mRNA processing;spliceosomal complex assembly | cytoplasm;nucleus |
| PF3D7_0520000 | RPS9 | 40S ribosomal protein S9, putative | 22.12 | no | 0 | 587.12 | 471.77 | 200.53 | 62.55 | 168.15 | 92.75 | 299.37 | R | 111.98 | 129.99 | 0 | RNA binding;rRNA binding;structural constituent of ribosome | positive regulation of translational fidelity | RNA N6-methyladenosine methyltransferase complex;cytosolic small ribosomal subunit;nucleus;small ribosomal subunit;small-subunit processome |
| PF3D7_0801800 | N/A | mannose-6-phosphate isomerase, putative | 117.66 | no | 1 | 151.07 | 351.16 | 97.39 | 17.91 | 8.38 | 28.18 | 104.44 | T | 105.15 | 38.70 | M | mannose-6-phosphate isomerase activity;zinc ion binding | GDP-mannose biosynthetic process;cell wall mannoprotein biosynthetic process;protein glycosylation | cytosol;nucleus |
| PF3D7_0709000 | CRT | chloroquine resistance transporter | 48.68 | no | 10 | 248.36 | 96.71 | 31.15 | 3.85 | 13.44 | 30.89 | 70.08 | R | 20.67 | 8.17 | M | ferric iron transmembrane transporter activity;ferrous iron transmembrane transporter activity;xenobiotic transmembrane transporter activity | glutathione transport;peptide transport;response to chloroquine;response to drug;response to xenobiotic stimulus | endoplasmic reticulum;food vacuole;membrane |
| PF3D7_1324900 | LDH | L-lactate dehydrogenase | 34.11 | no | 1 | 1444.93 | 553.00 | 1021.46 | 115.77 | 63.88 | 44.10 | 73.22 | R | 173.83 | 18.00 | M | L-lactate dehydrogenase activity;L-malate dehydrogenase activity;RNA binding;protein binding | glycolytic process | cell surface;cytoplasm;cytosol;extracellular vesicle;food vacuole;intracellular;mitochondrion;nucleus |
| PF3D7_1141800 | N/A | EELM2 domain-containing protein, putative | 46.67 | no | 0 | 6.41 | 0.52 | 39.04 | 28.37 | 3.98 | 19.09 | 46.26 | O | 6.31 | 33.32 | F | transcription corepressor activity;metal ion binding | negative regulation of transcription, DNA-templated | nucleus |
| PF3D7_1368100 | RPN11 | 26S proteasome regulatory subunit RPN11, putative | 35.21 | no | 0 | 50.34 | 39.56 | 135.99 | 77.42 | 20.37 | 19.41 | 96.83 | T | 31.30 | 38.72 | F | metallopeptidase activity;proteasome binding;thiol-dependent deubiquitinase | proteasome-mediated ubiquitin-dependent protein catabolic process;protein deubiquitination;ubiquitin-dependent protein catabolic process | nucleus;proteasome regulatory particle;proteasome regulatory particle, lid subcomplex |
| PF3D7_0910400 | N/A | selenide water dikinase, putative | 140.32 | no | 0 | 9.78 | 7.06 | 10.13 | 4.23 | 1.80 | 10.68 | 13.51 | O | 18.60 | 15.82 | 0 | selenide, water dikinase activity | selenocysteine biosynthetic process | cytoplasm;nucleus |
| PF3D7_0905900 | SEC27 | coatomer subunit beta, putative | 118.24 | no | 0 | 63.10 | 8.78 | 17.00 | 2.75 | 1.45 | 13.95 | 50.03 | R | 29.33 | 21.01 | M | protein binding | endoplasmic reticulum to Golgi vesicle-mediated transport;intra-Golgi vesicle-mediated transport;intracellular protein transport;retrograde vesicle-mediated transport, Golgi to endoplasmic reticulum | COPI vesicle coat |

| | | | | | | | | | | | | | | | | | | | |
|---|---|---|---|---|---|---|---|---|---|---|---|---|---|---|---|---|---|---|---|
| PF3D7_0918300 | EIF3F | eukaryotic translation initiation factor 3 subunit F, putative | 36.82 | no | 0 | 123.68 | 44.90 | 31.80 | 1.91 | 16.45 | 29.15 | 19.08 | R | 63.08 | 23.22 | M | translation initiation factor activity;translation initiation factor binding | translational initiation | eukaryotic translation initiation factor 3 complex;eukaryotic translation initiation factor 3 complex, eIF3m;nucleus |
| PF3D7_0822600 | SEC23 | protein transport protein SEC23 | 86.17 | no | 0 | 247.51 | 37.65 | 132.15 | 42.15 | 14.38 | 36.58 | 51.94 | R | 73.80 | 120.56 | F | GTPase activator activity | COPII-coated vesicle cargo loading;endoplasmic reticulum to Golgi vesicle-mediated transport | COPII vesicle coat;Maurer's cleft;endoplasmic reticulum exit site |
| PF3D7_0501300 | SBP1 | skeleton-binding protein 1 | 36.30 | no | 1 | 5630.61 | 1039.83 | 114.73 | 15.29 | 24.32 | 11.48 | 125.20 | R | 3.79 | 2.12 | 0 | protein binding | protein transport | Maurer's cleft;cytoplasm;extracellular vesicle;integral component of membrane;membrane;nucleus;symbiont-containing vacuolar space;symbiont-containing vacuole membrane |
| PF3D7_0811600 | MAP1 | protein MAP1, putative | 144.01 | yes | 0 | 4.92 | 2.93 | 32.26 | 15.04 | 5.94 | 23.45 | 13.07 | T | 63.90 | 9.75 | M | RNA binding | microtubuli complex | cell surface;nucleus;plasma membrane |
| PF3D7_1364100 | P92 | 6-cysteine protein | 29.86 | yes | 1 | 28642.00 | 0.42 | 55.06 | 13.78 | 36161.00 | 22.26 | 41.68 | GC II | 19845.00 | 44.31 | M | protein binding | cell adhesion | anchored component of plasma membrane;cell surface |
| PF3D7_1347100 | TOP3 | DNA topoisomerase 3 | 84.31 | no | 0 | 7.65 | 2.04 | 18.95 | 6.61 | 3.85 | 75.60 | 116.55 | O | 69.75 | 157.00 | F | DNA binding;DNA topoisomerase activity;protein binding | DNA topological change;DNA unwinding involved in DNA replication | mitochondrion;nucleus |
| PF3D7_0719300 | ARP6 | actin-related protein ARP6 | 121.13 | no | 0 | 67.57 | 25.65 | 22.10 | 2.16 | 4.58 | 24.11 | 558.24 | O | 77.78 | 90.84 | F | nucleosome binding;protein binding | histone exchange | Swr1 complex;nucleus |
| PF3D7_1446200 | LAP | M17 leucyl aminopeptidase | 67.82 | no | 0 | 476.97 | 60.11 | 58.18 | 7.19 | 18.06 | 48.48 | 57.01 | R | 12.78 | 0.41 | M | aminopeptidase activity;identical protein binding;metal ion binding;metalloaminopeptidase activity | proteolysis | apicoplast;cytosol;food vacuole |
| PF3D7_0933000 | N/A | CSTF domain-containing protein, putative | 88.93 | no | 0 | 67.99 | 16.78 | 17.78 | 3.75 | 2.26 | 18.44 | 52.44 | R | 117.99 | 23.74 | M | mRNA binding | pre-mRNA cleavage required for polyadenylation | mRNA cleavage and polyadenylation specificity factor complex;nucleus |
| PF3D7_1324100 | N/A | cyclin-like protein, putative | 61.55 | no | 5 | 4.61 | 0.88 | 15.75 | 5.26 | 2.67 | 10.38 | 8.13 | T | 10.20 | 3.10 | M | N/A | N/A | integral component of membrane;nucleus |
| PF3D7_1408400 | FANCJ | FANCJ-like helicase, putative | 135.54 | no | 0 | 10.87 | 14.94 | 10.88 | 2.72 | 0.98 | 13.05 | 43.48 | O | 278.60 | 10.44 | M | ATP binding;DNA helicase activity;DNA polymerase binding | DNA duplex unwinding;negative regulation of DNA recombination;negative regulation of t-circle formation;regulation of double-strand break repair via homologous recombination;telomeric loop disassembly | mitochondrion;nucleus |
| PF3D7_0722400 | OLA1 | Obg-like ATPase 1, putative | 45.16 | no | 0 | 196.76 | 106.14 | 119.82 | 19.13 | 6.13 | 6.39 | 85.92 | R | 12.21 | 6.80 | M | ATP hydrolysis activity;GTP binding | signal transduction | cytoplasm;nucleus |
| PF3D7_0321500 | N/A | peptidase, putative | 125.89 | no | 0 | 6.20 | 4.40 | 18.98 | 3.75 | 2.02 | 35.74 | 30.93 | GV | 19.39 | 100.30 | F | peptidase activity;serine-type endopeptidase activity | N/A | nucleus |
| PF3D7_1026800 | RPS2 | 40S ribosomal protein S2 | 29.86 | no | 0 | 1651.31 | 969.41 | 501.51 | 98.27 | 64.09 | 263.46 | 1652.19 | R | 123.54 | 290.32 | F | RNA binding;structural constituent of ribosome | ribosomal small subunit assembly;translation | ribosome;small ribosomal subunit |
| PF3D7_0929200 | N/A | RNA-binding protein, putative | 33.60 | no | 0 | 69.72 | 44.30 | 35.57 | 6.32 | 27.39 | 44.17 | 191.66 | O | 143.91 | 48.01 | M | mRNA binding | N/A | cytoplasm;nucleus |
| PF3D7_0829000 | N/A | conserved Plasmodium membrane protein, unknown function | 228.81 | no | 5 | 13.07 | 6.67 | 10.29 | 2.20 | 2.31 | 15.23 | 47.85 | O | 31.25 | 14.56 | M | N/A | N/A | integral component of membrane |
| PF3D7_0813300 | N/A | NPL domain-containing protein, putative | 36.26 | no | 0 | 511.86 | 1801.83 | 170.55 | 6.64 | 91.14 | 79.72 | 270.71 | T | 189.00 | 282.59 | F | RNA binding | N/A | food vacuole;nucleolus;nucleus |
| PF3D7_0405700 | N/A | amino acid decarboxylase AAD, putative | 280.73 | no | 0 | 8.39 | 7.94 | 6.26 | 0.58 | 1.88 | 16.23 | 102.43 | O | 74.43 | 28.70 | M | lysine decarboxylase activity | lysine catabolic process | cytoplasm |
| PF3D7_1010200 | N/A | DNA2/NAM7 helicase, putative | 217.16 | no | 0 | 185.97 | 60.34 | 12.11 | 0.93 | 2.12 | 12.33 | 25.60 | R | 13.03 | 5.14 | M | RNA binding;RNA helicase activity | nuclear-transcribed mRNA catabolic process, nonsense-mediated decay | cytoplasm;nucleus |
| PF3D7_0807600 | N/A | conserved Plasmodium protein, unknown function | 159.03 | no | 0 | 143.64 | 53.18 | 34.07 | 5.28 | 5.87 | 18.43 | 63.24 | R | 64.44 | 55.06 | M | N/A | N/A | nucleus |
| PF3D7_0807500 | N/A | proteasome subunit alpha type-6, putative | 29.50 | no | 0 | 34.26 | 17.51 | 57.18 | 10.35 | 8.95 | 24.86 | 79.68 | O | 50.95 | 62.74 | F | endopeptidase activity | proteasomal protein catabolic process;proteasomal ubiquitin-independent protein catabolic process;proteasome-mediated ubiquitin-dependent protein catabolic process;ubiquitin-dependent protein catabolic process | cytoplasm;nucleus;proteasome core complex;proteasome core complex, alpha-subunit complex |
| PF3D7_1110700 | ALP1 | actin-like protein, putative | 42.48 | no | 0 | 7.09 | 0.34 | 31.59 | 34.77 | 14.52 | 31.17 | 117.12 | O | 13.62 | 44.76 | F | protein binding | cytoskeleton organization; actin filament organization; actin filament-based movement; vesicle transport along actin filament | nucleus;cytoplasm |
| PF3D7_1341200 | N/A | 60S ribosomal protein L18, putative | 21.75 | no | 0 | 311.87 | 221.18 | 108.15 | 16.90 | 60.88 | 83.53 | 136.74 | R | 55.32 | 55.51 | 0 | structural constituent of ribosome | translation | cytosolic large ribosomal subunit;nucleus |
| PF3D7_1345700 | IDH | isocitrate dehydrogenase [NADP], mitochondrial | 53.59 | no | 0 | 21.79 | 14.16 | 51.14 | 17.21 | 4.32 | 8.80 | 25.36 | T | 39.99 | 44.92 | F | isocitrate dehydrogenase (NADP+) activity | NADP metabolic process;isocitrate metabolic process;tricarboxylic acid cycle | mitochondrion;nucleus |
| PF3D7_0924400 | N/A | conserved Plasmodium protein, unknown function | 144.68 | no | 0 | 41.70 | 18.03 | 49.95 | 9.80 | 16.39 | 60.81 | 557.61 | O | 270.77 | 95.40 | M | N/A | N/A | N/A |
| PF3D7_1302100 | G27/25 | gamete antigen 27/25 | 26.00 | no | 0 | 41.03 | 348.81 | 77.43 | 17.89 | 1238.79 | 648.85 | 268.30 | G II | 293.47 | 343.37 | F | protein dimerization activity | N/A | nucleus |
| PF3D7_0218700 | PRP45 | pre-mRNA-processing protein 45, putative | 57.53 | no | 0 | 35.23 | 18.41 | 10.35 | 1.41 | 4.59 | 27.48 | 164.60 | O | 233.68 | 76.18 | M | RNA binding | RNA processing | nucleus |
| PF3D7_0523000 | MDR1 | multidrug resistance protein 1 | 162.25 | no | 11 | 291.72 | 142.99 | 52.35 | 5.94 | 4.61 | 25.10 | 131.76 | R | 28.73 | 199.61 | F | ATPase-coupled transmembrane transporter activity;protein binding | response to drug;response to xenobiotic stimulus;transmembrane transport | food vacuole |
| PF3D7_1247400 | FKBP35 | peptidyl-prolyl cis-trans isomerase FKBP35 | 34.83 | no | 0 | 17.76 | 32.59 | 23.49 | 3.13 | 2.66 | 9.09 | 35.88 | O | 85.93 | 16.16 | M | FK506 binding;peptidyl-prolyl cis-trans isomerase activity;protein dimerization activity | chaperone-mediated protein folding;negative regulation of phosphoprotein phosphatase activity;protein folding;protein peptidyl-prolyl isomerization;response to xenobiotic stimulus | cytoplasm;nucleus |
| PF3D7_0727400 | N/A | proteasome subunit alpha type-5, putative | 28.39 | no | 0 | 168.66 | 69.91 | 228.43 | 89.03 | 56.18 | 47.95 | 89.49 | T | 22.26 | 33.70 | 0 | endopeptidase activity | proteasomal protein catabolic process;proteasomal ubiquitin-independent protein catabolic process;proteasome-mediated ubiquitin-dependent protein catabolic process;ubiquitin-dependent protein catabolic process | cytoplasm;extracellular vesicle;nucleus;proteasome core complex;proteasome core complex, alpha-subunit complex |
| PF3D7_1124300 | BDP7 | bromodomain protein 7 | 62.33 | no | 0 | 64.40 | 69.11 | 38.46 | 10.73 | 3.55 | 14.52 | 360.42 | O | 73.69 | 26.80 | M | chromatin binding;histone binding;protein binding | regulation of transcription | heterochromatin;nucleus |
| PF3D7_1145200 | N/A | serine/threonine protein kinase, putative | 238.76 | no | 0 | 14.70 | 1.72 | 82.11 | 115.78 | 12.23 | 165.29 | 1299.69 | O | 74.52 | 1053.09 | F | enzymatic activity | signal transduction | nucleus |
| PF3D7_1135900 | N/A | 3-oxo-5-alpha-steroid 4-dehydrogenase, putative | 35.47 | no | 5 | 26.16 | 2.60 | 57.58 | 23.88 | 13.04 | 115.06 | 126.58 | O | 60.52 | 165.09 | F | oxidoreductase activity | very long-chain fatty acid biosynthetic process | cytoplasm |
| PF3D7_0613700 | N/A | syntaxin-binding protein, putative | 82.77 | no | 0 | 10.68 | 10.44 | 21.72 | 10.42 | 5.22 | 55.31 | 206.15 | O | 25.80 | 143.86 | F | syntaxin binding | intracellular protein transport;protein secretion;vesicle docking involved in exocytosis;vesicle-mediated transport | plasma membrane;secretory granule |
| PF3D7_1420600 | PANK1 | pantothenate kinase 1 | 59.92 | no | 0 | 21.25 | 8.12 | 13.60 | 2.82 | 3.07 | 9.96 | 2.94 | R | 20.59 | 0.79 | M | 14-3-3 protein binding;pantothenate kinase activity;protein binding | coenzyme A biosynthetic process;response to xenobiotic stimulus | cytosol;nucleus |
| PF3D7_1006200 | ALBA3 | DNA/RNA-binding protein Alba 3 | 11.98 | no | 0 | 829.33 | 497.94 | 424.34 | 49.27 | 361.79 | 261.75 | 278.91 | R | 61.84 | 77.63 | F | DNA binding;RNA binding;endonuclease activity;mRNA binding;protein binding | | RNA N6-methyladenosine methyltransferase complex;extracellular vesicle;food vacuole;nucleus |
| PF3D7_1323400 | RPL23 | 60S ribosomal protein L23 | 22.09 | no | 0 | 294.72 | 206.52 | 100.00 | 19.62 | 40.97 | 57.18 | 166.47 | R | 97.25 | 39.60 | M | rRNA binding;structural constituent of ribosome | ribosomal large subunit assembly;translation | cytosolic large ribosomal subunit;nucleus |
| PF3D7_0907400 | ClpY | ATP-dependent protease ATPase subunit ClpY | 106.46 | no | 0 | 56.06 | 31.45 | 56.78 | 15.98 | 6.00 | 22.76 | 225.56 | O | 90.55 | 39.48 | M | mRNA binding;protein binding | N/A | mitochondrion;nucleus |
| PF3D7_1103100 | RPP1 | 60S acidic ribosomal protein P1, putative | 13.01 | no | 0 | 1624.59 | 1493.07 | 452.53 | 198.66 | 1479.73 | 2641.06 | 1067.62 | G V | 273.89 | 1049.36 | F | protein kinase activator activity;ribonucleoprotein complex binding;structural constituent of ribosome | cytoplasmic translation;translation | cytosolic large ribosomal subunit;large ribosomal subunit;nucleus |
| PF3D7_0319400 | N/A | kinesin-8X | 157.39 | no | 0 | 6.22 | 0.40 | 16.49 | 2.91 | 5.70 | 31.08 | 49.68 | O | 293.33 | 45.04 | M | ATP hydrolysis activity;microtubule binding;microtubule motor activity;plus-end-directed microtubule motor activity | microtubule depolymerization;microtubule-based movement | kinesin complex;microtubule;mitotic spindle;nucleus |
| PF3D7_1364200 | NUP205 | nucleoporin NUP205, putative | 207.29 | no | 0 | 28.39 | 6.64 | 9.97 | 2.03 | 1.15 | 9.24 | 17.17 | R | 22.05 | 8.27 | M | protein binding | nucleocytoplasmic transport | nuclear pore;nucleus |
| PF3D7_0307100 | RPS12 | 40S ribosomal protein S12, putative | 15.40 | no | 0 | 323.64 | 393.50 | 185.85 | 22.52 | 150.04 | 91.36 | 125.32 | T | 80.85 | 104.89 | F | structural constituent of ribosome | translation | cytosolic small ribosomal subunit;nucleus |
| PF3D7_1346400 | N/A | VPS13 domain-containing protein, putative | 697.04 | no | 0 | 44.58 | 16.01 | 22.43 | 6.53 | 3.75 | 37.64 | 167.02 | O | 42.82 | 59.33 | F | protein binding | protein retention in Golgi apparatus;protein targeting to vacuole | extrinsic component of membrane |
| PF3D7_1369700 | SNRPA1 | U2 small nuclear ribonucleoprotein A', putative | 20.42 | no | 0 | 60.45 | 7.55 | 5.27 | 1.24 | 16.53 | 34.73 | 18.06 | R | 13.52 | 9.74 | M | RNA binding | mRNA splicing, via spliceosome | spliceosomal complex |
| PF3D7_1008800 | NOP5 | nucleolar protein 5, putative | 53.41 | no | 0 | 127.27 | 62.97 | 23.33 | 1.00 | 10.00 | 34.74 | 182.09 | O | 98.41 | 141.36 | F | RNA binding;snoRNA binding | rRNA processing | box C/D RNP complex;nucleolus;nucleus;small-subunit processome |
| PF3D7_1456000 | AP2-HC | AP2 domain transcription factor AP2-HC | 161.51 | no | 0 | 4.11 | 0.68 | 15.05 | 3.24 | 2.56 | 11.86 | 30.98 | O | 82.54 | 8.94 | M | protein binding;sequence-specific DNA binding | regulation of transcription | heterochromatin;nucleus |
| PF3D7_1018200 | PPP8 | pseudophosphatase PPP8 | 254.11 | no | 0 | 16.40 | 1.78 | 46.07 | 79.30 | 6.66 | 48.53 | 259.74 | O | 7.75 | 184.96 | F | protein binding;protein serine/threonine phosphatase activity | | basal part of cell;cytoplasm;nucleus;protein-containing complex |
| PF3D7_1319400 | N/A | conserved protein, unknown function | 127.67 | no | 0 | 13.32 | 4.96 | 40.99 | 8.88 | 7.60 | 40.04 | 239.32 | O | 1028.50 | 132.26 | M | N/A | N/A | nucleus |
| PF3D7_1213200 | MED18 | mediator of RNA polymerase II transcription subunit 18, putative | 33.25 | no | 0 | 10.89 | 5.18 | 7.41 | 1.12 | 4.98 | 37.90 | 11.42 | GV | 13.84 | 25.14 | F | protein binding | regulation of transcription | mediator complex;nucleus |
| PF3D7_0806700 | N/A | conserved Plasmodium membrane protein, unknown function | 307.60 | no | 4 | 31.25 | 5.25 | 8.78 | 2.09 | 2.96 | 25.05 | 32.98 | O | 70.93 | 34.69 | M | N/A | N/A | N/A |
| PF3D7_1349200 | GluRS | glutamate--tRNA ligase | 101.42 | no | 0 | 177.94 | 42.40 | 38.72 | 6.00 | 3.73 | 25.33 | 140.01 | R | 25.32 | 29.44 | F | ATP binding;glutamate-tRNA ligase activity;glutamine-tRNA ligase activity | glutaminyl-tRNA aminoacylation;glutamyl-tRNA aminoacylation | cytosol;nucleus |
| PF3D7_1019400 | N/A | 60S ribosomal protein L30e, putative | 11.78 | no | 0 | 151.18 | 124.64 | 63.22 | 5.98 | 57.61 | 81.27 | 208.06 | O | 65.43 | 46.29 | M | RNA binding;structural constituent of ribosome | translation | cytosolic large ribosomal subunit;nucleus |
| PF3D7_0721600 | N/A | 40S ribosomal protein S5, putative | 21.86 | no | 0 | 1392.72 | 990.21 | 392.28 | 48.67 | 248.03 | 145.91 | 432.69 | R | 235.02 | 245.98 | F | RNA binding;mRNA binding;rRNA binding;structural constituent of ribosome | ribosomal small subunit assembly;translation | cytosolic small ribosomal subunit;nucleus;ribosome |
| PF3D7_1037500 | DYN2 | dynamin-like protein | 81.53 | no | 0 | 3.60 | 0.50 | 10.53 | 6.30 | 1.48 | 19.73 | 51.41 | O | 11.47 | 19.22 | F | GTPase activity;microtubule binding | mitochondrial fission;organelle fission | Golgi apparatus;apicoplast;cytoplasm;endoplasmic reticulum;membrane |
| PF3D7_1119400 | N/A | ubiquitin-protein ligase, putative | 256.82 | no | 2 | 24.82 | 10.01 | 13.14 | 2.89 | 1.23 | 14.73 | 129.00 | O | 177.54 | 45.73 | M | ubiquitin conjugating enzyme activity | protein polyubiquitination;ubiquitin-dependent protein catabolic process | |
| PF3D7_1414400 | PP1 | serine/threonine protein phosphatase PP1 | 34.90 | no | 0 | 22.21 | 26.86 | 170.82 | 90.06 | 50.23 | 195.76 | 310.63 | O | 155.58 | 92.23 | M | protein binding;protein serine/threonine phosphatase activity | entry into host;exit from host cell;protein dephosphorylation | Maurer's cleft;cell surface;cytoplasm;nucleus |
| PF3D7_0909400 | N/A | exoribonuclease, putative | 154.96 | no | 0 | 77.12 | 29.14 | 29.26 | 4.33 | 2.38 | 27.46 | 31.23 | R | 31.25 | 87.57 | F | 5'-3' exoribonuclease activity;RNA binding | nuclear-transcribed mRNA catabolic process | nucleus |
| PF3D7_0923900 | PABP2 | polyadenylate-binding protein 2, putative | 22.98 | no | 0 | 444.51 | 140.03 | 100.20 | 26.94 | 127.43 | 194.13 | 154.95 | R | 95.85 | 77.53 | M | RNA binding;poly(A) binding | posttranscriptional regulation | cytosol;nucleus |
| PF3D7_0626000 | N/A | conserved Plasmodium protein, unknown function | 209.43 | no | 0 | 38.26 | 20.15 | 6.90 | 1.22 | 1.46 | 61.20 | 7.91 | GV | 9.26 | 41.16 | F | N/A | N/A | N/A |
| PF3D7_1460800 | SNAPC3 | snRNA-activating protein complex subunit 3, putative | 75.93 | no | 0 | 5.30 | 3.41 | 5.87 | 1.86 | 6.42 | 18.05 | 109.34 | O | 68.12 | 32.63 | M | RNA polymerase III type 3 promoter sequence-specific DNA binding;bent DNA binding | snRNA transcription by RNA polymerase II;snRNA transcription by RNA polymerase III | nucleus;snRNA-activating protein complex |
| PF3D7_1019700 | N/A | conserved Plasmodium protein, unknown function | 128.21 | no | 0 | 7.24 | 2.36 | 9.76 | 3.61 | 2.82 | 19.95 | 217.76 | O | 199.14 | 339.55 | F | N/A | N/A | histone acetyltransferase complex;nucleus |
| PF3D7_0725300 | N/A | WD repeat-containing protein, putative | 93.45 | no | 0 | 113.99 | 75.72 | 14.43 | 2.86 | 4.48 | 5.01 | 63.19 | R | 11.10 | 7.02 | M | protein binding | ribosomal large subunit biogenesis | nucleolus;nucleus;preribosome, large subunit precursor |
| PF3D7_1455700 | N/A | conserved Plasmodium protein, unknown function | 126.12 | no | 0 | 3.90 | 0.78 | 9.91 | 3.03 | 2.84 | 14.22 | 45.05 | O | 33.18 | 17.59 | M | N/A | N/A | nucleus |
| PF3D7_0903300 | N/A | conserved protein, unknown function | 380.98 | no | 6 | 3.96 | 0.52 | 29.82 | 8.95 | 3.99 | 21.76 | 49.15 | O | 145.53 | 27.55 | M | N/A | N/A | integral component of membrane |
| PF3D7_0730300 | ApiAP2 | AP2 domain transcription factor AP2-L, putative | 155.13 | no | 0 | 227.37 | 137.47 | 40.04 | 3.01 | 1.76 | 9.32 | 83.57 | R | 53.53 | 9.26 | M | DNA binding | regulation of transcription | nucleus |
| PF3D7_1033500 | WDR70 | WD repeat-containing protein 70, putative | 83.23 | no | 0 | 39.56 | 10.58 | 12.49 | 3.79 | 2.37 | 42.59 | 107.81 | O | 30.97 | 101.63 | F | protein binding | regulation of DNA double-strand break processing;regulation of histone H2B conserved C-terminal lysine ubiquitination | nucleus;site of double-strand break |
| PF3D7_0312300 | RPN12 | 26S proteasome regulatory subunit RPN12, putative | 35.10 | no | 0 | 33.67 | 18.77 | 33.87 | 11.50 | 59.24 | 150.20 | 36.40 | GV | 110.44 | 37.74 | M | protein binding | proteasome-mediated ubiquitin-dependent protein catabolic process;ubiquitin-dependent protein catabolic process | cytosol;intracellular;nucleus;proteasome regulatory particle;proteasome regulatory particle, lid subcomplex |
| PF3D7_0629100 | NAPRT | nicotinate phosphoribosyltransferase, putative | 77.13 | no | 0 | 24.40 | 14.71 | 19.00 | 5.35 | 1.36 | 23.13 | 117.81 | O | 6.53 | 72.99 | F | nicotinate phosphoribosyltransferase activity | NAD biosynthetic process;NAD salvage | cytoplasm;cytosol |
| PF3D7_1136300 | TSN | tudor staphylococcal nuclease | 128.54 | no | 0 | 248.19 | 19.54 | 31.74 | 12.59 | 6.70 | 22.89 | 57.62 | R | 65.82 | 108.22 | F | RNA binding;calcium ion binding;nuclease activity;protein binding | RNA catabolic process | cytoplasm;nucleus |
| PF3D7_1326400 | eIF-2Bg | translation initiation factor eIF-2B subunit gamma, putative | 55.04 | no | 0 | 58.95 | 8.89 | 11.95 | 0.98 | 5.61 | 33.96 | 14.58 | R | 50.42 | 13.16 | M | protein binding | translational initiation | eukaryotic translation initiation factor 2B complex |
| PF3D7_0102500 | EBA181 | erythrocyte binding antigen-181 | 181.15 | yes | 2 | 10.91 | 0.66 | 28.65 | 7.13 | 2.24 | 4.87 | 136.60 | O | 4.50 | 0.08 | M | heparin binding;sialic acid binding | cell-cell adhesion;entry into host | extracellular vesicle;microneme |
| PF3D7_0918000 | GAP50 | glideosome-associated protein 50 | 44.60 | no | 2 | 17.85 | 1.74 | 201.56 | 118.61 | 48.52 | 77.83 | 248.51 | O | 72.91 | 77.35 | F | RNA binding;acid phosphatase activity;ferric iron binding;ferrous iron binding;protein binding;protein-membrane adaptor activity | | cell surface;endoplasmic reticulum;extracellular vesicle;food vacuole;inner membrane pellicle complex;nucleus;plasma membrane |
| PF3D7_1419900 | YTH1 | YTH domain-containing protein 1, putative | 67.17 | no | 0 | 26.52 | 5.62 | 6.22 | 0.53 | 3.82 | 26.27 | 6.89 | G V | 42.05 | 16.95 | M | N6-methyladenosine-containing RNA binding;mRNA binding | mRNA splicing, via spliceosome;regulation of mRNA splicing, via spliceosome | cytoplasm;nucleoplasm;nucleus |
| PF3D7_0527600 | N/A | conserved Plasmodium protein, unknown function | 52.97 | no | 0 | 9.83 | 11.53 | 30.94 | 6.43 | 5.55 | 47.32 | 116.42 | O | 44.48 | 184.91 | F | N/A | N/A | nucleus |
| PF3D7_0527500 | HIP | Hsc70-interacting protein | 51.13 | no | 0 | 818.66 | 971.48 | 631.26 | 656.09 | 90.01 | 59.83 | 311.89 | T | 290.55 | 320.92 | F | heat shock protein binding | chaperone cofactor-dependent protein refolding;chaperone-mediated protein folding;protein-containing complex assembly | nucleus |
| PF3D7_0919000 | NAPS | nucleosome assembly protein | 31.83 | no | 0 | 541.25 | 142.40 | 251.29 | 39.57 | 72.58 | 65.00 | 248.73 | R | 357.25 | 45.25 | M | chromatin binding;histone binding | | cytoplasm;histone acetyltransferase complex;nuclear periphery;nucleus |
| PF3D7_1438500 | CPSF3 | cleavage and polyadenylation specificity factor subunit 3, putative | 101.18 | no | 0 | 24.85 | 6.74 | 6.28 | 0.51 | 2.22 | 4.01 | 16.95 | R | 21.65 | 11.06 | M | 5'-3' exonuclease activity;RNA binding;endoribonuclease activity | mRNA 3'-end processing by stem-loop binding and cleavage;mRNA cleavage;mRNA polyadenylation;response to xenobiotic stimulus | mRNA cleavage and polyadenylation specificity factor complex;nucleus |
| PF3D7_0515700 | GAP40 | glideosome-associated protein 40 | 51.80 | no | 10 | 20.90 | 2.83 | 401.41 | 370.69 | 100.25 | 68.52 | 315.76 | T | 110.67 | 181.50 | F | protein binding | cell gliding;entry into host | cell periphery;cell surface;cytoplasm;cytosol;inner membrane pellicle complex |
| PF3D7_1364400 | N/A | conserved Plasmodium protein, unknown function | 386.33 | no | 2 | 27.67 | 21.32 | 7.00 | 1.10 | 2.35 | 10.21 | 21.20 | R | 18.36 | 6.72 | M | N/A | N/A | histone acetyltransferase complex;nucleus;cytoplasm |
| PF3D7_1014300 | N/A | SPRY domain-containing protein, putative | 283.13 | no | 8 | 7.90 | 7.20 | 35.86 | 8.75 | 7.47 | 45.11 | 54.03 | O | 728.81 | 139.25 | M | N/A | proteasome-mediated ubiquitin-dependent protein catabolic process | cytoplasm;membrane;nucleus |
| PF3D7_1360400 | N/A | conserved Plasmodium protein, unknown function | 36.77 | no | 0 | 12.99 | 8.33 | 4.72 | 0.13 | 15.24 | 59.82 | 5.96 | GV | 27.71 | 22.35 | M | N/A | N/A | N/A |
| PF3D7_0412200 | N/A | conserved Plasmodium protein, unknown function | 39.10 | no | 0 | 6.66 | 3.02 | 1.97 | 0.47 | 4.73 | 32.12 | 0.00 | GV | 6.17 | 5.49 | 0 | N/A | N/A | osmiophilic body |
| PF3D7_1133400 | AMA1 | apical membrane antigen 1 | 72.04 | yes | 1 | 8.85 | 1.36 | 55.84 | 33.44 | 5.04 | 6.96 | 110.80 | O | 1.98 | 3.26 | 0 | host cell surface binding;protein binding | entry into host | apical complex;microneme;plasma membrane |
| PF3D7_1126100 | ATG7 | autophagy-related protein 7, putative | 156.63 | no | 0 | 5.19 | 3.44 | 6.87 | 0.88 | 1.47 | 7.71 | 105.32 | O | 0.75 | 6.77 | F | Atg12 activating enzyme activity;Atg8 activating enzyme activity | C-terminal protein lipidation;autophagosome assembly;autophagy of mitochondrion;cellular response to nitrogen starvation;late nucleophagy;piecemeal microautophagy of the nucleus;plastid organization;protein modification by small protein conjugation | cytoplasm;phagophore assembly site |
| PF3D7_1347500 | ALBA4 | DNA/RNA-binding protein Alba 4 | 42.16 | no | 0 | 628.83 | 120.68 | 135.98 | 12.40 | 18.97 | 63.69 | 221.29 | R | 94.80 | 65.11 | M | DNA binding;RNA binding;mRNA binding;protein-containing complex binding | regulation of translation | cytoplasm;cytosol;food vacuole;nucleus |
| PF3D7_1443000 | SRPK2 | serine/threonine protein kinase | 150.09 | no | 0 | 3.42 | 0.61 | 4.03 | 1.69 | 1.19 | 9.32 | 7.90 | GV | 6.59 | 15.58 | F | protein kinase activity;protein serine/threonine kinase activity;protein tyrosine kinase activity | peptidyl-serine phosphorylation;peptidyl-threonine phosphorylation;protein phosphorylation | Maurer's cleft;cytoplasm;nucleus |
| PF3D7_0103300 | N/A | conserved protein, unknown function | 108.38 | no | 0 | 53.35 | 19.25 | 13.64 | 2.91 | 2.33 | 14.32 | 40.92 | R | 56.65 | 32.14 | M | N/A | N/A | N/A |

| | | | | | | | | | | | | | | | | | | |
|---|---|---|---|---|---|---|---|---|---|---|---|---|---|---|---|---|---|---|
| PF3D7_0827900 | PDI8 | protein disulfide-isomerase | 55.51 | yes | 1 | 574.16 | 553.96 | 866.53 | 290.00 | 227.10 | 140.70 | 904.20 | O | 132.18 | 225.60 | F | RNA binding;protein binding;protein disulfide isomerase activity | protein folding;response to endoplasmic reticulum stress | cell surface;endoplasmic reticulum;endoplasmic reticulum lumen;food vacuole;microneme;nucleus;symbiont-containing vacuole |
| PF3D7_1316600 | CCT | choline-phosphate cytidylyltransferase | 104.78 | no | 0 | 56.47 | 37.32 | 41.62 | 22.49 | 6.61 | 15.91 | 90.53 | O | 35.14 | 17.65 | M | choline-phosphate cytidylyltransferase activity;identical protein binding;phosphatidylcholine binding | phosphatidylcholine biosynthetic process | cytoplasm;nucleus |
| PF3D7_0110400 | RPB9 | DNA-directed RNA polymerase II subunit RPB9, putative | 29.69 | no | 0 | 45.15 | 31.60 | 30.74 | 4.60 | 13.05 | 119.22 | 167.78 | O | 121.76 | 135.17 | F | DNA-directed 5'-3' RNA polymerase activity | maintenance of transcriptional fidelity during DNA-templated transcription elongation from RNA polymerase II promoter;transcription by RNA polymerase II;transcription initiation from RNA polymerase II promoter;transcription-coupled nucleotide-excision repair | RNA polymerase II, core complex;nucleus |
| PF3D7_0714500 | N/A | transcription elongation factor s-II, putative | 47.35 | no | 0 | 32.83 | 12.19 | 11.63 | 1.69 | 2.43 | 11.63 | 92.88 | O | 112.43 | 12.57 | M | DNA binding | regulation of DNA-templated transcription, elongation;regulation of transcription by RNA polymerase II | nucleus;transcription elongation factor complex |
| PF3D7_1315800 | MYB1 | transcription factor MYB1 | 49.99 | no | 0 | 25.21 | 22.70 | 15.37 | 2.90 | 3.82 | 53.30 | 38.97 | GV | 40.54 | 16.77 | M | DNA-binding transcription factor activity | positive regulation of transcription, DNA-templated;regulation of mitotic cell cycle;regulation of transcription, DNA-templated | nucleus |
| PF3D7_1353300 | N/A | conserved Plasmodium protein, unknown function | 248.26 | no | 2 | 32.09 | 32.60 | 25.53 | 5.34 | 5.88 | 39.26 | 337.50 | O | 52.00 | 163.18 | F | N/A | N/A | apicoplast |
| PF3D7_1124600 | EK | ethanolamine kinase | 49.91 | no | 0 | 10.72 | 8.20 | 112.40 | 23.47 | 9.26 | 53.84 | 139.51 | O | 73.94 | 53.36 | M | ethanolamine kinase activity | lipid metabolic process;phosphatidylcholine biosynthetic process;phosphatidylethanolamine biosynthetic process | Maurer's cleft;cytoplasm |
| PF3D7_0831400 | N/A | Plasmodium exported protein, unknown function | 36.41 | yes | 1 | 32.59 | 264.78 | 29.70 | 0.43 | 1.08 | 2.32 | 21.38 | T | 2.81 | 0.59 | 0 | N/A | N/A | integral component of membrane |
| PF3D7_1129000 | SPDS | spermidine synthase | 36.60 | no | 0 | 70.49 | 91.94 | 72.07 | 39.97 | 18.47 | 36.92 | 79.27 | T | 15.03 | 57.74 | F | RNA binding;spermidine synthase activity | polyamine biosynthetic process;spermidine biosynthetic process | cytoplasm;cytosol;mitochondrion;nucleus |
| PF3D7_1316900 | N/A | conserved protein, unknown function | 114.27 | no | 0 | 59.60 | 24.81 | 16.82 | 3.21 | 5.55 | 28.94 | 19.40 | R | 63.12 | 9.14 | M | N/A | regulation of transcription by RNA polymerase II | Piccolo NuA4 histone acetyltransferase complex;nucleus |
| PF3D7_0813900 | RPS16 | 40S ribosomal protein S16, putative | 16.29 | no | 0 | 231.51 | 168.64 | 92.97 | 16.33 | 125.34 | 100.88 | 62.77 | R | 142.94 | 112.88 | M | RNA binding;structural constituent of ribosome | maturation of SSU-rRNA from tricistronic rRNA transcript (SSU-rRNA, 5.8S rRNA, LSU-rRNA);translation | cytosolic small ribosomal subunit;nucleus;small ribosomal subunit |
| PF3D7_0718500 | N/A | prefoldin subunit 3, putative | 22.64 | no | 0 | 23.17 | 24.48 | 33.37 | 7.36 | 114.48 | 94.99 | 145.46 | O | 116.37 | 70.29 | M | tubulin binding;unfolded protein binding | 'de novo' protein folding;microtubule-based process;tubulin complex assembly | cytoplasm;polysome;prefoldin complex |
| PF3D7_1426100 | BTF3 | transcription factor BTF3, putative | 19.39 | no | 0 | 87.26 | 83.37 | 30.28 | 1.86 | 25.60 | 91.62 | 65.00 | GV | 45.91 | 56.56 | F | DNA binding | regulation of transcription | cytosol;nascent polypeptide-associated complex;nucleus;polysomal ribosome;transcription regulator complex |
| PF3D7_0801700 | SENP2 | sentrin-specific protease 2, putative | 210.88 | no | 0 | 15.44 | 20.41 | 17.08 | 2.12 | 2.68 | 16.21 | 41.33 | O | 21.59 | 53.21 | F | cysteine-type peptidase activity | proteolysis | nucleus |
| PF3D7_1438900 | Trx-Px1 | thioredoxin peroxidase 1 | 21.81 | no | 0 | 528.37 | 147.40 | 188.59 | 52.36 | 319.18 | 169.42 | 66.49 | R | 56.09 | 139.88 | F | RNA binding;antioxidant activity;double-stranded DNA binding;double-stranded RNA binding;long noncoding RNA binding;protein binding;protein dimerization activity;thioredoxin peroxidase activity | DNA protection;cell redox homeostasis;protein folding;response to heat;response to oxidative stress | Maurer's cleft;cell surface;cytoplasm;cytosol;nucleolus;nucleus |
| PF3D7_1251200 | N/A | coronin | 69.01 | no | 0 | 19.93 | 1.22 | 63.45 | 66.96 | 1.06 | 6.29 | 256.48 | O | 103.61 | 10.37 | M | actin binding;actin filament binding | actin filament organization;cell motility;microtubule-based process;response to xenobiotic stimulus | nucleus;pellicle |
| PF3D7_1027300 | nPrx | peroxiredoxin | 43.93 | no | 0 | 236.88 | 45.44 | 438.32 | 815.09 | 92.97 | 54.92 | 774.05 | S | 111.48 | 1667.71 | F | thioredoxin peroxidase activity | cell redox homeostasis;cellular response to oxidative stress;entry into host | cytoplasm;nucleus |
| PF3D7_1346300 | ALBA2 | DNA/RNA-binding protein Alba 2 | 24.98 | no | 0 | 914.99 | 182.62 | 122.63 | 22.05 | 50.48 | 77.26 | 56.49 | R | 50.55 | 87.92 | F | DNA binding;RNA binding;mRNA binding | regulation of translation | cytoplasm;nucleus |
| PF3D7_1229400 | MIF | macrophage migration inhibitory factor | 12.84 | no | 0 | 178.74 | 77.56 | 63.25 | 13.16 | 35.78 | 151.71 | 330.04 | O | 12.71 | 19.65 | F | cytokine activity;deoxyribonuclease activity;host cell surface receptor binding;oxidoreductase activity | evasion of host immune response;suppression by symbiont of host innate immune response | Maurer's cleft;cytoplasm;host cell cytoplasm |
| PF3D7_1420400 | GlyRS | glycine--tRNA ligase | 104.30 | yes | 0 | 140.93 | 52.74 | 23.07 | 2.03 | 2.66 | 13.90 | 61.80 | R | 13.86 | 64.12 | F | glycine-tRNA ligase activity | glycyl-tRNA aminoacylation;mitochondrial glycyl-tRNA aminoacylation | apicoplast;cytoplasm;mitochondrion |
| PF3D7_1228600 | MSP9 | Merozoite surface protein 9 | 86.62 | yes | 0 | 50.84 | 24990.00 | 1245.35 | 1997.72 | 44.24 | 23.39 | 993.68 | T | 149.07 | 187.25 | F | protein binding | cell adhesion | extracellular vesicle;host cell cytoplasm;membrane;symbiont-containing vacuole |
| PF3D7_1115400 | FP3 | cysteine proteinase falcipain 3 | 56.67 | no | 1 | 130.27 | 51.08 | 143.54 | 27.71 | 18.03 | 45.64 | 92.28 | T | 97.15 | 52.24 | M | cysteine-type endopeptidase activity | hemoglobin catabolic process;proteolysis;proteolysis involved in cellular protein catabolic process | extracellular space;food vacuole;lysosome |
| PF3D7_0517300 | SR1 | serine/arginine-rich splicing factor 1 | 34.56 | no | 0 | 404.84 | 179.12 | 92.82 | 18.50 | 15.00 | 43.87 | 133.57 | R | 93.83 | 76.82 | M | RNA binding;phosphorylated histone binding | RNA splicing;cellular response to DNA damage stimulus;mRNA cis splicing, via spliceosome;mRNA processing;regulation of alternative mRNA splicing, via spliceosome;regulation of mRNA splicing, via spliceosome | cytoplasm;nuclear speck;nucleus;spliceosomal complex |
| PF3D7_1437400 | PANK2 | pantothenate kinase 2 | 91.04 | no | 0 | 10.06 | 9.72 | 15.29 | 3.89 | 1.12 | 6.15 | 22.92 | O | 10.76 | 3.20 | M | 14-3-3 protein binding;pantothenate kinase activity;protein binding | coenzyme A biosynthetic process | cytosol;nucleus |
| PF3D7_1110900 | N/A | ES2 protein, putative | 69.02 | no | 0 | 35.87 | 14.38 | 15.75 | 2.01 | 2.15 | 17.12 | 47.50 | O | 59.20 | 29.81 | M | protein binding;RNA binding | RNA splicing | catalytic step 2 spliceosome |
| PF3D7_0311300 | N/A | phosphatidylinositol 3- and 4-kinase, putative | 110.56 | no | 0 | 6.17 | 1.72 | 45.19 | 20.67 | 11.52 | 35.88 | 147.36 | O | 159.71 | 49.59 | M | enzymatic activity | signal transduction | nucleus;cytoplasm |
| PF3D7_0719900 | N/A | conserved Plasmodium membrane protein, unknown function | 402.70 | no | 4 | 53.44 | 26.28 | 34.75 | 23.27 | 5.67 | 26.90 | 41.61 | R | 37.14 | 46.57 | F | N/A | N/A | trans-Golgi network |
| PF3D7_1340600 | DBR1 | RNA lariat debranching enzyme, putative | 67.55 | no | 0 | 44.62 | 41.84 | 34.26 | 2.84 | 2.13 | 8.42 | 109.60 | O | 49.08 | 21.59 | M | hydrolase activity, acting on ester bonds | mRNA processing | nucleus |
| PF3D7_0621800 | N/A | nascent polypeptide-associated complex subunit alpha | 20.62 | no | 0 | 727.07 | 431.74 | 116.63 | 8.14 | 243.72 | 158.76 | 103.51 | R | 52.90 | 52.33 | 0 | unfolded protein binding | 'de novo' cotranslational protein folding;protein targeting to membrane | cytoplasm;nascent polypeptide-associated complex;nucleus |
| PF3D7_0923000 | RPB3 | DNA-directed RNA polymerase II subunit RPB3, putative | 38.35 | no | 0 | 75.74 | 13.31 | 9.78 | 0.61 | 4.20 | 36.68 | 11.96 | R | 25.76 | 12.27 | M | DNA binding | regulation of transcription | RNA polymerase II, core complex;nucleus |
| PF3D7_0822800 | SNRNP40 | U5 small nuclear ribonucleoprotein 40 kDa protein, putative | 37.09 | no | 0 | 24.10 | 6.32 | 8.54 | 0.55 | 1.71 | 11.55 | 25.15 | O | 19.01 | 20.98 | 0 | RNA binding | RNA splicing, via transesterification reactions | catalytic step 2 spliceosome;sno(s)RNA-containing ribonucleoprotein complex |
| PF3D7_0707700 | N/A | E3 ubiquitin-protein ligase, putative | 110.78 | no | 0 | 136.99 | 65.52 | 57.95 | 34.11 | 37.56 | 96.70 | 133.19 | R | 167.10 | 67.43 | M | chaperone binding;ubiquitin protein ligase activity | cellular response to misfolded protein;positive regulation of proteolysis;proteasome-mediated ubiquitin-dependent protein catabolic process;protein polyubiquitination;protein quality control for misfolded or incompletely synthesized proteins | cytoplasm;nucleus |
| PF3D7_1107300 | PAIP1 | polyadenylate-binding protein-interacting protein 1, putative | 381.87 | no | 0 | 265.97 | 114.16 | 49.70 | 12.61 | 11.21 | 67.65 | 173.07 | R | 608.32 | 166.19 | M | mRNA binding;mRNA cap binding;translation initiation factor activity | | eukaryotic translation initiation factor 4F complex;nucleus |
| PF3D7_0936800 | N/A | Plasmodium exported protein (PHISTc), unknown function | 45.50 | no | 1 | 880.47 | 81.09 | 22.22 | 0.19 | 4.25 | 2.46 | 27.89 | R | 3.80 | 1.25 | 0 | protein binding | N/A | Maurer's cleft;host cell cytosol;host cell membrane;host cell plasma membrane |
| PF3D7_1417500 | CBF5 | H/ACA ribonucleoprotein complex subunit 4, putative | 52.94 | no | 0 | 111.44 | 39.78 | 22.12 | 2.50 | 4.97 | 10.72 | 21.22 | R | 182.67 | 23.21 | M | pseudouridine synthase activity | box H/ACA RNA 3'-end processing;mRNA pseudouridine synthesis;rRNA pseudouridine synthesis;snRNA pseudouridine synth | box H/ACA snoRNP complex;nucleus |
| PF3D7_1105700 | RTCB | RNA-splicing ligase RtcB homolog | 56.32 | no | 0 | 100.05 | 36.31 | 45.84 | 28307.00 | 14.66 | 46722.00 | 34.82 | GC V | 36.12 | 37.41 | 0 | RNA ligase (ATP) activity | tRNA exon ligation utilizing 2',3' cyclic phosphate of 5'-exon as source of linkage phosphate;tRNA splicing, via endonucleolytic cleavage and ligation | nucleus;tRNA-splicing ligase complex |
| PF3D7_1126900 | SNRPF | small nuclear ribonucleoprotein F, putative | 9.89 | no | 0 | 55.70 | 29.11 | 30.48 | 5.18 | 28.30 | 39.12 | 46.85 | R | 61.42 | 126.09 | F | RNA binding | mRNA splicing, via spliceosome | U1 snRNP;U4/U6 x U5 tri-snRNP complex;U5 snRNP;catalytic step 2 spliceosome;nucleus |
| PF3D7_0309500 | AS | asparagine synthetase [glutamine-hydrolyzing], putative | 69.86 | no | 0 | 311.94 | 131.53 | 149.76 | 97.72 | 9.10 | 8.84 | 72.42 | R | 69.37 | 39.67 | M | asparagine synthase (glutamine-hydrolyzing) activity | asparagine biosynthetic process | cytosol;nucleus |
| PF3D7_0802200 | 1-CysPxn | 1-cys peroxiredoxin | 25.16 | no | 0 | 8.86 | 33.62 | 135.46 | 10.47 | 19.91 | 74.28 | 440.74 | O | 17.39 | 13.63 | 0 | antioxidant activity;peroxidase activity | cell redox homeostasis;response to oxidative stress | cytosol;mitochondrion |
| PF3D7_1315900 | N/A | exportin-T, putative | 143.00 | no | 0 | 80.07 | 40.03 | 36.51 | 5.38 | 5.98 | 27.25 | 44.65 | R | 26.75 | 9.39 | M | tRNA binding | tRNA export from nucleus | nucleus;cytoplasm |
| PF3D7_0705300 | ORC2 | origin recognition complex subunit 2 | 98.01 | no | 0 | 6.27 | 3.00 | 19.70 | 7.46 | 2.50 | 28.83 | 81.22 | O | 575.95 | 55.65 | M | DNA replication origin binding | DNA replication initiation | endoplasmic reticulum;mitochondrion;nuclear origin of replication recognition complex;nuclear periphery;nucleus |
| PF3D7_0308200 | CCT7 | T-complex protein 1 subunit eta | 59.58 | no | 0 | 177.06 | 106.03 | 66.15 | 11.71 | 11.42 | 13.35 | 147.58 | R | 76.58 | 54.96 | M | unfolded protein binding | protein folding | chaperonin-containing T-complex;cytosol;nucleus |
| PF3D7_1338200 | RPL6 | 60S ribosomal protein L6, putative | 25.53 | no | 0 | 178.02 | 111.44 | 44.82 | 3.22 | 28.96 | 59.05 | 72.65 | R | 74.97 | 17.81 | M | RNA binding;structural constituent of ribosome | cytoplasmic translation | cytosolic large ribosomal subunit;extracellular vesicle;nucleus |
| PF3D7_0610200 | RBM25 | RNA-binding protein 25, putative | 115.74 | no | 0 | 50.95 | 13.91 | 7.35 | 1.10 | 1.69 | 14.32 | 54.20 | O | 385.44 | 127.39 | M | RNA binding | N/A | nucleus |
| PF3D7_0318100 | STOML | stomatin-like protein, putative | 43.08 | no | 0 | 12.48 | 21.15 | 15.44 | 1.41 | 1.22 | 4.71 | 16.17 | T | 8.54 | 7.14 | 0 | N/A | N/A | mitochondrion;nucleus |
| PF3D7_0929000 | TAF7 | transcription initiation factor TFIID subunit 7, putative | 46.80 | no | 0 | 54.96 | 12.90 | 12.12 | 1.98 | 14.15 | 48.04 | 34.10 | R | 72.91 | 36.34 | M | RNA binding | RNA polymerase II preinitiation complex assembly;transcription by RNA polymerase II | transcription factor TFIID complex |
| PF3D7_0518600 | WDR26 | WD repeat-containing protein 26, putative | 150.70 | no | 0 | 7.89 | 0.92 | 20.13 | 4.01 | 12.71 | 54.98 | 113.80 | O | 375.61 | 116.80 | M | protein binding | N/A | N/A |
| PF3D7_1419200 | ATrx1 | thioredoxin-like protein, putative | 80.24 | yes | 0 | 6.98 | 12.30 | 13.96 | 1.82 | 2.13 | 11.20 | 1.19 | T | 13.61 | 6.85 | M | N/A | N/A | apicoplast;nucleus |
| PF3D7_0904900 | CuTP | copper-transporting ATPase | 299.39 | no | 7 | 7.90 | 4.09 | 46.30 | 7.87 | 3.41 | 7.99 | 46.51 | O | 74.22 | 43.35 | M | ATP hydrolysis activity;P-type calcium transporter activity;P-type proton-exporting transporter activity;copper ion binding | calcium ion transmembrane transport;cellular calcium ion homeostasis;copper ion transport | cytoplasmic vesicle;host cell plasma membrane;plasma membrane |
| PF3D7_1015400 | N/A | RMI1 domain-containing protein, putative | 92.88 | no | 0 | 8.03 | 1.80 | 14.13 | 4.94 | 1.50 | 10.98 | 10.11 | T | 4.06 | 10.95 | F | N/A | N/A | nucleus |
| PF3D7_0907700 | PA28 | proteasome activator 28 | 33.13 | no | 0 | 154.12 | 31.80 | 37.26 | 4.88 | 2.65 | 35.03 | 108.52 | R | 39.48 | 17.56 | M | endopeptidase activator activity | positive regulation of endopeptidase activity;regulation of G1/S transition of mitotic cell cycle;regulation of proteasomal protein catabolic process;response to xenobiotic stimulus | cytoplasm;nucleoplasm;nucleus;proteasome activator complex |
| PF3D7_0306900 | N/A | 40S ribosomal protein S23, putative | 16.13 | no | 0 | 591.61 | 623.27 | 260.25 | 109.84 | 284.46 | 149.34 | 165.61 | T | 156.23 | 128.94 | M | structural constituent of ribosome | translation | cytosolic small ribosomal subunit;nucleus;ribosome |
| PF3D7_0614400 | CWF7 | pre-mRNA-splicing factor CWF7, putative | 30.85 | no | 0 | 26.99 | 15.23 | 5.44 | 0.65 | 2.77 | 24.46 | 432.49 | O | 12.48 | 51.49 | F | RNA binding | mRNA processing | Prp19 complex;catalytic step 2 spliceosome |
| PF3D7_1412600 | DHS | deoxyhypusine synthase | 57.24 | no | 1 | 60.03 | 32.74 | 9.75 | 0.55 | 15.84 | 9.77 | 9.00 | R | 41.11 | 3.72 | M | deoxyhypusine synthase activity | peptidyl-lysine modification to peptidyl-hypusine | cytoplasm |
| PF3D7_1116100 | N/A | serine esterase, putative | 217.07 | no | 0 | 6.16 | 1.76 | 34.14 | 13.55 | 2.86 | 16.13 | 102.14 | O | 82.97 | 26.02 | M | enzymatic activity | cellular lipid metabolic process | food vacuole |
| PF3D7_1320800 | N/A | dihydrolipoyllysine-residue succinyltransferase component of 2-oxoglutarate dehydrogenase | 47.63 | no | 0 | 8.94 | 2.69 | 21.55 | 1.78 | 3.67 | 16.61 | 86.97 | O | 3.37 | 50.02 | F | RNA binding;dihydrolipoyllysine-residue succinyltransferase activity | 2-oxoglutarate metabolic process | mitochondrion |
| PF3D7_1407300 | PRP38B | pre-mRNA-splicing factor 38B, putative | 81.76 | no | 0 | 108.25 | 64.89 | 55.56 | 5.46 | 7.76 | 26.88 | 223.88 | O | 197.66 | 55.67 | M | RNA binding | mRNA processing | precatalytic spliceosome;nucleus |
| PF3D7_1353900 | N/A | proteasome subunit alpha type-7, putative | 27.23 | no | 0 | 65.78 | 46.70 | 104.96 | 25.63 | 49.91 | 83.15 | 33.60 | T | 20.10 | 39.53 | F | endopeptidase activity | proteasomal protein catabolic process;proteasomal ubiquitin-independent protein catabolic process;proteasome-mediated ubiquitin-dependent protein catabolic process;ubiquitin-dependent protein catabolic process | cytoplasm;nucleus;proteasome core complex;proteasome core complex, alpha-subunit complex |
| PF3D7_0606800 | VFT1 | VFT protein | 34.50 | yes | 0 | 12.40 | 1.86 | 13.75 | 2.49 | 7.43 | 34.29 | 49.17 | O | 11.77 | 11.82 | 0 | N/A | N/A | cell surface |
| PF3D7_1410600 | eIF2gamma | eukaryotic translation initiation factor 2 subunit gamma, putative | 51.24 | no | 0 | 937.06 | 198.24 | 211.93 | 42.44 | 34.53 | 34.65 | 105.36 | R | 20.39 | 51.97 | F | GTP binding;protein binding;translation initiation factor activity | formation of translation preinitiation complex;positive regulation of translational fidelity;translational initiation | cytoplasm;cytosol;eukaryotic translation initiation factor 2 complex;nucleus |
| PF3D7_0310500 | DHX57 | ATP-dependent RNA helicase DHX57, putative | 267.23 | no | 0 | 22.97 | 20.27 | 15.48 | 1.98 | 2.34 | 18.60 | 38.18 | O | 26.64 | 15.82 | M | RNA binding;helicase activity | maturation of SSU-rRNA from tricistronic rRNA transcript (SSU-rRNA, 5.8S rRNA, LSU-rRNA) | nucleolus;nucleus |
| PF3D7_1368200 | ABCE1 | ABC transporter E family member 1, putative | 70.27 | no | 0 | 283.55 | 70.15 | 83.17 | 21.73 | 10.31 | 16.54 | 31.28 | R | 18.17 | 15.02 | 0 | ATP binding;RNA binding;iron ion binding;ribosomal small subunit binding | ribosomal subunit export from nucleus;translational initiation;translational termination | nucleus |
| PF3D7_0205700 | N/A | RNA-binding protein, putative | 24.33 | no | 0 | 47.93 | 14.13 | 43344.00 | 34700.00 | 21.14 | 266.53 | 61.8 | T | 49.23 | 66.93 | F | RNA binding | N/A | nucleus;cytoplasm |
| PF3D7_1465200 | MED4 | mediator of RNA polymerase II transcription subunit 4, putative | 44.68 | no | 0 | 52.44 | 11.71 | 9.66 | 0.75 | 3.05 | 12.84 | 0.00 | R | 11.70 | 6.52 | M | protein binding | regulation of transcription | mediator complex;nucleus |
| PF3D7_1248200 | RBM22 | Pre-mRNA-splicing factor RBM22, putative | 41.97 | no | 0 | 38.85 | 24.29 | 13.53 | 2.71 | 17.94 | 110.22 | 121.74 | O | 65.51 | 145.18 | R | U6 snRNA binding;pre-mRNA binding | mRNA cis splicing, via spliceosome | Prp19 complex;U2-type catalytic step 1 spliceosome;U2-type catalytic step 2 spliceosome |
| PF3D7_1239000 | N/A | HD superfamily phosphohydrolase protein, putative | 96.00 | no | 0 | 21.90 | 10.32 | 20.64 | 7.42 | 9.07 | 87.35 | 85.49 | GV | 2.92 | 57.12 | F | dGTPase activity | dGTP catabolic process | nucleus |
| PF3D7_0109200 | CFIM25 | cleavage and polyadenylation specificity factor subunit 5, putative | 27.32 | no | 0 | 21.09 | 11.97 | 19.67 | 3.38 | 46.97 | 124.90 | 70.98 | GV | 22.24 | 16.42 | M | RNA binding;mRNA binding | mRNA processing | mRNA cleavage factor complex |
| PF3D7_0216200 | N/A | Alba domain-containing protein, putative | 27.21 | no | 0 | 28.93 | 13.50 | 7.69 | 0.54 | 3.68 | 23.34 | 37.48 | O | 14.96 | 29.60 | F | protein binding | N/A | nucleus;cytoplasm |
| PF3D7_1447700 | N/A | conserved protein, unknown function | 40.70 | no | 3 | 9.49 | 3.33 | 13.54 | 5.70 | 8.20 | 82.38 | 8.57 | GV | 13.10 | 12.32 | 0 | RNA binding | N/A | nucleus;cytoplasm |
| PF3D7_1203700 | NAPL | nucleosome assembly protein | 40.49 | no | 0 | 456.51 | 180.45 | 265.44 | 31.80 | 44.21 | 75.80 | 516.99 | O | 363.93 | 98.04 | M | chromatin binding;histone binding;protein binding | nucleosome assembly | cytoplasm;nucleus |
| PF3D7_0506500 | N/A | conserved Plasmodium protein, unknown function | 469.87 | no | 0 | 6.40 | 10.50 | 6.77 | 1.71 | 2.13 | 22.20 | 212.22 | O | 706.74 | 125.88 | M | N/A | N/A | N/A |
| PF3D7_1010100 | N/A | PI31 domain-containing protein, putative | 35.60 | no | 0 | 77.53 | 8.35 | 20.93 | 3.94 | 41.32 | 263.44 | 374.13 | O | 230.09 | 134.73 | M | N/A | ubiquitin-dependent protein catabolic process | N/A |
| PF3D7_1237700 | N/A | conserved protein, unknown function | 23.66 | no | 5 | 77.47 | 51.48 | 134.58 | 57.98 | 103.33 | 527.14 | 129.80 | GV | 13.39 | 117.33 | F | N/A | N/A | cell surface;food vacuole;mitochondrion;nucleus |
| PF3D7_1309100 | N/A | 60S ribosomal protein L24, putative | 19.25 | no | 0 | 302.41 | 241.43 | 125.89 | 19.63 | 149.63 | 251.51 | 648.75 | O | 712.32 | 554.36 | M | RNA binding;structural constituent of ribosome | assembly of large subunit precursor of preribosome;ribosomal large subunit assembly;translation | cytosolic large ribosomal subunit;nucleus |
| PF3D7_1142100 | N/A | DnaJ domain-containing protein, putative | 324.82 | no | 0 | 92.19 | 10.52 | 24.05 | 5.73 | 14.57 | 42.45 | 45.85 | R | 229.05 | 48.90 | M | protein binding | chaperone complex | nucleus |
| PF3D7_1016400 | FIKK10.1 | Serine/threonine protein kinase, FIKK family | 72.42 | no | 1 | 38.85 | 24.29 | 13.53 | 25965.00 | 17.94 | 110.22 | 121.74 | S | 65.51 | 145.18 | F | protein serine/threonine kinase activity | intracellular signal transduction;modulation by symbiont of host process;protein phosphorylation | Maurer's cleft;cytoplasm;nucleus |
| PF3D7_0501200 | PIESP2 | parasite-infected erythrocyte surface protein | 48.72 | no | 3 | 57.67 | 365.94 | 123.20 | 5.85 | 1.63 | 1.71 | 11.18 | T | 6.79 | 2.29 | 0 | host cell surface binding;protein binding | cell adhesion | Maurer's cleft;cell surface;host cell cytoplasm;host cell surface;nucleus |
| PF3D7_1104400 | Trx-mero | thioredoxin-like mero protein | 49.27 | yes | 1 | 120.76 | 75.63 | 173.84 | 52.72 | 17.36 | 47.08 | 79.91 | T | 43.99 | 51.29 | F | host cell surface binding | entry into host | cell surface;nucleus;symbiont-containing vacuole |
| PF3D7_0604500 | N/A | GYF domain-containing protein, putative | 334.41 | no | 0 | 230.63 | 68.12 | 42.18 | 8.77 | 6.13 | 53.06 | 139.04 | R | 358.20 | 182.15 | M | mRNA binding | N/A | nucleus |
| PF3D7_1005500 | UPF1 | regulator of nonsense transcripts 1, putative | 183.96 | no | 0 | 31.20 | 22.88 | 46.85 | 7.51 | 10.15 | 113.61 | 139.54 | O | 165.50 | 175.06 | F | RNA binding;RNA helicase activity | nuclear-transcribed mRNA catabolic process, nonsense-mediated decay | cytoplasm;nucleus |

| | | | | | | | | | | | | | | | | | | |
|---|---|---|---|---|---|---|---|---|---|---|---|---|---|---|---|---|---|---|
| PF3D7_1364500 | RRP45 | exosome complex component RRP45, putative | 55.03 | no | 0 | 22.56 | 6.17 | 6.95 | 0.98 | 1.29 | 10.20 | 6.90 | R | 31.77 | 10.00 | M | 3'-5'-exoribonuclease activity | RNA processing;U1 snRNA 3'-end processing;U4 snRNA 3'-end processing;U5 snRNA 3'-end processing;exonucleolytic catabolism of deadenylated mRNA;exonucleolytic trimming to generate mature 3'-end of 5.8S rRNA from tricistronic rRNA transcript (SSU-rRNA, 5.8S rRNA, LSU-rRNA);nuclear mRNA surveillance;nu | cytoplasmic exosome (RNase complex);exosome (RNase complex);nuclear exosome (RNase complex) |
| PF3D7_1119900 | SEC16 | protein transport protein SEC16, putative | 208.59 | no | 0 | 66.61 | 10.63 | 14.08 | 4.23 | 0.85 | 10.73 | 14.38 | R | 20.21 | 19.88 | 0 | N/A | vesicle-mediated transport | endoplasmic reticulum |
| PF3D7_0705500 | N/A | inositol-phosphate phosphatase, putative | 330.70 | no | 0 | 148.21 | 24.50 | 39.78 | 16.13 | 4.99 | 29.47 | 174.48 | O | 80.04 | 32.22 | M | phosphatidylinositol-4,5-bisphosphate 5-phosphatase activity | inositol phosphate dephosphorylation;phosphatidylinositol dephosphorylation | N/A |
| PF3D7_0803700 | g-tub | tubulin gamma chain | 51.64 | no | 0 | 13.30 | 7.24 | 26.45 | 4.84 | 16.41 | 42.20 | 93.24 | O | 246.66 | 83.70 | M | GTP binding;RNA binding;structural constituent of cytoskeleton | meiotic spindle organization;microtubule cytoskeleton organization;microtubule nucleation;microtubule-based process;mitotic cell cycle;mitotic sister chromatid segregation;mitotic spindle organization | centrosome;cytoplasm;gamma-tubulin complex;microtubule;microtubule organizing center;nucleus;spindle |
| PF3D7_0420300 | ApiAP2 | AP2 domain transcription factor, putative | 400.16 | no | 0 | 115.68 | 92.16 | 57.10 | 28.79 | 4.65 | 20.37 | 743.23 | O | 51.70 | 52.09 | 0 | protein binding;sequence-specific DNA binding | regulation of transcription | nucleus |
| PF3D7_0532400 | LyMP | lysine-rich membrane-associated PHISTb protein | 61.09 | no | 1 | 123.16 | 207.91 | 37.88 | 3.00 | 0.72 | 1.58 | 27.27 | T | 20.91 | 1.18 | M | protein binding | cytoadherence to microvasculature, mediated by symbiont protein | Maurer's cleft;host cell cytoplasm;host cell cytosol;host cell membrane;host cell periphery;host cell surface;infected host cell surface knob;nucleus |
| PF3D7_1130100 | RPL38 | 60S ribosomal protein L38 | 10.31 | no | 0 | 57.29 | 57.51 | 28.38 | 7.12 | 63.28 | 368.45 | 76.23 | GV | 96.72 | 195.43 | F | structural constituent of ribosome | ribonucleoprotein complex assembly;translation | cytosolic large ribosomal subunit;nucleus |
| PF3D7_0925700 | HDAC1 | histone deacetylase 1 | 51.38 | no | 0 | 156.78 | 62.14 | 85.17 | 62.49 | 20.60 | 21.37 | 76.19 | R | 89.71 | 34.53 | M | histone deacetylase activity;protein binding;sphingolipid binding | cellular response to drug;histone deacetylation;response to xenobiotic stimulus | nucleus |
| PF3D7_0822100 | MED7 | mediator of RNA polymerase II transcription subunit 4, putative | 28.20 | no | 0 | 20.92 | 7.64 | 5.30 | 0.86 | 59.08 | 126.74 | 31.95 | GV | 54.32 | 34.45 | M | protein binding | regulation of transcription | mediator complex;nucleus |
| PF3D7_1128100 | N/A | prefoldin subunit 5, putative | 29.14 | no | 0 | 42.67 | 38.52 | 48.66 | 6.62 | 14.10 | 65.25 | 182.62 | O | 91.39 | 64.33 | M | unfolded protein binding | protein folding | cytoplasm;prefoldin complex;nucleus |
| PF3D7_0617200 | N/A | BFR1 domain-containing protein, putative | 52.98 | no | 0 | 214.03 | 87.02 | 54.24 | 13.86 | 11.02 | 41.13 | 102.72 | R | 306.61 | 196.38 | M | protein binding | intracellular mRNA localization | endoplasmic reticulum;nuclear outer membrane-endoplasmic reticulum membrane network;nucleus;ribonucleoprotein complex |
| PF3D7_1464500 | N/A | conserved membrane protein, unknown function | 388.81 | no | 4 | 10.28 | 15.01 | 13.78 | 3.90 | 1.29 | 9.70 | 34.92 | O | 15.91 | 13.69 | 0 | N/A | N/A | integral component of membrane |
| PF3D7_1361900 | PCNA1 | Proliferating cell nuclear antigen | 30.59 | no | 0 | 18810.00 | 12267.00 | 97.41 | 17.16 | 16.89 | 46.54 | 75.08 | R | 421.82 | 13.97 | M | DNA polymerase processivity factor activity | leading strand elongation;mismatch repair;translesion synthesis | PCNA complex;cytosol;nucleus |
| PF3D7_1014100 | MSA180 | merozoite surface protein MSA180 | 173.27 | yes | 0 | 3.97 | 0.23 | 3.84 | 4.28 | 0.28 | 3.58 | 23.18 | O | 4.06 | 2.88 | 0 | protein binding | entry into host;exit from host cell | cell surface;cytoplasm;extracellular vesicle;symbiont-containing vacuole |
| PF3D7_1338700 | N/A | conserved protein, unknown function | 73.21 | no | 0 | 4.79 | 3.13 | 32.35 | 5.96 | 1.08 | 6.91 | 422.25 | O | 10.81 | 56.42 | F | N/A | N/A | nucleus |
| PF3D7_1303400 | LisH | LisH domain-containing protein, putative | 118.62 | no | 0 | 78.90 | 48.90 | 23.20 | 6.94 | 3.36 | 9.71 | 67.47 | R | 115.69 | 39.57 | M | protein binding | N/A | nucleus |
| PF3D7_1353400 | N/A | Ran-binding protein, putative | 140.42 | no | 0 | 41.34 | 16.96 | 8.04 | 2.05 | 2.75 | 8.49 | 14.89 | R | 59.80 | 15.90 | M | RNA-binding;metal ion binding | N/A | N/A |
| PF3D7_0926400 | N/A | monocarboxylate transporter, putative | 60.71 | no | 11 | 16.71 | 1.57 | 19.01 | 3.43 | 13.42 | 543.35 | 457.27 | GV | 19.77 | 594.79 | F | monocarboxylic acid transmembrane transporter activity | monocarboxylic acid transport | apicoplast;integral component of membrane;membrane |
| PF3D7_0904400 | SPC3 | signal peptidase complex subunit 3, putative | 22.21 | no | 1 | 5.51 | 2.71 | 18.50 | 5.19 | 22.25 | 109.04 | 26.19 | GV | 16.08 | 31.12 | F | peptidase activity | protein targeting to ER;signal peptide processing | osmiophilic body;signal peptidase complex |
| PF3D7_1111100 | RFC5 | replication factor C subunit 5, putative | 40.35 | no | 0 | 9.89 | 5.14 | 37.32 | 5.46 | 10.62 | 61.42 | 119.20 | O | 155.28 | 61.78 | M | RNA binding | DNA repair;DNA replication;DNA strand elongation involved in DNA replication;DNA-dependent DNA replication | DNA replication factor C complex;nucleus |
| PF3D7_0803000 | CYP81 | peptidyl-prolyl cis-trans isomerase | 80.92 | no | 0 | 59.73 | 16.02 | 22.71 | 2.80 | 4.92 | 15.30 | 126.97 | O | 96.24 | 75.26 | M | peptidyl-prolyl cis-trans isomerase activity;protein dimerization activity | chaperone-mediated protein folding;negative regulation of phosphoprotein phosphatase activity;protein folding;protein peptidyl-prolyl isomerization;response to xenobiotic stimulus | cytoplasm;nucleus |
| PF3D7_0722500 | CWC15 | pre-mRNA-splicing factor CWC15, putative | 34.15 | no | 0 | 378.24 | 104.64 | 57.66 | 114.25 | 22.65 | 56.05 | 157.58 | R | 177.63 | 250.92 | F | RNA binding | RNA splicing;mRNA cis splicing, via spliceosome;mRNA splicing, via spliceosome;regulation of cell cycle | U2-type spliceosomal complex;catalytic step 2 spliceosome;spliceosomal complex;nucleus |
| PF3D7_1409300 | DDI1 | DNA damage-inducible protein 1 | 43.84 | no | 0 | 4.92 | 1.51 | 22.34 | 11.29 | 8.18 | 50.85 | 84.60 | O | 75.60 | 45.29 | 0 | aspartic-type endopeptidase activity | proteolysis;response to xenobiotic stimulus | cytoplasm;nucleus |
| PF3D7_1146600 | ORP1 | oocyst rupture protein 1, putative | 150.98 | no | 0 | 4.11 | 0.06 | 12.50 | 0.11 | 3.69 | 16.93 | 56.42 | O | 18.46 | 34.39 | F | DNA binding | regulation of transcription, DNA-templated | cell periphery;nucleus |
| PF3D7_0514900 | N/A | conserved Plasmodium protein, unknown function | 51.90 | no | 0 | 64.91 | 10.96 | 15.01 | 2.40 | 12.96 | 81.06 | 42.68 | GV | 20.60 | 22.05 | 0 | N/A | N/A | cytoplasm;nucleus;transcription regulator complex |
| PF3D7_1229100 | MRP2 | multidrug resistance-associated protein 2 | 248.30 | no | 11 | 494.10 | 52.95 | 24.07 | 5.81 | 3.89 | 10.57 | 104.92 | R | 21.34 | 6.28 | M | ATPase-coupled transmembrane transporter activity | response to drug;response to xenobiotic stimulus;schizogony;transmembrane transport | integral component of plasma membrane;membrane;plasma membrane |
| PF3D7_0112200 | MRP1 | multidrug resistance-associated protein 1 | 214.44 | no | 11 | 5.61 | 22.88 | 16.47 | 2.58 | 3.78 | 4.53 | 2.63 | T | 0.25 | 0.14 | 0 | ATPase-coupled transmembrane transporter activity;glutathione binding | response to drug;response to xenobiotic stimulus;transmembrane transport | Maurer's cleft;integral component of plasma membrane;membrane;plasma membrane |
| PF3D7_0628600 | N/A | DNA methyltransferase 1-associated protein 1, putative | 46.56 | no | 0 | 32.66 | 16.54 | 11.66 | 1.03 | 7.50 | 38.86 | 101.09 | O | 29.91 | 20.19 | M | transcription corepressor activity | histone H2A acetylation;histone H4 acetylation;histone exchange;negative regulation of transcription by RNA polymerase II | NuA4 histone acetyltransferase complex;Swr1 complex;nucleus |
| PF3D7_0518300 | N/A | proteasome subunit beta type-1, putative | 27.27 | no | 0 | 59.43 | 33.09 | 89.90 | 21.12 | 71.34 | 91.04 | 40.74 | GV | 41.43 | 13.20 | M | endopeptidase activity | proteasomal protein catabolic process;proteasomal ubiquitin-independent protein catabolic process;proteasome-mediated ubiquitin-dependent protein catabolic process;ubiquitin-dependent protein catabolic process | cytoplasm;extracellular vesicle;nucleus;proteasome core complex |
| PF3D7_1331800 | N/A | 60S ribosomal protein L23, putative | 14.99 | no | 0 | 268.05 | 367.47 | 121.43 | 76.67 | 219.53 | 135.09 | 163.31 | T | 133.90 | 226.26 | F | RNA binding;large ribosomal subunit rRNA binding;structural constituent of ribosome | | cytosolic large ribosomal subunit;nucleus |
| PF3D7_0903200 | RAB7 | ras-related protein RAB7 | 23.79 | no | 0 | 26.95 | 77.56 | 144.27 | 30.07 | 86.71 | 92.76 | 61.79 | T | 61.14 | 39.61 | M | GTPase activity;protein binding;retromer complex binding | N/A | endosome;membrane;vacuole |
| PF3D7_1473200 | N/A | DnaJ protein, putative | 52.45 | no | 0 | 12.11 | 48.15 | 95.86 | 26.66 | 6.34 | 10.24 | 117.91 | O | 128.29 | 44.19 | M | protein binding | response to heat;response to unfolded protein | Maurer's cleft;nucleus |
| PF3D7_1473700 | NUP116 | nucleoporin NUP116/NSP116, putative | 142.21 | no | 0 | 7.91 | 3.13 | 4.78 | 0.26 | 1.72 | 5.92 | 52.39 | O | 396.00 | 18.66 | M | double-stranded DNA binding;structural constituent of nuclear pore;transcription coregulator activity | protein import into nucleus;regulation of transcription by RNA polymerase II | nuclear pore central transport channel;nucleus |
| PF3D7_1225800 | UBA1 | ubiquitin-activating enzyme E1 | 131.78 | no | 0 | 50.93 | 19.32 | 50.76 | 12.50 | 4.07 | 14.84 | 29.95 | R | 39.60 | 28.32 | M | ubiquitin activating enzyme activity | cellular response to DNA damage stimulus;protein modification by small protein conjugation;protein ubiquitination;response to xenobiotic stimulus;ubiquitin-dependent protein catabolic process | cytoplasm;nucleus |
| PF3D7_0317000 | N/A | proteasome subunit alpha type-3, putative | 29.29 | no | 0 | 49.71 | 19.84 | 48.63 | 22.49 | 129.68 | 215.14 | 67.31 | GV | 23.66 | 27.83 | 0 | endopeptidase activity | proteasomal protein catabolic process;proteasomal ubiquitin-independent protein catabolic process;proteasome-mediated ubiquitin-dep | cytoplasm;nucleus;proteasome core complex;proteasome core complex, alpha-subunit complex |
| PF3D7_1361100 | SEC24A | protein transport protein Sec24A | 106.68 | no | 0 | 303.85 | 28.64 | 146.81 | 112.42 | 28.47 | 24.40 | 105.46 | R | 109.38 | 37.73 | M | RNA binding;SNARE binding;zinc ion binding | COPII-coated vesicle cargo loading;endoplasmic reticulum to Golgi vesicle-mediated transport;protein transport by the Sec complex | COPII vesicle coat;Maurer's cleft;endoplasmic reticulum;endoplasmic reticulum exit site;nucleus |
| PF3D7_0726400 | N/A | conserved Plasmodium membrane protein | 599.32 | no | 8 | 14.59 | 4.99 | 10.86 | 2.74 | 2.25 | 28.13 | 13.14 | GV | 43.22 | 13.70 | M | 5'-flap endonuclease activity;flap endonuclease activity | DNA repair | integral component of membrane;nucleus |
| PF3D7_1419700 | EIF3H | eukaryotic translation initiation factor 3 subunit H, putative | 59.69 | no | 0 | 301.05 | 106.68 | 60.81 | 7.43 | 12.45 | 42.27 | 112.22 | R | 180.46 | 98.93 | M | mRNA binding | initiation of translation | eukaryotic translation initiation factor 3 complex;nucleolus;nucleus;cytoplasm |
| PF3D7_1244600 | ARFGAP1 | ADP-ribosylation factor GTPase-activating protein 1 | 37.29 | no | 0 | 27.62 | 8.07 | 26.61 | 7.23 | 10.72 | 40.17 | 82.97 | O | 53.54 | 69.73 | F | GTPase activator activity | COPI coating of Golgi vesicle;regulation of endocytosis | cytosol |
| PF3D7_1021800 | SEA1 | schizont egress antigen-1 | 244.90 | no | 0 | 10.76 | 3.46 | 36.90 | 20.75 | 6.79 | 25.71 | 818.39 | O | 183.77 | 74.22 | M | N/A | exit from host cell;nuclear chromosome segregation;schizogony | Maurer's cleft;chromosome, centromeric region;host cell membrane;nucleus;symbiont-containing vacuole membrane |
| PF3D7_0321600 | DDX42 | ATP-dependent RNA helicase DDX42, putative | 86.65 | no | 0 | 13.26 | 14.71 | 18.84 | 8.28 | 4.52 | 25.40 | 61.68 | O | 71.84 | 89.96 | F | RNA binding;RNA helicase activity | regulation of transcription | cytoplasm;nucleus;ribonucleoprotein complex |
| PF3D7_1453000 | N/A | conserved Plasmodium protein, unknown function | 110.62 | no | 0 | 24.80 | 8.24 | 5.80 | 1.09 | 1.56 | 12.65 | 10.44 | R | 9.64 | 21.94 | F | N/A | N/A | N/A |
| PF3D7_1112500 | N/A | RNA transcription, translation and transport factor protein, putative | 33.66 | no | 0 | 16.63 | 4.92 | 4.37 | 1.13 | 6.51 | 32.14 | 12.97 | GV | 31.72 | 6.80 | M | N/A | N/A | nucleus |
| PF3D7_1334100 | ORC4 | origin recognition complex subunit 4, putative | 117.79 | no | 0 | 4.80 | 2.93 | 13.36 | 2.56 | 3.68 | 11.85 | 44.21 | O | 204.94 | 26.96 | M | DNA binding | regulation of replication | nucleus |
| PF3D7_0709700 | PARE | prodrug activation and resistance esterase | 42.36 | no | 0 | 19.60 | 12.92 | 60.12 | 73.95 | 40.58 | 620.60 | 173.06 | GV | 18.67 | 830.89 | F | acylglycerol lipase activity;hydrolase activity;hydrolase activity, acting on ester bonds;lipase activity | monoacylglycerol catabolic process;phospholipid metabolic process;response to xenobiotic stimulus | cytoplasm;membrane |
| PF3D7_1420700 | P113 | surface protein P113 | 112.57 | yes | 1 | 553.41 | 343.81 | 141.56 | 10.51 | 15.53 | 16.57 | 132.15 | R | 77.50 | 8.23 | M | protein binding | vacuole organization | cell surface;plasma membrane;symbiont-containing vacuole;symbiont-containing vacuole membrane |
| PF3D7_1139800 | N/A | conserved Plasmodium protein, unknown function | 265.26 | no | 0 | 10.89 | 3.13 | 8.33 | 1.86 | 1.77 | 20.65 | 11.09 | GV | 7.28 | 7.85 | 0 | N/A | N/A | N/A |
| PF3D7_1336400 | N/A | RanBPM and CLTH-like protein, putative | 33.73 | no | 0 | 5.55 | 1.02 | 4.89 | 0.94 | 6.85 | 40.03 | 20.63 | GV | 50.96 | 22.19 | M | protein binding | proteasome-mediated ubiquitin-dependent protein catabolic process | cytoplasm;nucleus |
| PF3D7_1143200 | N/A | DnaJ protein, putative | 39.36 | yes | 2 | 2.89 | 1.20 | 14.20 | 8.05 | 2.29 | 30.91 | 52.43 | O | 8.67 | 29.66 | F | protein binding | protein folding | endoplasmic reticulum membrane |
| PF3D7_0824800 | N/A | conserved Plasmodium membrane protein | 154.87 | no | 4 | 33.19 | 14.08 | 8.48 | 2.05 | 2.34 | 22.54 | 39.11 | O | 43.49 | 23.65 | M | N/A | N/A | transcription regulator complex;nucleus;integral component of membrane |
| PF3D7_1409800 | CELF2 | CUGBP Elav-like family member 2, putative | 59.27 | no | 0 | 5.96 | 31.04 | 20.57 | 1.59 | 19.66 | 39.45 | 52.01 | O | 63.42 | 75.90 | F | RNA binding;mRNA binding | regulation of translation | P granule;cytoplasm;nucleus;ribonucleoprotein complex |
| PF3D7_1459400 | N/A | conserved protein, unknown function | 40.07 | no | 1 | 57.25 | 3.15 | 19.53 | 9.52 | 7.99 | 43.79 | 8.68 | R | 0.78 | 4.09 | F | RNA binding | entry into host | nucleus |
| PF3D7_1343900 | PRPF4 | U4/U6 small nuclear ribonucleoprotein PRP4, putative | 76.19 | no | 0 | 14.18 | 5.48 | 8.13 | 2.06 | 2.09 | 12.27 | 7.10 | R | 37.98 | 6.38 | M | U4 snRNA binding;U6 snRNA binding | mRNA splicing, via spliceosome | U4/U6 x U5 tri-snRNP complex |
| PF3D7_1242800 | rabGDI | rab specific GDP dissociation inhibitor | 52.34 | no | 0 | 154.52 | 15.74 | 47.64 | 15.08 | 26.68 | 36.90 | 86.14 | R | 7.24 | 26.68 | F | GTPase activator activity;Rab GDP-dissociation inhibitor activity | vesicle-mediated transport | cytoplasm;nucleus |
| PF3D7_1015600 | HSP60 | heat shock protein 60 | 62.55 | no | 0 | 33.35 | 234.06 | 124.14 | 25.15 | 8.70 | 13.12 | 33.20 | T | 67.10 | 86.13 | F | ATP hydrolysis activity;protein binding | protein folding;protein targeting to mitochondrion;response to unfolded protein | Maurer's cleft;cytoplasm;food vacuole;mitochondrion;nucleus |
| PF3D7_1224000 | GCH1 | GTP cyclohydrolase 1 | 45.88 | no | 0 | 30.08 | 15.28 | 18.06 | 1.41 | 6.59 | 39.64 | 36.17 | GV | 82.07 | 18.30 | M | GTP binding;GTP cyclohydrolase I activity;zinc ion binding | folic acid-containing compound biosynthetic process;one-carbon metabolic process;response to xenobiotic stimulus;tetrahydrobiopterin biosynthetic process | cytoplasm;mitochondrion;nucleus |
| PF3D7_1332900 | cyto-IRS | isoleucine--tRNA ligase, putative | 151.31 | no | 0 | 70.15 | 27.52 | 34.03 | 6.90 | 2.42 | 9.67 | 43.49 | R | 27.65 | 22.51 | 0 | isoleucine-tRNA ligase activity | isoleucyl-tRNA aminoacylation;response to xenobiotic stimulus | cytoplasm;nucleus |
| PF3D7_1317800 | RPS19 | 40S ribosomal protein S19 | 16.76 | no | 0 | 284.85 | 310.85 | 207.25 | 24.19 | 182.37 | 237.39 | 185.00 | T | 153.69 | 108.15 | M | RNA binding;structural constituent of ribosome | ribosomal small subunit assembly;translation | cytosolic small ribosomal subunit;nucleus |
| PF3D7_1470900 | N/A | proteasome subunit beta type-2, putative | 22.86 | no | 0 | 23.46 | 23.56 | 66.26 | 14.27 | 45.07 | 169.73 | 118.03 | GV | 34.92 | 63.97 | F | endopeptidase activity | proteasomal protein catabolic process;proteasomal ubiquitin-independent protein catabolic process;proteasome-mediated ubiquitin-dependent protein catabolic process;ubiquitin-dependent protein catabolic process | cytoplasm;extracellular vesicle;nucleus;proteasome core complex |
| PF3D7_1139300 | AP2-G5 | AP2 domain transcription factor AP2-G5 | 309.44 | no | 0 | 64.79 | 41.42 | 12.07 | 0.88 | 1.99 | 22.23 | 190.04 | O | 120.04 | 21.26 | M | DNA-binding transcription factor activity;protein binding;sequence-specific DNA binding | regulation of transcription, DNA-templated | heterochromatin;nucleus |
| PF3D7_1036900 | N/A | conserved Plasmodium protein, unknown function | 192.68 | no | 0 | 225.83 | 57.69 | 32.04 | 4.98 | 21.91 | 169.45 | 201.15 | R | 313.27 | 168.45 | M | mRNA binding | N/A | RNA N6-methyladenosine methyltransferase complex;nucleus |
| PF3D7_1004000 | N/A | 60S ribosomal protein L13, putative | 23.75 | no | 0 | 303.74 | 246.76 | 124.08 | 8.83 | 112.30 | 94.36 | 260.90 | R | 253.53 | 64.17 | M | mRNA binding;structural constituent of ribosome | negative regulation of translation;translation | cytosolic large ribosomal subunit;nucleus;ribosome |
| PF3D7_1322100 | SET2 | histone-lysine N-methyltransferase SET2 | 300.71 | no | 0 | 303.74 | 17.82 | 6.80 | 0.55 | 2.71 | 17.71 | 60.61 | R | 7.37 | 33.13 | F | chromatin binding;histone methyltransferase activity (H3-K36 specific);histone-lysine N-methyltransferase activity;nucleosomal histone binding;protein binding | antigenic variation;histone H3-K36 methylation;histone H3-K36 trimethylation;regulation of transcription, DNA-templated | chromatin;chromosome;nuclear periphery;nucleus |
| PF3D7_0316300 | Ppase | Inorganic pyrophosphatase, putative | 51.31 | yes | 1 | 32813.00 | 25020.00 | 193.11 | 68.21 | 17.34 | 107.1 | 597.98 | R | 213.82 | 292.53 | F | inorganic diphosphatase activity;magnesium ion binding;pyrophosphatase activity | phosphate-containing compound metabolic process | apicoplast;cytosol |
| PF3D7_1459200 | N/A | WD repeat-containing protein, putative | 113.23 | no | 0 | 303.74 | 15.66 | 10.64 | 1.63 | 5.33 | 14.63 | 32.88 | R | 21.53 | 12.09 | M | protein binding | N/A | N/A |
| PF3D7_0707400 | ATAD3A | ATPase family AAA domain-containing protein 3A, putative | 78.19 | no | 0 | 10.60 | 32.11 | 17.27 | 1.77 | 4.18 | 31.99 | 85.93 | O | 45.42 | 61.66 | F | zinc ion binding | mitochondrion organization | mitochondrion;nucleus |
| PF3D7_1440700 | AP3M | AP-3 complex subunit mu, putative | 87.17 | no | 0 | 11.79 | 16.65 | 17.41 | 4.00 | 2.93 | 14.31 | 4.29 | T | 4.18 | 5.04 | 0 | protein binding | endocytosis;vesicle-mediated transport | AP-3 adaptor complex;cytoplasmic vesicle;intracellular membrane-bounded organelle |
| PF3D7_1439500 | ORP2 | oocyst rupture protein 2, putative | 123.02 | no | 0 | 6.12 | 0.56 | 10.69 | 6.11 | 3.91 | 20.15 | 10.89 | GV | 74.44 | 7.59 | M | DNA-binding transcription activator activity, RNA polymerase II-specific;RNA polymerase II cis-regulatory region sequence-specific DNA binding | regulation of transcription, DNA-templated | cell periphery;cytoplasm;nucleus |
| PF3D7_1363400 | DOA1 | polyubiquitin binding protein, putative | 107.61 | no | 0 | 19.56 | 3.38 | 11.03 | 2.44 | 4.07 | 29.05 | 22.04 | GV | 54.40 | 15.11 | M | protein binding;ubiquitin binding | proteasome-mediated ubiquitin-dependent protein catabolic process | cytoplasm |
| PF3D7_1448300 | N/A | conserved protein, unknown function | 191.02 | no | 0 | 25.46 | 26.85 | 41.64 | 16.79 | 8.09 | 25.04 | 27.92 | T | 46.06 | 36.43 | M | N/A | N/A | nucleus |
| PF3D7_1220100 | PRP17 | pre-mRNA-processing factor 17, putative | 72.54 | no | 0 | 23.80 | 6.83 | 6.56 | 0.80 | 2.46 | 16.04 | 83.11 | O | 28.74 | 13.83 | M | RNA binding | RNA splicing, via transesterification reactions;generation of catalytic spliceosome for second transesterification step;mRNA 3'-splice site recognition | spliceosomal complex;nucleus |
| PF3D7_1121700 | GCN20 | protein GCN20 | 95.37 | no | 0 | 52.14 | 34.67 | 13.09 | 0.80 | 4.85 | 9.01 | 25.14 | R | 14.82 | 11.87 | | ATP binding;ATPase-coupled transmembrane transporter activity | regulation of translation | nucleus |
| PF3D7_0413500 | PGM2 | phosphoglucomutase-2 | 35.23 | no | 1 | 6.84 | 0.61 | 41.36 | 15.59 | 6.93 | 24.15 | 28.04 | T | 51.32 | 44.82 | M | phosphoglycerate mutase activity | glycolytic process | cytoplasm;mitochondrion |
| PF3D7_0932300 | M18AAP | M18 aspartyl aminopeptidase | 65.63 | no | 0 | 22.90 | 14.45 | 15.17 | 2.43 | 1.59 | 9.75 | 51.93 | O | 47.20 | 82.35 | F | metallopeptidase activity;zinc ion binding | proteolysis | cytosol;nucleus |
| PF3D7_0211800 | AsnRS | asparagine--tRNA ligase | 70.50 | no | 0 | 206.62 | 66.39 | 69.96 | 9.22 | 9.83 | 11.56 | 80.93 | R | 114.67 | 86.61 | M | asparagine-tRNA ligase activity | asparaginyl-tRNA aminoacylation | nucleus |
| PF3D7_0411100 | MED8 | mediator of RNA polymerase II transcription subunit 8 | 23.97 | no | 0 | 34.82 | 2.70 | 5.48 | 0.45 | 4.61 | 105.53 | 52.18 | GV | 39.21 | 54.38 | F | protein binding | regulation of transcription | mediator complex;nucleus |
| PF3D7_0403600 | N/A | conserved Plasmodium protein, unknown function | 89.43 | no | 0 | 5.93 | 2.34 | 49.84 | 69.47 | 2.65 | 26.00 | 450.28 | O | 24.32 | 35.56 | F | N/A | N/A | N/A |

| | | | | | | | | | | | | | | | | | | | |
|---|---|---|---|---|---|---|---|---|---|---|---|---|---|---|---|---|---|---|---|
| PF3D7_0102200 | RESA | ring-infected erythrocyte surface antigen | 126.52 | no | 0 | 1397.96 | 11.00 | 166.61 | 23.76 | 1.59 | 1.73 | 422.52 | R | 0.46 | 0.19 | O | protein binding | regulation of immune response | extracellular vesicle;food vacuole;host cell plasma membrane;host cell surface;merozoite dense granule;symbiont-containing vacuole |
| PF3D7_1244200 | TFB2 | RNA polymerase II transcription factor B subunit 2, putative | 111.87 | no | 0 | 73.96 | 19.46 | 11.81 | 1.23 | 2.67 | 21.02 | 38.76 | R | 14.92 | 18.47 | F | double-stranded DNA binding | nucleotide-excision repair;phosphorylation of RNA polymerase II C-terminal domain;transcription by RNA polymerase II | nucleus;transcription factor TFIIH core complex;transcription factor TFIIH holo complex |
| PF3D7_1352500 | N/A | thioredoxin-related protein, putative | 23.99 | yes | 2 | 69.07 | 41.33 | 201.48 | 185.08 | 23.87 | 36.67 | 73.45 | T | 156.49 | 78.31 | M | RNA binding;protein disulfide isomerase activity | N/A | food vacuole;nucleus |
| PF3D7_1317300 | N/A | RNA-binding protein, putative | 34.86 | no | 0 | 13.03 | 28.33 | 29.23 | 7.61 | 16.02 | 68.32 | 42.74 | GV | 144.93 | 56.17 | M | N/A | N/A | nucleus |
| PF3D7_1368800 | ERCC4 | DNA repair endonuclease XPF, putative | 204.53 | no | 0 | 4.91 | 1.45 | 10.48 | 2.34 | 2.25 | 22.83 | 77.44 | O | 38.48 | 19.24 | M | DNA binding;damaged DNA binding;endodeoxyribonuclease activity;single-stranded DNA binding;single-stranded DNA endodeoxyribonuclease activity | DNA repair;double-strand break repair via homologous recombination;nucleotide-excision repair involved in interstrand cross-link repair;nucleotide-excision repair, DNA incision, 5'-to lesion;resolution of meiotic recombination intermediates | nucleotide-excision repair factor 1 complex |
| PF3D7_0405600 | N/A | TMEM33 domain-containing protein, putative | 37.81 | no | 4 | 29.88 | 9.66 | 15.74 | 2.50 | 28.31 | 91.48 | 51.26 | GV | 29.87 | 45.59 | F | protein binding | endoplasmic reticulum tubular network organization;membrane organization | endoplasmic reticulum |
| PF3D7_0516900 | RPL2 | 60S ribosomal protein L2 | 28.00 | no | 0 | 1604.47 | 3148.30 | 873.98 | 471.41 | 68.91 | 50.94 | 352.43 | T | 194.22 | 193.81 | O | RNA binding;structural constituent of ribosome | cytoplasmic translation;translation | cytosolic large ribosomal subunit;nucleus |
| PF3D7_0810600 | DBP1 | ATP-dependent RNA helicase DBP1, putative | 108.63 | no | 0 | 122.19 | 50.03 | 29.03 | 2.43 | 14.18 | 87.40 | 88.86 | R | 132.97 | 88.01 | M | RNA binding;RNA helicase activity;mRNA binding | regulation of transcription | P granule;nucleus |
| PF3D7_0111300 | N/A | P-loop containing nucleoside triphosphate hydrolase | 139.21 | no | 0 | 4.91 | 2.18 | 12.37 | 2.73 | 2.33 | 21.09 | 24.92 | O | 171.23 | 27.02 | M | DNA binding | DNA replication | DNA replication factor C complex;nucleus |
| PF3D7_1230700 | SEC13 | protein transport protein SEC13 | 90.75 | no | 0 | 486.15 | 167.29 | 448.51 | 624.52 | 33.84 | 16.17 | 97.63 | S | 92.83 | 56.97 | M | protein binding | COPII-coated vesicle budding | COPII vesicle coat;Maurer's cleft;cytosol;endoplasmic reticulum exit site;nuclear pore;nuclear pore outer ring;nucleoplasm |
| PF3D7_1129900 | MFR5 | major facilitator superfamily-related transporter, putative | 70.14 | no | 12 | 87.18 | 178.41 | 19.30 | 4.38 | 29.80 | 32.06 | 185.51 | O | 78.19 | 28.17 | M | N/A | N/A | plasma membrane;integral component of membrane |
| PF3D7_1460700 | RPL27 | 60S ribosomal protein L27 | 16.75 | no | 0 | 232.90 | 242.16 | 76.26 | 12.19 | 80.06 | 73.49 | 234.32 | T | 350.76 | 353.44 | O | RNA binding;structural constituent of ribosome | translation | cytosolic large ribosomal subunit;nucleus |
| PF3D7_1330400 | N/A | ER lumen protein retaining receptor 1, putative | 32.81 | no | 8 | 123.88 | 12.22 | 74.45 | 32.88 | 27.04 | 66.69 | 189.70 | O | 214.09 | 21.40 | M | ER retention sequence binding;HDEL sequence binding | endoplasmic reticulum to Golgi vesicle-mediated transport;protein retention in ER lumen | Golgi apparatus;cis-Golgi network;endoplasmic reticulum |
| PF3D7_0513300 | PNP | purine nucleoside phosphorylase | 26.86 | no | 0 | 215.33 | 280.18 | 266.95 | 27.55 | 43.75 | 25.49 | 333.54 | O | 6.92 | 84.51 | F | purine-nucleoside phosphorylase activity | inosine catabolic process;purine nucleoside catabolic process;response to xenobiotic stimulus | nucleus |
| PF3D7_1135100 | PPM8 | protein phosphatase PPM8, putative | 79.93 | no | 0 | 61.84 | 12.93 | 21.60 | 13.23 | 3.33 | 51.75 | 15.64 | R | 42.47 | 125.43 | F | magnesium-dependent protein serine/threonine phosphatase activity;protein serine/threonine phosphatase activity | | cytoplasm;nucleus |
| PF3D7_0604600 | N/A | DNA helicase, putative | 206.43 | no | 0 | 4.44 | 1.15 | 1.88 | 0.27 | 0.86 | 5.31 | 20.11 | O | 9.57 | 4.24 | M | ATP-dependent DNA/DNA annealing activity;ATP-dependent activity, acting on DNA;DNA helicase activity;endodeoxyribonuclease activity | DNA repair;DNA rewinding;cellular response to DNA damage stimulus;replication fork processing;replication fork protection | nuclear replication fork;nucleus |
| PF3D7_0934800 | PKAc | cAMP-dependent protein kinase catalytic subunit | 40.20 | no | 0 | 11.11 | 1.24 | 56.80 | 63.13 | 4.29 | 18.80 | 154.44 | O | 145.68 | 25.66 | M | cAMP-dependent protein kinase activity;protein binding | cell differentiation;entry into host;peptidyl-serine phosphorylation;signal transduction | cAMP-dependent protein kinase complex;nucleus |
| PF3D7_0218500 | SNRPD2 | small nuclear ribonucleoprotein Sm D2, putative | 11.71 | no | 0 | 70.47 | 20.33 | 14.76 | 1.80 | 55.51 | 57.77 | 38.27 | R | 61.34 | 33.14 | M | RNA binding | RNA splicing;RNA splicing, via transesterification reactions;spliceosomal snRNP assembly | U1 snRNP;U2 snRNP;U4/U6 x U5 tri-snRNP complex;U5 snRNP;catalytic step 2 spliceosome;precatalytic spliceosome;spliceosomal complex;nucleus |
| PF3D7_1363600 | N/A | conserved Plasmodium protein, unknown function | 27.36 | no | 0 | 5.20 | 4.07 | 7.46 | 1.76 | 14.44 | 35.92 | 0.00 | GV | 24.62 | 8.88 | M | N/A | N/A | mediator complex;nucleus |
| PF3D7_1009200 | N/A | ribonuclease, putative | 106.46 | no | 0 | 32.31 | 5.41 | 3.43 | 0.64 | 2.47 | 15.81 | 48.29 | O | 9.86 | 17.02 | F | enzymatic activity;RNA binding | RNA processing | nucleus |
| PF3D7_1426800 | N/A | conserved Plasmodium protein, unknown function | 248.50 | no | 0 | 41.44 | 16.25 | 7.34 | 0.72 | 1.84 | 16.93 | 16.58 | R | 80.40 | 14.97 | M | N/A | N/A | N/A |
| PF3D7_1446700 | AKIT4 | apicomplexan kinetochore protein 4, putative | 42.63 | no | 0 | 20210.00 | 45047.00 | 25.67 | 26481.00 | 31686.00 | 89.37 | 62.00 | T | 62.14 | 68.67 | F | DNA binding | DNA replication | kinetochore;nucleus |
| PF3D7_1427900 | N/A | Leucine-rich repeat protein | 33.01 | no | 0 | 15.73 | 45348.00 | 73.32 | 13.72 | 13.88 | 37.15 | 122.83 | T | 304.46 | 45554.00 | F | histone binding | N/A | nucleus |
| PF3D7_1356100 | N/A | conserved Plasmodium protein, unknown function | 113.75 | no | 0 | 7.55 | 13.51 | 31.60 | 17.96 | 13.50 | 137.73 | 121.61 | GV | 98.95 | 875.90 | F | N/A | N/A | nucleus |
| PF3D7_0927600 | N/A | RNA-binding protein, putative | 69.55 | no | 0 | 3.37 | 2.49 | 22.11 | 7.55 | 5.67 | 117.60 | 217.73 | O | 76.28 | 555.34 | F | histone pre-mRNA stem-loop binding;mRNA binding | mRNA 3'-end processing by stem-loop binding and cleavage;mRNA transport | cytoplasm;histone pre-mRNA 3'end processing complex;nucleus |
| PF3D7_1235500 | MT-A70.2 | N6-adenosine-methyltransferase, putative | 75.05 | no | 0 | 29.19 | 22.46 | 23.97 | 3.11 | 9.87 | 13.82 | 74.37 | O | 54.81 | 69.24 | F | mRNA binding;protein binding | mRNA methylation | RNA N6-methyladenosine methyltransferase complex;nucleus |
| PF3D7_0530200 | PPT | phosphoenolpyruvate/phosphate translocator | 59.79 | yes | 7 | 6.52 | 1.41 | 30.78 | 5.98 | 13.38 | 26.30 | 72.22 | O | 18.70 | 14.17 | M | GDP-fucose transmembrane transporter activity;antiporter activity;glucose 6-phosphate:inorganic phosphate antiporter activity;nucleotide-sugar transmembrane transporter activity;transmembrane transporter activity;triose-phosphate:phosphate antiporter activity | glucose import | Golgi apparatus;apicoplast;integral component of plastid inner membrane;plastid inner membrane |
| PF3D7_1014900 | KIC8 | protein KIC8 | 267.98 | no | 0 | 6.48 | 0.81 | 13.39 | 11.94 | 1.25 | 57.32 | 227.16 | O | 12.04 | 357.97 | F | N/A | N/A | nucleus |
| PF3D7_1119700 | N/A | conserved Plasmodium protein, unknown function | 41.77 | no | 0 | 7.34 | 1.02 | 3.58 | 1.38 | 3.27 | 220.25 | 10.15 | GV | 16.87 | 362.10 | F | N/A | N/A | N/A |
| PF3D7_0519800 | N/A | EELM2 domain-containing protein, putative | 78.29 | no | 0 | 162.14 | 132.95 | 33.53 | 7.72 | 14.95 | 22.80 | 1124.78 | O | 128.29 | 63.31 | M | lysine-acetylated histone binding | | nucleus |
| PF3D7_1462400 | N/A | conserved Plasmodium protein, unknown function | 395.67 | no | 0 | 68.54 | 16.91 | 5.10 | 0.31 | 2.07 | 15.03 | 16.52 | R | 11.57 | 23.84 | F | N/A | N/A | N/A |
| PF3D7_1121300 | TKL2 | tyrosine kinase-like protein | 200.42 | no | 0 | 4.40 | 0.32 | 21.82 | 7.67 | 2.40 | 3.27 | 13.05 | T | 2.06 | 0.97 | O | enzymatic activity | signal transduction | host cell cytosol |
| PF3D7_1359400 | CELF1 | CUGBP Elav-like family member 1 | 57.11 | no | 0 | 171.39 | 220.50 | 137.97 | 29.61 | 25.06 | 72.25 | 76.36 | T | 174.59 | 62.10 | M | RNA binding;mRNA binding | N/A | cytoplasm;nucleus;ribonucleoprotein complex |
| PF3D7_1329500 | N/A | conserved protein, unknown function | 218.44 | no | 0 | 527.52 | 33.64 | 25.15 | 4.57 | 5.28 | 20.58 | 337.46 | R | 14.02 | 32.26 | F | N/A | N/A | nucleus |
| PF3D7_0726300 | PMS1 | DNA mismatch repair protein PMS1, putative | 156.84 | no | 0 | 22.25 | 2.79 | 10.82 | 3.61 | 1.81 | 10.37 | 51.67 | O | 114.31 | 7.48 | M | ATP binding;ATP hydrolysis activity;DNA binding;damaged DNA binding | DNA replication;mismatch repair | MutLalpha complex;mismatch repair complex;nucleus |
| PF3D7_0912900 | RPN8 | 26S proteasome regulatory subunit RPN8, putative | 39.14 | no | 0 | 108.90 | 25.82 | 57.04 | 13.57 | 17.04 | 66.53 | 193.85 | O | 39.88 | 25.37 | M | protein binding | proteasome-mediated ubiquitin-dependent protein catabolic process | proteasome complex; nucleus; cytoplasm |
| PF3D7_0822300 | SNRPG | small nuclear ribonucleoprotein G, putative | 9.51 | no | 0 | 84.40 | 17.64 | 13.19 | 2.06 | 58.39 | 259.25 | 63.08 | GV | 24.96 | 27.20 | O | RNA binding | RNA splicing, via transesterification reactions;mRNA splicing, via spliceosome | P granule;SMN-Sm protein complex;U1 snRNP;U12-type spliceosomal complex;U2 snRNP;U2-type prespliceosome;U4 snRNP;U4/U6 x U5 tri-snRNP complex;U5 snRNP;catalytic step 2 spliceosome;precatalytic spliceosome;ribonucleoprotein complex;sno(s)RNA-containing ribonucleoprotein complex;spliceosomal tri-snRNP |
| PF3D7_0817300 | N/A | conserved Plasmodium protein, unknown function | 254.75 | no | 0 | 20.79 | 106.98 | 59.01 | 3.53 | 2.99 | 16.04 | 62.13 | T | 17.29 | 23.85 | F | N/A | N/A | nucleus |
| PF3D7_1369500 | N/A | nuclear cap-binding protein subunit 1, putative | 131.37 | no | 0 | 69.83 | 17.18 | 7.99 | 0.77 | 0.74 | 7.36 | 11.75 | R | 19.26 | 13.68 | M | mRNA cap binding | regulation of translation | nucleus |
| PF3D7_0105200 | RAP1 | RAP protein RAP1 | 67.16 | no | 0 | 10.85 | 15.12 | 17.66 | 1.86 | 5.71 | 34.28 | 51.63 | O | 29.17 | 20.47 | M | mRNA binding;mitochondrial ribosome binding;rRNA binding | N/A | mitochondrion |
| PF3D7_1123800 | SMC5 | structural maintenance of chromosomes protein 5, putative | 152.39 | no | 0 | 5.28 | 2.99 | 10.68 | 3.11 | 1.86 | 11.98 | 58.11 | O | 14.60 | 32.68 | F | chromatin binding | N/A | mitochondrion |
| PF3D7_1304900 | RPB11 | DNA-directed RNA polymerase II subunit RPB11, putative | 14.13 | no | 0 | 30.48 | 15.12 | 15.53 | 2.29 | 13.84 | 18.36 | 8.41 | R | 20.42 | 5.66 | M | DNA-directed 5'-3' RNA polymerase activity | transcription by RNA polymerase II | RNA polymerase II, core complex;RNA polymerase II, holoenzyme;nucleus |
| PF3D7_1425900 | N/A | protein HGH1, putative | 43.16 | no | 0 | 97.07 | 44.36 | 10.04 | 1.19 | 16.18 | 45.26 | 42.05 | R | 10.29 | 22.48 | F | protein binding | regulation of transcription: TF chaperone | nucleus |
| PF3D7_1409600 | N/A | histidine phosphatase, putative | 92.75 | no | 2 | 17.84 | 18.04 | 24.28 | 4.70 | 1.77 | 9.91 | 5.50 | T | 15.11 | 3.46 | M | intramolecular transferase activity, phosphotransferases | N/A | cytoplasm |
| PF3D7_1307000 | RRP40 | exosome complex component RRP40, putative | 28.29 | no | 0 | 44.01 | 14.02 | 11.76 | 1.32 | 18.46 | 19.31 | 8.05 | R | 11.04 | 6.87 | M | RNA binding | CUT catabolic process;U4 snRNA 3'-end processing;exonucleolytic catabolism of deadenylated mRNA;exonucleolytic trimming to generate mature 3'-end of 5.8S rRNA from tricistronic rRNA transcript (SSU-rRNA, 5.8S rRNA, LSU-rRNA);nuclear polyadenylation-dependent rRNA catabolic process;nuclear polyadenyl | cytoplasmic exosome (RNase complex);exosome (RNase complex);nuclear exosome (RNase complex) |
| PF3D7_1129800 | N/A | Brr1 domain-containing protein | 33.66 | no | 2 | 10.37 | 1.65 | 6.28 | 0.37 | 4.78 | 18.31 | 24.56 | O | 6.05 | 6.73 | O | RNA binding | spliceosomal complex activity | nucleus |
| PF3D7_1125500 | SNRPD1 | small nuclear ribonucleoprotein Sm D1, putative | 14.34 | no | 0 | 75.14 | 24.58 | 11.72 | 1.08 | 30.60 | 89.63 | 54.68 | GV | 13.90 | 10.38 | O | RNA binding;protein binding | RNA splicing;RNA splicing, via transesterification reactions;mRNA splicing, via spliceosome;spliceosomal snRNP assembly | SMN-Sm protein complex;U1 snRNP;U12-type spliceosomal complex;U2 snRNP;U4 snRNP;U5 snRNP;catalytic step 2 spliceosome;commitment complex;nucleus;pICln-Sm protein complex;precatalytic spliceosome;small nuclear ribonucleoprotein complex;spliceosomal tri-snRNP complex |
| PF3D7_0210100 | N/A | ribosomal protein L37ae, putative | 10.80 | no | 0 | 550.93 | 482.78 | 198.68 | 72.4 | 527.03 | 818.94 | 293.43 | GC V | 118.29 | 475.14 | F | RNA binding;structural constituent of ribosome | translation | cytosolic large ribosomal subunit;nucleus |
| PF3D7_0808400 | SEC28 | coatomer subunit epsilon, putative | 33.09 | no | 0 | 28.61 | 4.26 | 8.06 | 4.46 | 18.59 | 73.04 | 24.60 | GV | 61.17 | 27.04 | M | protein binding | endoplasmic reticulum to Golgi vesicle-mediated transport;intra-Golgi vesicle-mediated transport | COPI vesicle coat |
| PF3D7_1446600 | CEN2 | centrin-2 | 19.31 | no | 0 | 6.49 | 2.71 | 102.92 | 55.53 | 79.45 | 450.14 | 625.36 | O | 213.42 | 717.72 | F | RNA binding;calcium ion binding | centriole replication;mitotic cell cycle | centriole;centrosome;nucleus |
| PF3D7_0303300 | RPB6 | DNA-directed RNA polymerases I, II, and III subunit RPABC2, putative | 17.83 | no | 0 | 26.80 | 9.18 | 8.42 | 0.74 | 19.62 | 37.33 | 33.93 | GV | 19.80 | 10.59 | M | DNA-directed 5'-3' RNA polymerase activity | transcription by RNA polymerase II | RNA polymerase I complex;RNA polymerase II, core complex;RNA polymerase III complex |
| PF3D7_0621900 | SRP68 | signal recognition particle subunit SRP68, putative | 92.45 | no | 0 | 136.08 | 29.71 | 35.45 | 16.11 | 8.15 | 15.34 | 18.22 | R | 32.73 | 24.74 | M | RNA binding;signal recognition particle binding | protein processing | signal recognition particle, endoplasmic reticulum targeting |
| PF3D7_0503200 | CDC6 | cell division control protein 6, putative | 115.09 | no | 0 | 12.88 | 4.90 | 10.67 | 2.32 | 1.87 | 83.12 | 330.08 | O | 570.22 | 504.11 | M | DNA replication origin binding | DNA replication initiation;mitotic DNA replication checkpoint signaling;mitotic cell cycle | mitochondrion;nuclear origin of replication recognition complex |
| PF3D7_1469800 | MED22 | mediator of RNA polymerase II transcription subunit 22 | 25.17 | no | 0 | 7.43 | 10.00 | 10.90 | 3.11 | 113.91 | 815.29 | 172.11 | GV | 200.55 | 434.25 | F | protein binding | regulation of transcription | mediator complex;nucleus |
| PF3D7_0921200 | N/A | conserved Plasmodium membrane protein | 276.94 | no | 6 | 57.04 | 31.00 | 7.06 | 0.53 | 1.62 | 12.75 | 4.20 | R | 16.81 | 5.67 | M | N/A | N/A | integral component of membrane |
| PF3D7_1006800 | GBP2 | G-strand-binding protein 2 | 29.53 | no | 0 | 705.47 | 275.18 | 206.92 | 48.93 | 46.83 | 44.27 | 150.79 | R | 221.83 | 116.06 | M | G-quadruplex DNA binding;G-rich strand telomeric DNA binding;RNA binding;mRNA binding;telomeric DNA binding | | cytoplasm;food vacuole;nucleus;ribonucleoprotein complex |
| PF3D7_0215800 | ORC5 | origin recognition complex subunit 5 | 103.99 | no | 0 | 7.16 | 1.23 | 11.68 | 1.93 | 1.70 | 12.83 | 24.33 | O | 234.01 | 8.45 | M | ATP binding;ATP hydrolysis activity;DNA replication origin binding | DNA replication initiation | nuclear origin of replication recognition complex;nuclear pre-replicative complex |
| PF3D7_1132300 | N/A | nucleic acid binding protein, putative | 31.73 | no | 0 | 3.88 | 7.62 | 11.31 | 3.93 | 11.03 | 74.42 | 187.67 | O | 589.80 | 122.40 | M | RNA binding | | nucleus |
| PF3D7_0512600 | RAB1b | ras-related protein Rab-1B | 22.89 | no | 0 | 88.65 | 16.96 | 107.52 | 31.55 | 153.72 | 399.53 | 151.61 | GV | 84.53 | 358.94 | F | protein binding | protein transport | Maurer's cleft;cis-Golgi network;endoplasmic reticulum;endoplasmic reticulum-Golgi intermediate compartment;food vacuole |
| PF3D7_1022400 | SRSF4 | serine/arginine-rich splicing factor 4 | 62.12 | no | 0 | 259.04 | 261.66 | 121.92 | 24.13 | 9.22 | 12.27 | 101.14 | T | 81.36 | 160.91 | F | RNA binding | RNA splicing;RNA splicing, via transesterification reactions | nucleus |
| PF3D7_1360900 | N/A | RNA-binding protein, putative | 44.81 | no | 0 | 10.41 | 5.12 | 119.79 | 31.44 | 22.50 | 27.40 | 65.92 | T | 301.08 | 75.45 | M | RNA binding | N/A | nucleus;ribonucleoprotein complex |
| PF3D7_0503400 | ADF1 | actin-depolymerizing factor 1 | 13.74 | no | 0 | 18.60 | 30.75 | 314.90 | 127.30 | 272.77 | 236.67 | 598.47 | O | 20.83 | 296.92 | F | actin filament binding;actin monomer binding | actin cytoskeleton organization;actin filament depolymerization;actin filament severing;cell motility;cytoplasmic actin-based contraction involved in cell motility | actin cytoskeleton;cytoplasm;cytosol;nucleus |
| PF3D7_1453700 | P23 | HSP90 co-chaperone p23 | 30.49 | no | 0 | 518.31 | 508.28 | 263.06 | 18.22 | 311.77 | 314.46 | 326.45 | R | 482.70 | 332.69 | M | Hsp90 protein binding;chaperone binding | chaperone-mediated protein complex assembly;negative regulation of ATP-dependent activity;protein folding | cytoplasm;cytosol;nucleus |
| PF3D7_1401800 | CK | choline kinase | 52.21 | no | 0 | 11.05 | 8.95 | 41.04 | 12.01 | 10.95 | 33.22 | 39.24 | T | 33.89 | 29.66 | O | choline binding;choline kinase activity | phosphatidylcholine biosynthetic process;response to xenobiotic stimulus | Maurer's cleft;cytosol |
| PF3D7_1208100 | N/A | conserved Plasmodium protein, unknown function | 533.64 | no | 2 | 21.52 | 11.86 | 2.48 | 0.23 | 1.08 | 12.13 | 8.65 | R | 11.63 | 7.18 | O | N/A | N/A | 90S preribosome;nucleolus;small-subunit processome |
| PF3D7_1438100 | SEC62 | translocation protein SEC62, putative | 43.94 | no | 3 | 70.95 | 18.34 | 98.25 | 130.26 | 23.87 | 18.16 | 41.11 | S | 139.91 | 22.16 | M | protein binding | posttranslational protein targeting to membrane, translocation;protein targeting to ER;protein transport by the Sec complex | Sec61 translocon complex;endoplasmic reticulum;endoplasmic reticulum membrane;membrane;nucleus |
| PF3D7_1145100 | SEC21 | coatomer subunit gamma, putative | 124.37 | no | 0 | 37.73 | 10.65 | 25.41 | 5.15 | 4.12 | 19.59 | 29.51 | R | 65.24 | 24.45 | M | protein binding | endoplasmic reticulum to Golgi vesicle-mediated transport;intra-Golgi vesicle-mediated transport;protein secretion;retrograde vesicle-mediated transport, Golgi to endoplasmic reticulum;vesicle-mediated transport | COPI vesicle coat;Golgi apparatus;Golgi membrane;cytosol;endoplasmic reticulum;endoplasmic reticulum-Golgi intermediate compartment;membrane;nucleus |
| PF3D7_0616600 | N/A | HROB domain-containing protein | 73.90 | no | 0 | 6.02 | 1.53 | 11.94 | 3.40 | 2.47 | 52.95 | 627.19 | O | 24.05 | 249.27 | F | DNA binding | recombinational repair, DNA repair | nucleus |
| PF3D7_1351700 | IMC1f | inner membrane complex protein 1f | 151.52 | no | 0 | 5.72 | 1.83 | 19.12 | 9.95 | 0.42 | 1.55 | 338.62 | O | 1.02 | 0.98 | O | N/A | structural complex | basal part of cell;inner membrane pellicle complex |
| PF3D7_1461600 | SF3B2 | splicing factor 3B subunit 2, putative | 79.34 | no | 0 | 152.59 | 71.38 | 41.67 | 6.14 | 24.04 | 20.53 | 106.24 | R | 99.31 | 38.84 | M | RNA binding | mRNA splicing, via spliceosome | U2 snRNP;U2-type spliceosomal complex;catalytic step 2 spliceosome;nucleus;precatalytic spliceosome;spliceosomal complex |
| PF3D7_0803800 | N/A | proteasome subunit beta type-4 | 30.87 | no | 0 | 211.58 | 102.81 | 184.94 | 66.31 | 123.91 | 238.56 | 243.48 | O | 95.98 | 189.70 | F | endopeptidase activity | proteasomal protein catabolic process;proteasomal ubiquitin-independent protein catabolic process;proteasome-mediated ubiquitin-dependent protein catabolic process | cytoplasm;nucleus;proteasome core complex |
| PF3D7_1003500 | N/A | 40S ribosomal protein S20e, putative | 13.51 | no | 0 | 117.84 | 105.43 | 49.56 | 6.35 | 59.69 | 75.21 | 83.53 | R | 82.17 | 50.56 | M | structural constituent of ribosome | translation | cytosolic small ribosomal subunit;nucleus;small ribosomal subunit |
| PF3D7_1031500 | DDX3X | ATP-dependent DNA helicase DDX3X | 107.80 | no | 0 | 71.32 | 12.20 | 39.01 | 8.12 | 8.93 | 59.66 | 156.10 | O | 87.61 | 150.99 | F | ATP hydrolysis activity;DNA helicase activity | mRNA splicing, via spliceosome | cytoplasm;nucleus |
| PF3D7_0921800 | RPF1 | ribosome production factor 1, putative | 39.17 | no | 0 | 64.18 | 14.20 | 5.10 | 0.41 | 6.26 | 64.51 | 93.79 | O | 4.74 | 40.86 | F | protein binding | maturation of 5.8S rRNA;maturation of LSU-rRNA;rRNA processing;ribosomal large subunit assembly | nucleolus;preribosome, large subunit precursor |
| PF3D7_1108600 | ERC | endoplasmic reticulum-resident calcium binding protein | 39.37 | yes | 0 | 126.40 | 61.09 | 462.04 | 147.94 | 42.65 | 287.99 | 700.63 | O | 37.99 | 488.08 | F | calcium ion binding;protein binding | exit from host cell;intracellular protein transport;proteolysis | endoplasmic reticulum;food vacuole |
| PF3D7_1354400 | N/A | V-type proton ATPase 21 kDa proteolipid subunit, putative | 19.12 | no | 4 | 77.87 | 64.26 | 51.65 | 24.37 | 35.24 | 52.65 | 30.09 | R | 35.66 | 20.14 | M | proton-transporting ATPase activity, rotational mechanism | | integral component of membrane;vacuolar proton-transporting V-type ATPase complex |
| PF3D7_0729500 | MT-A70 | N6-adenosine-methyltransferase MT-A70 | 89.63 | no | 0 | 31.54 | 17.97 | 11.09 | 1.30 | 1.71 | 8.81 | 19.92 | R | 61.92 | 25.32 | M | mRNA (2'-O-methyladenosine-N6-)-methyltransferase activity;mRNA (N6-adenosine)-methyltransferase activity;methyltransferase activity;protein binding | RNA methylation;RNA modification;mRNA methylation | RNA N6-methyladenosine methyltransferase complex;cytoplasm;nuclear speck;nucleus |
| PF3D7_1126200 | RPS18 | 40S ribosomal protein S18, putative | 17.89 | no | 0 | 973.78 | 448.53 | 220.40 | 70.31 | 572.01 | 287.04 | 371.84 | R | 71.95 | 135.15 | F | RNA binding;structural constituent of ribosome | translation | cytosol;nucleus;small ribosomal subunit |
| PF3D7_1460300 | RPL29 | 60S ribosomal protein L29, putative | 8.01 | no | 0 | 112.73 | 112.98 | 32.31 | 6.01 | 60.47 | 125.83 | 321.90 | O | 129.56 | 177.44 | F | structural constituent of ribosome | cytoplasmic translation | cytosolic large ribosomal subunit;nucleus |

| | | | | | | | | | | | | | | | | | | | |
|---|---|---|---|---|---|---|---|---|---|---|---|---|---|---|---|---|---|---|---|
| PF3D7_1136800 | N/A | DnaJ protein, putative | 44.53 | no | 1 | 14.88 | 7.70 | 6.84 | 1.45 | 3.48 | 10.24 | 21.86 | O | 18.33 | 6.72 | M | Hsp70 protein binding | cellular response to misfolded protein;chaperone cofactor-dependent protein refolding;ubiquitin-dependent ERAD pathway | endoplasmic reticulum membrane |
| PF3D7_1451200 | N/A | conserved Plasmodium protein, unknown function | 176.23 | no | 0 | 288.80 | 101.55 | 56.25 | 4.96 | 2.47 | 12.68 | 239.53 | R | 46.39 | 66.70 | F | protein binding | N/A | nucleus |
| PF3D7_1250800 | N/A | DNA repair protein rhp16, putative | 192.17 | no | 0 | 58.81 | 38.20 | 25.28 | 5.26 | 6.29 | 7.57 | 88.79 | O | 188.32 | 29.03 | M | ATP binding;helicase activity | nucleotide-excision repair | nucleotide-excision repair complex |
| PF3D7_0912000 | N/A | conserved Plasmodium protein, unknown function | 188.68 | no | 3 | 18.55 | 5.23 | 7.53 | 1.27 | 1.14 | 13.14 | 9.28 | R | 28.68 | 12.36 | M | N/A | N/A | N/A |
| PF3D7_0906100 | VPS46 | vacuolar protein sorting-associated protein 46, putative | 21.85 | no | 0 | 7.57 | 9.48 | 115.18 | 65.30 | 252.60 | 151.01 | 259.27 | O | 15.10 | 385.61 | F | protein binding | endosome transport via multivesicular body sorting pathway;late endosome to vacuole transport;protein transport | ESCRT III complex;multivesicular body |
| PF3D7_0824400 | NT2 | nucleoside transporter 2 | 67.64 | no | 10 | 10.33 | 52.67 | 29.07 | 5.54 | 21.53 | 37.20 | 39.04 | T | 11.57 | 7.47 | M | nucleoside transmembrane transporter activity | nucleoside transport | endoplasmic reticulum;membrane |
| PF3D7_0416300 | MCM9 | DNA helicase MCM9, putative | 171.31 | no | 0 | 9.40 | 5.04 | 16.46 | 3.63 | 1.93 | 12.57 | 624.75 | O | 22.80 | 79.50 | F | ATP-dependent activity, acting on DNA;DNA replication origin binding;single-stranded DNA binding | DNA replication initiation;double-strand break repair via homologous recombination | MCM complex;nucleus |
| PF3D7_1323100 | N/A | 60S ribosomal protein L6, putative | 21.60 | no | 0 | 213.70 | 245.30 | 104.30 | 17.28 | 49.63 | 63.79 | 246.40 | O | 49.94 | 101.63 | F | RNA binding;structural constituent of ribosome | cytoplasmic translation;translation | cytosolic large ribosomal subunit;large ribosomal subunit;nucleus |
| PF3D7_0805500 | N/A | conserved Plasmodium protein, unknown function | 128.69 | no | 0 | 18.66 | 6.69 | 5.40 | 1.84 | 1.01 | 7.51 | 4.97 | R | 21.67 | 10.02 | M | N/A | N/A | N/A |
| PF3D7_0914800 | N/A | GINS complex subunit Psf3, putative | 24.84 | no | 0 | 12.20 | 1.17 | 10.89 | 2.80 | 10.18 | 141.91 | 47.86 | GV | 35.12 | 128.51 | F | N/A | DNA replication | nucleus |
| PF3D7_0312400 | GSK3beta | glycogen synthase kinase-3 beta | 51.62 | no | 0 | 9.52 | 15.50 | 79.38 | 27.57 | 38.99 | 128.17 | 257.17 | O | 246.11 | 303.98 | F | protein serine/threonine kinase activity | cell maturation;entry into host;response to xenobiotic stimulus;signal transduction | Maurer's cleft;cytoplasm;nuclear periphery |
| PF3D7_0412000 | N/A | LITAF-like zinc finger protein, putative | 92.20 | no | 2 | 3.81 | 6.23 | 32.81 | 8.14 | 6.41 | 12.44 | 19.90 | T | 134.21 | 27.38 | M | protein binding | N/A | integral component of membrane |
| PF3D7_1474500 | SF3A1 | splicing factor 3A subunit 1, putative | 83.75 | no | 0 | 33.66 | 31.90 | 27.39 | 2.85 | 1.54 | 16.44 | 145.85 | O | 93.11 | 19.31 | M | RNA binding | mRNA splicing, via spliceosome | U2 snRNP;U2-type prespliceosome;catalytic step 2 spliceosome;nucleus;spliceosomal complex |
| PF3D7_0904500 | N/A | prefoldin subunit 4, putative | 15.33 | no | 0 | 9.39 | 5.53 | 6.01 | 1.50 | 18.88 | 86.11 | 34.47 | GV | 15.25 | 17.75 | 0 | unfolded protein binding | protein folding | cytoplasm;prefoldin complex;nucleus |
| PF3D7_0415900 | N/A | 60S ribosomal protein L15, putative | 24.13 | no | 0 | 1135.97 | 793.88 | 245.39 | 54.51 | 319.48 | 238.70 | 588.77 | R | 125.52 | 273.77 | F | RNA binding;structural constituent of ribosome | cytoplasmic translation;translation | cytosolic large ribosomal subunit;nucleus;ribosome |
| PF3D7_1304700 | CYC4 | cyclin | 32.61 | no | 0 | 6.77 | 2.13 | 6.02 | 1.20 | 4.81 | 73.07 | 61.74 | GV | 14.98 | 57.16 | F | cyclin-dependent protein serine/threonine kinase regulator activity | regulation of cell cycle;regulation of cyclin-dependent protein serine/threonine kinase activity;regulation of transcription by RNA polymerase II | nucleus |
| PF3D7_1451500 | N/A | pre-mRNA-splicing factor CWF18, putative | 17.26 | no | 0 | 11.63 | 3.85 | 2.37 | 0.12 | 8.48 | 30.41 | 0.00 | GV | 7.47 | 1.73 | M | | | U2-type spliceosomal complex;nucleus;post-mRNA release spliceosomal complex |
| PF3D7_0719700 | RPS10 | 40S ribosomal protein S10, putative | 16.48 | no | 0 | 184.47 | 244.87 | 111.26 | 25.18 | 97.83 | 121.65 | 246.83 | O | 42.03 | 159.70 | F | RNA binding;structural constituent of ribosome | ribosomal small subunit assembly;translation | cytosolic small ribosomal subunit;nucleus |
| PF3D7_1416900 | N/A | prefoldin subunit 2, putative | 16.65 | no | 0 | 21.07 | 38.82 | 60.79 | 6.54 | 585.05 | 130.72 | 86.15 | G II | 123.21 | 42.28 | M | protein folding chaperone | protein folding | cytoplasm;nucleus |
| PF3D7_1246800 | SRPRB | signal recognition particle receptor subunit beta, putative | 30.62 | no | 1 | 8.66 | 11.75 | 28.76 | 5.54 | 16.93 | 68.25 | 67.99 | O | 38.34 | 22.95 | M | RNA binding | protein targeting to ER | signal recognition particle receptor complex |
| PF3D7_0512000 | PFD6 | prefoldin subunit 6 | 13.67 | no | 0 | 58.22 | 76.45 | 39.56 | 13.50 | 269.32 | 456.37 | 375.15 | O | 238.44 | 197.12 | M | chaperone binding;protein binding | chaperone-mediated protein complex assembly;protein folding | cytoplasm;prefoldin complex;symbiont-containing vacuole membrane |
| PF3D7_1302500 | N/A | AAA domain-containing protein, putative | 325.17 | no | 0 | 16.39 | 4.55 | 9.87 | 1.26 | 2.03 | 19.46 | 16.37 | O | 78.68 | 16.89 | M | RNA binding | N/A | nucleus |
| PF3D7_0935000 | SNRPB2 | U2 small nuclear ribonucleoprotein B'', putative | 22.29 | no | 0 | 118.96 | 35.61 | 40.86 | 7.70 | 28.30 | 100.78 | 58.85 | R | 27.78 | 30.07 | 0 | U1 snRNA binding;mRNA binding | mRNA splicing, via spliceosome | U1 snRNP;nucleus |
| PF3D7_1134100 | PDI-11 | protein disulfide-isomerase, putative | 49.19 | yes | 0 | 15.72 | 7.67 | 28.63 | 11.47 | 4.67 | 17.04 | 13.32 | T | 4.46 | 6.68 | 0 | RNA binding;electron transfer activity;protein disulfide isomerase activity | oxidation-reduction process;protein folding | endoplasmic reticulum lumen;nucleus |
| PF3D7_1213700 | RPB8 | DNA-directed RNA polymerases I, II, and III subunit RPABC3, putative | 16.90 | no | 0 | 36.18 | 14.63 | 7.88 | 0.44 | 11.18 | 14.97 | 60.92 | O | 6.89 | 10.34 | F | DNA-directed 5'-3' RNA polymerase activity | DNA-templated transcription, initiation | RNA polymerase I complex;RNA polymerase II, core complex;RNA polymerase III complex;nucleus |
| PF3D7_1313800 | N/A | conserved Plasmodium membrane protein | 408.84 | no | 8 | 71.00 | 21.78 | 9.08 | 4.05 | 2.68 | 9.58 | 10.43 | R | 20.17 | 6.11 | M | N/A | N/A | integral component of membrane |
| PF3D7_0621500 | RPP1 | ribonuclease P/MRP protein subunit RPP1, putative | 40.29 | no | 0 | 4.95 | 8.79 | 7.05 | 2.47 | 23.78 | 71.71 | 16.16 | GV | 18.24 | 8.59 | M | RNA binding;ribonuclease activity | RNA phosphodiester bond hydrolysis, endonucleolytic;tRNA processing | nucleolar ribonuclease P complex;ribonuclease MRP complex;ribonuclease P complex |
| PF3D7_1341900 | N/A | V-type proton ATPase subunit D, putative | 27.80 | no | 0 | 83.26 | 57.44 | 46.24 | 16.30 | 129.53 | 309.41 | 84.38 | GV | 99.47 | 175.84 | F | proton-transporting ATPase activity, rotational mechanism | proton transmembrane transport;vacuolar acidification | food vacuole;proton-transporting V-type ATPase complex;vacuolar membrane;vacuolar proton-transporting V-type ATPase, V0 domain;vacuolar proton-transporting V-type ATPase, V1 domain |
| PF3D7_1437000 | NAT8 | N-acetyltransferase, GNAT family | 64.19 | no | 2 | 125.63 | 22.58 | 34.74 | 7.46 | 3.67 | 21.82 | 55.54 | R | 48.69 | 44.44 | 0 | N-acetyltransferase activity | schizogony | endoplasmic reticulum;nucleus |
| PF3D7_0627900 | POP4 | ribonuclease P protein subunit p29, putative | 34.32 | no | 0 | 24.49 | 10.50 | 7.53 | 1.45 | 2.86 | 8.41 | 4.17 | R | 17.75 | 7.48 | M | ribonuclease P RNA binding;ribonuclease activity | rRNA processing | ribonuclease MRP complex;ribonuclease P complex |
| PF3D7_1112100 | N/A | protein kinase, putative | 296.62 | no | 3 | 9.79 | 3.79 | 12.81 | 6.30 | 1.98 | 18.47 | 170.18 | O | 67.12 | 67.44 | 0 | protein kinase activity | meiotic sister chromatid cohesion, centromeric;mitotic spindle assembly checkpoint signaling | condensed nuclear chromosome kinetochore;nucleus |
| PF3D7_0415000 | GET3 | ATPase GET3 | 43.32 | no | 0 | 86.90 | 10.81 | 23.00 | 1.98 | 9.07 | 31.48 | 4.77 | R | 12.91 | 55.18 | F | ATP hydrolysis activity;ATPase-coupled arsenite transmembrane transporter activity;protein binding | ion transport;posttranslational protein targeting to endoplasmic reticulum membrane | GET complex;cytosol;endoplasmic reticulum;membrane |
| PF3D7_0206000 | RAD2 | DNA repair protein RAD2, putative | 178.73 | no | 0 | 22.89 | 10.97 | 8.28 | 3.50 | 1.29 | 11.82 | 321.85 | O | 71.69 | 15.63 | M | 5'-flap endonuclease activity;damaged DNA binding;endonuclease activity;flap endonuclease activity;single-stranded DNA endodeoxyribonuclease activity | DNA repair;nucleotide-excision repair, DNA incision, 3'-to lesion | chromosome, telomeric region;nucleotide-excision repair factor 3 complex |
| PF3D7_1034900 | MRScyt | methionine--tRNA ligase | 104.17 | no | 0 | 97.54 | 9.09 | 96.73 | 18.28 | 1.95 | 9.22 | 21.02 | R | 18.29 | 65.71 | F | ATP binding;methionine-tRNA ligase activity | methionyl-tRNA aminoacylation | cytoplasm;nucleus |
| PF3D7_0618300 | N/A | 60S ribosomal protein L27a, putative | 16.72 | no | 0 | 465.64 | 215.55 | 132.00 | 96.46 | 72.34 | 99.81 | 161.29 | R | 99.37 | 210.25 | F | structural constituent of ribosome | translation | cytosolic large ribosomal subunit;large ribosomal subunit;nucleus |
| PF3D7_1307100 | UTP6 | U3 small nucleolar RNA-associated protein 6, putative | 120.72 | no | 1 | 46.66 | 24.11 | 7.20 | 1.15 | 0.96 | 4.78 | 0.85 | R | 12.92 | 14.03 | 0 | snoRNA binding | maturation of SSU-rRNA from tricistronic rRNA transcript (SSU-rRNA, 5.8S rRNA, LSU-rRNA) | Pwp2p-containing subcomplex of 90S preribosome;small-subunit processome;nucleus |
| PF3D7_1468200 | TLS1 | telomere length and silencing protein 1, putative | 27.10 | no | 0 | 12.85 | 9.45 | 14.74 | 1.60 | 6.99 | 40.19 | 19.61 | GV | 191.31 | 27.19 | M | DNA binding | regulation of mRNA splicing, via spliceosome | nucleus;spliceosomal complex |
| PF3D7_1317600 | N/A | conserved Plasmodium protein, unknown function | 73.87 | no | 0 | 4.09 | 3.26 | 9.19 | 1.28 | 1.10 | 5.73 | 9.07 | T | 2.04 | 13.59 | F | N/A | N/A | nucleus |
| PF3D7_1433300 | RBAP46 | histone-binding protein RBBP7, putative | 50.17 | no | 0 | 8.36 | 11.57 | 22.55 | 12.72 | 9.02 | 22.30 | 28.17 | O | 26.36 | 26.76 | 0 | histone binding;unfolded protein binding | chromatin assembly;chromatin assembly or disassembly;chromatin remodeling;nucleosome assembly | nucleus |
| PF3D7_1134700 | RPA2 | DNA-directed RNA polymerase I subunit RPA2, putative | 175.53 | no | 0 | 100.82 | 61.29 | 9.83 | 0.49 | 4.59 | 12.21 | 17.07 | R | 27.44 | 18.67 | M | DNA binding | regulation of transcription | nucleus |
| PF3D7_0505600 | N/A | BCNT domain-containing protein, putative | 40.20 | no | 0 | 17.18 | 5.44 | 9.63 | 1.40 | 6.66 | 275.39 | 54.41 | GV | 45.00 | 45.99 | 0 | N/A | N/A | nucleus |
| PF3D7_0813200 | N/A | CS domain protein, putative | 37.12 | no | 0 | 23.07 | 13.64 | 21.72 | 8.80 | 6.58 | 39.57 | 107.79 | O | 14.26 | 69.48 | F | unfolded protein binding | developmental process | cytoplasm;osmiophilic body |
| PF3D7_0522600 | N/A | magnesium transporter NIPA, putative | 55.56 | no | 8 | 5.32 | 4.55 | 16.93 | 20.28 | 3.26 | 9.35 | 36.64 | O | 51.06 | 11.14 | 0 | N/A | N/A | inner membrane pellicle complex |
| PF3D7_1031600 | GEXP15 | protein GEXP15 | 106.69 | no | 0 | 42.55 | 35.51 | 26.57 | 7.09 | 9.95 | 27.51 | 34.96 | R | 59.41 | 58.76 | 0 | protein binding;ribosome binding;protein phosphatase regulator activity | N/A | U5 snRNP;nucleus;cytoplasm |
| PF3D7_0818700 | N/A | DNA helicase, putative | 143.66 | no | 0 | 154.41 | 20.67 | 46.75 | 11.91 | 9.22 | 29.57 | 96.04 | R | 359.16 | 21.96 | M | ATP-dependent activity, acting on DNA | regulation of replication | nucleus |
| PF3D7_1144500 | AOS1 | SUMO-activating enzyme subunit 1 | 39.69 | no | 0 | 18.28 | 7.10 | 7.01 | 1.56 | 4.33 | 27.50 | 34.27 | O | 19.71 | 7.76 | M | SUMO activating enzyme activity | protein modification by small protein conjugation | N/A |
| PF3D7_0209100 | PATPL1 | patatin-like phospholipase 1 | 78.30 | no | 0 | 54.45 | 10.34 | 28.25 | 10.08 | 3.93 | 11.09 | 81.02 | O | 52.24 | 32.18 | M | phospholipase A2 activity | lipid metabolic process | cytoplasm |
| PF3D7_1004400 | N/A | RNA-binding protein, putative | 106.30 | no | 0 | 59.09 | 175.50 | 55.95 | 9.92 | 8.40 | 14.17 | 179.28 | O | 232.06 | 80.97 | M | mRNA binding | N/A | cytoplasm;nuclues |
| PF3D7_1105600 | PTEX88 | translocon component PTEX88 | 90.79 | yes | 0 | 101.42 | 42.70 | 19.06 | 1.26 | 2.09 | 4.26 | 28.80 | R | 17.91 | 5.24 | M | RNA binding;protein binding | translocation of peptides or proteins into host cell cytoplasm | merozoite dense granule;symbiont-containing vacuole membrane |
| PF3D7_1247500 | N/A | serine/threonine protein kinase, putative | 105.54 | no | 0 | 3.56 | 1.50 | 8.26 | 4.41 | 2.73 | 40.34 | 67.03 | O | 36.78 | 125.03 | F | protein phosphorylation;protein serine threonine kinase activity;ATP binding | protein phosphorylation | cytoplasm;nucleus |
| PF3D7_1202200 | PIC | mitochondrial phosphate carrier protein | 36.80 | no | 1 | 8.53 | 1.01 | 45.89 | 15.61 | 17.57 | 41.63 | 53.70 | O | 160.87 | 43.35 | M | inorganic phosphate transmembrane transporter activity | phosphate ion transmembrane transport | integral component of mitochondrial inner membrane |
| PF3D7_0909800 | SNRPD3 | small nuclear ribonucleoprotein Sm D3, putative | 12.98 | no | 0 | 60.21 | 13.00 | 10.68 | 0.92 | 26.75 | 51.99 | 45.11 | R | 50.35 | 21.34 | M | RNA binding | spliceosomal snRNP complex activity;mRNA splicing, via spliceosome | spliceosomal complex;nucleus |
| PF3D7_0525100 | ACS10 | acyl-CoA synthetase | 76.85 | no | 0 | 124.01 | 51.70 | 119.43 | 10.82 | 5.03 | 28.93 | 459.74 | O | 37.75 | 167.13 | F | long-chain fatty acid-CoA ligase activity | lipid metabolic process | membrane;endoplasmic reticuluum |
| PF3D7_1462100 | N/A | conserved Plasmodium protein, unknown function | 161.20 | no | 0 | 3.04 | 1.13 | 9.95 | 2.31 | 1.05 | 6.55 | 3.37 | T | 8.73 | 4.81 | M | N/A | N/A | N/A |
| PF3D7_1119300 | U2AF1 | splicing factor U2AF small subunit, putative | 35.31 | no | 0 | 276.90 | 112.99 | 103.24 | 34.40 | 8.39 | 17.26 | 357.34 | O | 467.43 | 101.63 | M | RNA binding;pre-mRNA 3'-splice site binding | mRNA splicing, via spliceosome | U2 snRNP;U2AF complex;nucleus;spliceosomal complex |
| PF3D7_1347200 | NT1 | nucleoside transporter 1 | 47.63 | no | 9 | 575.20 | 422.75 | 186.24 | 10.94 | 31.86 | 74.12 | 99.76 | R | 37.64 | 44.38 | F | nucleoside transmembrane transporter activity;purine nucleoside transmembrane transporter activity | adenosine transport;purine nucleobase transport;purine nucleoside transmembrane transport;response to xenobiotic stimulus | Maurer's cleft;integral component of membrane;integral component of plasma membrane;nucleus;plasma membrane |
| PF3D7_0804900 | N/A | GTPase-activating protein, putative | 56.50 | no | 0 | 50.73 | 17.52 | 28.25 | 7.30 | 2.83 | 21.42 | 265.52 | O | 39.93 | 23.51 | M | enzyme activity | COPI coating of Golgi vesicle | nucleus;cytoplasm |
| PF3D7_0822900 | PIC2 | PhIL1-interacting candidate PIC2 | 138.24 | no | 0 | 8.87 | 0.33 | 18.51 | 14.58 | 0.60 | 3.53 | 73.96 | O | 0.19 | 2.44 | | protein binding | structural complex | inner membrane pellicle complex |
| PF3D7_0916700 | HoMu | RNA-binding protein musashi, putative | 45.14 | no | 0 | 231.78 | 31.55 | 17.12 | 2.64 | 7.94 | 34.75 | 47.06 | R | 5.05 | 20.34 | F | RNA binding;mRNA binding | N/A | nucleoplasm;nucleus |
| PF3D7_0505800 | SUMO | small ubiquitin-related modifier | 11.06 | no | 0 | 265.07 | 145.70 | 157.66 | 46.42 | 313.85 | 164.82 | 64.16 | G II | 160.33 | 37.68 | M | protein binding;protein tag;ubiquitin-like protein ligase binding | modification-dependent protein catabolic process;protein sumoylation;ubiquitin-dependent protein catabolic process | nucleus |
| PF3D7_0523400 | N/A | DnaJ protein, putative | 61.90 | no | 2 | 17.11 | 22.15 | 36.86 | 7.55 | 4.19 | 23.48 | 332.79 | O | 125.88 | 70.85 | M | RNA binding;unfolded protein binding | chaperone cofactor-dependent protein refolding;protein refolding | cytoplasm;food vacuole;nucleus |
| PF3D7_1315300 | N/A | conserved protein, unknown function | 56.34 | no | 0 | 2.13 | 0.67 | 11.78 | 6.66 | 2.75 | 206.52 | 25.79 | GV | 7.24 | 176.71 | F | N/A | N/A | nucleus |
| PF3D7_0730200 | N/A | AP-4 complex subunit beta, putative | 100.38 | no | 0 | 7.44 | 3.19 | 8.18 | 0.98 | 1.24 | 12.79 | 6.43 | GV | 18.49 | 6.59 | M | protein binding | vesicle-mediated transport | clathrin coat of trans-Golgi network vesicle;cytoplasm |
| PF3D7_0614100 | N/A | filamin domain-containing protein, putative | 112.57 | no | 0 | 7.05 | 1.64 | 6.93 | 5.83 | 3.63 | 36.29 | 42.15 | O | 27.91 | 66.77 | F | N/A | N/A | N/A |
| PF3D7_0912800 | GCD10 | tRNA (adenine(58)-N(1))-methyltransferase non-catalytic subunit TRM6, putative | 72.84 | no | 0 | 29.11 | 18.96 | 9.14 | 2.07 | 1.95 | 14.50 | 23.75 | R | 5.44 | 9.99 | F | methyltransferase activity | N/A | nucleus;tRNA (m1A) methyltransferase complex |
| PF3D7_0509400 | RNAPI | RNA polymerase I | 340.67 | no | 0 | 106.77 | 76.52 | 12.86 | 1.42 | 2.56 | 11.12 | 26.32 | R | 5.62 | 14.88 | F | RNA binding | regulation of transcription | RNA polymerase I complex;nucleolus |
| PF3D7_0522200 | TAF10 | transcription initiation factor TFIID subunit 10, putative | 13.63 | no | 0 | 34.26 | 4.30 | 6.39 | 1.51 | 26.85 | 70.37 | 13.85 | GV | 17.25 | 11.92 | M | protein binding | transcription by RNA polymerase II | transcription regulator complex |
| PF3D7_0213100 | SIS1 | protein SIS1 | 37.36 | no | 0 | 48.31 | 104.03 | 46.29 | 5.24 | 16.77 | 106.80 | 353.92 | O | 117.59 | 217.98 | F | chaperone binding;unfolded protein binding | chaperone cofactor-dependent protein refolding;protein folding;response to heat;response to unfolded protein | Maurer's cleft;cytosol;nucleus |
| PF3D7_0829200 | PHB1 | prohibitin 1 | 30.56 | no | 0 | 37.63 | 44.93 | 20.63 | 1.59 | 45.89 | 193.59 | 158.23 | GV | 41.65 | 109.24 | F | protein binding | mitochondrion organization;regulation of cell population proliferation;response to xenobiotic stimulus | mitochondrion |
| PF3D7_1364800 | RPB5 | DNA-directed RNA polymerases I, II, and III subunit RPABC1, putative | 24.15 | no | 0 | 185.49 | 68.80 | 33.03 | 1.20 | 81.54 | 74.57 | 61.96 | R | 53.98 | 90.20 | F | DNA binding;DNA-directed 5'-3' RNA polymerase activity | transcription by RNA polymerase II | RNA polymerase I complex;RNA polymerase II, core complex;RNA polymerase III complex;nucleus |
| PF3D7_1033700 | BDP1 | bromodomain protein 1 | 55.94 | no | 0 | 143.33 | 49.86 | 47.36 | 9.11 | 3.16 | 50.71 | 356.93 | O | 62.30 | 100.83 | F | chromatin binding;histone binding;protein binding | entry into host;regulation of transcription, DNA-templated | chromosome, telomeric region;nucleus |
| PF3D7_0113000 | GARP | glutamic acid-rich protein GARP | 79.79 | yes | 1 | 712.21 | 1602.48 | 64.11 | 4.19 | 15.18 | 4.65 | 129.16 | T | 82.69 | 5.01 | M | host cell surface receptor binding | modulation by symbiont of host immune response;programmed cell death | host cell periphery;host cell surface |
| PF3D7_1033400 | HAD1 | haloacid dehalogenase-like hydrolase | 32.84 | no | 0 | 286.80 | 208.06 | 68.73 | 5.98 | 7.28 | 6.44 | 58.50 | R | 48.91 | 6.06 | M | sugar-phosphatase activity | dephosphorylation;negative regulation of isopentenyl diphosphate biosynthetic process, methylerythritol 4-phosphate pathway | cytoplasm |
| PF3D7_0510200 | CYP87 | peptidyl-prolyl cis-trans isomerase | 87.04 | no | 0 | 17.64 | 11.19 | 12.27 | 5.09 | 6.67 | 16.07 | 20.97 | O | 42.43 | 19.17 | M | cyclosporin A binding;peptidyl-prolyl cis-trans isomerase activity | protein folding;protein peptidyl-prolyl isomerization | |
| PF3D7_1358700 | YOP1L | YOP1-like protein, putative | 18.42 | no | 3 | 60.59 | 8.18 | 14.36 | 4.68 | 15.45 | 75.09 | 48.99 | GV | 44.72 | 29.75 | M | protein binding | endoplasmic reticulum organization;endoplasmic reticulum tubular network organization | integral component of membrane |
| PF3D7_1250600 | N/A | translation initiation factor eIF-2B subunit beta, putative | 78.75 | no | 0 | 42.49 | 7.64 | 13.97 | 2.88 | 4.61 | 20.47 | 100.78 | O | 65.77 | 23.09 | M | GTP binding;translation initiation factor activity | regulation of translational initiation;translational initiation | eukaryotic translation initiation factor 2B complex |
| PF3D7_1100800 | MC-2TM | Pfmc-2TM Maurer's cleft two transmembrane protein | 27.18 | yes | 2 | 1.91 | 10.87 | 0.40 | 0.15 | 0.00 | 1.30 | 55.96 | O | 0.63 | 0.00 | | | translocation of peptides or proteins into host | Maurer's cleft;integral component of membrane;membrane |
| PF3D7_1342400 | CK2beta2 | casein kinase II beta chain | 45.28 | no | 0 | 189.41 | 281.64 | 188.38 | 37.43 | 16.92 | 20.24 | 142.90 | T | 80.81 | 68.01 | M | protein binding;protein kinase regulator activity | protein phosphorylation | cytoplasm;nucleus;protein kinase CK2 complex |
| PF3D7_0720700 | PX1 | phosphoinositide-binding protein PX1 | 258.42 | no | 4 | 207.09 | 24.78 | 39.92 | 33.67 | 15.73 | 37.34 | 545.76 | O | 169.86 | 57.66 | M | phosphatidylinositol binding;phosphatidylinositol-3-phosphate binding | intracellular protein transport;response to xenobiotic stimulus;retrograde transport, endosome to plasma membrane | early endosome;food vacuole;membrane;vesicle |
| PF3D7_1031300 | N/A | SAE2 domain-containing protein, putative | 109.28 | no | 0 | 5.52 | 4.83 | 6.30 | 0.81 | 3.62 | 34.43 | 104.05 | O | 26.17 | 20.23 | M | N/A | N/A | nucleus |
| PF3D7_0819900 | LSM3 | U6 snRNA-associated Sm-like protein LSm3, putative | 10.70 | no | 0 | 23.44 | 7.05 | 4.52 | 1.25 | 14.44 | 47.45 | 71.71 | O | 163.54 | 86.32 | M | RNA binding | P-body assembly;RNA splicing;RNA splicing, via transesterification reactions;mRNA splicing, via spliceosome | Lsm1-7-Pat1 complex;P-body;U4/U6 x U5 tri-snRNP complex;U6 snRNP;catalytic step 2 spliceosome;nucleus;precatalytic spliceosome;sno(s)RNA-containing ribonucleoprotein complex |
| PF3D7_1037600 | XPB | TFIIH basal transcription factor complex helicase XPB subunit, putative | 102.87 | no | 0 | 53.16 | 19.24 | 21.04 | 4.81 | 1.70 | 41.13 | 138.14 | O | 77.09 | 257.49 | F | 3'-5' DNA helicase activity;ATP hydrolysis activity;DNA binding;DNA helicase activity;damaged DNA binding;helicase activity;transcription factor binding | nucleotide-excision repair, DNA duplex unwinding;nucleotide-excision repair, DNA incision;response to UV;transcription by RNA polymerase II;transcription initiation from RNA polymerase II promoter | nucleotide-excision repair factor 3 complex;transcription factor TFIIH holo complex;transcription preinitiation complex |
| PF3D7_0109800 | cPheRS | phenylalanine--tRNA ligase alpha subunit | 67.72 | no | 0 | 74.74 | 25.31 | 22.84 | 2.89 | 4.88 | 12.76 | 144.71 | O | 18.96 | 23.23 | F | phenylalanine-tRNA ligase activity | phenylalanyl-tRNA aminoacylation;response to xenobiotic stimulus | cytoplasm;cytosol;nucleus;phenylalanine-tRNA ligase complex |
| PF3D7_1127600 | N/A | CRAL/TRIO domain-containing protein, putative | 34.97 | no | 0 | 8.61 | 11.08 | 32.68 | 7.39 | 35.70 | 350.97 | 266.48 | GV | 381.57 | 811.67 | F | N/A | N/A | nucleus |
| PF3D7_1414800 | SMB1 | small nuclear ribonucleoprotein-associated protein B | 22.33 | no | 0 | 56.39 | 34.30 | 36.42 | 4.03 | 12.02 | 14.63 | 23.66 | R | 12.73 | 20.39 | F | | RNA splicing;mRNA splicing, via spliceosome | U1 snRNP;U2 snRNP;U2-type prespliceosome;U4 snRNP;U4/U6 x U5 tri-snRNP complex;U5 snRNP;catalytic step 2 spliceosome;cytoplasm;nucleus;spliceosomal complex |
| PF3D7_1144900 | RAB6 | ras-related protein Rab-6 | 23.61 | no | 0 | 35.18 | 14.48 | 61.44 | 42.92 | 37.71 | 62.85 | 217.66 | O | 53.20 | 47.06 | M | GTPase activity | intra-Golgi vesicle-mediated transport;intracellular protein transport;retrograde transport, endosome to Golgi;retrograde vesicle-mediated transport, Golgi to endoplasmic reticulum | Golgi apparatus;cytosol;endomembrane system;trans-Golgi network |
| PF3D7_0823200 | N/A | RNA-binding protein, putative | 32.44 | no | 0 | 550.79 | 719.66 | 320.03 | 151.33 | 100.30 | 85.49 | 70.74 | T | 132.07 | 63.23 | M | RNA binding;mRNA binding | regulation of translation | nucleus;ribonucleoprotein complex |
| PF3D7_0526800 | N/A | conserved Plasmodium protein, unknown function | 43.78 | no | 0 | 236.44 | 60.21 | 48.77 | 4.83 | 20.03 | 45.70 | 38.42 | R | 164.30 | 68.03 | M | N/A | N/A | mediator complex;nucleus |

| | | | | | | | | | | | | | | | | | | | |
|---|---|---|---|---|---|---|---|---|---|---|---|---|---|---|---|---|---|---|---|
| PF3D7_1103600 | ALP2a | actin-like protein, putative | 57.97 | no | 0 | 20.27 | 12.52 | 7.99 | 0.86 | 3.01 | 12.02 | 19.64 | R | 15.88 | 3.26 | M | chromatin binding | ATP-dependent chromatin remodeling;histone H4 acetylation;regulation of transcription by RNA polymerase II | NuA4 histone acetyltransferase complex;SWI/SNF complex;nucleus |
| PF3D7_1321700 | SF1 | splicing factor 1 | 100.94 | no | 0 | 230.22 | 148.06 | 70.38 | 8.37 | 15.54 | 16.39 | 340.97 | O | 368.51 | 172.11 | M | mRNA binding | RNA splicing;RNA splicing, via transesterification reactions | nucleus |
| PF3D7_0719600 | N/A | 60S ribosomal protein L11a, putative | 20.23 | no | 0 | 251.71 | 413.57 | 149.95 | 46.70 | 58.45 | 50.53 | 137.89 | T | 46.08 | 48.84 | O | RNA binding;structural constituent of ribosome | ribosomal large subunit assembly;translation | cytosolic large ribosomal subunit;nucleus |
| PF3D7_0611800 | N/A | conserved Plasmodium protein, unknown function | 374.89 | no | 0 | 16.29 | 23.57 | 13.72 | 3.46 | 9.43 | 77.65 | 395.36 | O | 23.84 | 83.65 | F | N/A | N/A | nucleus |
| PF3D7_0917600 | PRP43 | pre-mRNA-splicing factor ATP-dependent RNA helicase PRP43, putative | 94.14 | no | 0 | 60.85 | 25.10 | 26.07 | 2.55 | 1.14 | 3.76 | 9.19 | R | 47.08 | 15.27 | M | RNA binding | N/A | nucleus |
| PF3D7_1147500 | FTB | protein farnesyltransferase subunit beta | 109.82 | no | 0 | 10.99 | 5.65 | 1.87 | 0.05 | 1.01 | 6.12 | 10.02 | R | 9.42 | 2.25 | M | protein farnesyltransferase activity;zinc ion binding | protein farnesylation;protein prenylation;response to xenobiotic stimulus | protein farnesyltransferase complex |
| PF3D7_0913600 | N/A | conserved Plasmodium protein, unknown function | 147.87 | no | 0 | 4.71 | 1.18 | 9.33 | 1.77 | 2.10 | 33.83 | 124.09 | P | 41.15 | 86.68 | F | N/A | N/A | N/A |
| PF3D7_1318400 | SMC2 | structural maintenance of chromosomes protein 2, putative | 143.06 | no | 0 | 11.66 | 4.23 | 22.11 | 7.44 | 3.26 | 22.35 | 195.74 | O | 961.78 | 136.31 | M | protein binding;DNA binding | mitotic chromosome condensation;mitotic sister chromatid segregation | chromosome, centromeric region;condensin complex;nucleus |
| PF3D7_0307700 | AKIT10 | apicomplexan kinetochore protein 10, putative | 216.26 | no | 0 | 13.01 | 44.81 | 224.44 | 169.17 | 9.51 | 37.12 | 98.73 | T | 245.33 | 56.40 | M | DNA binding | structural complex | kinetochore;nucleus |
| PF3D7_1450500 | N/A | conserved Plasmodium protein, unknown function | 199.74 | no | 1 | 9.24 | 1.34 | 26.12 | 12.37 | 5.36 | 21.82 | 47.40 | O | 49.33 | 48.88 | O | N/A | N/A | N/A |
| PF3D7_0308000 | N/A | DNA polymerase delta small subunit, putative | 57.72 | no | 0 | 4.63 | 8.80 | 14.98 | 1.67 | 6.89 | 22.70 | 21.25 | GV | 257.69 | 23.61 | M | DNA-directed DNA polymerase activity | DNA biosynthetic process;DNA strand elongation involved in DNA replication | DNA polymerase complex;delta DNA polymerase complex |
| PF3D7_1212900 | BDP2 | bromodomain protein 2, putative | 125.45 | no | 0 | 107.19 | 70.60 | 57.02 | 6.58 | 14.80 | 54.82 | 207.44 | O | 99.72 | 43.82 | M | protein binding | regulation of transcription | nucleus |
| PF3D7_1029600 | ADA | adenosine deaminase | 42.47 | no | 0 | 120.64 | 40.09 | 218.08 | 17.40 | 28.17 | 69.37 | 95.48 | T | 27.01 | 16.02 | M | adenosine deaminase activity | adenosine catabolic process;hypoxanthine salvage;inosine biosynthetic process;negative regulation of adenosine receptor signaling pathway;response to drug;response to xenobiotic stimulus | Maurer's cleft;cytosol;external side of plasma membrane |
| PF3D7_1116800 | HSP101 | heat shock protein 101 | 102.87 | yes | 1 | 600.08 | 336.05 | 66.33 | 16.10 | 4.75 | 13.30 | 623.97 | O | 25.67 | 20.47 | M | ATP hydrolysis activity;RNA binding;protein binding | cellular response to heat;protein unfolding;response to heat;response to unfolded protein;translocation of peptides or proteins into host cell cytoplasm | Maurer's cleft;PTEX complex;apical complex;cytoplasm;endoplasmic reticulum;extracellular vesicle;merozoite dense granule;nucleus;symbiont-containing vacuole;symbiont-containing vacuole membrane |
| PF3D7_1142500 | RPL28 | 60S ribosomal protein L28 | 14.41 | no | 0 | 138.01 | 232.06 | 77.25 | 12.08 | 123.19 | 50.59 | 158.39 | T | 39.35 | 66.94 | F | | | cytosolic large ribosomal subunit;nucleus |
| PF3D7_1121600 | EXP1 | exported protein 1 | 17.30 | yes | 1 | 1727.19 | 3544.12 | 1251.48 | 402.34 | 689.59 | 61.04 | 263.95 | T | 61.98 | 37.71 | M | channel activity;glutathione transferase activity;protein binding | glutathione metabolic process;response to drug;response to xenobiotic stimulus | extracellular vesicle;merozoite dense granule;nucleus;symbiont-containing vacuole membrane |
| PF3D7_1447800 | N/A | calponin homology domain-containing protein, putative | 238.05 | no | 0 | 9.76 | 6.72 | 4.88 | 0.97 | 6.01 | 31.78 | 26.49 | GV | 29.14 | 37.71 | F | protein binding | N/A | N/A |
| PF3D7_0524500 | N/A | conserved Plasmodium protein, unknown function | 14.51 | no | 0 | 21.83 | 4.15 | 6.98 | 1.61 | 8.23 | 26.06 | 10.28 | GV | 7.57 | 7.20 | O | N/A | N/A | nucleus |
| PF3D7_1306900 | SNRPA | U1 small nuclear ribonucleoprotein A, putative | 51.94 | no | 0 | 31.95 | 13.15 | 14.94 | 2.93 | 1.04 | 5.07 | 42.49 | O | 33.01 | 25.13 | M | U1 snRNA binding | mRNA splicing, via spliceosome | U1 snRNP;nucleus;spliceosomal complex |
| PF3D7_1118600 | MYST | histone acetyltransferase MYST | 71.13 | no | 0 | 64.05 | 16.93 | 10.55 | 2.51 | 2.39 | 16.11 | 353.78 | O | 83.51 | 54.94 | M | histone acetyltransferase activity;histone binding;protein binding;transcription coregulator activity | histone H4-K12 acetylation;histone H4-K16 acetylation;histone H4-K5 acetylation;histone H4-K8 acetylation;negative regulation of transcription, DNA-templated;positive regulation of transcription by RNA polymerase II;response to xenobiotic stimulus | cytoplasm;host cell periphery;nucleus;symbiont-containing vacuole |
| PF3D7_1013100 | UTP13 | U3 small nucleolar RNA-associated protein 13, putative | 144.60 | no | 0 | 74.13 | 20.58 | 8.35 | 0.80 | 3.72 | 39.75 | 30.24 | R | 15.33 | 80.61 | F | U3 snoRNA binding | endonucleolytic cleavage in 5'-ETS of tricistronic rRNA transcript (SSU-rRNA, 5.8S rRNA, LSU-rRNA);endonucleolytic cleavage to generate mature 5'-end of SSU-rRNA from (SSU-rRNA, 5.8S rRNA, LSU-rRNA) | 90S preribosome;nucleolus;small-subunit processome |
| PF3D7_1231600 | PRP2 | pre-mRNA-splicing factor ATP-dependent RNA helicase PRP2, putative | 137.20 | no | 0 | 54.29 | 14.50 | 22.31 | 2.19 | 6.33 | 24.26 | 46.16 | R | 179.16 | 29.09 | M | ATP-dependent activity, acting on RNA;RNA binding;RNA helicase activity | RNA splicing;mRNA splicing, via spliceosome;regulation of cell cycle | spliceosomal complex |
| PF3D7_0423000 | N/A | conserved Plasmodium protein, unknown function | 31.01 | no | 0 | 21.72 | 3.69 | 9.61 | 3.39 | 14.03 | 214.57 | 269.76 | O | 15.31 | 74.33 | F | N/A | N/A | N/A |
| PF3D7_1358800 | RPS15 | 40S ribosomal protein S15 | 17.25 | no | 0 | 189.89 | 207.18 | 86.98 | 5.99 | 74.09 | 75.84 | 152.83 | T | 142.65 | 136.41 | M | RNA binding;small ribosomal subunit rRNA binding;structural constituent of ribosome | translation | Maurer's cleft;cytosolic small ribosomal subunit;nucleolus;nucleus |
| PF3D7_1105100 | H2B | histone H2B | 13.13 | no | 0 | 309.53 | 379.32 | 2104.54 | 810.05 | 2957.94 | 3275.46 | 21511.98 | O | 3802.21 | 854.97 | M | DNA binding;RNA binding | chromosome organization;nucleosome assembly | nucleosome;nucleus |
| PF3D7_1310700 | PIC5 | PhIL1-interacting candidate PIC5 | 16.53 | no | 0 | 10.08 | 1.07 | 83.41 | 64.66 | 182.78 | 1099.54 | 2354.12 | O | 19.35 | 511.64 | F | protein binding | entry into host;schizogony;structural complex | inner membrane pellicle complex |
| PF3D7_1415400 | N/A | Btz domain-containing protein, putative | 178.50 | no | 0 | 13.21 | 35.55 | 33.82 | 4.41 | 8.76 | 33.77 | 47.12 | O | 159.31 | 48.38 | M | mRNA binding | N/A | nucleus |
| PF3D7_0406000 | N/A | conserved protein, unknown function | 60.61 | no | 0 | 21.03 | 12.36 | 26.17 | 12.52 | 9.31 | 51.61 | 47.61 | GV | 60.98 | 31.43 | M | N/A | N/A | cytoplasm |
| PF3D7_0111900 | N/A | mitochondrial distribution and morphology protein, putative | 9.58 | no | 0 | 21.04 | 26.57 | 47.82 | 5.77 | 40.70 | 83.27 | 220.67 | O | 206.69 | 70.87 | M | protein binding | N/A | mitochondrion |
| PF3D7_1211600 | LSD1 | lysine-specific histone demethylase 1, putative | 350.58 | no | 0 | 22.35 | 12.62 | 7.45 | 1.39 | 1.10 | 10.96 | 106.93 | O | 143.65 | 46.81 | M | oxidoreductase activity;polyamine oxidase activity | chromatin remodeling | nuclear periphery;nucleus |
| PF3D7_1107400 | RAD51 | DNA repair protein RAD51 | 38.59 | no | 0 | 37.19 | 12.00 | 71.34 | 14.19 | 32.42 | 202.44 | 711.67 | O | 143.07 | 928.22 | F | ATP binding;ATP-dependent activity, acting on DNA;DNA binding;DNA strand exchange activity;damaged DNA binding;double-stranded DNA binding;protein binding;single-stranded DNA binding | DNA recombinase assembly;DNA recombination;DNA repair;chromosome organization involved in meiotic cell cycle;mitochondrial double-strand break repair;mitotic recombination;reciprocal meiotic recombination;response to ionizing radiation;strand invasion | condensed nuclear chromosome;mitochondrion;nucleus |
| PF3D7_1107000 | LSM4 | U6 snRNA-associated Sm-like protein LSm4, putative | 14.20 | no | 0 | 124.50 | 88.08 | 123.78 | 164.71 | 222.49 | 137.82 | 84.79 | GV | 31.98 | 34.04 | O | RNA binding;U6 snRNA binding | P-body assembly;mRNA splicing, via spliceosome;spliceosomal snRNP assembly | P-body;U4/U6 x U5 tri-snRNP complex;U6 snRNP;nucleus;sno(s)RNA-containing ribonucleoprotein complex;spliceosomal tri-snRNP complex |
| PF3D7_0316500 | NUF2 | kinetochore protein NUF2, putative | 54.68 | no | 0 | 4.57 | 2.09 | 21.73 | 10.29 | 10.28 | 55.04 | 148.98 | O | 59.78 | 34.59 | M | protein-containing complex binding | attachment of mitotic spindle microtubules to kinetochore;chromosome segregation;kinetochore organization;meiotic chromosome segregation;mitotic spindle organization | Ndc80 complex;chromosome, centromeric region;condensed nuclear chromosome kinetochore;kinetochore;nucleus |
| PF3D7_0202000 | KAHRP | knob-associated histidine-rich protein | 71.30 | no | 0 | 4606.05 | 41.48 | 14.70 | 2.55 | 68.44 | 16.21 | 574.90 | R | 17.29 | 5.81 | M | RNA binding;actin binding;protein binding | cytoadherence to microvasculature, mediated by symbiont protein | Maurer's cleft;cytoplasm;extracellular vesicle;infected host cell surface knob |
| PF3D7_1324200 | N/A | micro-fibrillar-associated protein, putative | 59.63 | no | 0 | 220.15 | 128.47 | 84.99 | 37.19 | 29.09 | 24.42 | 187.92 | R | 258.46 | 589.70 | F | extracellular matrix structural constituent | mRNA splicing, via spliceosome | U2-type spliceosomal complex;extracellular matrix |
| PF3D7_1460600 | ISP3 | inner membrane complex sub-compartment protein 3 | 17.12 | no | 0 | 6.65 | 1.54 | 4.66 | 3.89 | 3.80 | 95.65 | 46.70 | GV | 6.75 | 59.88 | F | protein binding | structural complex | cell periphery;inner membrane pellicle complex |
| PF3D7_1237000 | UBA2 | SUMO-activating enzyme subunit 2 | 81.12 | no | 0 | 11.47 | 12.93 | 14.83 | 2.14 | 0.62 | 3.11 | 74.07 | O | 46.12 | 1.29 | M | SUMO activating enzyme activity;ubiquitin activating enzyme activity | protein modification by small protein conjugation;protein sumoylation | SUMO activating enzyme complex;cytoplasm |
| PF3D7_1365000 | N/A | conserved Plasmodium protein, unknown function | 42.22 | no | 0 | 26.37 | 5.27 | 7.84 | 0.78 | 1.82 | 25.01 | 24.98 | R | 60.92 | 78.88 | F | N/A | N/A | N/A |
| PF3D7_0913200 | EF-1beta | elongation factor 1-beta | 32.03 | no | 0 | 265.51 | 100.51 | 77.46 | 6.89 | 13.71 | 38.56 | 150.89 | R | 112.18 | 31.41 | M | guanyl-nucleotide exchange factor activity | translational elongation | Maurer's cleft;cytosol;eukaryotic translation elongation factor 1 complex;nucleus |
| PF3D7_1407900 | PMI | Plasmepsin I | 51.46 | no | 1 | 316.79 | 45.87 | 25.65 | 46388.00 | 47150.00 | 27120.00 | 41760.00 | GC II | 30225.00 | 0.35 | M | aspartic-type endopeptidase activity | hemoglobin catabolic process | food vacuole |
| PF3D7_0410300 | PPM1 | protein phosphatase PPM1, putative | 105.20 | no | 0 | 26.85 | 12.52 | 12.51 | 3.32 | 3.59 | 9.10 | 73.24 | O | 54.56 | 18.39 | M | magnesium-dependent protein serine/threonine phosphatase activity;phosphoprotein phosphatase activity;protein serine/threonine phosphatase activity | protein dephosphorylation | cytoplasm;nucleus |
| PF3D7_1337900 | N/A | conserved Plasmodium protein, unknown function | 25.63 | no | 0 | 22.55 | 4.05 | 7.07 | 0.81 | 5.72 | 49.93 | 14.40 | GV | 3.64 | 18.57 | F | N/A | N/A | N/A |
| PF3D7_1330800 | N/A | RNA-binding protein, putative | 68.01 | no | 0 | 404.82 | 484.55 | 63.89 | 11.47 | 24.20 | 54.54 | 1335.48 | O | 97.42 | 902.73 | F | RNA binding;mRNA binding | RNA splicing;mRNA splicing, via spliceosome;regulation of alternative mRNA splicing, via spliceosome | ASAP complex;cytoplasm;nucleoplasm;nucleus |
| PF3D7_0934100 | XPD | TFIIH transcription factor complex helicase XPD SU | 122.84 | no | 0 | 52.70 | 25.47 | 18.32 | 3.21 | 4.92 | 18.59 | 32.07 | R | 19.99 | 15.03 | M | 5'-3' DNA helicase activity;DNA helicase activity;damaged DNA binding;protein binding;single-stranded DNA helicase activity | nucleotide-excision repair, DNA duplex unwinding;nucleotide-excision repair, DNA incision;positive regulation of mitotic recombination;regulation of mitotic recombination;response to UV;response to oxidative stress;transcription by RNA polymerase II | cytoplasm;nucleus |
| PF3D7_1458800 | N/A | DNA-directed RNA polymerase III subunit RPC5, putative | 103.36 | no | 0 | 8.36 | 3.60 | 2.21 | 0.15 | 2.33 | 11.75 | 9.45 | GV | 22.51 | 4.24 | M | RNA binding | regulation of transcription | RNA polymerase III complex |
| PF3D7_1019600 | N/A | conserved Plasmodium protein, unknown function | 145.83 | no | 0 | 5.67 | 0.25 | 12.86 | 8.90 | 4.56 | 98.44 | 122.47 | O | 39.72 | 111.75 | F | N/A | N/A | N/A |
| PF3D7_0815600 | EIF3G | eukaryotic translation initiation factor 3 subunit G, putative | 31.25 | no | 0 | 186.04 | 119.45 | 51.26 | 19.72 | 24.30 | 59.82 | 64.99 | R | 158.59 | 441.14 | F | mRNA binding;translation initiation factor activity | translation;translational initiation | nucleus |
| PF3D7_1216900 | N/A | DNA-binding chaperone, putative | 111.06 | no | 0 | 220.24 | 178.03 | 59.42 | 5.91 | 6.12 | 29.52 | 76.35 | R | 118.79 | 49.72 | M | DNA binding;protein binding | protein folding | nucleus |
| PF3D7_1347700 | ECT | ethanolamine-phosphate cytidylyltransferase | 66.04 | no | 0 | 133.49 | 5.05 | 25.73 | 3.83 | 4.07 | 22.72 | 59.07 | R | 50.91 | 115.51 | F | ethanolamine-phosphate cytidylyltransferase activity | phospholipid biosynthetic process | cytosol |
| PF3D7_1145800 | N/A | conserved Plasmodium protein, unknown function | 108.95 | no | 0 | 32.44 | 12.89 | 4.48 | 0.85 | 1.86 | 15.25 | 30.31 | R | 31.80 | 17.65 | M | N/A | N/A | N/A |
| PF3D7_1250300 | VPS26 | Vacuolar protein sorting-associated protein 26, putative | 34.51 | no | 0 | 43800.00 | 14.51 | 47.71 | 13058.00 | 23.35 | 58.04 | 393.18 | R | 122.13 | 216.04 | F | protein binding | intracellular protein transport;retrograde transport, endosome to Golgi | endosome;retromer complex |
| PF3D7_0912400 | N/A | alkaline phosphatase, putative | 52.70 | yes | 1 | 27.39 | 39.52 | 46.53 | 8.81 | 21.87 | 34.01 | 30.26 | T | 31.21 | 22.98 | M | enzyme activity | N/A | apicoplast;nucleus;plasma membrane;symbiont-containing vacuole |
| PF3D7_0708100 | RPB10 | DNA-directed RNA polymerases I, II, and III subunit RPABC5, putative | 8.24 | no | 0 | 71.20 | 23.14 | 13.25 | 0.61 | 41.22 | 177.64 | 30.92 | GV | 51.46 | 51.89 | O | DNA-directed 5'-3' RNA polymerase activity;zinc ion binding | tRNA transcription by RNA polymerase III;transcription by RNA polymerase I;transcription by RNA polymerase II | RNA polymerase I complex;RNA polymerase II, core complex;RNA polymerase III complex;nucleus |
| PF3D7_1447900 | MDR2 | multidrug resistance protein 2 | 119.01 | no | 10 | 11.93 | 35.04 | 9.88 | 1.07 | 1.95 | 11.37 | 9.42 | T | 14.29 | 14.64 | O | ATPase-coupled transmembrane transporter activity;metal ion binding;metal ion transmembrane transporter activity | response to xenobiotic stimulus;transmembrane transport;xenobiotic transmembrane transport | symbiont-containing vacuole;integral component of membrane |
| PF3D7_1404000 | RPB4 | DNA-directed RNA polymerase II subunit RPB4, putative | 20.05 | no | 0 | 14.20 | 9.21 | 6.56 | 1.07 | 44.29 | 865.01 | 61.34 | GV | 247.78 | 378.31 | F | RNA binding;DNA binding | regulation of transcription | nucleus |
| PF3D7_1426200 | PRMT1 | protein arginine N-methyltransferase 1 | 47.45 | no | 0 | 88.70 | 62.27 | 26.83 | 4.28 | 3.73 | 11.61 | 63.39 | R | 14.90 | 20.57 | F | protein-arginine N-methyltransferase activity | RNA metabolic process;RNA processing;histone methylation;ribosome biogenesis | cytoplasm;nucleus |
| PF3D7_0522800 | BUD31 | pre-mRNA-splicing factor BUD31, putative | 17.39 | no | 0 | 34.39 | 16.03 | 6.69 | 1.24 | 11.43 | 27.64 | 120.15 | O | 39.71 | 5.58 | M | RNA binding | RNA splicing;alternative mRNA splicing, via spliceosome;mRNA splicing, via spliceosome | U2 snRNP;nucleus;spliceosomal complex |
| PF3D7_0723400 | N/A | conserved Plasmodium protein, unknown function | 167.92 | no | 0 | 192.48 | 79.38 | 22.21 | 2.19 | 4.65 | 21.64 | 144.23 | R | 237.30 | 81.59 | M | N/A | N/A | nucleus |
| PF3D7_1014700 | PHB2 | prohibitin 2 | 34.79 | no | 0 | 22.44 | 37.69 | 10.00 | 0.79 | 25.59 | 90.63 | 58.68 | GV | 16.53 | 59.92 | F | protein binding | cellular process;mitochondrion organization;response to xenobiotic stimulus | mitochondrion |
| PF3D7_0715300 | N/A | calcium/calmodulin-dependent protein kinase, putative | 61.33 | no | 0 | 3.56 | 0.14 | 10.76 | 2.99 | 7.06 | 40.67 | 26.17 | GV | 6.03 | 32.67 | F | protein kinase activity | protein phosphorylation | cytoplasm |
| PF3D7_0704200 | NSUN1 | RNA cytosine C(5)-methyltransferase, putative | 141.23 | no | 0 | 17.70 | 12.83 | 4.69 | 0.51 | 0.66 | 7.06 | 14.21 | R | 1.49 | 6.38 | F | methyltransferase activity | RNA methylation | nucleus |
| PF3D7_1426700 | PEPC | phosphoenolpyruvate carboxylase | 133.96 | no | 0 | 26.11 | 10.19 | 25.91 | 9.77 | 6.99 | 63.94 | 105.02 | O | 217.02 | 79.21 | M | phosphoenolpyruvate carboxylase activity | carbon fixation | cytosol;nucleus |
| PF3D7_1117800 | MLH | DNA mismatch repair protein MLH | 118.31 | no | 0 | 13.04 | 3.70 | 10.35 | 2.81 | 2.66 | 12.61 | 20.56 | O | 85.60 | 11.79 | M | ATP binding;ATP hydrolysis activity;endonuclease activity;protein binding | mismatch repair | MutLalpha complex;chiasma;chromosome, telomeric region;mismatch repair complex |
| PF3D7_0728600 | N/A | RING zinc finger protein, putative | 253.97 | no | 0 | 7.82 | 1.93 | 18.21 | 6.73 | 8.82 | 86.84 | 270.45 | O | 914.65 | 299.48 | M | RNA binding | N/A | nucleus;cytoplasm |
| PF3D7_0727800 | N/A | cation transporting ATPase, putative | 225.07 | no | 12 | 27.54 | 32.28 | 30.24 | 7.68 | 3.40 | 27.99 | 58.33 | O | 21.65 | 37.62 | F | transporter activity | cellular calcium ion homeostasis | integral component of membrane;membrane |
| PF3D7_0604300 | N/A | conserved Plasmodium protein, unknown function | 91.57 | no | 0 | 11.18 | 9.64 | 16.87 | 3.43 | 8.93 | 81.80 | 17.35 | GV | 22.29 | 6.29 | M | N/A | N/A | N/A |
| PF3D7_1205500 | N/A | zinc finger protein, putative | 204.85 | no | 0 | 230.33 | 15.87 | 84.84 | 23.09 | 15.07 | 113.73 | 119.58 | R | 200.69 | 86.48 | M | ubiquitin protein ligase activity | protein ubiquitination | cytoplasm;nucleus |
| PF3D7_0807300 | RAB18 | ras-related protein Rab-18 | 23.16 | no | 0 | 166.44 | 89.06 | 92.04 | 19.05 | 134.25 | 44.56 | 33.79 | R | 38.90 | 14.27 | M | GTP binding;GTPase activity | intracellular protein transport;small GTPase mediated signal transduction | Golgi apparatus;endomembrane system;membrane;nucleus |
| PF3D7_0407700 | N/A | TH1 domain-containing protein, putative | 197.41 | no | 0 | 6.11 | 0.33 | 13.33 | 9.64 | 3.60 | 21.13 | 51.24 | O | 208.91 | 61.67 | M | RNA binding | negative regulation of transcription elongation from RNA polymerase II promoter | NELF complex;nucleus |
| PF3D7_1010600 | eIF2beta | eukaryotic translation initiation factor 2 subunit beta | 25.32 | no | 0 | 429.26 | 92.40 | 93.89 | 18.73 | 39.05 | 106.08 | 58.49 | R | 91.58 | 89.95 | O | RNA binding;mRNA binding;protein binding;translation initiation factor activity;translation initiation factor binding | formation of cytoplasmic translation initiation complex;formation of translation preinitiation complex;translational initiation | cytoplasm;eukaryotic translation initiation factor 2 complex;nucleus |
| PF3D7_1107200 | N/A | vacuolar fusion protein CCZ1, putative | 84.21 | no | 0 | 17.89 | 8.27 | 15.47 | 2.32 | 1.39 | 15.97 | 23.15 | O | 49.17 | 41.96 | M | protein binding | vesicle-mediated transport | intracellular membrane-bounded organelle |
| PF3D7_1136400 | SRP72 | signal recognition particle subunit SRP72, putative | 107.11 | no | 0 | 41.29 | 10.97 | 13.25 | 3.07 | 3.43 | 17.82 | 8.16 | R | 29.05 | 14.37 | M | 7S RNA binding;signal sequence binding | SRP-dependent cotranslational protein targeting to membrane | signal recognition particle, endoplasmic reticulum targeting |
| PF3D7_0917500 | N/A | conserved protein, unknown function | 79.67 | no | 0 | 5.61 | 1.47 | 28.42 | 13.25 | 5.88 | 22.82 | 44.10 | O | 174.40 | 35.60 | M | N/A | N/A | N/A |
| PF3D7_1407100 | NOP1 | rRNA 2'-O-methyltransferase fibrillarin, putative | 34.13 | no | 0 | 246.22 | 243.57 | 61.48 | 23.22 | 14.98 | 18.28 | 129.25 | R | 100.49 | 69.74 | M | RNA binding;histone-glutamine methyltransferase activity;rRNA methyltransferase activity | box C/D RNA 3'-end processing;histone glutamine methylation;rRNA methylation;rRNA processing | Cajal body;box C/D RNP complex;nucleolus;nucleus;small-subunit processome;sno(s)RNA-containing ribonucleoprotein complex |
| PF3D7_1331200 | N/A | conserved Plasmodium protein, unknown function | 48.62 | no | 0 | 56.43 | 24.03 | 11.76 | 2.96 | 6.94 | 50.28 | 105.66 | O | 25.60 | 9.83 | M | N/A | N/A | N/A |
| PF3D7_1002900 | N/A | conserved Plasmodium protein, unknown function | 14.52 | no | 1 | 50.36 | 4.26 | 23.26 | 6.07 | 120.56 | 244.38 | 91.19 | GV | 41.53 | 41.30 | O | | | |
| PF3D7_0323700 | SART1 | U4/U6.U5 tri-snRNP-associated protein 1, putative | 82.90 | no | 0 | 35.40 | 16.66 | 13.93 | 2.00 | 2.24 | 14.30 | 77.78 | O | 94.06 | 66.24 | M | RNA binding | mRNA cis splicing, via spliceosome;maturation of 5S rRNA | U4/U6 x U5 tri-snRNP complex;nucleus |
| PF3D7_0926100 | N/A | protein kinase, putative | 366.96 | no | 0 | 119.10 | 25.64 | 28.40 | 3.99 | 4.52 | 33.34 | 64.48 | R | 64.92 | 17.67 | M | protein kinase activity | protein phosphorylation | N/A |
| PF3D7_0801000 | N/A | Plasmodium exported protein (PHISTc) | 147.07 | no | 1 | 178.02 | 162.69 | 23.28 | 0.77 | 0.71 | 3.38 | 1036.96 | O | 1.15 | 13.76 | F | protein binding | entry into host | Maurer's cleft;extracellular vesicle;nucleus;protein-containing complex;rhoptry |
| PF3D7_1331700 | N/A | glutamine--tRNA ligase, putative | 108.53 | no | 0 | 125.78 | 111.41 | 71.69 | 18.35 | 5.16 | 18.90 | 52.00 | R | 13.81 | 25.91 | F | glutamine-tRNA ligase activity | glutaminyl-tRNA aminoacylation | cytosol;nucleus |
| PF3D7_0320500 | Nico | nicotinamidase, putative | 50.15 | no | 0 | 14.81 | 25.95 | 22.97 | 8.21 | 4.17 | 77.65 | 68.32 | GV | 18.72 | 60.13 | F | enzyme activity | N/A | cytoplasm;nucleus |
| PF3D7_1305300 | GCN1 | translational activator GCN1, putative | 418.58 | no | 0 | 57.07 | 26.86 | 11.37 | 0.93 | 3.17 | 27.57 | 15.41 | R | 39.26 | 12.22 | M | protein binding | regulation of translation | nucleus;cytoplasm |
| PF3D7_1010700 | N/A | dolichyl-phosphate-mannose--protein mannosyltransferase, putative | 25.68 | yes | 0 | 260.52 | 111.96 | 147.15 | 32.08 | 290.95 | 239.73 | 99.78 | G II | 247.09 | 172.69 | M | enzyme activity | N/A | nucleus |
| PF3D7_1118200 | N/A | heat shock protein 90, putative | 108.46 | | 0 | 35.68 | 74.78 | 27.10 | 1.05 | 6.37 | 28.80 | 106.59 | O | 57.53 | 167.41 | F | ATP binding;ATP hydrolysis activity;mRNA binding;protein kinase binding;unfolded protein binding | protein folding;response to heat;response to unfolded protein | mitochondrial inner membrane;mitochondrion;nucleus |
| PF3D7_0528100 | N/A | AP-1/2 complex subunit beta, putative | 106.15 | no | 0 | 115.66 | 50.41 | 59.66 | 20.46 | 6.30 | 21.67 | 81.95 | R | 82.01 | 46.56 | M | protein binding;clathrin binding | vesicle-mediated transport | clathrin adapter complex;membran coat |
| PF3D7_1238600 | NSM | sphingomyelin phosphodiesterase | 46.01 | no | 0 | 6.00 | 0.74 | 25.52 | 17.68 | 4.84 | 16.35 | 72.73 | O | 12.80 | 8.17 | M | nuclease activity;phospholipase activity;sphingomyelin phosphodiesterase activity | sphingomyelin metabolic process | cytoplasm;nucleus |
| PF3D7_1147300 | N/A | conserved Plasmodium protein, unknown function | 158.90 | no | 0 | 7.42 | 5.90 | 18.37 | 3.37 | 4.97 | 56.97 | 212.49 | O | 66.77 | 92.87 | F | N/A | N/A | nucleus |
| PF3D7_0824200 | N/A | conserved Plasmodium protein, unknown function | 133.80 | no | 0 | 4.10 | 1.68 | 4.40 | 0.65 | 1.68 | 9.74 | 27.78 | O | 65.97 | 3.04 | M | N/A | N/A | N/A |

| | | | | | | | | | | | | | | | | | | | |
|---|---|---|---|---|---|---|---|---|---|---|---|---|---|---|---|---|---|---|---|
| PF3D7_1438600 | GET4 | Golgi to ER traffic protein 4 | 33.15 | no | 0 | 16.55 | 2.30 | 5.21 | 1.09 | 7.41 | 42.12 | 0.00 | GV | 9.57 | 8.61 | 0 | protein binding | protein insertion into ER membrane | BAT3 complex;cytosol |
| PF3D7_1119800 | ASF1 | alternative splicing factor ASF-1, putative | 23.65 | no | 0 | 25.48 | 9.96 | 10.18 | 3.59 | 17.60 | 59.49 | 10.61 | GV | 35.15 | 24.44 | M | RNA binding;mRNA binding | alternative mRNA splicing, via spliceosome | Maurer's cleft;nucleus |
| PF3D7_1225600 | N/A | Uncharacterized protein | 93.05 | no | 0 | 43800.00 | 14.51 | 47.71 | 13058.00 | 23.35 | 58.04 | 393.18 | R | 122.13 | 216.04 | F | N/A | N/A | N/A |
| PF3D7_0318600 | N/A | cleavage and polyadenylation specificity factor, putative | 118.85 | no | 0 | 39.76 | 16.85 | 7.15 | 2.21 | 1.22 | 9.22 | 4.83 | R | 20.95 | 10.82 | M | mRNA binding | mRNA cleavage;mRNA polyadenylation;snRNA processing | mRNA cleavage and polyadenylation specificity factor complex;nucleus |
| PF3D7_0909900 | N/A | helicase SKI2W, putative | 160.37 | no | 0 | 28.73 | 7.92 | 11.19 | 1.63 | 2.02 | 9.06 | 8.98 | R | 20.33 | 10.27 | M | RNA helicase activity | RNA catabolic process;nuclear-transcribed mRNA catabolic process, 3'-5' exonucleolytic nonsense-mediated decay | Ski complex |
| PF3D7_1123200 | LRR11 | leucine-rich repeat protein | 93.33 | no | 0 | 10.22 | 0.77 | 17.35 | 7.05 | 6.67 | 13.40 | 30.64 | O | 394.72 | 15.99 | M | protein binding | SCF-dependent proteasomal ubiquitin-dependent protein catabolic process;ubiquitin-dependent protein catabolic process | SCF ubiquitin ligase complex;cytoplasm;nucleus |
| PF3D7_1363100 | N/A | conserved Plasmodium protein, unknown function | 124.75 | no | 0 | 12.49 | 1.69 | 6.82 | 1.85 | 1.33 | 15.08 | 29.31 | O | 22.38 | 9.13 | M | N/A | N/A | N/A |
| PF3D7_1345600 | N/A | inner membrane complex protein | 43.22 | no | 0 | 2.87 | 0.43 | 31.89 | 30.07 | 23.42 | 26.26 | 109.96 | O | 25.79 | 14.82 | M | protein binding | structural complex | inner membrane pellicle complex |
| PF3D7_1238800 | ACS11 | acyl-CoA synthetase | 92.09 | no | 0 | 60.81 | 21.66 | 22.81 | 5.05 | 1.21 | 19.16 | 60.66 | R | 17.87 | 222.03 | F | long-chain fatty acid-CoA ligase activity | fatty acid metabolic process;response to xenobiotic stimulus | endoplasmic reticulum;membrane;nucleus |
| PF3D7_1469600 | ACC | acetyl-CoA carboxylase | 394.88 | yes | 0 | 4.70 | 0.45 | 3.53 | 0.62 | 1.71 | 21.81 | 117.71 | O | 7.36 | 51.35 | F | acetyl-CoA carboxylase activity;biotin binding;biotin carboxylase activity | fatty acid biosynthetic process | acetyl-CoA carboxylase complex;apicoplast;biotin carboxylase complex |
| PF3D7_1126000 | ThrRS | threonine--tRNA ligase | 119.54 | yes | 0 | 97.23 | 75.56 | 39.69 | 12.63 | 9.31 | 10.24 | 58.64 | R | 36.62 | 36.04 | 0 | ATP binding;aminoacyl-tRNA ligase activity;threonine-tRNA ligase activity | response to drug;response to xenobiotic stimulus;tRNA aminoacylation for protein translation;threonyl-tRNA aminoacylation | apicoplast;cytosol;mitochondrion;nucleus |
| PF3D7_0705700 | N/A | 40S ribosomal protein S29, putative | 6.42 | no | 0 | 307.36 | 185.31 | 107.65 | 20.54 | 100.80 | 118.34 | 80.20 | R | 185.40 | 134.98 | M | structural constituent of ribosome;zinc ion binding | | cytosolic small ribosomal subunit;nucleus |
| PF3D7_1116700 | DPAP1 | Dipeptidyl aminopeptidase 1 | 80.41 | yes | 1 | 460.35 | 641.9 | 162.52 | 21.16 | 16316.00 | 16.24 | 295.04 | GC II | 19.38 | 34.67 | F | cysteine-type endopeptidase activity | proteolysis involved in cellular protein catabolic process;response to xenobiotic stimulus | extracellular space;food vacuole;lysosome;symbiont-containing vacuole |
| PF3D7_0812100 | N/A | proteasome activator complex subunit 4, putative | 314.49 | no | 2 | 19.79 | 8.86 | 16.15 | 3.48 | 2.39 | 18.84 | 18.55 | R | 21.74 | 7.75 | M | lysine-acetylated histone binding;peptidase activator activity;proteasome binding | proteasomal ubiquitin-independent protein catabolic process | cytosol;nucleus |
| PF3D7_1304200 | MIT2 | CorA-like Mg2+ transporter protein, putative | 55.01 | no | 2 | 45325.00 | 42767.00 | 11841.00 | 41730.00 | 42461.00 | 31017.00 | 66.53 | R | 45490.00 | 15.44 | M | magnesium ion transmembrane transporter activity | magnesium ion transport | integral component of membrane |
| PF3D7_0217900 | N/A | thioesterase/thiol ester dehydrase-isomerase, putative | 60.60 | no | 0 | 23.42 | 3.07 | 34.86 | 5.31 | 2.73 | 16.23 | 36.09 | O | 15.26 | 8.31 | M | acyl-CoA hydrolase activity | acyl-CoA metabolic process | mitochondrion |
| PF3D7_0718100 | EST | exported serine/threonine protein kinase | 317.69 | no | 0 | 5.74 | 0.79 | 9.74 | 1.62 | 2.67 | 18.85 | 28.62 | O | 286.85 | 23.21 | M | protein serine/threonine kinase activity | protein phosphorylation | Maurer's cleft;extracellular vesicle;membrane |
| PF3D7_0920400 | N/A | conserved Plasmodium protein, unknown function | 224.49 | no | 0 | 7.20 | 4.58 | 5.47 | 1.05 | 1.17 | 8.74 | 11.57 | O | 18.47 | 8.44 | M | N/A | N/A | N/A |
| PF3D7_0416400 | HAT1 | histone acetyltransferase, putative | 146.72 | no | 0 | 23.91 | 32.37 | 90.95 | 27.34 | 6.12 | 10.98 | 652.73 | O | 6.29 | 71.97 | F | H4 histone acetyltransferase activity | histone H4 acetylation | nucleus |
| PF3D7_0920800 | IMPDH | inosine-5'-monophosphate dehydrogenase | 56.15 | no | 0 | 234.36 | 64.70 | 21.60 | 4.09 | 3.00 | 8.36 | 21.86 | R | 21.88 | 18.36 | 0 | enzyme activity | N/A | food vacuole;nucleus |
| PF3D7_1146000 | RSA4 | ribosome assembly protein 4, putative | 73.13 | no | 0 | 47.40 | 21.69 | 12.96 | 4.85 | 6.60 | 12.82 | 98.11 | O | 7.16 | 27.04 | F | protein binding | ribosomal large subunit assembly | nucleolus;nucleus |
| PF3D7_0523600 | N/A | conserved protein, unknown function | 24.56 | no | 0 | 5.15 | 4.45 | 34.45 | 21.57 | 53.67 | 28.49 | 34.37 | G II | 190.88 | 29.64 | M | N/A | N/A | N/A |
| PF3D7_0423500 | GAPM2 | glideosome associated protein with multiple membrane spans 2 | 42.62 | no | 6 | 28.31 | 4.21 | 323.15 | 618.06 | 155.84 | 285.46 | 373.58 | S | 190.46 | 254.38 | F | RNA binding;protein binding | entry into host | cell surface;inner membrane pellicle complex;protein-containing complex |
| PF3D7_0405100 | SEC24B | protein transport protein Sec24B, putative | 155.94 | no | 0 | 39.22 | 11.12 | 37.10 | 12.71 | 6.75 | 17.57 | 25.41 | R | 70.97 | 32.45 | M | SNARE binding;zinc ion binding | COPII-coated vesicle cargo loading | COPII vesicle coat;Maurer's cleft;endoplasmic reticulum exit site |
| PF3D7_0920900 | PRP24 | U4/U6 snRNA-associated-splicing factor, putative | 56.65 | no | 0 | 39.86 | 7.84 | 12.14 | 1.72 | 2.14 | 24.75 | 6.31 | R | 38.25 | 13.46 | M | RNA binding | mRNA splicing | ribonucleoprotein complex |
| PF3D7_1201000 | N/A | Plasmodium exported protein (PHISTb), unknown function | 77.12 | yes | 0 | 69.59 | 454.13 | 54.46 | 0.93 | 1.70 | 2.17 | 34.54 | T | 49.86 | 3.58 | M | protein binding | N/A | host cell periphery;host cell surface;infected host cell surface knob;nucleus;symbiont-containing vacuole |
| PF3D7_0209200 | MTR3 | exosome complex component MTR3, putative | 31.25 | no | 0 | 35.68 | 10.80 | 8.20 | 0.85 | 7.36 | 13.71 | 177.10 | O | 8.70 | 13.51 | F | protein binding | U4 snRNA 3'-end processing;nuclear mRNA surveillance;nuclear-transcribed mRNA catabolic process, exonucleolytic, 3'-5';polyadenylation-dependent snoRNA 3'-end processing;rRNA catabolic process | cytoplasmic exosome (RNase complex);exosome (RNase complex);nuclear exosome (RNase complex);nucleolus;osmiophilic body |
| PF3D7_0507900 | N/A | conserved Plasmodium protein, unknown function | 13.51 | no | 0 | 131.62 | 28.67 | 21.53 | 2.19 | 36.68 | 18.34 | 20.13 | R | 1.52 | 36.63 | F | N/A | N/A | N/A |
| PF3D7_1368400 | N/A | ribosomal protein L1, putative | 38.26 | no | 0 | 24.67 | 8.11 | 13.08 | 2.93 | 7.44 | 84.45 | 40.56 | GV | 17.78 | 101.47 | F | RNA binding;structural constituent of ribosome | translation | cytosol;cytosolic small ribosomal subunit;nucleus |
| PF3D7_0505100 | TRS85 | trafficking protein particle complex subunit 8, putative | 328.67 | no | 0 | 22.85 | 17.38 | 18.45 | 7.85 | 2.48 | 12.13 | 169.71 | O | 26.61 | 25.56 | 0 | protein binding | autophagosome assembly;autophagy of peroxisome;endoplasmic reticulum to Golgi vesicle-mediated transport;protein localization to phagophore assembly site | TRAPPIII protein complex;cytoplasmic vesicle;nucleus;phagophore assembly site |
| PF3D7_0404300 | N/A | Ran-binding protein, putative | 74.18 | no | 0 | 160.64 | 113.12 | 83.13 | 17.64 | 22.05 | 11.10 | 205.79 | O | 769.74 | 277.07 | M | protein binding | intracellular transport | nucleus |
| PF3D7_1345100 | TRX2 | thioredoxin 2 | 18.63 | yes | 0 | 210.98 | 41.22 | 18.97 | 1.53 | 23.29 | 110.97 | 54.77 | R | 17.76 | 7.38 | M | oxidoreductase activity, acting on a sulfur group of donors, disulfide as acceptor;protein binding;protein-disulfide reductase (NAD(P)) activity;protein-disulfide reductase activity | cell redox homeostasis | PTEX complex;cytoplasm;mitochondrion;symbiont-containing vacuole |
| PF3D7_1239700 | FTSH1 | ATP-dependent zinc metalloprotease FTSH 1 | 100.96 | no | 1 | 12.57 | 49.67 | 106.81 | 22.50 | 1.32 | 5.74 | 243.07 | O | 48.59 | 19.17 | M | ATP binding;ATP-dependent peptidase activity;metalloendopeptidase activity | plastid organization;proteolysis;response to xenobiotic stimulus | apicoplast;mitochondrion |
| PF3D7_1408100 | PMIII | plasmepsin III | 51.69 | no | 1 | 331.47 | 158.86 | 69.58 | 5.84 | 12.64 | 12.54 | 48.62 | R | 37.03 | 9.53 | M | aspartic-type endopeptidase activity | response to xenobiotic stimulus | food vacuole |
| PF3D7_0520300 | LSM2 | U6 snRNA-associated Sm-like protein LSm2, putative | 12.06 | no | 0 | 86.80 | 33.23 | 16.22 | 2.09 | 28.04 | 46.39 | 38.90 | R | 40.96 | 20.43 | M | RNA binding | mRNA splicing, via spliceosome | Lsm1-7-Pat1 complex;P-body;U4/U6 x U5 tri-snRNP complex;U6 snRNP;catalytic step 2 spliceosome;intracellular;nucleus;precatalytic spliceosome;sno(s)RNA-containing ribonucleoprotein complex |
| PF3D7_0602600 | N/A | SAC3 domain-containing protein, putative | 153.72 | no | 0 | 63.37 | 11.71 | 8.11 | 1.23 | 2.24 | 17.14 | 11.54 | R | 34.02 | 12.93 | M | RNA binding | mRNA export from nucleus | cytoplasm;nucleus;transcription export complex 2 |
| PF3D7_1021400 | N/A | endomembrane protein 70, putative | 72.53 | yes | 9 | 87.87 | 11.41 | 32.77 | 8.86 | 9.06 | 20.44 | 29.38 | R | 16.17 | 17.21 | 0 | N/A | protein localization to membrane | integral component of membrane |
| PF3D7_0827000 | DBP10 | ATP-dependent RNA helicase DBP10, putative | 151.10 | no | 0 | 70.36 | 41.06 | 15.42 | 1.18 | 0.91 | 8.26 | 71.34 | O | 164.04 | 65.86 | M | ATP binding;RNA binding | assembly of large subunit precursor of preribosome | nucleolus |
| PF3D7_1033200 | ETRAMP10.2 | early transcribed membrane protein 10.2 | 38.93 | yes | 2 | 56.51 | 469.05 | 113.15 | 43.66 | 44.92 | 13.95 | 333.37 | T | 4.22 | 3.51 | 0 | N/A | N/A | Maurer's cleft;symbiont-containing vacuole membrane |
| PF3D7_0520700 | N/A | CDC73 domain-containing protein, putative | 118.00 | no | 0 | 17.35 | 11.54 | 4.41 | 0.54 | 1.68 | 16.96 | 2.06 | R | 4.91 | 7.37 | F | RNA polymerase II complex binding | positive regulation of transcription elongation from RNA polymerase II promoter;recruitment of 3'-end processing factors to RNA polymerase II holoenzyme complex | Cdc73/Paf1 complex |
| PF3D7_1428800 | N/A | Transcription initiation TFIID-like, putative | 41.98 | no | 1 | 42583.00 | 43556.00 | 30072.00 | 35431.00 | 15281.00 | 107.79 | 45319.00 | O | 24.15 | 67.6 | F | RNA polymerase II general transcription initiation factor activity | DNA-templated transcription, initiation | nucleus |
| PF3D7_1419000 | N/A | conserved Plasmodium protein, unknown function | 136.94 | no | 0 | 10.25 | 3.42 | 24.40 | 5.62 | 5.15 | 25.57 | 77.83 | O | 362.77 | 113.47 | M | N/A | N/A | nucleus |
| PF3D7_1328500 | N/A | alpha/beta-hydrolase, putative | 115.89 | no | 0 | 12.80 | 33.77 | 19.42 | 2.03 | 1.84 | 9.48 | 60.10 | O | 17.41 | 11.37 | M | enzyme activity | N/A | N/A |
| PF3D7_1324800 | DHFS-FPGS | dihydrofolate synthase/folylpolyglutamate synthase | 60.06 | no | 0 | 27.12 | 7.88 | 34.46 | 5.42 | 3.89 | 14.16 | 1014.84 | O | 25.35 | 3.65 | M | ATP binding;dihydrofolate synthase activity;tetrahydrofolylpolyglutamate synthase activity | folic acid-containing compound biosynthetic process | cytoplasm;cytosol;mitochondrion |
| PF3D7_0903500 | NUP138 | nucleoporin NUP138, putative | 155.15 | no | 0 | 72.56 | 28.49 | 20.67 | 5.55 | 1.90 | 6.39 | 33.44 | R | 61.61 | 18.41 | M | protein binding | nucleocytoplasmic transport | nuclear pore;nucleus |
| PF3D7_1342800 | PEPCK | phosphoenolpyruvate carboxykinase | 66.21 | no | 0 | 48.08 | 81.13 | 25.73 | 1.75 | 3.23 | 84.88 | 206.04 | O | 16.63 | 437.73 | F | phosphoenolpyruvate carboxykinase (ATP) activity | gluconeogenesis | cytosol;mitochondrion;nucleus |
| PF3D7_1435700.2 | N/A | Ataxin-2 like protein, putative | 105.19 | no | 0 | 34790.00 | 15.75 | 34.21 | 34790.00 | 19.97 | 42.6 | 96.12 | S | 180.07 | 68.06 | M | RNA binding | regulation of cytoplasmic mRNA processing body assembly;stress granule assembly | cytoplasmic stress granule;mitochondrion |
| PF3D7_1021200 | N/A | conserved Plasmodium protein, unknown function | 82.85 | no | 0 | 5.73 | 2.18 | 33.86 | 12.13 | 8.98 | 41.80 | 1142.78 | O | 714.78 | 99.01 | M | N/A | N/A | N/A |
| PF3D7_1464000 | N/A | YL1 nuclear protein, putative | 84.61 | no | 0 | 75.86 | 39.27 | 26.34 | 8.06 | 3.41 | 17.98 | 15.92 | R | 50.61 | 71.71 | F | protein binding | histone exchange | nucleus |
| PF3D7_0924700 | SF3A3 | splicing factor 3A subunit 3, putative | 71.38 | no | 0 | 35.49 | 11.86 | 12.14 | 1.21 | 9.08 | 142.54 | 246.09 | O | 46.13 | 27.27 | M | RNA binding | mRNA splicing, via spliceosome | spliceosomal complex |
| PF3D7_1463400 | N/A | DNA-directed RNA polymerase III subunit RPC4, putative | 39.12 | no | 0 | 25.05 | 7.33 | 3.86 | 0.16 | 4.16 | 19.51 | 5.41 | R | 34.50 | 7.62 | M | DNA binding;RNA binding | regulation of transcription | RNA polymerase III complex |
| PF3D7_1105800 | N/A | conserved Apicomplexan protein, unknown function | 30.66 | no | 1 | 92.54 | 99.60 | 106.58 | 36.70 | 48.22 | 91.20 | 175.82 | O | 42.58 | 56.12 | F | N/A | N/A | food vacuole;mitochondrion |
| PF3D7_0828500 | N/A | translation initiation factor eIF-2B subunit alpha, putative | 40.03 | no | 0 | 3.41 | 0.43 | 22.89 | 6.28 | 10.97 | 61.12 | 37.60 | GV | 19.20 | 55.65 | F | translation initiation factor activity | regulation of translational initiation;translational initiation | eukaryotic translation initiation factor 2B complex |
| PF3D7_0407900 | N/A | AAA family ATPase, putative | 77.81 | no | 0 | 10.99 | 4.06 | 9.67 | 1.95 | 1.34 | 9.13 | 51.57 | O | 4.51 | 49.00 | F | ATP hydrolysis activity;microtubule-severing ATPase activity | cytoplasmic microtubule organization;regulation of double-strand break repair via homologous recombination | nucleus |
| PF3D7_1459000 | DBP5 | ATP-dependent RNA helicase DBP5 | 84.31 | no | 0 | 219.61 | 30.01 | 45.93 | 5.12 | 8.44 | 27.29 | 267.93 | O | 73.85 | 66.19 | M | 3'-5' DNA/RNA helicase activity;5'-3' DNA/RNA helicase activity;ATP hydrolysis activity;RNA binding;RNA helicase activity;single-stranded DNA helicase activity | DNA duplex unwinding;poly(A)+ mRNA export from nucleus | cytoplasm;cytoplasmic stress granule;nucleus |
| PF3D7_1202600 | NEMF | nuclear export mediator factor NEMF, putative | 254.29 | no | 0 | 217.88 | 133.11 | 94.30 | 76.86 | 14.44 | 44.10 | 660.64 | O | 234.67 | 357.87 | F | ribosomal large subunit binding;tRNA binding | nuclear export;rescue of stalled ribosome;ribosome-associated ubiquitin-dependent protein catabolic process | RQC complex;nucleus |
| PF3D7_1103700 | CK2beta1 | casein kinase II beta chain | 28.37 | no | 0 | 60.92 | 20.73 | 30.85 | 3.93 | 24.82 | 34.67 | 32.46 | R | 12.28 | 21.31 | F | protein kinase regulator activity;protein serine/threonine kinase activity | protein phosphorylation | cytoplasm;nucleus;protein kinase CK2 complex |
| PF3D7_0316600 | FNT | formate-nitrite transporter | 34.46 | no | 6 | 189.02 | 171.80 | 80.43 | 15.33 | 14.45 | 27.68 | 119.68 | R | 54.01 | 15.98 | M | high-affinity secondary active nitrite transmembrane transporter activity;lactate:proton symporter activity | acetate transport;formate transport;lactate transmembrane transport;nitrite transport;pyruvate transport;response to xenobiotic stimulus | cell surface;integral component of plasma membrane;plasma membrane |
| PF3D7_0205400 | N/A | PCI domain-containing protein, putative | 54.35 | no | 0 | 49.55 | 8.48 | 7.44 | 0.71 | 7.44 | 35.13 | 20.83 | R | 42.01 | 27.67 | M | RNA binding;double-stranded DNA binding | nuclear retention of pre-mRNA at the site of transcription;poly(A)+ mRNA export from nucleus;posttranscriptional tethering of RNA polymerase II gene DNA at nuclear periphery;transcription elongation from RNA polymerase II promoter | transcription export complex 2 |
| PF3D7_0531400 | N/A | G-patch domain-containing protein, putative | 142.14 | no | 0 | 14.80 | 5.01 | 2.84 | 0.47 | 1.48 | 17.14 | 3.31 | GV | 19.23 | 5.64 | M | protein binding | spliceosomal complex disassembly | U2-type post-mRNA release spliceosomal complex |
| PF3D7_1414000 | RPN13 | 26S proteasome regulatory subunit RPN13, putative | 29.49 | no | 0 | 11.34 | 4.56 | 40.36 | 7.59 | 15.47 | 60.24 | 104.54 | O | 133.30 | 141.30 | F | endopeptidase activator activity;proteasome binding;ubiquitin binding | ubiquitin-dependent protein catabolic process | proteasome regulatory particle, lid subcomplex |
| PF3D7_1310400 | N/A | conserved Plasmodium protein, unknown function | 132.86 | no | 0 | 33.40 | 25.94 | 8.93 | 1.10 | 1.35 | 16.19 | 117.32 | O | 31.13 | 11.61 | M | N/A | N/A | N/A |
| PF3D7_0607700 | N/A | conserved Plasmodium protein, unknown function | 325.01 | no | 0 | 4.96 | 0.25 | 18.52 | 10.55 | 4.38 | 302.97 | 126.05 | GV | 116.32 | 670.10 | F | N/A | N/A | N/A |
| PF3D7_0728000 | eIF2alpha | eukaryotic translation initiation factor 2 subunit alpha | 37.82 | no | 0 | 334.37 | 102.99 | 141.94 | 30.15 | 9.70 | 21.15 | 96.56 | R | 162.16 | 71.68 | M | RNA binding;ribosome binding;translation initiation factor activity | translational initiation | eukaryotic 48S preinitiation complex;eukaryotic translation initiation factor 2 complex;eukaryotic translation initiation factor 2B complex;multi-eIF complex;nucleus |
| PF3D7_0528700 | CYP23 | peptidyl-prolyl cis-trans isomerase | 23.20 | no | 0 | 78.15 | 32.21 | 42.23 | 5.18 | 205.66 | 236.21 | 104.00 | GV | 75.00 | 93.27 | F | cyclosporin A binding;peptidyl-prolyl cis-trans isomerase activity | protein folding;protein peptidyl-prolyl isomerization | nucleus |
| PF3D7_1475000 | MED31 | mediator of RNA polymerase II transcription subunit 31 | 16.61 | no | 0 | 7.30 | 4.62 | 5.12 | 0.39 | 1.42 | 20.92 | 0.00 | GV | 12.76 | 22.07 | F | protein binding | regulation of transcription by RNA polymerase II | core mediator complex;mediator complex;nucleus |
| PF3D7_1419300 | GST | Glutathione S-transferase | 24.79 | no | 0 | 31.76 | 32356.00 | 24624.00 | 45352.00 | 15.28 | 21.68 | 0.00 | S | 41821.00 | 43586.00 | F | glutathione transferase activity | glutathione metabolic process;response to xenobiotic stimulus | cytoplasm |
| PF3D7_1104000 | N/A | phenylalanine--tRNA ligase beta subunit | 72.50 | no | 0 | 50.92 | 21.22 | 13.09 | 2.05 | 2.02 | 11.11 | 12.40 | R | 2.36 | 5.74 | 0 | phenylalanine-tRNA ligase activity | phenylalanyl-tRNA aminoacylation | intracellular;nucleus;phenylalanine-tRNA ligase complex |
| PF3D7_1442300 | tRIP | tRNA import protein tRIP | 46.44 | no | 0 | 111.43 | 41.12 | 30.21 | 3.30 | 3.69 | 20.39 | 42.18 | R | 12.43 | 32.74 | F | RNA binding;aminoacyl-tRNA ligase activity | tRNA aminoacylation for protein translation | cell surface;nucleus;plasma membrane |
| PF3D7_0101300 | MC-2TM | Pfmc-2TM Maurers cleft two transmembrane protein | 27.56 | no | 3 | 13881.00 | 28246.00 | 0.17 | 0.00 | 0.33 | 0.11 | 25051.00 | T | 0.00 | 0.00 | 0 | N/A | translocation of peptides or proteins into host | Maurer's cleft;integral component of membrane;membrane |
| PF3D7_1305900 | N/A | conserved Plasmodium protein, unknown function | 57.92 | no | 0 | 5.63 | 7.72 | 9.65 | 1.77 | 1.81 | 12.75 | 17.62 | O | 28.59 | 23.87 | M | N/A | N/A | N/A |
| PF3D7_1442100 | RPA3 | replication factor A protein 3, putative | 12.61 | no | 0 | 5.70 | 6.24 | 16.13 | 1.98 | 23.62 | 51.00 | 45.38 | GV | 165.38 | 24.30 | M | damaged DNA binding;single-stranded DNA binding | DNA replication;base-excision repair;double-strand break repair via homologous recombination;mismatch repair;nucleotide-excision repair | DNA replication factor A complex;site of double-strand break |
| PF3D7_1357700 | UTP21 | U3 small nucleolar RNA-associated protein 21, putative | 143.36 | no | 0 | 87.49 | 25.91 | 5.22 | 0.22 | 1.44 | 17.50 | 19.58 | R | 22.04 | 20.00 | 0 | RNA binding | mRNA splicing, via splicosome | nucleolus;nucleus |
| PF3D7_0828900 | N/A | conserved protein, unknown function | 21.07 | no | 0 | 6.10 | 0.39 | 13.90 | 7.64 | 6.00 | 50.52 | 137.22 | O | 4.69 | 22.43 | F | N/A | N/A | N/A |
| PF3D7_1126400 | MED21 | mediator of RNA polymerase II transcription subunit 21 | 21.72 | no | 0 | 36.61 | 8.48 | 3.95 | 0.45 | 16.07 | 52.03 | 144.76 | O | 43.52 | 55.74 | F | protein binding | regulation of transcription by RNA polymerase II | core mediator complex;mediator complex;nucleus |
| PF3D7_0207500 | SERA6 | serine repeat antigen 6 | 118.83 | yes | 0 | 12.36 | 1.19 | 245.09 | 91.55 | 3.94 | 1.16 | 61.72 | T | 0.54 | 1.24 | 0 | cysteine-type endopeptidase activity;cysteine-type peptidase activity;protein binding | exit from host cell;immunoglobulin production;proteolysis;proteolysis involved in cellular protein catabolic process;regulation of immune response | extracellular space;lysosome;nucleus;symbiont-containing vacuole |
| PF3D7_0812700 | SNRPC | U1 small nuclear ribonucleoprotein C, putative | 27.88 | no | 0 | 64.39 | 16.92 | 22.12 | 3.78 | 24.51 | 44.96 | 35.79 | R | 23.94 | 9.56 | M | RNA binding;pre-mRNA 5'-splice site binding | mRNA 5'-splice site recognition;mRNA splicing, via spliceosome | U1 snRNP |
| PF3D7_1417100 | N/A | conserved Plasmodium protein, unknown function | 32.34 | no | 0 | 3.90 | 0.62 | 3.28 | 1.31 | 6.72 | 23.72 | 7.54 | GV | 3.65 | 10.64 | F | N/A | N/A | N/A |
| PF3D7_1032700 | N/A | conserved Plasmodium protein, unknown function | 65.54 | no | 0 | 18.42 | 9.37 | 11.47 | 2.46 | 4.30 | 31.28 | 76.93 | 76.93 | 186.51 | 49.94 | M | | | cytoplasm;nucleus;transcription regulator complex;vesicle |
| PF3D7_1003600 | IMC1c | inner membrane complex protein 1c | 32.65 | no | 0 | 8.30 | 3.78 | 48.20 | 38.07 | 4.65 | 251.30 | 1407.69 | O | 24.58 | 953.96 | F | protein binding | structural complex | inner membrane pellicle complex |
| PF3D7_0319100 | RBX1 | E3 ubiquitin-protein ligase RBX1, putative | 12.42 | no | 0 | 20.46 | 16.05 | 27.96 | 6.92 | 72.02 | 202.05 | 119.05 | GV | 27.69 | 183.26 | F | cullin family protein binding;ubiquitin protein ligase activity;ubiquitin-protein transferase activity | protein ubiquitination;ubiquitin-dependent protein catabolic process | SCF ubiquitin ligase complex;cullin-RING ubiquitin ligase complex;nucleus;ubiquitin ligase complex |
| PF3D7_0928300 | N/A | conserved Plasmodium protein, unknown function | 23.39 | no | 0 | 24.88 | 3.42 | 5.00 | 0.38 | 22.47 | 82.25 | 22.01 | GV | 5.10 | 15.33 | F | N/A | N/A | N/A |
| PF3D7_1311400 | AP1M1 | AP-1 complex subunit mu-1 | 50.81 | no | 0 | 91.11 | 39.88 | 48.84 | 20.93 | 9.88 | 27.98 | 35.57 | R | 60.26 | 15.12 | M | protein binding | intracellular protein transport;vesicle-mediated transport | Golgi apparatus;clathrin adaptor complex;clathrin vesicle coat;cytoplasmic vesicle;intracellular membrane-bounded organelle;rhoptry |
| PF3D7_1241900 | N/A | tetratricopeptide repeat protein, putative | 79.80 | no | 1 | 38.95 | 9.97 | 10.79 | 3.77 | 2.63 | 33.84 | 55.70 | O | 116.99 | 81.94 | M | protein binding | N/A | integral component of membrane |
| PF3D7_0616200 | NDC80 | kinetochore protein NDC80 | 69.87 | no | 0 | 8.34 | 9.60 | 36.73 | 18.91 | 25.49 | 30.36 | 438.89 | O | 310.06 | 68.54 | M | microtubule binding | attachment of mitotic spindle microtubules to kinetochore;chromosome segregation | Ndc80 complex;chromosome, centromeric region;condensed nuclear chromosome kinetochore;microtubule;nucleus |
| PF3D7_1137300 | N/A | CLPTM1 domain-containing protein, putative | 81.61 | no | 6 | 62.25 | 33.12 | 26.16 | 2.05 | 11.92 | 52.08 | 64.28 | O | 93.69 | 14.94 | M | N/A | N/A | endomembrane system;membrane |
| PF3D7_1469700 | MED6 | mediator of RNA polymerase II transcription subunit 6 | 24.57 | no | 0 | 3.19 | 2.42 | 7.14 | 1.20 | 17.37 | 183.73 | 34.02 | GV | 38.06 | 63.23 | F | protein binding | regulation of transcription by RNA polymerase II | core mediator complex;mediator complex;nucleus |
| PF3D7_1471200 | SulP | inorganic anion exchanger, inorganic anion antiporter | 75.47 | no | 11 | 18.15 | 19.23 | 97.98 | 25.39 | 8.04 | 19.07 | 511.89 | O | 208.40 | 46.59 | M | transporter activity | response to xenobiotic stimulus | integral component of plasma membrane;plasma membrane |

| | | | | | | | | | | | | | | | | | | | |
|---|---|---|---|---|---|---|---|---|---|---|---|---|---|---|---|---|---|---|---|
| PF3D7_0309200 | ARK2 | aurora-related kinase ARK2, putative | 193.78 | no | 0 | 10.07 | 3.95 | 18.87 | 3.88 | 7.55 | 53.61 | 792.94 | O | 723.17 | 142.91 | M | histone serine kinase activity;protein serine/threonine kinase activity | mitotic spindle organization;protein phosphorylation;regulation of cytokinesis | chromosome passenger complex;kinetochore;mitotic spindle;spindle microtubule;spindle midzone;spindle pole centrosome |
| PF3D7_0627700 | N/A | transportin | 132.84 | no | 0 | 66.34 | 15.77 | 34.44 | 6.78 | 4.25 | 16.27 | 23.98 | R | 37.44 | 16.26 | M | nuclear import signal receptor activity;nuclear localization sequence binding;protein transmembrane transporter activity | protein import into nucleus | apicoplast;cytoplasm;nuclear pore;nucleus |
| PF3D7_1033100 | AdoMetDC/ODC | S-adenosylmethionine decarboxylase/ornithine decarboxylase | 168.17 | no | 0 | 577.00 | 307.54 | 187.19 | 58.86 | 11.95 | 51.45 | 323.43 | R | 110.74 | 299.82 | F | adenosylmethionine decarboxylase activity;ornithine decarboxylase activity | polyamine biosynthetic process;putrescine biosynthetic process from ornithine;putrescine metabolic process | cytoplasm;nucleus |
| PF3D7_0622900 | AP2Tel | AP2 domain transcription factor AP2Tel | 237.36 | no | 0 | 69.50 | 12.07 | 8.39 | 1.68 | 2.51 | 18.98 | 114.12 | O | 34.73 | 15.43 | M | DNA binding;telomeric DNA binding | regulation of transcription, DNA-templated | chromosome, telomeric region;nuclear periphery |
| PF3D7_1309300 | PRPF3 | U4/U6 small nuclear ribonucleoprotein PRP3, putative | 86.49 | no | 0 | 76.30 | 34.22 | 28.94 | 6.41 | 2.74 | 25.25 | 101.22 | O | 156.58 | 97.81 | M | RNA binding | mRNA splicing, via spliceosome | U4/U6 x U5 tri-snRNP complex |
| PF3D7_0411800 | N/A | conserved Plasmodium protein, unknown function | 229.69 | no | 0 | 3.77 | 0.61 | 5.13 | 1.48 | 2.28 | 27.20 | 24.95 | GV | 23.85 | 14.84 | M | N/A | N/A | N/A |
| PF3D7_1211200 | N/A | conserved Plasmodium protein, unknown function | 219.65 | no | 0 | 23.55 | 17.19 | 7.79 | 0.62 | 2.12 | 21.16 | 51.70 | O | 59.94 | 26.79 | M | N/A | N/A | cytoplasm;nucleus |
| PF3D7_1355300 | SET6 | histone-lysine N-methyltransferase, putative | 60.02 | no | 0 | 25.03 | 3.61 | 3.00 | 0.25 | 1.97 | 10.37 | 7.26 | R | 4.63 | 12.20 | F | histone-lysine N-methyltransferase activity | histone lysine methylation | nuclear periphery;nucleus |
| PF3D7_1212000 | TPx(Gl) | glutathione peroxidase-like thioredoxin peroxidase | 23.95 | no | 0 | 6.17 | 4.16 | 22.40 | 4.06 | 29.78 | 51.04 | 18.13 | GV | 22.04 | 14.19 | M | peroxidase activity;thioredoxin peroxidase activity | Peroxidation complex | apicoplast;cytosol;mitochondrion |
| PF3D7_1418000 | UFD1 | ubiquitin fusion degradation protein 1, putative | 32.48 | no | 0 | 42.97 | 19.86 | 50.79 | 20.87 | 37.54 | 38.73 | 38.06 | T | 329.58 | 134.98 | M | polyubiquitin modification-dependent protein binding;ubiquitin binding | ER-associated misfolded protein catabolic process;ubiquitin-dependent ERAD pathway;ubiquitin-dependent protein catabolic process | VCP-NPL4-UFD1 AAA ATPase complex;intracellular;nucleus |
| PF3D7_1361000 | PRMT5 | protein arginine N-methyltransferase 5, putative | 85.55 | no | 0 | 38.47 | 10.13 | 4.47 | 1.83 | 1.59 | 5.94 | 1.40 | R | 14.28 | 3.21 | M | histone-arginine N-methyltransferase activity;protein binding;protein-arginine N-methyltransferase activity | histone arginine methylation;regulation of transcription, DNA-templated | cytosol;nucleus |
| PF3D7_1114900 | N/A | filamin domain-containing protein, putative | 295.38 | no | 0 | 4.67 | 0.22 | 3.98 | 3.01 | 1.05 | 23.03 | 28.62 | O | 6.91 | 52.78 | F | N/A | plastid organization | N/A |
| PF3D7_1335700 | COG3 | conserved oligomeric Golgi complex subunit 3, putative | 193.64 | no | 0 | 7.97 | 6.77 | 13.11 | 3.99 | 2.30 | 15.30 | 28.90 | O | 16.99 | 16.23 | 0 | N/A | intra-Golgi vesicle-mediated transport | cytoplasm |
| PF3D7_0605300 | ARK1 | aurora-related kinase ARK1 | 40.91 | no | 0 | 8.72 | 0.83 | 38.46 | 9.27 | 57.44 | 196.97 | 135.55 | GV | 395.58 | 288.46 | M | kinase activity | schizogony | nucleus;spindle pole body |
| PF3D7_1235300 | NOT4 | CCR4-NOT transcription complex subunit 4, putative | 194.67 | no | 0 | 201.91 | 27.26 | 19.57 | 6.46 | 7.38 | 38.25 | 234.99 | O | 1104.27 | 245.68 | M | mRNA binding;ubiquitin-protein transferase activity | protein ubiquitination | CCR4-NOT complex;nucleus |
| PF3D7_1126300 | N/A | DnaJ protein, putative | 73.53 | no | 0 | 9.93 | 18.34 | 21.58 | 4.86 | 5.74 | 62.87 | 355.02 | O | 22.75 | 35.36 | F | RNA binding;unfolded protein binding | chaperone cofactor-dependent protein refolding;protein refolding | cytoplasm;nucleus |
| PF3D7_1210400 | N/A | general transcription factor 3C polypeptide 5, putative | 107.05 | no | 0 | 26.23 | 6.67 | 7.28 | 0.65 | 1.80 | 8.29 | 59.71 | O | 5.26 | 15.47 | F | DNA binding | regulation of transcription | transcription factor TFIIIC complex;nucleus |
| PF3D7_1450100 | SRP54 | signal recognition particle subunit SRP54 | 55.98 | no | 0 | 366.28 | 59.81 | 53.48 | 29.49 | 4.92 | 15.38 | 78.53 | R | 46.98 | 36.77 | M | 7S RNA binding;endoplasmic reticulum signal peptide binding;protein binding;ribosome binding;signal sequence binding | SRP-dependent cotranslational protein targeting to membrane;SRP-dependent cotranslational protein targeting to membrane, translocation | cytosol;endoplasmic reticulum membrane;signal recognition particle, endoplasmic reticulum targeting |
| PF3D7_1226800 | ATX3 | ataxin-3, putative | 44.98 | no | 0 | 7.87 | 4.23 | 18.27 | 7.63 | 8.88 | 51.13 | 134.24 | O | 125.83 | 121.07 | M | thiol-dependent deubiquitinase | protein deubiquitination | nucleus |
| PF3D7_0918900 | gammaGCS | gamma-glutamylcysteine synthetase | 124.46 | no | 0 | 64.48 | 48.03 | 19.36 | 1.84 | 7.05 | 58.31 | 202.16 | O | 51.19 | 98.44 | F | glutamate-cysteine ligase activity | glutathione biosynthetic process | cytoplasm |
| PF3D7_0106000 | N/A | conserved protein, unknown function | 109.28 | no | 0 | 10.37 | 3.36 | 16.45 | 8.63 | 0.87 | 12.90 | 40.59 | O | 3.23 | 16.36 | F | N/A | N/A | N/A |
| PF3D7_1250900 | N/A | conserved protein, unknown function | 38.89 | no | 0 | 6.80 | 5.57 | 5.60 | 0.68 | 1.70 | 15.55 | 84.26 | O | 18.37 | 32.24 | F | N/A | N/A | N/A |
| PF3D7_0520400 | N/A | single-stranded DNA-binding protein, putative | 35.66 | no | 0 | 3.87 | 4.12 | 10.75 | 2.23 | 2.66 | 14.66 | 3.79 | GV | 2.59 | 15.50 | F | DNA binding | DNA replication | intracellular;nucleus |
| PF3D7_0608900 | N/A | conserved Plasmodium protein, unknown function | 120.67 | no | 3 | 8.77 | 15.56 | 10.70 | 2.95 | 6.53 | 44.72 | 16.31 | GV | 20.37 | 46.23 | F | N/A | N/A | N/A |
| PF3D7_1403900 | CPPED1 | serine/threonine protein phosphatase CPPED1, putative | 35.22 | no | 0 | 12.11 | 26.30 | 10.55 | 1.40 | 21.28 | 101.91 | 71.88 | GV | 9.30 | 35.26 | F | enzyme activity | phosphorylation process | nucleus |
| PF3D7_1209200 | LSM7 | U6 snRNA-associated Sm-like protein LSm7, putative | 12.00 | no | 0 | 68.19 | 29.48 | 26.74 | 6.28 | 106.14 | 38.69 | 0.00 | G II | 22.19 | 19.11 | 0 | RNA binding | RNA splicing, via transesterification reactions;mRNA splicing, via spliceosome | Lsm1-7-Pat1 complex;U12-type spliceosomal complex;U2-type prespliceosome;U6 snRNP;catalytic step 2 spliceosome;nucleus;ribonucleoprotein complex;sno(s)RNA-containing ribonucleoprotein complex;spliceosomal complex;spliceosomal tri-snRNP complex |
| PF3D7_0701600 | MC-2TM | Pfmc-2TM Maurer's cleft two transmembrane protein | 59.79 | no | 3 | 1.65 | 7.68 | 0.26 | 0.00 | 0.00 | 0.40 | 0.00 | T | 0.08 | 0.12 | 0 | N/A | translocation of peptides or proteins into host | Maurer's cleft;membrane |
| PF3D7_0203200 | N/A | conserved Plasmodium protein, unknown function | 28.26 | no | 0 | 17.13 | 4.77 | 3.55 | 0.87 | 2.10 | 54.97 | 51.04 | GV | 80.61 | 114.31 | F | N/A | N/A | N/A |
| PF3D7_1024800 | EXP3 | exported protein 3 | 35.02 | yes | 2 | 518.84 | 211.13 | 110.18 | 12.91 | 68.00 | 158.36 | 694.69 | O | 2065.30 | 619.42 | M | protein binding;protein-containing complex binding | N/A | cytoplasm;symbiont-containing vacuole;symbiont-containing vacuole membrane |
| PF3D7_1301700 | CBP2 | CX3CL1-binding protein 2 | 28.50 | no | 3 | 451.57 | 95.57 | 8.58 | 0.41 | 2.47 | 1.93 | 21.28 | 451.57 | 0.36 | 0.24 | 0 | ATP binding;chemokine binding;nucleic acid binding;protein binding | cytoadherence to microvasculature, mediated by symbiont protein | Maurer's cleft;host cell surface;nucleus |
| PF3D7_1329600 | N/A | conserved Plasmodium protein, unknown function | 270.47 | no | 0 | 102.67 | 19.31 | 17.38 | 1.42 | 21.06 | 97.10 | 52.49 | R | 86.40 | 22.93 | M | N/A | N/A | N/A |
| PF3D7_1135400 | N/A | HotDog domain-containing protein, putative | 33.41 | no | 0 | 124.02 | 12.75 | 21.98 | 5.68 | 6.41 | 25.83 | 0.00 | R | 46.92 | 9.60 | M | RNA binding | N/A | symbiont-containing vacuole |
| PF3D7_1436200 | BCP1 | basal complex protein BCP1 | 271.49 | no | 0 | 6.82 | 0.39 | 8.12 | 9.58 | 0.97 | 6.24 | 35.28 | O | 3.50 | 4.09 | 0 | protein binding | structural complex | basal part of cell;inner membrane pellicle complex |
| PF3D7_1471100 | EXP2 | exported protein 2 | 63.11 | yes | 0 | 384.53 | 1445.82 | 386.76 | 37.45 | 51.11 | 19.77 | 209.28 | T | 15.95 | 7.71 | M | RNA binding;channel activity;protein binding | entry into host;pore formation in membrane of host by symbiont | PTEX complex;extracellular vesicle;host cell cytoplasmic vesicle;merozoite dense granule;nucleus;symbiont-containing vacuole membrane |
| PF3D7_0204100 | N/A | Sel1 repeat-containing protein, putative | 23.99 | yes | 0 | 9.43 | 3.30 | 47.10 | 71.80 | 5.60 | 40.20 | 111.36 | O | 72.03 | 46.20 | M | protein binding | N/A | cytosol |
| PF3D7_1007900 | EIF3D | eukaryotic translation initiation factor 3 subunit D, putative | 244.67 | no | 0 | 149.30 | 37.91 | 38.38 | 3.47 | 4.60 | 16.20 | 49.13 | R | 27.12 | 15.79 | M | translation initiation factor activity | translational initiation | eukaryotic translation initiation factor 3 complex;nucleus |
| PF3D7_1455200 | N/A | methyltransferase AAMT | 70.62 | no | 0 | 3.63 | 4.47 | 9.05 | 1.58 | 8.51 | 32.44 | 25.57 | GV | 146.54 | 38.43 | M | methyltransferase activity | N/A | apical part of cell |
| PF3D7_0609900 | N/A | tetratricopeptide repeat protein, putative | 50.76 | no | 0 | 34.32 | 5.61 | 11.49 | 12.07 | 4.48 | 12.04 | 8.41 | R | 30.53 | 11.04 | M | DNA binding | transcription by RNA polymerase III | transcription factor TFIIIC complex |
| PF3D7_1346500 | N/A | conserved Plasmodium protein, unknown function | 27.68 | no | 0 | 49.88 | 11.80 | 4.62 | 0.41 | 0.96 | 12.62 | 20.44 | R | 54.49 | 32.20 | M | N/A | N/A | N/A |
| PF3D7_1236300 | N/A | conserved protein, unknown function | 70.66 | no | 0 | 2.73 | 0.45 | 4.52 | 1.22 | 3.63 | 18.21 | 64.48 | O | 46.00 | 2.66 | M | endopeptidase activity | N/A | N/A |
| PF3D7_0619900 | SF3A2 | splicing factor 3A subunit 2, putative | 33.07 | no | 0 | 13.49 | 5.61 | 9.37 | 0.89 | 5.28 | 32.01 | 85.16 | O | 15.85 | 24.45 | F | RNA binding | RNA splicing;spliceosomal complex assembly | U2 snRNP;U2-type prespliceosome;catalytic step 2 spliceosome;spliceosomal complex |
| PF3D7_1334500 | MSRP6 | MSP7-like protein | 220.45 | yes | 0 | 5.80 | 15.24 | 11.13 | 1.29 | 0.59 | 0.68 | 6.82 | T | 7.00 | 0.31 | M | protein binding | cell adhesion | Maurer's cleft |
| PF3D7_1455400 | HlyIII | hemolysin III | 120.15 | no | 7 | 45.63 | 16.56 | 19.15 | 2.28 | 36.24 | 76.36 | 50.04 | GV | 41.07 | 75.55 | F | signaling receptor activity | hemolysis in other organism | food vacuole;membrane |
| PF3D7_0703000 | N/A | conserved Plasmodium protein, unknown function | 149.48 | no | 1 | 3.07 | 1.34 | 2.95 | 0.62 | 0.92 | 9.41 | 8.03 | GV | 3.99 | 2.72 | 0 | N/A | N/A | N/A |
| PF3D7_1106700 | DNA2 | DNA replication ATP-dependent helicase/nuclease DNA2, putative | 25.38 | no | 0 | 25.64 | 9.15 | 10.23 | 1.40 | 2.10 | 9.03 | 58.53 | O | 88.77 | 19.55 | M | nuclease activity | DNA-dependent DNA replication | nucleus |
| PF3D7_0306200 | AHA1 | activator of Hsp90 ATPase | 191.38 | no | 0 | 16.95 | 6.92 | 6.77 | 0.37 | 8.56 | 322.52 | 244.72 | GV | 18.11 | 172.53 | F | ATPase activator activity;Hsp90 protein binding | positive regulation of ATP-dependent activity;protein folding | cytosol |
| PF3D7_1020700 | N/A | N-acetyltransferase, GNAT family, putative | 53.67 | no | 0 | 38.87 | 20.19 | 4.93 | 0.37 | 1.27 | 14.24 | 17.16 | R | 14.99 | 21.05 | F | rRNA cytidine N-acetyltransferase activity;tRNA binding | rRNA acetylation involved in maturation of SSU-rRNA | nucleolus;nucleus |
| PF3D7_1149400 | N/A | Plasmodium exported protein, unknown function | 210.71 | yes | 0 | 14.57 | 76.44 | 25.21 | 0.35 | 0.84 | 2.09 | 0.00 | T | 0.14 | 0.07 | 0 | N/A | N/A | host cell |
| PF3D7_0820900 | N/A | conserved Plasmodium protein, unknown function | 323.20 | no | 2 | 18.59 | 2.65 | 9.37 | 4.73 | 2.01 | 17.88 | 16.22 | R | 80.61 | 27.56 | M | N/A | N/A | N/A |
| PF3D7_0210600 | CERLI1 | protein CERLI1 | 17.84 | no | 0 | 7.10 | 1.27 | 31.71 | 27.79 | 6.73 | 197.47 | 409.32 | O | 10.85 | 214.63 | F | lipid binding | entry into host | apical part of cell;rhoptry;rhoptry membrane |
| PF3D7_1435600 | N/A | conserved Plasmodium protein, unknown function | 175.57 | no | 0 | 7.96 | 0.59 | 11.62 | 11.95 | 3.42 | 140.60 | 545.53 | O | 204.39 | 597.76 | F | N/A | N/A | nucleus |
| PF3D7_0615400 | N/A | ribonuclease, putative | 24.26 | no | 1 | 47.62 | 16.97 | 22.89 | 4.16 | 4.65 | 21.91 | 102.84 | O | 232.36 | 91.26 | M | 3'-5'-exoribonuclease activity;ribonuclease activity | RNA processing;mRNA catabolic process | P-body;exosome (RNase complex);nucleus |
| PF3D7_0719000 | N/A | translation machinery-associated protein, putative | 168.93 | no | 0 | 4.30 | 1.88 | 2.28 | 0.00 | 15.78 | 51.42 | 14.04 | GV | 91.69 | 17.03 | M | protein-macromolecule adaptor activity | proteasome regulatory particle assembly | cytoplasm;nucleus |
| PF3D7_0803500 | N/A | AAA family ATPase, putative | 50.81 | no | 1 | 17.13 | 13.10 | 13.20 | 4.13 | 3.22 | 17.93 | 12.54 | GV | 15.02 | 9.33 | M | ATP-dependent peptidase activity | proteolysis | mitochondrion |
| PF3D7_1004500 | AKIT2 | apicomplexan kinetochore protein 2, putative | 111.44 | no | 0 | 3.09 | 0.75 | 5.41 | 2.69 | 4.67 | 17.29 | 20.89 | O | 23.69 | 12.24 | M | DNA binding | DNA-dependent DNA replication | kinetochore;nucleus |
| PF3D7_1454200 | N/A | conserved Plasmodium protein, unknown function | 51.93 | no | 2 | 24.51 | 16.77 | 3.84 | 0.26 | 2.64 | 19.59 | 17.07 | R | 5.31 | 4.30 | 0 | N/A | transcription, DNA-templated | nucleus |
| PF3D7_1031800 | N/A | conserved protein, unknown function | 28.95 | no | 0 | 2.76 | 0.20 | 13.39 | 5.22 | 0.83 | 6.75 | 85.04 | O | 12.19 | 40.80 | F | N/A | N/A | N/A |
| PF3D7_1453900 | N/A | conserved Plasmodium protein, unknown function | 138.86 | no | 0 | 49.43 | 7.80 | 4.62 | 0.66 | 2.77 | 17.11 | 15.94 | R | 21.79 | 22.03 | 0 | N/A | N/A | N/A |
| PF3D7_0826300 | N/A | SPRY domain, putative | 47.58 | no | 0 | 3.77 | 1.93 | 14.07 | 2.25 | 4.19 | 35.85 | 42.08 | O | 43.14 | 27.60 | M | transcription cis-regulatory region binding | histone H3-K4 methylation | Set1C/COMPASS complex;nucleus |
| PF3D7_1370300 | MAHRP1 | membrane associated histidine-rich protein 1 | 70.43 | no | 1 | 3028.12 | 1030.67 | 37.48 | 9.28 | 23.01 | 19.99 | 40.61 | R | 15.22 | 24.85 | F | protein binding | cell adhesion | Maurer's cleft;extracellular vesicle |
| PF3D7_1360800 | FLN | falcilysin | 61.68 | no | 0 | 152.60 | 67.64 | 64.01 | 11.08 | 5.79 | 28.64 | 235.69 | O | 26.33 | 40.96 | F | metalloendopeptidase activity | hemoglobin catabolic process;protein processing | apicoplast;cytoplasm;food vacuole |
| PF3D7_1457000 | SPP | signal peptide peptidase | 54.72 | no | 8 | 126.33 | 90.99 | 157.60 | 41.91 | 43.96 | 300.13 | 241.98 | GV | 93.24 | 503.18 | F | aspartic endopeptidase activity, intramembrane cleaving;peptidase activity;protein homodimerization activity | membrane protein proteolysis;response to xenobiotic stimulus;signal peptide processing | cell surface;endoplasmic reticulum;integral component of cytoplasmic side of endoplasmic reticulum membrane;integral component of lumenal side of endoplasmic reticulum membrane;microneme |
| PF3D7_0623600 | N/A | splicing factor-like protein 1, putative | 220.40 | no | 0 | 120.96 | 42.53 | 58.06 | 21.17 | 20.03 | 15.51 | 113.26 | R | 53.65 | 34.30 | M | RNA binding;mRNA binding | RNA splicing;RNA splicing, via transesterification reactions;mRNA splicing, via spliceosome;regulation of alternative mRNA splicing, via spliceosome | nucleus |
| PF3D7_1121900 | PDK1 | 3-phosphoinositide-dependent protein kinase 1 | 120.38 | no | 0 | 2.72 | 0.23 | 12.55 | 3.49 | 0.47 | 9.70 | 6.95 | T | 6.10 | 16.40 | F | protein kinase activity;protein serine/threonine kinase activity | activation of protein kinase activity;intracellular signal transduction;peptidyl-serine phosphorylation;protein phosphorylation;response to xenobiotic stimulus | cytosol;nucleus |
| PF3D7_1218400 | N/A | triose or hexose phosphate/phosphate translocator | 191.46 | no | 10 | 256.69 | 29.63 | 43.13 | 21.58 | 33.76 | 84.90 | 107.90 | R | 43.84 | 27.43 | M | antiporter activity;transmembrane transporter activity | transporter complex | Golgi apparatus;membrane |
| PF3D7_1450400 | N/A | E3 ubiquitin-protein ligase ZNF598, putative | 165.75 | no | 0 | 45.10 | 15.26 | 12.46 | 4.72 | 1.95 | 17.93 | 47.98 | O | 28.25 | 19.20 | M | ligase activity | protein ubiquitination complex | nucleus |
| PF3D7_1141300 | Apr-01 | apical polar ring protein APR1, putative | 46.02 | no | 0 | 4.09 | 0.38 | 19.89 | 8.76 | 11.23 | 50.97 | 29.19 | GV | 44.67 | 117.88 | F | protein binding | structural complex | apical complex;microneme;plasma membrane |
| PF3D7_1358200 | N/A | conserved Plasmodium protein, unknown function | 76.68 | no | 0 | 31.70 | 10.80 | 9.00 | 1.63 | 1.92 | 17.00 | 12.05 | R | 43.25 | 24.84 | M | N/A | N/A | N/A |
| PF3D7_1344300 | N/A | zinc finger protein, putative | 140.26 | no | 0 | 5.22 | 5.33 | 22.17 | 14.25 | 4.14 | 32.99 | 73.95 | O | 42.20 | 28.75 | M | protein binding | N/A | cytoplasm;nucleus |
| PF3D7_1336800 | NUDC | nuclear movement protein, putative | 282.22 | no | 0 | 51.63 | 28.53 | 12.42 | 0.59 | 3.89 | 19.55 | 40.44 | 51.63 | 86.84 | 45.79 | M | unfolded protein binding | developmental process;protein folding | cytoplasm;nucleus |
| PF3D7_0502100 | N/A | HCNGP-like protein | 36.61 | no | 0 | 3.17 | 0.99 | 2.83 | 1.16 | 3.76 | 17.93 | 16.63 | GV | 10.13 | 24.73 | F | N/A | N/A | nucleus |
| PF3D7_0408500 | FEN1 | flap endonuclease 1 | 63.85 | no | 0 | 9.73 | 1.55 | 51.59 | 26.82 | 10.07 | 9.97 | 121.16 | O | 1072.17 | 59.74 | M | 5'-3' exonuclease activity;5'-flap endonuclease activity;endodeoxyribonuclease activity;flap endonuclease activity | | nucleus |
| PF3D7_1211900 | ATP4 | non-SERCA-type Ca2+ -transporting P-ATPase | 69.68 | no | 8 | 97.79 | 104.68 | 73.60 | 17.69 | 3.79 | 7.62 | 65.62 | T | 27.52 | 32.97 | F | ATP hydrolysis activity;P-type calcium transporter activity;P-type proton-exporting transporter activity | calcium ion transmembrane transport;cellular calcium ion homeostasis;regulation of sodium ion transport;response to xenobiotic stimulus | |
| PF3D7_1008000 | HDA2 | histone deacetylase 2 | 198.28 | no | 0 | 70.23 | 35.07 | 26.53 | 6.58 | 2.90 | 21.36 | 45.15 | R | 150.69 | 35.45 | M | deacetylase activity | inositol phosphate biosynthetic process;regulation of gene expression;chromatin remodeling | nucleus |
| PF3D7_0319500 | N/A | RNA-binding protein, putative | 132.61 | no | 0 | 163.64 | 56.62 | 58.19 | 6.41 | 18.29 | 29.71 | 59.94 | R | 137.53 | 117.33 | M | RNA binding | mRNA splicing, via spliceosome | U2 snRNP;precatalytic spliceosome |
| PF3D7_1242600 | N/A | protein farnesyltransferase subunit alpha | 112.70 | no | 0 | 11.78 | 12.05 | 5.04 | 0.65 | 2.87 | 15.03 | 22.38 | O | 6.06 | 8.04 | 0 | enzyme activity | protein farnesylation;protein geranylgeranylation | CAAX-protein geranylgeranyltransferase complex;cytoplasm;protein farnesyltransferase complex |
| PF3D7_0420600 | AKIT6 | apicomplexan kinetochore protein 6, putative | 419.08 | no | 0 | 5.59 | 2.69 | 19.00 | 10.88 | 6.52 | 62.42 | 0.00 | GV | 210.81 | 82.79 | M | DNA binding | DNA replication | kinetochore;nucleus |
| PF3D7_0627100 | N/A | ankyrin-repeat protein, putative | 16.01 | no | 0 | 120.81 | 40.20 | 29.06 | 7.37 | 4.43 | 45.03 | 74.09 | R | 98.11 | 18.43 | M | N/A | N/A | N/A |
| PF3D7_1211300 | MCM8 | DNA helicase MCM8, putative | 98.54 | no | 0 | 5.73 | 0.50 | 6.88 | 2.18 | 2.52 | 12.78 | 13.22 | O | 32.44 | 36.01 | 0 | ATP binding;ATP-dependent activity, acting on DNA;DNA helicase activity;DNA replication origin binding;single-stranded DNA binding | DNA replication initiation;DNA strand elongation involved in DNA replication | MCM complex;nuclear pre-replicative complex;nucleus |
| PF3D7_0505700 | N/A | conserved Plasmodium membrane protein | 96.86 | yes | 4 | 30.50 | 12.43 | 26.21 | 5.45 | 3.19 | 12.11 | 97.52 | O | 117.64 | 148.99 | F | N/A | N/A | apicoplast |
| PF3D7_1325400 | N/A | CRWN-like protein, putative | 32.78 | no | 0 | 5.49 | 0.34 | 12.48 | 7.52 | 3.74 | 50.96 | 79.13 | O | 80.14 | 103.24 | F | chromatin binding | N/A | heterochromatin;nucleus |
| PF3D7_1123900 | SNU13 | 13 kDa ribonucleoprotein-associated protein, putative | 48.06 | no | 0 | 495.51 | 199.33 | 81.42 | 8.88 | 257.61 | 82.42 | 104.38 | O | 99.29 | 64.75 | M | RNA binding;U3 snoRNA binding;U4 snRNA binding | mRNA splicing, via spliceosome;maturation of LSU-rRNA;maturation of SSU-rRNA | U4/U6 x U5 tri-snRNP complex;box C/D RNP complex;nucleolus;nucleus;precatalytic spliceosome;small-subunit processome |
| PF3D7_1028500 | N/A | partial CSTF domain-containing protein, putative | 76.83 | no | 0 | 56.23 | 18.95 | 12.54 | 3.07 | 2.95 | 53.76 | 186.77 | O | 64.36 | 104.16 | F | N/A | N/A | nucleus |
| PF3D7_0906600 | N/A | zinc finger protein, putative | 14.60 | no | 0 | 383.60 | 154.92 | 24.61 | 2.13 | 4.11 | 32.47 | 160.02 | R | 28.60 | 10.60 | M | mRNA binding | N/A | nucleus |
| PF3D7_1353100 | N/A | Plasmodium exported protein, unknown function | 326.04 | no | 2 | 237.88 | 272.75 | 12.47 | 0.80 | 4.29 | 3.93 | 5.04 | T | 52.53 | 0.90 | M | protein binding | N/A | Maurer's cleft;extracellular vesicle;nucleus |
| PF3D7_0212300 | ERF1 | eukaryotic peptide chain release factor subunit 1, putative | 73.03 | no | 0 | 165.17 | 109.79 | 74.15 | 7.41 | 4.51 | 10.01 | 94.37 | R | 63.49 | 364.13 | F | sequence-specific mRNA binding;translation release factor activity;translation release factor activity, codon specific | cytoplasmic translational termination;regulation of translational termination | cytosol;nucleus;translation release factor complex |
| PF3D7_1438000 | N/A | eukaryotic translation initiation factor eIF2A, putative | 11.00 | no | 0 | 122.85 | 24.59 | 19.10 | 2.61 | 4.51 | 32.68 | 44.27 | R | 57.73 | 44.99 | M | mRNA binding;ribosome binding;tRNA binding;translation initiation factor activity | | cytosolic small ribosomal subunit;nucleus |
| PF3D7_0528000 | UMP1 | proteasome maturation factor UMP1, putative | 41.93 | no | 0 | 18.46 | 9.13 | 12.79 | 2.87 | 19.73 | 142.73 | 50.36 | GV | 38.48 | 175.22 | F | protein binding | proteasome assembly | cytoplasm;nucleus;proteasome complex |
| PF3D7_1414500 | ABCk2 | atypical protein kinase, ABC-1 family, putative | 78.57 | no | 0 | 52.58 | 9.77 | 16.99 | 8.83 | 5.74 | 54.44 | 413.23 | O | 75.31 | 193.62 | F | kinase activity | N/A | N/A |
| PF3D7_0111800 | N/A | eukaryotic translation initiation factor 4E, putative | 51.00 | no | 0 | 11.27 | 13.05 | 11.53 | 1.16 | 2.28 | 14.91 | 40.76 | O | 57.32 | 49.14 | M | RNA 7-methylguanosine cap binding;translation initiation factor activity | | eukaryotic translation initiation factor 4F complex;nucleus |
| PF3D7_1415200 | N/A | DNA-directed RNA polymerases I and III subunit RPAC2 | 202.14 | no | 0 | 66.20 | 17.72 | 4.32 | 0.67 | 34.34 | 29.47 | 0.00 | R | 23.61 | 21.00 | 0 | DNA-directed 5'-3' RNA polymerase activity | transcription, DNA-templated | RNA polymerase I complex;RNA polymerase III complex |
| PF3D7_1445700 | N/A | conserved Plasmodium protein, unknown function | 46.03 | no | 0 | 86.60 | 51.32 | 29.15 | 2.46 | 4.66 | 42.68 | 105.68 | O | 108.27 | 92.40 | M | N/A | N/A | Maurer's cleft;cytosol;inner membrane pellicle complex |
| PF3D7_1366900 | N/A | conserved protein, unknown function | 76.56 | no | 0 | 367.91 | 87.94 | 97.50 | 19.49 | 14.75 | 48.64 | 158.75 | R | 207.94 | 46.16 | M | mRNA binding | N/A | cytoplasm;nucleus |
| PF3D7_0910800 | NBP35 | cytosolic Fe-S cluster assembly factor NBP35, putative | 37.28 | no | 1 | 25.35 | 19.92 | 17.23 | 8.19 | 2.63 | 3.71 | 19.95 | R | 9.35 | 10.45 | 0 | iron-sulfur cluster binding | iron-sulfur cluster assembly | apicoplast;cytosol;mitochondrion |
| PF3D7_0308100 | N/A | zinc finger protein, putative | 100.17 | no | 0 | 6.16 | 0.29 | 11.98 | 10.29 | 1.32 | 5.98 | 147.37 | O | 6.05 | 20.83 | F | protein binding | N/A | cytoplasm;membrane;nucleus |
| PF3D7_1301400 | HYP12 | Plasmodium exported protein (hyp12), unknown function | 26.04 | yes | 1 | 53.54 | 173.95 | 2.54 | 0.13 | 0.30 | 1.43 | 8.88 | T | 2.55 | 1.00 | 0 | N/A | N/A | Maurer's cleft;food vacuole;host cell cytoplasm |
| PF3D7_1354900 | N/A | conserved Plasmodium protein, unknown function | 15.26 | no | 0 | 10.52 | 6.89 | 5.90 | 2.19 | 4.32 | 25.22 | 17.59 | GV | 49.14 | 18.85 | M | N/A | N/A | N/A |
| PF3D7_0716800 | EIF3I | eukaryotic translation initiation factor 3 subunit I, putative | 91.14 | no | 0 | 355.70 | 209.51 | 132.52 | 11.21 | 25.76 | 86.80 | 304.59 | R | 113.70 | 111.58 | 0 | RNA binding;translation initiation factor activity | cytoplasmic translational initiation;regulation of translational initiation;translational initiation | eukaryotic translation initiation factor 3 complex;eukaryotic translation initiation factor 3 complex, eIF3m;nucleus |
| PF3D7_1207100 | N/A | pre-rRNA-processing protein ESF1, putative | 134.57 | no | 0 | 62.49 | 31.83 | 15.73 | 1.73 | 4.28 | 24.31 | 157.61 | O | 307.93 | 85.73 | M | RNA binding | rRNA processing | nucleus |
| PF3D7_0316700 | YOP1 | protein YOP1, putative | 74.44 | no | 3 | 23.93 | 34.97 | 44.22 | 38.80 | 9.61 | 16.32 | 48.41 | O | 43.64 | 6.45 | M | protein binding | endoplasmic reticulum organization;endoplasmic reticulum tubular network organization | cell surface;endoplasmic reticulum |
| PF3D7_1117300 | N/A | conserved protein, unknown function | 53.72 | no | 3 | 119.04 | 6.87 | 25.33 | 10.21 | 14.37 | 79.50 | 10.41 | R | 19.94 | 64.03 | F | N/A | N/A | N/A |
| PF3D7_0623100 | NAB2 | nuclear polyadenylated RNA-binding protein NAB2, putative | 178.07 | no | 0 | 561.35 | 175.35 | 96.22 | 18.52 | 22.36 | 31.62 | 252.45 | R | 247.02 | 118.44 | M | RNA binding;poly(A) binding | negative regulation of mRNA polyadenylation;regulation of mRNA stability | cytoplasm;nuclear periphery;nucleus |
| PF3D7_1364300 | PRP16 | pre-mRNA-splicing factor RNA helicase PRP16 | 124.94 | no | 0 | 81.65 | 22.90 | 9.05 | 2.51 | 1.69 | 18.83 | 17.96 | R | 26.93 | 12.71 | M | RNA helicase activity | regulation of transcription | nucleus |

| | | | | | | | | | | | | | | | | | | |
|---|---|---|---|---|---|---|---|---|---|---|---|---|---|---|---|---|---|---|
| PF3D7_0424500 | FIKK4.1 | serine/threonine protein kinase, FIKK family | 45.99 | no | 1 | 106.28 | 1.15 | 0.68 | 0.00 | 0.22 | 2.79 | 0.87 | R | 5.69 | 0.12 | 0 | protein serine/threonine kinase activity | cytoadherence to microvasculature, mediated by symbiont protein;intracellular signal transduction;modulation by symbiont of host process;protein phosphorylation | Maurer's cleft;cytoplasm;host cell periphery |
| PF3D7_1420000 | SF3B4 | splicing factor 3B subunit 4, putative | 75.16 | no | 0 | 271.19 | 120.67 | 109.97 | 11.56 | 23.14 | 12.97 | 120.37 | R | 129.99 | 133.99 | F | RNA binding | RNA splicing;RNA splicing, via transesterification reactions;mRNA splicing, via spliceosome | U2 snRNP;ribonucleoprotein complex;spliceosomal complex |
| PF3D7_1225700 | N/A | VAC14 domain-containing protein, putative | 245.04 | no | 1 | 51.82 | 14.58 | 11.51 | 0.97 | 2.03 | 7.25 | 9.44 | R | 13.80 | 7.18 | M | protein binding | phosphatidylinositol biosynthetic process;positive regulation of kinase activity | PAS complex;endosome membrane;extrinsic component of vacuolar membrane;vacuolar membrane |
| PF3D7_1228300 | NEK1 | NIMA related kinase 1 | 31.25 | no | 0 | 6.99 | 2.82 | 49.89 | 20.53 | 4.58 | 12.14 | 475.52 | O | 355.09 | 42.46 | M | kinase activity | mitotic cell cycle;protein phosphorylation;response to xenobiotic stimulus | nucleus |
| PF3D7_1314900 | P44 | general transcription factor IIH subunit 2 | 190.94 | no | 0 | 60.35 | 18.63 | 22.33 | 2.70 | 2.10 | 6.00 | 3.44 | R | 137.43 | 10.72 | M | protein binding;DNA binding | nucleotide-excision repair;regulation of transcription by RNA polymerase II | cytoplasm;nucleus;transcription factor TFIIH holo complex |
| PF3D7_0625600 | PAP | poly(A) polymerase PAP, putative | 157.31 | no | 0 | 34.22 | 5.48 | 2.29 | 0.42 | 2.68 | 13.77 | 4.01 | R | 13.68 | 9.61 | M | polynucleotide adenylyltransferase activity | RNA polyadenylation;mRNA polyadenylation | nucleus |
| PF3D7_1133200 | N/A | conserved Plasmodium protein, unknown function | 23.57 | no | 0 | 9.18 | 1.78 | 41.01 | 10.84 | 8.38 | 24.27 | 238.59 | O | 20.21 | 73.52 | F | N/A | N/A | nucleus |
| PF3D7_1340100 | RRP42 | exosome complex component RRP42, putative | 16.54 | no | 0 | 46.63 | 12.00 | 8.09 | 1.07 | 6.51 | 125.61 | 109.72 | GV | 14.16 | 50.47 | F | RNA binding | rRNA catabolic process | exosome (RNase complex) |
| PF3D7_0423600 | N/A | conserved Plasmodium protein, unknown function | 74.37 | no | 0 | 39.89 | 22.44 | 9.79 | 1.99 | 4.85 | 29.82 | 182.06 | O | 94.00 | 58.21 | M | N/A | N/A | N/A |
| PF3D7_0602100 | MTR4 | ATP-dependent RNA helicase MTR4 | 60.19 | no | 0 | 39.50 | 17.73 | 8.33 | 0.29 | 2.59 | 23.23 | 102.05 | O | 42.07 | 32.80 | M | RNA helicase activity;protein binding | RNA catabolic process;maturation of 5.8S rRNA | TRAMP complex;nuclear periphery;nucleus |
| PF3D7_0936200 | N/A | Plasmodium exported protein (hyp11), unknown function | 77.22 | yes | 0 | 12.81 | 29.15 | 3.78 | 0.24 | 23.00 | 197.48 | 13.94 | GV | 8.55 | 17.53 | F | N/A | N/A | Maurer's cleft;host cell |
| PF3D7_1203900 | UBC | ubiquitin-conjugating enzyme E2 | 149.32 | no | 0 | 243.54 | 225.32 | 212.64 | 92.18 | 1228.58 | 1828.85 | 1225.87 | GV | 993.79 | 2149.05 | F | ubiquitin conjugating enzyme activity;ubiquitin-protein transferase activity | cellular protein modification process;protein polyubiquitination | cytoplasm;nucleus |
| PF3D7_1129400 | NSUN3 | RNA cytosine C(5)-methyltransferase, putative | 137.53 | no | 0 | 51.00 | 41.04 | 13.34 | 0.79 | 5.54 | 10.60 | 62.24 | O | 9.49 | 33.55 | F | rRNA (cytosine-C5-)-methyltransferase activity | RNA methylation;maturation of LSU-rRNA;rRNA base methylation | nucleolus;nucleus |
| PF3D7_0532300 | N/A | Plasmodium exported protein (PHISTb), unknown function | 159.36 | no | 0 | 238.76 | 433.92 | 23.72 | 0.46 | 0.89 | 1.19 | 19.20 | T | 13.47 | 0.10 | M | protein binding | N/A | Maurer's cleft;host cell cytoplasm;host cell surface |
| PF3D7_0606500 | PTB | polypyrimidine tract-binding protein, putative | 61.42 | no | 0 | 70.90 | 8.85 | 13.32 | 1.82 | 4.21 | 28.39 | 65.98 | R | 51.31 | 17.20 | M | mRNA binding;poly-pyrimidine tract binding | RNA splicing;regulation of RNA splicing;regulation of translation | nucleus |
| PF3D7_1448000 | UTP12 | U3 small nucleolar RNA-associated protein 12, putative | 69.35 | no | 0 | 63.61 | 16.09 | 5.94 | 0.27 | 1.55 | 14.25 | 14.40 | R | 14.61 | 18.09 | F | snoRNA binding | maturation of SSU-rRNA | Pwp2p-containing subcomplex of 90S preribosome;nucleolus;small-subunit processome |
| PF3D7_1308900 | DCP2 | mRNA-decapping enzyme 2, putative | 10.11 | no | 0 | 181.57 | 78.75 | 30.88 | 5.29 | 3.39 | 21.18 | 472.16 | O | 151.21 | 60.83 | M | hydrolase activity;m7G(5')pppN diphosphatase activity | deadenylation-dependent decapping of nuclear-transcribed mRNA | cytoplasm;nucleus |
| PF3D7_0621100 | N/A | tetratricopeptide repeat protein, putative | 39.14 | no | 0 | 8.32 | 1.56 | 22.23 | 17.58 | 2.55 | 38.08 | 51.04 | O | 7.35 | 84.37 | F | protein binding | N/A | N/A |
| PF3D7_0528200 | EIF3E | eukaryotic translation initiation factor 3 subunit E, putative | 102.00 | no | 2 | 146.91 | 55.97 | 43.09 | 4.03 | 11.95 | 60.74 | 27.87 | R | 32.63 | 19.57 | M | mRNA binding | translational initiation | eukaryotic translation initiation factor 3 complex;nucleus |
| PF3D7_1445600 | N/A | RNA-binding protein, putative | 60.80 | no | 0 | 132.96 | 23.29 | 19.46 | 3.38 | 7.11 | 32.08 | 80.16 | R | 54.83 | 15.09 | M | RNA binding | N/A | nucleus |
| PF3D7_1355800 | SF3B5 | splicing factor 3B subunit 5, putative | 671.11 | no | 0 | 36.43 | 14.21 | 9.89 | 1.42 | 20.39 | 22.03 | 12.41 | R | 52.57 | 8.22 | M | RNA binding | mRNA splicing, via spliceosome;spliceosomal complex assembly | U2 snRNP;nucleus;precatalytic spliceosome |
| PF3D7_1439700 | N/A | leucine carboxyl methyltransferase, putative | 59.69 | no | 0 | 2.23 | 0.56 | 9.97 | 1.80 | 8.09 | 17.56 | 13.33 | GV | 34.90 | 3.81 | M | methyltransferase activity | N/A | N/A |
| PF3D7_1226400 | N/A | regulator of MON1-CCZ1 complex, putative | 38.76 | no | 0 | 42.55 | 22.21 | 26.85 | 6.15 | 9.95 | 57.57 | 97.97 | O | 42.45 | 123.06 | F | protein binding | regulation of autophagy | Mon1-Ccz1 complex;late endosome membrane |
| PF3D7_0217500 | CDPK1 | Calcium-dependent protein kinase 1 | 60.80 | no | 0 | 19906.00 | 0.85 | 60.65 | 51.87 | 31564.00 | 19.97 | 163.66 | GC II | 1341.63 | 638.05 | M | 14-3-3 protein binding;calcium-dependent protein serine/threonine kinase activity;calmodulin binding;protein binding;protein serine/threonine kinase activity | entry into host;exit from host cell;intracellular signal transduction;peptidyl-serine phosphorylation;protein autophosphorylation;protein phosphorylation;response to drug;response to xenobiotic stimulus | cytoplasm;membrane;nucleus;plasma membrane;symbiont-containing vacuolar space;symbiont-containing vacuole;symbiont-containing vacuole membrane |
| PF3D7_0923800 | TRXR | Thioredoxin reductase | 68.70 | no | 0 | 45436.00 | 26785.00 | 117.94 | 62.9 | 45748.00 | 16193.00 | 440.71 | GC II | 69.7 | 135.07 | F | thioredoxin-disulfide reductase activity | cell redox homeostasis | cytoplasm;mitochondrion |
| PF3D7_0410400 | RRP4 | exosome complex component RRP4 | 12.26 | no | 0 | 52.17 | 13.87 | 6.14 | 1.39 | 32.30 | 115.72 | 102.61 | GV | 98.27 | 27.55 | M | 3'-5'-exoribonuclease activity;RNA binding;protein binding | CUT catabolic process;U4 snRNA 3'-end processing;exonucleolytic catabolism of deadenylated mRNA;exonucleolytic trimming to generate mature 3'-end of 5.8S rRNA from tricistronic rRNA transcript (SSU-rRNA, 5.8S rRNA, LSU-rRNA);nuclear polyadenylation-dependent rRNA catabolic process;nuclear polyadenyl | TRAMP complex;cytoplasm;cytoplasmic exosome (RNase complex);exosome (RNase complex);nuclear exosome (RNase complex) |
| PF3D7_1329400 | AMPD | AMP deaminase | 277.69 | no | 0 | 23.91 | 9.22 | 18.66 | 5.93 | 7.10 | 6.27 | 9.86 | R | 23.01 | 40.18 | F | AMP deaminase activity | AMP catabolic process;AMP metabolic process;IMP biosynthetic process;purine nucleotide metabolic process | Maurer's cleft;cytosol |
| PF3D7_1224900 | SF3B6 | splicing factor 3B subunit 6, putative | 52.16 | no | 0 | 34.59 | 19.11 | 12.88 | 2.08 | 80.75 | 148.88 | 138.90 | GV | 128.99 | 78.97 | M | | mRNA splicing, via spliceosome | U12-type spliceosomal complex;U2 snRNP;U2-type spliceosomal complex;catalytic step 2 spliceosome;precatalytic spliceosome |
| PF3D7_1312800 | AAP2 | protein AAP2 | 63.94 | no | 0 | 7.18 | 2.06 | 17.96 | 9.43 | 3.52 | 34.23 | 206.12 | O | 271.11 | 320.34 | F | protein binding | structural complex | apical part of cell |
| PF3D7_1349800 | N/A | GPN-loop GTPase, putative | 212.53 | no | 0 | 19.92 | 3.85 | 2.71 | 0.59 | 2.75 | 18.48 | 6.76 | R | 29.25 | 15.34 | M | ATP binding;GTPase activity | N/A | N/A |
| PF3D7_0910900 | N/A | DNA primase large subunit, putative | 460.08 | no | 0 | 4.22 | 0.71 | 18.95 | 2.64 | 2.37 | 7.12 | 17.90 | T | 158.78 | 17.75 | M | DNA primase activity | DNA replication, synthesis of RNA primer | alpha DNA polymerase:primase complex;nucleus |
| PF3D7_1349300 | TKL3 | tyrosine kinase-like protein | 29.76 | no | 0 | 13.45 | 2.16 | 9.86 | 2.98 | 1.80 | 23.53 | 80.68 | O | 30.24 | 32.49 | 0 | protein kinase activity | protein phosphorylation | cytoplasm;microtubule cytoskeleton |
| PF3D7_0403400 | N/A | zinc finger protein, putative | 100.27 | no | 2 | 18.02 | 7.39 | 7.87 | 1.92 | 2.57 | 17.41 | 47.42 | O | 22.34 | 14.54 | M | protein binding | N/A | cytoplasm;nucleus |
| PF3D7_1124800 | RRS1 | ribosome biogenesis regulatory protein, putative | 62.88 | no | 0 | 122.18 | 36.73 | 16.31 | 2.89 | 17.11 | 117.72 | 331.35 | O | 171.60 | 308.15 | F | | endonucleolytic cleavage in ITS1 to separate SSU-rRNA from 5.8S rRNA and LSU-rRNA from tricistronic rRNA transcript (SSU-rRNA, 5.8S rRNA, LSU-rRNA);ribosomal large subunit biogenesis;ribosomal large subunit export from nucleus | nucleolus;preribosome, large subunit precursor |
| PF3D7_0811000 | CUL1 | cullin-1, putative | 118.33 | no | 0 | 3.06 | 0.27 | 13.37 | 5.72 | 3.63 | 18.00 | 17.24 | GV | 27.67 | 18.79 | M | protein binding;ubiquitin protein ligase binding;ubiquitin-protein transferase activity | SCF-dependent proteasomal ubiquitin-dependent protein catabolic process;protein ubiquitination | SCF ubiquitin ligase complex;cullin-RING ubiquitin ligase complex;nucleus;cytoplasm |
| PF3D7_1123400 | ERF3 | eukaryotic peptide chain release factor GTP-binding subunit, putative | 73.48 | no | 0 | 347.06 | 148.87 | 133.98 | 17.18 | 9.71 | 25.08 | 98.22 | R | 72.39 | 112.45 | F | GTP binding;GTPase activity;translation elongation factor activity;translation release factor activity | cytoplasmic translational termination;translation;translational elongation | eukaryotic translation elongation factor 1 complex;translation release factor complex |
| PF3D7_1130500 | COG2 | conserved oligomeric Golgi complex subunit 2, putative | 51.57 | no | 0 | 6.20 | 2.03 | 12.39 | 3.89 | 2.41 | 16.22 | 90.14 | O | 47.25 | 7.73 | M | | Golgi organization;endoplasmic reticulum to Golgi vesicle-mediated transport;intra-Golgi vesicle-mediated transport | Golgi transport complex |
| PF3D7_1305400 | N/A | AAR2 protein, putative | 75.92 | no | 0 | 34.05 | 6.80 | 7.72 | 0.15 | 2.59 | 33.19 | 71.69 | O | 26.63 | 34.42 | F | N/A | spliceosomal tri-snRNP complex assembly | nucleus |
| PF3D7_0724200 | TAP42 | type 2A phosphatase-associated protein 42, putative | 94.40 | no | 0 | 33.08 | 13.75 | 10.88 | 1.09 | 3.03 | 17.64 | 20.43 | R | 76.54 | 40.89 | M | protein phosphatase 2A binding | regulation of dephosphorylation | cytosol |
| PF3D7_0409400 | DnaJ | chaperone protein DnaJ | 27.96 | no | 0 | 43.71 | 31.24 | 15.11 | 1.16 | 1.55 | 8.61 | 43.05 | O | 74.35 | 30.77 | M | DNA binding;DNA replication origin binding;unfolded protein binding | chaperone cofactor-dependent protein refolding;protein refolding;response to heat | apicoplast;cytoplasm;mitochondrion |
| PF3D7_1120500 | N/A | tRNA nucleotidyltransferase, putative | 97.76 | yes | 0 | 25.12 | 8.93 | 9.11 | 1.17 | 1.78 | 15.61 | 19.21 | 25.12 | 11.17 | 19.20 | F | ATP:3'-cytidine-cytidine-tRNA adenylyltransferase activity;CTP:3'-cytidine-tRNA cytidylyltransferase activity;nucleotidyltransferase activity | tRNA 3'-terminal CCA addition | apicoplast |
| PF3D7_0206800 | MSP2 | Merozoite surface protein 2 | 27.96 | no | 0 | 0.22 | 0.00 | 0.22 | 0.37 | 0.02 | 0.2 | 0.00 | S | 45325.00 | 45445.00 | F | protein binding | protein adhesion | anchored component of plasma membrane;integral component of membrane;integral component of plasma membrane;membrane;nucleus;plasma membrane |
| PF3D7_0926700 | NADSYN | glutamine-dependent NAD(+) synthetase, putative | 107.24 | no | 0 | 23.31 | 13.52 | 15.26 | 3.18 | 0.84 | 6.85 | 180.97 | O | 8.11 | 14.15 | F | NAD+ synthase (glutamine-hydrolyzing) activity;glutaminase activity | NAD biosynthetic process | cytoplasm |
| PF3D7_0821100 | PK1 | protein kinase 1 | 242.55 | no | 0 | 48.05 | 6.52 | 17.08 | 3.47 | 2.15 | 9.98 | 20.98 | R | 9.59 | 14.93 | F | protein serine/threonine kinase activity | signal transduction | cytoplasm;cytosol |
| PF3D7_1362400 | Pcalp | calpain | 419.39 | no | 0 | 39.77 | 10.04 | 13.37 | 3.39 | 3.55 | 57.28 | 54.57 | GV | 68.70 | 171.79 | F | protease activity | signal transduction | cytoplasm;nucleolus |
| PF3D7_1457900 | N/A | conserved Plasmodium protein, unknown function | 154.09 | no | 5 | 12.63 | 4.67 | 9.44 | 3.98 | 2.58 | 17.97 | 10.56 | GV | 51.63 | 47.58 | M | N/A | N/A | N/A |
| PF3D7_0310400 | PIESP1 | parasite-infected erythrocyte surface protein | 153.70 | yes | 0 | 10.99 | 1.36 | 63.26 | 23.05 | 1.46 | 23.74 | 139.59 | O | 8.58 | 123.72 | F | protein binding | structural complex | cell surface;host cell plasma membrane;nucleus;tubovesicular membrane network |
| PF3D7_1123500 | GP2 | golgi protein 2 | 63.62 | yes | 2 | 70.85 | 24.94 | 29.86 | 5.94 | 3.23 | 17.24 | 53.65 | R | 31.21 | 22.83 | M | protein binding;protein-containing complex binding | golgi organization | Golgi apparatus;apicoplast;symbiont-containing vacuole |
| PF3D7_0810300 | PPM5 | protein phosphatase PPM5, putative | 67.79 | no | 0 | 10.73 | 13.05 | 70.65 | 31.75 | 22.62 | 85.93 | 556.33 | O | 804.95 | 255.60 | M | phosphatase activity | phosphorylation process | nucleus |
| PF3D7_0309000 | YVH1 | dual specificity protein phosphatase | 41.72 | no | 0 | 60.42 | 9.65 | 11.08 | 2.10 | 6.34 | 30.80 | 26.81 | R | 51.76 | 16.26 | M | phosphatase activity;protein binding;protein tyrosine/serine/threonine phosphatase activity;structural constituent of nuclear pore;zinc ion binding | negative regulation of nucleocytoplasmic transport;protein dephosphorylation;protein phosphorylation;regulation of cell development | cytoplasm;nucleus |
| PF3D7_1472900 | N/A | dihydroorotase, putative | 36.51 | no | 0 | 14.26 | 5.62 | 13.95 | 7.35 | 124.16 | 33.27 | 122.01 | G II | 14.14 | 18.27 | F | dihydroorotase activity | 'de novo' pyrimidine nucleobase biosynthetic process;pyrimidine nucleotide biosynthetic process | cytoplasm;nucleus |
| PF3D7_1032800 | LRR1 | leucine-rich repeat protein 1 | 39.42 | no | 0 | 3.85 | 1.33 | 8.74 | 2.47 | 5.88 | 36.16 | 30.88 | GV | 11.04 | 10.29 | 0 | protein binding;protein serine/threonine phosphatase inhibitor activity | N/A | protein phosphatase type 1 complex |
| PF3D7_1113300 | UGT | UDP-galactose transporter, putative | 51.69 | no | 8 | 14.05 | 2.35 | 28.57 | 10.78 | 9.66 | 80.44 | 21.35 | GV | 6.61 | 56.51 | F | UDP-galactose transmembrane transporter activity;UDP-glucose transmembrane transporter activity;transmembrane transporter activity | UDP-galactose transmembrane transport;response to xenobiotic stimulus | endoplasmic reticulum;integral component of Golgi membrane;integral component of endoplasmic reticulum membrane;membrane |
| PF3D7_0829400 | N/A | prolyl 4-hydroxylase subunit alpha, putative | 529.36 | no | 0 | 4.40 | 0.13 | 11.76 | 6.22 | 4.20 | 26.62 | 50.01 | O | 240.89 | 51.61 | M | N/A | peptidyl-proline hydroxylation to 4-hydroxy-L-proline | endoplasmic reticulum |
| PF3D7_0703900 | N/A | conserved Plasmodium membrane protein | 129.20 | no | 11 | 66.37 | 25.85 | 3.78 | 0.47 | 1.54 | 25.05 | 51.77 | R | 36.54 | 12.81 | M | WW domain binding | N/A | membrane;nucleus |
| PF3D7_1227800 | ELP3 | elongator complex protein 3, putative | 311.42 | no | 1 | 79.99 | 63.53 | 12.51 | 0.95 | 2.49 | 5.07 | 79.10 | R | 15.93 | 13.07 | 0 | histone acetyltransferase activity | tRNA wobble base 5-methoxycarbonylmethyl-2-thiouridinylation | cytoplasm;elongator holoenzyme complex;mitochondrial outer membrane;mitochondrion;nucleus |
| PF3D7_1207000 | N/A | conserved Plasmodium protein, unknown function | 89.33 | no | 0 | 6.90 | 2.22 | 15.63 | 8.27 | 1.76 | 15.48 | 213.60 | O | 755.85 | 48.10 | M | N/A | N/A | mitochondrion;nucleus |
| PF3D7_0215500 | N/A | conserved Plasmodium protein, unknown function | 159.53 | no | 0 | 28.18 | 12.84 | 6.93 | 1.87 | 0.93 | 7.73 | 27.99 | R | 95.66 | 18.14 | M | N/A | N/A | N/A |
| PF3D7_0714200 | N/A | conserved Plasmodium protein, unknown function | 24.00 | no | 0 | 37.26 | 19.17 | 21.89 | 7.67 | 2.71 | 22.01 | 80.01 | O | 66.94 | 28.44 | M | N/A | N/A | nucleus |
| PF3D7_1472000 | ISY1 | pre-mRNA-splicing factor ISY1, putative | 135.82 | no | 0 | 29.46 | 19.38 | 12.22 | 3.31 | 82.25 | 67.17 | 26.31 | G II | 228.41 | 85.10 | M | RNA binding | generation of catalytic spliceosome for second transesterification step;mRNA 3'-splice site recognition | Prp19 complex;catalytic step 2 spliceosome;post-mRNA release spliceosomal complex;post-spliceosomal complex |
| PF3D7_0923400 | N/A | conserved Plasmodium protein, unknown function | 57.64 | no | 0 | 17.53 | 15.57 | 6.46 | 0.58 | 2.06 | 21.53 | 172.45 | O | 41.16 | 93.18 | F | N/A | N/A | N/A |
| PF3D7_1471400 | N/A | diacylglycerol kinase, putative | 129.05 | no | 0 | 4.28 | 1.04 | 26.01 | 12.51 | 2.74 | 17.93 | 110.87 | O | 68.16 | 25.67 | M | diacylglycerol kinase activity | protein kinase C-activating G protein-coupled receptor signaling pathway;second-messenger-mediated signaling | nucleus |
| PF3D7_0802300 | PWP2 | periodic tryptophan protein 2, putative | 93.75 | no | 0 | 113.16 | 54.11 | 16.96 | 1.74 | 6.04 | 12.09 | 66.54 | R | 45.73 | 39.97 | M | RNA binding | maturation of SSU-rRNA from tricistronic rRNA transcript (SSU-rRNA, 5.8S rRNA, LSU-rRNA);ribosomal small subunit assembly | Pwp2p-containing subcomplex of 90S preribosome;nucleolus;small-subunit processome |
| PF3D7_0914300 | N/A | met-10+ like protein, putative | 27.61 | no | 1 | 34.05 | 12.31 | 7.71 | 0.78 | 2.24 | 11.65 | 10.99 | R | 39.85 | 58.13 | F | tRNA (guanine-N1-)-methyltransferase activity;tRNA methyltransferase activity | tRNA N1-guanine methylation;tRNA methylation | apicoplast;cytoplasm |
| PF3D7_1008900 | AK1 | adenylate kinase | 154.90 | no | 0 | 93.97 | 78.85 | 85.11 | 11.95 | 23.24 | 32.91 | 132.43 | O | 18.20 | 34.44 | F | adenylate kinase activity | generation of precursor metabolites and energy;nucleoside diphosphate metabolic process;purine nucleotide metabolic process | cytoplasm;cytosol;mitochondrion;nucleus |
| PF3D7_1244100 | N/A | N-alpha-acetyltransferase 15, NatA auxiliary subunit, putative | 46.60 | no | 0 | 24.12 | 22.80 | 14.76 | 3.65 | 1.65 | 12.66 | 26.08 | O | 30.19 | 15.45 | M | enzyme activity | N-terminal peptidyl-methionine acetylation | cytoplasm;nucleus |
| PF3D7_1202100 | N/A | SRAP domain-containing protein, putative | 46.67 | no | 0 | 45992.00 | 45604.00 | 22.99 | 24473.00 | 28185.00 | 55.91 | 378.04 | R | 134.92 | 2993.15 | F | single-stranded DNA binding, peptidase activity | cellular response to DNA damage stimulus, protein-DNA covalent cross-linking | N/A |
| PF3D7_1234500 | N/A | conserved Plasmodium protein, unknown function | 46.68 | no | 0 | 29.16 | 5.97 | 5.28 | 0.59 | 2.65 | 23.77 | 15.36 | R | 67.05 | 12.33 | M | N/A | N/A | N/A |
| PF3D7_0619500 | ACS12 | acyl-CoA synthetase | 163.90 | no | 0 | 17.37 | 11.03 | 8.14 | 0.98 | 0.82 | 2.37 | 3.00 | R | 1.58 | 0.57 | 0 | medium-chain fatty acid-CoA ligase activity | fatty acid metabolic process | cytoplasm |
| PF3D7_0530600 | N/A | XAP-5 DNA binding protein, putative | 52.03 | no | 0 | 40.83 | 8.08 | 5.15 | 0.61 | 13.30 | 86.11 | 66.41 | GV | 82.14 | 24.41 | M | DNA binding | chromatin organization | nucleus |
| PF3D7_1364000 | N/A | conserved protein, unknown function | 97.40 | no | 0 | 10.99 | 3.34 | 8.71 | 2.19 | 3.19 | 17.87 | 33.18 | O | 17.80 | 9.14 | M | N/A | N/A | N/A |
| PF3D7_1467500 | N/A | DNA/RNA-binding protein KIN17, putative | 52.46 | no | 0 | 43.88 | 13.86 | 20.36 | 6.75 | 1.42 | 6.50 | 7.67 | R | 21.23 | 13.09 | M | double-stranded DNA binding | DNA replication;cellular response to DNA damage stimulus | nucleus |
| PF3D7_0505200 | ALP2b | actin-like protein, putative | 65.87 | no | 0 | 10.44 | 6.93 | 10.08 | 4.39 | 1.26 | 14.33 | 61.93 | O | 1.63 | 23.44 | F | chromatin binding;structural constituent of cytoskeleton | ATP-dependent chromatin remodeling;cytoskeleton organization;histone H4 acetylation;regulation of transcription by RNA polymerase II | NuA4 histone acetyltransferase complex;SWI/SNF complex;actin cytoskeleton;nucleus |
| PF3D7_1021600 | DPA | deoxyribose-phosphate aldolase, putative | 28.98 | no | 0 | 24.83 | 15.86 | 37.24 | 4.42 | 71.51 | 56.06 | 334.46 | O | 11.42 | 83.12 | F | deoxyribose-phosphate aldolase activity | carbohydrate catabolic process;deoxyribonucleotide catabolic process;pentose-phosphate shunt | intracellular |
| PF3D7_1363200 | PNKP | bifunctional polynucleotide phosphatase/kinase | 55.33 | no | 0 | 12.87 | 3.16 | 3.58 | 0.47 | 1.35 | 6.69 | 1.52 | R | 45.36 | 3.77 | M | damaged DNA binding;double-stranded DNA binding;polydeoxyribonucleotide 5'-hydroxyl-kinase activity;polynucleotide 3'-phosphatase activity | DNA repair;cellular response to DNA damage stimulus;nucleotide phosphorylation | nucleus |
| PF3D7_1315700 | GCD14 | tRNA (adenine(58)-N(1))-methyltransferase catalytic subunit TRM61, putative | 37.20 | no | 0 | 34.31 | 11.04 | 5.22 | 1.65 | 5.22 | 96.27 | 27.32 | GV | 3.81 | 91.57 | F | methyltransferase activity | tRNA methylation | nucleus;tRNA (m1A) methyltransferase complex |
| PF3D7_1448100 | N/A | conserved Plasmodium protein, unknown function | 23.44 | no | 0 | 10.42 | 2.91 | 2.09 | 0.27 | 6.93 | 27.17 | 6.25 | GV | 24.30 | 9.68 | M | N/A | N/A | N/A |
