## Supplementary material for "*Plasmodium falciparum* SET10 is a histone H3 lysine K18 methyltransferase that participates in a chromatin modulation network crucial for intraerythrocytic development": Table S4

Table S4. Selected groups of *Pf* SET10 interactors, sorted by peptide count ≥ 50 or by functions in transcription and DNA replication

### Peptide count 50+

| | | | | | | Expression | | | | | | | | Sex | | | GO Terms | | |
|---|---|---|---|---|---|---|---|---|---|---|---|---|---|---|---|---|---|---|---|
| Gene ID | Short name | Product Description | Mol. weight [kDa] | Signal Peptide | # TM Domains | Ring | Early Trophozoite | Late Trophozoite | Schizont | Gametocyte II | Gametocyte V | Ookinete | Peak expression | male gametocyte | female gametocyte | Sex specificity | Curated GO Functions (Molecular) | Curated GO Processes (Biological) | Curated GO Components (Cellular) |
| PF3D7_1233600 | AARP1 | asparagine and aspartate rich protein 1 | 646.33 | no | 0 | 50.78 | 15.99 | 24.42 | 3.49 | 10.84 | 46.00 | 181.42 | O | 125.79 | 43.18 | M | zinc ion binding | N/A | plasma membrane |
| PF3D7_0613800 | ApiAP2 | AP2 domain transcription factor, putative | 485.58 | no | 0 | 9.78 | 1.34 | 89.95 | 46.14 | 11.78 | 17.07 | 97.31 | O | 85.41 | 6.69 | M | sequence-specific DNA binding, DNA-binding transcription factor activity, DNA binding | response to xenobiotic stimulus, regulation of transcription, DNA-templated | nucleus |
| PF3D7_1468100 | MORC | MORC family protein | 295.83 | no | 0 | 170.77 | 129.57 | 91.30 | 24.26 | 7.88 | 27.78 | 110.92 | R | 155.68 | 84.96 | M | protein binding | regulation of cell cycle | nucleus |
| PF3D7_0704600 | UT | HECT-type E3 ubiquitin ligase UT | 460.41 | no | 0 | 134.30 | 57.58 | 41.68 | 13.08 | 14.52 | 61.20 | 373.27 | O | 561.82 | 158.74 | M | ubiquitin-protein transferase activity | response to drug | endoplasmic reticulum, Golgi apparatus |
| PF3D7_0910200 | N/A | conserved Plasmodium protein, unknown function | 368.34 | no | 0 | 18.62 | 6.72 | 13.52 | 4.05 | 3.60 | 11.87 | 40.91 | O | 55.00 | 33.69 | M | N/A | N/A | nucleus |
| PF3D7_1008100 | PHD1 | PHD finger protein PHD1 | 453.00 | no | 0 | 86.17 | 53.30 | 21.26 | 8.32 | 3.22 | 17.25 | 68.53 | R | 35.99 | 17.39 | M | methylated histone binding, metal ion binding | entry into host, histone H3-K9 acetylation, histone H3-K4 trimethylation | histone acetyltransferase complex, nucleus |
| PF3D7_0317300 | NUP335 | nucleoporin NUP335, putative | 402.95 | no | 0 | 53.47 | 39.23 | 25.60 | 8.19 | 6.84 | 23.42 | 78.67 | O | 166.65 | 27.89 | M | DNA-binding transcription factor activity, protein binding, sequence-specific DNA binding | protein autoprocessing, positive regulation of transcription, DNA-templated | nucleus, cytoplasm, endoplasmic reticulum membrane, nuclear periphery, membrane |
| PF3D7_1221000 | SET10 | histone-lysine N-methyltransferase, H3 lysine-4 specific | 271.07 | no | 0 | 5.25 | 3.05 | 21.93 | 5.72 | 2.16 | 12.63 | 50.60 | O | 9.49 | 12.36 | 0 | histone methyltransferase activity (H3-K4 specific), metal ion binding | chromatin remodelling;regulation of transcription | nuclear periphery, nucleus |
| PF3D7_0915400 | PFK9 | ATP-dependent 6-phosphofructokinase | 159.45 | no | 0 | 431.40 | 123.27 | 115.68 | 12.05 | 4.03 | 6.74 | 60.44 | R | 112.38 | 18.75 | M | 6-phosphofructokinase activity; ATP binding, phosphofructokinase activity, metal ion binding, diphosphate-fructose-6-phosphate 1-phosphotransferase activity | response to glucose, response to xenobiotic stimulus, glycolytic process, fructose 6-phosphate metabolic process | cytoplasm, cytosol, 6-phosphofructokinase complex |
| PF3D7_0405400 | PRPF8 | pre-mRNA-processing-splicing factor 8, putative | 366.39 | no | 0 | 156.78 | 102.89 | 99.78 | 16.79 | 6.56 | 15.84 | 402.18 | O | 85.17 | 40.86 | M | U1 snRNA binding, U6 snRNA binding, U2 snRNA binding, U5 snRNA binding, U1 snRNA binding, U6 snRNA binding, pre-mRNA intronic binding | spliceosomal tri-snRNP complex assembly, RNA splicing, via transesterification reactions, RNA splicing | nucleus, cytoplasm, spliceosomal complex, U5 snRNP, apicoplast |
| PF3D7_0317700 | N/A | CPSF (cleavage and polyadenylation specific factor), subunit A, putative | 338.53 | no | 0 | 35.07 | 13.47 | 13.36 | 2.84 | 5.65 | 38.85 | 85.22 | O | 166.90 | 117.96 | M | damaged DNA binding; nucleic acid binding; protein binding | nucleotide-excision repair; DNA repair; proteasome-mediated ubiquitin-dependent protein catabolic process | nucleus; cullin-RING ubiquitin ligase complex |
| PF3D7_0609000 | NUP637 | nucleoporin NUP637, putative | 720.51 | no | 0 | 86.14 | 62.26 | 38.45 | 12.58 | 8.74 | 47.35 | 51.50 | R | 79.23 | 33.93 | M | protein binding | nucleocytoplasmic transport | nuclear periphery; nucleus |
| PF3D7_1239200 | ApiAP2 | AP2 domain transcription factor, putative | 299.38 | no | 0 | 16650.00 | 28095.00 | 61.7 | 45480.00 | 34029.00 | 28.91 | 223.22 | S | 69.00 | 39.57 | M | DNA binding, DNA-binding transcription factor activity | regulation of transcription, DNA-templated, | heterochromatin, nucleus |
| PF3D7_0605800 | RAD50 | DNA repair protein RAD50, putative | 267.95 | no | 0 | 20.12 | 6.26 | 12.65 | 2.62 | 1.82 | 9.21 | 35.80 | O | 19.13 | 14.92 | M | G-quadruplex DNA binding; metal ion binding; single-stranded telomeric DNA binding adenylate kinase activity; double-stranded telomeric DNA binding; single-stranded DNA endodeoxyribonuclease activity | telomere maintenance via recombination; double-strand break repair; telomere maintenance via telomerase; DNA duplex unwinding; chromosome organization involved in meiotic cell cycle | condensed nuclear chromosome; nucleus; Mre11 complex; site of double-strand break |
| PF3D7_0629700 | SET1 | SET domain protein, putative | 796.04 | no | 0 | 45.67 | 31.36 | 25.30 | 4.30 | 3.13 | 23.45 | 102.36 | O | 82.65 | 38.44 | M | histone methyltransferase activity (H3-K4 specific) | positive regulation of transcription, DNA-templated, histone H3-K4 methylation | nucleus;histone methyltransferase complex |
| PF3D7_0922800 | N/A | conserved Plasmodium protein, unknown function | 516.45 | no | 2 | 10.86 | 3.03 | 7.97 | 0.99 | 3.65 | 34.85 | 35.96 | O | 61.71 | 21.16 | M | protein kinase binding | signal transduction | integral component of membrane |
| PF3D7_1252100 | RON3 | rhoptry neck protein 3 | 263.15 | yes | 3 | 5.87 | 0.67 | 56.25 | 101.19 | 3.59 | 1.98 | 98.22 | S | 1.39 | 0.38 | 0 | RNA binding, host cell surface binding; protein binding | entry into host;symbiont intracellular protein transport in host | rhoptry; extracellular vesicle |
| PF3D7_0628100 | HECT1 | HECT domain-containing protein 1, putative | 1206.00 | no | 2 | 8.24 | 0.65 | 21.71 | 36.80 | 2.47 | 28.22 | 948.15 | O | 8.36 | 58.58 | F | ubiquitin protein ligase activity, protein binding, ubiquitin-protein transferase activity | N/A | integral component of membrane |
| PF3D7_1219100 | CHC | clathrin heavy chain, putative | 232.95 | no | 0 | 39.47 | 9.25 | 27.53 | 6.29 | 1.92 | 28.74 | 51.03 | O | 14.72 | 23.77 | F | clathrin light chain binding, structural molecule activity, protein binding | receptor-mediated endocytosis, intracellular protein transport, vesicle-mediated transport | clathrin complex; extracellular vesicle, clathrin coat of trans-Golgi network vesicle |
| PF3D7_1472200 | HDA1 | histone deacetylase, putative | 268.92 | no | 3 | 127.77 | 27.82 | 14.49 | 4.13 | 5.76 | 46.66 | 129.46 | O | 122.03 | 38.33 | M | NAD-dependent histone deacetylase activity (H3-K14 specific) | chromatin remodelling;regulation of transcription | nucleus, integral component of membrane |
| PF3D7_1303800 | N/A | conserved Plasmodium protein, unknown function | 1111.80 | no | 2 | 5.85 | 3.71 | 11.23 | 1.84 | 2.07 | 33.23 | 81.14 | O | 109.15 | 24.81 | M | N/A | N/A | nucleus, integral component of membrane |
| PF3D7_0516800 | ApiAP2 | AP2 domain transcription factor AP2-O2, putative | 276.42 | no | 0 | 12.31 | 1.70 | 43.02 | 9.63 | 6.21 | 80.08 | 288.62 | O | 122.19 | 204.96 | F | sequence-specific DNA binding, DNA-binding transcription factor activity, DNA binding | regulation of transcription, DNA-templated | heterochromatin, nucleus |
| PF3D7_1434500 | N/A | dynein-related AAA-type ATPase, putative | 971.00 | no | 9 | 46.73 | 45.83 | 7.08 | 0.54 | 1.98 | 20.47 | 18.71 | R | 64.58 | 16.90 | M | ATP binding; ATP hydrolysis activity | rRNA processing; ribosomal large subunit assembly | nucleus; membrane; integral component of membrane; preribosome, large subunit precursor |
| PF3D7_1308400 | N/A | conserved Plasmodium protein, unknown function | 737.69 | no | 3 | 15.92 | 18.24 | 15.04 | 3.71 | 2.80 | 16.91 | 31.38 | O | 30.75 | 13.91 | M | N/A | N/A | N/A |
| PF3D7_0726500 | N/A | ubiquitin carboxyl-terminal hydrolase, putative | 373.16 | no | 0 | 76.66 | 27.76 | 41.70 | 8.34 | 12.86 | 90.14 | 137.99 | O | 112.68 | 49.59 | M | cysteine-type endopeptidase activity; thiol-dependent deubiquitinase; protein binding | protein deubiquitination; ubiquitin-dependent protein catabolic process | nucleus; cytosol |
| PF3D7_1433400 | PHD2 | PHD finger protein PHD2, putative | 682.45 | no | 12 | 16.22 | 42.77 | 44.79 | 13.13 | 3.44 | 35.46 | 76.44 | O | 157.85 | 59.14 | M | metal ion binding | N/A | histone acetyltransferase complex; nucleus; membrane; integral component of membrane |
| PF3D7_0422500 | BRR2 | pre-mRNA-splicing helicase BRR2, putative | 337.90 | no | 0 | 105.44 | 72.81 | 27.09 | 4.35 | 8.08 | 18.99 | 136.41 | O | 92.14 | 36.50 | M | RNA helicase activity; nucleic acid binding; mRNA binding | regulation of translation; RNA splicing, via transesterification reactions | spliceosomal complex; cytoplasmic stress granule;nucleus |
| PF3D7_1410300 | N/A | WD repeat-containing protein, putative | 526.73 | no | 0 | 26.36 | 4.05 | 9.03 | 1.22 | 3.16 | 22.31 | 15.23 | R | 128.48 | 24.03 | M | U3 snoRNA binding | endonucleolytic cleavage in 5'-ETS of tricistronic rRNA transcript (SSU-rRNA, 5.8S rRNA, LSU-rRNA) | nucleus;cytoplasm |
| PF3D7_0826100 | HEUL | HECT-like E3 ubiquitin ligase, putative | 1005.10 | no | 2 | 99.48 | 75.67 | 24.99 | 3.25 | 6.02 | 43.36 | 68.66 | R | 119.77 | 52.32 | M | ubiquitin protein ligase activity; ubiquitin-protein transferase activity | proteasome-mediated ubiquitin-dependent protein catabolic process; positive regulation of protein catabolic process; protein ubiquitination; ubiquitin-dependent protein catabolic process | cytoplasm; integral component of membrane |
| PF3D7_0710200 | N/A | conserved Plasmodium protein, unknown function | 350.70 | no | 0 | 8.14 | 11.06 | 10.76 | 10.36 | 4.34 | 56.98 | 330.97 | O | 3852.79 | 1172.20 | M | N/A | N/A | nucleus |

nuclear localization
with signal peptide

### Transcription

| | | | | | | Expression | | | | | | | | Sex | | | GO Terms | | |
|---|---|---|---|---|---|---|---|---|---|---|---|---|---|---|---|---|---|---|---|
| Gene ID | Short name | Product Description | Mol. weight [kDa] | Signal Peptide | # TM Domains | Ring | Early Trophozoite | Late Trophozoite | Schizont | Gametocyte II | Gametocyte V | Ookinete | Peak expression | male gametocyte | female gametocyte | Sex specificity | Curated GO Functions (Molecular) | Curated GO Processes (Biological) | Curated GO Components (Cellular) |
| **AP2 transcription factors** | | | | | | | | | | | | | | | | | | | |
| PF3D7_0420300 | ApiAP2 | AP2 domain transcription factor, putative | 400.16 | no | 0 | 115.68 | 92.16 | 57.10 | 28.79 | 4.65 | 20.37 | 743.23 | O | 51.70 | 52.09 | 0 | protein binding;sequence-specific DNA binding | regulation of transcription | nucleus |
| PF3D7_0516800 | ApiAP2-O2 | AP2 domain transcription factor AP2-O2, putative | 276.42 | no | 0 | 12.31 | 1.70 | 43.02 | 9.63 | 6.21 | 80.08 | 288.62 | O | 122.19 | 204.96 | F | sequence-specific DNA binding, DNA-binding transcription factor activity, DNA binding | ? regulation of transcription, DNA-templated | heterochromatin, nucleus |
| PF3D7_0604100 | SIP2 | AP2 domain transcription factor | 229.62 | no | 0 | 5.73 | 0.57 | 29.83 | 15.16 | 6.58 | 5.73 | 40.37 | O | 45.16 | 1.87 | M | sequence-specific DNA binding;telomeric DNA binding | N/A | nuclear periphery;nucleus |
| PF3D7_0613800 | ApiAP2 | AP2 domain transcription factor, putative | 485.58 | no | 0 | 9.78 | 1.34 | 89.95 | 46.14 | 11.78 | 17.07 | 97.31 | O | 85.41 | 6.69 | M | sequence-specific DNA binding, DNA-binding transcription factor activity, DNA binding | response to xenobiotic stimulus, regulation of transcription, DNA-templated | nucleus |
| PF3D7_0622900 | AP2Tel | AP2 domain transcription factor AP2Tel | 237.36 | no | 0 | 69.50 | 12.07 | 8.39 | 1.68 | 2.51 | 18.98 | 114.12 | O | 34.73 | 15.43 | M | DNA binding;telomeric DNA binding | regulation of transcription, DNA-templated | chromosome, telomeric region;nuclear periphery |
| PF3D7_0730300 | ApiAP2-L | AP2 domain transcription factor AP2-L, putative | 155.13 | no | 0 | 227.37 | 137.47 | 40.04 | 3.01 | 1.76 | 9.32 | 83.57 | R | 53.53 | 9.26 | M | DNA binding | regulation of transcription | nucleus |
| PF3D7_0802100 | AP2-LT | AP2 domain transcription factor AP2-LT | 299.29 | no | 0 | 7.72 | 1.64 | 39.37 | 26.49 | 7.16 | 32.48 | 220.82 | O | 142.23 | 54.00 | M | sequence-specific DNA binding | regulation of transcription | histone acetyltransferase complex;nucleus |
| PF3D7_1007700 | AP2-I | AP2 domain transcription factor AP2-I | 182.66 | no | 0 | 35.10 | 204.32 | 285.73 | 99.61 | 2.49 | 14.17 | 189.67 | T | 299.60 | 65.11 | M | protein binding;sequence-specific DNA binding | response to xenobiotic stimulus | Maurer's cleft;nucleus;protein-containing complex |
| PF3D7_1107800 | ApiAP2 | AP2 domain transcription factor, putative | 206.87 | no | 0 | 39.31 | 93.68 | 44.62 | 35.76 | 10.53 | 72.68 | 200.31 | O | 29.60 | 146.15 | F | sequence-specific DNA binding; DNA-binding transcription factor activity; DNA binding | regulation of transcription, DNA-templated | heterochromatin; nucleus |
| PF3D7_1139300 | AP2-G5 | AP2 domain transcription factor AP2-G5 | 309.44 | no | 0 | 64.79 | 41.42 | 12.07 | 0.88 | 1.99 | 22.23 | 190.04 | O | 120.04 | 21.26 | M | DNA-binding transcription factor activity;protein binding;sequence-specific DNA binding | regulation of transcription, DNA-templated | heterochromatin;nucleus |
| PF3D7_1222600 | AP2-G | AP2 domain transcription factor AP2-G | 284.06 | no | 0 | 6.31 | 0.44 | 0.95 | 0.04 | 0.84 | 2.23 | 2.59 | R | 0.16 | 0.17 | 0 | DNA-binding transcription activator activity;protein binding;sequence-specific DNA binding | response to xenobiotic stimulus | nucleus |
| PF3D7_1239200 | ApiAP2 | AP2 domain transcription factor, putative | 299.38 | no | 0 | 16650.00 | 28095.00 | 61.7 | 45480.00 | 34029.00 | 28.91 | 223.22 | S | 69.00 | 39.57 | M | DNA binding, DNA-binding transcription factor activity | regulation of transcription, DNA-templated, | heterochromatin, nucleus |
| PF3D7_1449500 | ApiAP2-O5 | AP2 domain transcription factor AP2-O5, putative | 84.43 | no | 0 | 12.33 | 6.91 | 15.43 | 2.38 | 4.29 | 27.87 | 41.68 | O | 77.68 | 62.28 | M | DNA-binding transcription factor activity | regulation of transcription, DNA-templated;cell motility | heterochromatin;nucleus |
| PF3D7_1456000 | AP2-HC | AP2 domain transcription factor AP2-HC | 161.51 | no | 0 | 4.11 | 0.68 | 15.05 | 3.24 | 2.56 | 11.86 | 30.98 | O | 82.54 | 8.94 | M | protein binding;sequence-specific DNA binding | regulation of transcription | heterochromatin;nucleus |
| PF3D7_1466400 | ApiAP2-EXP | AP2 domain transcription factor AP2-EXP | 92.09 | no | 0 | 200.36 | 29.73 | 20.00 | 2.82 | 3.59 | 14.47 | 15.52 | R | 64.32 | 9.47 | M | sequence-specific DNA binding | positive regulation of gene expression;response to xenobiotic stimulus | heterochromatin;nucleus |
| **Trancription factors, other** | | | | | | | | | | | | | | | | | | | |
| PF3D7_0522200 | TAF10 | transcription initiation factor TFIID subunit 10, putative | 13.63 | no | 0 | 34.26 | 4.30 | 6.39 | 1.51 | 26.85 | 70.37 | 13.85 | GV | 17.25 | 11.92 | M | protein binding | transcription by RNA polymerase II | transcription regulator complex |
| PF3D7_1014600 | ADA2 | transcriptional coactivator ADA2 | 300.25 | no | 1 | 176.38 | 156.32 | 115.15 | 33.94 | 6.81 | 17.88 | 125.06 | R | 123.81 | 63.55 | M | DNA-binding transcription factor activity;chromatin binding;protein binding;transcription coactivator activity | chromatin remodeling;positive regulation of histone acetylation;regulation of transcription by RNA polymerase II;regulation of transcription, DNA-templated | histone acetyltransferase complex;nucleus |
| PF3D7_1210400 | N/A | general transcription factor 3C polypeptide 5, putative | 107.05 | no | 0 | 26.23 | 6.67 | 7.28 | 0.65 | 1.80 | 8.29 | 59.71 | O | 5.26 | 15.47 | F | DNA binding | regulation of transcription | transcription factor TFIIIC complex;nucleus |
| PF3D7_1244200 | TFB2 | RNA polymerase II transcription factor B subunit 2, putative | 111.87 | no | 0 | 73.96 | 19.46 | 11.81 | 1.23 | 2.67 | 21.02 | 38.76 | R | 14.92 | 18.47 | F | double-stranded DNA binding | nucleotide-excision repair;phosphorylation of RNA polymerase II C-terminal domain;transcription by RNA polymerase II | nucleus;transcription factor TFIIH core complex;transcription factor TFIIH holo complex |
| PF3D7_1314900 | P44 | general transcription factor IIH subunit 2 | 190.94 | no | 0 | 60.35 | 18.63 | 22.33 | 2.70 | 2.10 | 6.00 | 3.44 | R | 137.43 | 10.72 | M | protein binding;DNA binding | nucleotide-excision repair;regulation of transcription by RNA polymerase II | cytoplasm;nucleus;transcription factor TFIIH holo complex |
| PF3D7_1315800 | MYB1 | transcription factor MYB1 | 49.99 | no | 0 | 25.21 | 22.70 | 15.37 | 2.90 | 3.82 | 53.30 | 38.97 | GV | 40.54 | 16.77 | M | DNA-binding transcription factor activity | positive regulation of transcription, DNA-templated;regulation of mitotic cell cycle;regulation of transcription, DNA-templated | nucleus |
| PF3D7_1426100 | BTF3 | transcription factor BTF3, putative | 19.39 | no | 0 | 87.26 | 83.37 | 30.28 | 1.86 | 25.60 | 91.62 | 65.00 | GV | 45.91 | 56.56 | F | DNA binding | regulation of transcription | cytosol;nascent polypeptide-associated complex;nucleus;polysomal ribosome;transcription regulator complex |
| PF3D7_1428800 | N/A | Transcription initiation TFIID-like, putative | 41.98 | no | 1 | 42583.00 | 43556.00 | 30072.00 | 35431.00 | 15281.00 | 107.79 | 45319.00 | O | 24.15 | 67.6 | F | RNA polymerase II general transcription initiation factor activity | DNA-templated transcription, initiation | nucleus |
| **Epigenetic regulators** | | | | | | | | | | | | | | | | | | | |
| PF3D7_0110500 | BDP3 | bromodomain protein 3, putative | 268.20 | no | 0 | 20.61 | 32.68 | 12.96 | 1.68 | 9.50 | 34.61 | 111.42 | O | 99.22 | 48.99 | M | N/A | histone acetylation | NuA4 histone acetyltransferase complex;nucleus |
| PF3D7_0416400 | HAT1 | histone acetyltransferase, putative | 146.72 | no | 0 | 23.91 | 32.37 | 90.95 | 27.34 | 6.12 | 10.98 | 652.73 | O | 6.29 | 71.97 | F | H4 histone acetyltransferase activity | histone H4 acetylation | nucleus |
| PF3D7_0628600 | N/A | DNA methyltransferase 1-associated protein 1, putative | 46.56 | no | 0 | 32.66 | 16.54 | 11.66 | 1.03 | 7.50 | 38.86 | 101.09 | O | 29.91 | 20.19 | M | transcription corepressor activity | histone H2A acetylation;histone H4 acetylation;histone exchange;negative regulation of transcription by RNA polymerase II | NuA4 histone acetyltransferase complex;Swr1 complex;nucleus |
| PF3D7_0629700 | SET1 | SET domain protein, putative | 796.04 | no | 0 | 45.67 | 31.36 | 25.30 | 4.30 | 3.13 | 23.45 | 102.36 | O | 82.65 | 38.44 | M | histone methyltransferase activity (H3-K4 specific) | positive regulation of transcription, DNA-templated, histone H3-K4 methylation | nucleus;histone methyltransferase complex |
| PF3D7_0724700 | BDP6 | bromodomain protein 6, putative | 420.67 | no | 0 | 42.01 | 19.54 | 10.52 | 2.44 | 4.24 | 19.55 | 32.71 | R | 24.66 | 18.50 | M | protein binding | regulation of transcription | nucleus;transcription regulator complex |
| PF3D7_0823300 | GCN5 | histone acetyltransferase GCN5 | 170.92 | no | 0 | 138.06 | 181.90 | 71.62 | 12.74 | 7.97 | 24.22 | 59.65 | T | 59.27 | 31.03 | M | H3 histone acetyltransferase activity | chromatin remodeling | nucleus |
| PF3D7_0827800 | SET3 | SET domain protein, putative | 283.55 | no | 0 | 64.54 | 10.91 | 23.63 | 8.84 | 6.71 | 49.68 | 167.97 | O | 119.89 | 325.00 | F | protein binding; histone-lysine N-methyltransferase activity | chromatin remodelling;regulation of transcription | nuclear periphery; ? nucleus |
| PF3D7_0925700 | HDAC1 | histone deacetylase 1 | 51.38 | no | 0 | 156.78 | 62.14 | 85.17 | 62.49 | 20.60 | 21.37 | 76.19 | R | 89.71 | 34.53 | M | histone deacetylase activity;protein binding;sphingolipid binding | cellular response to drug;histone deacetylation;response to xenobiotic stimulus | nucleus |
| PF3D7_1008000 | HDA2 | histone deacetylase 2 | 198.28 | no | 0 | 70.23 | 35.07 | 26.53 | 6.58 | 2.90 | 21.36 | 45.15 | R | 150.69 | 35.45 | M | deacetylase activity | inositol phosphate biosynthetic process;regulation of gene expression;chromatin remodeling | nucleus |
| PF3D7_1008100 | PHD1 | PHD finger protein PHD1 | 453.00 | no | 0 | 86.17 | 53.30 | 21.26 | 8.32 | 3.22 | 17.25 | 68.53 | R | 35.99 | 17.39 | M | methylated histone binding, ? metal ion binding | entry into host, histone H3-K9 acetylation, histone H3-K4 trimethylation | ? histone acetyltransferase complex, nucleus |
| PF3D7_1033700 | BDP1 | bromodomain protein 1 | 55.94 | no | 0 | 143.33 | 49.86 | 47.36 | 9.11 | 3.16 | 50.71 | 356.93 | O | 62.30 | 100.83 | F | chromatin binding;histone binding;protein binding | entry into host;regulation of transcription, DNA-templated | chromosome, telomeric region;nucleus |
| PF3D7_1118600 | MYST | histone acetyltransferase MYST | 71.13 | no | 0 | 64.05 | 16.93 | 10.55 | 2.51 | 2.39 | 16.11 | 353.78 | O | 83.51 | 54.94 | M | histone acetyltransferase activity;histone binding;protein binding;transcription coregulator activity | histone H4-K12 acetylation;histone H4-K16 acetylation;histone H4-K5 acetylation;histone H4-K8 acetylation;negative regulation of transcription, DNA-templated;positive regulation of transcription by RNA polymerase II;response to xenobiotic stimulus | cytoplasm;host cell periphery;nucleus;symbiont-containing vacuole |
| PF3D7_1124300 | BDP7 | bromodomain protein 7 | 62.33 | no | 0 | 64.40 | 69.11 | 38.46 | 10.73 | 3.55 | 14.52 | 360.42 | O | 73.69 | 26.80 | M | chromatin binding;histone binding;protein binding | regulation of transcription | heterochromatin;nucleus |
| PF3D7_1212900 | BDP2 | bromodomain protein 2, putative | 125.45 | no | 0 | 107.19 | 70.60 | 57.02 | 6.58 | 14.80 | 54.82 | 207.44 | O | 99.72 | 43.82 | M | protein binding | regulation of transcription | nucleus |
| PF3D7_1221000 | SET10 | histone-lysine N-methyltransferase, H3 lysine-4 specific | 271.07 | no | 0 | 5.25 | 3.05 | 21.93 | 5.72 | 2.16 | 12.63 | 50.60 | O | 9.49 | 12.36 | 0 | histone methyltransferase activity (H3-K4 specific), metal ion binding | chromatin remodelling;regulation of transcription | nuclear periphery, nucleus |
| PF3D7_1234100 | BDP5 | bromodomain protein 5 | 466.01 | no | 0 | 68.35 | 28.88 | 11.75 | 2.61 | 2.92 | 31.91 | 83.41 | O | 100.42 | 56.57 | M | protein binding | N/A | nucleus;transcription regulator complex |

| | | | | | | | | | | | | | | | | | | | |
|---|---|---|---|---|---|---|---|---|---|---|---|---|---|---|---|---|---|---|---|
| PF3D7_1322100 | SET2 | histone-lysine N-methyltransferase SET2 | 300.71 | no | 0 | 303.74 | 17.82 | 6.80 | 0.55 | 2.71 | 17.71 | 60.61 | R | 7.37 | 33.13 | F | chromatin binding;histone methyltransferase activity (H3-K36 specific);histone-lysine N-methyltransferase activity;nucleosomal histone binding;protein binding | antigenic variation;histone H3-K36 methylation;histone H3-K36 trimethylation;regulation of transcription, DNA-templated | chromatin;chromosome;nuclear periphery;nucleus |
| PF3D7_1355300 | SET6 | histone-lysine N-methyltransferase, putative | 60.02 | no | 0 | 25.03 | 3.61 | 3.00 | 0.25 | 1.97 | 10.37 | 7.26 | R | 4.63 | 12.20 | F | histone-lysine N-methyltransferase activity | histone lysine methylation | nuclear periphery;nucleus |
| PF3D7_1433400 | PHD2 | PHD finger protein PHD2, putative | 682.45 | no | 12 | 16.22 | 42.77 | 44.79 | 13.13 | 3.44 | 35.46 | 76.44 | O | 157.85 | 59.14 | M | metal ion binding | N/A | histone acetyltransferase complex; nucleus; membrane; integral component of membrane |
| PF3D7_1468100 | MORC | MORC family protein | 295.83 | no | 0 | 170.77 | 129.57 | 91.30 | 24.26 | 7.88 | 27.78 | 110.92 | R | 155.68 | 84.96 | M | protein binding | regulation of cell cycle | nucleus |
| PF3D7_1472200 | HDA1 | histone deacetylase, putative | 268.92 | no | 3 | 127.77 | 27.82 | 14.49 | 4.13 | 5.76 | 46.66 | 129.46 | O | 122.03 | 38.33 | M | NAD-dependent histone deacetylase activity (H3-K14 specific) | chromatin remodelling;regulation of transcription | nucleus, ⍰ integral component of membrane |
| PF3D7_1475600 | BDP4 | bromodomain protein 4, putative | 85.80 | no | 0 | 54.56 | 18.05 | 13.16 | 2.03 | 2.69 | 13.61 | 21.76 | R | 33.04 | 29.24 | M | protein binding | regulation of transcription | nucleus |
| **RNA polymerases** | | | | | | | | | | | | | | | | | | | |
| PF3D7_0110400 | RPB9 | DNA-directed RNA polymerase II subunit RPB9, putative | 29.69 | no | 0 | 45.15 | 31.60 | 30.74 | 4.60 | 13.05 | 119.22 | 167.78 | O | 121.76 | 135.17 | F | DNA-directed 5'-3' RNA polymerase activity | maintenance of transcriptional fidelity during DNA-templated transcription elongation from RNA polymerase II promoter;transcription by RNA polymerase II;transcription initiation from RNA polymerase II promoter;transcription-coupled nucleotide-excision repair | RNA polymerase II, core complex;nucleus |
| PF3D7_0215700 | RPB2 | DNA-directed RNA polymerase II subunit RPB2, putative | 151.72 | no | 0 | 99.34 | 58.36 | 67.24 | 8.90 | 1.43 | 4.46 | 56.77 | R | 202.29 | 68.53 | M | DNA-directed 5'-3' RNA polymerase activity | transcription by RNA polymerase II | RNA polymerase II, core complex |
| PF3D7_0303300 | RPB6 | DNA-directed RNA polymerases I, II, and III subunit RPABC2, putative | 17.83 | no | 0 | 26.80 | 9.18 | 8.42 | 0.74 | 19.62 | 37.33 | 33.93 | GV | 19.80 | 10.59 | M | DNA-directed 5'-3' RNA polymerase activity | transcription by RNA polymerase II | RNA polymerase I complex;RNA polymerase II, core complex;RNA polymerase III complex |
| PF3D7_0318200 | RPB1 | DNA-directed RNA polymerase II subunit RPB1 | 278.67 | no | 0 | 227.11 | 99.19 | 104.71 | 27.79 | 14.63 | 10.43 | 78.93 | R | 39.53 | 34.43 | 0 | DNA-directed 5'-3' RNA polymerase activity; protein binding; metal ion binding | transcription, DNA-templated; transcription by RNA polymerase II | transcription by RNA polymerase II; transcription, DNA-templated |
| PF3D7_0509400 | RNAPI | RNA polymerase I | 340.67 | no | 0 | 106.77 | 76.52 | 12.86 | 1.42 | 2.56 | 11.12 | 26.32 | R | 5.62 | 14.88 | F | RNA binding | regulation of transcription | RNA polymerase I complex;nucleolus |
| PF3D7_0708100 | RPB10 | DNA-directed RNA polymerases I, II, and III subunit RPABC5, putative | 8.24 | no | 0 | 71.20 | 23.14 | 13.25 | 0.61 | 41.22 | 177.64 | 30.92 | GV | 51.46 | 51.89 | 0 | DNA-directed 5'-3' RNA polymerase activity;zinc ion binding | tRNA transcription by RNA polymerase III;transcription by RNA polymerase I;transcription by RNA polymerase II | RNA polymerase I complex;RNA polymerase II, core complex;RNA polymerase III complex;nucleus |
| PF3D7_0923000 | RPB3 | DNA-directed RNA polymerase II subunit RPB3, putative | 38.35 | no | 0 | 75.74 | 13.31 | 9.78 | 0.61 | 4.20 | 36.68 | 11.96 | R | 25.76 | 12.27 | M | DNA binding | regulation of transcription | RNA polymerase II, core complex;nucleus |
| PF3D7_1134700 | RPA2 | DNA-directed RNA polymerase I subunit RPA2, putative | 175.53 | no | 0 | 100.82 | 61.29 | 9.83 | 0.49 | 4.59 | 12.21 | 17.07 | R | 27.44 | 18.67 | M | DNA binding | regulation of transcription | nucleus |
| PF3D7_1143300 | RPC40 | DNA-directed RNA polymerases I and III subunit RPAC1 | 38.96 | no | 0 | 105.20 | 31.73 | 12.05 | 2.45 | 5.62 | 56.14 | 102.24 | R | 55.67 | 50.72 | M | | tRNA transcription by RNA polymerase III;transcription by RNA polymerase I | RNA polymerase I complex;RNA polymerase III complex |
| PF3D7_1206600 | RPC2 | DNA-directed RNA polymerase III subunit RPC2, putative | 167.29 | no | 0 | 95.46 | 39.26 | 14.03 | 1.06 | 2.24 | 23.32 | 71.43 | R | 38.00 | 44.63 | F | DNA-directed 5'-3' RNA polymerase activity | transcription by RNA polymerase III | RNA polymerase III complex |
| PF3D7_1213700 | RPB8 | DNA-directed RNA polymerases I, II, and III subunit RPABC3, putative | 16.90 | no | 0 | 36.18 | 14.63 | 7.88 | 0.44 | 11.18 | 14.97 | 60.92 | O | 6.89 | 10.34 | F | DNA-directed 5'-3' RNA polymerase activity | DNA-templated transcription, initiation | RNA polymerase I complex;RNA polymerase II, core complex;RNA polymerase III complex;nucleus |
| PF3D7_1329000 | N/A | DNA-directed RNA polymerase III subunit RPC1, putative | 274.83 | no | 0 | 71.87 | 38.90 | 13.44 | 3.58 | 2.22 | 14.13 | 53.81 | R | 19.77 | 19.71 | 0 | DNA-directed 5'-3' RNA polymerase activity | transcription by RNA polymerase III | RNA polymerase III complex |
| PF3D7_1364800 | RPB5 | DNA-directed RNA polymerases I, II, and III subunit RPABC1, putative | 24.15 | no | 0 | 185.49 | 68.80 | 33.03 | 1.20 | 81.54 | 74.57 | 61.96 | R | 53.98 | 90.20 | F | DNA binding;DNA-directed 5'-3' RNA polymerase activity | transcription by RNA polymerase II | RNA polymerase I complex;RNA polymerase II, core complex;RNA polymerase III complex;nucleus |
| PF3D7_1404000 | RPB4 | DNA-directed RNA polymerase II subunit RPB4, putative | 20.05 | no | 0 | 14.20 | 9.21 | 6.56 | 1.07 | 44.29 | 865.01 | 61.34 | GV | 247.78 | 378.31 | F | RNA binding;DNA binding | regulation of transcription | nucleus |
| PF3D7_1415200 | N/A | DNA-directed RNA polymerases I and III subunit RPAC2 | 202.14 | no | 0 | 66.20 | 17.72 | 4.32 | 0.67 | 34.34 | 29.47 | 0.00 | R | 23.61 | 21.00 | 0 | DNA-directed 5'-3' RNA polymerase activity | transcription, DNA-templated | RNA polymerase I complex;RNA polymerase III complex |
| PF3D7_1458800 | N/A | DNA-directed RNA polymerase III subunit RPC5, putative | 103.36 | no | 0 | 8.36 | 3.60 | 2.21 | 0.15 | 2.33 | 11.75 | 9.45 | GV | 22.51 | 4.24 | M | RNA binding | regulation of transcription | RNA polymerase III complex |
| PF3D7_1463400 | N/A | DNA-directed RNA polymerase III subunit RPC4, putative | 39.12 | no | 0 | 25.05 | 7.33 | 3.86 | 0.16 | 4.16 | 19.51 | 5.41 | R | 34.50 | 7.62 | M | DNA binding;RNA binding | regulation of transcription | RNA polymerase III complex |
| **Mediators of RNA polymerase II** | | | | | | | | | | | | | | | | | | | |
| PF3D7_0411100 | MED8 | mediator of RNA polymerase II transcription subunit 8 | 23.97 | no | 0 | 34.82 | 2.70 | 5.48 | 0.45 | 4.61 | 105.53 | 52.18 | GV | 39.21 | 54.38 | F | protein binding | regulation of transcription | mediator complex;nucleus |
| PF3D7_0520200 | MED17 | mediator of RNA polymerase II transcription subunit 17, putative | 82.39 | no | 0 | 9.15 | 11.46 | 10.71 | 1.28 | 1.51 | 6.68 | 40.65 | O | 36.17 | 13.04 | M | N/A | N/A | mediator complex;nucleus |
| PF3D7_0709300 | MED14 | mediator of RNA polymerase II transcription subunit 14 | 325.49 | no | 0 | 19.66 | 9.05 | 8.28 | 1.61 | 1.74 | 11.70 | 17.26 | R | 30.96 | 12.10 | M | protein binding | regulation of transcription | mediator complex;nucleolus;nucleus |
| PF3D7_0822100 | MED7 | mediator of RNA polymerase II transcription subunit 4, putative | 28.20 | no | 0 | 20.92 | 7.64 | 5.30 | 0.86 | 59.08 | 126.74 | 31.95 | GV | 54.32 | 34.45 | M | protein binding | regulation of transcription | mediator complex;nucleus |
| PF3D7_1126400 | MED21 | mediator of RNA polymerase II transcription subunit 21 | 21.72 | no | 0 | 36.61 | 8.48 | 3.95 | 0.45 | 16.07 | 52.03 | 144.76 | O | 43.52 | 55.74 | F | protein binding | regulation of transcription by RNA polymerase II | core mediator complex;mediator complex;nucleus |
| PF3D7_1213200 | MED18 | mediator of RNA polymerase II transcription subunit 18, putative | 33.25 | no | 0 | 10.89 | 5.18 | 7.41 | 1.12 | 4.98 | 37.90 | 11.42 | GV | 13.84 | 25.14 | F | protein binding | regulation of transcription | mediator complex;nucleus |
| PF3D7_1465200 | MED4 | mediator of RNA polymerase II transcription subunit 4, putative | 44.68 | no | 0 | 52.44 | 11.71 | 9.66 | 0.75 | 3.05 | 12.84 | 0.00 | R | 11.70 | 6.52 | M | protein binding | regulation of transcription | mediator complex;nucleus |
| PF3D7_1469700 | MED6 | mediator of RNA polymerase II transcription subunit 6 | 24.57 | no | 0 | 3.19 | 2.42 | 7.14 | 1.20 | 17.37 | 183.73 | 34.02 | GV | 38.06 | 63.23 | F | protein binding | regulation of transcription by RNA polymerase II | core mediator complex;mediator complex;nucleus |
| PF3D7_1469800 | MED22 | mediator of RNA polymerase II transcription subunit 22 | 25.17 | no | 0 | 7.43 | 10.00 | 10.90 | 3.11 | 113.91 | 815.29 | 172.11 | GV | 200.55 | 434.25 | F | protein binding | regulation of transcription | mediator complex;nucleus |
| PF3D7_1475000 | MED31 | mediator of RNA polymerase II transcription subunit 31 | 16.61 | no | 0 | 7.30 | 4.62 | 5.12 | 0.39 | 1.42 | 20.92 | 0.00 | GV | 12.76 | 22.07 | F | protein binding | regulation of transcription by RNA polymerase II | core mediator complex;mediator complex;nucleus |
| **RNA helicases** | | | | | | | | | | | | | | | | | | | |
| PF3D7_0209800 | UAP56 | ATP-dependent RNA helicase UAP56 | 52.22 | no | 0 | 435.76 | 272.99 | 202.38 | 49.10 | 24.66 | 75.83 | 356.27 | R | 115.16 | 216.34 | F | RNA binding;RNA helicase activity | N/A | RNA N6-methyladenosine methyltransferase complex;cytoplasm;nucleus |
| PF3D7_0310500 | DHX57 | ATP-dependent RNA helicase DHX57, putative | 267.23 | no | 0 | 22.97 | 20.27 | 15.48 | 1.98 | 2.34 | 18.60 | 38.18 | O | 26.64 | 15.82 | M | RNA binding;helicase activity | maturation of SSU-rRNA from tricistronic rRNA transcript (SSU-rRNA, 5.8S rRNA, LSU-rRNA) | nucleolus;nucleus |
| PF3D7_0320800 | DOZI | ATP-dependent RNA helicase DDX6 | 49.41 | no | 0 | 6.50 | 20.89 | 54.61 | 11.23 | 47.25 | 163.76 | 237.14 | O | 110.82 | 263.97 | F | RNA helicase activity;mRNA binding;protein binding | P-body assembly;stress granule assembly | P-body;cytoplasm;cytoplasmic stress granule;nucleus |
| PF3D7_0321600 | DDX42 | ATP-dependent RNA helicase DDX42, putative | 86.65 | no | 0 | 13.26 | 14.71 | 18.84 | 8.28 | 4.52 | 25.40 | 61.68 | O | 71.84 | 89.96 | F | RNA binding;RNA helicase activity | regulation of transcription | cytoplasm;nucleus;ribonucleoprotein complex |
| PF3D7_0422500 | BRR2 | pre-mRNA-splicing helicase BRR2, putative | 337.90 | no | 0 | 105.44 | 72.81 | 27.09 | 4.35 | 8.08 | 18.99 | 136.41 | O | 92.14 | 36.50 | M | RNA helicase activity; nucleic acid binding; mRNA binding | ⍰ regulation of translation; RNA splicing, via transesterification reactions | spliceosomal complex; cytoplasmic stress granule;nucleus |
| PF3D7_0521700 | DDX1 | ATP-dependent RNA helicase DDX1, putative | 97.34 | no | 0 | 15.61 | 3.91 | 4.11 | 0.65 | 1.15 | 4.03 | 6.91 | R | 11.86 | 6.42 | M | RNA binding | rRNA processing | nucleus |
| PF3D7_0602100 | MTR4 | ATP-dependent RNA helicase MTR4 | 60.19 | no | 0 | 39.50 | 17.73 | 8.33 | 0.29 | 2.59 | 23.23 | 102.05 | O | 42.07 | 32.80 | M | RNA helicase activity;protein binding | RNA catabolic process;maturation of 5.8S rRNA | TRAMP complex;nuclear periphery;nucleus |
| PF3D7_0810600 | DBP1 | ATP-dependent RNA helicase DBP1, putative | 108.63 | no | 0 | 122.19 | 50.03 | 29.03 | 2.43 | 14.18 | 87.40 | 88.86 | R | 132.97 | 88.01 | M | RNA binding;RNA helicase activity;mRNA binding | regulation of transcription | P granule;nucleus |
| PF3D7_0827000 | DBP10 | ATP-dependent RNA helicase DBP10, putative | 151.10 | no | 0 | 70.36 | 41.06 | 15.42 | 1.18 | 0.91 | 8.26 | 71.34 | O | 164.04 | 65.86 | M | ATP binding;RNA binding | assembly of large subunit precursor of preribosome | nucleolus |
| PF3D7_0909900 | N/A | helicase SKI2W, putative | 160.37 | no | 0 | 28.73 | 7.92 | 11.19 | 1.63 | 2.02 | 9.06 | 8.98 | R | 20.33 | 10.27 | M | RNA helicase activity | RNA catabolic process;nuclear-transcribed mRNA catabolic process, 3'-5' exonucleolytic nonsense-mediated decay | Ski complex |
| PF3D7_0917600 | PRP43 | pre-mRNA-splicing factor ATP-dependent RNA helicase PRP43, putative | 94.14 | no | 0 | 60.85 | 25.10 | 26.07 | 2.55 | 1.14 | 3.76 | 9.19 | R | 47.08 | 15.27 | M | RNA binding | N/A | nucleus |
| PF3D7_0934100 | XPD | TFIIH transcription factor complex helicase XPD SU | 122.84 | no | 0 | 52.70 | 25.47 | 18.32 | 3.21 | 4.92 | 18.59 | 32.07 | R | 19.99 | 15.03 | M | 5'-3' DNA helicase activity;DNA helicase activity;damaged DNA binding;protein binding;single-stranded DNA helicase activity | nucleotide-excision repair, DNA duplex unwinding;nucleotide-excision repair, DNA incision;positive regulation of mitotic recombination;regulation of mitotic recombination;response to UV;response to oxidative stress;transcription by RNA polymerase II | cytoplasm;nucleus |
| PF3D7_1005500 | UPF1 | regulator of nonsense transcripts 1, putative | 183.96 | no | 0 | 31.20 | 22.88 | 46.85 | 7.51 | 10.15 | 113.61 | 139.54 | O | 165.50 | 175.06 | F | RNA binding;RNA helicase activity | nuclear-transcribed mRNA catabolic process, nonsense-mediated decay | cytoplasm;nucleus |
| PF3D7_1010200 | N/A | DNA2/NAM7 helicase, putative | 217.16 | no | 0 | 185.97 | 60.34 | 12.11 | 0.93 | 2.12 | 12.33 | 25.60 | R | 13.03 | 5.14 | M | RNA binding;RNA helicase activity | nuclear-transcribed mRNA catabolic process, nonsense-mediated decay | cytoplasm;nucleus |
| PF3D7_1030100 | PRP22 | pre-mRNA-splicing factor ATP-dependent RNA helicase PRP22, putative | 149.37 | no | 0 | 89.11 | 45.94 | 47.19 | 9.66 | 5.17 | 9.10 | 128.89 | O | 60.58 | 101.03 | F | RNA binding;RNA helicase activity | RNA splicing;spliceosomal complex disassembly | catalytic step 2 spliceosome |

| | | | | | | | | | | | | | | | | | | | |
|---|---|---|---|---|---|---|---|---|---|---|---|---|---|---|---|---|---|---|---|
| PF3D7_1037600 | XPB | TFIIH basal transcription factor complex helicase XPB subunit, putative | 102.87 | no | 0 | 53.16 | 19.24 | 21.04 | 4.81 | 1.70 | 41.13 | 138.14 | O | 77.09 | 257.49 | F | 3'-5' DNA helicase activity;ATP hydrolysis activity;DNA binding;DNA helicase activity;damaged DNA binding;helicase activity;transcription factor binding | nucleotide-excision repair, DNA duplex unwinding;nucleotide-excision repair, DNA incision;response to UV;transcription by RNA polymerase II;transcription initiation from RNA polymerase II promoter | nucleotide-excision repair factor 3 complex;transcription factor TFIIH holo complex;transcription preinitiation complex |
| PF3D7_1106000 | RUVB2 | RuvB-like helicase 2 | 53.41 | no | 0 | 51.92 | 24.07 | 16.98 | 1.03 | 1.78 | 19.30 | 32.70 | R | 3.61 | 20.86 | F | 5'-3' DNA helicase activity;ATP hydrolysis activity;DNA helicase activity;RNA binding;protein binding | DNA recombination;DNA repair;box C/D snoRNP assembly;chromatin remodeling;histone acetylation;regulation of transcription by RNA polymerase II | Ino80 complex;NuA4 histone acetyltransferase complex;R2TP complex;Swr1 complex;cytoplasm |
| PF3D7_1231600 | PRP2 | pre-mRNA-splicing factor ATP-dependent RNA helicase PRP2, putative | 137.20 | no | 0 | 54.29 | 14.50 | 22.31 | 2.19 | 6.33 | 24.26 | 46.16 | R | 179.16 | 29.09 | M | ATP-dependent activity, acting on RNA;RNA binding;RNA helicase activity | RNA splicing;mRNA splicing, via spliceosome;regulation of cell cycle | spliceosomal complex |
| PF3D7_1364300 | PRP16 | pre-mRNA-splicing factor RNA helicase PRP16 | 124.94 | no | 0 | 81.65 | 22.90 | 9.05 | 2.51 | 1.69 | 18.83 | 17.96 | R | 26.93 | 12.71 | M | RNA helicase activity | regulation of transcription | nucleus |
| PF3D7_1445900 | DDX17 | ATP-dependent RNA helicase DDX17 | 60.04 | no | 0 | 425.42 | 127.30 | 77.55 | 7.59 | 21.68 | 59.39 | 267.15 | R | 240.31 | 119.07 | M | ATP hydrolysis activity;RNA binding;RNA helicase activity;helicase activity;nucleic acid binding | | cytoplasm;host cell membrane;nucleus;ribonucleoprotein complex |
| PF3D7_1459000 | DBP5 | ATP-dependent RNA helicase DBP5 | 84.31 | no | 0 | 219.61 | 30.01 | 45.93 | 5.12 | 8.44 | 27.29 | 267.93 | O | 73.85 | 66.19 | M | 3'-5' DNA/RNA helicase activity;5'-3' DNA/RNA helicase activity;ATP hydrolysis activity;RNA binding;RNA helicase activity;single-stranded DNA helicase activity | DNA duplex unwinding;poly(A)+ mRNA export from nucleus | cytoplasm;cytoplasmic stress granule;nucleus |
| **Others** | | | | | | | | | | | | | | | | | | | |
| PF3D7_0205400 | N/A | PCI domain-containing protein, putative | 54.35 | no | 0 | 49.55 | 8.48 | 7.44 | 0.71 | 7.44 | 35.13 | 20.83 | R | 42.01 | 27.67 | M | RNA binding;double-stranded DNA binding | nuclear retention of pre-mRNA at the site of transcription;poly(A)+ mRNA export from nucleus;posttranscriptional tethering of RNA polymerase II gene DNA at nuclear periphery;transcription elongation from RNA polymerase II promoter | transcription export complex 2 |
| PF3D7_0407700 | N/A | TH1 domain-containing protein, putative | 197.41 | no | 0 | 6.11 | 0.33 | 13.33 | 9.64 | 3.60 | 21.13 | 51.24 | O | 208.91 | 61.67 | M | RNA binding | negative regulation of transcription elongation from RNA polymerase II promoter | NELF complex;nucleus |
| PF3D7_0422700 | EIF4A3 | eukaryotic initiation factor 4A-III, putative | 44.80 | no | 0 | 63.93 | 31.93 | 38.19 | 2.07 | 5.63 | 14.41 | 86.68 | O | 68.36 | 27.70 | M | RNA binding;RNA helicase activity;mRNA binding;translation initiation factor activity | regulation of translation;regulation of translational initiation | catalytic step 2 spliceosome;eukaryotic translation initiation factor 4F complex;nucleolus;nucleus |
| PF3D7_0517400 | FACT-L | FACT complex subunit SPT16, putative | 132.68 | no | 0 | 139.58 | 175.55 | 174.46 | 143.00 | 27.06 | 23.39 | 1056.64 | O | 484.10 | 74.42 | M | nucleosome binding | DNA replication-independent chromatin organization;positive regulation of transcription elongation from RNA polymerase II promoter;transcription elongation from RNA polymerase II promoter;transcription, DNA-templated | FACT complex;nucleus |
| PF3D7_0520700 | N/A | CDC73 domain-containing protein, putative | 118.00 | no | 0 | 17.35 | 11.54 | 4.41 | 0.54 | 1.68 | 16.96 | 2.06 | R | 4.91 | 7.37 | F | RNA polymerase II complex binding | positive regulation of transcription elongation from RNA polymerase II promoter;recruitment of 3'-end processing factors to RNA polymerase II holoenzyme complex | Cdc73/Paf1 complex |
| PF3D7_0609900 | N/A | tetratricopeptide repeat protein, putative | 50.76 | no | 0 | 34.32 | 5.61 | 11.49 | 12.07 | 4.48 | 12.04 | 8.41 | R | 30.53 | 11.04 | M | DNA binding | transcription by RNA polymerase III | transcription factor TFIIIC complex |
| PF3D7_0610900 | SPT5 | transcription elongation factor SPT5, putative | 149.36 | no | 0 | 144.61 | 74.80 | 24.35 | 3.25 | 15.75 | 32.14 | 146.31 | O | 213.18 | 73.26 | M | mRNA binding | regulation of transcription by RNA polymerase II;transcription elongation from RNA polymerase II promoter;transcription, DNA-templated | DSIF complex;nucleus |
| PF3D7_0611400 | SWIB | SWI/SNF-related matrix-associated actin-dependent regulator of chromatin | 97.79 | no | 0 | 14.57 | 3.36 | 5.49 | 1.01 | 1.07 | 9.72 | 10.54 | R | 29.07 | 17.75 | M | protein binding | regulation of gene expression;regulation of transcription by RNA polymerase II | cytoplasm;nucleus |
| PF3D7_0714500 | N/A | transcription elongation factor s-II, putative | 47.35 | no | 0 | 32.83 | 12.19 | 11.63 | 1.69 | 2.43 | 11.63 | 92.88 | O | 112.43 | 12.57 | M | DNA binding | regulation of DNA-templated transcription, elongation;regulation of transcription by RNA polymerase II | nucleus;transcription elongation factor complex |
| PF3D7_0820000 | SRCAP | Snf2-related CBP activator, putative | 246.75 | no | 0 | 41.42 | 33.94 | 15.68 | 1.21 | 2.94 | 11.26 | 93.84 | O | 39.79 | 33.45 | M | ATP binding;ATP hydrolysis activity;helicase activity;histone binding;transcription coactivator activity | ATP-dependent chromatin remodeling;chromatin remodeling;gene silencing;histone exchange | Swr1 complex;cytoplasm;membrane;nucleus |
| PF3D7_1012700 | NIF4 | NLI interacting factor-like phosphatase, putative | 171.01 | no | 0 | 16.21 | 6.12 | 11.28 | 2.63 | 2.17 | 7.39 | 29.05 | O | 4.24 | 13.92 | F | RNA polymerase II CTD heptapeptide repeat phosphatase activity | dephosphorylation of RNA polymerase II C-terminal domain | nucleus |
| PF3D7_1316900 | N/A | conserved protein, unknown function | 114.27 | no | 0 | 59.60 | 24.81 | 16.82 | 3.21 | 5.55 | 28.94 | 19.40 | R | 63.12 | 9.14 | M | N/A | regulation of transcription by RNA polymerase II | Piccolo NuA4 histone acetyltransferase complex;nucleus |
| PF3D7_1352700 | AQR | intron-binding protein aquarius, putative | 304.30 | no | 0 | 54.96 | 21.31 | 9.91 | 1.34 | 1.54 | 8.94 | 5.34 | R | 36.03 | 6.85 | M | RNA binding;RNA helicase activity | nuclear-transcribed mRNA catabolic process, nonsense-mediated decay;mRNA splicing, via spliceosome | spliceosomal complex;catalytic step 2 spliceosome;nucleus |
| PF3D7_1355700 | NIF3 | NLI interacting factor-like phosphatase, putative | 151.54 | no | 0 | 63.86 | 32.67 | 22.22 | 5.36 | 8.66 | 42.63 | 25.36 | R | 19.83 | 18.81 | 0 | RNA polymerase II CTD heptapeptide repeat phosphatase activity | dephosphorylation of RNA polymerase II C-terminal domain | nucleus |
| PF3D7_1406200 | SPT6 | transcription elongation factor SPT6, putative | 338.38 | no | 0 | 102.71 | 61.35 | 21.10 | 2.44 | 5.82 | 29.94 | 60.85 | O | 224.56 | 96.85 | M | histone binding;nucleosome binding | mRNA transcription by RNA polymerase II;nucleosome organization;obsolete chromatin maintenance;positive regulation of transcription elongation from RNA polymerase II promoter;regulation of mRNA processing;transcription elongation from RNA polymerase II promoter | nucleus;transcription elongation factor complex;transcriptionally active chromatin |
| PF3D7_1439500 | ORP2 | oocyst rupture protein 2, putative | 123.02 | no | 0 | 6.12 | 0.56 | 10.69 | 6.11 | 3.91 | 20.15 | 10.89 | GV | 74.44 | 7.59 | M | DNA-binding transcription activator activity, RNA polymerase II-specific;RNA polymerase II cis-regulatory region sequence-specific DNA binding | regulation of transcription, DNA-templated | cell periphery;cytoplasm;nucleus |

### DNA replication

| | | | | | | Expression | | | | | | | | Sex | | | GO Terms | | |
|---|---|---|---|---|---|---|---|---|---|---|---|---|---|---|---|---|---|---|---|
| Gene ID | Short name | Product Description | Mol. weight [kDa] | Signal Peptide | # TM Domains | Ring | Early Trophozoite | Late Trophozoite | Schizont | Gametocyte II | Gametocyte V | Ookinete | Peak expression | male gametocyte | female gametocyte | Sex specificity | Curated GO Functions (Molecular) | Curated GO Processes (Biological) | Curated GO Components (Cellular) |
| **DNA polymerases** | | | | | | | | | | | | | | | | | | | |
| PF3D7_0308000 | N/A | DNA polymerase delta small subunit, putative | 57.72 | no | 0 | 4.63 | 8.80 | 14.98 | 1.67 | 6.89 | 22.70 | 21.25 | GV | 257.69 | 23.61 | M | DNA-directed DNA polymerase activity | DNA biosynthetic process;DNA strand elongation involved in DNA replication | DNA polymerase complex;delta DNA polymerase complex |
| PF3D7_0411900 | N/A | DNA polymerase alpha catalytic subunit A | 225.40 | no | 0 | 5.61 | 3.94 | 47.52 | 14.94 | 6.49 | 42.82 | 46.11 | T | 467.94 | 35.04 | M | 3'-5' exonuclease activity;DNA replication origin binding;DNA-directed DNA polymerase activity;chromatin binding;purine nucleotide binding;pyrimidine nucleotide binding;single-stranded DNA binding | DNA replication;lagging strand elongation;leading strand elongation;mitotic DNA replication initiation | alpha DNA polymerase:primase complex |
| PF3D7_0630300 | N/A | DNA polymerase epsilon catalytic subunit A, putative | 344.60 | no | 0 | 4.33 | 1.08 | 11.56 | 1.66 | 2.02 | 24.46 | 16.96 | GV | 248.47 | 9.46 | M | DNA binding;DNA-directed DNA polymerase activity;single-stranded DNA 3'-5' exodeoxyribonuclease activity | DNA replication proofreading;DNA-dependent DNA replication;base-excision repair, gap-filling;leading strand elongation;mitotic cell cycle;nucleotide-excision repair, DNA gap filling | epsilon DNA polymerase complex |
| **DNA helicases** | | | | | | | | | | | | | | | | | | | |
| PF3D7_0514100 | UvrD | ATP-dependent DNA helicase UvrD | 170.29 | no | 1 | 6.20 | 3.20 | 17.97 | 6.27 | 6.44 | 29.73 | 89.45 | O | 27.77 | 33.96 | F | 3'-5' DNA helicase activity; single-stranded DNA helicase activity; ATP binding; DNA helicase activity; protein binding | recombinational repair, DNA repair | nucleus |
| PF3D7_0604600 | N/A | DNA helicase, putative | 206.43 | no | 0 | 4.44 | 1.15 | 1.88 | 0.27 | 0.86 | 5.31 | 20.11 | O | 9.57 | 4.24 | M | ATP-dependent DNA/DNA annealing activity;ATP-dependent activity, acting on DNA;DNA helicase activity;endodeoxyribonuclease activity | DNA repair;DNA rewinding;cellular response to DNA damage stimulus;replication fork processing;replication fork protection | nuclear replication fork;nucleus |
| PF3D7_0807100 | PSH3 | DNA helicase PSH3 | 157.20 | no | 0 | 16.63 | 13.93 | 9.47 | 2.40 | 4.95 | 45.14 | 34.89 | GV | 66.75 | 31.31 | M | 3'-5' DNA helicase activity;ATP hydrolysis activity;RNA binding;RNA helicase activity;single-stranded DNA helicase activity | DNA duplex unwinding | cytoplasm;nucleus |
| PF3D7_0818700 | N/A | DNA helicase, putative | 143.66 | no | 0 | 154.41 | 20.67 | 46.75 | 11.91 | 9.22 | 29.57 | 96.04 | R | 359.16 | 21.96 | M | ATP-dependent activity, acting on DNA | regulation of replication | nucleus |
| PF3D7_1031500 | DDX3X | ATP-dependent DNA helicase DDX3X | 107.80 | no | 0 | 71.32 | 12.20 | 39.01 | 8.12 | 8.93 | 59.66 | 156.10 | O | 87.61 | 150.99 | F | ATP hydrolysis activity;DNA helicase activity | mRNA splicing, via spliceosome | cytoplasm;nucleus |
| PF3D7_1106000 | RUVB2 | RuvB-like helicase 2 | 53.41 | no | 0 | 51.92 | 24.07 | 16.98 | 1.03 | 1.78 | 19.30 | 32.70 | R | 3.61 | 20.86 | F | 5'-3' DNA helicase activity;ATP hydrolysis activity;DNA helicase activity;RNA binding;protein binding | DNA recombination;DNA repair;box C/D snoRNP assembly;chromatin remodeling;histone acetylation;regulation of transcription by RNA polymerase II | Ino80 complex;NuA4 histone acetyltransferase complex;R2TP complex;Swr1 complex;cytoplasm |
| PF3D7_1106700 | DNA2 | DNA replication ATP-dependent helicase/nuclease DNA2, | 25.38 | no | 0 | 25.64 | 9.15 | 10.23 | 1.40 | 2.10 | 9.03 | 58.53 | O | 88.77 | 19.55 | M | nuclease activity | DNA-dependent DNA replication | nucleus |
| PF3D7_1250800 | N/A | DNA repair protein rhp16, putative | 192.17 | no | 0 | 58.81 | 38.20 | 25.28 | 5.26 | 6.29 | 7.57 | 88.79 | O | 188.32 | 29.03 | M | ATP binding;helicase activity | nucleotide-excision repair | nucleotide-excision repair complex |
| PF3D7_1357500 | N/A | DNA helicase, putative | 106.29 | no | 0 | 10.13 | 5.57 | 23.90 | 8.13 | 2.70 | 5.64 | 23.43 | O | 131.45 | 10.02 | M | ATP-dependent DNA/DNA annealing activity;ATP-dependent activity, acting on DNA;DNA helicase activity | DNA repair;DNA rewinding;cellular response to DNA damage stimulus;replication fork processing;replication fork protection | nuclear replication fork;nucleus |
| PF3D7_1362200 | RUVB3 | RuvB-like helicase 3 | 54.63 | no | 0 | 104.80 | 60.60 | 25.50 | 4.05 | 7.16 | 8.75 | 30.94 | R | 148.76 | 19.58 | M | ATP hydrolysis activity;DNA helicase activity;protein binding | box C/D snoRNP assembly;chromatin remodeling;histone acetylation;regulation of transcription by RNA polymerase II | Ino80 complex;NuA4 histone acetyltransferase complex;R2TP complex;Swr1 complex;host cell periphery;nucleus;symbiont-containing vacuole |
| PF3D7_1408400 | FANCJ | FANCJ-like helicase, putative | 135.54 | no | 0 | 10.87 | 14.94 | 10.88 | 2.72 | 0.98 | 13.05 | 43.48 | O | 278.60 | 10.44 | M | ATP binding;DNA helicase activity;DNA polymerase binding | DNA duplex unwinding;negative regulation of DNA recombination;negative regulation of t-circle formation;regulation of double-strand break repair via homologous recombination;telomeric loop disassembly | mitochondrion;nucleus |
| PF3D7_1429900 | WRN | ADP-dependent DNA helicase RecQ | 169.54 | no | 0 | 11.82 | 5.98 | 11.13 | 2.72 | 0.79 | 25.42 | 66.19 | O | 165.82 | 101.35 | M | 3'-5' DNA helicase activity;DNA helicase activity;four-way junction helicase activity;protein binding | DNA duplex unwinding;DNA recombination;DNA repair;DNA unwinding involved in DNA replication;double-strand break repair via homologous recombination | chromosome;chromosome, telomeric region;cytoplasm;nucleus |
| **Replication factors** | | | | | | | | | | | | | | | | | | | |
| PF3D7_0218000 | RFC2 | replication factor C subunit 2, putative | 37.92 | no | 0 | 8.69 | 6.06 | 58.69 | 10.39 | 13.35 | 40.20 | 886.79 | O | 241.59 | 33.82 | M | RNA binding | DNA repair;DNA replication;DNA-dependent DNA replication | DNA replication factor C complex;nucleus |
| PF3D7_0219600 | RFC1 | replication factor C subunit 1 | 104.18 | no | 0 | 6.02 | 2.72 | 24.02 | 5.31 | 3.69 | 11.36 | 106.06 | O | 282.45 | 17.91 | M | DNA binding | DNA repair;DNA replication | vacuolar acidification;vacuolar transport |
| PF3D7_1111100 | RFC5 | replication factor C subunit 5, putative | 40.35 | no | 0 | 9.89 | 5.14 | 37.32 | 5.46 | 10.62 | 61.42 | 119.20 | O | 155.28 | 61.78 | M | RNA binding | DNA repair;DNA replication;DNA strand elongation involved in DNA replication;DNA-dependent DNA replication | DNA replication factor C complex;nucleus |
| PF3D7_1241700 | RFC4 | replication factor C subunit 4, putative | 37.64 | no | 0 | 8.96 | 5.46 | 35.56 | 11.77 | 8.63 | 24.56 | 49.68 | O | 130.71 | 54.65 | M | ATP binding;DNA clamp loader activity | DNA repair;DNA replication;DNA-dependent DNA replication;leading strand elongation;response to xenobiotic stimulus | DNA replication factor C complex;nucleus |
| PF3D7_1442100 | RPA3 | replication factor A protein 3, putative | 12.61 | no | 0 | 5.70 | 6.24 | 16.13 | 1.98 | 23.62 | 51.00 | 45.38 | GV | 165.38 | 24.30 | M | damaged DNA binding;single-stranded DNA binding | DNA replication;base-excision repair;double-strand break repair via homologous recombination;mismatch repair;nucleotide-excision repair | DNA replication factor A complex;site of double-strand break |
| PF3D7_1463200 | RFC3 | replication factor C subunit 3, putative | 39.23 | no | 0 | 4.54 | 3.54 | 45.00 | 6.50 | 6.03 | 11.15 | 8.77 | T | 141.14 | 15.91 | M | DNA binding | DNA repair;DNA replication;DNA strand elongation involved in DNA replication;DNA-dependent DNA replication;leading strand elongation | DNA replication factor C complex;PCNA complex;nucleus |
| **DNA replication licensing factors** | | | | | | | | | | | | | | | | | | | |
| PF3D7_0416300 | MCM9 | DNA helicase MCM9, putative | 171.31 | no | 0 | 9.40 | 5.04 | 16.46 | 3.63 | 1.93 | 12.57 | 624.75 | O | 22.80 | 79.50 | F | ATP-dependent activity, acting on DNA;DNA replication origin binding;single-stranded DNA binding | DNA replication initiation;double-strand break repair via homologous recombination | MCM complex;nucleus |
| PF3D7_0527000 | MCM3 | DNA replication licensing factor MCM3, putative | 109.80 | no | 0 | 53.07 | 30.87 | 511.11 | 273.79 | 16.69 | 48.61 | 269.17 | T | 2853.14 | 372.70 | M | ATP hydrolysis activity | DNA replication | nucleus |
| PF3D7_0705400 | MCM7 | DNA replication licensing factor MCM7 | 94.24 | no | 0 | 11.77 | 5.78 | 133.58 | 36.61 | 6.29 | 12.61 | 117.12 | T | 730.03 | 91.54 | M | ATP-dependent activity, acting on DNA; ATP binding; single-stranded DNA binding; DNA replication origin binding; DNA binding | DNA duplex unwinding; DNA strand elongation involved in DNA replication; pre-replicative complex assembly involved in nuclear cell cycle DNA replication; double-strand break repair via break-induced replication; DNA replication | nucleus; cytoplasm; MCM complex |
| PF3D7_1211300 | MCM8 | DNA helicase MCM8, putative | 98.54 | no | 0 | 5.73 | 0.50 | 6.88 | 2.18 | 2.52 | 12.78 | 13.22 | O | 32.44 | 36.01 | 0 | ATP binding;ATP-dependent activity, acting on DNA;DNA helicase activity;DNA replication origin binding;single-stranded DNA binding | DNA replication initiation;DNA strand elongation involved in DNA replication | MCM complex;nuclear pre-replicative complex;nucleus |
| PF3D7_1211700 | MCM5 | DNA replication licensing factor MCM5, putative | 85.86 | no | 0 | 11.75 | 4.56 | 194.84 | 67.29 | 5.70 | 18.80 | 37.43 | T | 1362.90 | 99.36 | M | ATP binding;ATP-dependent activity, acting on DNA;DNA helicase activity;DNA replication origin binding;RNA binding;single-stranded DNA binding | DNA replication initiation;DNA strand elongation involved in DNA replication;double-strand break repair via break-induced replication;pre-replicative complex assembly involved in nuclear cell cycle DNA replication | MCM complex;nuclear pre-replicative complex;nucleus |

| | | | | | | | | | | | | | | | | | | | |
|---|---|---|---|---|---|---|---|---|---|---|---|---|---|---|---|---|---|---|---|
| PF3D7_1317100 | MCM4 | DNA replication licensing factor MCM4 | 115.22 | no | 0 | 10.44 | 6.05 | 148.92 | 45.16 | 5.69 | 21.55 | 67.84 | T | 785.94 | 69.19 | M | DNA replication origin binding; single-stranded DNA binding; ATP binding | double-strand break repair via break-induced replication; □<br>DNA replication; pre-replicative complex assembly involved in nuclear cell cycle DNA replication; DNA unwinding involved in DNA replication; □<br>DNA duplex unwinding; mitotic DNA replication initiation | MCM complex; □<br>nucleus |
| PF3D7_1355100 | MCM6 | DNA replication licensing factor MCM6 | 105.67 | no | 0 | 9.80 | 7.08 | 110.38 | 42.81 | 1.91 | 8.18 | 57.04 | T | 1124.74 | 56.30 | M | ATP-dependent activity, acting on DNA;DNA binding;DNA replication origin binding;protein binding;single-stranded DNA binding | DNA replication initiation;DNA unwinding involved in DNA replication;double-strand break repair via break-induced replication;mitotic DNA replication;pre-replicative complex assembly involved in nuclear cell cycle DNA replication | MCM complex;cytoplasm;nucleus |
| PF3D7_1412100 | MCMBP | mini-chromosome maintenance complex-binding protein | 112.77 | no | 0 | 8.12 | 5.38 | 15.08 | 6.86 | 1.21 | 7.49 | 16.46 | O | 27.05 | 5.82 | M | chromatin binding | DNA-dependent DNA replication;schizogony | MCM complex;cytoplasm;nucleus |
| Others | | | | | | | | | | | | | | | | | | | |
| PF3D7_0111300 | N/A | P-loop containing nucleoside triphosphate hydrolase | 139.21 | no | 0 | 4.91 | 2.18 | 12.37 | 2.73 | 2.33 | 21.09 | 24.92 | O | 171.23 | 27.02 | M | DNA binding | DNA replication | DNA replication factor C complex;nucleus |
| PF3D7_0206000 | RAD2 | DNA repair protein RAD2, putative | 178.73 | no | 0 | 22.89 | 10.97 | 8.28 | 3.50 | 1.29 | 11.82 | 321.85 | O | 71.69 | 15.63 | M | 5'-flap endonuclease activity;damaged DNA binding;endonuclease activity;flap endonuclease activity;single-stranded DNA endodeoxyribonuclease activity | DNA repair;nucleotide-excision repair, DNA incision, 3'-to lesion | chromosome, telomeric region;nucleotide-excision repair factor 3 complex |
| PF3D7_0215800 | ORC5 | origin recognition complex subunit 5 | 103.99 | no | 0 | 7.16 | 1.23 | 11.68 | 1.93 | 1.70 | 12.83 | 24.33 | O | 234.01 | 8.45 | M | ATP binding;ATP hydrolysis activity;DNA replication origin binding | DNA replication initiation | nuclear origin of replication recognition complex;nuclear pre-replicative complex |
| PF3D7_0317200 | CRK4 | cdc2-related protein kinase 4 | 182.22 | no | 0 | 10.39 | 5.75 | 73.63 | 29.01 | 3.48 | 8.00 | 243.56 | O | 43.80 | 24.98 | M | protein kinase activity; cyclin-dependent protein serine/threonine kinase activity; □<br>ATP binding | mitotic cell cycle; DNA replication; protein phosphorylation; peptidyl-serine phosphorylation; regulation of cell cycle | nucleus;cytosol;protein kinase CK2 complex |
| PF3D7_0317400 | N/A | DNA replication complex GINS protein, putative | 61.49 | no | 0 | 8.65 | 8.98 | 12.81 | 2.64 | 5.32 | 25.80 | 45.50 | O | 8.44 | 45.64 | F | N/A | double-strand break repair via break-induced replication | GINS complex;replication fork protection complex |
| PF3D7_0408500 | FEN1 | flap endonuclease 1 | 63.85 | no | 0 | 9.73 | 1.55 | 51.59 | 26.82 | 10.07 | 9.97 | 121.16 | O | 1072.17 | 59.74 | M | 5'-3' exonuclease activity;5'-flap endonuclease activity;endodeoxyribonuclease activity;flap endonuclease activity | | nucleus |
| PF3D7_0409600 | RPA1 | replication protein A1, large subunit | 134.20 | no | 0 | 8.16 | 10.81 | 24.70 | 5.94 | 1.44 | 12.79 | 52.50 | O | 45.15 | 31.82 | M | damaged DNA binding;sequence-specific DNA binding;single-stranded DNA binding;single-stranded telomeric DNA binding | DNA repair;DNA replication;DNA unwinding involved in DNA replication;DNA-dependent DNA replication;double-strand break repair via homologous recombination;meiotic cell cycle;nucleotide-excision repair;regulation of DNA metabolic process;telomere maintenance;telomere maintenance via telomerase | DNA replication factor A complex;nucleus |
| PF3D7_0420600 | AKIT6 | apicomplexan kinetochore protein 6, putative | 419.08 | no | 0 | 5.59 | 2.69 | 19.00 | 10.88 | 6.52 | 62.42 | 0.00 | GV | 210.81 | 82.79 | M | DNA binding | DNA replication | kinetochore;nucleus |
| PF3D7_0501800 | CAF1A | chromatin assembly factor 1 subunit A | 125.35 | no | 0 | 4.88 | 4.06 | 27.53 | 3.78 | 2.10 | 9.17 | 63.82 | O | 236.91 | 34.05 | M | histone binding;protein binding;transcription corepressor activity | DNA replication-independent chromatin assembly;mitotic sister chromatid segregation;obsolete regulation of chromatin silencing | HIR complex;nucleus |
| PF3D7_0503200 | CDC6 | cell division control protein 6, putative | 115.09 | no | 0 | 12.88 | 4.90 | 10.67 | 2.32 | 1.87 | 83.12 | 330.08 | O | 570.22 | 504.11 | M | DNA replication origin binding | DNA replication initiation;mitotic DNA replication checkpoint signaling;mitotic cell cycle | mitochondrion;nuclear origin of replication recognition complex |
| PF3D7_0505500 | MSH6 | DNA mismatch repair protein MSH6, putative | 156.44 | no | 0 | 9.34 | 6.55 | 55.50 | 9.61 | 2.49 | 7.86 | 43.93 | T | 219.05 | 23.24 | M | ATP-dependent activity, acting on DNA; mismatched DNA binding; □<br>four-way junction DNA binding | meiotic mismatch repair; DNA repair; pyrimidine dimer repair; maintenance of DNA repeat elements; negative regulation of DNA recombination | nucleus; mismatch repair complex; MutSalpha complex |
| PF3D7_0517400 | FACT-L | FACT complex subunit SPT16, putative | 132.68 | no | 0 | 139.58 | 175.55 | 174.46 | 143.00 | 27.06 | 23.39 | 1056.64 | O | 484.10 | 74.42 | M | nucleosome binding | DNA replication-independent chromatin organization;positive regulation of transcription elongation from RNA polymerase II promoter;transcription elongation from RNA polymerase II promoter;transcription, DNA-templated | FACT complex;nucleus |
| PF3D7_0520400 | N/A | single-stranded DNA-binding protein, putative | 35.66 | no | 0 | 3.87 | 4.12 | 10.75 | 2.23 | 2.66 | 14.66 | 3.79 | GV | 2.59 | 15.50 | F | DNA binding | DNA replication | intracellular;nucleus |
| PF3D7_0605800 | RAD50 | DNA repair protein RAD50, putative | 267.95 | no | 0 | 20.12 | 6.26 | 12.65 | 2.62 | 1.82 | 9.21 | 35.80 | O | 19.13 | 14.92 | M | G-quadruplex DNA binding; □<br>metal ion binding; single-stranded telomeric DNA binding<br>□adenylate kinase activity; double-stranded telomeric DNA binding; single-stranded DNA endodeoxyribonuclease activity | telomere maintenance via recombination; double-strand break repair; telomere maintenance via telomerase; □<br>DNA duplex unwinding; □<br>chromosome organization involved in meiotic cell cycle | condensed nuclear chromosome; □<br>nucleus; Mre11 complex; site of double-strand break |
| PF3D7_0611400 | SWIB | SWI/SNF-related matrix-associated actin-dependent regul | 97.79 | no | 0 | 14.57 | 3.36 | 5.49 | 1.01 | 1.07 | 9.72 | 10.54 | R | 29.07 | 17.75 | M | protein binding | regulation of gene expression;regulation of transcription by RNA polymerase II | cytoplasm;nucleus |
| PF3D7_0624600 | ISWI | ISWI chromatin-remodeling complex ATPase | 315.62 | no | 0 | 236.36 | 81.38 | 73.90 | 18.07 | 8.80 | 44.49 | 197.63 | R | 451.10 | 62.63 | M | ATP-dependent activity, acting on DNA; transcription cis-regulatory region binding; DNA binding; DNA helicase activity; helicase activity | positive regulation of transcription, DNA-templated; chromatin remodeling | nucleus |
| PF3D7_0705300 | ORC2 | origin recognition complex subunit 2 | 98.01 | no | 0 | 6.27 | 3.00 | 19.70 | 7.46 | 2.50 | 28.83 | 81.22 | O | 575.95 | 55.65 | M | DNA replication origin binding | DNA replication initiation | endoplasmic reticulum;mitochondrion;nuclear origin of replication recognition complex;nuclear periphery;nucleus |
| PF3D7_0726300 | PMS1 | DNA mismatch repair protein PMS1, putative | 156.84 | no | 0 | 22.25 | 2.79 | 10.82 | 3.61 | 1.81 | 10.37 | 51.67 | O | 114.31 | 7.48 | M | ATP binding;ATP hydrolysis activity;DNA binding;damaged DNA binding | DNA replication;mismatch repair | MutLalpha complex;mismatch repair complex;nucleus |
| PF3D7_0904800 | RPA1 | replication protein A1, small fragment | 56.23 | no | 0 | 7.42 | 7.10 | 30.90 | 6.31 | 2.47 | 15.07 | 35.85 | O | 360.05 | 45.89 | M | RNA binding;damaged DNA binding;sequence-specific DNA binding;single-stranded DNA binding;single-stranded telomeric DNA binding | DNA repair;DNA unwinding involved in DNA replication;DNA-dependent DNA replication;double-strand break repair via homologous recombination;meiotic cell cycle;nucleotide-excision repair;telomere maintenance;telomere maintenance via telomerase | DNA replication factor A complex;nucleus |
| PF3D7_0910900 | N/A | DNA primase large subunit, putative | 460.08 | no | 0 | 4.22 | 0.71 | 18.95 | 2.64 | 2.37 | 7.12 | 17.90 | T | 158.78 | 17.75 | M | DNA primase activity | DNA replication, synthesis of RNA primer | alpha DNA polymerase:primase complex;nucleus |
| PF3D7_0914800 | N/A | GINS complex subunit Psf3, putative | 24.84 | no | 0 | 12.20 | 1.17 | 10.89 | 2.80 | 10.18 | 141.91 | 47.86 | GV | 35.12 | 128.51 | F | N/A | DNA replication | nucleus |
| PF3D7_1004500 | AKIT2 | apicomplexan kinetochore protein 2, putative | 111.44 | no | 0 | 3.09 | 0.75 | 5.41 | 2.69 | 4.67 | 17.29 | 20.89 | O | 23.69 | 12.24 | M | DNA binding | DNA-dependent DNA replication | kinetochore;nucleus |
| PF3D7_1015800 | N/A | ribonucleoside-diphosphate reductase small chain, putati | 40.39 | no | 0 | 7.61 | 3.65 | 56.77 | 10.65 | 17.43 | 55.43 | 26.92 | T | 206.75 | 49.49 | M | ribonucleoside-diphosphate reductase activity, thioredoxin disulfide as acceptor | DNA replication;deoxyribonucleoside diphosphate metabolic process;deoxyribonucleotide biosynthetic process | ribonucleoside-diphosphate reductase complex |
| PF3D7_1023900 | CHD1 | chromodomain-helicase-DNA-binding protein 1 homolog, | 381.25 | no | 0 | 626.71 | 717.52 | 387.05 | 195.77 | 20.21 | 57.73 | 783.10 | O | 575.01 | 126.36 | M | protein binding; DNA binding; DNA helicase activity; ATP binding | N/A | nucleus |
| PF3D7_1104200 | SNF2L | chromatin remodeling protein | 167.44 | no | 0 | 185.35 | 54.00 | 35.85 | 6.97 | 3.54 | 23.34 | 164.78 | R | 22.56 | 21.17 | M | ATP binding; DNA binding; helicase activity | regulation of transcription, DNA-templated; nucleosome positioning; chromatin remodeling | nucleus |

| | | | | | | | | | | | | | | | | | | | |
|---|---|---|---|---|---|---|---|---|---|---|---|---|---|---|---|---|---|---|---|
| PF3D7_1107400 | RAD51 | DNA repair protein RAD51 | 38.59 | no | 0 | 37.19 | 12.00 | 71.34 | 14.19 | 32.42 | 202.44 | 711.67 | O | 143.07 | 928.22 | F | ATP binding;ATP-dependent activity, acting on DNA;DNA binding;DNA strand exchange activity;damaged DNA binding;double-stranded DNA binding;protein binding;single-stranded DNA binding | DNA recombinase assembly;DNA recombination;DNA repair;chromosome organization involved in meiotic cell cycle;mitochondrial double-strand break repair;mitotic recombination;reciprocal meiotic recombination;response to ionizing radiation;strand invasion | condensed nuclear chromosome;mitochondrion;nucleus |
| PF3D7_1117800 | MLH | DNA mismatch repair protein MLH | 118.31 | no | 0 | 13.04 | 3.70 | 10.35 | 2.81 | 2.66 | 12.61 | 20.56 | O | 85.60 | 11.79 | M | ATP binding;ATP hydrolysis activity;endonuclease activity;protein binding | mismatch repair | MutLalpha complex;chiasma;chromosome, telomeric region;mismatch repair complex |
| PF3D7_1250800 | Rhp16 | DNA repair protein rhp16, putative | 192.17 | no | 0 | 58.81 | 38.20 | 25.28 | 5.26 | 6.29 | 7.57 | 88.79 | O | 188.32 | 29.03 | M | ATP binding;helicase activity | nucleotide-excision repair | nucleotide-excision repair complex |
| PF3D7_1305000 | N/A | MCL1 domain-containing protein, putative | 174.47 | no | 0 | 11.28 | 7.44 | 16.91 | 5.13 | 2.51 | 14.42 | 30.60 | O | 2.01 | 21.63 | F | chromatin binding | DNA repair;DNA-dependent DNA replication;cellular response to DNA damage stimulus;mitotic cell cycle | nuclear replication fork;nucleus;replication fork protection complex |
| PF3D7_1329300 | CAF1B | chromatin assembly factor 1 subunit B, putative | 66.51 | no | 0 | 7.86 | 1.35 | 36.46 | 6.21 | 8.40 | 23.98 | 40.73 | O | 143.86 | 25.16 | M | chromatin binding;histone binding;protein binding;unfolded protein binding | chromatin assembly;chromatin assembly or disassembly | nucleus |
| PF3D7_1334100 | ORC4 | origin recognition complex subunit 4, putative | 117.79 | no | 0 | 4.80 | 2.93 | 13.36 | 2.56 | 3.68 | 11.85 | 44.21 | O | 204.94 | 26.96 | M | DNA binding | regulation of replication | nucleus |
| PF3D7_1347100 | TOP3 | DNA topoisomerase 3 | 84.31 | no | 0 | 7.65 | 2.04 | 18.95 | 6.61 | 3.85 | 75.60 | 116.55 | O | 69.75 | 157.00 | F | DNA binding;DNA topoisomerase activity;protein binding | DNA topological change;DNA unwinding involved in DNA replication | mitochondrion;nucleus |
| PF3D7_1361900 | PCNA1 | Proliferating cell nuclear antigen | 30.59 | no | 0 | 18810.00 | 12267.00 | 97.41 | 17.16 | 16.89 | 46.54 | 75.08 | R | 421.82 | 13.97 | M | DNA polymerase processivity factor activity | leading strand elongation;mismatch repair;translesion synthesis | PCNA complex;cytosol;nucleus |
| PF3D7_1427500 | MSH2-1 | DNA mismatch repair protein MSH2, putative | 94.74 | no | 0 | 3.85 | 2.31 | 17.95 | 2.51 | 0.90 | 3.35 | 9.92 | T | 3.36 | 2.14 | 0 | ATP-dependent activity, acting on DNA;DNA binding;damaged DNA binding;double-strand/single-strand DNA junction binding | DNA recombination;maintenance of DNA repeat elements;mismatch repair;negative regulation of DNA recombination;postreplication repair | MutSalpha complex;MutSbeta complex;mismatch repair complex;nucleus |
| PF3D7_1437200 | N/A | ribonucleoside-diphosphate reductase large subunit | 96.90 | no | 0 | 23.72 | 35.89 | 205.89 | 65.75 | 8.03 | 31.85 | 156.84 | T | 1037.72 | 215.42 | M | ATP binding;ribonucleoside-diphosphate reductase activity, thioredoxin disulfide as acceptor | DNA replication;deoxyribonucleotide biosynthetic process | apicoplast;nucleus;ribonucleoside-diphosphate reductase complex |
| PF3D7_1438700 | N/A | DNA primase small subunit | 53.49 | no | 0 | 21002.00 | 22282.00 | 42.4 | 45505.00 | 34455.00 | 27.69 | 55.58 | S | 326.36 | 70.91 | M | DNA primase activity;RNA binding | DNA replication, synthesis of RNA primer | alpha DNA polymerase:primase complex |
| PF3D7_1446600 | CEN2 | centrin-2 | 19.31 | no | 0 | 6.49 | 2.71 | 102.92 | 55.53 | 79.45 | 450.14 | 625.36 | O | 213.42 | 717.72 | F | RNA binding;calcium ion binding | centriole replication;mitotic cell cycle | centriole;centrosome;nucleus |
| PF3D7_1446700 | AKIT4 | apicomplexan kinetochore protein 4, putative | 42.63 | no | 0 | 20210.00 | 45047.00 | 25.67 | 26481.00 | 31686.00 | 89.37 | 62.00 | T | 62.14 | 68.67 | F | DNA binding | DNA replication | kinetochore;nucleus |
| PF3D7_1467500 | KIN17 | DNA/RNA-binding protein KIN17, putative | 52.46 | no | 0 | 43.88 | 13.86 | 20.36 | 6.75 | 1.42 | 6.50 | 7.67 | R | 21.23 | 13.09 | M | double-stranded DNA binding | DNA replication;cellular response to DNA damage stimulus | nucleus |
