## Supplementary material for "*Plasmodium falciparum* SET10 is a histone H3 lysine K18 methyltransferase that participates in a chromatin modulation network crucial for intraerythrocytic development": Table S5

Table S5. STRING analysis of *Pf* SET10 interactors with nuclear localization, sorted by cluster

### Nuclear proteins

| | | | | | | Expression | | | | | | | | Sex | | | GO Terms | | |
|---|---|---|---|---|---|---|---|---|---|---|---|---|---|---|---|---|---|---|---|
| Gene ID | Short name | Product Description | Mol. weight [kDa] | Signal Peptide | # TM Domains | Ring | Early Trophozoite | Late Trophozoite | Schizont | Gametocyte II | Gametocyte V | Ookinete | Peak expression | male gametocyte | female gametocyte | Sex specificity | Curated GO Functions (Molecular) | Curated GO Processes (Biological) | Curated GO Components (Cellular) |
| PF3D7_0613800 | ApiAP2 | AP2 domain transcription factor, putative | 485.58 | no | 0 | 9.78 | 1.34 | 89.95 | 46.14 | 11.78 | 17.07 | 97.31 | O | 85.41 | 6.69 | M | sequence-specific DNA binding, DNA-binding transcription factor activity, DNA binding | response to xenobiotic stimulus, regulation of transcription, DNA-templated | nucleus |
| PF3D7_1468100 | MORC | MORC family protein | 295.83 | no | 0 | 170.77 | 129.57 | 91.30 | 24.26 | 7.88 | 27.78 | 110.92 | R | 155.68 | 84.96 | M | protein binding | regulation of cell cycle | nucleus |
| PF3D7_0910200 | N/A | conserved Plasmodium protein, unknown function | 368.34 | no | 0 | 18.62 | 6.72 | 13.52 | 4.05 | 3.60 | 11.87 | 40.91 | O | 55.00 | 33.69 | M | N/A | N/A | nucleus |
| PF3D7_1008100 | PHD1 | PHD finger protein PHD1 | 453.00 | no | 0 | 86.17 | 53.30 | 21.26 | 8.32 | 3.22 | 17.25 | 68.53 | R | 35.99 | 17.39 | M | methylated histone binding, metal ion binding | entry into host, histone H3-K9 acetylation, histone H3-K4 trimethylation | histone acetyltransferase complex, nucleus |
| PF3D7_1221000 | SET10 | histone-lysine N-methyltransferase, H3 lysine-4 specific | 271.07 | no | 0 | 5.25 | 3.05 | 21.93 | 5.72 | 2.16 | 12.63 | 50.60 | O | 9.49 | 12.36 | 0 | histone methyltransferase activity (H3-K4 specific), metal ion binding | chromatin remodelling;regulation of transcription | nuclear periphery, nucleus |
| PF3D7_0405400 | PRPF8 | pre-mRNA-processing-splicing factor 8, putative | 366.39 | no | 0 | 156.78 | 102.89 | 99.78 | 16.79 | 6.56 | 15.84 | 402.18 | O | 85.17 | 40.86 | M | U1 snRNA binding, U6 snRNA binding, U2 snRNA binding, U5 snRNA binding, U1 snRNA binding, U6 snRNA binding, pre-mRNA intronic binding | spliceosomal tri-snRNP complex assembly, RNA splicing, via transesterification reactions, RNA splicing | nucleus, cytoplasm, spliceosomal complex, U5 snRNP, apicoplast |
| PF3D7_0317700 | N/A | CPSF (cleavage and polyadenylation specific factor), subunit A, putative | 338.53 | no | 0 | 35.07 | 13.47 | 13.36 | 2.84 | 5.65 | 38.85 | 85.22 | O | 166.90 | 117.96 | M | damaged DNA binding; nucleic acid binding; protein binding | nucleotide-excision repair; DNA repair; proteasome-mediated ubiquitin-dependent protein catabolic process | nucleus; cullin-RING ubiquitin ligase complex |
| PF3D7_0609000 | NUP637 | nucleoporin NUP637, putative | 720.51 | no | 0 | 86.14 | 62.26 | 38.45 | 12.58 | 8.74 | 47.35 | 51.50 | R | 79.23 | 33.93 | M | protein binding | nucleocytoplasmic transport | nuclear periphery; nucleus |
| PF3D7_1239200 | ApiAP2 | AP2 domain transcription factor, putative | 299.38 | no | 0 | 16650.00 | 28095.00 | 61.7 | 45480.00 | 34029.00 | 28.91 | 223.22 | S | 69.00 | 39.57 | M | DNA binding, DNA-binding transcription factor activity | regulation of transcription, DNA-templated. | heterochromatin, nucleus |
| PF3D7_0818900 | HSP70 | heat shock protein 70 | 73.91 | no | 0 | 3426.39 | 3329.13 | 3055.78 | 1342.83 | 388.88 | 170.78 | 1845.88 | R | 1425.79 | 301.35 | M | ATP binding; ATP hydrolysis activity; heat shock protein binding; phosphatidylinositol-3-phosphate binding; protein folding chaperone; unfolded protein binding; misfolded protein binding | response to unfolded protein; response to heat; cellular response to unfolded protein; | nucleus; cytoplasm; cytosol; cell surface; food vacuole; extracellular vesicle |
| PF3D7_0605800 | RAD50 | DNA repair protein RAD50, putative | 267.95 | no | 0 | 20.12 | 6.26 | 12.65 | 2.62 | 1.82 | 9.21 | 35.80 | O | 19.13 | 14.92 | M | G-quadruplex DNA binding; metal ion binding; single-stranded telomeric DNA binding adenylate kinase activity; double-stranded telomeric DNA binding; single-stranded DNA endodeoxyribonuclease activity | telomere maintenance via recombination; double-strand break repair; telomere maintenance via telomerase; DNA duplex unwinding; chromosome organization involved in meiotic cell cycle | condensed nuclear chromosome; nucleus; Mre11 complex; site of double-strand break |
| PF3D7_0629700 | SET1 | SET domain protein, putative | 796.04 | no | 0 | 45.67 | 31.36 | 25.30 | 4.30 | 3.13 | 23.45 | 102.36 | O | 82.65 | 38.44 | M | histone methyltransferase activity (H3-K4 specific) | positive regulation of transcription, DNA-templated, histone H3-K4 methylation | nucleus;histone methyltransferase complex |
| PF3D7_1472200 | HDA1 | histone deacetylase, putative | 268.92 | no | 3 | 127.77 | 27.82 | 14.49 | 4.13 | 5.76 | 46.66 | 129.46 | O | 122.03 | 38.33 | M | NAD-dependent histone deacetylase activity (H3-K14 specific) | chromatin remodelling;regulation of transcription | nucleus, integral component of membrane |
| PF3D7_1303800 | N/A | conserved Plasmodium protein, unknown function | 1111.80 | no | 2 | 5.85 | 3.71 | 11.23 | 1.84 | 2.07 | 33.23 | 81.14 | O | 109.15 | 24.81 | M | N/A | N/A | nucleus, integral component of membrane |
| PF3D7_0516800 | ApiAP2 | AP2 domain transcription factor AP2-O2, putative | 276.42 | no | 0 | 12.31 | 1.70 | 43.02 | 9.63 | 6.21 | 80.08 | 288.62 | O | 122.19 | 204.96 | F | sequence-specific DNA binding, DNA-binding transcription factor activity, DNA binding | regulation of transcription, DNA-templated | heterochromatin, nucleus |
| PF3D7_1322300 | N/A | BRCT domain-containing protein, putative | 178.35 | no | 0 | 5.93 | 3.97 | 8.03 | 1.79 | 4.15 | 27.02 | 75.21 | O | 508.88 | 10.67 | M | N/A | N/A | nucleus |
| PF3D7_1434500 | N/A | dynein-related AAA-type ATPase, putative | 971.00 | no | 9 | 46.73 | 45.83 | 7.08 | 0.54 | 1.98 | 20.47 | 18.71 | R | 64.58 | 16.90 | M | ATP binding; ATP hydrolysis activity | rRNA processing; ribosomal large subunit assembly | nucleus; membrane; integral component of membrane; preribosome, large subunit precursor |
| PF3D7_0524000 | KASbeta | karyopherin beta | 127.35 | no | 0 | 347.01 | 93.65 | 99.26 | 14.61 | 7.36 | 26.71 | 97.28 | R | 73.58 | 16.93 | M | RNA binding; nuclear localization sequence binding; protein domain specific binding; protein heterodimerization activity; nuclear import signal receptor activity | protein import into nucleus; NLS-bearing protein import into nucleus; nucleocytoplasmic transport | nucleus; nuclear pore; cytoplasm; food vacuole; RNA N6-methyladenosine methyltransferase complex |
| PF3D7_0726500 | N/A | ubiquitin carboxyl-terminal hydrolase, putative | 373.16 | no | 0 | 76.66 | 27.76 | 41.70 | 8.34 | 12.86 | 90.14 | 137.99 | O | 112.68 | 49.59 | M | cysteine-type endopeptidase activity; thiol-dependent deubiquitinase; protein binding | protein deubiquitination; ubiquitin-dependent protein catabolic process | nucleus; cytosol |
| PF3D7_1433400 | PHD2 | PHD finger protein PHD2, putative | 682.45 | no | 12 | 16.22 | 42.77 | 44.79 | 13.13 | 3.44 | 35.46 | 76.44 | O | 157.85 | 59.14 | M | metal ion binding | N/A | histone acetyltransferase complex; nucleus; membrane; integral component of membrane |
| PF3D7_0422500 | BRR2 | pre-mRNA-splicing helicase BRR2, putative | 337.90 | no | 0 | 105.44 | 72.81 | 27.09 | 4.35 | 8.08 | 18.99 | 136.41 | O | 92.14 | 36.50 | M | RNA helicase activity; nucleic acid binding; mRNA binding | regulation of translation; RNA splicing, via transesterification reactions | spliceosomal complex; cytoplasmic stress granule;nucleus |
| PF3D7_1003800 | EFTUD2 | U5 small nuclear ribonucleoprotein component, putative | 142.94 | no | 0 | 77.87 | 48.98 | 25.68 | 2.72 | 8.20 | 15.49 | 24.12 | R | 81.59 | 18.30 | M | GTPase activity; U5 snRNA binding, RNA binding GTP binding; | RNA splicing, via transesterification reactions; mRNA splicing, via spliceosome | U4/U6 x U5 tri-snRNP complex; U2-type catalytic step 2 spliceosome; ribonucleoprotein complex |
| PF3D7_0318200 | RPB1 | DNA-directed RNA polymerase II subunit RPB1 | 278.67 | no | 0 | 227.11 | 99.19 | 104.71 | 27.79 | 14.63 | 10.43 | 78.93 | R | 39.53 | 34.43 | 0 | DNA-directed 5'-3' RNA polymerase activity; protein binding; metal ion binding | transcription, DNA-templated; transcription by RNA polymerase II | transcription by RNA polymerase II; transcription, DNA-templated |
| PF3D7_1460500 | N/A | conserved Plasmodium protein, unknown function | 192.01 | no | 0 | 4.68 | 2.11 | 7.58 | 1.74 | 2.35 | 14.39 | 14.84 | O | 36.53 | 2.71 | M | N/A | N/A | nucleus |
| PF3D7_1410300 | N/A | WD repeat-containing protein, putative | 526.73 | no | 0 | 26.36 | 4.05 | 9.03 | 1.22 | 3.16 | 22.31 | 15.23 | R | 128.48 | 24.03 | M | U3 snoRNA binding | endonucleolytic cleavage in 5'-ETS of tricistronic rRNA transcript (SSU-rRNA, 5.8S rRNA, LSU-rRNA) | nucleus;cytoplasm |
| PF3D7_1246200 | ActI | Actin-1 | 41.87 | no | 0 | 55.25 | 77.34 | 1088.75 | 970.19 | 191.01 | 748.05 | 2474.74 | O | 240.73 | 2466.7 | F | protein binding | structural complex | cytoskeleton;cytoplasm;nucleus |
| PF3D7_0317500 | EG5 | kinesin-5 | 191.78 | no | 0 | 42.54 | 25.61 | 21.30 | 5.23 | 5.77 | 17.14 | 44.40 | O | 110.89 | 21.40 | M | ATP hydrolysis activity, microtubule motor activity; microtubule binding, plus-end-directed microtubule motor activity | microtubule-based movement, response to xenobiotic stimulus | nucleus, kinesin complex |
| PF3D7_1308200 | cpsSII | carbamoyl phosphate synthetase | 273.43 | no | 0 | 131.81 | 92.56 | 20.97 | 1.88 | 1.73 | 7.79 | 30.79 | R | 21.64 | 9.93 | M | carbamoyl-phosphate synthase (glutamine-hydrolyzing) activity, ATP binding. | de novo' pyrimidine nucleobase biosynthetic process, nitrogen compound metabolic process, glutamine metabolic process, glutamate catabolic process | nucleus, cytoplasm |
| PF3D7_1012700 | NIF4 | NLI interacting factor-like phosphatase, putative | 171.01 | no | 0 | 16.21 | 6.12 | 11.28 | 2.63 | 2.17 | 7.39 | 29.05 | O | 4.24 | 13.92 | F | RNA polymerase II CTD heptapeptide repeat phosphatase activity | dephosphorylation of RNA polymerase II C-terminal domain | nucleus |
| PF3D7_0619400 | N/A | cell division cycle protein 48 homologue, putative | 92.39 | no | 3 | 60.44 | 96.67 | 256.32 | 91.56 | 19.79 | 18.16 | 284.25 | O | 199.42 | 68.50 | M | ATP hydrolysis activity; polyubiquitin modification-dependent protein binding, hydrolase activity, RNA binding | regulation of cell cycle process, macroautophagy, ubiquitin-dependent ERAD pathway, retrograde protein transport, ER to cytosol; mitotic spindle disassembly | nucleus; cytoplasm;endoplasmic reticulum, cytosol |
| PF3D7_0321800 | N/A | WD repeat-containing protein, putative | 322.24 | no | 0 | 77.86 | 29.97 | 17.73 | 10.61 | 2.66 | 15.76 | 43.51 | R | 184.29 | 15.87 | M | protein binding | RNA splicing | nucleus; precatalytic spliceosome |
| PF3D7_1357100 | N/A | elongation factor 1-alpha | 48.96 | no | 0 | 119.25 | 251.23 | 99.01 | 20.40 | 18.26 | 13.44 | 5.82 | T | 14.96 | 20.46 | F | GTPase activity; translation elongation factor activity; RNA binding GTP binding | translation; translational elongation | nucleus; mitochondrion; eukaryotic translation elongation factor 1 complex; extracellular vesicle |
| PF3D7_0410900 | AKIT7 | apicomplexan kinetochore protein 7, putative | 117.53 | no | 0 | 22.49 | 12.17 | 8.33 | 1.00 | 3.33 | 22.24 | 189.56 | O | 133.30 | 46.13 | M | N/A | N/A | kinetochore; nucleus |
| PF3D7_1451100 | eEF2 | elongation factor 2 | 93.52 | no | 0 | 1381.80 | 1093.50 | 653.01 | 116.96 | 38.30 | 31.65 | 275.30 | R | 123.94 | 185.42 | F | GTP binding; GTPase activity; translation elongation factor activity; RNA binding | translational elongation; response to xenobiotic stimulus | nucleus; ribonucleoprotein complex |
| PF3D7_0827800 | SET3 | SET domain protein, putative | 283.55 | no | 0 | 64.54 | 10.91 | 23.63 | 8.84 | 6.71 | 49.68 | 167.97 | O | 119.89 | 325.00 | F | protein binding; histone-lysine N-methyltransferase activity | chromatin remodelling;regulation of transcription | nuclear periphery; nucleus |
| PF3D7_0710200 | N/A | conserved Plasmodium protein, unknown function | 350.70 | no | 0 | 8.14 | 11.06 | 10.76 | 10.36 | 4.34 | 56.98 | 330.97 | O | 3852.79 | 1172.20 | M | N/A | N/A | nucleus |
| PF3D7_1317100 | MCM4 | DNA replication licensing factor MCM4 | 115.22 | no | 0 | 10.44 | 6.05 | 148.92 | 45.16 | 5.69 | 21.55 | 67.84 | T | 785.94 | 69.19 | M | DNA replication origin binding; single-stranded DNA binding; ATP binding | double-strand break repair via break-induced replication; DNA replication; pre-replicative complex assembly involved in nuclear cell cycle DNA replication; DNA unwinding involved in DNA replication; DNA duplex unwinding; mitotic DNA replication initiation | MCM complex; nucleus |
| PF3D7_0420000 | N/A | zinc finger protein, putative | 401.14 | no | 0 | 64.52 | 36.23 | 11.67 | 1.96 | 2.55 | 21.96 | 96.31 | O | 83.95 | 39.48 | M | zinc ion binding | N/A | cytoplasm; membrane; nucleus |
| PF3D7_0629800 | N/A | cullin-2, putative | 136.34 | no | 0 | 69.76 | 7.61 | 10.68 | 1.65 | 2.34 | 30.56 | 11.76 | R | 48.74 | 24.90 | M | protein binding; ubiquitin-protein transferase activity; ubiquitin protein ligase binding | G1/S transition of mitotic cell cycle; ubiquitin-dependent protein catabolic process; protein ubiquitination; regulation of cell cycle | cullin-RING ubiquitin ligase complex;nucleus |
| PF3D7_0205900 | RPN1 | 26S proteasome regulatory subunit RPN1, putative | 108.36 | no | 0 | 95.37 | 81.94 | 202.97 | 133.40 | 48.64 | 59.85 | 270.09 | O | 37.03 | 121.60 | F | enzyme regulator activity | proteasome-mediated ubiquitin-dependent protein catabolic process; ubiquitin-dependent protein catabolic process | nucleus;cytoplasm;proteasome complex; proteasome regulatory particle; proteasome storage granule |
| PF3D7_1107800 | ApiAP2 | AP2 domain transcription factor, putative | 206.87 | no | 0 | 39.31 | 93.68 | 44.62 | 35.76 | 10.53 | 72.68 | 200.31 | O | 29.60 | 146.15 | F | sequence-specific DNA binding; DNA-binding transcription factor activity; DNA binding | regulation of transcription, DNA-templated | heterochromatin; nucleus |
| PF3D7_1130700 | SMC1 | structural maintenance of chromosomes protein 1 | 214.67 | no | 0 | 6.89 | 8.26 | 26.27 | 5.38 | 5.10 | 45.26 | 97.20 | O | 196.05 | 131.32 | M | ATP hydrolysis activity;ATP binding; double-stranded DNA binding; protein binding | mitotic sister chromatid segregation; chromosome organization | chromosome;cohesin complex; nuclear mitotic cohesin complex |
| PF3D7_1228800 | N/A | WD repeat-containing protein, putative | 378.91 | no | 1 | 37.43 | 30.41 | 51.02 | 14.24 | 7.36 | 39.98 | 253.23 | O | 225.37 | 160.00 | M | protein binding | nucleosome assembly | CAF-1 complex; nucleus; integral component of membrane |
| PF3D7_0624600 | ISWI | ISWI chromatin-remodeling complex ATPase | 315.62 | no | 0 | 236.36 | 81.38 | 73.90 | 18.07 | 8.80 | 44.49 | 197.63 | R | 451.10 | 62.63 | M | ATP-dependent activity, acting on DNA; transcription cis-regulatory region binding; DNA binding; DNA helicase activity; helicase activity | positive regulation of transcription, DNA-templated; chromatin remodeling | nucleus |
| PF3D7_1023900 | CHD1 | chromodomain-helicase-DNA-binding protein 1 homolog, putative | 381.25 | no | 0 | 626.71 | 717.52 | 387.05 | 195.77 | 20.21 | 57.73 | 783.10 | O | 575.01 | 126.36 | M | protein binding; DNA binding; DNA helicase activity; ATP binding | N/A | nucleus |
| PF3D7_1357800 | CCT4 | T-complex protein 1 subunit delta | 57.98 | no | 0 | 328.28 | 188.56 | 112.06 | 31.14 | 9.69 | 11.12 | 36.31 | R | 43.45 | 26.43 | M | unfolded protein binding; ATP binding | protein folding; response to xenobiotic stimulus | nucleus; cytosol; chaperonin-containing T-complex |
| PF3D7_0302900 | N/A | exportin-1, putative | 147.94 | no | 0 | 70.88 | 14.79 | 42.43 | 10.66 | 8.60 | 14.64 | 23.16 | R | 57.21 | 18.62 | M | nuclear export signal receptor activity | protein export from nucleus, regulation of protein export from nucleus, ribosomal small subunit export from nucleus | cytoplasm; nucleus |
| PF3D7_1148000 | N/A | serine/threonine protein kinase, putative | 170.70 | no | 0 | 6.11 | 1.34 | 12.99 | 2.78 | 4.63 | 22.21 | 81.72 | O | 59.36 | 41.96 | M | ATP binding, protein serine/threonine kinase activity, protein kinase activity | regulation of chromatin assembly or disassembly, protein phosphorylation, chromosome segregation, peptidyl-serine phosphorylation, intracellular signal transduction | nucleus |
| PF3D7_1234800 | SF3B3 | splicing factor 3B subunit 3, putative | 151.81 | no | 0 | 77.19 | 39.63 | 36.93 | 3.75 | 9.01 | 18.18 | 91.56 | O | 46.12 | 37.97 | M | nucleic acid binding, protein binding | RNA splicing, mRNA splicing, via spliceosome, RNA splicing, via transesterification reactions, spliceosomal complex assembly | spliceosomal complex, U2 snRNP nucleus, cytoplasm |
| PF3D7_0514100 | UvrD | ATP-dependent DNA helicase UvrD | 170.29 | no | 1 | 6.20 | 3.20 | 17.97 | 6.27 | 6.44 | 29.73 | 89.45 | O | 27.77 | 33.96 | F | 3'-5' DNA helicase activity; single-stranded DNA helicase activity; ATP binding; DNA helicase activity; protein binding | recombinational repair, DNA repair | nucleus |
| PF3D7_1468700 | eIF4A | eukaryotic initiation factor 4A | 45.31 | no | 0 | 1715.49 | 1152.94 | 678.76 | 92.91 | 79.47 | 83.48 | 368.49 | R | 299.10 | 214.37 | M | 3'-5' DNA/RNA helicase activity; ATP hydrolysis activity; helicase activity; translation initiation factor activity; nucleic acid binding | cytoplasmic translational initiation; regulation of translational initiation | nucleus; cytoplasm; cytosol; cytoplasmic stress granule; eukaryotic translation initiation factor 4F complex; food vacuole |
| PF3D7_1104200 | SNF2L | chromatin remodeling protein | 167.44 | no | 0 | 185.35 | 54.00 | 35.85 | 6.97 | 3.54 | 23.34 | 164.78 | R | 22.56 | 21.17 | M | ATP binding; DNA binding; helicase activity | regulation of transcription, DNA-templated; nucleosome positioning; chromatin remodeling | nucleus |
| PF3D7_1145400 | DYN1 | dynamin-like protein | 96.37 | no | 0 | 104.32 | 24.52 | 141.95 | 195.44 | 14.85 | 47.22 | 125.00 | S | 70.70 | 413.90 | F | GTPase activity; RNA binding; microtubule binding | mitochondrial fission; organelle fission; | nucleus; cytoplasm; plasma membrane; symbiont-containing vacuole membrane; food vacuole |
| PF3D7_0705400 | MCM7 | DNA replication licensing factor MCM7 | 94.24 | no | 0 | 11.77 | 5.78 | 133.58 | 36.61 | 6.29 | 12.61 | 117.12 | T | 730.03 | 91.54 | M | ATP-dependent activity, acting on DNA; ATP binding; single-stranded DNA binding; DNA replication origin binding; DNA binding | DNA duplex unwinding; DNA strand elongation involved in DNA replication; pre-replicative complex assembly involved in nuclear cell cycle DNA replication; double-strand break repair via break-induced replication; DNA replication | nucleus; cytoplasm; MCM complex |
| PF3D7_1311900 | vapA | V-type proton ATPase catalytic subunit A | 68.58 | no | 0 | 215.18 | 98.82 | 149.85 | 52.77 | 11.05 | 17.98 | 44.13 | R | 125.94 | 38.91 | M | proton-transporting ATPase activity, rotational mechanism; ATP binding | proton transmembrane transport; ATP metabolic process; vacuolar acidification | vacuolar proton-transporting V-type ATPase, V1 domain; vacuolar membrane; plasma membrane; food vacuole; proton-transporting V-type ATPase, V1 domain |
| PF3D7_0706000 | N/A | importin-7, putative | 145.51 | no | 0 | 87.79 | 27.74 | 41.24 | 10.56 | 14.46 | 84.34 | 38.58 | R | 77.58 | 44.48 | M | N/A | protein import into nucleus; intracellular protein transport | cytoplasm; cytosol; nuclear envelope; nucleus; nuclear pore |
| PF3D7_1458600 | N/A | zinc finger protein, putative | 139.59 | no | 0 | 67.96 | 8.21 | 7.98 | 1.50 | 3.18 | 13.13 | 7.82 | R | 31.50 | 10.95 | M | protein binding | N/A | nucleus; cytoplasm |
| PF3D7_1208900 | PPM11 | protein phosphatase PPM11, putative | 166.86 | no | 0 | 10.69 | 16.35 | 63.35 | 21.44 | 14.28 | 51.73 | 81.43 | O | 199.06 | 27.11 | M | phosphoprotein phosphatase activity;phosphatase activity | N/A | nucleus |
| PF3D7_0627800 | ACAS | acetyl-CoA synthetase | 113.88 | no | 0 | 121.57 | 49.14 | 54.06 | 13.24 | 3.01 | 7.58 | 34.99 | R | 23.00 | 17.90 | M | acetate-CoA ligase activity | acetyl-CoA biosynthetic process; generation of precursor metabolites and energy; chromatin organization; response to xenobiotic stimulus; acetyl-CoA biosynthetic process from acetate; histone acetylation | nucleus;cytoplasm |
| PF3D7_0317200 | CRK4 | cdc2-related protein kinase 4 | 182.22 | no | 0 | 10.39 | 5.75 | 73.63 | 29.01 | 3.48 | 8.00 | 243.56 | O | 43.80 | 24.98 | M | protein kinase activity; cyclin-dependent protein serine/threonine kinase activity; ATP binding | mitotic cell cycle; DNA replication; protein phosphorylation; peptidyl-serine phosphorylation; regulation of cell cycle | nucleus;cytosol;protein kinase CK2 complex |
| PF3D7_0505500 | MSH6 | DNA mismatch repair protein MSH6, putative | 156.44 | no | 0 | 9.34 | 6.55 | 55.50 | 9.61 | 2.49 | 7.86 | 43.93 | T | 219.05 | 23.24 | M | ATP-dependent activity, acting on DNA; mismatched DNA binding; four-way junction DNA binding | meiotic mismatch repair; DNA repair; pyrimidine dimer repair; maintenance of DNA repeat elements; negative regulation of DNA recombination | nucleus; mismatch repair complex; MutSalpha complex |
| PF3D7_0922200 | SAMS | S-adenosylmethionine synthetase | 44.84 | no | 0 | 315.15 | 102.02 | 890.99 | 258.09 | 11.44 | 83.26 | 194.91 | T | 44.53 | 171.00 | F | methionine adenosyltransferase activity | protein modification | nucleus;cytoplasm |
| PF3D7_1248900 | RPT6 | 26S protease regulatory subunit 8, putative | 49.55 | no | 0 | 80.34 | 66.05 | 178.09 | 57.21 | 26.44 | 35.09 | 140.92 | T | 79.96 | 175.91 | F | ATP hydrolysis activity; hydrolase activity; peptidase activity; transcription coregulator activity | proteasome regulatory particle assembly; proteasome-mediated ubiquitin-dependent protein catabolic process; | proteasome complex; nucleus; cytoplasm; proteasome regulatory particle, base subcomplex |
| PF3D7_0513200 | N/A | CPSF domain-containing protein, putative | 332.13 | no | 0 | 76.27 | 13.90 | 11.28 | 4.06 | 2.27 | 18.49 | 76.44 | O | 64.45 | 16.96 | M | polynucleotide adenylyltransferase activity;nucleic acid binding; protein binding | histone mRNA catabolic process; sno(s)RNA polyadenylation | nucleus; nucleolus; TRAMP complex |
| PF3D7_0303000 | NSF | N-ethylmaleimide-sensitive fusion protein | 89.13 | no | 0 | 13.10 | 3.04 | 8.97 | 1.09 | 2.59 | 4.81 | 20.57 | O | 11.59 | 33.59 | F | ATP hydrolysis activity; metal ion binding; ATP binding | Golgi to plasma membrane protein transport; endoplasmic reticulum to Golgi vesicle-mediated transport;SNARE complex disassembly | Golgi stack; food vacuole; host cell cytoplasm |
| PF3D7_1329100 | MyoF | myosin F, putative | 250.29 | no | 0 | 19.44 | 66.45 | 51.14 | 22.61 | 5.62 | 19.54 | 106.53 | O | 79.52 | 39.69 | M | actin filament binding; microfilament motor activity; cytoskeletal motor activity; protein binding; ATP binding; actin-dependent ATPase activity | cytoskeleton organization; actin filament organization; actin filament-based movement; vesicle transport along actin filament | cytoplasm; cytosol; nuclear envelope; nucleus; nuclear pore; actin cytoskeleton; myosin complex; vesicle; apical part of cell |

| | | | | | | | | | | | | | | | | | | | |
|---|---|---|---|---|---|---|---|---|---|---|---|---|---|---|---|---|---|---|---|
| PF3D7_0526500 | N/A | Suf domain-containing protein, putative | 165.38 | no | 0 | 88.79 | 24.83 | 16.01 | 1.62 | 3.76 | 15.92 | 41.01 | R | 80.94 | 45.64 | M | mRNA binding | RNA 3'-end processing | nucleus |
| PF3D7_0105800 | N/A | cyclin-dependent kinases regulatory subunit, putative | 99.31 | no | 0 | 5.42 | 1.25 | 24.78 | 12.70 | 4.33 | 26.24 | 56.62 | O | 28.40 | 36.31 | F | kinase activity | signal transduction | nucleus |
| PF3D7_0823300 | GCN5 | histone acetyltransferase GCN5 | 170.92 | no | 0 | 138.06 | 181.90 | 71.62 | 12.74 | 7.97 | 24.22 | 59.65 | T | 59.27 | 31.03 | M | H3 histone acetyltransferase activity | chromatin remodeling | nucleus |
| PF3D7_0527000 | MCM3 | DNA replication licensing factor MCM3, putative | 109.80 | no | 0 | 53.07 | 30.87 | 511.11 | 273.79 | 16.69 | 48.61 | 269.17 | T | 2853.14 | 372.70 | M | ATP hydrolysis activity | DNA replication | nucleus |
| PF3D7_1311500 | RPT1 | 26S protease regulatory subunit 7, putative | 46.84 | no | 0 | 81.96 | 50.75 | 189.27 | 46.12 | 7.85 | 12.95 | 126.39 | T | 378.22 | 148.07 | M | TBP-class protein binding | proteasome regulatory particle assembly; proteasome-mediated ubiquitin-dependent protein catabolic process; | proteasome complex; nucleus; cytoplasm; proteasome regulatory particle, base subcomplex |
| PF3D7_0215700 | RPB2 | DNA-directed RNA polymerase II subunit RPB2, putative | 151.72 | no | 0 | 99.34 | 58.36 | 67.24 | 8.90 | 1.43 | 4.46 | 56.77 | R | 202.29 | 68.53 | M | DNA-directed 5'-3' RNA polymerase activity | transcription by RNA polymerase II | RNA polymerase II, core complex |
| PF3D7_1437900 | HSP40 | HSP40, subfamily A | 48.47 | no | 0 | 217.89 | 220.27 | 89.47 | 8.74 | 27.79 | 75.89 | 222.01 | O | 178.64 | 320.86 | F | ATP hydrolysis activity;heat shock protein binding | protein folding;response to heat;response to xenobiotic stimulus | nucleus;cytosol;host cell plasma membrane |
| PF3D7_1466300 | RPN2 | 26S proteasome regulatory subunit RPN2, putative | 132.77 | no | 0 | 225.33 | 118.29 | 193.13 | 100.95 | 9.41 | 13.39 | 180.04 | R | 74.16 | 107.13 | F | endopeptidase activity | proteasome regulatory particle assembly; proteasome-mediated ubiquitin-dependent protein catabolic process; | proteasome complex; nucleus; cytoplasm; proteasome regulatory particle, base subcomplex |
| PF3D7_1352700 | AQR | intron-binding protein aquarius, putative | 304.30 | no | 0 | 54.96 | 21.31 | 9.91 | 1.34 | 1.54 | 8.94 | 5.34 | R | 36.03 | 6.85 | M | RNA binding;RNA helicase activity | nuclear-transcribed mRNA catabolic process, nonsense-mediated decay;mRNA splicing, via spliceosome | spliceosomal complex;catalytic step 2 spliceosome;nucleus |
| PF3D7_0818200 | 14-3-3I | 14-3-3 protein | 30.19 | no | 0 | 316.38 | 298.08 | 1144.07 | 511.61 | 242.14 | 88.32 | 323.33 | T | 267.35 | 91.82 | M | RNA binding;protein binding;histone binding | entry into host | nucleus;cytoplasm;food vacuole |
| PF3D7_0107800 | MRE11 | double-strand break repair protein MRE11 | 146.15 | no | 0 | 6.20 | 4.86 | 7.78 | 1.16 | 1.93 | 18.27 | 39.47 | O | 15.12 | 12.23 | 0 | nuclease activity;protein binding;single-stranded DNA endodeoxyribonuclease activity | DNA repair;double-strand break repair via homologous recombination;double-strand break repair via nonhomologous end joining;meiotic DNA double-strand break formation;mitochondrial double-strand break repair via homologous recombination;mitotic G2 DNA damage checkpoint signaling;telomere maintenance | Mre11 complex;nucleus;site of double-strand break |
| PF3D7_0322900 | RPS3A | 40S ribosomal protein S3A, putative | 30.05 | no | 0 | 414.72 | 482.85 | 209.93 | 44.12 | 40.69 | 29.20 | 132.31 | T | 93.24 | 60.94 | M | RNA binding;structural constituent of ribosome | translation | cytosol;cytosolic small ribosomal subunit;extracellular vesicle;nucleus |
| PF3D7_0604100 | SIP2 | AP2 domain transcription factor | 229.62 | no | 0 | 5.73 | 0.57 | 29.83 | 15.16 | 6.58 | 5.73 | 40.37 | O | 45.16 | 1.87 | M | sequence-specific DNA binding;telomeric DNA binding | N/A | nuclear periphery;nucleus |
| PF3D7_1132200 | TCP1 | T-complex protein 1 subunit alpha | 60.26 | no | 0 | 295.57 | 252.35 | 125.91 | 21.29 | 30.46 | 59.02 | 174.64 | R | 328.72 | 202.50 | M | ATP binding;unfolded protein binding | protein folding | chaperonin-containing T-complex;cytosol;intracellular;nucleus |
| PF3D7_1366300 | N/A | conserved Plasmodium protein, unknown function | 494.68 | no | 0 | 75.37 | 41.91 | 14.07 | 1.33 | 3.05 | 27.38 | 94.33 | O | 58.40 | 32.96 | M | N/A | N/A | RNA N6-methyladenosine methyltransferase complex;nucleus |
| PF3D7_0932800 | CSE1 | importin alpha re-exporter, putative | 159.11 | no | 0 | 48.05 | 33.94 | 46.16 | 12.39 | 15.18 | 32.16 | 16.80 | R | 58.79 | 27.77 | M | nuclear export signal receptor activity | protein export from nucleus | cytosol;nuclear envelope |
| PF3D7_1464700 | N/A | ATP synthase (C/AC39) subunit, putative | 44.53 | no | 0 | 51.00 | 19.16 | 23.83 | 6.78 | 4.64 | 36.85 | 83.72 | O | 99.58 | 32.97 | M | P-type proton-exporting transporter activity;proton-transporting ATPase activity, rotational mechanism | vacuolar acidification;vacuolar transport | food vacuole;lysosomal membrane;nucleus;vacuolar proton-transporting V-type ATPase complex;vacuolar proton-transporting V-type ATPase, V0 domain |
| PF3D7_0219600 | RFC1 | replication factor C subunit 1 | 104.18 | no | 0 | 6.02 | 2.72 | 24.02 | 5.31 | 3.69 | 11.36 | 106.06 | O | 282.45 | 17.91 | M | DNA binding | DNA repair;DNA replication | vacuolar acidification;vacuolar transport |
| PF3D7_0820000 | SRCAP | Snf2-related CBP activator, putative | 246.75 | no | 0 | 41.42 | 33.94 | 15.68 | 1.21 | 2.94 | 11.26 | 93.84 | O | 39.79 | 33.45 | M | ATP binding;ATP hydrolysis activity;helicase activity;histone binding;transcription coactivator activity | ATP-dependent chromatin remodeling;chromatin remodeling;gene silencing;histone exchange | Swr1 complex;cytoplasm;membrane;nucleus |
| PF3D7_1033600 | CEF1 | pre-mRNA-splicing factor CEF1, putative | 108.41 | no | 0 | 194.63 | 116.32 | 60.04 | 17.35 | 7.74 | 14.41 | 248.40 | O | 176.84 | 91.41 | M | DNA binding;protein binding | mRNA splicing, via spliceosome;regulation of transcription, DNA-templated | ATP-dependent chromatin remodeling;chromatin remodeling;gene silencing;histone exchange |
| PF3D7_0727900 | N/A | conserved Plasmodium protein, unknown function | 230.43 | no | 2 | 38.79 | 4.71 | 4.81 | 1.50 | 1.94 | 24.82 | 8.71 | R | 7.54 | 7.14 | 0 | DNA-binding transcription factor activity;sequence-specific DNA binding | positive regulation of transcription, DNA-templated;protein autoprocessing | endoplasmic reticulum membrane;nucleus |
| PF3D7_0303100 | N/A | CLP1 P-loop domain-containing protein, putative | 170.17 | no | 0 | 3.82 | 2.53 | 4.30 | 1.18 | 1.16 | 8.17 | 11.88 | O | 18.64 | 7.25 | M | polynucleotide 5'-hydroxyl-kinase activity | RNA processing;cleavage in ITS2 between 5.8S rRNA and LSU-rRNA of tricistronic rRNA transcript (SSU-rRNA, 5.8S rRNA, LSU-rRNA) | cytosol;nucleus |
| PF3D7_1440100 | RAD21 | cohesin complex subunit RAD21 | 118.50 | no | 0 | 3.85 | 1.40 | 26.43 | 10.09 | 1.34 | 5.96 | 78.05 | O | 163.27 | 1.93 | M | N/A | N/A | cohesin complex;mitochondrion;nucleus |
| PF3D7_1138500 | PPM2 | protein phosphatase PPM2, protein phosphatase 2C | 105.40 | no | 0 | 138.71 | 30.49 | 78.17 | 16.89 | 10.96 | 57.72 | 293.37 | O | 65.16 | 61.85 | M | magnesium-dependent protein serine/threonine phosphatase activity;protein serine/threonine phosphatase activity | protein dephosphorylation | cytoplasm;nucleus |
| PF3D7_0709300 | MED14 | mediator of RNA polymerase II transcription subunit 14 | 325.49 | no | 0 | 19.66 | 9.05 | 8.28 | 1.61 | 1.74 | 11.70 | 17.26 | R | 30.96 | 12.10 | M | protein binding | regulation of transcription | mediator complex;nucleolus;nucleus |
| PF3D7_0513600 | N/A | deoxyribodipyrimidine photo-lyase, putative | 129.20 | no | 0 | 13.47 | 26.26 | 47.79 | 5.71 | 17.26 | 94.76 | 1193.82 | O | 100.80 | 190.88 | F | deoxyribodipyrimidine photo-lyase activity | DNA repair | cytostome;food vacuole;mediator complex;nucleolus;symbiont-containing vacuole membrane |
| PF3D7_1304000 | CAPH | condensin complex subunit 2, putative | 119.22 | no | 0 | 4.84 | 0.86 | 29.56 | 18.96 | 11.09 | 12.11 | 74.38 | O | 140.41 | 26.73 | M | DNA topoisomerase binding;DNA topoisomerase type II (double strand cut, ATP-hydrolyzing) activator activity;chromatin binding | mitotic chromosome condensation | condensin complex;nucleus |
| PF3D7_0802000 | GDH3 | glutamate dehydrogenase, putative | 160.42 | no | 0 | 170.50 | 430.05 | 200.66 | 20.77 | 11.31 | 26.03 | 501.36 | O | 115.71 | 37.02 | M | glutamate dehydrogenase (NAD+) activity | glutamate catabolic process;glutamate metabolic process | cytosol;mitochondrion;nucleus |
| PF3D7_1355100 | MCM6 | DNA replication licensing factor MCM6 | 105.67 | no | 0 | 9.80 | 7.08 | 110.38 | 42.81 | 1.91 | 8.18 | 57.04 | T | 1124.74 | 56.30 | M | ATP-dependent activity, acting on DNA;DNA binding;DNA replication origin binding;protein binding;single-stranded DNA binding | DNA replication initiation;DNA unwinding involved in DNA replication;double-strand break repair via break-induced replication;mitotic DNA replication;pre-replicative complex assembly involved in nuclear cell cycle DNA replication | MCM complex;cytoplasm;nucleus |
| PF3D7_1248700 | N/A | conserved protein, unknown function | 231.32 | no | 0 | 312.04 | 166.94 | 96.03 | 21.10 | 16.74 | 35.34 | 527.35 | O | 589.21 | 211.21 | M | N/A | chromatin remodeling | nucleus |
| PF3D7_0411900 | N/A | DNA polymerase alpha catalytic subunit A | 225.40 | no | 0 | 5.61 | 3.94 | 47.52 | 14.94 | 6.49 | 42.82 | 46.11 | T | 467.94 | 35.04 | M | 3'-5' exonuclease activity;DNA replication origin binding;DNA-directed DNA polymerase activity;chromatin binding;purine nucleotide binding;pyrimidine nucleotide binding;single-stranded DNA binding | DNA replication;lagging strand elongation;leading strand elongation;mitotic DNA replication initiation | alpha DNA polymerase:primase complex |
| PF3D7_1008700 | N/A | Tubulin beta chain | 49.75 | no | 0 | 33.02 | 39.26 | 637.59 | 248.09 | 237.32 | 201.96 | 312.03 | T | 586.64 | 270.26 | M | GTP binding;protein binding;structural constituent of cytoskeleton | microtubule cytoskeleton organization;microtubule-based process;mitotic cell cycle;response to drug;response to xenobiotic stimulus | apical part of cell;cell surface;cytoplasm;food vacuole;microtubule;nucleus |
| PF3D7_0611400 | SWIB | SWI/SNF-related matrix-associated actin-dependent regulator of chromatin | 97.79 | no | 0 | 14.57 | 3.36 | 5.49 | 1.01 | 1.07 | 9.72 | 10.54 | R | 29.07 | 17.75 | M | protein binding | regulation of gene expression;regulation of transcription by RNA polymerase II | cytoplasm;nucleus |
| PF3D7_0403700 | CLF1 | pre-mRNA-splicing factor CLF1, putative | 94.02 | no | 0 | 80.84 | 48.89 | 31.01 | 7.52 | 5.83 | 13.21 | 161.17 | O | 589.82 | 61.49 | M | RNA binding;chromatin binding | DNA replication initiation;cis assembly of pre-catalytic spliceosome;mRNA splicing, via spliceosome;spliceosomal complex assembly | Prp19 complex;U2-type catalytic step 2 spliceosome;U2-type prespliceosome;catalytic step 2 spliceosome;post-mRNA release spliceosomal complex |
| PF3D7_0413600 | RPT3 | 26S protease regulatory subunit 6B, putative | 44.67 | no | 0 | 81.92 | 46.03 | 175.75 | 69.06 | 19.18 | 33.49 | 333.87 | O | 95.74 | 139.05 | F | ATP hydrolysis activity | positive regulation of RNA polymerase II transcription preinitiation complex assembly;proteasome regulatory particle assembly;ubiquitin-dependent protein catabolic process | cytosolic proteasome complex;nucleus;proteasome regulatory particle;proteasome regulatory particle, base subcomplex |
| PF3D7_1249800 | THO2 | THO complex subunit 2, putative | 348.67 | no | 2 | 80.68 | 47.06 | 30.83 | 8.05 | 5.43 | 20.81 | 43.22 | R | 141.68 | 51.21 | M | mRNA binding | mRNA export from nucleus | THO complex part of transcription export complex;nucleus |
| PF3D7_1352400 | NUP176 | nucleoporin NUP176, putative | 190.29 | no | 0 | 21.45 | 11.81 | 11.75 | 3.17 | 1.49 | 11.76 | 19.88 | R | 29.90 | 19.17 | M | | | nuclear periphery;nucleus |
| PF3D7_1447000 | N/A | 40S ribosomal protein S5 | 29.96 | no | 0 | 740.02 | 1270.54 | 396.40 | 181.44 | 67.76 | 94.22 | 436.80 | T | 105.97 | 119.57 | F | RNA binding;structural constituent of ribosome | translation | cytosolic small ribosomal subunit;nucleus |
| PF3D7_0306800 | CCT2 | T-complex protein 1 subunit beta | 59.20 | no | 0 | 208.93 | 137.08 | 82.22 | 4.81 | 5.79 | 10.28 | 45.60 | R | 16.29 | 46.58 | F | unfolded protein binding | protein folding | chaperonin-containing T-complex;cytosol;nucleus |
| PF3D7_1213800 | N/A | Proline--tRNA ligase | 86.65 | no | 0 | 155.35 | 47.67 | 44.04 | 17958.00 | 45387.00 | 45448.00 | 45460.00 | O | 45349.00 | 42644.00 | M | Ala-tRNA(Pro) hydrolase activity;proline-tRNA ligase activity | prolyl-tRNA aminoacylation;response to drug;response to xenobiotic stimulus;tRNA aminoacylation for protein translation | aminoacyl-tRNA synthetase multienzyme complex;cytoplasm;nucleus |
| PF3D7_0511500 | N/A | RNA pseudouridylate synthase, putative | 1187.60 | no | 0 | 123.39 | 80.80 | 23.46 | 4.89 | 3.70 | 34.06 | 95.55 | R | 124.35 | 50.82 | M | mRNA binding | N/A | nucleus |
| PF3D7_0724700 | BDP6 | bromodomain protein 6, putative | 420.67 | no | 0 | 42.01 | 19.54 | 10.52 | 2.44 | 4.24 | 19.55 | 32.71 | R | 24.66 | 18.50 | M | protein binding | regulation of transcription | nucleus;transcription regulator complex |
| PF3D7_1433500 | TOP2 | DNA topoisomerase 2 | 169.23 | no | 0 | 111.47 | 15.78 | 36.65 | 6.46 | 2.06 | 9.82 | 72.07 | R | 254.67 | 3.28 | M | DNA topoisomerase type II (double strand cut, ATP-hydrolyzing) activity | DNA topological change;mitotic DNA integrity checkpoint signaling;resolution of meiotic recombination intermediates;sister chromatid segregation | nucleus |
| PF3D7_1229500 | CCT3 | T-complex protein 1 subunit gamma | 60.66 | no | 0 | 234.12 | 125.81 | 56.76 | 22.11 | 7.81 | 17.33 | 345.08 | O | 34.13 | 92.49 | F | ATP binding;unfolded protein binding | protein folding | RNA N6-methyladenosine methyltransferase complex;chaperonin-containing T-complex;cytosol;nucleus |
| PF3D7_1306400 | RPT4 | 26S protease regulatory subunit 10B, putative | 44.68 | no | 0 | 59.18 | 26.24 | 86.30 | 19.83 | 15.15 | 35.78 | 120.16 | O | 204.00 | 88.24 | F | endopeptidase activity | positive regulation of RNA polymerase II transcription preinitiation complex assembly;proteasome regulatory particle assembly;ubiquitin-dependent ERAD pathway;ubiquitin-dependent protein catabolic process | cytosolic proteasome complex;extracellular vesicle;food vacuole;nucleus;proteasome regulatory particle;proteasome regulatory particle, base subcomplex |
| PF3D7_1008400 | RPT2 | 26S protease regulatory subunit 4, putative | 49.84 | no | 0 | 134.38 | 43.87 | 143.21 | 30.59 | 38.35 | 38.42 | 233.03 | O | 123.79 | 109.29 | M | TBP-class protein binding;endopeptidase activity;proteasome-activating activity | proteasome regulatory particle assembly;proteolysis;ubiquitin-dependent protein catabolic process | nucleus;proteasome regulatory particle;proteasome regulatory particle, base subcomplex |
| PF3D7_1235900 | XAB2 | pre-mRNA-splicing factor SYF1, putative | 125.43 | no | 1 | 76.06 | 22.68 | 17.74 | 2.69 | 5.63 | 30.58 | 67.57 | R | 108.46 | 30.05 | M | | generation of catalytic spliceosome for first transesterification step;mRNA splicing, via spliceosome | Prp19 complex;U2-type catalytic step 2 spliceosome;U2-type prespliceosome;catalytic step 2 spliceosome;post-mRNA release spliceosomal complex |
| PF3D7_0709600 | POP1 | ribonucleases P/MRP protein subunit POP1, putative | 138.30 | no | 0 | 32.52 | 20.67 | 5.42 | 0.77 | 1.54 | 5.16 | 7.96 | R | 0.90 | 5.04 | F | ribonuclease MRP activity;ribonuclease P activity | RNA phosphodiester bond hydrolysis, endonucleolytic;tRNA processing | nucleolar ribonuclease P complex;ribonuclease MRP complex;ribonuclease P complex |
| PF3D7_0409600 | RPA1 | replication protein A1, large subunit | 134.20 | no | 0 | 8.16 | 10.81 | 24.70 | 5.94 | 1.44 | 12.79 | 52.50 | O | 45.15 | 31.82 | M | damaged DNA binding;sequence-specific DNA binding;single-stranded DNA binding;single-stranded telomeric DNA binding | DNA repair;DNA replication;DNA unwinding involved in DNA replication;DNA-dependent DNA replication;double-strand break repair via homologous recombination;meiotic cell cycle;nucleotide-excision repair;regulation of DNA metabolic process;telomere maintenance;telomere maintenance via telomerase | DNA replication factor A complex;nucleus |
| PF3D7_1130200 | PfP0 | 60S ribosomal protein P0 | 34.97 | no | 0 | 1163.17 | 995.01 | 428.93 | 172.28 | 87.91 | 103.68 | 343.91 | R | 253.24 | 230.29 | M | RNA binding;large ribosomal subunit rRNA binding;structural constituent of ribosome | cytoplasmic translation;ribosomal large subunit assembly;translation | cytosolic large ribosomal subunit;extracellular vesicle;intracellular;nucleus |
| PF3D7_1224300 | PABP1 | polyadenylate-binding protein 1, putative | 97.23 | no | 0 | 1875.90 | 685.21 | 499.54 | 234.63 | 24.10 | 64.04 | 249.12 | R | 322.42 | 162.99 | M | RNA binding;mRNA 3'-UTR binding;mRNA binding;poly(A) binding;poly(U) RNA binding | RNA processing | P granule;cell surface;cytoplasm;cytosol;nuclear periphery;nucleus;ribonucleoprotein complex |
| PF3D7_0904800 | RPA1 | replication protein A1, small fragment | 56.23 | no | 0 | 7.42 | 7.10 | 30.90 | 6.31 | 2.47 | 15.07 | 35.85 | O | 360.05 | 45.89 | M | RNA binding;damaged DNA binding;sequence-specific DNA binding;single-stranded DNA binding;single-stranded telomeric DNA binding | DNA repair;DNA unwinding involved in DNA replication;DNA-dependent DNA replication;double-strand break repair via homologous recombination;meiotic cell cycle;nucleotide-excision repair;telomere maintenance;telomere maintenance via telomerase | DNA replication factor A complex;nucleus |
| PF3D7_1241200 | NUP269 | nucleoporin NUP269, putative | 276.53 | no | 2 | 17.81 | 11.52 | 8.80 | 2.09 | 1.91 | 15.78 | 7.04 | R | 244.28 | 13.90 | M | protein kinase activity | protein phosphorylation | nuclear periphery |
| PF3D7_1402300 | RPN6 | 26S proteasome regulatory subunit RPN6 | 78.38 | no | 0 | 33.29 | 15.83 | 33.70 | 8.60 | 3.12 | 22.47 | 28.63 | T | 32.61 | 20.03 | M | endopeptidase activity;structural molecule activity | proteasome assembly;proteolysis;ubiquitin-dependent protein catabolic process | cytoplasm;proteasome regulatory particle;proteasome regulatory particle, lid subcomplex |
| PF3D7_0516700 | N/A | ubiquitin carboxyl-terminal hydrolase 2, putative | 156.33 | no | 0 | 19.53 | 10.38 | 18.47 | 7.23 | 3.51 | 8.23 | 40.98 | O | 52.68 | 10.59 | M | thiol-dependent deubiquitinase | protein deubiquitination | nucleus;cytoplasm |
| PF3D7_0903700 | N/A | alpha tubulin 1 | 50.30 | no | 0 | 22.74 | 44.85 | 384.81 | 289.76 | 119.99 | 198.01 | 803.96 | O | 117.12 | 823.04 | F | GTP binding;protein binding;structural constituent of cytoskeleton | cytoplasmic microtubule organization;entry into host;microtubule cytoskeleton organization;microtubule-based process;mitotic cell cycle | apical part of cell;cell surface;cytoplasm;cytoplasmic microtubule;microtubule;nucleus |
| PF3D7_0812400 | KARalpha | karyopherin alpha | 61.16 | no | 0 | 96.50 | 76.80 | 129.72 | 22.27 | 9.20 | 30.64 | 206.57 | O | 128.54 | 58.17 | M | RNA binding;nuclear import signal receptor activity;nuclear localization sequence binding;protein domain specific binding;protein heterodimerization activity | NLS-bearing protein import into nucleus;nucleocytoplasmic transport;response to xenobiotic stimulus | nuclear pore;nucleus |
| PF3D7_0214000 | CCT8 | T-complex protein 1 subunit theta | 60.96 | no | 0 | 110.98 | 59.89 | 40.19 | 3.45 | 7.99 | 15.28 | 94.86 | R | 61.26 | 78.00 | F | protein binding;unfolded protein binding | protein folding | Maurer's cleft;apical part of cell;chaperonin-containing T-complex;cytosol;host cell cytoplasm;nucleus |
| PF3D7_0110500 | BDP3 | bromodomain protein 3, putative | 268.20 | no | 0 | 20.61 | 32.68 | 12.96 | 1.68 | 9.50 | 34.61 | 111.42 | O | 99.22 | 48.99 | M | N/A | histone acetylation | NuA4 histone acetyltransferase complex;nucleus |
| PF3D7_1362200 | RUVB3 | RuvB-like helicase 3 | 54.63 | no | 0 | 104.80 | 60.60 | 25.50 | 4.05 | 7.16 | 8.75 | 30.94 | R | 148.76 | 19.58 | M | ATP hydrolysis activity;DNA helicase activity;protein binding | box C/D snoRNP assembly;chromatin remodeling;histone acetylation;regulation of transcription by RNA polymerase II | Ino80 complex;NuA4 histone acetyltransferase complex;R2TP complex;Swr1 complex;host cell periphery;nucleus;symbiont-containing vacuole |
| PF3D7_1437200 | N/A | ribonucleoside-diphosphate reductase large subunit | 96.90 | no | 0 | 23.72 | 35.89 | 205.89 | 65.75 | 8.03 | 31.85 | 156.84 | T | 1037.72 | 215.42 | M | ATP binding;ribonucleoside-diphosphate reductase activity, thioredoxin disulfide as acceptor | DNA replication;deoxyribonucleotide biosynthetic process | apicoplast;nucleus;ribonucleoside-diphosphate reductase complex |
| PF3D7_0716000 | N/A | RNA-binding protein, putative | 132.71 | no | 0 | 148.99 | 75.92 | 37.64 | 3.54 | 21.92 | 36.91 | 172.10 | O | 560.82 | 121.67 | M | poly-pyrimidine tract binding;pre-mRNA 3'-splice site binding | N/A | U2-type prespliceosome;U2AF complex;commitment complex;nuclear speck |
| PF3D7_0630300 | N/A | DNA polymerase epsilon catalytic subunit A, putative | 344.60 | no | 0 | 4.33 | 1.08 | 11.56 | 1.66 | 2.02 | 24.46 | 16.96 | GV | 248.47 | 9.46 | M | DNA binding;DNA-directed DNA polymerase activity;single-stranded DNA 3'-5' exodeoxyribonuclease activity | DNA replication proofreading;DNA-dependent DNA replication;base-excision repair, gap-filling;leading strand elongation;mitotic cell cycle;nucleotide-excision repair, DNA gap filling | epsilon DNA polymerase complex |
| PF3D7_1030100 | PRP22 | pre-mRNA-splicing factor ATP-dependent RNA helicase PRP22, putative | 149.37 | no | 0 | 89.11 | 45.94 | 47.19 | 9.66 | 5.17 | 9.10 | 128.89 | O | 60.58 | 101.03 | F | RNA binding;RNA helicase activity | RNA splicing;spliceosomal complex disassembly | catalytic step 2 spliceosome |
| PF3D7_0922500 | PGK | phosphoglycerate kinase | 45.43 | no | 0 | 406.37 | 176.26 | 195.58 | 22.40 | 43.13 | 36.44 | 63.16 | R | 129.82 | 48.07 | M | ADP binding;ATP binding;phosphoglycerate kinase activity | gluconeogenesis;glycolytic process;positive regulation of oxidative phosphorylation | Maurer's cleft;cell surface;cytosol;extracellular vesicle;food vacuole;nucleus |
| PF3D7_1432100 | VAC | voltage-dependent anion-selective channel protein | 33.27 | no | 0 | 54.25 | 32.13 | 20.59 | 2.67 | 32.04 | 82.83 | 61.29 | GV | 60.91 | 71.44 | F | voltage-gated anion channel activity | N/A | cell periphery;mitochondrial outer membrane;mitochondrion;nucleus;symbiont-containing vacuole membrane |
| PF3D7_1423700 | N/A | conserved Plasmodium protein, unknown function | 183.57 | no | 0 | 290.42 | 132.36 | 44.41 | 5.71 | 7.51 | 69.34 | 152.92 | R | 223.74 | 67.63 | M | N/A | N/A | nucleus |
| PF3D7_1448500 | N/A | conserved Plasmodium protein, unknown function | 443.07 | no | 0 | 6.57 | 9.25 | 24.35 | 7.30 | 5.58 | 27.75 | 31.37 | O | 170.52 | 42.02 | M | N/A | N/A | nucleus |
| PF3D7_0706500 | N/A | conserved Plasmodium protein, unknown function | 196.57 | no | 0 | 230.24 | 65.83 | 27.68 | 5.79 | 7.52 | 50.37 | 101.37 | R | 86.47 | 42.42 | M | RNA binding;mRNA binding | | nucleus |
| PF3D7_1444800 | FBPA | Fructose-bisphosphate aldolase | 40.11 | no | 0 | 984.18 | 1910.1 | 1106.79 | 609.73 | 63.2 | 46.24 | 504.93 | T | 2266.46 | 147.86 | M | RNA binding;actin binding;fructose-bisphosphate aldolase activity | actin polymerization or depolymerization;fructose 1,6-bisphosphate metabolic process;glycolytic process | cell surface;cytoplasm;cytosol;extracellular vesicle;nucleus |

| | | | | | | | | | | | | | | | | | | |
|---|---|---|---|---|---|---|---|---|---|---|---|---|---|---|---|---|---|---|
| PF3D7_0723800 | AKIT1 | apicomplexan kinetochore protein 1, putative | 260.76 | no | 0 | 10.95 | 11.84 | 46.81 | 42.10 | 8.02 | 36.54 | 166.60 | O | 294.12 | 97.91 | M | WW domain binding | | kinetochore;nucleus |
| PF3D7_1305000 | N/A | MCL1 domain-containing protein, putative | 174.47 | no | 0 | 11.28 | 7.44 | 16.91 | 5.13 | 2.51 | 14.42 | 30.60 | O | 2.01 | 21.63 | F | chromatin binding | DNA repair;DNA-dependent DNA replication;cellular response to DNA damage stimulus;mitotic cell cycle | nuclear replication fork;nucleus;replication fork protection complex |
| PF3D7_0826700 | RACK1 | receptor for activated c kinase | 35.69 | no | 0 | 1995.06 | 865.88 | 479.73 | 48.03 | 106.13 | 110.78 | 224.72 | R | 44.18 | 64.90 | F | RNA binding;enzyme binding;protein kinase C binding;ribosome binding | positive regulation of protein phosphorylation;regulation of cell division;rescue of stalled ribosome | cytoplasm;cytosol;cytosolic small ribosomal subunit;extracellular vesicle;nucleus |
| PF3D7_0212500 | NUP434 | nucleoporin NUP434, putative | 477.85 | no | 2 | 70.81 | 41.72 | 22.97 | 4.31 | 3.90 | 19.31 | 92.59 | O | 68.86 | 30.60 | M | N/A | N/A | nuclear periphery |
| PF3D7_1130400 | RPT5 | 26S protease regulatory subunit 6A, putative | 49.54 | no | 0 | 111.69 | 60.79 | 162.07 | 59.91 | 21.55 | 31.01 | 61.73 | T | 87.43 | 136.32 | F | ATP hydrolysis activity;endopeptidase activity | positive regulation of RNA polymerase II transcription preinitiation complex assembly;proteasome regulatory particle assembly;proteolysis;ubiquitin-dependent protein catabolic process | nucleus;proteasome complex;proteasome regulatory particle, base subcomplex |
| PF3D7_0703500 | N/A | erythrocyte membrane-associated antigen | 264.91 | no | 0 | 229.90 | 253.73 | 93.48 | 14.28 | 21.44 | 69.16 | 67.43 | T | 127.83 | 69.48 | M | RNA binding;mRNA binding;helicase activity | N/A | membrane;nucleus |
| PF3D7_1021900 | N/A | PHAX domain-containing protein, putative | 267.38 | no | 0 | 27.79 | 31.93 | 71.51 | 11.88 | 9.75 | 77.18 | 222.86 | O | 276.89 | 369.11 | F | mRNA binding | N/A | nucleus |
| PF3D7_1441400 | FACT-S | FACT complex subunit SSRP1, putative | 58.79 | no | 0 | 38.32 | 114.71 | 63.91 | 24.41 | 47.03 | 20.72 | 36.31 | T | 130.87 | 23.21 | M | DNA binding;RNA binding;histone binding;nucleosome binding;single-stranded DNA binding | | FACT complex;nucleus |
| PF3D7_0320300 | CCT5 | T-complex protein 1 subunit epsilon | 59.18 | no | 0 | 113.31 | 82.48 | 44.30 | 5.62 | 7.69 | 19.46 | 77.57 | R | 60.63 | 36.12 | M | unfolded protein binding | protein folding | cell surface;chaperonin-containing T-complex;cytosol;nucleus |
| PF3D7_1014600 | ADA2 | transcriptional coactivator ADA2 | 300.25 | no | 1 | 176.38 | 156.32 | 115.15 | 33.94 | 6.81 | 17.88 | 125.06 | R | 123.81 | 63.55 | M | DNA-binding transcription factor activity;chromatin binding;protein binding;transcription coactivator activity | chromatin remodeling;positive regulation of histone acetylation;regulation of transcription by RNA polymerase II;regulation of transcription, DNA-templated | histone acetyltransferase complex;nucleus |
| PF3D7_0627500 | DJ1 | protein DJ-1 | 20.29 | no | 0 | 299.23 | 270.15 | 191.03 | 38.95 | 146.23 | 67.62 | 330.11 | O | 85.46 | 33.53 | M | peptidase activity;protein deglycase activity | guanine deglycation, glyoxal removal;protein deglycation;protein deglycation, glyoxal removal;protein folding;thiamine biosynthetic process | cell surface;cytosol;nucleus |
| PF3D7_0814300 | N/A | AAA family ATPase, putative | 139.67 | no | 0 | 28.83 | 10.41 | 4.15 | 0.61 | 1.26 | 6.08 | 21.25 | R | 2.39 | 5.76 | 0 | ATP hydrolysis activity;preribosome binding | positive regulation of telomerase activity;ribosome biogenesis | nucleus |
| PF3D7_1237500 | N/A | conserved Plasmodium protein, unknown function | 192.79 | no | 0 | 59.41 | 55.92 | 13.24 | 3.77 | 3.55 | 16.58 | 37.37 | R | 72.37 | 20.54 | M | N/A | N/A | nucleus |
| PF3D7_1225200 | N/A | DNA-binding protein, putative | 138.14 | no | 0 | 35.67 | 11.27 | 14.29 | 3.22 | 1.84 | 19.75 | 10.66 | R | 56.24 | 14.40 | M | N/A | N/A | nucleus |
| PF3D7_0624000 | HK | hexokinase | 55.26 | no | 0 | 190.58 | 185.98 | 224.54 | 73.63 | 5.34 | 7.04 | 88.78 | T | 112.02 | 47.33 | M | fructokinase activity | fructokinase activity;glucokinase activity;hexokinase activity;mannokinase activity | Maurer's cleft;RNA N6-methyladenosine methyltransferase complex;cytosol;mitochondrion;nucleus |
| PF3D7_1106000 | RUVB2 | RuvB-like helicase 2 | 53.41 | no | 0 | 51.92 | 24.07 | 16.98 | 1.03 | 1.78 | 19.30 | 32.70 | R | 3.61 | 20.86 | F | 5'-3' DNA helicase activity;ATP hydrolysis activity;DNA helicase activity;RNA binding;protein binding | DNA recombination;DNA repair;box C/D snoRNP assembly;chromatin remodeling;histone acetylation;regulation of transcription by RNA polymerase II | Ino80 complex;NuA4 histone acetyltransferase complex;R2TP complex;Swr1 complex;cytoplasm |
| PF3D7_1238500 | N/A | conserved Plasmodium protein, unknown function | 212.76 | no | 0 | 13.47 | 21.80 | 24.29 | 4.48 | 3.14 | 18.86 | 115.71 | O | 220.70 | 27.87 | M | N/A | N/A | nucleus |
| PF3D7_1408700 | N/A | conserved protein, unknown function | 851.88 | no | 0 | 37.27 | 10.13 | 14.03 | 3.19 | 3.36 | 46.01 | 87.60 | O | 88.85 | 43.04 | M | K63-linked polyubiquitin modification-dependent protein binding;thiol-dependent deubiquitinase | plastid organization;protein deubiquitination involved in ubiquitin-dependent protein catabolic process | cytoplasm;nucleus |
| PF3D7_1213900 | N/A | W2 domain-containing protein, putative | 134.55 | no | 0 | 101.63 | 31.40 | 16.90 | 1.13 | 4.11 | 21.58 | 48.64 | R | 37.46 | 48.80 | F | guanyl-nucleotide exchange factor activity;translation initiation factor activity;translation initiation factor binding | | eukaryotic translation initiation factor 2B complex |
| PF3D7_1222600 | AP2-G | AP2 domain transcription factor AP2-G | 284.06 | no | 0 | 6.31 | 0.44 | 0.95 | 0.04 | 0.84 | 2.23 | 2.59 | R | 0.16 | 0.17 | 0 | DNA-binding transcription activator activity;protein binding;sequence-specific DNA binding | response to xenobiotic stimulus | nucleus |
| PF3D7_1211700 | MCM5 | DNA replication licensing factor MCM5, putative | 85.86 | no | 0 | 11.75 | 4.56 | 194.84 | 67.29 | 5.70 | 18.80 | 37.43 | T | 1362.90 | 99.36 | M | ATP binding;ATP-dependent activity, acting on DNA;DNA helicase activity;DNA replication origin binding;RNA binding;single-stranded DNA binding | DNA replication initiation;DNA strand elongation involved in DNA replication;double-strand break repair via break-induced replication;pre-replicative complex assembly involved in nuclear cell cycle DNA replication | MCM complex;nuclear pre-replicative complex;nucleus |
| PF3D7_0102900 | N/A | aspartate--tRNA ligase | 73.45 | no | 0 | 59.33 | 45.30 | 24.36 | 3.75 | 7.32 | 20.19 | 47.74 | R | 26.44 | 71.25 | F | RNA binding;aspartate-tRNA ligase activity;protein homodimerization activity | aspartyl-tRNA aminoacylation | aminoacyl-tRNA synthetase multienzyme complex;cytosol;nucleus |
| PF3D7_1234100 | BDP5 | bromodomain protein 5 | 466.01 | no | 0 | 68.35 | 28.88 | 11.75 | 2.61 | 2.92 | 31.91 | 83.41 | O | 100.42 | 56.57 | M | protein binding | N/A | nucleus;transcription regulator complex |
| PF3D7_1019000 | eIF2A | eukaryotic translation initiation factor subunit eIF2A | 280.75 | no | 2 | 225.70 | 93.84 | 47.82 | 21.01 | 11.12 | 21.07 | 293.04 | O | 54.48 | 31.32 | M | mRNA binding;ribosome binding;tRNA binding;translation initiation factor activity | | cytosolic small ribosomal subunit;nucleus |
| PF3D7_0317400 | N/A | DNA replication complex GINS protein, putative | 61.49 | no | 0 | 8.65 | 8.98 | 12.81 | 2.64 | 5.32 | 25.80 | 45.50 | O | 8.44 | 45.64 | F | N/A | double-strand break repair via break-induced replication | GINS complex;replication fork protection complex |
| PF3D7_1205600 | N/A | tetratricopeptide repeat protein, putative | 39.02 | no | 0 | 18.28 | 50.17 | 85.34 | 19.59 | 20.31 | 47.55 | 177.97 | O | 2047.55 | 140.64 | M | histone binding | CENP-A containing chromatin assembly;DNA replication-dependent chromatin assembly | nucleoplasm;nucleus |
| PF3D7_0725000 | N/A | exonuclease I, putative | 157.93 | no | 0 | 10.51 | 6.57 | 15.78 | 4.83 | 1.75 | 18.82 | 115.46 | O | 9.15 | 25.87 | F | 5'-3' exonuclease activity;5'-flap endonuclease activity;flap endonuclease activity | DNA recombination;mismatch repair | nucleus |
| PF3D7_0703200 | N/A | conserved Plasmodium protein, unknown function | 182.54 | no | 0 | 11.53 | 17.90 | 48.64 | 42.06 | 12.08 | 44.49 | 162.80 | O | 42.79 | 60.42 | F | N/A | N/A | nucleus |
| PF3D7_1012400 | HGPRT | hypoxanthine-guanine phosphoribosyltransferase | 26.36 | no | 0 | 752.59 | 1396.42 | 877.39 | 110.33 | 95.30 | 127.31 | 309.40 | T | 37.91 | 118.91 | F | RNA binding;hypoxanthine phosphoribosyltransferase activity;magnesium ion binding | GMP salvage;IMP salvage;adenine salvage;guanine salvage;hypoxanthine metabolic process;purine ribonucleoside salvage;response to xenobiotic stimulus | |
| PF3D7_1143300 | RPC40 | DNA-directed RNA polymerases I and III subunit RPAC1 | 38.96 | no | 0 | 105.20 | 31.73 | 12.05 | 2.45 | 5.62 | 56.14 | 102.24 | R | 55.67 | 50.72 | M | | tRNA transcription by RNA polymerase III;transcription by RNA polymerase I | RNA polymerase I complex;RNA polymerase III complex |
| PF3D7_0708800 | HSP110c | heat shock protein 110 | 99.96 | no | 1 | 303.12 | 163.20 | 71.91 | 8.00 | 18.46 | 42.18 | 256.20 | R | 96.20 | 105.75 | F | ATP hydrolysis activity;protein binding | protein folding;response to heat | cytoplasm;cytosol;inner membrane pellicle complex;nucleus |
| PF3D7_0823800 | N/A | DnaJ protein, putative | 76.68 | no | 0 | 25.29 | 15.54 | 28.59 | 7.15 | 3.60 | 22.20 | 12.31 | T | 20.43 | 17.83 | M | RNA binding;unfolded protein binding | chaperone cofactor-dependent protein refolding;protein refolding | cytoplasm;food vacuole;nucleus |
| PF3D7_0917000 | MOP | merozoite organizing protein | 215.35 | no | 3 | 9.08 | 4.80 | 13.43 | 2.87 | 0.67 | 7.49 | 17.43 | O | 66.93 | 12.66 | M | N/A | cell division;mitotic chromosome condensation | apical part of cell;condensed chromosome;condensin complex |
| PF3D7_1249300 | PPM4 | protein phosphatase PPM4, putative | 119.60 | no | 0 | 29.40 | 13.49 | 25.15 | 4.69 | 2.70 | 11.17 | 124.95 | O | 48.51 | 30.66 | M | magnesium-dependent protein serine/threonine phosphatase activity;protein serine/threonine phosphatase activity | | nucleus |
| PF3D7_0309300 | N/A | N2227-like protein, putative | 112.25 | no | 0 | 37.83 | 11.42 | 24.53 | 9.60 | 4.12 | 7.50 | 29.73 | R | 24.17 | 15.36 | M | S-adenosylmethionine-dependent methyltransferase activity | N/A | cytoplasm;nucleus |
| PF3D7_1309400 | N/A | HORMA domain protein, putative | 158.12 | no | 0 | 24.72 | 22.75 | 59.20 | 28.26 | 10.48 | 158.85 | 346.56 | O | 139.32 | 2078.18 | F | N/A | N/A | nucleus |
| PF3D7_0805700 | FIKK8 | serine/threonine protein kinase, FIKK family | 171.80 | no | 0 | 7.04 | 1.11 | 16.65 | 6.97 | 2.04 | 10.72 | 87.91 | O | 13.35 | 12.90 | 0 | protein serine/threonine kinase activity | intracellular signal transduction;protein phosphorylation | cytoplasm;cytosol;nucleus |
| PF3D7_0729100 | AKIT8 | apicomplexan kinetochore protein 8, putative | 304.52 | no | 0 | 7.69 | 3.83 | 13.36 | 1.61 | 3.38 | 21.43 | 63.27 | O | 53.85 | 20.87 | M | allantoicase activity | allantoin catabolic process | kinetochore;nucleus |
| PF3D7_0306100 | N/A | conserved Plasmodium protein, unknown function | 74.33 | no | 0 | 33.32 | 10.03 | 10.27 | 1.63 | 2.25 | 27.52 | 54.24 | O | 22.87 | 32.48 | F | N/A | N/A | nucleus |
| PF3D7_1011800 | PREBP | PRE-binding protein | 131.62 | no | 0 | 393.91 | 383.43 | 88.77 | 14.46 | 23.65 | 184.56 | 573.99 | O | 377.24 | 2674.04 | F | DNA-binding transcription factor activity;RNA binding;double-stranded DNA binding;mRNA binding | regulation of gene expression;regulation of transcription, DNA-templated | cytoplasm;nucleus |
| PF3D7_1015800 | N/A | ribonucleoside-diphosphate reductase small chain, putative | 40.39 | no | 0 | 7.61 | 3.65 | 56.77 | 10.65 | 17.43 | 55.43 | 26.92 | T | 206.75 | 49.49 | M | ribonucleoside-diphosphate reductase activity, thioredoxin disulfide as acceptor | DNA replication;deoxyribonucleoside diphosphate metabolic process;deoxyribonucleotide biosynthetic process | ribonucleoside-diphosphate reductase complex |
| PF3D7_1224500 | ASF1 | histone chaperone ASF1 | 31.55 | no | 0 | 4.62 | 2.91 | 31.66 | 6.35 | 19.20 | 52.48 | 29.64 | GV | 35.66 | 23.46 | M | chromatin binding;histone binding;unfolded protein binding | DNA replication-dependent chromatin assembly;DNA replication-independent chromatin assembly;chromatin assembly;gene silencing;histone exchange | nucleus |
| PF3D7_1212700 | EIF3A | eukaryotic translation initiation factor 3 subunit A, putative | 166.06 | no | 0 | 1252.39 | 541.64 | 142.06 | 22.49 | 14.34 | 34.28 | 377.21 | R | 205.89 | 277.96 | F | mRNA binding;translation initiation factor activity | formation of cytoplasmic translation initiation complex;regulation of translational initiation;translation reinitiation;translational initiation | eukaryotic translation initiation factor 3 complex;eukaryotic translation initiation factor 3 complex, eIF3e;eukaryotic translation initiation factor 3 complex, eIF3m;multi-eIF complex;nucleus |
| PF3D7_1138800 | N/A | WD repeat-containing protein, putative | 222.08 | no | 0 | 78.02 | 18.27 | 13.61 | 4.09 | 2.74 | 18.55 | 102.40 | O | 48.83 | 28.15 | M | protein binding | N/A | nucleus |
| PF3D7_1325100 | N/A | phosphoribosylpyrophosphate synthetase | 49.38 | no | 0 | 651.23 | 226.86 | 147.30 | 57.81 | 16.22 | 77.72 | 47.56 | R | 12.65 | 47.31 | F | RNA binding;ribose phosphate diphosphokinase activity | 5-phosphoribose 1-diphosphate biosynthetic process;nucleotide biosynthetic process;pentose-phosphate shunt;purine nucleotide biosynthetic process;purine ribonucleoside salvage | cytoplasm;mitochondrion;nucleus;ribose phosphate diphosphokinase complex |
| PF3D7_1108400 | CK2alpha | casein kinase 2, alpha subunit | 39.89 | no | 0 | 101.82 | 47.36 | 63.15 | 17.65 | 6.41 | 27.58 | 137.34 | O | 19.73 | 33.65 | F | RNA binding;protein binding;protein serine/threonine kinase activity | peptidyl-serine phosphorylation;peptidyl-threonine phosphorylation;protein autophosphorylation;protein phosphorylation;regulation of cell cycle | cytoplasm;cytosol;extracellular vesicle;nucleus;protein kinase CK2 complex |
| PF3D7_0308600 | PRPF19 | pre-mRNA-processing factor 19, putative | 60.75 | no | 0 | 87.33 | 38.22 | 41.68 | 4.64 | 3.19 | 6.89 | 87.33 | O | 17.59 | 7.95 | M | protein binding;ubiquitin-protein transferase activity | generation of catalytic spliceosome for first transesterification step | Prp19 complex;U2-type catalytic step 1 spliceosome;catalytic step 2 spliceosome;nuclear matrix;nucleus |
| PF3D7_0209800 | UAP56 | ATP-dependent RNA helicase UAP56 | 52.22 | no | 0 | 435.76 | 272.99 | 202.38 | 49.10 | 24.66 | 75.83 | 356.27 | R | 115.16 | 216.34 | F | RNA binding;RNA helicase activity | N/A | RNA N6-methyladenosine methyltransferase complex;cytoplasm;nucleus |
| PF3D7_0206700 | ADSL | adenylosuccinate lyase | 55.18 | no | 0 | 1.31 | 0.30 | 0.14 | 0.00 | 0.14 | 0.16 | 0.00 | R | 4.22 | 21.68 | F | N6-(1,2-dicarboxyethyl)AMP AMP-lyase (fumarate-forming) activity | purine nucleotide metabolic process | Maurer's cleft;nucleus |
| PF3D7_1223100 | PKAr | cAMP-dependent protein kinase regulatory subunit | 50.83 | no | 0 | 10.63 | 1.61 | 151.01 | 346.12 | 7.79 | 21.53 | 315.60 | S | 306.72 | 87.24 | M | 14-3-3 protein binding;RNA binding;cAMP-dependent protein kinase regulator activity;protein binding | | cAMP-dependent protein kinase complex;extracellular vesicle;nucleus |
| PF3D7_0205600 | N/A | conserved Plasmodium protein, unknown function | 76.45 | no | 0 | 14.59 | 11.45 | 23.75 | 5.60 | 6.45 | 11.15 | 61.75 | O | 192.46 | 20.89 | M | N/A | N/A | nucleus |
| PF3D7_1438700 | N/A | DNA primase small subunit | 53.49 | no | 0 | 21002.00 | 22282.00 | 42.4 | 45505.00 | 34455.00 | 27.69 | 55.58 | S | 326.36 | 70.91 | M | DNA primase activity;RNA binding | DNA replication, synthesis of RNA primer | alpha DNA polymerase:primase complex |
| PF3D7_0520200 | MED17 | mediator of RNA polymerase II transcription subunit 17, putative | 82.39 | no | 0 | 9.15 | 11.46 | 10.71 | 1.28 | 1.51 | 6.68 | 40.65 | O | 36.17 | 13.04 | M | N/A | N/A | mediator complex;nucleus |
| PF3D7_0501800 | CAF1A | chromatin assembly factor 1 subunit A | 125.35 | no | 0 | 4.88 | 4.06 | 27.53 | 3.78 | 2.10 | 9.17 | 63.82 | O | 236.91 | 34.05 | M | histone binding;protein binding;transcription corepressor activity | DNA replication-independent chromatin assembly;mitotic sister chromatid segregation;obsolete regulation of chromatin silencing | HIR complex;nucleus |
| PF3D7_0517000 | N/A | 60S ribosomal protein L12, putative | 18.11 | no | 0 | 305.23 | 372.68 | 146.88 | 18.76 | 112.87 | 133.48 | 270.47 | T | 72.26 | 111.64 | F | large ribosomal subunit rRNA binding;rRNA binding;structural constituent of ribosome | ribosomal large subunit assembly;translation | cytosolic large ribosomal subunit;large ribosomal subunit;nucleus;ribosome |
| PF3D7_0422700 | EIF4A3 | eukaryotic initiation factor 4A-III, putative | 44.80 | no | 0 | 63.93 | 31.93 | 38.19 | 2.07 | 5.63 | 14.41 | 86.68 | O | 68.36 | 27.70 | M | RNA binding;RNA helicase activity;mRNA binding;translation initiation factor activity | regulation of translation;regulation of translational initiation | catalytic step 2 spliceosome;eukaryotic translation initiation factor 4F complex;nucleolus;nucleus |
| PF3D7_0815200 | N/A | importin subunit beta, putative | 99.96 | no | 0 | 80.94 | 34.55 | 40.57 | 4.74 | 7.58 | 25.51 | 61.61 | R | 79.78 | 22.95 | M | nuclear import signal receptor activity;nuclear localization sequence binding | NLS-bearing protein import into nucleus;protein import into nucleus | cytoplasm;nuclear pore;nucleus |
| PF3D7_1465900 | N/A | 40S ribosomal protein S3 | 24.67 | no | 0 | 256.94 | 318.02 | 161.44 | 15.43 | 79.36 | 127.59 | 573.21 | O | 208.09 | 546.87 | F | RNA binding;structural constituent of ribosome | translation | cytosolic small ribosomal subunit;extracellular vesicle;nucleus |
| PF3D7_1206200 | EIF3C | eukaryotic translation initiation factor 3 subunit C, putative | 116.01 | no | 0 | 352.53 | 186.86 | 66.18 | 21.86 | 15.58 | 19.57 | 102.70 | R | 156.82 | 56.50 | M | mRNA binding;translation initiation factor activity;translation initiation factor binding | regulation of translational initiation;translational initiation | eukaryotic translation initiation factor 3 complex;nucleus |
| PF3D7_1429900 | WRN | ADP-dependent DNA helicase RecQ | 169.54 | no | 0 | 11.82 | 5.98 | 11.13 | 2.72 | 0.79 | 25.42 | 66.19 | O | 165.82 | 101.35 | M | 3'-5' DNA helicase activity;DNA helicase activity;four-way junction helicase activity;protein binding | DNA duplex unwinding;DNA recombination;DNA repair;DNA unwinding involved in DNA replication;double-strand break repair via homologous recombination | chromosome;chromosome, telomeric region;cytoplasm;nucleus |
| PF3D7_1338100 | RPN3 | 26S proteasome regulatory subunit RPN3, putative | 59.07 | no | 0 | 98.98 | 64.68 | 135.80 | 130.06 | 12.21 | 33.34 | 96.02 | T | 103.08 | 90.99 | M | enzyme regulator activity | regulation of protein catabolic process | proteasome complex; nucleus; cytoplasm |
| PF3D7_1329000 | N/A | DNA-directed RNA polymerase III subunit RPC1, putative | 274.83 | no | 0 | 71.87 | 38.90 | 13.44 | 3.58 | 2.22 | 14.13 | 53.81 | R | 19.77 | 19.71 | 0 | DNA-directed 5'-3' RNA polymerase activity | transcription by RNA polymerase III | RNA polymerase III complex |
| PF3D7_0320800 | DOZI | ATP-dependent RNA helicase DDX6 | 49.41 | no | 0 | 6.50 | 20.89 | 54.61 | 11.23 | 47.25 | 163.76 | 237.14 | O | 110.82 | 263.97 | F | RNA helicase activity;mRNA binding;protein binding | P-body assembly;stress granule assembly | P-body;cytoplasm;cytoplasmic stress granule;nucleus |
| PF3D7_0510100 | N/A | KH domain-containing protein, putative | 297.57 | no | 0 | 6.52 | 7.19 | 38.61 | 10.19 | 2.95 | 17.42 | 30.71 | T | 91.56 | 10.18 | M | RNA binding | N/A | nucleus |
| PF3D7_0410800 | AKIT3 | apicomplexan kinetochore protein 3, putative | 150.27 | no | 0 | 10.68 | 11.15 | 55.02 | 17.66 | 11.24 | 38.06 | 107.14 | O | 322.61 | 77.92 | M | protein binding | cell division | kinetochore;nucleus |
| PF3D7_0814200 | ALBA1 | DNA/RNA-binding protein Alba 1 | 27.26 | no | 0 | 3045.36 | 2028.17 | 1052.00 | 492.03 | 221.15 | 158.23 | 1025.18 | R | 161.56 | 545.58 | F | DNA binding;RNA binding;mRNA binding;translation repressor activity, mRNA regulatory element binding | | cytoplasm;extracellular vesicle;food vacuole;nucleus |
| PF3D7_0620500 | N/A | cleavage stimulation factor subunit 1, putative | 76.49 | no | 0 | 31.44 | 15.11 | 9.28 | 2.22 | 2.95 | 12.55 | 20.34 | R | 5.21 | 17.95 | F | RNA binding | RNA processing | mRNA cleavage stimulating factor complex |
| PF3D7_0630600 | N/A | deubiquitinating enzyme MINDY, putative | 110.12 | no | 0 | 16.52 | 7.76 | 38.11 | 9.58 | 5.62 | 33.99 | 105.26 | O | 74.44 | 39.73 | M | thiol-dependent deubiquitinase;Lys48-specific deubiquitinase activity;Lys48-specific deubiquitinase activity | protein K48-linked deubiquitination | cytoplasm;nucleus |
| PF3D7_0515000 | CWC2 | pre-mRNA-splicing factor CWC2, putative | 98.38 | no | 0 | 80.85 | 64.26 | 86.26 | 54.79 | 6.78 | 29.88 | 138.15 | O | 133.81 | 57.80 | M | U6 snRNA binding;mRNA binding;pre-mRNA binding;protein binding | mRNA splicing, via spliceosome;positive regulation of cell cycle;spliceosomal snRNP assembly | Prp19 complex;U2-type catalytic step 1 spliceosome;U2-type catalytic step 2 spliceosome;nucleus;spliceosomal complex |
| PF3D7_1449500 | ApiAP2 | AP2 domain transcription factor AP2-O5, putative | 84.43 | no | 0 | 12.33 | 6.91 | 15.43 | 2.38 | 4.29 | 27.87 | 41.68 | O | 77.68 | 62.28 | M | DNA-binding transcription factor activity | regulation of transcription, DNA-templated;cell motility | heterochromatin;nucleus |
| PF3D7_0418200 | EIF3M | eukaryotic translation initiation factor 3 subunit M, putative | 51.03 | no | 0 | 94.39 | 40.37 | 29.60 | 4.56 | 10.95 | 32.90 | 33.66 | R | 140.78 | 21.84 | M | translation initiation factor activity | cytoplasmic translational initiation | eukaryotic translation initiation factor 3 complex;nucleus |
| PF3D7_1134000 | HSP70-3 | heat shock protein 70 | 73.30 | no | 0 | 95.08 | 272.42 | 123.14 | 7.53 | 20.29 | 21.48 | 163.92 | T | 371.73 | 193.18 | M | ATP binding;ATP hydrolysis activity;heat shock protein binding;misfolded protein binding;protein folding chaperone;unfolded protein binding | cellular response to unfolded protein;chaperone cofactor-dependent protein refolding;protein refolding;response to heat;response to unfolded protein | cytoplasm;food vacuole;mitochondrion;nucleus |
| PF3D7_1117700 | RAN | GTP-binding nuclear protein RAN/TC4 | 24.88 | no | 0 | 645.89 | 348.61 | 516.00 | 167.18 | 256.89 | 470.32 | 436.49 | R | 180.16 | 308.94 | F | GTP binding;GTPase activity | RNA export from nucleus;intracellular protein transport;protein import into nucleus;regulation of cell cycle;ribosomal subunit export from nucleus | cytoplasm;extracellular vesicle;food vacuole;nucleus |
| PF3D7_1205800 | HMGB3 | high mobility group protein B3, putative | 267.02 | no | 0 | 111.87 | 55.52 | 45.64 | 13.44 | 4.28 | 30.02 | 208.46 | O | 60.29 | 63.24 | 0 | DNA binding | N/A | nucleus |
| PF3D7_1007700 | AP2-I | AP2 domain transcription factor AP2-I | 182.66 | no | 0 | 35.10 | 204.32 | 285.73 | 99.61 | 2.49 | 14.17 | 189.67 | T | 299.60 | 65.11 | M | protein binding;sequence-specific DNA binding | response to xenobiotic stimulus | Maurer's cleft;nucleus;protein-containing complex |
| PF3D7_0218000 | RFC2 | replication factor C subunit 2, putative | 37.92 | no | 0 | 8.69 | 6.06 | 58.69 | 10.39 | 13.35 | 40.20 | 886.79 | O | 241.59 | 33.82 | M | RNA binding | DNA repair;DNA replication;DNA-dependent DNA replication | DNA replication factor C complex;nucleus |
| PF3D7_0507700 | NPL4 | nuclear protein localization protein 4, putative | 62.57 | no | 0 | 56.39 | 19.95 | 28.39 | 4.74 | 5.56 | 32.48 | 166.21 | O | 48.98 | 38.85 | M | ubiquitin binding;ubiquitin protein ligase binding | ubiquitin-dependent ERAD pathway;ubiquitin-dependent protein catabolic process | nucleus |
| PF3D7_1017900 | RPN5 | 26S proteasome regulatory subunit p55, putative | 55.21 | no | 0 | 81.76 | 28.08 | 53.35 | 20.16 | 6.70 | 38.10 | 62.76 | 81.76 | 26.05 | 48.36 | F | endopeptidase activity | proteasome-mediated ubiquitin-dependent protein catabolic process;ubiquitin-dependent protein catabolic process | cytoplasm;nuclear proteasome complex;nucleus;proteasome regulatory particle;proteasome regulatory particle, lid subcomplex |

| | | | | | | | | | | | | | | | | | | | |
|---|---|---|---|---|---|---|---|---|---|---|---|---|---|---|---|---|---|---|---|
| PF3D7_0517400 | FACT-L | FACT complex subunit SPT16, putative | 132.68 | no | 0 | 139.58 | 175.55 | 174.46 | 143.00 | 27.06 | 23.39 | 1056.64 | O | 484.10 | 74.42 | M | nucleosome binding | DNA replication-independent chromatin organization;positive regulation of transcription elongation from RNA polymerase II promoter;transcription elongation from RNA polymerase II promoter;transcription, DNA-templated | FACT complex;nucleus |
| PF3D7_1456500 | N/A | STAG domain-containing protein, putative | 283.20 | no | 0 | 5.28 | 2.08 | 13.94 | 2.25 | 2.51 | 88.83 | 66.84 | GV | 134.03 | 118.31 | M | protein binding | N/A | cohesin complex;nucleus |
| PF3D7_0406100 | N/A | V-type proton ATPase subunit B | 55.79 | no | 0 | 327.22 | 176.18 | 222.98 | 35.39 | 39.39 | 60.60 | 320.99 | R | 257.44 | 218.04 | 0 | protein binding;proton-transporting ATPase activity, rotational mechanism | proton-transporting V-type ATPase complex assembly;vacuolar acidification | cell periphery;food vacuole;nucleus |
| PF3D7_0309600 | PfP2 | 60S acidic ribosomal protein P2 | 11.95 | no | 0 | 317.26 | 634.57 | 246.28 | 172.14 | 305.70 | 170.07 | 584.90 | T | 301.99 | 315.73 | F | RNA binding | entry into host;protein tetramerization | Maurer's cleft;cell surface;host cell surface;nucleus |
| PF3D7_1103800 | NOT1-G | CCR4-NOT transcription complex subunit NOT1-G, putative | 391.94 | no | 4 | 7.06 | 8.95 | 17.42 | 2.72 | 5.79 | 50.75 | 124.88 | O | 380.35 | 147.15 | M | mRNA binding;molecular adaptor activity | negative regulation of translation;nuclear-transcribed mRNA catabolic process, deadenylation-dependent decay | CCR4-NOT complex;CCR4-NOT core complex;P-body;cytosol;nucleus |
| PF3D7_1328200 | N/A | BRCA2 protein, putative | 320.13 | no | 0 | 11.49 | 1.29 | 7.23 | 2.38 | 1.15 | 10.80 | 100.26 | O | 29.68 | 11.97 | M | N/A | DNA repair | nucleus |
| PF3D7_0610900 | SPT5 | transcription elongation factor SPT5, putative | 149.36 | no | 0 | 144.61 | 74.80 | 24.35 | 3.25 | 15.75 | 32.14 | 146.31 | O | 213.18 | 73.26 | M | mRNA binding | regulation of transcription by RNA polymerase II;transcription elongation from RNA polymerase II promoter;transcription, DNA-templated | DSIF complex;nucleus |
| PF3D7_0605100 | KH1 | KH domain-containing protein, putative | 85.63 | no | 0 | 94.39 | 30.37 | 24.80 | 21.69 | 8.18 | 31.37 | 150.30 | O | 110.44 | 21.84 | M | N6-methyladenosine-containing RNA binding;RNA binding;mRNA binding | regulation of gene expression | cytoplasm;nucleus |
| PF3D7_1322200 | STU2 | protein STU2, putative | 402.41 | no | 0 | 7.42 | 6.79 | 22.70 | 9.88 | 9.18 | 46.69 | 172.04 | O | 302.30 | 31.79 | M | N/A | N/A | kinetochore;nucleus |
| PF3D7_1417200 | NOT1 | CCR4-NOT transcription complex subunit NOT1, putative | 519.82 | no | 0 | 89.22 | 42.65 | 19.22 | 3.54 | 6.25 | 51.71 | 134.31 | O | 53.47 | 50.80 | 0 | mRNA binding;molecular adaptor activity | negative regulation of translation;nuclear-transcribed mRNA catabolic process, deadenylation-dependent decay | CCR4-NOT complex;CCR4-NOT core complex;P-body;cytosol;nucleus |
| PF3D7_0501000 | N/A | Plasmodium exported protein, unknown function | 30.72 | no | 1 | 97.26 | 295.98 | 76.18 | 2.87 | 30.52 | 7.85 | 15.94 | T | 7.29 | 0.63 | M | N/A | N/A | Maurer's cleft;extracellular vesicle;nucleus |
| PF3D7_0302000 | PRP46 | pre-mRNA-splicing factor PRP46, putative | 69.57 | no | 0 | 81.27 | 35.17 | 22.89 | 2.60 | 7.81 | 22.93 | 47.13 | R | 95.92 | 40.16 | M | RNA binding | mRNA splicing, via spliceosome | Cul4-RING E3 ubiquitin ligase complex;Prp19 complex;catalytic step 2 spliceosome;spliceosomal complex |
| PF3D7_1230800 | WTAP | pre-mRNA-splicing regulator WTAP, putative | 71.31 | no | 0 | 108.15 | 43.58 | 19.74 | 2.49 | 6.32 | 15.78 | 168.80 | O | 242.04 | 120.20 | M | protein binding | mRNA methylation;regulation of alternative mRNA splicing, via spliceosome | RNA N6-methyladenosine methyltransferase complex;nucleus |
| PF3D7_0810800 | PPPK-DHPS | hydroxymethyldihydropterin pyrophosphokinase-dihydropteroate synthase | 83.37 | no | 0 | 20.91 | 15.29 | 10.16 | 0.87 | 0.96 | 1.90 | 7.46 | R | 14.14 | 2.96 | M | dihydropteroate synthase activity | dihydrofolate biosynthetic process;response to xenobiotic stimulus;tetrahydrofolate biosynthetic process | mitochondrial envelope;nucleus |
| PF3D7_1421200 | RPS25 | 40S ribosomal protein S25 | 11.66 | no | 0 | 273.22 | 496.15 | 161.45 | 100.96 | 318.88 | 181.82 | 299.03 | T | 163.83 | 244.34 | F | structural constituent of ribosome | translation | cytosolic small ribosomal subunit;nucleus |
| PF3D7_1030500 | RPN9 | 26S proteasome regulatory subunit RPN9, putative | 46.80 | no | 0 | 18.61 | 9.68 | 15.02 | 4.31 | 2.13 | 17.23 | 10.99 | R | 8.72 | 9.74 | 0 | endopeptidase activity;structural molecule activity | proteasome assembly;ubiquitin-dependent protein catabolic process | cytosol;nucleus;proteasome regulatory particle;proteasome regulatory particle, lid subcomplex |
| PF3D7_0507100 | RPL4 | 60S ribosomal protein L4 | 46.21 | no | 0 | 1015.63 | 1201.12 | 449.42 | 114.39 | 19.70 | 33.01 | 1682.00 | O | 103.13 | 153.31 | F | RNA binding;structural constituent of ribosome | translation | cytosolic large ribosomal subunit;nucleus |
| PF3D7_0629400 | PABP3 | polyadenylate-binding protein 3, putative | 90.78 | no | 0 | 7.40 | 22.89 | 26.01 | 1.72 | 4.84 | 27.03 | 8.94 | GV | 74.20 | 42.02 | M | RNA binding;mRNA binding | N/A | cytosol;nucleus |
| PF3D7_0317600 | RPS11 | 40S ribosomal protein S11, putative | 18.76 | no | 0 | 719.09 | 557.12 | 229.51 | 44.62 | 128.79 | 161.81 | 689.78 | R | 72.78 | 405.01 | F | structural constituent of ribosome | translation | cytosolic small ribosomal subunit;extracellular vesicle;nucleus |
| PF3D7_0305600 | APE1 | DNA-(apurinic or apyrimidinic site) endonuclease | 72.44 | no | 0 | 7.32 | 1.45 | 22.35 | 7.53 | 3.58 | 102.14 | 172.82 | O | 368.56 | 508.80 | F | 3'-5' exonuclease activity;DNA-(apurinic or apyrimidinic site) endonuclease activity;double-stranded DNA 3'-5' exodeoxyribonuclease activity;phosphatase activity;phosphodiesterase I activity;ribonuclease activity | DNA repair;base-excision repair;entry into host | mitochondrion;nucleus |
| PF3D7_1427500 | MSH2-1 | DNA mismatch repair protein MSH2, putative | 94.74 | no | 0 | 3.85 | 2.31 | 17.95 | 2.51 | 0.90 | 3.35 | 9.92 | T | 3.36 | 2.14 | 0 | ATP-dependent activity, acting on DNA;DNA binding;damaged DNA binding;double-strand/single-strand DNA junction binding | DNA recombination;maintenance of DNA repeat elements;mismatch repair;negative regulation of DNA recombination;postreplication repair | MutSalpha complex;MutSbeta complex;mismatch repair complex;nucleus |
| PF3D7_1333600 | UTP4 | U3 small nucleolar RNA-associated protein 4, putative | 159.34 | no | 1 | 118.79 | 83.82 | 11.71 | 1.78 | 4.58 | 20.87 | 22.24 | R | 41.72 | 24.81 | F | protein binding | maturation of SSU-rRNA from tricistronic rRNA transcript (SSU-rRNA, 5.8S rRNA, LSU-rRNA) | 90S preribosome;membrane;nucleolus;small-subunit processome;t-UTP complex |
| PF3D7_1474800 | N/A | proteasome subunit alpha type-1, putative | 28.84 | no | 0 | 102.83 | 39.66 | 90.00 | 14.65 | 29.03 | 67.83 | 201.62 | O | 150.12 | 52.39 | F | endopeptidase activity | proteasomal protein catabolic process;proteasomal ubiquitin-independent protein catabolic process;proteasome-mediated ubiquitin-dependent protein catabolic process;ubiquitin-dependent protein catabolic process | cytoplasm;nucleus;proteasome core complex;proteasome core complex, alpha-subunit complex |
| PF3D7_1245100 | KLP8 | kinesin-13, putative | 153.12 | no | 0 | 3.53 | 0.56 | 11.61 | 2.52 | 2.73 | 30.72 | 63.98 | O | 112.10 | 28.28 | 0 | cytoskeletal motor activity | axoneme assembly;microtubule-based movement | cytoplasm;kinesin complex;kinetochore;nucleus |
| PF3D7_1027800 | RPL3 | 60S ribosomal protein L3 | 44.22 | no | 0 | 770.37 | 705.27 | 466.74 | 64.19 | 52.59 | 45.18 | 446.97 | R | 211.93 | 219.59 | F | RNA binding;structural constituent of ribosome | ribosomal large subunit assembly;translation | cytosolic large ribosomal subunit;intracellular;nucleus |
| PF3D7_1424100 | RPL5 | 60S ribosomal protein L5, putative | 34.00 | no | 0 | 504.89 | 589.18 | 262.21 | 53.34 | 13.25 | 26.40 | 437.26 | T | 146.76 | 106.81 | M | 5S rRNA binding;RNA binding;structural constituent of ribosome | ribosomal large subunit assembly;translation | cytosolic large ribosomal subunit;extracellular vesicle;nucleus |
| PF3D7_0212400 | NUP390 | nucleoporin NUP390, putative | 494.21 | no | 7 | 18.97 | 13.56 | 9.58 | 1.65 | 3.32 | 21.83 | 26.29 | O | 18.83 | 7.73 | M | protein binding | nucleocytoplasmic transport | nuclear periphery |
| PF3D7_0608500 | N/A | proteasome subunit alpha type-2, putative | 26.53 | no | 0 | 27.23 | 14.55 | 40.38 | 9.35 | 14.22 | 57.75 | 75.37 | O | 61.22 | 53.59 | M | endopeptidase activity | proteasomal protein catabolic process;proteasomal ubiquitin-independent protein catabolic process;proteasome-mediated ubiquitin-dependent protein catabolic process;ubiquitin-dependent protein catabolic process | cytoplasm;extracellular vesicle;nucleus;proteasome core complex;proteasome core complex, alpha-subunit complex |
| PF3D7_0802100 | AP2-LT | AP2 domain transcription factor AP2-LT | 299.29 | no | 0 | 7.72 | 1.64 | 39.37 | 26.49 | 7.16 | 32.48 | 220.82 | O | 142.23 | 54.00 | M | sequence-specific DNA binding | regulation of transcription | histone acetyltransferase complex;nucleus |
| PF3D7_0821600 | CLP1 | polyribonucleotide 5'-hydroxyl-kinase Clp1, putative | 79.06 | no | 0 | 51.37 | 17.43 | 11.49 | 0.87 | 3.16 | 21.91 | 81.13 | O | 12.29 | 27.11 | F | polydeoxyribonucleotide kinase activity;polynucleotide 5'-hydroxyl-kinase activity | RNA processing;mRNA polyadenylation;tRNA splicing, via endonucleolytic cleavage and ligation | nucleus;tRNA-intron endonuclease complex |
| PF3D7_1129200 | RPN7 | 26S proteasome regulatory subunit RPN7, putative | 46.62 | no | 0 | 73.85 | 21.92 | 67.53 | 14.76 | 9.22 | 42.44 | 107.29 | O | 23.62 | 65.43 | F | endopeptidase activity | proteasome-mediated ubiquitin-dependent protein catabolic process;proteolysis | nucleus;proteasome complex |
| PF3D7_0811500 | CARM1 | histone-arginine methyltransferase CARM1, putative | 108.43 | no | 0 | 48.98 | 10.74 | 3.63 | 0.68 | 1.74 | 7.54 | 5.48 | R | 3.06 | 3.00 | 0 | histone methyltransferase activity | histone methylation | nucleus |
| PF3D7_1338300 | EF-1gamma | elongation factor 1-gamma, putative | 47.78 | no | 0 | 488.79 | 165.11 | 106.59 | 26.11 | 13.81 | 67.05 | 138.72 | R | 10.21 | 114.70 | F | glutathione transferase activity;translation elongation factor activity | translational elongation | eukaryotic translation elongation factor 1 complex;nucleus |
| PF3D7_0316800 | N/A | 40S ribosomal protein S15A, putative | 14.87 | no | 0 | 237.36 | 212.36 | 135.61 | 19.70 | 225.81 | 167.35 | 275.29 | O | 159.49 | 273.05 | F | RNA binding;structural constituent of ribosome | translation | cytosolic small ribosomal subunit;nucleus |
| PF3D7_0819000 | N/A | ribonuclease, putative | 157.67 | no | 0 | 77.37 | 25.93 | 9.81 | 0.73 | 2.69 | 22.02 | 93.05 | O | 54.18 | 27.29 | M | RNA binding | mRNA 3'-end processing by stem-loop binding and cleavage;mRNA polyadenylation;pre-mRNA cleavage required for polyadenylation | mRNA cleavage and polyadenylation specificity factor complex |
| PF3D7_1006700 | N/A | conserved Plasmodium protein, unknown function | 45.62 | no | 0 | 39.04 | 24.62 | 12.39 | 0.44 | 14.69 | 100.71 | 181.79 | O | 265.79 | 363.06 | F | N/A | N/A | nucleus |
| PF3D7_1110200 | PRPF6 | pre-mRNA-processing factor 6, putative | 152.15 | no | 0 | 172.13 | 70.74 | 43.29 | 7.94 | 3.00 | 12.88 | 157.43 | R | 72.23 | 49.06 | M | RNA bindding | mRNA splicing, via spliceosome;spliceosomal tri-snRNP complex assembly | U4/U6 x U5 tri-snRNP complex;catalytic step 2 spliceosome |
| PF3D7_1136500 | CK1 | Casein kinase 1 | 37.79 | no | 0 | 756.48 | 111.09 | 331.05 | 88.97 | 93.86 | 156.19 | 928.77 | O | 895.88 | 546.88 | M | protein serine/threonine kinase activity | endocytosis;peptidyl-serine phosphorylation | cytoplasm;nucleus |
| PF3D7_0811400 | N/A | conserved protein, unknown function | 71.37 | no | 0 | 179.07 | 76.29 | 53.67 | 13.10 | 10.31 | 12.55 | 63.45 | R | 149.38 | 25.93 | M | N/A | N/A | nucleus |
| PF3D7_1355700 | NIF3 | NLI interacting factor-like phosphatase, putative | 151.54 | no | 0 | 63.86 | 32.67 | 22.22 | 5.36 | 8.66 | 42.63 | 25.36 | R | 19.83 | 18.81 | 0 | RNA polymerase II CTD heptapeptide repeat phosphatase activity | dephosphorylation of RNA polymerase II C-terminal domain | nucleus |
| PF3D7_1013600 | N/A | Uncharacterized protein | 269.27 | no | 0 | 27485.00 | 28856.00 | 45363.00 | 27454.00 | 28856.00 | 15.38 | 29921.00 | T | 39.93 | 19.54 | M | | | nucleus |
| PF3D7_1241700 | RFC4 | replication factor C subunit 4, putative | 37.64 | no | 0 | 8.96 | 5.46 | 35.56 | 11.77 | 8.63 | 24.56 | 49.68 | O | 130.71 | 54.65 | M | ATP binding;DNA clamp loader activity | DNA repair;DNA replication;DNA-dependent DNA replication;leading strand elongation;response to xenobiotic stimulus | DNA replication factor C complex;nucleus |
| PF3D7_0529400 | AKIT5 | apicomplexan kinetochore protein 5, putative | 185.37 | no | 0 | 11.01 | 18.82 | 62.00 | 31.27 | 9.27 | 38.11 | 461.55 | O | 288.64 | 93.96 | M | N/A | N/A | nucleus |
| PF3D7_1422400 | N/A | nucleolar RNA-associated protein, putative | 195.36 | no | 0 | 36.18 | 14.53 | 3.87 | 0.89 | 2.11 | 14.64 | 30.65 | O | 36.45 | 18.82 | M | RNA binding | rRNA processing;tRNA export from nucleus | CURI complex;UTP-C complex;nucleolus;small-subunit processome |
| PF3D7_0730500 | N/A | conserved Plasmodium protein, unknown function | 329.46 | no | 0 | 5.58 | 0.92 | 10.46 | 3.78 | 2.68 | 26.83 | 67.63 | O | 19.23 | 47.57 | F | N/A | N/A | nucleus |
| PF3D7_1004300 | RFUL | RING finger E3 ubiquitin-protein ligase | 131.70 | no | 0 | 166.81 | 29.31 | 59.63 | 14.23 | 9.65 | 88.75 | 244.47 | O | 180.51 | 739.51 | F | ubiquitin protein ligase activity | protein ubiquitination;ubiquitin-dependent protein catabolic process | cytoplasm;nucleus |
| PF3D7_0607000 | N/A | translation initiation factor IF-2, putative | 112.36 | no | 0 | 262.17 | 174.02 | 64.16 | 15.05 | 9.03 | 23.82 | 207.45 | R | 175.02 | 100.50 | M | translation initiation factor activity | translational initiation | nucleus |
| PF3D7_0910100 | N/A | exportin-7, putative | 139.18 | no | 0 | 37.56 | 8.20 | 37.48 | 8.39 | 4.83 | 7.55 | 13.01 | R | 40.83 | 8.46 | M | nuclear export signal receptor activity | plastid organization;protein export from nucleus | cytoplasm;nuclear pore;nucleus |
| PF3D7_1343300 | N/A | CDT1-like protein, putative | 114.58 | no | 0 | 8.77 | 13.07 | 34.37 | 6.99 | 7.79 | 46.36 | 129.45 | O | 342.31 | 59.51 | M | N/A | N/A | nucleus |
| PF3D7_1422800 | ARP4 | actin-related protein ARP4 | 60.66 | no | 0 | 15.09 | 5.60 | 8.36 | 0.84 | 5.33 | 13.50 | 43.63 | O | 38.95 | 9.25 | M | chromatin binding;protein binding;structural constituent of cytoskeleton | ATP-dependent chromatin remodeling;chromatin remodeling;cytoskeleton organization;endocytosis;exocytosis;histone H4 acetylation;regulation of gene expression;regulation of transcription by RNA polymerase II;schizogony | NuA4 histone acetyltransferase complex;SWI/SNF complex;actin filament;nucleus |
| PF3D7_1118500 | NOP56 | nucleolar protein 56, putative | 69.39 | no | 0 | 411.95 | 138.07 | 42.30 | 11.69 | 21.43 | 15.69 | 130.05 | R | 401.74 | 99.98 | M | snoRNA binding | rRNA processing | box C/D RNP complex;nucleus;small-subunit processome |
| PF3D7_1457300 | N/A | MA3 domain-containing protein, putative | 75.99 | no | 0 | 182.45 | 36.58 | 26.15 | 4.41 | 4.98 | 44.93 | 31.83 | R | 9.68 | 17.11 | F | translation initiation factor binding | translation | apicoplast;nucleus |
| PF3D7_0212100 | N/A | conserved Plasmodium protein, unknown function | 214.68 | no | 0 | 49.63 | 26.06 | 45.75 | 11.10 | 8.00 | 14.18 | 180.40 | O | 75.13 | 26.74 | M | N/A | N/A | nuclear periphery;nucleus |
| PF3D7_1406200 | SPT6 | transcription elongation factor SPT6, putative | 338.38 | no | 0 | 102.71 | 61.35 | 21.10 | 2.44 | 5.82 | 29.94 | 60.85 | O | 224.56 | 96.85 | M | histone binding;nucleosome binding | mRNA transcription by RNA polymerase II;nucleosome organization;obsolete chromatin maintenance;positive regulation of transcription elongation from RNA polymerase II promoter;regulation of mRNA processing;transcription elongation from RNA polymerase II promoter | nucleus;transcription elongation factor complex;transcriptionally active chromatin |
| PF3D7_0407800 | CINCH | protein CINCH | 230.12 | no | 0 | 5.14 | 0.48 | 36.03 | 22.44 | 4.14 | 43.47 | 225.65 | O | 46.25 | 176.58 | F | protein binding | cell division | basal part of cell;inner membrane pellicle complex;nucleus |
| PF3D7_1105400 | RPS4 | 40S ribosomal protein S4, putative | 29.77 | no | 0 | 288.16 | 391.25 | 236.48 | 50.29 | 86.13 | 104.03 | 334.58 | T | 86.41 | 114.11 | F | RNA binding;structural constituent of ribosome | translation | cytosolic small ribosomal subunit;extracellular vesicle;nucleus;small ribosomal subunit |
| PF3D7_0612900 | NOG1 | nucleolar GTP-binding protein 1, putative | 79.53 | no | 0 | 424.38 | 254.88 | 72.02 | 12.96 | 26.81 | 55.44 | 284.10 | R | 52.09 | 385.59 | F | GTP binding | ribosomal large subunit biogenesis | nucleolus;preribosome, large subunit precursor |
| PF3D7_1446500 | NUP313 | nucleoporin NUP313, putative | 346.92 | no | 0 | 69.27 | 31.12 | 23.23 | 1.91 | 3.37 | 27.38 | 52.11 | R | 63.87 | 19.05 | M | protein binding | nucleocytoplasmic transport | nuclear pore;nucleus |
| PF3D7_1463200 | RFC3 | replication factor C subunit 3, putative | 39.23 | no | 0 | 4.54 | 3.54 | 45.00 | 6.50 | 6.03 | 11.15 | 8.77 | T | 141.14 | 15.91 | M | DNA binding | DNA repair;DNA replication;DNA strand elongation involved in DNA replication;DNA-dependent DNA replication;leading strand elongation | DNA replication factor C complex;PCNA complex;nucleus |
| PF3D7_1441200 | N/A | 60S ribosomal protein L1, putative | 24.81 | no | 0 | 66.45 | 52.63 | 24.83 | 3.54 | 14.53 | 36.07 | 21.44 | R | 30.17 | 15.71 | M | RNA binding;structural constituent of ribosome | maturation of LSU-rRNA;translation | cytosolic large ribosomal subunit;intracellular;nucleus |
| PF3D7_1466400 | ApiAP2 | AP2 domain transcription factor AP2-EXP | 92.09 | no | 0 | 200.36 | 29.73 | 20.00 | 2.82 | 3.59 | 14.47 | 15.52 | R | 64.32 | 9.47 | M | sequence-specific DNA binding | positive regulation of gene expression;response to xenobiotic stimulus | heterochromatin;nucleus |
| PF3D7_0704400 | N/A | phosphoinositide-binding protein, putative | 118.84 | no | 0 | 7.37 | 2.67 | 15.19 | 4.91 | 1.25 | 23.51 | 7.79 | GV | 22.36 | 56.25 | F | transcription coactivator activity | histone H3 acetylation;regulation of transcription, DNA-templated;transcription initiation from RNA polymerase II promoter | nucleus;transcription factor TFIID complex |
| PF3D7_1412100 | MCMBP | mini-chromosome maintenance complex-binding protein | 112.77 | no | 0 | 8.12 | 5.38 | 15.08 | 6.86 | 1.21 | 7.49 | 16.46 | O | 27.05 | 5.82 | M | chromatin binding | DNA-dependent DNA replication;schizogony | MCM complex;cytoplasm;nucleus |
| PF3D7_1345800 | N/A | conserved Plasmodium protein, unknown function | 118.92 | no | 0 | 33.18 | 23.32 | 21.71 | 4.63 | 6.31 | 24.30 | 17.32 | R | 109.60 | 17.73 | M | N/A | N/A | nucleus |
| PF3D7_0210200 | N/A | Uncharacterized protein | 310.02 | no | 0 | 22586.00 | 22859.00 | 29.31 | 32721.00 | 19603.00 | 45.48 | 53.73 | S | 548.18 | 52.03 | M | N/A | N/A | nucleolus;nucleus |
| PF3D7_1206600 | RPC2 | DNA-directed RNA polymerase III subunit RPC2, putative | 167.29 | no | 0 | 95.46 | 39.26 | 14.03 | 1.06 | 2.24 | 23.32 | 71.43 | R | 38.00 | 44.63 | F | DNA-directed 5'-3' RNA polymerase activity | transcription by RNA polymerase III | RNA polymerase III complex |
| PF3D7_1405600 | RNR | ribonucleoside-diphosphate reductase small chain, putative | 40.60 | no | 0 | 6.45 | 6.44 | 43.54 | 10.75 | 27.90 | 32.33 | 118.87 | O | 494.07 | 46.18 | M | ribonucleoside-diphosphate reductase activity, thioredoxin disulfide as acceptor | deoxyribonucleotide biosynthetic process | intracellular;ribonucleoside-diphosphate reductase complex;cytoplasm;nucleus |
| PF3D7_1367100 | SNP1 | U1 small nuclear ribonucleoprotein 70 kDa homolog, putative | 51.84 | no | 0 | 411.70 | 108.40 | 99.27 | 49.07 | 15.74 | 12.26 | 268.47 | R | 262.26 | 101.69 | M | RNA binding;U1 snRNA binding;mRNA binding;snRNA binding | RNA processing;mRNA splicing, via spliceosome | U1 snRNP;U2-type prespliceosome;spliceosomal complex |
| PF3D7_1466800 | N/A | NOC3 domain-containing protein, putative | 105.45 | no | 0 | 25.90 | 12.41 | 4.08 | 0.23 | 47.43 | 116.99 | 628.46 | O | 19.05 | 307.57 | F | chromatin binding | N/A | nucleus |
| PF3D7_0509100 | SMC4 | structural maintenance of chromosomes protein 4, putative | 201.67 | no | 0 | 12.70 | 5.30 | 49.19 | 17.16 | 9.97 | 12.86 | 132.53 | O | 749.06 | 46.70 | M | chromatin binding | mitotic chromosome condensation | chromosome, centromeric region;condensin complex;nucleus |
| PF3D7_1211400 | PfJ4 | heat shock protein DNAJ homologue Pfj4 | 28.06 | no | 0 | 170.37 | 50.43 | 83.14 | 8.07 | 162.81 | 95.63 | 127.01 | O | 213.38 | 92.31 | M | Hsp70 protein binding;unfolded protein binding | response to heat;response to unfolded protein | Maurer's cleft;cytoplasm;host cell cytoplasm;nucleus |
| PF3D7_1302800 | N/A | 40S ribosomal protein S7, putative | 22.48 | no | 0 | 99.48 | 117.31 | 46.10 | 4.25 | 26.45 | 145.33 | 204.69 | O | 90.68 | 82.38 | M | RNA binding;structural constituent of ribosome | rRNA processing;ribosomal small subunit biogenesis;translation | cytosolic small ribosomal subunit;nucleus;small-subunit processome |
| PF3D7_1445900 | DDX17 | ATP-dependent RNA helicase DDX17 | 60.04 | no | 0 | 425.42 | 127.30 | 77.55 | 7.59 | 21.68 | 59.39 | 267.15 | R | 240.31 | 119.07 | M | ATP hydrolysis activity;RNA binding;RNA helicase activity;helicase activity;nucleic acid binding | | cytoplasm;host cell membrane;nucleus;ribonucleoprotein complex |
| PF3D7_0711500 | N/A | regulator of chromosome condensation, putative | 78.90 | no | 0 | 27.37 | 30.63 | 28.12 | 2.71 | 4.71 | 18.73 | 50.38 | O | 71.90 | 35.29 | M | chromatin binding | Ran protein signal transduction;chromosome condensation;chromosome segregation;regulation of mitotic spindle assembly | Maurer's cleft;cytoplasm;nucleus |
| PF3D7_0521700 | DDX1 | ATP-dependent RNA helicase DDX1, putative | 97.34 | no | 0 | 15.61 | 3.91 | 4.11 | 0.65 | 1.15 | 4.03 | 6.91 | R | 11.86 | 6.42 | M | RNA binding | rRNA processing | nucleus |
| PF3D7_0807100 | PSH3 | DNA helicase PSH3 | 157.20 | no | 0 | 16.63 | 13.93 | 9.47 | 2.40 | 4.95 | 45.14 | 34.89 | GV | 66.75 | 31.31 | M | 3'-5' DNA helicase activity;ATP hydrolysis activity;RNA binding;RNA helicase activity;single-stranded DNA helicase activity | DNA duplex unwinding | cytoplasm;nucleus |
| PF3D7_1004200 | N/A | WD repeat-containing protein, putative | 358.07 | no | 4 | 12.85 | 12.49 | 23.40 | 8.26 | 2.93 | 21.24 | 37.30 | O | 25.21 | 43.36 | F | protein binding | RNA splicing | nucleus;precatalytic spliceosome;integral component of membrane |
| PF3D7_1475600 | BDP4 | bromodomain protein 4, putative | 85.80 | no | 0 | 54.56 | 18.05 | 13.16 | 2.03 | 2.69 | 13.61 | 21.76 | R | 33.04 | 29.24 | M | protein binding | regulation of transcription | nucleus |
| PF3D7_1357500 | N/A | DNA helicase, putative | 106.29 | no | 0 | 10.13 | 5.57 | 23.90 | 8.13 | 2.70 | 5.64 | 23.43 | O | 131.45 | 10.02 | M | ATP-dependent DNA/DNA annealing activity;ATP-dependent activity, acting on DNA;DNA helicase activity | DNA repair;DNA rewinding;cellular response to DNA damage stimulus;replication fork processing;replication fork protection | nuclear replication fork;nucleus |
| PF3D7_1120100 | PGM1 | phosphoglycerate mutase, putative | 28.77 | no | 0 | 1236.80 | 235.64 | 358.89 | 60.81 | 38.32 | 33.55 | 82.78 | R | 197.09 | 52.49 | M | phosphoglycerate mutase activity;protein binding | glycolytic process | Maurer's cleft;cytosol;nucleus |
| PF3D7_1242700 | N/A | 40S ribosomal protein S17, putative | 16.07 | no | 0 | 273.91 | 363.31 | 204.11 | 14.89 | 277.19 | 324.95 | 425.63 | O | 243.60 | 196.71 | M | RNA binding;structural constituent of ribosome | translation | cytosolic small ribosomal subunit;mitochondrion;nucleus |
| PF3D7_0811300 | CAF1 | CCR4-associated factor 1 | 200.21 | no | 0 | 402.20 | 70.06 | 117.56 | 38.84 | 12.49 | 69.82 | 206.03 | R | 129.59 | 48.34 | M | poly(A)-specific ribonuclease activity | exonucleolytic catabolism of deadenylated mRNA;negative regulation of translation | CCR4-NOT complex;CCR4-NOT core complex;P-body;cytosol;nucleus |

| | | | | | | | | | | | | | | | | | | | |
|---|---|---|---|---|---|---|---|---|---|---|---|---|---|---|---|---|---|---|---|
| PF3D7_0517700 | EIF3B | eukaryotic translation initiation factor 3 subunit B, putative | 84.06 | no | 0 | 162.41 | 84.94 | 55.68 | 3.89 | 11.43 | 23.64 | 136.98 | R | 62.62 | 55.88 | M | protein binding;RNA binding; translation initiation factor activity | translational initiation | eukaryotic translation initiation factor 3 complex;nucleus |
| PF3D7_0612100 | EIF3L | eukaryotic translation initiation factor 3 subunit L, putative | 78.13 | no | 2 | 117.96 | 23.59 | 30.02 | 4.76 | 1.47 | 9.55 | 19.84 | R | 31.75 | 21.04 | M | protein binding;RNA binding; translation initiation factor activity | translational initiation | eukaryotic translation initiation factor 3 complex;nucleus |
| PF3D7_1330600 | N/A | elongation factor Tu, putative | 57.63 | no | 0 | 7.98 | 6.81 | 48.77 | 8.67 | 5.87 | 31.84 | 200.76 | O | 45.31 | 199.91 | F | GTP binding;mRNA binding;translation elongation factor activity | mitochondrial translational elongation;translational elongation | mitochondrion;nucleus |
| PF3D7_1211800 | PfpUB | polyubiquitin | 14.62 | no | 0 | 69.15 | 43.1 | 111.26 | 64.84 | 45615.00 | 46.64 | 72.64 | GC II | 1283.13 | 1053.03 | M | protein tag;ubiquitin protein ligase binding | modification-dependent protein catabolic process;protein polyubiquitination;protein ubiquitination;ubiquitin-dependent protein catabolic process | cell surface;cytoplasm;nucleus;ubiquitin ligase complex |
| PF3D7_1316500 | PRP40 | pre-mRNA-processing factor 40, putative | 106.12 | no | 0 | 30.88 | 26.83 | 22.97 | 2.56 | 2.73 | 10.38 | 58.69 | O | 47.37 | 27.64 | M | protein binding;RNA binding | mRNA cis splicing, via spliceosome | spliceosomal complex;nucleus |
| PF3D7_0516200 | RPS11 | 40S ribosomal protein S11 | 16.08 | no | 0 | 390.82 | 422.05 | 175.46 | 68.46 | 193.51 | 161.36 | 117.78 | T | 110.70 | 172.59 | F | RNA binding | translation | cytosolic small ribosomal subunit;extracellular vesicle;nucleus |
| PF3D7_0807800 | RPN10 | 26S proteasome regulatory subunit RPN10, putative | 55.07 | no | 0 | 204.37 | 39.57 | 80.09 | 26.26 | 11.01 | 103.98 | 331.06 | O | 149.45 | 159.35 | F | polyubiquitin modification-dependent protein binding | proteasome assembly;proteasome-mediated ubiquitin-dependent protein catabolic process | cytosol;nucleus;proteasome regulatory particle, base subcomplex |
| PF3D7_1342000 | N/A | 40S ribosomal protein S6 | 35.38 | no | 0 | 373.69 | 306.10 | 149.68 | 19.30 | 14.43 | 36.14 | 266.82 | R | 391.35 | 213.34 | M | structural constituent of ribosome | translation | cytosolic small ribosomal subunit;nucleus |
| PF3D7_0414000 | SMC3 | structural maintenance of chromosomes protein 3 | 141.22 | no | 0 | 4.75 | 6.70 | 16.43 | 2.39 | 4.70 | 42.00 | 98.77 | O | 167.87 | 111.69 | M | ATP binding;centromeric DNA binding;chromatin binding;minor groove of adenine-thymine-rich DNA binding;protein binding | chromosome organization;mitotic sister chromatid segregation;regulation of DNA replication | chromatin;chromosome, centromeric region;cohesin complex;nuclear chromosome;nuclear mitotic cohesin complex;nuclear periphery;nucleus |
| PF3D7_0422400 | RPS19 | 40S ribosomal protein S19 | 19.72 | no | 0 | 265.03 | 189.45 | 98.34 | 8.91 | 81.63 | 161.57 | 229.58 | R | 80.80 | 83.79 | 0 | RNA binding;structural constituent of ribosome | ribosomal small subunit assembly;translation | cytosolic small ribosomal subunit;extracellular vesicle;nucleus |
| PF3D7_1329300 | CAF1B | chromatin assembly factor 1 subunit B, putative | 66.51 | no | 0 | 7.86 | 1.35 | 36.46 | 6.21 | 8.40 | 23.98 | 40.73 | O | 143.86 | 25.16 | M | chromatin binding;histone binding;protein binding;unfolded protein binding | chromatin assembly;chromatin assembly or disassembly | nucleus |
| PF3D7_1359600 | N/A | conserved Plasmodium protein, unknown function | 235.14 | no | 0 | 7.69 | 49.44 | 30.86 | 4.06 | 2.98 | 18.14 | 35.41 | T | 20.51 | 32.19 | F | N/A | N/A | nucleus |
| PF3D7_1403100 | CAPD2 | condensin complex subunit 1, putative | 229.53 | no | 1 | 8.74 | 5.50 | 41.88 | 13.10 | 15.82 | 33.23 | 615.29 | O | 256.00 | 109.38 | M | chromatin binding;histone binding | chromosome separation;meiotic chromosome condensation;mitotic chromosome condensation | condensed chromosome, centromeric region;condensin complex |
| PF3D7_0315100 | eIF4E | eukaryotic translation initiation factor 4E | 26.95 | no | 0 | 32.14 | 15.40 | 12.51 | 2.04 | 13.42 | 19.80 | 48.77 | O | 15.76 | 22.43 | F | RNA 7-methylguanosine cap binding;RNA cap binding;mRNA cap binding;protein binding;translation initiation factor activity | translational initiation | P granule;eukaryotic translation initiation factor 4F complex;nucleus |
| PF3D7_0302100 | SRPK1 | serine/threonine protein kinase | 157.37 | no | 0 | 38.29 | 33.71 | 24.41 | 2.20 | 3.78 | 26.91 | 68.03 | O | 259.41 | 38.10 | M | MAP kinase activity;protein serine/threonine kinase activity | intracellular signal transduction;protein phosphorylation;regulation of gene expression;regulation of mRNA processing;spliceosomal complex assembly | cytoplasm;nucleus |
| PF3D7_0520000 | RPS9 | 40S ribosomal protein S9, putative | 22.12 | no | 0 | 587.12 | 471.77 | 200.53 | 62.55 | 168.15 | 92.75 | 299.37 | R | 111.98 | 129.99 | 0 | RNA binding;rRNA binding;structural constituent of ribosome | positive regulation of translational fidelity | RNA N6-methyladenosine methyltransferase complex;cytosolic small ribosomal subunit;nucleus;small ribosomal subunit;small-subunit processome |
| PF3D7_1141800 | N/A | EELM2 domain-containing protein, putative | 46.67 | no | 0 | 6.41 | 0.52 | 39.04 | 28.37 | 3.98 | 19.09 | 46.26 | O | 6.31 | 33.32 | F | transcription corepressor activity;metal ion binding | negative regulation of transcription, DNA-templated | nucleus |
| PF3D7_1368100 | RPN11 | 26S proteasome regulatory subunit RPN11, putative | 35.21 | no | 0 | 50.34 | 39.56 | 135.99 | 77.42 | 20.37 | 19.41 | 96.83 | T | 31.30 | 38.72 | F | metallopeptidase activity;proteasome binding;thiol-dependent deubiquitinase | proteasome-mediated ubiquitin-dependent protein catabolic process;protein deubiquitination;ubiquitin-dependent protein catabolic process | nucleus;proteasome regulatory particle;proteasome regulatory particle, lid subcomplex |
| PF3D7_0910400 | N/A | selenide water dikinase, putative | 140.32 | no | 0 | 9.78 | 7.06 | 10.13 | 4.23 | 1.80 | 10.68 | 13.51 | O | 18.60 | 15.82 | 0 | selenide, water dikinase activity | selenocysteine biosynthetic process | cytoplasm;nucleus |
| PF3D7_0918300 | EIF3F | eukaryotic translation initiation factor 3 subunit F, putative | 36.82 | no | 0 | 123.68 | 44.90 | 31.80 | 1.91 | 16.45 | 29.15 | 19.08 | R | 63.08 | 23.22 | M | translation initiation factor activity;translation initiation factor binding | translational initiation | eukaryotic translation initiation factor 3 complex;eukaryotic translation initiation factor 3 complex, eIF3m;nucleus |
| PF3D7_1347100 | TOP3 | DNA topoisomerase 3 | 84.31 | no | 0 | 7.65 | 2.04 | 18.95 | 6.61 | 3.85 | 75.60 | 116.55 | O | 69.75 | 157.00 | F | DNA binding;DNA topoisomerase activity;protein binding | DNA topological change;DNA unwinding involved in DNA replication | mitochondrion;nucleus |
| PF3D7_0719300 | ARP6 | actin-related protein ARP6 | 121.13 | no | 0 | 67.57 | 25.65 | 22.10 | 2.16 | 4.58 | 24.11 | 558.24 | O | 77.78 | 90.84 | F | nucleosome binding;protein binding | histone exchange | Swr1 complex;nucleus |
| PF3D7_0933000 | N/A | CSTF domain-containing protein, putative | 88.93 | no | 0 | 67.99 | 16.78 | 17.78 | 3.75 | 2.26 | 18.44 | 52.44 | R | 117.99 | 23.74 | M | mRNA binding | pre-mRNA cleavage required for polyadenylation | mRNA cleavage and polyadenylation specificity factor complex;nucleus |
| PF3D7_1324100 | N/A | cyclin-like protein, putative | 61.55 | no | 5 | 4.61 | 0.88 | 15.75 | 5.26 | 2.67 | 10.38 | 8.13 | T | 10.20 | 3.10 | M | N/A | N/A | integral component of membrane;nucleus |
| PF3D7_1408400 | FANCJ | FANCJ-like helicase, putative | 135.54 | no | 0 | 10.87 | 14.94 | 10.88 | 2.72 | 0.98 | 13.05 | 43.48 | O | 278.60 | 10.44 | M | ATP binding;DNA helicase activity;DNA polymerase binding | DNA duplex unwinding;negative regulation of DNA recombination;negative regulation of t-circle formation;regulation of double-strand break repair via homologous recombination;telomeric loop disassembly | mitochondrion;nucleus |
| PF3D7_0722400 | OLA1 | Obg-like ATPase 1, putative | 45.16 | no | 0 | 196.76 | 106.14 | 119.82 | 19.13 | 6.13 | 6.39 | 85.92 | R | 12.21 | 6.80 | M | ATP hydrolysis activity;GTP binding | signal transduction | cytoplasm;nucleus |
| PF3D7_0321500 | N/A | peptidase, putative | 125.89 | no | 0 | 6.20 | 4.40 | 18.98 | 3.75 | 2.02 | 35.74 | 30.93 | GV | 19.39 | 100.30 | F | peptidase activity;serine-type endopeptidase activity | N/A | nucleus |
| PF3D7_1026800 | RPS2 | 40S ribosomal protein S2 | 29.86 | no | 0 | 1651.31 | 969.41 | 501.51 | 98.27 | 64.09 | 263.46 | 1652.19 | R | 123.54 | 290.32 | F | RNA binding;structural constituent of ribosome | ribosomal small subunit assembly;translation | ribosome;small ribosomal subunit |
| PF3D7_0929200 | N/A | RNA-binding protein, putative | 33.60 | no | 0 | 69.72 | 44.30 | 35.57 | 6.32 | 27.39 | 44.17 | 191.66 | O | 143.91 | 48.01 | M | mRNA binding | N/A | cytoplasm;nucleus |
| PF3D7_0813300 | N/A | NPL domain-containing protein, putative | 36.26 | no | 0 | 511.86 | 1801.83 | 170.55 | 6.64 | 91.14 | 79.72 | 270.71 | T | 189.00 | 282.59 | F | RNA binding | N/A | food vacuole;nucleolus;nucleus |
| PF3D7_1010200 | N/A | DNA2/NAM7 helicase, putative | 217.16 | no | 0 | 185.97 | 60.34 | 12.11 | 0.93 | 2.12 | 12.33 | 25.60 | R | 13.03 | 5.14 | M | RNA binding;RNA helicase activity | nuclear-transcribed mRNA catabolic process, nonsense-mediated decay | cytoplasm;nucleus |
| PF3D7_0807600 | N/A | conserved Plasmodium protein, unknown function | 159.03 | no | 0 | 143.64 | 53.18 | 34.07 | 5.28 | 5.87 | 18.43 | 63.24 | R | 64.44 | 55.06 | M | N/A | N/A | nucleus |
| PF3D7_0807500 | N/A | proteasome subunit alpha type-6, putative | 29.50 | no | 0 | 34.26 | 17.51 | 57.18 | 10.35 | 8.95 | 24.86 | 79.68 | O | 50.95 | 62.74 | F | endopeptidase activity | proteasomal protein catabolic process;proteasomal ubiquitin-independent protein catabolic process;proteasome-mediated ubiquitin-dependent protein catabolic process;ubiquitin-dependent protein catabolic process | cytoplasm;nucleus;proteasome core complex;proteasome core complex, alpha-subunit complex |
| PF3D7_1110700 | ALP1 | actin-like protein, putative | 42.48 | no | 0 | 7.09 | 0.34 | 31.59 | 34.77 | 14.52 | 31.17 | 117.12 | O | 13.62 | 44.76 | F | protein binding | cytoskeleton organization; actin filament organization; actin filament-based movement; vesicle transport along actin filament | nucleus;cytoplasm |
| PF3D7_1341200 | N/A | 60S ribosomal protein L18, putative | 21.75 | no | 0 | 311.87 | 221.18 | 108.15 | 16.90 | 60.88 | 83.53 | 136.74 | R | 55.32 | 55.51 | 0 | structural constituent of ribosome | translation | cytosolic large ribosomal subunit;nucleus |
| PF3D7_1345700 | IDH | isocitrate dehydrogenase [NADP], mitochondrial | 53.59 | no | 0 | 21.79 | 14.16 | 51.14 | 17.21 | 4.32 | 8.80 | 25.36 | T | 39.99 | 44.92 | F | isocitrate dehydrogenase (NADP+) activity | NADP metabolic process;isocitrate metabolic process;tricarboxylic acid cycle | mitochondrion;nucleus |
| PF3D7_1302100 | G27/25 | gamete antigen 27/25 | 26.00 | no | 0 | 41.03 | 348.81 | 77.43 | 17.89 | 1238.79 | 648.85 | 268.30 | G II | 293.47 | 343.37 | F | protein dimerization activity | N/A | nucleus |
| PF3D7_0218700 | PRP45 | pre-mRNA-processing protein 45, putative | 57.53 | no | 0 | 35.23 | 18.41 | 10.35 | 1.41 | 4.59 | 27.48 | 164.60 | O | 233.68 | 76.18 | M | RNA binding | RNA processing | nucleus |
| PF3D7_1247400 | FKBP35 | peptidyl-prolyl cis-trans isomerase FKBP35 | 34.83 | no | 0 | 17.76 | 32.59 | 23.49 | 3.13 | 2.66 | 9.09 | 35.88 | O | 85.93 | 16.16 | M | FK506 binding;peptidyl-prolyl cis-trans isomerase activity;protein dimerization activity | chaperone-mediated protein folding;negative regulation of phosphoprotein phosphatase activity;protein folding;protein peptidyl-prolyl isomerization;response to xenobiotic stimulus | cytoplasm;nucleus |
| PF3D7_0727400 | N/A | proteasome subunit alpha type-5, putative | 28.39 | no | 0 | 168.66 | 69.91 | 228.43 | 89.03 | 56.18 | 47.95 | 89.49 | T | 22.26 | 33.70 | 0 | endopeptidase activity | proteasomal protein catabolic process;proteasomal ubiquitin-independent protein catabolic process;proteasome-mediated ubiquitin-dependent protein catabolic process;ubiquitin-dependent protein catabolic process | cytoplasm;extracellular vesicle;nucleus;proteasome core complex;proteasome core complex, alpha-subunit complex |
| PF3D7_1124300 | BDP7 | bromodomain protein 7 | 62.33 | no | 0 | 64.40 | 69.11 | 38.46 | 10.73 | 3.55 | 14.52 | 360.42 | O | 73.69 | 26.80 | M | chromatin binding;histone binding;protein binding | regulation of transcription | heterochromatin;nucleus |
| PF3D7_1145200 | N/A | serine/threonine protein kinase, putative | 238.76 | no | 0 | 14.70 | 1.72 | 82.11 | 115.78 | 12.23 | 165.29 | 1299.69 | O | 74.52 | 1053.09 | F | enzymatic activity | signal transduction | nucleus |
| PF3D7_1420600 | PANK1 | pantothenate kinase 1 | 59.92 | no | 0 | 21.25 | 8.12 | 13.60 | 2.82 | 3.07 | 9.96 | 2.94 | R | 20.59 | 0.79 | M | 14-3-3 protein binding;pantothenate kinase activity;protein binding | coenzyme A biosynthetic process;response to xenobiotic stimulus | cytosol;nucleus |
| PF3D7_1006200 | ALBA3 | DNA/RNA-binding protein Alba 3 | 11.98 | no | 0 | 829.33 | 497.94 | 424.34 | 49.27 | 361.79 | 261.75 | 278.91 | R | 61.84 | 77.63 | F | DNA binding;RNA binding;endonuclease activity;mRNA binding;protein binding | | RNA N6-methyladenosine methyltransferase complex;extracellular vesicle;food vacuole;nucleus |
| PF3D7_1323400 | RPL23 | 60S ribosomal protein L23 | 22.09 | no | 0 | 294.72 | 206.52 | 100.00 | 19.62 | 40.97 | 57.18 | 166.47 | R | 97.25 | 39.60 | M | rRNA binding;structural constituent of ribosome | ribosomal large subunit assembly;translation | cytosolic large ribosomal subunit;nucleus |
| PF3D7_0907400 | ClpY | ATP-dependent protease ATPase subunit ClpY | 106.46 | no | 0 | 56.06 | 31.45 | 56.78 | 15.98 | 6.00 | 22.76 | 225.56 | O | 90.55 | 39.48 | M | mRNA binding;protein binding | N/A | mitochondrion;nucleus |
| PF3D7_1103100 | RPP1 | 60S acidic ribosomal protein P1, putative | 13.01 | no | 0 | 1624.59 | 1493.07 | 452.53 | 198.66 | 1479.73 | 2641.06 | 1067.62 | G V | 273.89 | 1049.36 | F | protein kinase activator activity;ribonucleoprotein complex binding;structural constituent of ribosome | cytoplasmic translation;translation | cytosolic large ribosomal subunit;large ribosomal subunit;nucleus |
| PF3D7_0319400 | N/A | kinesin-8X | 157.39 | no | 0 | 6.22 | 0.40 | 16.49 | 2.91 | 5.70 | 31.08 | 49.68 | O | 293.33 | 45.04 | M | ATP hydrolysis activity;microtubule binding;microtubule motor activity;plus-end-directed microtubule motor activity | microtubule depolymerization;microtubule-based movement | kinesin complex;microtubule;mitotic spindle;nucleus |
| PF3D7_1364200 | NUP205 | nucleoporin NUP205, putative | 207.29 | no | 0 | 28.39 | 6.64 | 9.97 | 2.03 | 1.15 | 9.24 | 17.17 | R | 22.05 | 8.27 | M | protein binding | nucleocytoplasmic transport | nuclear pore;nucleus |
| PF3D7_0307100 | RPS12 | 40S ribosomal protein S12, putative | 15.40 | no | 0 | 323.64 | 393.50 | 185.85 | 22.52 | 150.04 | 91.36 | 125.32 | T | 80.85 | 104.89 | F | structural constituent of ribosome | translation | cytosolic small ribosomal subunit;nucleus |
| PF3D7_1369700 | SNRPA1 | U2 small nuclear ribonucleoprotein A', putative | 20.42 | no | 0 | 60.45 | 7.55 | 5.27 | 1.24 | 16.53 | 34.73 | 18.06 | R | 13.52 | 9.74 | M | RNA binding | mRNA splicing, via spliceosome | spliceosomal complex |
| PF3D7_1008800 | NOP5 | nucleolar protein 5, putative | 53.41 | no | 0 | 127.27 | 62.97 | 23.33 | 1.00 | 10.00 | 34.74 | 182.09 | O | 98.41 | 141.36 | F | RNA binding;snoRNA binding | rRNA processing | box C/D RNP complex;nucleolus;nucleus;small-subunit processome |
| PF3D7_1456000 | AP2-HC | AP2 domain transcription factor AP2-HC | 161.51 | no | 0 | 4.11 | 0.68 | 15.05 | 3.24 | 2.56 | 11.86 | 30.98 | O | 82.54 | 8.94 | M | protein binding;sequence-specific DNA binding | regulation of transcription | heterochromatin;nucleus |
| PF3D7_1018200 | PPP8 | pseudophosphatase PPP8 | 254.11 | no | 0 | 16.40 | 1.78 | 46.07 | 79.30 | 6.66 | 48.53 | 259.74 | O | 7.75 | 184.96 | F | protein binding;protein serine/threonine phosphatase activity | | basal part of cell;cytoplasm;nucleus;protein-containing complex |
| PF3D7_1319400 | N/A | conserved protein, unknown function | 127.67 | no | 0 | 13.32 | 4.96 | 40.99 | 8.88 | 7.60 | 40.04 | 239.32 | O | 1028.50 | 132.26 | M | N/A | N/A | nucleus |
| PF3D7_1213200 | MED18 | mediator of RNA polymerase II transcription subunit 18, putative | 33.25 | no | 0 | 10.89 | 5.18 | 7.41 | 1.12 | 4.98 | 37.90 | 11.42 | GV | 13.84 | 25.14 | F | protein binding | regulation of transcription | mediator complex;nucleus |
| PF3D7_1349200 | GluRS | glutamate--tRNA ligase | 101.42 | no | 0 | 177.94 | 42.40 | 38.72 | 6.00 | 3.73 | 25.33 | 140.01 | R | 25.32 | 29.44 | F | ATP binding;glutamate-tRNA ligase activity;glutamine-tRNA ligase activity | glutaminyl-tRNA aminoacylation;glutamyl-tRNA aminoacylation | cytosol;nucleus |
| PF3D7_1019400 | N/A | 60S ribosomal protein L30e, putative | 11.78 | no | 0 | 151.18 | 124.64 | 63.22 | 5.98 | 57.61 | 81.27 | 208.06 | O | 65.43 | 46.29 | M | RNA binding;structural constituent of ribosome | translation | cytosolic large ribosomal subunit;nucleus |
| PF3D7_0721600 | N/A | 40S ribosomal protein S5, putative | 21.86 | no | 0 | 1392.72 | 990.21 | 392.28 | 48.67 | 248.03 | 145.91 | 432.69 | R | 235.02 | 245.98 | F | RNA binding;mRNA binding;rRNA binding;structural constituent of ribosome | ribosomal small subunit assembly;translation | cytosolic small ribosomal subunit;nucleus;ribosome |
| PF3D7_1037500 | DYN2 | dynamin-like protein | 81.53 | no | 0 | 3.60 | 0.50 | 10.53 | 6.30 | 1.48 | 19.73 | 51.41 | O | 11.47 | 19.22 | F | GTPase activity;microtubule binding | mitochondrial fission;organelle fission | Golgi apparatus;apicoplast;cytoplasm;endoplasmic reticulum;membrane |
| PF3D7_1414400 | PP1 | serine/threonine protein phosphatase PP1 | 34.90 | no | 0 | 22.21 | 26.86 | 170.82 | 90.06 | 50.23 | 195.76 | 310.63 | O | 155.58 | 92.23 | M | protein binding;protein serine/threonine phosphatase activity | entry into host;exit from host cell;protein dephosphorylation | Maurer's cleft;cell surface;cytoplasm;nucleus |
| PF3D7_0909400 | N/A | exoribonuclease, putative | 154.96 | no | 0 | 77.12 | 29.14 | 29.26 | 4.33 | 2.38 | 27.46 | 31.23 | R | 31.25 | 87.57 | F | 5'-3' exoribonuclease activity;RNA binding | nuclear-transcribed mRNA catabolic process | nucleus |
| PF3D7_0923900 | PABP2 | polyadenylate-binding protein 2, putative | 22.98 | no | 0 | 444.51 | 140.03 | 100.20 | 26.94 | 127.43 | 194.13 | 154.95 | R | 95.85 | 77.53 | M | RNA binding;poly(A) binding | posttranscriptional regulation | cytosol;nucleus |
| PF3D7_1460800 | SNAPC3 | snRNA-activating protein complex subunit 3, putative | 75.93 | no | 0 | 5.30 | 3.41 | 5.87 | 1.86 | 6.42 | 18.05 | 109.34 | O | 68.12 | 32.63 | M | RNA polymerase III type 3 promoter sequence-specific DNA binding;bent DNA binding | snRNA transcription by RNA polymerase II;snRNA transcription by RNA polymerase III | nucleus;snRNA-activating protein complex |
| PF3D7_1019700 | N/A | conserved Plasmodium protein, unknown function | 128.21 | no | 0 | 7.24 | 2.36 | 9.76 | 3.61 | 2.82 | 19.95 | 217.76 | O | 199.14 | 339.55 | F | N/A | N/A | histone acetyltransferase complex;nucleus |
| PF3D7_0725300 | N/A | WD repeat-containing protein, putative | 93.45 | no | 0 | 113.99 | 75.72 | 14.43 | 2.86 | 4.48 | 5.01 | 63.19 | R | 11.10 | 7.02 | M | protein binding | ribosomal large subunit biogenesis | nucleolus;nucleus;preribosome, large subunit precursor |
| PF3D7_0730300 | ApiAP2 | AP2 domain transcription factor AP2-L, putative | 155.13 | no | 0 | 227.37 | 137.47 | 40.04 | 3.01 | 1.76 | 9.32 | 83.57 | R | 53.53 | 9.26 | M | DNA binding | regulation of transcription | nucleus |
| PF3D7_1033500 | WDR70 | WD repeat-containing protein 70, putative | 83.23 | no | 0 | 39.56 | 10.58 | 12.49 | 3.79 | 2.37 | 42.59 | 107.81 | O | 30.97 | 101.63 | F | protein binding | regulation of DNA double-strand break processing;regulation of histone H2B conserved C-terminal lysine ubiquitination | nucleus;site of double-strand break |
| PF3D7_0312300 | RPN12 | 26S proteasome regulatory subunit RPN12, putative | 35.10 | no | 0 | 33.67 | 18.77 | 33.87 | 11.50 | 59.24 | 150.20 | 36.40 | GV | 110.44 | 37.74 | M | protein binding | proteasome-mediated ubiquitin-dependent protein catabolic process;ubiquitin-dependent protein catabolic process | cytosol;intracellular;nucleus;proteasome regulatory particle;proteasome regulatory particle, lid subcomplex |
| PF3D7_0629100 | NAPRT | nicotinate phosphoribosyltransferase, putative | 77.13 | no | 0 | 24.40 | 14.71 | 19.00 | 5.35 | 1.36 | 23.13 | 117.81 | O | 6.53 | 72.99 | F | nicotinate phosphoribosyltransferase activity | NAD biosynthetic process;NAD salvage | cytoplasm;cytosol |
| PF3D7_1136300 | TSN | tudor staphylococcal nuclease | 128.54 | no | 0 | 248.19 | 19.54 | 31.74 | 12.59 | 6.70 | 22.89 | 57.62 | R | 65.82 | 108.22 | F | RNA binding;calcium ion binding;nuclease activity;protein binding | RNA catabolic process | cytoplasm;nucleus |
| PF3D7_1326400 | eIF-2Bg | translation initiation factor eIF-2B subunit gamma, putative | 55.04 | no | 0 | 58.95 | 8.89 | 11.95 | 0.98 | 5.61 | 33.96 | 14.58 | R | 50.42 | 13.16 | M | protein binding | translational initiation | eukaryotic translation initiation factor 2B complex |
| PF3D7_1419900 | YTH1 | YTH domain-containing protein 1, putative | 67.17 | no | 0 | 26.52 | 5.62 | 6.22 | 0.53 | 3.82 | 26.27 | 6.89 | G V | 42.05 | 16.95 | M | N6-methyladenosine-containing RNA binding;mRNA binding | mRNA splicing, via spliceosome;regulation of mRNA splicing, via spliceosome | cytoplasm;nucleoplasm;nucleus |
| PF3D7_0527600 | N/A | conserved Plasmodium protein, unknown function | 52.97 | no | 0 | 9.83 | 11.53 | 30.94 | 6.43 | 5.55 | 47.32 | 116.42 | O | 44.48 | 184.91 | F | N/A | N/A | nucleus |
| PF3D7_0527500 | HIP | Hsc70-interacting protein | 51.13 | no | 0 | 818.66 | 971.48 | 631.26 | 656.09 | 90.01 | 59.83 | 311.89 | T | 290.55 | 320.92 | F | heat shock protein binding | chaperone cofactor-dependent protein refolding;chaperone-mediated protein folding;protein-containing complex assembly | nucleus |
| PF3D7_0919000 | NAPS | nucleosome assembly protein | 31.83 | no | 0 | 541.25 | 142.40 | 251.29 | 39.57 | 72.58 | 65.00 | 248.73 | R | 357.25 | 45.25 | M | chromatin binding;histone binding | | cytoplasm;histone acetyltransferase complex;nuclear periphery;nucleus |
| PF3D7_1438500 | CPSF3 | cleavage and polyadenylation specificity factor subunit 3, putative | 101.18 | no | 0 | 24.85 | 6.74 | 6.28 | 0.51 | 2.22 | 4.01 | 16.95 | R | 21.65 | 11.06 | M | 5'-3' exonuclease activity;RNA binding;endoribonuclease activity | mRNA 3'-end processing by stem-loop binding and cleavage;mRNA cleavage;mRNA polyadenylation;response to xenobiotic stimulus | mRNA cleavage and polyadenylation specificity factor complex;nucleus |
| PF3D7_1014300 | N/A | SPRY domain-containing protein, putative | 283.13 | no | 8 | 7.90 | 7.20 | 35.86 | 8.75 | 7.47 | 45.11 | 54.03 | O | 728.81 | 139.25 | M | N/A | proteasome-mediated ubiquitin-dependent protein catabolic process | cytoplasm;membrane;nucleus |
| PF3D7_1347500 | ALBA4 | DNA/RNA-binding protein Alba 4 | 42.16 | no | 0 | 628.83 | 120.68 | 135.98 | 12.40 | 18.97 | 63.69 | 221.29 | R | 94.80 | 65.11 | M | DNA binding;RNA binding;mRNA binding;protein-containing complex binding | regulation of translation | cytoplasm;cytosol;food vacuole;nucleus |
| PF3D7_1443000 | SRPK2 | serine/threonine protein kinase | 150.09 | no | 0 | 3.42 | 0.61 | 4.03 | 1.69 | 1.19 | 9.32 | 7.90 | GV | 6.59 | 15.58 | F | protein kinase activity;protein serine/threonine kinase activity;protein tyrosine kinase activity | peptidyl-serine phosphorylation;peptidyl-threonine phosphorylation;protein phosphorylation | Maurer's cleft;cytoplasm;nucleus |
| PF3D7_1316600 | CCT | choline-phosphate cytidylyltransferase | 104.78 | no | 0 | 56.47 | 37.32 | 41.62 | 22.49 | 6.61 | 15.91 | 90.53 | O | 35.14 | 17.65 | M | choline-phosphate cytidylyltransferase activity;identical protein binding;phosphatidylcholine binding | phosphatidylcholine biosynthetic process | cytoplasm;nucleus |
| PF3D7_0110400 | RPB9 | DNA-directed RNA polymerase II subunit RPB9, putative | 29.69 | no | 0 | 45.15 | 31.60 | 30.74 | 4.60 | 13.05 | 119.22 | 167.78 | O | 121.76 | 135.17 | F | DNA-directed 5'-3' RNA polymerase activity | maintenance of transcriptional fidelity during DNA-templated transcription elongation from RNA polymerase II promoter;transcription by RNA polymerase II;transcription initiation from RNA polymerase II promoter;transcription-coupled nucleotide-excision repair | RNA polymerase II, core complex;nucleus |
| PF3D7_0714500 | N/A | transcription elongation factor s-II, putative | 47.35 | no | 0 | 32.83 | 12.19 | 11.63 | 1.69 | 2.43 | 11.63 | 92.88 | O | 112.43 | 12.57 | M | DNA binding | regulation of DNA-templated transcription, elongation;regulation of transcription by RNA polymerase II | nucleus;transcription elongation factor complex |

| | | | | | | | | | | | | | | | | | | |
|---|---|---|---|---|---|---|---|---|---|---|---|---|---|---|---|---|---|---|
| PF3D7_1315800 | MYB1 | transcription factor MYB1 | 49.99 | no | 0 | 25.21 | 22.70 | 15.37 | 2.90 | 3.82 | 53.30 | 38.97 | GV | 40.54 | 16.77 | M | DNA-binding transcription factor activity | positive regulation of transcription, DNA-templated;regulation of mitotic cell cycle;regulation of transcription, DNA-templated | nucleus |
| PF3D7_1129000 | SPDS | spermidine synthase | 36.60 | no | 0 | 70.49 | 91.94 | 72.07 | 39.97 | 18.47 | 36.92 | 79.27 | T | 15.03 | 57.74 | F | RNA binding;spermidine synthase activity | polyamine biosynthetic process;spermidine biosynthetic process | cytoplasm;cytosol;mitochondrion;nucleus |
| PF3D7_1316900 | N/A | conserved protein, unknown function | 114.27 | no | 0 | 59.60 | 24.81 | 16.82 | 3.21 | 5.55 | 28.94 | 19.40 | R | 63.12 | 9.14 | M | N/A | regulation of transcription by RNA polymerase II | Piccolo NuA4 histone acetyltransferase complex;nucleus |
| PF3D7_0813900 | RPS16 | 40S ribosomal protein S16, putative | 16.29 | no | 0 | 231.51 | 168.64 | 92.97 | 16.33 | 125.34 | 100.88 | 62.77 | R | 142.94 | 112.88 | M | RNA binding;structural constituent of ribosome | maturation of SSU-rRNA from tricistronic rRNA transcript (SSU-rRNA, 5.8S rRNA, LSU-rRNA);translation | cytosolic small ribosomal subunit;nucleus;small ribosomal subunit |
| PF3D7_0718500 | N/A | prefoldin subunit 3, putative | 22.64 | no | 0 | 23.17 | 24.48 | 33.37 | 7.36 | 114.48 | 94.99 | 145.46 | O | 116.37 | 70.29 | M | tubulin binding;unfolded protein binding | 'de novo' protein folding;microtubule-based process;tubulin complex assembly | cytoplasm;polysome;prefoldin complex |
| PF3D7_1426100 | BTF3 | transcription factor BTF3, putative | 19.39 | no | 0 | 87.26 | 83.37 | 30.28 | 1.86 | 25.60 | 91.62 | 65.00 | GV | 45.91 | 56.56 | F | DNA binding | regulation of transcription | cytosol;nascent polypeptide-associated complex;nucleus;polysomal ribosome;transcription regulator complex |
| PF3D7_0801700 | SENP2 | sentrin-specific protease 2, putative | 210.88 | no | 0 | 15.44 | 20.41 | 17.08 | 2.12 | 2.68 | 16.21 | 41.33 | O | 21.59 | 53.21 | F | cysteine-type peptidase activity | proteolysis | nucleus |
| PF3D7_1438900 | Trx-Px1 | thioredoxin peroxidase 1 | 21.81 | no | 0 | 528.37 | 147.40 | 188.59 | 52.36 | 319.18 | 169.42 | 66.49 | R | 56.09 | 139.88 | F | RNA binding;antioxidant activity;double-stranded DNA binding;double-stranded RNA binding;long noncoding RNA binding;protein binding;protein dimerization activity;thioredoxin peroxidase activity | DNA protection;cell redox homeostasis;protein folding;response to heat;response to oxidative stress | Maurer's cleft;cell surface;cytoplasm;cytosol;nucleolus;nucleus |
| PF3D7_1251200 | N/A | coronin | 69.01 | no | 0 | 19.93 | 1.22 | 63.45 | 66.96 | 1.06 | 6.29 | 256.48 | O | 103.61 | 10.37 | M | actin binding;actin filament binding | actin filament organization;cell motility;microtubule-based process;response to xenobiotic stimulus | nucleus;pellicle |
| PF3D7_1027300 | nPrx | peroxiredoxin | 43.93 | no | 0 | 236.88 | 45.44 | 438.32 | 815.09 | 92.97 | 54.92 | 774.05 | S | 111.48 | 1667.71 | F | thioredoxin peroxidase activity | cell redox homeostasis;cellular response to oxidative stress;entry into host | cytoplasm;nucleus |
| PF3D7_1346300 | ALBA2 | DNA/RNA-binding protein Alba 2 | 24.98 | no | 0 | 914.99 | 182.62 | 122.63 | 22.05 | 50.48 | 77.26 | 56.49 | R | 50.55 | 87.92 | F | DNA binding;RNA binding;mRNA binding | regulation of translation | cytoplasm;nucleus |
| PF3D7_0517300 | SR1 | serine/arginine-rich splicing factor 1 | 34.56 | no | 0 | 404.84 | 179.12 | 92.82 | 18.50 | 15.00 | 43.87 | 133.57 | R | 93.83 | 76.82 | M | RNA binding;phosphorylated histone binding | RNA splicing;cellular response to DNA damage stimulus;mRNA cis splicing, via spliceosome;mRNA processing;regulation of alternative mRNA splicing, via spliceosome;regulation of mRNA splicing, via spliceosome | cytoplasm;nuclear speck;nucleus;spliceosomal complex |
| PF3D7_1437400 | PANK2 | pantothenate kinase 2 | 91.04 | no | 0 | 10.06 | 9.72 | 15.29 | 3.89 | 1.12 | 6.15 | 22.92 | O | 10.76 | 3.20 | M | 14-3-3 protein binding;pantothenate kinase activity;protein binding | coenzyme A biosynthetic process | cytosol;nucleus |
| PF3D7_1110900 | N/A | ES2 protein, putative | 69.02 | no | 0 | 35.87 | 14.38 | 15.75 | 2.01 | 2.15 | 17.12 | 47.50 | O | 59.20 | 29.81 | M | protein binding;RNA binding | RNA splicing | catalytic step 2 spliceosome |
| PF3D7_0311300 | N/A | phosphatidylinositol 3- and 4-kinase, putative | 110.56 | no | 0 | 6.17 | 1.72 | 45.19 | 20.67 | 11.52 | 35.88 | 147.36 | O | 159.71 | 49.59 | M | enzymatic activity | signal transduction | nucleus;cytoplasm |
| PF3D7_1340600 | DBR1 | RNA lariat debranching enzyme, putative | 67.55 | no | 0 | 44.62 | 41.84 | 34.26 | 2.84 | 2.13 | 8.42 | 109.60 | O | 49.08 | 21.59 | M | hydrolase activity, acting on ester bonds | mRNA processing | nucleus |
| PF3D7_0621800 | N/A | nascent polypeptide-associated complex subunit alpha | 20.62 | no | 0 | 727.07 | 431.74 | 116.63 | 8.14 | 243.72 | 158.76 | 103.51 | R | 52.90 | 52.33 | 0 | unfolded protein binding | 'de novo' cotranslational protein folding;protein targeting to membrane | cytoplasm;nascent polypeptide-associated complex;nucleus |
| PF3D7_0923000 | RPB3 | DNA-directed RNA polymerase II subunit RPB3, putative | 38.35 | no | 0 | 75.74 | 13.31 | 9.78 | 0.61 | 4.20 | 36.68 | 11.96 | R | 25.76 | 12.27 | M | DNA binding | regulation of transcription | RNA polymerase II, core complex;nucleus |
| PF3D7_0822800 | SNRNP40 | U5 small nuclear ribonucleoprotein 40 kDa protein, putative | 37.09 | no | 0 | 24.10 | 6.32 | 8.54 | 0.55 | 1.71 | 11.55 | 25.15 | O | 19.01 | 20.98 | 0 | RNA binding | RNA splicing, via transesterification reactions | catalytic step 2 spliceosome;sno(s)RNA-containing ribonucleoprotein complex |
| PF3D7_0707700 | N/A | E3 ubiquitin-protein ligase, putative | 110.78 | no | 0 | 136.99 | 65.52 | 57.95 | 34.11 | 37.56 | 96.70 | 133.19 | R | 167.10 | 67.43 | M | chaperone binding;ubiquitin protein ligase activity | cellular response to misfolded protein;positive regulation of proteolysis;proteasome-mediated ubiquitin-dependent protein catabolic process;protein polyubiquitination;protein quality control for misfolded or incompletely synthesized proteins | cytoplasm;nucleus |
| PF3D7_1107300 | PAIP1 | polyadenylate-binding protein-interacting protein 1, putative | 381.87 | no | 0 | 265.97 | 114.16 | 49.70 | 12.61 | 11.21 | 67.65 | 173.07 | R | 608.32 | 166.19 | M | mRNA binding;mRNA cap binding;translation initiation factor activity | | eukaryotic translation initiation factor 4F complex;nucleus |
| PF3D7_1417500 | CBF5 | H/ACA ribonucleoprotein complex subunit 4, putative | 52.94 | no | 0 | 111.44 | 39.78 | 22.12 | 2.50 | 4.97 | 10.72 | 21.22 | R | 182.67 | 23.21 | M | pseudouridine synthase activity | box H/ACA RNA 3'-end processing;mRNA pseudouridine synthesis;rRNA pseudouridine synthesis;snRNA pseudouridine synth | box H/ACA snoRNP complex;nucleus |
| PF3D7_1105700 | RTCB | RNA-splicing ligase RtcB homolog | 56.32 | no | 0 | 100.05 | 36.31 | 45.84 | 28307.00 | 14.66 | 46722.00 | 34.82 | GC V | 36.12 | 37.41 | 0 | RNA ligase (ATP) activity | tRNA exon ligation utilizing 2',3' cyclic phosphate of 5'-exon as source of linkage phosphate;tRNA splicing, via endonucleolytic cleavage and ligation | nucleus;tRNA-splicing ligase complex |
| PF3D7_1126900 | SNRPF | small nuclear ribonucleoprotein F, putative | 9.89 | no | 0 | 55.70 | 29.11 | 30.48 | 5.18 | 28.30 | 39.12 | 46.85 | R | 61.42 | 126.09 | F | RNA binding | mRNA splicing, via spliceosome | U1 snRNP;U4/U6 x U5 tri-snRNP complex;U5 snRNP;catalytic step 2 spliceosome;nucleus |
| PF3D7_0309500 | AS | asparagine synthetase [glutamine-hydrolyzing], putative | 69.86 | no | 0 | 311.94 | 131.53 | 149.76 | 97.72 | 9.10 | 8.84 | 72.42 | R | 69.37 | 39.67 | M | asparagine synthase (glutamine-hydrolyzing) activity | asparagine biosynthetic process | cytosol;nucleus |
| PF3D7_1315900 | N/A | exportin-T, putative | 143.00 | no | 0 | 80.07 | 40.03 | 36.51 | 5.38 | 5.98 | 27.25 | 44.65 | R | 26.75 | 9.39 | M | tRNA binding | tRNA export from nucleus | nucleus;cytoplasm |
| PF3D7_0705300 | ORC2 | origin recognition complex subunit 2 | 98.01 | no | 0 | 6.27 | 3.00 | 19.70 | 7.46 | 2.50 | 28.83 | 81.22 | O | 575.95 | 55.65 | M | DNA replication origin binding | DNA replication initiation | endoplasmic reticulum;mitochondrion;nuclear origin of replication recognition complex;nuclear periphery;nucleus |
| PF3D7_0308200 | CCT7 | T-complex protein 1 subunit eta | 59.58 | no | 0 | 177.06 | 106.03 | 66.15 | 11.71 | 11.42 | 13.35 | 147.58 | R | 76.58 | 54.96 | M | unfolded protein binding | protein folding | chaperonin-containing T-complex;cytosol;nucleus |
| PF3D7_1338200 | RPL6 | 60S ribosomal protein L6, putative | 25.53 | no | 0 | 178.02 | 111.44 | 44.82 | 3.22 | 28.96 | 59.05 | 72.65 | R | 74.97 | 17.81 | M | RNA binding;structural constituent of ribosome | cytoplasmic translation | cytosolic large ribosomal subunit;extracellular vesicle;nucleus |
| PF3D7_0610200 | RBM25 | RNA-binding protein 25, putative | 115.74 | no | 0 | 50.95 | 13.91 | 7.35 | 1.10 | 1.69 | 14.32 | 54.20 | O | 385.44 | 127.39 | M | RNA binding | N/A | nucleus |
| PF3D7_0318100 | STOML | stomatin-like protein, putative | 43.08 | no | 0 | 12.48 | 21.15 | 15.44 | 1.41 | 1.22 | 4.71 | 16.17 | T | 8.54 | 7.14 | 0 | N/A | N/A | mitochondrion;nucleus |
| PF3D7_0929000 | TAF7 | transcription initiation factor TFIID subunit 7, putative | 46.80 | no | 0 | 54.96 | 12.90 | 12.12 | 1.98 | 14.15 | 48.04 | 34.10 | R | 72.91 | 36.34 | M | RNA binding | RNA polymerase II preinitiation complex assembly;transcription by RNA polymerase II | transcription factor TFIID complex |
| PF3D7_1015400 | N/A | RMI1 domain-containing protein, putative | 92.88 | no | 0 | 8.03 | 1.80 | 14.13 | 4.94 | 1.50 | 10.98 | 10.11 | T | 4.06 | 10.95 | F | N/A | N/A | nucleus |
| PF3D7_0907700 | PA28 | proteasome activator 28 | 33.13 | no | 0 | 154.12 | 31.80 | 37.26 | 4.88 | 2.65 | 35.03 | 108.52 | R | 39.48 | 17.56 | M | endopeptidase activator activity | positive regulation of endopeptidase activity;regulation of G1/S transition of mitotic cell cycle;regulation of proteasomal protein catabolic process;response to xenobiotic stimulus | cytoplasm;nucleoplasm;nucleus;proteasome activator complex |
| PF3D7_0306900 | N/A | 40S ribosomal protein S23, putative | 16.13 | no | 0 | 591.61 | 623.27 | 260.25 | 109.84 | 284.46 | 149.34 | 165.61 | T | 156.23 | 128.94 | M | structural constituent of ribosome | translation | cytosolic small ribosomal subunit;nucleus;ribosome |
| PF3D7_0614400 | CWF7 | pre-mRNA-splicing factor CWF7, putative | 30.85 | no | 0 | 26.99 | 15.23 | 5.44 | 0.65 | 2.77 | 24.46 | 432.49 | O | 12.48 | 51.49 | F | RNA binding | mRNA processing | Prp19 complex;catalytic step 2 spliceosome |
| PF3D7_1407300 | PRP38B | pre-mRNA-splicing factor 38B, putative | 81.76 | no | 0 | 108.25 | 64.89 | 55.56 | 5.46 | 7.76 | 26.88 | 223.88 | O | 197.66 | 55.67 | M | RNA binding | mRNA processing | precatalytic spliceosome;nucleus |
| PF3D7_1353900 | N/A | proteasome subunit alpha type-7, putative | 27.23 | no | 0 | 65.78 | 46.70 | 104.96 | 25.63 | 49.91 | 83.15 | 33.60 | T | 20.10 | 39.53 | F | endopeptidase activity | proteasomal protein catabolic process;proteasomal ubiquitin-independent protein catabolic process;proteasome-mediated ubiquitin-dependent protein catabolic process;ubiquitin-dependent protein catabolic process | cytoplasm;nucleus;proteasome core complex;proteasome core complex, alpha-subunit complex |
| PF3D7_1410600 | eIF2gamma | eukaryotic translation initiation factor 2 subunit gamma, putative | 51.24 | no | 0 | 937.06 | 198.24 | 211.93 | 42.44 | 34.53 | 34.65 | 105.36 | R | 20.39 | 51.97 | F | GTP binding;protein binding;translation initiation factor activity | formation of translation preinitiation complex;positive regulation of translational fidelity;translational initiation | cytoplasm;cytosol;eukaryotic translation initiation factor 2 complex;nucleus |
| PF3D7_0310500 | DHX57 | ATP-dependent RNA helicase DHX57, putative | 267.23 | no | 0 | 22.97 | 20.27 | 15.48 | 1.98 | 2.34 | 18.60 | 38.18 | O | 26.64 | 15.82 | M | RNA binding;helicase activity | maturation of SSU-rRNA from tricistronic rRNA transcript (SSU-rRNA, 5.8S rRNA, LSU-rRNA) | nucleolus;nucleus |
| PF3D7_1368200 | ABCE1 | ABC transporter E family member 1, putative | 70.27 | no | 0 | 283.55 | 70.15 | 83.17 | 21.73 | 10.31 | 16.54 | 31.28 | R | 18.17 | 15.02 | 0 | ATP binding;RNA binding;iron ion binding;ribosomal small subunit binding | ribosomal subunit export from nucleus;translational initiation;translational termination | nucleus |
| PF3D7_0205700 | N/A | RNA-binding protein, putative | 24.33 | no | 0 | 47.93 | 14.13 | 43344.00 | 34700.00 | 21.14 | 266.53 | 61.8 | T | 49.23 | 66.93 | F | RNA binding | N/A | nucleus;cytoplasm |
| PF3D7_1465200 | MED4 | mediator of RNA polymerase II transcription subunit 4, putative | 44.68 | no | 0 | 52.44 | 11.71 | 9.66 | 0.75 | 3.05 | 12.84 | 0.00 | R | 11.70 | 6.52 | M | protein binding | regulation of transcription | mediator complex;nucleus |
| PF3D7_1248200 | RBM22 | Pre-mRNA-splicing factor RBM22, putative | 41.97 | no | 0 | 38.85 | 24.29 | 13.53 | 2.71 | 17.94 | 110.22 | 121.74 | O | 65.51 | 145.18 | R | U6 snRNA binding;pre-mRNA binding | mRNA cis splicing, via spliceosome | Prp19 complex;U2-type catalytic step 1 spliceosome;U2-type catalytic step 2 spliceosome |
| PF3D7_1239000 | N/A | HD superfamily phosphohydrolase protein, putative | 96.00 | no | 0 | 21.90 | 10.32 | 20.64 | 7.42 | 9.07 | 87.35 | 85.49 | GV | 2.92 | 57.12 | F | dGTPase activity | dGTP catabolic process | nucleus |
| PF3D7_0109200 | CFIM25 | cleavage and polyadenylation specificity factor subunit 5, putative | 27.32 | no | 0 | 21.09 | 11.97 | 19.67 | 3.38 | 46.97 | 124.90 | 70.98 | GV | 22.24 | 16.42 | M | RNA binding;mRNA binding | mRNA processing | mRNA cleavage factor complex |
| PF3D7_0216200 | N/A | Alba domain-containing protein, putative | 27.21 | no | 0 | 28.93 | 13.50 | 7.69 | 0.54 | 3.68 | 23.34 | 37.48 | O | 14.96 | 29.60 | F | protein binding | N/A | nucleus;cytoplasm |
| PF3D7_1447700 | N/A | conserved protein, unknown function | 40.70 | no | 3 | 9.49 | 3.33 | 13.54 | 5.70 | 8.20 | 82.38 | 8.57 | GV | 13.10 | 12.32 | 0 | RNA binding | N/A | nucleus;cytoplasm |
| PF3D7_1203700 | NAPL | nucleosome assembly protein | 40.49 | no | 0 | 456.51 | 180.45 | 265.44 | 31.80 | 44.21 | 75.80 | 516.99 | O | 363.93 | 98.04 | M | chromatin binding;histone binding;protein binding | nucleosome assembly | cytoplasm;nucleus |
| PF3D7_1237700 | N/A | conserved protein, unknown function | 23.66 | no | 5 | 77.47 | 51.48 | 134.58 | 57.98 | 103.33 | 527.14 | 129.80 | GV | 13.39 | 117.33 | F | N/A | N/A | cell surface;food vacuole;mitochondrion;nucleus |
| PF3D7_1309100 | N/A | 60S ribosomal protein L24, putative | 19.25 | no | 0 | 302.41 | 241.43 | 125.89 | 19.63 | 149.63 | 251.51 | 648.75 | O | 712.32 | 554.36 | M | RNA binding;structural constituent of ribosome | assembly of large subunit precursor of preribosome;ribosomal large subunit assembly;translation | cytosolic large ribosomal subunit;nucleus |
| PF3D7_1142100 | N/A | DnaJ domain-containing protein, putative | 324.82 | no | 0 | 92.19 | 10.52 | 24.05 | 5.73 | 14.57 | 42.45 | 45.85 | R | 229.05 | 48.90 | M | protein binding | chaperone complex | nucleus |
| PF3D7_1016400 | FIKK10.1 | Serine/threonine protein kinase, FIKK family | 72.42 | no | 1 | 38.85 | 24.29 | 13.53 | 25965.00 | 17.94 | 110.22 | 121.74 | S | 65.51 | 145.18 | F | protein serine/threonine kinase activity | intracellular signal transduction;modulation by symbiont of host process;protein phosphorylation | Maurer's cleft;cytoplasm;nucleus |
| PF3D7_1104400 | Trx-mero | thioredoxin-like mero protein | 49.27 | yes | 1 | 120.76 | 75.63 | 173.84 | 52.72 | 17.36 | 47.08 | 79.91 | T | 43.99 | 51.29 | F | host cell surface binding | entry into host | cell surface;nucleus;symbiont-containing vacuole |
| PF3D7_0604500 | N/A | GYF domain-containing protein, putative | 334.41 | no | 0 | 230.63 | 68.12 | 42.18 | 8.77 | 6.13 | 53.06 | 139.04 | R | 358.20 | 182.15 | M | mRNA binding | N/A | nucleus |
| PF3D7_1005500 | UPF1 | regulator of nonsense transcripts 1, putative | 183.96 | no | 0 | 31.20 | 22.88 | 46.85 | 7.51 | 10.15 | 113.61 | 139.54 | O | 165.50 | 175.06 | F | RNA binding;RNA helicase activity | nuclear-transcribed mRNA catabolic process, nonsense-mediated decay | cytoplasm;nucleus |
| PF3D7_1364500 | RRP45 | exosome complex component RRP45, putative | 55.03 | no | 0 | 22.56 | 6.17 | 6.95 | 0.98 | 1.29 | 10.20 | 6.90 | R | 31.77 | 10.00 | M | 3'-5'-exoribonuclease activity | RNA processing;U1 snRNA 3'-end processing;U4 snRNA 3'-end processing;U5 snRNA 3'-end processing;exonucleolytic catabolism of deadenylated mRNA;exonucleolytic trimming to generate mature 3'-end of 5.8S rRNA from tricistronic rRNA transcript (SSU-rRNA, 5.8S rRNA, LSU-rRNA);nuclear mRNA surveillance;nu | cytoplasmic exosome (RNase complex);exosome (RNase complex);nuclear exosome (RNase complex) |
| PF3D7_0803700 | g-tub | tubulin gamma chain | 51.64 | no | 0 | 13.30 | 7.24 | 26.45 | 4.84 | 16.41 | 42.20 | 93.24 | O | 246.66 | 83.70 | M | GTP binding;RNA binding;structural constituent of cytoskeleton | meiotic spindle organization;microtubule cytoskeleton organization;microtubule nucleation;microtubule-based process;mitotic cell cycle;mitotic sister chromatid segregation;mitotic spindle organization | centrosome;cytoplasm;gamma-tubulin complex;microtubule;microtubule organizing center;nucleus;spindle |
| PF3D7_0420300 | ApiAP2 | AP2 domain transcription factor, putative | 400.16 | no | 0 | 115.68 | 92.16 | 57.10 | 28.79 | 4.65 | 20.37 | 743.23 | O | 51.70 | 52.09 | 0 | protein binding;sequence-specific DNA binding | regulation of transcription | nucleus |
| PF3D7_1130100 | RPL38 | 60S ribosomal protein L38 | 10.31 | no | 0 | 57.29 | 57.51 | 28.38 | 7.12 | 63.28 | 368.45 | 76.23 | GV | 96.72 | 195.43 | F | structural constituent of ribosome | ribonucleoprotein complex assembly;translation | cytosolic large ribosomal subunit;nucleus |
| PF3D7_0925700 | HDAC1 | histone deacetylase 1 | 51.38 | no | 0 | 156.78 | 62.14 | 85.17 | 62.49 | 20.60 | 21.37 | 76.19 | R | 89.71 | 34.53 | M | histone deacetylase activity;protein binding;sphingolipid binding | cellular response to drug;histone deacetylation;response to xenobiotic stimulus | nucleus |
| PF3D7_0822100 | MED7 | mediator of RNA polymerase II transcription subunit 4, putative | 28.20 | no | 0 | 20.92 | 7.64 | 5.30 | 0.86 | 59.08 | 126.74 | 31.95 | GV | 54.32 | 34.45 | M | protein binding | regulation of transcription | mediator complex;nucleus |
| PF3D7_1128100 | N/A | prefoldin subunit 5, putative | 29.14 | no | 0 | 42.67 | 38.52 | 48.66 | 6.62 | 14.10 | 65.25 | 182.62 | O | 91.39 | 64.33 | M | unfolded protein binding | protein folding | cytoplasm;prefoldin complex;nucleus |
| PF3D7_1361900 | PCNA1 | Proliferating cell nuclear antigen | 30.59 | no | 0 | 18810.00 | 12267.00 | 97.41 | 17.16 | 16.89 | 46.54 | 75.08 | R | 421.82 | 13.97 | M | DNA polymerase processivity factor activity | leading strand elongation;mismatch repair;translesion synthesis | PCNA complex;cytosol;nucleus |
| PF3D7_1338700 | N/A | conserved protein, unknown function | 73.21 | no | 0 | 4.79 | 3.13 | 32.35 | 5.96 | 1.08 | 6.91 | 422.25 | O | 10.81 | 56.42 | F | N/A | N/A | nucleus |
| PF3D7_1303400 | LisH | LisH domain-containing protein, putative | 118.62 | no | 0 | 78.90 | 48.90 | 23.20 | 6.94 | 3.36 | 9.71 | 67.47 | R | 115.69 | 39.57 | M | protein binding | N/A | nucleus |
| PF3D7_1111100 | RFC5 | replication factor C subunit 5, putative | 40.35 | no | 0 | 9.89 | 5.14 | 37.32 | 5.46 | 10.62 | 61.42 | 119.20 | O | 155.28 | 61.78 | M | RNA binding | DNA repair;DNA replication;DNA strand elongation involved in DNA replication;DNA-dependent DNA replication | DNA replication factor C complex;nucleus |
| PF3D7_0803000 | CYP81 | peptidyl-prolyl cis-trans isomerase | 80.92 | no | 0 | 59.73 | 16.02 | 22.71 | 2.80 | 4.92 | 15.30 | 126.97 | O | 96.24 | 75.26 | M | peptidyl-prolyl cis-trans isomerase activity;protein dimerization activity | chaperone-mediated protein folding;negative regulation of phosphoprotein phosphatase activity;protein folding;protein peptidyl-prolyl isomerization;response to xenobiotic stimulus | cytoplasm;nucleus |
| PF3D7_0722500 | CWC15 | pre-mRNA-splicing factor CWC15, putative | 34.15 | no | 0 | 378.24 | 104.64 | 57.66 | 114.25 | 22.65 | 56.05 | 157.58 | R | 177.63 | 250.92 | F | RNA binding | RNA splicing;mRNA cis splicing, via spliceosome;mRNA splicing, via spliceosome;regulation of cell cycle | U2-type spliceosomal complex;catalytic step 2 spliceosome;spliceosomal complex;nucleus |
| PF3D7_1409300 | DDI1 | DNA damage-inducible protein 1 | 43.84 | no | 0 | 4.92 | 1.51 | 22.34 | 11.29 | 8.18 | 50.85 | 84.60 | O | 75.60 | 45.29 | 0 | aspartic-type endopeptidase activity | proteolysis;response to xenobiotic stimulus | cytoplasm;nucleus |
| PF3D7_1146600 | ORP1 | oocyst rupture protein 1, putative | 150.98 | no | 0 | 4.11 | 0.06 | 12.50 | 0.11 | 3.69 | 16.93 | 56.42 | O | 18.46 | 34.39 | F | DNA binding | regulation of transcription, DNA-templated | cell periphery;nucleus |
| PF3D7_0514900 | N/A | conserved Plasmodium protein, unknown function | 51.90 | no | 0 | 64.91 | 10.96 | 15.01 | 2.40 | 12.96 | 81.06 | 42.68 | GV | 20.60 | 22.05 | 0 | N/A | N/A | cytoplasm;nucleus;transcription regulator complex |
| PF3D7_0628600 | N/A | DNA methyltransferase 1-associated protein 1, putative | 46.56 | no | 0 | 32.66 | 16.54 | 11.66 | 1.03 | 7.50 | 38.86 | 101.09 | O | 29.91 | 20.19 | M | transcription corepressor activity | histone H2A acetylation;histone H4 acetylation;histone exchange;negative regulation of transcription by RNA polymerase II | NuA4 histone acetyltransferase complex;Swr1 complex;nucleus |
| PF3D7_0518300 | N/A | proteasome subunit beta type-1, putative | 27.27 | no | 0 | 59.43 | 33.09 | 89.90 | 21.12 | 71.34 | 91.04 | 40.74 | GV | 41.43 | 13.20 | M | endopeptidase activity | proteasomal protein catabolic process;proteasomal ubiquitin-independent protein catabolic process;proteasome-mediated ubiquitin-dependent protein catabolic process;ubiquitin-dependent protein catabolic process | cytoplasm;extracellular vesicle;nucleus;proteasome core complex |
| PF3D7_1331800 | N/A | 60S ribosomal protein L23, putative | 14.99 | no | 0 | 268.05 | 367.47 | 121.43 | 76.67 | 219.53 | 135.09 | 163.31 | T | 133.90 | 226.26 | F | RNA binding;large ribosomal subunit rRNA binding;structural constituent of ribosome | | cytosolic large ribosomal subunit;nucleus |
| PF3D7_1473200 | N/A | DnaJ protein, putative | 52.45 | no | 0 | 12.11 | 48.15 | 95.86 | 26.66 | 6.34 | 10.24 | 117.91 | O | 128.29 | 44.19 | M | protein binding | response to heat;response to unfolded protein | Maurer's cleft;nucleus |
| PF3D7_1473700 | NUP116 | nucleoporin NUP116/NSP116, putative | 142.21 | no | 0 | 7.91 | 3.13 | 4.78 | 0.26 | 1.72 | 5.92 | 52.39 | O | 396.00 | 18.66 | M | double-stranded DNA binding;structural constituent of nuclear pore;transcription coregulator activity | protein import into nucleus;regulation of transcription by RNA polymerase II | nuclear pore central transport channel;nucleus |
| PF3D7_1225800 | UBA1 | ubiquitin-activating enzyme E1 | 131.78 | no | 0 | 50.93 | 19.32 | 50.76 | 12.50 | 4.07 | 14.84 | 29.95 | R | 39.60 | 28.32 | M | ubiquitin activating enzyme activity | cellular response to DNA damage stimulus;protein modification by small protein conjugation;protein ubiquitination;response to xenobiotic stimulus;ubiquitin-dependent protein catabolic process | cytoplasm;nucleus |
| PF3D7_0317000 | N/A | proteasome subunit alpha type-3, putative | 29.29 | no | 0 | 49.71 | 19.84 | 48.63 | 22.49 | 129.68 | 215.14 | 67.31 | GV | 23.66 | 27.83 | 0 | endopeptidase activity | proteasomal protein catabolic process;proteasomal ubiquitin-independent protein catabolic process;proteasome-mediated ubiquitin-dep | cytoplasm;nucleus;proteasome core complex;proteasome core complex, alpha-subunit complex |
| PF3D7_1361100 | SEC24A | protein transport protein Sec24A | 106.68 | no | 0 | 303.85 | 28.64 | 146.81 | 112.42 | 28.47 | 24.40 | 105.46 | R | 109.38 | 37.73 | M | RNA binding;SNARE binding;zinc ion binding | COPII-coated vesicle cargo loading;endoplasmic reticulum to Golgi vesicle-mediated transport;protein transport by the Sec complex | COPII vesicle coat;Maurer's cleft;endoplasmic reticulum;endoplasmic reticulum exit site;nucleus |
| PF3D7_0726400 | N/A | conserved Plasmodium membrane protein | 599.32 | no | 8 | 14.59 | 4.99 | 10.86 | 2.74 | 2.25 | 28.13 | 13.14 | GV | 43.22 | 13.70 | M | 5'-flap endonuclease activity;flap endonuclease activity | DNA repair | integral component of membrane;nucleus |

| | | | | | | | | | | | | | | | | | | |
|---|---|---|---|---|---|---|---|---|---|---|---|---|---|---|---|---|---|---|
| PF3D7_1419700 | EIF3H | eukaryotic translation initiation factor 3 subunit H, putative | 59.69 | no | 0 | 301.05 | 106.68 | 60.81 | 7.43 | 12.45 | 42.27 | 112.22 | R | 180.46 | 98.93 | M | mRNA binding | initiation of translation | eukaryotic translation initiation factor 3 complex;nucleolus;nucleus;cytoplasm |
| PF3D7_1021800 | SEA1 | schizont egress antigen-1 | 244.90 | no | 0 | 10.76 | 3.46 | 36.90 | 20.75 | 6.79 | 25.71 | 818.39 | O | 183.77 | 74.22 | M | N/A | exit from host cell;nuclear chromosome segregation;schizogony | Maurer's cleft;chromosome, centromeric region;host cell membrane;nucleus;symbiont-containing vacuole membrane |
| PF3D7_0321600 | DDX42 | ATP-dependent RNA helicase DDX42, putative | 86.65 | no | 0 | 13.26 | 14.71 | 18.84 | 8.28 | 4.52 | 25.40 | 61.68 | O | 71.84 | 89.96 | F | RNA binding;RNA helicase activity | regulation of transcription | cytoplasm;nucleus;ribonucleoprotein complex |
| PF3D7_1112500 | N/A | RNA transcription, translation and transport factor protein, putative | 33.66 | no | 0 | 16.63 | 4.92 | 4.37 | 1.13 | 6.51 | 32.14 | 12.97 | GV | 31.72 | 6.80 | M | N/A | N/A | nucleus |
| PF3D7_1334100 | ORC4 | origin recognition complex subunit 4, putative | 117.79 | no | 0 | 4.80 | 2.93 | 13.36 | 2.56 | 3.68 | 11.85 | 44.21 | O | 204.94 | 26.96 | M | DNA binding | regulation of replication | nucleus |
| PF3D7_1336400 | N/A | RanBPM and CLTH-like protein, putative | 33.73 | no | 0 | 5.55 | 1.02 | 4.89 | 0.94 | 6.85 | 40.03 | 20.63 | GV | 50.96 | 22.19 | M | protein binding | proteasome-mediated ubiquitin-dependent protein catabolic process | cytoplasm;nucleus |
| PF3D7_0824800 | N/A | conserved Plasmodium membrane protein | 154.87 | no | 4 | 33.19 | 14.08 | 8.48 | 2.05 | 2.34 | 22.54 | 39.11 | O | 43.49 | 23.65 | M | N/A | N/A | transcription regulator complex;nucleus;integral component of membrane |
| PF3D7_1409800 | CELF2 | CUGBP Elav-like family member 2, putative | 59.27 | no | 0 | 5.96 | 31.04 | 20.57 | 1.59 | 19.66 | 39.45 | 52.01 | O | 63.42 | 75.90 | F | RNA binding;mRNA binding | regulation of translation | P granule;cytoplasm;nucleus;ribonucleoprotein complex |
| PF3D7_1459400 | N/A | conserved protein, unknown function | 40.07 | no | 1 | 57.25 | 3.15 | 19.53 | 9.52 | 7.99 | 43.79 | 8.68 | R | 0.78 | 4.09 | F | RNA binding | entry into host | nucleus |
| PF3D7_1343900 | PRPF4 | U4/U6 small nuclear ribonucleoprotein PRP4, putative | 76.19 | no | 0 | 14.18 | 5.48 | 8.13 | 2.06 | 2.09 | 12.27 | 7.10 | R | 37.98 | 6.38 | M | U4 snRNA binding;U6 snRNA binding | mRNA splicing, via spliceosome | U4/U6 x U5 tri-snRNP complex |
| PF3D7_1242800 | rabGDI | rab specific GDP dissociation inhibitor | 52.34 | no | 0 | 154.52 | 15.74 | 47.64 | 15.08 | 26.68 | 36.90 | 86.14 | R | 7.24 | 26.68 | F | GTPase activator activity;Rab GDP-dissociation inhibitor activity | vesicle-mediated transport | cytoplasm;nucleus |
| PF3D7_1015600 | HSP60 | heat shock protein 60 | 62.55 | no | 0 | 33.35 | 234.06 | 124.14 | 25.15 | 8.70 | 13.12 | 33.20 | T | 67.10 | 86.13 | F | ATP hydrolysis activity;protein binding | protein folding;protein targeting to mitochondrion;response to unfolded protein | Maurer's cleft;cytoplasm;food vacuole;mitochondrion;nucleus |
| PF3D7_1224000 | GCH1 | GTP cyclohydrolase 1 | 45.88 | no | 0 | 30.08 | 15.28 | 18.06 | 1.41 | 6.59 | 39.64 | 36.17 | GV | 82.07 | 18.30 | M | GTP binding;GTP cyclohydrolase I activity;zinc ion binding | folic acid-containing compound biosynthetic process;one-carbon metabolic process;response to xenobiotic stimulus;tetrahydrobiopterin biosynthetic process | cytoplasm;mitochondrion;nucleus |
| PF3D7_1332900 | cyto-IRS | isoleucine--tRNA ligase, putative | 151.31 | no | 0 | 70.15 | 27.52 | 34.03 | 6.90 | 2.42 | 9.67 | 43.49 | R | 27.65 | 22.51 | 0 | isoleucine-tRNA ligase activity | isoleucyl-tRNA aminoacylation;response to xenobiotic stimulus | cytoplasm;nucleus |
| PF3D7_1317800 | RPS19 | 40S ribosomal protein S19 | 16.76 | no | 0 | 284.85 | 310.85 | 207.25 | 24.19 | 182.37 | 237.39 | 185.00 | T | 153.69 | 108.15 | M | RNA binding;structural constituent of ribosome | ribosomal small subunit assembly;translation | cytosolic small ribosomal subunit;nucleus |
| PF3D7_1470900 | N/A | proteasome subunit beta type-2, putative | 22.86 | no | 0 | 23.46 | 23.56 | 66.26 | 14.27 | 45.07 | 169.73 | 118.03 | GV | 34.92 | 63.97 | F | endopeptidase activity | proteasomal protein catabolic process;proteasomal ubiquitin-independent protein catabolic process;proteasome-mediated ubiquitin-dependent protein catabolic process;ubiquitin-dependent protein catabolic process | cytoplasm;extracellular vesicle;nucleus;proteasome core complex |
| PF3D7_1139300 | AP2-G5 | AP2 domain transcription factor AP2-G5 | 309.44 | no | 0 | 64.79 | 41.42 | 12.07 | 0.88 | 1.99 | 22.23 | 190.04 | O | 120.04 | 21.26 | M | DNA-binding transcription factor activity;protein binding;sequence-specific DNA binding | regulation of transcription, DNA-templated | heterochromatin;nucleus |
| PF3D7_1036900 | N/A | conserved Plasmodium protein, unknown function | 192.68 | no | 0 | 225.83 | 57.69 | 32.04 | 4.98 | 21.91 | 169.45 | 201.15 | R | 313.27 | 168.45 | M | mRNA binding | N/A | RNA N6-methyladenosine methyltransferase complex;nucleus |
| PF3D7_1004000 | N/A | 60S ribosomal protein L13, putative | 23.75 | no | 0 | 303.74 | 246.76 | 124.08 | 8.83 | 112.30 | 94.36 | 260.90 | R | 253.53 | 64.17 | M | mRNA binding;structural constituent of ribosome | negative regulation of translation;translation | cytosolic large ribosomal subunit;nucleus;ribosome |
| PF3D7_1322100 | SET2 | histone-lysine N-methyltransferase SET2 | 300.71 | no | 0 | 303.74 | 17.82 | 6.80 | 0.55 | 2.71 | 17.71 | 60.61 | R | 7.37 | 33.13 | F | chromatin binding;histone methyltransferase activity (H3-K36 specific);histone-lysine N-methyltransferase activity;nucleosomal histone binding;protein binding | antigenic variation;histone H3-K36 methylation;histone H3-K36 trimethylation;regulation of transcription, DNA-templated | chromatin;chromosome;nuclear periphery;nucleus |
| PF3D7_0707400 | ATAD3A | ATPase family AAA domain-containing protein 3A, putative | 78.19 | no | 0 | 10.60 | 32.11 | 17.27 | 1.77 | 4.18 | 31.99 | 85.93 | O | 45.42 | 61.66 | F | zinc ion binding | mitochondrion organization | mitochondrion;nucleus |
| PF3D7_1439500 | ORP2 | oocyst rupture protein 2, putative | 123.02 | no | 0 | 6.12 | 0.56 | 10.69 | 6.11 | 3.91 | 20.15 | 10.89 | GV | 74.44 | 7.59 | M | DNA-binding transcription activator activity, RNA polymerase II-specific;RNA polymerase II cis-regulatory region sequence-specific DNA binding | regulation of transcription, DNA-templated | cell periphery;cytoplasm;nucleus |
| PF3D7_1448300 | N/A | conserved protein, unknown function | 191.02 | no | 0 | 25.46 | 26.85 | 41.64 | 16.79 | 8.09 | 25.04 | 27.92 | T | 46.06 | 36.43 | M | N/A | N/A | nucleus |
| PF3D7_1220100 | PRP17 | pre-mRNA-processing factor 17, putative | 72.54 | no | 0 | 23.80 | 6.83 | 6.56 | 0.80 | 2.46 | 16.04 | 83.11 | O | 28.74 | 13.83 | M | RNA binding | RNA splicing, via transesterification reactions;generation of catalytic spliceosome for second transesterification step;mRNA 3'-splice site recognition | spliceosomal complex;nucleus |
| PF3D7_1121700 | GCN20 | protein GCN20 | 95.37 | no | 0 | 52.14 | 34.67 | 13.09 | 0.80 | 4.85 | 9.01 | 25.14 | R | 14.82 | 11.87 | 0 | ATP binding;ATPase-coupled transmembrane transporter activity | regulation of translation | nucleus |
| PF3D7_0932300 | M18AAP | M18 aspartyl aminopeptidase | 65.63 | no | 0 | 22.90 | 14.45 | 15.17 | 2.43 | 1.59 | 9.75 | 51.93 | O | 47.20 | 82.35 | F | metallopeptidase activity;zinc ion binding | proteolysis | cytosol;nucleus |
| PF3D7_0211800 | AsnRS | asparagine--tRNA ligase | 70.50 | no | 0 | 206.62 | 66.39 | 69.96 | 9.22 | 9.83 | 11.56 | 80.93 | R | 114.67 | 86.61 | M | asparagine-tRNA ligase activity | asparaginyl-tRNA aminoacylation | nucleus |
| PF3D7_0411100 | MED8 | mediator of RNA polymerase II transcription subunit 8 | 23.97 | no | 0 | 34.82 | 2.70 | 5.48 | 0.45 | 4.61 | 105.53 | 52.18 | GV | 39.21 | 54.38 | F | protein binding | regulation of transcription | mediator complex;nucleus |
| PF3D7_1244200 | TFB2 | RNA polymerase II transcription factor B subunit 2, putative | 111.87 | no | 0 | 73.96 | 19.46 | 11.81 | 1.23 | 2.67 | 21.02 | 38.76 | R | 14.92 | 18.47 | F | double-stranded DNA binding | nucleotide-excision repair;phosphorylation of RNA polymerase II C-terminal domain;transcription by RNA polymerase II | nucleus;transcription factor TFIIH core complex;transcription factor TFIIH holo complex |
| PF3D7_1317300 | N/A | RNA-binding protein, putative | 34.86 | no | 0 | 13.03 | 28.33 | 29.23 | 7.61 | 16.02 | 68.32 | 42.74 | GV | 144.93 | 56.17 | M | N/A | N/A | nucleus |
| PF3D7_1368800 | ERCC4 | DNA repair endonuclease XPF, putative | 204.53 | no | 0 | 4.91 | 1.45 | 10.48 | 2.34 | 2.25 | 22.83 | 77.44 | O | 38.48 | 19.24 | M | DNA binding;damaged DNA binding;endodeoxyribonuclease activity;single-stranded DNA binding;single-stranded DNA endodeoxyribonuclease activity | DNA repair;double-strand break repair via homologous recombination;nucleotide-excision repair involved in interstrand cross-link repair;nucleotide-excision repair, DNA incision, 5'-to lesion;resolution of meiotic recombination intermediates | nucleotide-excision repair factor 1 complex |
| PF3D7_0516900 | RPL2 | 60S ribosomal protein L2 | 28.00 | no | 0 | 1604.47 | 3148.30 | 873.98 | 471.41 | 68.91 | 50.94 | 352.43 | T | 194.22 | 193.81 | 0 | RNA binding;structural constituent of ribosome | cytoplasmic translation;translation | cytosolic large ribosomal subunit;nucleus |
| PF3D7_0810600 | DBP1 | ATP-dependent RNA helicase DBP1, putative | 108.63 | no | 0 | 122.19 | 50.03 | 29.03 | 2.43 | 14.18 | 87.40 | 88.86 | R | 132.97 | 88.01 | M | RNA binding;RNA helicase activity;mRNA binding | regulation of transcription | P granule;nucleus |
| PF3D7_0111300 | N/A | P-loop containing nucleoside triphosphate hydrolase | 139.21 | no | 0 | 4.91 | 2.18 | 12.37 | 2.73 | 2.33 | 21.09 | 24.92 | O | 171.23 | 27.02 | M | DNA binding | DNA replication | DNA replication factor C complex;nucleus |
| PF3D7_1230700 | SEC13 | protein transport protein SEC13 | 90.75 | no | 0 | 486.15 | 167.29 | 448.51 | 624.52 | 33.84 | 16.17 | 97.63 | S | 92.83 | 56.97 | M | protein binding | COPII-coated vesicle budding | COPII vesicle coat;Maurer's cleft;cytosol;endoplasmic reticulum exit site;nuclear pore;nuclear pore outer ring;nucleoplasm |
| PF3D7_1460700 | RPL27 | 60S ribosomal protein L27 | 16.75 | no | 0 | 232.90 | 242.16 | 76.26 | 12.19 | 80.06 | 73.49 | 234.32 | T | 350.76 | 353.44 | 0 | RNA binding;structural constituent of ribosome | translation | cytosolic large ribosomal subunit;nucleus |
| PF3D7_0513300 | PNP | purine nucleoside phosphorylase | 26.86 | no | 0 | 215.33 | 280.18 | 266.95 | 27.55 | 43.75 | 25.49 | 333.54 | O | 6.92 | 84.51 | F | purine-nucleoside phosphorylase activity | inosine catabolic process;purine nucleoside catabolic process;response to xenobiotic stimulus | nucleus |
| PF3D7_1135100 | PPM8 | protein phosphatase PPM8, putative | 79.93 | no | 0 | 61.84 | 12.93 | 21.60 | 13.23 | 3.33 | 51.75 | 15.64 | R | 42.47 | 125.43 | F | magnesium-dependent protein serine/threonine phosphatase activity;protein serine/threonine phosphatase activity | | cytoplasm;nucleus |
| PF3D7_0604600 | N/A | DNA helicase, putative | 206.43 | no | 0 | 4.44 | 1.15 | 1.88 | 0.27 | 0.86 | 5.31 | 20.11 | O | 9.57 | 4.24 | M | ATP-dependent DNA/DNA annealing activity;ATP-dependent activity, acting on DNA;DNA helicase activity;endodeoxyribonuclease activity | DNA repair;DNA rewinding;cellular response to DNA damage stimulus;replication fork processing;replication fork protection | nuclear replication fork;nucleus |
| PF3D7_0934800 | PKAc | cAMP-dependent protein kinase catalytic subunit | 40.20 | no | 0 | 11.11 | 1.24 | 56.80 | 63.13 | 4.29 | 18.80 | 154.44 | O | 145.68 | 25.66 | M | cAMP-dependent protein kinase activity;protein binding | cell differentiation;entry into host;peptidyl-serine phosphorylation;signal transduction | cAMP-dependent protein kinase complex;nucleus |
| PF3D7_0218500 | SNRPD2 | small nuclear ribonucleoprotein Sm D2, putative | 11.71 | no | 0 | 70.47 | 20.33 | 14.76 | 1.80 | 55.51 | 57.77 | 38.27 | R | 61.34 | 33.14 | M | RNA binding | RNA splicing;RNA splicing, via transesterification reactions;spliceosomal snRNP assembly | U1 snRNP;U2 snRNP;U4/U6 x U5 tri-snRNP complex;U5 snRNP;catalytic step 2 spliceosome;precatalytic spliceosome;spliceosomal complex;nucleus |
| PF3D7_1363600 | N/A | conserved Plasmodium protein, unknown function | 27.36 | no | 0 | 5.20 | 4.07 | 7.46 | 1.76 | 14.44 | 35.92 | 0.00 | GV | 24.62 | 8.88 | M | N/A | N/A | mediator complex;nucleus |
| PF3D7_1009200 | N/A | ribonuclease, putative | 106.46 | no | 0 | 32.31 | 5.41 | 3.43 | 0.64 | 2.47 | 15.81 | 48.29 | O | 9.86 | 17.02 | F | enzymatic activity;RNA binding | RNA processing | nucleus |
| PF3D7_1446700 | AKIT4 | apicomplexan kinetochore protein 4, putative | 42.63 | no | 0 | 20210.00 | 45047.00 | 25.67 | 26481.00 | 31686.00 | 89.37 | 62.00 | T | 62.14 | 68.67 | F | DNA binding | DNA replication | kinetochore;nucleus |
| PF3D7_1427900 | N/A | Leucine-rich repeat protein | 33.01 | no | 0 | 15.73 | 45348.00 | 73.32 | 13.72 | 13.88 | 37.15 | 122.83 | T | 304.46 | 45554.00 | F | histone binding | N/A | nucleus |
| PF3D7_1356100 | N/A | conserved Plasmodium protein, unknown function | 113.75 | no | 0 | 7.55 | 13.51 | 31.60 | 17.96 | 13.50 | 137.73 | 121.61 | GV | 98.95 | 875.90 | F | N/A | N/A | nucleus |
| PF3D7_0927600 | N/A | RNA-binding protein, putative | 69.55 | no | 0 | 3.37 | 2.49 | 22.11 | 7.55 | 5.67 | 117.60 | 217.73 | O | 76.28 | 555.34 | F | histone pre-mRNA stem-loop binding;mRNA binding | mRNA 3'-end processing by stem-loop binding and cleavage;mRNA transport | cytoplasm;histone pre-mRNA 3'end processing complex;nucleus |
| PF3D7_1235500 | MT-A70.2 | N6-adenosine-methyltransferase, putative | 75.05 | no | 0 | 29.19 | 22.46 | 23.97 | 3.11 | 9.87 | 13.82 | 74.37 | O | 54.81 | 69.24 | F | mRNA binding;protein binding | mRNA methylation | RNA N6-methyladenosine methyltransferase complex;nucleus |
| PF3D7_1014900 | KIC8 | protein KIC8 | 267.98 | no | 0 | 6.48 | 0.81 | 13.39 | 11.94 | 1.25 | 57.32 | 227.16 | O | 12.04 | 357.97 | F | N/A | N/A | nucleus |
| PF3D7_0519800 | N/A | EELM2 domain-containing protein, putative | 78.29 | no | 0 | 162.14 | 132.95 | 33.53 | 7.72 | 14.95 | 22.80 | 1124.78 | O | 128.29 | 63.31 | M | lysine-acetylated histone binding | | nucleus |
| PF3D7_1359400 | CELF1 | CUGBP Elav-like family member 1 | 57.11 | no | 0 | 171.39 | 220.50 | 137.97 | 29.61 | 25.06 | 72.25 | 76.36 | T | 174.59 | 62.10 | M | RNA binding;mRNA binding | N/A | cytoplasm;nucleus;ribonucleoprotein complex |
| PF3D7_1329500 | N/A | conserved protein, unknown function | 218.44 | no | 0 | 527.52 | 33.64 | 25.15 | 4.57 | 5.28 | 20.58 | 337.46 | R | 14.02 | 32.26 | F | N/A | N/A | nucleus |
| PF3D7_0726300 | PMS1 | DNA mismatch repair protein PMS1, putative | 156.84 | no | 0 | 22.25 | 2.79 | 10.82 | 3.61 | 1.81 | 10.37 | 51.67 | O | 114.31 | 7.48 | M | ATP binding;ATP hydrolysis activity;DNA binding;damaged DNA binding | DNA replication;mismatch repair | MutLalpha complex;mismatch repair complex;nucleus |
| PF3D7_0912900 | RPN8 | 26S proteasome regulatory subunit RPN8, putative | 39.14 | no | 0 | 108.90 | 25.82 | 57.04 | 13.57 | 17.04 | 66.53 | 193.85 | O | 39.88 | 25.37 | M | protein binding | proteasome-mediated ubiquitin-dependent protein catabolic process | proteasome complex; nucleus; cytoplasm |
| PF3D7_0822300 | SNRPG | small nuclear ribonucleoprotein G, putative | 9.51 | no | 0 | 84.40 | 17.64 | 13.19 | 2.06 | 58.39 | 259.25 | 63.08 | GV | 24.96 | 27.20 | 0 | RNA binding | RNA splicing, via transesterification reactions;mRNA splicing, via spliceosome | P granule;SMN-Sm protein complex;U1 snRNP;U12-type spliceosomal complex;U2 snRNP;U2-type prespliceosome;U4 snRNP;U4/U6 x U5 tri-snRNP complex;U5 snRNP;catalytic step 2 spliceosome;precatalytic spliceosome;ribonucleoprotein complex;sno(s)RNA-containing ribonucleoprotein complex;spliceosomal tri-snRNP |
| PF3D7_0817300 | N/A | conserved Plasmodium protein, unknown function | 254.75 | no | 0 | 20.79 | 106.98 | 59.01 | 3.53 | 2.99 | 16.04 | 62.13 | T | 17.29 | 23.85 | F | N/A | N/A | nucleus |
| PF3D7_1369500 | N/A | nuclear cap-binding protein subunit 1, putative | 131.37 | no | 0 | 69.83 | 17.18 | 7.99 | 0.77 | 0.74 | 7.36 | 11.75 | R | 19.26 | 13.68 | M | mRNA cap binding | regulation of translation | nucleus |
| PF3D7_1425900 | N/A | protein HGH1, putative | 43.16 | no | 0 | 97.07 | 44.36 | 10.04 | 1.19 | 16.18 | 45.26 | 42.05 | R | 10.29 | 22.48 | F | protein binding | regulation of transcription; TF chaperone | nucleus |
| PF3D7_1307000 | RRP40 | exosome complex component RRP40, putative | 28.29 | no | 0 | 44.01 | 14.02 | 11.76 | 1.32 | 18.46 | 19.31 | 8.05 | R | 11.04 | 6.87 | M | RNA binding | CUT catabolic process;U4 snRNA 3'-end processing;exonucleolytic catabolism of deadenylated mRNA;exonucleolytic trimming to generate mature 3'-end of 5.8S rRNA from tricistronic rRNA transcript (SSU-rRNA, 5.8S rRNA, LSU-rRNA);nuclear polyadenylation-dependent rRNA catabolic process;nuclear polyadenyl | cytoplasmic exosome (RNase complex);exosome (RNase complex);nuclear exosome (RNase complex) |
| PF3D7_1125500 | SNRPD1 | small nuclear ribonucleoprotein Sm D1, putative | 14.34 | no | 0 | 75.14 | 24.58 | 11.72 | 1.08 | 30.60 | 89.63 | 54.68 | GV | 13.90 | 10.38 | 0 | RNA binding;protein binding | RNA splicing;RNA splicing, via transesterification reactions;mRNA splicing, via spliceosome;spliceosomal snRNP assembly | SMN-Sm protein complex;U1 snRNP;U12-type spliceosomal complex;U2 snRNP;U4 snRNP;U5 snRNP;catalytic step 2 spliceosome;commitment complex;nucleus;pICln-Sm protein complex;precatalytic spliceosome;small nuclear ribonucleoprotein complex;spliceosomal tri-snRNP complex |
| PF3D7_0210100 | N/A | ribosomal protein L37ae, putative | 10.80 | no | 0 | 550.93 | 482.78 | 198.68 | 72.4 | 527.03 | 818.94 | 293.43 | GC V | 118.29 | 475.14 | F | RNA binding;structural constituent of ribosome | translation | cytosolic large ribosomal subunit;nucleus |
| PF3D7_1446600 | CEN2 | centrin-2 | 19.31 | no | 0 | 6.49 | 2.71 | 102.92 | 55.53 | 79.45 | 450.14 | 625.36 | O | 213.42 | 717.72 | F | RNA binding;calcium ion binding | centriole replication;mitotic cell cycle | centriole;centrosome;nucleus |
| PF3D7_0303300 | RPB6 | DNA-directed RNA polymerases I, II, and III subunit RPABC2, putative | 17.83 | no | 0 | 26.80 | 9.18 | 8.42 | 0.74 | 19.62 | 37.33 | 33.93 | GV | 19.80 | 10.59 | M | DNA-directed 5'-3' RNA polymerase activity | transcription by RNA polymerase II | RNA polymerase I complex;RNA polymerase II, core complex;RNA polymerase III complex |
| PF3D7_0503200 | CDC6 | cell division control protein 6, putative | 115.09 | no | 0 | 12.88 | 4.90 | 10.67 | 2.32 | 1.87 | 83.12 | 330.08 | O | 570.22 | 504.11 | M | DNA replication origin binding | DNA replication initiation;mitotic DNA replication checkpoint signaling;mitotic cell cycle | mitochondrion;nuclear origin of replication recognition complex |
| PF3D7_1469800 | MED22 | mediator of RNA polymerase II transcription subunit 22 | 25.17 | no | 0 | 7.43 | 10.00 | 10.90 | 3.11 | 113.91 | 815.29 | 172.11 | GV | 200.55 | 434.25 | F | protein binding | regulation of transcription | mediator complex;nucleus |
| PF3D7_1006800 | GBP2 | G-strand-binding protein 2 | 29.53 | no | 0 | 705.47 | 275.18 | 206.92 | 48.93 | 46.83 | 44.27 | 150.79 | R | 221.83 | 116.06 | M | G-quadruplex DNA binding;G-rich strand telomeric DNA binding;RNA binding;mRNA binding;telomeric DNA binding | | cytoplasm;food vacuole;nucleus;ribonucleoprotein complex |
| PF3D7_0215800 | ORC5 | origin recognition complex subunit 5 | 103.99 | no | 0 | 7.16 | 1.23 | 11.68 | 1.93 | 1.70 | 12.83 | 24.33 | O | 234.01 | 8.45 | M | ATP binding;ATP hydrolysis activity;DNA replication origin binding | DNA replication initiation | nuclear origin of replication recognition complex;nuclear pre-replicative complex |
| PF3D7_1132300 | N/A | nucleic acid binding protein, putative | 31.73 | no | 0 | 3.88 | 7.62 | 11.31 | 3.93 | 11.03 | 74.42 | 187.67 | O | 589.80 | 122.40 | M | RNA binding | | nucleus |
| PF3D7_1022400 | SRSF4 | serine/arginine-rich splicing factor 4 | 62.12 | no | 0 | 259.04 | 261.66 | 121.92 | 24.13 | 9.22 | 12.27 | 101.14 | T | 81.36 | 160.91 | F | RNA binding | RNA splicing;RNA splicing, via transesterification reactions | nucleus |
| PF3D7_1360900 | N/A | RNA-binding protein, putative | 44.81 | no | 0 | 10.41 | 5.12 | 119.79 | 31.44 | 22.50 | 27.40 | 65.92 | T | 301.08 | 75.45 | M | RNA binding | N/A | nucleus;ribonucleoprotein complex |
| PF3D7_0503400 | ADF1 | actin-depolymerizing factor 1 | 13.74 | no | 0 | 18.60 | 30.75 | 314.90 | 127.30 | 272.77 | 236.67 | 598.47 | O | 20.83 | 296.92 | F | actin filament binding;actin monomer binding | actin cytoskeleton organization;actin filament depolymerization;actin filament severing;cell motility;cytoplasmic actin-based contraction involved in cell motility | actin cytoskeleton;cytoplasm;cytosol;nucleus |
| PF3D7_1453700 | P23 | HSP90 co-chaperone p23 | 30.49 | no | 0 | 518.31 | 508.28 | 263.06 | 18.22 | 311.77 | 314.46 | 326.45 | R | 482.70 | 332.69 | M | Hsp90 protein binding;chaperone binding | chaperone-mediated protein complex assembly;negative regulation of ATP-dependent activity;protein folding | cytoplasm;cytosol;nucleus |
| PF3D7_1208100 | N/A | conserved Plasmodium protein, unknown function | 533.64 | no | 2 | 21.52 | 11.86 | 2.48 | 0.23 | 1.08 | 12.13 | 8.65 | R | 11.63 | 7.18 | 0 | N/A | N/A | 90S preribosome;nucleolus;small-subunit processome |
| PF3D7_1145100 | SEC21 | coatomer subunit gamma, putative | 124.37 | no | 0 | 37.73 | 10.65 | 25.41 | 5.15 | 4.12 | 19.59 | 29.51 | R | 65.24 | 24.45 | M | protein binding | endoplasmic reticulum to Golgi vesicle-mediated transport;intra-Golgi vesicle-mediated transport;protein secretion;retrograde vesicle-mediated transport, Golgi to endoplasmic reticulum;vesicle-mediated transport | COPI vesicle coat;Golgi apparatus;Golgi membrane;cytosol;endoplasmic reticulum;endoplasmic reticulum-Golgi intermediate compartment;membrane;nucleus |
| PF3D7_0616600 | N/A | HROB domain-containing protein | 73.90 | no | 0 | 6.02 | 1.53 | 11.94 | 3.40 | 2.47 | 52.95 | 627.19 | O | 24.05 | 249.27 | F | DNA binding | recombinational repair, DNA repair | nucleus |
| PF3D7_1461600 | SF3B2 | splicing factor 3B subunit 2, putative | 79.34 | no | 0 | 152.59 | 71.38 | 41.67 | 6.14 | 24.04 | 20.53 | 106.24 | R | 99.31 | 38.84 | M | RNA binding | mRNA splicing, via spliceosome | U2 snRNP;U2-type spliceosomal complex;catalytic step 2 spliceosome;nucleus;precatalytic spliceosome;spliceosomal complex |
| PF3D7_0803800 | N/A | proteasome subunit beta type-4 | 30.87 | no | 0 | 211.58 | 102.81 | 184.94 | 66.31 | 123.91 | 238.56 | 243.48 | O | 95.98 | 189.70 | F | endopeptidase activity | proteasomal protein catabolic process;proteasomal ubiquitin-independent protein catabolic process;proteasome-mediated ubiquitin-dependent protein catabolic process | cytoplasm;nucleus;proteasome core complex |

| | | | | | | | | | | | | | | | | | | | |
|---|---|---|---|---|---|---|---|---|---|---|---|---|---|---|---|---|---|---|---|
| PF3D7_1003500 | N/A | 40S ribosomal protein S20e, putative | 13.51 | no | 0 | 117.84 | 105.43 | 49.56 | 6.35 | 59.69 | 75.21 | 83.53 | R | 82.17 | 50.56 | M | structural constituent of ribosome | translation | cytosolic small ribosomal subunit;nucleus;small ribosomal subunit |
| PF3D7_1031500 | DDX3X | ATP-dependent DNA helicase DDX3X | 107.80 | no | 0 | 71.32 | 12.20 | 39.01 | 8.12 | 8.93 | 59.66 | 156.10 | O | 87.61 | 150.99 | F | ATP hydrolysis activity;DNA helicase activity | mRNA splicing, via spliceosome | cytoplasm;nucleus |
| PF3D7_0921800 | RPF1 | ribosome production factor 1, putative | 39.17 | no | 0 | 64.18 | 14.20 | 5.10 | 0.41 | 6.26 | 64.51 | 93.79 | O | 4.74 | 40.86 | F | protein binding | maturation of 5.8S rRNA;maturation of LSU-rRNA;rRNA processing;ribosomal large subunit assembly | nucleolus;preribosome, large subunit precursor |
| PF3D7_0729500 | MT-A70 | N6-adenosine-methyltransferase MT-A70 | 89.63 | no | 0 | 31.54 | 17.97 | 11.09 | 1.30 | 1.71 | 8.81 | 19.92 | R | 61.92 | 25.32 | M | mRNA (2'-O-methyladenosine-N6-)-methyltransferase activity;mRNA (N6-adenosine)-methyltransferase activity;methyltransferase activity;protein binding | RNA methylation;RNA modification;mRNA methylation | RNA N6-methyladenosine methyltransferase complex;cytoplasm;nuclear speck;nucleus |
| PF3D7_1126200 | RPS18 | 40S ribosomal protein S18, putative | 17.89 | no | 0 | 973.78 | 448.53 | 220.40 | 70.31 | 572.01 | 287.04 | 371.84 | R | 71.95 | 135.15 | F | RNA binding;structural constituent of ribosome | translation | cytosol;nucleus;small ribosomal subunit |
| PF3D7_1460300 | RPL29 | 60S ribosomal protein L29, putative | 8.01 | no | 0 | 112.73 | 112.98 | 32.31 | 6.01 | 60.47 | 125.83 | 321.90 | O | 129.56 | 177.44 | F | structural constituent of ribosome | cytoplasmic translation | cytosolic large ribosomal subunit;nucleus |
| PF3D7_1451200 | N/A | conserved Plasmodium protein, unknown function | 176.23 | no | 0 | 288.80 | 101.55 | 56.25 | 4.96 | 2.47 | 12.68 | 239.53 | R | 46.39 | 66.70 | F | protein binding | N/A | nucleus |
| PF3D7_1250800 | N/A | DNA repair protein rhp16, putative | 192.17 | no | 0 | 58.81 | 38.20 | 25.28 | 5.26 | 6.29 | 7.57 | 88.79 | O | 188.32 | 29.03 | M | ATP binding;helicase activity | nucleotide-excision repair | nucleotide-excision repair complex |
| PF3D7_0906100 | VPS46 | vacuolar protein sorting-associated protein 46, putative | 21.85 | no | 0 | 7.57 | 9.48 | 115.18 | 65.30 | 252.60 | 151.01 | 259.27 | O | 15.10 | 385.61 | F | protein binding | endosome transport via multivesicular body sorting pathway;late endosome to vacuole transport;protein transport | ESCRT III complex;multivesicular body |
| PF3D7_0416300 | MCM9 | DNA helicase MCM9, putative | 171.31 | no | 0 | 9.40 | 5.04 | 16.46 | 3.63 | 1.93 | 12.57 | 624.75 | O | 22.80 | 79.50 | F | ATP-dependent activity, acting on DNA;DNA replication origin binding;single-stranded DNA binding | DNA replication initiation;double-strand break repair via homologous recombination | MCM complex;nucleus |
| PF3D7_1323100 | N/A | 60S ribosomal protein L6, putative | 21.60 | no | 0 | 213.70 | 245.30 | 104.30 | 17.28 | 49.63 | 63.79 | 246.40 | O | 49.94 | 101.63 | F | RNA binding;structural constituent of ribosome | cytoplasmic translation;translation | cytosolic large ribosomal subunit;large ribosomal subunit;nucleus |
| PF3D7_0914800 | N/A | GINS complex subunit Psf3, putative | 24.84 | no | 0 | 12.20 | 1.17 | 10.89 | 2.80 | 10.18 | 141.91 | 47.86 | GV | 35.12 | 128.51 | F | N/A | DNA replication | nucleus |
| PF3D7_1474500 | SF3A1 | splicing factor 3A subunit 1, putative | 83.75 | no | 0 | 33.66 | 31.90 | 27.39 | 2.85 | 1.54 | 16.44 | 145.85 | O | 93.11 | 19.31 | M | RNA binding | mRNA splicing, via spliceosome | U2 snRNP;U2-type prespliceosome;catalytic step 2 spliceosome;nucleus;spliceosomal complex |
| PF3D7_0904500 | N/A | prefoldin subunit 4, putative | 15.33 | no | 0 | 9.39 | 5.53 | 6.01 | 1.50 | 18.88 | 86.11 | 34.47 | GV | 15.25 | 17.75 | 0 | unfolded protein binding | protein folding | cytoplasm;prefoldin complex;nucleus |
| PF3D7_0415900 | N/A | 60S ribosomal protein L15, putative | 24.13 | no | 0 | 1135.97 | 793.88 | 245.39 | 54.51 | 319.48 | 238.70 | 588.77 | R | 125.52 | 273.77 | F | RNA binding;structural constituent of ribosome | cytoplasmic translation;translation | cytosolic large ribosomal subunit;nucleus;ribosome |
| PF3D7_1304700 | CYC4 | cyclin | 32.61 | no | 0 | 6.77 | 2.13 | 6.02 | 1.20 | 4.81 | 73.07 | 61.74 | GV | 14.98 | 57.16 | F | cyclin-dependent protein serine/threonine kinase regulator activity | regulation of cell cycle;regulation of cyclin-dependent protein serine/threonine kinase activity;regulation of transcription by RNA polymerase II | nucleus |
| PF3D7_1451500 | N/A | pre-mRNA-splicing factor CWF18, putative | 17.26 | no | 0 | 11.63 | 3.85 | 2.37 | 0.12 | 8.48 | 30.41 | 0.00 | GV | 7.47 | 1.73 | M | | | U2-type spliceosomal complex;nucleus;post-mRNA release spliceosomal complex |
| PF3D7_0719700 | RPS10 | 40S ribosomal protein S10, putative | 16.48 | no | 0 | 184.47 | 244.87 | 111.26 | 25.18 | 97.83 | 121.65 | 246.83 | O | 42.03 | 159.70 | F | RNA binding;structural constituent of ribosome | ribosomal small subunit assembly;translation | cytosolic small ribosomal subunit;nucleus |
| PF3D7_1416900 | N/A | prefoldin subunit 2, putative | 16.65 | no | 0 | 21.07 | 38.82 | 60.79 | 6.54 | 585.05 | 130.72 | 86.15 | G II | 123.21 | 42.28 | M | protein folding chaperone | protein folding | cytoplasm;nucleus |
| PF3D7_1302500 | N/A | AAA domain-containing protein, putative | 325.17 | no | 0 | 16.39 | 4.55 | 9.87 | 1.26 | 2.03 | 19.46 | 16.37 | O | 78.68 | 16.89 | M | RNA binding | N/A | nucleus |
| PF3D7_0935000 | SNRPB2 | U2 small nuclear ribonucleoprotein B'', putative | 22.29 | no | 0 | 118.96 | 35.61 | 40.86 | 7.70 | 28.30 | 100.78 | 58.85 | R | 27.78 | 30.07 | 0 | U1 snRNA binding;mRNA binding | mRNA splicing, via spliceosome | U1 snRNP;nucleus |
| PF3D7_1213700 | RPB8 | DNA-directed RNA polymerases I, II, and III subunit RPABC3, putative | 16.90 | no | 0 | 36.18 | 14.63 | 7.88 | 0.44 | 11.18 | 14.97 | 60.92 | O | 6.89 | 10.34 | F | DNA-directed 5'-3' RNA polymerase activity | DNA-templated transcription, initiation | RNA polymerase I complex;RNA polymerase II, core complex;RNA polymerase III complex;nucleus |
| PF3D7_0621500 | RPP1 | ribonuclease P/MRP protein subunit RPP1, putative | 40.29 | no | 0 | 4.95 | 8.79 | 7.05 | 2.47 | 23.78 | 71.71 | 16.16 | GV | 18.24 | 8.59 | M | RNA binding;ribonuclease activity | RNA phosphodiester bond hydrolysis, endonucleolytic;tRNA processing | nucleolar ribonuclease P complex;ribonuclease MRP complex;ribonuclease P complex |
| PF3D7_1341900 | N/A | V-type proton ATPase subunit D, putative | 27.80 | no | 0 | 83.26 | 57.44 | 46.24 | 16.30 | 129.53 | 309.41 | 84.38 | GV | 99.47 | 175.84 | F | proton-transporting ATPase activity, rotational mechanism | proton transmembrane transport;vacuolar acidification | food vacuole;proton-transporting V-type ATPase complex;vacuolar membrane;vacuolar proton-transporting V-type ATPase, V0 domain;vacuolar proton-transporting V-type ATPase, V1 domain |
| PF3D7_1437000 | NAT8 | N-acetyltransferase, GNAT family | 64.19 | no | 2 | 125.63 | 22.58 | 34.74 | 7.46 | 3.67 | 21.82 | 55.54 | R | 48.69 | 44.44 | 0 | N-acetyltransferase activity | schizogony | endoplasmic reticulum;nucleus |
| PF3D7_0627900 | POP4 | ribonuclease P protein subunit p29, putative | 34.32 | no | 0 | 24.49 | 10.50 | 7.53 | 1.45 | 2.86 | 8.41 | 4.17 | R | 17.75 | 7.48 | M | ribonuclease P RNA binding;ribonuclease activity | rRNA processing | ribonuclease MRP complex;ribonuclease P complex |
| PF3D7_1112100 | N/A | protein kinase, putative | 296.62 | no | 3 | 9.79 | 3.79 | 12.81 | 6.30 | 1.98 | 18.47 | 170.18 | O | 67.12 | 67.44 | 0 | protein kinase activity | meiotic sister chromatid cohesion, centromeric;mitotic spindle assembly checkpoint signaling | condensed nuclear chromosome kinetochore;nucleus |
| PF3D7_0206000 | RAD2 | DNA repair protein RAD2, putative | 178.73 | no | 0 | 22.89 | 10.97 | 8.28 | 3.50 | 1.29 | 11.82 | 321.85 | O | 71.69 | 15.63 | M | 5'-flap endonuclease activity;damaged DNA binding;endonuclease activity;flap endonuclease activity;single-stranded DNA endodeoxyribonuclease activity | DNA repair;nucleotide-excision repair, DNA incision, 3'-to lesion | chromosome, telomeric region;nucleotide-excision repair factor 3 complex |
| PF3D7_1034900 | MRScyt | methionine--tRNA ligase | 104.17 | no | 0 | 97.54 | 9.09 | 96.73 | 18.28 | 1.95 | 9.22 | 21.02 | R | 18.29 | 65.71 | F | ATP binding;methionine-tRNA ligase activity | methionyl-tRNA aminoacylation | cytoplasm;nucleus |
| PF3D7_0618300 | N/A | 60S ribosomal protein L27a, putative | 16.72 | no | 0 | 465.64 | 215.55 | 132.00 | 96.46 | 72.34 | 99.81 | 161.29 | R | 99.37 | 210.25 | F | structural constituent of ribosome | translation | cytosolic large ribosomal subunit;large ribosomal subunit;nucleus |
| PF3D7_1307100 | UTP6 | U3 small nucleolar RNA-associated protein 6, putative | 120.72 | no | 1 | 46.66 | 24.11 | 7.20 | 1.15 | 0.96 | 4.78 | 0.85 | R | 12.92 | 14.03 | 0 | snoRNA binding | maturation of SSU-rRNA from tricistronic rRNA transcript (SSU-rRNA, 5.8S rRNA, LSU-rRNA) | Pwp2p-containing subcomplex of 90S preribosome;small-subunit processome;nucleus |
| PF3D7_1468200 | TLS1 | telomere length and silencing protein 1, putative | 27.10 | no | 0 | 12.85 | 9.45 | 14.74 | 1.60 | 6.99 | 40.19 | 19.61 | GV | 191.31 | 27.19 | M | DNA binding | regulation of mRNA splicing, via spliceosome | nucleus;spliceosomal complex |
| PF3D7_1317600 | N/A | conserved Plasmodium protein, unknown function | 73.87 | no | 0 | 4.09 | 3.26 | 9.19 | 1.28 | 1.10 | 5.73 | 9.07 | T | 2.04 | 13.59 | F | N/A | N/A | nucleus |
| PF3D7_1433300 | RBAP46 | histone-binding protein RBBP7, putative | 50.17 | no | 0 | 8.36 | 11.57 | 22.55 | 12.72 | 9.02 | 22.30 | 28.17 | O | 26.36 | 26.76 | 0 | histone binding;unfolded protein binding | chromatin assembly;chromatin assembly or disassembly;chromatin remodeling;nucleosome assembly | nucleus |
| PF3D7_1134700 | RPA2 | DNA-directed RNA polymerase I subunit RPA2, putative | 175.53 | no | 0 | 100.82 | 61.29 | 9.83 | 0.49 | 4.59 | 12.21 | 17.07 | R | 27.44 | 18.67 | M | DNA binding | regulation of transcription | nucleus |
| PF3D7_0505600 | N/A | BCNT domain-containing protein, putative | 40.20 | no | 0 | 17.18 | 5.44 | 9.63 | 1.40 | 6.66 | 275.39 | 54.41 | GV | 45.00 | 45.99 | 0 | N/A | N/A | nucleus |
| PF3D7_1031600 | GEXP15 | protein GEXP15 | 106.69 | no | 0 | 42.55 | 35.51 | 26.57 | 7.09 | 9.95 | 27.51 | 34.96 | R | 59.41 | 58.76 | 0 | protein binding;ribosome binding;protein phosphatase regulator activity | N/A | U5 snRNP;nucleus;cytoplasm |
| PF3D7_0818700 | N/A | DNA helicase, putative | 143.66 | no | 0 | 154.41 | 20.67 | 46.75 | 11.91 | 9.22 | 29.57 | 96.04 | R | 359.16 | 21.96 | M | ATP-dependent activity, acting on DNA | regulation of replication | nucleus |
| PF3D7_1004400 | N/A | RNA-binding protein, putative | 106.30 | no | 0 | 59.09 | 175.50 | 55.95 | 9.92 | 8.40 | 14.17 | 179.28 | O | 232.06 | 80.97 | M | mRNA binding | N/A | cytoplasm;nucleus |
| PF3D7_1247500 | N/A | serine/threonine protein kinase, putative | 105.54 | no | 0 | 3.56 | 1.50 | 8.26 | 4.41 | 2.73 | 40.34 | 67.03 | O | 36.78 | 125.03 | F | protein phosphorylation;protein serine threonine kinase activity;ATP binding | protein phosphorylation | cytoplasm;nucleus |
| PF3D7_0909800 | SNRPD3 | small nuclear ribonucleoprotein Sm D3, putative | 12.98 | no | 0 | 60.21 | 13.00 | 10.68 | 0.92 | 26.75 | 51.99 | 45.11 | R | 50.35 | 21.34 | M | RNA binding | spliceosomal snRNP complex activity;mRNA splicing, via spliceosome | spliceosomal complex;nucleus |
| PF3D7_1119300 | U2AF1 | splicing factor U2AF small subunit, putative | 35.31 | no | 0 | 276.90 | 112.99 | 103.24 | 34.40 | 8.39 | 17.26 | 357.34 | O | 467.43 | 101.63 | M | RNA binding;pre-mRNA 3'-splice site binding | mRNA splicing, via spliceosome | U2 snRNP;U2AF complex;nucleus;spliceosomal complex |
| PF3D7_1347200 | NT1 | nucleoside transporter 1 | 47.63 | no | 9 | 575.20 | 422.75 | 186.24 | 10.94 | 31.86 | 74.12 | 99.76 | R | 37.64 | 44.38 | F | nucleoside transmembrane transporter activity;purine nucleoside transmembrane transporter activity | adenosine transport;purine nucleobase transport;purine nucleoside transmembrane transport;response to xenobiotic stimulus | Maurer's cleft;integral component of membrane;integral component of plasma membrane;nucleus;plasma membrane |
| PF3D7_0804900 | N/A | GTPase-activating protein, putative | 56.50 | no | 0 | 50.73 | 17.52 | 28.25 | 7.30 | 2.83 | 21.42 | 265.52 | O | 39.93 | 23.51 | M | enzyme activity | COPI coating of Golgi vesicle | nucleus;cytoplasm |
| PF3D7_0916700 | HoMu | RNA-binding protein musashi, putative | 45.14 | no | 0 | 231.78 | 31.55 | 17.12 | 2.64 | 7.94 | 34.75 | 47.06 | R | 5.05 | 20.34 | F | RNA binding;mRNA binding | N/A | nucleoplasm;nucleus |
| PF3D7_0505800 | SUMO | small ubiquitin-related modifier | 11.06 | no | 0 | 265.07 | 145.70 | 157.66 | 46.42 | 313.85 | 164.82 | 64.16 | G II | 160.33 | 37.68 | M | protein binding;protein tag;ubiquitin-like protein ligase binding | modification-dependent protein catabolic process;protein sumoylation;ubiquitin-dependent protein catabolic process | nucleus |
| PF3D7_0523400 | N/A | DnaJ protein, putative | 61.90 | no | 2 | 17.11 | 22.15 | 36.86 | 7.55 | 4.19 | 23.48 | 332.79 | O | 125.88 | 70.85 | M | RNA binding;unfolded protein binding | chaperone cofactor-dependent protein refolding;protein refolding | cytoplasm;food vacuole;nucleus |
| PF3D7_1315300 | N/A | conserved protein, unknown function | 56.34 | no | 0 | 2.13 | 0.67 | 11.78 | 6.66 | 2.75 | 206.52 | 25.79 | GV | 7.24 | 176.71 | F | N/A | N/A | nucleus |
| PF3D7_0912800 | GCD10 | tRNA (adenine(58)-N(1))-methyltransferase non-catalytic subunit TRM6, putative | 72.84 | no | 0 | 29.11 | 18.96 | 9.14 | 2.07 | 1.95 | 14.50 | 23.75 | R | 5.44 | 9.99 | F | methyltransferase activity | N/A | nucleus;tRNA (m1A) methyltransferase complex |
| PF3D7_0509400 | RNAPI | RNA polymerase I | 340.67 | no | 0 | 106.77 | 76.52 | 12.86 | 1.42 | 2.56 | 11.12 | 26.32 | R | 5.62 | 14.88 | F | RNA binding | regulation of transcription | RNA polymerase I complex;nucleolus |
| PF3D7_0522200 | TAF10 | transcription initiation factor TFIID subunit 10, putative | 13.63 | no | 0 | 34.26 | 4.30 | 6.39 | 1.51 | 26.85 | 70.37 | 13.85 | GV | 17.25 | 11.92 | M | protein binding | transcription by RNA polymerase II | transcription regulator complex |
| PF3D7_0213100 | SIS1 | protein SIS1 | 37.36 | no | 0 | 48.31 | 104.03 | 46.29 | 5.24 | 16.77 | 106.80 | 353.92 | O | 117.59 | 217.98 | F | chaperone binding;unfolded protein binding | chaperone cofactor-dependent protein refolding;protein folding;response to heat;response to unfolded protein | Maurer's cleft;cytosol;nucleus |
| PF3D7_1364800 | RPB5 | DNA-directed RNA polymerases I, II, and III subunit RPABC1, putative | 24.15 | no | 0 | 185.49 | 68.80 | 33.03 | 1.20 | 81.54 | 74.57 | 61.96 | R | 53.98 | 90.20 | F | DNA binding;DNA-directed 5'-3' RNA polymerase activity | transcription by RNA polymerase II | RNA polymerase I complex;RNA polymerase II, core complex;RNA polymerase III complex;nucleus |
| PF3D7_1033700 | BDP1 | bromodomain protein 1 | 55.94 | no | 0 | 143.33 | 49.86 | 47.36 | 9.11 | 3.16 | 50.71 | 356.93 | O | 62.30 | 100.83 | F | chromatin binding;histone binding;protein binding | entry into host;regulation of transcription, DNA-templated | chromosome, telomeric region;nucleus |
| PF3D7_0510200 | CYP87 | peptidyl-prolyl cis-trans isomerase | 87.04 | no | 0 | 17.64 | 11.19 | 12.27 | 5.09 | 6.67 | 16.07 | 20.97 | O | 42.43 | 19.17 | M | cyclosporin A binding;peptidyl-prolyl cis-trans isomerase activity | protein folding;protein peptidyl-prolyl isomerization | nucleus; cytoplasm |
| PF3D7_1250600 | N/A | translation initiation factor eIF-2B subunit beta, putative | 78.75 | no | 0 | 42.49 | 7.64 | 13.97 | 2.88 | 4.61 | 20.47 | 100.78 | O | 65.77 | 23.09 | M | GTP binding;translation initiation factor activity | regulation of translational initiation;translational initiation | eukaryotic translation initiation factor 2B complex |
| PF3D7_1342400 | CK2beta2 | casein kinase II beta chain | 45.28 | no | 0 | 189.41 | 281.64 | 188.38 | 37.43 | 16.92 | 20.24 | 142.90 | T | 80.81 | 68.01 | M | protein binding;protein kinase regulator activity | protein phosphorylation | cytoplasm;nucleus;protein kinase CK2 complex |
| PF3D7_1031300 | N/A | SAE2 domain-containing protein, putative | 109.28 | no | 0 | 5.52 | 4.83 | 6.30 | 0.81 | 3.62 | 34.43 | 104.05 | O | 26.17 | 20.23 | M | N/A | N/A | nucleus |
| PF3D7_0819900 | LSM3 | U6 snRNA-associated Sm-like protein LSm3, putative | 10.70 | no | 0 | 23.44 | 7.05 | 4.52 | 1.25 | 14.44 | 47.45 | 71.71 | O | 163.54 | 86.32 | M | RNA binding | P-body assembly;RNA splicing;RNA splicing, via transesterification reactions;mRNA splicing, via spliceosome | Lsm1-7-Pat1 complex;P-body;U4/U6 x U5 tri-snRNP complex;U6 snRNP;catalytic step 2 spliceosome;nucleus;precatalytic spliceosome;sno(s)RNA-containing ribonucleoprotein complex |
| PF3D7_1037600 | XPB | TFIIH basal transcription factor complex helicase XPB subunit, putative | 102.87 | no | 0 | 53.16 | 19.24 | 21.04 | 4.81 | 1.70 | 41.13 | 138.14 | O | 77.09 | 257.49 | F | 3'-5' DNA helicase activity;ATP hydrolysis activity;DNA binding;DNA helicase activity;damaged DNA binding;helicase activity;transcription factor binding | nucleotide-excision repair, DNA duplex unwinding;nucleotide-excision repair, DNA incision;response to UV;transcription by RNA polymerase II;transcription initiation from RNA polymerase II promoter | nucleotide-excision repair factor 3 complex;transcription factor TFIIH holo complex;transcription preinitiation complex |
| PF3D7_0109800 | cPheRS | phenylalanine--tRNA ligase alpha subunit | 67.72 | no | 0 | 74.74 | 25.31 | 22.84 | 2.89 | 4.88 | 12.76 | 144.71 | O | 18.96 | 23.23 | F | phenylalanine-tRNA ligase activity | phenylalanyl-tRNA aminoacylation;response to xenobiotic stimulus | cytoplasm;cytosol;nucleus;phenylalanine-tRNA ligase complex |
| PF3D7_1127600 | N/A | CRAL/TRIO domain-containing protein, putative | 34.97 | no | 0 | 8.61 | 11.08 | 32.68 | 7.39 | 35.70 | 350.97 | 266.48 | GV | 381.57 | 811.67 | F | N/A | N/A | nucleus |
| PF3D7_1414800 | SMB1 | small nuclear ribonucleoprotein-associated protein B | 22.33 | no | 0 | 56.39 | 34.30 | 36.42 | 4.03 | 12.02 | 14.63 | 23.66 | R | 12.73 | 20.39 | F | | RNA splicing;mRNA splicing, via spliceosome | U1 snRNP;U2 snRNP;U2-type prespliceosome;U4 snRNP;U4/U6 x U5 tri-snRNP complex;U5 snRNP;catalytic step 2 spliceosome;cytoplasm;nucleus;spliceosomal complex |
| PF3D7_0823200 | N/A | RNA-binding protein, putative | 32.44 | no | 0 | 550.79 | 719.66 | 320.03 | 151.33 | 100.30 | 85.49 | 70.74 | T | 132.07 | 63.23 | M | RNA binding;mRNA binding | regulation of translation | nucleus;ribonucleoprotein complex |
| PF3D7_0526800 | N/A | conserved Plasmodium protein, unknown function | 43.78 | no | 0 | 236.44 | 60.21 | 48.77 | 4.83 | 20.03 | 45.70 | 38.42 | R | 164.30 | 68.03 | M | N/A | N/A | mediator complex;nucleus |
| PF3D7_1103600 | ALP2a | actin-like protein, putative | 57.97 | no | 0 | 20.27 | 12.52 | 7.99 | 0.86 | 3.01 | 12.02 | 19.64 | R | 15.88 | 3.26 | M | chromatin binding | ATP-dependent chromatin remodeling;histone H4 acetylation;regulation of transcription by RNA polymerase II | NuA4 histone acetyltransferase complex;SWI/SNF complex;nucleus |
| PF3D7_1321700 | SF1 | splicing factor 1 | 100.94 | no | 0 | 230.22 | 148.06 | 70.38 | 8.37 | 15.54 | 16.39 | 340.97 | O | 368.51 | 172.11 | M | mRNA binding | RNA splicing;RNA splicing, via transesterification reactions | nucleus |
| PF3D7_0719600 | N/A | 60S ribosomal protein L11a, putative | 20.23 | no | 0 | 251.71 | 413.57 | 149.95 | 46.70 | 58.45 | 50.53 | 137.89 | T | 46.08 | 48.84 | 0 | RNA binding;structural constituent of ribosome | ribosomal large subunit assembly;translation | cytosolic large ribosomal subunit;nucleus |
| PF3D7_0611800 | N/A | conserved Plasmodium protein, unknown function | 374.89 | no | 0 | 16.29 | 23.57 | 13.72 | 3.46 | 9.43 | 77.65 | 395.36 | O | 23.84 | 83.65 | F | N/A | N/A | nucleus |
| PF3D7_0917600 | PRP43 | pre-mRNA-splicing factor ATP-dependent RNA helicase PRP43, putative | 94.14 | no | 0 | 60.85 | 25.10 | 26.07 | 2.55 | 1.14 | 3.76 | 9.19 | R | 47.08 | 15.27 | M | RNA binding | N/A | nucleus |
| PF3D7_1147500 | FTB | protein farnesyltransferase subunit beta | 109.82 | no | 0 | 10.99 | 5.65 | 1.87 | 0.05 | 1.01 | 6.12 | 10.02 | R | 9.42 | 2.25 | M | protein farnesyltransferase activity;zinc ion binding | protein farnesylation;protein prenylation;response to xenobiotic stimulus | protein farnesyltransferase complex |
| PF3D7_1318400 | SMC2 | structural maintenance of chromosomes protein 2, putative | 143.06 | no | 0 | 11.66 | 4.23 | 22.11 | 7.44 | 3.26 | 22.35 | 195.74 | O | 961.78 | 136.31 | M | protein binding;DNA binding | mitotic chromosome condensation;mitotic sister chromatid segregation | chromosome, centromeric region;condensin complex;nucleus |
| PF3D7_0307700 | AKIT10 | apicomplexan kinetochore protein 10, putative | 216.26 | no | 0 | 13.01 | 44.81 | 224.44 | 169.17 | 9.51 | 37.12 | 98.73 | T | 245.33 | 56.40 | M | DNA binding | structural complex | kinetochore;nucleus |
| PF3D7_0308000 | N/A | DNA polymerase delta small subunit, putative | 57.72 | no | 0 | 4.63 | 8.80 | 14.98 | 1.67 | 6.89 | 22.70 | 21.25 | GV | 257.69 | 23.61 | M | DNA-directed DNA polymerase activity | DNA biosynthetic process;DNA strand elongation involved in DNA replication | DNA polymerase complex;delta DNA polymerase complex |
| PF3D7_1212900 | BDP2 | bromodomain protein 2, putative | 125.45 | no | 0 | 107.19 | 70.60 | 57.02 | 6.58 | 14.80 | 54.82 | 207.44 | O | 99.72 | 43.82 | M | protein binding | regulation of transcription | nucleus |
| PF3D7_1029600 | ADA | adenosine deaminase | 42.47 | no | 0 | 120.64 | 40.09 | 218.08 | 17.40 | 28.17 | 69.37 | 95.48 | T | 27.01 | 16.02 | M | adenosine deaminase activity | adenosine catabolic process;hypoxanthine salvage;inosine biosynthetic process;negative regulation of adenosine receptor signaling pathway;response to drug;response to xenobiotic stimulus | Maurer's cleft;cytosol;external side of plasma membrane |
| PF3D7_1142500 | RPL28 | 60S ribosomal protein L28 | 14.41 | no | 0 | 138.01 | 232.06 | 77.25 | 12.08 | 123.19 | 50.59 | 158.39 | T | 39.35 | 66.94 | F | | | cytosolic large ribosomal subunit;nucleus |
| PF3D7_0524500 | N/A | conserved Plasmodium protein, unknown function | 14.51 | no | 0 | 21.83 | 4.15 | 6.98 | 1.61 | 8.23 | 26.06 | 10.28 | GV | 7.57 | 7.20 | 0 | N/A | N/A | nucleus |
| PF3D7_1306900 | SNRPA | U1 small nuclear ribonucleoprotein A, putative | 51.94 | no | 0 | 31.95 | 13.15 | 14.94 | 2.93 | 1.04 | 5.07 | 42.49 | O | 33.01 | 25.13 | M | U1 snRNA binding | mRNA splicing, via spliceosome | U1 snRNP;nucleus;spliceosomal complex |
| PF3D7_1118600 | MYST | histone acetyltransferase MYST | 71.13 | no | 0 | 64.05 | 16.93 | 10.55 | 2.51 | 2.39 | 16.11 | 353.78 | O | 83.51 | 54.94 | M | histone acetyltransferase activity;histone binding;protein binding;transcription coregulator activity | histone H4-K12 acetylation;histone H4-K16 acetylation;histone H4-K5 acetylation;histone H4-K8 acetylation;negative regulation of transcription, DNA-templated;positive regulation of transcription by RNA polymerase II;response to xenobiotic stimulus | cytoplasm;host cell periphery;nucleus;symbiont-containing vacuole |
| PF3D7_1013100 | UTP13 | U3 small nucleolar RNA-associated protein 13, putative | 144.60 | no | 0 | 74.13 | 20.58 | 8.35 | 0.80 | 3.72 | 39.75 | 30.24 | R | 15.33 | 80.61 | F | U3 snoRNA binding | endonucleolytic cleavage in 5'-ETS of tricistronic rRNA transcript (SSU-rRNA, 5.8S rRNA, LSU-rRNA);endonucleolytic cleavage to generate mature 5'-end of SSU-rRNA from (SSU-rRNA, 5.8S rRNA, LSU-rRNA) | 90S preribosome;nucleolus;small-subunit processome |
| PF3D7_1231600 | PRP2 | pre-mRNA-splicing factor ATP-dependent RNA helicase PRP2, putative | 137.20 | no | 0 | 54.29 | 14.50 | 22.31 | 2.19 | 6.33 | 24.26 | 46.16 | R | 179.16 | 29.09 | M | ATP-dependent activity, acting on RNA;RNA binding;RNA helicase activity | RNA splicing;mRNA splicing, via spliceosome;regulation of cell cycle | spliceosomal complex |
| PF3D7_1358800 | RPS15 | 40S ribosomal protein S15 | 17.25 | no | 0 | 189.89 | 207.18 | 86.98 | 5.99 | 74.09 | 75.84 | 152.83 | T | 142.65 | 136.41 | M | RNA binding;small ribosomal subunit rRNA binding;structural constituent of ribosome | translation | Maurer's cleft;cytosolic small ribosomal subunit;nucleolus;nucleus |
| PF3D7_1105100 | H2B | histone H2B | 13.13 | no | 0 | 309.53 | 379.32 | 2104.54 | 810.05 | 2957.94 | 3275.46 | 21511.98 | O | 3802.21 | 854.97 | M | DNA binding;RNA binding | chromosome organization;nucleosome assembly | nucleosome;nucleus |
| PF3D7_1415400 | N/A | Btz domain-containing protein, putative | 178.50 | no | 0 | 13.21 | 35.55 | 33.82 | 4.41 | 8.76 | 33.77 | 47.12 | O | 159.31 | 48.38 | M | mRNA binding | N/A | nucleus |
| PF3D7_1211600 | LSD1 | lysine-specific histone demethylase 1, putative | 350.58 | no | 0 | 22.35 | 12.62 | 7.45 | 1.39 | 1.10 | 10.96 | 106.93 | O | 143.65 | 46.81 | M | oxidoreductase activity;polyamine oxidase activity | chromatin remodeling | nuclear periphery;nucleus |
| PF3D7_1107400 | RAD51 | DNA repair protein RAD51 | 38.59 | no | 0 | 37.19 | 12.00 | 71.34 | 14.19 | 32.42 | 202.44 | 711.67 | O | 143.07 | 928.22 | F | ATP binding;ATP-dependent activity, acting on DNA;DNA binding;DNA strand exchange activity;damaged DNA binding;double-stranded DNA binding;protein binding;single-stranded DNA binding | DNA recombinase assembly;DNA recombination;DNA repair;chromosome organization involved in meiotic cell cycle;mitochondrial double-strand break repair;mitotic recombination;reciprocal meiotic recombination;response to ionizing radiation;strand invasion | condensed nuclear chromosome;mitochondrion;nucleus |

| | | | | | | | | | | | | | | | | | | | |
|---|---|---|---|---|---|---|---|---|---|---|---|---|---|---|---|---|---|---|---|
| PF3D7_1107000 | LSM4 | U6 snRNA-associated Sm-like protein LSm4, putative | 14.20 | no | 0 | 124.50 | 88.08 | 123.78 | 164.71 | 222.49 | 137.82 | 84.79 | GV | 31.98 | 34.04 | O | RNA binding;U6 snRNA binding | P-body assembly;mRNA splicing, via spliceosome;spliceosomal snRNP assembly | P-body;U4/U6 x U5 tri-snRNP complex;U6 snRNP;nucleus;sno(s)RNA-containing ribonucleoprotein complex;spliceosomal tri-snRNP complex |
| PF3D7_0316500 | NUF2 | kinetochore protein NUF2, putative | 54.68 | no | 0 | 4.57 | 2.09 | 21.73 | 10.29 | 10.28 | 55.04 | 148.98 | O | 59.78 | 34.59 | M | protein-containing complex binding | attachment of mitotic spindle microtubules to kinetochore;chromosome segregation;kinetochore organization;meiotic chromosome segregation;mitotic spindle organization | Ndc80 complex;chromosome, centromeric region;condensed nuclear chromosome kinetochore;kinetochore;nucleus |
| PF3D7_1324200 | N/A | micro-fibrillar-associated protein, putative | 59.63 | no | 0 | 220.15 | 128.47 | 84.99 | 37.19 | 29.09 | 24.42 | 187.92 | R | 258.46 | 589.70 | F | extracellular matrix structural constituent | mRNA splicing, via spliceosome | U2-type spliceosomal complex;extracellular matrix |
| PF3D7_0913200 | EF-1beta | elongation factor 1-beta | 32.03 | no | 0 | 265.51 | 100.51 | 77.46 | 6.89 | 13.71 | 38.56 | 150.89 | R | 112.18 | 31.41 | M | guanyl-nucleotide exchange factor activity | translational elongation | Maurer's cleft;cytosol;eukaryotic translation elongation factor 1 complex;nucleus |
| PF3D7_0410300 | PPM1 | protein phosphatase PPM1, putative | 105.20 | no | 0 | 26.85 | 12.52 | 12.51 | 3.32 | 3.59 | 9.10 | 73.24 | O | 54.56 | 18.39 | M | magnesium-dependent protein serine/threonine phosphatase activity;phosphoprotein phosphatase activity;protein serine/threonine phosphatase activity | protein dephosphorylation | cytoplasm;nucleus |
| PF3D7_1330800 | N/A | RNA-binding protein, putative | 68.01 | no | 0 | 404.82 | 484.55 | 63.89 | 11.47 | 24.20 | 54.54 | 1335.48 | O | 97.42 | 902.73 | F | RNA binding;mRNA binding | RNA splicing;mRNA splicing, via spliceosome;regulation of alternative mRNA splicing, via spliceosome | ASAP complex;cytoplasm;nucleoplasm;nucleus |
| PF3D7_0934100 | XPD | TFIIH transcription factor complex helicase XPD SU | 122.84 | no | 0 | 52.70 | 25.47 | 18.32 | 3.21 | 4.92 | 18.59 | 32.07 | R | 19.99 | 15.03 | M | 5'-3' DNA helicase activity;DNA helicase activity;damaged DNA binding;protein binding;single-stranded DNA helicase activity | nucleotide-excision repair, DNA duplex unwinding;nucleotide-excision repair, DNA incision;positive regulation of mitotic recombination;regulation of mitotic recombination;response to UV;response to oxidative stress;transcription by RNA polymerase II | cytoplasm;nucleus |
| PF3D7_1458800 | N/A | DNA-directed RNA polymerase III subunit RPC5, putative | 103.36 | no | 0 | 8.36 | 3.60 | 2.21 | 0.15 | 2.33 | 11.75 | 9.45 | GV | 22.51 | 4.24 | M | RNA binding | regulation of transcription | RNA polymerase III complex |
| PF3D7_0815600 | EIF3G | eukaryotic translation initiation factor 3 subunit G, putative | 31.25 | no | 0 | 186.04 | 119.45 | 51.26 | 19.72 | 24.30 | 59.82 | 64.99 | R | 158.59 | 441.14 | F | mRNA binding;translation initiation factor activity | translation;translational initiation | nucleus |
| PF3D7_1216900 | N/A | DNA-binding chaperone, putative | 111.06 | no | 0 | 220.24 | 178.03 | 59.42 | 5.91 | 6.12 | 29.52 | 76.35 | R | 118.79 | 49.72 | M | DNA binding;protein binding | protein folding | nucleus |
| PF3D7_0708100 | RPB10 | DNA-directed RNA polymerases I, II, and III subunit RPABC5, putative | 8.24 | no | 0 | 71.20 | 23.14 | 13.25 | 0.61 | 41.22 | 177.64 | 30.92 | GV | 51.46 | 51.89 | O | DNA-directed 5'-3' RNA polymerase activity;zinc ion binding | tRNA transcription by RNA polymerase III;transcription by RNA polymerase I;transcription by RNA polymerase II | RNA polymerase I complex;RNA polymerase II, core complex;RNA polymerase III complex;nucleus |
| PF3D7_1404000 | RPB4 | DNA-directed RNA polymerase II subunit RPB4, putative | 20.05 | no | 0 | 14.20 | 9.21 | 6.56 | 1.07 | 44.29 | 865.01 | 61.34 | GV | 247.78 | 378.31 | F | RNA binding;DNA binding | regulation of transcription | nucleus |
| PF3D7_1426200 | PRMT1 | protein arginine N-methyltransferase 1 | 47.45 | no | 0 | 88.70 | 62.27 | 26.83 | 4.28 | 3.73 | 11.61 | 63.39 | R | 14.90 | 20.57 | F | protein-arginine N-methyltransferase activity | RNA metabolic process;RNA processing;histone methylation;ribosome biogenesis | cytoplasm;nucleus |
| PF3D7_0522800 | BUD31 | pre-mRNA-splicing factor BUD31, putative | 17.39 | no | 0 | 34.39 | 16.03 | 6.69 | 1.24 | 11.43 | 27.64 | 120.15 | O | 39.71 | 5.58 | M | RNA binding | RNA splicing;alternative mRNA splicing, via spliceosome;mRNA splicing, via spliceosome | U2 snRNP;nucleus;spliceosomal complex |
| PF3D7_0723400 | N/A | conserved Plasmodium protein, unknown function | 167.92 | no | 0 | 192.48 | 79.38 | 22.21 | 2.19 | 4.65 | 21.64 | 144.23 | R | 237.30 | 81.59 | M | N/A | N/A | nucleus |
| PF3D7_0704200 | NSUN1 | RNA cytosine C(5)-methyltransferase, putative | 141.23 | no | 0 | 17.70 | 12.83 | 4.69 | 0.51 | 0.66 | 7.06 | 14.21 | R | 1.49 | 6.38 | F | methyltransferase activity | RNA methylation | nucleus |
| PF3D7_1426700 | PEPC | phosphoenolpyruvate carboxylase | 133.96 | no | 0 | 26.11 | 10.19 | 25.91 | 9.77 | 6.99 | 63.94 | 105.02 | O | 217.02 | 79.21 | M | phosphoenolpyruvate carboxylase activity | carbon fixation | cytosol;nucleus |
| PF3D7_1117800 | MLH | DNA mismatch repair protein MLH | 118.31 | no | 0 | 13.04 | 3.70 | 10.35 | 2.81 | 2.66 | 12.61 | 20.56 | O | 85.60 | 11.79 | M | ATP binding;ATP hydrolysis activity;endonuclease activity;protein binding | mismatch repair | MutLalpha complex;chiasma;chromosome, telomeric region;mismatch repair complex |
| PF3D7_0728600 | N/A | RING zinc finger protein, putative | 253.97 | no | 0 | 7.82 | 1.93 | 18.21 | 6.73 | 8.82 | 86.84 | 270.45 | O | 914.65 | 299.48 | M | RNA binding | N/A | nucleus;cytoplasm |
| PF3D7_1205500 | N/A | zinc finger protein, putative | 204.85 | no | 0 | 230.33 | 15.87 | 84.84 | 23.09 | 15.07 | 113.73 | 119.58 | R | 200.69 | 86.48 | M | ubiquitin protein ligase activity | protein ubiquitination | cytoplasm;nucleus |
| PF3D7_0807300 | RAB18 | ras-related protein Rab-18 | 23.16 | no | 0 | 166.44 | 89.06 | 92.04 | 19.05 | 134.25 | 44.56 | 33.79 | R | 38.90 | 14.27 | M | GTP binding;GTPase activity | intracellular protein transport;small GTPase mediated signal transduction | Golgi apparatus;endomembrane system;membrane;nucleus |
| PF3D7_0407700 | N/A | TH1 domain-containing protein, putative | 197.41 | no | 0 | 6.11 | 0.33 | 13.33 | 9.64 | 3.60 | 21.13 | 51.24 | O | 208.91 | 61.67 | M | RNA binding | negative regulation of transcription elongation from RNA polymerase II promoter | NELF complex;nucleus |
| PF3D7_1010600 | eIF2beta | eukaryotic translation initiation factor 2 subunit beta | 25.32 | no | 0 | 429.26 | 92.40 | 93.89 | 18.73 | 39.05 | 106.08 | 58.49 | R | 91.58 | 89.95 | O | RNA binding;mRNA binding;protein binding;translation initiation factor activity;translation initiation factor binding | formation of cytoplasmic translation initiation complex;formation of translation preinitiation complex;translational initiation | cytoplasm;eukaryotic translation initiation factor 2 complex;nucleus |
| PF3D7_1407100 | NOP1 | rRNA 2'-O-methyltransferase fibrillarin, putative | 34.13 | no | 0 | 246.22 | 243.57 | 61.48 | 23.22 | 14.98 | 18.28 | 129.25 | R | 100.49 | 69.74 | M | RNA binding;histone-glutamine methyltransferase activity;rRNA methyltransferase activity | box C/D RNA 3'-end processing;histone glutamine methylation;rRNA methylation;rRNA processing | Cajal body;box C/D RNP complex;nucleolus;nucleus;small-subunit processome;sno(s)RNA-containing ribonucleoprotein complex |
| PF3D7_0323700 | SART1 | U4/U6.U5 tri-snRNP-associated protein 1, putative | 82.90 | no | 0 | 35.40 | 16.66 | 13.93 | 2.00 | 2.24 | 14.30 | 77.78 | O | 94.06 | 66.24 | M | RNA binding | mRNA cis splicing, via spliceosome;maturation of 5S rRNA | U4/U6 x U5 tri-snRNP complex;nucleus |
| PF3D7_1331700 | N/A | glutamine--tRNA ligase, putative | 108.53 | no | 0 | 125.78 | 111.41 | 71.69 | 18.35 | 5.16 | 18.90 | 52.00 | R | 13.81 | 25.91 | F | glutamine-tRNA ligase activity | glutaminyl-tRNA aminoacylation | cytosol;nucleus |
| PF3D7_0320500 | Nico | nicotinamidase, putative | 50.15 | no | 0 | 14.81 | 25.95 | 22.97 | 8.21 | 4.17 | 77.65 | 68.32 | GV | 18.72 | 60.13 | F | enzyme activity | N/A | cytoplasm;nucleus |
| PF3D7_1305300 | GCN1 | translational activator GCN1, putative | 418.58 | no | 0 | 57.07 | 26.86 | 11.37 | 0.93 | 3.17 | 27.57 | 15.41 | R | 39.26 | 12.22 | M | protein binding | regulation of translation | nucleus;cytoplasm |
| PF3D7_1118200 | N/A | heat shock protein 90, putative | 108.46 | | 0 | 35.68 | 74.78 | 27.10 | 1.05 | 6.37 | 28.80 | 106.59 | O | 57.53 | 167.41 | F | ATP binding;ATP hydrolysis activity;mRNA binding;protein kinase binding;unfolded protein binding | protein folding;response to heat;response to unfolded protein | mitochondrial inner membrane;mitochondrion;nucleus |
| PF3D7_1238600 | NSM | sphingomyelin phosphodiesterase | 46.01 | no | 0 | 6.00 | 0.74 | 25.52 | 17.68 | 4.84 | 16.35 | 72.73 | O | 12.80 | 8.17 | M | nuclease activity;phospholipase activity;sphingomyelin phosphodiesterase activity | sphingomyelin metabolic process | cytoplasm;nucleus |
| PF3D7_1147300 | N/A | conserved Plasmodium protein, unknown function | 158.90 | no | 0 | 7.42 | 5.90 | 18.37 | 3.37 | 4.97 | 56.97 | 212.49 | O | 66.77 | 92.87 | F | N/A | N/A | nucleus |
| PF3D7_1119800 | ASF1 | alternative splicing factor ASF-1, putative | 23.65 | no | 0 | 25.48 | 9.96 | 10.18 | 3.59 | 17.60 | 59.49 | 10.61 | GV | 35.15 | 24.44 | M | RNA binding;mRNA binding | alternative mRNA splicing, via spliceosome | Maurer's cleft;nucleus |
| PF3D7_0318600 | N/A | cleavage and polyadenylation specificity factor, putative | 118.85 | no | 0 | 39.76 | 16.85 | 7.15 | 2.21 | 1.22 | 9.22 | 4.83 | R | 20.95 | 10.82 | M | mRNA binding | mRNA cleavage;mRNA polyadenylation;snRNA processing | mRNA cleavage and polyadenylation specificity factor complex;nucleus |
| PF3D7_0909900 | N/A | helicase SKI2W, putative | 160.37 | no | 0 | 28.73 | 7.92 | 11.19 | 1.63 | 2.02 | 9.06 | 8.98 | R | 20.33 | 10.27 | M | RNA helicase activity | RNA catabolic process;nuclear-transcribed mRNA catabolic process, 3'-5' exonucleolytic nonsense-mediated decay | Ski complex |
| PF3D7_1123200 | LRR11 | leucine-rich repeat protein | 93.33 | no | 0 | 10.22 | 0.77 | 17.35 | 7.05 | 6.67 | 13.40 | 30.64 | O | 394.72 | 15.99 | M | protein binding | SCF-dependent proteasomal ubiquitin-dependent protein catabolic process;ubiquitin-dependent protein catabolic process | SCF ubiquitin ligase complex;cytoplasm;nucleus |
| PF3D7_1238800 | ACS11 | acyl-CoA synthetase | 92.09 | no | 0 | 60.81 | 21.66 | 22.81 | 5.05 | 1.21 | 19.16 | 60.66 | R | 17.87 | 222.03 | F | long-chain fatty acid-CoA ligase activity | fatty acid metabolic process;response to xenobiotic stimulus | endoplasmic reticulum;membrane;nucleus |
| PF3D7_0705700 | N/A | 40S ribosomal protein S29, putative | 6.42 | no | 0 | 307.36 | 185.31 | 107.65 | 20.54 | 100.80 | 118.34 | 80.20 | R | 185.40 | 134.98 | M | structural constituent of ribosome;zinc ion binding | translation | cytosolic small ribosomal subunit;nucleus |
| PF3D7_0812100 | N/A | proteasome activator complex subunit 4, putative | 314.49 | no | 2 | 19.79 | 8.86 | 16.15 | 3.48 | 2.39 | 18.84 | 18.55 | R | 21.74 | 7.75 | M | lysine-acetylated histone binding;peptidase activator activity;proteasome binding | proteasomal ubiquitin-independent protein catabolic process | cytosol;nucleus |
| PF3D7_0416400 | HAT1 | histone acetyltransferase, putative | 146.72 | no | 0 | 23.91 | 32.37 | 90.95 | 27.34 | 6.12 | 10.98 | 652.73 | O | 6.29 | 71.97 | F | H4 histone acetyltransferase activity | histone H4 acetylation | nucleus |
| PF3D7_0920800 | IMPDH | inosine-5'-monophosphate dehydrogenase | 56.15 | no | 0 | 234.36 | 64.70 | 21.60 | 4.09 | 3.00 | 8.36 | 21.86 | R | 21.88 | 18.36 | O | enzyme activity | N/A | food vacuole;nucleus |
| PF3D7_1146000 | RSA4 | ribosome assembly protein 4, putative | 73.13 | no | 0 | 47.40 | 21.69 | 12.96 | 4.85 | 6.60 | 12.82 | 98.11 | O | 7.16 | 27.04 | F | protein binding | ribosomal large subunit assembly | nucleolus;nucleus |
| PF3D7_0920900 | PRP24 | U4/U6 snRNA-associated-splicing factor, putative | 56.65 | no | 0 | 39.86 | 7.84 | 12.14 | 1.72 | 2.14 | 24.75 | 6.31 | R | 38.25 | 13.46 | M | RNA binding | mRNA splicing | ribonucleoprotein complex |
| PF3D7_0209200 | MTR3 | exosome complex component MTR3, putative | 31.25 | no | 0 | 35.68 | 10.80 | 8.20 | 0.85 | 7.36 | 13.71 | 177.10 | O | 8.70 | 13.51 | F | protein binding | U4 snRNA 3'-end processing;nuclear mRNA surveillance;nuclear-transcribed mRNA catabolic process, exonucleolytic, 3'-5';polyadenylation-dependent snoRNA 3'-end processing;rRNA catabolic process | cytoplasmic exosome (RNase complex);exosome (RNase complex);nuclear exosome (RNase complex);nucleolus;osmiophilic body |
| PF3D7_1368400 | N/A | ribosomal protein L1, putative | 38.26 | no | 0 | 24.67 | 8.11 | 13.08 | 2.93 | 7.44 | 84.45 | 40.56 | GV | 17.78 | 101.47 | F | RNA binding;structural constituent of ribosome | translation | cytosol;cytosolic small ribosomal subunit;nucleus |
| PF3D7_0505100 | TRS85 | trafficking protein particle complex subunit 8, putative | 328.67 | no | 0 | 22.85 | 17.38 | 18.45 | 7.85 | 2.48 | 12.13 | 169.71 | O | 26.61 | 25.56 | O | protein binding | autophagosome assembly;autophagy of peroxisome;endoplasmic reticulum to Golgi vesicle-mediated transport;protein localization to phagophore assembly site | TRAPPIII protein complex;cytoplasmic vesicle;nucleus;phagophore assembly site |
| PF3D7_0404300 | N/A | Ran-binding protein, putative | 74.18 | no | 0 | 160.64 | 113.12 | 83.13 | 17.64 | 22.05 | 11.10 | 205.79 | O | 769.74 | 277.07 | M | protein binding | intracellular transport | nucleus |
| PF3D7_0520300 | LSM2 | U6 snRNA-associated Sm-like protein LSm2, putative | 12.06 | no | 0 | 86.80 | 33.23 | 16.22 | 2.09 | 28.04 | 46.39 | 38.90 | R | 40.96 | 20.43 | M | RNA binding | mRNA splicing, via spliceosome | Lsm1-7-Pat1 complex;P-body;U4/U6 x U5 tri-snRNP complex;U6 snRNP;catalytic step 2 spliceosome;intracellular;nucleus;precatalytic spliceosome;sno(s)RNA-containing ribonucleoprotein complex |
| PF3D7_0602600 | N/A | SAC3 domain-containing protein, putative | 153.72 | no | 0 | 63.37 | 11.71 | 8.11 | 1.23 | 2.24 | 17.14 | 11.54 | R | 34.02 | 12.93 | M | RNA binding | mRNA export from nucleus | cytoplasm;nucleus;transcription export complex 2 |
| PF3D7_0827000 | DBP10 | ATP-dependent RNA helicase DBP10, putative | 151.10 | no | 0 | 70.36 | 41.06 | 15.42 | 1.18 | 0.91 | 8.26 | 71.34 | O | 164.04 | 65.86 | M | ATP binding;RNA binding | assembly of large subunit precursor of preribosome | nucleolus |
| PF3D7_0520700 | N/A | CDC73 domain-containing protein, putative | 118.00 | no | 0 | 17.35 | 11.54 | 4.41 | 0.54 | 1.68 | 16.96 | 2.06 | R | 4.91 | 7.37 | F | RNA polymerase II complex binding | positive regulation of transcription elongation from RNA polymerase II promoter;recruitment of 3'-end processing factors to RNA polymerase II holoenzyme complex | Cdc73/Paf1 complex |
| PF3D7_1428800 | N/A | Transcription initiation TFIID-like, putative | 41.98 | no | 1 | 42583.00 | 43556.00 | 30072.00 | 35431.00 | 15281.00 | 107.79 | 45319.00 | O | 24.15 | 67.6 | F | RNA polymerase II general transcription initiation factor activity | DNA-templated transcription, initiation | nucleus |
| PF3D7_1419000 | N/A | conserved Plasmodium protein, unknown function | 136.94 | no | 0 | 10.25 | 3.42 | 24.40 | 5.62 | 5.15 | 25.57 | 77.83 | O | 362.77 | 113.47 | M | N/A | N/A | nucleus |
| PF3D7_0903500 | NUP138 | nucleoporin NUP138, putative | 155.15 | no | 0 | 72.56 | 28.49 | 20.67 | 5.55 | 1.90 | 6.39 | 33.44 | R | 61.61 | 18.41 | M | protein binding | nucleocytoplasmic transport | nuclear pore;nucleus |
| PF3D7_1342800 | PEPCK | phosphoenolpyruvate carboxykinase | 66.21 | no | 0 | 48.08 | 81.13 | 25.73 | 1.75 | 3.23 | 84.88 | 206.04 | O | 16.63 | 437.73 | F | phosphoenolpyruvate carboxykinase (ATP) activity | gluconeogenesis | cytosol;mitochondrion;nucleus |
| PF3D7_1464000 | N/A | YL1 nuclear protein, putative | 84.61 | no | 0 | 75.86 | 39.27 | 26.34 | 8.06 | 3.41 | 17.98 | 15.92 | R | 50.61 | 71.71 | F | protein binding | histone exchange | nucleus |
| PF3D7_0924700 | SF3A3 | splicing factor 3A subunit 3, putative | 71.38 | no | 0 | 35.49 | 11.86 | 12.14 | 1.21 | 9.08 | 142.54 | 246.09 | O | 46.13 | 27.27 | M | RNA binding | mRNA splicing, via spliceosome | spliceosomal complex |
| PF3D7_1463400 | N/A | DNA-directed RNA polymerase III subunit RPC4, putative | 39.12 | no | 0 | 25.05 | 7.33 | 3.86 | 0.16 | 4.16 | 19.51 | 5.41 | R | 34.50 | 7.62 | M | DNA binding;RNA binding | regulation of transcription | RNA polymerase III complex |
| PF3D7_0828500 | N/A | translation initiation factor eIF-2B subunit alpha, putative | 40.03 | no | 0 | 3.41 | 0.43 | 22.89 | 6.28 | 10.97 | 61.12 | 37.60 | GV | 19.20 | 55.65 | F | translation initiation factor activity | regulation of translational initiation;translational initiation | eukaryotic translation initiation factor 2B complex |
| PF3D7_0407900 | N/A | AAA family ATPase, putative | 77.81 | no | 0 | 10.99 | 4.06 | 9.67 | 1.95 | 1.34 | 9.13 | 51.57 | O | 4.51 | 49.00 | F | ATP hydrolysis activity;microtubule-severing ATPase activity | cytoplasmic microtubule organization;regulation of double-strand break repair via homologous recombination | nucleus |
| PF3D7_1459000 | DBP5 | ATP-dependent RNA helicase DBP5 | 84.31 | no | 0 | 219.61 | 30.01 | 45.93 | 5.12 | 8.44 | 27.29 | 267.93 | O | 73.85 | 66.19 | M | 3'-5' DNA/RNA helicase activity;5'-3' DNA/RNA helicase activity;ATP hydrolysis activity;RNA binding;RNA helicase activity;single-stranded DNA helicase activity | DNA duplex unwinding;poly(A)+ mRNA export from nucleus | cytoplasm;cytoplasmic stress granule;nucleus |
| PF3D7_1202600 | NEMF | nuclear export mediator factor NEMF, putative | 254.29 | no | 0 | 217.88 | 133.11 | 94.30 | 76.86 | 14.44 | 44.10 | 660.64 | O | 234.67 | 357.87 | F | ribosomal large subunit binding;tRNA binding | nuclear export;rescue of stalled ribosome;ribosome-associated ubiquitin-dependent protein catabolic process | RQC complex;nucleus |
| PF3D7_1103700 | CK2beta1 | casein kinase II beta chain | 28.37 | no | 0 | 60.92 | 20.73 | 30.85 | 3.93 | 24.82 | 34.67 | 32.46 | R | 12.28 | 21.31 | F | protein kinase regulator activity;protein serine/threonine kinase activity | protein phosphorylation | cytoplasm;nucleus;protein kinase CK2 complex |
| PF3D7_0205400 | N/A | PCI domain-containing protein, putative | 54.35 | no | 0 | 49.55 | 8.48 | 7.44 | 0.71 | 7.44 | 35.13 | 20.83 | R | 42.01 | 27.67 | M | RNA binding;double-stranded DNA binding | nuclear retention of pre-mRNA at the site of transcription;poly(A)+ mRNA export from nucleus;posttranscriptional tethering of RNA polymerase II gene DNA at nuclear periphery;transcription elongation from RNA polymerase II promoter | transcription export complex 2 |
| PF3D7_0531400 | N/A | G-patch domain-containing protein, putative | 142.14 | no | 0 | 14.80 | 5.01 | 2.84 | 0.47 | 1.48 | 17.14 | 3.31 | GV | 19.23 | 5.64 | M | protein binding | spliceosomal complex disassembly | U2-type post-mRNA release spliceosomal complex |
| PF3D7_1414000 | RPN13 | 26S proteasome regulatory subunit RPN13, putative | 29.49 | no | 0 | 11.34 | 4.56 | 40.36 | 7.59 | 15.47 | 60.24 | 104.54 | O | 133.30 | 141.30 | F | endopeptidase activator activity;proteasome binding;ubiquitin binding | ubiquitin-dependent protein catabolic process | proteasome regulatory particle, lid subcomplex |
| PF3D7_0728000 | eIF2alpha | eukaryotic translation initiation factor 2 subunit alpha | 37.82 | no | 0 | 334.37 | 102.99 | 141.94 | 30.15 | 9.70 | 21.15 | 96.56 | R | 162.16 | 71.68 | M | RNA binding;ribosome binding;translation initiation factor activity | translational initiation | eukaryotic 48S preinitiation complex;eukaryotic translation initiation factor 2 complex;eukaryotic translation initiation factor 2B complex;multi-eIF complex;nucleus |
| PF3D7_0528700 | CYP23 | peptidyl-prolyl cis-trans isomerase | 23.20 | no | 0 | 78.15 | 32.21 | 42.23 | 5.18 | 205.66 | 236.21 | 104.00 | GV | 75.00 | 93.27 | F | cyclosporin A binding;peptidyl-prolyl cis-trans isomerase activity | protein folding;protein peptidyl-prolyl isomerization | nucleus |
| PF3D7_1475000 | MED31 | mediator of RNA polymerase II transcription subunit 31 | 16.61 | no | 0 | 7.30 | 4.62 | 5.12 | 0.39 | 1.42 | 20.92 | 0.00 | GV | 12.76 | 22.07 | F | protein binding | regulation of transcription by RNA polymerase II | core mediator complex;mediator complex;nucleus |
| PF3D7_1104000 | N/A | phenylalanine--tRNA ligase beta subunit | 72.50 | no | 0 | 50.92 | 21.22 | 13.09 | 2.05 | 2.02 | 11.11 | 12.40 | R | 2.36 | 5.74 | O | phenylalanine-tRNA ligase activity | phenylalanyl-tRNA aminoacylation | intracellular;nucleus;phenylalanine-tRNA ligase complex |
| PF3D7_1442300 | tRIP | tRNA import protein tRIP | 46.44 | no | 0 | 111.43 | 41.12 | 30.21 | 3.30 | 3.69 | 20.39 | 42.18 | R | 12.43 | 32.74 | F | RNA binding;aminoacyl-tRNA ligase activity | tRNA aminoacylation for protein translation | cell surface;nucleus;plasma membrane |
| PF3D7_1442100 | RPA3 | replication factor A protein 3, putative | 12.61 | no | 0 | 5.70 | 6.24 | 16.13 | 1.98 | 23.62 | 51.00 | 45.38 | GV | 165.38 | 24.30 | M | damaged DNA binding;single-stranded DNA binding | DNA replication;base-excision repair;double-strand break repair via homologous recombination;mismatch repair;nucleotide-excision repair | DNA replication factor A complex;site of double-strand break |
| PF3D7_1357700 | UTP21 | U3 small nucleolar RNA-associated protein 21, putative | 143.36 | no | 0 | 87.49 | 25.91 | 5.22 | 0.22 | 1.44 | 17.50 | 19.58 | R | 22.04 | 20.00 | O | RNA binding | mRNA splicing, via spliceosome | nucleolus;nucleus |
| PF3D7_1126400 | MED21 | mediator of RNA polymerase II transcription subunit 21 | 21.72 | no | 0 | 36.61 | 8.48 | 3.95 | 0.45 | 16.07 | 52.03 | 144.76 | O | 43.52 | 55.74 | F | protein binding | regulation of transcription by RNA polymerase II | core mediator complex;mediator complex;nucleus |
| PF3D7_0812700 | SNRPC | U1 small nuclear ribonucleoprotein C, putative | 27.88 | no | 0 | 64.39 | 16.92 | 22.12 | 3.78 | 24.51 | 44.96 | 35.79 | R | 23.94 | 9.56 | M | RNA binding;pre-mRNA 5'-splice site binding | mRNA 5'-splice site recognition;mRNA splicing, via spliceosome | U1 snRNP |
| PF3D7_1032700 | N/A | conserved Plasmodium protein, unknown function | 65.54 | no | 0 | 18.42 | 9.37 | 11.47 | 2.46 | 4.30 | 31.28 | 76.93 | 76.93 | 186.51 | 49.94 | M | | | cytoplasm;nucleus;transcription regulator complex;vesicle |
| PF3D7_0319100 | RBX1 | E3 ubiquitin-protein ligase RBX1, putative | 12.42 | no | 0 | 20.46 | 16.05 | 27.96 | 6.92 | 72.02 | 202.05 | 119.05 | GV | 27.69 | 183.26 | F | cullin family protein binding;ubiquitin protein ligase activity;ubiquitin-protein transferase activity | protein ubiquitination;ubiquitin-dependent protein catabolic process | SCF ubiquitin ligase complex;cullin-RING ubiquitin ligase complex;nucleus;ubiquitin ligase complex |
| PF3D7_0616200 | NDC80 | kinetochore protein NDC80 | 69.87 | no | 0 | 8.34 | 9.60 | 36.73 | 18.91 | 25.49 | 30.36 | 438.89 | O | 310.06 | 68.54 | M | microtubule binding | attachment of mitotic spindle microtubules to kinetochore;chromosome segregation | Ndc80 complex;chromosome, centromeric region;condensed nuclear chromosome kinetochore;microtubule;nucleus |
| PF3D7_1469700 | MED6 | mediator of RNA polymerase II transcription subunit 6 | 24.57 | no | 0 | 3.19 | 2.42 | 7.14 | 1.20 | 17.37 | 183.73 | 34.02 | GV | 38.06 | 63.23 | F | protein binding | regulation of transcription by RNA polymerase II | core mediator complex;mediator complex;nucleus |
| PF3D7_0309200 | ARK2 | aurora-related kinase ARK2, putative | 193.78 | no | 0 | 10.07 | 3.95 | 18.87 | 3.88 | 7.55 | 53.61 | 792.94 | O | 723.17 | 142.91 | M | histone serine kinase activity;protein serine/threonine kinase activity | mitotic spindle organization;protein phosphorylation;regulation of cytokinesis | chromosome passenger complex;kinetochore;mitotic spindle;spindle microtubule;spindle midzone;spindle pole centrosome |
| PF3D7_0627700 | N/A | transportin | 132.84 | no | 0 | 66.34 | 15.77 | 34.44 | 6.78 | 4.25 | 16.27 | 23.98 | R | 37.44 | 16.26 | M | nuclear import signal receptor activity;nuclear localization sequence binding;protein transmembrane transporter activity | protein import into nucleus | apicoplast;cytoplasm;nuclear pore;nucleus |
| PF3D7_1033100 | AdoMetDC/ODC | S-adenosylmethionine decarboxylase/ornithine decarboxylase | 168.17 | no | 0 | 577.00 | 307.54 | 187.19 | 58.86 | 11.95 | 51.45 | 323.43 | R | 110.74 | 299.82 | F | adenosylmethionine decarboxylase activity;ornithine decarboxylase activity | polyamine biosynthetic process;putrescine biosynthetic process from ornithine;putrescine metabolic process | cytoplasm;nucleus |
| PF3D7_0622900 | AP2Tel | AP2 domain transcription factor AP2Tel | 237.36 | no | 0 | 69.50 | 12.07 | 8.39 | 1.68 | 2.51 | 18.98 | 114.12 | O | 34.73 | 15.43 | M | DNA binding;telomeric DNA binding | regulation of transcription, DNA-templated | chromosome, telomeric region;nuclear periphery |
| PF3D7_1309300 | PRPF3 | U4/U6 small nuclear ribonucleoprotein PRP3, putative | 86.49 | no | 0 | 76.30 | 34.22 | 28.94 | 6.41 | 2.74 | 25.25 | 101.22 | O | 156.58 | 97.81 | M | RNA binding | mRNA splicing, via spliceosome | U4/U6 x U5 tri-snRNP complex |
| PF3D7_1211200 | N/A | conserved Plasmodium protein, unknown function | 219.65 | no | 0 | 23.55 | 17.19 | 7.79 | 0.62 | 2.12 | 21.16 | 51.70 | O | 59.94 | 26.79 | M | N/A | N/A | cytoplasm;nucleus |

| | | | | | | | | | | | | | | | | | | | |
|---|---|---|---|---|---|---|---|---|---|---|---|---|---|---|---|---|---|---|---|
| PF3D7_1355300 | SET6 | histone-lysine N-methyltransferase, putative | 60.02 | no | 0 | 25.03 | 3.61 | 3.00 | 0.25 | 1.97 | 10.37 | 7.26 | R | 4.63 | 12.20 | F | histone-lysine N-methyltransferase activity | histone lysine methylation | nuclear periphery;nucleus |
| PF3D7_1418000 | UFD1 | ubiquitin fusion degradation protein 1, putative | 32.48 | no | 0 | 42.97 | 19.86 | 50.79 | 20.87 | 37.54 | 38.73 | 38.06 | T | 329.58 | 134.98 | M | polyubiquitin modification-dependent protein binding;ubiquitin binding | ER-associated misfolded protein catabolic process;ubiquitin-dependent ERAD pathway;ubiquitin-dependent protein catabolic process | VCP-NPL4-UFD1 AAA ATPase complex;intracellular;nucleus |
| PF3D7_1361000 | PRMT5 | protein arginine N-methyltransferase 5, putative | 85.55 | no | 0 | 38.47 | 10.13 | 4.47 | 1.83 | 1.59 | 5.94 | 1.40 | R | 14.28 | 3.21 | M | histone-arginine N-methyltransferase activity;protein binding;protein-arginine N-methyltransferase activity | histone arginine methylation;regulation of transcription, DNA-templated | cytosol;nucleus |
| PF3D7_0605300 | ARK1 | aurora-related kinase ARK1 | 40.91 | no | 0 | 8.72 | 0.83 | 38.46 | 9.27 | 57.44 | 196.97 | 135.55 | GV | 395.58 | 288.46 | M | kinase activity | schizogony | nucleus;spindle pole body |
| PF3D7_1235300 | NOT4 | CCR4-NOT transcription complex subunit 4, putative | 194.67 | no | 0 | 201.91 | 27.26 | 19.57 | 6.46 | 7.38 | 38.25 | 234.99 | O | 1104.27 | 245.68 | M | mRNA binding;ubiquitin-protein transferase activity | protein ubiquitination | CCR4-NOT complex;nucleus |
| PF3D7_1126300 | N/A | DnaJ protein, putative | 73.53 | no | 0 | 9.93 | 18.34 | 21.58 | 4.86 | 5.74 | 62.87 | 355.02 | O | 22.75 | 35.36 | F | RNA binding;unfolded protein binding | chaperone cofactor-dependent protein refolding;protein refolding | cytoplasm;nucleus |
| PF3D7_1210400 | N/A | general transcription factor 3C polypeptide 5, putative | 107.05 | no | 0 | 26.23 | 6.67 | 7.28 | 0.65 | 1.80 | 8.29 | 59.71 | O | 5.26 | 15.47 | F | DNA binding | regulation of transcription | transcription factor TFIIIC complex;nucleus |
| PF3D7_1226800 | ATX3 | ataxin-3, putative | 44.98 | no | 0 | 7.87 | 4.23 | 18.27 | 7.63 | 8.88 | 51.13 | 134.24 | O | 125.83 | 121.07 | M | thiol-dependent deubiquitinase | protein deubiquitination | nucleus |
| PF3D7_0520400 | N/A | single-stranded DNA-binding protein, putative | 35.66 | no | 0 | 3.87 | 4.12 | 10.75 | 2.23 | 2.66 | 14.66 | 3.79 | GV | 2.59 | 15.50 | F | DNA binding | DNA replication | intracellular;nucleus |
| PF3D7_1403900 | CPPED1 | serine/threonine protein phosphatase CPPED1, putative | 35.22 | no | 0 | 12.11 | 26.30 | 10.55 | 1.40 | 21.28 | 101.91 | 71.88 | GV | 9.30 | 35.26 | F | enzyme activity | phosphorylation process | nucleus |
| PF3D7_1209200 | LSM7 | U6 snRNA-associated Sm-like protein LSm7, putative | 12.00 | no | 0 | 68.19 | 29.48 | 26.74 | 6.28 | 106.14 | 38.69 | 0.00 | G II | 22.19 | 19.11 | 0 | RNA binding | RNA splicing, via transesterification reactions;mRNA splicing, via spliceosome | Lsm1-7-Pat1 complex;U12-type spliceosomal complex;U2-type prespliceosome;U6 snRNP;catalytic step 2 spliceosome;nucleus;ribonucleoprotein complex;sno(s)RNA-containing ribonucleoprotein complex;spliceosomal complex;spliceosomal tri-snRNP complex |
| PF3D7_1301700 | CBP2 | CX3CL1-binding protein 2 | 28.50 | no | 3 | 451.57 | 95.57 | 8.58 | 0.41 | 2.47 | 1.93 | 21.28 | 451.57 | 0.36 | 0.24 | 0 | ATP binding;chemokine binding;nucleic acid binding;protein binding | cytoadherence to microvasculature, mediated by symbiont protein | Maurer's cleft;host cell surface;nucleus |
| PF3D7_1329600 | N/A | conserved Plasmodium protein, unknown function | 270.47 | no | 0 | 102.67 | 19.31 | 17.38 | 1.42 | 21.06 | 97.10 | 52.49 | R | 86.40 | 22.93 | M | N/A | N/A | N/A |
| PF3D7_1007900 | EIF3D | eukaryotic translation initiation factor 3 subunit D, putative | 244.67 | no | 0 | 149.30 | 37.91 | 38.38 | 3.47 | 4.60 | 16.20 | 49.13 | R | 27.12 | 15.79 | M | translation initiation factor activity | translational initiation | eukaryotic translation initiation factor 3 complex;nucleus |
| PF3D7_0609900 | N/A | tetratricopeptide repeat protein, putative | 50.76 | no | 0 | 34.32 | 5.61 | 11.49 | 12.07 | 4.48 | 12.04 | 8.41 | R | 30.53 | 11.04 | M | DNA binding | transcription by RNA polymerase III | transcription factor TFIIIC complex |
| PF3D7_0619900 | SF3A2 | splicing factor 3A subunit 2, putative | 33.07 | no | 0 | 13.49 | 5.61 | 9.37 | 0.89 | 5.28 | 32.01 | 85.16 | O | 15.85 | 24.45 | F | RNA binding | RNA splicing;spliceosomal complex assembly | U2 snRNP;U2-type prespliceosome;catalytic step 2 spliceosome;spliceosomal complex |
| PF3D7_1106700 | DNA2 | DNA replication ATP-dependent helicase/nuclease DNA2, putative | 25.38 | no | 0 | 25.64 | 9.15 | 10.23 | 1.40 | 2.10 | 9.03 | 58.53 | O | 88.77 | 19.55 | M | nuclease activity | DNA-dependent DNA replication | nucleus |
| PF3D7_1020700 | N/A | N-acetyltransferase, GNAT family, putative | 53.67 | no | 0 | 38.87 | 20.19 | 4.93 | 0.37 | 1.27 | 14.24 | 17.16 | R | 14.99 | 21.05 | F | rRNA cytidine N-acetyltransferase activity;tRNA binding | rRNA acetylation involved in maturation of SSU-rRNA | nucleolus;nucleus |
| PF3D7_1435600 | N/A | conserved Plasmodium protein, unknown function | 175.57 | no | 0 | 7.96 | 0.59 | 11.62 | 11.95 | 3.42 | 140.60 | 545.53 | O | 204.39 | 597.76 | F | N/A | N/A | nucleus |
| PF3D7_0719000 | N/A | translation machinery-associated protein, putative | 168.93 | no | 0 | 4.30 | 1.88 | 2.28 | 0.00 | 15.78 | 51.42 | 14.04 | GV | 91.69 | 17.03 | M | protein-macromolecule adaptor activity | proteasome regulatory particle assembly | cytoplasm;nucleus |
| PF3D7_1004500 | AKIT2 | apicomplexan kinetochore protein 2, putative | 111.44 | no | 0 | 3.09 | 0.75 | 5.41 | 2.69 | 4.67 | 17.29 | 20.89 | O | 23.69 | 12.24 | M | DNA binding | DNA-dependent DNA replication | kinetochore;nucleus |
| PF3D7_1454200 | N/A | conserved Plasmodium protein, unknown function | 51.93 | no | 2 | 24.51 | 16.77 | 3.84 | 0.26 | 2.64 | 19.59 | 17.07 | R | 5.31 | 4.30 | 0 | N/A | transcription, DNA-templated | nucleus |
| PF3D7_0826300 | N/A | SPRY domain, putative | 47.58 | no | 0 | 3.77 | 1.93 | 14.07 | 2.25 | 4.19 | 35.85 | 42.08 | O | 43.14 | 27.60 | M | transcription cis-regulatory region binding | histone H3-K4 methylation | Set1C/COMPASS complex;nucleus |
| PF3D7_0623600 | N/A | splicing factor-like protein 1, putative | 220.40 | no | 0 | 120.96 | 42.53 | 58.06 | 21.17 | 20.03 | 15.51 | 113.26 | R | 53.65 | 34.30 | M | RNA binding;mRNA binding | RNA splicing;RNA splicing, via transesterification reactions;mRNA splicing, via spliceosome;regulation of alternative mRNA splicing, via spliceosome | nucleus |
| PF3D7_1121900 | PDK1 | 3-phosphoinositide-dependent protein kinase 1 | 120.38 | no | 0 | 2.72 | 0.23 | 12.55 | 3.49 | 0.47 | 9.70 | 6.95 | T | 6.10 | 16.40 | F | protein kinase activity;protein serine/threonine kinase activity | activation of protein kinase activity;intracellular signal transduction;peptidyl-serine phosphorylation;protein phosphorylation;response to xenobiotic stimulus | cytosol;nucleus |
| PF3D7_1450400 | N/A | E3 ubiquitin-protein ligase ZNF598, putative | 165.75 | no | 0 | 45.10 | 15.26 | 12.46 | 4.72 | 1.95 | 17.93 | 47.98 | O | 28.25 | 19.20 | M | ligase activity | protein ubiquitination complex | nucleus |
| PF3D7_1344300 | N/A | zinc finger protein, putative | 140.26 | no | 0 | 5.22 | 5.33 | 22.17 | 14.25 | 4.14 | 32.99 | 73.95 | O | 42.20 | 28.75 | M | protein binding | N/A | cytoplasm;nucleus |
| PF3D7_1336800 | NUDC | nuclear movement protein, putative | 282.22 | no | 0 | 51.63 | 28.53 | 12.42 | 0.59 | 3.89 | 19.55 | 40.44 | 51.63 | 86.84 | 45.79 | M | unfolded protein binding | developmental process;protein folding | cytoplasm;nucleus |
| PF3D7_0502100 | N/A | HCNGP-like protein | 36.61 | no | 0 | 3.17 | 0.99 | 2.83 | 1.16 | 3.76 | 17.93 | 16.63 | GV | 10.13 | 24.73 | F | N/A | N/A | nucleus |
| PF3D7_0408500 | FEN1 | flap endonuclease 1 | 63.85 | no | 0 | 9.73 | 1.55 | 51.59 | 26.82 | 10.07 | 9.97 | 121.16 | O | 1072.17 | 59.74 | M | 5'-3' exonuclease activity;5'-flap endonuclease activity;endodeoxyribonuclease activity;flap endonuclease activity | | nucleus |
| PF3D7_1008000 | HDA2 | histone deacetylase 2 | 198.28 | no | 0 | 70.23 | 35.07 | 26.53 | 6.58 | 2.90 | 21.36 | 45.15 | R | 150.69 | 35.45 | M | deacetylase activity | inositol phosphate biosynthetic process;regulation of gene expression;chromatin remodeling | nucleus |
| PF3D7_0319500 | N/A | RNA-binding protein, putative | 132.61 | no | 0 | 163.64 | 56.62 | 58.19 | 6.41 | 18.29 | 29.71 | 59.94 | R | 137.53 | 117.33 | M | RNA binding | mRNA splicing, via spliceosome | U2 snRNP;precatalytic spliceosome |
| PF3D7_0420600 | AKIT6 | apicomplexan kinetochore protein 6, putative | 419.08 | no | 0 | 5.59 | 2.69 | 19.00 | 10.88 | 6.52 | 62.42 | 0.00 | GV | 210.81 | 82.79 | M | DNA binding | DNA replication | kinetochore;nucleus |
| PF3D7_1211300 | MCM8 | DNA helicase MCM8, putative | 98.54 | no | 0 | 5.73 | 0.50 | 6.88 | 2.18 | 2.52 | 12.78 | 13.22 | O | 32.44 | 36.01 | 0 | ATP binding;ATP-dependent activity, acting on DNA;DNA helicase activity;DNA replication origin binding;single-stranded DNA binding | DNA replication initiation;DNA strand elongation involved in DNA replication | MCM complex;nuclear pre-replicative complex;nucleus |
| PF3D7_1325400 | N/A | CRWN-like protein, putative | 32.78 | no | 0 | 5.49 | 0.34 | 12.48 | 7.52 | 3.74 | 50.96 | 79.13 | O | 80.14 | 103.24 | F | chromatin binding | N/A | heterochromatin;nucleus |
| PF3D7_1123900 | SNU13 | 13 kDa ribonucleoprotein-associated protein, putative | 48.06 | no | 0 | 495.51 | 199.33 | 81.42 | 8.88 | 257.61 | 82.42 | 104.38 | O | 99.29 | 64.75 | M | RNA binding;U3 snoRNA binding;U4 snRNA binding | mRNA splicing, via spliceosome;maturation of LSU-rRNA;maturation of SSU-rRNA | U4/U6 x U5 tri-snRNP complex;box C/D RNP complex;nucleolus;nucleus;precatalytic spliceosome;small-subunit processome |
| PF3D7_1028500 | N/A | partial CSTF domain-containing protein, putative | 76.83 | no | 0 | 56.23 | 18.95 | 12.54 | 3.07 | 2.95 | 53.76 | 186.77 | O | 64.36 | 104.16 | F | N/A | N/A | nucleus |
| PF3D7_0906600 | N/A | zinc finger protein, putative | 14.60 | no | 0 | 383.60 | 154.92 | 24.61 | 2.13 | 4.11 | 32.47 | 160.02 | R | 28.60 | 10.60 | M | mRNA binding | N/A | nucleus |
| PF3D7_0212300 | ERF1 | eukaryotic peptide chain release factor subunit 1, putative | 73.03 | no | 0 | 165.17 | 109.79 | 74.15 | 7.41 | 4.51 | 10.01 | 94.37 | R | 63.49 | 364.13 | F | sequence-specific mRNA binding;translation release factor activity;translation release factor activity, codon specific | cytoplasmic translational termination;regulation of translational termination | cytosol;nucleus;translation release factor complex |
| PF3D7_1438000 | N/A | eukaryotic translation initiation factor eIF2A, putative | 11.00 | no | 0 | 122.85 | 24.59 | 19.10 | 2.61 | 4.51 | 32.68 | 44.27 | R | 57.73 | 44.99 | M | mRNA binding;ribosome binding;tRNA binding;translation initiation factor activity | | cytosolic small ribosomal subunit;nucleus |
| PF3D7_0528000 | UMP1 | proteasome maturation factor UMP1, putative | 41.93 | no | 0 | 18.46 | 9.13 | 12.79 | 2.87 | 19.73 | 142.73 | 50.36 | GV | 38.48 | 175.22 | F | protein binding | proteasome assembly | cytoplasm;nucleus;proteasome complex |
| PF3D7_0111800 | N/A | eukaryotic translation initiation factor 4E, putative | 51.00 | no | 0 | 11.27 | 13.05 | 11.53 | 1.16 | 2.28 | 14.91 | 40.76 | O | 57.32 | 49.14 | M | RNA 7-methylguanosine cap binding;translation initiation factor activity | | eukaryotic translation initiation factor 4F complex;nucleus |
| PF3D7_1415200 | N/A | DNA-directed RNA polymerases I and III subunit RPAC2 | 202.14 | no | 0 | 66.20 | 17.72 | 4.32 | 0.67 | 34.34 | 29.47 | 0.00 | R | 23.61 | 21.00 | 0 | DNA-directed 5'-3' RNA polymerase activity | transcription, DNA-templated | RNA polymerase I complex;RNA polymerase III complex |
| PF3D7_1366900 | N/A | conserved protein, unknown function | 76.56 | no | 0 | 367.91 | 87.94 | 97.50 | 19.49 | 14.75 | 48.64 | 158.75 | R | 207.94 | 46.16 | M | mRNA binding | N/A | cytoplasm;nucleus |
| PF3D7_0308100 | N/A | zinc finger protein, putative | 100.17 | no | 0 | 6.16 | 0.29 | 11.98 | 10.29 | 1.32 | 5.98 | 147.37 | O | 6.05 | 20.83 | F | protein binding | N/A | cytoplasm;membrane;nucleus |
| PF3D7_0716800 | EIF3I | eukaryotic translation initiation factor 3 subunit I, putative | 91.14 | no | 0 | 355.70 | 209.51 | 132.52 | 11.21 | 25.76 | 86.80 | 304.59 | R | 113.70 | 111.58 | 0 | RNA binding;translation initiation factor activity | cytoplasmic translational initiation;regulation of translational initiation;translational initiation | eukaryotic translation initiation factor 3 complex;eukaryotic translation initiation factor 3 complex, eIF3m;nucleus |
| PF3D7_1207100 | N/A | pre-rRNA-processing protein ESF1, putative | 134.57 | no | 0 | 62.49 | 31.83 | 15.73 | 1.73 | 4.28 | 24.31 | 157.61 | O | 307.93 | 85.73 | M | RNA binding | rRNA processing | nucleus |
| PF3D7_0623100 | NAB2 | nuclear polyadenylated RNA-binding protein NAB2, putative | 178.07 | no | 0 | 561.35 | 175.35 | 96.22 | 18.52 | 22.36 | 31.62 | 252.45 | R | 247.02 | 118.44 | M | RNA binding;poly(A) binding | negative regulation of mRNA polyadenylation;regulation of mRNA stability | cytoplasm;nuclear periphery;nucleus |
| PF3D7_1364300 | PRP16 | pre-mRNA-splicing factor RNA helicase PRP16 | 124.94 | no | 0 | 81.65 | 22.90 | 9.05 | 2.51 | 1.69 | 18.83 | 17.96 | R | 26.93 | 12.71 | M | RNA helicase activity | regulation of transcription | nucleus |
| PF3D7_1420000 | SF3B4 | splicing factor 3B subunit 4, putative | 75.16 | no | 0 | 271.19 | 120.67 | 109.97 | 11.56 | 23.14 | 12.97 | 120.37 | R | 129.99 | 133.99 | F | RNA binding | RNA splicing;RNA splicing, via transesterification reactions;mRNA splicing, via spliceosome | U2 snRNP;ribonucleoprotein complex;spliceosomal complex |
| PF3D7_1228300 | NEK1 | NIMA related kinase 1 | 31.25 | no | 0 | 6.99 | 2.82 | 49.89 | 20.53 | 4.58 | 12.14 | 475.52 | O | 355.09 | 42.46 | M | kinase activity | mitotic cell cycle;protein phosphorylation;response to xenobiotic stimulus | nucleus |
| PF3D7_1314900 | P44 | general transcription factor IIH subunit 2 | 190.94 | no | 0 | 60.35 | 18.63 | 22.33 | 2.70 | 2.10 | 6.00 | 3.44 | R | 137.43 | 10.72 | M | protein binding;DNA binding | nucleotide-excision repair;regulation of transcription by RNA polymerase II | cytoplasm;nucleus;transcription factor TFIIH holo complex |
| PF3D7_0625600 | PAP | poly(A) polymerase PAP, putative | 157.31 | no | 0 | 34.22 | 5.48 | 2.29 | 0.42 | 2.68 | 13.77 | 4.01 | R | 13.68 | 9.61 | M | polynucleotide adenylyltransferase activity | RNA polyadenylation;mRNA polyadenylation | nucleus |
| PF3D7_1133200 | N/A | conserved Plasmodium protein, unknown function | 23.57 | no | 0 | 9.18 | 1.78 | 41.01 | 10.84 | 8.38 | 24.27 | 238.59 | O | 20.21 | 73.52 | F | N/A | N/A | nucleus |
| PF3D7_1340100 | RRP42 | exosome complex component RRP42, putative | 16.54 | no | 0 | 46.63 | 12.00 | 8.09 | 1.07 | 6.51 | 125.61 | 109.72 | GV | 14.16 | 50.47 | F | RNA binding | rRNA catabolic process | exosome (RNase complex) |
| PF3D7_0602100 | MTR4 | ATP-dependent RNA helicase MTR4 | 60.19 | no | 0 | 39.50 | 17.73 | 8.33 | 0.29 | 2.59 | 23.23 | 102.05 | O | 42.07 | 32.80 | M | RNA helicase activity;protein binding | RNA catabolic process;maturation of 5.8S rRNA | TRAMP complex;nuclear periphery;nucleus |
| PF3D7_1203900 | UBC | ubiquitin-conjugating enzyme E2 | 149.32 | no | 0 | 243.54 | 225.32 | 212.64 | 92.18 | 1228.58 | 1828.85 | 1225.87 | GV | 993.79 | 2149.05 | F | ubiquitin conjugating enzyme activity;ubiquitin-protein transferase activity | cellular protein modification process;protein polyubiquitination | cytoplasm;nucleus |
| PF3D7_1129400 | NSUN3 | RNA cytosine C(5)-methyltransferase, putative | 137.53 | no | 0 | 51.00 | 41.04 | 13.34 | 0.79 | 5.54 | 10.60 | 62.24 | O | 9.49 | 33.55 | F | rRNA (cytosine-C5-)-methyltransferase activity | RNA methylation;maturation of LSU-rRNA;rRNA base methylation | nucleolus;nucleus |
| PF3D7_0606500 | PTB | polypyrimidine tract-binding protein, putative | 61.42 | no | 0 | 70.90 | 8.85 | 13.32 | 1.82 | 4.21 | 28.39 | 65.98 | R | 51.31 | 17.20 | M | mRNA binding;poly-pyrimidine tract binding | RNA splicing;regulation of RNA splicing;regulation of translation | nucleus |
| PF3D7_1448000 | UTP12 | U3 small nucleolar RNA-associated protein 12, putative | 69.35 | no | 0 | 63.61 | 16.09 | 5.94 | 0.27 | 1.55 | 14.25 | 14.40 | R | 14.61 | 18.09 | F | snoRNA binding | maturation of SSU-rRNA | Pwp2p-containing subcomplex of 90S preribosome;nucleolus;small-subunit processome |
| PF3D7_1308900 | DCP2 | mRNA-decapping enzyme 2, putative | 10.11 | no | 0 | 181.57 | 78.75 | 30.88 | 5.29 | 3.39 | 21.18 | 472.16 | O | 151.21 | 60.83 | M | hydrolase activity;m7G(5')pppN diphosphatase activity | deadenylation-dependent decapping of nuclear-transcribed mRNA | cytoplasm;nucleus |
| PF3D7_0528200 | EIF3E | eukaryotic translation initiation factor 3 subunit E, putative | 102.00 | no | 2 | 146.91 | 55.97 | 43.09 | 4.03 | 11.95 | 60.74 | 27.87 | R | 32.63 | 19.57 | M | mRNA binding | translational initiation | eukaryotic translation initiation factor 3 complex;nucleus |
| PF3D7_1445600 | N/A | RNA-binding protein, putative | 60.80 | no | 0 | 132.96 | 23.29 | 19.46 | 3.38 | 7.11 | 32.08 | 80.16 | R | 54.83 | 15.09 | M | RNA binding | N/A | nucleus |
| PF3D7_1355800 | SF3B5 | splicing factor 3B subunit 5, putative | 671.11 | no | 0 | 36.43 | 14.21 | 9.89 | 1.42 | 20.39 | 22.03 | 12.41 | R | 52.57 | 8.22 | M | RNA binding | mRNA splicing, via spliceosome;spliceosomal complex assembly | U2 snRNP;nucleus;precatalytic spliceosome |
| PF3D7_1439700 | N/A | leucine carboxyl methyltransferase, putative | 59.69 | no | 0 | 2.23 | 0.56 | 9.97 | 1.80 | 8.09 | 17.56 | 13.33 | GV | 34.90 | 3.81 | M | methyltransferase activity | N/A | N/A |
| PF3D7_0217500 | CDPK1 | Calcium-dependent protein kinase 1 | 60.80 | no | 0 | 19906.00 | 0.85 | 60.65 | 51.87 | 31564.00 | 19.97 | 163.66 | GC II | 1341.63 | 638.05 | M | 14-3-3 protein binding;calcium-dependent protein serine/threonine kinase activity;calmodulin binding;protein binding;protein serine/threonine kinase activity | entry into host;exit from host cell;intracellular signal transduction;peptidyl-serine phosphorylation;protein autophosphorylation;protein phosphorylation;response to drug;response to xenobiotic stimulus | cytoplasm;membrane;nucleus;plasma membrane;symbiont-containing vacuolar space;symbiont-containing vacuole;symbiont-containing vacuole membrane |
| PF3D7_0410400 | RRP4 | exosome complex component RRP4 | 12.26 | no | 0 | 52.17 | 13.87 | 6.14 | 1.39 | 32.30 | 115.72 | 102.61 | GV | 98.27 | 27.55 | M | 3'-5'-exoribonuclease activity;RNA binding;protein binding | CUT catabolic process;U4 snRNA 3'-end processing;exonucleolytic catabolism of deadenylated mRNA;exonucleolytic trimming to generate mature 3'-end of 5.8S rRNA from tricistronic rRNA transcript (SSU-rRNA, 5.8S rRNA, LSU-rRNA);nuclear polyadenylation-dependent rRNA catabolic process;nuclear polyadenyl | TRAMP complex;cytoplasm;cytoplasmic exosome (RNase complex);exosome (RNase complex);nuclear exosome (RNase complex) |
| PF3D7_1224900 | SF3B6 | splicing factor 3B subunit 6, putative | 52.16 | no | 0 | 34.59 | 19.11 | 12.88 | 2.08 | 80.75 | 148.88 | 138.90 | GV | 128.99 | 78.97 | M | | mRNA splicing, via spliceosome | U12-type spliceosomal complex;U2 snRNP;U2-type spliceosomal complex;catalytic step 2 spliceosome;precatalytic spliceosome |
| PF3D7_0910900 | N/A | DNA primase large subunit, putative | 460.08 | no | 0 | 4.22 | 0.71 | 18.95 | 2.64 | 2.37 | 7.12 | 17.90 | T | 158.78 | 17.75 | M | DNA primase activity | DNA replication, synthesis of RNA primer | alpha DNA polymerase:primase complex;nucleus |
| PF3D7_0403400 | N/A | zinc finger protein, putative | 100.27 | no | 2 | 18.02 | 7.39 | 7.87 | 1.92 | 2.57 | 17.41 | 47.42 | O | 22.34 | 14.54 | M | protein binding | N/A | cytoplasm;nucleus |
| PF3D7_1124800 | RRS1 | ribosome biogenesis regulatory protein, putative | 62.88 | no | 0 | 122.18 | 36.73 | 16.31 | 2.89 | 17.11 | 117.72 | 331.35 | O | 171.60 | 308.15 | F | | endonucleolytic cleavage in ITS1 to separate SSU-rRNA from 5.8S rRNA and LSU-rRNA from tricistronic rRNA transcript (SSU-rRNA, 5.8S rRNA, LSU-rRNA);ribosomal large subunit biogenesis;ribosomal large subunit export from nucleus | nucleolus;preribosome, large subunit precursor |
| PF3D7_0811000 | CUL1 | cullin-1, putative | 118.33 | no | 0 | 3.06 | 0.27 | 13.37 | 5.72 | 3.63 | 18.00 | 17.24 | GV | 27.67 | 18.79 | M | protein binding;ubiquitin protein ligase binding;ubiquitin-protein transferase activity | SCF-dependent proteasomal ubiquitin-dependent protein catabolic process;protein ubiquitination | SCF ubiquitin ligase complex;cullin-RING ubiquitin ligase complex;nucleus;cytoplasm |
| PF3D7_1123400 | ERF3 | eukaryotic peptide chain release factor GTP-binding subunit, putative | 73.48 | no | 0 | 347.06 | 148.87 | 133.98 | 17.18 | 9.71 | 25.08 | 98.22 | R | 72.39 | 112.45 | F | GTP binding;GTPase activity;translation elongation factor activity;translation release factor activity | cytoplasmic translational termination;translation;translational elongation | eukaryotic translation elongation factor 1 complex;translation release factor complex |
| PF3D7_1305400 | N/A | AAR2 protein, putative | 75.92 | no | 0 | 34.05 | 6.80 | 7.72 | 0.15 | 2.59 | 33.19 | 71.69 | O | 26.63 | 34.42 | F | N/A | spliceosomal tri-snRNP complex assembly | nucleus |
| PF3D7_1362400 | Pcalp | calpain | 419.39 | no | 0 | 39.77 | 10.04 | 13.37 | 3.39 | 3.55 | 57.28 | 54.57 | GV | 68.70 | 171.79 | F | protease activity | signal transduction | cytoplasm;nucleolus |
| PF3D7_0810300 | PPM5 | protein phosphatase PPM5, putative | 67.79 | no | 0 | 10.73 | 13.05 | 70.65 | 31.75 | 22.62 | 85.93 | 556.33 | O | 804.95 | 255.60 | M | phosphatase activity | phosphorylation process | nucleus |
| PF3D7_0309000 | YVH1 | dual specificity protein phosphatase | 41.72 | no | 0 | 60.42 | 9.65 | 11.08 | 2.10 | 6.34 | 30.80 | 26.81 | R | 51.76 | 16.26 | M | phosphatase activity;protein binding;protein tyrosine/serine/threonine phosphatase activity;structural constituent of nuclear pore;zinc ion binding | negative regulation of nucleocytoplasmic transport;protein dephosphorylation;protein phosphorylation;regulation of cell development | cytoplasm;nucleus |
| PF3D7_1472900 | N/A | dihydroorotase, putative | 36.51 | no | 0 | 14.26 | 5.62 | 13.95 | 7.35 | 124.16 | 33.27 | 122.01 | G II | 14.14 | 18.27 | F | dihydroorotase activity | 'de novo' pyrimidine nucleobase biosynthetic process;pyrimidine nucleotide biosynthetic process | cytoplasm;nucleus |
| PF3D7_0703900 | N/A | conserved Plasmodium membrane protein | 129.20 | no | 11 | 66.37 | 25.85 | 3.78 | 0.47 | 1.54 | 25.05 | 51.77 | R | 36.54 | 12.81 | M | WW domain binding | N/A | membrane;nucleus |
| PF3D7_1227800 | ELP3 | elongator complex protein 3, putative | 311.42 | no | 1 | 79.99 | 63.53 | 12.51 | 0.95 | 2.49 | 5.07 | 79.10 | R | 15.93 | 13.07 | 0 | histone acetyltransferase activity | tRNA wobble base 5-methoxycarbonylmethyl-2-thiouridinylation | cytoplasm;elongator holoenzyme complex;mitochondrial outer membrane;mitochondrion;nucleus |
| PF3D7_1207000 | N/A | conserved Plasmodium protein, unknown function | 89.33 | no | 0 | 6.90 | 2.22 | 15.63 | 8.27 | 1.76 | 15.48 | 213.60 | O | 755.85 | 48.10 | M | N/A | N/A | mitochondrion;nucleus |
| PF3D7_0714200 | N/A | conserved Plasmodium protein, unknown function | 24.00 | no | 0 | 37.26 | 19.17 | 21.89 | 7.67 | 2.71 | 22.01 | 80.01 | O | 66.94 | 28.44 | M | N/A | N/A | nucleus |
| PF3D7_1472000 | ISY1 | pre-mRNA-splicing factor ISY1, putative | 135.82 | no | 0 | 29.46 | 19.38 | 12.22 | 3.31 | 82.25 | 67.17 | 26.31 | G II | 228.41 | 85.10 | M | RNA binding | generation of catalytic spliceosome for second transesterification step;mRNA 3'-splice site recognition | Prp19 complex;catalytic step 2 spliceosome;post-mRNA release spliceosomal complex;post-spliceosomal complex |
| PF3D7_1471400 | N/A | diacylglycerol kinase, putative | 129.05 | no | 0 | 4.28 | 1.04 | 26.01 | 12.51 | 2.74 | 17.93 | 110.87 | O | 68.16 | 25.67 | M | diacylglycerol kinase activity | protein kinase C-activating G protein-coupled receptor signaling pathway;second-messenger-mediated signaling | nucleus |
| PF3D7_0802300 | PWP2 | periodic tryptophan protein 2, putative | 93.75 | no | 0 | 113.16 | 54.11 | 16.96 | 1.74 | 6.04 | 12.09 | 66.54 | R | 45.73 | 39.97 | M | RNA binding | maturation of SSU-rRNA from tricistronic rRNA transcript (SSU-rRNA, 5.8S rRNA, LSU-rRNA);ribosomal small subunit assembly | Pwp2p-containing subcomplex of 90S preribosome;nucleolus;small-subunit processome |
| PF3D7_1008900 | AK1 | adenylate kinase | 154.90 | no | 0 | 93.97 | 78.85 | 85.11 | 11.95 | 23.24 | 32.91 | 132.43 | O | 18.20 | 34.44 | F | adenylate kinase activity | generation of precursor metabolites and energy;nucleoside diphosphate metabolic process;purine nucleotide metabolic process | cytoplasm;cytosol;mitochondrion;nucleus |
| PF3D7_1244100 | N/A | N-alpha-acetyltransferase 15, NatA auxiliary subunit, putative | 46.60 | no | 0 | 24.12 | 22.80 | 14.76 | 3.65 | 1.65 | 12.66 | 26.08 | O | 30.19 | 15.45 | M | enzyme activity | N-terminal peptidyl-methionine acetylation | cytoplasm;nucleus |

| | | | | | | | | | | | | | | | | | | | |
|---|---|---|---|---|---|---|---|---|---|---|---|---|---|---|---|---|---|---|---|
| PF3D7_1202100 | N/A | SRAP domain-containing protein, putative | 46.67 | no | 0 | 45992.00 | 45604.00 | 22.99 | 24473.00 | 28185.00 | 55.91 | 378.04 | R | 134.92 | 2993.15 | F | single-stranded DNA binding, peptidase activity | cellular response to DNA damage stimulus, protein-DNA covalent cross-linking | N/A |
| PF3D7_0530600 | N/A | XAP-5 DNA binding protein, putative | 52.03 | no | 0 | 40.83 | 8.08 | 5.15 | 0.61 | 13.30 | 86.11 | 66.41 | GV | 82.14 | 24.41 | M | DNA binding | chromatin organization | nucleus |
| PF3D7_1467500 | N/A | DNA/RNA-binding protein KIN17, putative | 52.46 | no | 0 | 43.88 | 13.86 | 20.36 | 6.75 | 1.42 | 6.50 | 7.67 | R | 21.23 | 13.09 | M | double-stranded DNA binding | DNA replication;cellular response to DNA damage stimulus | nucleus |
| PF3D7_0505200 | ALP2b | actin-like protein, putative | 65.87 | no | 0 | 10.44 | 6.93 | 10.08 | 4.39 | 1.26 | 14.33 | 61.93 | O | 1.63 | 23.44 | F | chromatin binding;structural constituent of cytoskeleton | ATP-dependent chromatin remodeling;cytoskeleton organization;histone H4 acetylation;regulation of transcription by RNA polymerase II | NuA4 histone acetyltransferase complex;SWI/SNF complex;actin cytoskeleton;nucleus |
| PF3D7_1363200 | PNKP | bifunctional polynucleotide phosphatase/kinase | 55.33 | no | 0 | 12.87 | 3.16 | 3.58 | 0.47 | 1.35 | 6.69 | 1.52 | R | 45.36 | 3.77 | M | damaged DNA binding;double-stranded DNA binding;polydeoxyribonucleotide 5'-hydroxyl-kinase activity;polynucleotide 3'-phosphatase activity | DNA repair;cellular response to DNA damage stimulus;nucleotide phosphorylation | nucleus |
| PF3D7_1315700 | GCD14 | tRNA (adenine(58)-N(1))-methyltransferase catalytic subunit TRM61, putative | 37.20 | no | 0 | 34.31 | 11.04 | 5.22 | 1.65 | 5.22 | 96.27 | 27.32 | GV | 3.81 | 91.57 | F | methyltransferase activity | tRNA methylation | nucleus;tRNA (m1A) methyltransferase complex |

Highest confidence (0.9000)

**k-means**

43 cluster

cluster with ≥7 proteins analyzed

disconnected nodes excluded

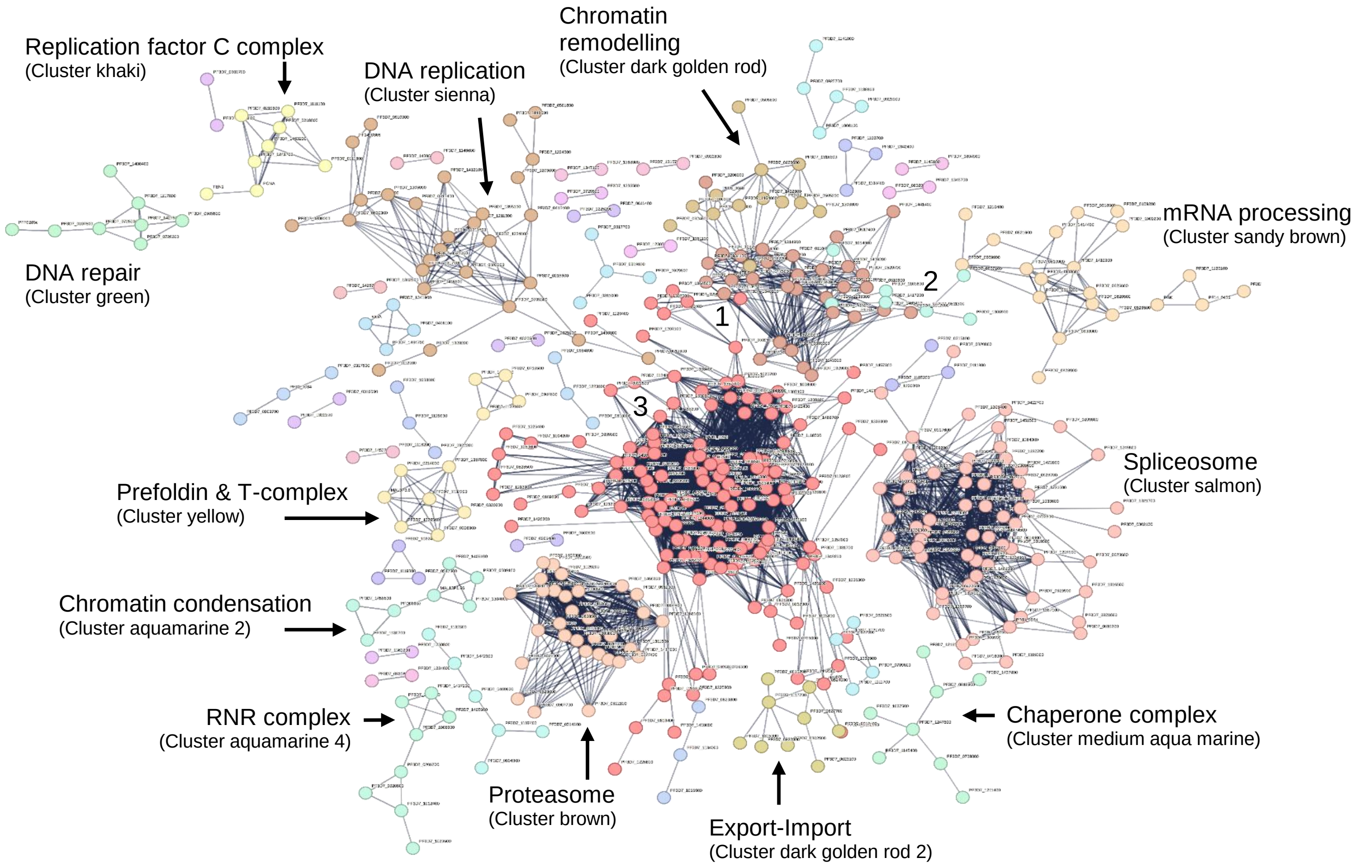

1 - Transcription (Cluster fire brick)
2 - CCR-NOT complex (Cluster cyan)
3 - Ribosome and translation (Cluster red)

| Ribosome and translation | Cluster number | Cluster color | Gene ID | Protein identifier | Protein description |
|---|---|---|---|---|---|
| | 1 | Red | PF3D7_0102900 | 36329.Q8I2B1 | Aspartate--tRNA ligase. |
| | 1 | Red | PF3D7_0109800 | 36329.Q8I246 | Phenylalanine--tRNA ligase alpha subunit. |
| | 1 | Red | PF3D7_0209200 | 36329.O96177 | Exosome complex component MTR3, putative. |
| | 1 | Red | PF3D7_0212300 | 36329.O96203 | Peptide chain release factor subunit 1, putative. |
| | 1 | Red | PF3D7_0217800 | 36329.O96258 | 40S ribosomal protein S26; Belongs to the eukaryotic ribosomal protein eS26 family. |
| | 1 | Red | PF3D7_0306900 | 36329.O97248 | 40S ribosomal protein S23, putative; Belongs to the universal ribosomal protein uS12 family. |
| | 1 | Red | PF3D7_0307100 | 36329.O97249 | 40S ribosomal protein S12; Belongs to the eukaryotic ribosomal protein eS12 family. |
| | 1 | Red | PF3D7_0309600 | 36329.O00806 | 60S acidic ribosomal protein P2; Plays an important role in the elongation step of protein synthesis. |
| | 1 | Red | PF3D7_0316800 | 36329.O77395 | 40S ribosomal protein S15A, putative; Belongs to the universal ribosomal protein uS8 family. |
| | 1 | Red | PF3D7_0317600 | 36329.O77381 | 40S ribosomal protein S11, putative; Belongs to the universal ribosomal protein uS17 family. |
| | 1 | Red | PF3D7_0322900 | 36329.O97313 | 40S ribosomal protein S3a; Belongs to the eukaryotic ribosomal protein eS1 family. |
| | 1 | Red | PF3D7_0410400 | 36329.Q9U0I4 | Exosome complex component RRP4. |
| | 1 | Red | PF3D7_0415900 | 36329.C0H4A6 | Ribosomal protein L15; Belongs to the eukaryotic ribosomal protein eL15 family. |
| | 1 | Red | PF3D7_0418200 | 36329.Q8I1Q5 | Eukaryotic translation initiation factor 3 subunit M; Component of the eukaryotic translation initiation factor 3 (eIF-3) complex, which is involved in protein synthesis of a specialized repertoire of mRNAs and, together with other initiation factors, stimulates binding of mRNA and methionyl-tRNAi to the 40S ribosome. The eIF-3 complex specifically targets and initiates translation of a subset of mRNAs involved in cell proliferation. |
| | 1 | Red | PF3D7_0422400 | 36329.Q8IFP2 | 40S ribosomal protein S19. |
| | 1 | Red | PF3D7_0507100 | 36329.Q8I431 | 60S ribosomal protein L4. |
| | 1 | Red | PF3D7_0507700 | 36329.Q8I426 | Nuclear protein localization protein 4, putative. |
| | 1 | Red | PF3D7_0516200 | 36329.A0A5K1K925 | 40S ribosomal protein S11; Belongs to the universal ribosomal protein uS11 family. |
| | 1 | Red | PF3D7_0516900 | 36329.Q8I3T9 | 60S ribosomal protein L2. |
| | 1 | Red | PF3D7_0517000 | 36329.Q8I3T8 | 60S ribosomal protein L12, putative; Belongs to the universal ribosomal protein uL11 family. |
| | 1 | Red | PF3D7_0517700 | 36329.Q8I3T1 | Eukaryotic translation initiation factor 3 subunit B; Component of the eukaryotic translation initiation factor 3 (eIF-3) complex, which is involved in protein synthesis and, together with other initiation factors, stimulates binding of mRNA and methionyl-tRNAi to the 40S ribosome; Belongs to the eIF-3 subunit B family. |
| | 1 | Red | PF3D7_0519400 | 36329.Q8I3R6 | 40S ribosomal protein S24; Belongs to the eukaryotic ribosomal protein eS24 family. |
| | 1 | Red | PF3D7_0520000 | 36329.Q8I3R0 | 40S ribosomal protein S9, putative; Belongs to the universal ribosomal protein uS4 family. |
| | 1 | Red | PF3D7_0528200 | 36329.Q8I3I5 | Eukaryotic translation initiation factor 3 subunit E; Component of the eukaryotic translation initiation factor 3 (eIF-3) complex, which is involved in protein synthesis of a specialized repertoire of mRNAs and, together with other initiation factors, stimulates binding of mRNA and methionyl-tRNAi to the 40S ribosome. The eIF-3 complex specifically targets and initiates translation of a subset of mRNAs involved in cell proliferation. |
| | 1 | Red | PF3D7_0602100 | 36329.C6KSM1 | ATP-dependent RNA helicase, putative. |
| | 1 | Red | PF3D7_0607000 | 36329.C6KSR8 | Translation initiation factor IF-2, putative. |
| | 1 | Red | PF3D7_0612100 | 36329.C6KSW5 | Eukaryotic translation initiation factor 3 subunit L; Component of the eukaryotic translation initiation factor 3 (eIF-3) complex, which is involved in protein synthesis of a specialized repertoire of mRNAs and, together with other initiation factors, stimulates binding of mRNA and methionyl-tRNAi to the 40S ribosome. The eIF-3 complex specifically targets and initiates translation of a subset of mRNAs involved in cell proliferation. |
| | 1 | Red | PF3D7_0618300 | 36329.C6KT23 | 60S ribosomal protein L27a, putative; Belongs to the universal ribosomal protein uL15 family. |
| | 1 | Red | PF3D7_0619400 | 36329.C6KT34 | Cell division cycle protein 48 homologue, putative. |
| | 1 | Red | PF3D7_0621800 | 36329.C6KT55 | Nascent polypeptide-associated complex subunit alpha, putative. |
| | 1 | Red | PF3D7_0705700 | 36329.C0H4K8 | 40S ribosomal protein S29, putative. |
| | 1 | Red | PF3D7_0706000 | 36329.C0H4L1 | Importin-7, putative. |
| | 1 | Red | PF3D7_0716800 | 36329.Q8IBT2 | Eukaryotic translation initiation factor 3 subunit I; Component of the eukaryotic translation initiation factor 3 (eIF-3) complex, which is involved in protein synthesis of a specialized repertoire of mRNAs and, together with other initiation factors, stimulates binding of mRNA and methionyl-tRNAi to the 40S ribosome. The eIF-3 complex specifically targets and initiates translation of a subset of mRNAs involved in cell proliferation. |
| | 1 | Red | PF3D7_0719600 | 36329.Q8IBQ6 | 60S ribosomal protein L11a, putative; Belongs to the universal ribosomal protein uL5 family. |
| | 1 | Red | PF3D7_0719700 | 36329.Q8IBQ5 | 40S ribosomal protein S10, putative. |
| | 1 | Red | PF3D7_0721600 | 36329.Q8IBN5 | 40S ribosomal protein S5, putative; Belongs to the universal ribosomal protein uS7 family. |
| | 1 | Red | PF3D7_0726500 | 36329.Q8IBJ1 | Ubiquitin carboxyl-terminal hydrolase, putative; Belongs to the peptidase C19 family. |
| | 1 | Red | PF3D7_0728000 | 36329.Q8IBH7 | Eukaryotic translation initiation factor 2 subunit alpha. |
| | 1 | Red | PF3D7_0802300 | 36329.Q8IAM3 | Periodic tryptophan protein 2, putative. |
| | 1 | Red | PF3D7_0813900 | 36329.Q8IAX5 | 40S ribosomal protein S16, putative; Belongs to the universal ribosomal protein uS9 family. |
| | 1 | Red | PF3D7_0815600 | 36329.Q8IAZ3 | Eukaryotic translation initiation factor 3 subunit G; RNA-binding component of the eukaryotic translation initiation factor 3 (eIF-3) complex, which is involved in protein synthesis of a specialized repertoire of mRNAs and, together with other initiation factors, stimulates binding of mRNA and methionyl-tRNAi to the 40S ribosome. The eIF-3 complex specifically targets and initiates translation of a subset of mRNAs involved in cell proliferation. This subunit can bind 18S rRNA. |
| | 1 | Red | PF3D7_0826300 | 36329.Q8IB96 | SPRY domain, putative. |
| | 1 | Red | PF3D7_0826700 | 36329.Q8IBA0 | Receptor for activated c kinase. |
| | 1 | Red | PF3D7_0828500 | 36329.Q8IBB6 | Translation initiation factor eIF-2B subunit alpha, putative; Belongs to the eIF-2B alpha/beta/delta subunits family. |
| | 1 | Red | PF3D7_0909900 | 36329.Q8I353 | Helicase SKI2W, putative. |
| | 1 | Red | PF3D7_0913200 | 36329.Q8I320 | Elongation factor 1-beta. |
| | 1 | Red | PF3D7_0918300 | 36329.Q8I2X0 | Eukaryotic translation initiation factor 3 subunit F, putative; Belongs to the eIF-3 subunit F family. |
| | 1 | Red | PF3D7_0921800 | 36329.C0H549 | Ribosome production factor 1, putative. |
| | 1 | Red | PF3D7_1003500 | 36329.Q8IK02 | 40S ribosomal protein S20e, putative. |
| | 1 | Red | PF3D7_1004000 | 36329.Q8IJZ7 | 60S ribosomal protein L13, putative. |
| | 1 | Red | PF3D7_1004200 | 36329.Q8IJZ5 | WD repeat-containing protein, putative. |
| | 1 | Red | PF3D7_1007900 | 36329.Q8IJW4 | Eukaryotic translation initiation factor 3 subunit D; mRNA cap-binding component of the eukaryotic translation initiation factor 3 (eIF-3) complex, which is involved in protein synthesis of a specialized repertoire of mRNAs and, together with other initiation factors, stimulates binding of mRNA and methionyl-tRNAi to the 40S ribosome. The eIF-3 complex specifically targets and initiates translation of a subset of mRNAs involved in cell proliferation. In the eIF-3 complex, eif3d specifically recognizes and binds the 7- methylguanosine cap of a subset of mRNAs. |
| | 1 | Red | PF3D7_1008800 | 36329.Q8IJV7 | Nucleolar protein 5, putative. |
| | 1 | Red | PF3D7_1010600 | 36329.Q8IJT9 | Eukaryotic translation initiation factor 2 subunit beta. |

| | | | | | |
|---|---|---|---|---|---|
| | 1 | Red | PF3D7_1013100 | 36329.Q8IJR4 | U3 small nucleolar RNA-associated protein 13, putative. |
| | 1 | Red | PF3D7_1019400 | 36329.Q8IJK8 | 60S ribosomal protein L30e, putative. |
| | 1 | Red | PF3D7_1020700 | 36329.Q8IJJ5 | RNA cytidine acetyltransferase; RNA cytidine acetyltransferase with specificity toward both 18S rRNA and tRNAs. Catalyzes the formation of N(4)-acetylcytidine (ac4C) in 18S rRNA. Required for early nucleolar cleavages of precursor rRNA at sites A0, A1 and A2 during 18S rRNA synthesis. Catalyzes the formation of ac4C in serine and leucine tRNAs. Requires a tRNA-binding adapter protein for full tRNA acetyltransferase activity but not for 18S rRNA acetylation. |
| | 1 | Red | PF3D7_1026800 | 36329.Q8IJD4 | 40S ribosomal protein SA; Required for the assembly and/or stability of the 40S ribosomal subunit. Required for the processing of the 20S rRNA- precursor to mature 18S rRNA in a late step of the maturation of 40S ribosomal subunits. |
| | 1 | Red | PF3D7_1027800 | 36329.Q8IJC6 | 60S ribosomal protein L3; Belongs to the universal ribosomal protein uL3 family. |
| | 1 | Red | PF3D7_1103100 | 36329.Q8IIX0 | 60S acidic ribosomal protein P1, putative; Belongs to the eukaryotic ribosomal protein P1/P2 family. |
| | 1 | Red | PF3D7_1104000 | 36329.Q8IIW2 | Phenylalanine--tRNA ligase beta subunit. |
| | 1 | Red | PF3D7_1105400 | 36329.Q8IIU8 | 40S ribosomal protein S4; Belongs to the eukaryotic ribosomal protein eS4 family. |
| | 1 | Red | PF3D7_1118500 | 36329.Q8III3 | Nucleolar protein 56, putative. |
| | 1 | Red | PF3D7_1121700 | 36329.Q8IIE9 | Protein GCN20. |
| | 1 | Red | PF3D7_1123400 | 36329.Q8IIC9 | Translation elongation factor EF-1, subunit alpha, putative. |
| | 1 | Red | PF3D7_1123900 | 36329.Q8IIC4 | Ribonucleoprotein; Common component of the spliceosome and rRNA processing machinery; Belongs to the eukaryotic ribosomal protein eL8 family. |
| | 1 | Red | PF3D7_1124800 | 36329.Q8IIB5 | Ribosome biogenesis regulatory protein; Involved in ribosomal large subunit assembly. Belongs to the RRS1 family. |
| | 1 | Red | PF3D7_1126200 | 36329.Q8IIA2 | 40S ribosomal protein S18, putative; Belongs to the universal ribosomal protein uS13 family. |
| | 1 | Red | PF3D7_1129400 | 36329.Q8II69 | rRNA (Cytosine-C(5))-methyltransferase, putative; Belongs to the class I-like SAM-binding methyltransferase superfamily. RsmB/NOP family. |
| | 1 | Red | PF3D7_1130100 | 36329.Q8II62 | 60S ribosomal protein L38; Belongs to the eukaryotic ribosomal protein eL38 family. |
| | 1 | Red | PF3D7_1130200 | 36329.Q8II61 | 60S acidic ribosomal protein P0; Ribosomal protein P0 is the functional equivalent of E.coli protein L10. |
| | 1 | Red | PF3D7_1142500 | 36329.Q8IHU0 | 60S ribosomal protein L28. |
| | 1 | Red | PF3D7_1144000 | 36329.Q8IHS5 | 40S ribosomal protein S21; Belongs to the eukaryotic ribosomal protein eS21 family. |
| | 1 | Red | PF3D7_1203900 | 36329.Q8I607 | Ubiquitin-conjugating enzyme E2; Belongs to the ubiquitin-conjugating enzyme family. |
| | 1 | Red | PF3D7_1206200 | 36329.Q8I5Y3 | Eukaryotic translation initiation factor 3 subunit C; Component of the eukaryotic translation initiation factor 3 (eIF-3) complex, which is involved in protein synthesis of a specialized repertoire of mRNAs and, together with other initiation factors, stimulates binding of mRNA and methionyl-tRNAi to the 40S ribosome. The eIF-3 complex specifically targets and initiates translation of a subset of mRNAs involved in cell proliferation. |
| | 1 | Red | PF3D7_1207100 | 36329.Q8I5X4 | Small subunit rRNA processing factor, putative. |
| | 1 | Red | PF3D7_1208100 | 36329.Q8I5W6 | Uncharacterized protein. |
| | 1 | Red | PF3D7_1211800 | 36329.Q7KQK2 | Polyubiquitin. |
| | 1 | Red | PF3D7_1212700 | 36329.Q8I5S6 | Eukaryotic translation initiation factor 3 subunit A; RNA-binding component of the eukaryotic translation initiation factor 3 (eIF-3) complex, which is involved in protein synthesis of a specialized repertoire of mRNAs and, together with other initiation factors, stimulates binding of mRNA and methionyl-tRNAi to the 40S ribosome. The eIF-3 complex specifically targets and initiates translation of a subset of mRNAs involved in cell proliferation. |
| | 1 | Red | PF3D7_1213800 | 36329.Q8I5R7 | Proline--tRNA ligase; Catalyzes the attachment of proline to tRNA(Pro) in a two- step reaction: proline is first activated by ATP to form Pro-AMP and then transferred to the acceptor end of tRNA(Pro) (By similarity). Functions in trans to edit the amino acid moiety from incorrectly charged Ala-tRNA(Pro). Has no activity on correctly charged Pro- tRNA(Pro) or Ala-tRNA(Ala). |
| | 1 | Red | PF3D7_1213900 | 36329.A0A144A0U7 | W2 domain-containing protein. |
| | 1 | Red | PF3D7_1225800 | 36329.Q8I5F9 | Ubiquitin-activating enzyme E1. |
| | 1 | Red | PF3D7_1226800 | 36329.Q8I5F0 | Ataxin-3, putative. |
| | 1 | Red | PF3D7_1242700 | 36329.Q8I502 | 40S ribosomal protein S17, putative. |
| | 1 | Red | PF3D7_1250600 | 36329.Q8I4S8 | Translation initiation factor eIF-2B subunit beta, putative; Belongs to the eIF-2B alpha/beta/delta subunits family. |
| | 1 | Red | PF3D7_1302800 | 36329.Q8IET7 | 40S ribosomal protein S7; Belongs to the eukaryotic ribosomal protein eS7 family. |
| | 1 | Red | PF3D7_1305300 | 36329.C0H5A0 | Translational activator GCN1, putative. |
| | 1 | Red | PF3D7_1307000 | 36329.Q8IEP5 | Exosome complex component RRP40, putative. |
| | 1 | Red | PF3D7_1307100 | 36329.Q8IEP4 | U3 small nucleolar RNA-associated protein 6, putative. |
| | 1 | Red | PF3D7_1308300 | 36329.Q8IEN2 | 40S ribosomal protein S27. |
| | 1 | Red | PF3D7_1309100 | 36329.Q8IEM3 | 60S ribosomal protein L24, putative. |
| | 1 | Red | PF3D7_1317800 | 36329.C0H5C2 | 40S ribosomal protein S19; Belongs to the universal ribosomal protein uS19 family. |
| | 1 | Red | PF3D7_1323100 | 36329.Q8IE85 | 60S ribosomal protein L6, putative. |
| | 1 | Red | PF3D7_1323400 | 36329.Q8IE82 | 60S ribosomal protein L23; Belongs to the universal ribosomal protein uL23 family. |
| | 1 | Red | PF3D7_1326400 | 36329.A0A5K1K977 | Translation initiation factor eIF-2B subunit gamma, putative. |
| | 1 | Red | PF3D7_1330600 | 36329.Q8IE20 | Elongation factor Tu. |
| | 1 | Red | PF3D7_1331700 | 36329.Q8IE10 | Glutamine--tRNA ligase, putative; Belongs to the class-I aminoacyl-tRNA synthetase family. |
| | 1 | Red | PF3D7_1331800 | 36329.Q8IE09 | 60S ribosomal protein L23, putative; Belongs to the universal ribosomal protein uL14 family. |
| | 1 | Red | PF3D7_1332900 | 36329.Q8IDZ9 | Isoleucine--tRNA ligase, putative; Belongs to the class-I aminoacyl-tRNA synthetase family. |
| | 1 | Red | PF3D7_1333600 | 36329.Q8IDZ2 | U3 small nucleolar RNA-associated protein 4, putative. |
| | 1 | Red | PF3D7_1338200 | 36329.A0A5K1K8V8 | 60S ribosomal protein L6, putative. |
| | 1 | Red | PF3D7_1338300 | 36329.A0A5K1K967 | Elongation factor 1-gamma, putative. |
| | 1 | Red | PF3D7_1340100 | 36329.A0A5K1K8M9 | Exosome complex component RRP42, putative. |
| | 1 | Red | PF3D7_1341200 | 36329.Q8IDS6 | 60S ribosomal protein L18a; Belongs to the eukaryotic ribosomal protein eL20 family. |
| | 1 | Red | PF3D7_1342000 | 36329.Q8IDR9 | 40S ribosomal protein S6. |
| | 1 | Red | PF3D7_1349200 | 36329.Q8IDK7 | Glutamate--tRNA ligase, putative; Belongs to the class-I aminoacyl-tRNA synthetase family. |
| | 1 | Red | PF3D7_1357000 | 36329.Q8I0P6 | Elongation factor 1-alpha; This protein promotes the GTP-dependent binding of aminoacyl- tRNA to the A-site of ribosomes during protein biosynthesis. |
| | 1 | Red | PF3D7_1357700 | 36329.A0A5K1K903 | U3 small nucleolar RNA-associated protein 21, putative. |
| | 1 | Red | PF3D7_1358800 | 36329.Q8IDB0 | 40S ribosomal protein S15; Belongs to the universal ribosomal protein uS15 family. |
| | 1 | Red | PF3D7_1364500 | 36329.Q8ID62 | Exosome complex component RRP45, putative. |
| | 1 | Red | PF3D7_1365900 | 36329.Q8ID50 | Ubiquitin-60S ribosomal protein L40. |
| | 1 | Red | PF3D7_1368200 | 36329.Q8I6Z4 | ABC transporter E family member 1, putative. |
| | 1 | Red | PF3D7_1368400 | 36329.Q8ID26 | Ribosomal protein L1, putative. |
| | 1 | Red | PF3D7_1402500 | 36329.Q8IM64 | Ribosomal protein S27a, putative. |

| | | | | | |
|---|---|---|---|---|---|
| | 1 | Red | PF3D7_1407100 | 36329.Q8IM23 | rRNA 2'-O-methyltransferase fibrillarin, putative. |
| | 1 | Red | PF3D7_1408600 | 36329.Q8IM10 | 40S ribosomal protein S8; Belongs to the eukaryotic ribosomal protein eS8 family. |
| | 1 | Red | PF3D7_1410600 | 36329.Q8ILY9 | Eukaryotic translation initiation factor 2 subunit gamma, putative. |
| | 1 | Red | PF3D7_1417500 | 36329.Q8ILS0 | H/ACA ribonucleoprotein complex subunit 4, putative. |
| | 1 | Red | PF3D7_1418000 | 36329.Q8ILR6 | Ubiquitin fusion degradation protein 1, putative. |
| | 1 | Red | PF3D7_1419700 | 36329.Q8ILQ3 | Uncharacterized protein. |
| | 1 | Red | PF3D7_1421200 | 36329.Q8ILN8 | 40S ribosomal protein S25; Belongs to the eukaryotic ribosomal protein eS25 family. |
| | 1 | Red | PF3D7_1422400 | 36329.Q8ILM9 | Uncharacterized protein. |
| | 1 | Red | PF3D7_1424100 | 36329.Q8ILL3 | 60S ribosomal protein L5, putative. |
| | 1 | Red | PF3D7_1426100 | 36329.Q8ILK2 | Nascent polypeptide-associated complex subunit beta. |
| | 1 | Red | PF3D7_1426200 | 36329.Q8ILK1 | Protein arginine N-methyltransferase 1; Belongs to the class I-like SAM-binding methyltransferase superfamily. Protein arginine N-methyltransferase family. |
| | 1 | Red | PF3D7_1441200 | 36329.Q8IL58 | 60S ribosomal protein L1, putative. |
| | 1 | Red | PF3D7_1442300 | 36329.Q8IL48 | tRNA import protein tRIP. |
| | 1 | Red | PF3D7_1447000 | 36329.Q8IL02 | 40S ribosomal protein S5; Belongs to the universal ribosomal protein uS5 family. |
| | 1 | Red | PF3D7_1448000 | 36329.Q8IKZ5 | U3 small nucleolar RNA-associated protein 12, putative. |
| | 1 | Red | PF3D7_1451100 | 36329.Q8IKW5 | Elongation factor 2. |
| | 1 | Red | PF3D7_1457300 | 36329.Q8IKQ7 | MA3 domain-containing protein, putative. |
| | 1 | Red | PF3D7_1460300 | 36329.C6S3J6 | 60S ribosomal protein L29; Belongs to the eukaryotic ribosomal protein eL29 family. |
| | 1 | Red | PF3D7_1460700 | 36329.Q8IKM5 | 60S ribosomal protein L27. |
| | 1 | Red | PF3D7_1461300 | 36329.Q8IKL9 | 40S ribosomal protein S28e, putative. |
| | 1 | Red | PF3D7_1465900 | 36329.Q8IKH8 | 40S ribosomal protein S3. |
| | 1 | Red | PF3D7_1468700 | 36329.Q8IKF0 | Eukaryotic initiation factor 4A; Belongs to the DEAD box helicase family. |

| Spliceosome | Cluster number | Cluster color | Gene ID | Protein identifier | Protein description |
|---|---|---|---|---|---|
| | 2 | Salmon | PF3D7_0209800 | 36329.Q9TY94 | ATP-dependent RNA helicase UAP56. |
| | 2 | Salmon | PF3D7_0218500 | 36329.O96265 | Small nuclear ribonucleoprotein Sm D2. |
| | 2 | Salmon | PF3D7_0218700 | 36329.O96267 | Pre-mRNA-processing protein 45, putative. |
| | 2 | Salmon | PF3D7_0302000 | 36329.O97334 | Pre-mRNA-splicing factor PRP46, putative. |
| | 2 | Salmon | PF3D7_0302100 | 36329.O77306 | Serine/threonine protein kinase. |
| | 2 | Salmon | PF3D7_0308600 | 36329.O77325 | Pre-mRNA-processing factor 19, putative. |
| | 2 | Salmon | PF3D7_0319500 | 36329.O97318 | RNA-binding protein, putative. |
| | 2 | Salmon | PF3D7_0320800 | 36329.O97285 | ATP-dependent RNA helicase DDX6; Belongs to the DEAD box helicase family. |
| | 2 | Salmon | PF3D7_0321600 | 36329.O97290 | ATP-dependent RNA helicase DDX42, putative. |
| | 2 | Salmon | PF3D7_0323700 | 36329.O97303 | U4/U6.U5 tri-snRNP-associated protein 1, putative. |
| | 2 | Salmon | PF3D7_0403700 | 36329.Q8I1Z2 | Pre-mRNA-splicing factor CLF1, putative. |
| | 2 | Salmon | PF3D7_0405400 | 36329.Q8I1X5 | Pre-mRNA-processing-splicing factor 8, putative. |
| | 2 | Salmon | PF3D7_0422500 | 36329.Q8IFP1 | Pre-mRNA-splicing helicase BRR2, putative. |
| | 2 | Salmon | PF3D7_0422700 | 36329.Q8IFN9 | Eukaryotic initiation factor 4A-III, putative; Belongs to the DEAD box helicase family. |
| | 2 | Salmon | PF3D7_0520300 | 36329.Q8I3Q7 | U6 snRNA-associated Sm-like protein LSm2; Binds specifically to the 3'-terminal U-tract of U6 snRNA. |
| | 2 | Salmon | PF3D7_0522800 | 36329.Q8I3N5 | Pre-mRNA-splicing factor BUD31, putative. |
| | 2 | Salmon | PF3D7_0528700 | 36329.Q8I3I0 | Peptidyl-prolyl cis-trans isomerase; PPIases accelerate the folding of proteins. It catalyzes the cis-trans isomerization of proline imidic peptide bonds in oligopeptides; Belongs to the cyclophilin-type PPIase family. |
| | 2 | Salmon | PF3D7_0610200 | 36329.C6KSU9 | RNA-binding protein 25, putative. |
| | 2 | Salmon | PF3D7_0614400 | 36329.C6KSY5 | Pre-mRNA-splicing factor CWF7, putative. |
| | 2 | Salmon | PF3D7_0619900 | 36329.C6KT39 | Splicing factor 3A subunit 2, putative. |
| | 2 | Salmon | PF3D7_0623600 | 36329.C6KT72 | Transcription or splicing factor-like protein, putative. |
| | 2 | Salmon | PF3D7_0716000 | 36329.Q8IBU0 | RNA-binding protein, putative. |
| | 2 | Salmon | PF3D7_0722500 | 36329.Q8IBM8 | Pre-mRNA-splicing factor CWC15, putative. |
| | 2 | Salmon | PF3D7_0812700 | 36329.Q8IAW3 | U1 small nuclear ribonucleoprotein C; Component of the spliceosomal U1 snRNP, which is essential for recognition of the pre-mRNA 5' splice-site and the subsequent assembly of the spliceosome. U1-C is directly involved in initial 5' splice-site recognition for both constitutive and regulated alternative splicing. The |
| | 2 | Salmon | PF3D7_0819900 | 36329.C0H4W2 | U6 snRNA-associated Sm-like protein LSm3; Binds specifically to the 3'-terminal U-tract of U6 snRNA. |
| | 2 | Salmon | PF3D7_0822300 | 36329.Q8IB57 | Small nuclear ribonucleoprotein G. |
| | 2 | Salmon | PF3D7_0822800 | 36329.C0H4X1 | U5 small nuclear ribonucleoprotein 40 kDa protein, putative. |
| | 2 | Salmon | PF3D7_0909800 | 36329.C0H529 | Small nuclear ribonucleoprotein Sm D3. |
| | 2 | Salmon | PF3D7_0917600 | 36329.Q8I2X7 | Pre-mRNA-splicing factor ATP-dependent RNA helicase PRP43, putative. |
| | 2 | Salmon | PF3D7_0924700 | 36329.Q8I2R0 | Splicing factor 3A subunit 3, putative. |
| | 2 | Salmon | PF3D7_0935000 | 36329.Q8I2G9 | U2 small nuclear ribonucleoprotein B'', putative. |
| | 2 | Salmon | PF3D7_1003800 | 36329.Q8IJZ9 | U5 small nuclear ribonucleoprotein component, putative. |
| | 2 | Salmon | PF3D7_1033600 | 36329.Q7KQL1 | Pre-mRNA-splicing factor CEF1, putative. |
| | 2 | Salmon | PF3D7_1107000 | 36329.Q8IIT3 | U6 snRNA-associated Sm-like protein LSm4; Binds specifically to the 3'-terminal U-tract of U6 snRNA. |
| | 2 | Salmon | PF3D7_1110200 | 36329.Q8IIR0 | Pre-mRNA-processing factor 6, putative. |
| | 2 | Salmon | PF3D7_1119300 | 36329.Q8IIH4 | Splicing factor U2AF small subunit, putative. |
| | 2 | Salmon | PF3D7_1125500 | 36329.Q8IIA8 | Small nuclear ribonucleoprotein Sm D1; Essential for pre-mRNA splicing. Implicated in the formation of stable, |
| | 2 | Salmon | PF3D7_1126900 | 36329.Q8II94 | Small nuclear ribonucleoprotein F, putative. |
| | 2 | Salmon | PF3D7_1209200 | 36329.Q8I5V6 | U6 snRNA-associated Sm-like protein LSm7, putative. |
| | 2 | Salmon | PF3D7_1220100 | 36329.Q8I5L1 | Pre-mRNA-processing factor 17, putative. |
| | 2 | Salmon | PF3D7_1224900 | 36329.Q8I5G8 | Splicing factor 3B subunit 6, putative. |
| | 2 | Salmon | PF3D7_1230800 | 36329.Q8I5B2 | Pre-mRNA-splicing regulator, putative. |
| | 2 | Salmon | PF3D7_1231600 | 36329.Q8I5A4 | Pre-mRNA-splicing factor ATP-dependent RNA helicase PRP2, putative. |

| | | | | | |
|---|---|---|---|---|---|
| | 2 | Salmon | PF3D7_1234800 | 36329.Q8I574 | Splicing factor 3B subunit 3, putative. |
| | 2 | Salmon | PF3D7_1235900 | 36329.A0A144A2S8 | Pre-mRNA-splicing factor SYF1, putative. |
| | 2 | Salmon | PF3D7_1249800 | 36329.Q8I4T6 | THO complex subunit 2, putative. |
| | 2 | Salmon | PF3D7_1305400 | 36329.Q8IER1 | AAR2 protein, putative. |
| | 2 | Salmon | PF3D7_1306900 | 36329.C0H5A4 | U1 small nuclear ribonucleoprotein A, putative. |
| | 2 | Salmon | PF3D7_1309300 | 36329.Q8IEM1 | U4/U6 small nuclear ribonucleoprotein PRP3, putative. |
| | 2 | Salmon | PF3D7_1316500 | 36329.A0A5K1K8A8 | Pre-mRNA-processing factor 40, putative. |
| | 2 | Salmon | PF3D7_1321700 | 36329.Q8IE99 | Splicing factor 1. |
| | 2 | Salmon | PF3D7_1324200 | 36329.Q8IE75 | Micro-fibrillar-associated protein, putative. |
| | 2 | Salmon | PF3D7_1343900 | 36329.C0H5G7 | U4/U6 small nuclear ribonucleoprotein PRP4, putative. |
| | 2 | Salmon | PF3D7_1352700 | 36329.Q8IDH3 | Intron-binding protein aquarius, putative. |
| | 2 | Salmon | PF3D7_1355800 | 36329.C0H5I5 | Splicing factor subunit; Belongs to the SF3B5 family. |
| | 2 | Salmon | PF3D7_1364300 | 36329.A0A5K1K9B8 | Pre-mRNA-splicing factor ATP-dependent RNA helicase PRP16. |
| | 2 | Salmon | PF3D7_1367100 | 36329.Q8ID37 | U1 small nuclear ribonucleoprotein 70 kDa homolog, putative. |
| | 2 | Salmon | PF3D7_1369700 | 36329.C0H5L8 | U2 small nuclear ribonucleoprotein A', putative. |
| | 2 | Salmon | PF3D7_1414800 | 36329.Q8ILU8 | Small nuclear ribonucleoprotein-associated protein B, putative. |
| | 2 | Salmon | PF3D7_1420000 | 36329.Q8ILQ0 | Splicing factor 3B subunit 4, putative. |
| | 2 | Salmon | PF3D7_1451500 | 36329.Q8IKW1 | Pre-mRNA-splicing factor CWF18, putative. |
| | 2 | Salmon | PF3D7_1461600 | 36329.Q8IKL7 | Splicing factor 3B subunit 2, putative. |
| | 2 | Salmon | PF3D7_1472000 | 36329.Q8IKB8 | Pre-mRNA-splicing factor ISY1, putative. |
| | 2 | Salmon | PF3D7_1474500 | 36329.Q8IK93 | Splicing factor 3A subunit 1, putative. |

| Transcription | Cluster number | Cluster color | Gene ID | Protein identifier | Protein description |
|---|---|---|---|---|---|
| | 3 | Fire Brick | PF3D7_0110400 | 36329.Q8I241 | DNA-directed RNA polymerase II subunit RPB9, putative. |
| | 3 | Fire Brick | PF3D7_0206000 | 36329.O96154 | DNA repair protein RAD2, putative. |
| | 3 | Fire Brick | PF3D7_0215700 | 36329.O96236 | DNA-directed RNA polymerase subunit beta; DNA-dependent RNA polymerase catalyzes the transcription of DNA into RNA using the four ribonucleoside triphosphates as substrates. |
| | 3 | Fire Brick | PF3D7_0303300 | 36329.O77315 | DNA-directed RNA polymerases I, II, and III subunit RPABC2, putative. |
| | 3 | Fire Brick | PF3D7_0318200 | 36329.O77375 | DNA-directed RNA polymerase subunit; DNA-dependent RNA polymerase catalyzes the transcription of DNA into RNA using the four ribonucleoside triphosphates as substrates. |
| | 3 | Fire Brick | PF3D7_0509400 | 36329.Q8I410 | DNA-directed RNA polymerase subunit; DNA-dependent RNA polymerase catalyzes the transcription of DNA into RNA using the four ribonucleoside triphosphates as substrates. |
| | 3 | Fire Brick | PF3D7_0517400 | 36329.Q8I3T4 | FACT complex subunit SPT16, putative. |
| | 3 | Fire Brick | PF3D7_0520700 | 36329.Q8I3Q4 | CDC73 domain-containing protein, putative. |
| | 3 | Fire Brick | PF3D7_0610900 | 36329.C6KSV4 | Transcription elongation factor SPT5, putative. |
| | 3 | Fire Brick | PF3D7_0708100 | 36329.Q8IC08 | DNA-directed RNA polymerases I, II, and III subunit RPABC5, putative. |
| | 3 | Fire Brick | PF3D7_0714500 | 36329.Q8IBV2 | Transcription elongation factor s-II, putative. |
| | 3 | Fire Brick | PF3D7_0822100 | 36329.Q8IB55 | Mediator of RNA polymerase II transcription subunit 7; Component of the Mediator complex, a coactivator involved in the regulated transcription of nearly all RNA polymerase II-dependent genes. Mediator functions as a bridge to convey information from gene- specific regulatory proteins to the basal RNA polymerase II transcription machinery. |
| | 3 | Fire Brick | PF3D7_0923000 | 36329.Q8I2S6 | DNA-directed RNA polymerase II subunit RPB3, putative. |
| | 3 | Fire Brick | PF3D7_0934100 | 36329.Q8I2H7 | TFIIH basal transcription factor complex helicase XPD subunit. |
| | 3 | Fire Brick | PF3D7_1012700 | 36329.Q8IJR8 | NLI interacting factor-like phosphatase, putative. |
| | 3 | Fire Brick | PF3D7_1037600 | 36329.Q8IJ31 | TFIIH basal transcription factor complex helicase XPB subunit, putative. |
| | 3 | Fire Brick | PF3D7_1134700 | 36329.Q8II17 | DNA-directed RNA polymerase subunit beta; DNA-dependent RNA polymerase catalyzes the transcription of DNA into RNA using the four ribonucleoside triphosphates as substrates. |
| | 3 | Fire Brick | PF3D7_1143300 | 36329.Q8IHT3 | DNA-directed RNA polymerases I and III subunit RPAC1, putative. |
| | 3 | Fire Brick | PF3D7_1206600 | 36329.Q8I5X9 | DNA-directed RNA polymerase subunit beta; DNA-dependent RNA polymerase catalyzes the transcription of DNA into RNA using the four ribonucleoside triphosphates as substrates. |
| | 3 | Fire Brick | PF3D7_1213700 | 36329.Q8I5R8 | DNA-directed RNA polymerases I, II, and III subunit RPABC3; DNA-dependent RNA polymerase catalyzes the transcription of DNA into RNA using the four ribonucleoside triphosphates as substrates. Common component of RNA polymerases I, II and III which synthesize ribosomal RNA precursors, mRNA precursors and many functional non- coding RNAs, and small RNAs, such as 5S rRNA and tRNAs, respectively. |
| | 3 | Fire Brick | PF3D7_1244200 | 36329.Q8I4Y8 | General transcription factor IIH subunit 4; Component of the general transcription and DNA repair factor IIH (TFIIH) core complex which is involved in general and transcription-coupled nucleotide excision repair (NER) of damaged DNA. Belongs to the TFB2 family. |
| | 3 | Fire Brick | PF3D7_1314900 | 36329.Q8IEG6 | General transcription factor IIH subunit. |
| | 3 | Fire Brick | PF3D7_1329000 | 36329.Q8IE49 | DNA-directed RNA polymerase subunit; DNA-dependent RNA polymerase catalyzes the transcription of DNA into RNA using the four ribonucleoside triphosphates as substrates. |
| | 3 | Fire Brick | PF3D7_1364800 | 36329.Q8ID59 | DNA-directed RNA polymerases I, II, and III subunit RPABC1, putative. |
| | 3 | Fire Brick | PF3D7_1406200 | 36329.Q8IM32 | Transcription elongation factor SPT6, putative. |
| | 3 | Fire Brick | PF3D7_1415200 | 36329.Q8ILU4 | DNA-directed RNA polymerases I and III subunit RPAC2, putative. |
| | 3 | Fire Brick | PF3D7_1441400 | 36329.Q8IL56 | FACT complex subunit SSRP1; Component of the FACT complex, a general chromatin factor that acts to reorganize nucleosomes. The FACT complex is involved in multiple processes that require DNA as a template such as mRNA elongation, DNA replication and DNA repair. During transcription elongation the FACT complex acts as a histone chaperone that both destabilizes and restores nucleosomal structure. It facilitates the passage of RNA polymerase II and transcription by promoting the dissociation of one histone H2A-H2B dimer from the nucleosome, then subsequently promotes the reestablishment o [...] |
| | 3 | Fire Brick | PF3D7_1458800 | 36329.Q8IKP3 | DNA-directed RNA polymerase III subunit RPC5, putative. |
| | 3 | Fire Brick | PF3D7_1463400 | 36329.Q8IKK2 | DNA-directed RNA polymerase III subunit RPC4, putative. |
| | 3 | Fire Brick | PF3D7_1469700 | 36329.Q8IKE0 | Mediator of RNA polymerase II transcription subunit 6; Component of the Mediator complex, a coactivator involved in the regulated transcription of nearly all RNA polymerase II-dependent genes. Mediator functions as a bridge to convey information from gene- specific regulatory proteins to the basal RNA polymerase II transcription machinery. Mediator is recruited to promoters by direct interactions with regulatory proteins and serves as a scaffold for the assembly of a functional preinitiation complex with RNA polymerase II and the general transcription factors. |
| | 3 | Fire Brick | PF3D7_1475000 | 36329.Q8IK88 | Mediator of RNA polymerase II transcription subunit 31 |

| Cluster number | Cluster color | Gene ID | Protein identifier | Protein description |
|---|---|---|---|---|
| 4 | Brown | PF3D7_0205900 | 36329.O96153 | 26S proteasome regulatory subunit RPN1, putative. |
| 4 | Brown | PF3D7_0312300 | 36329.O77345 | 26S proteasome regulatory subunit RPN12, putative. |
| 4 | Brown | PF3D7_0317000 | 36329.O77396 | Proteasome subunit alpha type-3, putative. |
| 4 | Brown | PF3D7_0413600 | 36329.Q8I1V1 | 26S protease regulatory subunit 6B, putative; Belongs to the AAA ATPase family. |
| 4 | Brown | PF3D7_0518300 | 36329.A0A5K1K7U1 | Proteasome subunit beta; The proteasome is a multicatalytic proteinase complex which is characterized by its ability to cleave peptides with Arg, Phe, Tyr, Leu, and Glu adjacent to the leaving group at neutral or slightly basic pH; Belongs to the peptidase T1B family. |
| 4 | Brown | PF3D7_0528000 | 36329.A0A5K1K941 | Proteasome maturation factor UMP1, putative. |
| 4 | Brown | PF3D7_0608500 | 36329.C6KST3 | Proteasome endopeptidase complex; Belongs to the peptidase T1A family. |
| 4 | Brown | PF3D7_0727400 | 36329.Q8IBI3 | Proteasome subunit alpha type. |
| 4 | Brown | PF3D7_0803800 | 36329.Q7K6A9 | Proteasome subunit beta; Belongs to the peptidase T1B family. |
| 4 | Brown | PF3D7_0807500 | 36329.Q8IAR3 | Proteasome endopeptidase complex; Belongs to the peptidase T1A family. |
| 4 | Brown | PF3D7_0807800 | 36329.Q8IAR6 | 26S proteasome regulatory subunit RPN10, putative. |
| 4 | Brown | PF3D7_0812100 | 36329.Q8IAV6 | Proteasome activator complex subunit 4, putative. |
| 4 | Brown | PF3D7_0907700 | 36329.Q8I374 | Proteasome activator 28 subunit beta, putative. |
| 4 | Brown | PF3D7_0912900 | 36329.Q8I323 | 26S proteasome regulatory subunit RPN8, putative. |
| 4 | Brown | PF3D7_1008400 | 36329.Q8IJW0 | 26S protease regulatory subunit 4, putative; Belongs to the AAA ATPase family. |
| 4 | Brown | PF3D7_1017900 | 36329.Q8IJM0 | 26S proteasome regulatory subunit p55, putative. |
| 4 | Brown | PF3D7_1021900 | 36329.Q8IJI4 | PHAX domain-containing protein, putative. |
| 4 | Brown | PF3D7_1030500 | 36329.Q8IJA0 | 26S proteasome regulatory subunit RPN9, putative. |
| 4 | Brown | PF3D7_1129200 | 36329.Q8II71 | 26S proteasome regulatory subunit RPN7, putative. |
| 4 | Brown | PF3D7_1130400 | 36329.Q8II60 | 26S protease regulatory subunit 6A, putative; Belongs to the AAA ATPase family. |
| 4 | Brown | PF3D7_1248900 | 36329.Q8I4U5 | 26S protease regulatory subunit 8, putative; Belongs to the AAA ATPase family. |
| 4 | Brown | PF3D7_1306400 | 36329.Q8IEQ1 | 26S protease regulatory subunit 10B, putative; Belongs to the AAA ATPase family. |
| 4 | Brown | PF3D7_1311500 | 36329.Q8IEK3 | 26S protease regulatory subunit 7, putative; Belongs to the AAA ATPase family. |
| 4 | Brown | PF3D7_1338100 | 36329.A0A5K1K9F3 | 26S proteasome regulatory subunit RPN3, putative. |
| 4 | Brown | PF3D7_1353900 | 36329.Q8IDG2 | Proteasome subunit alpha type. |
| 4 | Brown | PF3D7_1368100 | 36329.Q8ID28 | 26S proteasome regulatory subunit RPN11, putative. |
| 4 | Brown | PF3D7_1402300 | 36329.Q8IM66 | 26S proteasome regulatory subunit RPN6. |
| 4 | Brown | PF3D7_1414000 | 36329.Q8ILV5 | 26S proteasome regulatory subunit RPN13, putative. |
| 4 | Brown | PF3D7_1466300 | 36329.Q8IKH3 | 26S proteasome regulatory subunit RPN2, putative; Belongs to the proteasome subunit S1 family. |
| 4 | Brown | PF3D7_1470900 | 36329.Q8IKC9 | Proteasome subunit beta; The proteasome is a multicatalytic proteinase complex which is characterized by its ability to cleave peptides with Arg, Phe, Tyr, Leu, and Glu adjacent to the leaving group at neutral or slightly basic pH; Belongs to the peptidase T1B family. |
| 4 | Brown | PF3D7_1474800 | 36329.Q8IK90 | Proteasome endopeptidase complex; Belongs to the peptidase T1A family. |

| DNA replication | Cluster number | Cluster color | Gene ID | Protein identifier | Protein description |
|---|---|---|---|---|---|
| | 5 | Sienna | PF3D7_0215800 | 36329.O96237 | Origin recognition complex subunit 5. |
| | 5 | Sienna | PF3D7_0308000 | 36329.O77321 | DNA polymerase delta small subunit, putative. |
| | 5 | Sienna | PF3D7_0317400 | 36329.O77383 | DNA replication complex GINS protein, putative. |
| | 5 | Sienna | PF3D7_0411900 | 36329.Q9U0H1 | DNA polymerase. |
| | 5 | Sienna | PF3D7_0416300 | 36329.Q8I1S4 | DNA helicase MCM9, putative; Belongs to the MCM family. |
| | 5 | Sienna | PF3D7_0416400 | 36329.C0H4A7 | Histone acetyltransferase, putative. |
| | 5 | Sienna | PF3D7_0501800 | 36329.Q8I482 | Protein HIRA; Required for replication-independent chromatin assembly and for the periodic repression of histone gene transcription during the cell cycle; Belongs to the WD repeat HIR1 family. |
| | 5 | Sienna | PF3D7_0503200 | 36329.Q8I469 | ATPase_AAA_core domain-containing protein. |
| | 5 | Sienna | PF3D7_0527000 | 36329.Q8I3J5 | DNA replication licensing factor MCM3, putative; Belongs to the MCM family. |
| | 5 | Sienna | PF3D7_0612900 | 36329.C6KSX2 | Nucleolar GTP-binding protein 1; Involved in the biogenesis of the 60S ribosomal subunit. Belongs to the TRAFAC class OBG-HflX-like GTPase superfamily. OBG GTPase family. NOG subfamily. |
| | 5 | Sienna | PF3D7_0630300 | 36329.C6KTD8 | DNA polymerase epsilon catalytic subunit A, putative. |
| | 5 | Sienna | PF3D7_0705300 | 36329.Q8IC17 | Origin recognition complex subunit 2. |
| | 5 | Sienna | PF3D7_0705400 | 36329.Q8IC16 | DNA replication licensing factor MCM7; Acts as component of the mcm2-7 complex (mcm complex) which is the putative replicative helicase essential for 'once per cell cycle' DNA replication initiation and elongation in eukaryotic cells. The active ATPase sites in the mcm2-7 ring are formed through the interaction surfaces of two neighboring subunits such that a critical structure of a conserved arginine finger motif is provided in trans relative to the ATP-binding site of the Walker A box of the adjacent subunit. The six ATPase active sites, however, are likely to contribute differential [...] |
| | 5 | Sienna | PF3D7_0910900 | 36329.C0H531 | DNA primase large subunit; DNA primase is the polymerase that synthesizes small RNA primers for the Okazaki fragments made during discontinuous DNA replication; Belongs to the eukaryotic-type primase large subunit family. |
| | 5 | Sienna | PF3D7_1112100 | 36329.Q8IIP2 | Protein kinase domain-containing protein. |
| | 5 | Sienna | PF3D7_1148000 | 36329.Q8IHP1 | Serine/threonine protein kinase, putative. |
| | 5 | Sienna | PF3D7_1205600 | 36329.Q8I5Y9 | Tetratricopeptide repeat protein, putative. |
| | 5 | Sienna | PF3D7_1211300 | 36329.Q8I5T7 | DNA helicase MCM8, putative; Belongs to the MCM family. |
| | 5 | Sienna | PF3D7_1211700 | 36329.Q8I5T4 | DNA replication licensing factor MCM5, putative; Belongs to the MCM family. |
| | 5 | Sienna | PF3D7_1224500 | 36329.Q8I5H2 | Histone chaperone ASF1, putative. |
| | 5 | Sienna | PF3D7_1305000 | 36329.Q8IER6 | MCL1 domain-containing protein, putative. |
| | 5 | Sienna | PF3D7_1317100 | 36329.Q8IEE5 | DNA replication licensing factor MCM4; Belongs to the MCM family. |
| | 5 | Sienna | PF3D7_1328200 | 36329.A0A5K1K982 | Uncharacterized protein. |
| | 5 | Sienna | PF3D7_1334100 | 36329.A0A5K1K8V1 | Uncharacterized protein. |
| | 5 | Sienna | PF3D7_1355100 | 36329.Q8IDF0 | DNA helicase; Belongs to the MCM family. |
| | 5 | Sienna | PF3D7_1412100 | 36329.Q8ILX3 | Mini-chromosome maintenance complex-binding protein, putative. |
| | 5 | Sienna | PF3D7_1438700 | 36329.Q7KQM1 | DNA primase small subunit; DNA primase is the polymerase that synthesizes small RNA primers for the Okazaki fragments made during discontinuous DNA replication. |
| | 5 | Sienna | PF3D7_1466800 | 36329.Q8IKG9 | NOC3p domain-containing protein. |

| mRNA processing | Cluster number | Cluster color | Gene ID | Protein identifier | Protein description |
|---|---|---|---|---|---|
| | 6 | Sandy Brown | PF3D7_0109200 | 36329.Q8I0V9 | Cleavage and polyadenylation specificity factor subunit 5; Component of the cleavage factor Im (CFIm) complex that functions as an activator of the pre-mRNA 3'-end cleavage and polyadenylation processing required for the maturation of pre-mRNA into functional mRNAs. CFIm contributes to the recruitment of multiprotein complexes on specific sequences on the pre-mRNA 3'-end, so called cleavage and polyadenylation signals (pA signals). Most pre-mRNAs contain multiple pA signals, resulting in alternative cleavage and polyadenylation (APA) producing mRNAs with variable 3'-end formation. The [...] |
| | 6 | Sandy Brown | PF3D7_0318600 | 36329.O77371 | Cleavage and polyadenylation specificity factor, putative. |
| | 6 | Sandy Brown | PF3D7_0513200 | 36329.Q8I3X5 | CPSF_A domain-containing protein. |
| | 6 | Sandy Brown | PF3D7_0526500 | 36329.Q8I3K0 | Suf domain-containing protein. |
| | 6 | Sandy Brown | PF3D7_0609900 | 36329.C6KSU6 | Uncharacterized protein. |
| | 6 | Sandy Brown | PF3D7_0620500 | 36329.C6KT45 | Cleavage stimulation factor subunit 1, putative. |
| | 6 | Sandy Brown | PF3D7_0624000 | 36329.C6KT76 | Phosphotransferase. |
| | 6 | Sandy Brown | PF3D7_0625600 | 36329.C6KT92 | Poly(A) polymerase; Polymerase that creates the 3'-poly(A) tail of mRNA's. |
| | 6 | Sandy Brown | PF3D7_0819000 | 36329.Q8IB25 | Cleavage and polyadenylation specificity factor subunit 2. |
| | 6 | Sandy Brown | PF3D7_0821600 | 36329.Q8IB50 | Polyribonucleotide 5'-hydroxyl-kinase Clp1, putative. |
| | 6 | Sandy Brown | PF3D7_0922500 | 36329.P27362 | Phosphoglycerate kinase. |
| | 6 | Sandy Brown | PF3D7_0923900 | 36329.Q8I2R8 | Polyadenylate-binding protein 2, putative. |
| | 6 | Sandy Brown | PF3D7_0933000 | 36329.Q8I0W1 | CSTF domain-containing protein, putative. |
| | 6 | Sandy Brown | PF3D7_1009200 | 36329.Q8IJV3 | Ribonuclease, putative. |
| | 6 | Sandy Brown | PF3D7_1120100 | 36329.Q8IIG6 | Phosphoglycerate mutase. |
| | 6 | Sandy Brown | PF3D7_1210400 | 36329.Q8I5U5 | General transcription factor 3C polypeptide 5, putative. |
| | 6 | Sandy Brown | PF3D7_1414400 | 36329.Q8ILV1 | Serine/threonine-protein phosphatase. |
| | 6 | Sandy Brown | PF3D7_1419900 | 36329.Q8ILQ1 | Cleavage and polyadenylation specificity factor subunit 4, putative. |
| | 6 | Sandy Brown | PF3D7_1438500 | 36329.Q8IL83 | Cleavage and polyadenylation specificity factor subunit 3, putative. |
| | 6 | Sandy Brown | PF3D7_1444800 | 36329.Q7KQL9 | Fructose-bisphosphate aldolase. |

| Prefoldin and T-complex | Cluster number | Cluster color | Gene ID | Protein identifier | Protein description |
|---|---|---|---|---|---|
| | 8 | Yellow | PF3D7_0214000 | 36329.O96220 | T-complex protein 1 subunit theta. |
| | 8 | Yellow | PF3D7_0306800 | 36329.O97247 | T-complex protein 1 subunit beta. |
| | 8 | Yellow | PF3D7_0308200 | 36329.O77323 | T-complex protein 1 subunit eta; Molecular chaperone; assists the folding of proteins upon ATP hydrolysis. Known to play a role, in vitro, in the folding of actin and tubulin (By similarity). |
| | 8 | Yellow | PF3D7_0320300 | 36329.O97282 | T-complex protein 1 subunit epsilon. |
| | 8 | Yellow | PF3D7_0718500 | 36329.Q8IBR6 | Prefoldin subunit 3; Binds specifically to cytosolic chaperonin (c-CPN) and transfers target proteins to it. Binds to nascent polypeptide chain and promotes folding in an environment in which there are many competing pathways for nonnative proteins; Belongs to the prefoldin subunit alpha family. |
| | 8 | Yellow | PF3D7_0904500 | 36329.Q8I3A4 | Prefoldin subunit 4; Binds specifically to cytosolic chaperonin (c-CPN) and transfers target proteins to it. Binds to nascent polypeptide chain and promotes folding in an environment in which there are many competing pathways for nonnative proteins; Belongs to the prefoldin subunit beta family. |
| | 8 | Yellow | PF3D7_1128100 | 36329.Q8II82 | Prefoldin subunit 5, putative. |
| | 8 | Yellow | PF3D7_1132200 | 36329.Q8II43 | T-complex protein 1 subunit alpha. |
| | 8 | Yellow | PF3D7_1229500 | 36329.Q8I5C4 | T-complex protein 1 subunit gamma. |
| | 8 | Yellow | PF3D7_1357800 | 36329.C0H5I7 | T-complex protein 1 subunit delta. |
| | 8 | Yellow | PF3D7_1416900 | 36329.Q8ILS7 | Prefoldin subunit 2, putative. |

| Replication factor C complex | Cluster number | Cluster color | Gene ID | Protein identifie | Protein description |
|---|---|---|---|---|---|
| | 10 | Khaki | PF3D7_0111300 | 36329.Q8I233 | P-loop containing nucleoside triphosphate hydrolase, putative. |
| | 10 | Khaki | PF3D7_0218000 | 36329.O96260 | Replication factor C subunit 2, putative. |
| | 10 | Khaki | PF3D7_0219600 | 36329.O96271 | Replication factor C subunit 1. |
| | 10 | Khaki | PF3D7_0408500 | 36329.Q7K734 | FEN1 Flap endonuclease 1; Structure-specific nuclease with 5'-flap endonuclease and 5'- 3' exonuclease activities involved in DNA replication and repair. During DNA replication, cleaves the 5'-overhanging flap structure that is generated by displacement synthesis when DNA polymerase encounters the 5'-end of a downstream Okazaki fragment. It enters the flap from the 5'-end and then tracks to cleave the flap base, leaving a nick for ligation. Also involved in the long patch base excision repair (LP-BER) pathway, by cleaving within the apurinic/apyrimidinic (AP) site- terminated flap. Acts as [...] |
| | 10 | Khaki | PF3D7_1111100 | 36329.Q8IIQ1 | Replication factor C subunit 5, putative. |
| | 10 | Khaki | PF3D7_1241700 | 36329.Q8I512 | Replication factor C subunit 4, putative. |
| | 10 | Khaki | PF3D7_1361900 | 36329.P61074 | PCNA1 Proliferating cell nuclear antigen; This protein is an auxiliary protein of DNA polymerase delta and is involved in the control of eukaryotic DNA replication by increasing the polymerase's processibility during elongation of the leading strand. |
| | 10 | Khaki | PF3D7_1463200 | 36329.Q8IKK4 | Replication factor C subunit 3, putative. |

| Export-Import | Cluster number | Cluster color | Gene ID | Protein identifier | Protein description |
|---|---|---|---|---|---|
| | 9 | Dark Golden Rod 2 | PF3D7_0302900 | 36329.O77312 | Exportin-1, putative. |
| | 9 | Dark Golden Rod 2 | PF3D7_0524000 | 36329.Q8I3M5 | Karyopherin beta. |
| | 9 | Dark Golden Rod 2 | PF3D7_0623100 | 36329.C6KT67 | Nuclear polyadenylated RNA-binding protein NAB2, putative. |
| | 9 | Dark Golden Rod 2 | PF3D7_0627700 | 36329.C6KTB3 | Transportin. |
| | 9 | Dark Golden Rod 2 | PF3D7_0812400 | 36329.Q8IAW0 | Importin subunit alpha; Belongs to the importin alpha family. |
| | 9 | Dark Golden Rod 2 | PF3D7_0815200 | 36329.Q8IAY9 | Importin subunit beta, putative. |
| | 9 | Dark Golden Rod 2 | PF3D7_0932800 | 36329.Q8I2I8 | Importin alpha re-exporter, putative. |
| | 9 | Dark Golden Rod 2 | PF3D7_1117700 | 36329.Q7KQK6 | GTP-binding nuclear protein; GTP-binding protein involved in nucleocytoplasmic transport. Required for the import of protein into the nucleus and also for RNA export. Involved in chromatin condensation and control of cell cycle. Belongs to the small GTPase superfamily. Ran family. |
| | 9 | Dark Golden Rod 2 | PF3D7_1315900 | 36329.A0A5K1K8Q1 | Exportin-T; tRNA nucleus export receptor which facilitates tRNA translocation across the nuclear pore complex. Belongs to the exportin family. |

| DNA repair | Cluster number | Cluster color | Gene ID | Protein identifier | Protein description |
|---|---|---|---|---|---|
| | 11 | Green | PF3D7_0107800 | 36329.A0A143ZUM0 | Double-strand break repair protein MRE11. |
| | 11 | Green | PF3D7_0505500 | 36329.Q8I447 | DNA mismatch repair protein; Component of the post-replicative DNA mismatch repair system (MMR). |
| | 11 | Green | PF3D7_0605800 | 36329.C6KSQ6 | Probable DNA repair protein RAD50; Essential component of the MRN complex, a complex that possesses single-stranded DNA endonuclease and 3' to 5' exonuclease activities, and plays a central role in double-strand break (DSB) repair, chromosome morphogenesis, DNA repair and meiosis. In the complex, it mediates the ATP-binding and is probably required to bind DNA ends and hold them in close proximity (By similarity). Belongs to the SMC family. RAD50 subfamily. |
| | 11 | Green | PF3D7_0725000 | 36329.Q8IBK1 | Exonuclease I, putative. |
| | 11 | Green | PF3D7_0726300 | 36329.Q8IBJ3 | DNA mismatch repair protein PMS1, putative. |
| | 11 | Green | PF3D7_1117800 | 36329.Q8IIJ0 | DNA mismatch repair protein MLH. |
| | 11 | Green | PF3D7_1408400 | 36329.Q8IM12 | FANCJ-like helicase, putative. |
| | 11 | Green | PF3D7_1427500 | 36329.Q8ILI9 | DNA mismatch repair protein MSH2, putative. |

| Chaperone complex | Cluster number | Cluster color | Gene ID | Protein identifier | Protein description |
|---|---|---|---|---|---|
| | 12 | Medium Aqua Marine | PF3D7_0708800 | 36329.Q8IC01 | Heat shock protein 110. |
| | 12 | Medium Aqua Marine | PF3D7_0818900 | 36329.Q8IB24 | Heat shock protein 70; Belongs to the heat shock protein 70 family. |
| | 12 | Medium Aqua Marine | PF3D7_1037500 | 36329.Q8IJ32 | Dynamin-like protein; Belongs to the TRAFAC class dynamin-like GTPase superfamily. Dynamin/Fzo/YdjA family. |
| | 12 | Medium Aqua Marine | PF3D7_1145400 | 36329.Q8IHR4 | Dynamin-like protein; Belongs to the TRAFAC class dynamin-like GTPase superfamily. Dynamin/Fzo/YdjA family. |
| | 12 | Medium Aqua Marine | PF3D7_1211400 | 36329.Q7KQK3 | Heat shock protein DNAJ homologue Pfj4. |
| | 12 | Medium Aqua Marine | PF3D7_1216900 | 36329.Q8I5N9 | DNA-binding chaperone, putative. |
| | 12 | Medium Aqua Marine | PF3D7_1247500 | 36329.Q8I4V7 | Serine/threonine protein kinase, putative. |
| | 12 | Medium Aqua Marine | PF3D7_1437900 | 36329.Q8IL88 | HSP40, subfamily A. |

| Ribonucleotide reductase complex | Cluster number | Cluster color | Gene ID | Protein identifier | Protein description |
|---|---|---|---|---|---|
| | 13 | Aquamarine 4 | PF3D7_0206700 | 36329.Q7KWJ4 | Adenylosuccinate lyase; Belongs to the lyase 1 family. Adenylosuccinate lyase subfamily. |
| | 13 | Aquamarine 4 | PF3D7_0920800 | 36329.Q8I2U5 | Inosine-5'-monophosphate dehydrogenase; Catalyzes the conversion of inosine 5'-phosphate (IMP) to xanthosine 5'-phosphate (XMP), the first committed and rate-limiting step in the de novo synthesis of guanine nucleotides, and therefore plays an important role in the regulation of cell growth. |
| | 13 | Aquamarine 4 | PF3D7_1008900 | 36329.Q8IJV6 | Adenylate kinase; Belongs to the adenylate kinase family. |
| | 13 | Aquamarine 4 | PF3D7_1012400 | 36329.Q8IJS1 | Hypoxanthine phosphoribosyltransferase; Belongs to the purine/pyrimidine phosphoribosyltransferase family. |
| | 13 | Aquamarine 4 | PF3D7_1015800 | 36329.Q8IJN8 | Ribonucleoside-diphosphate reductase small chain, putative. |
| | 13 | Aquamarine 4 | PF3D7_1029600 | 36329.Q8IJA9 | Adenosine deaminase. |
| | 13 | Aquamarine 4 | PF3D7_1405600 | 36329.Q8IM38 | Ribonucleoside-diphosphate reductase small chain, putative. |
| | 13 | Aquamarine 4 | PF3D7_1437200 | 36329.Q8IL94 | Ribonucleoside-diphosphate reductase; Provides the precursors necessary for DNA synthesis. Catalyzes the biosynthesis of deoxyribonucleotides from the corresponding ribonucleotides. |

| Chromatin condensation | Cluster number | Cluster color | Gene ID | Protein identifier | Protein description |
|---|---|---|---|---|---|
| | 14 | Aquamarine 2 | PF3D7_1318400 | 36329.Q8IED2 | Structural maintenance of chromosomes protein 2; May play a role in the conversion of interphase chromatin into condensed chromosomes; Belongs to the SMC family. SMC2 subfamily. |
| | 14 | Aquamarine 2 | PF3D7_0509100 | 36329.Q8I413 | Structural maintenance of chromosomes protein 4, putative. |
| | 14 | Aquamarine 2 | PF3D7_1130700 | 36329.Q8II57 | Structural maintenance of chromosomes protein 1, putative. |
| | 14 | Aquamarine 2 | PF3D7_1304000 | 36329.C0H598 | Condensin complex subunit 2, putative. |
| | 14 | Aquamarine 2 | PF3D7_1403100 | 36329.C6S3H2 | Condensin complex subunit 1, putative. |
| | 14 | Aquamarine 2 | PF3D7_1456500 | 36329.Q8IKR4 | SCD domain-containing protein. |
| | 14 | Aquamarine 2 | PF3D7_0414000 | 36329.Q8I1U7 | Structural maintenance of chromosomes protein 3 homolog; Central component of cohesin, a complex required for chromosome cohesion during the cell cycle. The cohesin complex may form a large proteinaceous ring within which sister chromatids can be trapped. At anaphase, the complex is cleaved and dissociates from chromatin, allowing sister chromatids to segregate. Cohesion is coupled to DNA replication and is involved in DNA repair. The cohesin complex plays also an important role in spindle pole assembly during mitosis and in chromosomes movement (By similarity); Belongs to the SMC family |

| CCR-NOT complex | Cluster number | Cluster color | Gene ID | Protein identifier | Protein description |
|---|---|---|---|---|---|
| | 15 | Cyan | PF3D7_0807100 | 36329.Q8IAR1 | DNA helicase PSH3. |
| | 15 | Cyan | PF3D7_0811300 | 36329.C0H4T9 | CCR4-associated factor 1. |
| kmeans | 15 | Cyan | PF3D7_1014600 | 36329.Q8IJP9 | Transcriptional coactivator ADA2. |
| kmeans | 15 | Cyan | PF3D7_1103800 | 36329.Q8IIW4 | CCR4-NOT transcription complex subunit 1, putative. |
| kmeans | 15 | Cyan | PF3D7_1235300 | 36329.Q8I569 | CCR4-NOT transcription complex subunit 4, putative. |
| kmeans | 15 | Cyan | PF3D7_1308900 | 36329.Q8IEM5 | mRNA-decapping enzyme 2, putative. |
| kmeans | 15 | Cyan | PF3D7_1417200 | 36329.Q8ILS4 | NOT family protein, putative. |

| #Clustering method | Cluster number | Cluster color | Gene count | Gene ID | Protein identifier | Protein description |
|---|---|---|---|---|---|---|
| kmeans | 1 | Red | 138 | MAL3P3.19 | 36329.O00806 | 60S acidic ribosomal protein P2; Plays an important role in the elongation step of protein synthesis. |
| kmeans | 1 | Red | 138 | MAL3P7.35 | 36329.O97313 | 40S ribosomal protein S3a; Belongs to the eukaryotic ribosomal protein eS1 family. |
| kmeans | 1 | Red | 138 | PF10_0264 | 36329.Q8IJD4 | 40S ribosomal protein SA; Required for the assembly and/or stability of the 40S ribosomal subunit. Required for the processing of the 20S rRNA- precursor to mature 18S rRNA in a late step of the maturation of 40S ribosomal subunits. |
| kmeans | 1 | Red | 138 | PF3D7_0102900 | 36329.Q8I2B1 | Aspartate--tRNA ligase. |
| kmeans | 1 | Red | 138 | PF3D7_0109800 | 36329.Q8I246 | Phenylalanine--tRNA ligase alpha subunit. |
| kmeans | 1 | Red | 138 | PF3D7_0209200 | 36329.O96177 | Exosome complex component MTR3, putative. |
| kmeans | 1 | Red | 138 | PF3D7_0212300 | 36329.O96203 | Peptide chain release factor subunit 1, putative. |
| kmeans | 1 | Red | 138 | PF3D7_0217800 | 36329.O96258 | 40S ribosomal protein S26; Belongs to the eukaryotic ribosomal protein eS26 family. |
| kmeans | 1 | Red | 138 | PF3D7_0306900 | 36329.O97248 | 40S ribosomal protein S23, putative; Belongs to the universal ribosomal protein uS12 family. |
| kmeans | 1 | Red | 138 | PF3D7_0316800 | 36329.O77395 | 40S ribosomal protein S15A, putative; Belongs to the universal ribosomal protein uS8 family. |
| kmeans | 1 | Red | 138 | PF3D7_0317600 | 36329.O77381 | 40S ribosomal protein S11, putative; Belongs to the universal ribosomal protein uS17 family. |
| kmeans | 1 | Red | 138 | PF3D7_0410400 | 36329.Q9U0I4 | Exosome complex component RRP4. |
| kmeans | 1 | Red | 138 | PF3D7_0415900 | 36329.C0H4A6 | Ribosomal protein L15; Belongs to the eukaryotic ribosomal protein eL15 family. |
| kmeans | 1 | Red | 138 | PF3D7_0418200 | 36329.Q8I1Q5 | Eukaryotic translation initiation factor 3 subunit M; Component of the eukaryotic translation initiation factor 3 (eIF-3) complex, which is involved in protein synthesis of a specialized repertoire of mRNAs and, together with other initiation factors, stimulates binding of mRNA and methionyl-tRNAi to the 40S ribosome. The eIF-3 complex specifically targets and initiates translation of a subset of mRNAs involved in cell proliferation. |
| kmeans | 1 | Red | 138 | PF3D7_0422400 | 36329.Q8IFP2 | 40S ribosomal protein S19. |
| kmeans | 1 | Red | 138 | PF3D7_0507100 | 36329.Q8I431 | 60S ribosomal protein L4. |
| kmeans | 1 | Red | 138 | PF3D7_0507700 | 36329.Q8I426 | Nuclear protein localization protein 4, putative. |
| kmeans | 1 | Red | 138 | PF3D7_0516200 | 36329.A0A5K1K925 | 40S ribosomal protein S11; Belongs to the universal ribosomal protein uS11 family. |
| kmeans | 1 | Red | 138 | PF3D7_0516900 | 36329.Q8I3T9 | 60S ribosomal protein L2. |
| kmeans | 1 | Red | 138 | PF3D7_0517000 | 36329.Q8I3T8 | 60S ribosomal protein L12, putative; Belongs to the universal ribosomal protein uL11 family. |
| kmeans | 1 | Red | 138 | PF3D7_0517700 | 36329.Q8I3T1 | Eukaryotic translation initiation factor 3 subunit B; Component of the eukaryotic translation initiation factor 3 (eIF-3) complex, which is involved in protein synthesis and, together with other initiation factors, stimulates binding of mRNA and methionyl-tRNAi to the 40S ribosome; Belongs to the eIF-3 subunit B family. |
| kmeans | 1 | Red | 138 | PF3D7_0519400 | 36329.Q8I3R6 | 40S ribosomal protein S24; Belongs to the eukaryotic ribosomal protein eS24 family. |
| kmeans | 1 | Red | 138 | PF3D7_0520000 | 36329.Q8I3R0 | 40S ribosomal protein S9, putative; Belongs to the universal ribosomal protein uS4 family. |
| kmeans | 1 | Red | 138 | PF3D7_0528200 | 36329.Q8I3I5 | Eukaryotic translation initiation factor 3 subunit E; Component of the eukaryotic translation initiation factor 3 (eIF-3) complex, which is involved in protein synthesis of a specialized repertoire of mRNAs and, together with other initiation factors, stimulates binding of mRNA and methionyl-tRNAi to the 40S ribosome. The eIF-3 complex specifically targets and initiates translation of a subset of mRNAs involved in cell proliferation. |
| kmeans | 1 | Red | 138 | PF3D7_0602100 | 36329.C6KSM1 | ATP-dependent RNA helicase, putative. |
| kmeans | 1 | Red | 138 | PF3D7_0607000 | 36329.C6KSR8 | Translation initiation factor IF-2, putative. |
| kmeans | 1 | Red | 138 | PF3D7_0612100 | 36329.C6KSW5 | Eukaryotic translation initiation factor 3 subunit L; Component of the eukaryotic translation initiation factor 3 (eIF-3) complex, which is involved in protein synthesis of a specialized repertoire of mRNAs and, together with other initiation factors, stimulates binding of mRNA and methionyl-tRNAi to the 40S ribosome. The eIF-3 complex specifically targets and initiates translation of a subset of mRNAs involved in cell proliferation. |
| kmeans | 1 | Red | 138 | PF3D7_0618300 | 36329.C6KT23 | 60S ribosomal protein L27a, putative; Belongs to the universal ribosomal protein uL15 family. |
| kmeans | 1 | Red | 138 | PF3D7_0619400 | 36329.C6KT34 | Cell division cycle protein 48 homologue, putative. |
| kmeans | 1 | Red | 138 | PF3D7_0621800 | 36329.C6KT55 | Nascent polypeptide-associated complex subunit alpha, putative. |
| kmeans | 1 | Red | 138 | PF3D7_0705700 | 36329.C0H4K8 | 40S ribosomal protein S29, putative. |
| kmeans | 1 | Red | 138 | PF3D7_0706000 | 36329.C0H4L1 | Importin-7, putative. |
| kmeans | 1 | Red | 138 | PF3D7_0716800 | 36329.Q8IBT2 | Eukaryotic translation initiation factor 3 subunit I; Component of the eukaryotic translation initiation factor 3 (eIF-3) complex, which is involved in protein synthesis of a specialized repertoire of mRNAs and, together with other initiation factors, stimulates binding of mRNA and methionyl-tRNAi to the 40S ribosome. The eIF-3 complex specifically targets and initiates translation of a subset of mRNAs involved in cell proliferation. |
| kmeans | 1 | Red | 138 | PF3D7_0719600 | 36329.Q8IBQ6 | 60S ribosomal protein L11a, putative; Belongs to the universal ribosomal protein uL5 family. |
| kmeans | 1 | Red | 138 | PF3D7_0719700 | 36329.Q8IBQ5 | 40S ribosomal protein S10, putative. |
| kmeans | 1 | Red | 138 | PF3D7_0721600 | 36329.Q8IBN5 | 40S ribosomal protein S5, putative; Belongs to the universal ribosomal protein uS7 family. |
| kmeans | 1 | Red | 138 | PF3D7_0726500 | 36329.Q8IBJ1 | Ubiquitin carboxyl-terminal hydrolase, putative; Belongs to the peptidase C19 family. |
| kmeans | 1 | Red | 138 | PF3D7_0728000 | 36329.Q8IBH7 | Eukaryotic translation initiation factor 2 subunit alpha. |
| kmeans | 1 | Red | 138 | PF3D7_0802300 | 36329.Q8IAM3 | Periodic tryptophan protein 2, putative. |
| kmeans | 1 | Red | 138 | PF3D7_0813900 | 36329.Q8IAX5 | 40S ribosomal protein S16, putative; Belongs to the universal ribosomal protein uS9 family. |
| kmeans | 1 | Red | 138 | PF3D7_0815600 | 36329.Q8IAZ3 | Eukaryotic translation initiation factor 3 subunit G; RNA-binding component of the eukaryotic translation initiation factor 3 (eIF-3) complex, which is involved in protein synthesis of a specialized repertoire of mRNAs and, together with other initiation factors, stimulates binding of mRNA and methionyl-tRNAi to the 40S ribosome. The eIF-3 complex specifically targets and initiates translation of a subset of mRNAs involved in cell proliferation. This subunit can bind 18S rRNA. |
| kmeans | 1 | Red | 138 | PF3D7_0826300 | 36329.Q8IB96 | SPRY domain, putative. |
| kmeans | 1 | Red | 138 | PF3D7_0826700 | 36329.Q8IBA0 | Receptor for activated c kinase. |
| kmeans | 1 | Red | 138 | PF3D7_0828500 | 36329.Q8IBB6 | Translation initiation factor eIF-2B subunit alpha, putative; Belongs to the eIF-2B alpha/beta/delta subunits family. |
| kmeans | 1 | Red | 138 | PF3D7_0909900 | 36329.Q8I353 | Helicase SKI2W, putative. |
| kmeans | 1 | Red | 138 | PF3D7_0913200 | 36329.Q8I320 | Elongation factor 1-beta. |
| kmeans | 1 | Red | 138 | PF3D7_0918300 | 36329.Q8I2X0 | Eukaryotic translation initiation factor 3 subunit F, putative; Belongs to the eIF-3 subunit F family. |
| kmeans | 1 | Red | 138 | PF3D7_0921800 | 36329.C0H549 | Ribosome production factor 1, putative. |
| kmeans | 1 | Red | 138 | PF3D7_1003500 | 36329.Q8IK02 | 40S ribosomal protein S20e, putative. |

| | | | | | | |
|---|---|---|---|---|---|---|
| kmeans | 1 | Red | 138 | PF3D7_1004000 | 36329.Q8IJZ7 | 60S ribosomal protein L13, putative. |
| kmeans | 1 | Red | 138 | PF3D7_1004200 | 36329.Q8IJZ5 | WD repeat-containing protein, putative. |
| kmeans | 1 | Red | 138 | PF3D7_1007900 | 36329.Q8IJW4 | Eukaryotic translation initiation factor 3 subunit D; mRNA cap-binding component of the eukaryotic translation initiation factor 3 (eIF-3) complex, which is involved in protein synthesis of a specialized repertoire of mRNAs and, together with other initiation factors, stimulates binding of mRNA and methionyl-tRNAi to the 40S ribosome. The eIF-3 complex specifically targets and initiates translation of a subset of mRNAs involved in cell proliferation. In the eIF-3 complex, eif3d specifically recognizes and binds the 7- methylguanosine cap of a subset of mRNAs. |
| kmeans | 1 | Red | 138 | PF3D7_1008800 | 36329.Q8IJV7 | Nucleolar protein 5, putative. |
| kmeans | 1 | Red | 138 | PF3D7_1010600 | 36329.Q8IJT9 | Eukaryotic translation initiation factor 2 subunit beta. |
| kmeans | 1 | Red | 138 | PF3D7_1013100 | 36329.Q8IJR4 | U3 small nucleolar RNA-associated protein 13, putative. |
| kmeans | 1 | Red | 138 | PF3D7_1019400 | 36329.Q8IJK8 | 60S ribosomal protein L30e, putative. |
| kmeans | 1 | Red | 138 | PF3D7_1020700 | 36329.Q8IJJ5 | RNA cytidine acetyltransferase; RNA cytidine acetyltransferase with specificity toward both 18S rRNA and tRNAs. Catalyzes the formation of N(4)-acetylcytidine (ac4C) in 18S rRNA. Required for early nucleolar cleavages of precursor rRNA at sites A0, A1 and A2 during 18S rRNA synthesis. Catalyzes the formation of ac4C in serine and leucine tRNAs. Requires a tRNA-binding adapter protein for full tRNA acetyltransferase activity but not for 18S rRNA acetylation. |
| kmeans | 1 | Red | 138 | PF3D7_1027800 | 36329.Q8IJC6 | 60S ribosomal protein L3; Belongs to the universal ribosomal protein uL3 family. |
| kmeans | 1 | Red | 138 | PF3D7_1103100 | 36329.Q8IIX0 | 60S acidic ribosomal protein P1, putative; Belongs to the eukaryotic ribosomal protein P1/P2 family. |
| kmeans | 1 | Red | 138 | PF3D7_1104000 | 36329.Q8IIW2 | Phenylalanine--tRNA ligase beta subunit. |
| kmeans | 1 | Red | 138 | PF3D7_1105400 | 36329.Q8IIU8 | 40S ribosomal protein S4; Belongs to the eukaryotic ribosomal protein eS4 family. |
| kmeans | 1 | Red | 138 | PF3D7_1118500 | 36329.Q8III3 | Nucleolar protein 56, putative. |
| kmeans | 1 | Red | 138 | PF3D7_1121700 | 36329.Q8IIE9 | Protein GCN20. |
| kmeans | 1 | Red | 138 | PF3D7_1123400 | 36329.Q8IIC9 | Translation elongation factor EF-1, subunit alpha, putative. |
| kmeans | 1 | Red | 138 | PF3D7_1123900 | 36329.Q8IIC4 | Ribonucleoprotein; Common component of the spliceosome and rRNA processing machinery; Belongs to the eukaryotic ribosomal protein eL8 family. |
| kmeans | 1 | Red | 138 | PF3D7_1124800 | 36329.Q8IIB5 | Ribosome biogenesis regulatory protein; Involved in ribosomal large subunit assembly. Belongs to the RRS1 family. |
| kmeans | 1 | Red | 138 | PF3D7_1126200 | 36329.Q8IIA2 | 40S ribosomal protein S18, putative; Belongs to the universal ribosomal protein uS13 family. |
| kmeans | 1 | Red | 138 | PF3D7_1129400 | 36329.Q8II69 | rRNA (Cytosine-C(5))-methyltransferase, putative; Belongs to the class I-like SAM-binding methyltransferase superfamily. RsmB/NOP family. |
| kmeans | 1 | Red | 138 | PF3D7_1130100 | 36329.Q8II62 | 60S ribosomal protein L38; Belongs to the eukaryotic ribosomal protein eL38 family. |
| kmeans | 1 | Red | 138 | PF3D7_1130200 | 36329.Q8II61 | 60S acidic ribosomal protein P0; Ribosomal protein P0 is the functional equivalent of E.coli protein L10. |
| kmeans | 1 | Red | 138 | PF3D7_1142500 | 36329.Q8IHU0 | 60S ribosomal protein L28. |
| kmeans | 1 | Red | 138 | PF3D7_1144000 | 36329.Q8IHS5 | 40S ribosomal protein S21; Belongs to the eukaryotic ribosomal protein eS21 family. |
| kmeans | 1 | Red | 138 | PF3D7_1203900 | 36329.Q8I607 | Ubiquitin-conjugating enzyme E2; Belongs to the ubiquitin-conjugating enzyme family. |
| kmeans | 1 | Red | 138 | PF3D7_1206200 | 36329.Q8I5Y3 | Eukaryotic translation initiation factor 3 subunit C; Component of the eukaryotic translation initiation factor 3 (eIF-3) complex, which is involved in protein synthesis of a specialized repertoire of mRNAs and, together with other initiation factors, stimulates binding of mRNA and methionyl-tRNAi to the 40S ribosome. The eIF-3 complex specifically targets and initiates translation of a subset of mRNAs involved in cell proliferation. |
| kmeans | 1 | Red | 138 | PF3D7_1207100 | 36329.Q8I5X4 | Small subunit rRNA processing factor, putative. |
| kmeans | 1 | Red | 138 | PF3D7_1208100 | 36329.Q8I5W6 | Uncharacterized protein. |
| kmeans | 1 | Red | 138 | PF3D7_1211800 | 36329.Q7KQK2 | Polyubiquitin. |
| kmeans | 1 | Red | 138 | PF3D7_1212700 | 36329.Q8I5S6 | Eukaryotic translation initiation factor 3 subunit A; RNA-binding component of the eukaryotic translation initiation factor 3 (eIF-3) complex, which is involved in protein synthesis of a specialized repertoire of mRNAs and, together with other initiation factors, stimulates binding of mRNA and methionyl-tRNAi to the 40S ribosome. The eIF-3 complex specifically targets and initiates translation of a subset of mRNAs involved in cell proliferation. |
| kmeans | 1 | Red | 138 | PF3D7_1213900 | 36329.A0A144A0U7 | W2 domain-containing protein. |
| kmeans | 1 | Red | 138 | PF3D7_1225800 | 36329.Q8I5F9 | Ubiquitin-activating enzyme E1. |
| kmeans | 1 | Red | 138 | PF3D7_1226800 | 36329.Q8I5F0 | Ataxin-3, putative. |
| kmeans | 1 | Red | 138 | PF3D7_1242700 | 36329.Q8I502 | 40S ribosomal protein S17, putative. |
| kmeans | 1 | Red | 138 | PF3D7_1250600 | 36329.Q8I4S8 | Translation initiation factor eIF-2B subunit beta, putative; Belongs to the eIF-2B alpha/beta/delta subunits family. |
| kmeans | 1 | Red | 138 | PF3D7_1302800 | 36329.Q8IET7 | 40S ribosomal protein S7; Belongs to the eukaryotic ribosomal protein eS7 family. |
| kmeans | 1 | Red | 138 | PF3D7_1305300 | 36329.C0H5A0 | Translational activator GCN1, putative. |
| kmeans | 1 | Red | 138 | PF3D7_1307000 | 36329.Q8IEP5 | Exosome complex component RRP40, putative. |
| kmeans | 1 | Red | 138 | PF3D7_1307100 | 36329.Q8IEP4 | U3 small nucleolar RNA-associated protein 6, putative. |
| kmeans | 1 | Red | 138 | PF3D7_1308300 | 36329.Q8IEN2 | 40S ribosomal protein S27. |
| kmeans | 1 | Red | 138 | PF3D7_1309100 | 36329.Q8IEM3 | 60S ribosomal protein L24, putative. |
| kmeans | 1 | Red | 138 | PF3D7_1317800 | 36329.C0H5C2 | 40S ribosomal protein S19; Belongs to the universal ribosomal protein uS19 family. |
| kmeans | 1 | Red | 138 | PF3D7_1323100 | 36329.Q8IE85 | 60S ribosomal protein L6, putative. |
| kmeans | 1 | Red | 138 | PF3D7_1323400 | 36329.Q8IE82 | 60S ribosomal protein L23; Belongs to the universal ribosomal protein uL23 family. |
| kmeans | 1 | Red | 138 | PF3D7_1326400 | 36329.A0A5K1K977 | Translation initiation factor eIF-2B subunit gamma, putative. |
| kmeans | 1 | Red | 138 | PF3D7_1330600 | 36329.Q8IE20 | Elongation factor Tu. |
| kmeans | 1 | Red | 138 | PF3D7_1331700 | 36329.Q8IE10 | Glutamine--tRNA ligase, putative; Belongs to the class-I aminoacyl-tRNA synthetase family. |
| kmeans | 1 | Red | 138 | PF3D7_1331800 | 36329.Q8IE09 | 60S ribosomal protein L23, putative; Belongs to the universal ribosomal protein uL14 family. |
| kmeans | 1 | Red | 138 | PF3D7_1332900 | 36329.Q8IDZ9 | Isoleucine--tRNA ligase, putative; Belongs to the class-I aminoacyl-tRNA synthetase family. |
| kmeans | 1 | Red | 138 | PF3D7_1333600 | 36329.Q8IDZ2 | U3 small nucleolar RNA-associated protein 4, putative. |
| kmeans | 1 | Red | 138 | PF3D7_1338200 | 36329.A0A5K1K8V8 | 60S ribosomal protein L6, putative. |
| kmeans | 1 | Red | 138 | PF3D7_1338300 | 36329.A0A5K1K967 | Elongation factor 1-gamma, putative. |
| kmeans | 1 | Red | 138 | PF3D7_1340100 | 36329.A0A5K1K8M9 | Exosome complex component RRP42, putative. |
| kmeans | 1 | Red | 138 | PF3D7_1341200 | 36329.Q8IDS6 | 60S ribosomal protein L18a; Belongs to the eukaryotic ribosomal protein eL20 family. |

| | | | | | | |
|---|---|---|---|---|---|---|
| kmeans | 1 | Red | 138 | PF3D7_1342000 | 36329.Q8IDR9 | 40S ribosomal protein S6. |
| kmeans | 1 | Red | 138 | PF3D7_1349200 | 36329.Q8IDK7 | Glutamate--tRNA ligase, putative; Belongs to the class-I aminoacyl-tRNA synthetase family. |
| kmeans | 1 | Red | 138 | PF3D7_1357000 | 36329.Q8I0P6 | Elongation factor 1-alpha; This protein promotes the GTP-dependent binding of aminoacyl- tRNA to the A-site of ribosomes during protein biosynthesis. |
| kmeans | 1 | Red | 138 | PF3D7_1357700 | 36329.A0A5K1K903 | U3 small nucleolar RNA-associated protein 21, putative. |
| kmeans | 1 | Red | 138 | PF3D7_1358800 | 36329.Q8IDB0 | 40S ribosomal protein S15; Belongs to the universal ribosomal protein uS15 family. |
| kmeans | 1 | Red | 138 | PF3D7_1364500 | 36329.Q8ID62 | Exosome complex component RRP45, putative. |
| kmeans | 1 | Red | 138 | PF3D7_1365900 | 36329.Q8ID50 | Ubiquitin-60S ribosomal protein L40. |
| kmeans | 1 | Red | 138 | PF3D7_1368200 | 36329.Q8I6Z4 | ABC transporter E family member 1, putative. |
| kmeans | 1 | Red | 138 | PF3D7_1368400 | 36329.Q8ID26 | Ribosomal protein L1, putative. |
| kmeans | 1 | Red | 138 | PF3D7_1402500 | 36329.Q8IM64 | Ribosomal protein S27a, putative. |
| kmeans | 1 | Red | 138 | PF3D7_1407100 | 36329.Q8IM23 | rRNA 2'-O-methyltransferase fibrillarin, putative. |
| kmeans | 1 | Red | 138 | PF3D7_1408600 | 36329.Q8IM10 | 40S ribosomal protein S8; Belongs to the eukaryotic ribosomal protein eS8 family. |
| kmeans | 1 | Red | 138 | PF3D7_1410600 | 36329.Q8ILY9 | Eukaryotic translation initiation factor 2 subunit gamma, putative. |
| kmeans | 1 | Red | 138 | PF3D7_1417500 | 36329.Q8ILS0 | H/ACA ribonucleoprotein complex subunit 4, putative. |
| kmeans | 1 | Red | 138 | PF3D7_1418000 | 36329.Q8ILR6 | Ubiquitin fusion degradation protein 1, putative. |
| kmeans | 1 | Red | 138 | PF3D7_1419700 | 36329.Q8ILQ3 | Uncharacterized protein. |
| kmeans | 1 | Red | 138 | PF3D7_1421200 | 36329.Q8ILN8 | 40S ribosomal protein S25; Belongs to the eukaryotic ribosomal protein eS25 family. |
| kmeans | 1 | Red | 138 | PF3D7_1422400 | 36329.Q8ILM9 | Uncharacterized protein. |
| kmeans | 1 | Red | 138 | PF3D7_1424100 | 36329.Q8ILL3 | 60S ribosomal protein L5, putative. |
| kmeans | 1 | Red | 138 | PF3D7_1426100 | 36329.Q8ILK2 | Nascent polypeptide-associated complex subunit beta. |
| kmeans | 1 | Red | 138 | PF3D7_1426200 | 36329.Q8ILK1 | Protein arginine N-methyltransferase 1; Belongs to the class I-like SAM-binding methyltransferase superfamily. Protein arginine N-methyltransferase family. |
| kmeans | 1 | Red | 138 | PF3D7_1441200 | 36329.Q8IL58 | 60S ribosomal protein L1, putative. |
| kmeans | 1 | Red | 138 | PF3D7_1442300 | 36329.Q8IL48 | tRNA import protein tRIP. |
| kmeans | 1 | Red | 138 | PF3D7_1447000 | 36329.Q8IL02 | 40S ribosomal protein S5; Belongs to the universal ribosomal protein uS5 family. |
| kmeans | 1 | Red | 138 | PF3D7_1448000 | 36329.Q8IKZ5 | U3 small nucleolar RNA-associated protein 12, putative. |
| kmeans | 1 | Red | 138 | PF3D7_1451100 | 36329.Q8IKW5 | Elongation factor 2. |
| kmeans | 1 | Red | 138 | PF3D7_1457300 | 36329.Q8IKQ7 | MA3 domain-containing protein, putative. |
| kmeans | 1 | Red | 138 | PF3D7_1460300 | 36329.C6S3J6 | 60S ribosomal protein L29; Belongs to the eukaryotic ribosomal protein eL29 family. |
| kmeans | 1 | Red | 138 | PF3D7_1460700 | 36329.Q8IKM5 | 60S ribosomal protein L27. |
| kmeans | 1 | Red | 138 | PF3D7_1461300 | 36329.Q8IKL9 | 40S ribosomal protein S28e, putative. |
| kmeans | 1 | Red | 138 | PF3D7_1465900 | 36329.Q8IKH8 | 40S ribosomal protein S3. |
| kmeans | 1 | Red | 138 | PF3D7_1468700 | 36329.Q8IKF0 | Eukaryotic initiation factor 4A; Belongs to the DEAD box helicase family. |
| kmeans | 1 | Red | 138 | RPL44 | 36329.O97231 | 60S ribosomal protein L44; Belongs to the eukaryotic ribosomal protein eL42 family. |
| kmeans | 1 | Red | 138 | RPS12 | 36329.O97249 | 40S ribosomal protein S12; Belongs to the eukaryotic ribosomal protein eS12 family. |
| kmeans | 1 | Red | 138 | RPS30 | 36329.O96269 | 40S ribosomal protein S30; Belongs to the eukaryotic ribosomal protein eS30 family. |
| kmeans | 1 | Red | 138 | proRS | 36329.Q8I5R7 | Proline--tRNA ligase; Catalyzes the attachment of proline to tRNA(Pro) in a two- step reaction: proline is first activated by ATP to form Pro-AMP and then transferred to the acceptor end of tRNA(Pro) (By similarity). Functions in trans to edit the amino acid moiety from incorrectly charged Ala-tRNA(Pro). Has no activity on correctly charged Pro-tRNA(Pro) or Ala-tRNA(Ala). |
| kmeans | 2 | Salmon | 65 | LSM3 | 36329.C0H4W2 | U6 snRNA-associated Sm-like protein LSm3; Binds specifically to the 3'-terminal U-tract of U6 snRNA. |
| kmeans | 2 | Salmon | 65 | LSM4 | 36329.Q8IIT3 | U6 snRNA-associated Sm-like protein LSm4; Binds specifically to the 3'-terminal U-tract of U6 snRNA. |
| kmeans | 2 | Salmon | 65 | PF08_0084 | 36329.Q8IAW3 | U1 small nuclear ribonucleoprotein C; Component of the spliceosomal U1 snRNP, which is essential for recognition of the pre-mRNA 5' splice-site and the subsequent assembly of the spliceosome. U1-C is directly involved in initial 5' splice-site recognition for both constitutive and regulated alternative splicing. The interaction with the 5' splice-site seems to precede base-pairing between the pre-mRNA and the U1 snRNA. Stimulates commitment or early (E) complex formation by stabilizing the base pairing of the 5' end of the U1 snRNA and the 5' splice-site region. |
| kmeans | 2 | Salmon | 65 | PF3D7_0209800 | 36329.Q9TY94 | ATP-dependent RNA helicase UAP56. |
| kmeans | 2 | Salmon | 65 | PF3D7_0218500 | 36329.O96265 | Small nuclear ribonucleoprotein Sm D2. |
| kmeans | 2 | Salmon | 65 | PF3D7_0218700 | 36329.O96267 | Pre-mRNA-processing protein 45, putative. |
| kmeans | 2 | Salmon | 65 | PF3D7_0302000 | 36329.O97334 | Pre-mRNA-splicing factor PRP46, putative. |
| kmeans | 2 | Salmon | 65 | PF3D7_0302100 | 36329.O77306 | Serine/threonine protein kinase. |
| kmeans | 2 | Salmon | 65 | PF3D7_0308600 | 36329.O77325 | Pre-mRNA-processing factor 19, putative. |
| kmeans | 2 | Salmon | 65 | PF3D7_0319500 | 36329.O97318 | RNA-binding protein, putative. |
| kmeans | 2 | Salmon | 65 | PF3D7_0320800 | 36329.O97285 | ATP-dependent RNA helicase DDX6; Belongs to the DEAD box helicase family. |
| kmeans | 2 | Salmon | 65 | PF3D7_0321600 | 36329.O97290 | ATP-dependent RNA helicase DDX42, putative. |
| kmeans | 2 | Salmon | 65 | PF3D7_0323700 | 36329.O97303 | U4/U6.U5 tri-snRNP-associated protein 1, putative. |
| kmeans | 2 | Salmon | 65 | PF3D7_0403700 | 36329.Q8I1Z2 | Pre-mRNA-splicing factor CLF1, putative. |
| kmeans | 2 | Salmon | 65 | PF3D7_0405400 | 36329.Q8I1X5 | Pre-mRNA-processing-splicing factor 8, putative. |
| kmeans | 2 | Salmon | 65 | PF3D7_0422500 | 36329.Q8IFP1 | Pre-mRNA-splicing helicase BRR2, putative. |
| kmeans | 2 | Salmon | 65 | PF3D7_0422700 | 36329.Q8IFN9 | Eukaryotic initiation factor 4A-III, putative; Belongs to the DEAD box helicase family. |
| kmeans | 2 | Salmon | 65 | PF3D7_0520300 | 36329.Q8I3Q7 | U6 snRNA-associated Sm-like protein LSm2; Binds specifically to the 3'-terminal U-tract of U6 snRNA. |
| kmeans | 2 | Salmon | 65 | PF3D7_0522800 | 36329.Q8I3N5 | Pre-mRNA-splicing factor BUD31, putative. |
| kmeans | 2 | Salmon | 65 | PF3D7_0528700 | 36329.Q8I3I0 | Peptidyl-prolyl cis-trans isomerase; PPIases accelerate the folding of proteins. It catalyzes the cis-trans isomerization of proline imidic peptide bonds in oligopeptides; Belongs to the cyclophilin-type PPIase family. |

| | | | | | | |
|---|---|---|---|---|---|---|
| kmeans | 2 | Salmon | 65 | PF3D7_0610200 | 36329.C6KSU9 | RNA-binding protein 25, putative. |
| kmeans | 2 | Salmon | 65 | PF3D7_0614400 | 36329.C6KSY5 | Pre-mRNA-splicing factor CWF7, putative. |
| kmeans | 2 | Salmon | 65 | PF3D7_0619900 | 36329.C6KT39 | Splicing factor 3A subunit 2, putative. |
| kmeans | 2 | Salmon | 65 | PF3D7_0623600 | 36329.C6KT72 | Transcription or splicing factor-like protein, putative. |
| kmeans | 2 | Salmon | 65 | PF3D7_0716000 | 36329.Q8IBU0 | RNA-binding protein, putative. |
| kmeans | 2 | Salmon | 65 | PF3D7_0722500 | 36329.Q8IBM8 | Pre-mRNA-splicing factor CWC15, putative. |
| kmeans | 2 | Salmon | 65 | PF3D7_0822300 | 36329.Q8IB57 | Small nuclear ribonucleoprotein G. |
| kmeans | 2 | Salmon | 65 | PF3D7_0822800 | 36329.C0H4X1 | U5 small nuclear ribonucleoprotein 40 kDa protein, putative. |
| kmeans | 2 | Salmon | 65 | PF3D7_0909800 | 36329.C0H529 | Small nuclear ribonucleoprotein Sm D3. |
| kmeans | 2 | Salmon | 65 | PF3D7_0917600 | 36329.Q8I2X7 | Pre-mRNA-splicing factor ATP-dependent RNA helicase PRP43, putative. |
| kmeans | 2 | Salmon | 65 | PF3D7_0924700 | 36329.Q8I2R0 | Splicing factor 3A subunit 3, putative. |
| kmeans | 2 | Salmon | 65 | PF3D7_0935000 | 36329.Q8I2G9 | U2 small nuclear ribonucleoprotein B'', putative. |
| kmeans | 2 | Salmon | 65 | PF3D7_1003800 | 36329.Q8IJZ9 | U5 small nuclear ribonucleoprotein component, putative. |
| kmeans | 2 | Salmon | 65 | PF3D7_1033600 | 36329.Q7KQL1 | Pre-mRNA-splicing factor CEF1, putative. |
| kmeans | 2 | Salmon | 65 | PF3D7_1110200 | 36329.Q8IIR0 | Pre-mRNA-processing factor 6, putative. |
| kmeans | 2 | Salmon | 65 | PF3D7_1119300 | 36329.Q8IIH4 | Splicing factor U2AF small subunit, putative. |
| kmeans | 2 | Salmon | 65 | PF3D7_1125500 | 36329.Q8IIA8 | Small nuclear ribonucleoprotein Sm D1; Essential for pre-mRNA splicing. Implicated in the formation of stable, biologically active snRNP structures. Belongs to the snRNP core protein family. |
| kmeans | 2 | Salmon | 65 | PF3D7_1126900 | 36329.Q8II94 | Small nuclear ribonucleoprotein F, putative. |
| kmeans | 2 | Salmon | 65 | PF3D7_1209200 | 36329.Q8I5V6 | U6 snRNA-associated Sm-like protein LSm7, putative. |
| kmeans | 2 | Salmon | 65 | PF3D7_1220100 | 36329.Q8I5L1 | Pre-mRNA-processing factor 17, putative. |
| kmeans | 2 | Salmon | 65 | PF3D7_1224900 | 36329.Q8I5G8 | Splicing factor 3B subunit 6, putative. |
| kmeans | 2 | Salmon | 65 | PF3D7_1230800 | 36329.Q8I5B2 | Pre-mRNA-splicing regulator, putative. |
| kmeans | 2 | Salmon | 65 | PF3D7_1231600 | 36329.Q8I5A4 | Pre-mRNA-splicing factor ATP-dependent RNA helicase PRP2, putative. |
| kmeans | 2 | Salmon | 65 | PF3D7_1234800 | 36329.Q8I574 | Splicing factor 3B subunit 3, putative. |
| kmeans | 2 | Salmon | 65 | PF3D7_1235900 | 36329.A0A144A2S8 | Pre-mRNA-splicing factor SYF1, putative. |
| kmeans | 2 | Salmon | 65 | PF3D7_1249800 | 36329.Q8I4T6 | THO complex subunit 2, putative. |
| kmeans | 2 | Salmon | 65 | PF3D7_1305400 | 36329.Q8IER1 | AAR2 protein, putative. |
| kmeans | 2 | Salmon | 65 | PF3D7_1306900 | 36329.C0H5A4 | U1 small nuclear ribonucleoprotein A, putative. |
| kmeans | 2 | Salmon | 65 | PF3D7_1309300 | 36329.Q8IEM1 | U4/U6 small nuclear ribonucleoprotein PRP3, putative. |
| kmeans | 2 | Salmon | 65 | PF3D7_1316500 | 36329.A0A5K1K8A8 | Pre-mRNA-processing factor 40, putative. |
| kmeans | 2 | Salmon | 65 | PF3D7_1321700 | 36329.Q8IE99 | Splicing factor 1. |
| kmeans | 2 | Salmon | 65 | PF3D7_1324200 | 36329.Q8IE75 | Micro-fibrillar-associated protein, putative. |
| kmeans | 2 | Salmon | 65 | PF3D7_1343900 | 36329.C0H5G7 | U4/U6 small nuclear ribonucleoprotein PRP4, putative. |
| kmeans | 2 | Salmon | 65 | PF3D7_1352700 | 36329.Q8IDH3 | Intron-binding protein aquarius, putative. |
| kmeans | 2 | Salmon | 65 | PF3D7_1355800 | 36329.C0H5I5 | Splicing factor subunit; Belongs to the SF3B5 family. |
| kmeans | 2 | Salmon | 65 | PF3D7_1364300 | 36329.A0A5K1K9B8 | Pre-mRNA-splicing factor ATP-dependent RNA helicase PRP16. |
| kmeans | 2 | Salmon | 65 | PF3D7_1367100 | 36329.Q8ID37 | U1 small nuclear ribonucleoprotein 70 kDa homolog, putative. |
| kmeans | 2 | Salmon | 65 | PF3D7_1369700 | 36329.C0H5L8 | U2 small nuclear ribonucleoprotein A', putative. |
| kmeans | 2 | Salmon | 65 | PF3D7_1414800 | 36329.Q8ILU8 | Small nuclear ribonucleoprotein-associated protein B, putative. |
| kmeans | 2 | Salmon | 65 | PF3D7_1420000 | 36329.Q8ILQ0 | Splicing factor 3B subunit 4, putative. |
| kmeans | 2 | Salmon | 65 | PF3D7_1451500 | 36329.Q8IKW1 | Pre-mRNA-splicing factor CWF18, putative. |
| kmeans | 2 | Salmon | 65 | PF3D7_1461600 | 36329.Q8IKL7 | Splicing factor 3B subunit 2, putative. |
| kmeans | 2 | Salmon | 65 | PF3D7_1472000 | 36329.Q8IKB8 | Pre-mRNA-splicing factor ISY1, putative. |
| kmeans | 2 | Salmon | 65 | PF3D7_1474500 | 36329.Q8IK93 | Splicing factor 3A subunit 1, putative. |
| kmeans | 2 | Salmon | 65 | PFL2310w | 36329.Q8I4V2 | Pre-mRNA-splicing factor PFL2310w; Involved in pre-mRNA splicing. Binds RNA (By similarity). |
| kmeans | 3 | Fire Brick | 31 | MED6 | 36329.Q8IKE0 | Mediator of RNA polymerase II transcription subunit 6; Component of the Mediator complex, a coactivator involved in the regulated transcription of nearly all RNA polymerase II-dependent genes. Mediator functions as a bridge to convey information from gene- specific regulatory proteins to the basal RNA polymerase II transcription machinery. Mediator is recruited to promoters by direct interactions with regulatory proteins and serves as a scaffold for the assembly of a functional preinitiation complex with RNA polymerase II and the general transcription factors. |
| kmeans | 3 | Fire Brick | 31 | PF3D7_0110400 | 36329.Q8I241 | DNA-directed RNA polymerase II subunit RPB9, putative. |
| kmeans | 3 | Fire Brick | 31 | PF3D7_0206000 | 36329.O96154 | DNA repair protein RAD2, putative. |
| kmeans | 3 | Fire Brick | 31 | PF3D7_0215700 | 36329.O96236 | DNA-directed RNA polymerase subunit beta; DNA-dependent RNA polymerase catalyzes the transcription of DNA into RNA using the four ribonucleoside triphosphates as substrates. |
| kmeans | 3 | Fire Brick | 31 | PF3D7_0303300 | 36329.O77315 | DNA-directed RNA polymerases I, II, and III subunit RPABC2, putative. |
| kmeans | 3 | Fire Brick | 31 | PF3D7_0318200 | 36329.O77375 | DNA-directed RNA polymerase subunit; DNA-dependent RNA polymerase catalyzes the transcription of DNA into RNA using the four ribonucleoside triphosphates as substrates. |
| kmeans | 3 | Fire Brick | 31 | PF3D7_0509400 | 36329.Q8I410 | DNA-directed RNA polymerase subunit; DNA-dependent RNA polymerase catalyzes the transcription of DNA into RNA using the four ribonucleoside triphosphates as substrates. |
| kmeans | 3 | Fire Brick | 31 | PF3D7_0517400 | 36329.Q8I3T4 | FACT complex subunit SPT16, putative. |
| kmeans | 3 | Fire Brick | 31 | PF3D7_0520700 | 36329.Q8I3Q4 | CDC73 domain-containing protein, putative. |
| kmeans | 3 | Fire Brick | 31 | PF3D7_0610900 | 36329.C6KSV4 | Transcription elongation factor SPT5, putative. |
| kmeans | 3 | Fire Brick | 31 | PF3D7_0708100 | 36329.Q8IC08 | DNA-directed RNA polymerases I, II, and III subunit RPABC5, putative. |
| kmeans | 3 | Fire Brick | 31 | PF3D7_0714500 | 36329.Q8IBV2 | Transcription elongation factor s-II, putative. |

| | | | | | | |
|---|---|---|---|---|---|---|
| kmeans | 3 | Fire Brick | 31 | PF3D7_0822100 | 36329.Q8IB55 | Mediator of RNA polymerase II transcription subunit 7; Component of the Mediator complex, a coactivator involved in the regulated transcription of nearly all RNA polymerase II-dependent genes. Mediator functions as a bridge to convey information from gene- specific regulatory proteins to the basal RNA polymerase II transcription machinery. |
| kmeans | 3 | Fire Brick | 31 | PF3D7_0923000 | 36329.Q8I2S6 | DNA-directed RNA polymerase II subunit RPB3, putative. |
| kmeans | 3 | Fire Brick | 31 | PF3D7_0934100 | 36329.Q8I2H7 | TFIIH basal transcription factor complex helicase XPD subunit. |
| kmeans | 3 | Fire Brick | 31 | PF3D7_1012700 | 36329.Q8IJR8 | NLI interacting factor-like phosphatase, putative. |
| kmeans | 3 | Fire Brick | 31 | PF3D7_1037600 | 36329.Q8IJ31 | TFIIH basal transcription factor complex helicase XPB subunit, putative. |
| kmeans | 3 | Fire Brick | 31 | PF3D7_1134700 | 36329.Q8II17 | DNA-directed RNA polymerase subunit beta; DNA-dependent RNA polymerase catalyzes the transcription of DNA into RNA using the four ribonucleoside triphosphates as substrates. |
| kmeans | 3 | Fire Brick | 31 | PF3D7_1143300 | 36329.Q8IHT3 | DNA-directed RNA polymerases I and III subunit RPAC1, putative. |
| kmeans | 3 | Fire Brick | 31 | PF3D7_1206600 | 36329.Q8I5X9 | DNA-directed RNA polymerase subunit beta; DNA-dependent RNA polymerase catalyzes the transcription of DNA into RNA using the four ribonucleoside triphosphates as substrates. |
| kmeans | 3 | Fire Brick | 31 | PF3D7_1213700 | 36329.Q8I5R8 | DNA-directed RNA polymerases I, II, and III subunit RPABC3; DNA-dependent RNA polymerase catalyzes the transcription of DNA into RNA using the four ribonucleoside triphosphates as substrates. Common component of RNA polymerases I, II and III which synthesize ribosomal RNA precursors, mRNA precursors and many functional non-coding RNAs, and small RNAs, such as 5S rRNA and tRNAs, respectively. |
| kmeans | 3 | Fire Brick | 31 | PF3D7_1244200 | 36329.Q8I4Y8 | General transcription factor IIH subunit 4; Component of the general transcription and DNA repair factor IIH (TFIIH) core complex which is involved in general and transcription-coupled nucleotide excision repair (NER) of damaged DNA. Belongs to the TFB2 family. |
| kmeans | 3 | Fire Brick | 31 | PF3D7_1314900 | 36329.Q8IEG6 | General transcription factor IIH subunit. |
| kmeans | 3 | Fire Brick | 31 | PF3D7_1329000 | 36329.Q8IE49 | DNA-directed RNA polymerase subunit; DNA-dependent RNA polymerase catalyzes the transcription of DNA into RNA using the four ribonucleoside triphosphates as substrates. |
| kmeans | 3 | Fire Brick | 31 | PF3D7_1364800 | 36329.Q8ID59 | DNA-directed RNA polymerases I, II, and III subunit RPABC1, putative. |
| kmeans | 3 | Fire Brick | 31 | PF3D7_1406200 | 36329.Q8IM32 | Transcription elongation factor SPT6, putative. |
| kmeans | 3 | Fire Brick | 31 | PF3D7_1415200 | 36329.Q8ILU4 | DNA-directed RNA polymerases I and III subunit RPAC2, putative. |
| kmeans | 3 | Fire Brick | 31 | PF3D7_1441400 | 36329.Q8IL56 | FACT complex subunit SSRP1; Component of the FACT complex, a general chromatin factor that acts to reorganize nucleosomes. The FACT complex is involved in multiple processes that require DNA as a template such as mRNA elongation, DNA replication and DNA repair. During transcription elongation the FACT complex acts as a histone chaperone that both destabilizes and restores nucleosomal structure. It facilitates the passage of RNA polymerase II and transcription by promoting the dissociation of one histone H2A-H2B dimer from the nucleosome, then subsequently promotes the reestablishment o [...] |
| kmeans | 3 | Fire Brick | 31 | PF3D7_1458800 | 36329.Q8IKP3 | DNA-directed RNA polymerase III subunit RPC5, putative. |
| kmeans | 3 | Fire Brick | 31 | PF3D7_1463400 | 36329.Q8IKK2 | DNA-directed RNA polymerase III subunit RPC4, putative. |
| kmeans | 3 | Fire Brick | 31 | PF3D7_1475000 | 36329.Q8IK88 | Mediator of RNA polymerase II transcription subunit 31; Component of the Mediator complex, a coactivator involved in the regulated transcription of nearly all RNA polymerase II-dependent genes. Mediator functions as a bridge to convey information from gene- specific regulatory proteins to the basal RNA polymerase II transcription machinery. Mediator is recruited to promoters by direct interactions with regulatory proteins and serves as a scaffold for the assembly of a functional preinitiation complex with RNA polymerase II and the general transcription factors. |
| kmeans | 4 | Brown | 31 | PF3D7_0205900 | 36329.O96153 | 26S proteasome regulatory subunit RPN1, putative. |
| kmeans | 4 | Brown | 31 | PF3D7_0312300 | 36329.O77345 | 26S proteasome regulatory subunit RPN12, putative. |
| kmeans | 4 | Brown | 31 | PF3D7_0317000 | 36329.O77396 | Proteasome subunit alpha type-3, putative. |
| kmeans | 4 | Brown | 31 | PF3D7_0413600 | 36329.Q8I1V1 | 26S protease regulatory subunit 6B, putative; Belongs to the AAA ATPase family. |
| kmeans | 4 | Brown | 31 | PF3D7_0518300 | 36329.A0A5K1K7U1 | Proteasome subunit beta; The proteasome is a multicatalytic proteinase complex which is characterized by its ability to cleave peptides with Arg, Phe, Tyr, Leu, and Glu adjacent to the leaving group at neutral or slightly basic pH; Belongs to the peptidase T1B family. |
| kmeans | 4 | Brown | 31 | PF3D7_0528000 | 36329.A0A5K1K941 | Proteasome maturation factor UMP1, putative. |
| kmeans | 4 | Brown | 31 | PF3D7_0608500 | 36329.C6KST3 | Proteasome endopeptidase complex; Belongs to the peptidase T1A family. |
| kmeans | 4 | Brown | 31 | PF3D7_0727400 | 36329.Q8IBI3 | Proteasome subunit alpha type. |
| kmeans | 4 | Brown | 31 | PF3D7_0803800 | 36329.Q7K6A9 | Proteasome subunit beta; Belongs to the peptidase T1B family. |
| kmeans | 4 | Brown | 31 | PF3D7_0807500 | 36329.Q8IAR3 | Proteasome endopeptidase complex; Belongs to the peptidase T1A family. |
| kmeans | 4 | Brown | 31 | PF3D7_0807800 | 36329.Q8IAR6 | 26S proteasome regulatory subunit RPN10, putative. |
| kmeans | 4 | Brown | 31 | PF3D7_0812100 | 36329.Q8IAV6 | Proteasome activator complex subunit 4, putative. |
| kmeans | 4 | Brown | 31 | PF3D7_0907700 | 36329.Q8I374 | Proteasome activator 28 subunit beta, putative. |
| kmeans | 4 | Brown | 31 | PF3D7_0912900 | 36329.Q8I323 | 26S proteasome regulatory subunit RPN8, putative. |
| kmeans | 4 | Brown | 31 | PF3D7_1008400 | 36329.Q8IJW0 | 26S protease regulatory subunit 4, putative; Belongs to the AAA ATPase family. |
| kmeans | 4 | Brown | 31 | PF3D7_1017900 | 36329.Q8IJM0 | 26S proteasome regulatory subunit p55, putative. |
| kmeans | 4 | Brown | 31 | PF3D7_1021900 | 36329.Q8IJI4 | PHAX domain-containing protein, putative. |
| kmeans | 4 | Brown | 31 | PF3D7_1030500 | 36329.Q8IJA0 | 26S proteasome regulatory subunit RPN9, putative. |
| kmeans | 4 | Brown | 31 | PF3D7_1129200 | 36329.Q8II71 | 26S proteasome regulatory subunit RPN7, putative. |
| kmeans | 4 | Brown | 31 | PF3D7_1130400 | 36329.Q8II60 | 26S protease regulatory subunit 6A, putative; Belongs to the AAA ATPase family. |
| kmeans | 4 | Brown | 31 | PF3D7_1248900 | 36329.Q8I4U5 | 26S protease regulatory subunit 8, putative; Belongs to the AAA ATPase family. |
| kmeans | 4 | Brown | 31 | PF3D7_1306400 | 36329.Q8IEQ1 | 26S protease regulatory subunit 10B, putative; Belongs to the AAA ATPase family. |
| kmeans | 4 | Brown | 31 | PF3D7_1311500 | 36329.Q8IEK3 | 26S protease regulatory subunit 7, putative; Belongs to the AAA ATPase family. |
| kmeans | 4 | Brown | 31 | PF3D7_1338100 | 36329.A0A5K1K9F3 | 26S proteasome regulatory subunit RPN3, putative. |
| kmeans | 4 | Brown | 31 | PF3D7_1353900 | 36329.Q8IDG2 | Proteasome subunit alpha type. |
| kmeans | 4 | Brown | 31 | PF3D7_1368100 | 36329.Q8ID28 | 26S proteasome regulatory subunit RPN11, putative. |
| kmeans | 4 | Brown | 31 | PF3D7_1402300 | 36329.Q8IM66 | 26S proteasome regulatory subunit RPN6. |
| kmeans | 4 | Brown | 31 | PF3D7_1414000 | 36329.Q8ILV5 | 26S proteasome regulatory subunit RPN13, putative. |
| kmeans | 4 | Brown | 31 | PF3D7_1466300 | 36329.Q8IKH3 | 26S proteasome regulatory subunit RPN2, putative; Belongs to the proteasome subunit S1 family. |

| | | | | | | |
|---|---|---|---|---|---|---|
| kmeans | 4 | Brown | 31 | PF3D7_1470900 | 36329.Q8IKC9 | Proteasome subunit beta; The proteasome is a multicatalytic proteinase complex which is characterized by its ability to cleave peptides with Arg, Phe, Tyr, Leu, and Glu adjacent to the leaving group at neutral or slightly basic pH; Belongs to the peptidase T1B family. |
| kmeans | 4 | Brown | 31 | PF3D7_1474800 | 36329.Q8IK90 | Proteasome endopeptidase complex; Belongs to the peptidase T1A family. |
| kmeans | 5 | Sienna | 28 | MCM7 | 36329.Q8IC16 | DNA replication licensing factor MCM7; Acts as component of the mcm2-7 complex (mcm complex) which is the putative replicative helicase essential for 'once per cell cycle' DNA replication initiation and elongation in eukaryotic cells. The active ATPase sites in the mcm2-7 ring are formed through the interaction surfaces of two neighboring subunits such that a critical structure of a conserved arginine finger motif is provided in trans relative to the ATP-binding site of the Walker A box of the adjacent subunit. The six ATPase active sites, however, are likely to contribute differential [...] |
| kmeans | 5 | Sienna | 28 | PF14_0366 | 36329.Q7KQM1 | DNA primase small subunit; DNA primase is the polymerase that synthesizes small RNA primers for the Okazaki fragments made during discontinuous DNA replication. |
| kmeans | 5 | Sienna | 28 | PF3D7_0215800 | 36329.O96237 | Origin recognition complex subunit 5. |
| kmeans | 5 | Sienna | 28 | PF3D7_0308000 | 36329.O77321 | DNA polymerase delta small subunit, putative. |
| kmeans | 5 | Sienna | 28 | PF3D7_0317400 | 36329.O77383 | DNA replication complex GINS protein, putative. |
| kmeans | 5 | Sienna | 28 | PF3D7_0411900 | 36329.Q9U0H1 | DNA polymerase. |
| kmeans | 5 | Sienna | 28 | PF3D7_0416300 | 36329.Q8I1S4 | DNA helicase MCM9, putative; Belongs to the MCM family. |
| kmeans | 5 | Sienna | 28 | PF3D7_0416400 | 36329.C0H4A7 | Histone acetyltransferase, putative. |
| kmeans | 5 | Sienna | 28 | PF3D7_0501800 | 36329.Q8I482 | Protein HIRA; Required for replication-independent chromatin assembly and for the periodic repression of histone gene transcription during the cell cycle; Belongs to the WD repeat HIR1 family. |
| kmeans | 5 | Sienna | 28 | PF3D7_0503200 | 36329.Q8I469 | ATPase_AAA_core domain-containing protein. |
| kmeans | 5 | Sienna | 28 | PF3D7_0527000 | 36329.Q8I3J5 | DNA replication licensing factor MCM3, putative; Belongs to the MCM family. |
| kmeans | 5 | Sienna | 28 | PF3D7_0612900 | 36329.C6KSX2 | Nucleolar GTP-binding protein 1; Involved in the biogenesis of the 60S ribosomal subunit. Belongs to the TRAFAC class OBG-HflX-like GTPase superfamily. OBG GTPase family. NOG subfamily. |
| kmeans | 5 | Sienna | 28 | PF3D7_0630300 | 36329.C6KTD8 | DNA polymerase epsilon catalytic subunit A, putative. |
| kmeans | 5 | Sienna | 28 | PF3D7_0705300 | 36329.Q8IC17 | Origin recognition complex subunit 2. |
| kmeans | 5 | Sienna | 28 | PF3D7_0910900 | 36329.C0H531 | DNA primase large subunit; DNA primase is the polymerase that synthesizes small RNA primers for the Okazaki fragments made during discontinuous DNA replication; Belongs to the eukaryotic-type primase large subunit family. |
| kmeans | 5 | Sienna | 28 | PF3D7_1112100 | 36329.Q8IIP2 | Protein kinase domain-containing protein. |
| kmeans | 5 | Sienna | 28 | PF3D7_1148000 | 36329.Q8IHP1 | Serine/threonine protein kinase, putative. |
| kmeans | 5 | Sienna | 28 | PF3D7_1205600 | 36329.Q8I5Y9 | Tetratricopeptide repeat protein, putative. |
| kmeans | 5 | Sienna | 28 | PF3D7_1211300 | 36329.Q8I5T7 | DNA helicase MCM8, putative; Belongs to the MCM family. |
| kmeans | 5 | Sienna | 28 | PF3D7_1211700 | 36329.Q8I5T4 | DNA replication licensing factor MCM5, putative; Belongs to the MCM family. |
| kmeans | 5 | Sienna | 28 | PF3D7_1224500 | 36329.Q8I5H2 | Histone chaperone ASF1, putative. |
| kmeans | 5 | Sienna | 28 | PF3D7_1305000 | 36329.Q8IER6 | MCL1 domain-containing protein, putative. |
| kmeans | 5 | Sienna | 28 | PF3D7_1317100 | 36329.Q8IEE5 | DNA replication licensing factor MCM4; Belongs to the MCM family. |
| kmeans | 5 | Sienna | 28 | PF3D7_1328200 | 36329.A0A5K1K982 | Uncharacterized protein. |
| kmeans | 5 | Sienna | 28 | PF3D7_1334100 | 36329.A0A5K1K8V1 | Uncharacterized protein. |
| kmeans | 5 | Sienna | 28 | PF3D7_1355100 | 36329.Q8IDF0 | DNA helicase; Belongs to the MCM family. |
| kmeans | 5 | Sienna | 28 | PF3D7_1412100 | 36329.Q8ILX3 | Mini-chromosome maintenance complex-binding protein, putative. |
| kmeans | 5 | Sienna | 28 | PF3D7_1466800 | 36329.Q8IKG9 | NOC3p domain-containing protein. |
| kmeans | 6 | Sandy Brown | 20 | PF14_0425 | 36329.Q7KQL9 | Fructose-bisphosphate aldolase. |
| kmeans | 6 | Sandy Brown | 20 | PF3D7_0109200 | 36329.Q8I0V9 | Cleavage and polyadenylation specificity factor subunit 5; Component of the cleavage factor Im (CFIm) complex that functions as an activator of the pre-mRNA 3'-end cleavage and polyadenylation processing required for the maturation of pre-mRNA into functional mRNAs. CFIm contributes to the recruitment of multiprotein complexes on specific sequences on the pre-mRNA 3'-end, so called cleavage and polyadenylation signals (pA signals). Most pre-mRNAs contain multiple pA signals, resulting in alternative cleavage and polyadenylation (APA) producing mRNAs with variable 3'-end formation. The [...] |
| kmeans | 6 | Sandy Brown | 20 | PF3D7_0318600 | 36329.O77371 | Cleavage and polyadenylation specificity factor, putative. |
| kmeans | 6 | Sandy Brown | 20 | PF3D7_0513200 | 36329.Q8I3X5 | CPSF_A domain-containing protein. |
| kmeans | 6 | Sandy Brown | 20 | PF3D7_0526500 | 36329.Q8I3K0 | Suf domain-containing protein. |
| kmeans | 6 | Sandy Brown | 20 | PF3D7_0609900 | 36329.C6KSU6 | Uncharacterized protein. |
| kmeans | 6 | Sandy Brown | 20 | PF3D7_0620500 | 36329.C6KT45 | Cleavage stimulation factor subunit 1, putative. |
| kmeans | 6 | Sandy Brown | 20 | PF3D7_0624000 | 36329.C6KT76 | Phosphotransferase. |
| kmeans | 6 | Sandy Brown | 20 | PF3D7_0625600 | 36329.C6KT92 | Poly(A) polymerase; Polymerase that creates the 3'-poly(A) tail of mRNA's. |
| kmeans | 6 | Sandy Brown | 20 | PF3D7_0819000 | 36329.Q8IB25 | Cleavage and polyadenylation specificity factor subunit 2. |
| kmeans | 6 | Sandy Brown | 20 | PF3D7_0821600 | 36329.Q8IB50 | Polyribonucleotide 5'-hydroxyl-kinase Clp1, putative. |
| kmeans | 6 | Sandy Brown | 20 | PF3D7_0923900 | 36329.Q8I2R8 | Polyadenylate-binding protein 2, putative. |
| kmeans | 6 | Sandy Brown | 20 | PF3D7_0933000 | 36329.Q8I0W1 | CSTF domain-containing protein, putative. |
| kmeans | 6 | Sandy Brown | 20 | PF3D7_1009200 | 36329.Q8IJV3 | Ribonuclease, putative. |
| kmeans | 6 | Sandy Brown | 20 | PF3D7_1120100 | 36329.Q8IIG6 | Phosphoglycerate mutase. |
| kmeans | 6 | Sandy Brown | 20 | PF3D7_1210400 | 36329.Q8I5U5 | General transcription factor 3C polypeptide 5, putative. |
| kmeans | 6 | Sandy Brown | 20 | PF3D7_1414400 | 36329.Q8ILV1 | Serine/threonine-protein phosphatase. |
| kmeans | 6 | Sandy Brown | 20 | PF3D7_1419900 | 36329.Q8ILQ1 | Cleavage and polyadenylation specificity factor subunit 4, putative. |
| kmeans | 6 | Sandy Brown | 20 | PF3D7_1438500 | 36329.Q8IL83 | Cleavage and polyadenylation specificity factor subunit 3, putative. |
| kmeans | 6 | Sandy Brown | 20 | PGK | 36329.P27362 | Phosphoglycerate kinase. |
| kmeans | 7 | Dark Golden Rod | 13 | PF08_0048 | 36329.C0H4W3 | Probable ATP-dependent helicase PF08_0048; Catalytic component of a chromatin remodeling complex. Belongs to the SNF2/RAD54 helicase family. SWR1 subfamily. |

| | | | | | | |
|---|---|---|---|---|---|---|
| kmeans | 7 | Dark Golden Rod | 13 | PF3D7_0505200 | 36329.Q8I450 | Actin-like protein, putative; Belongs to the actin family. |
| kmeans | 7 | Dark Golden Rod | 13 | PF3D7_0505600 | 36329.Q8I446 | BCNT-C domain-containing protein. |
| kmeans | 7 | Dark Golden Rod | 13 | PF3D7_0624600 | 36329.C6KT82 | SNF2 helicase, putative. |
| kmeans | 7 | Dark Golden Rod | 13 | PF3D7_0628600 | 36329.C6KTC1 | DNA methyltransferase 1-associated protein 1, putative. |
| kmeans | 7 | Dark Golden Rod | 13 | PF3D7_0719300 | 36329.Q8IBQ9 | Actin-related protein, putative; Belongs to the actin family. |
| kmeans | 7 | Dark Golden Rod | 13 | PF3D7_1103600 | 36329.Q8IIW6 | Actin-like protein, putative; Belongs to the actin family. |
| kmeans | 7 | Dark Golden Rod | 13 | PF3D7_1106000 | 36329.Q8IIU3 | RuvB-like helicase; Belongs to the RuvB family. |
| kmeans | 7 | Dark Golden Rod | 13 | PF3D7_1303800 | 36329.Q8IES7 | Uncharacterized protein. |
| kmeans | 7 | Dark Golden Rod | 13 | PF3D7_1316900 | 36329.C0H5C0 | Uncharacterized protein. |
| kmeans | 7 | Dark Golden Rod | 13 | PF3D7_1362200 | 36329.Q8ID85 | RuvB-like helicase; Belongs to the RuvB family. |
| kmeans | 7 | Dark Golden Rod | 13 | PF3D7_1422800 | 36329.Q8ILM5 | Actin-related protein, putative; Belongs to the actin family. |
| kmeans | 7 | Dark Golden Rod | 13 | PF3D7_1464000 | 36329.Q8IKJ7 | YL1 nuclear protein, putative. |
| kmeans | 8 | Yellow | 11 | MAL3P3.6 | 36329.O77323 | T-complex protein 1 subunit eta; Molecular chaperone; assists the folding of proteins upon ATP hydrolysis. Known to play a role, in vitro, in the folding of actin and tubulin (By similarity). |
| kmeans | 8 | Yellow | 11 | PF3D7_0214000 | 36329.O96220 | T-complex protein 1 subunit theta. |
| kmeans | 8 | Yellow | 11 | PF3D7_0306800 | 36329.O97247 | T-complex protein 1 subunit beta. |
| kmeans | 8 | Yellow | 11 | PF3D7_0320300 | 36329.O97282 | T-complex protein 1 subunit epsilon. |
| kmeans | 8 | Yellow | 11 | PF3D7_0718500 | 36329.Q8IBR6 | Prefoldin subunit 3; Binds specifically to cytosolic chaperonin (c-CPN) and transfers target proteins to it. Binds to nascent polypeptide chain and promotes folding in an environment in which there are many competing pathways for nonnative proteins; Belongs to the prefoldin subunit alpha family. |
| kmeans | 8 | Yellow | 11 | PF3D7_0904500 | 36329.Q8I3A4 | Prefoldin subunit 4; Binds specifically to cytosolic chaperonin (c-CPN) and transfers target proteins to it. Binds to nascent polypeptide chain and promotes folding in an environment in which there are many competing pathways for nonnative proteins; Belongs to the prefoldin subunit beta family. |
| kmeans | 8 | Yellow | 11 | PF3D7_1128100 | 36329.Q8II82 | Prefoldin subunit 5, putative. |
| kmeans | 8 | Yellow | 11 | PF3D7_1132200 | 36329.Q8II43 | T-complex protein 1 subunit alpha. |
| kmeans | 8 | Yellow | 11 | PF3D7_1229500 | 36329.Q8I5C4 | T-complex protein 1 subunit gamma. |
| kmeans | 8 | Yellow | 11 | PF3D7_1357800 | 36329.C0H5I7 | T-complex protein 1 subunit delta. |
| kmeans | 8 | Yellow | 11 | PF3D7_1416900 | 36329.Q8ILS7 | Prefoldin subunit 2, putative. |
| kmeans | 9 | Dark Golden Rod 2 | 9 | PF3D7_0302900 | 36329.O77312 | Exportin-1, putative. |
| kmeans | 9 | Dark Golden Rod 2 | 9 | PF3D7_0524000 | 36329.Q8I3M5 | Karyopherin beta. |
| kmeans | 9 | Dark Golden Rod 2 | 9 | PF3D7_0623100 | 36329.C6KT67 | Nuclear polyadenylated RNA-binding protein NAB2, putative. |
| kmeans | 9 | Dark Golden Rod 2 | 9 | PF3D7_0627700 | 36329.C6KTB3 | Transportin. |
| kmeans | 9 | Dark Golden Rod 2 | 9 | PF3D7_0812400 | 36329.Q8IAW0 | Importin subunit alpha; Belongs to the importin alpha family. |
| kmeans | 9 | Dark Golden Rod 2 | 9 | PF3D7_0815200 | 36329.Q8IAY9 | Importin subunit beta, putative. |
| kmeans | 9 | Dark Golden Rod 2 | 9 | PF3D7_0932800 | 36329.Q8I2I8 | Importin alpha re-exporter, putative. |
| kmeans | 9 | Dark Golden Rod 2 | 9 | PF3D7_1117700 | 36329.Q7KQK6 | GTP-binding nuclear protein; GTP-binding protein involved in nucleocytoplasmic transport. Required for the import of protein into the nucleus and also for RNA export. Involved in chromatin condensation and control of cell cycle. Belongs to the small GTPase superfamily. Ran family. |
| kmeans | 9 | Dark Golden Rod 2 | 9 | PF3D7_1315900 | 36329.A0A5K1K8Q1 | Exportin-T; tRNA nucleus export receptor which facilitates tRNA translocation across the nuclear pore complex. Belongs to the exportin family. |
| kmeans | 10 | Khaki | 8 | FEN1 | 36329.Q7K734 | Flap endonuclease 1; Structure-specific nuclease with 5'-flap endonuclease and 5'- 3' exonuclease activities involved in DNA replication and repair. During DNA replication, cleaves the 5'-overhanging flap structure that is generated by displacement synthesis when DNA polymerase encounters the 5'-end of a downstream Okazaki fragment. It enters the flap from the 5'-end and then tracks to cleave the flap base, leaving a nick for ligation. Also involved in the long patch base excision repair (LP-BER) pathway, by cleaving within the apurinic/apyrimidinic (AP) site- terminated flap. Acts as [...] |
| kmeans | 10 | Khaki | 8 | PCNA | 36329.P61074 | Proliferating cell nuclear antigen; This protein is an auxiliary protein of DNA polymerase delta and is involved in the control of eukaryotic DNA replication by increasing the polymerase's processibility during elongation of the leading strand. |

| | | | | | | |
|---|---|---|---|---|---|---|
| kmeans | 10 | Khaki | 8 | PF3D7_0111300 | 36329.Q8I233 | P-loop containing nucleoside triphosphate hydrolase, putative. |
| kmeans | 10 | Khaki | 8 | PF3D7_0218000 | 36329.O96260 | Replication factor C subunit 2, putative. |
| kmeans | 10 | Khaki | 8 | PF3D7_0219600 | 36329.O96271 | Replication factor C subunit 1. |
| kmeans | 10 | Khaki | 8 | PF3D7_1111100 | 36329.Q8IIQ1 | Replication factor C subunit 5, putative. |
| kmeans | 10 | Khaki | 8 | PF3D7_1241700 | 36329.Q8I512 | Replication factor C subunit 4, putative. |
| kmeans | 10 | Khaki | 8 | PF3D7_1463200 | 36329.Q8IKK4 | Replication factor C subunit 3, putative. |
| kmeans | 11 | Green | 8 | PF3D7_0107800 | 36329.A0A143ZUM0 | Double-strand break repair protein MRE11. |
| kmeans | 11 | Green | 8 | PF3D7_0505500 | 36329.Q8I447 | DNA mismatch repair protein; Component of the post-replicative DNA mismatch repair system (MMR). |
| kmeans | 11 | Green | 8 | PF3D7_0725000 | 36329.Q8IBK1 | Exonuclease I, putative. |
| kmeans | 11 | Green | 8 | PF3D7_0726300 | 36329.Q8IBJ3 | DNA mismatch repair protein PMS1, putative. |
| kmeans | 11 | Green | 8 | PF3D7_1117800 | 36329.Q8IIJ0 | DNA mismatch repair protein MLH. |
| kmeans | 11 | Green | 8 | PF3D7_1408400 | 36329.Q8IM12 | FANCJ-like helicase, putative. |
| kmeans | 11 | Green | 8 | PF3D7_1427500 | 36329.Q8ILI9 | DNA mismatch repair protein MSH2, putative. |
| kmeans | 11 | Green | 8 | PFF0285c | 36329.C6KSQ6 | Probable DNA repair protein RAD50; Essential component of the MRN complex, a complex that possesses single-stranded DNA endonuclease and 3' to 5' exonuclease activities, and plays a central role in double-strand break (DSB) repair, chromosome morphogenesis, DNA repair and meiosis. In the complex, it mediates the ATP-binding and is probably required to bind DNA ends and hold them in close proximity (By similarity). Belongs to the SMC family. RAD50 subfamily. |
| kmeans | 12 | Medium Aqua Marine | 8 | PF3D7_0708800 | 36329.Q8IC01 | Heat shock protein 110. |
| kmeans | 12 | Medium Aqua Marine | 8 | PF3D7_0818900 | 36329.Q8IB24 | Heat shock protein 70; Belongs to the heat shock protein 70 family. |
| kmeans | 12 | Medium Aqua Marine | 8 | PF3D7_1037500 | 36329.Q8IJ32 | Dynamin-like protein; Belongs to the TRAFAC class dynamin-like GTPase superfamily. Dynamin/Fzo/YdjA family. |
| kmeans | 12 | Medium Aqua Marine | 8 | PF3D7_1145400 | 36329.Q8IHR4 | Dynamin-like protein; Belongs to the TRAFAC class dynamin-like GTPase superfamily. Dynamin/Fzo/YdjA family. |
| kmeans | 12 | Medium Aqua Marine | 8 | PF3D7_1211400 | 36329.Q7KQK3 | Heat shock protein DNAJ homologue Pfj4. |
| kmeans | 12 | Medium Aqua Marine | 8 | PF3D7_1216900 | 36329.Q8I5N9 | DNA-binding chaperone, putative. |
| kmeans | 12 | Medium Aqua Marine | 8 | PF3D7_1247500 | 36329.Q8I4V7 | Serine/threonine protein kinase, putative. |
| kmeans | 12 | Medium Aqua Marine | 8 | PF3D7_1437900 | 36329.Q8IL88 | HSP40, subfamily A. |
| kmeans | 13 | Aquamarine 4 | 8 | PF3D7_0206700 | 36329.Q7KWJ4 | Adenylosuccinate lyase; Belongs to the lyase 1 family. Adenylosuccinate lyase subfamily. |
| kmeans | 13 | Aquamarine 4 | 8 | PF3D7_0920800 | 36329.Q8I2U5 | Inosine-5'-monophosphate dehydrogenase; Catalyzes the conversion of inosine 5'-phosphate (IMP) to xanthosine 5'-phosphate (XMP), the first committed and rate-limiting step in the de novo synthesis of guanine nucleotides, and therefore plays an important role in the regulation of cell growth. |
| kmeans | 13 | Aquamarine 4 | 8 | PF3D7_1008900 | 36329.Q8IJV6 | Adenylate kinase; Belongs to the adenylate kinase family. |
| kmeans | 13 | Aquamarine 4 | 8 | PF3D7_1012400 | 36329.Q8IJS1 | Hypoxanthine phosphoribosyltransferase; Belongs to the purine/pyrimidine phosphoribosyltransferase family. |
| kmeans | 13 | Aquamarine 4 | 8 | PF3D7_1015800 | 36329.Q8IJN8 | Ribonucleoside-diphosphate reductase small chain, putative. |
| kmeans | 13 | Aquamarine 4 | 8 | PF3D7_1029600 | 36329.Q8IJA9 | Adenosine deaminase. |
| kmeans | 13 | Aquamarine 4 | 8 | PF3D7_1405600 | 36329.Q8IM38 | Ribonucleoside-diphosphate reductase small chain, putative. |
| kmeans | 13 | Aquamarine 4 | 8 | PF3D7_1437200 | 36329.Q8IL94 | Ribonucleoside-diphosphate reductase; Provides the precursors necessary for DNA synthesis. Catalyzes the biosynthesis of deoxyribonucleotides from the corresponding ribonucleotides. |
| kmeans | 14 | Aquamarine 2 | 7 | MAL13P1.96 | 36329.Q8IED2 | Structural maintenance of chromosomes protein 2; May play a role in the conversion of interphase chromatin into condensed chromosomes; Belongs to the SMC family. SMC2 subfamily. |
| kmeans | 14 | Aquamarine 2 | 7 | PF3D7_0509100 | 36329.Q8I413 | Structural maintenance of chromosomes protein 4, putative. |
| kmeans | 14 | Aquamarine 2 | 7 | PF3D7_1130700 | 36329.Q8II57 | Structural maintenance of chromosomes protein 1, putative. |
| kmeans | 14 | Aquamarine 2 | 7 | PF3D7_1304000 | 36329.C0H598 | Condensin complex subunit 2, putative. |
| kmeans | 14 | Aquamarine 2 | 7 | PF3D7_1403100 | 36329.C6S3H2 | Condensin complex subunit 1, putative. |
| kmeans | 14 | Aquamarine 2 | 7 | PF3D7_1456500 | 36329.Q8IKR4 | SCD domain-containing protein. |
| kmeans | 14 | Aquamarine 2 | 7 | PFD0685c | 36329.Q8I1U7 | Structural maintenance of chromosomes protein 3 homolog; Central component of cohesin, a complex required for chromosome cohesion during the cell cycle. The cohesin complex may form a large proteinaceous ring within which sister chromatids can be trapped. At anaphase, the complex is cleaved and dissociates from chromatin, allowing sister chromatids to segregate. Cohesion is coupled to DNA replication and is involved in DNA repair. The cohesin complex plays also an important role in spindle pole assembly during mitosis and in chromosomes movement (By similarity); Belongs to the SMC fami [...] |
| kmeans | 15 | Cyan | 7 | PF3D7_0807100 | 36329.Q8IAR1 | DNA helicase PSH3. |
| kmeans | 15 | Cyan | 7 | PF3D7_0811300 | 36329.C0H4T9 | CCR4-associated factor 1. |
| kmeans | 15 | Cyan | 7 | PF3D7_1014600 | 36329.Q8IJP9 | Transcriptional coactivator ADA2. |
| kmeans | 15 | Cyan | 7 | PF3D7_1103800 | 36329.Q8IIW4 | CCR4-NOT transcription complex subunit 1, putative. |
| kmeans | 15 | Cyan | 7 | PF3D7_1235300 | 36329.Q8I569 | CCR4-NOT transcription complex subunit 4, putative. |
| kmeans | 15 | Cyan | 7 | PF3D7_1308900 | 36329.Q8IEM5 | mRNA-decapping enzyme 2, putative. |
| kmeans | 15 | Cyan | 7 | PF3D7_1417200 | 36329.Q8ILS4 | NOT family protein, putative. |
| kmeans | 16 | Aquamarine | 6 | PF3D7_0409600 | 36329.Q9U0J0 | Replication protein A1, large subunit. |

| | | | | | | |
|---|---|---|---|---|---|---|
| kmeans | 16 | Aquamarine | 6 | PF3D7_0514100 | 36329.Q8I3W6 | ATP-dependent DNA helicase UvrD. |
| kmeans | 16 | Aquamarine | 6 | PF3D7_0904800 | 36329.Q8I3A1 | Replication protein A1, small. |
| kmeans | 16 | Aquamarine | 6 | PF3D7_1107400 | 36329.Q8IIS8 | DNA repair protein RAD51 homolog; Binds to single and double-stranded DNA and exhibits DNA- dependent ATPase activity. Underwinds duplex DNA. Belongs to the RecA family. RAD51 subfamily. |
| kmeans | 16 | Aquamarine | 6 | PF3D7_1132300 | 36329.Q8II42 | Nucleic acid binding protein, putative. |
| kmeans | 16 | Aquamarine | 6 | PF3D7_1442100 | 36329.C6S3I6 | Replication factor A protein 3, putative. |
| kmeans | 17 | Aquamarine 3 | 5 | PF3D7_0925700 | 36329.Q7K6A1 | Histone deacetylase; Belongs to the histone deacetylase family. HD Type 1 subfamily. |
| kmeans | 17 | Aquamarine 3 | 5 | PF3D7_0929000 | 36329.Q8I2M0 | Transcription initiation factor TFIID subunit 7, putative. |
| kmeans | 17 | Aquamarine 3 | 5 | PF3D7_1008100 | 36329.Q8IJW2 | Zinc finger protein, putative. |
| kmeans | 17 | Aquamarine 3 | 5 | PF3D7_1138800 | 36329.Q8IHX6 | WD repeat-containing protein, putative. |
| kmeans | 17 | Aquamarine 3 | 5 | PF3D7_1141800 | 36329.Q8IHU7 | EELM2 domain-containing protein, putative. |
| kmeans | 18 | Sky Blue 4 | 4 | PF3D7_0621500 | 36329.C6KT53 | Ribonuclease P/MRP protein subunit RPP1, putative. |
| kmeans | 18 | Sky Blue 4 | 4 | PF3D7_0627900 | 36329.C6KTB5 | Ribonuclease P protein subunit p29, putative. |
| kmeans | 18 | Sky Blue 4 | 4 | PF3D7_0709600 | 36329.C0H4M3 | Ribonucleases P/MRP protein subunit POP1, putative. |
| kmeans | 18 | Sky Blue 4 | 4 | PF3D7_1111700 | 36329.Q8IIP6 | Uncharacterized protein. |
| kmeans | 19 | Sky Blue 2 | 4 | PF3D7_0317700 | 36329.O77380 | CPSF (Cleavage and polyadenylation specific factor), subunit A, putative. |
| kmeans | 19 | Sky Blue 2 | 4 | PF3D7_0319100 | 36329.O77367 | E3 ubiquitin-protein ligase RBX1, putative. |
| kmeans | 19 | Sky Blue 2 | 4 | PF3D7_0629800 | 36329.C6KTD3 | Cullin-like protein, putative; Belongs to the cullin family. |
| kmeans | 19 | Sky Blue 2 | 4 | PF3D7_0811000 | 36329.Q8IAU5 | Cullin-1, putative; Belongs to the cullin family. |
| kmeans | 20 | Sky Blue | 4 | PF3D7_0406100 | 36329.Q6ZMA8 | Vacuolar proton pump subunit B; Non-catalytic subunit of the peripheral V1 complex of vacuolar ATPase; Belongs to the ATPase alpha/beta chains family. |
| kmeans | 20 | Sky Blue | 4 | PF3D7_1341900 | 36329.Q8IDS0 | V-type proton ATPase subunit D, putative. |
| kmeans | 20 | Sky Blue | 4 | PF3D7_1464700 | 36329.Q8IKJ0 | V-type proton ATPase subunit; Subunit of the integral membrane V0 complex of vacuolar ATPase. Vacuolar ATPase is responsible for acidifying a variety of intracellular compartments in eukaryotic cells, thus providing most of the energy required for transport processes in the vacuolar system. Belongs to the V-ATPase V0D/AC39 subunit family. |
| kmeans | 20 | Sky Blue | 4 | vapA | 36329.Q76NM6 | V-type proton ATPase catalytic subunit A; Catalytic subunit of the peripheral V1 complex of vacuolar ATPase. V-ATPase vacuolar ATPase is responsible for acidifying a variety of intracellular compartments in eukaryotic cells; Belongs to the ATPase alpha/beta chains family. |
| kmeans | 21 | Sky Blue 3 | 3 | PF3D7_0810300 | 36329.C0H4T6 | Protein phosphatase PPM5, putative. |
| kmeans | 21 | Sky Blue 3 | 3 | PF3D7_0934800 | 36329.Q7K6A0 | cAMP-dependent protein kinase catalytic subunit; Belongs to the protein kinase superfamily. |
| kmeans | 21 | Sky Blue 3 | 3 | PF3D7_1223100 | 36329.Q7KQK0 | cAMP-dependent protein kinase regulatory subunit. |
| kmeans | 22 | Cornflower Blue 2 | 3 | PF10_0084 | 36329.Q7KQL5 | Tubulin beta chain; Tubulin is the major constituent of microtubules. It binds two moles of GTP, one at an exchangeable site on the beta chain and one at a non-exchangeable site on the alpha chain. |
| kmeans | 22 | Cornflower Blue 2 | 3 | PF3D7_0317500 | 36329.O77382 | Kinesin-5; Belongs to the TRAFAC class myosin-kinesin ATPase superfamily. Kinesin family. |
| kmeans | 22 | Cornflower Blue 2 | 3 | PF3D7_0903700 | 36329.Q6ZLZ9 | Tubulin alpha chain; Tubulin is the major constituent of microtubules. It binds two moles of GTP, one at an exchangeable site on the beta chain and one at a non-exchangeable site on the alpha chain. |
| kmeans | 23 | Cornflower Blue | 3 | PF3D7_0823800 | 36329.Q8IB72 | DnaJ protein, putative. |
| kmeans | 23 | Cornflower Blue | 3 | PF3D7_1015600 | 36329.Q8IJN9 | Heat shock protein 60; Belongs to the chaperonin (HSP60) family. |
| kmeans | 23 | Cornflower Blue | 3 | PF3D7_1134000 | 36329.Q8II24 | Heat shock protein 70; Belongs to the heat shock protein 70 family. |
| kmeans | 24 | Blue | 3 | PF3D7_0922200 | 36329.Q7K6A4 | S-adenosylmethionine synthase; Catalyzes the formation of S-adenosylmethionine from methionine and ATP. |
| kmeans | 24 | Blue | 3 | PF3D7_1033100 | 36329.Q8IJ77 | S-adenosylmethionine decarboxylase/ornithine decarboxylase. |
| kmeans | 24 | Blue | 3 | PF3D7_1129000 | 36329.Q8II73 | Spermidine synthase; Belongs to the spermidine/spermine synthase family. |
| kmeans | 25 | Medium Slate Blue 3 | 3 | PF3D7_1103700 | 36329.Q8IIW5 | Casein kinase II subunit beta; Plays a complex role in regulating the basal catalytic activity of the alpha subunit; Belongs to the casein kinase 2 subunit beta family. |
| kmeans | 25 | Medium Slate Blue 3 | 3 | PF3D7_1108400 | 36329.Q8IIR9 | Casein kinase 2, alpha subunit; Belongs to the protein kinase superfamily. |
| kmeans | 25 | Medium Slate Blue 3 | 3 | PF3D7_1342400 | 36329.Q8IDR5 | Casein kinase II subunit beta; Plays a complex role in regulating the basal catalytic activity of the alpha subunit; Belongs to the casein kinase 2 subunit beta family. |
| kmeans | 26 | Medium Slate Blue 2 | 3 | PF3D7_0111800 | 36329.B9ZSJ0 | Eukaryotic translation initiation factor 4E, putative; Belongs to the eukaryotic initiation factor 4E family. |
| kmeans | 26 | Medium Slate Blue 2 | 3 | PF3D7_0315100 | 36329.O97266 | Eukaryotic translation initiation factor 4E; Belongs to the eukaryotic initiation factor 4E family. |
| kmeans | 26 | Medium Slate Blue 2 | 3 | PF3D7_1107300 | 36329.Q8IIS9 | Polyadenylate-binding protein-interacting protein 1, putative. |
| kmeans | 27 | Medium Slate Blue | 3 | PF3D7_0517300 | 36329.Q8I3T5 | Serine/arginine-rich splicing factor 1. |
| kmeans | 27 | Medium Slate Blue | 3 | PF3D7_1022400 | 36329.Q8IJI0 | Serine/arginine-rich splicing factor 4. |
| kmeans | 27 | Medium Slate Blue | 3 | PF3D7_1119800 | 36329.Q8IIG9 | Alternative splicing factor ASF-1, putative. |
| kmeans | 28 | Medium Purple | 2 | PF3D7_0205400 | 36329.O96149 | PCI domain-containing protein, putative. |

| | | | | | | |
|---|---|---|---|---|---|---|
| kmeans | 28 | Medium Purple | 2 | PF3D7_0602600 | 36329.C6KSM6 | SAC3 domain-containing protein, putative. |
| kmeans | 29 | Medium Purple 2 | 2 | PF3D7_0611400 | 36329.C6KSV9 | SWIB/MDM2 domain-containing protein. |
| kmeans | 29 | Medium Purple 2 | 2 | PF3D7_1225200 | 36329.Q8I5G5 | Uncharacterized protein. |
| kmeans | 30 | Medium Purple 3 | 2 | PF3D7_0320500 | 36329.O97284 | Nicotinamidase, putative. |
| kmeans | 30 | Medium Purple 3 | 2 | PF3D7_0629100 | 36329.C6KTC6 | Nicotinate phosphoribosyltransferase, putative. |
| kmeans | 31 | Medium Orchid | 2 | PF3D7_0316500 | 36329.O77390 | Kinetochore protein NUF2, putative. |
| kmeans | 31 | Medium Orchid | 2 | PF3D7_1322200 | 36329.A0A5K1K9E8 | Uncharacterized protein. |
| kmeans | 32 | Orchid 4 | 2 | PF3D7_0505800 | 36329.Q8I444 | Small ubiquitin-related modifier. |
| kmeans | 32 | Orchid 4 | 2 | PF3D7_0801700 | 36329.C0H4Y4 | Sentrin-specific protease 2, putative. |
| kmeans | 33 | Orchid 2 | 2 | PF3D7_1250800 | 36329.Q8I4S6 | DNA repair protein rhp16, putative. |
| kmeans | 33 | Orchid 2 | 2 | PF3D7_1363200 | 36329.Q8ID74 | Bifunctional polynucleotide phosphatase/kinase. |
| kmeans | 34 | Violet 2 | 2 | PF3D7_0729500 | 36329.Q8I6Z2 | mRNA (N6-adenosine)-methyltransferase, putative; Belongs to the MT-A70-like family. |
| kmeans | 34 | Violet 2 | 2 | PF3D7_1235500 | 36329.Q8I567 | mRNA methyltransferase, putative; Belongs to the MT-A70-like family. |
| kmeans | 35 | Purple | 2 | PF3D7_1230700 | 36329.Q8I5B3 | Protein transport protein SEC13; Belongs to the WD repeat SEC13 family. |
| kmeans | 35 | Purple | 2 | PF3D7_1361100 | 36329.C0H5J6 | Protein transport protein Sec24A. |
| kmeans | 36 | Violet | 2 | PF3D7_0802000 | 36329.Q8IAM0 | Glutamate dehydrogenase, putative. |
| kmeans | 36 | Violet | 2 | PF3D7_1345700 | 36329.Q8I6T2 | Isocitrate dehydrogenase [NADP]; Belongs to the isocitrate and isopropylmalate dehydrogenases family. |
| kmeans | 37 | Orchid | 2 | PF3D7_0804900 | 36329.Q8IAP4 | GTPase-activating protein, putative. |
| kmeans | 37 | Orchid | 2 | PF3D7_1145100 | 36329.Q8IHR6 | Coatomer subunit gamma; The coatomer is a cytosolic protein complex that binds to dilysine motifs and reversibly associates with Golgi non-clathrin- coated vesicles, which further mediate biosynthetic protein transport from the ER, via the Golgi up to the trans Golgi network. Coatomer complex is required for budding from Golgi membranes, and is essential for the retrograde Golgi-to-ER transport of dilysine-tagged proteins. |
| kmeans | 38 | Orchid 3 | 2 | PF3D7_0810800 | 36329.Q8IAU3 | Hydroxymethyldihydropterin pyrophosphokinase-dihydropteroate synthase. |
| kmeans | 38 | Orchid 3 | 2 | PF3D7_1224000 | 36329.Q8I5H7 | GTP cyclohydrolase 1. |
| kmeans | 39 | Hot Pink | 2 | PF3D7_1347100 | 36329.Q8IDM7 | DNA topoisomerase; Introduces a single-strand break via transesterification at a target site in duplex DNA. Releases the supercoiling and torsional tension of DNA introduced during the DNA replication and transcription by transiently cleaving and rejoining one strand of the DNA duplex. The scissile phosphodiester is attacked by the catalytic tyrosine of the enzyme, resulting in the formation of a DNA-(5'-phosphotyrosyl)-enzyme intermediate and the expulsion of a 3'-OH DNA strand. Belongs to the type IA topoisomerase family. |
| kmeans | 39 | Hot Pink | 2 | PF3D7_1368800 | 36329.Q8ID22 | DNA repair endonuclease XPF, putative. |
| kmeans | 40 | Hot Pink 2 | 2 | PF3D7_0912800 | 36329.Q8I324 | tRNA (Adenine(58)-N(1))-methyltransferase non-catalytic subunit TRM6, putative. |
| kmeans | 40 | Hot Pink 2 | 2 | PF3D7_1315700 | 36329.Q8IEF7 | tRNA (adenine(58)-N(1))-methyltransferase. |
| kmeans | 41 | Pale Violet Red | 2 | PF3D7_1118200 | 36329.Q8III6 | Heat shock protein 90, putative. |
| kmeans | 41 | Pale Violet Red | 2 | PF3D7_1453700 | 36329.Q8IKU1 | HSP90 co-chaperone p23. |
| kmeans | 42 | Pink | 2 | PF3D7_1146600 | 36329.Q8IHQ2 | Oocyst rupture protein 1, putative. |
| kmeans | 42 | Pink | 2 | PF3D7_1439500 | 36329.Q8IL74 | Oocyst rupture protein 2, putative. |
| kmeans | 43 | Light Coral | 2 | PF3D7_1342800 | 36329.Q8IDR1 | Phosphoenolpyruvate carboxykinase. |
| kmeans | 43 | Light Coral | 2 | PF3D7_1426700 | 36329.Q8ILJ7 | Phosphoenolpyruvate carboxylase. |
