## Supplementary material for "*Plasmodium falciparum* SET10 is a histone H3 lysine K18 methyltransferase that participates in a chromatin modulation network crucial for intraerythrocytic development": Table S6

Highest confidence (0.9000)
**k-means**
29 cluster
cluster with ≥4 proteins analyzed
disconnected nodes excluded

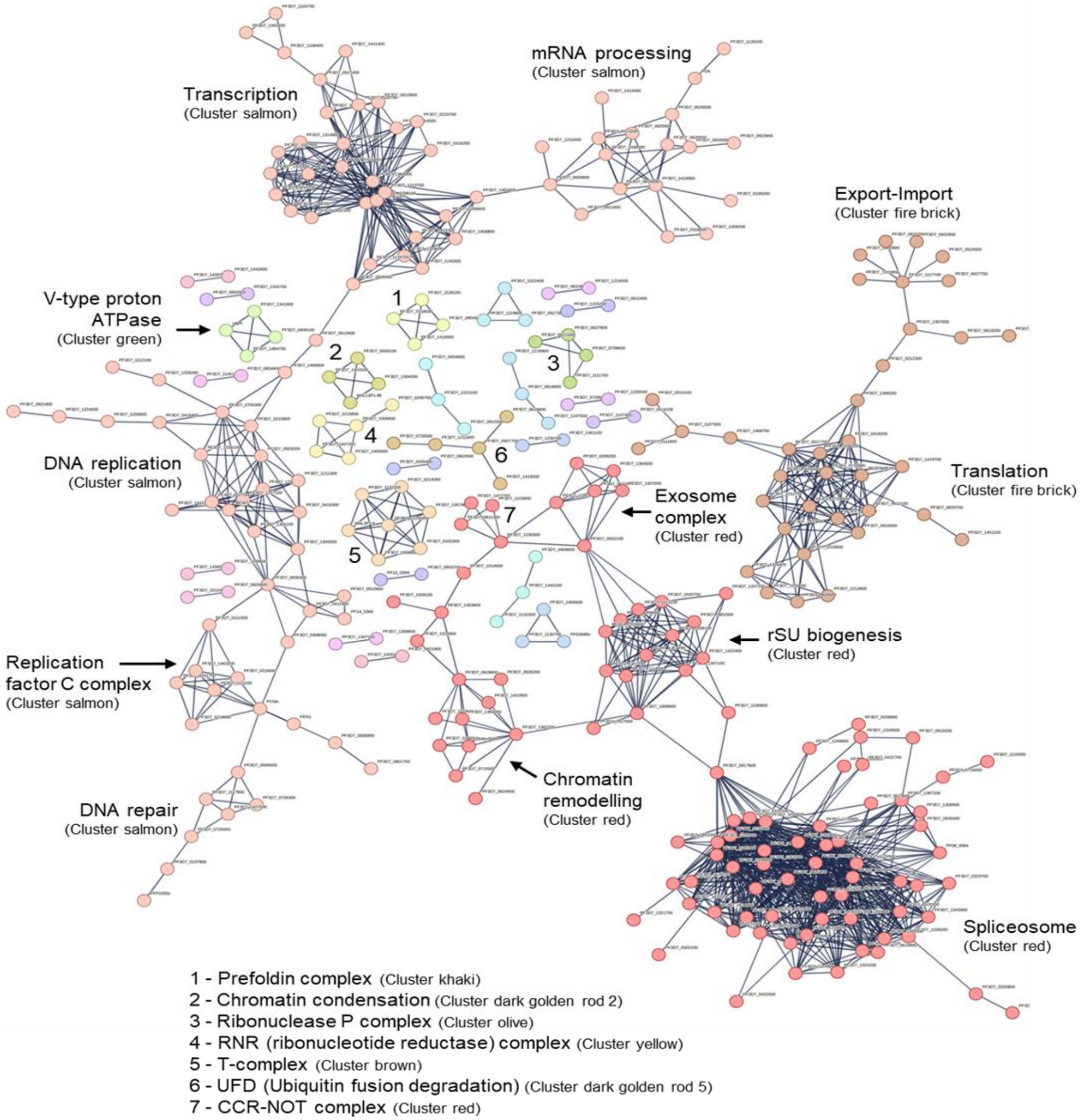

| Cluster red | Cluster number | Cluster color | Gene ID | Protein identifier | Protein description |
|---|---|---|---|---|---|
| | 1 | Red | PF3D7_0209200 | 36329.O96177 | Exosome complex component MTR3, putative. |
| | 1 | Red | PF3D7_0209800 | 36329.Q9TY94 | ATP-dependent RNA helicase UAP56. |
| | 1 | Red | PF3D7_0218500 | 36329.O96265 | Small nuclear ribonucleoprotein Sm D2. |
| | 1 | Red | PF3D7_0218700 | 36329.O96267 | Pre-mRNA-processing protein 45, putative. |
| | 1 | Red | PF3D7_0302000 | 36329.O97334 | Pre-mRNA-splicing factor PRP46, putative. |
| | 1 | Red | PF3D7_0302100 | 36329.O77306 | Serine/threonine protein kinase. |
| | 1 | Red | PF3D7_0308600 | 36329.O77325 | Pre-mRNA-processing factor 19, putative. |
| | 1 | Red | PF3D7_0319500 | 36329.O97318 | RNA-binding protein, putative. |
| | 1 | Red | PF3D7_0320800 | 36329.O97285 | ATP-dependent RNA helicase DDX6; Belongs to the DEAD box helicase family. |
| | 1 | Red | PF3D7_0323700 | 36329.O97303 | U4/U6.U5 tri-snRNP-associated protein 1, putative. |
| | 1 | Red | PF3D7_0403700 | 36329.Q8I1Z2 | Pre-mRNA-splicing factor CLF1, putative. |
| | 1 | Red | PF3D7_0405400 | 36329.Q8I1X5 | Pre-mRNA-processing-splicing factor 8, putative. |
| | 1 | Red | PF3D7_0410400 | 36329.Q9U0I4 | Exosome complex component RRP4. |
| | 1 | Red | PF3D7_0422500 | 36329.Q8IFP1 | Pre-mRNA-splicing helicase BRR2, putative. |
| | 1 | Red | PF3D7_0422700 | 36329.Q8IFN9 | Eukaryotic initiation factor 4A-III, putative; Belongs to the DEAD box helicase family. |
| | 1 | Red | PF3D7_0505200 | 36329.Q8I450 | Actin-like protein, putative; Belongs to the actin family. |
| | 1 | Red | PF3D7_0520300 | 36329.Q8I3Q7 | U6 snRNA-associated Sm-like protein LSm2; Binds specifically to the 3'-terminal U-tract of U6 snRNA. |
| | 1 | Red | PF3D7_0522800 | 36329.Q8I3N5 | Pre-mRNA-splicing factor BUD31, putative. |
| | 1 | Red | PF3D7_0528700 | 36329.Q8I3I0 | Peptidyl-prolyl cis-trans isomerase; PPIases accelerate the folding of proteins. It catalyzes the cis-trans isomerization of proline imidic peptide bonds in oligopeptides; Belongs to the cyclophilin-type PPIase family. |
| | 1 | Red | PF3D7_0602100 | 36329.C6KSM1 | ATP-dependent RNA helicase, putative. |
| | 1 | Red | PF3D7_0610200 | 36329.C6KSU9 | RNA-binding protein 25, putative. |
| | 1 | Red | PF3D7_0614400 | 36329.C6KSY5 | Pre-mRNA-splicing factor CWF7, putative. |
| | 1 | Red | PF3D7_0619900 | 36329.C6KT39 | Splicing factor 3A subunit 2, putative. |
| | 1 | Red | PF3D7_0623600 | 36329.C6KT72 | Transcription or splicing factor-like protein, putative. |
| | 1 | Red | PF3D7_0624600 | 36329.C6KT82 | SNF2 helicase, putative. |
| | 1 | Red | PF3D7_0628600 | 36329.C6KTC1 | DNA methyltransferase 1-associated protein 1, putative. |
| | 1 | Red | PF3D7_0716000 | 36329.Q8IBU0 | RNA-binding protein, putative. |
| | 1 | Red | PF3D7_0719300 | 36329.Q8IBQ9 | Actin-related protein, putative; Belongs to the actin family. |
| | 1 | Red | PF3D7_0722500 | 36329.Q8IBM8 | Pre-mRNA-splicing factor CWC15, putative. |
| | 1 | Red | PF3D7_0802300 | 36329.Q8IAM3 | Periodic tryptophan protein 2, putative. |
| | 1 | Red | PF3D7_0811300 | 36329.C0H4T9 | CCR4-associated factor 1. |
| | 1 | Red | PF3D7_0812700 | 36329.Q8IAW3 | U1 small nuclear ribonucleoprotein C; Component of the spliceosomal U1 snRNP, which is essential for recognition of the pre-mRNA 5' splice-site and the subsequent assembly of the spliceosome. U1-C is directly involved in initial 5' splice-site recognition for both constitutive and regulated alternative splicing. The interaction with the 5' splice-site seems to precede base-pairing between the pre-mRNA and the U1 snRNA. Stimulates commitment or early (E) complex formation by stabilizing the base pairing of the 5' end of the U1 snRNA and the 5' splice-site region. |
| | 1 | Red | PF3D7_0819900 | 36329.C0H4W2 | U6 snRNA-associated Sm-like protein LSm3; Binds specifically to the 3'-terminal U-tract of U6 snRNA. |
| | 1 | Red | PF3D7_0820000 | 36329.C0H4W3 | Snf2-related CBP activator, putative |
| | 1 | Red | PF3D7_0822300 | 36329.Q8IB57 | Small nuclear ribonucleoprotein G. |
| | 1 | Red | PF3D7_0822800 | 36329.C0H4X1 | U5 small nuclear ribonucleoprotein 40 kDa protein, putative. |
| | 1 | Red | PF3D7_0909800 | 36329.C0H529 | Small nuclear ribonucleoprotein Sm D3. |
| | 1 | Red | PF3D7_0917600 | 36329.Q8I2X7 | Pre-mRNA-splicing factor ATP-dependent RNA helicase PRP43, putative. |
| | 1 | Red | PF3D7_0924700 | 36329.Q8I2R0 | Splicing factor 3A subunit 3, putative. |
| | 1 | Red | PF3D7_0935000 | 36329.Q8I2G9 | U2 small nuclear ribonucleoprotein B'', putative. |
| | 1 | Red | PF3D7_1003800 | 36329.Q8IJZ9 | U5 small nuclear ribonucleoprotein component, putative. |
| | 1 | Red | PF3D7_1008100 | 36329.Q8IJW2 | Zinc finger protein, putative. |
| | 1 | Red | PF3D7_1008800 | 36329.Q8IJV7 | Nucleolar protein 5, putative. |
| | 1 | Red | PF3D7_1013100 | 36329.Q8IJR4 | U3 small nucleolar RNA-associated protein 13, putative. |
| | 1 | Red | PF3D7_1014600 | 36329.Q8IJP9 | Transcriptional coactivator ADA2. |
| | 1 | Red | PF3D7_1020700 | 36329.Q8IJJ5 | RNA cytidine acetyltransferase; RNA cytidine acetyltransferase with specificity toward both 18S rRNA and tRNAs. Catalyzes the formation of N(4)-acetylcytidine (ac4C) in 18S rRNA. Required for early nucleolar cleavages of precursor rRNA at sites A0, A1 and A2 during 18S rRNA synthesis. Catalyzes the formation of ac4C in serine and leucine tRNAs. Requires a tRNA-binding adapter protein for full tRNA acetyltransferase activity but not for 18S rRNA acetylation. |
| | 1 | Red | PF3D7_1033600 | 36329.Q7KQL1 | Pre-mRNA-splicing factor CEF1, putative. |
| | 1 | Red | PF3D7_1103600 | 36329.Q8IIW6 | Actin-like protein, putative; Belongs to the actin family. |
| | 1 | Red | PF3D7_1103800 | 36329.Q8IIW4 | CCR4-NOT transcription complex subunit 1, putative. |
| | 1 | Red | PF3D7_1106000 | 36329.Q8IIU3 | RuvB-like helicase; Belongs to the RuvB family. |
| | 1 | Red | PF3D7_1107000 | 36329.Q8IIT3 | U6 snRNA-associated Sm-like protein LSm4; Binds specifically to the 3'-terminal U-tract of U6 snRNA. |
| | 1 | Red | PF3D7_1110200 | 36329.Q8IIR0 | Pre-mRNA-processing factor 6, putative. |

| | | | | | |
|---|---|---|---|---|---|
| | 1 | Red | PF3D7_1118500 | 36329.Q8III3 | Nucleolar protein 56, putative. |
| | 1 | Red | PF3D7_1119300 | 36329.Q8IIH4 | Splicing factor U2AF small subunit, putative. |
| | 1 | Red | PF3D7_1125500 | 36329.Q8IIA8 | Small nuclear ribonucleoprotein Sm D1; Essential for pre-mRNA splicing. Implicated in the formation of stable, biologically active snRNP structures. Belongs to the snRNP core protein family. |
| | 1 | Red | PF3D7_1126900 | 36329.Q8II94 | Small nuclear ribonucleoprotein F, putative. |
| | 1 | Red | PF3D7_1207100 | 36329.Q8I5X4 | Small subunit rRNA processing factor, putative. |
| | 1 | Red | PF3D7_1208100 | 36329.Q8I5W6 | Uncharacterized protein. |
| | 1 | Red | PF3D7_1209200 | 36329.Q8I5V6 | U6 snRNA-associated Sm-like protein LSm7, putative. |
| | 1 | Red | PF3D7_1220100 | 36329.Q8I5L1 | Pre-mRNA-processing factor 17, putative. |
| | 1 | Red | PF3D7_1224900 | 36329.Q8I5G8 | Splicing factor 3B subunit 6, putative. |
| | 1 | Red | PF3D7_1230800 | 36329.Q8I5B2 | Pre-mRNA-splicing regulator, putative. |
| | 1 | Red | PF3D7_1231600 | 36329.Q8I5A4 | Pre-mRNA-splicing factor ATP-dependent RNA helicase PRP2, putative. |
| | 1 | Red | PF3D7_1234800 | 36329.Q8I574 | Splicing factor 3B subunit 3, putative. |
| | 1 | Red | PF3D7_1235300 | 36329.Q8I569 | CCR4-NOT transcription complex subunit 4, putative. |
| | 1 | Red | PF3D7_1235900 | 36329.A0A144A2S8 | Pre-mRNA-splicing factor SYF1, putative. |
| | 1 | Red | PF3D7_1248200 | 36329.Q8I4V2 | Pre-mRNA-splicing factor PFL2310w; Involved in pre-mRNA splicing. Binds RNA (By similarity). |
| | 1 | Red | PF3D7_1249800 | 36329.Q8I4T6 | THO complex subunit 2, putative. |
| | 1 | Red | PF3D7_1303800 | 36329.Q8IES7 | Uncharacterized protein. |
| | 1 | Red | PF3D7_1305400 | 36329.Q8IER1 | AAR2 protein, putative. |
| | 1 | Red | PF3D7_1306900 | 36329.C0H5A4 | U1 small nuclear ribonucleoprotein A, putative. |
| | 1 | Red | PF3D7_1307000 | 36329.Q8IEP5 | Exosome complex component RRP40, putative. |
| | 1 | Red | PF3D7_1307100 | 36329.Q8IEP4 | U3 small nucleolar RNA-associated protein 6, putative. |
| | 1 | Red | PF3D7_1308900 | 36329.Q8IEM5 | mRNA-decapping enzyme 2, putative. |
| | 1 | Red | PF3D7_1309300 | 36329.Q8IEM1 | U4/U6 small nuclear ribonucleoprotein PRP3, putative. |
| | 1 | Red | PF3D7_1316500 | 36329.A0A5K1K8A8 | Pre-mRNA-processing factor 40, putative. |
| | 1 | Red | PF3D7_1316900 | 36329.C0H5C0 | Uncharacterized protein. |
| | 1 | Red | PF3D7_1321700 | 36329.Q8IE99 | Splicing factor 1. |
| | 1 | Red | PF3D7_1324200 | 36329.Q8IE75 | Micro-fibrillar-associated protein, putative. |
| | 1 | Red | PF3D7_1333600 | 36329.Q8IDZ2 | U3 small nucleolar RNA-associated protein 4, putative. |
| | 1 | Red | PF3D7_1340100 | 36329.A0A5K1K8M9 | Exosome complex component RRP42, putative. |
| | 1 | Red | PF3D7_1343900 | 36329.C0H5G7 | U4/U6 small nuclear ribonucleoprotein PRP4, putative. |
| | 1 | Red | PF3D7_1352700 | 36329.Q8IDH3 | Intron-binding protein aquarius, putative. |
| | 1 | Red | PF3D7_1355800 | 36329.C0H5I5 | Splicing factor subunit; Belongs to the SF3B5 family. |
| | 1 | Red | PF3D7_1357700 | 36329.A0A5K1K903 | U3 small nucleolar RNA-associated protein 21, putative. |
| | 1 | Red | PF3D7_1362200 | 36329.Q8ID85 | RuvB-like helicase; Belongs to the RuvB family. |
| | 1 | Red | PF3D7_1364300 | 36329.A0A5K1K9B8 | Pre-mRNA-splicing factor ATP-dependent RNA helicase PRP16. |
| | 1 | Red | PF3D7_1364500 | 36329.Q8ID62 | Exosome complex component RRP45, putative. |
| | 1 | Red | PF3D7_1367100 | 36329.Q8ID37 | U1 small nuclear ribonucleoprotein 70 kDa homolog, putative. |
| | 1 | Red | PF3D7_1407100 | 36329.Q8IM23 | rRNA 2'-O-methyltransferase fibrillarin, putative. |
| | 1 | Red | PF3D7_1414800 | 36329.Q8ILU8 | Small nuclear ribonucleoprotein-associated protein B, putative. |
| | 1 | Red | PF3D7_1417200 | 36329.Q8ILS4 | NOT family protein, putative. |
| | 1 | Red | PF3D7_1417500 | 36329.Q8ILS0 | H/ACA ribonucleoprotein complex subunit 4, putative. |
| | 1 | Red | PF3D7_1420000 | 36329.Q8ILQ0 | Splicing factor 3B subunit 4, putative. |
| | 1 | Red | PF3D7_1422400 | 36329.Q8ILM9 | Uncharacterized protein. |
| | 1 | Red | PF3D7_1422800 | 36329.Q8ILM5 | Actin-related protein, putative; Belongs to the actin family. |
| | 1 | Red | PF3D7_1448000 | 36329.Q8IKZ5 | U3 small nucleolar RNA-associated protein 12, putative. |
| | 1 | Red | PF3D7_1451500 | 36329.Q8IKW1 | Pre-mRNA-splicing factor CWF18, putative. |
| | 1 | Red | PF3D7_1461600 | 36329.Q8IKL7 | Splicing factor 3B subunit 2, putative. |
| | 1 | Red | PF3D7_1464000 | 36329.Q8IKJ7 | YL1 nuclear protein, putative. |
| | 1 | Red | PF3D7_1472000 | 36329.Q8IKB8 | Pre-mRNA-splicing factor ISY1, putative. |
| | 1 | Red | PF3D7_1474500 | 36329.Q8IK93 | Splicing factor 3A subunit 1, putative. |

| Cluster salmon | Cluster number | Cluster color | Gene ID | Protein identifier | Protein description |
|---|---|---|---|---|---|
| | 2 | Salmon | PF3D7_0107800 | 36329.A0A143ZUM0 | Double-strand break repair protein MRE11. |
| | 2 | Salmon | PF3D7_0109200 | 36329.Q8I0V9 | Cleavage and polyadenylation specificity factor subunit 5; Component of the cleavage factor Im (CFIm) complex that functions as an activator of the pre-mRNA 3'-end cleavage and polyadenylation processing required for the maturation of pre-mRNA into functional mRNAs. CFIm contributes to the recruitment of multiprotein complexes on specific sequences on the pre-mRNA 3'-end, so called cleavage and polyadenylation signals (pA signals). Most pre-mRNAs contain multiple pA signals, resulting in alternative cleavage and polyadenylation (APA) producing mRNAs with variable 3'-end formation. The [...] |
| | 2 | Salmon | PF3D7_0110400 | 36329.Q8I241 | DNA-directed RNA polymerase II subunit RPB9, putative. |
| | 2 | Salmon | PF3D7_0111300 | 36329.Q8I233 | P-loop containing nucleoside triphosphate hydrolase, putative. |
| | 2 | Salmon | PF3D7_0215700 | 36329.O96236 | DNA-directed RNA polymerase subunit beta; DNA-dependent RNA polymerase catalyzes the transcription of DNA into RNA using the four ribonucleoside triphosphates as substrates. |
| | 2 | Salmon | PF3D7_0215800 | 36329.O96237 | Origin recognition complex subunit 5. |
| | 2 | Salmon | PF3D7_0218000 | 36329.O96260 | Replication factor C subunit 2, putative. |
| | 2 | Salmon | PF3D7_0219600 | 36329.O96271 | Replication factor C subunit 1. |
| | 2 | Salmon | PF3D7_0303300 | 36329.O77315 | DNA-directed RNA polymerases I, II, and III subunit RPABC2, putative. |
| | 2 | Salmon | PF3D7_0308000 | 36329.O77321 | DNA polymerase delta small subunit, putative. |
| | 2 | Salmon | PF3D7_0318200 | 36329.O77375 | DNA-directed RNA polymerase subunit; DNA-dependent RNA polymerase catalyzes the transcription of DNA into RNA using the four ribonucleoside triphosphates as substrates. |
| | 2 | Salmon | PF3D7_0318600 | 36329.O77371 | Cleavage and polyadenylation specificity factor, putative. |
| | 2 | Salmon | PF3D7_0408500 | 36329.Q7K734 | Flap endonuclease 1; Structure-specific nuclease with 5'-flap endonuclease and 5'- 3' exonuclease activities involved in DNA replication and repair. During DNA replication, cleaves the 5'-overhanging flap structure that is generated by displacement synthesis when DNA polymerase encounters the 5'-end of a downstream Okazaki fragment. It enters the flap from the 5'-end and then tracks to cleave the flap base, leaving a nick for ligation. Also involved in the long patch base excision repair (LP-BER) pathway, by cleaving within the apurinic/apyrimidinic (AP) site- terminated flap. Acts as [...] |
| | 2 | Salmon | PF3D7_0411900 | 36329.Q9U0H1 | DNA polymerase. |
| | 2 | Salmon | PF3D7_0416300 | 36329.Q8I1S4 | DNA helicase MCM9, putative; Belongs to the MCM family. |
| | 2 | Salmon | PF3D7_0416400 | 36329.C0H4A7 | Histone acetyltransferase, putative. |
| | 2 | Salmon | PF3D7_0501800 | 36329.Q8I482 | Protein HIRA; Required for replication-independent chromatin assembly and for the periodic repression of histone gene transcription during the cell cycle; Belongs to the WD repeat HIR1 family. |
| | 2 | Salmon | PF3D7_0503200 | 36329.Q8I469 | ATPase_AAA_core domain-containing protein. |
| | 2 | Salmon | PF3D7_0505500 | 36329.Q8I447 | DNA mismatch repair protein; Component of the post-replicative DNA mismatch repair system (MMR). |
| | 2 | Salmon | PF3D7_0505800 | 36329.Q8I444 | Small ubiquitin-related modifier. |
| | 2 | Salmon | PF3D7_0509400 | 36329.Q8I410 | DNA-directed RNA polymerase subunit; DNA-dependent RNA polymerase catalyzes the transcription of DNA into RNA using the four ribonucleoside triphosphates as substrates. |
| | 2 | Salmon | PF3D7_0513200 | 36329.Q8I3X5 | CPSF_A domain-containing protein. |
| | 2 | Salmon | PF3D7_0517400 | 36329.Q8I3T4 | FACT complex subunit SPT16, putative. |
| | 2 | Salmon | PF3D7_0520700 | 36329.Q8I3Q4 | CDC73 domain-containing protein, putative. |
| | 2 | Salmon | PF3D7_0526500 | 36329.Q8I3K0 | Suf domain-containing protein. |
| | 2 | Salmon | PF3D7_0527000 | 36329.Q8I3J5 | DNA replication licensing factor MCM3, putative; Belongs to the MCM family. |
| | 2 | Salmon | PF3D7_0605800 | 36329.C6KSQ6 | Probable DNA repair protein RAD50; Essential component of the MRN complex, a complex that possesses single-stranded DNA endonuclease and 3' to 5' exonuclease activities, and plays a central role in double-strand break (DSB) repair, chromosome morphogenesis, DNA repair and meiosis. In the complex, it mediates the ATP-binding and is probably required to bind DNA ends and hold them in close proximity (By similarity). Belongs to the SMC family. RAD50 subfamily. |
| | 2 | Salmon | PF3D7_0609900 | 36329.C6KSU6 | Uncharacterized protein. |
| | 2 | Salmon | PF3D7_0610900 | 36329.C6KSV4 | Transcription elongation factor SPT5, putative. |
| | 2 | Salmon | PF3D7_0612900 | 36329.C6KSX2 | Nucleolar GTP-binding protein 1; Involved in the biogenesis of the 60S ribosomal subunit. Belongs to the TRAFAC class OBG-HflX-like GTPase superfamily. OBG GTPase family. NOG subfamily. |
| | 2 | Salmon | PF3D7_0620500 | 36329.C6KT45 | Cleavage stimulation factor subunit 1, putative. |
| | 2 | Salmon | PF3D7_0625600 | 36329.C6KT92 | Poly(A) polymerase; Polymerase that creates the 3'-poly(A) tail of mRNA's. |
| | 2 | Salmon | PF3D7_0630300 | 36329.C6KTD8 | DNA polymerase epsilon catalytic subunit A, putative. |
| | 2 | Salmon | PF3D7_0705300 | 36329.Q8IC17 | Origin recognition complex subunit 2. |
| | 2 | Salmon | PF3D7_0705400 | 36329.Q8IC16 | DNA replication licensing factor MCM7; Acts as component of the mcm2-7 complex (mcm complex) which is the putative replicative helicase essential for 'once per cell cycle' DNA replication initiation and elongation in eukaryotic cells. The active ATPase sites in the mcm2-7 ring are formed through the interaction surfaces of two neighboring subunits such that a critical structure of a conserved arginine finger motif is provided in trans relative to the ATP-binding site of the Walker A box of the adjacent subunit. The six ATPase active sites, however, are likely to contribute differential [...] |
| | 2 | Salmon | PF3D7_0708100 | 36329.Q8IC08 | DNA-directed RNA polymerases I, II, and III subunit RPABC5, putative. |
| | 2 | Salmon | PF3D7_0714500 | 36329.Q8IBV2 | Transcription elongation factor s-II, putative. |
| | 2 | Salmon | PF3D7_0725000 | 36329.Q8IBK1 | Exonuclease I, putative. |
| | 2 | Salmon | PF3D7_0726300 | 36329.Q8IBJ3 | DNA mismatch repair protein PMS1, putative. |
| | 2 | Salmon | PF3D7_0801700 | 36329.C0H4Y4 | Sentrin-specific protease 2, putative. |
| | 2 | Salmon | PF3D7_0819000 | 36329.Q8IB25 | Cleavage and polyadenylation specificity factor subunit 2. |
| | 2 | Salmon | PF3D7_0821600 | 36329.Q8IB50 | Polyribonucleotide 5'-hydroxyl-kinase Clp1, putative. |
| | 2 | Salmon | PF3D7_0822100 | 36329.Q8IB55 | Mediator of RNA polymerase II transcription subunit 7; Component of the Mediator complex, a coactivator involved in the regulated transcription of nearly all RNA polymerase II-dependent genes. Mediator functions as a bridge to convey information from gene- specific regulatory proteins to the basal RNA polymerase II transcription machinery. |
| | 2 | Salmon | PF3D7_0910900 | 36329.C0H531 | DNA primase large subunit; DNA primase is the polymerase that synthesizes small RNA primers for the Okazaki fragments made during discontinuous DNA replication; Belongs to the eukaryotic-type primase large subunit family. |
| | 2 | Salmon | PF3D7_0922500 | 36329.P27362 | Phosphoglycerate kinase. |
| | 2 | Salmon | PF3D7_0923000 | 36329.Q8I2S6 | DNA-directed RNA polymerase II subunit RPB3, putative. |
| | 2 | Salmon | PF3D7_0923900 | 36329.Q8I2R8 | Polyadenylate-binding protein 2, putative. |
| | 2 | Salmon | PF3D7_0933000 | 36329.Q8I0W1 | CSTF domain-containing protein, putative. |
| | 2 | Salmon | PF3D7_0934100 | 36329.Q8I2H7 | TFIIH basal transcription factor complex helicase XPD subunit. |
| | 2 | Salmon | PF3D7_1009200 | 36329.Q8IJV3 | Ribonuclease, putative. |

| | | | | | |
|---|---|---|---|---|---|
| | 2 | Salmon | PF3D7_1037600 | 36329.Q8IJ31 | TFIIH basal transcription factor complex helicase XPB subunit, putative. |
| | 2 | Salmon | PF3D7_1103700 | 36329.Q8IIW5 | Casein kinase II subunit beta; Plays a complex role in regulating the basal catalytic activity of the alpha subunit; Belongs to the casein kinase 2 subunit beta family. |
| | 2 | Salmon | PF3D7_1108400 | 36329.Q8IIR9 | Casein kinase 2, alpha subunit; Belongs to the protein kinase superfamily. |
| | 2 | Salmon | PF3D7_1111100 | 36329.Q8IIQ1 | Replication factor C subunit 5, putative. |
| | 2 | Salmon | PF3D7_1112100 | 36329.Q8IIP2 | Protein kinase domain-containing protein. |
| | 2 | Salmon | PF3D7_1117800 | 36329.Q8IIJ0 | DNA mismatch repair protein MLH. |
| | 2 | Salmon | PF3D7_1120100 | 36329.Q8IIG6 | Phosphoglycerate mutase. |
| | 2 | Salmon | PF3D7_1134700 | 36329.Q8II17 | DNA-directed RNA polymerase subunit beta; DNA-dependent RNA polymerase catalyzes the transcription of DNA into RNA using the four ribonucleoside triphosphates as substrates. |
| | 2 | Salmon | PF3D7_1143300 | 36329.Q8IHT3 | DNA-directed RNA polymerases I and III subunit RPAC1, putative. |
| | 2 | Salmon | PF3D7_1205600 | 36329.Q8I5Y9 | Tetratricopeptide repeat protein, putative. |
| | 2 | Salmon | PF3D7_1206600 | 36329.Q8I5X9 | DNA-directed RNA polymerase subunit beta; DNA-dependent RNA polymerase catalyzes the transcription of DNA into RNA using the four ribonucleoside triphosphates as substrates. |
| | 2 | Salmon | PF3D7_1210400 | 36329.Q8I5U5 | General transcription factor 3C polypeptide 5, putative. |
| | 2 | Salmon | PF3D7_1211300 | 36329.Q8I5T7 | DNA helicase MCM8, putative; Belongs to the MCM family. |
| | 2 | Salmon | PF3D7_1211700 | 36329.Q8I5T4 | DNA replication licensing factor MCM5, putative; Belongs to the MCM family. |
| | 2 | Salmon | PF3D7_1213700 | 36329.Q8I5R8 | DNA-directed RNA polymerases I, II, and III subunit RPABC3; DNA-dependent RNA polymerase catalyzes the transcription of DNA into RNA using the four ribonucleoside triphosphates as substrates. Common component of RNA polymerases I, II and III which synthesize ribosomal RNA precursors, mRNA precursors and many functional non- coding RNAs, and small RNAs, such as 5S rRNA and tRNAs, respectively. |
| | 2 | Salmon | PF3D7_1224500 | 36329.Q8I5H2 | Histone chaperone ASF1, putative. |
| | 2 | Salmon | PF3D7_1241700 | 36329.Q8I512 | Replication factor C subunit 4, putative. |
| | 2 | Salmon | PF3D7_1244200 | 36329.Q8I4Y8 | General transcription factor IIH subunit 4; Component of the general transcription and DNA repair factor IIH (TFIIH) core complex which is involved in general and transcription-coupled nucleotide excision repair (NER) of damaged DNA. Belongs to the TFB2 family. |
| | 2 | Salmon | PF3D7_1305000 | 36329.Q8IER6 | MCL1 domain-containing protein, putative. |
| | 2 | Salmon | PF3D7_1314900 | 36329.Q8IEG6 | General transcription factor IIH subunit. |
| | 2 | Salmon | PF3D7_1317100 | 36329.Q8IEE5 | DNA replication licensing factor MCM4; Belongs to the MCM family. |
| | 2 | Salmon | PF3D7_1328200 | 36329.A0A5K1K982 | Uncharacterized protein. |
| | 2 | Salmon | PF3D7_1329000 | 36329.Q8IE49 | DNA-directed RNA polymerase subunit; DNA-dependent RNA polymerase catalyzes the transcription of DNA into RNA using the four ribonucleoside triphosphates as substrates. |
| | 2 | Salmon | PF3D7_1334100 | 36329.A0A5K1K8V1 | Uncharacterized protein. |
| | 2 | Salmon | PF3D7_1342400 | 36329.Q8IDR5 | Casein kinase II subunit beta; Plays a complex role in regulating the basal catalytic activity of the alpha subunit; Belongs to the casein kinase 2 subunit beta family. |
| | 2 | Salmon | PF3D7_1355100 | 36329.Q8IDF0 | DNA helicase; Belongs to the MCM family. |
| | 2 | Salmon | PF3D7_1361900 | 36329.P61074 | Proliferating cell nuclear antigen; This protein is an auxiliary protein of DNA polymerase delta and is involved in the control of eukaryotic DNA replication by increasing the polymerase's processibility during elongation of the leading strand. |
| | 2 | Salmon | PF3D7_1364800 | 36329.Q8ID59 | DNA-directed RNA polymerases I, II, and III subunit RPABC1, putative. |
| | 2 | Salmon | PF3D7_1406200 | 36329.Q8IM32 | Transcription elongation factor SPT6, putative. |
| | 2 | Salmon | PF3D7_1412100 | 36329.Q8ILX3 | Mini-chromosome maintenance complex-binding protein, putative. |
| | 2 | Salmon | PF3D7_1414400 | 36329.Q8ILV1 | Serine/threonine-protein phosphatase. |
| | 2 | Salmon | PF3D7_1415200 | 36329.Q8ILU4 | DNA-directed RNA polymerases I and III subunit RPAC2, putative. |
| | 2 | Salmon | PF3D7_1419900 | 36329.Q8ILQ1 | Cleavage and polyadenylation specificity factor subunit 4, putative. |
| | 2 | Salmon | PF3D7_1427500 | 36329.Q8ILI9 | DNA mismatch repair protein MSH2, putative. |
| | 2 | Salmon | PF3D7_1438500 | 36329.Q8IL83 | Cleavage and polyadenylation specificity factor subunit 3, putative. |
| | 2 | Salmon | PF3D7_1438700 | 36329.Q7KQM1 | DNA primase small subunit; DNA primase is the polymerase that synthesizes small RNA primers for the Okazaki fragments made during discontinuous DNA replication. |
| | 2 | Salmon | PF3D7_1441400 | 36329.Q8IL56 | FACT complex subunit SSRP1; Component of the FACT complex, a general chromatin factor that acts to reorganize nucleosomes. The FACT complex is involved in multiple processes that require DNA as a template such as mRNA elongation, DNA replication and DNA repair. During transcription elongation the FACT complex acts as a histone chaperone that both destabilizes and restores nucleosomal structure. It facilitates the passage of RNA polymerase II and transcription by promoting the dissociation of one histone H2A-H2B dimer from the nucleosome, then subsequently promotes the reestablishment o [...] |
| | 2 | Salmon | PF3D7_1458800 | 36329.Q8IKP3 | DNA-directed RNA polymerase III subunit RPC5, putative. |
| | 2 | Salmon | PF3D7_1463200 | 36329.Q8IKK4 | Replication factor C subunit 3, putative. |
| | 2 | Salmon | PF3D7_1463400 | 36329.Q8IKK2 | DNA-directed RNA polymerase III subunit RPC4, putative. |
| | 2 | Salmon | PF3D7_1466800 | 36329.Q8IKG9 | NOC3p domain-containing protein. |
| | 2 | Salmon | PF3D7_1469700 | 36329.Q8IKE0 | Mediator of RNA polymerase II transcription subunit 6; Component of the Mediator complex, a coactivator involved in the regulated transcription of nearly all RNA polymerase II-dependent genes. Mediator functions as a bridge to convey information from gene- specific regulatory proteins to the basal RNA polymerase II transcription machinery. Mediator is recruited to promoters by direct interactions with regulatory proteins and serves as a scaffold for the assembly of a functional preinitiation complex with RNA polymerase II and the general transcription factors. |
| | 2 | Salmon | PF3D7_1475000 | 36329.Q8IK88 | Mediator of RNA polymerase II transcription subunit 31; Component of the Mediator complex, a coactivator involved in the regulated transcription of nearly all RNA polymerase II-dependent genes. Mediator functions as a bridge to convey information from gene- specific regulatory proteins to the basal RNA polymerase II transcription machinery. Mediator is recruited to promoters by direct interactions with regulatory proteins and serves as a scaffold for the assembly of a functional preinitiation complex with RNA polymerase II and the general transcription factors. |

| Cluster fire brick | Cluster number | Cluster color | Gene ID | Protein identifier | Protein description |
|---|---|---|---|---|---|
| | 3 | Fire Brick | PF3D7_0111800 | 36329.B9ZSJ0 | Eukaryotic translation initiation factor 4E, putative; Belongs to the eukaryotic initiation factor 4E family. |
| | 3 | Fire Brick | PF3D7_0212300 | 36329.O96203 | Peptide chain release factor subunit 1, putative. |
| | 3 | Fire Brick | PF3D7_0302900 | 36329.O77312 | Exportin-1, putative. |
| | 3 | Fire Brick | PF3D7_0315100 | 36329.O97266 | Eukaryotic translation initiation factor 4E; Belongs to the eukaryotic initiation factor 4E family. |
| | 3 | Fire Brick | PF3D7_0418200 | 36329.Q8I1Q5 | Eukaryotic translation initiation factor 3 subunit M; Component of the eukaryotic translation initiation factor 3 (eIF-3) complex, which is involved in protein synthesis of a specialized repertoire of mRNAs and, together with other initiation factors, stimulates binding of mRNA and methionyl-tRNAi to the 40S ribosome. The eIF-3 complex specifically targets and initiates translation of a subset of mRNAs involved in cell proliferation. |
| | 3 | Fire Brick | PF3D7_0517700 | 36329.Q8I3T1 | Eukaryotic translation initiation factor 3 subunit B; Component of the eukaryotic translation initiation factor 3 (eIF-3) complex, which is involved in protein synthesis and, together with other initiation factors, stimulates binding of mRNA and methionyl-tRNAi to the 40S ribosome; Belongs to the eIF-3 subunit B family. |
| | 3 | Fire Brick | PF3D7_0524000 | 36329.Q8I3M5 | Karyopherin beta. |
| | 3 | Fire Brick | PF3D7_0528200 | 36329.Q8I3I5 | Eukaryotic translation initiation factor 3 subunit E; Component of the eukaryotic translation initiation factor 3 (eIF-3) complex, which is involved in protein synthesis of a specialized repertoire of mRNAs and, together with other initiation factors, stimulates binding of mRNA and methionyl-tRNAi to the 40S ribosome. The eIF-3 complex specifically targets and initiates translation of a subset of mRNAs involved in cell proliferation. |
| | 3 | Fire Brick | PF3D7_0612100 | 36329.C6KSW5 | Eukaryotic translation initiation factor 3 subunit L; Component of the eukaryotic translation initiation factor 3 (eIF-3) complex, which is involved in protein synthesis of a specialized repertoire of mRNAs and, together with other initiation factors, stimulates binding of mRNA and methionyl-tRNAi to the 40S ribosome. The eIF-3 complex specifically targets and initiates translation of a subset of mRNAs involved in cell proliferation. |
| | 3 | Fire Brick | PF3D7_0627700 | 36329.C6KTB3 | Transportin. |
| | 3 | Fire Brick | PF3D7_0716800 | 36329.Q8IBT2 | Eukaryotic translation initiation factor 3 subunit I; Component of the eukaryotic translation initiation factor 3 (eIF-3) complex, which is involved in protein synthesis of a specialized repertoire of mRNAs and, together with other initiation factors, stimulates binding of mRNA and methionyl-tRNAi to the 40S ribosome. The eIF-3 complex specifically targets and initiates translation of a subset of mRNAs involved in cell proliferation. |
| | 3 | Fire Brick | PF3D7_0728000 | 36329.Q8IBH7 | Eukaryotic translation initiation factor 2 subunit alpha. |
| | 3 | Fire Brick | PF3D7_0815200 | 36329.Q8IAY9 | Importin subunit beta, putative. |
| | 3 | Fire Brick | PF3D7_0815600 | 36329.Q8IAZ3 | Eukaryotic translation initiation factor 3 subunit G; RNA-binding component of the eukaryotic translation initiation factor 3 (eIF-3) complex, which is involved in protein synthesis of a specialized repertoire of mRNAs and, together with other initiation factors, stimulates binding of mRNA and methionyl-tRNAi to the 40S ribosome. The eIF-3 complex specifically targets and initiates translation of a subset of mRNAs involved in cell proliferation. This subunit can bind 18S rRNA. |
| | 3 | Fire Brick | PF3D7_0826700 | 36329.Q8IBA0 | Receptor for activated c kinase. |
| | 3 | Fire Brick | PF3D7_0828500 | 36329.Q8IBB6 | Translation initiation factor eIF-2B subunit alpha, putative; Belongs to the eIF-2B alpha/beta/delta subunits family. |
| | 3 | Fire Brick | PF3D7_0913200 | 36329.Q8I320 | Elongation factor 1-beta. |
| | 3 | Fire Brick | PF3D7_0918300 | 36329.Q8I2X0 | Eukaryotic translation initiation factor 3 subunit F, putative; Belongs to the eIF-3 subunit F family. |
| | 3 | Fire Brick | PF3D7_0932800 | 36329.Q8I2I8 | Importin alpha re-exporter, putative. |
| | 3 | Fire Brick | PF3D7_1007900 | 36329.Q8IJW4 | Eukaryotic translation initiation factor 3 subunit D; mRNA cap-binding component of the eukaryotic translation initiation factor 3 (eIF-3) complex, which is involved in protein synthesis of a specialized repertoire of mRNAs and, together with other initiation factors, stimulates binding of mRNA and methionyl-tRNAi to the 40S ribosome. The eIF-3 complex specifically targets and initiates translation of a subset of mRNAs involved in cell proliferation. In the eIF-3 complex, eif3d specifically recognizes and binds the 7- methylguanosine cap of a subset of mRNAs. |
| | 3 | Fire Brick | PF3D7_1010600 | 36329.Q8IJT9 | Eukaryotic translation initiation factor 2 subunit beta. |
| | 3 | Fire Brick | PF3D7_1107300 | 36329.Q8IIS9 | Polyadenylate-binding protein-interacting protein 1, putative. |
| | 3 | Fire Brick | PF3D7_1117700 | 36329.Q7KQK6 | GTP-binding nuclear protein; GTP-binding protein involved in nucleocytoplasmic transport. Required for the import of protein into the nucleus and also for RNA export. Involved in chromatin condensation and control of cell cycle. Belongs to the small GTPase superfamily. Ran family. |
| | 3 | Fire Brick | PF3D7_1206200 | 36329.Q8I5Y3 | Eukaryotic translation initiation factor 3 subunit C; Component of the eukaryotic translation initiation factor 3 (eIF-3) complex, which is involved in protein synthesis of a specialized repertoire of mRNAs and, together with other initiation factors, stimulates binding of mRNA and methionyl-tRNAi to the 40S ribosome. The eIF-3 complex specifically targets and initiates translation of a subset of mRNAs involved in cell proliferation. |
| | 3 | Fire Brick | PF3D7_1212700 | 36329.Q8I5S6 | Eukaryotic translation initiation factor 3 subunit A; RNA-binding component of the eukaryotic translation initiation factor 3 (eIF-3) complex, which is involved in protein synthesis of a specialized repertoire of mRNAs and, together with other initiation factors, stimulates binding of mRNA and methionyl-tRNAi to the 40S ribosome. The eIF-3 complex specifically targets and initiates translation of a subset of mRNAs involved in cell proliferation. |
| | 3 | Fire Brick | PF3D7_1213900 | 36329.A0A144A0U7 | W2 domain-containing protein. |
| | 3 | Fire Brick | PF3D7_1250600 | 36329.Q8I4S8 | Translation initiation factor eIF-2B subunit beta, putative; Belongs to the eIF-2B alpha/beta/delta subunits family. |
| | 3 | Fire Brick | PF3D7_1315900 | 36329.A0A5K1K8Q1 | Exportin-T; tRNA nucleus export receptor which facilitates tRNA translocation across the nuclear pore complex. Belongs to the exportin family. |
| | 3 | Fire Brick | PF3D7_1326400 | 36329.A0A5K1K977 | Translation initiation factor eIF-2B subunit gamma, putative. |
| | 3 | Fire Brick | PF3D7_1338300 | 36329.A0A5K1K967 | Elongation factor 1-gamma, putative. |
| | 3 | Fire Brick | PF3D7_1357000 | 36329.Q8I0P6 | Elongation factor 1-alpha; This protein promotes the GTP-dependent binding of aminoacyl- tRNA to the A-site of ribosomes during protein biosynthesis. |
| | 3 | Fire Brick | PF3D7_1368200 | 36329.Q8I6Z4 | ABC transporter E family member 1, putative. |
| | 3 | Fire Brick | PF3D7_1410600 | 36329.Q8ILY9 | Eukaryotic translation initiation factor 2 subunit gamma, putative. |
| | 3 | Fire Brick | PF3D7_1419700 | 36329.Q8ILQ3 | Uncharacterized protein. |
| | 3 | Fire Brick | PF3D7_1451100 | 36329.Q8IKW5 | Elongation factor 2. |
| | 3 | Fire Brick | PF3D7_1468700 | 36329.Q8IKF0 | Eukaryotic initiation factor 4A; Belongs to the DEAD box helicase family. |

| T-complex | Cluster number | Cluster color | Gene ID | Protein identifier | Protein description |
|---|---|---|---|---|---|
| | 4 | Brown | PF3D7_0214000 | 36329.O96220 | T-complex protein 1 subunit theta. |
| | 4 | Brown | PF3D7_0306800 | 36329.O97247 | T-complex protein 1 subunit beta. |
| | 4 | Brown | PF3D7_0308200 | 36329.O77323 | T-complex protein 1 subunit eta; Molecular chaperone; assists the folding of proteins upon ATP hydrolysis. Known to play a role, in vitro, in the folding of actin and tubulin (By similarity). |
| | 4 | Brown | PF3D7_0320300 | 36329.O97282 | T-complex protein 1 subunit epsilon. |
| | 4 | Brown | PF3D7_1132200 | 36329.Q8II43 | T-complex protein 1 subunit alpha. |
| | 4 | Brown | PF3D7_1229500 | 36329.Q8I5C4 | T-complex protein 1 subunit gamma. |
| | 4 | Brown | PF3D7_1357800 | 36329.C0H5I7 | T-complex protein 1 subunit delta. |

| Ubiquitin fusion degradation | Cluster number | Cluster color | Gene ID | Protein identifier | Protein description |
|---|---|---|---|---|---|
| | 5 | Dark Golden Rod | PF3D7_0507700 | 36329.Q8I426 | Nuclear protein localization protein 4, putative. |
| | 5 | Dark Golden Rod | PF3D7_0619400 | 36329.C6KT34 | Cell division cycle protein 48 homologue, putative. |
| | 5 | Dark Golden Rod | PF3D7_0726500 | 36329.Q8IBJ1 | Ubiquitin carboxyl-terminal hydrolase, putative; Belongs to the peptidase C19 family. |
| | 5 | Dark Golden Rod | PF3D7_1211800 | 36329.Q7KQK2 | Polyubiquitin. |
| | 5 | Dark Golden Rod | PF3D7_1418000 | 36329.Q8ILR6 | Ubiquitin fusion degradation protein 1, putative. |

| Ribonucleotide reductase complex | Cluster number | Cluster color | Gene ID | Protein identifier | Protein description |
|---|---|---|---|---|---|
| | 6 | Yellow | PF3D7_0206700 | 36329.Q7KWJ4 | Adenylosuccinate lyase; Belongs to the lyase 1 family. Adenylosuccinate lyase subfamily. |
| | 6 | Yellow | PF3D7_1008900 | 36329.Q8IJV6 | Adenylate kinase; Belongs to the adenylate kinase family. |
| | 6 | Yellow | PF3D7_1015800 | 36329.Q8IJN8 | Ribonucleoside-diphosphate reductase small chain, putative. |
| | 6 | Yellow | PF3D7_1405600 | 36329.Q8IM38 | Ribonucleoside-diphosphate reductase small chain, putative. |
| | 6 | Yellow | PF3D7_1437200 | 36329.Q8IL94 | Ribonucleoside-diphosphate reductase; Provides the precursors necessary for DNA synthesis. Catalyzes the biosynthesis of deoxyribonucleotides from the corresponding ribonucleotides. |

| Chromatin condensation | Cluster number | Cluster color | Gene ID | Protein identifier | Protein description |
|---|---|---|---|---|---|
| | 7 | Dark Golden Rod 2 | PF3D7_0509100 | 36329.Q8I413 | Structural maintenance of chromosomes protein 4, putative. |
| | 7 | Dark Golden Rod 2 | PF3D7_1304000 | 36329.C0H598 | Condensin complex subunit 2, putative. |
| | 7 | Dark Golden Rod 2 | PF3D7_1318400 | 36329.Q8IED2 | Structural maintenance of chromosomes protein 2; May play a role in the conversion of interphase chromatin into condensed chromosomes; Belongs to the SMC family. SMC2 subfamily. |
| | 7 | Dark Golden Rod 2 | PF3D7_1403100 | 36329.C6S3H2 | Condensin complex subunit 1, putative. |

| Prefoldin complex | Cluster number | Cluster color | Gene ID | Protein identifier | Protein description |
|---|---|---|---|---|---|
| | 8 | Khaki | PF3D7_0718500 | 36329.Q8IBR6 | Prefoldin subunit 3; Binds specifically to cytosolic chaperonin (c-CPN) and transfers target proteins to it. Binds to nascent polypeptide chain and promotes folding in an environment in which there are many competing pathways for nonnative proteins; Belongs to the prefoldin subunit alpha family. |
| | 8 | Khaki | PF3D7_0904500 | 36329.Q8I3A4 | Prefoldin subunit 4; Binds specifically to cytosolic chaperonin (c-CPN) and transfers target proteins to it. Binds to nascent polypeptide chain and promotes folding in an environment in which there are many competing pathways for nonnative proteins; Belongs to the prefoldin subunit beta family. |
| | 8 | Khaki | PF3D7_1128100 | 36329.Q8II82 | Prefoldin subunit 5, putative. |
| | 8 | Khaki | PF3D7_1416900 | 36329.Q8ILS7 | Prefoldin subunit 2, putative. |

| ribonuclease P complex | Cluster number | Cluster color | Gene ID | Protein identifier | Protein description |
|---|---|---|---|---|---|
| | 9 | Olive | PF3D7_0621500 | 36329.C6KT53 | Ribonuclease P/MRP protein subunit RPP1, putative. |
| | 9 | Olive | PF3D7_0627900 | 36329.C6KTB5 | Ribonuclease P protein subunit p29, putative. |
| | 9 | Olive | PF3D7_0709600 | 36329.C0H4M3 | Ribonucleases P/MRP protein subunit POP1, putative. |
| | 9 | Olive | PF3D7_1111700 | 36329.Q8IIP6 | Uncharacterized protein. |

| V-type proton ATPase | Cluster number | Cluster color | Gene ID | Protein identifier | Protein description |
| --- | --- | --- | --- | --- | --- |
| | 10 | Green | PF3D7_0406100 | 36329.Q6ZMA8 | Vacuolar proton pump subunit B; Non-catalytic subunit of the peripheral V1 complex of vacuolar ATPase; Belongs to the ATPase alpha/beta chains family. |
| | 10 | Green | PF3D7_1311900 | 36329.Q76NM6 | V-type proton ATPase catalytic subunit A; Catalytic subunit of the peripheral V1 complex of vacuolar ATPase. V-ATPase vacuolar ATPase is responsible for acidifying a variety of intracellular compartments in eukaryotic cells; Belongs to the ATPase alpha/beta chains family. |
| | 10 | Green | PF3D7_1341900 | 36329.Q8IDS0 | V-type proton ATPase subunit D, putative. |
| | 10 | Green | PF3D7_1464700 | 36329.Q8IKJ0 | V-type proton ATPase subunit; Subunit of the integral membrane V0 complex of vacuolar ATPase. Vacuolar ATPase is responsible for acidifying a variety of intracellular compartments in eukaryotic cells, thus providing most of the energy required for transport processes in the vacuolar system. Belongs to the V-ATPase V0D/AC39 subunit family. |

| #Clustering method | Cluster number | Cluster color | Gene count | Gene ID | Protein identifier | Protein description |
|---|---|---|---|---|---|---|
| kmeans | 1 | Red | 102 | LSM3 | 36329.C0H4W2 | U6 snRNA-associated Sm-like protein LSm3; Binds specifically to the 3'-terminal U-tract of U6 snRNA. |
| kmeans | 1 | Red | 102 | LSM4 | 36329.Q8IIT3 | U6 snRNA-associated Sm-like protein LSm4; Binds specifically to the 3'-terminal U-tract of U6 snRNA. |
| kmeans | 1 | Red | 102 | PF08_0048 | 36329.C0H4W3 | Probable ATP-dependent helicase PF08_0048; Catalytic component of a chromatin remodeling complex. Belongs to the SNF2/RAD54 helicase family. SWR1 subfamily. |
| kmeans | 1 | Red | 102 | PF08_0084 | 36329.Q8IAW3 | U1 small nuclear ribonucleoprotein C; Component of the spliceosomal U1 snRNP, which is essential for recognition of the pre-mRNA 5' splice-site and the subsequent assembly of the spliceosome. U1-C is directly involved in initial 5' splice-site recognition for both constitutive and regulated alternative splicing. The interaction with the 5' splice-site seems to precede base-pairing between the pre-mRNA and the U1 snRNA. Stimulates commitment or early (E) complex formation by stabilizing the base pairing of the 5' end of the U1 snRNA and the 5' splice-site region. |
| kmeans | 1 | Red | 102 | PF3D7_0209200 | 36329.O96177 | Exosome complex component MTR3, putative. |
| kmeans | 1 | Red | 102 | PF3D7_0209800 | 36329.Q9TY94 | ATP-dependent RNA helicase UAP56. |
| kmeans | 1 | Red | 102 | PF3D7_0218500 | 36329.O96265 | Small nuclear ribonucleoprotein Sm D2. |
| kmeans | 1 | Red | 102 | PF3D7_0218700 | 36329.O96267 | Pre-mRNA-processing protein 45, putative. |
| kmeans | 1 | Red | 102 | PF3D7_0302000 | 36329.O97334 | Pre-mRNA-splicing factor PRP46, putative. |
| kmeans | 1 | Red | 102 | PF3D7_0302100 | 36329.O77306 | Serine/threonine protein kinase. |
| kmeans | 1 | Red | 102 | PF3D7_0308600 | 36329.O77325 | Pre-mRNA-processing factor 19, putative. |
| kmeans | 1 | Red | 102 | PF3D7_0319500 | 36329.O97318 | RNA-binding protein, putative. |
| kmeans | 1 | Red | 102 | PF3D7_0320800 | 36329.O97285 | ATP-dependent RNA helicase DDX6; Belongs to the DEAD box helicase family. |
| kmeans | 1 | Red | 102 | PF3D7_0323700 | 36329.O97303 | U4/U6.U5 tri-snRNP-associated protein 1, putative. |
| kmeans | 1 | Red | 102 | PF3D7_0403700 | 36329.Q8I1Z2 | Pre-mRNA-splicing factor CLF1, putative. |
| kmeans | 1 | Red | 102 | PF3D7_0405400 | 36329.Q8I1X5 | Pre-mRNA-processing-splicing factor 8, putative. |
| kmeans | 1 | Red | 102 | PF3D7_0410400 | 36329.Q9U0I4 | Exosome complex component RRP4. |
| kmeans | 1 | Red | 102 | PF3D7_0422500 | 36329.Q8IFP1 | Pre-mRNA-splicing helicase BRR2, putative. |
| kmeans | 1 | Red | 102 | PF3D7_0422700 | 36329.Q8IFN9 | Eukaryotic initiation factor 4A-III, putative; Belongs to the DEAD box helicase family. |
| kmeans | 1 | Red | 102 | PF3D7_0505200 | 36329.Q8I450 | Actin-like protein, putative; Belongs to the actin family. |
| kmeans | 1 | Red | 102 | PF3D7_0520300 | 36329.Q8I3Q7 | U6 snRNA-associated Sm-like protein LSm2; Binds specifically to the 3'-terminal U-tract of U6 snRNA. |
| kmeans | 1 | Red | 102 | PF3D7_0522800 | 36329.Q8I3N5 | Pre-mRNA-splicing factor BUD31, putative. |
| kmeans | 1 | Red | 102 | PF3D7_0528700 | 36329.Q8I3I0 | Peptidyl-prolyl cis-trans isomerase; PPIases accelerate the folding of proteins. It catalyzes the cis-trans isomerization of proline imidic peptide bonds in oligopeptides; Belongs to the cyclophilin-type PPIase family. |
| kmeans | 1 | Red | 102 | PF3D7_0602100 | 36329.C6KSM1 | ATP-dependent RNA helicase, putative. |
| kmeans | 1 | Red | 102 | PF3D7_0610200 | 36329.C6KSU9 | RNA-binding protein 25, putative. |
| kmeans | 1 | Red | 102 | PF3D7_0614400 | 36329.C6KSY5 | Pre-mRNA-splicing factor CWF7, putative. |
| kmeans | 1 | Red | 102 | PF3D7_0619900 | 36329.C6KT39 | Splicing factor 3A subunit 2, putative. |
| kmeans | 1 | Red | 102 | PF3D7_0623600 | 36329.C6KT72 | Transcription or splicing factor-like protein, putative. |
| kmeans | 1 | Red | 102 | PF3D7_0624600 | 36329.C6KT82 | SNF2 helicase, putative. |
| kmeans | 1 | Red | 102 | PF3D7_0628600 | 36329.C6KTC1 | DNA methyltransferase 1-associated protein 1, putative. |
| kmeans | 1 | Red | 102 | PF3D7_0716000 | 36329.Q8IBU0 | RNA-binding protein, putative. |
| kmeans | 1 | Red | 102 | PF3D7_0719300 | 36329.Q8IBQ9 | Actin-related protein, putative; Belongs to the actin family. |
| kmeans | 1 | Red | 102 | PF3D7_0722500 | 36329.Q8IBM8 | Pre-mRNA-splicing factor CWC15, putative. |
| kmeans | 1 | Red | 102 | PF3D7_0802300 | 36329.Q8IAM3 | Periodic tryptophan protein 2, putative. |
| kmeans | 1 | Red | 102 | PF3D7_0811300 | 36329.C0H4T9 | CCR4-associated factor 1. |
| kmeans | 1 | Red | 102 | PF3D7_0822300 | 36329.Q8IB57 | Small nuclear ribonucleoprotein G. |
| kmeans | 1 | Red | 102 | PF3D7_0822800 | 36329.C0H4X1 | U5 small nuclear ribonucleoprotein 40 kDa protein, putative. |
| kmeans | 1 | Red | 102 | PF3D7_0909800 | 36329.C0H529 | Small nuclear ribonucleoprotein Sm D3. |
| kmeans | 1 | Red | 102 | PF3D7_0917600 | 36329.Q8I2X7 | Pre-mRNA-splicing factor ATP-dependent RNA helicase PRP43, putative. |
| kmeans | 1 | Red | 102 | PF3D7_0924700 | 36329.Q8I2R0 | Splicing factor 3A subunit 3, putative. |
| kmeans | 1 | Red | 102 | PF3D7_0935000 | 36329.Q8I2G9 | U2 small nuclear ribonucleoprotein B'', putative. |
| kmeans | 1 | Red | 102 | PF3D7_1003800 | 36329.Q8IJZ9 | U5 small nuclear ribonucleoprotein component, putative. |
| kmeans | 1 | Red | 102 | PF3D7_1008100 | 36329.Q8IJW2 | Zinc finger protein, putative. |
| kmeans | 1 | Red | 102 | PF3D7_1008800 | 36329.Q8IJV7 | Nucleolar protein 5, putative. |
| kmeans | 1 | Red | 102 | PF3D7_1013100 | 36329.Q8IJR4 | U3 small nucleolar RNA-associated protein 13, putative. |
| kmeans | 1 | Red | 102 | PF3D7_1014600 | 36329.Q8IJP9 | Transcriptional coactivator ADA2. |
| kmeans | 1 | Red | 102 | PF3D7_1020700 | 36329.Q8IJJ5 | RNA cytidine acetyltransferase; RNA cytidine acetyltransferase with specificity toward both 18S rRNA and tRNAs. Catalyzes the formation of N(4)-acetylcytidine (ac4C) in 18S rRNA. Required for early nucleolar cleavages of precursor rRNA at sites A0, A1 and A2 during 18S rRNA synthesis. Catalyzes the formation of ac4C in serine and leucine tRNAs. Requires a tRNA-binding adapter protein for full tRNA acetyltransferase activity but not for 18S rRNA acetylation. |
| kmeans | 1 | Red | 102 | PF3D7_1033600 | 36329.Q7KQL1 | Pre-mRNA-splicing factor CEF1, putative. |
| kmeans | 1 | Red | 102 | PF3D7_1103600 | 36329.Q8IIW6 | Actin-like protein, putative; Belongs to the actin family. |
| kmeans | 1 | Red | 102 | PF3D7_1103800 | 36329.Q8IIW4 | CCR4-NOT transcription complex subunit 1, putative. |
| kmeans | 1 | Red | 102 | PF3D7_1106000 | 36329.Q8IIU3 | RuvB-like helicase; Belongs to the RuvB family. |
| kmeans | 1 | Red | 102 | PF3D7_1110200 | 36329.Q8IIR0 | Pre-mRNA-processing factor 6, putative. |
| kmeans | 1 | Red | 102 | PF3D7_1118500 | 36329.Q8III3 | Nucleolar protein 56, putative. |
| kmeans | 1 | Red | 102 | PF3D7_1119300 | 36329.Q8IIH4 | Splicing factor U2AF small subunit, putative. |
| kmeans | 1 | Red | 102 | PF3D7_1125500 | 36329.Q8IIA8 | Small nuclear ribonucleoprotein Sm D1; Essential for pre-mRNA splicing. Implicated in the formation of stable, biologically active snRNP structures. Belongs to the snRNP core protein family. |
| kmeans | 1 | Red | 102 | PF3D7_1126900 | 36329.Q8II94 | Small nuclear ribonucleoprotein F, putative. |
| kmeans | 1 | Red | 102 | PF3D7_1207100 | 36329.Q8I5X4 | Small subunit rRNA processing factor, putative. |
| kmeans | 1 | Red | 102 | PF3D7_1208100 | 36329.Q8I5W6 | Uncharacterized protein. |
| kmeans | 1 | Red | 102 | PF3D7_1209200 | 36329.Q8I5V6 | U6 snRNA-associated Sm-like protein LSm7, putative. |
| kmeans | 1 | Red | 102 | PF3D7_1220100 | 36329.Q8I5L1 | Pre-mRNA-processing factor 17, putative. |

| | | | | | | |
|---|---|---|---|---|---|---|
| kmeans | 1 | Red | 102 | PF3D7_1224900 | 36329.Q8I5G8 | Splicing factor 3B subunit 6, putative. |
| kmeans | 1 | Red | 102 | PF3D7_1230800 | 36329.Q8I5B2 | Pre-mRNA-splicing regulator, putative. |
| kmeans | 1 | Red | 102 | PF3D7_1231600 | 36329.Q8I5A4 | Pre-mRNA-splicing factor ATP-dependent RNA helicase PRP2, putative. |
| kmeans | 1 | Red | 102 | PF3D7_1234800 | 36329.Q8I574 | Splicing factor 3B subunit 3, putative. |
| kmeans | 1 | Red | 102 | PF3D7_1235300 | 36329.Q8I569 | CCR4-NOT transcription complex subunit 4, putative. |
| kmeans | 1 | Red | 102 | PF3D7_1235900 | 36329.A0A144A2S8 | Pre-mRNA-splicing factor SYF1, putative. |
| kmeans | 1 | Red | 102 | PF3D7_1249800 | 36329.Q8I4T6 | THO complex subunit 2, putative. |
| kmeans | 1 | Red | 102 | PF3D7_1303800 | 36329.Q8IES7 | Uncharacterized protein. |
| kmeans | 1 | Red | 102 | PF3D7_1305400 | 36329.Q8IER1 | AAR2 protein, putative. |
| kmeans | 1 | Red | 102 | PF3D7_1306900 | 36329.C0H5A4 | U1 small nuclear ribonucleoprotein A, putative. |
| kmeans | 1 | Red | 102 | PF3D7_1307000 | 36329.Q8IEP5 | Exosome complex component RRP40, putative. |
| kmeans | 1 | Red | 102 | PF3D7_1307100 | 36329.Q8IEP4 | U3 small nucleolar RNA-associated protein 6, putative. |
| kmeans | 1 | Red | 102 | PF3D7_1308900 | 36329.Q8IEM5 | mRNA-decapping enzyme 2, putative. |
| kmeans | 1 | Red | 102 | PF3D7_1309300 | 36329.Q8IEM1 | U4/U6 small nuclear ribonucleoprotein PRP3, putative. |
| kmeans | 1 | Red | 102 | PF3D7_1316500 | 36329.A0A5K1K8A8 | Pre-mRNA-processing factor 40, putative. |
| kmeans | 1 | Red | 102 | PF3D7_1316900 | 36329.C0H5C0 | Uncharacterized protein. |
| kmeans | 1 | Red | 102 | PF3D7_1321700 | 36329.Q8IE99 | Splicing factor 1. |
| kmeans | 1 | Red | 102 | PF3D7_1324200 | 36329.Q8IE75 | Micro-fibrillar-associated protein, putative. |
| kmeans | 1 | Red | 102 | PF3D7_1333600 | 36329.Q8IDZ2 | U3 small nucleolar RNA-associated protein 4, putative. |
| kmeans | 1 | Red | 102 | PF3D7_1340100 | 36329.A0A5K1K8M9 | Exosome complex component RRP42, putative. |
| kmeans | 1 | Red | 102 | PF3D7_1343900 | 36329.C0H5G7 | U4/U6 small nuclear ribonucleoprotein PRP4, putative. |
| kmeans | 1 | Red | 102 | PF3D7_1352700 | 36329.Q8IDH3 | Intron-binding protein aquarius, putative. |
| kmeans | 1 | Red | 102 | PF3D7_1355800 | 36329.C0H5I5 | Splicing factor subunit; Belongs to the SF3B5 family. |
| kmeans | 1 | Red | 102 | PF3D7_1357700 | 36329.A0A5K1K903 | U3 small nucleolar RNA-associated protein 21, putative. |
| kmeans | 1 | Red | 102 | PF3D7_1362200 | 36329.Q8ID85 | RuvB-like helicase; Belongs to the RuvB family. |
| kmeans | 1 | Red | 102 | PF3D7_1364300 | 36329.A0A5K1K9B8 | Pre-mRNA-splicing factor ATP-dependent RNA helicase PRP16. |
| kmeans | 1 | Red | 102 | PF3D7_1364500 | 36329.Q8ID62 | Exosome complex component RRP45, putative. |
| kmeans | 1 | Red | 102 | PF3D7_1367100 | 36329.Q8ID37 | U1 small nuclear ribonucleoprotein 70 kDa homolog, putative. |
| kmeans | 1 | Red | 102 | PF3D7_1407100 | 36329.Q8IM23 | rRNA 2'-O-methyltransferase fibrillarin, putative. |
| kmeans | 1 | Red | 102 | PF3D7_1414800 | 36329.Q8ILU8 | Small nuclear ribonucleoprotein-associated protein B, putative. |
| kmeans | 1 | Red | 102 | PF3D7_1417200 | 36329.Q8ILS4 | NOT family protein, putative. |
| kmeans | 1 | Red | 102 | PF3D7_1417500 | 36329.Q8ILS0 | H/ACA ribonucleoprotein complex subunit 4, putative. |
| kmeans | 1 | Red | 102 | PF3D7_1420000 | 36329.Q8ILQ0 | Splicing factor 3B subunit 4, putative. |
| kmeans | 1 | Red | 102 | PF3D7_1422400 | 36329.Q8ILM9 | Uncharacterized protein. |
| kmeans | 1 | Red | 102 | PF3D7_1422800 | 36329.Q8ILM5 | Actin-related protein, putative; Belongs to the actin family. |
| kmeans | 1 | Red | 102 | PF3D7_1448000 | 36329.Q8IKZ5 | U3 small nucleolar RNA-associated protein 12, putative. |
| kmeans | 1 | Red | 102 | PF3D7_1451500 | 36329.Q8IKW1 | Pre-mRNA-splicing factor CWF18, putative. |
| kmeans | 1 | Red | 102 | PF3D7_1461600 | 36329.Q8IKL7 | Splicing factor 3B subunit 2, putative. |
| kmeans | 1 | Red | 102 | PF3D7_1464000 | 36329.Q8IKJ7 | YL1 nuclear protein, putative. |
| kmeans | 1 | Red | 102 | PF3D7_1472000 | 36329.Q8IKB8 | Pre-mRNA-splicing factor ISY1, putative. |
| kmeans | 1 | Red | 102 | PF3D7_1474500 | 36329.Q8IK93 | Splicing factor 3A subunit 1, putative. |
| kmeans | 1 | Red | 102 | PFL2310w | 36329.Q8I4V2 | Pre-mRNA-splicing factor PFL2310w; Involved in pre-mRNA splicing. Binds RNA (By similarity). |
| kmeans | 2 | Salmon | 93 | FEN1 | 36329.Q7K734 | Flap endonuclease 1; Structure-specific nuclease with 5'-flap endonuclease and 5'- 3' exonuclease activities involved in DNA replication and repair. During DNA replication, cleaves the 5'-overhanging flap structure that is generated by displacement synthesis when DNA polymerase encounters the 5'-end of a downstream Okazaki fragment. It enters the flap from the 5'-end and then tracks to cleave the flap base, leaving a nick for ligation. Also involved in the long patch base excision repair (LP-BER) pathway, by cleaving within the apurinic/apyrimidinic (AP) site-terminated flap. Acts as [...] |
| kmeans | 2 | Salmon | 93 | MCM7 | 36329.Q8IC16 | DNA replication licensing factor MCM7; Acts as component of the mcm2-7 complex (mcm complex) which is the putative replicative helicase essential for 'once per cell cycle' DNA replication initiation and elongation in eukaryotic cells. The active ATPase sites in the mcm2-7 ring are formed through the interaction surfaces of two neighboring subunits such that a critical structure of a conserved arginine finger motif is provided in trans relative to the ATP-binding site of the Walker A box of the adjacent subunit. The six ATPase active sites, however, are likely to contribute differential [...] |
| kmeans | 2 | Salmon | 93 | MED6 | 36329.Q8IKE0 | Mediator of RNA polymerase II transcription subunit 6; Component of the Mediator complex, a coactivator involved in the regulated transcription of nearly all RNA polymerase II-dependent genes. Mediator functions as a bridge to convey information from gene- specific regulatory proteins to the basal RNA polymerase II transcription machinery. Mediator is recruited to promoters by direct interactions with regulatory proteins and serves as a scaffold for the assembly of a functional preinitiation complex with RNA polymerase II and the general transcription factors. |
| kmeans | 2 | Salmon | 93 | PCNA | 36329.P61074 | Proliferating cell nuclear antigen; This protein is an auxiliary protein of DNA polymerase delta and is involved in the control of eukaryotic DNA replication by increasing the polymerase's processibility during elongation of the leading strand. |
| kmeans | 2 | Salmon | 93 | PF14_0366 | 36329.Q7KQM1 | DNA primase small subunit; DNA primase is the polymerase that synthesizes small RNA primers for the Okazaki fragments made during discontinuous DNA replication. |
| kmeans | 2 | Salmon | 93 | PF3D7_0107800 | 36329.A0A143ZUM0 | Double-strand break repair protein MRE11. |
| kmeans | 2 | Salmon | 93 | PF3D7_0109200 | 36329.Q8I0V9 | Cleavage and polyadenylation specificity factor subunit 5; Component of the cleavage factor Im (CFIm) complex that functions as an activator of the pre-mRNA 3'-end cleavage and polyadenylation processing required for the maturation of pre-mRNA into functional mRNAs. CFIm contributes to the recruitment of multiprotein complexes on specific sequences on the pre-mRNA 3'-end, so called cleavage and polyadenylation signals (pA signals). Most pre-mRNAs contain multiple pA signals, resulting in alternative cleavage and polyadenylation (APA) producing mRNAs with variable 3'-end formation. The [...] |
| kmeans | 2 | Salmon | 93 | PF3D7_0110400 | 36329.Q8I241 | DNA-directed RNA polymerase II subunit RPB9, putative. |
| kmeans | 2 | Salmon | 93 | PF3D7_0111300 | 36329.Q8I233 | P-loop containing nucleoside triphosphate hydrolase, putative. |
| kmeans | 2 | Salmon | 93 | PF3D7_0215700 | 36329.O96236 | DNA-directed RNA polymerase subunit beta; DNA-dependent RNA polymerase catalyzes the transcription of DNA into RNA using the four ribonucleoside triphosphates as substrates. |
| kmeans | 2 | Salmon | 93 | PF3D7_0215800 | 36329.O96237 | Origin recognition complex subunit 5. |
| kmeans | 2 | Salmon | 93 | PF3D7_0218000 | 36329.O96260 | Replication factor C subunit 2, putative. |

| | | | | | | |
|---|---|---|---|---|---|---|
| kmeans | 2 | Salmon | 93 | PF3D7_0219600 | 36329.O96271 | Replication factor C subunit 1. |
| kmeans | 2 | Salmon | 93 | PF3D7_0303300 | 36329.O77315 | DNA-directed RNA polymerases I, II, and III subunit RPABC2, putative. |
| kmeans | 2 | Salmon | 93 | PF3D7_0308000 | 36329.O77321 | DNA polymerase delta small subunit, putative. |
| kmeans | 2 | Salmon | 93 | PF3D7_0318200 | 36329.O77375 | DNA-directed RNA polymerase subunit; DNA-dependent RNA polymerase catalyzes the transcription of DNA into RNA using the four ribonucleoside triphosphates as substrates. |
| kmeans | 2 | Salmon | 93 | PF3D7_0318600 | 36329.O77371 | Cleavage and polyadenylation specificity factor, putative. |
| kmeans | 2 | Salmon | 93 | PF3D7_0411900 | 36329.Q9U0H1 | DNA polymerase. |
| kmeans | 2 | Salmon | 93 | PF3D7_0416300 | 36329.Q8I1S4 | DNA helicase MCM9, putative; Belongs to the MCM family. |
| kmeans | 2 | Salmon | 93 | PF3D7_0416400 | 36329.C0H4A7 | Histone acetyltransferase, putative. |
| kmeans | 2 | Salmon | 93 | PF3D7_0501800 | 36329.Q8I482 | Protein HIRA; Required for replication-independent chromatin assembly and for the periodic repression of histone gene transcription during the cell cycle; Belongs to the WD repeat HIR1 family. |
| kmeans | 2 | Salmon | 93 | PF3D7_0503200 | 36329.Q8I469 | ATPase_AAA_core domain-containing protein. |
| kmeans | 2 | Salmon | 93 | PF3D7_0505500 | 36329.Q8I447 | DNA mismatch repair protein; Component of the post-replicative DNA mismatch repair system (MMR). |
| kmeans | 2 | Salmon | 93 | PF3D7_0505800 | 36329.Q8I444 | Small ubiquitin-related modifier. |
| kmeans | 2 | Salmon | 93 | PF3D7_0509400 | 36329.Q8I410 | DNA-directed RNA polymerase subunit; DNA-dependent RNA polymerase catalyzes the transcription of DNA into RNA using the four ribonucleoside triphosphates as substrates. |
| kmeans | 2 | Salmon | 93 | PF3D7_0513200 | 36329.Q8I3X5 | CPSF_A domain-containing protein. |
| kmeans | 2 | Salmon | 93 | PF3D7_0517400 | 36329.Q8I3T4 | FACT complex subunit SPT16, putative. |
| kmeans | 2 | Salmon | 93 | PF3D7_0520700 | 36329.Q8I3Q4 | CDC73 domain-containing protein, putative. |
| kmeans | 2 | Salmon | 93 | PF3D7_0526500 | 36329.Q8I3K0 | Suf domain-containing protein. |
| kmeans | 2 | Salmon | 93 | PF3D7_0527000 | 36329.Q8I3J5 | DNA replication licensing factor MCM3, putative; Belongs to the MCM family. |
| kmeans | 2 | Salmon | 93 | PF3D7_0609900 | 36329.C6KSU6 | Uncharacterized protein. |
| kmeans | 2 | Salmon | 93 | PF3D7_0610900 | 36329.C6KSV4 | Transcription elongation factor SPT5, putative. |
| kmeans | 2 | Salmon | 93 | PF3D7_0612900 | 36329.C6KSX2 | Nucleolar GTP-binding protein 1; Involved in the biogenesis of the 60S ribosomal subunit. Belongs to the TRAFAC class OBG-HflX-like GTPase superfamily. OBG GTPase family. NOG subfamily. |
| kmeans | 2 | Salmon | 93 | PF3D7_0620500 | 36329.C6KT45 | Cleavage stimulation factor subunit 1, putative. |
| kmeans | 2 | Salmon | 93 | PF3D7_0625600 | 36329.C6KT92 | Poly(A) polymerase; Polymerase that creates the 3'-poly(A) tail of mRNA's. |
| kmeans | 2 | Salmon | 93 | PF3D7_0630300 | 36329.C6KTD8 | DNA polymerase epsilon catalytic subunit A, putative. |
| kmeans | 2 | Salmon | 93 | PF3D7_0705300 | 36329.Q8IC17 | Origin recognition complex subunit 2. |
| kmeans | 2 | Salmon | 93 | PF3D7_0708100 | 36329.Q8IC08 | DNA-directed RNA polymerases I, II, and III subunit RPABC5, putative. |
| kmeans | 2 | Salmon | 93 | PF3D7_0714500 | 36329.Q8IBV2 | Transcription elongation factor s-II, putative. |
| kmeans | 2 | Salmon | 93 | PF3D7_0725000 | 36329.Q8IBK1 | Exonuclease I, putative. |
| kmeans | 2 | Salmon | 93 | PF3D7_0726300 | 36329.Q8IBJ3 | DNA mismatch repair protein PMS1, putative. |
| kmeans | 2 | Salmon | 93 | PF3D7_0801700 | 36329.C0H4Y4 | Sentrin-specific protease 2, putative. |
| kmeans | 2 | Salmon | 93 | PF3D7_0819000 | 36329.Q8IB25 | Cleavage and polyadenylation specificity factor subunit 2. |
| kmeans | 2 | Salmon | 93 | PF3D7_0821600 | 36329.Q8IB50 | Polyribonucleotide 5'-hydroxyl-kinase Clp1, putative. |
| kmeans | 2 | Salmon | 93 | PF3D7_0822100 | 36329.Q8IB55 | Mediator of RNA polymerase II transcription subunit 7; Component of the Mediator complex, a coactivator involved in the regulated transcription of nearly all RNA polymerase II-dependent genes. Mediator functions as a bridge to convey information from gene- specific regulatory proteins to the basal RNA polymerase II transcription machinery. |
| kmeans | 2 | Salmon | 93 | PF3D7_0910900 | 36329.C0H531 | DNA primase large subunit; DNA primase is the polymerase that synthesizes small RNA primers for the Okazaki fragments made during discontinuous DNA replication; Belongs to the eukaryotic-type primase large subunit family. |
| kmeans | 2 | Salmon | 93 | PF3D7_0923000 | 36329.Q8I2S6 | DNA-directed RNA polymerase II subunit RPB3, putative. |
| kmeans | 2 | Salmon | 93 | PF3D7_0923900 | 36329.Q8I2R8 | Polyadenylate-binding protein 2, putative. |
| kmeans | 2 | Salmon | 93 | PF3D7_0933000 | 36329.Q8I0W1 | CSTF domain-containing protein, putative. |
| kmeans | 2 | Salmon | 93 | PF3D7_0934100 | 36329.Q8I2H7 | TFIIH basal transcription factor complex helicase XPD subunit. |
| kmeans | 2 | Salmon | 93 | PF3D7_1009200 | 36329.Q8IJV3 | Ribonuclease, putative. |
| kmeans | 2 | Salmon | 93 | PF3D7_1037600 | 36329.Q8IJ31 | TFIIH basal transcription factor complex helicase XPB subunit, putative. |
| kmeans | 2 | Salmon | 93 | PF3D7_1103700 | 36329.Q8IIW5 | Casein kinase II subunit beta; Plays a complex role in regulating the basal catalytic activity of the alpha subunit; Belongs to the casein kinase 2 subunit beta family. |
| kmeans | 2 | Salmon | 93 | PF3D7_1108400 | 36329.Q8IIR9 | Casein kinase 2, alpha subunit; Belongs to the protein kinase superfamily. |
| kmeans | 2 | Salmon | 93 | PF3D7_1111100 | 36329.Q8IIQ1 | Replication factor C subunit 5, putative. |
| kmeans | 2 | Salmon | 93 | PF3D7_1112100 | 36329.Q8IIP2 | Protein kinase domain-containing protein. |
| kmeans | 2 | Salmon | 93 | PF3D7_1117800 | 36329.Q8IIJ0 | DNA mismatch repair protein MLH. |
| kmeans | 2 | Salmon | 93 | PF3D7_1120100 | 36329.Q8IIG6 | Phosphoglycerate mutase. |
| kmeans | 2 | Salmon | 93 | PF3D7_1134700 | 36329.Q8II17 | DNA-directed RNA polymerase subunit beta; DNA-dependent RNA polymerase catalyzes the transcription of DNA into RNA using the four ribonucleoside triphosphates as substrates. |
| kmeans | 2 | Salmon | 93 | PF3D7_1143300 | 36329.Q8IHT3 | DNA-directed RNA polymerases I and III subunit RPAC1, putative. |
| kmeans | 2 | Salmon | 93 | PF3D7_1205600 | 36329.Q8I5Y9 | Tetratricopeptide repeat protein, putative. |
| kmeans | 2 | Salmon | 93 | PF3D7_1206600 | 36329.Q8I5X9 | DNA-directed RNA polymerase subunit beta; DNA-dependent RNA polymerase catalyzes the transcription of DNA into RNA using the four ribonucleoside triphosphates as substrates. |
| kmeans | 2 | Salmon | 93 | PF3D7_1210400 | 36329.Q8I5U5 | General transcription factor 3C polypeptide 5, putative. |
| kmeans | 2 | Salmon | 93 | PF3D7_1211300 | 36329.Q8I5T7 | DNA helicase MCM8, putative; Belongs to the MCM family. |
| kmeans | 2 | Salmon | 93 | PF3D7_1211700 | 36329.Q8I5T4 | DNA replication licensing factor MCM5, putative; Belongs to the MCM family. |
| kmeans | 2 | Salmon | 93 | PF3D7_1213700 | 36329.Q8I5R8 | DNA-directed RNA polymerases I, II, and III subunit RPABC3; DNA-dependent RNA polymerase catalyzes the transcription of DNA into RNA using the four ribonucleoside triphosphates as substrates. Common component of RNA polymerases I, II and III which synthesize ribosomal RNA precursors, mRNA precursors and many functional non- coding RNAs, and small RNAs, such as 5S rRNA and tRNAs, respectively. |
| kmeans | 2 | Salmon | 93 | PF3D7_1224500 | 36329.Q8I5H2 | Histone chaperone ASF1, putative. |
| kmeans | 2 | Salmon | 93 | PF3D7_1241700 | 36329.Q8I512 | Replication factor C subunit 4, putative. |

| | | | | | | |
|---|---|---|---|---|---|---|
| kmeans | 2 | Salmon | 93 | PF3D7_1244200 | 36329.Q8I4Y8 | General transcription factor IIH subunit 4; Component of the general transcription and DNA repair factor IIH (TFIIH) core complex which is involved in general and transcription-coupled nucleotide excision repair (NER) of damaged DNA. Belongs to the TFB2 family. |
| kmeans | 2 | Salmon | 93 | PF3D7_1305000 | 36329.Q8IER6 | MCL1 domain-containing protein, putative. |
| kmeans | 2 | Salmon | 93 | PF3D7_1314900 | 36329.Q8IEG6 | General transcription factor IIH subunit. |
| kmeans | 2 | Salmon | 93 | PF3D7_1317100 | 36329.Q8IEE5 | DNA replication licensing factor MCM4; Belongs to the MCM family. |
| kmeans | 2 | Salmon | 93 | PF3D7_1328200 | 36329.A0A5K1K982 | Uncharacterized protein. |
| kmeans | 2 | Salmon | 93 | PF3D7_1329000 | 36329.Q8IE49 | DNA-directed RNA polymerase subunit; DNA-dependent RNA polymerase catalyzes the transcription of DNA into RNA using the four ribonucleoside triphosphates as substrates. |
| kmeans | 2 | Salmon | 93 | PF3D7_1334100 | 36329.A0A5K1K8V1 | Uncharacterized protein. |
| kmeans | 2 | Salmon | 93 | PF3D7_1342400 | 36329.Q8IDR5 | Casein kinase II subunit beta; Plays a complex role in regulating the basal catalytic activity of the alpha subunit; Belongs to the casein kinase 2 subunit beta family. |
| kmeans | 2 | Salmon | 93 | PF3D7_1355100 | 36329.Q8IDF0 | DNA helicase; Belongs to the MCM family. |
| kmeans | 2 | Salmon | 93 | PF3D7_1364800 | 36329.Q8ID59 | DNA-directed RNA polymerases I, II, and III subunit RPABC1, putative. |
| kmeans | 2 | Salmon | 93 | PF3D7_1406200 | 36329.Q8IM32 | Transcription elongation factor SPT6, putative. |
| kmeans | 2 | Salmon | 93 | PF3D7_1412100 | 36329.Q8ILX3 | Mini-chromosome maintenance complex-binding protein, putative. |
| kmeans | 2 | Salmon | 93 | PF3D7_1414400 | 36329.Q8ILV1 | Serine/threonine-protein phosphatase. |
| kmeans | 2 | Salmon | 93 | PF3D7_1415200 | 36329.Q8ILU4 | DNA-directed RNA polymerases I and III subunit RPAC2, putative. |
| kmeans | 2 | Salmon | 93 | PF3D7_1419900 | 36329.Q8ILQ1 | Cleavage and polyadenylation specificity factor subunit 4, putative. |
| kmeans | 2 | Salmon | 93 | PF3D7_1427500 | 36329.Q8ILI9 | DNA mismatch repair protein MSH2, putative. |
| kmeans | 2 | Salmon | 93 | PF3D7_1438500 | 36329.Q8IL83 | Cleavage and polyadenylation specificity factor subunit 3, putative. |
| kmeans | 2 | Salmon | 93 | PF3D7_1441400 | 36329.Q8IL56 | FACT complex subunit SSRP1; Component of the FACT complex, a general chromatin factor that acts to reorganize nucleosomes. The FACT complex is involved in multiple processes that require DNA as a template such as mRNA elongation, DNA replication and DNA repair. During transcription elongation the FACT complex acts as a histone chaperone that both destabilizes and restores nucleosomal structure. It facilitates the passage of RNA polymerase II and transcription by promoting the dissociation of one histone H2A-H2B dimer from the nucleosome, then subsequently promotes the reestablishment o [...] |
| kmeans | 2 | Salmon | 93 | PF3D7_1458800 | 36329.Q8IKP3 | DNA-directed RNA polymerase III subunit RPC5, putative. |
| kmeans | 2 | Salmon | 93 | PF3D7_1463200 | 36329.Q8IKK4 | Replication factor C subunit 3, putative. |
| kmeans | 2 | Salmon | 93 | PF3D7_1463400 | 36329.Q8IKK2 | DNA-directed RNA polymerase III subunit RPC4, putative. |
| kmeans | 2 | Salmon | 93 | PF3D7_1466800 | 36329.Q8IKG9 | NOC3p domain-containing protein. |
| kmeans | 2 | Salmon | 93 | PF3D7_1475000 | 36329.Q8IK88 | Mediator of RNA polymerase II transcription subunit 31; Component of the Mediator complex, a coactivator involved in the regulated transcription of nearly all RNA polymerase II-dependent genes. Mediator functions as a bridge to convey information from gene- specific regulatory proteins to the basal RNA polymerase II transcription machinery. Mediator is recruited to promoters by direct interactions with regulatory proteins and serves as a scaffold for the assembly of a functional preinitiation complex with RNA polymerase II and the general transcription factors. |
| kmeans | 2 | Salmon | 93 | PFF0285c | 36329.C6KSQ6 | Probable DNA repair protein RAD50; Essential component of the MRN complex, a complex that possesses single-stranded DNA endonuclease and 3' to 5' exonuclease activities, and plays a central role in double-strand break (DSB) repair, chromosome morphogenesis, DNA repair and meiosis. In the complex, it mediates the ATP-binding and is probably required to bind DNA ends and hold them in close proximity (By similarity). Belongs to the SMC family. RAD50 subfamily. |
| kmeans | 2 | Salmon | 93 | PGK | 36329.P27362 | Phosphoglycerate kinase. |
| kmeans | 3 | Fire Brick | 36 | PF3D7_0111800 | 36329.B9ZSJ0 | Eukaryotic translation initiation factor 4E, putative; Belongs to the eukaryotic initiation factor 4E family. |
| kmeans | 3 | Fire Brick | 36 | PF3D7_0212300 | 36329.O96203 | Peptide chain release factor subunit 1, putative. |
| kmeans | 3 | Fire Brick | 36 | PF3D7_0302900 | 36329.O77312 | Exportin-1, putative. |
| kmeans | 3 | Fire Brick | 36 | PF3D7_0315100 | 36329.O97266 | Eukaryotic translation initiation factor 4E; Belongs to the eukaryotic initiation factor 4E family. |
| kmeans | 3 | Fire Brick | 36 | PF3D7_0418200 | 36329.Q8I1Q5 | Eukaryotic translation initiation factor 3 subunit M; Component of the eukaryotic translation initiation factor 3 (eIF-3) complex, which is involved in protein synthesis of a specialized repertoire of mRNAs and, together with other initiation factors, stimulates binding of mRNA and methionyl-tRNAi to the 40S ribosome. The eIF-3 complex specifically targets and initiates translation of a subset of mRNAs involved in cell proliferation. |
| kmeans | 3 | Fire Brick | 36 | PF3D7_0517700 | 36329.Q8I3T1 | Eukaryotic translation initiation factor 3 subunit B; Component of the eukaryotic translation initiation factor 3 (eIF-3) complex, which is involved in protein synthesis and, together with other initiation factors, stimulates binding of mRNA and methionyl-tRNAi to the 40S ribosome; Belongs to the eIF-3 subunit B family. |
| kmeans | 3 | Fire Brick | 36 | PF3D7_0524000 | 36329.Q8I3M5 | Karyopherin beta. |
| kmeans | 3 | Fire Brick | 36 | PF3D7_0528200 | 36329.Q8I3I5 | Eukaryotic translation initiation factor 3 subunit E; Component of the eukaryotic translation initiation factor 3 (eIF-3) complex, which is involved in protein synthesis of a specialized repertoire of mRNAs and, together with other initiation factors, stimulates binding of mRNA and methionyl-tRNAi to the 40S ribosome. The eIF-3 complex specifically targets and initiates translation of a subset of mRNAs involved in cell proliferation. |
| kmeans | 3 | Fire Brick | 36 | PF3D7_0612100 | 36329.C6KSW5 | Eukaryotic translation initiation factor 3 subunit L; Component of the eukaryotic translation initiation factor 3 (eIF-3) complex, which is involved in protein synthesis of a specialized repertoire of mRNAs and, together with other initiation factors, stimulates binding of mRNA and methionyl-tRNAi to the 40S ribosome. The eIF-3 complex specifically targets and initiates translation of a subset of mRNAs involved in cell proliferation. |
| kmeans | 3 | Fire Brick | 36 | PF3D7_0627700 | 36329.C6KTB3 | Transportin. |
| kmeans | 3 | Fire Brick | 36 | PF3D7_0716800 | 36329.Q8IBT2 | Eukaryotic translation initiation factor 3 subunit I; Component of the eukaryotic translation initiation factor 3 (eIF-3) complex, which is involved in protein synthesis of a specialized repertoire of mRNAs and, together with other initiation factors, stimulates binding of mRNA and methionyl-tRNAi to the 40S ribosome. The eIF-3 complex specifically targets and initiates translation of a subset of mRNAs involved in cell proliferation. |
| kmeans | 3 | Fire Brick | 36 | PF3D7_0728000 | 36329.Q8IBH7 | Eukaryotic translation initiation factor 2 subunit alpha. |
| kmeans | 3 | Fire Brick | 36 | PF3D7_0815200 | 36329.Q8IAY9 | Importin subunit beta, putative. |
| kmeans | 3 | Fire Brick | 36 | PF3D7_0815600 | 36329.Q8IAZ3 | Eukaryotic translation initiation factor 3 subunit G; RNA-binding component of the eukaryotic translation initiation factor 3 (eIF-3) complex, which is involved in protein synthesis of a specialized repertoire of mRNAs and, together with other initiation factors, stimulates binding of mRNA and methionyl-tRNAi to the 40S ribosome. The eIF-3 complex specifically targets and initiates translation of a subset of mRNAs involved in cell proliferation. This subunit can bind 18S rRNA. |
| kmeans | 3 | Fire Brick | 36 | PF3D7_0826700 | 36329.Q8IBA0 | Receptor for activated c kinase. |
| kmeans | 3 | Fire Brick | 36 | PF3D7_0828500 | 36329.Q8IBB6 | Translation initiation factor eIF-2B subunit alpha, putative; Belongs to the eIF-2B alpha/beta/delta subunits family. |
| kmeans | 3 | Fire Brick | 36 | PF3D7_0913200 | 36329.Q8I320 | Elongation factor 1-beta. |
| kmeans | 3 | Fire Brick | 36 | PF3D7_0918300 | 36329.Q8I2X0 | Eukaryotic translation initiation factor 3 subunit F, putative; Belongs to the eIF-3 subunit F family. |
| kmeans | 3 | Fire Brick | 36 | PF3D7_0932800 | 36329.Q8I2I8 | Importin alpha re-exporter, putative. |

| | | | | | | |
|---|---|---|---|---|---|---|
| kmeans | 3 | Fire Brick | 36 | PF3D7_1007900 | 36329.Q8IJW4 | Eukaryotic translation initiation factor 3 subunit D; mRNA cap-binding component of the eukaryotic translation initiation factor 3 (eIF-3) complex, which is involved in protein synthesis of a specialized repertoire of mRNAs and, together with other initiation factors, stimulates binding of mRNA and methionyl-tRNAi to the 40S ribosome. The eIF-3 complex specifically targets and initiates translation of a subset of mRNAs involved in cell proliferation. In the eIF-3 complex, eif3d specifically recognizes and binds the 7- methylguanosine cap of a subset of mRNAs. |
| kmeans | 3 | Fire Brick | 36 | PF3D7_1010600 | 36329.Q8IJT9 | Eukaryotic translation initiation factor 2 subunit beta. |
| kmeans | 3 | Fire Brick | 36 | PF3D7_1107300 | 36329.Q8IIS9 | Polyadenylate-binding protein-interacting protein 1, putative. |
| kmeans | 3 | Fire Brick | 36 | PF3D7_1117700 | 36329.Q7KQK6 | GTP-binding nuclear protein; GTP-binding protein involved in nucleocytoplasmic transport. Required for the import of protein into the nucleus and also for RNA export. Involved in chromatin condensation and control of cell cycle. Belongs to the small GTPase superfamily. Ran family. |
| kmeans | 3 | Fire Brick | 36 | PF3D7_1206200 | 36329.Q8I5Y3 | Eukaryotic translation initiation factor 3 subunit C; Component of the eukaryotic translation initiation factor 3 (eIF-3) complex, which is involved in protein synthesis of a specialized repertoire of mRNAs and, together with other initiation factors, stimulates binding of mRNA and methionyl-tRNAi to the 40S ribosome. The eIF-3 complex specifically targets and initiates translation of a subset of mRNAs involved in cell proliferation. |
| kmeans | 3 | Fire Brick | 36 | PF3D7_1212700 | 36329.Q8I5S6 | Eukaryotic translation initiation factor 3 subunit A; RNA-binding component of the eukaryotic translation initiation factor 3 (eIF-3) complex, which is involved in protein synthesis of a specialized repertoire of mRNAs and, together with other initiation factors, stimulates binding of mRNA and methionyl-tRNAi to the 40S ribosome. The eIF-3 complex specifically targets and initiates translation of a subset of mRNAs involved in cell proliferation. |
| kmeans | 3 | Fire Brick | 36 | PF3D7_1213900 | 36329.A0A144A0U7 | W2 domain-containing protein. |
| kmeans | 3 | Fire Brick | 36 | PF3D7_1250600 | 36329.Q8I4S8 | Translation initiation factor eIF-2B subunit beta, putative; Belongs to the eIF-2B alpha/beta/delta subunits family. |
| kmeans | 3 | Fire Brick | 36 | PF3D7_1315900 | 36329.A0A5K1K8Q1 | Exportin-T; tRNA nucleus export receptor which facilitates tRNA translocation across the nuclear pore complex. Belongs to the exportin family. |
| kmeans | 3 | Fire Brick | 36 | PF3D7_1326400 | 36329.A0A5K1K977 | Translation initiation factor eIF-2B subunit gamma, putative. |
| kmeans | 3 | Fire Brick | 36 | PF3D7_1338300 | 36329.A0A5K1K967 | Elongation factor 1-gamma, putative. |
| kmeans | 3 | Fire Brick | 36 | PF3D7_1357000 | 36329.Q8I0P6 | Elongation factor 1-alpha; This protein promotes the GTP-dependent binding of aminoacyl- tRNA to the A-site of ribosomes during protein biosynthesis. |
| kmeans | 3 | Fire Brick | 36 | PF3D7_1368200 | 36329.Q8I6Z4 | ABC transporter E family member 1, putative. |
| kmeans | 3 | Fire Brick | 36 | PF3D7_1410600 | 36329.Q8ILY9 | Eukaryotic translation initiation factor 2 subunit gamma, putative. |
| kmeans | 3 | Fire Brick | 36 | PF3D7_1419700 | 36329.Q8ILQ3 | Uncharacterized protein. |
| kmeans | 3 | Fire Brick | 36 | PF3D7_1451100 | 36329.Q8IKW5 | Elongation factor 2. |
| kmeans | 3 | Fire Brick | 36 | PF3D7_1468700 | 36329.Q8IKF0 | Eukaryotic initiation factor 4A; Belongs to the DEAD box helicase family. |
| kmeans | 4 | Brown | 7 | MAL3P3.6 | 36329.O77323 | T-complex protein 1 subunit eta; Molecular chaperone; assists the folding of proteins upon ATP hydrolysis. Known to play a role, in vitro, in the folding of actin and tubulin (By similarity). |
| kmeans | 4 | Brown | 7 | PF3D7_0214000 | 36329.O96220 | T-complex protein 1 subunit theta. |
| kmeans | 4 | Brown | 7 | PF3D7_0306800 | 36329.O97247 | T-complex protein 1 subunit beta. |
| kmeans | 4 | Brown | 7 | PF3D7_0320300 | 36329.O97282 | T-complex protein 1 subunit epsilon. |
| kmeans | 4 | Brown | 7 | PF3D7_1132200 | 36329.Q8II43 | T-complex protein 1 subunit alpha. |
| kmeans | 4 | Brown | 7 | PF3D7_1229500 | 36329.Q8I5C4 | T-complex protein 1 subunit gamma. |
| kmeans | 4 | Brown | 7 | PF3D7_1357800 | 36329.C0H5I7 | T-complex protein 1 subunit delta. |
| kmeans | 5 | Dark Golden Rod | 5 | PF3D7_0507700 | 36329.Q8I426 | Nuclear protein localization protein 4, putative. |
| kmeans | 5 | Dark Golden Rod | 5 | PF3D7_0619400 | 36329.C6KT34 | Cell division cycle protein 48 homologue, putative. |
| kmeans | 5 | Dark Golden Rod | 5 | PF3D7_0726500 | 36329.Q8IBJ1 | Ubiquitin carboxyl-terminal hydrolase, putative; Belongs to the peptidase C19 family. |
| kmeans | 5 | Dark Golden Rod | 5 | PF3D7_1211800 | 36329.Q7KQK2 | Polyubiquitin. |
| kmeans | 5 | Dark Golden Rod | 5 | PF3D7_1418000 | 36329.Q8ILR6 | Ubiquitin fusion degradation protein 1, putative. |
| kmeans | 6 | Yellow | 5 | PF3D7_0206700 | 36329.Q7KWJ4 | Adenylosuccinate lyase; Belongs to the lyase 1 family. Adenylosuccinate lyase subfamily. |
| kmeans | 6 | Yellow | 5 | PF3D7_1008900 | 36329.Q8IJV6 | Adenylate kinase; Belongs to the adenylate kinase family. |
| kmeans | 6 | Yellow | 5 | PF3D7_1015800 | 36329.Q8IJN8 | Ribonucleoside-diphosphate reductase small chain, putative. |
| kmeans | 6 | Yellow | 5 | PF3D7_1405600 | 36329.Q8IM38 | Ribonucleoside-diphosphate reductase small chain, putative. |
| kmeans | 6 | Yellow | 5 | PF3D7_1437200 | 36329.Q8IL94 | Ribonucleoside-diphosphate reductase; Provides the precursors necessary for DNA synthesis. Catalyzes the biosynthesis of deoxyribonucleotides from the corresponding ribonucleotides. |
| kmeans | 7 | Dark Golden Rod 2 | 4 | MAL13P1.96 | 36329.Q8IED2 | Structural maintenance of chromosomes protein 2; May play a role in the conversion of interphase chromatin into condensed chromosomes; Belongs to the SMC family. SMC2 subfamily. |
| kmeans | 7 | Dark Golden Rod 2 | 4 | PF3D7_0509100 | 36329.Q8I413 | Structural maintenance of chromosomes protein 4, putative. |
| kmeans | 7 | Dark Golden Rod 2 | 4 | PF3D7_1304000 | 36329.C0H598 | Condensin complex subunit 2, putative. |
| kmeans | 7 | Dark Golden Rod 2 | 4 | PF3D7_1403100 | 36329.C6S3H2 | Condensin complex subunit 1, putative. |
| kmeans | 8 | Khaki | 4 | PF3D7_0718500 | 36329.Q8IBR6 | Prefoldin subunit 3; Binds specifically to cytosolic chaperonin (c-CPN) and transfers target proteins to it. Binds to nascent polypeptide chain and promotes folding in an environment in which there are many competing pathways for nonnative proteins; Belongs to the prefoldin subunit alpha family. |
| kmeans | 8 | Khaki | 4 | PF3D7_0904500 | 36329.Q8I3A4 | Prefoldin subunit 4; Binds specifically to cytosolic chaperonin (c-CPN) and transfers target proteins to it. Binds to nascent polypeptide chain and promotes folding in an environment in which there are many competing pathways for nonnative proteins; Belongs to the prefoldin subunit beta family. |
| kmeans | 8 | Khaki | 4 | PF3D7_1128100 | 36329.Q8II82 | Prefoldin subunit 5, putative. |
| kmeans | 8 | Khaki | 4 | PF3D7_1416900 | 36329.Q8ILS7 | Prefoldin subunit 2, putative. |
| kmeans | 9 | Olive | 4 | PF3D7_0621500 | 36329.C6KT53 | Ribonuclease P/MRP protein subunit RPP1, putative. |
| kmeans | 9 | Olive | 4 | PF3D7_0627900 | 36329.C6KTB5 | Ribonuclease P protein subunit p29, putative. |
| kmeans | 9 | Olive | 4 | PF3D7_0709600 | 36329.C0H4M3 | Ribonucleases P/MRP protein subunit POP1, putative. |
| kmeans | 9 | Olive | 4 | PF3D7_1111700 | 36329.Q8IIP6 | Uncharacterized protein. |
| kmeans | 10 | Green | 4 | PF3D7_0406100 | 36329.Q6ZMA8 | Vacuolar proton pump subunit B; Non-catalytic subunit of the peripheral V1 complex of vacuolar ATPase; Belongs to the ATPase alpha/beta chains family. |
| kmeans | 10 | Green | 4 | PF3D7_1341900 | 36329.Q8IDS0 | V-type proton ATPase subunit D, putative. |
| kmeans | 10 | Green | 4 | PF3D7_1464700 | 36329.Q8IKJ0 | V-type proton ATPase subunit; Subunit of the integral membrane V0 complex of vacuolar ATPase. Vacuolar ATPase is responsible for acidifying a variety of intracellular compartments in eukaryotic cells, thus providing most of the energy required for transport processes in the vacuolar system. Belongs to the V-ATPase V0D/AC39 subunit family. |
| kmeans | 10 | Green | 4 | vapA | 36329.Q76NM6 | V-type proton ATPase catalytic subunit A; Catalytic subunit of the peripheral V1 complex of vacuolar ATPase. V-ATPase vacuolar ATPase is responsible for acidifying a variety of intracellular compartments in eukaryotic cells; Belongs to the ATPase alpha/beta chains family. |
| kmeans | 11 | Cyan | 3 | PF3D7_0409600 | 36329.Q9U0J0 | Replication protein A1, large subunit. |
| kmeans | 11 | Cyan | 3 | PF3D7_1132300 | 36329.Q8II42 | Nucleic acid binding protein, putative. |

| | | | | | | |
|---|---|---|---|---|---|---|
| kmeans | 11 | Cyan | 3 | PF3D7_1442100 | 36329.C6S3I6 | Replication factor A protein 3, putative. |
| kmeans | 12 | Sky Blue 3 | 3 | PF3D7_0810300 | 36329.C0H4T6 | Protein phosphatase PPM5, putative. |
| kmeans | 12 | Sky Blue 3 | 3 | PF3D7_0934800 | 36329.Q7K6A0 | cAMP-dependent protein kinase catalytic subunit; Belongs to the protein kinase superfamily. |
| kmeans | 12 | Sky Blue 3 | 3 | PF3D7_1223100 | 36329.Q7KQK0 | cAMP-dependent protein kinase regulatory subunit. |
| kmeans | 13 | Sky Blue | 3 | PF3D7_0517300 | 36329.Q8I3T5 | Serine/arginine-rich splicing factor 1. |
| kmeans | 13 | Sky Blue | 3 | PF3D7_1022400 | 36329.Q8IJI0 | Serine/arginine-rich splicing factor 4. |
| kmeans | 13 | Sky Blue | 3 | PF3D7_1119800 | 36329.Q8IIG9 | Alternative splicing factor ASF-1, putative. |
| kmeans | 14 | Sky Blue 2 | 3 | PF3D7_0818900 | 36329.Q8IB24 | Heat shock protein 70; Belongs to the heat shock protein 70 family. |
| kmeans | 14 | Sky Blue 2 | 3 | PF3D7_1216900 | 36329.Q8I5N9 | DNA-binding chaperone, putative. |
| kmeans | 14 | Sky Blue 2 | 3 | PF3D7_1247500 | 36329.Q8I4V7 | Serine/threonine protein kinase, putative. |
| kmeans | 15 | Cornflower Blue 2 | 3 | PF3D7_1130700 | 36329.Q8II57 | Structural maintenance of chromosomes protein 1, putative. |
| kmeans | 15 | Cornflower Blue 2 | 3 | PF3D7_1456500 | 36329.Q8IKR4 | SCD domain-containing protein. |
| kmeans | 15 | Cornflower Blue 2 | 3 | PFD0685c | 36329.Q8I1U7 | Structural maintenance of chromosomes protein 3 homolog; Central component of cohesin, a complex required for chromosome cohesion during the cell cycle. The cohesin complex may form a large proteinaceous ring within which sister chromatids can be trapped. At anaphase, the complex is cleaved and dissociates from chromatin, allowing sister chromatids to segregate. Cohesion is coupled to DNA replication and is involved in DNA repair. The cohesin complex plays also an important role in spindle pole assembly during mitosis and in chromosomes movement (By similarity); Belongs to the SMC fami [...] |
| kmeans | 16 | Blue | 2 | PF3D7_1230700 | 36329.Q8I5B3 | Protein transport protein SEC13; Belongs to the WD repeat SEC13 family. |
| kmeans | 16 | Blue | 2 | PF3D7_1361100 | 36329.C0H5J6 | Protein transport protein Sec24A. |
| kmeans | 17 | Cornflower Blue | 2 | PF3D7_0205400 | 36329.O96149 | PCI domain-containing protein, putative. |
| kmeans | 17 | Cornflower Blue | 2 | PF3D7_0602600 | 36329.C6KSM6 | SAC3 domain-containing protein, putative. |
| kmeans | 18 | Medium Slate Blue | 2 | PF3D7_0611400 | 36329.C6KSV9 | SWIB/MDM2 domain-containing protein. |
| kmeans | 18 | Medium Slate Blue | 2 | PF3D7_1225200 | 36329.Q8I5G5 | Uncharacterized protein. |
| kmeans | 19 | Medium Slate Blue 2 | 2 | PF10_0084 | 36329.Q7KQL5 | Tubulin beta chain; Tubulin is the major constituent of microtubules. It binds two moles of GTP, one at an exchangeable site on the beta chain and one at a non-exchangeable site on the alpha chain. |
| kmeans | 19 | Medium Slate Blue 2 | 2 | PF3D7_0903700 | 36329.Q6ZLZ9 | Tubulin alpha chain; Tubulin is the major constituent of microtubules. It binds two moles of GTP, one at an exchangeable site on the beta chain and one at a non-exchangeable site on the alpha chain. |
| kmeans | 20 | Medium Purple | 2 | PF3D7_0514100 | 36329.Q8I3W6 | ATP-dependent DNA helicase UvrD. |
| kmeans | 20 | Medium Purple | 2 | PF3D7_1107400 | 36329.Q8IIS8 | DNA repair protein RAD51 homolog; Binds to single and double-stranded DNA and exhibits DNA- dependent ATPase activity. Underwinds duplex DNA. Belongs to the RecA family. RAD51 subfamily. |
| kmeans | 21 | Medium Purple 2 | 2 | PF3D7_0802000 | 36329.Q8IAM0 | Glutamate dehydrogenase, putative. |
| kmeans | 21 | Medium Purple 2 | 2 | PF3D7_1345700 | 36329.Q8I6T2 | Isocitrate dehydrogenase [NADP]; Belongs to the isocitrate and isopropylmalate dehydrogenases family. |
| kmeans | 22 | Orchid 4 | 2 | PF3D7_0729500 | 36329.Q8I6Z2 | mRNA (N6-adenosine)-methyltransferase, putative; Belongs to the MT-A70-like family. |
| kmeans | 22 | Orchid 4 | 2 | PF3D7_1235500 | 36329.Q8I567 | mRNA methyltransferase, putative; Belongs to the MT-A70-like family. |
| kmeans | 23 | Orchid 2 | 2 | PF3D7_0810800 | 36329.Q8IAU3 | Hydroxymethyldihydropterin pyrophosphokinase-dihydropteroate synthase. |
| kmeans | 23 | Orchid 2 | 2 | PF3D7_1224000 | 36329.Q8I5H7 | GTP cyclohydrolase 1. |
| kmeans | 24 | Purple | 2 | PF3D7_1347100 | 36329.Q8IDM7 | DNA topoisomerase; Introduces a single-strand break via transesterification at a target site in duplex DNA. Releases the supercoiling and torsional tension of DNA introduced during the DNA replication and transcription by transiently cleaving and rejoining one strand of the DNA duplex. The scissile phosphodiester is attacked by the catalytic tyrosine of the enzyme, resulting in the formation of a DNA-(5'-phosphotyrosyl)-enzyme intermediate and the expulsion of a 3'-OH DNA strand. Belongs to the type IA topoisomerase family. |
| kmeans | 24 | Purple | 2 | PF3D7_1368800 | 36329.Q8ID22 | DNA repair endonuclease XPF, putative. |
| kmeans | 25 | Orchid | 2 | PF3D7_0804900 | 36329.Q8IAP4 | GTPase-activating protein, putative. |
| kmeans | 25 | Orchid | 2 | PF3D7_1145100 | 36329.Q8IHR6 | Coatomer subunit gamma; The coatomer is a cytosolic protein complex that binds to dilysine motifs and reversibly associates with Golgi non-clathrin- coated vesicles, which further mediate biosynthetic protein transport from the ER, via the Golgi up to the trans Golgi network. Coatomer complex is required for budding from Golgi membranes, and is essential for the retrograde Golgi-to-ER transport of dilysine-tagged proteins. |
| kmeans | 26 | Orchid 3 | 2 | PF3D7_0920800 | 36329.Q8I2U5 | Inosine-5'-monophosphate dehydrogenase; Catalyzes the conversion of inosine 5'-phosphate (IMP) to xanthosine 5'-phosphate (XMP), the first committed and rate-limiting step in the de novo synthesis of guanine nucleotides, and therefore plays an important role in the regulation of cell growth. |
| kmeans | 26 | Orchid 3 | 2 | PF3D7_1012400 | 36329.Q8IJS1 | Hypoxanthine phosphoribosyltransferase; Belongs to the purine/pyrimidine phosphoribosyltransferase family. |
| kmeans | 27 | Pink | 2 | PF3D7_1146600 | 36329.Q8IHQ2 | Oocyst rupture protein 1, putative. |
| kmeans | 27 | Pink | 2 | PF3D7_1439500 | 36329.Q8IL74 | Oocyst rupture protein 2, putative. |
| kmeans | 28 | Pale Violet Red | 2 | PF3D7_1342800 | 36329.Q8IDR1 | Phosphoenolpyruvate carboxykinase. |
| kmeans | 28 | Pale Violet Red | 2 | PF3D7_1426700 | 36329.Q8ILJ7 | Phosphoenolpyruvate carboxylase. |
| kmeans | 29 | Light Coral | 2 | PF3D7_0621800 | 36329.C6KT55 | Nascent polypeptide-associated complex subunit alpha, putative. |
| kmeans | 29 | Light Coral | 2 | PF3D7_1426100 | 36329.Q8ILK2 | Nascent polypeptide-associated complex subunit beta. |
