## Supplementary material for "*Plasmodium falciparum* SET10 is a histone H3 lysine K18 methyltransferase that participates in a chromatin modulation network crucial for intraerythrocytic development": Table S7

Table S7. STRING analysis of interactors shared between *Pf* SET10, *Pf* MORC, *Pf* AP2-P and PfISWI

### Interactors by bait

| PfSET10 interactors, nuclear proteins curated (Musabyimana et al., this study) | PfMORC interactors (Chahine et al., 2023) | PfMORC interactors (Singh et al., 2024) | PfISWI interactors (Bryant et al., 2020) | PfAP2-P 40 hpi interactors (Subudhi et al., 2023) | PfMORC interactors combined |
|---|---|---|---|---|---|
| **PF3D7_1221000** | **PF3D7_1468100** | **PF3D7_1468100** | **PF3D7_0624600** | **PF3D7_1107800** | |
| PF3D7_0105800 | PF3D7_0110500 | PF3D7_0102900 | PF3D7_0212300 | PF3D7_0102900 | PF3D7_0102900 |
| PF3D7_0107800 | PF3D7_0115600 | PF3D7_0103200 | PF3D7_0511800 | PF3D7_0103200 | PF3D7_0103200 |
| PF3D7_0109200 | PF3D7_0212500 | PF3D7_0105200 | PF3D7_0517400 | PF3D7_0103900 | PF3D7_0105200 |
| PF3D7_0110400 | PF3D7_0218500 | PF3D7_0108300 | PF3D7_0621800 | PF3D7_0104400 | PF3D7_0108300 |
| PF3D7_0110500 | PF3D7_0303400 | PF3D7_0109200 | PF3D7_0624600 | PF3D7_0105200 | PF3D7_0109200 |
| PF3D7_0111300 | PF3D7_0307700 | PF3D7_0110700 | PF3D7_0705400 | PF3D7_0105800 | PF3D7_0110500 |
| PF3D7_0111800 | PF3D7_0311800 | PF3D7_0112200 | PF3D7_0716800 | PF3D7_0106300 | PF3D7_0110700 |
| PF3D7_0205400 | PF3D7_0320300 | PF3D7_0202000 | PF3D7_0722400 | PF3D7_0107000 | PF3D7_0112200 |
| PF3D7_0205600 | PF3D7_0403600 | PF3D7_0214100 | PF3D7_0727400 | PF3D7_0108300 | PF3D7_0115600 |
| PF3D7_0205700 | PF3D7_0410800 | PF3D7_0215700 | PF3D7_0812400 | PF3D7_0108700 | PF3D7_0202000 |
| PF3D7_0206000 | PF3D7_0420300 | PF3D7_0217500 | PF3D7_0925700 | PF3D7_0110500 | PF3D7_0212500 |
| PF3D7_0206700 | PF3D7_0501500 | PF3D7_0304200 | PF3D7_1011800 | PF3D7_0111500 | PF3D7_0214100 |
| PF3D7_0209200 | PF3D7_0506500 | PF3D7_0308000 | PF3D7_1107800 | PF3D7_0111800 | PF3D7_0215700 |
| PF3D7_0209800 | PF3D7_0516800 | PF3D7_0308200 | PF3D7_1118500 | PF3D7_0205600 | PF3D7_0217500 |
| PF3D7_0210200 | PF3D7_0604100 | PF3D7_0310600 | PF3D7_1136500 | PF3D7_0205900 | PF3D7_0218500 |
| PF3D7_0212100 | PF3D7_0609000 | PF3D7_0318200 | PF3D7_1138500 | PF3D7_0209800 | PF3D7_0303400 |
| PF3D7_0212300 | PF3D7_0611400 | PF3D7_0318500 | PF3D7_1211700 | PF3D7_0210100 | PF3D7_0304200 |
| PF3D7_0212400 | PF3D7_0613800 | PF3D7_0401800 | PF3D7_1242800 | PF3D7_0210900 | PF3D7_0307700 |
| PF3D7_0212500 | PF3D7_0619900 | PF3D7_0405400 | PF3D7_1311800 | PF3D7_0211800 | PF3D7_0308000 |
| PF3D7_0213100 | PF3D7_0624600 | PF3D7_0419900 | PF3D7_1317100 | PF3D7_0212100 | PF3D7_0308200 |
| PF3D7_0214000 | PF3D7_0627800 | PF3D7_0420300 | PF3D7_1353800 | PF3D7_0212300 | PF3D7_0310600 |
| PF3D7_0215700 | PF3D7_0628100 | PF3D7_0420400 | PF3D7_1355100 | PF3D7_0213100 | PF3D7_0311800 |
| PF3D7_0215800 | PF3D7_0630600 | PF3D7_0505500 | PF3D7_1357800 | PF3D7_0214000 | PF3D7_0318200 |
| PF3D7_0216200 | PF3D7_0708800 | PF3D7_0505900 | PF3D7_1358800 | PF3D7_0214100 | PF3D7_0318500 |
| PF3D7_0217500 | PF3D7_0803700 | PF3D7_0513600 | PF3D7_1361900 | PF3D7_0215700 | PF3D7_0320300 |
| PF3D7_0218000 | PF3D7_0816600 | PF3D7_0519400 | PF3D7_1417800 | PF3D7_0217100 | PF3D7_0401800 |
| PF3D7_0218500 | PF3D7_0827800 | PF3D7_0519800 | PF3D7_1441400 | PF3D7_0217500 | PF3D7_0403600 |
| PF3D7_0218700 | PF3D7_0903500 | PF3D7_0522300 | PF3D7_1468100 | PF3D7_0217800 | PF3D7_0405400 |
| PF3D7_0219600 | PF3D7_0903700 | PF3D7_0527000 | PF3D7_1474800 | PF3D7_0218200 | PF3D7_0410800 |
| PF3D7_0302000 | PF3D7_0907400 | PF3D7_0528100 | | PF3D7_0219400 | PF3D7_0419900 |
| PF3D7_0302100 | PF3D7_0910200 | PF3D7_0528200 | | PF3D7_0219600 | PF3D7_0420300 |
| PF3D7_0302900 | PF3D7_0923000 | PF3D7_0604500 | | PF3D7_0301600 | PF3D7_0420400 |
| PF3D7_0303000 | PF3D7_0925700 | PF3D7_0607000 | | PF3D7_0302500 | PF3D7_0501500 |
| PF3D7_0303100 | PF3D7_0926100 | PF3D7_0610800 | | PF3D7_0302900 | PF3D7_0505500 |
| PF3D7_0303300 | PF3D7_0935400 | PF3D7_0610900 | | PF3D7_0303700 | PF3D7_0505900 |
| PF3D7_0305600 | PF3D7_1002400 | PF3D7_0613800 | | PF3D7_0304100 | PF3D7_0506500 |
| PF3D7_0306100 | PF3D7_1003600 | PF3D7_0621800 | | PF3D7_0304400 | PF3D7_0513600 |
| PF3D7_0306800 | PF3D7_1008400 | PF3D7_0621900 | | PF3D7_0304500 | PF3D7_0516800 |
| PF3D7_0307700 | PF3D7_1014100 | PF3D7_0624600 | | PF3D7_0305500 | PF3D7_0519400 |
| PF3D7_0308000 | PF3D7_1020700 | PF3D7_0627700 | | PF3D7_0305600 | PF3D7_0519800 |
| PF3D7_0308100 | PF3D7_1106000 | PF3D7_0706000 | | PF3D7_0306400 | PF3D7_0522300 |
| PF3D7_0308200 | PF3D7_1107800 | PF3D7_0709000 | | PF3D7_0306800 | PF3D7_0527000 |
| PF3D7_0308600 | PF3D7_1135400 | PF3D7_0714500 | | PF3D7_0306900 | PF3D7_0528100 |
| PF3D7_0309000 | PF3D7_1139300 | PF3D7_0717700 | | PF3D7_0307100 | PF3D7_0528200 |
| PF3D7_0309200 | PF3D7_1148000 | PF3D7_0801000 | | PF3D7_0307200 | PF3D7_0604100 |
| PF3D7_0309300 | PF3D7_1202200 | PF3D7_0802000 | | PF3D7_0308200 | PF3D7_0604500 |
| PF3D7_0309500 | PF3D7_1211700 | PF3D7_0803700 | | PF3D7_0308600 | PF3D7_0607000 |
| PF3D7_0310500 | PF3D7_1212000 | PF3D7_0811600 | | PF3D7_0309500 | PF3D7_0609000 |
| PF3D7_0311300 | PF3D7_1225200 | PF3D7_0813400 | | PF3D7_0309600 | PF3D7_0610800 |
| PF3D7_0315100 | PF3D7_1228800 | PF3D7_0814200 | | PF3D7_0310300 | PF3D7_0610900 |
| PF3D7_0316500 | PF3D7_1234800 | PF3D7_0815200 | | PF3D7_0312400 | PF3D7_0611400 |
| PF3D7_0317200 | PF3D7_1239200 | PF3D7_0817500 | | PF3D7_0312800 | PF3D7_0613800 |
| PF3D7_0317400 | PF3D7_1246400 | PF3D7_0818800 | | PF3D7_0315100 | PF3D7_0619900 |
| PF3D7_0317500 | PF3D7_1249300 | PF3D7_0821200 | | PF3D7_0315600 | PF3D7_0621800 |
| PF3D7_0317700 | PF3D7_1303800 | PF3D7_0823200 | | PF3D7_0316700 | PF3D7_0621900 |
| PF3D7_0318100 | PF3D7_1314500 | PF3D7_0823900 | | PF3D7_0316800 | PF3D7_0624600 |
| PF3D7_0318200 | PF3D7_1329500 | PF3D7_0824600 | | PF3D7_0317200 | PF3D7_0627700 |
| PF3D7_0318600 | PF3D7_1330800 | PF3D7_0904000 | | PF3D7_0317400 | PF3D7_0627800 |
| PF3D7_0319100 | PF3D7_1355800 | PF3D7_0910100 | | PF3D7_0317600 | PF3D7_0628100 |
| PF3D7_0319400 | PF3D7_1368400 | PF3D7_0918900 | | PF3D7_0318200 | PF3D7_0630600 |
| PF3D7_0319500 | PF3D7_1408200 | PF3D7_0920800 | | PF3D7_0319600 | PF3D7_0706000 |
| PF3D7_0320300 | PF3D7_1409400 | PF3D7_0931800 | | PF3D7_0320100 | PF3D7_0708800 |
| PF3D7_0320500 | PF3D7_1412500 | PF3D7_0935800 | | PF3D7_0320300 | PF3D7_0709000 |
| PF3D7_0320800 | PF3D7_1415300 | PF3D7_1006800 | | PF3D7_0320800 | PF3D7_0714500 |
| PF3D7_0321500 | PF3D7_1428900 | PF3D7_1007700 | | PF3D7_0320900 | PF3D7_0717700 |
| PF3D7_0321600 | PF3D7_1433400 | PF3D7_1012500 | | PF3D7_0321500 | PF3D7_0801000 |
| PF3D7_0321800 | PF3D7_1433500 | PF3D7_1015600 | | PF3D7_0322000 | PF3D7_0802000 |
| PF3D7_0323700 | PF3D7_1434500 | PF3D7_1019000 | | PF3D7_0322900 | PF3D7_0803700 |
| PF3D7_0403400 | PF3D7_1438900 | PF3D7_1019400 | | PF3D7_0401800 | PF3D7_0811600 |
| PF3D7_0403700 | PF3D7_1440500 | PF3D7_1021900 | | PF3D7_0402000 | PF3D7_0813400 |
| PF3D7_0404300 | PF3D7_1447800 | PF3D7_1023900 | | PF3D7_0402400 | PF3D7_0814200 |
| PF3D7_0405400 | PF3D7_1451200 | PF3D7_1027800 | | PF3D7_0405400 | PF3D7_0815200 |
| PF3D7_0406100 | PF3D7_1455700 | PF3D7_1032500 | | PF3D7_0406100 | PF3D7_0816600 |
| PF3D7_0407700 | PF3D7_1468100 | PF3D7_1033100 | | PF3D7_0406400 | PF3D7_0817500 |
| PF3D7_0407800 | | PF3D7_1034700 | | PF3D7_0407800 | PF3D7_0818800 |
| PF3D7_0407900 | | PF3D7_1104000 | | PF3D7_0409400 | PF3D7_0821200 |
| PF3D7_0408500 | | PF3D7_1107300 | | PF3D7_0409600 | PF3D7_0823200 |
| PF3D7_0409600 | | PF3D7_1107800 | | PF3D7_0413600 | PF3D7_0823900 |
| PF3D7_0410300 | | PF3D7_1108500 | | PF3D7_0414000 | PF3D7_0824600 |
| PF3D7_0410400 | | PF3D7_1109900 | | PF3D7_0415300 | PF3D7_0827800 |
| PF3D7_0410800 | | PF3D7_1116500 | | PF3D7_0415500 | PF3D7_0903500 |
| PF3D7_0410900 | | PF3D7_1118300 | | PF3D7_0415600 | PF3D7_0903700 |
| PF3D7_0411100 | | PF3D7_1124900 | | PF3D7_0415900 | PF3D7_0904000 |
| PF3D7_0411900 | | PF3D7_1128100 | | PF3D7_0416800 | PF3D7_0907400 |
| PF3D7_0414000 | | PF3D7_1128500 | | PF3D7_0417500 | PF3D7_0910100 |
| PF3D7_0416300 | | PF3D7_1130400 | | PF3D7_0418200 | PF3D7_0910200 |
| PF3D7_0416400 | | PF3D7_1132200 | | PF3D7_0419600 | PF3D7_0918900 |
| PF3D7_0418200 | | PF3D7_1133200 | | PF3D7_0422400 | PF3D7_0920800 |
| PF3D7_0420000 | | PF3D7_1134800 | | PF3D7_0422500 | PF3D7_0923000 |
| PF3D7_0420300 | | PF3D7_1139300 | | PF3D7_0422700 | PF3D7_0925700 |

PF3D7_0420600
PF3D7_0422500
PF3D7_0422700
PF3D7_0501000
PF3D7_0501800
PF3D7_0502100
PF3D7_0503200
PF3D7_0503400
PF3D7_0505100
PF3D7_0505200
PF3D7_0505500
PF3D7_0505600
PF3D7_0505800
PF3D7_0507700
PF3D7_0509100
PF3D7_0509400
PF3D7_0510100
PF3D7_0510200
PF3D7_0511500
PF3D7_0513200
PF3D7_0513300
PF3D7_0513600
PF3D7_0514100
PF3D7_0514900
PF3D7_0515000
PF3D7_0516700
PF3D7_0516800
PF3D7_0517300
PF3D7_0517400
PF3D7_0517700
PF3D7_0519800
PF3D7_0520200
PF3D7_0520300
PF3D7_0520400
PF3D7_0520700
PF3D7_0521700
PF3D7_0522200
PF3D7_0522800
PF3D7_0523400
PF3D7_0524000
PF3D7_0524500
PF3D7_0526500
PF3D7_0526800
PF3D7_0527000
PF3D7_0527500
PF3D7_0527600
PF3D7_0528200
PF3D7_0528700
PF3D7_0529400
PF3D7_0530600
PF3D7_0531400
PF3D7_0602100
PF3D7_0602600
PF3D7_0604100
PF3D7_0604500
PF3D7_0604600
PF3D7_0605100
PF3D7_0605300
PF3D7_0605800
PF3D7_0606500
PF3D7_0607000
PF3D7_0609000
PF3D7_0609900
PF3D7_0610200
PF3D7_0610900
PF3D7_0611400
PF3D7_0611800
PF3D7_0612100
PF3D7_0612900
PF3D7_0613800
PF3D7_0614400
PF3D7_0616200
PF3D7_0616600
PF3D7_0619400
PF3D7_0619900
PF3D7_0620500
PF3D7_0621500
PF3D7_0621800
PF3D7_0622900
PF3D7_0623100
PF3D7_0623600
PF3D7_0624000
PF3D7_0624600
PF3D7_0625600
PF3D7_0627500
PF3D7_0627700
PF3D7_0627800
PF3D7_0627900
PF3D7_0628600
PF3D7_0629100
PF3D7_0629400
PF3D7_0629700
PF3D7_0629800

PF3D7_1144000
PF3D7_1145100
PF3D7_1201000
PF3D7_1203700
PF3D7_1206200
PF3D7_1211900
PF3D7_1216900
PF3D7_1227100
PF3D7_1228600
PF3D7_1230700
PF3D7_1232100
PF3D7_1239200
PF3D7_1248900
PF3D7_1249100
PF3D7_1252100
PF3D7_1308300
PF3D7_1310000
PF3D7_1314700
PF3D7_1315300
PF3D7_1315700
PF3D7_1318800
PF3D7_1323400
PF3D7_1330800
PF3D7_1338200
PF3D7_1340600
PF3D7_1341300
PF3D7_1347700
PF3D7_1364100
PF3D7_1370300
PF3D7_1402300
PF3D7_1407100
PF3D7_1409800
PF3D7_1412100
PF3D7_1416100
PF3D7_1418000
PF3D7_1419700
PF3D7_1419800
PF3D7_1428300
PF3D7_1431700
PF3D7_1442100
PF3D7_1445900
PF3D7_1449500
PF3D7_1451900
PF3D7_1457200
PF3D7_1457300
PF3D7_1459000
PF3D7_1460500
PF3D7_1461900
PF3D7_1465900
PF3D7_1466300
PF3D7_1468100
PF3D7_1471400
PF3D7_1473200

PF3D7_0423000
PF3D7_0423500
PF3D7_0424600
PF3D7_0500800
PF3D7_0501000
PF3D7_0501200
PF3D7_0501500
PF3D7_0501600
PF3D7_0503300
PF3D7_0503400
PF3D7_0503800
PF3D7_0504600
PF3D7_0505500
PF3D7_0505700
PF3D7_0505800
PF3D7_0506500
PF3D7_0506900
PF3D7_0507100
PF3D7_0507700
PF3D7_0509000
PF3D7_0510100
PF3D7_0511000
PF3D7_0511800
PF3D7_0512600
PF3D7_0513300
PF3D7_0513600
PF3D7_0515000
PF3D7_0515700
PF3D7_0516200
PF3D7_0516800
PF3D7_0516900
PF3D7_0517000
PF3D7_0517300
PF3D7_0517400
PF3D7_0517700
PF3D7_0518200
PF3D7_0519400
PF3D7_0519700
PF3D7_0519800
PF3D7_0520000
PF3D7_0520400
PF3D7_0520900
PF3D7_0523000
PF3D7_0523100
PF3D7_0523600
PF3D7_0524000
PF3D7_0524400
PF3D7_0525100
PF3D7_0525800
PF3D7_0526500
PF3D7_0527000
PF3D7_0527500
PF3D7_0528100
PF3D7_0528200
PF3D7_0529400
PF3D7_0530100
PF3D7_0532400
PF3D7_0601200
PF3D7_0922100
PF3D7_0922200
PF3D7_0922500
PF3D7_0923900
PF3D7_0924700
PF3D7_0925700
PF3D7_0927300
PF3D7_0927600
PF3D7_0929200
PF3D7_0929400
PF3D7_0930300
PF3D7_0931400
PF3D7_0931800
PF3D7_0932200
PF3D7_0933600
PF3D7_0934500
PF3D7_0934700
PF3D7_0934800
PF3D7_0935800
PF3D7_1001600
PF3D7_1002400
PF3D7_1002900
PF3D7_1003500
PF3D7_1003600
PF3D7_1003800
PF3D7_1004000
PF3D7_1004400
PF3D7_1005500
PF3D7_1006200
PF3D7_1006800
PF3D7_1007700
PF3D7_1007900
PF3D7_1008400
PF3D7_1008700
PF3D7_1008800

PF3D7_0926100
PF3D7_0931800
PF3D7_0935400
PF3D7_0935800
PF3D7_1002400
PF3D7_1003600
PF3D7_1006800
PF3D7_1007700
PF3D7_1008400
PF3D7_1012500
PF3D7_1014100
PF3D7_1015600
PF3D7_1019000
PF3D7_1019400
PF3D7_1020700
PF3D7_1021900
PF3D7_1023900
PF3D7_1027800
PF3D7_1032500
PF3D7_1033100
PF3D7_1034700
PF3D7_1104000
PF3D7_1106000
PF3D7_1107300
PF3D7_1107800
PF3D7_1108500
PF3D7_1109900
PF3D7_1116500
PF3D7_1118300
PF3D7_1124900
PF3D7_1128100
PF3D7_1128500
PF3D7_1130400
PF3D7_1132200
PF3D7_1133200
PF3D7_1134800
PF3D7_1135400
PF3D7_1139300
PF3D7_1144000
PF3D7_1145100
PF3D7_1148000
PF3D7_1201000
PF3D7_1202200
PF3D7_1203700
PF3D7_1206200
PF3D7_1211700
PF3D7_1211900
PF3D7_1212000
PF3D7_1216900
PF3D7_1225200
PF3D7_1227100
PF3D7_1228600
PF3D7_1228800
PF3D7_1230700
PF3D7_1232100
PF3D7_1234800
PF3D7_1239200
PF3D7_1246400
PF3D7_1248900
PF3D7_1249100
PF3D7_1249300
PF3D7_1252100
PF3D7_1303800
PF3D7_1308300
PF3D7_1310000
PF3D7_1314500
PF3D7_1314700
PF3D7_1315300
PF3D7_1315700
PF3D7_1318800
PF3D7_1323400
PF3D7_1329500
PF3D7_1330800
PF3D7_1338200
PF3D7_1340600
PF3D7_1341300
PF3D7_1347700
PF3D7_1355800
PF3D7_1364100
PF3D7_1368400
PF3D7_1370300
PF3D7_1402300
PF3D7_1407100
PF3D7_1408200
PF3D7_1409400
PF3D7_1409800
PF3D7_1412100
PF3D7_1412500
PF3D7_1415300
PF3D7_1416100
PF3D7_1418000
PF3D7_1419700
PF3D7_1419800

PF3D7_0630300
PF3D7_0630600
PF3D7_0703200
PF3D7_0703500
PF3D7_0703900
PF3D7_0704200
PF3D7_0704400
PF3D7_0705300
PF3D7_0705400
PF3D7_0706000
PF3D7_0706500
PF3D7_0707400
PF3D7_0707700
PF3D7_0708100
PF3D7_0708800
PF3D7_0709300
PF3D7_0709600
PF3D7_0710200
PF3D7_0711500
PF3D7_0714200
PF3D7_0714500
PF3D7_0716000
PF3D7_0716800
PF3D7_0718500
PF3D7_0719000
PF3D7_0719300
PF3D7_0722400
PF3D7_0722500
PF3D7_0723400
PF3D7_0723800
PF3D7_0724700
PF3D7_0725000
PF3D7_0725300
PF3D7_0726300
PF3D7_0726400
PF3D7_0726500
PF3D7_0727900
PF3D7_0728000
PF3D7_0728600
PF3D7_0729100
PF3D7_0729500
PF3D7_0730300
PF3D7_0730500
PF3D7_0801700
PF3D7_0802000
PF3D7_0802100
PF3D7_0802300
PF3D7_0803000
PF3D7_0803700
PF3D7_0804900
PF3D7_0805700
PF3D7_0807100
PF3D7_0807300
PF3D7_0807600
PF3D7_0810300
PF3D7_0810600
PF3D7_0810800
PF3D7_0811000
PF3D7_0811300
PF3D7_0811400
PF3D7_0811500
PF3D7_0812400
PF3D7_0812700
PF3D7_0813300
PF3D7_0814200
PF3D7_0814300
PF3D7_0815200
PF3D7_0815600
PF3D7_0817300
PF3D7_0818200
PF3D7_0818700
PF3D7_0818900
PF3D7_0819000
PF3D7_0819900
PF3D7_0820000
PF3D7_0821600
PF3D7_0822100
PF3D7_0822300
PF3D7_0822800
PF3D7_0823200
PF3D7_0823300
PF3D7_0823800
PF3D7_0824800
PF3D7_0826300
PF3D7_0826700
PF3D7_0827000
PF3D7_0827800
PF3D7_0828500
PF3D7_0903500
PF3D7_0903700
PF3D7_0904500
PF3D7_0904800
PF3D7_0906100

PF3D7_1008900
PF3D7_1010600
PF3D7_1011400
PF3D7_1011800
PF3D7_1012400
PF3D7_1012900
PF3D7_1013300
PF3D7_1014900
PF3D7_1015200
PF3D7_1015600
PF3D7_1015900
PF3D7_1016300
PF3D7_1018200
PF3D7_1019000
PF3D7_1019400
PF3D7_1020900
PF3D7_1021900
PF3D7_1022400
PF3D7_1023900
PF3D7_1024800
PF3D7_1025000
PF3D7_1025300
PF3D7_1026000
PF3D7_1026800
PF3D7_1027300
PF3D7_1027700
PF3D7_1027800
PF3D7_1029600
PF3D7_1030100
PF3D7_1032500
PF3D7_1033200
PF3D7_1033400
PF3D7_1033700
PF3D7_1034000
PF3D7_1034400
PF3D7_1034900
PF3D7_1036900
PF3D7_1037300
PF3D7_1037700
PF3D7_1103100
PF3D7_1103700
PF3D7_1104000
PF3D7_1104100
PF3D7_1104200
PF3D7_1104400
PF3D7_1105000
PF3D7_1105100
PF3D7_1105400
PF3D7_1105600
PF3D7_1105700
PF3D7_1105800
PF3D7_1107300
PF3D7_1107800
PF3D7_1108400
PF3D7_1108500
PF3D7_1108600
PF3D7_1108700
PF3D7_1109900
PF3D7_1110200
PF3D7_1110400
PF3D7_1111100
PF3D7_1111500
PF3D7_1115800
PF3D7_1116200
PF3D7_1116800
PF3D7_1117300
PF3D7_1117700
PF3D7_1118100
PF3D7_1118200
PF3D7_1118500
PF3D7_1119000
PF3D7_1119800
PF3D7_1120000
PF3D7_1120100
PF3D7_1121100
PF3D7_1121600
PF3D7_1121700
PF3D7_1123400
PF3D7_1123900
PF3D7_1124600
PF3D7_1124700
PF3D7_1124900
PF3D7_1125500
PF3D7_1126200
PF3D7_1127600
PF3D7_1128200
PF3D7_1129000
PF3D7_1129100
PF3D7_1130100
PF3D7_1130200
PF3D7_1130400
PF3D7_1130700
PF3D7_1132200

PF3D7_1428300
PF3D7_1428900
PF3D7_1431700
PF3D7_1433400
PF3D7_1433500
PF3D7_1434500
PF3D7_1438900
PF3D7_1440500
PF3D7_1442100
PF3D7_1445900
PF3D7_1447800
PF3D7_1449500
PF3D7_1451200
PF3D7_1451900
PF3D7_1455700
PF3D7_1457200
PF3D7_1457300
PF3D7_1459000
PF3D7_1460500
PF3D7_1461900
PF3D7_1465900
PF3D7_1466300
PF3D7_1468100
PF3D7_1471400
PF3D7_1473200

PF3D7_0906600
PF3D7_0907400
PF3D7_0909400
PF3D7_0909800
PF3D7_0909900
PF3D7_0910100
PF3D7_0910200
PF3D7_0910400
PF3D7_0910900
PF3D7_0913200
PF3D7_0914800
PF3D7_0916700
PF3D7_0917000
PF3D7_0917600
PF3D7_0918300
PF3D7_0919000
PF3D7_0920800
PF3D7_0920900
PF3D7_0922200
PF3D7_0922500
PF3D7_0923000
PF3D7_0923900
PF3D7_0924700
PF3D7_0925700
PF3D7_0927600
PF3D7_0929000
PF3D7_0929200
PF3D7_0932300
PF3D7_0932800
PF3D7_0933000
PF3D7_0934100
PF3D7_0934800
PF3D7_0935000
PF3D7_1003800
PF3D7_1004200
PF3D7_1004300
PF3D7_1004400
PF3D7_1004500
PF3D7_1005500
PF3D7_1006200
PF3D7_1006700
PF3D7_1006800
PF3D7_1007700
PF3D7_1007900
PF3D7_1008000
PF3D7_1008100
PF3D7_1008700
PF3D7_1008800
PF3D7_1008900
PF3D7_1009200
PF3D7_1010200
PF3D7_1010600
PF3D7_1011800
PF3D7_1012400
PF3D7_1012700
PF3D7_1013100
PF3D7_1013600
PF3D7_1014300
PF3D7_1014600
PF3D7_1014900
PF3D7_1015400
PF3D7_1015600
PF3D7_1015800
PF3D7_1016400
PF3D7_1018200
PF3D7_1019000
PF3D7_1019700
PF3D7_1020700
PF3D7_1021800
PF3D7_1021900
PF3D7_1022400
PF3D7_1023900
PF3D7_1027300
PF3D7_1028500
PF3D7_1029600
PF3D7_1030100
PF3D7_1031300
PF3D7_1031500
PF3D7_1031600
PF3D7_1032700
PF3D7_1033100
PF3D7_1033500
PF3D7_1033600
PF3D7_1033700
PF3D7_1036900
PF3D7_1037500
PF3D7_1037600
PF3D7_1103600
PF3D7_1103700
PF3D7_1103800
PF3D7_1104200
PF3D7_1104400
PF3D7_1105100

PF3D7_1132300
PF3D7_1133800
PF3D7_1134000
PF3D7_1134100
PF3D7_1134600
PF3D7_1134800
PF3D7_1136300
PF3D7_1136500
PF3D7_1137400
PF3D7_1138500
PF3D7_1140200
PF3D7_1140400
PF3D7_1141800
PF3D7_1142100
PF3D7_1142500
PF3D7_1142600
PF3D7_1143200
PF3D7_1143400
PF3D7_1144000
PF3D7_1144900
PF3D7_1145100
PF3D7_1145400
PF3D7_1202900
PF3D7_1203700
PF3D7_1204300
PF3D7_1205600
PF3D7_1206200
PF3D7_1207000
PF3D7_1208900
PF3D7_1211400
PF3D7_1211700
PF3D7_1211900
PF3D7_1212000
PF3D7_1212700
PF3D7_1215000
PF3D7_1216200
PF3D7_1216300
PF3D7_1216900
PF3D7_1219100
PF3D7_1220900
PF3D7_1221000
PF3D7_1222300
PF3D7_1223100
PF3D7_1223300
PF3D7_1224000
PF3D7_1224300
PF3D7_1224500
PF3D7_1225800
PF3D7_1226300
PF3D7_1227100
PF3D7_1228800
PF3D7_1229300
PF3D7_1229400
PF3D7_1229500
PF3D7_1230400
PF3D7_1230700
PF3D7_1230800
PF3D7_1231100
PF3D7_1231600
PF3D7_1232100
PF3D7_1234500
PF3D7_1234800
PF3D7_1235300
PF3D7_1235600
PF3D7_1236100
PF3D7_1237700
PF3D7_1238100
PF3D7_1238800
PF3D7_1239200
PF3D7_1239500
PF3D7_1239600
PF3D7_1239700
PF3D7_1241700
PF3D7_1242700
PF3D7_1242800
PF3D7_1243600
PF3D7_1244100
PF3D7_1244800
PF3D7_1245100
PF3D7_1246200
PF3D7_1246800
PF3D7_1247400
PF3D7_1248900
PF3D7_1249300
PF3D7_1249800
PF3D7_1252100
PF3D7_1302100
PF3D7_1302800
PF3D7_1304400
PF3D7_1304500
PF3D7_1305300
PF3D7_1305900
PF3D7_1306400

PF3D7 1105700
PF3D7 1106000
PF3D7 1106700
PF3D7 1107000
PF3D7 1107300
PF3D7 1107400
PF3D7 1107800
PF3D7 1108400
PF3D7 1110200
PF3D7 1110700
PF3D7 1110900
PF3D7_1111100
PF3D7 1112100
PF3D7 1112500
PF3D7 1117700
PF3D7 1117800
PF3D7 1118200
PF3D7 1118500
PF3D7 1118600
PF3D7 1119300
PF3D7 1119800
PF3D7 1120100
PF3D7 1121700
PF3D7 1121900
PF3D7 1123200
PF3D7 1124300
PF3D7 1125500
PF3D7 1126300
PF3D7 1126400
PF3D7 1126900
PF3D7_1127600
PF3D7 1128100
PF3D7 1129000
PF3D7 1129400
PF3D7 1130700
PF3D7 1132200
PF3D7 1132300
PF3D7 1133200
PF3D7 1134000
PF3D7 1134700
PF3D7 1135100
PF3D7 1136300
PF3D7 1136500
PF3D7 1138500
PF3D7 1138800
PF3D7 1139300
PF3D7 1141800
PF3D7_1142100
PF3D7 1143300
PF3D7 1145100
PF3D7 1145200
PF3D7 1145400
PF3D7 1146600
PF3D7 1147300
PF3D7 1147500
PF3D7 1148000
PF3D7 1202100
PF3D7 1202600
PF3D7 1203700
PF3D7 1203900
PF3D7 1205500
PF3D7 1205600
PF3D7 1205800
PF3D7 1206200
PF3D7_1206600
PF3D7 1207000
PF3D7 1207100
PF3D7 1208100
PF3D7 1208900
PF3D7 1209200
PF3D7 1210400
PF3D7 1211200
PF3D7 1211300
PF3D7 1211400
PF3D7 1211600
PF3D7 1211700
PF3D7 1211800
PF3D7 1212700
PF3D7 1212900
PF3D7 1213200
PF3D7 1213700
PF3D7 1213900
PF3D7 1216900
PF3D7_1220100
PF3D7 1221000
PF3D7 1222600
PF3D7 1223100
PF3D7 1224000
PF3D7 1224300
PF3D7 1224500
PF3D7 1224900
PF3D7 1225200
PF3D7 1225800

PF3D7 1306900
PF3D7 1308200
PF3D7 1308300
PF3D7 1309100
PF3D7 1309500
PF3D7 1310700
PF3D7 1311400
PF3D7 1311500
PF3D7 1311800
PF3D7 1311900
PF3D7 1312600
PF3D7_1312900
PF3D7 1314700
PF3D7 1315300
PF3D7 1315400
PF3D7 1316800
PF3D7 1317100
PF3D7 1317400
PF3D7 1317800
PF3D7 1318800
PF3D7 1320600
PF3D7 1320800
PF3D7 1321700
PF3D7 1322200
PF3D7 1322300
PF3D7 1323100
PF3D7 1323200
PF3D7 1323400
PF3D7 1323700
PF3D7 1324900
PF3D7_1325100
PF3D7 1326300
PF3D7 1327800
PF3D7 1328100
PF3D7 1328300
PF3D7 1329100
PF3D7 1330300
PF3D7 1330600
PF3D7 1330800
PF3D7 1331800
PF3D7 1332900
PF3D7 1333300
PF3D7 1334200
PF3D7 1336900
PF3D7 1338200
PF3D7 1338300
PF3D7 1339700
PF3D7_1340600
PF3D7 1341200
PF3D7 1341300
PF3D7 1342000
PF3D7 1342100
PF3D7 1342600
PF3D7 1343000
PF3D7 1343700
PF3D7 1345600
PF3D7 1345700
PF3D7 1346100
PF3D7 1346300
PF3D7 1347500
PF3D7 1349200
PF3D7 1350100
PF3D7 1351400
PF3D7 1352500
PF3D7_1353100
PF3D7 1353200
PF3D7 1353800
PF3D7 1353900
PF3D7 1354500
PF3D7 1355100
PF3D7 1355300
PF3D7 1357100
PF3D7 1357800
PF3D7 1357900
PF3D7 1358200
PF3D7 1358800
PF3D7 1358900
PF3D7 1359400
PF3D7 1359600
PF3D7 1360800
PF3D7 1360900
PF3D7 1361100
PF3D7 1361800
PF3D7_1361900
PF3D7 1362700
PF3D7 1365500
PF3D7 1365900
PF3D7 1366500
PF3D7 1366900
PF3D7 1367100
PF3D7 1368200
PF3D7 1368400
PF3D7 1369500

PF3D7_1226800
PF3D7_1227800
PF3D7_1228300
PF3D7_1228800
PF3D7_1229500
PF3D7_1230700
PF3D7_1230800
PF3D7_1231600
PF3D7_1234100
PF3D7_1234800
PF3D7_1235300
PF3D7_1235500
PF3D7_1235900
PF3D7_1237500
PF3D7_1237700
PF3D7_1238500
PF3D7_1238600
PF3D7_1238800
PF3D7_1239000
PF3D7_1239200
PF3D7_1241200
PF3D7_1241700
PF3D7_1242800
PF3D7_1244100
PF3D7_1244200
PF3D7_1245100
PF3D7_1246200
PF3D7_1247400
PF3D7_1247500
PF3D7_1248200
PF3D7_1248700
PF3D7_1249300
PF3D7_1249800
PF3D7_1250600
PF3D7_1250800
PF3D7_1251200
PF3D7_1301700
PF3D7_1302100
PF3D7_1302500
PF3D7_1303400
PF3D7_1303800
PF3D7_1304000
PF3D7_1304700
PF3D7_1305000
PF3D7_1305300
PF3D7_1305400
PF3D7_1306900
PF3D7_1307000
PF3D7_1307100
PF3D7_1308200
PF3D7_1308900
PF3D7_1309300
PF3D7_1309400
PF3D7_1311900
PF3D7_1314900
PF3D7_1315300
PF3D7_1315800
PF3D7_1315900
PF3D7_1316500
PF3D7_1316600
PF3D7_1316900
PF3D7_1317100
PF3D7_1317300
PF3D7_1317600
PF3D7_1318400
PF3D7_1319400
PF3D7_1321700
PF3D7_1322100
PF3D7_1322200
PF3D7_1322300
PF3D7_1324100
PF3D7_1324200
PF3D7_1325100
PF3D7_1325400
PF3D7_1326400
PF3D7_1328200
PF3D7_1329000
PF3D7_1329100
PF3D7_1329300
PF3D7_1329500
PF3D7_1329600
PF3D7_1330600
PF3D7_1330800
PF3D7_1333600
PF3D7_1334100
PF3D7_1336400
PF3D7_1336800
PF3D7_1338300
PF3D7_1338700
PF3D7_1340100
PF3D7_1340600
PF3D7_1341900
PF3D7_1342400

PF3D7_1370300
PF3D7_1401600
PF3D7_1402500
PF3D7_1405600
PF3D7_1407100
PF3D7_1407800
PF3D7_1408600
PF3D7_1408700
PF3D7_1409400
PF3D7_1409600
PF3D7_1409800
PF3D7_1410400
PF3D7_1410600
PF3D7_1411300
PF3D7_1412500
PF3D7_1412800
PF3D7_1414300
PF3D7_1414400
PF3D7_1414800
PF3D7_1415300
PF3D7_1415400
PF3D7_1416100
PF3D7_1416900
PF3D7_1417500
PF3D7_1417800
PF3D7_1418000
PF3D7_1419700
PF3D7_1421200
PF3D7_1423700
PF3D7_1424100
PF3D7_1424400
PF3D7_1426000
PF3D7_1426100
PF3D7_1427500
PF3D7_1427900
PF3D7_1428300
PF3D7_1429800
PF3D7_1431600
PF3D7_1431700
PF3D7_1433500
PF3D7_1434200
PF3D7_1434300
PF3D7_1434600
PF3D7_1434800
PF3D7_1435500
PF3D7_1435700
PF3D7_1437200
PF3D7_1437700
PF3D7_1437900
PF3D7_1438000
PF3D7_1438100
PF3D7_1438900
PF3D7_1439800
PF3D7_1439900
PF3D7_1441200
PF3D7_1442300
PF3D7_1444800
PF3D7_1445100
PF3D7_1445400
PF3D7_1445700
PF3D7_1445900
PF3D7_1446200
PF3D7_1446600
PF3D7_1447000
PF3D7_1447700
PF3D7_1449500
PF3D7_1450100
PF3D7_1451100
PF3D7_1451800
PF3D7_1452000
PF3D7_1453700
PF3D7_1454700
PF3D7_1455500
PF3D7_1455700
PF3D7_1456800
PF3D7_1457000
PF3D7_1457300
PF3D7_1459000
PF3D7_1459400
PF3D7_1459600
PF3D7_1460300
PF3D7_1460500
PF3D7_1460600
PF3D7_1460700
PF3D7_1461300
PF3D7_1462300
PF3D7_1462800
PF3D7_1463200
PF3D7_1464700
PF3D7_1465900
PF3D7_1466400
PF3D7_1466900
PF3D7_1468100

PF3D7_1342800
PF3D7_1343300
PF3D7_1343900
PF3D7_1344300
PF3D7_1345700
PF3D7_1345800
PF3D7_1346300
PF3D7_1347100
PF3D7_1347200
PF3D7_1347500
PF3D7_1352400
PF3D7_1352700
PF3D7_1355100
PF3D7_1355300
PF3D7_1355700
PF3D7_1355800
PF3D7_1356100
PF3D7_1357100
PF3D7_1357500
PF3D7_1357700
PF3D7_1357800
PF3D7_1359400
PF3D7_1359600
PF3D7_1360900
PF3D7_1361000
PF3D7_1361100
PF3D7_1361900
PF3D7_1362200
PF3D7_1362400
PF3D7_1363200
PF3D7_1363600
PF3D7_1364200
PF3D7_1364300
PF3D7_1364500
PF3D7_1364800
PF3D7_1366300
PF3D7_1366900
PF3D7_1367100
PF3D7_1368200
PF3D7_1368800
PF3D7_1369500
PF3D7_1369700
PF3D7_1403100
PF3D7_1403900
PF3D7_1404000
PF3D7_1405600
PF3D7_1406200
PF3D7_1407100
PF3D7_1407300
PF3D7_1408400
PF3D7_1408700
PF3D7_1409300
PF3D7_1409800
PF3D7_1410300
PF3D7_1410600
PF3D7_1412100
PF3D7_1414400
PF3D7_1414800
PF3D7_1415200
PF3D7_1415400
PF3D7_1416900
PF3D7_1417200
PF3D7_1417500
PF3D7_1418000
PF3D7_1419000
PF3D7_1419700
PF3D7_1419900
PF3D7_1420000
PF3D7_1420600
PF3D7_1422400
PF3D7_1422800
PF3D7_1423700
PF3D7_1425900
PF3D7_1426100
PF3D7_1426700
PF3D7_1427500
PF3D7_1427900
PF3D7_1428800
PF3D7_1429900
PF3D7_1432100
PF3D7_1433300
PF3D7_1433400
PF3D7_1433500
PF3D7_1434500
PF3D7_1435600
PF3D7_1437000
PF3D7_1437200
PF3D7_1437400
PF3D7_1437900
PF3D7_1438000
PF3D7_1438500
PF3D7_1438700
PF3D7_1438900

PF3D7_1468700
PF3D7_1468900
PF3D7_1471100
PF3D7_1472200
PF3D7_1473200
PF3D7_1473700
PF3D7_1474500
PF3D7_1474900

PF3D7_1439500
PF3D7_1439700
PF3D7_1440100
PF3D7_1441400
PF3D7_1442100
PF3D7_1443000
PF3D7_1444800
PF3D7_1445600
PF3D7_1445900
PF3D7_1446500
PF3D7_1446600
PF3D7_1446700
PF3D7_1447700
PF3D7_1448000
PF3D7_1448300
PF3D7_1448500
PF3D7_1449500
PF3D7_1450400
PF3D7_1451100
PF3D7_1451200
PF3D7_1451500
PF3D7_1453700
PF3D7_1454200
PF3D7_1456000
PF3D7_1456500
PF3D7_1457300
PF3D7_1458600
PF3D7_1458800
PF3D7_1459000
PF3D7_1459400
PF3D7_1460500
PF3D7_1460800
PF3D7_1461600
PF3D7_1463200
PF3D7_1463400
PF3D7_1464000
PF3D7_1464700
PF3D7_1465200
PF3D7_1466400
PF3D7_1466800
PF3D7_1467500
PF3D7_1468100
PF3D7_1468200
PF3D7_1468700
PF3D7_1469700
PF3D7_1469800
PF3D7_1471400
PF3D7_1472000
PF3D7_1472200
PF3D7_1472900
PF3D7_1473200
PF3D7_1473700
PF3D7_1474500
PF3D7_1475000
PF3D7_1475600

### List of interactors in total

| Short name | Product description | Gene IDs | Interactors of |
|---|---|---|---|
| N/A | aspartate--tRNA ligase | PF3D7_0102900 | PfAP2-P, PfMORC |
| NT4 | nucleoside transporter 4 | PF3D7_0103200 | PfAP2-P, PfMORC |
| | | PF3D7_0103900 | |
| | | PF3D7_0104400 | |
| RAP1 | RAP protein RAP1 | PF3D7_0105200 | PfAP2-P, PfMORC |
| N/A | cyclin-dependent kinases regulatory subunit, putative | PF3D7_0105800 | PfAP2-P, PfSET10 |
| | | PF3D7_0106300 | |
| | | PF3D7_0107000 | |
| | | PF3D7_0107800 | |
| ARP | conserved Plasmodium protein, unknown function | PF3D7_0108300 | PfAP2-P, PfMORC |
| | | PF3D7_0108700 | |
| CFIM25 | cleavage and polyadenylation specificity factor subunit 5, putative | PF3D7_0109200 | PfSET10, PfMORC |
| | | PF3D7_0110400 | |
| BDP3 | bromodomain protein 3, putative | PF3D7_0110500 | PfAP2-P, PfSET10, PfMORC |
| | | PF3D7_0110700 | |
| | | PF3D7_0111300 | |
| | | PF3D7_0111500 | |
| N/A | eukaryotic translation initiation factor 4E, putative | PF3D7_0111800 | PfAP2-P, PfSET10 |
| | | PF3D7_0112200 | |
| | | PF3D7_0115600 | |
| | | PF3D7_0202000 | |
| | | PF3D7_0205400 | |
| N/A | conserved Plasmodium protein, unknown function | PF3D7_0205600 | PfAP2-P, PfSET10 |
| | | PF3D7_0205700 | |
| | | PF3D7_0205900 | |
| | | PF3D7_0206000 | |
| | | PF3D7_0206700 | |
| | | PF3D7_0209200 | |
| UAP56 | ATP-dependent RNA helicase UAP56 | PF3D7_0209800 | PfAP2-P, PfSET10 |
| | | PF3D7_0210100 | |
| | | PF3D7_0210200 | |
| | | PF3D7_0210900 | |
| | | PF3D7_0211800 | |
| N/A | conserved Plasmodium protein, unknown function | PF3D7_0212100 | PfAP2-P, PfSET10 |
| ERF1 | eukaryotic peptide chain release factor subunit 1, putative | PF3D7_0212300 | PfAP2-P, PfSET10, PfISWI |
| | | PF3D7_0212400 | |
| NUP434 | nucleoporin NUP434, putative | PF3D7_0212500 | PfSET10, PfMORC |
| SIS1 | protein SIS1 | PF3D7_0213100 | PfAP2-P, PfSET10 |
| CCT8 | T-complex protein 1 subunit theta | PF3D7_0214000 | PfAP2-P, PfSET10 |
| SEC31 | protein transport protein SEC31 | PF3D7_0214100 | PfAP2-P, PfMORC |
| RPB2 | DNA-directed RNA polymerase II subunit RPB2, putative | PF3D7_0215700 | PfAP2-P, PfSET10, PfMORC |
| | | PF3D7_0215800 | |
| | | PF3D7_0216200 | |
| | | PF3D7_0217100 | |

| | |
|---|---|
| Interactions total | 328 |
| PfAP2-P, PfMORC | 46 |
| PfAP2-P, PfSET10 | 161 |
| PfSET10, PfMORC | 50 |
| PfSET10, PfISWI | 6 |
| PfAP2-P, PfISWI | 5 |
| PfAP2-P, PfSET10, PfMORC | 43 |
| PfAP2-P, PfSET10, PfISWI | 11 |
| PfSET10, PfMORC, PfISWI | 2 |
| PfAP2-P, PfSET10, PfMORC, PfISWI | 4 |

| | | | |
|---|---|---|---|
| CDPK1 | Calcium-dependent protein kinase 1 | PF3D7_0217500 | PfAP2-P, PfSET10, PfMORC |
| | | PF3D7_0217800 | |
| | | PF3D7_0218000 | |
| | | PF3D7_0218200 | |
| SNRPD2 | small nuclear ribonucleoprotein Sm D2, putative | PF3D7_0218500 | PfSET10, PfMORC |
| | | PF3D7_0218700 | |
| | | PF3D7_0219400 | |
| RFC1 | replication factor C subunit 1 | PF3D7_0219600 | PfAP2-P, PfSET10 |
| | | PF3D7_0301600 | |
| | | PF3D7_0302000 | |
| | | PF3D7_0302100 | |
| | | PF3D7_0302500 | |
| N/A | exportin-1, putative | PF3D7_0302900 | PfAP2-P, PfSET10 |
| | | PF3D7_0303000 | |
| | | PF3D7_0303100 | |
| | | PF3D7_0303300 | |
| | | PF3D7_0303400 | |
| | | PF3D7_0303700 | |
| | | PF3D7_0304100 | |
| | | PF3D7_0304200 | |
| | | PF3D7_0304400 | |
| | | PF3D7_0304500 | |
| | | PF3D7_0305500 | |
| APE1 | DNA-(apurinic or apyrimidinic site) endonuclease | PF3D7_0305600 | PfAP2-P, PfSET10 |
| | | PF3D7_0306100 | |
| | | PF3D7_0306400 | |
| CCT2 | T-complex protein 1 subunit beta | PF3D7_0306800 | PfAP2-P, PfSET10 |
| | | PF3D7_0306900 | |
| | | PF3D7_0307100 | |
| | | PF3D7_0307200 | |
| AKIT10 | apicomplexan kinetochore protein 10, putative | PF3D7_0307700 | PfSET10, PfMORC |
| N/A | DNA polymerase delta small subunit, putative | PF3D7_0308000 | PfSET10, PfMORC |
| | | PF3D7_0308100 | |
| CCT7 | T-complex protein 1 subunit eta | PF3D7_0308200 | PfAP2-P, PfSET10, PfMORC |
| PRPF19 | pre-mRNA-processing factor 19, putative | PF3D7_0308600 | PfAP2-P, PfSET10 |
| | | PF3D7_0309000 | |
| | | PF3D7_0309200 | |
| | | PF3D7_0309300 | |
| AS | asparagine synthetase [glutamine-hydrolyzing], putative | PF3D7_0309500 | PfAP2-P, PfSET10 |
| | | PF3D7_0309600 | |
| | | PF3D7_0310300 | |
| | | PF3D7_0310500 | |
| | | PF3D7_0310600 | |
| | | PF3D7_0311300 | |
| | | PF3D7_0311800 | |
| | | PF3D7_0312400 | |
| | | PF3D7_0312800 | |

| | | | |
|---|---|---|---|
| eIF4E | eukaryotic translation initiation factor 4E | PF3D7_0315100 | PfAP2-P, PfSET10 |
| | | PF3D7_0315600 | |
| | | PF3D7_0316500 | |
| | | PF3D7_0316700 | |
| | | PF3D7_0316800 | |
| CRK4 | cdc2-related protein kinase 4 | PF3D7_0317200 | PfAP2-P, PfSET10 |
| N/A | DNA replication complex GINS protein, putative | PF3D7_0317400 | PfAP2-P, PfSET10 |
| | | PF3D7_0317500 | |
| | | PF3D7_0317600 | |
| | | PF3D7_0317700 | |
| | | PF3D7_0318100 | |
| RPB1 | DNA-directed RNA polymerase II subunit RPB1 | PF3D7_0318200 | PfAP2-P, PfSET10, PfMORC |
| | | PF3D7_0318500 | |
| | | PF3D7_0318600 | |
| | | PF3D7_0319100 | |
| | | PF3D7_0319400 | |
| | | PF3D7_0319500 | |
| | | PF3D7_0319600 | |
| | | PF3D7_0320100 | |
| CCT5 | T-complex protein 1 subunit epsilon | PF3D7_0320300 | PfAP2-P, PfSET10, PfMORC |
| | | PF3D7_0320500 | |
| DOZI | ATP-dependent RNA helicase DDX6 | PF3D7_0320800 | PfAP2-P, PfSET10 |
| | | PF3D7_0320900 | |
| N/A | peptidase, putative | PF3D7_0321500 | PfAP2-P, PfSET10 |
| | | PF3D7_0321600 | |
| | | PF3D7_0321800 | |
| | | PF3D7_0322000 | |
| | | PF3D7_0322900 | |
| | | PF3D7_0323700 | |
| N/A | Plasmodium exported protein (PHISTb), unknown function | PF3D7_0401800 | PfAP2-P, PfMORC |
| | | PF3D7_0402000 | |
| | | PF3D7_0402400 | |
| | | PF3D7_0403400 | |
| | | PF3D7_0403600 | |
| | | PF3D7_0403700 | |
| | | PF3D7_0404300 | |
| PRPF8 | pre-mRNA-processing-splicing factor 8, putative | PF3D7_0405400 | PfAP2-P, PfSET10, PfMORC |
| N/A | V-type proton ATPase subunit B | PF3D7_0406100 | PfAP2-P, PfSET10 |
| | | PF3D7_0406400 | |
| | | PF3D7_0407700 | |
| CINCH | protein CINCH | PF3D7_0407800 | PfAP2-P, PfSET10 |
| | | PF3D7_0407900 | |
| | | PF3D7_0408500 | |
| | | PF3D7_0409400 | |
| RPA1 | replication protein A1, large subunit | PF3D7_0409600 | PfAP2-P, PfSET10 |
| | | PF3D7_0410300 | |
| | | PF3D7_0410400 | |

| | | | |
|---|---|---|---|
| AKIT3 | apicomplexan kinetochore protein 3, putative | PF3D7_0410800 | PfSET10, PfMORC |
| | | PF3D7_0410900 | |
| | | PF3D7_0411100 | |
| | | PF3D7_0411900 | |
| | | PF3D7_0413600 | |
| SMC3 | structural maintenance of chromosomes protein 3 | PF3D7_0414000 | PfAP2-P, PfSET10 |
| | | PF3D7_0415300 | |
| | | PF3D7_0415500 | |
| | | PF3D7_0415600 | |
| | | PF3D7_0415900 | |
| | | PF3D7_0416300 | |
| | | PF3D7_0416400 | |
| | | PF3D7_0416800 | |
| | | PF3D7_0417500 | |
| EIF3M | eukaryotic translation initiation factor 3 subunit M, putative | PF3D7_0418200 | PfAP2-P, PfSET10 |
| | | PF3D7_0419600 | |
| | | PF3D7_0419900 | |
| | | PF3D7_0420000 | |
| ApiAP2 | AP2 domain transcription factor, putative | PF3D7_0420300 | PfSET10, PfMORC |
| | | PF3D7_0420400 | |
| | | PF3D7_0420600 | |
| | | PF3D7_0422400 | |
| BRR2 | pre-mRNA-splicing helicase BRR2, putative | PF3D7_0422500 | PfAP2-P, PfSET10 |
| EIF4A3 | eukaryotic initiation factor 4A-III, putative | PF3D7_0422700 | PfAP2-P, PfSET10 |
| | | PF3D7_0423000 | |
| | | PF3D7_0423500 | |
| | | PF3D7_0424600 | |
| | | PF3D7_0500800 | |
| N/A | Plasmodium exported protein, unknown function | PF3D7_0501000 | PfAP2-P, PfSET10 |
| | | PF3D7_0501200 | |
| ROP3 | rhoptry-associated protein 3 | PF3D7_0501500 | PfAP2-P, PfMORC |
| | | PF3D7_0501600 | |
| | | PF3D7_0501800 | |
| | | PF3D7_0502100 | |
| | | PF3D7_0503200 | |
| | | PF3D7_0503300 | |
| ADF1 | actin-depolymerizing factor 1 | PF3D7_0503400 | PfAP2-P, PfSET10 |
| | | PF3D7_0503800 | |
| | | PF3D7_0504600 | |
| | | PF3D7_0505100 | |
| | | PF3D7_0505200 | |
| MSH6 | DNA mismatch repair protein MSH6, putative | PF3D7_0505500 | PfAP2-P, PfSET10, PfMORC |
| | | PF3D7_0505600 | |
| | | PF3D7_0505700 | |
| SUMO | small ubiquitin-related modifier | PF3D7_0505800 | PfAP2-P, PfSET10 |
| | | PF3D7_0505900 | |
| N/A | conserved Plasmodium protein, unknown function | PF3D7_0506500 | PfAP2-P, PfMORC |

| | | | |
|---|---|---|---|
| | | PF3D7_0506900 | |
| | | PF3D7_0507100 | |
| NPL4 | nuclear protein localization protein 4, putative | PF3D7_0507700 | PfAP2-P, PfSET10 |
| | | PF3D7_0509000 | |
| | | PF3D7_0509100 | |
| | | PF3D7_0509400 | |
| | | PF3D7_0510100 | |
| | | PF3D7_0510100 | |
| | | PF3D7_0510200 | |
| | | PF3D7_0511000 | |
| | | PF3D7_0511500 | |
| INO1 | inositol-3-phosphate synthase | PF3D7_0511800 | PfAP2-P, PfISWI |
| | | PF3D7_0512600 | |
| | | PF3D7_0513200 | |
| PNP | purine nucleoside phosphorylase | PF3D7_0513300 | PfAP2-P, PfSET10 |
| N/A | deoxyribodipyrimidine photo-lyase, putative | PF3D7_0513600 | PfAP2-P, PfSET10, PfMORC |
| | | PF3D7_0514100 | |
| | | PF3D7_0514900 | |
| CWC2 | pre-mRNA-splicing factor CWC2, putative | PF3D7_0515000 | PfAP2-P, PfSET10 |
| | | PF3D7_0515700 | |
| | | PF3D7_0516200 | |
| | | PF3D7_0516700 | |
| ApiAP2 | AP2 domain transcription factor AP2-O2, putative | PF3D7_0516800 | PfAP2-P, PfSET10, PfMORC |
| | | PF3D7_0516900 | |
| | | PF3D7_0517000 | |
| | | PF3D7_0517300 | |
| FACT-L | FACT complex subunit SPT16, putative | PF3D7_0517400 | PfAP2-P, PfSET10, PfISWI |
| EIF3B | eukaryotic translation initiation factor 3 subunit B, putative | PF3D7_0517700 | PfAP2-P, PfSET10 |
| | | PF3D7_0518200 | |
| RPS24 | 40S ribosomal protein S24 | PF3D7_0519400 | PfAP2-P, PfMORC |
| | | PF3D7_0519700 | |
| N/A | EELM2 domain-containing protein, putative | PF3D7_0519800 | PfAP2-P, PfSET10, PfMORC |
| | | PF3D7_0520000 | |
| | | PF3D7_0520200 | |
| | | PF3D7_0520300 | |
| N/A | single-stranded DNA-binding protein, putative | PF3D7_0520400 | PfAP2-P, PfSET10 |
| | | PF3D7_0520700 | |
| | | PF3D7_0520900 | |
| | | PF3D7_0521700 | |
| | | PF3D7_0522200 | |
| | | PF3D7_0522300 | |
| | | PF3D7_0522800 | |
| | | PF3D7_0523000 | |
| | | PF3D7_0523100 | |
| | | PF3D7_0523400 | |
| | | PF3D7_0523600 | |
| KASbeta | karyopherin beta | PF3D7_0524000 | PfAP2-P, PfSET10 |

| | | | |
|---|---|---|---|
| | | PF3D7_0524400 | |
| | | PF3D7_0524500 | |
| | | PF3D7_0525100 | |
| | | PF3D7_0525800 | |
| N/A | Suf domain-containing protein, putative | PF3D7_0526500 | PfAP2-P, PfSET10 |
| | | PF3D7_0526800 | |
| MCM3 | DNA replication licensing factor MCM3, putative | PF3D7_0527000 | PfAP2-P, PfSET10, PfMORC |
| HIP | Hsc70-interacting protein | PF3D7_0527500 | PfAP2-P, PfSET10 |
| | | PF3D7_0527600 | |
| N/A | AP-1/2 complex subunit beta, putative | PF3D7_0528100 | PfAP2-P, PfMORC |
| EIF3E | eukaryotic translation initiation factor 3 subunit E, putative | PF3D7_0528200 | PfAP2-P, PfSET10, PfMORC |
| | | PF3D7_0528700 | |
| AKIT5 | apicomplexan kinetochore protein 5, putative | PF3D7_0529400 | PfAP2-P, PfSET10 |
| | | PF3D7_0530100 | |
| | | PF3D7_0530600 | |
| | | PF3D7_0531400 | |
| | | PF3D7_0532400 | |
| | | PF3D7_0601200 | |
| | | PF3D7_0602100 | |
| | | PF3D7_0602600 | |
| SIP2 | AP2 domain transcription factor | PF3D7_0604100 | PfSET10, PfMORC |
| N/A | GYF domain-containing protein, putative | PF3D7_0604500 | PfSET10, PfMORC |
| | | PF3D7_0604600 | |
| | | PF3D7_0605100 | |
| | | PF3D7_0605300 | |
| | | PF3D7_0605800 | |
| | | PF3D7_0606500 | |
| N/A | translation initiation factor IF-2, putative | PF3D7_0607000 | PfSET10, PfMORC |
| NUP637 | nucleoporin NUP637, putative | PF3D7_0609000 | PfSET10, PfMORC |
| | | PF3D7_0609900 | |
| | | PF3D7_0610200 | |
| | | PF3D7_0610800 | |
| SPT5 | transcription elongation factor SPT5, putative | PF3D7_0610900 | PfSET10, PfMORC |
| SPT5 | transcription elongation factor SPT5, putative | PF3D7_0611400 | PfSET10, PfMORC |
| | | PF3D7_0611800 | |
| | | PF3D7_0612100 | |
| | | PF3D7_0612900 | |
| ApiAP2 | AP2 domain transcription factor, putative | PF3D7_0613800 | PfSET10, PfMORC (2x) |
| | | PF3D7_0614400 | |
| | | PF3D7_0616200 | |
| | | PF3D7_0616600 | |
| | | PF3D7_0619400 | |
| SF3A2 | splicing factor 3A subunit 2, putative | PF3D7_0619900 | PfSET10, PfMORC |
| | | PF3D7_0620500 | |
| | | PF3D7_0621500 | |
| N/A | nascent polypeptide-associated complex subunit alpha | PF3D7_0621800 | PfSET10, PfMORC, PfISWI |
| | | PF3D7_0621900 | |

| | | | |
|---|---|---|---|
| | | PF3D7_0622900 | |
| | | PF3D7_0623100 | |
| | | PF3D7_0623600 | |
| | | PF3D7_0624000 | |
| ISWI | ISWI chromatin-remodeling complex ATPase | PF3D7_0624600 | PfSET10, PfMORC (2x), ISWI |
| | | PF3D7_0625600 | |
| | | PF3D7_0627500 | |
| N/A | transportin | PF3D7_0627700 | PfSET10, PfMORC |
| ACAS | acetyl-CoA synthetase | PF3D7_0627800 | PfSET10, PfMORC |
| | | PF3D7_0627900 | |
| | | PF3D7_0628100 | |
| | | PF3D7_0628600 | |
| | | PF3D7_0629100 | |
| | | PF3D7_0629400 | |
| | | PF3D7_0629700 | |
| | | PF3D7_0629800 | |
| | | PF3D7_0630300 | |
| N/A | deubiquitinating enzyme MINDY, putative | PF3D7_0630600 | PfSET10, PfMORC |
| | | PF3D7_0703200 | |
| | | PF3D7_0703500 | |
| | | PF3D7_0703900 | |
| | | PF3D7_0704200 | |
| | | PF3D7_0704400 | |
| | | PF3D7_0705300 | |
| MCM7 | DNA replication licensing factor MCM7 | PF3D7_0705400 | PfSET10, ISWI |
| N/A | importin-7, putative | PF3D7_0706000 | PfSET10, PfMORC |
| | | PF3D7_0706500 | |
| | | PF3D7_0707400 | |
| | | PF3D7_0707700 | |
| | | PF3D7_0708100 | |
| HSP110c | heat shock protein 110 | PF3D7_0708800 | PfSET10, PfMORC |
| | | PF3D7_0709000 | |
| | | PF3D7_0709300 | |
| | | PF3D7_0709600 | |
| | | PF3D7_0710200 | |
| | | PF3D7_0711500 | |
| | | PF3D7_0714200 | |
| N/A | transcription elongation factor s-II, putative | PF3D7_0714500 | PfSET10, PfMORC |
| | | PF3D7_0716000 | |
| EIF3I | eukaryotic translation initiation factor 3 subunit I, putative | PF3D7_0716800 | PfSET10, ISWI |
| | | PF3D7_0717700 | |
| | | PF3D7_0718500 | |
| | | PF3D7_0719000 | |
| | | PF3D7_0719300 | |
| OLA1 | Obg-like ATPase 1, putative | PF3D7_0722400 | PfSET10, ISWI |
| | | PF3D7_0722500 | |
| | | PF3D7_0723400 | |

| | | | |
|---|---|---|---|
| | | PF3D7_0723800 | |
| | | PF3D7_0724700 | |
| | | PF3D7_0725000 | |
| | | PF3D7_0725300 | |
| | | PF3D7_0726300 | |
| | | PF3D7_0726400 | |
| | | PF3D7_0726500 | |
| | | PF3D7_0727400 | |
| | | PF3D7_0727900 | |
| | | PF3D7_0728000 | |
| | | PF3D7_0728600 | |
| | | PF3D7_0729100 | |
| | | PF3D7_0729500 | |
| | | PF3D7_0730300 | |
| | | PF3D7_0730500 | |
| | | PF3D7_0801000 | |
| | | PF3D7_0801700 | |
| GDH3 | glutamate dehydrogenase, putative | PF3D7_0802000 | PfSET10, PfMORC |
| | | PF3D7_0802100 | |
| | | PF3D7_0802300 | |
| | | PF3D7_0803000 | |
| g-tub | tubulin gamma chain | PF3D7_0803700 | PfSET10, PfMORC (2x) |
| | | PF3D7_0804900 | |
| | | PF3D7_0805700 | |
| | | PF3D7_0807100 | |
| | | PF3D7_0807300 | |
| | | PF3D7_0807600 | |
| | | PF3D7_0810300 | |
| | | PF3D7_0810600 | |
| | | PF3D7_0810800 | |
| | | PF3D7_0811000 | |
| | | PF3D7_0811300 | |
| | | PF3D7_0811400 | |
| | | PF3D7_0811500 | |
| | | PF3D7_0811600 | |
| KARalpha | karyopherin alpha | PF3D7_0812400 | PfSET10, ISWI |
| | | PF3D7_0812700 | |
| | | PF3D7_0813300 | |
| | | PF3D7_0813400 | |
| ALBA1 | DNA/RNA-binding protein Alba 1 | PF3D7_0814200 | PfSET10, PfMORC |
| | | PF3D7_0814300 | |
| N/A | importin subunit beta, putative | PF3D7_0815200 | PfSET10, PfMORC |
| | | PF3D7_0815600 | |
| | | PF3D7_0816600 | |
| | | PF3D7_0817300 | |
| | | PF3D7_0817500 | |
| | | PF3D7_0818200 | |

| | | | |
|---|---|---|---|
| | | PF3D7_0818700 | |
| | | PF3D7_0818800 | |
| | | PF3D7_0818900 | |
| | | PF3D7_0819000 | |
| | | PF3D7_0819900 | |
| | | PF3D7_0820000 | |
| | | PF3D7_0821200 | |
| | | PF3D7_0821600 | |
| | | PF3D7_0822100 | |
| | | PF3D7_0822300 | |
| | | PF3D7_0822800 | |
| N/A | RNA-binding protein, putative | PF3D7_0823200 | PfSET10, PfMORC |
| | | PF3D7_0823300 | |
| | | PF3D7_0823800 | |
| | | PF3D7_0823900 | |
| | | PF3D7_0824600 | |
| | | PF3D7_0824800 | |
| | | PF3D7_0826300 | |
| | | PF3D7_0826700 | |
| | | PF3D7_0827000 | |
| SET3 | SET domain protein, putative | PF3D7_0827800 | PfSET10, PfMORC |
| | | PF3D7_0828500 | |
| NUP138 | nucleoporin NUP138, putative | PF3D7_0903500 | PfSET10, PfMORC |
| N/A | alpha tubulin 1 | PF3D7_0903700 | PfSET10, PfMORC |
| | | PF3D7_0904000 | |
| | | PF3D7_0904500 | |
| | | PF3D7_0904800 | |
| | | PF3D7_0906100 | |
| | | PF3D7_0906600 | |
| ClpY | ATP-dependent protease ATPase subunit ClpY | PF3D7_0907400 | PfSET10, PfMORC |
| | | PF3D7_0909400 | |
| | | PF3D7_0909800 | |
| | | PF3D7_0909900 | |
| N/A | exportin-7, putative | PF3D7_0910100 | PfSET10, PfMORC |
| N/A | conserved Plasmodium protein, unknown function | PF3D7_0910200 | PfSET10, PfMORC |
| | | PF3D7_0910400 | |
| | | PF3D7_0910900 | |
| | | PF3D7_0913200 | |
| | | PF3D7_0914800 | |
| | | PF3D7_0916700 | |
| | | PF3D7_0917000 | |
| | | PF3D7_0917600 | |
| | | PF3D7_0918300 | |
| | | PF3D7_0918900 | |
| | | PF3D7_0919000 | |
| IMPDH | inosine-5'-monophosphate dehydrogenase | PF3D7_0920800 | PfSET10, PfMORC |
| | | PF3D7_0920900 | |

| | | | |
|---|---|---|---|
| | | PF3D7_0922100 | |
| SAMS | S-adenosylmethionine synthetase | PF3D7_0922200 | PfAP2-P, PfSET10 |
| PGK | phosphoglycerate kinase | PF3D7_0922500 | PfAP2-P, PfSET10 |
| RPB3 | DNA-directed RNA polymerase II subunit RPB3, putative | PF3D7_0923000 | PfSET10, PfMORC |
| PABP2 | polyadenylate-binding protein 2, putative | PF3D7_0923900 | PfAP2-P, PfSET10 |
| SF3A3 | splicing factor 3A subunit 3, putative | PF3D7_0924700 | PfAP2-P, PfSET10 |
| HDAC1 | histone deacetylase 1 | PF3D7_0925700 | PfAP2-P, PfSET10, PfMORC, ISWI |
| | | PF3D7_0926100 | |
| | | PF3D7_0927300 | |
| N/A | RNA-binding protein, putative | PF3D7_0927600 | PfAP2-P, PfSET10 |
| | | PF3D7_0929000 | |
| N/A | RNA-binding protein, putative | PF3D7_0929200 | PfAP2-P, PfSET10 |
| | | PF3D7_0929400 | |
| | | PF3D7_0930300 | |
| | | PF3D7_0931400 | |
| N/A | proteasome subunit beta type-6, putative | PF3D7_0931800 | PfAP2-P, PfMORC |
| | | PF3D7_0932200 | |
| | | PF3D7_0932300 | |
| | | PF3D7_0932800 | |
| | | PF3D7_0933000 | |
| | | PF3D7_0933600 | |
| | | PF3D7_0934100 | |
| | | PF3D7_0934500 | |
| | | PF3D7_0934700 | |
| PKAc | cAMP-dependent protein kinase catalytic subunit | PF3D7_0934800 | PfAP2-P, PfSET10 |
| | | PF3D7_0935000 | |
| | | PF3D7_0935400 | |
| CLAG9 | cytoadherence linked asexual protein 9 | PF3D7_0935800 | PfAP2-P, PfMORC |
| | | PF3D7_1001600 | |
| TRAB2 | transformer-2 protein homolog beta, putative | PF3D7_1002400 | PfAP2-P, PfMORC |
| | | PF3D7_1002900 | |
| | | PF3D7_1003500 | |
| IMC1c | inner membrane complex protein 1c | PF3D7_1003600 | PfAP2-P, PfMORC |
| EFTUD2 | U5 small nuclear ribonucleoprotein component, putative | PF3D7_1003800 | PfAP2-P, PfSET10 |
| | | PF3D7_1004000 | |
| | | PF3D7_1004200 | |
| | | PF3D7_1004300 | |
| N/A | RNA-binding protein, putative | PF3D7_1004400 | PfAP2-P, PfSET10 |
| | | PF3D7_1004500 | |
| UPF1 | regulator of nonsense transcripts 1, putative | PF3D7_1005500 | PfAP2-P, PfSET10 |
| ALBA3 | DNA/RNA-binding protein Alba 3 | PF3D7_1006200 | PfAP2-P, PfSET10 |
| | | PF3D7_1006700 | |
| GBP2 | G-strand-binding protein 2 | PF3D7_1006800 | PfAP2-P, PfSET10, PfMORC |
| AP2-I | AP2 domain transcription factor AP2-I | PF3D7_1007700 | PfAP2-P, PfSET10, PfMORC |
| EIF3D | eukaryotic translation initiation factor 3 subunit D, putative | PF3D7_1007900 | PfAP2-P, PfSET10 |
| | | PF3D7_1008000 | |
| | | PF3D7_1008100 | |

| | | | |
|---|---|---|---|
| RPT2 | 26S protease regulatory subunit 4, putative | PF3D7_1008400 | PfAP2-P, PfMORC |
| N/A | Tubulin beta chain | PF3D7_1008700 | PfAP2-P, PfSET10 |
| NOP5 | nucleolar protein 5, putative | PF3D7_1008800 | PfAP2-P, PfSET10 |
| AK1 | adenylate kinase | PF3D7_1008900 | PfAP2-P, PfSET10 |
| | | PF3D7_1009200 | |
| | | PF3D7_1010200 | |
| eIF2beta | eukaryotic translation initiation factor 2 subunit beta | PF3D7_1010600 | PfAP2-P, PfSET10 |
| | | PF3D7_1011400 | |
| PREBP | PRE-binding protein | PF3D7_1011800 | PfAP2-P, PfSET10, ISWI |
| HGPRT | hypoxanthine-guanine phosphoribosyltransferase | PF3D7_1012400 | PfAP2-P, PfSET10 |
| | | PF3D7_1012500 | |
| | | PF3D7_1012700 | |
| | | PF3D7_1012900 | |
| | | PF3D7_1013100 | |
| | | PF3D7_1013300 | |
| | | PF3D7_1013600 | |
| | | PF3D7_1014100 | |
| | | PF3D7_1014300 | |
| | | PF3D7_1014600 | |
| KIC8 | protein KIC8 | PF3D7_1014900 | PfAP2-P, PfSET10 |
| | | PF3D7_1015200 | |
| | | PF3D7_1015400 | |
| HSP60 | heat shock protein 60 | PF3D7_1015600 | PfAP2-P, PfSET10, PfMORC |
| | | PF3D7_1015800 | |
| | | PF3D7_1015900 | |
| | | PF3D7_1016300 | |
| | | PF3D7_1016400 | |
| PPP8 | pseudophosphatase PPP8 | PF3D7_1018200 | PfAP2-P, PfSET10 |
| eIF2A | eukaryotic translation initiation factor subunit eIF2A | PF3D7_1019000 | PfAP2-P, PfSET10, PfMORC |
| N/A | 60S ribosomal protein L30e, putative | PF3D7_1019400 | PfAP2-P, PfMORC |
| | | PF3D7_1019700 | |
| N/A | N-acetyltransferase, GNAT family, putative | PF3D7_1020700 | PfSET10, PfMORC |
| | | PF3D7_1020900 | |
| | | PF3D7_1021800 | |
| N/A | PHAX domain-containing protein, putative | PF3D7_1021900 | PfAP2-P, PfSET10, PfMORC |
| SRSF4 | serine/arginine-rich splicing factor 4 | PF3D7_1022400 | PfAP2-P, PfSET10 |
| CHD1 | chromodomain-helicase-DNA-binding protein 1 homolog, putative | PF3D7_1023900 | PfAP2-P, PfSET10, PfMORC |
| | | PF3D7_1024800 | |
| | | PF3D7_1025000 | |
| | | PF3D7_1025300 | |
| | | PF3D7_1026000 | |
| | | PF3D7_1026800 | |
| nPrx | peroxiredoxin | PF3D7_1027300 | PfAP2-P, PfSET10 |
| | | PF3D7_1027700 | |
| RPL3 | 60S ribosomal protein L3 | PF3D7_1027800 | PfAP2-P, PfMORC |
| | | PF3D7_1028500 | |
| ADA | adenosine deaminase | PF3D7_1029600 | PfAP2-P, PfSET10 |

| | | | |
|---|---|---|---|
| PRP22 | pre-mRNA-splicing factor ATP-dependent RNA helicase PRP22, put | PF3D7_1030100 | PfAP2-P, PfSET10 |
| | | PF3D7_1031300 | |
| | | PF3D7_1031500 | |
| | | PF3D7_1031600 | |
| Der1-2 | DER1-like protein, putative | PF3D7_1032500 | PfAP2-P, PfMORC |
| | | PF3D7_1032700 | |
| AdoMetDC/ODC | S-adenosylmethionine decarboxylase/ornithine decarboxylase | PF3D7_1033100 | PfSET10, PfMORC |
| | | PF3D7_1033200 | |
| | | PF3D7_1033400 | |
| | | PF3D7_1033500 | |
| | | PF3D7_1033600 | |
| BDP1 | bromodomain protein 1 | PF3D7_1033700 | PfAP2-P, PfSET10 |
| | | PF3D7_1034000 | |
| | | PF3D7_1034400 | |
| | | PF3D7_1034700 | |
| | | PF3D7_1034900 | |
| N/A | conserved Plasmodium protein, unknown function | PF3D7_1036900 | PfAP2-P, PfSET10 |
| | | PF3D7_1037300 | |
| | | PF3D7_1037500 | |
| | | PF3D7_1037600 | |
| | | PF3D7_1037700 | |
| | | PF3D7_1103100 | |
| | | PF3D7_1103600 | |
| | | PF3D7_1103700 | |
| | | PF3D7_1103700 | |
| | | PF3D7_1103800 | |
| N/A | phenylalanine--tRNA ligase beta subunit | PF3D7_1104000 | PfAP2-P, PfMORC |
| | | PF3D7_1104100 | |
| SNF2L | chromatin remodeling protein | PF3D7_1104200 | PfAP2-P, PfSET10 |
| Trx-mero | thioredoxin-like mero protein | PF3D7_1104400 | PfAP2-P, PfSET10 |
| | | PF3D7_1105000 | |
| H2B | histone H2B | PF3D7_1105100 | PfAP2-P, PfSET10 |
| | | PF3D7_1105400 | |
| | | PF3D7_1105600 | |
| RTCB | tRNA-splicing ligase RtcB, putative | PF3D7_1105700 | PfAP2-P, PfSET10 |
| | | PF3D7_1105800 | |
| RUVB2 | RuvB-like helicase 2 | PF3D7_1106000 | PfSET10, PfMORC |
| | | PF3D7_1106700 | |
| | | PF3D7_1107000 | |
| PAIP1 | polyadenylate-binding protein-interacting protein 1, putative | PF3D7_1107300 | PfAP2-P, PfSET10, PfMORC |
| | | PF3D7_1107400 | |
| AP2-P | AP2 domain transcription factor, putative | PF3D7_1107800 | PfAP2-P, PfSET10, PfMORC, ISWI |
| CK2alpha | casein kinase 2, alpha subunit | PF3D7_1108400 | PfAP2-P, PfSET10 |
| SCS-alpha | succinate--CoA ligase [ADP-forming] subunit alpha, putative | PF3D7_1108500 | PfAP2-P, PfMORC |
| | | PF3D7_1108600 | |
| | | PF3D7_1108700 | |
| RPL36 | 60S ribosomal protein L36 | PF3D7_1109900 | PfAP2-P, PfMORC |

| | | | |
|---|---|---|---|
| PRPF6 | pre-mRNA-processing factor 6, putative | PF3D7_1110200 | PfAP2-P, PfSET10 |
| | | PF3D7_1110400 | |
| | | PF3D7_1110700 | |
| | | PF3D7_1110900 | |
| RFC5 | replication factor C subunit 5, putative | PF3D7_1111100 | PfAP2-P, PfSET10 |
| | | PF3D7_1111500 | |
| | | PF3D7_1112100 | |
| | | PF3D7_1112500 | |
| | | PF3D7_1115800 | |
| | | PF3D7_1116200 | |
| | | PF3D7_1116500 | |
| | | PF3D7_1116800 | |
| | | PF3D7_1117300 | |
| RAN | GTP-binding nuclear protein RAN/TC4 | PF3D7_1117700 | PfAP2-P, PfSET10 |
| | | PF3D7_1117800 | |
| | | PF3D7_1118100 | |
| N/A | heat shock protein 90, putative | PF3D7_1118200 | PfAP2-P, PfSET10 |
| | | PF3D7_1118300 | |
| NOP56 | nucleolar protein 56, putative | PF3D7_1118500 | PfAP2-P, PfSET10, ISWI |
| | | PF3D7_1118600 | |
| | | PF3D7_1119000 | |
| | | PF3D7_1119300 | |
| ASF1 | alternative splicing factor ASF-1, putative | PF3D7_1119800 | PfAP2-P, PfSET10 |
| | | PF3D7_1120000 | |
| PGM1 | phosphoglycerate mutase, putative | PF3D7_1120100 | PfAP2-P, PfSET10 |
| | | PF3D7_1121100 | |
| | | PF3D7_1121600 | |
| GCN20 | protein GCN20 | PF3D7_1121700 | PfAP2-P, PfSET10 |
| | | PF3D7_1121900 | |
| | | PF3D7_1123200 | |
| | | PF3D7_1123400 | |
| | | PF3D7_1123900 | |
| | | PF3D7_1124300 | |
| | | PF3D7_1124600 | |
| | | PF3D7_1124700 | |
| RPL35 | 60S ribosomal protein L35, putative | PF3D7_1124900 | PfAP2-P, PfMORC |
| SNRPD1 | small nuclear ribonucleoprotein Sm D1, putative | PF3D7_1125500 | PfAP2-P, PfSET10 |
| | | PF3D7_1126200 | |
| | | PF3D7_1126300 | |
| | | PF3D7_1126400 | |
| | | PF3D7_1126900 | |
| N/A | CRAL/TRIO domain-containing protein, putative | PF3D7_1127600 | PfAP2-P, PfSET10 |
| PCS5 | prefoldin subunit 5, putative | PF3D7_1128100 | PfSET10, PfMORC |
| | | PF3D7_1128200 | |
| | | PF3D7_1128500 | |
| SPDS | spermidine synthase | PF3D7_1129000 | PfAP2-P, PfSET10 |
| | | PF3D7_1129100 | |

| | | | |
|---|---|---|---|
| | | PF3D7_1129400 | |
| | | PF3D7_1130100 | |
| | | PF3D7_1130200 | |
| RPT5 | 26S protease regulatory subunit 6A, putative | PF3D7_1130400 | PfAP2-P, PfMORC |
| SMC1 | structural maintenance of chromosomes protein 1 | PF3D7_1130700 | PfAP2-P, PfSET10 |
| TCP1 | T-complex protein 1 subunit alpha | PF3D7_1132200 | PfAP2-P, PfSET10, PfMORC |
| N/A | nucleic acid binding protein, putative | PF3D7_1132300 | PfAP2-P, PfSET10 |
| N/A | conserved Plasmodium protein, unknown function | PF3D7_1133200 | PfSET10, PfMORC |
| | | PF3D7_1133800 | |
| HSP70-3 | heat shock protein 70 | PF3D7_1134000 | PfAP2-P, PfSET10 |
| | | PF3D7_1134100 | |
| | | PF3D7_1134600 | |
| | | PF3D7_1134700 | |
| N/A | coatomer subunit delta | PF3D7_1134800 | PfAP2-P, PfMORC |
| | | PF3D7_1135100 | |
| | | PF3D7_1135400 | |
| TSN | tudor staphylococcal nuclease | PF3D7_1136300 | PfAP2-P, PfSET10 |
| CK1 | casein kinase 1 | PF3D7_1136500 | PfAP2-P, PfSET10, ISWI |
| | | PF3D7_1137400 | |
| PPM2 | protein phosphatase PPM2, protein phosphatase 2C | PF3D7_1138500 | PfAP2-P, PfSET10, ISWI |
| | | PF3D7_1138800 | |
| AP2-G5 | AP2 domain transcription factor AP2-G5 | PF3D7_1139300 | PfSET10, PfMORC (2x) |
| | | PF3D7_1140200 | |
| | | PF3D7_1140400 | |
| N/A | EELM2 domain-containing protein, putative | PF3D7_1141800 | PfAP2-P, PfSET10 |
| N/A | DnaJ domain-containing protein, putative | PF3D7_1142100 | PfAP2-P, PfSET10 |
| | | PF3D7_1142500 | |
| | | PF3D7_1142600 | |
| | | PF3D7_1143200 | |
| | | PF3D7_1143300 | |
| | | PF3D7_1143400 | |
| RPS21 | 40S ribosomal protein S21 | PF3D7_1144000 | PfAP2-P, PfMORC |
| | | PF3D7_1144900 | |
| SEC21 | coatomer subunit gamma, putative | PF3D7_1145100 | PfAP2-P, PfSET10, PfMORC |
| | | PF3D7_1145200 | |
| DYN1 | dynamin-like protein | PF3D7_1145400 | PfAP2-P, PfSET10 |
| | | PF3D7_1146600 | |
| | | PF3D7_1147300 | |
| | | PF3D7_1147500 | |
| N/A | serine/threonine protein kinase, putative | PF3D7_1148000 | PfSET10, PfMORC |
| | | PF3D7_1201000 | |
| | | PF3D7_1202100 | |
| | | PF3D7_1202200 | |
| | | PF3D7_1202600 | |
| | | PF3D7_1202900 | |
| NAPL | nucleosome assembly protein | PF3D7_1203700 | PfAP2-P, PfSET10, PfMORC |
| | | PF3D7_1203900 | |

| | | | |
|---|---|---|---|
| | | PF3D7_1204300 | |
| | | PF3D7_1205500 | |
| N/A | tetratricopeptide repeat protein, putative | PF3D7_1205600 | PfAP2-P, PfSET10 |
| | | PF3D7_1205800 | |
| EIF3C | eukaryotic translation initiation factor 3 subunit C, putative | PF3D7_1206200 | PfAP2-P, PfSET10, PfMORC |
| | | PF3D7_1206600 | |
| N/A | conserved Plasmodium protein, unknown function | PF3D7_1207000 | PfAP2-P, PfSET10 |
| | | PF3D7_1207100 | |
| | | PF3D7_1208100 | |
| PPM11 | protein phosphatase PPM11, putative | PF3D7_1208900 | PfAP2-P, PfSET10 |
| | | PF3D7_1209200 | |
| | | PF3D7_1210400 | |
| | | PF3D7_1211200 | |
| | | PF3D7_1211300 | |
| PfJ4 | heat shock protein DNAJ homologue Pfj4 | PF3D7_1211400 | PfAP2-P, PfSET10 |
| | | PF3D7_1211600 | |
| MCM5 | DNA replication licensing factor MCM5, putative | PF3D7_1211700 | PfAP2-P, PfSET10, PfMORC, PfISWI |
| | | PF3D7_1211800 | |
| ATP4 | non-SERCA-type Ca2+ -transporting P-ATPase | PF3D7_1211900 | PfAP2-P, PfMORC |
| TPx(Gl) | glutathione peroxidase-like thioredoxin peroxidase | PF3D7_1212000 | PfAP2-P, PfMORC |
| EIF3A | eukaryotic translation initiation factor 3 subunit A, putative | PF3D7_1212700 | PfAP2-P, PfSET10 |
| | | PF3D7_1212900 | |
| | | PF3D7_1213200 | |
| | | PF3D7_1213700 | |
| | | PF3D7_1213900 | |
| | | PF3D7_1215000 | |
| | | PF3D7_1216200 | |
| | | PF3D7_1216300 | |
| N/A | DNA-binding chaperone, putative | PF3D7_1216900 | PfAP2-P, PfSET10, PfMORC |
| | | PF3D7_1219100 | |
| | | PF3D7_1220100 | |
| | | PF3D7_1220900 | |
| SET10 | histone-lysine N-methyltransferase, H3 lysine-4 specific | PF3D7_1221000 | PfAP2-P, PfSET10 |
| | | PF3D7_1222300 | |
| | | PF3D7_1222600 | |
| PKAr | cAMP-dependent protein kinase regulatory subunit | PF3D7_1223100 | PfAP2-P, PfSET10 |
| | | PF3D7_1223300 | |
| GCH1 | GTP cyclohydrolase 1 | PF3D7_1224000 | PfAP2-P, PfSET10 |
| PABP1 | polyadenylate-binding protein 1, putative | PF3D7_1224300 | PfAP2-P, PfSET10 |
| ASF1 | histone chaperone ASF1 | PF3D7_1224500 | PfAP2-P, PfSET10 |
| | | PF3D7_1224900 | |
| N/A | DNA-binding protein, putative | PF3D7_1225200 | PfSET10, PfMORC |
| UBA1 | ubiquitin-activating enzyme E1 | PF3D7_1225800 | PfAP2-P, PfSET10 |
| | | PF3D7_1226300 | |
| | | PF3D7_1226800 | |
| DH60 | DNA helicase 60 | PF3D7_1227100 | PfAP2-P, PfMORC |
| | | PF3D7_1227800 | |

| | | | |
|---|---|---|---|
| | | PF3D7_1228300 | |
| | | PF3D7_1228600 | |
| N/A | WD repeat-containing protein, putative | PF3D7_1228800 | PfAP2-P, PfSET10, PfMORC |
| | | PF3D7_1229300 | |
| | | PF3D7_1229400 | |
| CCT3 | T-complex protein 1 subunit gamma | PF3D7_1229500 | PfAP2-P, PfSET10 |
| | | PF3D7_1230400 | |
| SEC13 | protein transport protein SEC13 | PF3D7_1230700 | PfAP2-P, PfSET10, PfMORC |
| WTAP | pre-mRNA-splicing regulator WTAP, putative | PF3D7_1230800 | PfAP2-P, PfSET10 |
| | | PF3D7_1231100 | |
| PRP2 | pre-mRNA-splicing factor ATP-dependent RNA helicase PRP2, puta | PF3D7_1231600 | PfAP2-P, PfSET10 |
| CPN60 | 60 kDa chaperonin | PF3D7_1232100 | PfAP2-P, PfMORC |
| | | PF3D7_1234100 | |
| | | PF3D7_1234500 | |
| SF3B3 | splicing factor 3B subunit 3, putative | PF3D7_1234800 | PfAP2-P, PfSET10, PfMORC |
| NOT4 | CCR4-NOT transcription complex subunit 4, putative | PF3D7_1235300 | PfAP2-P, PfSET10 |
| | | PF3D7_1235500 | |
| | | PF3D7_1235600 | |
| | | PF3D7_1235900 | |
| | | PF3D7_1236100 | |
| | | PF3D7_1237500 | |
| N/A | conserved protein, unknown function | PF3D7_1237700 | PfAP2-P, PfSET10 |
| | | PF3D7_1238100 | |
| | | PF3D7_1238500 | |
| | | PF3D7_1238600 | |
| ACS11 | acyl-CoA synthetase | PF3D7_1238800 | PfAP2-P, PfSET10 |
| | | PF3D7_1239000 | |
| ApiAP2 | AP2 domain transcription factor, putative | PF3D7_1239200 | PfAP2-P, PfSET10, PfMORC (2x) |
| | | PF3D7_1239500 | |
| | | PF3D7_1239600 | |
| | | PF3D7_1239700 | |
| | | PF3D7_1241200 | |
| RFC4 | replication factor C subunit 4, putative | PF3D7_1241700 | PfAP2-P, PfSET10 |
| | | PF3D7_1242700 | |
| rabGDI | rab specific GDP dissociation inhibitor | PF3D7_1242800 | PfAP2-P, PfSET10, PfISWI |
| | | PF3D7_1243600 | |
| N/A | N-alpha-acetyltransferase 15, NatA auxiliary subunit, putative | PF3D7_1244100 | PfAP2-P, PfSET10 |
| | | PF3D7_1244200 | |
| | | PF3D7_1244800 | |
| KLP8 | kinesin-13, putative | PF3D7_1245100 | PfAP2-P, PfSET10 |
| ACT1 | actin I | PF3D7_1246200 | PfAP2-P, PfSET10 |
| | | PF3D7_1246400 | |
| | | PF3D7_1246800 | |
| FKBP35 | peptidyl-prolyl cis-trans isomerase FKBP35 | PF3D7_1247400 | PfAP2-P, PfSET10 |
| | | PF3D7_1247500 | |
| | | PF3D7_1248200 | |
| | | PF3D7_1248700 | |

| | | | |
|---|---|---|---|
| RPT6 | 26S protease regulatory subunit 8, putative | PF3D7_1248900 | PfAP2-P, PfMORC |
| | | PF3D7_1249100 | |
| PPM4 | protein phosphatase PPM4, putative | PF3D7_1249300 | PfAP2-P, PfSET10, PfMORC (2x) |
| THO2 | THO complex subunit 2, putative | PF3D7_1249800 | PfAP2-P, PfSET10 |
| | | PF3D7_1250600 | |
| | | PF3D7_1250800 | |
| | | PF3D7_1251200 | |
| RON3 | rhoptry neck protein 3 | PF3D7_1252100 | PfAP2-P, PfMORC |
| | | PF3D7_1301700 | |
| G27/25 | gamete antigen 27/25 | PF3D7_1302100 | PfAP2-P, PfSET10 |
| | | PF3D7_1302500 | |
| | | PF3D7_1302800 | |
| | | PF3D7_1303400 | |
| N/A | conserved Plasmodium protein, unknown function | PF3D7_1303800 | PfSET10, PfMORC |
| | | PF3D7_1304000 | |
| | | PF3D7_1304400 | |
| | | PF3D7_1304500 | |
| | | PF3D7_1304700 | |
| | | PF3D7_1305000 | |
| GCN1 | translational activator GCN1, putative | PF3D7_1305300 | PfAP2-P, PfSET10 |
| | | PF3D7_1305400 | |
| | | PF3D7_1305900 | |
| | | PF3D7_1306400 | |
| SNRPA | U1 small nuclear ribonucleoprotein A, putative | PF3D7_1306900 | PfAP2-P, PfSET10 |
| | | PF3D7_1307000 | |
| | | PF3D7_1307100 | |
| cpsSII | carbamoyl phosphate synthetase | PF3D7_1308200 | PfAP2-P, PfSET10 |
| RPS27 | 40S ribosomal protein S27 | PF3D7_1308300 | PfAP2-P, PfMORC |
| | | PF3D7_1308900 | |
| | | PF3D7_1309100 | |
| | | PF3D7_1309300 | |
| | | PF3D7_1309400 | |
| | | PF3D7_1309500 | |
| | | PF3D7_1310000 | |
| | | PF3D7_1310700 | |
| | | PF3D7_1311400 | |
| | | PF3D7_1311500 | |
| M1AAP | M1-family alanyl aminopeptidase | PF3D7_1311800 | PfAP2-P, PfISWI |
| vapA | V-type proton ATPase catalytic subunit A | PF3D7_1311900 | PfAP2-P, PfSET10 |
| | | PF3D7_1312600 | |
| | | PF3D7_1312900 | |
| | | PF3D7_1314500 | |
| | | PF3D7_1314700 | |
| | | PF3D7_1314700 | |
| | | PF3D7_1314900 | |
| | | PF3D7_1315300 | |
| N/A | conserved protein, unknown function | PF3D7_1315300 | PfAP2-P, PfMORC |

| | | | |
|---|---|---|---|
| | | PF3D7_1315400 | |
| | | PF3D7_1315700 | |
| | | PF3D7_1315800 | |
| | | PF3D7_1315900 | |
| | | PF3D7_1316500 | |
| | | PF3D7_1316600 | |
| | | PF3D7_1316800 | |
| | | PF3D7_1316900 | |
| MCM4 | DNA replication licensing factor MCM4 | PF3D7_1317100 | PfAP2-P, PfSET10, PfISWI |
| | | PF3D7_1317300 | |
| | | PF3D7_1317400 | |
| | | PF3D7_1317600 | |
| | | PF3D7_1317800 | |
| | | PF3D7_1318400 | |
| SEC63 | translocation protein SEC63, putative | PF3D7_1318800 | PfAP2-P, PfMORC |
| | | PF3D7_1319400 | |
| | | PF3D7_1320600 | |
| | | PF3D7_1320800 | |
| SF1 | splicing factor 1 | PF3D7_1321700 | PfAP2-P, PfSET10 |
| | | PF3D7_1322100 | |
| STU2 | protein STU2, putative | PF3D7_1322200 | PfAP2-P, PfSET10 |
| N/A | BRCT domain-containing protein, putative | PF3D7_1322300 | PfAP2-P, PfSET10 |
| | | PF3D7_1323100 | |
| | | PF3D7_1323200 | |
| RPL23 | 60S ribosomal protein L23 | PF3D7_1323400 | PfAP2-P, PfMORC |
| | | PF3D7_1323700 | |
| | | PF3D7_1324100 | |
| | | PF3D7_1324200 | |
| | | PF3D7_1324900 | |
| N/A | phosphoribosylpyrophosphate synthetase | PF3D7_1325100 | PfAP2-P, PfSET10 |
| | | PF3D7_1325400 | |
| | | PF3D7_1326300 | |
| | | PF3D7_1326400 | |
| | | PF3D7_1327800 | |
| | | PF3D7_1328100 | |
| | | PF3D7_1328200 | |
| | | PF3D7_1328300 | |
| | | PF3D7_1329000 | |
| MyoF | myosin F, putative | PF3D7_1329100 | PfAP2-P, PfSET10 |
| | | PF3D7_1329300 | |
| N/A | conserved protein, unknown function | PF3D7_1329500 | PfSET10, PfMORC |
| | | PF3D7_1329600 | |
| | | PF3D7_1330300 | |
| N/A | elongation factor Tu, putative | PF3D7_1330600 | PfAP2-P, PfSET10 |
| N/A | RNA-binding protein, putative | PF3D7_1330800 | PfAP2-P, PfSET10, PfMORC (2x) |
| | | PF3D7_1331800 | |
| | | PF3D7_1332900 | |

| | | | |
|---|---|---|---|
| | | PF3D7_1333300 | |
| | | PF3D7_1333600 | |
| | | PF3D7_1334100 | |
| | | PF3D7_1334200 | |
| | | PF3D7_1336400 | |
| | | PF3D7_1336800 | |
| | | PF3D7_1336900 | |
| RPL6 | 60S ribosomal protein L6, putative | PF3D7_1338200 | PfAP2-P, PfMORC |
| EF-1gamma | elongation factor 1-gamma, putative | PF3D7_1338300 | PfAP2-P, PfSET10 |
| | | PF3D7_1338700 | |
| | | PF3D7_1339700 | |
| | | PF3D7_1340100 | |
| DBR1 | RNA lariat debranching enzyme, putative | PF3D7_1340600 | PfAP2-P, PfSET10, PfMORC |
| | | PF3D7_1341200 | |
| N/A | 60S ribosomal protein L18-2, putative | PF3D7_1341300 | PfAP2-P, PfMORC |
| | | PF3D7_1341900 | |
| | | PF3D7_1342000 | |
| | | PF3D7_1342100 | |
| | | PF3D7_1342400 | |
| | | PF3D7_1342600 | |
| | | PF3D7_1342800 | |
| | | PF3D7_1343000 | |
| | | PF3D7_1343300 | |
| | | PF3D7_1343700 | |
| | | PF3D7_1343900 | |
| | | PF3D7_1344300 | |
| | | PF3D7_1345600 | |
| IDH | isocitrate dehydrogenase [NADP], mitochondrial | PF3D7_1345700 | PfAP2-P, PfSET10 |
| | | PF3D7_1345800 | |
| | | PF3D7_1346100 | |
| ALBA2 | DNA/RNA-binding protein Alba 2 | PF3D7_1346300 | PfAP2-P, PfSET10 |
| | | PF3D7_1347100 | |
| | | PF3D7_1347200 | |
| ALBA4 | DNA/RNA-binding protein Alba 4 | PF3D7_1347500 | PfAP2-P, PfSET10 |
| | | PF3D7_1347700 | |
| | | PF3D7_1349200 | |
| | | PF3D7_1350100 | |
| | | PF3D7_1351400 | |
| | | PF3D7_1352400 | |
| | | PF3D7_1352500 | |
| | | PF3D7_1352700 | |
| | | PF3D7_1353100 | |
| | | PF3D7_1353200 | |
| N/A | proteasome subunit alpha type-4, putative | PF3D7_1353800 | PfAP2-P, PfISWI |
| | | PF3D7_1353900 | |
| | | PF3D7_1354500 | |
| MCM6 | DNA replication licensing factor MCM6 | PF3D7_1355100 | PfAP2-P, PfSET10, PfISWI |

| | | | |
|---|---|---|---|
| SET6 | histone-lysine N-methyltransferase, putative | PF3D7_1355300 | PfAP2-P, PfSET10 |
| | | PF3D7_1355700 | |
| SF3B5 | splicing factor 3B subunit 5, putative | PF3D7_1355800 | PfSET10, PfMORC |
| | | PF3D7_1356100 | |
| N/A | elongation factor 1-alpha | PF3D7_1357100 | PfAP2-P, PfSET10 |
| | | PF3D7_1357500 | |
| | | PF3D7_1357700 | |
| CCT4 | T-complex protein 1 subunit delta | PF3D7_1357800 | PfAP2-P, PfSET10, PfISWI |
| | | PF3D7_1357900 | |
| | | PF3D7_1358200 | |
| RPS15 | 40S ribosomal protein S15 | PF3D7_1358800 | PfAP2-P, PfISWI |
| | | PF3D7_1358900 | |
| CELF1 | CUGBP Elav-like family member 1 | PF3D7_1359400 | PfAP2-P, PfSET10 |
| N/A | conserved Plasmodium protein, unknown function | PF3D7_1359600 | PfAP2-P, PfSET10 |
| | | PF3D7_1360800 | |
| N/A | RNA-binding protein, putative | PF3D7_1360900 | PfAP2-P, PfSET10 |
| | | PF3D7_1361000 | |
| SEC24A | protein transport protein Sec24A | PF3D7_1361100 | PfAP2-P, PfSET10 |
| | | PF3D7_1361800 | |
| PCNA1 | proliferating cell nuclear antigen 1 | PF3D7_1361900 | PfAP2-P, PfSET10, PfISWI |
| | | PF3D7_1362200 | |
| | | PF3D7_1362400 | |
| | | PF3D7_1362700 | |
| | | PF3D7_1363200 | |
| | | PF3D7_1363600 | |
| | | PF3D7_1364100 | |
| | | PF3D7_1364200 | |
| | | PF3D7_1364300 | |
| | | PF3D7_1364500 | |
| | | PF3D7_1364800 | |
| | | PF3D7_1365500 | |
| | | PF3D7_1365900 | |
| | | PF3D7_1366300 | |
| | | PF3D7_1366500 | |
| N/A | conserved protein, unknown function | PF3D7_1366900 | PfAP2-P, PfSET10 |
| SNP1 | U1 small nuclear ribonucleoprotein 70 kDa homolog, putative | PF3D7_1367100 | PfAP2-P, PfSET10 |
| ABCE1 | ABC transporter E family member 1, putative | PF3D7_1368200 | PfAP2-P, PfSET10 |
| N/A | ribosomal protein L1, putative | PF3D7_1368400 | PfAP2-P, PfMORC |
| | | PF3D7_1368800 | |
| N/A | nuclear cap-binding protein subunit 1, putative | PF3D7_1369500 | PfAP2-P, PfSET10 |
| | | PF3D7_1369700 | |
| MAHRP1 | membrane associated histidine-rich protein 1 | PF3D7_1370300 | PfAP2-P, PfMORC |
| | | PF3D7_1401600 | |
| | | PF3D7_1402300 | |
| | | PF3D7_1402500 | |
| | | PF3D7_1403100 | |
| | | PF3D7_1403900 | |

| | | | |
|---|---|---|---|
| | | PF3D7_1404000 | |
| RNR | ribonucleoside-diphosphate reductase small chain, putative | PF3D7_1405600 | PfAP2-P, PfSET10 |
| | | PF3D7_1406200 | |
| NOP1 | rRNA 2'-O-methyltransferase fibrillarin, putative | PF3D7_1407100 | PfAP2-P, PfSET10, PfMORC |
| | | PF3D7_1407300 | |
| | | PF3D7_1407800 | |
| | | PF3D7_1408200 | |
| | | PF3D7_1408400 | |
| | | PF3D7_1408600 | |
| N/A | conserved protein, unknown function | PF3D7_1408700 | PfAP2-P, PfSET10 |
| | | PF3D7_1409300 | |
| N/A | conserved protein, unknown function | PF3D7_1409400 | PfAP2-P, PfMORC |
| | | PF3D7_1409600 | |
| CELF2 | CUGBP Elav-like family member 2, putative | PF3D7_1409800 | PfAP2-P, PfSET10, PfMORC |
| | | PF3D7_1410300 | |
| | | PF3D7_1410400 | |
| eIF2gamma | eukaryotic translation initiation factor 2 subunit gamma, putative | PF3D7_1410600 | PfAP2-P, PfSET10 |
| | | PF3D7_1411300 | |
| MCMBP | mini-chromosome maintenance complex-binding protein | PF3D7_1412100 | PfSET10, PfMORC |
| ACT2 | actin II | PF3D7_1412500 | PfAP2-P, PfMORC |
| | | PF3D7_1412800 | |
| | | PF3D7_1414300 | |
| PP1 | serine/threonine protein phosphatase PP1 | PF3D7_1414400 | PfAP2-P, PfSET10 |
| SMB1 | small nuclear ribonucleoprotein-associated protein B, putative | PF3D7_1414800 | PfAP2-P, PfSET10 |
| | | PF3D7_1415200 | |
| N/A | RNA-binding protein Nova-1, putative | PF3D7_1415300 | PfAP2-P, PfMORC |
| N/A | Btz domain-containing protein, putative | PF3D7_1415400 | PfAP2-P, PfSET10 |
| SEY1 | protein SEY1, putative | PF3D7_1416100 | PfAP2-P, PfMORC |
| N/A | prefoldin subunit 2, putative | PF3D7_1416900 | PfAP2-P, PfSET10 |
| | | PF3D7_1417200 | |
| CBF5 | H/ACA ribonucleoprotein complex subunit 4, putative | PF3D7_1417500 | PfAP2-P, PfSET10 |
| MCM2 | DNA replication licensing factor MCM2 | PF3D7_1417800 | PfAP2-P, PfISWI |
| UFD1 | ubiquitin fusion degradation protein 1, putative | PF3D7_1418000 | PfAP2-P, PfSET10, PfMORC |
| | | PF3D7_1419000 | |
| EIF3H | eukaryotic translation initiation factor 3 subunit H, putative | PF3D7_1419700 | PfAP2-P, PfSET10, PfMORC |
| | | PF3D7_1419800 | |
| | | PF3D7_1419900 | |
| | | PF3D7_1420000 | |
| | | PF3D7_1420600 | |
| | | PF3D7_1421200 | |
| | | PF3D7_1422400 | |
| | | PF3D7_1422800 | |
| N/A | NTF2-like domain-containing protein, putative | PF3D7_1423700 | PfAP2-P, PfSET10 |
| | | PF3D7_1424100 | |
| | | PF3D7_1424400 | |
| | | PF3D7_1425900 | |
| | | PF3D7_1426000 | |

| | | | |
|---|---|---|---|
| BTF3 | transcription factor BTF3, putative | PF3D7_1426100 | PfAP2-P, PfSET10 |
| | | PF3D7_1426700 | |
| MSH2-1 | DNA mismatch repair protein MSH2, putative | PF3D7_1427500 | PfAP2-P, PfSET10 |
| N/A | leucine-rich repeat protein | PF3D7_1427900 | PfAP2-P, PfSET10 |
| N/A | proliferation-associated protein 2g4, putative | PF3D7_1428300 | PfAP2-P, PfMORC |
| | | PF3D7_1428800 | |
| | | PF3D7_1428900 | |
| | | PF3D7_1429800 | |
| | | PF3D7_1429900 | |
| | | PF3D7_1431600 | |
| N/A | 60S ribosomal protein L14, putative | PF3D7_1431700 | PfAP2-P, PfMORC |
| | | PF3D7_1432100 | |
| | | PF3D7_1433300 | |
| PHD2 | PHD finger protein PHD2, putative | PF3D7_1433400 | PfSET10, PfMORC |
| TOP2 | DNA topoisomerase 2 | PF3D7_1433500 | PfAP2-P, PfSET10, PfMORC |
| | | PF3D7_1434200 | |
| | | PF3D7_1434300 | |
| N/A | dynein-related AAA-type ATPase, putative | PF3D7_1434500 | PfSET10, PfMORC |
| | | PF3D7_1434600 | |
| | | PF3D7_1434800 | |
| | | PF3D7_1435500 | |
| | | PF3D7_1435600 | |
| | | PF3D7_1435700 | |
| | | PF3D7_1437000 | |
| N/A | ribonucleoside-diphosphate reductase large subunit, putative | PF3D7_1437200 | PfAP2-P, PfSET10 |
| | | PF3D7_1437400 | |
| | | PF3D7_1437700 | |
| HSP40 | HSP40, subfamily A | PF3D7_1437900 | PfAP2-P, PfSET10 |
| N/A | eukaryotic translation initiation factor eIF2A, putative | PF3D7_1438000 | PfAP2-P, PfSET10 |
| | | PF3D7_1438100 | |
| | | PF3D7_1438500 | |
| | | PF3D7_1438700 | |
| | | PF3D7_1438900 | |
| Trx-Px1 | thioredoxin peroxidase 1 | PF3D7_1438900 | PfAP2-P, PfSET10, PfMORC |
| | | PF3D7_1439700 | |
| | | PF3D7_1439800 | |
| | | PF3D7_1439900 | |
| | | PF3D7_1440100 | |
| | | PF3D7_1440500 | |
| | | PF3D7_1441200 | |
| FACT-S | FACT complex subunit SSRP1, putative | PF3D7_1441400 | PfSET10, PfISWI |
| RPA3 | replication factor A protein 3, putative | PF3D7_1442100 | PfSET10, PfMORC |
| | | PF3D7_1442300 | |
| | | PF3D7_1443000 | |
| FBPA | fructose-bisphosphate aldolase | PF3D7_1444800 | PfAP2-P, PfSET10 |
| | | PF3D7_1445100 | |
| | | PF3D7_1445400 | |

| | | | |
|---|---|---|---|
| | | PF3D7_1445600 | |
| | | PF3D7_1445700 | |
| DDX17 | ATP-dependent RNA helicase DDX17 | PF3D7_1445900 | PfAP2-P, PfSET10, PfMORC |
| | | PF3D7_1446200 | |
| | | PF3D7_1446500 | |
| CEN2 | centrin-2 | PF3D7_1446600 | PfAP2-P, PfSET10 |
| | | PF3D7_1446700 | |
| | | PF3D7_1447000 | |
| N/A | conserved protein, unknown function | PF3D7_1447700 | PfAP2-P, PfSET10 |
| | | PF3D7_1447800 | |
| | | PF3D7_1448000 | |
| | | PF3D7_1448300 | |
| | | PF3D7_1448500 | |
| ApiAP2 | AP2 domain transcription factor AP2-O5, putative | PF3D7_1449500 | PfAP2-P, PfSET10, PfMORC |
| | | PF3D7_1450100 | |
| | | PF3D7_1450400 | |
| eEF2 | elongation factor 2 | PF3D7_1451100 | PfAP2-P, PfSET10 |
| N/A | conserved protein, unknown function | PF3D7_1451200 | PfSET10, PfMORC |
| | | PF3D7_1451500 | |
| | | PF3D7_1451800 | |
| | | PF3D7_1451900 | |
| | | PF3D7_1452000 | |
| P23 | HSP90 co-chaperone p23 | PF3D7_1453700 | PfAP2-P, PfSET10 |
| | | PF3D7_1454200 | |
| | | PF3D7_1454700 | |
| | | PF3D7_1455500 | |
| N/A | conserved Plasmodium protein, unknown function | PF3D7_1455700 | PfAP2-P, PfMORC |
| | | PF3D7_1456000 | |
| | | PF3D7_1456500 | |
| | | PF3D7_1456800 | |
| | | PF3D7_1457000 | |
| | | PF3D7_1457200 | |
| N/A | MA3 domain-containing protein, putative | PF3D7_1457300 | PfAP2-P, PfSET10, PfMORC |
| | | PF3D7_1458600 | |
| | | PF3D7_1458800 | |
| DBP5 | ATP-dependent RNA helicase DBP5 | PF3D7_1459000 | PfAP2-P, PfSET10, PfMORC |
| N/A | conserved Plasmodium protein, unknown function | PF3D7_1459400 | PfAP2-P, PfSET10 |
| | | PF3D7_1459600 | |
| | | PF3D7_1460300 | |
| N/A | conserved Plasmodium protein, unknown function | PF3D7_1460500 | PfAP2-P, PfSET10, PfMORC |
| | | PF3D7_1460600 | |
| | | PF3D7_1460700 | |
| | | PF3D7_1460800 | |
| | | PF3D7_1461300 | |
| | | PF3D7_1461600 | |
| | | PF3D7_1461900 | |
| | | PF3D7_1462300 | |

| | | | |
|---|---|---|---|
| | | PF3D7_1462800 | |
| RFC3 | replication factor C subunit 3, putative | PF3D7_1463200 | PfAP2-P, PfSET10 |
| | | PF3D7_1463400 | |
| | | PF3D7_1464000 | |
| N/A | ATP synthase (C/AC39) subunit, putative | PF3D7_1464700 | PfAP2-P, PfSET10 |
| | | PF3D7_1465200 | |
| | | PF3D7_1465900 | |
| | | PF3D7_1465900 | |
| | | PF3D7_1466300 | |
| ApiAP2 | AP2 domain transcription factor AP2-EXP | PF3D7_1466400 | PfAP2-P, PfSET10 |
| | | PF3D7_1466800 | |
| | | PF3D7_1466900 | |
| | | PF3D7_1467500 | |
| MORC | MORC family protein | PF3D7_1468100 | PfAP2-P, PfSET10, PfMORC (2x), PfISWI |
| | | PF3D7_1468200 | |
| eIF4A | eukaryotic initiation factor 4A | PF3D7_1468700 | PfAP2-P, PfSET10 |
| | | PF3D7_1468900 | |
| | | PF3D7_1469700 | |
| | | PF3D7_1469800 | |
| | | PF3D7_1471100 | |
| N/A | diacylglycerol kinase, putative | PF3D7_1471400 | PfSET10, PfMORC |
| | | PF3D7_1472000 | |
| HDA1 | histone deacetylase, putative | PF3D7_1472200 | PfAP2-P, PfSET10 |
| | | PF3D7_1472900 | |
| N/A | DnaJ protein, putative | PF3D7_1473200 | PfAP2-P, PfSET10 |
| | | PF3D7_1473200 | |
| NUP116 | nucleoporin NUP116/NSP116, putative | PF3D7_1473700 | PfAP2-P, PfSET10 |
| SF3A1 | splicing factor 3A subunit 1, putative | PF3D7_1474500 | PfAP2-P, PfSET10 |
| | | PF3D7_1474800 | |
| | | PF3D7_1474900 | |
| | | PF3D7_1475000 | |
| | | PF3D7_1475600 | |

| Shared interactors |
|---|
| **GeneIDs** |
| PF3D7_0102900 |
| PF3D7_0103200 |
| PF3D7_0105200 |
| PF3D7_0105800 |
| PF3D7_0108300 |
| PF3D7_0109200 |
| PF3D7_0110500 |
| PF3D7_0111800 |
| PF3D7_0205600 |
| PF3D7_0209800 |
| PF3D7_0212100 |
| PF3D7_0212300 |
| PF3D7_0212500 |
| PF3D7_0213100 |
| PF3D7_0214000 |
| PF3D7_0214100 |
| PF3D7_0215700 |
| PF3D7_0217500 |
| PF3D7_0218500 |
| PF3D7_0219600 |
| PF3D7_0302900 |
| PF3D7_0305600 |
| PF3D7_0306800 |
| PF3D7_0307700 |
| PF3D7_0308000 |
| PF3D7_0308200 |
| PF3D7_0308600 |
| PF3D7_0309500 |
| PF3D7_0315100 |

PF3D7_0317200
PF3D7_0317400
PF3D7_0318200
PF3D7_0320300
PF3D7_0320800
PF3D7_0321500
PF3D7_0401800
PF3D7_0405400
PF3D7_0406100
PF3D7_0407800
PF3D7_0409600
PF3D7_0410800
PF3D7_0414000
PF3D7_0418200
PF3D7_0420300
PF3D7_0422500
PF3D7_0422700
PF3D7_0501000
PF3D7_0501500
PF3D7_0503400
PF3D7_0505500
PF3D7_0505800
PF3D7_0506500
PF3D7_0507700
PF3D7_0511800
PF3D7_0513300
PF3D7_0513600
PF3D7_0515000
PF3D7_0516800
PF3D7_0517400
PF3D7_0517700

PF3D7_0519400
PF3D7_0519800
PF3D7_0520400
PF3D7_0524000
PF3D7_0526500
PF3D7_0527000
PF3D7_0527500
PF3D7_0528100
PF3D7_0528200
PF3D7_0529400
PF3D7_0604100
PF3D7_0604500
PF3D7_0607000
PF3D7_0609000
PF3D7_0610900
PF3D7_0611400
PF3D7_0613800
PF3D7_0619900
PF3D7_0621800
PF3D7_0624600
PF3D7_0627700
PF3D7_0627800
PF3D7_0630600
PF3D7_0705400
PF3D7_0706000
PF3D7_0708800
PF3D7_0714500
PF3D7_0717700
PF3D7_0722400
PF3D7_0802000
PF3D7_0803700

PF3D7_0812400
PF3D7_0814200
PF3D7_0815200
PF3D7_0823200
PF3D7_0827800
PF3D7_0903500
PF3D7_0903700
PF3D7_0907400
PF3D7_0910100
PF3D7_0910200
PF3D7_0920800
PF3D7_0922200
PF3D7_0922500
PF3D7_0923000
PF3D7_0923900
PF3D7_0924700
PF3D7_0925700
PF3D7_0927600
PF3D7_0929200
PF3D7_0931800
PF3D7_0934800
PF3D7_0935800
PF3D7_1002400
PF3D7_1003600
PF3D7_1003800
PF3D7_1004400
PF3D7_1005500
PF3D7_1006200
PF3D7_1006800
PF3D7_1007700
PF3D7_1007900

PF3D7_1008400
PF3D7_1008700
PF3D7_1008800
PF3D7_1008900
PF3D7_1010600
PF3D7_1011800
PF3D7_1012400
PF3D7_1014900
PF3D7_1015600
PF3D7_1018200
PF3D7_1019000
PF3D7_1019400
PF3D7_1020700
PF3D7_1021900
PF3D7_1022400
PF3D7_1023900
PF3D7_1027300
PF3D7_1027800
PF3D7_1029600
PF3D7_1030100
PF3D7_1032500
PF3D7_1033100
PF3D7_1033700
PF3D7_1036900
PF3D7_1104000
PF3D7_1104200
PF3D7_1104400
PF3D7_1105100
PF3D7_1105700
PF3D7_1106000
PF3D7_1107300

PF3D7_1107800
PF3D7_1108400
PF3D7_1108500
PF3D7_1109900
PF3D7_1110200
PF3D7_1111100
PF3D7_1117700
PF3D7_1118200
PF3D7_1118500
PF3D7_1119800
PF3D7_1120100
PF3D7_1121700
PF3D7_1124900
PF3D7_1125500
PF3D7_1127600
PF3D7_1128100
PF3D7_1129000
PF3D7_1130400
PF3D7_1130700
PF3D7_1132200
PF3D7_1132300
PF3D7_1133200
PF3D7_1134000
PF3D7_1134800
PF3D7_1136300
PF3D7_1136500
PF3D7_1138500
PF3D7_1139300
PF3D7_1141800
PF3D7_1142100
PF3D7_1144000

PF3D7_1145100
PF3D7_1145400
PF3D7_1148000
PF3D7_1203700
PF3D7_1205600
PF3D7_1206200
PF3D7_1207000
PF3D7_1208900
PF3D7_1211400
PF3D7_1211700
PF3D7_1211900
PF3D7_1212000
PF3D7_1212700
PF3D7_1216900
PF3D7_1221000
PF3D7_1223100
PF3D7_1224000
PF3D7_1224300
PF3D7_1224500
PF3D7_1225200
PF3D7_1225800
PF3D7_1227100
PF3D7_1228800
PF3D7_1229500
PF3D7_1230700
PF3D7_1230800
PF3D7_1231600
PF3D7_1232100
PF3D7_1234800
PF3D7_1235300
PF3D7_1237700

PF3D7_1238800
PF3D7_1239200
PF3D7_1241700
PF3D7_1242800
PF3D7_1244100
PF3D7_1245100
PF3D7_1246200
PF3D7_1247400
PF3D7_1248900
PF3D7_1249300
PF3D7_1249800
PF3D7_1252100
PF3D7_1302100
PF3D7_1303800
PF3D7_1305300
PF3D7_1306900
PF3D7_1308200
PF3D7_1308300
PF3D7_1311800
PF3D7_1311900
PF3D7_1315300
PF3D7_1317100
PF3D7_1318800
PF3D7_1321700
PF3D7_1322200
PF3D7_1322300
PF3D7_1323400
PF3D7_1325100
PF3D7_1329100
PF3D7_1329500
PF3D7_1330600

PF3D7_1330800
PF3D7_1338200
PF3D7_1338300
PF3D7_1340600
PF3D7_1341300
PF3D7_1345700
PF3D7_1346300
PF3D7_1347500
PF3D7_1353800
PF3D7_1355100
PF3D7_1355300
PF3D7_1355800
PF3D7_1357100
PF3D7_1357800
PF3D7_1358800
PF3D7_1359400
PF3D7_1359600
PF3D7_1360900
PF3D7_1361100
PF3D7_1361900
PF3D7_1366900
PF3D7_1367100
PF3D7_1368200
PF3D7_1368400
PF3D7_1369500
PF3D7_1370300
PF3D7_1405600
PF3D7_1407100
PF3D7_1408700
PF3D7_1409400
PF3D7_1409800

PF3D7_1410600
PF3D7_1412100
PF3D7_1412500
PF3D7_1414400
PF3D7_1414800
PF3D7_1415300
PF3D7_1415400
PF3D7_1416100
PF3D7_1416900
PF3D7_1417500
PF3D7_1417800
PF3D7_1418000
PF3D7_1419700
PF3D7_1423700
PF3D7_1426100
PF3D7_1427500
PF3D7_1427900
PF3D7_1428300
PF3D7_1431700
PF3D7_1433400
PF3D7_1433500
PF3D7_1434500
PF3D7_1437200
PF3D7_1437900
PF3D7_1438000
PF3D7_1438900
PF3D7_1441400
PF3D7_1442100
PF3D7_1444800
PF3D7_1445900
PF3D7_1446600

PF3D7_1447700
PF3D7_1449500
PF3D7_1451100
PF3D7_1451200
PF3D7_1453700
PF3D7_1455700
PF3D7_1457300
PF3D7_1459000
PF3D7_1459400
PF3D7_1460500
PF3D7_1463200
PF3D7_1464700
PF3D7_1466400
PF3D7_1468100
PF3D7_1468700
PF3D7_1471400
PF3D7_1472200
PF3D7_1473200
PF3D7_1473700
PF3D7_1474500

Highest confidence (0.9000)

**k-means**

25 cluster

cluster with ≥3 proteins analyzed

disconnected nodes excluded

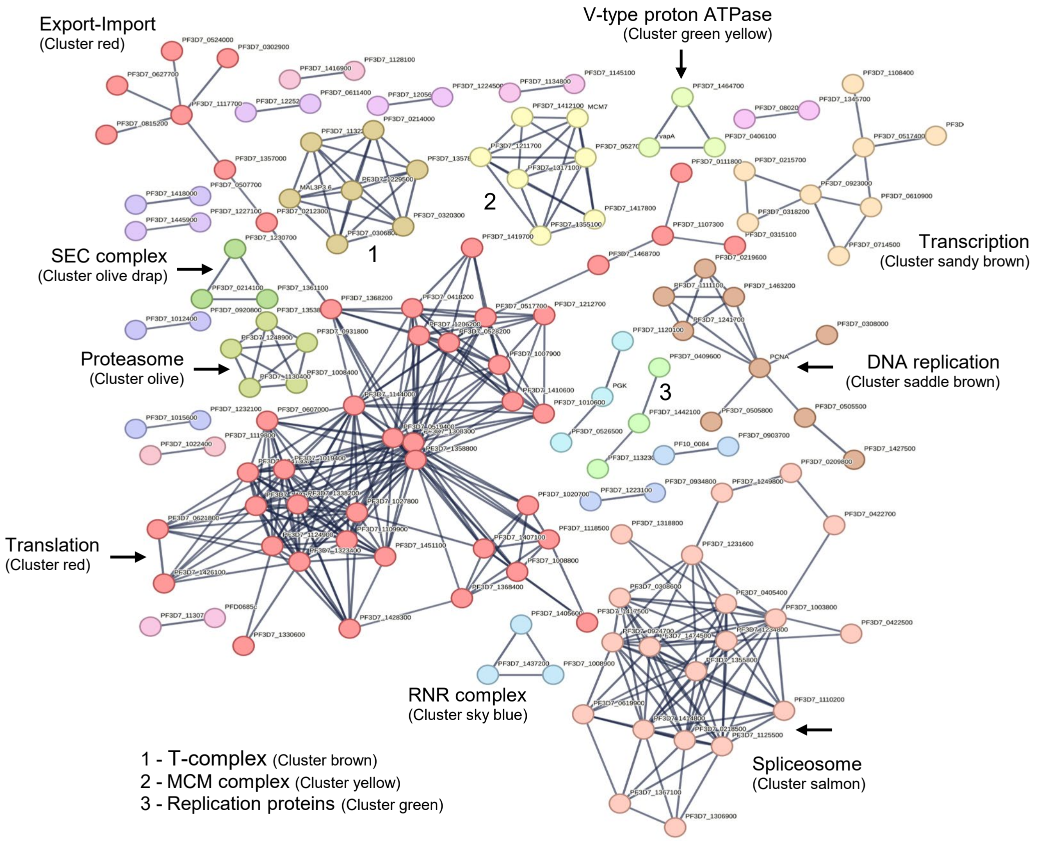

| Cluster red | Cluster number | Cluster color | Gene ID | Protein identifier | Protein description |
|---|---|---|---|---|---|
| | 1 | Red | PF3D7_0111800 | 36329.B9ZSJ0 | Eukaryotic translation initiation factor 4E, putative; Belongs to the eukaryotic initiation factor 4E family. |
| | 1 | Red | PF3D7_0212300 | 36329.O96203 | Peptide chain release factor subunit 1, putative. |
| | 1 | Red | PF3D7_0302900 | 36329.O77312 | Exportin-1, putative. |
| | 1 | Red | PF3D7_0315100 | 36329.O97266 | Eukaryotic translation initiation factor 4E; Belongs to the eukaryotic initiation factor 4E family. |
| | 1 | Red | PF3D7_0418200 | 36329.Q8I1Q5 | Eukaryotic translation initiation factor 3 subunit M; Component of the eukaryotic translation initiation factor 3 (eIF-3) complex, which is involved in protein synthesis of a specialized repertoire of mRNAs and, together with other initiation factors, stimulates binding of mRNA and methionyl-tRNAi to the 40S ribosome. The eIF-3 complex specifically targets and initiates translation of a subset of mRNAs involved in cell proliferation. |
| | 1 | Red | PF3D7_0517700 | 36329.Q8I3T1 | Eukaryotic translation initiation factor 3 subunit B; Component of the eukaryotic translation initiation factor 3 (eIF-3) complex, which is involved in protein synthesis and, together with other initiation factors, stimulates binding of mRNA and methionyl-tRNAi to the 40S ribosome; Belongs to the eIF-3 subunit B family. |
| | 1 | Red | PF3D7_0519400 | 36329.Q8I3R6 | 40S ribosomal protein S24; Belongs to the eukaryotic ribosomal protein eS24 family. |
| | 1 | Red | PF3D7_0524000 | 36329.Q8I3M5 | Karyopherin beta. |
| | 1 | Red | PF3D7_0528200 | 36329.Q8I3I5 | Eukaryotic translation initiation factor 3 subunit E; Component of the eukaryotic translation initiation factor 3 (eIF-3) complex, which is involved in protein synthesis of a specialized repertoire of mRNAs and, together with other initiation factors, stimulates binding of mRNA and methionyl-tRNAi to the 40S ribosome. The eIF-3 complex specifically targets and initiates translation of a subset of mRNAs involved in cell proliferation. |
| | 1 | Red | PF3D7_0607000 | 36329.C6KSR8 | Translation initiation factor IF-2, putative. |
| | 1 | Red | PF3D7_0621800 | 36329.C6KT55 | Nascent polypeptide-associated complex subunit alpha, putative. |
| | 1 | Red | PF3D7_0627700 | 36329.C6KTB3 | Transportin. |
| | 1 | Red | PF3D7_0815200 | 36329.Q8IAY9 | Importin subunit beta, putative. |
| | 1 | Red | PF3D7_1007900 | 36329.Q8IJW4 | Eukaryotic translation initiation factor 3 subunit D; mRNA cap-binding component of the eukaryotic translation initiation factor 3 (eIF-3) complex, which is involved in protein synthesis of a specialized repertoire of mRNAs and, together with other initiation factors, stimulates binding of mRNA and methionyl-tRNAi to the 40S ribosome. The eIF-3 complex specifically targets and initiates translation of a subset of mRNAs involved in cell proliferation. In the eIF-3 complex, eif3d specifically recognizes and binds the 7- methylguanosine cap of a subset of mRNAs. |
| | 1 | Red | PF3D7_1008800 | 36329.Q8IJV7 | Nucleolar protein 5, putative. |
| | 1 | Red | PF3D7_1010600 | 36329.Q8IJT9 | Eukaryotic translation initiation factor 2 subunit beta. |
| | 1 | Red | PF3D7_1019400 | 36329.Q8IJK8 | 60S ribosomal protein L30e, putative. |
| | 1 | Red | PF3D7_1020700 | 36329.Q8IJJ5 | RNA cytidine acetyltransferase; RNA cytidine acetyltransferase with specificity toward both 18S rRNA and tRNAs. Catalyzes the formation of N(4)-acetylcytidine (ac4C) in 18S rRNA. Required for early nucleolar cleavages of precursor rRNA at sites A0, A1 and A2 during 18S rRNA synthesis. Catalyzes the formation of ac4C in serine and leucine tRNAs. Requires a tRNA-binding adapter protein for full tRNA acetyltransferase activity but not for 18S rRNA acetylation. |
| | 1 | Red | PF3D7_1027800 | 36329.Q8IJC6 | 60S ribosomal protein L3; Belongs to the universal ribosomal protein uL3 family. |
| | 1 | Red | PF3D7_1107300 | 36329.Q8IIS9 | Polyadenylate-binding protein-interacting protein 1, putative. |
| | 1 | Red | PF3D7_1109900 | 36329.Q8I713 | 60S ribosomal protein L36; Belongs to the eukaryotic ribosomal protein eL36 family. |

| | | | | |
|---|---|---|---|---|
| 1 | Red | PF3D7_1117700 | 36329.Q7KQK6 | GTP-binding nuclear protein; GTP-binding protein involved in nucleocytoplasmic transport. Required for the import of protein into the nucleus and also for RNA export. Involved in chromatin condensation and control of cell cycle. Belongs to the small GTPase superfamily. Ran family. |
| 1 | Red | PF3D7_1118500 | 36329.Q8III3 | Nucleolar protein 56, putative. |
| 1 | Red | PF3D7_1124900 | 36329.Q8IIB4 | 60S ribosomal protein L35, putative. |
| 1 | Red | PF3D7_1144000 | 36329.Q8IHS5 | 40S ribosomal protein S21; Belongs to the eukaryotic ribosomal protein eS21 family. |
| 1 | Red | PF3D7_1206200 | 36329.Q8I5Y3 | Eukaryotic translation initiation factor 3 subunit C; Component of the eukaryotic translation initiation factor 3 (eIF-3) complex, which is involved in protein synthesis of a specialized repertoire of mRNAs and, together with other initiation factors, stimulates binding of mRNA and methionyl-tRNAi to the 40S ribosome. The eIF-3 complex specifically targets and initiates translation of a subset of mRNAs involved in cell proliferation. |
| 1 | Red | PF3D7_1212700 | 36329.Q8I5S6 | Eukaryotic translation initiation factor 3 subunit A; RNA-binding component of the eukaryotic translation initiation factor 3 (eIF-3) complex, which is involved in protein synthesis of a specialized repertoire of mRNAs and, together with other initiation factors, stimulates binding of mRNA and methionyl-tRNAi to the 40S ribosome. The eIF-3 complex specifically targets and initiates translation of a subset of mRNAs involved in cell proliferation. |
| 1 | Red | PF3D7_1308300 | 36329.Q8IEN2 | 40S ribosomal protein S27. |
| 1 | Red | PF3D7_1323400 | 36329.Q8IE82 | 60S ribosomal protein L23; Belongs to the universal ribosomal protein uL23 family. |
| 1 | Red | PF3D7_1330600 | 36329.Q8IE20 | Elongation factor Tu. |
| 1 | Red | PF3D7_1338200 | 36329.A0A5K1K8V8 | 60S ribosomal protein L6, putative. |
| 1 | Red | PF3D7_1341300 | 36329.C0H5G3 | 60S ribosomal protein L18-2, putative. |
| 1 | Red | PF3D7_1357000 | 36329.Q8I0P6 | Elongation factor 1-alpha; This protein promotes the GTP-dependent binding of aminoacyl- tRNA to the A-site of ribosomes during protein biosynthesis. |
| 1 | Red | PF3D7_1358800 | 36329.Q8IDB0 | 40S ribosomal protein S15; Belongs to the universal ribosomal protein uS15 family. |
| 1 | Red | PF3D7_1368200 | 36329.Q8I6Z4 | ABC transporter E family member 1, putative. |
| 1 | Red | PF3D7_1368400 | 36329.Q8ID26 | Ribosomal protein L1, putative. |
| 1 | Red | PF3D7_1407100 | 36329.Q8IM23 | rRNA 2'-O-methyltransferase fibrillarin, putative. |
| 1 | Red | PF3D7_1410600 | 36329.Q8ILY9 | Eukaryotic translation initiation factor 2 subunit gamma, putative. |
| 1 | Red | PF3D7_1417500 | 36329.Q8ILS0 | H/ACA ribonucleoprotein complex subunit 4, putative. |
| 1 | Red | PF3D7_1419700 | 36329.Q8ILQ3 | Uncharacterized protein. |
| 1 | Red | PF3D7_1426100 | 36329.Q8ILK2 | Nascent polypeptide-associated complex subunit beta. |
| 1 | Red | PF3D7_1428300 | 36329.Q8ILI2 | Proliferation-associated protein 2g4, putative. |
| 1 | Red | PF3D7_1431700 | 36329.Q8ILE8 | 60S ribosomal protein L14, putative. |
| 1 | Red | PF3D7_1451100 | 36329.Q8IKW5 | Elongation factor 2. |
| 1 | Red | PF3D7_1468700 | 36329.Q8IKF0 | Eukaryotic initiation factor 4A; Belongs to the DEAD box helicase family. |

| DNA replication | Cluster number | Cluster color | Gene ID | Protein identifier | Protein description |
|---|---|---|---|---|---|
| | 3 | Saddle Brown | PF3D7_1361900 | 36329.P61074 | Proliferating cell nuclear antigen; This protein is an auxiliary protein of DNA polymerase delta and is involved in the control of eukaryotic DNA replication by increasing the polymerase's processibility during elongation of the leading strand. |
| | 3 | Saddle Brown | PF3D7_0219600 | 36329.O96271 | Replication factor C subunit 1. |
| | 3 | Saddle Brown | PF3D7_0308000 | 36329.O77321 | DNA polymerase delta small subunit, putative. |
| | 3 | Saddle Brown | PF3D7_0505500 | 36329.Q8I447 | DNA mismatch repair protein; Component of the post-replicative DNA mismatch repair system (MMR). |
| | 3 | Saddle Brown | PF3D7_0505800 | 36329.Q8I444 | Small ubiquitin-related modifier. |
| | 3 | Saddle Brown | PF3D7_1111100 | 36329.Q8IIQ1 | Replication factor C subunit 5, putative. |
| | 3 | Saddle Brown | PF3D7_1241700 | 36329.Q8I512 | Replication factor C subunit 4, putative. |
| | 3 | Saddle Brown | PF3D7_1427500 | 36329.Q8ILI9 | DNA mismatch repair protein MSH2, putative. |
| | 3 | Saddle Brown | PF3D7_1463200 | 36329.Q8IKK4 | Replication factor C subunit 3, putative. |

| Transcription | Cluster number | Cluster color | Gene ID | Protein identifier | Protein description |
|---|---|---|---|---|---|
| | 4 | Sandy Brown | PF3D7_0215700 | 36329.O96236 | DNA-directed RNA polymerase subunit beta; DNA-dependent RNA polymerase catalyzes the transcription of DNA into RNA using the four ribonucleoside triphosphates as substrates. |
| | 4 | Sandy Brown | PF3D7_0318200 | 36329.O77375 | DNA-directed RNA polymerase subunit; DNA-dependent RNA polymerase catalyzes the transcription of DNA into RNA using the four ribonucleoside triphosphates as substrates. |
| | 4 | Sandy Brown | PF3D7_0517400 | 36329.Q8I3T4 | FACT complex subunit SPT16, putative. |
| | 4 | Sandy Brown | PF3D7_0610900 | 36329.C6KSV4 | Transcription elongation factor SPT5, putative. |
| | 4 | Sandy Brown | PF3D7_0714500 | 36329.Q8IBV2 | Transcription elongation factor s-II, putative. |
| | 4 | Sandy Brown | PF3D7_0923000 | 36329.Q8I2S6 | DNA-directed RNA polymerase II subunit RPB3, putative. |
| | 4 | Sandy Brown | PF3D7_1108400 | 36329.Q8IIR9 | Casein kinase 2, alpha subunit; Belongs to the protein kinase superfamily. |
| | 4 | Sandy Brown | PF3D7_1441400 | 36329.Q8IL56 | FACT complex subunit SSRP1; Component of the FACT complex, a general chromatin factor that acts to reorganize nucleosomes. |

| T-complex | Cluster number | Cluster color | Gene ID | Protein identifier | Protein description |
|---|---|---|---|---|---|
| | 5 | Brown | PF3D7_0308200 | 36329.O77323 | T-complex protein 1 subunit eta; Molecular chaperone; assists the folding of proteins upon ATP hydrolysis. Known to play a role, in vitro, in the folding of actin and tubulin (By similarity). |
| | 5 | Brown | PF3D7_0214000 | 36329.O96220 | T-complex protein 1 subunit theta. |
| | 5 | Brown | PF3D7_0306800 | 36329.O97247 | T-complex protein 1 subunit beta. |
| | 5 | Brown | PF3D7_0320300 | 36329.O97282 | T-complex protein 1 subunit epsilon. |
| | 5 | Brown | PF3D7_1132200 | 36329.Q8II43 | T-complex protein 1 subunit alpha. |
| | 5 | Brown | PF3D7_1229500 | 36329.Q8I5C4 | T-complex protein 1 subunit gamma. |
| | 5 | Brown | PF3D7_1357800 | 36329.C0H5I7 | T-complex protein 1 subunit delta. |

| MCM complex | Cluster numb | Cluster color | Gene ID | Protein identifier | Protein description |
|---|---|---|---|---|---|
| | 6 | Yellow | PF3D7_0705400 | 36329.Q8IC16 | DNA replication licensing factor MCM7; Acts as component of the mcm2-7 complex (mcm complex) which is the putative replicative helicase essential for 'once per cell cycle' DNA replication initiation and elongation in eukaryotic cells. The active ATPase sites in the mcm2-7 ring are formed through the interaction surfaces of two neighboring subunits such that a critical structure of a conserved arginine finger motif is provided in trans relative to the ATP-binding site of the Walker A box of the adjacent subunit. The six ATPase active sites, however, are likely to contribute differential [...] |
| | 6 | Yellow | PF3D7_0527000 | 36329.Q8I3J5 | DNA replication licensing factor MCM3, putative; Belongs to the MCM family. |
| | 6 | Yellow | PF3D7_1211700 | 36329.Q8I5T4 | DNA replication licensing factor MCM5, putative; Belongs to the MCM family. |
| | 6 | Yellow | PF3D7_1317100 | 36329.Q8IEE5 | DNA replication licensing factor MCM4; Belongs to the MCM family. |
| | 6 | Yellow | PF3D7_1355100 | 36329.Q8IDF0 | DNA helicase; Belongs to the MCM family. |
| | 6 | Yellow | PF3D7_1412100 | 36329.Q8ILX3 | Mini-chromosome maintenance complex-binding protein, putative. |
| | 6 | Yellow | PF3D7_1417800 | 36329.Q8ILR7 | DNA helicase. |

| Proteasome | Cluster number | Cluster color | Gene ID | Protein identifier | Protein description |
|---|---|---|---|---|---|
| | 7 | Olive | PF3D7_0931800 | 36329.Q8I0U7 | Proteasome subunit beta type-6, putative. |
| | 7 | Olive | PF3D7_1008400 | 36329.Q8IJW0 | 26S protease regulatory subunit 4, putative; Belongs to the AAA ATPase family. |
| | 7 | Olive | PF3D7_1130400 | 36329.Q8II60 | 26S protease regulatory subunit 6A, putative; Belongs to the AAA ATPase family. |
| | 7 | Olive | PF3D7_1248900 | 36329.Q8I4U5 | 26S protease regulatory subunit 8, putative; Belongs to the AAA ATPase family. |
| | 7 | Olive | PF3D7_1353800 | 36329.Q8IDG3 | Proteasome subunit alpha type. |

| V-type ATPase | Cluster number | Cluster color | Gene ID | Protein identifier | Protein description |
|---|---|---|---|---|---|
| | 8 | Green Yellow | PF3D7_0406100 | 36329.Q6ZMA8 | Vacuolar proton pump subunit B; Non-catalytic subunit of the peripheral V1 complex of vacuolar ATPase; Belongs to the ATPase alpha/beta chains family. |
| | 8 | Green Yellow | PF3D7_1464700 | 36329.Q8IKJ0 | V-type proton ATPase subunit; Subunit of the integral membrane V0 complex of vacuolar ATPase. Vacuolar ATPase is responsible for acidifying a variety of intracellular compartments in eukaryotic cells, thus providing most of the energy required for transport processes in the vacuolar system. Belongs to the V-ATPase V0D/AC39 subunit family. |
| | 8 | Green Yellow | PF3D7_1311900 | 36329.Q76NM6 | V-type proton ATPase catalytic subunit A; Catalytic subunit of the peripheral V1 complex of vacuolar ATPase. V-ATPase vacuolar ATPase is responsible for acidifying a variety of intracellular compartments in eukaryotic cells; Belongs to the ATPase alpha/beta chains family. |

| SEC complex | Cluster number | Cluster color | Gene ID | Protein identifier | Protein description |
|---|---|---|---|---|---|
| | 9 | Olive Drab | PF3D7_0214100 | 36329.O96221 | Protein transport protein SEC31. |
| | 9 | Olive Drab | PF3D7_1230700 | 36329.Q8I5B3 | Protein transport protein SEC13; Belongs to the WD repeat SEC13 family. |
| | 9 | Olive Drab | PF3D7_1361100 | 36329.C0H5J6 | Protein transport protein Sec24A. |

| Replication proteins | Cluster number | Cluster color | Gene ID | Protein identifier | Protein description |
|---|---|---|---|---|---|
| | 10 | Green | PF3D7_0409600 | 36329.Q9U0J0 | Replication protein A1, large subunit. |
| | 10 | Green | PF3D7_1132300 | 36329.Q8II42 | Nucleic acid binding protein, putative. |
| | 10 | Green | PF3D7_1442100 | 36329.C6S3I6 | Replication factor A protein 3, putative. |

| RNR complex | Cluster number | Cluster color | Gene ID | Protein identifier | Protein description |
|---|---|---|---|---|---|
| | 12 | Sky Blue | PF3D7_1008900 | 36329.Q8IJV6 | Adenylate kinase; Belongs to the adenylate kinase family. |
| | 12 | Sky Blue | PF3D7_1405600 | 36329.Q8IM38 | Ribonucleoside-diphosphate reductase small chain, putative. |
| | 12 | Sky Blue | PF3D7_1437200 | 36329.Q8IL94 | Ribonucleoside-diphosphate reductase; Provides the precursors necessary for DNA synthesis. Catalyzes the biosynthesis of deoxyribonucleotides from the corresponding ribonucleotides. |

| #Clustering method | Cluster number | Cluster color | Gene count | Gene ID | Protein identifier | Protein description |
|---|---|---|---|---|---|---|
| kmeans | 1 | Red | 45 | PF3D7_01118 | 36329.B9ZSJ0 | Eukaryotic translation initiation factor 4E, putative; Belongs to the eukaryotic initiation factor 4E family. |
| kmeans | 1 | Red | 45 | PF3D7_02123 | 36329.O96203 | Peptide chain release factor subunit 1, putative. |
| kmeans | 1 | Red | 45 | PF3D7_03029 | 36329.O77312 | Exportin-1, putative. |
| kmeans | 1 | Red | 45 | PF3D7_03151 | 36329.O97266 | Eukaryotic translation initiation factor 4E; Belongs to the eukaryotic initiation factor 4E family. |
| kmeans | 1 | Red | 45 | PF3D7_04182 | 36329.Q8I1Q5 | Eukaryotic translation initiation factor 3 subunit M; Component of the eukaryotic translation initiation factor 3 (eIF-3) complex, which is involved in protein synthesis of a specialized repertoire of mRNAs and, together with other initiation factors, stimulates binding of mRNA and methionyl-tRNAi to the 40S ribosome. The eIF-3 complex specifically targets and initiates translation of a subset of mRNAs involved in cell proliferation. |
| kmeans | 1 | Red | 45 | PF3D7_05177 | 36329.Q8I3T1 | Eukaryotic translation initiation factor 3 subunit B; Component of the eukaryotic translation initiation factor 3 (eIF-3) complex, which is involved in protein synthesis and, together with other initiation factors, stimulates binding of mRNA and methionyl-tRNAi to the 40S ribosome; Belongs to the eIF-3 subunit B family. |
| kmeans | 1 | Red | 45 | PF3D7_05194 | 36329.Q8I3R6 | 40S ribosomal protein S24; Belongs to the eukaryotic ribosomal protein eS24 family. |
| kmeans | 1 | Red | 45 | PF3D7_05240 | 36329.Q8I3M5 | Karyopherin beta. |
| kmeans | 1 | Red | 45 | PF3D7_05282 | 36329.Q8I3I5 | Eukaryotic translation initiation factor 3 subunit E; Component of the eukaryotic translation initiation factor 3 (eIF-3) complex, which is involved in protein synthesis of a specialized repertoire of mRNAs and, together with other initiation factors, stimulates binding of mRNA and methionyl-tRNAi to the 40S ribosome. The eIF-3 complex specifically targets and initiates translation of a subset of mRNAs involved in cell proliferation. |
| kmeans | 1 | Red | 45 | PF3D7_06070 | 36329.C6KSR8 | Translation initiation factor IF-2, putative. |
| kmeans | 1 | Red | 45 | PF3D7_06218 | 36329.C6KT55 | Nascent polypeptide-associated complex subunit alpha, putative. |
| kmeans | 1 | Red | 45 | PF3D7_06277 | 36329.C6KTB3 | Transportin. |
| kmeans | 1 | Red | 45 | PF3D7_08152 | 36329.Q8IAY9 | Importin subunit beta, putative. |
| kmeans | 1 | Red | 45 | PF3D7_10079 | 36329.Q8IJW4 | Eukaryotic translation initiation factor 3 subunit D; mRNA cap-binding component of the eukaryotic translation initiation factor 3 (eIF-3) complex, which is involved in protein synthesis of a specialized repertoire of mRNAs and, together with other initiation factors, stimulates binding of mRNA and methionyl-tRNAi to the 40S ribosome. The eIF-3 complex specifically targets and initiates translation of a subset of mRNAs involved in cell proliferation. In the eIF-3 complex, eif3d specifically recognizes and binds the 7-methylguanosine cap of a subset of mRNAs. |
| kmeans | 1 | Red | 45 | PF3D7_10088 | 36329.Q8IJV7 | Nucleolar protein 5, putative. |
| kmeans | 1 | Red | 45 | PF3D7_10106 | 36329.Q8IJT9 | Eukaryotic translation initiation factor 2 subunit beta. |
| kmeans | 1 | Red | 45 | PF3D7_10194 | 36329.Q8IJK8 | 60S ribosomal protein L30e, putative. |
| kmeans | 1 | Red | 45 | PF3D7_10207 | 36329.Q8IJJ5 | RNA cytidine acetyltransferase; RNA cytidine acetyltransferase with specificity toward both 18S rRNA and tRNAs. Catalyzes the formation of N(4)-acetylcytidine (ac4C) in 18S rRNA. Required for early nucleolar cleavages of precursor rRNA at sites A0, A1 and A2 during 18S rRNA synthesis. Catalyzes the formation of ac4C in serine and leucine tRNAs. Requires a tRNA-binding adapter protein for full tRNA acetyltransferase activity but not for 18S rRNA acetylation. |
| kmeans | 1 | Red | 45 | PF3D7_10278 | 36329.Q8IJC6 | 60S ribosomal protein L3; Belongs to the universal ribosomal protein uL3 family. |
| kmeans | 1 | Red | 45 | PF3D7_11073 | 36329.Q8IIS9 | Polyadenylate-binding protein-interacting protein 1, putative. |
| kmeans | 1 | Red | 45 | PF3D7_11099 | 36329.Q8I713 | 60S ribosomal protein L36; Belongs to the eukaryotic ribosomal protein eL36 family. |
| kmeans | 1 | Red | 45 | PF3D7_11177 | 36329.Q7KQK6 | GTP-binding nuclear protein; GTP-binding protein involved in nucleocytoplasmic transport. Required for the import of protein into the nucleus and also for RNA export. Involved in chromatin condensation and control of cell cycle. Belongs to the small GTPase superfamily. Ran family. |
| kmeans | 1 | Red | 45 | PF3D7_11185 | 36329.Q8III3 | Nucleolar protein 56, putative. |
| kmeans | 1 | Red | 45 | PF3D7_11249 | 36329.Q8IIB4 | 60S ribosomal protein L35, putative. |
| kmeans | 1 | Red | 45 | PF3D7_11440 | 36329.Q8IHS5 | 40S ribosomal protein S21; Belongs to the eukaryotic ribosomal protein eS21 family. |

| | | | | |
|---|---|---|---|---|
| kmeans | 1 Red | 45 | PF3D7_12062 36329.Q8I5Y3 | Eukaryotic translation initiation factor 3 subunit C; Component of the eukaryotic translation initiation factor 3 (eIF-3) complex, which is involved in protein synthesis of a specialized repertoire of mRNAs and, together with other initiation factors, stimulates binding of mRNA and methionyl-tRNAi to the 40S ribosome. The eIF-3 complex specifically targets and initiates translation of a subset of mRNAs involved in cell proliferation. |
| kmeans | 1 Red | 45 | PF3D7_12127 36329.Q8I5S6 | Eukaryotic translation initiation factor 3 subunit A; RNA-binding component of the eukaryotic translation initiation factor 3 (eIF-3) complex, which is involved in protein synthesis of a specialized repertoire of mRNAs and, together with other initiation factors, stimulates binding of mRNA and methionyl-tRNAi to the 40S ribosome. The eIF-3 complex specifically targets and initiates translation of a subset of mRNAs involved in cell proliferation. |
| kmeans | 1 Red | 45 | PF3D7_13083 36329.Q8IEN2 | 40S ribosomal protein S27. |
| kmeans | 1 Red | 45 | PF3D7_13234 36329.Q8IE82 | 60S ribosomal protein L23; Belongs to the universal ribosomal protein uL23 family. |
| kmeans | 1 Red | 45 | PF3D7_13306 36329.Q8IE20 | Elongation factor Tu. |
| kmeans | 1 Red | 45 | PF3D7_13382 36329.A0A5K1K8V8 | 60S ribosomal protein L6, putative. |
| kmeans | 1 Red | 45 | PF3D7_13413 36329.C0H5G3 | 60S ribosomal protein L18-2, putative. |
| kmeans | 1 Red | 45 | PF3D7_13570 36329.Q8I0P6 | Elongation factor 1-alpha; This protein promotes the GTP-dependent binding of aminoacyl- tRNA to the A-site of ribosomes during protein biosynthesis. |
| kmeans | 1 Red | 45 | PF3D7_13588 36329.Q8IDB0 | 40S ribosomal protein S15; Belongs to the universal ribosomal protein uS15 family. |
| kmeans | 1 Red | 45 | PF3D7_13682 36329.Q8I6Z4 | ABC transporter E family member 1, putative. |
| kmeans | 1 Red | 45 | PF3D7_13684 36329.Q8ID26 | Ribosomal protein L1, putative. |
| kmeans | 1 Red | 45 | PF3D7_14071 36329.Q8IM23 | rRNA 2'-O-methyltransferase fibrillarin, putative. |
| kmeans | 1 Red | 45 | PF3D7_14106 36329.Q8ILY9 | Eukaryotic translation initiation factor 2 subunit gamma, putative. |
| kmeans | 1 Red | 45 | PF3D7_14175 36329.Q8ILS0 | H/ACA ribonucleoprotein complex subunit 4, putative. |
| kmeans | 1 Red | 45 | PF3D7_14197 36329.Q8ILQ3 | Uncharacterized protein. |
| kmeans | 1 Red | 45 | PF3D7_14261 36329.Q8ILK2 | Nascent polypeptide-associated complex subunit beta. |
| kmeans | 1 Red | 45 | PF3D7_14283 36329.Q8ILI2 | Proliferation-associated protein 2g4, putative. |
| kmeans | 1 Red | 45 | PF3D7_14317 36329.Q8ILE8 | 60S ribosomal protein L14, putative. |
| kmeans | 1 Red | 45 | PF3D7_14511 36329.Q8IKW5 | Elongation factor 2. |
| kmeans | 1 Red | 45 | PF3D7_14687 36329.Q8IKF0 | Eukaryotic initiation factor 4A; Belongs to the DEAD box helicase family. |
| kmeans | 2 Salmon | 20 | PF3D7_02098 36329.Q9TY94 | ATP-dependent RNA helicase UAP56. |
| kmeans | 2 Salmon | 20 | PF3D7_02185 36329.O96265 | Small nuclear ribonucleoprotein Sm D2. |
| kmeans | 2 Salmon | 20 | PF3D7_03086 36329.O77325 | Pre-mRNA-processing factor 19, putative. |
| kmeans | 2 Salmon | 20 | PF3D7_04054 36329.Q8I1X5 | Pre-mRNA-processing-splicing factor 8, putative. |
| kmeans | 2 Salmon | 20 | PF3D7_04225 36329.Q8IFP1 | Pre-mRNA-splicing helicase BRR2, putative. |
| kmeans | 2 Salmon | 20 | PF3D7_04227 36329.Q8IFN9 | Eukaryotic initiation factor 4A-III, putative; Belongs to the DEAD box helicase family. |
| kmeans | 2 Salmon | 20 | PF3D7_06199 36329.C6KT39 | Splicing factor 3A subunit 2, putative. |
| kmeans | 2 Salmon | 20 | PF3D7_09247 36329.Q8I2R0 | Splicing factor 3A subunit 3, putative. |
| kmeans | 2 Salmon | 20 | PF3D7_10038 36329.Q8IJZ9 | U5 small nuclear ribonucleoprotein component, putative. |
| kmeans | 2 Salmon | 20 | PF3D7_11102 36329.Q8IIR0 | Pre-mRNA-processing factor 6, putative. |
| kmeans | 2 Salmon | 20 | PF3D7_11255 36329.Q8IIA8 | Small nuclear ribonucleoprotein Sm D1; Essential for pre-mRNA splicing. Implicated in the formation of stable, biologically active snRNP structures. Belongs to the snRNP core protein family. |
| kmeans | 2 Salmon | 20 | PF3D7_12316 36329.Q8I5A4 | Pre-mRNA-splicing factor ATP-dependent RNA helicase PRP2, putative. |
| kmeans | 2 Salmon | 20 | PF3D7_12348 36329.Q8I574 | Splicing factor 3B subunit 3, putative. |
| kmeans | 2 Salmon | 20 | PF3D7_12498 36329.Q8I4T6 | THO complex subunit 2, putative. |
| kmeans | 2 Salmon | 20 | PF3D7_13069 36329.C0H5A4 | U1 small nuclear ribonucleoprotein A, putative. |
| kmeans | 2 Salmon | 20 | PF3D7_13188 36329.Q8IEC8 | Translocation protein SEC63, putative. |
| kmeans | 2 Salmon | 20 | PF3D7_13558 36329.C0H5I5 | Splicing factor subunit; Belongs to the SF3B5 family. |
| kmeans | 2 Salmon | 20 | PF3D7_13671 36329.Q8ID37 | U1 small nuclear ribonucleoprotein 70 kDa homolog, putative. |
| kmeans | 2 Salmon | 20 | PF3D7_14148 36329.Q8ILU8 | Small nuclear ribonucleoprotein-associated protein B, putative. |
| kmeans | 2 Salmon | 20 | PF3D7_14745 36329.Q8IK93 | Splicing factor 3A subunit 1, putative. |

| | | | | | |
|---|---|---|---|---|---|
| kmeans | 3 | Saddle Brown | 9 | PCNA | 36329.P61074 | Proliferating cell nuclear antigen; This protein is an auxiliary protein of DNA polymerase delta and is involved in the control of eukaryotic DNA replication by increasing the polymerase's processibility during elongation of the leading strand. |
| kmeans | 3 | Saddle Brown | 9 | PF3D7_02196 | 36329.O96271 | Replication factor C subunit 1. |
| kmeans | 3 | Saddle Brown | 9 | PF3D7_03080 | 36329.O77321 | DNA polymerase delta small subunit, putative. |
| kmeans | 3 | Saddle Brown | 9 | PF3D7_05055 | 36329.Q8I447 | DNA mismatch repair protein; Component of the post-replicative DNA mismatch repair system (MMR). |
| kmeans | 3 | Saddle Brown | 9 | PF3D7_05058 | 36329.Q8I444 | Small ubiquitin-related modifier. |
| kmeans | 3 | Saddle Brown | 9 | PF3D7_11111 | 36329.Q8IIQ1 | Replication factor C subunit 5, putative. |
| kmeans | 3 | Saddle Brown | 9 | PF3D7_12417 | 36329.Q8I512 | Replication factor C subunit 4, putative. |
| kmeans | 3 | Saddle Brown | 9 | PF3D7_14275 | 36329.Q8ILI9 | DNA mismatch repair protein MSH2, putative. |
| kmeans | 3 | Saddle Brown | 9 | PF3D7_14632 | 36329.Q8IKK4 | Replication factor C subunit 3, putative. |
| kmeans | 4 | Sandy Brown | 8 | PF3D7_02157 | 36329.O96236 | DNA-directed RNA polymerase subunit beta; DNA-dependent RNA polymerase catalyzes the transcription of DNA into RNA using the four ribonucleoside triphosphates as substrates. |
| kmeans | 4 | Sandy Brown | 8 | PF3D7_03182 | 36329.O77375 | DNA-directed RNA polymerase subunit; DNA-dependent RNA polymerase catalyzes the transcription of DNA into RNA using the four ribonucleoside triphosphates as substrates. |
| kmeans | 4 | Sandy Brown | 8 | PF3D7_05174 | 36329.Q8I3T4 | FACT complex subunit SPT16, putative. |
| kmeans | 4 | Sandy Brown | 8 | PF3D7_06109 | 36329.C6KSV4 | Transcription elongation factor SPT5, putative. |
| kmeans | 4 | Sandy Brown | 8 | PF3D7_07145 | 36329.Q8IBV2 | Transcription elongation factor s-II, putative. |
| kmeans | 4 | Sandy Brown | 8 | PF3D7_09230 | 36329.Q8I2S6 | DNA-directed RNA polymerase II subunit RPB3, putative. |
| kmeans | 4 | Sandy Brown | 8 | PF3D7_11084 | 36329.Q8IIR9 | Casein kinase 2, alpha subunit; Belongs to the protein kinase superfamily. |
| kmeans | 4 | Sandy Brown | 8 | PF3D7_14414 | 36329.Q8IL56 | FACT complex subunit SSRP1; Component of the FACT complex, a general chromatin factor that acts to reorganize nucleosomes. The FACT complex is involved in multiple processes that require DNA as a template such as mRNA elongation, DNA replication and DNA repair. During transcription elongation the FACT complex acts as a histone chaperone that both destabilizes and restores nucleosomal structure. It facilitates the passage of RNA polymerase II and transcription by promoting the dissociation of one histone H2A-H2B dimer from the nucleosome, then subsequently promotes the reestablishment o [...] |
| kmeans | 5 | Brown | 7 | MAL3P3.6 | 36329.O77323 | T-complex protein 1 subunit eta; Molecular chaperone; assists the folding of proteins upon ATP hydrolysis. Known to play a role, in vitro, in the folding of actin and tubulin (By similarity). |
| kmeans | 5 | Brown | 7 | PF3D7_02140 | 36329.O96220 | T-complex protein 1 subunit theta. |
| kmeans | 5 | Brown | 7 | PF3D7_03068 | 36329.O97247 | T-complex protein 1 subunit beta. |
| kmeans | 5 | Brown | 7 | PF3D7_03203 | 36329.O97282 | T-complex protein 1 subunit epsilon. |
| kmeans | 5 | Brown | 7 | PF3D7_11322 | 36329.Q8II43 | T-complex protein 1 subunit alpha. |
| kmeans | 5 | Brown | 7 | PF3D7_12295 | 36329.Q8I5C4 | T-complex protein 1 subunit gamma. |
| kmeans | 5 | Brown | 7 | PF3D7_13578 | 36329.C0H5I7 | T-complex protein 1 subunit delta. |
| kmeans | 6 | Yellow | 7 | MCM7 | 36329.Q8IC16 | DNA replication licensing factor MCM7; Acts as component of the mcm2-7 complex (mcm complex) which is the putative replicative helicase essential for 'once per cell cycle' DNA replication initiation and elongation in eukaryotic cells. The active ATPase sites in the mcm2-7 ring are formed through the interaction surfaces of two neighboring subunits such that a critical structure of a conserved arginine finger motif is provided in trans relative to the ATP-binding site of the Walker A box of the adjacent subunit. The six ATPase active sites, however, are likely to contribute differential [...] |
| kmeans | 6 | Yellow | 7 | PF3D7_05270 | 36329.Q8I3J5 | DNA replication licensing factor MCM3, putative; Belongs to the MCM family. |
| kmeans | 6 | Yellow | 7 | PF3D7_12117 | 36329.Q8I5T4 | DNA replication licensing factor MCM5, putative; Belongs to the MCM family. |
| kmeans | 6 | Yellow | 7 | PF3D7_13171 | 36329.Q8IEE5 | DNA replication licensing factor MCM4; Belongs to the MCM family. |
| kmeans | 6 | Yellow | 7 | PF3D7_13551 | 36329.Q8IDF0 | DNA helicase; Belongs to the MCM family. |
| kmeans | 6 | Yellow | 7 | PF3D7_14121 | 36329.Q8ILX3 | Mini-chromosome maintenance complex-binding protein, putative. |
| kmeans | 6 | Yellow | 7 | PF3D7_14178 | 36329.Q8ILR7 | DNA helicase. |
| kmeans | 7 | Olive | 5 | PF3D7_09318 | 36329.Q8I0U7 | Proteasome subunit beta type-6, putative. |
| kmeans | 7 | Olive | 5 | PF3D7_10084 | 36329.Q8IJW0 | 26S protease regulatory subunit 4, putative; Belongs to the AAA ATPase family. |

| | | | | | | |
|---|---|---|---|---|---|---|
| kmeans | 7 | Olive | 5 | PF3D7_11304 | 36329.Q8II60 | 26S protease regulatory subunit 6A, putative; Belongs to the AAA ATPase family. |
| kmeans | 7 | Olive | 5 | PF3D7_12489 | 36329.Q8I4U5 | 26S protease regulatory subunit 8, putative; Belongs to the AAA ATPase family. |
| kmeans | 7 | Olive | 5 | PF3D7_13538 | 36329.Q8IDG3 | Proteasome subunit alpha type. |
| kmeans | 8 | Green Yellow | 3 | PF3D7_04061 | 36329.Q6ZMA8 | Vacuolar proton pump subunit B; Non-catalytic subunit of the peripheral V1 complex of vacuolar ATPase; Belongs to the ATPase alpha/beta chains family. |
| kmeans | 8 | Green Yellow | 3 | PF3D7_14647 | 36329.Q8IKJ0 | V-type proton ATPase subunit; Subunit of the integral membrane V0 complex of vacuolar ATPase. Vacuolar ATPase is responsible for acidifying a variety of intracellular compartments in eukaryotic cells, thus providing most of the energy required for transport processes in the vacuolar system. Belongs to the V-ATPase V0D/AC39 subunit family. |
| kmeans | 8 | Green Yellow | 3 | vapA | 36329.Q76NM6 | V-type proton ATPase catalytic subunit A; Catalytic subunit of the peripheral V1 complex of vacuolar ATPase. V-ATPase vacuolar ATPase is responsible for acidifying a variety of intracellular compartments in eukaryotic cells; Belongs to the ATPase alpha/beta chains family. |
| kmeans | 9 | Olive Drab | 3 | PF3D7_02141 | 36329.O96221 | Protein transport protein SEC31. |
| kmeans | 9 | Olive Drab | 3 | PF3D7_12307 | 36329.Q8I5B3 | Protein transport protein SEC13; Belongs to the WD repeat SEC13 family. |
| kmeans | 9 | Olive Drab | 3 | PF3D7_13611 | 36329.C0H5J6 | Protein transport protein Sec24A. |
| kmeans | 10 | Green | 3 | PF3D7_04096 | 36329.Q9U0J0 | Replication protein A1, large subunit. |
| kmeans | 10 | Green | 3 | PF3D7_11323 | 36329.Q8II42 | Nucleic acid binding protein, putative. |
| kmeans | 10 | Green | 3 | PF3D7_14421 | 36329.C6S3I6 | Replication factor A protein 3, putative. |
| kmeans | 11 | Sky Blue 2 | 3 | PF3D7_05265 | 36329.Q8I3K0 | Suf domain-containing protein. |
| kmeans | 11 | Sky Blue 2 | 3 | PF3D7_11201 | 36329.Q8IIG6 | Phosphoglycerate mutase. |
| kmeans | 11 | Sky Blue 2 | 3 | PGK | 36329.P27362 | Phosphoglycerate kinase. |
| kmeans | 12 | Sky Blue | 3 | PF3D7_10089 | 36329.Q8IJV6 | Adenylate kinase; Belongs to the adenylate kinase family. |
| kmeans | 12 | Sky Blue | 3 | PF3D7_14056 | 36329.Q8IM38 | Ribonucleoside-diphosphate reductase small chain, putative. |
| kmeans | 12 | Sky Blue | 3 | PF3D7_14372 | 36329.Q8IL94 | Ribonucleoside-diphosphate reductase; Provides the precursors necessary for DNA synthesis. Catalyzes the biosynthesis of deoxyribonucleotides from the corresponding ribonucleotides. |
| kmeans | 13 | Cornflower Blue | 2 | PF10_0084 | 36329.Q7KQL5 | Tubulin beta chain; Tubulin is the major constituent of microtubules. It binds two moles of GTP, one at an exchangeable site on the beta chain and one at a non-exchangeable site on the alpha chain. |
| kmeans | 13 | Cornflower Blue | 2 | PF3D7_09037 | 36329.Q6ZLZ9 | Tubulin alpha chain; Tubulin is the major constituent of microtubules. It binds two moles of GTP, one at an exchangeable site on the beta chain and one at a non-exchangeable site on the alpha chain. |
| kmeans | 14 | Blue | 2 | PF3D7_09348 | 36329.Q7K6A0 | cAMP-dependent protein kinase catalytic subunit; Belongs to the protein kinase superfamily. |
| kmeans | 14 | Blue | 2 | PF3D7_12231 | 36329.Q7KQK0 | cAMP-dependent protein kinase regulatory subunit. |
| kmeans | 15 | Medium Slate Blue | 2 | PF3D7_10156 | 36329.Q8IJN9 | Heat shock protein 60; Belongs to the chaperonin (HSP60) family. |
| kmeans | 15 | Medium Slate Blue | 2 | PF3D7_12321 | 36329.Q8I0V3 | 60 kDa chaperonin; Belongs to the chaperonin (HSP60) family. |
| kmeans | 16 | Purple | 2 | PF3D7_09208 | 36329.Q8I2U5 | Inosine-5'-monophosphate dehydrogenase; Catalyzes the conversion of inosine 5'-phosphate (IMP) to xanthosine 5'-phosphate (XMP), the first committed and rate-limiting step in the de novo synthesis of guanine nucleotides, and therefore plays an important role in the regulation of cell growth. |
| kmeans | 16 | Purple | 2 | PF3D7_10124 | 36329.Q8IJS1 | Hypoxanthine phosphoribosyltransferase; Belongs to the purine/pyrimidine phosphoribosyltransferase family. |
| kmeans | 17 | Medium Purple | 2 | PF3D7_05077 | 36329.Q8I426 | Nuclear protein localization protein 4, putative. |
| kmeans | 17 | Medium Purple | 2 | PF3D7_14180 | 36329.Q8ILR6 | Ubiquitin fusion degradation protein 1, putative. |
| kmeans | 18 | Medium Purple 2 | 2 | PF3D7_12271 | 36329.Q8I5E7 | DNA helicase 60; Belongs to the DEAD box helicase family. |
| kmeans | 18 | Medium Purple 2 | 2 | PF3D7_14459 | 36329.Q8IL13 | ATP-dependent RNA helicase DDX5, putative; Belongs to the DEAD box helicase family. |
| kmeans | 19 | Orchid 2 | 2 | PF3D7_06114 | 36329.C6KSV9 | SWIB/MDM2 domain-containing protein. |
| kmeans | 19 | Orchid 2 | 2 | PF3D7_12252 | 36329.Q8I5G5 | Uncharacterized protein. |
| kmeans | 20 | Violet 2 | 2 | PF3D7_12056 | 36329.Q8I5Y9 | Tetratricopeptide repeat protein, putative. |
| kmeans | 20 | Violet 2 | 2 | PF3D7_12245 | 36329.Q8I5H2 | Histone chaperone ASF1, putative. |
| kmeans | 21 | Violet | 2 | PF3D7_08020 | 36329.Q8IAM0 | Glutamate dehydrogenase, putative. |

| | | | | |
|---|---|---|---|---|
| kmeans | 21 Violet | 2 | PF3D7_13457 36329.Q8I6T2 | Isocitrate dehydrogenase [NADP]; Belongs to the isocitrate and isopropylmalate dehydrogenases family. |
| kmeans | 22 Orchid | 2 | PF3D7_11348 36329.Q8II16 | Coatomer subunit delta; The coatomer is a cytosolic protein complex that binds to dilysine motifs and reversibly associates with Golgi non-clathrin- coated vesicles, which further mediate biosynthetic protein transport from the ER, via the Golgi up to the trans Golgi network. Coatomer complex is required for budding from Golgi membranes, and is essential for the retrograde Golgi-to-ER transport of dilysine-tagged proteins. |
| kmeans | 22 Orchid | 2 | PF3D7_11451 36329.Q8IHR6 | Coatomer subunit gamma; The coatomer is a cytosolic protein complex that binds to dilysine motifs and reversibly associates with Golgi non-clathrin- coated vesicles, which further mediate biosynthetic protein transport from the ER, via the Golgi up to the trans Golgi network. Coatomer complex is required for budding from Golgi membranes, and is essential for the retrograde Golgi-to-ER transport of dilysine-tagged proteins. |
| kmeans | 23 Pink | 2 | PF3D7_11307 36329.Q8II57 | Structural maintenance of chromosomes protein 1, putative. |
| kmeans | 23 Pink | 2 | PFD0685c 36329.Q8I1U7 | Structural maintenance of chromosomes protein 3 homolog; Central component of cohesin, a complex required for chromosome cohesion during the cell cycle. The cohesin complex may form a large proteinaceous ring within which sister chromatids can be trapped. At anaphase, the complex is cleaved and dissociates from chromatin, allowing sister chromatids to segregate. Cohesion is coupled to DNA replication and is involved in DNA repair. The cohesin complex plays also an important role in spindle pole assembly during mitosis and in chromosomes movement (By similarity); Belongs to the SMC fami [...] |
| kmeans | 24 Pale Violet Red | 2 | PF3D7_11281 36329.Q8II82 | Prefoldin subunit 5, putative. |
| kmeans | 24 Pale Violet Red | 2 | PF3D7_14169 36329.Q8ILS7 | Prefoldin subunit 2, putative. |
| kmeans | 25 Light Coral | 2 | PF3D7_10224 36329.Q8IJI0 | Serine/arginine-rich splicing factor 4. |
| kmeans | 25 Light Coral | 2 | PF3D7_11198 36329.Q8IIG9 | Alternative splicing factor ASF-1, putative. |
