## Supplementary material for "*Plasmodium falciparum* SET10 is a histone H3 lysine K18 methyltransferase that participates in a chromatin modulation network crucial for intraerythrocytic development": Table S8

Table S8. List of primers for the generation and verification of the *Pf*SET10 HA-KD and *Pf*SET10-TurboID-GFP lines.

| **Function** | **Name** | **Sequence 5'-3'** |
| --- | --- | --- |
| **Primers for the generation and verification of the *Pf*SET10-HA-KD line** | | |
| Cloning primers | *Pf*SET10 pSLI-HA-*glmS* SacII FP | agatctCCGCGGTGGGAAATTACGAATGTCAGAA |
| | *Pf*SET10 pSLI-HA-*glmS* XhoI RP | agatctCTCGAGACTTGTAGACATAGTTCTTTTT CTTGTTTTTTTA |
| Vector integration primers | 5' Int *Pf*SET10 pSLI-HA-*glmS* (1) | AAAATTATCGGTTTGTTCAAATTGT |
| | 3' Int *Pf*SET10 pSLI-HA-*glmS* (2) | TTCATCTTGTTTTCCATTTATTTCC |
| | pSLI-HA-*glmS* FP (3) | GCTTTACACTTTATGCTTCCGGCTCG |
| | pSLI-HA-*glmS* RP (4) | TGTCTGTTGTGCCCAGTCAT |
| **Primers for the generation and verification of the *Pf*SET10-TurboID-GFP line** | | |
| Cloning primers | *Pf*SET10 pSLI-TurboID-GFP NotI FP | tcctccGCGGCCGCAGGTGAAGAAGAAGTGGGAAA |
| | *Pf*SET10 pSLI-TurboID- | ctttactACTAGTACTTGTAGACATAGTTCTTTTTCTTGT |

| | | |
|---|---|---|
| | GFP SpeI RP | |
| Vector integration primers | 5’ Int *Pf*SET10 pSLI-TurboID-GFP (1) | AAAATTATCGGTTTGTTCAAATTGT |
| | 3’ Int *Pf*SET10 pSLI-TurboID-GFP (2) | TTCATCTTGTTTTCCATTTATTTCC |
| | pSLI-HA-*glmS* FP (3) | GCTTTACACTTTATGCTTCCGGCTCG |
| | pSLI-TurboID GFP RP (4) | CAAGTGTTGGCCATGGAA |

FP, forward primer; RP, reverse primer. Underlined, sequences of restriction sites.
